# Human alveolar macrophage response to *Mycobacterium tuberculosis*: immune characteristics underlying large inter-individual variability

**DOI:** 10.1101/2022.03.14.484235

**Authors:** Wolfgang Sadee, Ian H. Cheeseman, Audrey Papp, Maciej Pietrzak, Michal Seweryn, Xiaofei Zhou, Shili Lin, Amanda M. Williams, Eusondia Arnett, Abul K. Azad, Larry S. Schlesinger

**Author notes:** These authors contributed equally to the work (Wolfgang Sadee, Ian H. Cheeseman).

## Abstract

*Mycobacterium tuberculosis* (*M.tb*) establishes residence and growth in human alveolar macrophages (AMs). Large inter-individual variation in *M.tb*-AM interactions is a potential early indicator of TB risk and efficacy of therapies and vaccines. Herein, we systematically analyze interactions of a virulent *M.tb* strain with freshly isolated human AMs from 28 healthy adult donors, measuring host RNA expression and secreted candidate proteins associated with TB pathogenesis over 72h. We observe large inter-individual differences in bacterial uptake and growth, with tenfold variation in *M.tb* load at 72h, reflected by large variation of gene expression programs. Systems analysis of differential and variable RNA and protein expression identifies TB-associated genes and networks (*e.g., IL1B* and *IDO1*). RNA time profiles document early stimulation of M1-type macrophage gene expression followed by emergence of an M2-type profile. The fine-scale resolution of this work enables the separation of genes and networks regulating early *M.tb* growth dynamics, and development of potential markers of individual susceptibility to *M.tb* infection and response to therapies.

## Introduction

One-quarter of the world’s population is approximated to be infected with *Mycobacterium tuberculosis* (*M.tb*), and nearly 1.3 million people die annually from tuberculosis (TB), a continuing worldwide public health problem(WHO, 2020). While a substantial portion of the human population gets infected with *M.tb*, only a fraction of infected individuals progress to active TB. To control TB, we must understand the immunopathogenesis of *M.tb* infection, including host susceptibility or resistance factors, to develop effective diagnostic, therapeutic and vaccine strategies for different individuals and populations(Azad et al., 2020).

Aerosolized *M.tb* enters the lung alveolar space where it is phagocytosed by alveolar macrophages (AMs), unique resident cells with a complex immunologic profile, to become an intracellular pathogen. *M.tb* infection activates several macrophage immunobiological pathways, including those involved in phagocytosis, phagosome-lysosome trafficking, and triggering of inflammatory cytokines, oxidants, and cell death pathways – all processes of innate immunity. Yet, critical factors that dictate *M.tb* infection and progression to TB remain uncertain - a roadblock to understanding an individual’s susceptibility to TB(Guirado et al., 2013, Rajaram et al., 2014, Schorey and Schlesinger, 2016).

The host response to *M.tb* infection ranges from complete clearance of infection to latent infection to active TB(Pai et al., 2016). Therapy and vaccine trials further highlight response variability across populations(Azad et al., 2020, Finan et al., 2008) with BCG vaccination trials attaining only 50% efficacy(Colditz et al., 1994). Evolutionary adaptation of *M.tb* to the host is considered a main factor modulating virulence, host response, and TB severity(Sousa et al., 2020, Liu et al., 2017). For example, evasion of immune surveillance by suppression of IL1B was proposed to be largely dictated by virulence of the *M.tb* strain(Sousa et al., 2020); however, expression of differentially expressed (DE) genes during infection with a single *M.tb* strain can vary >10-fold between individuals(Azad et al., 2013). Therefore, both genetic(Azad et al., 2012) and environmental/epigenetic(Thuong et al., 2008, Barreiro et al., 2012, Moores et al., 2017, Rajaram et al., 2011) host factors also play a role, while being less well understood.

Heritability estimates of susceptibility to TB range from 80% (twin and population studies) to 25-50% polygenic risk scores(Naranbhai, 2016); yet, GWAS-significant genetic variants alone fail to account for most of the estimated heritability(Naranbhai, 2016, Uren et al., 2017). Both *in vitro* and *in vivo* transcriptome studies of *M.tb*-infected macrophages and other immune cells reveal DE candidate genes(Lavalett et al., 2017, Roy et al., 2018, Rothchild et al., 2019, Moreira-Teixeira et al., 2020), resulting in gene networks (Rajaram et al., 2014, Barreiro et al., 2012, Blischak et al., 2015, Wu et al., 2014, Bragina et al., 2016), including type I IFN-associated signatures in active TB(Moreira-Teixeira et al., 2020). Similarly, protein studies invoke factors associated with TB status, including indoleamine oxidase 1 (IDO1), a marker of active TB(Adu-Gyamfi et al., 2017). Complexity of these interacting processes confound predictions of individual TB risk.

Our study addresses a critical gap in the field, the poorly understood variation of host innate immune responses between individuals during *M.tb* infection, by characterizing immune response genes and pathways that differ between individuals during infection of human AMs from healthy donors with a single virulent *M.tb* strain. We identify host genes and functional networks with highly variable expression between AM donors correlating with *M.tb* growth dynamics in AMs, some already recognized as potential biomarkers of active TB(Adu-Gyamfi et al., 2017, Gautam et al., 2018, Zak et al., 2016, Moreira-Teixeira et al., 2020, Sousa et al., 2020).

## Results

### Heterogeneity in *M.tb* uptake, adaptation and growth in human AMs among donors

We infected human AMs from 28 adult healthy donors over 2h with a virulent *M.tb* strain (H_37_R_v_) engineered to emit light (luminescence RLU values), and followed RLUs over 72h (**Figure 1A,B**). After batch effect correction, large variations in RLU values are detectable at each time point (**Figure 1B; Supplementary File 1**). RLUs correlate with *M.tb* colony forming units (CFUs), jointly reflecting the number of intracellular bacilli and microbial metabolic activity(Salunke et al., 2015), while early RLU levels at 2-24h post-infection largely reflect *M.tb* cellular uptake and early intracellular adaptation. AMs with the highest and lowest RLUs display a nearly tenfold difference at 72h. *M.tb* growth rates (24-72h post infection) are consistent with the expected *M.tb* doubling time (here referred to as generation time, in minutes required for doubling over 48-72h post infection), while varying 2-3 fold between donors. The assays presented throughout were conducted at a multiplicity of infection (MOI) of 2:1 (*M.tb*/AM cells). Bacterial growth rates (24-72h) were similar using a MOI of 10:1 (**Supplementary File 1**). To identify proteins and genes that influence *M.tb*-AM interactions, we focus on generation times (48-72h) in this study, since RLU levels prior to 48h post infection reflect *M.tb* uptake, adaptation, and early growth combined.

**Figure 1.**
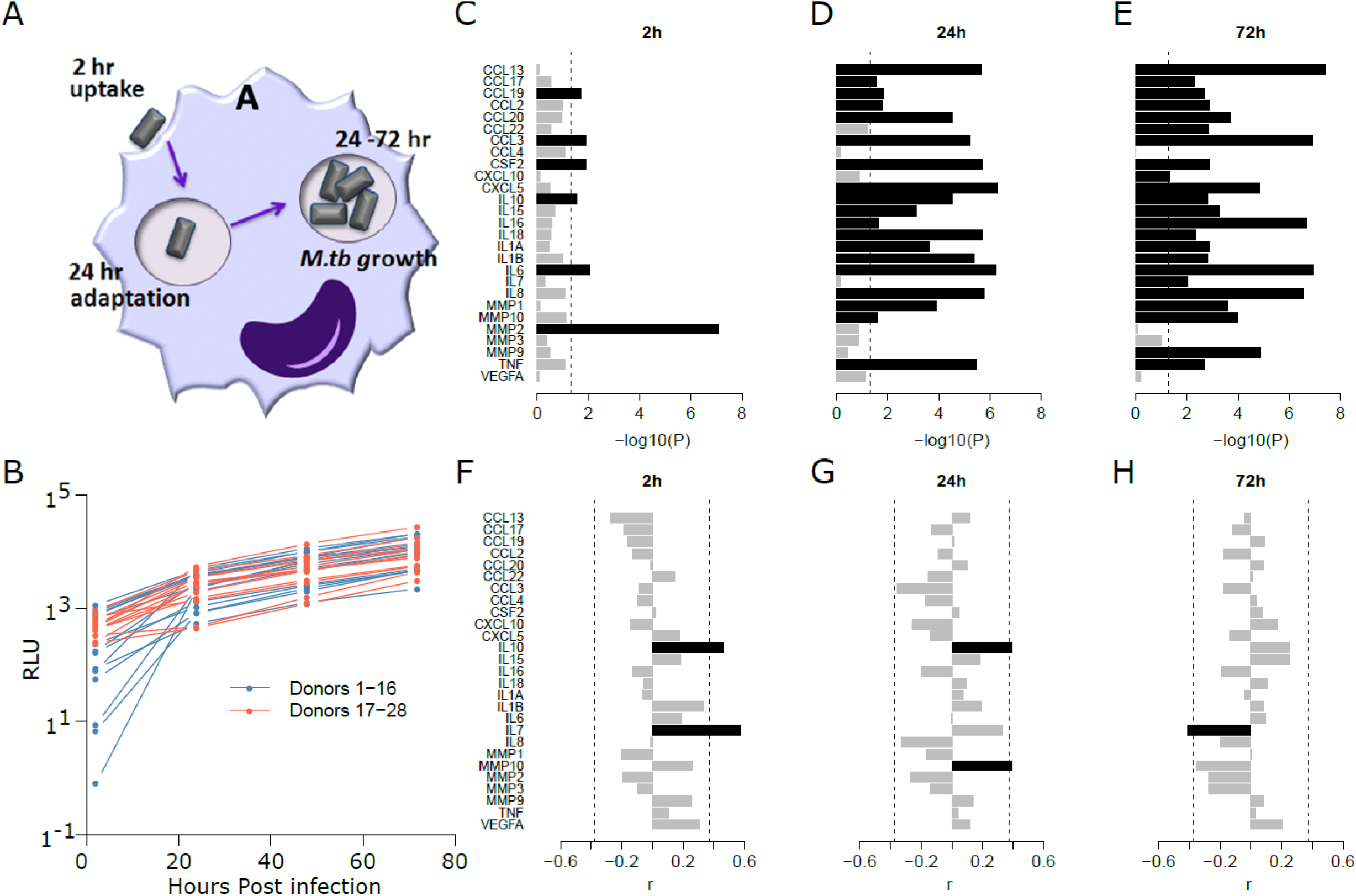
Panel A. Schematic of *M.tb* cell adhesion and uptake, adaptation and growth in human alveolar macrophages (AMs). **Panel B**. Changes in normalized luminescence RLU values over the course of *M.tb* infection. The x-axis shows hours post infection, with RLUs representing mostly uptake and adaptation at 2h and 24h, and growth rates (72h/48h) (data in Supplemental Table 2; results shown here are with MOI 2:1). **Panels C, D,** and **E**. Differentially secreted proteins post infection at 2, 24, and 72h respectively (infected/controls at each time point). The bars represent adjusted p values (−log10), with significant proteins shown with black bars. **Panels F, G,** and **H**. Correlations (R^2^) between protein levels and *M.tb* generation time (time for doubling during 48 to 72h period). Positive R^2^ values indicate that increased levels correlate with increased generation times, and *vice versa*. **Figure supplement 1**. The transcriptional response of 28 human AM cultures to *in vitro* incubation and to *M.tb* infection.

### Human AM-secreted proteins capture the immune response to *M.tb* infection

To assess the inflammatory mediator response by macrophages to *M.tb* infection, we measured 27 secreted proteins, previously implicated in cellular responses to *M.tb* infection, in all AMs over 72h (**Supplementary File 2**). At 2h, 6 of the 27 proteins (22.2%) are significantly differentially produced after *M.tb* infection (**Figure 1C**), increasing to 23 proteins (85.2%) at 72h (**Figure 1E**), all with increased secretion in infected cells. Differential secretion varies over time, with IL6 and MMP2 displaying highest levels at 2h, and IL6, CCL13, and CCL3 at 72h. Pronounced *M.tb*-stimulated secretion of MMP2 only at 2h (without stimulated mRNA expression; not shown) suggests a mechanism of rapid release from the cell rather than *de novo* transcription. Pro-inflammatory mediators including IL1β, TNF, and IL6 display significant differential secretion at all-time points, indicating their contribution to a pro-inflammatory reaction to infection.

We then tested whether protein levels secreted from infected cells correlate with *M.tb* generation times (**Figure 1F-H; Supplementary File 3**), with Pearson correlation reaching r=0.57 (IL7 at 2h). Three proteins are significant (FDR adj. p values <0.05): IL7 (2h, 72h), IL10 (2h, 24h), and MMP10 (24h). High IL10 levels correlate with longer generation times (slower growth) (r=0.46 and 0.49 at 2 and 24h, respectively), whereas IL7 has a positive r value (0.57) at 2h and a negative one (−0.41) at 72h (but is present at only low levels). A similar trend occurs with MMP10. Opposite effects at different time points highlight the changing *M.tb*-AM interactions occurring over time. These results indicate that protein expression profiles over the first 3 days of infection influence *M.tb*-AM interaction dynamics.

### Transcriptomes of uninfected control and *M.tb*-infected human AMs reveal differentially expressed gene profiles

Transcript profiles were assessed for each AM sample at 2, 24, and 72h post infection, using AmpliSeq(Papp et al., 2018). At each time point, 10,000-14,000 mRNAs were detectable, and reads per million (RPM) from replicate assays were highly correlated (r^2^≥0.99), enabling sensitive detection of differentially expressed (DE) genes. A principal component analysis (PCA) of all datasets from control and infected AMs reveals that 52% of the variance in gene expression between time points resulted from exposure to *ex vivo* culture conditions alone, while *M.tb*-induced RNA expression changes are smaller but increase over time (**Figure 1 - figure supplement 1**).

To identify significant DE genes (FDR adjusted p≤0.05) specific to infection, we analyzed RNA expression by comparing uninfected control cells to infected cells at each time point(Papp et al., 2018). This yielded 62 DE genes at 2h, 2,177 at 24h, and 3,662 at 72h post infection (**Figure 2A; Supplementary File 4**). Thirty-six DE genes significant at all-time points (**Figure 3**) include those encoding inflammatory cytokines (IL1A, IL1B, TNF) and chemokine receptors (*e.g.*, CCR7) characteristic of a spectrum representing “M1 type” cell states. Gene set enrichment analysis of 72h DE gene profiles, using gene co-expression modules previously identified during macrophage differentiation(Xue et al., 2014), shows significant enrichment of M1-like modules (**Figure 2B**). An independent assignment of M1- and M2-associated genes(Orecchioni et al., 2019) reveals high overlap of DE genes at each time point with either M1 or M2 genes (**Figure 2C-F**), accounting for 44.0% of M1 genes and 34.0% of M2 DE genes at 72h. Cell type deconvolution with Quantiseq on the RNAseq profiles(Finotello et al., 2019) supports a strong shift to M1 macrophages in infected samples, peaking at 24h, whereas the M2 cell type predominates at 72h in control AMs (**Figure 4A,B**). While significant correlations between M1/M2 profiles and *M.tb* uptake/growth dynamics are not detectable, individual donors vary significantly in the expression of several characteristic M1 or M2 genes (**Figure 5**). As one example, STAT1 (a transcription factor determining M1-like state) is one of the most differentially expressed genes between infected and control macrophages at both 24h and 72h with an average of >4-fold induction of gene expression over 72h, though induction varies between AM donors from a 2-fold reduction to a 25-fold increase.

**Figure 2.**
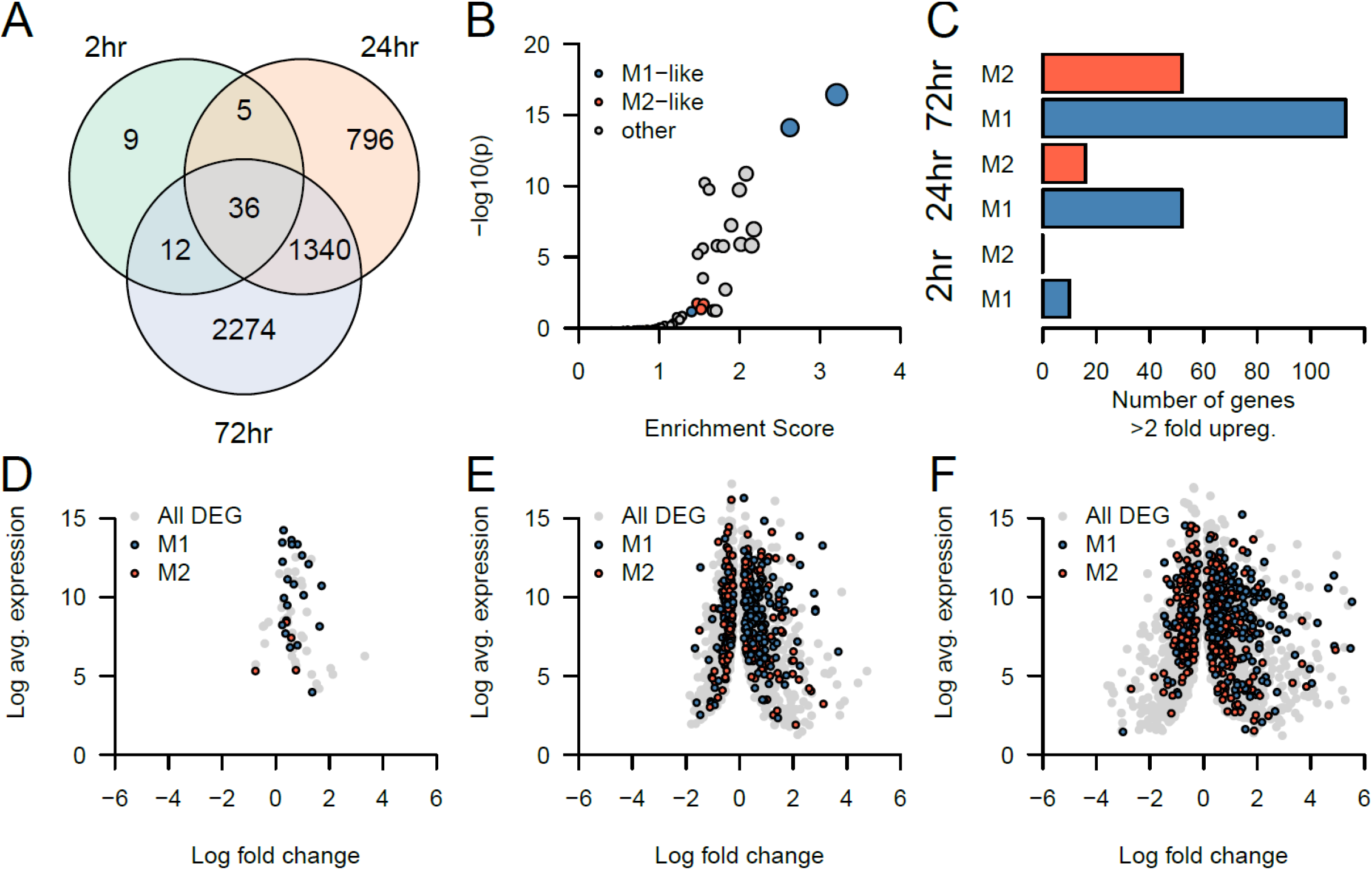
The transcriptional response of 28 human AMs to *M.tb* infection. **Panel A**. Venn diagram of differentially expressed (DE) genes between *M.tb*-treated and controls at three time points. Total number of genes with detectable expression (≥IRPM) was ~12,000. DE genes were detected with DESeq2 (adjusted p values <0.05): 62 at 2h, 2,177 at 24h, and 3,662 at 72h, with overlaps displayed in the Venn diagram. **Panel B.** Gene set enrichment analysis of DE genes from 72h using 48 gene expression modules previously determined as operational in macrophage differentiation. Modules annotated as ‘M1-like’ or ‘M2-like’ are shown in blue and red. **Panel C.** The number of DE genes with >2-fold upregulation defined as ‘M1-like’ or ‘M2-like’. This uses an independent definition of M1-like and M2-like genes to (B). **Pane D-F:** Log_2_ fold change and log_2_ average expression for DE genes at 2h (D), 24h (E) and 72h (F). M1 and M2 genes are shown in blue and red respectively.

**Figure 3.**
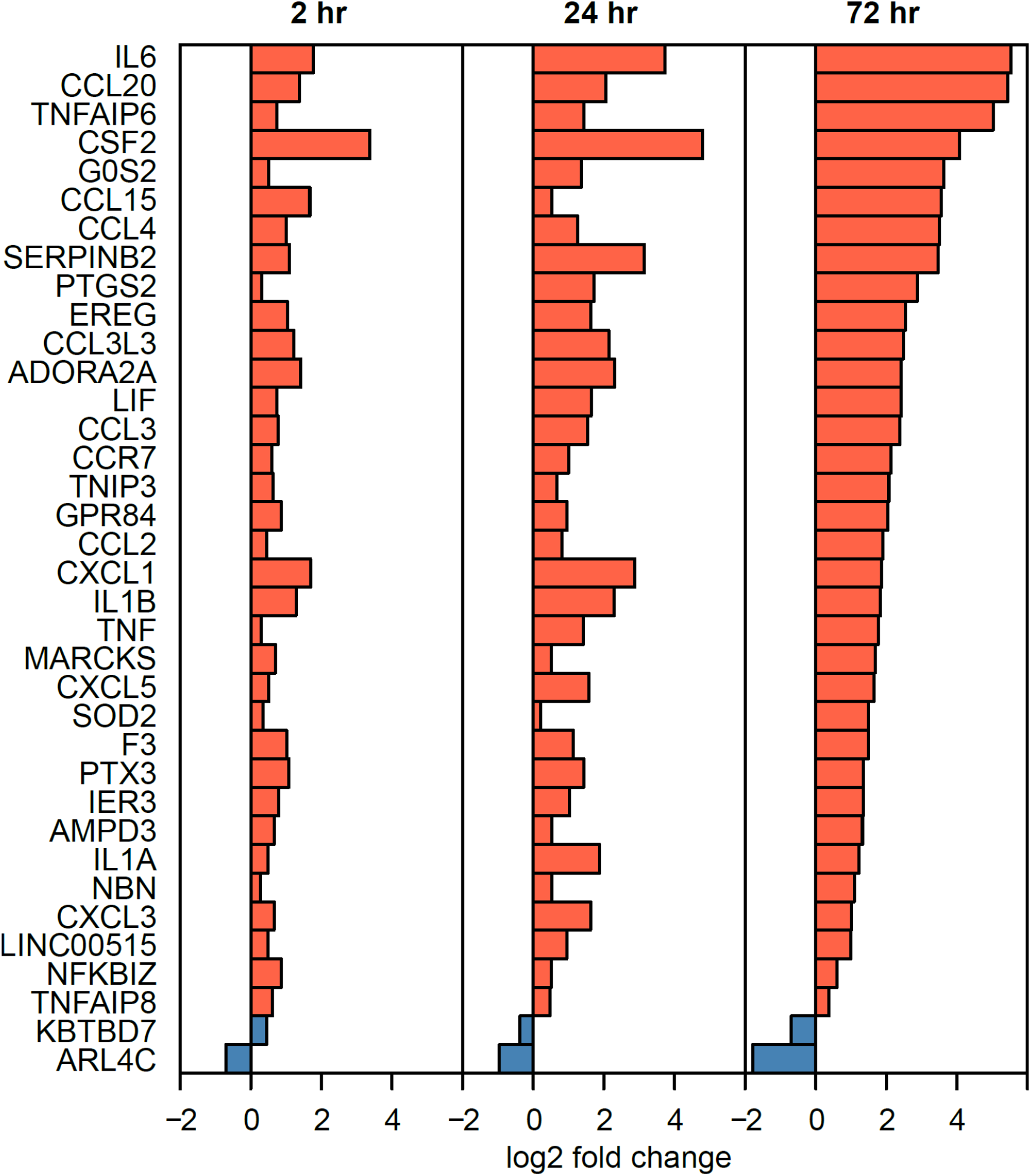
The transcriptional response of 28 human AMs to *M.tb* infection. Shown are the 36 significant DE genes common to all time points (see Figure 2A), and their log_2_ fold change.

**Figure 4.**
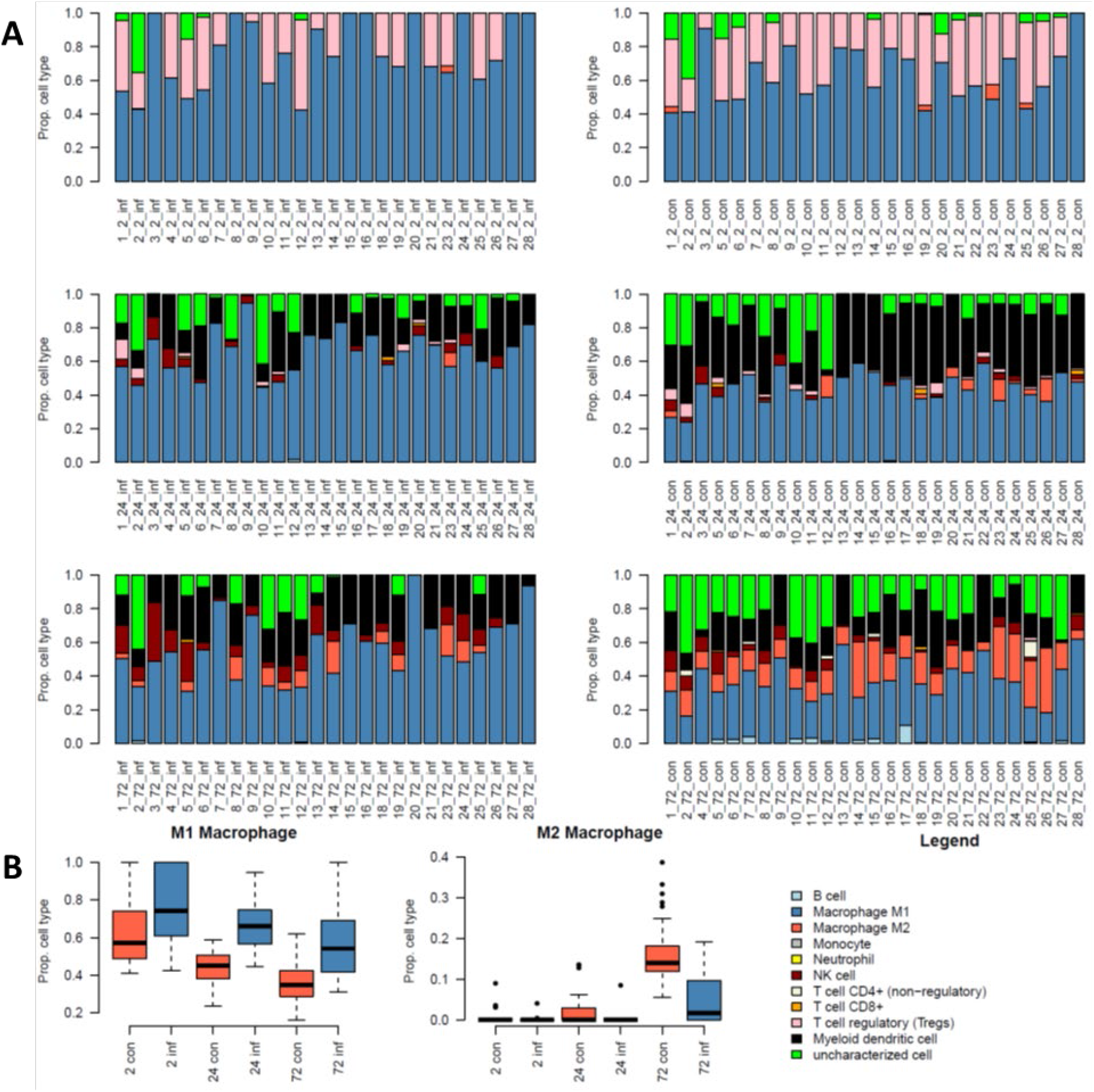
Cell type deconvolution with Quantiseq on RNAseq profiles at 2, 24 and 72h in infected and control cells. **Panel A.** Distribution of cell types visualized by the color code shown below. Note the preponderance of M1 type cells at 24h infected AMs, and of M2 type cells at 72h in controls. **Panel B.** Boxplots showing the proportion of M1 (left) and M2 (center) cells in infected and control cells at each time point, on the right is the legend for cell type proportions for the bar plots in (A).

**Figure 5.**
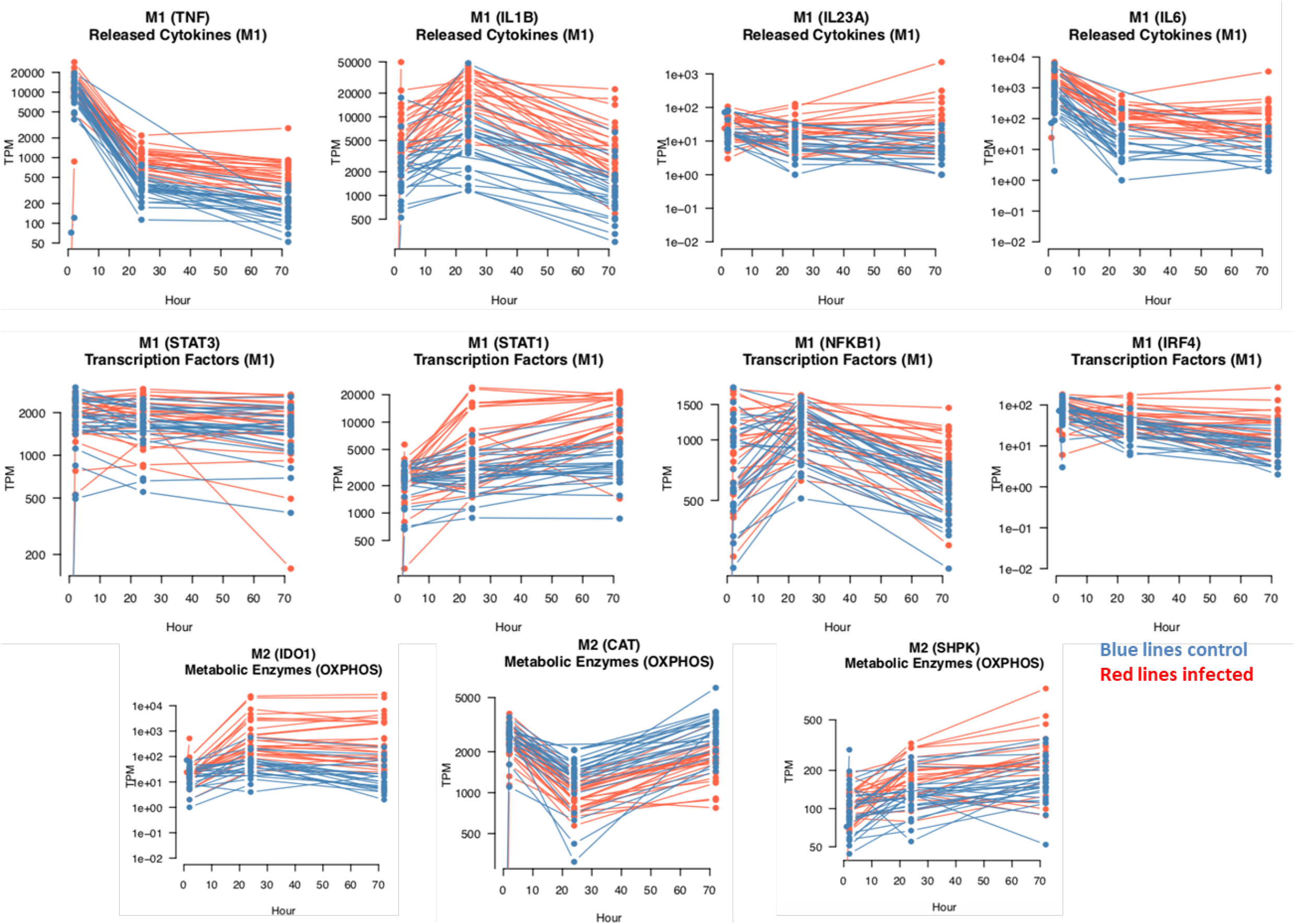
mRNA expression profiles over 72h in uninfected control and *M.tb*-infected AMs. The selected genes are marked as belonging to the M1 or M2 type macrophages, as annotated by other groups (Viola et al. 2019; Li et al. 2021), being involved in key metabolic differences between M1 and M2 types (categories: Transcription Factor, Surface Marker, Released Cytokine or Metabolic Enzyme, and sub categories such as M2A/M2B/M2C or OXPHOS/GLYCO.). https://www.ncbi.nlm.nih.gov/pmc/articles/PMC6618143/). See **Supplementary File 10.**

### Differentially expressed genes (DE genes) displaying the most variable expression among individual AMs (VE genes)

Genes with the most robust variable expression are of particular interest in understanding inter-subject differences. We performed variance analysis of gene expression in both control and infected AMs at all-time points, yielding a set of 324 genes with highly variable expression (VE genes; **Supplementary File 5A**). All 324 VE genes are specific to the infected AMs, upregulated in infected cells, and also significant DE genes (VE/DE genes). Examples include *IFI6, IL1B, CCL4, IDO1, GBP5, IRF1, JAK3, UBD, CXCL,5, CCL20, VDR, CD80, IFI44L, NLRP3*, and *IL7R*, several of which were previously implicated in TB pathogenesis. Correlations between expression of these genes with *M.tb* generation times are listed in **Supplementary File 5A**. Several genes (in RPMs) have strong correlations (r) with generation times, *e.g., MICAL2* (−0.69 at 24h), *RASSF2* (−0.73), *CD180* (−0.51), and *CNP* (0.49), the strongest correlations associated with shortened generations times.

The extent of variable expression of the VE/DE genes is exemplified by the RNA profiles across the 28 AMs (**Figure 6**). Among the VE/DE genes, expression of IDO1 mRNA is substantially stimulated by *M.tb* in only 10 of the 28 donor AMs (**Figure 6A**). IL1B is more broadly expressed, but with high expression mostly coinciding with high IDO1 expression (**Figure 6A**), similar to several other key genes (*e.g.*, IFNG, CXCL10, UBD, IFI44L; **Figure 6B**). These most variably expressed VE/DE genes are likely relevant to inter-individual variability following infection of AMs.

**Figure 6.**
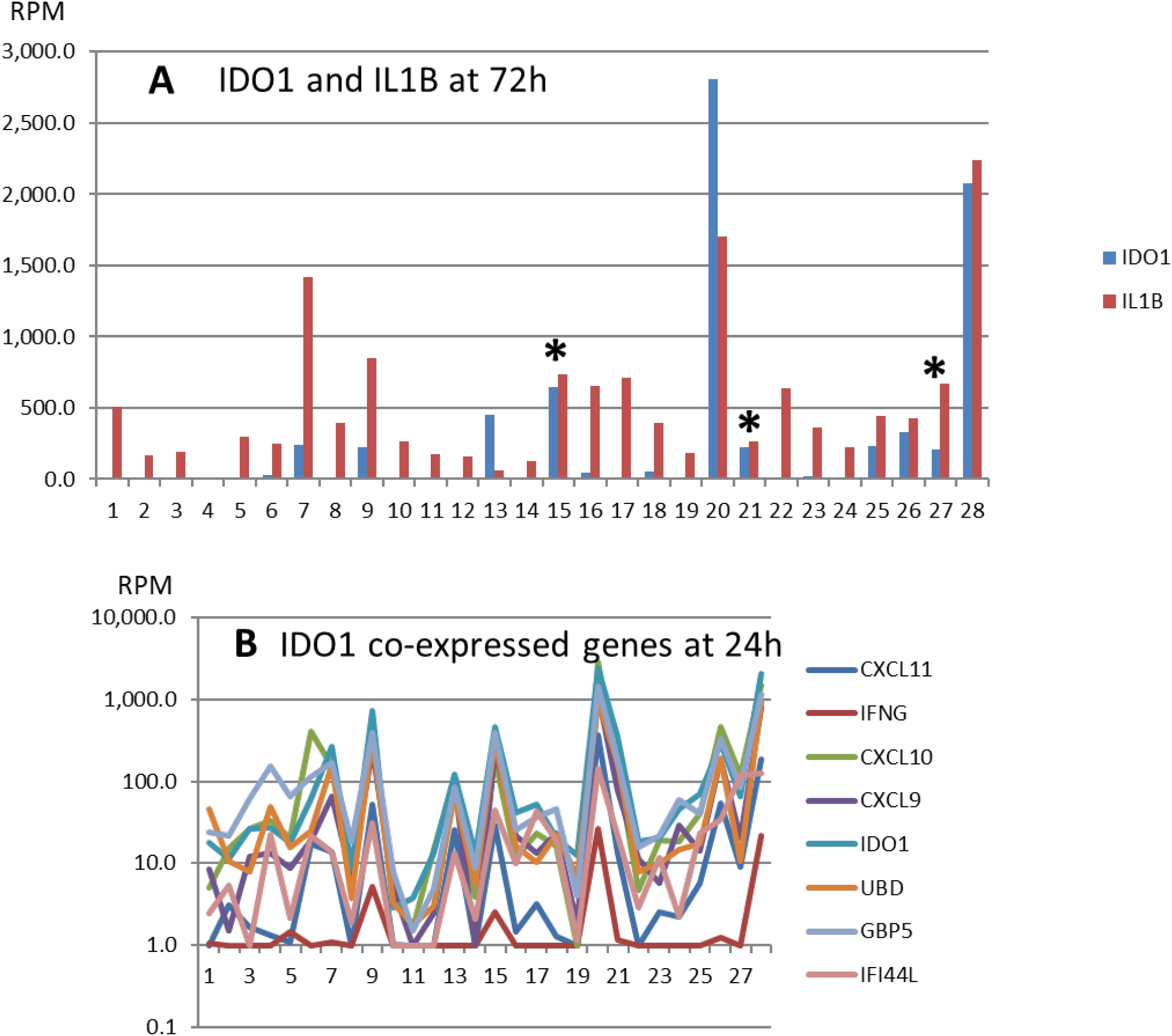
Expression of *IDO1, IL1B*, and co-expressed genes in AMs infected with *M.tb* over 72h. **Panel A.** Expression of *IDO1* and *IL1B.* Both are variably expressed with a different pattern across the 28 AMs. **Panel B**. Select mRNA transcripts with co-expression patterns similar to *IDO1. IDO1* is poorly expressed in 18 of 28 AMs (A), with tightly co-expressed genes representing the most variably expressed gene cluster (B).

Reactome analysis yields several significant pathways, including Immunological Diseases/Cellular Function and Maintenance/Inflammatory Response, and Function/Immune Cell Trafficking (**Supplementary File 5B**). Ingenuity Pathway Analysis (IPA) of the VE/DE genes yields the highest scoring hierarchical network with IL1B at the top, and with STAT1, IRF1, and IDO1 as connected nodes (**Fig. 7,** full gene annotation in **Figure 7 – figure supplement 1**). Both *STAT1* and *IRF1* interact with the *IDO1* promoter(Xue et al., 2012).

**Figure 7.**
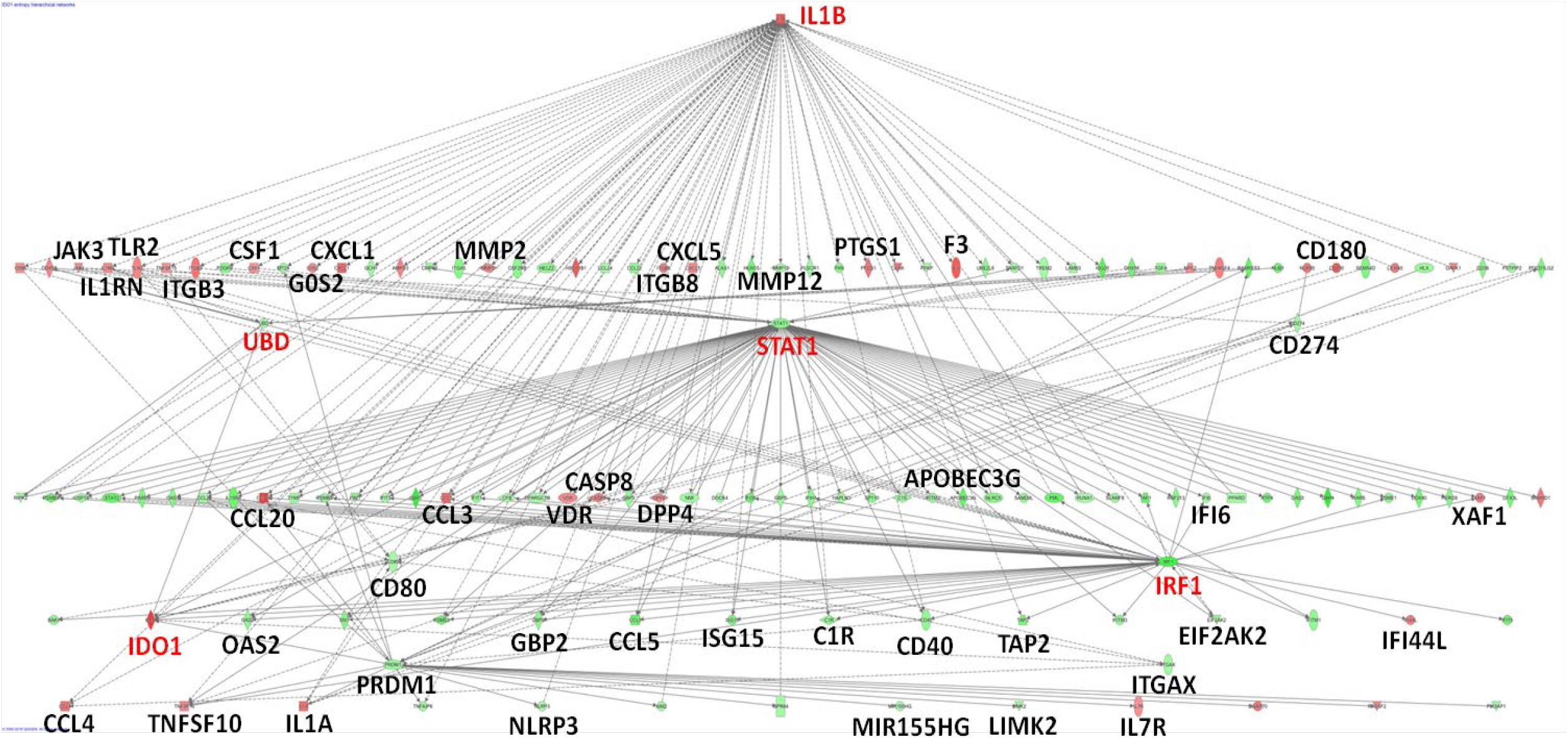
Network of DE genes with highly variable RNA expression (VE/DE genes). These VE/DE genes (n=324) were identified in the 28 AMs after *M.tb* infection across all time points. Standard pathway enrichment program, Ingenuity Pathway Analysis (IPA) (https://www.qiagenbioinformatics.com/products/ingenuity-pathway-analysis/), generates a top scoring gene network with *IL1B, UBD, STAT1*, and *IRF1* as key hub genes. *STAT1* and *IRF1* cooperatively bind to the promoter of *IDO1*, which is also highlighted (fully annotated network is depicted in Figure 7 - figure supplement 1). These key gene names are highlighted in red. **Figure supplement 1.** Top scoring network of DE genes with highly variable expression (VE/DE genes).

### Identification of human AM genes associated with *M.tb* growth

Single RNA profiles across the 28 donor AMs display high correlations with *M.tb* growth dynamics but fail to yield significance upon FDR corrections. Several candidate genes strongly correlate between RPMs at 24h and generation time, suggesting biologically relevant relationships. To explore this further, we tested correlations between *M.tb* generation times and gene expression modules, using WGCNA to build three consensus networks using batch-corrected RNA profiles from infected and control cells at 2, 24 and 72h, with RNAs having RPMs >35. This approach yielded 9,193, 9,098 and 9,115 genes, and identified 26, 27, and 37 modules for the 2, 24 and 72h time points, respectively (**Supplementary File 6A**). Correlation between network eigengene modules for infected cells and *M.tb* generation times (during 48-72h) at 2, 24 and 72h identified four significant associations between bacterial growth rate and eigengene modules, all in the 24h network (FDR-corrected p-value <0.05; blue r=-0.52, yellow r=-0.61, red r=-0.55 and tan r=0.52; **Supplementary File 7,** columns D,E, modules highlighted in red). Reactome pathway over-enrichment analysis (**Supplementary File 8**) showed Processing and Metabolism of RNA, and DNA Damage/Repair as significantly associated with these gene modules. Regulation of genes involved in DNA repair and recombination had been linked to *M.tb* regrowth from its non-replicating, persistent state(Du et al., 2016). This result supports a nexus between gene networks and *M.tb* generation times.

To prioritize genes within significantly associated modules, we measured correlations between generation time and gene expression for each gene within the four significant modules (**Supplementary File 6B**, columns Q-V). We lowered the stringency of our approach by performing multiple test correction based upon the module size, rather than for all expressed genes. This approach yielded 143 significant genes, with 139 genes belonging to the yellow module in the 24h network (97.2%). The yellow module genes have mostly negative correlations with generation times (117 of 139 genes, 84.2%); *i.e.*, elevated expression of these genes is associated with shorter generation times (higher growth rates), perhaps signaling a key *M.tb*-induced AM response favoring bacterial growth.

We performed hierarchical clustering of gene expression profiles and AM donors for genes within the yellow module, limiting to genes where we find a significant correlation between generation time and expression level (**Figure 8A**). This revealed four groups of genes, two of which contain genes with a positive correlation to generation time (clusters I and II), and two with a negative correlation to generation time (clusters III and IV). Hierarchical clustering also identifies three groups of AM donors, groups 1 and 2 have significantly longer generation times compared to group 3 (**Figure 8B**)

**Figure 8.**
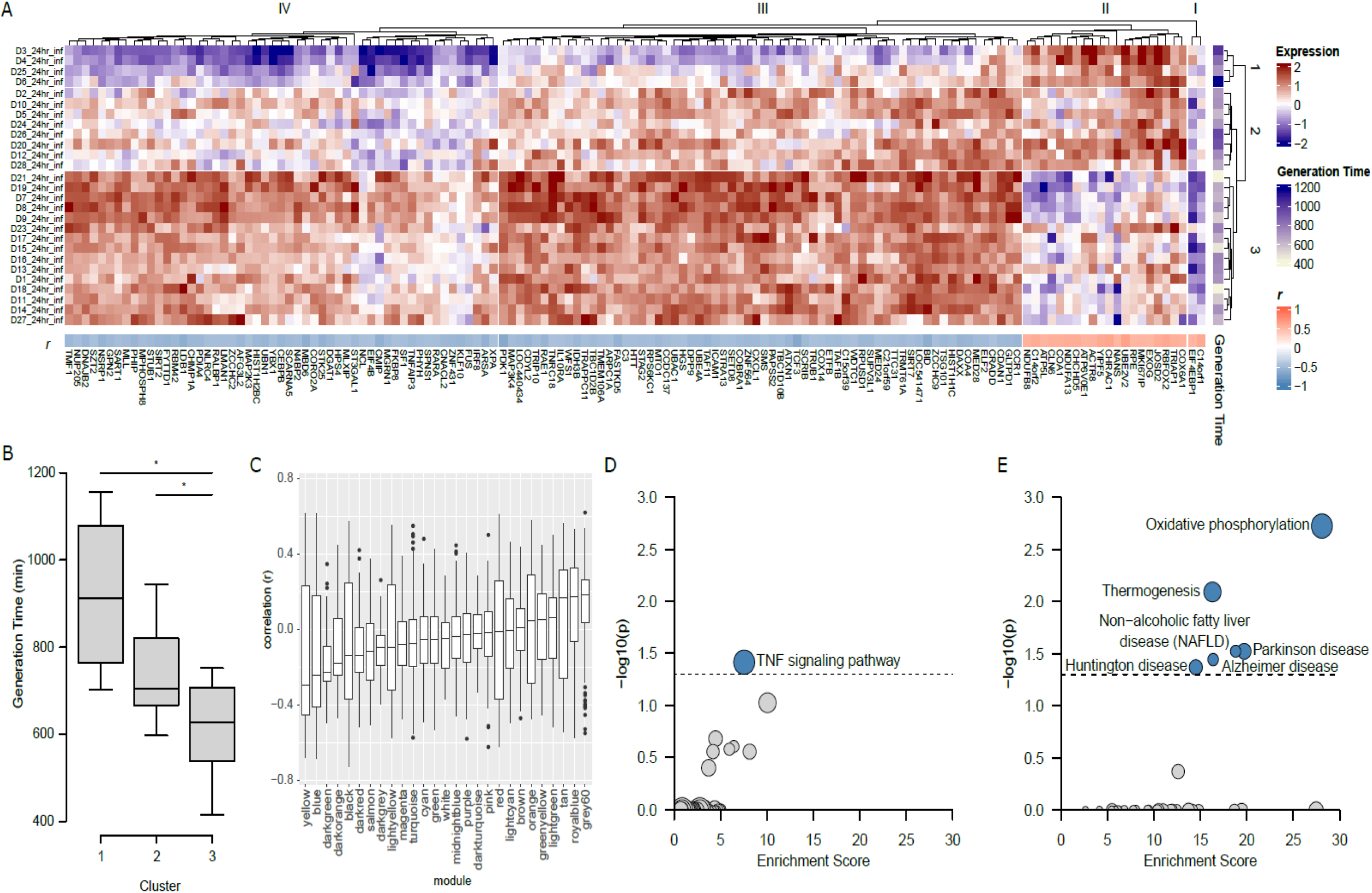
Association of the yellow gene module from 24h samples with generation time. (**A**) Gene expression profile of 139 genes from the yellow module which are correlated with generation time. Each gene is Z-normalized across all samples for ease of presentation (Expression). To the right of the heatmap is the generation time for each sample (Generation Time), and a dendrogram capturing the distance between samples. Below the heatmap are the r values for gene expression against generation time. Above the heatmap is a dendrogram capturing the distance between gene expression profiles. Three clusters of individuals were identified from the gene expression data (1-3). (**B**) The boxplot displays the generation time for individuals within each cluster; both cluster 1 and cluster 2 have significantly higher generation times than cluster 3. (**C**) The boxplot shows the distribution of r^2^ values for all modules from 24h expression data. (**D**) KEGG pathway enrichment for negatively correlated genes (clusters III and IV). (**E**) KEGG pathway enrichment for positively correlated genes (clusters I and II).

Gene expression sub-groups display opposite profiles between groups 1 and 3, with group 2 in between. We performed KEGG pathway enrichment of gene expression clusters I and II (positively correlated with generation time) and clusters III and IV (negatively correlated with generation time). This identified the TNF signaling pathway as significantly enriched in negatively correlated genes (**Figure 8D**), and oxidative phosphorylation and thermogenesis significantly enriched in positively correlated genes (**Figure 8E**). These results further support a biological relationship between gene expression profiles and *M.tb* generation times.

### Transcriptional modules coordinating the secreted protein response to *M.tb* infection in human AMs

Among the measured proteins implicated in TB pathogenesis, IL7, IL10 and MMP10 are significantly correlated with bacterial growth (Figure 1). In addition, the other measured proteins have also been shown to impact disease progression through interactions with the immune system not captured in our *in vitro* system. To identify the transcriptional modules that may regulate or respond to these proteins, we assessed correlations between eigengene modules and secreted protein levels (**Supplementary File 9**), identifying 29 significant correlations after Bonferroni correction. Only 6/29 significant correlations are in non-infected control AMs, 3 of which are at the 2h time point where there is little divergence in gene expression between control and infected conditions. Of these, CCL3 expression is strongly correlated with the royal blue module [**Supplementary Files 8** (column b) **and 9**], suggesting that the preexisting expression profile in non-infected AMs may affect *M.tb* generation times.

In the 24h network, we find three modules with a significant link to both growth rate and protein expression **(Supplementary Files 6B and 7)**. These are CCL3 (blue, red and yellow modules) and IL10 (red module). Significant correlations between these gene modules and individual proteins also suggest a causative relationship between gene expression and *M.tb* growth.

At 72h, a single gene expression module (tan) correlates with the expression of multiple proteins (IL15, IL18, IL1A and TNF). This module overlaps substantially with our highest scoring network from IPA analysis of variably expressed VE/DE genes (**Figure 7 – figure supplement 1**) with 47 of 184 module eigengenes (25.4%) falling within this network, including *IDO1, IL1A* and *IL1B*.

## Discussion

This study confirms large inter-individual variation in interactions between human AMs and a single virulent strain of *M.tb* (Figures 1 and 2), and for the first time dissects biological mechanisms underlying individual differences. Sousa et al.(Sousa et al., 2020) had demonstrated that secretion of IL1B is a surrogate marker distinguishing between TB cases with mild disease and severe TB, attributing differences in IL1B induction to virulence of the strains tested, independent of the host. However, here we show that a single virulent *M.tb* strain elicits substantial differences in IL1B expression between individual AMs, highlighting host factors, with IL1B one possible predictive marker of TB susceptibility.

The nexus between secreted TB candidate proteins and eigengene modules further supports the finding that gene expression networks, both in uninfected and *M.tb*-infected cells, affect early *M.tb*-AM interactions. While we have applied stringent FDR adjustments to reveal significant correlations, strong candidate genes and networks can serve to extract a deeper understanding of the various phases of *M.tb*-AM interactions and their possible significance in the pathogenesis of TB.

We first measured 27 candidate proteins that are secreted by *M.tb*-infected AMs and uninfected controls, finding robust stimulation by *M.tb* and inter-subject differences (Figure 1 C-H). Importantly, three proteins (IL7, IL10, and MMP10) are significantly correlated with bacterial growth over 48-72h (Supplementary File 3), supporting the hypothesis that proteins relevant to TB pathogenesis affect early *M.tb*-AM interactions, possibly presaging individual susceptibility to TB.

RNA expression profiles also appear to reflect or influence *M.tb*-AM interactions. The precision of the AmpliSeq RNA assay, together with use of control AM expression at each time point(Papp et al., 2018), enabled sensitive detection of numerous DE RNAs across the 28 AMs (Figure 2; Supplementary File 4). DE genes significant at each time point (Fig. 3) are consistent with previously reported results (*e.g.*, (Papp et al., 2018, Zak et al., 2016, Moreira-Teixeira et al., 2020)), including inflammatory cytokines CSF2, IL6, IL1B, and IFNG, but also anti-inflammatory factors such as CCL22. Gene Ontogeny (GO) Pathway analysis reveals the expected prevalence for Innate Immune System, Cytokine Signaling in Immune System, and more. The DE genes include both a classically activated M1 type induced by inflammatory agents (*e.g.*, IFNG) and alternatively activated M2 type induced by IL4 and IL13(Roy et al., 2018). While M1 and M2 markers vary between individuals, the M1 type is strongly activated at 24h post infection, while M2 markers increase at 72h, in particular in the uninfected cells (Figure 4). The balance of this mixed mode of activation may account in part for inter-subject differences in *M.tb*-AM interactions, but no significant trends are observed. In previous reports, *M.tb* ESAT-6-induced macrophage polarization to M1 phenotype occurred early, then switching to M2 phenotype at a later stage of infection(Refai et al., 2018). In our study, a substantial switch to the M2 phenotype at a later stage (72h) was observed in uninfected control macrophages compared to infected AMs, which could be due to increased expression of MMPs in control cultures. Upregulation of certain MMPs was found to be associated with macrophage polarization to both M1 and M2 phenotypes in *M.tb*-infected, cigarette smoke-exposed macrophages(Le et al., 2020).

Among the gene co-expression modules, one (yellow) significantly correlates with *M.tb* generation times and, in addition, contains numerous individual genes that also significantly correlate with *M.tb* growth (Supplementary File 6). As most correlations are negative, these genes appear to favor *M.tb* growth. The module GO terms indicate influence over translation processes, a potential mechanism by which *M.tb* usurps the cellular machinery to its advantage.

Variably expressed (VE) genes can reveal markers of individual susceptibility or resistance to disease phenotypes(Li et al., 2018), tested here for *M.tb* infection of AMs. We find that all of the most robust VE genes also classify as DE genes, likely contributing to the fate of *M.tb* in the cells. Prominent examples of VE/DE genes encode IL1B and IDO1 (Supplementary File 5), suggesting that VE/DE genes are relevant to infection and disease progression. IPA analysis of VE/DE genes yields Interferon Signaling as the top pathway, and a network with IL1B, STAT1, and IRF1 as dominant hubs including genes previously implicated in the cellular response to *M.tb* (e.g.,(Zak et al., 2016, Moreira-Teixeira et al., 2020, Rothchild et al., 2019)) (Figure 7; Figure 7 - figure supplement 1). This IL1B-dominated network connects IL1B, STAT1, and IRF1 to IDO1, consistent with previous reports that the IRF1/STAT1 transcription complex binds to the *IDO1* promoter(Xue et al., 2012, Du et al., 2000). IDO1 has been proposed recently as a sensitive and selective biomarker for active TB(Adu-Gyamfi et al., 2017). Its metabolic products, immunosuppressive kynurenins, act by preventing access of cytotoxic T cells to infected macrophages in TB lung granulomas(Gautam et al., 2018). IDO1 is robustly expressed in only 10 of 28 AMs after *M.tb* stimulation, with a similar expression profile for IL1B, CXCL9-11, IFNG, UBD, IFI44L, and GBP5 (an IFN/IL1B activated GTPase mediating antibacterial defense) (Figure 6). We propose that expression of both IL1B and IDO1 could be early predictive biomarkers of susceptibility to *M.tb* for a given individual.

The nexus we observe between secreted proteins reported relevant to TB and gene expression modules, with overlap to variably expressed VE/DE genes and, on the other hand, to *M.tb* generation times, strengthens our hypothesis that early *M.tb*-AM interactions could signal an individual’s susceptibility to infection and potentially risk of overt TB.

In summary, our results identify known and novel key genes and their encoded transcripts and proteins in the early phase of *M.tb* infection of freshly isolated human AMs. Previous work shows that infection of macrophages with *M.tb* stimulates multiple pathways, including both interferon type 1 and 2 pathways - thought to exert opposing effects on infections(Roy et al., 2018, Rothchild et al., 2019). We show here that variably and differentially expressed genes and their networks affecting *M.tb*-AM interactions can account for inter-individual variability during the early phase of infection. Results of this study provide a foundation for identifying predictive biomarkers of an individual’s susceptibility or resistance to *M.tb* infection, and response to vaccines and therapeutics(Tameris et al., 2013).

### Limitations

While the study group of 28 healthy AM donors was sufficient to reveal significant genes associated with *M.tb*-AM interactions, our results require replication. Larger cohorts will be needed to address contributions of single genes and networks to uptake, adaptation, and growth, reveal genetic factors, and assess ethnicity, sex and age, and environmental factors. Nevertheless, the study approach provides rich datasets suitable to accelerating the development of predictive biomarker panels of an individual’s risk for TB. The design of our study precluded repeat measures from each donor. However, our longstanding studies with healthy donor human monocyte-derived macrophages has demonstrated that individual donors provide consistent results for comparable measurements over time.

## Materials and Methods

### Measurements of uptake, adaptation, and growth rate of *M.tb* in infected AMs

Fresh AMs were obtained within 6h from 28 tuberculin skin test (TST)-negative healthy donors from Caucasians, Asians, Africans, according to the demographics of the Columbus, Ohio area (**Table 1**), under an approved IRB protocol at the Ohio State University Wexner Medical Center. Isolation and culture of human AMs from bronchoalveolar lavage (BAL) was done as described(Gaynor et al., 1995, Nguyen et al., 2012). Briefly, BAL fluid was centrifuged and washed once in cold RPMI at 4^0^C, and the cell pellet was re-suspended in RPMI medium. A portion of the cell suspension was subjected to cytospin followed by staining and microscopy to determine macrophage content (94 ± 5%; mean ± SD, N=10). AMs were adhered in either a 24-well plate (1.5×10^5^ cells/well) or 96-well plate (5×10^4^ cells/well) in RPMI containing 10% human AB serum and Penicillin G (10,000 U/ml) for 2h (99% pure) before collecting the supernatants and washing/lysing the adhered cell monolayer, in part used for protein and RNA analysis, respectively. Then, medium was replaced and incubations continued until 24 and 72h to repeat protein and RNA analysis.

**Table 1.** Donor demographics of AM samples used in this study (n=28)

| Donor | Sex | Age (Years) | Race | Donor | Sex | Age (Years) | Race |
| --- | --- | --- | --- | --- | --- | --- | --- |
| **D1** | Female | 24 | Asian (Chinese) | **D15** | Male | 18 | Hispanic |
| **D2** | Male | 24 | African-American | **D16** | Female | 25 | Caucasian |
| **D3** | Male | 27 | Caucasian | **D17** | Female | 20 | Caucasian |
| **D4** | Female | 26 | Caucasian | **D18** | Female | 26 | Caucasian |
| **D5** | Female | 26 | Caucasian | **D19** | Male | 19 | African-American |
| **D6** | Male | 21 | Caucasian | **D20** | Female | 22 | Hispanic |
| **D7** | Male | 24 | Caucasian | **D21** | Male | 20 | Asian (Chinese) |
| **D8** | Female | 22 | Caucasian | **D22** | Female | 19 | Not stated |
| **D9** | Male | 23 | Caucasian | **D23** | Female | 19 | Caucasian |
| **D10** | Male | 21 | Caucasian | **D24** | Male | 23 | Caucasian |
| **D11** | Female | 46 | Caucasian | **D25** | Male | 24 | Hispanic |
| **D12** | Female | 22 | African-American | **D26** | Male | 23 | Caucasian |
| **D13** | Male | 27 | Asian (Indian) | **D27** | Female | 20 | Caucasian |
| **D14** | Male | 33 | Asian (Chinese) | **D28** | Female | 19 | Caucasian |

Male/Female = 14/14; Caucasian = 60% (17/28), Asian = 14% (4/28), African-American = 11% (3/28), Hispanic = 11% (3/28)

Virulent *M.tb* H_37_R_v_ single cell suspensions were prepared(Schlesinger et al., 1990).

The virulent *M.tb* strain used contains the entire bacterial Lux operon cloned in a mycobacterial integrative expression vector (*M.tb* H_37_R_v_-Lux) as described(Salunke et al., 2015). *M.tb* cellular uptake, adaptation, and intracellular growth were assessed as relative luminescence units (RLUs) at 2, 24, 48, and 72h in 3-5 wells for each condition (mean values of 3-5 technical replicates are provided for each donor) using a multiwell plate reader Glomax (Promega). Each 2h incubation was performed at a multiplicity of infection (MOI) of both 2:1 and 10:1 (*M.tb*/AM cells for RLU assays); RNA and protein was measured only with MOI of 2:1.

For technical reasons, AM uptake and growth assays from donors 1-16 were incubated in 24 well plates, while AMs from donors 17-28 were incubated in 96 well plates. Use of the different plates resulted in ~3-fold difference in mean RLUs between the two sets of donor groups (directly related to a different number of AMs/well). For all analysis we performed batch correction of protein, transcript and growth rate measures using ComBAT(Johnson et al., 2007) to remove the impact of different plates used during culture as such we specified the batches as individuals 1-16 and 17-28.

### Measurement of secreted proteins in control and *M.tb*-infected AMs

Supernatants from culture wells of uninfected and infected AMs from 28 donors at 2, 24, and 72h were analyzed for secreted proteins relevant to *M.tb*-macrophage interactions using multiplex kits from Meso Scale Discovery (MSD) (Rockville, MD). The supernatants were collected at 2h and replaced with fresh medium, followed by continuing incubation until 24 or 72h. Twenty-seven secreted proteins were measured: 4 inflammatory mediators (TNF-α, IL-6, IL-1β, IL-10; V-PLEX kit), 6 cytokines [CSF2 (GM-CSF), IL-15, IL-16, IL-1α, IL-7, VEGFA] and 12 chemokines [IP-10 (CXCL10), MCP-1 (CCL2), MCP-4 (CCL13), MDC (CCL22), IL-8, TARC (CCL17), MIP-1α (CCL3; U-PLEX (customized multiplexing), MIP-1β (CCL4), ENA-78 (CXCL5), IL-18, MIP-3α (CCL20), MIP-3β (CCL19)], and 5 matrix metallo-proteinases (MMP-1, MMP-2, MMP-3, MMP-9, MMP-10; MMP 3-PLEX and MMP 2-PLEX).

### Measurement of RNA expression from uninfected and *M.tb*-infected human AMs

Expression of 20,804 RNAs, including 2,228 non-coding RNAs (ncRNAs), was measured with AmpliSeqTM (Whole transcriptome Human Gene Expression Kit, Life Technologies) for 28 donor AMs at 2, 24, and 72h after infection, for both uninfected controls and infected AMs at each time point (MOI 2:1). AmpliSeq transcriptome analysis incorporates a targeted, amplicon-based (~110 bps, spanning exons) workflow, and is quantitative over orders of magnitude. The precision of AmpliSeq analysis detects *M.tb*-induced expression changes with high sensitivity in human MDMs and AMs infected with *M.tb*(Papp et al., 2018). Genomic DNA and total RNA (TRIzol^®^ Reagent (Ambion™, Austin, TX)) were prepared from AMs using published procedures(Azad et al., 2013). RNA was purified, DNase-treated, RNA concentration measured, and RNA integrity assessed as described(Papp et al., 2018). After reverse transcription of 10 ng total RNA, using the AmpliSeq primers with the SuperScript® VILO™ cDNA Synthesis kit. the cDNAs were amplified for 12 cycles with Ion AmpliSeq™ primers and barcoded adapters, resulting libraries purified and pooled in equal amounts for emulsion PCR on an Ion OneTouch™ 2 instrument, followed by sequencing with the Ion Proton™ sequencer(Papp et al., 2018). Reads were aligned to BED (Browser Extensible Data) file specific for AmpliSeq amplicons. Typically, we obtain 5-9 million mappable reads per sample, with ~50-60% of RNA targets detected(Papp et al., 2018). Repeat experiments in the same sample yield correlation values of r^2^>0.99 (for both independent replicates and sequencing chip replicates).

The AM sample from D17 at 2h and 72h yielded <1 million reads and were excluded from the analyses. The AmpliSeq reads were normalized to mapped fragments per million reads for quantifying transcript expression levels(Mortazavi et al., 2008), yielding relative abundance for predicted transcripts in each AM.

### Differentially-expressed (DE) genes between control and *M.tb*-infected human AMs

To identify DE genes, we employed DESeq2(Love et al., 2014). FDR adjusted p values of 0.05 were used as cutoff for identifying DE genes at each time point. We estimated size factors using the “poscount” approach to correct for different sequencing depth and performed independent analysis at each time point, specifying the individual/donor and condition (infected or control) in the model.

### Weighted gene co-expression network analysis) (WGCNA)

We derived three gene co-expression works using WGCNA(Langfelder and Horvath, 2008) with the following parameters; power=8, TOMtype=signed, minModuleSize=30, mergeCutHeight=0.25, minKMEtoStay=0. These were blockwise consensus modules with the gene expression data from the control and infected cells grouped independently. We correlated the module eigengenes to either bacterial growth rate, or to proteins expressed/secreted from the same time point and performed FDR correction on p values.

### Detection of genes with variable expression in control and *M.tb*-infected AMs, characterized by variance measures

To identify the most variably expressed RNAs in control AMs and in those separately after *M.tb* infection, all AM transcriptome data at each time were subjected to Levene’s test(Levene, 1960), with ratios of variances as test statistics, reported adjusted p-value (FDR) for selected RNAs. A second variability test assesses whether the entropy of a given RNA’s expression is higher than expected given the total entropy in the *M.tb*-treated AMs(Handelman et al., 2015). A permutation test yields p-values for significance of the entropy computations.

### Gene pathway and ontogeny analysis

We performed over representation analysis of gene ontology (GO) terms and reactome pathways using WebGestalt(Liao et al., 2019) taking the relevant gene lists for each comparison as an input and using default parameters. We retained GO terms or pathways which reached an FDR corrected p value of 0.05. Comparison to genes associated with M1 and M2 states was performed using a curated list of genes from Viola et al.(Viola et al., 2019) and Li et al.(Li et al., 2021).

## Acknowledgements

This study was supported in part by a grant from the Bill & Melinda Gates Foundation, and a grant from the National Institutes of Health (NIH) General Medical Sciences U01 GM092655.

## Supplementary Files

**Supplementary File 1. Relative *M.tb* Luminescence Units (RLUs) and RLU ratios between time points.** All 28 AMs were incubated for 2h with *M.tb* at MOI 2:1 and 10:1, and luminescence was measured at 2, 24, 48, and 72h (n=3 per sample and time point). The RLU profiles were similar between MOI 2:1 and 10:1; therefore, only MOI 2:1 values are shown. To estimate *M.tb* generation times (equivalent to growth rates) between 24, 48, and 72h, RLU ratios between time points were calculated with unadjusted RLUs, yielding similar results with MOI 2:1 and 10:1, indicating the ratios are robust and reproducible.

**Supplementary File 2. Protein (n=27) levels secreted into the culture medium during 2, 24, and 72h incubation of uninfected AMs (Con) and after infection with *M.tb* (Mtb) from 0-2h incubation.** The incubation medium was changed once at 2h. Results are in picogram/ml, secreted into the medium by AMs over the incubation periods 0-2h, 2-24h, and 2-72h. Each value is the mean of results from duplicates of each of 2 independent samples per AM.

**Supplementary File 3. Correlations between secreted proteins and *M.tb* generation time during the 48 to 72h incubation period.** All values are r correlations, with + or - indicating whether the correlation is positive (longer generation time) or negative (shorter generation time).

**Supplementary File 4. Differential expression of RNAs control *versus M.tb* infected AMs in all 28 donor AMs at 2h, 24h and 72h (Suppl. Files 4A, B, and C), using separate non-infected control AMs for each time point.** baseMean: log2 RPMs in control cells at each time point; log2FoldChange: RPM ratios of infected/control. lfcSE: log fold-change standard error, stat: test statistic from DESeq2; pvalue: unadjusted p values. padj: FDR-adjusted p value.

**Supplementary File 5: A. Variably expressed genes (VE genes).** RNAs displaying the most variable expression between AMs based on variance and entropy in AM controls and *M.tb*-treated cells. All of the most variably expressed genes (VE Genes) were also significant DE genes (VE/DE genes), hence ordered in decreasing DE Expr Log2 Ratios (column C). All samples at all-time points were subjected to Levene’s test (using ratios of variances as a test statistic) (Levene, 1960) to find RNAs that are differentially variable between control samples and *M.tb* stimulated AMs, yielding RNAs with highest variance in cases that are induced (or inhibited) by *M.tb* stimulation only in a subset of individuals (encoded by VE genes). A second analysis asked whether the entropy of RNA expression in the *M.tb*-treated group is higher than expected given the total entropy over all AM samples (Handelman et al. 2015), using a permutation test to yield p-values for significance of the entropy computations. tau ratios are ratios of standard deviations (non-treated/treated). FDR Levene and FDR entropy are the respective adjusted p values. Columns I, J, and K present correlation values (r, red positive, blue negative) for each VE/DE gene with generation time over 48-72h (as in Suppl. File 3), using RPM expression values for each gene at 2, 24, and 72h in infected AMs. **B. Reactome Pathways significantly overrepresented among VE genes.** Results are from reactome.org/PathwayBrowser using 324 VE/DE genes.

**Supplementary File 6: A. Network module gene membership and correlations between RNA expression (log2 RPMs) at 2, 24 and 72h in infected cells, and *M.tb* generation times** (*aka* RLU ratios, growth rates, 48-72h) in *M.tb* infected and control AMs. **B. Genes with a significant association between growth rate and expression after FDR correction within modules.** Correlations are as indicated in Supplementary Files 3 and 5. The yellow module itself (24h RPMs) is also significantly correlated with generation time.

**Supplementary File 7. Summary of significant correlations between protein levels and gene modules in control and infected AMs at 2, 24 and 72 h.** Significant module correlations with generation time are also indicated (blue, red, tan, yellow; ‘yes’; column D). CCL3 is present in multiple modules. Negative correlations are highlighted in red. Adjusted p values reflect the number of proteins (n=27) and number of modules.

**Supplementary File 8. GO term enrichment for 72hr gene modules.** These four modules (blue, yellow, tan red) are significantly correlated with *M.tb* generation times (see Suppl. File 6). Results are from http://www.webgestalt.org/.

**Supplementary File 9: A-F. Correlations between protein levels and gene modules in control and infected AMs at 2, 24 and 72 h.** Shown are correlations (r, upper panels) and Adjusted p values (lower panels). Significant correlations are highlighted. A few strong correlations occur in the uninfected cells (for example at 2h: CCL3 with the Royalblue module (r=0.80, p=7.9e-7), CXCL5 with the Purple module (r=0.72, p=3.7e-5), and IL6 with the Royalblue module (r=0.70, p=6.6e-6). Positive correlations are highlighted in red, negative in blue (most in the infected AM correlations). Adjusted p values reflect the number of proteins (n=27).

**Supplementary File 10. Differentially expressed genes selected from a curated M1/M2 macrophage gene list** (Table 1 in https://www.ncbi.nlm.nih.gov/pmc/articles/PMC6618143/). Listed are DE in our dataset at any single time point. The genes are labelled as relevant to M1 or M2 state, the gene name, and the broad category (Transcription Factor, Surface Marker, Released Cytokine or Metabolic Enzyme) and sub categories such as M2A/M2B/M2C or OXPHOS/GLYCO. Fold change pval and Fold change data are from Supplementary Files 4A-C.

---

**Figure 1 - figure supplement 1.**
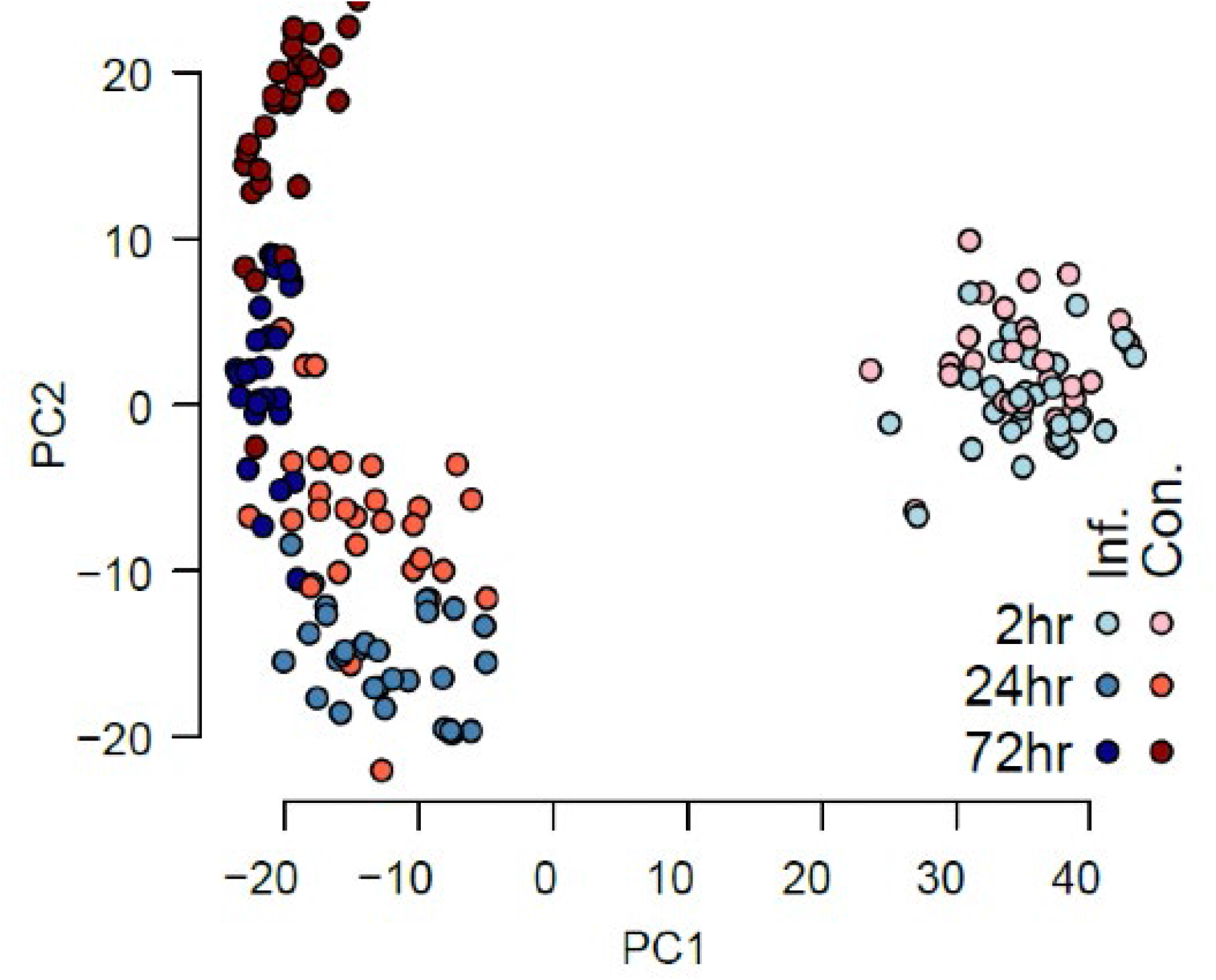
The transcriptional response of 28 human AM cultures to *in vitro* incubation and to *M.tb* infection. A principle component analysis of normalized gene expression reveals the impact of cell culture conditions with and without *M.tb* infection.

**Figure 7 - figure supplement 1.**
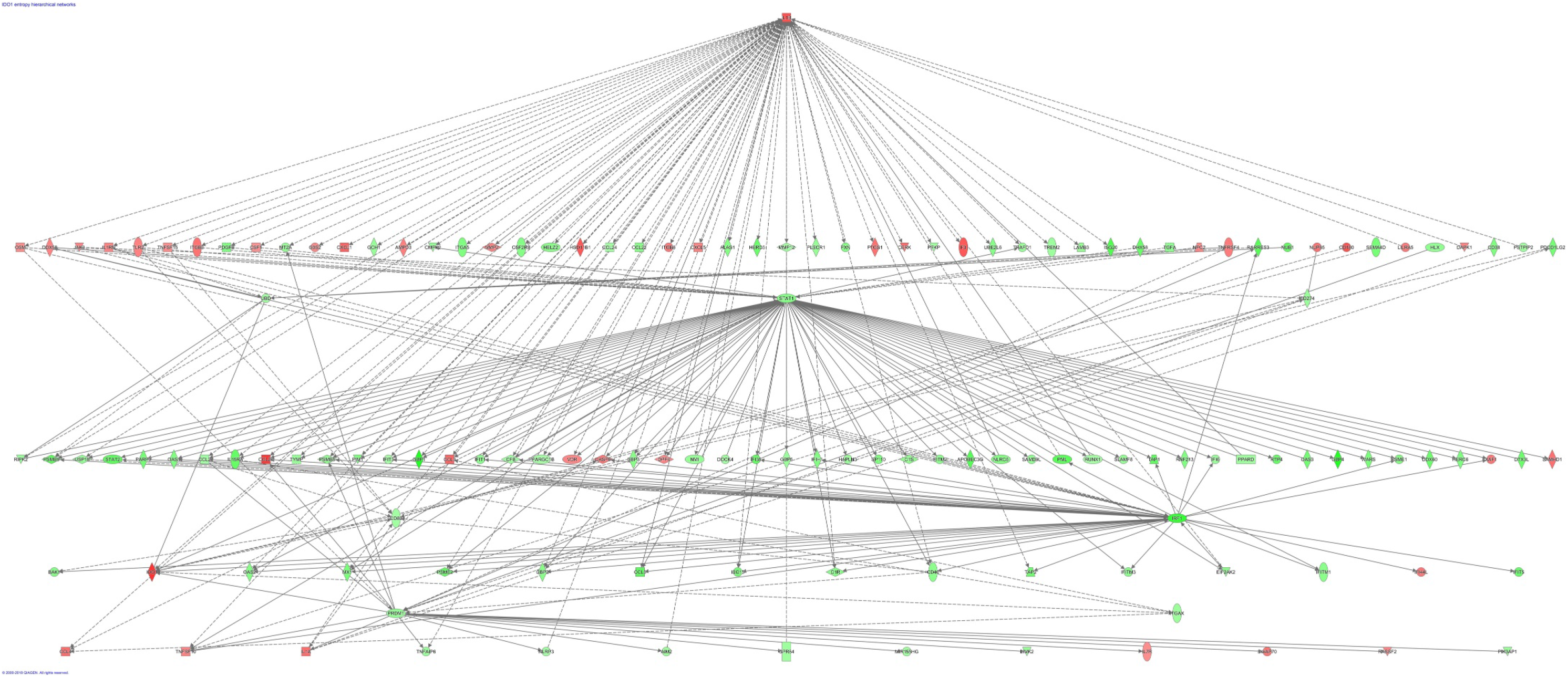
Top scoring network of DE genes with highly variable expression (VE/DE genes). The Ingenuity Pathway Analysis (IPA) (https://www.qiagenbioinformatics.com/products/ingenuity-pathway-analysis/) of VE/DE genes (n=324) generates a top scoring gene network, with IL1B on top. The same pathway is shown in Figure 7, while this graph displays all gene names.

## References

Adu-Gyamfi, C. G., Snyman, T., Hoffmann, C. J., Martinson, N. A., Chaisson, R. E., George, J. A. & Suchard, M. S. 2017. Plasma Indoleamine 2, 3-Dioxygenase, a Biomarker for Tuberculosis in Human Immunodeficiency Virus-Infected Patients. Clin Infect Dis, 65, 1356–1358.

Azad, A. K., Curtis, A., Papp, A., Webb, A., Knoell, D., Sadee, W. & Schlesinger, L. S. 2013. Allelic mRNA expression imbalance in C-type lectins reveals a frequent regulatory SNP in the human surfactant protein A (SP-A) gene. Genes Immun, 14, 99–106.

Azad, A. K., Lloyd, C., Sadee, W. & Schlesinger, L. S. 2020. Challenges of Immune Response Diversity in the Human Population Concerning New Tuberculosis Diagnostics, Therapies, and Vaccines. Front Cell Infect Microbiol, 10, 139.

Azad, A. K., Sadee, W. & Schlesinger, L. S. 2012. Innate immune gene polymorphisms in tuberculosis. Infect. Immun, 80, 3343–3359.

Barreiro, L. B., Tailleux, L., Pai, A. A., Gicquel, B., Marioni, J. C. & Gilad, Y. 2012. Deciphering the genetic architecture of variation in the immune response to Mycobacterium tuberculosis infection. Proc Natl Acad Sci U S A, 109, 1204–9.

Blischak, J. D., Tailleux, L., Mitrano, A., Barreiro, L. B. & Gilad, Y. 2015. Mycobacterial infection induces a specific human innate immune response. Sci Rep, 5, 16882.

Bragina, E. Y., Tiys, E. S., Rudko, A. A., Ivanisenko, V. A. & Freidin, M. B. 2016. Novel tuberculosis susceptibility candidate genes revealed by the reconstruction and analysis of associative networks. Infect Genet Evol, 46, 118–123.

Colditz, G. A., Brewer, T. F., Berkey, C. S., Wilson, M. E., Burdick, E., Fineberg, H. V. & Mosteller, F. 1994. Efficacy of BCG vaccine in the prevention of tuberculosis: Meta-analysis of the published literature. JAMA, 271, 698–702.

Du, M. X., Sotero-Esteva, W. D. & Taylor, M. W. 2000. Analysis of transcription factors regulating induction of indoleamine 2,3-dioxygenase by IFN-gamma. J Interferon Cytokine Res, 20, 133–42.

Du, P., Sohaskey, C. D. & Shi, L. 2016. Transcriptional and Physiological Changes during Mycobacterium tuberculosis Reactivation from Non-replicating Persistence. Front Microbiol, 7, 1346.

Finan, C., Ota, M. O., Marchant, A. & Newport, M. J. 2008. Natural variation in immune responses to neonatal Mycobacterium bovis Bacillus Calmette-Guerin (BCG) Vaccination in a Cohort of Gambian infants. PLoS One, 3, e3485.

Finotello, F., Mayer, C., Plattner, C., Laschober, G., Rieder, D., Hackl, H., Krogsdam, A., Loncova, Z., Posch, W., Wilflingseder, D., Sopper, S., Ijsselsteijn, M., Brouwer, T. P., Johnson, D., Xu, Y., Wang, Y., Sanders, M. E., Estrada, M. V., Ericsson-Gonzalez, P., Charoentong, P., Balko, J., De Miranda, N. & Trajanoski, Z. 2019. Molecular and pharmacological modulators of the tumor immune contexture revealed by deconvolution of RNA-seq data. Genome Med, 11, 34.

Gautam, U. S., Foreman, T. W., Bucsan, A. N., Veatch, A. V., Alvarez, X., Adekambi, T., Golden, N. A., Gentry, K. M., Doyle-Meyers, L. A., Russell-Lodrigue, K. E., Didier, P. J., Blanchard, J. L., Kousoulas, K. G., Lackner, A. A., Kalman, D., Rengarajan, J., Khader, S. A., Kaushal, D. & Mehra, S. 2018. In vivo inhibition of tryptophan catabolism reorganizes the tuberculoma and augments immune-mediated control of Mycobacterium tuberculosis. Proc Natl Acad Sci U S A, 115, E62–E71.

Gaynor, C. D., Mccormack, F. X., Voelker, D. R., Mcgowan, S. E. & Schlesinger, L. S. 1995. Pulmonary surfactant protein A mediates enhanced phagocytosis of Mycobacterium tuberculosis by a direct interaction with human macrophages. J Immunol, 155, 5343–51.

Guirado, E., Schlesinger, L. S. & Kaplan, G. 2013. Macrophages in tuberculosis: Friend or foe. Semin. Immunopathol, 35, 563–583.

Handelman, S. K., Seweryn, M., Smith, R. M., Hartmann, K., Wang, D., Pietrzak, M., Johnson, A. D., Kloczkowski, A. & Sadee, W. 2015. Conditional entropy in variation-adjusted windows detects selection signatures associated with expression quantitative trait loci (eQTLs). BMC Genomics, 16 Suppl 8, S8.

Johnson, W. E., Li, C. & Rabinovic, A. 2007. Adjusting batch effects in microarray expression data using empirical Bayes methods. Biostatistics, 8, 118–27.

Langfelder, P. & Horvath, S. 2008. WGCNA: an R package for weighted correlation network analysis. BMC Bioinformatics, 9, 559.

Lavalett, L., Rodriguez, H., Ortega, H., Sadee, W., Schlesinger, L. S. & Barrera, L. F. 2017. Alveolar macrophages from tuberculosis patients display an altered inflammatory gene expression profile. Tuberculosis (Edinb), 107, 156–167.

Le, Y., Cao, W., Zhou, L., Fan, X., Liu, Q., Liu, F., Gai, X., Chang, C., Xiong, J., Rao, Y., Li, A., Xu, W., Liu, B., Wang, T., Wang, B. & Sun, Y. 2020. Infection of Mycobacterium tuberculosis Promotes Both M1/M2 Polarization and MMP Production in Cigarette Smoke-Exposed Macrophages. Front Immunol, 11, 1902.

Levene, H. 1960. Robust tests for equality of variances. In: Ingram Olkin, H. H. E. A. (ed.) Contributions to Probability and Statistics. Stanford University Press.

Li, P., Hao, Z., Wu, J., Ma, C., Xu, Y., Li, J., Lan, R., Zhu, B., Ren, P., Fan, D. & Sun, S. 2021. Comparative Proteomic Analysis of Polarized Human THP-1 and Mouse RAW264.7 Macrophages. Front Immunol, 12, 700009.

Li, X., Fu, Y., Wang, X., Demeo, D. L., Tantisira, K., Weiss, S. T. & Qiu, W. 2018. Detecting Differentially Variable MicroRNAs via Model-Based Clustering. Int J Genomics, 2018, 6591634.

Liao, Y., Wang, J., Jaehnig, E. J., Shi, Z. & Zhang, B. 2019. WebGestalt 2019: gene set analysis toolkit with revamped UIs and APIs. Nucleic Acids Res, 47, W199–W205.

Liu, C. H., Liu, H. & Ge, B. 2017. Innate immunity in tuberculosis: host defense vs pathogen evasion. Cell Mol Immunol, 14, 963–975.

Love, M. I., Huber, W. & Anders, S. 2014. Moderated estimation of fold change and dispersion for RNA-seq data with DESeq2. Genome Biol, 15, 550.

Moores, R. C., Brilha, S., Schutgens, F., Elkington, P. T. & Friedland, J. S. 2017. Epigenetic Regulation of Matrix Metalloproteinase-1 and −3 Expression in Mycobacterium tuberculosis Infection. Front Immunol, 8, 602.

Moreira-Teixeira, L., Tabone, O., Graham, C. M., Singhania, A., Stavropoulos, E., Redford, P. S., Chakravarty, P., Priestnall, S. L., Suarez-Bonnet, A., Herbert, E., Mayer-Barber, K. D., Sher, A., Fonseca, K. L., Sousa, J., Ca, B., Verma, R., Haldar, P., Saraiva, M. & O’garra, A. 2020. Mouse transcriptome reveals potential signatures of protection and pathogenesis in human tuberculosis. Nat Immunol, 21, 464–476.

Mortazavi, A., Williams, B. A., Mccue, K., Schaeffer, L. & Wold, B. 2008. Mapping and quantifying mammalian transcriptomes by RNA-Seq. Nat. Methods, 5, 621–628.

Naranbhai, V. 2016. The Role of Host Genetics (and Genomics) in Tuberculosis. Microbiol Spectr, 4.

Nguyen, H. A., Rajaram, M. V., Meyer, D. A. & Schlesinger, L. S. 2012. Pulmonary surfactant protein A and surfactant lipids upregulate IRAK-M, a negative regulator of TLR-mediated inflammation in human macrophages. Am J Physiol Lung Cell Mol Physiol, 303, L608–16.

Orecchioni, M., Ghosheh, Y., Pramod, A. B. & Ley, K. 2019. Macrophage Polarization: Different Gene Signatures in M1(LPS+) vs. Classically and M2(LPS-) vs. Alternatively Activated Macrophages. Front Immunol, 10, 1084.

Pai, M., Behr, M. A., Dowdy, D., Dheda, K., Divangahi, M., Boehme, C. C., Ginsberg, A., Swaminathan, S., Spigelman, M., Getahun, H., Menzies, D. & Raviglione, M. 2016. Tuberculosis. Nat Rev Dis Primers, 2, 16076.

Papp, A. C., Azad, A. K., Pietrzak, M., Williams, A., Handelman, S. K., Igo, R. P., JR., Stein, C. M., Hartmann, K., Schlesinger, L. S. & Sadee, W. 2018. AmpliSeq transcriptome analysis of human alveolar and monocyte-derived macrophages over time in response to Mycobacterium tuberculosis infection. PLoS One, 13, e0198221.

Rajaram, M. V., Ni, B., Dodd, C. E. & Schlesinger, L. S. 2014. Macrophage immunoregulatory pathways in tuberculosis. Semin. Immunol, 26, 471–485.

Rajaram, M. V., Ni, B., Morris, J. D., Brooks, M. N., Carlson, T. K., Bakthavachalu, B., Schoenberg, D. R., Torrelles, J. B. & Schlesinger, L. S. 2011. *Mycobacterium tuberculosis* lipomannan blocks TNF biosynthesis by regulating macrophage MAPK-activated protein kinase 2 (MK2) and microRNA miR-125b. Proc Natl Acad Sci U S A, 108, 17408–13.

Refai, A., Gritli, S., Barbouche, M. R. & Essafi, M. 2018. Mycobacterium tuberculosis Virulent Factor ESAT-6 Drives Macrophage Differentiation Toward the Pro-inflammatory M1 Phenotype and Subsequently Switches It to the Anti-inflammatory M2 Phenotype. Front Cell Infect Microbiol, 8, 327.

Rothchild, A. C., Olson, G. S., Nemeth, J., Amon, L. M., Mai, D., Gold, E. S., Diercks, A. H. & Aderem, A. 2019. Alveolar macrophages generate a noncanonical NRF2-driven transcriptional response to Mycobacterium tuberculosis in vivo. Sci Immunol, 4.

Roy, S., Schmeier, S., Kaczkowski, B., Arner, E., Alam, T., Ozturk, M., Tamgue, O., Parihar, S. P., Kawaji, H., Itoh, M., Lassmann, T., Carninci, P., Hayashizaki, Y., Forrest, A. R. R., Guler, R., Bajic, V. B., Brombacher, F. & Suzuki, H. 2018. Transcriptional landscape of Mycobacterium tuberculosis infection in macrophages. Sci Rep, 8, 6758.

Salunke, S. B., Azad, A. K., Kapuriya, N. P., Balada-LLASAT, J. M., Pancholi, P., Schlesinger, L. S. & Chen, C. S. 2015. Design and synthesis of novel anti-tuberculosis agents from the celecoxib pharmacophore. Bioorg Med Chem, 23, 1935–43.

Schlesinger, L. S., Bellingerkawahara, C. G., Payne, N. R. & Horwitz, M. A. 1990. Phagocytosis of Mycobacterium-Tuberculosis Is Mediated by Human Monocyte Complement Receptors and Complement Component-C3. Journal of Immunology, 144, 2771–2780.

Schorey, J. S. & Schlesinger, L. S. 2016. Innate Immune Responses to Tuberculosis. Microbiol Spectr, 4.

Sousa, J., Ca, B., Maceiras, A. R., Simoes-Costa, L., Fonseca, K. L., Fernandes, A. I., Ramos, A., Carvalho, T., Barros, L., Magalhaes, C., Chiner-Oms, A., Machado, H., Veiga, M. I., Singh, A., Pereira, R., Amorim, A., Vieira, J., Vieira, C. P., Bhatt, A., Rodrigues, F., Rodrigues, P. N. S., Gagneux, S., Castro, A. G., Guimaraes, J. T., Bastos, H. N., Osorio, N. S., Comas, I. & Saraiva, M. 2020. Mycobacterium tuberculosis associated with severe tuberculosis evades cytosolic surveillance systems and modulates IL-1beta production. Nat Commun, 11, 1949.

Tameris, M. D., Hatherill, M., Landry, B. S., Scriba, T. J., Snowden, M. A., Lockhart, S., Shea, J. E., Mcclain, J. B., Hussey, G. D., Hanekom, W. A., Mahomed, H. & Mcshane, H. 2013. Safety and efficacy of MVA85A, a new tuberculosis vaccine, in infants previously vaccinated with BCG: a randomised, placebo-controlled phase 2b trial. Lancet, 381, 1021–1028.

Thuong, N. T., Dunstan, S. J., Chau, T. T., Thorsson, V., Simmons, C. P., Quyen, N. T., Thwaites, G. E., Thi Ngoc, L. N., Hibberd, M., Teo, Y. Y., Seielstad, M., Aderem, A., Farrar, J. J. & Hawn, T. R. 2008. Identification of tuberculosis susceptibility genes with human macrophage gene expression profiles. PLoS. Pathog, 4, e1000229.

Uren, C., Henn, B. M., Franke, A., Wittig, M., Van Helden, P. D., Hoal, E. G. & Moller, M. 2017. A post-GWAS analysis of predicted regulatory variants and tuberculosis susceptibility. PLoS One, 12, e0174738.

Viola, A., Munari, F., Sanchez-Rodriguez, R., Scolaro, T. & Castegna, A. 2019. The Metabolic Signature of Macrophage Responses. Front Immunol, 10, 1462.

WHO 2020. Global tuberculosis report 2020. Geneva: World Health Organization.

Wu, K., Fang, H., Lyu, L. D., Lowrie, D. B., Wong, K. W. & Fan, X. Y. 2014. A derived network-based interferon-related signature of human macrophages responding to Mycobacterium tuberculosis. Biomed Res Int, 2014, 713071.

Xue, J., Schmidt, S. V., Sander, J., Draffehn, A., Krebs, W., Quester, I., De Nardo, D., Gohel, T. D., Emde, M., Schmidleithner, L., Ganesan, H., Nino-Castro, A., Mallmann, M. R., Labzin, L., Theis, H., Kraut, M., Beyer, M., Latz, E., Freeman, T. C., Ulas, T. & Schultze, J. L. 2014. Transcriptome-based network analysis reveals a spectrum model of human macrophage activation. Immunity, 40, 274–88.

Xue, Z. T., Sjogren, H. O., Salford, L. G. & Widegren, B. 2012. An epigenetic mechanism for high, synergistic expression of indoleamine 2,3-dioxygenase 1 (IDO1) by combined treatment with zebularine and IFN-gamma: potential therapeutic use in autoimmune diseases. Mol Immunol, 51, 101–11.

Zak, D. E., Penn-Nicholson, A., Scriba, T. J., Thompson, E., Suliman, S., Amon, L. M., Mahomed, H., Erasmus, M., Whatney, W., Hussey, G. D., Abrahams, D., Kafaar, F., Hawkridge, T., Verver, S., Hughes, E. J., Ota, M., Sutherland, J., Howe, R., Dockrell, H. M., Boom, W. H., Thiel, B., Ottenhoff, T. H. M., Mayanja-Kizza, H., Crampin, A. C., Downing, K., Hatherill, M., Valvo, J., Shankar, S., Parida, S. K., Kaufmann, S. H. E., Walzl, G., Aderem, A., Hanekom, W. A., ACS & GROUPS, G. C. C. S. 2016. A blood RNA signature for tuberculosis disease risk: a prospective cohort study. Lancet, 387, 2312–2322.

