## Supplemental Table 1 for "Human alveolar macrophage response to *Mycobacterium tuberculosis*: immune characteristics underlying large inter-individual variability"

**Supplementary File 1: Relative M.tb Luminescence Units (RLUs) and RLU ratios between time points**

| AM | 2hr RLU avg MOI=2 | 24hr RLU avg MOI=2 | 48hr RLU avg MOI=2 | 72hr RLU avg MOI=2 | |
|---|---|---|---|---|---|
| **D4** | 21 | 116 | 387 | 750 | 1560 |
| **D21** | 4 | 62 | 416 | 885 | 1589 |
| **D18** | 18 | 145 | 397 | 933 | 2270 |
| **D13** | 28 | 268 | 842 | 1602 | 2904 |
| **D5** | 24 | 176 | 910 | 1732 | 3013 |
| **D15** | 13 | 120 | 774 | 1654 | 3090 |
| **D12** | 5 | 51 | 624 | 1499 | 3197 |
| **D28** | 12 | 93 | 939 | 1727 | 3333 |
| **D24** | 26 | 175 | 1411 | 2591 | 4125 |
| **D26** | 23 | 212 | 1609 | 2528 | 4472 |
| **D23** | 22 | 174 | 1123 | 2485 | 4665 |
| **D22** | 19 | 119 | 1498 | 2903 | 5827 |
| **D1** | 20 | 190 | 1896 | 3600 | 5968 |
| **D11** | 1 | 15 | 1402 | 3247 | 6175 |
| **D8** | 11 | 96 | 2001 | 2916 | 6567 |
| **D3** | 15 | 108 | 1177 | 2834 | 6652 |
| **D19** | 8 | 287 | 1713 | 3880 | 7169 |
| **D6** | 3 | 29 | 1588 | 4395 | 7249 |
| **D20** | 25 | 241 | 1942 | 3785 | 7342 |
| **D2** | 6 | 358 | 2714 | 5148 | 7704 |
| **D10** | 2 | 32 | 1663 | 4295 | 8301 |
| **D25** | 10 | 389 | 2662 | 4917 | 9274 |
| **D9** | 27 | 330 | 2427 | 4399 | 9587 |
| **D14** | 9 | 333 | 2548 | 5070 | 11656 |
| **D27** | 14 | 465 | 3243 | 6318 | 11872 |
| **D16** | 7 | 317 | 2878 | 6363 | 13299 |
| **D7** | 17 | 188 | 2870 | 7042 | 14918 |
| **D17** | 16 | 204 | 6673 | 15159 | 27105 |

| AM | 24/2 ratio MOI=10 | 24/2 ratio MOI=2 | 48/24 ratio MOI=10 | 48/24 ratio MOI=2 | 72/48 ratio MOI=10 | 72/48 ratio MOI=2 |
|---|---|---|---|---|---|---|
| **D1** | 140.6 | 52.0 | 2.92 | 2.55 | 1.87 | 1.87 |
| **D2** | 33.6 | 28.7 | 2.98 | 2.84 | 1.92 | 1.90 |
| **D3** | 43.4 | 30.4 | 2.68 | 3.05 | 1.82 | 1.62 |
| **D4** | 4.6 | 3.7 | 2.39 | 2.35 | 1.82 | 1.76 |
| **D5** | 9.3 | 6.9 | 2.79 | 2.65 | 2.01 | 2.09 |
| **D6** | 7.6 | 4.2 | 2.15 | 2.09 | 1.37 | 1.47 |
| **D7** | 9.9 | 5.1 | 2.42 | 2.43 | 1.98 | 2.05 |
| **D8** | 6.7 | 3.3 | 2.12 | 2.49 | 1.71 | 1.81 |
| **D9** | 7.5 | 4.3 | 2.35 | 2.19 | 1.87 | 2.26 |
| **D10** | 7.2 | 3.8 | 1.96 | 2.03 | 1.76 | 1.85 |
| **D11** | 15.3 | 11.6 | 1.39 | 1.60 | 1.86 | 2.21 |
| **D12** | 13.9 | 5.6 | 1.81 | 2.03 | 1.85 | 1.89 |
| **D13** | 4.8 | 3.6 | 2.52 | 2.35 | 1.96 | 1.83 |
| **D14** | 8.8 | 3.9 | 2.18 | 2.15 | 1.50 | 1.84 |
| **D15** | 20.5 | 10.9 | 2.13 | 2.41 | 2.48 | 2.35 |
| **D16** | 34.0 | 32.7 | 2.14 | 2.27 | 1.60 | 1.79 |
| **D17** | 17.8 | 15.3 | 2.50 | 2.45 | 2.02 | 2.12 |
| **D18** | 6.4 | 2.7 | 2.39 | 2.35 | 1.80 | 2.43 |
| **D19** | 15.3 | 12.6 | 2.20 | 1.94 | 1.83 | 2.01 |
| **D20** | 10.2 | 10.0 | 1.89 | 1.90 | 1.55 | 1.66 |
| **D21** | 5.6 | 3.3 | 1.81 | 1.94 | 2.12 | 2.08 |
| **D22** | 11.2 | 6.5 | 2.36 | 2.21 | 1.98 | 1.88 |
| **D23** | 16.6 | 7.6 | 1.79 | 1.57 | 1.92 | 1.77 |
| **D24** | 5.2 | 5.2 | 1.90 | 1.90 | 1.74 | 1.74 |
| **D25** | 8.1 | 8.1 | 2.21 | 1.95 | 1.89 | 1.94 |
| **D26** | 18.8 | 8.1 | 1.80 | 1.84 | 1.68 | 1.59 |
| **D27** | 10.1 | 7.4 | 1.88 | 1.81 | 2.08 | 2.18 |
| **D28** | 7.3 | 3.1 | 1.97 | 1.90 | 1.84 | 1.81 |
