## Supplemental Table 2 for "Human alveolar macrophage response to *Mycobacterium tuberculosis*: immune characteristics underlying large inter-individual variability"

**Supplementary File 2. Protein (n=27) levels secreted into the culture medium during 2, 24, and 72h incubation of uninfected AMs (Con) and after infection with M.tb (Mtb) from 0-2h incubation.**

| Analytes | D1.2h.Con | D1.2h.Mtb | D1.24h.Con | D1.24h.Mtb | D1.72h.Con | D1.72h.Mtb | D2.2h.Con | D2.2h.Mtb | D2.24h.Con | D2.24h.Mtb | D2.72h.Con | D2.72h.Mtb | D3.2h.Con | D3.2h.Mtb | D3.24h.Con |
|---|---|---|---|---|---|---|---|---|---|---|---|---|---|---|---|
| **ENA-78 (CXCL5)** | 22.6 | 19.8 | 5755.1 | 18221.8 | 16034.2 | 34643.5 | 26.8 | 30.1 | 6022.2 | 15115.5 | 14822.6 | 33316.3 | 59.1 | 64.5 | 9392.3 |
| **GM-CSF (CSF2)** | 0.0 | 0.2 | 1.7 | 55.5 | 1.0 | 50.8 | 0.0 | 0.1 | 1.6 | 18.5 | 1.1 | 12.0 | 0.1 | 0.4 | 1.6 |
| **IL-10** | 1.0 | 1.4 | 19.8 | 22.2 | 16.7 | 19.5 | 0.9 | 1.5 | 12.1 | 20.6 | 15.2 | 19.3 | 2.3 | 2.6 | 17.2 |
| **IL-15** | 0.1 | 0.1 | 0.7 | 0.6 | 1.2 | 1.3 | 0.1 | 0.1 | 0.5 | 0.5 | 0.7 | 0.6 | 0.0 | 0.0 | 0.3 |
| **IL-16** | 11.3 | 6.4 | 45.9 | 43.2 | 88.4 | 106.2 | 6.1 | 10.1 | 45.5 | 42.8 | 46.1 | 55.3 | 7.1 | 4.2 | 26.0 |
| **IL-18** | 43.6 | 42.8 | 63.3 | 69.3 | 46.5 | 87.7 | 25.5 | 37.6 | 80.4 | 118.7 | 53.2 | 126.9 | 36.8 | 39.3 | 99.1 |
| **IL-1a** | 1.0 | 0.4 | 14.6 | 24.5 | 90.6 | 181.3 | 0.4 | 0.5 | 4.7 | 17.2 | 25.1 | 85.0 | 0.7 | 0.3 | 5.6 |
| **IL-1b** | 1.2 | 1.2 | 16.5 | 47.3 | 19.1 | 65.6 | 0.8 | 3.0 | 10.9 | 54.6 | 12.6 | 54.5 | 1.0 | 1.5 | 18.4 |
| **IL-6** | 11.7 | 15.8 | 84.7 | 549.2 | 89.3 | 582.6 | 6.2 | 11.7 | 52.4 | 319.6 | 62.2 | 335.6 | 21.4 | 32.8 | 97.0 |
| **IL-7** | 0.0 | 0.1 | 0.1 | 0.1 | 0.3 | 0.3 | 0.1 | 0.1 | 0.1 | 0.2 | 0.2 | 0.1 | 0.1 | 0.1 | 0.1 |
| **IL-8** | 452.7 | 449.2 | 3094.2 | 3060.9 | 3041.4 | 3058.1 | 240.6 | 294.0 | 2953.1 | 2970.8 | 3031.2 | 3039.2 | 750.6 | 806.8 | 3131.9 |
| **IP-10 (CXCL10)** | 0.2 | 0.2 | 0.5 | 2.5 | 1.0 | 9.9 | 0.1 | 0.1 | 0.4 | 1.0 | 0.4 | 18.5 | 0.3 | 0.3 | 0.7 |
| **MCP-1 (CCL2)** | 1.2 | 1.1 | 41.8 | 83.7 | 196.6 | 328.4 | 0.5 | 0.4 | 59.3 | 106.3 | 381.1 | 509.4 | 0.8 | 0.9 | 24.9 |
| **MCP-4 (CCL13)** | 1.0 | 1.1 | 5.2 | 8.5 | 5.3 | 18.6 | 1.7 | 0.9 | 2.1 | 7.1 | 1.7 | 15.1 | 2.5 | 0.9 | 7.7 |
| **MDC (CCL22)** | 0.0 | 1.0 | 21.9 | 40.5 | 170.2 | 408.5 | 3.0 | 3.5 | 27.5 | 41.3 | 76.3 | 111.1 | 1.6 | 3.4 | 44.8 |
| **MIP-1a (CCL3)** | 6.5 | 8.9 | 680.3 | 3037.7 | 635.1 | 0.0 | 3.4 | 8.1 | 127.5 | 791.7 | 74.3 | 887.7 | 16.7 | 26.8 | 1140.5 |
| **MIP-1b (CCL4)** | 2.7 | 3.8 | 568.1 | 1119.2 | 640.1 | 10876.7 | 1.5 | 4.7 | 169.4 | 439.8 | 168.1 | 380.8 | 10.3 | 14.6 | 1412.1 |
| **MIP-3a (CCL20)** | 1.4 | 1.2 | 55.0 | 246.7 | 96.6 | 609.5 | 0.2 | 1.4 | 38.3 | 144.1 | 35.5 | 200.4 | 0.0 | 1.7 | 151.0 |
| **MIP-3b (CCL19)** | 0.0 | 0.1 | 7.0 | 8.0 | 8.5 | 10.4 | 0.0 | 0.0 | 15.1 | 16.6 | 11.7 | 12.5 | 0.0 | 0.0 | 16.9 |
| **MMP-1** | 30.7 | 32.2 | 4904.7 | 4690.2 | 4749.9 | 6733.2 | 27.2 | 7.6 | 1658.3 | 1177.0 | 731.0 | 971.5 | 7.7 | 8.4 | 1356.0 |
| **MMP-10** | 3.0 | 1.9 | 1169.5 | 1371.7 | 1391.3 | 2223.1 | 0.9 | 0.0 | 1477.4 | 1623.0 | 1430.3 | 1732.8 | 4.3 | 1.4 | 912.8 |
| **MMP-2** | 145.4 | 6260.9 | 1245.8 | 1465.2 | 1865.9 | 1737.8 | 152.6 | 7512.0 | 3244.2 | 2661.3 | 3557.8 | 3117.8 | 132.8 | 5751.6 | 3294.0 |
| **MMP-3** | 9.6 | 13.1 | 114.1 | 85.3 | 87.8 | 83.1 | 6.6 | 5.8 | 129.2 | 85.8 | 57.2 | 57.2 | 11.6 | 0.0 | 175.1 |
| **MMP-9** | 559.5 | 1118.3 | 2180.3 | 1398.7 | 8054.0 | 9018.7 | 444.0 | 0.0 | 1299.5 | 1040.4 | 2083.5 | 2080.5 | 33.0 | 33.0 | 1814.7 |
| **TARC (CCL17)** | 0.0 | 0.1 | 1.3 | 1.5 | 3.7 | 14.8 | 0.4 | 0.4 | 1.1 | 2.5 | 6.1 | 14.1 | 0.0 | 0.0 | 0.8 |
| **TNF-a** | 70.2 | 70.3 | 389.7 | 565.8 | 175.5 | 397.4 | 40.8 | 51.2 | 126.1 | 203.4 | 60.4 | 131.5 | 94.5 | 127.8 | 228.1 |
| **VEGFA** | 1.4 | 1.9 | 3.8 | 0.7 | 2.9 | 1.6 | 1.4 | 0.5 | 1.3 | 1.6 | 1.9 | 1.2 | 1.4 | 1.2 | 1.5 |

| D3.24h.Mtb | D3.72h.Con | D3.72h.Mtb | D4.2h.Con | D4.2h.Mtb | D4.24h.Con | D4.24h.Mtb | D4.72h.Con | D4.72h.Mtb | D5.2h.Con | D5.2h.Mtb | D5.24h.Con | D5.24h.Mtb | D5.72h.Con | D5.72h.Mtb | D6.2h.Con | D6.2h.Mtb |
|---|---|---|---|---|---|---|---|---|---|---|---|---|---|---|---|---|
| 32906.0 | 17889.7 | 35903.8 | 297.7 | 314.3 | 33852.5 | 36280.4 | 36045.2 | 38124.6 | 21.7 | 25.6 | 1812.0 | 7548.7 | 5026.3 | 25433.4 | 41.0 | 38.7 |
| 85.6 | 0.6 | 56.2 | 0.1 | 0.1 | 1.4 | 44.4 | 1.2 | 19.0 | 0.0 | 0.1 | 0.8 | 25.8 | 0.3 | 20.3 | 0.0 | 0.1 |
| 27.0 | 15.4 | 20.8 | 3.4 | 3.1 | 21.3 | 26.9 | 27.0 | 26.4 | 0.9 | 1.1 | 13.5 | 21.2 | 17.4 | 19.7 | 1.4 | 1.5 |
| 0.4 | 1.1 | 1.3 | 0.2 | 0.2 | 1.1 | 1.2 | 3.0 | 3.8 | 0.0 | 0.1 | 0.3 | 0.3 | 0.6 | 0.7 | 0.1 | 0.1 |
| 29.8 | 27.4 | 57.9 | 5.3 | 5.0 | 69.2 | 61.6 | 96.9 | 113.8 | 6.2 | 3.4 | 72.1 | 73.7 | 100.3 | 107.0 | 5.2 | 6.7 |
| 141.8 | 93.2 | 295.5 | 51.5 | 52.1 | 143.7 | 185.0 | 157.8 | 750.3 | 31.3 | 32.3 | 90.4 | 108.7 | 63.2 | 142.2 | 33.4 | 34.2 |
| 38.3 | 15.9 | 288.8 | 0.9 | 0.6 | 27.5 | 49.2 | 98.5 | 293.4 | 0.4 | 0.3 | 8.7 | 19.7 | 32.1 | 115.3 | 0.7 | 1.4 |
| 72.9 | 16.2 | 103.5 | 2.5 | 3.0 | 21.6 | 88.8 | 22.3 | 176.7 | 1.2 | 1.1 | 12.0 | 45.1 | 14.1 | 54.4 | 1.1 | 1.3 |
| 950.8 | 88.1 | 1010.5 | 57.7 | 63.7 | 280.9 | 1253.3 | 283.5 | 1255.8 | 13.9 | 18.7 | 100.3 | 682.9 | 102.9 | 714.6 | 9.6 | 11.6 |
| 0.2 | 0.3 | 0.2 | 0.0 | 0.1 | 0.1 | 0.1 | 0.3 | 0.2 | 0.0 | 0.1 | 0.2 | 0.1 | 0.2 | 0.2 | 0.1 | 0.1 |
| 3070.7 | 3135.3 | 3091.2 | 1739.5 | 1753.9 | 3086.6 | 3093.7 | 3109.2 | 3094.1 | 398.1 | 443.2 | 3046.7 | 3059.1 | 3059.8 | 3063.0 | 341.8 | 344.3 |
| 5.0 | 0.5 | 22.4 | 1.0 | 0.9 | 5.1 | 16.8 | 3.8 | 119.7 | 0.3 | 0.3 | 1.7 | 4.1 | 1.1 | 4.6 | 0.1 | 0.1 |
| 65.1 | 95.2 | 300.6 | 2.7 | 2.4 | 113.4 | 211.4 | 787.8 | 961.9 | 0.5 | 0.5 | 59.5 | 77.3 | 429.3 | 567.1 | 0.1 | 0.1 |
| 30.4 | 10.0 | 42.8 | 1.2 | 0.0 | 14.2 | 38.4 | 8.8 | 60.8 | 2.2 | 1.6 | 2.7 | 5.7 | 3.9 | 13.0 | 1.0 | 0.5 |
| 131.3 | 375.5 | 1900.0 | 1.8 | 3.5 | 272.9 | 335.6 | 1205.5 | 2554.2 | 0.7 | 3.7 | 53.0 | 89.5 | 357.3 | 805.1 | 2.4 | 8.6 |
| 0.0 | 1029.6 | 0.0 | 11.4 | 13.5 | 3304.8 | 0.0 | 167.9 | 0.0 | 5.7 | 8.3 | 262.8 | 1223.6 | 653.9 | 4878.3 | 0.2 | 2.9 |
| 0.0 | 1040.3 | 0.0 | 18.6 | 19.5 | 1897.2 | 0.0 | 2022.5 | 0.0 | 2.7 | 4.3 | 278.9 | 631.3 | 377.6 | 1017.6 | 1.8 | 2.5 |
| 761.9 | 168.6 | 1417.6 | 5.4 | 7.9 | 845.4 | 2367.0 | 894.5 | 3806.6 | 2.6 | 1.1 | 141.5 | 513.7 | 97.2 | 510.0 | 2.4 | 0.5 |
| 17.2 | 22.0 | 28.7 | 0.0 | 0.0 | 23.4 | 27.2 | 36.9 | 45.6 | 0.0 | 0.0 | 12.6 | 15.1 | 8.5 | 9.5 | 0.0 | 0.0 |
| 3764.9 | 1543.4 | 3958.6 | 10.1 | 0.0 | 3233.0 | 6927.2 | 11641.2 | 9990.0 | 19.6 | 32.2 | 1351.3 | 1597.1 | 1970.8 | 3440.6 | 10.3 | 2.1 |
| 1626.6 | 1359.8 | 2521.4 | 5.6 | 10.5 | 2187.7 | 3079.2 | 3082.0 | 0.0 | 3.2 | 3.5 | 656.7 | 892.9 | 694.8 | 1038.3 | 1.4 | 0.4 |
| 3531.1 | 3370.3 | 3431.8 | 158.9 | 2522.9 | 2887.4 | 3010.6 | 3420.3 | 167.4 | 242.5 | 6553.7 | 3477.6 | 3496.7 | 3618.9 | 3618.8 | 88.4 | 4506.2 |
| 276.0 | 182.6 | 198.0 | 1.7 | 4.3 | 119.8 | 177.8 | 423.4 | 227.0 | 0.0 | 17.6 | 193.3 | 188.9 | 241.8 | 355.8 | 4.7 | 2.9 |
| 3353.9 | 13622.0 | 38218.7 | 520.6 | 72.8 | 2981.9 | 4780.5 | 50710.0 | 47172.8 | 825.7 | 327.0 | 2276.5 | 2897.7 | 11794.0 | 25775.7 | 0.0 | 0.0 |
| 4.1 | 2.9 | 18.8 | 0.3 | 0.3 | 5.7 | 10.0 | 18.0 | 47.0 | 0.1 | 0.5 | 1.3 | 2.4 | 5.4 | 17.7 | 0.0 | 0.0 |
| 631.5 | 128.9 | 482.2 | 202.4 | 185.3 | 461.3 | 1096.4 | 261.2 | 693.1 | 57.0 | 55.8 | 206.4 | 411.1 | 127.2 | 279.3 | 48.8 | 71.7 |
| 1.9 | 2.1 | 1.2 | 2.7 | 2.2 | 1.8 | 0.7 | 5.2 | 1.0 | 2.5 | 1.9 | 1.8 | 1.4 | 2.1 | 1.4 | 1.6 | 1.2 |

| D6.24h.Con | D6.24h.Mtb | D6.72h.Con | D6.72h.Mtb | D7.2h.Con | D7.2h.Mtb | D7.24h.Con | D7.24h.Mtb | D7.72h.Con | D7.72h.Mtb | D8.2h.Con | D8.2h.Mtb | D8.24h.Con | D8.24h.Mtb | D8.72h.Con | D8.72h.Mtb | D9.2h.Con |
|---|---|---|---|---|---|---|---|---|---|---|---|---|---|---|---|---|
| 5820.4 | 10751.7 | 13818.9 | 26886.1 | 49.1 | 38.0 | 16918.5 | 43651.1 | 37522.3 | 46584.9 | 175.1 | 213.8 | 30441.4 | 47461.0 | 47291.3 | 49680.6 | 102.6 |
| 0.7 | 5.1 | 0.1 | 2.5 | 0.1 | 0.1 | 0.9 | 32.1 | 1.0 | 48.1 | 0.1 | 0.3 | 0.4 | 49.7 | 0.5 | 20.8 | 0.2 |
| 15.2 | 18.3 | 16.2 | 19.9 | 0.2 | 0.2 | 0.5 | 4.8 | 0.2 | 8.6 | 0.1 | 0.1 | 0.4 | 2.7 | 0.2 | 0.8 | 0.2 |
| 0.5 | 0.5 | 1.3 | 1.6 | 0.1 | 0.0 | 0.6 | 0.9 | 1.7 | 2.8 | 0.1 | 0.1 | 0.8 | 0.9 | 2.1 | 2.2 | 0.1 |
| 52.5 | 57.8 | 115.8 | 148.1 | 4.0 | 2.8 | 73.5 | 88.7 | 124.5 | 173.6 | 3.9 | 4.7 | 67.5 | 72.4 | 95.8 | 134.2 | 10.5 |
| 120.3 | 135.5 | 97.1 | 153.1 | 31.7 | 27.2 | 123.7 | 206.1 | 101.6 | 469.2 | 86.9 | 86.1 | 245.6 | 341.0 | 212.9 | 734.1 | 72.8 |
| 11.1 | 30.6 | 67.1 | 196.0 | 0.4 | 0.1 | 15.8 | 66.1 | 80.6 | 610.0 | 0.7 | 0.8 | 26.4 | 62.1 | 85.9 | 445.8 | 1.2 |
| 18.9 | 35.7 | 27.8 | 51.9 | 0.1 | 0.1 | 1.4 | 54.3 | 1.7 | 94.7 | 0.1 | 0.0 | 3.2 | 34.4 | 3.8 | 65.0 | 0.1 |
| 75.4 | 291.8 | 64.3 | 289.8 | 7.0 | 3.6 | 21.8 | 605.4 | 31.7 | 1081.4 | 3.0 | 0.0 | 35.4 | 1838.9 | 37.5 | 1892.0 | 16.3 |
| 0.1 | 0.2 | 0.5 | 0.3 | 0.0 | 0.0 | 0.1 | 0.1 | 0.2 | 0.1 | 0.1 | 0.0 | 0.2 | 0.1 | 0.7 | 0.2 | 0.0 |
| 3117.8 | 3069.5 | 3124.6 | 3073.9 | 97.6 | 69.6 | 11892.2 | 42164.8 | 17393.7 | 67344.5 | 117.6 | 170.7 | 80833.4 | 235658.2 | 110941.9 | 293632.6 | 764.6 |
| 1.4 | 41.2 | 3.9 | 295.8 | 0.2 | 0.2 | 3.0 | 118.7 | 5.3 | 787.6 | 0.2 | 0.2 | 7.7 | 18.2 | 8.5 | 211.6 | 1.0 |
| 33.9 | 113.3 | 458.7 | 1111.1 | 0.5 | 0.5 | 88.2 | 291.9 | 792.2 | 1740.7 | 0.8 | 0.7 | 167.5 | 850.8 | 760.6 | 3248.5 | 1.8 |
| 4.2 | 5.1 | 5.7 | 18.4 | 0.0 | 0.0 | 1.8 | 7.1 | 4.8 | 13.1 | 0.0 | 0.6 | 11.6 | 29.1 | 11.5 | 37.6 | 0.0 |
| 15.6 | 38.7 | 134.3 | 222.3 | 0.1 | 0.0 | 124.1 | 146.6 | 647.5 | 1768.5 | 0.8 | 0.0 | 108.6 | 181.3 | 547.5 | 1393.0 | 1.0 |
| 483.2 | 1618.7 | 109.9 | 2915.9 | 5.2 | 2.8 | 218.6 | 1780.2 | 24.0 | 1965.6 | 2.5 | 5.4 | 1818.4 | 7518.8 | 344.6 | 7881.0 | 64.7 |
| 740.6 | 1675.2 | 723.1 | 3557.0 | 4.0 | 2.5 | 675.4 | 1476.4 | 377.5 | 1600.2 | 2.5 | 4.4 | 4838.9 | 6092.3 | 5116.8 | 6408.6 | 33.8 |
| 331.1 | 648.5 | 379.9 | 760.3 | 2.1 | 0.9 | 350.4 | 1998.5 | 412.3 | 3887.7 | 2.2 | 4.7 | 958.3 | 4033.0 | 1258.1 | 8086.1 | 7.6 |
| 13.2 | 13.3 | 9.9 | 8.9 | 0.0 | 0.0 | 19.5 | 31.3 | 20.9 | 68.7 | 0.0 | 0.0 | 23.7 | 29.2 | 36.4 | 41.0 | 0.0 |
| 1687.7 | 2054.9 | 2678.7 | 3412.2 | 3.6 | 16.8 | 6941.4 | 11924.7 | 8055.6 | 18060.2 | 7.6 | 8.8 | 8105.3 | 12365.8 | 10316.0 | 18673.7 | 5.3 |
| 1558.7 | 1593.4 | 2134.0 | 2131.6 | 7.4 | 3.1 | 2979.9 | 3849.9 | 3944.6 | 5889.1 | 0.4 | 0.0 | 3295.7 | 4280.5 | 4348.9 | 6046.5 | 0.0 |
| 2691.7 | 3193.2 | 3091.1 | 3165.1 | 0.0 | 4770.8 | 5058.7 | 5315.9 | 5366.3 | 5349.9 | 60.8 | 3545.5 | 4280.0 | 4579.9 | 4928.1 | 4889.5 | 0.0 |
| 263.2 | 294.7 | 321.6 | 370.6 | 15.2 | 7.4 | 456.4 | 532.8 | 449.0 | 565.1 | 11.7 | 13.7 | 387.4 | 417.8 | 452.9 | 492.4 | 3.2 |
| 1665.8 | 1875.0 | 11859.1 | 20576.3 | 0.0 | 0.0 | 8565.3 | 9725.6 | 43684.8 | 67196.6 | 0.0 | 85.1 | 2528.0 | 5019.6 | 16723.4 | 54994.6 | 0.0 |
| 0.6 | 1.2 | 5.6 | 32.2 | 0.4 | 0.3 | 2.8 | 7.0 | 16.3 | 69.6 | 0.2 | 0.2 | 3.4 | 14.0 | 20.7 | 76.3 | 0.4 |
| 322.4 | 416.8 | 168.7 | 251.0 | 52.1 | 45.0 | 218.6 | 726.9 | 55.3 | 747.0 | 41.3 | 0.0 | 257.4 | 1478.3 | 91.8 | 789.5 | 55.3 |
| 2.3 | 1.7 | 1.5 | 1.9 | 0.4 | 0.4 | 0.7 | 0.0 | 5.4 | 6.0 | 0.3 | 0.4 | 0.0 | 0.0 | 0.9 | 0.0 | 0.3 |

| D9.2h.Mtb | D9.24h.Con | D9.24h.Mtb | D9.72h.Con | D9.72h.Mtb | D10.2h.Con | D10.2h.Mtb | D10.24h.Con | D10.24h.Mtb | D10.72h.Con | D10.72h.Mtb | D11.2h.Con | D11.2h.Mtb | D11.24h.Con | D11.24h.Mtb | D11.72h.Con |
|---|---|---|---|---|---|---|---|---|---|---|---|---|---|---|---|
| 91.3 | 46207.3 | 45911.0 | 48173.7 | 48785.2 | 21.5 | 23.1 | 9855.4 | 30616.3 | 43418.8 | 47872.5 | 78.6 | 86.7 | 9165.5 | 26783.9 | 21034.8 |
| 0.2 | 12.7 | 74.7 | 10.8 | 55.6 | 0.1 | 0.1 | 0.7 | 10.0 | 1.1 | 4.2 | 0.0 | 0.0 | 0.0 | 7.1 | 0.3 |
| 0.1 | 0.9 | 2.2 | 0.7 | 5.8 | 0.1 | 0.1 | 0.7 | 0.8 | 0.3 | 0.4 | 0.1 | 0.1 | 0.3 | 0.6 | 0.1 |
| 0.0 | 0.6 | 0.9 | 1.4 | 2.1 | 0.1 | 0.0 | 0.7 | 0.7 | 1.9 | 2.0 | 0.0 | 0.0 | 0.4 | 0.3 | 1.3 |
| 6.9 | 86.2 | 92.9 | 108.1 | 146.4 | 4.1 | 6.3 | 69.6 | 72.2 | 203.5 | 221.8 | 8.2 | 5.8 | 30.4 | 35.2 | 103.3 |
| 58.1 | 211.9 | 298.8 | 215.3 | 490.0 | 31.9 | 29.3 | 75.3 | 87.3 | 68.2 | 159.3 | 16.4 | 17.2 | 62.3 | 73.5 | 48.3 |
| 0.8 | 29.5 | 116.9 | 103.3 | 703.1 | 0.3 | 0.5 | 6.6 | 11.7 | 122.0 | 230.5 | 0.3 | 0.3 | 1.9 | 7.1 | 52.8 |
| 0.1 | 5.2 | 46.4 | 7.4 | 84.6 | 0.1 | 0.1 | 1.0 | 6.6 | 2.5 | 15.4 | 0.1 | 0.1 | 0.3 | 3.0 | 1.0 |
| 13.7 | 609.0 | 1380.0 | 842.3 | 1629.4 | 0.9 | 1.5 | 19.0 | 368.3 | 19.8 | 379.7 | 0.7 | 3.1 | 3.9 | 347.4 | 4.9 |
| 0.0 | 0.1 | 0.1 | 0.2 | 0.1 | 0.1 | 0.0 | 0.1 | 0.0 | 0.4 | 0.2 | 0.0 | 0.0 | 0.2 | 0.1 | 0.4 |
| 661.9 | 183224.7 | 243204.5 | 225766.7 | 283395.2 | 0.0 | 0.0 | 51210.7 | 132438.4 | 75105.6 | 165517.1 | 86.2 | 561.6 | 9323.1 | 59383.5 | 13383.8 |
| 1.0 | 35.3 | 494.3 | 42.7 | 1339.5 | 0.3 | 0.4 | 3.1 | 3.9 | 6.5 | 85.7 | 0.1 | 0.1 | 0.8 | 0.8 | 1.9 |
| 1.9 | 413.5 | 584.7 | 917.6 | 2327.0 | 0.9 | 1.1 | 84.3 | 186.3 | 667.1 | 1241.8 | 1.6 | 2.0 | 17.9 | 41.8 | 461.9 |
| 0.5 | 43.9 | 61.5 | 62.7 | 65.1 | 0.0 | 0.0 | 6.7 | 14.4 | 3.7 | 24.9 | 0.0 | 2.1 | 0.9 | 4.9 | 0.3 |
| 0.3 | 316.0 | 328.9 | 2266.2 | 3499.2 | 0.5 | 0.0 | 82.2 | 104.1 | 332.5 | 600.3 | 0.0 | 0.2 | 31.4 | 69.9 | 115.3 |
| 57.1 | 8325.3 | 8261.8 | 8535.4 | 8522.6 | 5.1 | 8.5 | 1344.7 | 5415.2 | 319.9 | 6581.1 | 3.1 | 9.0 | 22.4 | 822.1 | 20.1 |
| 32.1 | 6457.1 | 6547.6 | 6372.8 | 6533.8 | 8.9 | 10.8 | 1336.0 | 2908.4 | 1677.8 | 4544.2 | 6.2 | 10.0 | 164.7 | 588.7 | 128.8 |
| 5.5 | 2539.4 | 3061.4 | 3700.8 | 5518.2 | 0.0 | 2.1 | 228.9 | 766.9 | 258.8 | 1160.3 | 0.5 | 3.5 | 8.6 | 117.2 | 18.9 |
| 0.0 | 32.0 | 34.3 | 58.5 | 59.3 | 0.0 | 0.0 | 29.4 | 29.2 | 22.7 | 29.4 | 0.0 | 0.5 | 17.1 | 17.2 | 14.3 |
| 0.0 | 16683.9 | 21645.3 | 20622.0 | 25854.6 | 0.0 | 0.0 | 10135.2 | 10587.8 | 11215.5 | 14941.7 | 10.7 | 10.5 | 1075.9 | 1520.8 | 1509.1 |
| 0.0 | 5160.2 | 5810.9 | 7723.5 | 7591.9 | 1.5 | 0.0 | 5856.4 | 5180.7 | 8485.8 | 7350.3 | 0.0 | 1.6 | 1423.9 | 1556.3 | 1455.0 |
| 2570.4 | 4869.6 | 4715.4 | 5079.1 | 4962.1 | 0.0 | 3610.6 | 4695.6 | 4852.1 | 4936.7 | 5038.7 | 0.0 | 2487.8 | 3939.8 | 4644.1 | 4903.7 |
| 2.6 | 451.8 | 490.0 | 496.7 | 522.2 | 14.0 | 4.7 | 470.1 | 469.8 | 424.1 | 505.4 | 16.2 | 24.3 | 154.2 | 381.4 | 401.1 |
| 0.0 | 9938.1 | 10108.4 | 89012.0 | 85647.1 | 0.0 | 0.0 | 1606.3 | 2428.8 | 14318.7 | 29137.2 | 0.0 | 0.0 | 1026.2 | 2374.6 | 13103.8 |
| 0.3 | 16.9 | 16.2 | 40.4 | 64.0 | 0.3 | 0.4 | 2.4 | 3.9 | 6.0 | 19.1 | 0.4 | 0.4 | 1.0 | 1.2 | 6.1 |
| 55.2 | 651.9 | 1283.8 | 302.6 | 1081.3 | 51.8 | 54.5 | 446.7 | 715.7 | 149.0 | 265.4 | 39.3 | 48.5 | 216.0 | 471.9 | 67.5 |
| 0.5 | 0.0 | 0.0 | 0.0 | 0.2 | 0.3 | 0.3 | 0.7 | 0.2 | 13.4 | 0.4 | 0.2 | 0.0 | 0.0 | 0.0 | 0.0 |

| D11.72h.Mtb | D12.2h.Con | D12.2h.Mtb | D12.24h.Con | D12.24h.Mtb | D12.72h.Con | D12.72h.Mtb | D13.2h.Con | D13.2h.Mtb | D13.24h.Con | D13.24h.Mtb | D13.72h.Con | D13.72h.Mtb | D14.2h.Con | D14.2h.Mtb |
|---|---|---|---|---|---|---|---|---|---|---|---|---|---|---|
| 45996.4 | 127.5 | 125.6 | 23961.0 | 26454.5 | 47103.7 | 47121.6 | 84.1 | 46.1 | 12039.5 | 36943.2 | 29732.7 | 40790.0 | 95.8 | 117.3 |
| 4.2 | 0.2 | 0.2 | 4.0 | 11.9 | 3.6 | 11.5 | 0.2 | 0.5 | 2.8 | 62.4 | 1.0 | 23.8 | 0.0 | 0.1 |
| 0.3 | 0.2 | 0.2 | 0.3 | 0.5 | 0.2 | 0.3 | 0.1 | 0.1 | 0.1 | 1.5 | 0.1 | 1.7 | 0.0 | 0.0 |
| 1.2 | 0.1 | 0.0 | 0.2 | 0.2 | 0.8 | 0.9 | 0.1 | 0.1 | 0.5 | 0.8 | 1.3 | 3.2 | 0.1 | 0.1 |
| 149.8 | 5.2 | 4.1 | 44.2 | 41.7 | 67.4 | 95.5 | 3.1 | 3.2 | 29.0 | 39.7 | 49.3 | 103.3 | 4.8 | 3.7 |
| 95.0 | 36.2 | 34.8 | 99.3 | 112.0 | 105.9 | 150.9 | 23.3 | 21.5 | 87.1 | 165.5 | 88.6 | 262.8 | 16.1 | 17.2 |
| 116.1 | 0.6 | 0.6 | 7.9 | 33.1 | 42.8 | 274.2 | 0.2 | 0.3 | 4.2 | 54.0 | 33.9 | 271.0 | 1.1 | 0.3 |
| 6.3 | 0.4 | 0.4 | 1.7 | 7.7 | 2.7 | 31.0 | 0.0 | 0.1 | 0.4 | 30.3 | 0.6 | 36.2 | 0.0 | 0.0 |
| 394.5 | 22.8 | 22.1 | 138.1 | 237.7 | 191.0 | 347.6 | 5.0 | 7.1 | 72.6 | 1697.3 | 84.0 | 1494.1 | 0.7 | 2.4 |
| 0.1 | 0.0 | 0.0 | 0.1 | 0.1 | 0.1 | 0.2 | 0.1 | 0.1 | 0.1 | 0.1 | 0.2 | 0.2 | 0.1 | 0.1 |
| 92322.1 | 647.8 | 1181.9 | 43070.4 | 44416.7 | 95624.8 | 144766.6 | 627.0 | 650.1 | 68726.7 | 200329.4 | 94296.9 | 260515.3 | 0.0 | 0.0 |
| 136.8 | 0.3 | 0.4 | 2.5 | 6.7 | 7.3 | 208.8 | 0.8 | 0.8 | 8.9 | 105.9 | 31.7 | 4153.6 | 0.2 | 0.3 |
| 879.1 | 0.9 | 0.7 | 88.5 | 60.8 | 1221.9 | 1129.3 | 1.9 | 2.1 | 77.9 | 139.8 | 326.0 | 978.6 | 1.5 | 1.9 |
| 12.6 | 1.3 | 3.0 | 9.8 | 9.0 | 14.5 | 23.1 | 1.2 | 0.1 | 14.3 | 24.2 | 13.3 | 36.8 | 0.6 | 0.5 |
| 298.9 | 3.8 | 3.0 | 61.7 | 56.8 | 431.6 | 438.2 | 1.1 | 0.2 | 113.1 | 174.0 | 1009.1 | 3377.2 | 3.7 | 2.8 |
| 673.6 | 121.3 | 113.0 | 2283.6 | 3323.2 | 4256.9 | 7075.0 | 29.6 | 52.0 | 5666.9 | 6939.3 | 5216.6 | 7178.5 | 4.0 | 7.5 |
| 1051.2 | 26.4 | 24.3 | 848.6 | 991.8 | 1933.3 | 2847.7 | 14.3 | 25.7 | 5855.9 | 6181.6 | 5626.2 | 6325.8 | 5.0 | 8.0 |
| 241.3 | 7.0 | 8.8 | 347.3 | 447.3 | 565.8 | 857.5 | 3.6 | 1.8 | 163.6 | 696.7 | 330.7 | 1851.3 | 0.0 | 2.0 |
| 15.7 | 0.5 | 1.0 | 19.7 | 19.0 | 29.4 | 26.3 | 1.4 | 1.3 | 16.4 | 17.5 | 21.9 | 29.2 | 0.0 | 0.8 |
| 1973.1 | 0.0 | 2.8 | 1048.2 | 1353.5 | 2790.2 | 2818.9 | 30.1 | 34.0 | 3647.9 | 6779.3 | 3656.2 | 8239.3 | 3989.5 | 0.00000106277.181 |
| 1714.5 | 1.2 | 3.7 | 1271.5 | 1198.1 | 1650.1 | 1843.0 | 5.6 | 3.2 | 1811.1 | 4487.5 | 2566.9 | 6926.5 | 6.5 | 0.0 |
| 4707.3 | 0.0 | 3074.9 | 4578.4 | 4022.2 | 4177.2 | 4712.9 | 68.3 | 2261.7 | 4120.8 | 4227.8 | 5158.4 | 5893.3 | 384.1 | 6531.1 |
| 408.7 | 8.5 | 20.2 | 231.0 | 301.5 | 459.0 | 405.7 | 18.5 | 26.4 | 389.2 | 376.0 | 375.0 | 397.9 | 0.0 | 3.8 |
| 19280.3 | 0.0 | 0.0 | 1461.2 | 1879.5 | 14946.7 | 11965.0 | 0.0 | 0.0 | 0.0 | 0.0 | 65525.9 | 118101.4 | 0.0 | 0.0 |
| 7.8 | 0.6 | 0.8 | 4.4 | 3.0 | 17.3 | 20.6 | 0.2 | 0.2 | 2.7 | 4.9 | 9.9 | 51.5 | 0.9 | 0.8 |
| 193.1 | 68.9 | 56.5 | 145.4 | 181.3 | 63.7 | 127.9 | 49.8 | 53.1 | 328.1 | 1244.0 | 114.6 | 755.6 | 30.0 | 37.4 |
| 0.0 | 0.3 | 0.5 | 0.1 | 0.1 | 0.0 | 0.0 | 0.7 | 0.7 | 2.1 | 0.0 | 10.3 | 0.2 | 0.8 | 0.8 |

| D14.24h.Con | D14.24h.Mtb | D14.72h.Con | D14.72h.Mtb | D15.2h.Con | D15.2h.Mtb | D15.24h.Con | D15.24h.Mtb | D15.72h.Con | D15.72h.Mtb | D16.2h.Con | D16.2h.Mtb | D16.24h.Con | D16.24h.Mtb | D16.72h.Con |
|---|---|---|---|---|---|---|---|---|---|---|---|---|---|---|
| 36110.9 | 42422.5 | 41877.6 | 43883.1 | 47.3 | 62.8 | 11866.4 | 29727.5 | 24515.9 | 40147.2 | 77.2 | 87.4 | 21378.4 | 39938.3 | 40286.2 |
| 0.5 | 31.8 | 0.3 | 16.3 | 0.0 | 0.3 | 1.0 | 38.3 | 0.6 | 19.3 | 0.0 | 0.2 | 0.7 | 40.2 | 0.2 |
| 0.2 | 1.9 | 0.1 | 1.0 | 0.0 | 0.1 | 0.2 | 1.9 | 0.1 | 2.8 | 0.0 | 0.1 | 0.2 | 1.4 | 0.1 |
| 0.6 | 0.8 | 1.6 | 1.7 | 0.1 | 0.1 | 0.4 | 0.8 | 1.2 | 2.3 | 0.0 | 0.0 | 0.5 | 0.6 | 1.5 |
| 66.4 | 73.4 | 116.7 | 153.2 | 8.1 | 11.1 | 60.3 | 60.3 | 84.1 | 106.0 | 8.8 | 9.3 | 51.6 | 49.8 | 90.2 |
| 94.7 | 153.6 | 99.2 | 273.1 | 46.1 | 52.3 | 112.6 | 177.4 | 81.1 | 337.6 | 70.6 | 70.6 | 128.5 | 166.0 | 111.7 |
| 10.0 | 35.3 | 83.2 | 331.3 | 1.0 | 2.0 | 14.2 | 44.4 | 49.2 | 374.0 | 0.7 | 1.0 | 12.5 | 34.3 | 78.9 |
| 0.3 | 35.5 | 0.6 | 69.3 | 0.1 | 0.1 | 0.6 | 20.6 | 0.7 | 44.9 | 0.1 | 0.1 | 1.3 | 24.5 | 2.1 |
| 15.4 | 905.6 | 16.1 | 968.1 | 6.4 | 10.9 | 38.4 | 820.7 | 32.8 | 857.8 | 2.9 | 5.2 | 33.3 | 782.5 | 33.4 |
| 0.2 | 0.1 | 0.4 | 0.2 | 0.1 | 0.0 | 0.1 | 0.1 | 0.2 | 0.1 | 0.0 | 0.0 | 0.2 | 0.1 | 0.4 |
| 38832.4 | 166831.8 | 51314.6 | 218726.5 | 758.2 | 955.3 | 57591.6 | 170435.7 | 74089.9 | 239259.4 | 583.7 | 783.6 | 77179.3 | 200384.1 | 93450.4 |
| 2.3 | 10.4 | 4.5 | 287.7 | 1.5 | 1.6 | 21.2 | 927.8 | 36.0 | 18741.5 | 0.8 | 1.0 | 5.6 | 24.3 | 6.7 |
| 80.0 | 176.4 | 749.3 | 1090.5 | 1.4 | 1.7 | 128.2 | 266.2 | 617.6 | 2486.9 | 1.5 | 1.7 | 73.2 | 183.8 | 423.6 |
| 3.8 | 14.2 | 5.4 | 22.7 | 0.8 | 0.9 | 6.4 | 23.8 | 7.5 | 35.0 | 0.2 | 0.0 | 3.7 | 15.5 | 6.1 |
| 202.8 | 434.9 | 1106.9 | 3587.0 | 2.1 | 3.1 | 105.3 | 178.0 | 474.4 | 1620.3 | 2.4 | 3.0 | 127.5 | 188.3 | 824.6 |
| 814.8 | 6051.8 | 47.6 | 6467.7 | 15.6 | 37.8 | 2327.4 | 6712.1 | 282.2 | 6867.7 | 11.3 | 19.2 | 1489.4 | 6374.9 | 91.8 |
| 1751.1 | 4568.3 | 1112.9 | 5283.4 | 21.2 | 38.5 | 3812.9 | 6074.4 | 3304.1 | 6265.7 | 9.7 | 14.6 | 2877.5 | 5047.9 | 2373.3 |
| 138.2 | 824.8 | 177.8 | 1589.3 | 4.0 | 5.2 | 210.4 | 863.6 | 256.0 | 1873.1 | 2.7 | 3.7 | 115.1 | 571.6 | 187.2 |
| 21.2 | 24.9 | 31.7 | 35.8 | 0.2 | 1.0 | 16.8 | 21.8 | 22.7 | 51.5 | 0.3 | 1.3 | 16.0 | 21.0 | 29.4 |
| 3839.1 | 7943.4 | 19.1 | 5.2 | 1503.2 | 2948.2 | 1932.6 | 4238.4 | 31.8 | 30.3 | 6695.0 | 5901.4 | 6335.2 | 12419.2 | 75.8 |
| 2847.1 | 4252.5 | 2912.7 | 5236.1 | 0.0 | 0.0 | 930.1 | 1958.9 | 1473.6 | 3053.9 | 12.7 | 9.6 | 4629.8 | 5032.6 | 6678.2 |
| 3460.5 | 4104.9 | 4274.3 | 4723.4 | 0.0 | 2946.4 | 4811.6 | 5300.3 | 3582.1 | 3508.6 | 0.0 | 4232.8 | 3637.1 | 4143.0 | 4359.3 |
| 372.5 | 351.0 | 322.6 | 353.8 | 8.4 | 8.4 | 379.7 | 444.3 | 451.7 | 469.0 | 15.3 | 16.5 | 335.2 | 271.1 | 280.0 |
| 0.0 | 0.0 | 9744.6 | 29477.3 | 0.0 | 0.0 | 0.0 | 0.0 | 26691.2 | 61882.2 | 0.0 | 0.0 | 0.0 | 0.0 | 25278.3 |
| 2.7 | 9.0 | 9.3 | 107.7 | 0.4 | 0.5 | 3.3 | 8.2 | 8.4 | 66.5 | 0.1 | 0.2 | 1.2 | 3.3 | 6.7 |
| 393.0 | 1114.2 | 122.2 | 523.5 | 41.5 | 59.4 | 108.4 | 448.5 | 27.8 | 403.0 | 57.6 | 82.4 | 401.0 | 929.2 | 142.5 |
| 5.3 | 0.6 | 0.6 | 0.0 | 0.8 | 0.7 | 2.7 | 0.5 | 6.7 | 2.8 | 0.6 | 0.7 | 0.2 | 0.0 | 2.1 |

| D16.72h.Mtb | D17.2h.Con | D17.2h.Mtb | D17.24h.Con | D17.24h.Mtb | D17.72h.Con | D17.72h.Mtb | | D18.2h.Con | D18.2h.Mtb | D18.24h.Con | | D18.24h.Mtb | D18.72h.Con | D18.72h.Mtb |
|---|---|---|---|---|---|---|---|---|---|---|---|---|---|---|
| 43962.3 | 22.3 | 27.8 | 11658.7 | 39525.8 | 33769.1 | 43777.1 | | 36.6 | 39.5 | 21335.4 | | 36389.7 | 39618.9 | 42519.2 |
| 17.9 | 0.0 | 0.4 | 0.5 | 63.5 | 1.8 | 36.2 | | 0.0 | 0.1 | 1.2 | | 7.8 | 0.1 | 4.2 |
| 1.8 | 0.0 | 0.1 | 0.1 | 2.4 | 0.1 | 3.5 | | 0.0 | 0.0 | 0.2 | | 0.6 | 0.2 | 1.1 |
| 1.9 | 0.1 | 0.1 | 0.7 | 1.0 | 2.1 | 3.2 | | 0.1 | 0.0 | 0.6 | | 0.6 | 1.2 | 1.7 |
| 132.4 | 2.6 | 4.4 | 52.8 | 55.5 | 89.7 | 159.3 | | 2.9 | 3.7 | 55.8 | | 49.0 | 79.8 | 101.1 |
| 337.6 | 78.6 | 90.7 | 154.7 | 230.2 | 112.0 | 440.5 | | 30.1 | 37.6 | 95.9 | | 106.5 | 78.1 | 181.2 |
| 306.9 | 0.8 | 1.0 | 21.1 | 107.0 | 129.1 | 1133.8 | | 0.7 | 0.8 | 15.1 | | 30.5 | 86.2 | 248.0 |
| 48.9 | 0.1 | 0.2 | 1.0 | 64.6 | 1.7 | 149.7 | | 0.1 | 0.1 | 1.3 | | 10.9 | 1.8 | 21.9 |
| 859.5 | 2.2 | 4.9 | 29.2 | 1207.3 | 37.7 | 1568.6 | | 5.3 | 5.3 | 75.1 | | 242.3 | 71.2 | 307.2 |
| 0.2 | 0.1 | 0.0 | 0.2 | 0.1 | 0.5 | 0.2 | | 0.1 | 0.0 | 0.2 | | 0.1 | 0.2 | 0.2 |
| 247443.4 | 0.0 | 347.3 | 72528.7 | 231171.4 | 96478.3 | 304233.3 | | 411.7 | 523.6 | 74255.1 | | 122062.6 | 91655.1 | 175115.5 |
| 556.6 | 0.4 | 0.6 | 13.4 | 64.7 | 13.6 | 3009.6 | | 1.4 | 1.0 | 13.6 | | 86.7 | 14.3 | 523.4 |
| 1312.8 | 3.9 | 5.4 | 121.8 | 380.0 | 979.1 | 2216.4 | | 0.8 | 0.9 | 181.8 | | 294.7 | 871.0 | 1368.2 |
| 24.8 | 0.4 | 0.0 | 11.5 | 35.8 | 11.7 | 50.5 | | 0.0 | 1.0 | 7.2 | | 12.6 | 6.6 | 13.1 |
| 2573.4 | 2.7 | 3.5 | 390.5 | 588.3 | 1895.2 | 6257.0 | | 5.7 | 3.1 | 187.4 | | 183.8 | 863.9 | 2303.0 |
| 6614.1 | 8.6 | 22.8 | 3362.8 | 6896.8 | 1188.2 | 7154.6 | | 8.2 | 11.1 | 2385.2 | | 5103.2 | 806.2 | 6184.3 |
| 6081.9 | 8.0 | 19.4 | 5037.6 | 6079.8 | 4054.8 | 6229.8 | | 8.6 | 9.0 | 3417.3 | | 4287.6 | 2710.2 | 5276.7 |
| 1659.5 | 1.0 | 4.7 | 276.0 | 1825.3 | 441.3 | 6004.9 | | 3.3 | 4.4 | 380.9 | | 557.7 | 382.1 | 980.3 |
| 39.4 | 0.6 | 2.2 | 17.1 | 20.1 | 24.7 | 37.6 | | 0.3 | 0.8 | 14.8 | | 15.3 | 24.5 | 28.8 |
| 75.6 | 6453.6 | 13993.1 | 6915.7 | 17524.0 | 44.7 | 32.3 | | 8396.6 | 7316.1 | 7863.2 | | 10625.3 | | |
| 8116.4 | 6.6 | 1.3 | 3359.6 | 5493.7 | 4149.3 | 8389.7 | | 2.0 | 0.0 | 4336.2 | | 0.0 | 5017.1 | 6606.0 |
| 4606.7 | 0.0 | 3375.2 | 3480.0 | 4406.1 | 4503.8 | 4569.7 | 0.00000103523.5375 | | 2026.0 | 3953.7 | | 4157.0 | | |
| 421.9 | 53.0 | 58.2 | 324.8 | 426.7 | 298.1 | 365.3 | | 31.7 | 27.5 | 359.9 | | 304.4 | 281.0 | 297.1 |
| 67177.0 | 0.0 | 0.0 | 0.0 | 0.0 | 12346.8 | 35045.6 | | 0.0 | 0.0 | 27936.5 | 0.000001044914.46 | | | |
| 61.0 | 0.6 | 0.7 | 3.5 | 14.5 | 11.0 | 165.4 | | 0.9 | 0.9 | 5.4 | | 7.6 | 22.9 | 74.5 |
| 554.7 | 39.6 | 53.6 | 253.2 | 1008.3 | 85.7 | 724.7 | | 31.6 | 44.3 | 288.4 | | 354.1 | 73.6 | 176.4 |
| 0.2 | 0.7 | 0.8 | 1.4 | 0.2 | 12.2 | 11.0 | | 0.7 | 0.6 | 1.1 | | 0.1 | 52.5 | 6.8 |

| D19.2h.Con | D19.2h.Mtb | D19.24h.Con | D19.24h.Mtb | D19.72h.Con | D19.72h.Mtb | D20.2h.Con | D20.2h.Mtb | D20.24h.Con | D20.24h.Mtb | D20.72h.Con | D20.72h.Mtb | D21.2h.Con | D21.2h.Mtb | D21.24h.Con | D21.24h.Mtb |
|---|---|---|---|---|---|---|---|---|---|---|---|---|---|---|---|
| 37.5 | 38.7 | 9807.8 | 37372.0 | 36165.4 | 46349.8 | 253.8 | 289.3 | 40365.7 | 45556.0 | 42731.6 | 46360.8 | 45.5 | 61.0 | 21208.1 | 37855.4 |
| 0.1 | 0.6 | 0.7 | 103.6 | 0.5 | 62.5 | 0.2 | 2.2 | 3.0 | 165.1 | 1.7 | 210.5 | 0.1 | 0.5 | 1.3 | 26.7 |
| 0.1 | 0.1 | 0.4 | 2.5 | 0.2 | 1.0 | 0.1 | 0.5 | 0.5 | 6.7 | 0.5 | 23.5 | 0.1 | 0.2 | 0.3 | 1.9 |
| 0.1 | 0.1 | 0.3 | 0.4 | 0.8 | 1.0 | 0.1 | 0.1 | 0.8 | 1.9 | 2.0 | 5.7 | 0.1 | 0.1 | 0.4 | 0.6 |
| 13.7 | 12.5 | 119.7 | 115.0 | 203.3 | 239.6 | 9.1 | 10.5 | 100.7 | 118.8 | 141.9 | 295.8 | 14.9 | 14.1 | 102.8 | 105.0 |
| 68.0 | 67.9 | 123.0 | 159.2 | 89.5 | 311.2 | 38.5 | 51.7 | 119.6 | 342.2 | 114.9 | 2910.8 | 27.2 | 28.4 | 75.0 | 117.2 |
| 1.3 | 3.5 | 20.6 | 51.3 | 143.8 | 567.7 | 1.6 | 2.2 | 18.0 | 305.5 | 54.6 | 2992.9 | 1.0 | 0.7 | 14.6 | 49.3 |
| 0.1 | 0.1 | 0.9 | 30.8 | 1.5 | 75.8 | 0.1 | 0.3 | 0.8 | 147.9 | 0.8 | 575.1 | 0.0 | 0.1 | 0.4 | 22.1 |
| 3.5 | 8.1 | 35.9 | 1142.6 | 40.0 | 1093.2 | 7.5 | 25.5 | 49.7 | 2071.7 | 57.7 | 2326.2 | 1.9 | 4.4 | 20.0 | 364.0 |
| 0.0 | 0.0 | 0.1 | 0.2 | 0.3 | 0.2 | 0.0 | 0.1 | 0.1 | 0.2 | 0.1 | 0.4 | 0.0 | 0.0 | 0.1 | 0.1 |
| 109.5 | 177.5 | 27147.2 | 108951.1 | 37983.3 | 141931.1 | 471.1 | 703.5 | 68647.7 | 206548.5 | 69140.2 | 247860.0 | 126.5 | 232.1 | 20428.9 | 68622.7 |
| 0.2 | 0.2 | 1.8 | 0.7 | 2.5 | 72.3 | 1.9 | 2.1 | 45.4 | 0.0 | 168.5 | 0.0 | 0.7 | 0.8 | 18.6 | 424.2 |
| 5.7 | 6.3 | 335.4 | 685.9 | 2245.2 | 4616.4 | 25.8 | 29.3 | 1241.8 | 4708.1 | 4100.9 | 13730.9 | 10.5 | 13.3 | 654.8 | 1265.4 |
| 0.6 | 0.9 | 4.3 | 8.3 | 7.4 | 22.8 | 1.5 | 0.0 | 16.3 | 53.4 | 12.9 | 62.3 | 1.1 | 0.9 | 4.9 | 12.7 |
| 3.6 | 1.9 | 193.6 | 390.6 | 1891.6 | 7742.8 | 10.1 | 12.5 | 872.8 | 1314.9 | 4666.9 | 15348.2 | 6.4 | 7.8 | 651.9 | 659.6 |
| 7.2 | 16.9 | 1045.0 | 5624.2 | 964.6 | 7067.9 | 21.6 | 93.3 | 2675.7 | 7873.8 | 590.1 | 7877.0 | 1.8 | 12.4 | 1314.4 | 5598.0 |
| 12.1 | 24.7 | 11166.2 | 0.0 | 37470.5 | 0.0 | 79.0 | 234.1 | 0.0 | 0.0 | 0.0 | 0.0 | 7.2 | 27.8 | 0.0 | 0.0 |
| 0.2 | 4.1 | 175.1 | 829.5 | 242.3 | 1943.1 | 5.4 | 8.9 | 417.7 | 3833.5 | 620.1 | 12592.3 | 0.0 | 0.0 | 220.0 | 504.7 |
| 0.0 | 0.0 | 17.3 | 19.0 | 19.6 | 38.8 | 0.0 | 0.4 | 33.7 | 79.8 | 66.8 | 172.6 | 0.0 | 0.0 | 23.3 | 25.0 |
| 0.0 | 0.0 | 772.7 | 1462.9 | 978.8 | 3880.8 | 0.4 | 4.2 | 661.2 | 1857.0 | 474.3 | 1631.2 | 0.0 | 0.0 | 3274.8 | 3631.1 |
| 1.2 | 5.0 | 2272.4 | 3024.7 | 2533.5 | 3966.1 | 0.0 | 1.2 | 842.7 | 2367.5 | 896.1 | 3145.3 | 1.8 | 0.0 | 1771.2 | 2172.8 |
| 0.0 | 3270.1 | 4044.0 | 4383.8 | 4938.7 | 3961.3 | 0.0 | 1871.6 | 2435.7 | 2420.8 | 3292.9 | 2909.5 | 1.1 | 1805.5 | 3349.1 | 2458.7 |
| 0.0 | 0.0 | 772.7 | 1462.9 | 978.8 | 3880.8 | 0.4 | 4.2 | 661.2 | 1857.0 | 474.3 | 1631.2 | 0.0 | 0.0 | 3274.8 | 3631.1 |
| 0.0 | 0.0 | 0.0 | 356.7 | 2153.1 | 11936.5 | 0.0 | 26.7 | 2577.7 | 4628.7 | 7786.4 | 16588.4 | 0.0 | 0.0 | 2676.0 | 2044.3 |
| 3.8 | 3.8 | 13.3 | 40.5 | 85.4 | 643.9 | 8.2 | 7.9 | 66.6 | 212.9 | 164.3 | 1191.6 | 7.5 | 8.8 | 30.4 | 73.0 |
| 62.8 | 67.7 | 315.4 | 963.6 | 112.0 | 488.7 | 47.9 | 103.4 | 84.5 | 1824.7 | 39.0 | 4017.6 | 34.8 | 52.1 | 205.0 | 422.3 |
| 0.5 | 0.5 | 1.1 | 0.0 | 9.3 | 0.0 | 1.2 | 1.8 | 0.8 | 0.0 | 0.4 | 93.7 | 1.0 | 1.2 | 11.4 | 3.1 |

| D21.72h.Con | D21.72h.Mtb | D22.2h.Con | D22.2h.Mtb | D22.24h.Con | D22.24h.Mtb | D22.72h.Con | D22.72h.Mtb | D23.2h.Con | D23.2h.Mtb | D23.24h.Con | D23.24h.Mtb | D23.72h.Con | D23.72h.Mtb | D24.2h.Con | D24.2h.Mtb |
|---|---|---|---|---|---|---|---|---|---|---|---|---|---|---|---|
| 42544.3 | 46281.3 | 94.6 | 99.1 | 23574.6 | 44958.0 | 44667.4 | 48694.0 | 63.7 | 70.5 | 25627.4 | 44325.4 | 44314.2 | 48438.3 | 188.5 | 125.4 |
| 0.8 | 21.0 | 0.1 | 0.2 | 0.9 | 51.6 | 0.4 | 18.7 | 0.1 | 0.3 | 0.9 | 24.3 | 22.0 | 13.8 | 0.1 | 1.5 |
| 0.2 | 1.8 | 0.1 | 0.1 | 1.1 | 1.8 | 0.2 | 3.0 | 0.1 | 0.1 | 0.2 | 0.8 | 0.9 | 1.6 | 0.1 | 0.4 |
| 1.0 | 1.6 | 0.1 | 0.1 | 0.5 | 0.6 | 1.5 | 1.8 | 0.1 | 0.1 | 0.8 | 1.7 | 1.6 | 2.9 | 0.1 | 0.1 |
| 153.9 | 185.9 | 23.4 | 20.3 | 91.8 | 90.0 | 158.7 | 236.8 | 5.3 | 3.1 | 77.6 | 95.0 | 90.8 | 162.7 | 21.6 | 21.8 |
| 78.1 | 181.2 | 70.8 | 69.3 | 190.1 | 265.6 | 156.1 | 465.9 | 72.8 | 64.0 | 188.7 | 244.3 | 184.9 | 436.8 | 51.3 | 59.9 |
| 103.9 | 394.1 | 1.4 | 1.7 | 18.6 | 63.5 | 173.3 | 643.3 | 0.7 | 0.5 | 23.8 | 107.3 | 97.8 | 573.2 | 1.7 | 2.4 |
| 0.6 | 36.0 | 0.1 | 0.1 | 3.3 | 40.2 | 5.9 | 104.4 | 0.1 | 0.1 | 1.5 | 8.8 | 7.0 | 25.8 | 0.2 | 0.3 |
| 19.1 | 384.3 | 4.1 | 5.4 | 66.4 | 1033.6 | 67.3 | 1257.1 | 2.9 | 5.4 | 45.6 | 459.4 | 426.8 | 941.0 | 11.4 | 27.9 |
| 0.2 | 0.2 | 0.0 | 0.1 | 0.1 | 0.1 | 0.3 | 0.4 | 0.0 | 0.0 | 0.1 | 0.4 | 0.4 | 0.4 | 0.0 | 0.1 |
| 24035.6 | 99976.7 | 351.3 | 363.5 | 95025.6 | 157950.1 | 109525.9 | 165920.6 | 336.1 | 326.0 | 78245.4 | 140776.9 | 132084.3 | 196814.9 | 673.6 | 731.2 |
| 81.4 | 1258.2 | 2.1 | 1.9 | 13.7 | 12.2 | 22.0 | 144.3 | 0.6 | 0.5 | 13.6 | 24.0 | 15.1 | 198.6 | 1.8 | 1.9 |
| 3971.3 | 7450.9 | 13.1 | 13.5 | 444.9 | 1115.7 | 1478.9 | 3522.1 | 19.6 | 18.5 | 419.1 | 1071.5 | 915.6 | 1638.8 | 10.6 | 10.5 |
| 6.5 | 20.9 | 0.6 | 0.1 | 10.7 | 20.0 | 12.0 | 35.8 | 1.5 | 1.1 | 17.8 | 17.6 | 23.3 | 49.3 | 1.1 | 0.8 |
| 2544.7 | 7965.2 | 6.2 | 4.0 | 474.4 | 541.1 | 3152.7 | 6669.6 | 8.2 | 11.2 | 685.3 | 2510.3 | 2153.0 | 11098.2 | 5.2 | 5.9 |
| 68.8 | 5878.0 | 11.5 | 16.3 | 3251.3 | 6972.6 | 2127.3 | 7448.5 | 10.2 | 16.3 | 3618.7 | 4483.8 | 4262.6 | 7555.5 | 8.6 | 26.4 |
| 10202.6 | 0.0 | 23.8 | 27.2 | 0.0 | 0.0 | 0.0 | 0.0 | 21.7 | 32.4 | 0.0 | 0.0 | 0.0 | 0.0 | 25.8 | 67.6 |
| 269.1 | 1149.8 | 2.9 | 4.4 | 583.1 | 1345.1 | 858.3 | 3763.8 | 4.4 | 3.1 | 506.1 | 1969.0 | 948.0 | 4735.6 | 5.1 | 6.2 |
| 37.1 | 60.0 | 0.0 | 0.0 | 23.3 | 30.2 | 47.1 | 55.8 | 0.0 | 0.4 | 25.5 | 30.8 | 51.3 | 58.4 | 0.0 | 0.0 |
| 2799.1 | 3213.6 | 0.0 | 0.0 | 18545.9 | 22281.7 | 29022.1 | 39422.9 | 4.2 | 11.6 | 2971.7 | 1650.2 | 1504.6 | 3111.1 | 0.0 | 0.0 |
| 2756.3 | 3711.6 | 10.4 | 8.1 | 19717.2 | 18823.0 | 26402.8 | 28555.9 | 2.7 | 0.5 | 2442.9 | 2980.6 | 2977.0 | 4092.7 | 6.8 | 3.3 |
| 3620.0 | 3200.1 | 0.0 | 2286.2 | 2855.5 | 3390.1 | 3525.9 | 3282.6 | 0.0 | 746.7 | 2735.3 | 3448.0 | 3546.5 | 3282.8 | 0.0 | 1833.6 |
| 2799.1 | 3213.6 | 0.0 | 0.0 | 18545.9 | 22281.7 | 29022.1 | 39422.9 | 4.2 | 11.6 | 2971.7 | 1650.2 | 1504.6 | 3111.1 | 0.0 | 0.0 |
| 9776.5 | 11943.0 | 0.0 | 0.0 | 3719.3 | 5644.9 | 31002.4 | 54904.8 | 184.1 | 0.0 | 2771.5 | 1949.4 | 7390.8 | 23426.9 | 99.3 | 184.1 |
| 133.9 | 846.9 | 1.1 | 1.7 | 33.2 | 52.8 | 213.7 | 520.6 | 11.6 | 10.8 | 84.2 | 149.1 | 131.9 | 679.0 | 1.5 | 2.5 |
| 57.1 | 233.5 | 77.9 | 82.0 | 767.0 | 1188.1 | 246.8 | 557.4 | 29.2 | 58.8 | 261.4 | 304.3 | 272.5 | 323.1 | 61.4 | 149.5 |
| 30.7 | 10.4 | 0.7 | 0.8 | 0.9 | 0.1 | 42.7 | 28.9 | 0.7 | 0.6 | 2.5 | 15.8 | 14.1 | 14.8 | 1.9 | 1.9 |

| D24.24h.Con | D24.24h.Mtb | D24.72h.Con | D24.72h.Mtb | D25.2h.Con | D25.2h.Mtb | D25.24h.Con | D25.24h.Mtb | D25.72h.Con | D25.72h.Mtb | D26.2h.Con | D26.2h.Mtb | D26.24h.Con | D26.24h.Mtb | D26.72h.Con | D26.72h.Mtb |
|---|---|---|---|---|---|---|---|---|---|---|---|---|---|---|---|
| 37858.1 | 46646.7 | 45944.3 | 48057.0 | 47.6 | 68.2 | 14845.5 | 37043.4 | 37256.1 | 43069.9 | 133.5 | 148.1 | 38313.0 | 40966.2 | 40650.6 | 43806.6 |
| 1.1 | 65.2 | 1.2 | 20.1 | 0.0 | 0.5 | 0.4 | 30.9 | 0.2 | 24.2 | 0.1 | 0.0 | 2.4 | 16.5 | 0.6 | 12.7 |
| 0.0 | 0.2 | 0.2 | 1.2 | 0.1 | 0.2 | 0.2 | 1.4 | 0.0 | 1.0 | 0.1 | 0.1 | 0.4 | 2.0 | 0.1 | 4.5 |
| 0.6 | 0.9 | 2.0 | 3.4 | 0.2 | 0.2 | 0.6 | 0.8 | 1.6 | 2.6 | 0.1 | 0.1 | 0.9 | 2.0 | 2.8 | 6.5 |
| 68.2 | 72.5 | 114.9 | 183.0 | 9.4 | 8.3 | 40.8 | 36.1 | 62.8 | 94.5 | 4.3 | 4.1 | 34.4 | 37.7 | 42.8 | 99.4 |
| 135.4 | 202.3 | 152.8 | 517.1 | 17.5 | 19.8 | 58.1 | 80.7 | 54.4 | 177.7 | 13.1 | 14.2 | 74.1 | 129.9 | 79.6 | 380.4 |
| 12.6 | 49.8 | 80.6 | 309.6 | 0.6 | 1.5 | 8.7 | 32.5 | 46.1 | 265.8 | 0.4 | 0.2 | 6.9 | 36.4 | 23.4 | 198.2 |
| 0.3 | 0.4 | 2.2 | 43.8 | 0.0 | 0.1 | 0.3 | 19.1 | 0.4 | 32.8 | 0.0 | 0.0 | 0.8 | 21.2 | 0.8 | 39.6 |
| 0.1 | 1.1 | 72.9 | 1609.6 | 1.1 | 6.2 | 21.1 | 928.4 | 21.6 | 1077.2 | 7.3 | 6.5 | 23.9 | 233.7 | 30.6 | 322.0 |
| 0.1 | 0.2 | 0.2 | 0.2 | 0.0 | 0.1 | 0.1 | 0.1 | 0.3 | 0.2 | 0.1 | 0.1 | 0.1 | 0.1 | 0.4 | 0.3 |
| 62329.8 | 140132.5 | 83783.2 | 175998.0 | 396.5 | 288.1 | 22800.8 | 113032.8 | 30041.4 | 199190.1 | 1513.1 | 1444.2 | 64305.5 | 200470.0 | 83121.1 | 227343.7 |
| 20.8 | 112.5 | 26.4 | 561.4 | 0.6 | 0.7 | 5.8 | 88.8 | 5.7 | 884.1 | 1.2 | 1.0 | 32.2 | 2345.7 | 80.6 | 9415.2 |
| 318.5 | 698.1 | 1509.7 | 3315.1 | 1.2 | 1.3 | 28.4 | 77.4 | 285.6 | 812.8 | 1.4 | 1.3 | 166.5 | 676.0 | 584.1 | 2284.6 |
| 13.6 | 28.0 | 8.5 | 29.1 | 0.0 | 0.0 | 7.9 | 27.3 | 9.7 | 45.6 | 0.0 | 0.5 | 13.2 | 45.9 | 13.4 | 42.1 |
| 298.3 | 454.2 | 2520.5 | 22226.6 | 1.4 | 0.7 | 45.9 | 98.5 | 232.7 | 865.4 | 1.9 | 0.9 | 105.6 | 155.7 | 502.7 | 4660.3 |
| 3311.7 | 7412.8 | 305.9 | 7360.8 | 9.3 | 27.6 | 1206.8 | 6149.9 | 78.8 | 6305.8 | 10.7 | 7.0 | 1269.6 | 6416.2 | 72.7 | 6368.0 |
| 0.0 | 0.0 | 0.0 | 0.0 | 6.4 | 21.0 | 1725.7 | 4586.9 | 1387.6 | 5384.6 | 15.2 | 11.5 | 4632.9 | 5505.8 | 2637.3 | 5474.5 |
| 609.7 | 1764.4 | 1118.9 | 3825.9 | 3.8 | 1.7 | 79.5 | 515.9 | 149.0 | 1651.5 | 5.6 | 2.8 | 176.3 | 578.8 | 237.2 | 1767.3 |
| 31.9 | 40.4 | 51.1 | 61.8 | 0.1 | 0.0 | 17.6 | 19.2 | 25.7 | 39.3 | 0.0 | 0.1 | 26.6 | 25.7 | 40.9 | 68.2 |
| 1253.1 | 3077.0 | 2026.1 | 3926.0 | 27.6 | 10.6 | 1970.1 | 1637.0 | 3139.7 | 5083.1 | 14.9 | 9.4 | 6327.3 | 8211.3 | 4096.1 | 7737.3 |
| 2387.0 | 4235.8 | 2461.1 | 5416.2 | 0.0 | 0.0 | 845.3 | 1234.3 | 1169.5 | 1943.0 | 5.5 | 5.1 | 3563.7 | 5389.5 | 3550.4 | 7466.2 |
| 3373.2 | 2694.0 | 3595.9 | 3702.5 | 0.0 | 4377.1 | 3443.6 | 3545.5 | 3822.8 | 3950.0 | 2686.4 | 14.7 | 2725.3 | 2505.3 | 2786.0 | 3691.0 |
| 1253.1 | 3077.0 | 2026.1 | 3926.0 | 6.2 | 3.1 | 271.6 | 182.7 | 364.3 | 398.7 | 9.6 | 4.0 | 403.6 | 292.8 | 251.8 | 269.0 |
| 2599.6 | 4077.2 | 29026.6 | 68014.2 | 399.3 | 200.4 | 1950.3 | 1632.5 | 12836.3 | 31222.0 | 219.8 |  | 7141.9 | 6006.7 | 16587.7 | 43393.9 |
| 27.1 | 48.3 | 67.6 | 241.8 | 0.3 | 0.5 | 1.0 | 2.8 | 4.6 | 31.3 | 0.4 | 0.5 | 4.3 | 9.4 | 7.2 | 49.5 |
| 1.8 | 6.3 | 81.6 | 402.7 | 21.9 | 51.4 | 211.6 | 769.1 | 77.5 | 518.7 | 92.5 | 87.8 | 115.4 | 491.8 | 44.3 | 556.2 |
| 3.7 | 0.2 | 37.9 | 1.2 | 0.8 | 0.9 | 5.3 | 1.5 | 11.9 | 0.8 | 2.4 | 3.0 | 19.7 | 1.9 | 43.3 | 54.4 |

| D27.2h.Con | D27.2h.Mtb | D27.24h.Con | D27.24h.Mtb | D27.72h.Con | D27.72h.Mtb | D28.2h.Con | D28.2h.Mtb | D28.24h.Con | D28.24h.Mtb | D28.72h.Con | D28.72h.Mtb |
|---|---|---|---|---|---|---|---|---|---|---|---|
| 145.0 | 203.3 | 41329.5 | 43336.8 | 41862.8 | 45063.8 | 414.6 | 397.6 | 40572.6 | 41281.1 | 42686.5 | 43484.6 |
| 0.0 | 0.7 | 0.3 | 32.1 | 0.1 | 21.1 | 3.0 | 5.1 | 22.3 | 99.6 | 19.1 | 146.7 |
| 0.1 | 0.3 | 0.1 | 1.9 | 0.0 | 1.7 | 0.6 | 0.8 | 2.1 | 6.5 | 2.1 | 16.9 |
| 0.1 | 0.1 | 0.8 | 1.1 | 2.5 | 3.4 | 0.1 | 0.1 | 1.0 | 1.9 | 2.4 | 5.2 |
| 6.1 | 6.8 | 23.6 | 42.4 | 41.3 | 144.9 | 20.2 | 12.6 | 42.9 | 49.3 | 63.9 | 124.3 |
| 29.2 | 38.1 | 101.2 | 222.1 | 105.3 | 666.2 | 84.3 | 83.5 | 183.2 | 269.8 | 214.8 | 778.6 |
| 0.5 | 1.1 | 8.0 | 92.4 | 40.9 | 631.3 | 5.5 | 4.7 | 25.9 | 104.3 | 99.4 | 727.2 |
| 0.0 | 0.2 | 0.2 | 43.9 | 0.2 | 91.1 | 1.3 | 2.4 | 5.6 | 67.8 | 5.1 | 170.1 |
| 14.5 | 30.7 | 48.4 | 1078.5 | 39.3 | 1424.9 | 78.9 | 85.4 | 552.6 | 1051.9 | 490.6 | 1496.9 |
| 0.0 | 0.0 | 0.1 | 0.1 | 0.5 | 0.1 | 0.0 | 0.1 | 0.0 | 0.1 | 0.1 | 0.2 |
| 1216.9 | 1568.7 | 36168.5 | 165458.3 | 35255.0 | 237620.8 | 4878.0 | 4524.4 | 203371.6 | 259661.4 | 201462.5 | 323519.4 |
| 1.2 | 1.2 | 16.7 | 183.4 | 19.9 | 1836.8 | 4.2 | 3.6 | 244.5 | 6878.0 | 439.4 | |
| 2.4 | 3.0 | 64.6 | 127.7 | 208.2 | 693.0 | 15.5 | 16.4 | 436.6 | 1014.4 | 826.6 | 4778.9 |
| 0.0 | 1.4 | 10.3 | 24.5 | 8.8 | 39.6 | 4.0 | 4.0 | 49.7 | 58.9 | 37.6 | 88.9 |
| 0.2 | 1.8 | 87.9 | 208.3 | 305.4 | 2190.7 | 8.3 | 8.1 | 1074.2 | 952.7 | 14692.6 | 18761.2 |
| 4.0 | 24.1 | 393.6 | 5874.3 | 25.9 | 6138.7 | 290.8 | 316.8 | 6650.6 | 6815.9 | 5511.8 | 6919.5 |
| 12.5 | 30.0 | 5329.5 | 5493.1 | 4213.8 | 5490.2 | 152.2 | 172.3 | 5429.5 | 5517.0 | 4798.2 | 5526.5 |
| 7.5 | 8.1 | 324.4 | 2701.7 | 413.2 | 7989.0 | 39.1 | 40.3 | 1579.6 | 2149.2 | 1670.4 | 6731.8 |
| 0.1 | 0.0 | 36.0 | 44.5 | 52.0 | 78.9 | 0.0 | 0.5 | 41.5 | 46.9 | 172.5 | 239.3 |
| 11.8 | 9.4 | 1832.6 | 8488.1 | 3713.6 | 13049.7 | 23.5 | 19.8 | 5317.5 | 7033.6 | 4630.2 | 9204.2 |
| 4.4 | 2.9 | 2758.9 | 5413.9 | 3717.4 | 11631.7 | 44.6 | 36.0 | 3931.8 | 5043.5 | 5510.1 | 9256.6 |
| 37.7 | 3707.6 | 2889.5 | 2854.8 | 3529.0 | 3010.1 | 0.0 | 1672.3 | 2993.2 | 2751.3 | 2616.8 | 2611.9 |
| 0.4 | 0.9 | 201.7 | 373.9 | 414.2 | 413.0 | 1.8 | 2.2 | 324.7 | 309.4 | 332.8 | 460.4 |
| 322.1 | | 1031.0 | 2876.3 | 7999.1 | 22487.7 | 556.6 | 289.5 | 24149.8 | 18005.9 | 82533.9 | 83563.7 |
| 0.1 | 0.1 | 2.2 | 5.8 | 4.2 | 61.0 | 1.2 | 1.1 | 35.1 | 38.7 | 178.6 | 278.4 |
| 115.8 | 158.1 | 102.8 | 516.1 | 36.1 | 424.3 | 302.8 | 270.8 | 462.6 | 1090.7 | 188.7 | 1724.0 |
| 3.4 | 3.5 | 36.1 | 15.1 | 77.8 | 161.7 | 1.5 | 1.4 | 0.5 | 0.1 | 3.0 | 68.0 |
