## Supplemental Table 3 for "Human alveolar macrophage response to *Mycobacterium tuberculosis*: immune characteristics underlying large inter-individual variability"

**Supplementary File 3. Correlations between secreted proteins and M.tb generation time during the 48 to 72h incubation period (Protein vs. Growth Rate correlations)**

| | 2 Hours | | 24 Hours | | 72 Hours | |
|---|---|---|---|---|---|---|
| **Proteins** | **Correlation** | **P value** | **Correlation** | **P value** | **Correlation** | **P value** |
| **CCL13** | -0.269798907 | 0.164997643 | 0.116607257 | 0.554575598 | -0.038042855 | 0.847589272 |
| **CCL17** | -0.189736519 | 0.333524203 | -0.13315372 | 0.499372514 | -0.118479635 | 0.548190947 |
| **CCL19** | -0.162249431 | 0.409435731 | 0.014510437 | 0.94157931 | 0.090675281 | 0.64632014 |
| **CCL2** | -0.125763967 | 0.523682421 | -0.088708655 | 0.653512603 | -0.179502829 | 0.360722617 |
| **CCL20** | -0.011598741 | 0.95328794 | 0.099528036 | 0.614334604 | 0.083095613 | 0.674206354 |
| **CCL22** | 0.14745636 | 0.453978085 | -0.150759632 | 0.443815193 | 0.008475681 | 0.965857001 |
| **CCL3** | -0.091289125 | 0.644081468 | -0.351802057 | 0.066375664 | -0.180088968 | 0.359130416 |
| **CCL4** | -0.097536894 | 0.621471487 | -0.172685176 | 0.379546837 | 0.03989901 | 0.840245085 |
| **CSF2** | 0.022541329 | 0.909353867 | 0.047170389 | 0.811605944 | 0.075949646 | 0.700889194 |
| **CXCL10** | -0.14337731 | 0.466696169 | -0.258870666 | 0.18346251 | 0.171539689 | 0.39225414 |
| **CXCL5** | 0.174117717 | 0.375545053 | -0.137346456 | 0.485835291 | -0.142071993 | 0.47080493 |
| **IL10** | **0.464049148** | **0.012865159** | **0.393425808** | **0.038333683** | 0.252052778 | 0.195686116 |
| **IL15** | 0.184458395 | 0.347392628 | 0.1873198 | 0.339831925 | 0.253422209 | 0.19318697 |
| **IL16** | -0.128356344 | 0.515089812 | -0.201869748 | 0.302940866 | -0.190212411 | 0.332290538 |
| **IL18** | -0.056433721 | 0.775466605 | 0.094547856 | 0.632248173 | 0.106604149 | 0.589251949 |
| **IL1A** | -0.065155879 | 0.741850468 | 0.072624308 | 0.713427637 | -0.043272443 | 0.826931387 |
| **IL1B** | 0.331607713 | 0.084732403 | 0.190272179 | 0.332135798 | 0.085465254 | 0.665440752 |
| **IL6** | 0.188968813 | 0.335520191 | -0.000521199 | 0.997899826 | 0.09767182 | 0.620986802 |
| **IL7** | **0.57089279** | **0.001510547** | 0.330189256 | 0.086153793 | **-0.413428132** | **0.028756169** |
| **IL8** | -0.011749721 | 0.952680557 | -0.329045128 | 0.087313428 | -0.194322469 | 0.321751674 |
| **MMP1** | -0.201048307 | 0.304954139 | -0.167362614 | 0.394629986 | 0.003287128 | 0.987017244 |
| **MMP10** | 0.264383945 | 0.173975026 | **0.38989224** | **0.040264065** | -0.354437259 | 0.064227859 |
| **MMP2** | -0.19246302 | 0.326493896 | -0.271595113 | 0.162093559 | -0.277203961 | 0.161559406 |
| **MMP3** | -0.098166397 | 0.619211484 | -0.14146752 | 0.472713978 | -0.277501114 | 0.152800973 |
| **MMP9** | 0.253810327 | 0.210881697 | 0.140123432 | 0.485739551 | 0.081141386 | 0.687436088 |
| **TNF** | 0.104608285 | 0.596281575 | 0.040772982 | 0.836791558 | 0.030800875 | 0.876357181 |
| **VEGFA** | 0.304927897 | 0.114610458 | 0.12249261 | 0.534623327 | 0.205530419 | 0.294070293 |
