## Supplemental Table 4 for "Human alveolar macrophage response to *Mycobacterium tuberculosis*: immune characteristics underlying large inter-individual variability"

**Supplementary File 4. Differential expression of RNAs control versus M.tb infected AMs in all 28 donor AMs at 2h, 24h and 72h (Suppl. Tables 4A, B, and C), using separate non-infected control AMs for each time point**

**Suppl. File 4A: 2h**

| Gene | baseMean | log2FoldChange | lfcSE | stat | pvalue | padj |
|---|---|---|---|---|---|---|
| CSF2 | 78.61617599 | 3.377558739 | 0.308459944 | 10.94974827 | 6.66E-28 | 1.30E-23 |
| IL1B | 4561.340615 | 1.281010646 | 0.153130296 | 8.365494475 | 5.99E-17 | 3.91E-13 |
| IL6 | 1754.819037 | 1.766649188 | 0.210162717 | 8.406101778 | 4.24E-17 | 3.91E-13 |
| NFKBIZ | 10891.28273 | 0.865357358 | 0.10727297 | 8.066872382 | 7.21E-16 | 3.53E-12 |
| CCL3L3 | 554.2828804 | 1.21567708 | 0.153330598 | 7.928470208 | 2.22E-15 | 7.49E-12 |
| IER3 | 6173.41637 | 0.796628111 | 0.100531 | 7.9242036 | 2.30E-15 | 7.49E-12 |
| NFKBIA | 13099.77254 | 0.64427635 | 0.081623541 | 7.89326638 | 2.94E-15 | 8.24E-12 |
| CCL4 | 2174.05987 | 1.000850512 | 0.127588905 | 7.844338146 | 4.35E-15 | 1.07E-11 |
| PDE4B | 435.3264346 | 0.492166183 | 0.063434937 | 7.758598131 | 8.59E-15 | 1.87E-11 |
| SERPINB2 | 1158.791876 | 1.094106826 | 0.148989668 | 7.34350804 | 2.08E-13 | 4.07E-10 |
| CCL20 | 5657.404964 | 1.37738543 | 0.189266971 | 7.277473829 | 3.40E-13 | 6.06E-10 |
| CXCL1 | 293.5867136 | 1.690508385 | 0.23375014 | 7.232117107 | 4.76E-13 | 7.76E-10 |
| IL10 | 22.55436007 | 1.577335561 | 0.219243116 | 7.194458767 | 6.27E-13 | 9.45E-10 |
| TNFAIP8 | 571.8161196 | 0.615477644 | 0.086047654 | 7.152753359 | 8.51E-13 | 1.19E-09 |
| F3 | 194.8818206 | 1.014514315 | 0.142365133 | 7.126143135 | 1.03E-12 | 1.35E-09 |
| OR2B11 | 34.6220274 | 2.1849835 | 0.322617045 | 6.772684613 | 1.26E-11 | 1.55E-08 |
| OR5B21 | 220.0761845 | 0.676761855 | 0.1083418 | 6.246544315 | 4.20E-10 | 4.83E-07 |
| SOD2 | 20257.21645 | 0.345262748 | 0.056938651 | 6.063767619 | 1.33E-09 | 1.45E-06 |
| CXCL3 | 10829.49481 | 0.658200014 | 0.110076971 | 5.979452474 | 2.24E-09 | 2.31E-06 |
| TM4SF19 | 300.8473395 | -0.383283467 | 0.064325892 | -5.95846332 | 2.55E-09 | 2.49E-06 |
| NLRP3 | 99.18160534 | 1.20908037 | 0.212117121 | 5.700060255 | 1.20E-08 | 1.07E-05 |
| TNFAIP6 | 2211.433212 | 0.735287611 | 0.128969 | 5.70127406 | 1.19E-08 | 1.07E-05 |
| IFNG | 45.53170261 | 2.104536181 | 0.370249148 | 5.684108101 | 1.31E-08 | 1.12E-05 |
| CXCL2 | 6680.827698 | 1.038297782 | 0.184862224 | 5.616603316 | 1.95E-08 | 1.59E-05 |
| CCL3 | 3259.076378 | 0.781638751 | 0.141903382 | 5.508246115 | 3.62E-08 | 2.84E-05 |
| EREG | 1782.578432 | 1.039826355 | 0.19095602 | 5.445370893 | 5.17E-08 | 3.89E-05 |
| LINC00346 | 34.59202536 | 1.39651881 | 0.2617739 | 5.334828299 | 9.56E-08 | 6.94E-05 |
| G0S2 | 348.7209299 | 0.507174313 | 0.096322613 | 5.265371201 | 1.40E-07 | 9.78E-05 |
| KANK1 | 61.27531895 | 0.782343146 | 0.148876721 | 5.254972971 | 1.48E-07 | 1.00E-04 |
| PTX3 | 177.8767351 | 1.079788713 | 0.209980067 | 5.142339121 | 2.71E-07 | 0.000177121 |
| SPRY2 | 54.02424806 | -0.693740819 | 0.135486925 | -5.12035254 | 3.05E-07 | 0.000192649 |
| NBN | 669.883533 | 0.275081652 | 0.054010898 | 5.093076836 | 3.52E-07 | 0.000215596 |
| AGPAT9 | 293.6508332 | -0.416272788 | 0.081858303 | -5.08528486 | 3.67E-07 | 0.000217831 |
| ARL4C | 40.35486559 | -0.702707291 | 0.140786154 | -4.99130966 | 6.00E-07 | 0.000335548 |
| NINJ1 | 744.8114449 | 0.48311793 | 0.096723777 | 4.99482076 | 5.89E-07 | 0.000335548 |
| ZFX | 932.3222701 | 0.345148756 | 0.069228779 | 4.985625361 | 6.18E-07 | 0.000335967 |
| LYPD3 | 51.41533624 | 0.814078085 | 0.166209296 | 4.897909475 | 9.69E-07 | 0.00051266 |
| ADORA2A | 15.29732884 | 1.414681982 | 0.29324733 | 4.82419391 | 1.41E-06 | 0.00072442 |
| CCL15 | 18.06098791 | 1.671640377 | 0.349566206 | 4.782042289 | 1.74E-06 | 0.000871309 |
| NFKB2 | 1024.184741 | 0.361816224 | 0.075780392 | 4.774536178 | 1.80E-06 | 0.000881832 |
| AMPD3 | 77.79196104 | 0.6586112 | 0.139929028 | 4.706751752 | 2.52E-06 | 0.001202182 |
| GPR84 | 127.1423227 | 0.862070236 | 0.186342449 | 4.62626868 | 3.72E-06 | 0.00173595 |

| | | | | | | |
|---|---|---|---|---|---|---|
| JUN | 177.3666207 | 0.633162995 | 0.139386356 | 4.542503399 | 5.56E-06 | 0.002531677 |
| CXCL5 | 3228.808804 | 0.509871764 | 0.113426279 | 4.495181978 | 6.95E-06 | 0.00309369 |
| TNIP3 | 141.7185127 | 0.626646907 | 0.141886976 | 4.416521693 | 1.00E-05 | 0.004364913 |
| LIF | 134.431019 | 0.740554649 | 0.168128479 | 4.404694874 | 1.06E-05 | 0.004509747 |
| BIRC3 | 214.6478081 | 0.418423922 | 0.095474825 | 4.382557613 | 1.17E-05 | 0.004785356 |
| ZNF267 | 2984.265591 | 0.193882486 | 0.044224976 | 4.384004312 | 1.17E-05 | 0.004785356 |
| RASGRP3 | 41.90781682 | 0.80968604 | 0.186573927 | 4.339759862 | 1.43E-05 | 0.005700591 |
| KBTBD7 | 379.9258208 | 0.450743159 | 0.106120603 | 4.24746133 | 2.16E-05 | 0.008467943 |
| DOLK | 383.7466123 | 0.361777903 | 0.086605426 | 4.177312191 | 2.95E-05 | 0.011326425 |
| CCL22 | 352.2815844 | 0.451345075 | 0.109202588 | 4.133098712 | 3.58E-05 | 0.013478562 |
| LINC00515 | 723.6119851 | 0.495586769 | 0.120731247 | 4.104875756 | 4.05E-05 | 0.014947085 |
| CCR7 | 115.8091965 | 0.592005762 | 0.14853992 | 3.985499406 | 6.73E-05 | 0.024420126 |
| TNF | 11775.1251 | 0.300547417 | 0.076012312 | 3.953930732 | 7.69E-05 | 0.027372645 |
| IL1A | 2331.071554 | 0.495326282 | 0.125422711 | 3.949255103 | 7.84E-05 | 0.027414371 |
| PTGS2 | 5050.844177 | 0.309261181 | 0.078746274 | 3.927311934 | 8.59E-05 | 0.029512111 |
| NARF | 355.9792843 | -0.215836774 | 0.056236717 | -3.8380045 | 0.000124038 | 0.041880005 |
| TNFAIP3 | 1883.219639 | 0.727427951 | 0.19023876 | 3.823763102 | 0.00013143 | 0.043623689 |
| MARCKS | 107.9410563 | 0.699532428 | 0.183303321 | 3.816256163 | 0.000135492 | 0.044222279 |
| SLC2A6 | 312.7695863 | 0.297786305 | 0.078309852 | 3.802667166 | 0.000143147 | 0.045954756 |
| PLCXD1 | 137.7320976 | -0.354190488 | 0.093421477 | -3.79131758 | 0.00014985 | 0.047330908 |

**Supplementary File 4. Differential expression of RNAs control versus M.tb infected AMs in all 28 donor AMs at 2h, 24h and 72h (Suppl. Tables 4A, B, and C), using separate non-infected control AMs for each time point**

**Suppl. File 4B: 24h**

| Gene | baseMean | log2FoldChange | lfcSE | stat | pvalue | padj |
|---|---|---|---|---|---|---|
| INHBA | 4383.210884 | 2.305522311 | 0.140710317 | 16.38488465 | 2.45E-60 | 4.72E-56 |
| CCL3 | 5420.860212 | 1.549634818 | 0.098486329 | 15.73451705 | 8.77E-56 | 8.45E-52 |
| DCUN1D3 | 444.2160304 | 1.889492315 | 0.124646074 | 15.15885942 | 6.62E-52 | 4.25E-48 |
| SERPINB7 | 41.94863524 | 6.893715809 | 0.472819075 | 14.58002898 | 3.76E-48 | 1.81E-44 |
| IL1B | 15588.33476 | 2.288415746 | 0.157562824 | 14.52383048 | 8.56E-48 | 3.30E-44 |
| SERPINB2 | 10295.23192 | 3.146243555 | 0.217442547 | 14.46930968 | 1.89E-47 | 6.08E-44 |
| SCARF1 | 264.1116703 | 1.453547521 | 0.101956717 | 14.25651553 | 4.08E-46 | 1.12E-42 |
| CXCL3 | 1451.493281 | 1.622249558 | 0.117504581 | 13.80584093 | 2.35E-43 | 5.66E-40 |
| PPBP | 616.6665382 | 2.87816522 | 0.214310493 | 13.42988476 | 4.04E-41 | 8.65E-38 |
| CISH | 549.5478519 | 2.078644205 | 0.156076173 | 13.31813921 | 1.82E-40 | 3.50E-37 |
| CCL20 | 5070.755564 | 2.064866118 | 0.160444555 | 12.86965534 | 6.67E-38 | 1.17E-34 |
| EMP1 | 1796.72616 | 1.143433374 | 0.089992707 | 12.70584488 | 5.49E-37 | 8.81E-34 |
| ETS2 | 2133.574722 | 1.532595271 | 0.121474627 | 12.61658762 | 1.71E-36 | 2.54E-33 |
| SPRED1 | 967.6293434 | 1.521603749 | 0.121749361 | 12.49783768 | 7.67E-36 | 1.06E-32 |
| CCL3L3 | 260.3487733 | 2.138993191 | 0.172295668 | 12.41466615 | 2.18E-35 | 2.79E-32 |
| DPP4 | 137.3578149 | 2.985202916 | 0.241450083 | 12.36364418 | 4.11E-35 | 4.95E-32 |
| TLR2 | 1209.477317 | 1.517797548 | 0.12364074 | 12.27586912 | 1.22E-34 | 1.38E-31 |
| CXCL1 | 568.4383038 | 2.876177896 | 0.23562222 | 12.20673456 | 2.86E-34 | 3.06E-31 |
| IL6 | 96.03517463 | 3.734194443 | 0.306670189 | 12.17658115 | 4.14E-34 | 4.20E-31 |
| TMEM173 | 154.1522994 | 1.555625732 | 0.128810792 | 12.07682763 | 1.40E-33 | 1.35E-30 |
| MPEG1 | 291.6887392 | -1.242023373 | 0.10340441 | -12.01131923 | 3.10E-33 | 2.84E-30 |
| MS4A4A | 2213.279997 | -0.807965147 | 0.068364581 | -11.81847587 | 3.13E-32 | 2.74E-29 |
| TFPI2 | 366.847363 | 1.767377375 | 0.150077879 | 11.7764016 | 5.17E-32 | 4.33E-29 |
| CD274 | 1512.259315 | 1.039206836 | 0.088285311 | 11.77100497 | 5.51E-32 | 4.42E-29 |
| IL3RA | 402.4177024 | 1.858620529 | 0.15798544 | 11.76450517 | 5.95E-32 | 4.58E-29 |
| IDO1 | 1335.405852 | 3.862043365 | 0.331016914 | 11.66720855 | 1.87E-31 | 1.39E-28 |
| IL2RA | 76.77759614 | 3.34058494 | 0.286580762 | 11.65669642 | 2.12E-31 | 1.51E-28 |
| TNFAIP6 | 9552.540813 | 1.429510538 | 0.125118617 | 11.42524252 | 3.13E-30 | 2.15E-27 |
| SERPINE1 | 2570.914449 | 1.294873066 | 0.113584942 | 11.40004159 | 4.18E-30 | 2.78E-27 |
| IL7R | 5906.277628 | 1.403965146 | 0.123248522 | 11.39133457 | 4.62E-30 | 2.97E-27 |
| CCL4 | 9491.79488 | 1.25884123 | 0.111040491 | 11.33677646 | 8.63E-30 | 5.36E-27 |
| CXCL5 | 28416.86762 | 1.573049155 | 0.139418183 | 11.28295551 | 1.59E-29 | 9.59E-27 |
| CSF1 | 18649.64658 | 1.255211145 | 0.111990398 | 11.20820326 | 3.72E-29 | 2.17E-26 |
| CKB | 222.5548241 | 1.65846753 | 0.148858566 | 11.14123004 | 7.90E-29 | 4.48E-26 |
| TNF | 695.6472635 | 1.422489838 | 0.127754976 | 11.13451612 | 8.52E-29 | 4.69E-26 |
| ECSCR | 41.34211868 | 2.43428834 | 0.22367212 | 10.88328909 | 1.38E-27 | 7.41E-25 |
| FAM198B | 246.2650758 | -1.44300179 | 0.132995059 | -10.85004063 | 1.99E-27 | 1.04E-24 |
| NUPR1 | 3950.990398 | -1.410508719 | 0.131100354 | -10.75900008 | 5.38E-27 | 2.73E-24 |
| HSD11B1 | 456.9548496 | 1.674841493 | 0.156441209 | 10.70588436 | 9.55E-27 | 4.72E-24 |
| EEPD1 | 149.8600465 | -1.209777846 | 0.113110718 | -10.69551915 | 1.07E-26 | 5.14E-24 |
| PTGS2 | 882.4982746 | 1.714135245 | 0.16040048 | 10.68659674 | 1.18E-26 | 5.53E-24 |
| CSF2 | 40.2425273 | 4.801327111 | 0.449657032 | 10.67775386 | 1.29E-26 | 5.93E-24 |

| | | | | | | |
|---|---|---|---|---|---|---|
| ADORA2A | 51.04020773 | 2.303186601 | 0.215980508 | 10.66386323 | 1.50E-26 | 6.73E-24 |
| SCD | 4969.07607 | -0.650348234 | 0.061831633 | -10.51805047 | 7.13E-26 | 3.12E-23 |
| FAM124A | 62.08833886 | 2.300946865 | 0.219471631 | 10.48402863 | 1.02E-25 | 4.38E-23 |
| APOL4 | 179.3351866 | 1.971371481 | 0.189908651 | 10.38063021 | 3.04E-25 | 1.27E-22 |
| ABI3BP | 23.50386505 | 4.468801653 | 0.430883614 | 10.37124994 | 3.35E-25 | 1.37E-22 |
| KLHDC8B | 296.6513651 | -1.05775744 | 0.102589576 | -10.31057425 | 6.31E-25 | 2.53E-22 |
| CCL5 | 481.818689 | 1.611667319 | 0.157058885 | 10.26154818 | 1.05E-24 | 4.13E-22 |
| SLC2A3 | 1113.228612 | 1.126078782 | 0.109891345 | 10.24720174 | 1.22E-24 | 4.69E-22 |
| CD40 | 2377.879133 | 0.920809926 | 0.089984309 | 10.23300545 | 1.41E-24 | 5.28E-22 |
| ITGB3 | 661.3370218 | 1.528264981 | 0.149360245 | 10.23207336 | 1.42E-24 | 5.28E-22 |
| CYP27B1 | 297.2631181 | 2.255516207 | 0.220586297 | 10.22509666 | 1.53E-24 | 5.46E-22 |
| SLC7A2 | 35.4190351 | 3.373782648 | 0.32991313 | 10.22627577 | 1.51E-24 | 5.46E-22 |
| HCAR2 | 237.9792568 | 1.404968918 | 0.137844548 | 10.19241559 | 2.14E-24 | 7.51E-22 |
| PVALB | 80.99773061 | -1.539612374 | 0.151448906 | -10.16588642 | 2.82E-24 | 9.69E-22 |
| YPEL3 | 214.2203861 | -0.86238677 | 0.085195132 | -10.12248878 | 4.39E-24 | 1.48E-21 |
| IER3 | 2658.14964 | 1.030096079 | 0.102327009 | 10.06670758 | 7.75E-24 | 2.58E-21 |
| HCAR3 | 258.2970321 | 1.315036505 | 0.131125935 | 10.02880555 | 1.14E-23 | 3.72E-21 |
| FABP3 | 1344.004403 | -0.605600943 | 0.060925666 | -9.939997034 | 2.79E-23 | 8.95E-21 |
| CFP | 406.8055419 | -1.126690019 | 0.113554683 | -9.922004032 | 3.34E-23 | 1.04E-20 |
| SOCS3 | 142.1813865 | 1.42096971 | 0.143221334 | 9.921494705 | 3.36E-23 | 1.04E-20 |
| EDN1 | 440.0068244 | 1.63594468 | 0.165092516 | 9.909260096 | 3.79E-23 | 1.16E-20 |
| STON1 | 36.58323936 | 3.229256886 | 0.326252657 | 9.898024787 | 4.25E-23 | 1.28E-20 |
| VWA8 | 1566.48698 | 0.99255747 | 0.10030772 | 9.895125448 | 4.37E-23 | 1.30E-20 |
| RHCG | 72.02146111 | 1.774821915 | 0.179438231 | 9.890990946 | 4.55E-23 | 1.31E-20 |
| TRIB2 | 89.32916368 | 1.259785084 | 0.12735012 | 9.892296012 | 4.50E-23 | 1.31E-20 |
| SGPP2 | 56.20202721 | 2.007397521 | 0.203190593 | 9.87938216 | 5.11E-23 | 1.45E-20 |
| OCSTAMP | 96.82827258 | 2.061680435 | 0.20900044 | 9.864478916 | 5.93E-23 | 1.66E-20 |
| BMF | 185.4140474 | -0.958176809 | 0.097175129 | -9.860309152 | 6.19E-23 | 1.70E-20 |
| NFAT5 | 965.1782704 | 0.608037275 | 0.062119858 | 9.788130418 | 1.27E-22 | 3.44E-20 |
| CCL2 | 1725.676214 | 1.505114582 | 0.154357898 | 9.750810321 | 1.83E-22 | 4.90E-20 |
| RSPO3 | 186.0699737 | 2.107955624 | 0.216350603 | 9.743238968 | 1.97E-22 | 5.20E-20 |
| VSIG4 | 316.6768179 | -0.771935025 | 0.079298263 | -9.734576722 | 2.15E-22 | 5.59E-20 |
| CCL24 | 5936.662623 | 1.96069994 | 0.201503898 | 9.730332563 | 2.24E-22 | 5.75E-20 |
| FAM134B | 602.7622563 | -0.593736298 | 0.061213683 | -9.699404921 | 3.03E-22 | 7.69E-20 |
| IL36G | 21.79811789 | 3.68646998 | 0.380428505 | 9.690309556 | 3.32E-22 | 8.29E-20 |
| APBB2 | 396.3324651 | 1.07199517 | 0.111102645 | 9.648691733 | 4.98E-22 | 1.23E-19 |
| TGFA | 103.1491956 | 1.682377581 | 0.174468635 | 9.642865498 | 5.27E-22 | 1.29E-19 |
| IL8 | 73445.91326 | 1.373652784 | 0.142506773 | 9.639210481 | 5.46E-22 | 1.32E-19 |
| ARHGDIB | 7362.648567 | -0.612168131 | 0.063829364 | -9.590697718 | 8.75E-22 | 2.08E-19 |
| ALDH2 | 4415.862918 | -0.925731527 | 0.096624204 | -9.580741591 | 9.64E-22 | 2.26E-19 |
| UGDH | 355.1393097 | 0.601222407 | 0.062857336 | 9.564872559 | 1.12E-21 | 2.61E-19 |
| MRPS6 | 3506.715262 | 0.930377904 | 0.097903934 | 9.5029675 | 2.04E-21 | 4.68E-19 |
| MXI1 | 227.4106578 | -0.768157256 | 0.08118069 | -9.462314946 | 3.01E-21 | 6.83E-19 |
| MRC1 | 6152.855244 | 1.131985242 | 0.119784724 | 9.450163619 | 3.38E-21 | 7.58E-19 |
| CYP3A5 | 62.78415224 | 2.647865513 | 0.280505624 | 9.439616486 | 3.74E-21 | 8.29E-19 |
| PIM1 | 3406.275997 | 0.869892052 | 0.092575589 | 9.396559741 | 5.64E-21 | 1.23E-18 |
| LAMB3 | 98.568742 | 1.023441797 | 0.109982135 | 9.305527598 | 1.33E-20 | 2.89E-18 |

| | | | | | | |
|---|---|---|---|---|---|---|
| EREG | 517.1627249 | 1.638413775 | 0.176293303 | 9.293681282 | 1.49E-20 | 3.19E-18 |
| IL1A | 1681.836055 | 1.87851716 | 0.202199666 | 9.290406833 | 1.54E-20 | 3.25E-18 |
| KCNN4 | 1436.957974 | 0.82983376 | 0.089454793 | 9.276571203 | 1.75E-20 | 3.67E-18 |
| DUSP5 | 699.5476242 | 1.408019153 | 0.15231009 | 9.244424635 | 2.37E-20 | 4.90E-18 |
| C10orf128 | 586.1789223 | 0.968693433 | 0.105076773 | 9.218911162 | 3.00E-20 | 6.15E-18 |
| KDM6B | 1179.749715 | 0.730404598 | 0.080188398 | 9.108606906 | 8.34E-20 | 1.69E-17 |
| BHLHE40 | 2838.693937 | 0.824664891 | 0.09057045 | 9.105231216 | 8.61E-20 | 1.73E-17 |
| PIM3 | 2224.574305 | 0.489375083 | 0.053769752 | 9.101308167 | 8.93E-20 | 1.77E-17 |
| USF2 | 1528.3062 | -0.532153879 | 0.058646877 | -9.073865527 | 1.15E-19 | 2.26E-17 |
| APOC1 | 6841.869399 | -0.726707882 | 0.080112702 | -9.071069481 | 1.18E-19 | 2.29E-17 |
| ABTB2 | 69.30293268 | 1.010481359 | 0.111438477 | 9.067616392 | 1.22E-19 | 2.34E-17 |
| PPARD | 1342.650688 | 1.015641111 | 0.112122031 | 9.058354572 | 1.32E-19 | 2.53E-17 |
| MOB3B | 432.1815419 | 1.310743511 | 0.145832239 | 8.988022929 | 2.52E-19 | 4.75E-17 |
| MIR155HG | 161.0476688 | 1.276116376 | 0.143001431 | 8.923801436 | 4.51E-19 | 8.43E-17 |
| RUNX1 | 152.5779312 | 1.319402054 | 0.14790289 | 8.92073204 | 4.63E-19 | 8.58E-17 |
| MMP9 | 7830.13448 | 0.865835307 | 0.097383481 | 8.890987413 | 6.06E-19 | 1.11E-16 |
| GPNMB | 30408.40748 | -0.602630699 | 0.067846523 | -8.882263588 | 6.55E-19 | 1.18E-16 |
| HBEGF | 5126.974826 | 1.007873756 | 0.113458757 | 8.883172867 | 6.50E-19 | 1.18E-16 |
| CA12 | 103.1450885 | 1.452171208 | 0.164220668 | 8.842804194 | 9.33E-19 | 1.63E-16 |
| FAM129B | 3422.847187 | 0.714873021 | 0.080835638 | 8.843537768 | 9.27E-19 | 1.63E-16 |
| STAC | 641.7424579 | 1.028394608 | 0.116274439 | 8.844545838 | 9.19E-19 | 1.63E-16 |
| IL1RN | 30576.76402 | 0.973223129 | 0.110072167 | 8.841682308 | 9.43E-19 | 1.64E-16 |
| TNFRSF21 | 404.6832251 | -0.965864525 | 0.109262023 | -8.83989235 | 9.58E-19 | 1.65E-16 |
| APOC2 | 1711.42036 | -0.443753017 | 0.050213646 | -8.837299304 | 9.81E-19 | 1.67E-16 |
| CXCL2 | 765.1642031 | 1.5154421 | 0.172389104 | 8.790822987 | 1.48E-18 | 2.51E-16 |
| VDR | 948.9860568 | 1.127177201 | 0.128387697 | 8.779479855 | 1.64E-18 | 2.75E-16 |
| TREM2 | 655.7098908 | -0.914983729 | 0.10435021 | -8.768393785 | 1.81E-18 | 3.01E-16 |
| MSC | 1006.985327 | 0.683525889 | 0.078312593 | 8.72817336 | 2.59E-18 | 4.26E-16 |
| MMP1 | 403.1166381 | 1.353100235 | 0.155702346 | 8.690300883 | 3.61E-18 | 5.90E-16 |
| BAALC | 43.24587492 | 1.513036823 | 0.174168025 | 8.687225027 | 3.71E-18 | 6.01E-16 |
| TM4SF1 | 383.2404289 | 1.469368765 | 0.169990037 | 8.643852237 | 5.43E-18 | 8.73E-16 |
| TSLP | 27.40605291 | 2.444979819 | 0.284257905 | 8.601272923 | 7.88E-18 | 1.26E-15 |
| BTBD2 | 340.9104111 | -0.557055607 | 0.065026553 | -8.566586765 | 1.07E-17 | 1.68E-15 |
| LIF | 51.74317231 | 1.641818177 | 0.191787429 | 8.560614138 | 1.12E-17 | 1.76E-15 |
| FLT1 | 27.61470657 | 1.5365463 | 0.179618032 | 8.554521424 | 1.18E-17 | 1.84E-15 |
| DNASE2 | 1613.78923 | -0.780171522 | 0.091479179 | -8.528405386 | 1.48E-17 | 2.29E-15 |
| KIAA1462 | 115.1270937 | 1.220980041 | 0.143222135 | 8.525079151 | 1.53E-17 | 2.33E-15 |
| ISG20 | 63.99541402 | 2.004901284 | 0.235213627 | 8.523746321 | 1.54E-17 | 2.34E-15 |
| ELOVL7 | 486.0316433 | 0.875799656 | 0.103345118 | 8.474514082 | 2.36E-17 | 3.55E-15 |
| LSMD1 | 186.1069456 | -0.738754438 | 0.087221496 | -8.46986663 | 2.46E-17 | 3.67E-15 |
| CD226 | 185.2410131 | 1.258571588 | 0.148641094 | 8.467184649 | 2.51E-17 | 3.70E-15 |
| FUCA1 | 704.2317209 | -1.192004755 | 0.14077677 | -8.46733985 | 2.51E-17 | 3.70E-15 |
| NNMT | 86.43607432 | 2.321711819 | 0.274690251 | 8.452108544 | 2.86E-17 | 4.18E-15 |
| IGSF6 | 2420.41175 | 0.869280613 | 0.102964162 | 8.442555091 | 3.10E-17 | 4.50E-15 |
| PTPRF | 134.835711 | 1.857741136 | 0.220126991 | 8.439406391 | 3.19E-17 | 4.59E-15 |
| ADCK3 | 191.091426 | -0.727215231 | 0.086214103 | -8.43499161 | 3.31E-17 | 4.73E-15 |
| AMDHD2 | 344.8693566 | -0.571806487 | 0.068043833 | -8.403502016 | 4.33E-17 | 6.14E-15 |

| | | | | | | |
|---|---|---|---|---|---|---|
| RAB42 | 391.6695282 | -0.76050639 | 0.090872363 | -8.368951447 | 5.81E-17 | 8.18E-15 |
| GBP1P1 | 40.63224392 | 3.504945295 | 0.420293872 | 8.3392729 | 7.47E-17 | 1.03E-14 |
| IFI44L | 142.3235524 | 2.264205989 | 0.27153463 | 8.338553319 | 7.52E-17 | 1.03E-14 |
| ZNF589 | 374.1434581 | -0.564570771 | 0.067701235 | -8.339150196 | 7.48E-17 | 1.03E-14 |
| GLIPR2 | 2013.608808 | 0.900225408 | 0.108107046 | 8.327166824 | 8.28E-17 | 1.13E-14 |
| CASS4 | 187.3488037 | 1.216694658 | 0.146140465 | 8.325515174 | 8.40E-17 | 1.14E-14 |
| LDLRAP1 | 369.4280499 | 0.628240849 | 0.075748137 | 8.293812515 | 1.10E-16 | 1.48E-14 |
| ANKRD22 | 142.3760432 | 2.300348566 | 0.277712956 | 8.283187783 | 1.20E-16 | 1.60E-14 |
| PTX3 | 64.66898195 | 1.428395986 | 0.17253883 | 8.278692895 | 1.25E-16 | 1.65E-14 |
| ABTB1 | 549.6440272 | -0.640199691 | 0.077362957 | -8.275274359 | 1.28E-16 | 1.69E-14 |
| SLC7A11 | 4405.675767 | 0.878707082 | 0.106324707 | 8.264373397 | 1.40E-16 | 1.84E-14 |
| F3 | 158.873557 | 1.138511988 | 0.137948389 | 8.253173496 | 1.54E-16 | 2.01E-14 |
| DPEP2 | 452.3537472 | -0.719250878 | 0.08716624 | -8.251484513 | 1.56E-16 | 2.02E-14 |
| HLX | 341.015413 | 0.844944349 | 0.102414802 | 8.25021709 | 1.58E-16 | 2.03E-14 |
| KCNA3 | 454.9022324 | 0.916856373 | 0.111226224 | 8.243167293 | 1.68E-16 | 2.14E-14 |
| PARP12 | 876.2908751 | 0.396919438 | 0.048393418 | 8.201930314 | 2.37E-16 | 2.98E-14 |
| SAMSN1 | 395.7386876 | 0.766038511 | 0.093395048 | 8.202131949 | 2.36E-16 | 2.98E-14 |
| NEURL3 | 33.12430547 | 1.790193145 | 0.218539856 | 8.191609415 | 2.58E-16 | 3.22E-14 |
| SLC5A3 | 1238.335323 | 1.293208597 | 0.158266332 | 8.171090987 | 3.06E-16 | 3.80E-14 |
| C19orf59 | 3569.118224 | 1.08688508 | 0.133122305 | 8.16456026 | 3.23E-16 | 3.98E-14 |
| APOL6 | 754.0408314 | 0.731547887 | 0.089671512 | 8.158085799 | 3.40E-16 | 4.18E-14 |
| SLAMF6 | 66.58798935 | -1.099014283 | 0.134817913 | -8.151841703 | 3.58E-16 | 4.37E-14 |
| TRIM25 | 1296.470512 | 0.689930155 | 0.084703491 | 8.145238749 | 3.79E-16 | 4.59E-14 |
| GPR84 | 240.7160625 | 0.961413744 | 0.118345339 | 8.123798972 | 4.52E-16 | 5.44E-14 |
| CDKN1A | 5998.363369 | 0.746722517 | 0.092332709 | 8.087302165 | 6.10E-16 | 7.30E-14 |
| TNIP3 | 1723.128296 | 0.671065081 | 0.083098754 | 8.075513121 | 6.72E-16 | 7.99E-14 |
| PLAU | 1321.776097 | 0.788972513 | 0.097810106 | 8.066370101 | 7.24E-16 | 8.56E-14 |
| SORL1 | 111.4978758 | -1.609952852 | 0.19970977 | -8.061462655 | 7.54E-16 | 8.86E-14 |
| BTN2A1 | 817.3406476 | 0.514529172 | 0.064165594 | 8.018770491 | 1.07E-15 | 1.25E-13 |
| SMAD7 | 806.9068596 | 0.610762381 | 0.076319291 | 8.002726104 | 1.22E-15 | 1.41E-13 |
| MB21D2 | 166.1704759 | 0.772640157 | 0.096820021 | 7.980169279 | 1.46E-15 | 1.69E-13 |
| CCL1 | 10.01244276 | 3.960066835 | 0.497651172 | 7.957515337 | 1.76E-15 | 2.01E-13 |
| CHST7 | 1051.658933 | -0.543001741 | 0.068292106 | -7.951164092 | 1.85E-15 | 2.11E-13 |
| CYP27A1 | 23302.04811 | -0.346861195 | 0.044034143 | -7.877096461 | 3.35E-15 | 3.80E-13 |
| DAPK1 | 4273.411614 | -0.449232214 | 0.057095353 | -7.868104735 | 3.60E-15 | 4.06E-13 |
| APOL1 | 1033.40213 | 1.149635015 | 0.146301391 | 7.857991012 | 3.90E-15 | 4.37E-13 |
| CXCR4 | 503.6992894 | -1.182535255 | 0.150559021 | -7.854296933 | 4.02E-15 | 4.45E-13 |
| EGR2 | 6430.274089 | 0.856909916 | 0.10909453 | 7.854746827 | 4.01E-15 | 4.45E-13 |
| PKIA | 148.0856301 | 0.784779994 | 0.100081086 | 7.841441669 | 4.45E-15 | 4.88E-13 |
| RALGDS | 849.7774338 | 0.585879086 | 0.074714751 | 7.841545097 | 4.45E-15 | 4.88E-13 |
| HMOX1 | 4725.324884 | -0.411035858 | 0.052428488 | -7.839933404 | 4.51E-15 | 4.91E-13 |
| BHLHE41 | 2918.623049 | -0.724718497 | 0.092463083 | -7.837922725 | 4.58E-15 | 4.94E-13 |
| STARD8 | 271.9404132 | 0.655727371 | 0.083663937 | 7.837634665 | 4.59E-15 | 4.94E-13 |
| CSF3 | 8.192933408 | 3.800688902 | 0.486128024 | 7.818288007 | 5.35E-15 | 5.73E-13 |
| CYP3A7 | 14.53148398 | 2.743879302 | 0.351414576 | 7.808097591 | 5.81E-15 | 6.18E-13 |
| TSC22D1 | 1781.016296 | 0.59197167 | 0.075850213 | 7.804482671 | 5.97E-15 | 6.32E-13 |
| CD36 | 1400.985254 | -0.513474059 | 0.06592173 | -7.789147211 | 6.75E-15 | 7.10E-13 |

| | | | | | | |
|---|---|---|---|---|---|---|
| UBD | 655.4178825 | 2.354054524 | 0.302553806 | 7.780614492 | 7.22E-15 | 7.56E-13 |
| NFKBIZ | 1244.027095 | 0.515985508 | 0.066327364 | 7.779376086 | 7.29E-15 | 7.59E-13 |
| CSRNP1 | 136.9476971 | 0.915586937 | 0.118033562 | 7.757005069 | 8.70E-15 | 9.01E-13 |
| SLC28A3 | 224.550812 | 0.983089407 | 0.126752852 | 7.755954944 | 8.77E-15 | 9.03E-13 |
| CCR7 | 102.9616604 | 1.000337997 | 0.129232889 | 7.740583745 | 9.90E-15 | 1.01E-12 |
| TMEM45B | 59.25551738 | 1.586326775 | 0.20565773 | 7.713431322 | 1.22E-14 | 1.25E-12 |
| AMPD3 | 824.9104126 | 0.536457066 | 0.069755783 | 7.690503052 | 1.47E-14 | 1.49E-12 |
| SLAMF1 | 16.19907967 | 2.719368805 | 0.35407478 | 7.680210387 | 1.59E-14 | 1.60E-12 |
| PIK3IP1 | 62.58763328 | -1.075555797 | 0.140081494 | -7.67807202 | 1.62E-14 | 1.62E-12 |
| PLSCR1 | 1964.594083 | 0.525031121 | 0.068473818 | 7.66761856 | 1.75E-14 | 1.75E-12 |
| NT5E | 51.45348143 | 1.225110899 | 0.160115363 | 7.651426301 | 1.99E-14 | 1.97E-12 |
| WNT5A | 57.16615254 | 1.561503195 | 0.204685481 | 7.628793135 | 2.37E-14 | 2.34E-12 |
| VNN2 | 99.28632709 | -0.710027981 | 0.093106804 | -7.625951614 | 2.42E-14 | 2.38E-12 |
| LDLR | 285.8708468 | 0.857430436 | 0.112629848 | 7.612817104 | 2.68E-14 | 2.62E-12 |
| CYB5R3 | 90.0290541 | 0.974141559 | 0.128056981 | 7.607094519 | 2.80E-14 | 2.73E-12 |
| SLC22A1 | 33.52470948 | 1.854861456 | 0.244292872 | 7.592777632 | 3.13E-14 | 3.03E-12 |
| BCL6 | 1233.692227 | 0.59749604 | 0.078705923 | 7.591500329 | 3.16E-14 | 3.05E-12 |
| TTYH2 | 302.8376628 | -1.141926239 | 0.150502673 | -7.587415019 | 3.26E-14 | 3.13E-12 |
| CCL17 | 27.36880212 | 2.26670547 | 0.298816333 | 7.585614372 | 3.31E-14 | 3.16E-12 |
| MATK | 741.944356 | 0.66654044 | 0.088103135 | 7.565456534 | 3.87E-14 | 3.67E-12 |
| TMEM86A | 305.0255181 | -0.669600011 | 0.088622753 | -7.555621915 | 4.17E-14 | 3.94E-12 |
| FERMT2 | 45.70415138 | 1.730873417 | 0.229329971 | 7.547523809 | 4.44E-14 | 4.17E-12 |
| EAF1 | 1169.94604 | 0.510176751 | 0.067711794 | 7.534533095 | 4.90E-14 | 4.58E-12 |
| ENPP4 | 2106.23619 | 0.821542394 | 0.109057262 | 7.533128736 | 4.95E-14 | 4.60E-12 |
| SERINC5 | 1576.556457 | -0.367732449 | 0.048818277 | -7.532679818 | 4.97E-14 | 4.60E-12 |
| ARRDC4 | 1200.461809 | 0.793980163 | 0.105444024 | 7.529873483 | 5.08E-14 | 4.68E-12 |
| N4BP1 | 533.0340521 | 0.754950807 | 0.100605072 | 7.504102805 | 6.19E-14 | 5.67E-12 |
| EHD1 | 2277.130177 | 0.455642537 | 0.060766725 | 7.498224377 | 6.47E-14 | 5.91E-12 |
| ADO | 716.9003051 | 0.536866296 | 0.07176016 | 7.48139765 | 7.35E-14 | 6.68E-12 |
| STAT1 | 5168.896609 | 1.084570044 | 0.145357293 | 7.461407807 | 8.56E-14 | 7.74E-12 |
| PNPLA6 | 4102.736664 | 0.755968027 | 0.101473811 | 7.449883095 | 9.34E-14 | 8.41E-12 |
| CA13 | 82.13158125 | 0.80259779 | 0.107825083 | 7.44351657 | 9.80E-14 | 8.79E-12 |
| ITGA2 | 49.79368186 | 1.462619043 | 0.197310158 | 7.41279139 | 1.24E-13 | 1.10E-11 |
| CTNS | 625.3175282 | 0.753328512 | 0.101643691 | 7.411463563 | 1.25E-13 | 1.11E-11 |
| NBN | 1386.834673 | 0.536662008 | 0.072423953 | 7.41000717 | 1.26E-13 | 1.12E-11 |
| NFAM1 | 378.5780554 | -0.973861466 | 0.13150151 | -7.405705563 | 1.30E-13 | 1.15E-11 |
| LIMK2 | 316.899309 | 0.689876362 | 0.093210819 | 7.401247752 | 1.35E-13 | 1.18E-11 |
| ATP6V1B2 | 8375.346568 | 0.425226388 | 0.057521577 | 7.392467491 | 1.44E-13 | 1.26E-11 |
| RNASET2 | 1407.391052 | -0.521434352 | 0.070712791 | -7.37397496 | 1.66E-13 | 1.44E-11 |
| CCL22 | 3558.304694 | 0.811858233 | 0.11010845 | 7.373260049 | 1.67E-13 | 1.44E-11 |
| LYRM1 | 104.9937362 | 1.060670326 | 0.144143994 | 7.358408058 | 1.86E-13 | 1.60E-11 |
| MLLT4 | 410.7442541 | 0.789742107 | 0.107539185 | 7.343761295 | 2.08E-13 | 1.78E-11 |
| IRAK2 | 195.3092025 | 0.647755673 | 0.088296576 | 7.336135778 | 2.20E-13 | 1.87E-11 |
| G0S2 | 988.3754435 | 1.356846837 | 0.185189769 | 7.326791544 | 2.36E-13 | 2.00E-11 |
| METTL7A | 39.1718659 | -1.23490815 | 0.168714584 | -7.319510378 | 2.49E-13 | 2.10E-11 |
| CXCL10 | 1262.698126 | 2.83285329 | 0.388002908 | 7.301113552 | 2.85E-13 | 2.40E-11 |
| OPN3 | 4116.363433 | -0.493806557 | 0.067874207 | -7.275319716 | 3.46E-13 | 2.89E-11 |

| | | | | | | |
|---|---|---|---|---|---|---|
| LRRC8C | 545.1125858 | 0.75055238 | 0.103249839 | 7.269283801 | 3.61E-13 | 3.01E-11 |
| MOB3C | 800.2237475 | 0.50192176 | 0.069092459 | 7.264494117 | 3.74E-13 | 3.11E-11 |
| RIPK2 | 1437.766902 | 0.838609073 | 0.115528995 | 7.258862376 | 3.90E-13 | 3.23E-11 |
| DYRK4 | 54.09130687 | -0.995584939 | 0.137369503 | -7.2474961 | 4.25E-13 | 3.50E-11 |
| NLRC5 | 283.3207475 | 0.935278548 | 0.129136528 | 7.242556111 | 4.40E-13 | 3.61E-11 |
| APBA1 | 66.5378528 | -0.933832642 | 0.129527424 | -7.209536149 | 5.61E-13 | 4.58E-11 |
| TGM2 | 11146.04339 | 0.739864602 | 0.102870314 | 7.192207069 | 6.38E-13 | 5.18E-11 |
| SMOX | 119.1290431 | 1.313181081 | 0.183002028 | 7.175773389 | 7.19E-13 | 5.82E-11 |
| ID2 | 301.5841191 | 0.831627752 | 0.115970775 | 7.171011449 | 7.44E-13 | 6.00E-11 |
| FXYD6 | 228.3546024 | -0.909652717 | 0.127108402 | -7.156511299 | 8.28E-13 | 6.64E-11 |
| IFI35 | 1066.109675 | 0.78855813 | 0.110276109 | 7.150761281 | 8.63E-13 | 6.90E-11 |
| PI3 | 37.70799402 | 2.477698974 | 0.34682262 | 7.14399475 | 9.07E-13 | 7.22E-11 |
| SLC46A1 | 30.81370308 | -1.354715221 | 0.190196053 | -7.12273045 | 1.06E-12 | 8.39E-11 |
| ANKLE2 | 1638.422242 | 0.258791868 | 0.036382657 | 7.113055762 | 1.14E-12 | 8.96E-11 |
| DIRAS2 | 18.17361524 | 2.64416135 | 0.371932224 | 7.109255883 | 1.17E-12 | 9.14E-11 |
| DYRK3 | 145.4273563 | 0.720029122 | 0.101274306 | 7.109691996 | 1.16E-12 | 9.14E-11 |
| SCIMP | 2365.258652 | 0.65484942 | 0.092164074 | 7.1052569 | 1.20E-12 | 9.37E-11 |
| PLEKHA7 | 31.18176927 | 1.7774044 | 0.250387463 | 7.098615792 | 1.26E-12 | 9.79E-11 |
| HSD17B4 | 2874.084744 | -0.335771033 | 0.047488575 | -7.070564514 | 1.54E-12 | 1.19E-10 |
| PQLC3 | 783.7105593 | -0.550937308 | 0.078071474 | -7.056832394 | 1.70E-12 | 1.31E-10 |
| SEMA4D | 877.7415556 | 0.700937753 | 0.099364113 | 7.054234453 | 1.74E-12 | 1.33E-10 |
| SDC4 | 1863.734841 | 0.560921829 | 0.079596432 | 7.047072522 | 1.83E-12 | 1.40E-10 |
| SLC7A8 | 1136.236365 | 0.677242049 | 0.096356972 | 7.028469613 | 2.09E-12 | 1.59E-10 |
| SAMD1 | 232.0517837 | -0.578882139 | 0.08244844 | -7.021141186 | 2.20E-12 | 1.67E-10 |
| MPP1 | 3277.166127 | -0.436225096 | 0.062163266 | -7.01740951 | 2.26E-12 | 1.71E-10 |
| SLC27A1 | 159.8329987 | -0.855174168 | 0.122053699 | -7.006540356 | 2.44E-12 | 1.84E-10 |
| RPLP0P2 | 69.02091108 | 0.957621205 | 0.136706254 | 7.004955338 | 2.47E-12 | 1.85E-10 |
| GCH1 | 320.5811373 | 1.004647465 | 0.1435391 | 6.999120543 | 2.58E-12 | 1.92E-10 |
| TAP2 | 1292.833098 | 0.657454832 | 0.094000345 | 6.994174697 | 2.67E-12 | 1.98E-10 |
| ERMP1 | 441.7384846 | -0.48884703 | 0.069923374 | -6.991181964 | 2.73E-12 | 2.02E-10 |
| SPP1 | 1074.256079 | 0.615807709 | 0.088281452 | 6.975505013 | 3.05E-12 | 2.25E-10 |
| RHBDD2 | 1465.352774 | 0.833627753 | 0.119582331 | 6.971161604 | 3.14E-12 | 2.31E-10 |
| DNAJB5 | 238.3648114 | 0.761266227 | 0.109691737 | 6.940050792 | 3.92E-12 | 2.87E-10 |
| CYTIP | 893.3669648 | -0.399444516 | 0.057563238 | -6.939229407 | 3.94E-12 | 2.88E-10 |
| MPND | 132.9398583 | -0.585560775 | 0.084496253 | -6.930020624 | 4.21E-12 | 3.06E-10 |
| SLC41A2 | 506.4520012 | 0.73214493 | 0.105831727 | 6.918009867 | 4.58E-12 | 3.32E-10 |
| PSTPIP2 | 815.1099124 | 1.187456497 | 0.171780307 | 6.912646277 | 4.76E-12 | 3.43E-10 |
| SCARB1 | 524.7780493 | -0.960011118 | 0.138977196 | -6.907688071 | 4.93E-12 | 3.54E-10 |
| CLMN | 150.4916957 | -0.698867818 | 0.101332757 | -6.896761102 | 5.32E-12 | 3.81E-10 |
| FAM129A | 3132.574396 | 0.549243528 | 0.079656191 | 6.8951769 | 5.38E-12 | 3.84E-10 |
| CPNE8 | 214.2599879 | 0.556684618 | 0.080760623 | 6.893020336 | 5.46E-12 | 3.87E-10 |
| SOCS2 | 8.728950617 | 3.178367354 | 0.461112476 | 6.892824461 | 5.47E-12 | 3.87E-10 |
| SERPINF1 | 69.39349812 | -1.058944978 | 0.153836261 | -6.883584997 | 5.84E-12 | 4.12E-10 |
| FNIP2 | 2696.750497 | 0.551682063 | 0.080153206 | 6.882844618 | 5.87E-12 | 4.13E-10 |
| MICAL1 | 1107.367579 | -0.494637702 | 0.071937246 | -6.875961084 | 6.16E-12 | 4.31E-10 |
| RHOU | 772.9906028 | 0.583911645 | 0.085215783 | 6.852153703 | 7.27E-12 | 5.08E-10 |
| SLC19A3 | 337.6956912 | 1.025533017 | 0.149692336 | 6.850938702 | 7.34E-12 | 5.10E-10 |

| | | | | | | |
|---|---|---|---|---|---|---|
| CRYL1 | 242.8708629 | -0.493342347 | 0.072070691 | -6.845256256 | 7.63E-12 | 5.29E-10 |
| IL15RA | 467.0655212 | 0.817493477 | 0.119459905 | 6.84324569 | 7.74E-12 | 5.35E-10 |
| GPR68 | 242.8144155 | 0.907878518 | 0.132879137 | 6.832363153 | 8.35E-12 | 5.75E-10 |
| ANO5 | 634.4207868 | -0.49115306 | 0.071915534 | -6.829582369 | 8.52E-12 | 5.84E-10 |
| TP53BP2 | 1477.404074 | 0.346151692 | 0.050752488 | 6.820388619 | 9.08E-12 | 6.20E-10 |
| MPHOSPH | 594.5728062 | 0.497371583 | 0.072932457 | 6.81961917 | 9.13E-12 | 6.21E-10 |
| KRT23 | 144.367949 | 1.080436932 | 0.158626786 | 6.811188427 | 9.68E-12 | 6.57E-10 |
| MMP12 | 243.3681758 | 1.599323537 | 0.234941318 | 6.807331929 | 9.94E-12 | 6.72E-10 |
| KANK1 | 78.23864976 | 0.841461222 | 0.123625041 | 6.806559709 | 1.00E-11 | 6.73E-10 |
| DUSP6 | 1618.851732 | 0.480850009 | 0.070687915 | 6.802435899 | 1.03E-11 | 6.91E-10 |
| TAP1 | 2676.9272 | 0.92791735 | 0.136422085 | 6.801811822 | 1.03E-11 | 6.91E-10 |
| PXN | 259.1366425 | 0.74809912 | 0.110124206 | 6.793230578 | 1.10E-11 | 7.31E-10 |
| FAM105A | 171.2814605 | -0.684245363 | 0.100984 | -6.775779923 | 1.24E-11 | 8.22E-10 |
| VOPP1 | 1842.130357 | -0.502539907 | 0.074176035 | -6.774963211 | 1.24E-11 | 8.24E-10 |
| EMR3 | 112.8847923 | 1.219945399 | 0.180265598 | 6.767488703 | 1.31E-11 | 8.65E-10 |
| CD97 | 3787.976755 | 0.54269329 | 0.080331178 | 6.755699426 | 1.42E-11 | 9.35E-10 |
| GBP1 | 3684.740953 | 1.09129027 | 0.161589813 | 6.753459576 | 1.44E-11 | 9.46E-10 |
| IL23A | 15.83024036 | 2.528776364 | 0.375359782 | 6.736940095 | 1.62E-11 | 1.06E-09 |
| DUSP2 | 219.9843208 | 0.554743511 | 0.082386018 | 6.733466726 | 1.66E-11 | 1.08E-09 |
| HTRA4 | 199.5667877 | -0.613217361 | 0.091158382 | -6.72694433 | 1.73E-11 | 1.12E-09 |
| ITGA5 | 3902.80036 | 0.510383949 | 0.076035679 | 6.712427046 | 1.91E-11 | 1.24E-09 |
| CTPS2 | 103.929474 | -0.672527381 | 0.100214869 | -6.710854247 | 1.93E-11 | 1.25E-09 |
| ACP5 | 13996.76343 | -0.488058084 | 0.072800635 | -6.704036044 | 2.03E-11 | 1.30E-09 |
| SLAMF8 | 5377.814632 | 0.507484291 | 0.075915189 | 6.684884777 | 2.31E-11 | 1.48E-09 |
| PDCD4 | 1020.439082 | -0.4896687 | 0.073283885 | -6.681805998 | 2.36E-11 | 1.51E-09 |
| SAMD9L | 364.4766358 | 0.840191063 | 0.126130899 | 6.661262769 | 2.71E-11 | 1.73E-09 |
| YRDC | 219.3869652 | 0.51476668 | 0.077364705 | 6.653766488 | 2.86E-11 | 1.81E-09 |
| FBXO6 | 168.2185518 | 0.821917729 | 0.123577796 | 6.651014602 | 2.91E-11 | 1.84E-09 |
| ADAMTSL4 | 761.9307313 | 0.825494874 | 0.124275927 | 6.642435875 | 3.09E-11 | 1.94E-09 |
| HAPLN3 | 72.47387495 | 1.840712883 | 0.277126829 | 6.642131662 | 3.09E-11 | 1.94E-09 |
| CERK | 516.3637558 | -0.586537709 | 0.088391145 | -6.635706669 | 3.23E-11 | 2.02E-09 |
| PTPN7 | 182.6455356 | 0.746186137 | 0.112491459 | 6.63326927 | 3.28E-11 | 2.05E-09 |
| LMNB1 | 67.8836423 | 1.23961294 | 0.186971974 | 6.629939824 | 3.36E-11 | 2.09E-09 |
| TJP1 | 212.4942891 | 0.451934659 | 0.068266857 | 6.620118117 | 3.59E-11 | 2.22E-09 |
| KLF13 | 1185.178543 | -0.427905539 | 0.064780009 | -6.605518391 | 3.96E-11 | 2.45E-09 |
| NR3C2 | 32.96190053 | -0.816971261 | 0.124138633 | -6.581120177 | 4.67E-11 | 2.87E-09 |
| IL32 | 1009.453207 | 0.775883092 | 0.118051069 | 6.572435952 | 4.95E-11 | 3.04E-09 |
| MTF1 | 1109.075961 | 0.58467738 | 0.089137728 | 6.559258258 | 5.41E-11 | 3.30E-09 |
| NLRP3 | 94.70526072 | 0.74653189 | 0.113817553 | 6.55902248 | 5.42E-11 | 3.30E-09 |
| IMPA2 | 98.4881385 | -0.826238504 | 0.126185282 | -6.547819927 | 5.84E-11 | 3.54E-09 |
| MFSD2A | 345.2465704 | 0.873659696 | 0.133426824 | 6.547856497 | 5.84E-11 | 3.54E-09 |
| ANGEL1 | 469.6439437 | -0.5648584 | 0.086275242 | -6.54716681 | 5.86E-11 | 3.54E-09 |
| ATP6V0D2 | 490.8998773 | -0.531454065 | 0.08141797 | -6.527478662 | 6.69E-11 | 4.03E-09 |
| B3GNT5 | 804.7518749 | 0.579884081 | 0.089110535 | 6.507469433 | 7.64E-11 | 4.59E-09 |
| TSFM | 304.3834327 | 0.664266698 | 0.102167271 | 6.50175629 | 7.94E-11 | 4.75E-09 |
| ZDHHC3 | 407.6594637 | 0.315136447 | 0.04866646 | 6.475433977 | 9.45E-11 | 5.64E-09 |
| CTSF | 58.35824335 | -0.958572592 | 0.148133971 | -6.470984232 | 9.74E-11 | 5.79E-09 |

| | | | | | | |
|---|---|---|---|---|---|---|
| PSME4 | 1433.952691 | 0.637853944 | 0.098668146 | 6.464638982 | 1.02E-10 | 6.02E-09 |
| PCMTD2 | 185.2341658 | -0.538075444 | 0.083361222 | -6.454745169 | 1.08E-10 | 6.41E-09 |
| C15orf52 | 33.7504192 | -1.005351342 | 0.155810227 | -6.452409191 | 1.10E-10 | 6.49E-09 |
| SIPA1L1 | 163.4565388 | 0.72766712 | 0.112902862 | 6.445072379 | 1.16E-10 | 6.79E-09 |
| PDLIM4 | 94.75079859 | 0.81532209 | 0.126813266 | 6.429312293 | 1.28E-10 | 7.51E-09 |
| USP32 | 1300.671153 | 0.382029848 | 0.059528388 | 6.417607799 | 1.38E-10 | 8.08E-09 |
| SLC4A11 | 66.96429417 | -0.582538665 | 0.09099067 | -6.402180203 | 1.53E-10 | 8.92E-09 |
| NEDD4L | 216.1290598 | 0.559620668 | 0.087567598 | 6.390727661 | 1.65E-10 | 9.58E-09 |
| METTL9 | 439.6719515 | 0.707417915 | 0.111106354 | 6.367033819 | 1.93E-10 | 1.11E-08 |
| HSD17B14 | 114.8814947 | -0.56888449 | 0.089439903 | -6.360522186 | 2.01E-10 | 1.16E-08 |
| CEP170 | 1095.699715 | 0.520324502 | 0.081818344 | 6.35950907 | 2.02E-10 | 1.16E-08 |
| BTD | 187.8202984 | -0.571911699 | 0.090078533 | -6.349034337 | 2.17E-10 | 1.24E-08 |
| ASAH1 | 8939.08205 | -0.375586584 | 0.059231455 | -6.340998831 | 2.28E-10 | 1.31E-08 |
| ACSF2 | 222.8337915 | -0.597770996 | 0.094311221 | -6.338280716 | 2.32E-10 | 1.32E-08 |
| ITGA1 | 72.76062137 | 1.069818479 | 0.168930117 | 6.332905552 | 2.41E-10 | 1.37E-08 |
| CCDC71L | 410.4747232 | 0.865657338 | 0.13691866 | 6.322420471 | 2.57E-10 | 1.46E-08 |
| KLHL24 | 204.2891115 | -0.510764486 | 0.080987176 | -6.306732849 | 2.85E-10 | 1.61E-08 |
| NDP | 34.67706067 | 1.958219641 | 0.310535708 | 6.305940312 | 2.86E-10 | 1.61E-08 |
| CHST3 | 85.89771149 | 0.687877145 | 0.109110144 | 6.304428899 | 2.89E-10 | 1.62E-08 |
| ME1 | 639.4307693 | -0.728043337 | 0.115519874 | -6.302321064 | 2.93E-10 | 1.64E-08 |
| NFIA | 88.79914361 | -0.749886516 | 0.119182705 | -6.291907177 | 3.14E-10 | 1.75E-08 |
| LCP2 | 1428.484459 | 0.505960197 | 0.080673872 | 6.271673685 | 3.57E-10 | 1.99E-08 |
| RGCC | 4769.565172 | 0.499723444 | 0.079764452 | 6.26498939 | 3.73E-10 | 2.07E-08 |
| ACACB | 86.99577344 | -0.560713031 | 0.089563139 | -6.260533467 | 3.84E-10 | 2.12E-08 |
| TNFRSF11B | 4.931680335 | 3.791262035 | 0.605812679 | 6.258142436 | 3.90E-10 | 2.15E-08 |
| PLEKHN1 | 22.42064868 | 1.145170199 | 0.183188925 | 6.251306947 | 4.07E-10 | 2.24E-08 |
| KIAA0513 | 1146.07506 | -0.410815014 | 0.065877303 | -6.236062989 | 4.49E-10 | 2.46E-08 |
| ARHGAP10 | 824.5895915 | 0.478362718 | 0.076722593 | 6.234965474 | 4.52E-10 | 2.47E-08 |
| CLEC7A | 164.6860348 | 0.6318478 | 0.101347647 | 6.234459494 | 4.53E-10 | 2.47E-08 |
| PLTP | 501.0534685 | -0.752749516 | 0.120763412 | -6.233258113 | 4.57E-10 | 2.49E-08 |
| NR4A2 | 78.58223153 | 0.602257461 | 0.096647084 | 6.231511978 | 4.62E-10 | 2.51E-08 |
| SUCNR1 | 134.4749944 | 0.838628108 | 0.134934061 | 6.215095745 | 5.13E-10 | 2.78E-08 |
| SAV1 | 704.8982411 | 0.649145771 | 0.104609541 | 6.205416481 | 5.46E-10 | 2.94E-08 |
| CH25H | 64.60340734 | 2.064510867 | 0.332770806 | 6.204002359 | 5.50E-10 | 2.96E-08 |
| TEC | 132.4568504 | 0.552981697 | 0.089137964 | 6.203660848 | 5.52E-10 | 2.96E-08 |
| CKAP4 | 331.4210594 | 0.523601882 | 0.084423492 | 6.202087444 | 5.57E-10 | 2.98E-08 |
| SGK110 | 19.81876487 | 1.263914132 | 0.203856496 | 6.200018919 | 5.65E-10 | 3.01E-08 |
| RASSF5 | 1203.826244 | 0.435132738 | 0.070203873 | 6.198130054 | 5.71E-10 | 3.04E-08 |
| RHOH | 32.16830838 | 1.184418531 | 0.191187111 | 6.195075218 | 5.83E-10 | 3.08E-08 |
| SBNO2 | 743.0019975 | 0.468495905 | 0.075620561 | 6.195350831 | 5.82E-10 | 3.08E-08 |
| LYN | 2014.726634 | 0.340849371 | 0.05503424 | 6.19340558 | 5.89E-10 | 3.11E-08 |
| FABP4 | 12133.92014 | -0.741713022 | 0.119984649 | -6.181732626 | 6.34E-10 | 3.34E-08 |
| SUMO3 | 1170.344746 | -0.249204945 | 0.040381096 | -6.171326901 | 6.77E-10 | 3.55E-08 |
| MAP7 | 710.1697558 | 0.545885664 | 0.088561352 | 6.163926521 | 7.10E-10 | 3.72E-08 |
| INSR | 132.6103925 | -0.476946537 | 0.07758993 | -6.147015927 | 7.90E-10 | 4.12E-08 |
| ALAS1 | 2537.386398 | 0.527146519 | 0.085977055 | 6.131246546 | 8.72E-10 | 4.54E-08 |
| SASH3 | 333.8810468 | -0.663205276 | 0.108204007 | -6.129211797 | 8.83E-10 | 4.59E-08 |

| | | | | | | |
|---|---|---|---|---|---|---|
| RCN1 | 954.0992289 | 0.377524358 | 0.061608976 | 6.127749312 | 8.91E-10 | 4.62E-08 |
| LAP3 | 4477.401672 | 0.661641863 | 0.107985348 | 6.127144795 | 8.95E-10 | 4.62E-08 |
| WDR91 | 247.9575145 | -0.546112178 | 0.089262805 | -6.118026182 | 9.47E-10 | 4.88E-08 |
| SEMA4A | 425.3147935 | -0.530954744 | 0.08684865 | -6.113563576 | 9.74E-10 | 5.01E-08 |
| ASAP2 | 36.66956201 | 1.470150423 | 0.240655143 | 6.108950774 | 1.00E-09 | 5.14E-08 |
| TTC39B | 850.2495784 | 0.636589042 | 0.104490177 | 6.092333864 | 1.11E-09 | 5.69E-08 |
| IRF7 | 721.3731824 | 0.64423775 | 0.105871976 | 6.085064007 | 1.16E-09 | 5.93E-08 |
| KBTBD8 | 128.5221541 | 0.606887871 | 0.099772163 | 6.082737433 | 1.18E-09 | 6.01E-08 |
| SIRPB2 | 56.6148566 | -0.936459961 | 0.154324409 | -6.068126025 | 1.29E-09 | 6.56E-08 |
| EIF1B | 4105.995221 | 0.558876052 | 0.092192219 | 6.062074007 | 1.34E-09 | 6.80E-08 |
| CBX6 | 98.96186532 | 0.639450986 | 0.105581079 | 6.05649223 | 1.39E-09 | 7.02E-08 |
| HS3ST3A1 | 37.55724168 | 1.151467527 | 0.190376273 | 6.048377296 | 1.46E-09 | 7.34E-08 |
| PDK4 | 987.2668638 | -0.595184908 | 0.098402149 | -6.048495003 | 1.46E-09 | 7.34E-08 |
| SDS | 30.44329127 | -1.503746344 | 0.248725451 | -6.045808089 | 1.49E-09 | 7.44E-08 |
| IFIT3 | 138.9877153 | 1.444978932 | 0.239164633 | 6.041775135 | 1.52E-09 | 7.61E-08 |
| VAT1 | 18214.39702 | -0.39903987 | 0.066131458 | -6.034040103 | 1.60E-09 | 7.96E-08 |
| EIF4EBP1 | 475.0347666 | -0.473243896 | 0.078440152 | -6.033184302 | 1.61E-09 | 7.98E-08 |
| PTGFRN | 77.61756738 | -0.993596721 | 0.164760542 | -6.030550195 | 1.63E-09 | 8.09E-08 |
| ZC3H12A | 565.4799209 | 0.491138677 | 0.081462874 | 6.028987863 | 1.65E-09 | 8.15E-08 |
| SLC27A3 | 702.6120821 | 0.633983274 | 0.105342569 | 6.018300879 | 1.76E-09 | 8.68E-08 |
| CAMK1 | 155.685841 | 0.635284487 | 0.105690251 | 6.010814415 | 1.85E-09 | 9.07E-08 |
| CCDC69 | 503.4811382 | -0.647235588 | 0.107796219 | -6.004251281 | 1.92E-09 | 9.42E-08 |
| CD80 | 703.8998354 | 0.552799266 | 0.09230069 | 5.989113066 | 2.11E-09 | 1.03E-07 |
| ADAMTSL1 | 8.040782643 | 2.609912632 | 0.435959589 | 5.98659302 | 2.14E-09 | 1.05E-07 |
| MTHFD2 | 4181.957525 | 0.339322694 | 0.05677005 | 5.977142743 | 2.27E-09 | 1.10E-07 |
| AQP3 | 1200.819704 | 0.575414155 | 0.096291954 | 5.975724152 | 2.29E-09 | 1.11E-07 |
| HIP1 | 532.3029625 | 0.712734585 | 0.119303291 | 5.974140178 | 2.31E-09 | 1.12E-07 |
| MAP3K5 | 492.1963158 | 0.564827845 | 0.094781255 | 5.95927795 | 2.53E-09 | 1.22E-07 |
| TMEM171 | 13.77930664 | 2.072354475 | 0.347921179 | 5.956390695 | 2.58E-09 | 1.24E-07 |
| FAM40B | 95.79990112 | 0.75843075 | 0.127513487 | 5.94784733 | 2.72E-09 | 1.31E-07 |
| METTL21B | 145.2546008 | 0.728440474 | 0.122666133 | 5.938399275 | 2.88E-09 | 1.38E-07 |
| NCF4 | 1196.540062 | -0.351987327 | 0.059287846 | -5.936922192 | 2.90E-09 | 1.39E-07 |
| PLXND1 | 1509.891903 | -0.436850761 | 0.07358318 | -5.936829026 | 2.91E-09 | 1.39E-07 |
| GCHFR | 111.3707821 | -0.765346254 | 0.128929146 | -5.936177176 | 2.92E-09 | 1.39E-07 |
| MERTK | 13.20209715 | -1.802154918 | 0.303889592 | -5.930294968 | 3.02E-09 | 1.43E-07 |
| PIK3AP1 | 5194.732343 | 0.338748819 | 0.05713556 | 5.928861428 | 3.05E-09 | 1.44E-07 |
| FYB | 635.0122973 | -0.661312455 | 0.111929539 | -5.908292494 | 3.46E-09 | 1.63E-07 |
| TMEM170I | 166.5142858 | -0.488737907 | 0.082721102 | -5.908261571 | 3.46E-09 | 1.63E-07 |
| SLCO2B1 | 2467.023782 | 0.622443831 | 0.105384123 | 5.906428901 | 3.50E-09 | 1.64E-07 |
| CD47 | 1741.300853 | 0.42558707 | 0.072224617 | 5.892548651 | 3.80E-09 | 1.78E-07 |
| JAKMIP2 | 787.7240298 | 0.47914997 | 0.081309824 | 5.892891482 | 3.79E-09 | 1.78E-07 |
| EML4 | 2026.709319 | 0.58590863 | 0.099517252 | 5.887508122 | 3.92E-09 | 1.83E-07 |
| YES1 | 55.4523088 | 0.830488963 | 0.141134888 | 5.884363367 | 4.00E-09 | 1.86E-07 |
| SLIT3 | 62.36436815 | -0.961838429 | 0.163648444 | -5.877467608 | 4.17E-09 | 1.93E-07 |
| TSPAN4 | 212.1603511 | -0.712745522 | 0.121313091 | -5.875256475 | 4.22E-09 | 1.96E-07 |
| LTV1 | 276.0914399 | 0.331962241 | 0.056577542 | 5.86738534 | 4.43E-09 | 2.05E-07 |
| TNFSF15 | 88.44038134 | 1.263277104 | 0.215414036 | 5.864414067 | 4.51E-09 | 2.08E-07 |

| | | | | | | |
|---|---|---|---|---|---|---|
| BRI3BP | 81.66300415 | -0.600590925 | 0.102491074 | -5.859933973 | 4.63E-09 | 2.12E-07 |
| STON2 | 194.4719178 | 0.728051863 | 0.124241644 | 5.85996644 | 4.63E-09 | 2.12E-07 |
| SLC25A25 | 237.4825861 | 0.447736597 | 0.076438665 | 5.857462277 | 4.70E-09 | 2.15E-07 |
| GEM | 229.7454645 | 0.596096274 | 0.101815843 | 5.854651481 | 4.78E-09 | 2.18E-07 |
| PDE1B | 373.6837601 | 0.642713487 | 0.109863228 | 5.85012381 | 4.91E-09 | 2.24E-07 |
| ENTPD7 | 183.5291227 | 0.547147453 | 0.09370901 | 5.838792389 | 5.26E-09 | 2.39E-07 |
| NEK6 | 135.0684334 | 0.753976634 | 0.129244586 | 5.833719305 | 5.42E-09 | 2.46E-07 |
| GALNT2 | 1105.189462 | 0.334026396 | 0.057395225 | 5.81975928 | 5.89E-09 | 2.67E-07 |
| GSTK1 | 1100.566853 | -0.406391334 | 0.069858337 | -5.817363447 | 5.98E-09 | 2.70E-07 |
| SLC48A1 | 114.4860538 | -0.817628291 | 0.140573657 | -5.816369211 | 6.01E-09 | 2.71E-07 |
| CD93 | 28.12781885 | 1.125895345 | 0.193649361 | 5.814092733 | 6.10E-09 | 2.74E-07 |
| PPT1 | 5885.720174 | -0.232274869 | 0.039988022 | -5.808611142 | 6.30E-09 | 2.82E-07 |
| EBI3 | 21.63758476 | 1.739378183 | 0.299699106 | 5.803748317 | 6.48E-09 | 2.90E-07 |
| DCAF7 | 753.8559339 | 0.354206608 | 0.061082635 | 5.798810179 | 6.68E-09 | 2.98E-07 |
| PPP1R15B | 1262.475436 | 0.373075786 | 0.064353703 | 5.79726988 | 6.74E-09 | 3.00E-07 |
| BNIP3L | 1055.703886 | -0.484789517 | 0.083705874 | -5.79158298 | 6.97E-09 | 3.10E-07 |
| PPFIBP1 | 123.5055521 | 0.624715265 | 0.107941938 | 5.787512028 | 7.14E-09 | 3.16E-07 |
| ARL5B | 250.625536 | 0.490859093 | 0.084844454 | 5.785399838 | 7.23E-09 | 3.19E-07 |
| RBPJ | 1143.795165 | 0.410942508 | 0.071028331 | 5.785614006 | 7.22E-09 | 3.19E-07 |
| ALDH1A2 | 383.4328011 | 0.892938701 | 0.154803392 | 5.768211482 | 8.01E-09 | 3.52E-07 |
| CD9 | 9661.791192 | -0.425153612 | 0.073748054 | -5.764946834 | 8.17E-09 | 3.58E-07 |
| MICAL2 | 210.5735555 | 1.027250288 | 0.178615073 | 5.751195968 | 8.86E-09 | 3.88E-07 |
| DOT1L | 498.9049204 | 0.455102679 | 0.079137911 | 5.750754243 | 8.88E-09 | 3.88E-07 |
| PBXIP1 | 179.3315481 | -0.598430186 | 0.104294169 | -5.737906487 | 9.59E-09 | 4.18E-07 |
| SPATS2L | 246.5279954 | 0.469300982 | 0.081874522 | 5.73195383 | 9.93E-09 | 4.32E-07 |
| GUCY1B3 | 14.67200552 | 1.292418264 | 0.225675418 | 5.726889854 | 1.02E-08 | 4.44E-07 |
| RBMS1 | 1233.467556 | 0.59470634 | 0.103875965 | 5.725158271 | 1.03E-08 | 4.47E-07 |
| CUEDC1 | 261.6716346 | -0.335934451 | 0.058689745 | -5.72390374 | 1.04E-08 | 4.50E-07 |
| FBXO10 | 141.7011098 | -0.521151609 | 0.091104209 | -5.720390012 | 1.06E-08 | 4.58E-07 |
| SBSN | 4.822356375 | 2.736595206 | 0.478464548 | 5.719535993 | 1.07E-08 | 4.59E-07 |
| OSBPL1A | 502.6643853 | -0.431350211 | 0.075432211 | -5.718382182 | 1.08E-08 | 4.61E-07 |
| ATP2C1 | 981.4978093 | 0.454714721 | 0.079586141 | 5.713491248 | 1.11E-08 | 4.74E-07 |
| IFI44 | 293.1977069 | 0.920225975 | 0.161109039 | 5.711820884 | 1.12E-08 | 4.77E-07 |
| C7orf41 | 50.06598937 | -0.655692035 | 0.114849628 | -5.709135062 | 1.14E-08 | 4.83E-07 |
| NPAS2 | 72.25399026 | 0.897480151 | 0.157193254 | 5.709406279 | 1.13E-08 | 4.83E-07 |
| NRIP3 | 584.911305 | 0.451306408 | 0.079149841 | 5.701924382 | 1.18E-08 | 5.03E-07 |
| STEAP3 | 899.1482994 | -0.498408518 | 0.087460626 | -5.698661655 | 1.21E-08 | 5.11E-07 |
| CD101 | 295.4131788 | -0.620575615 | 0.10905169 | -5.690655626 | 1.27E-08 | 5.35E-07 |
| BMP2 | 9.808725736 | 2.437510115 | 0.428575519 | 5.687469319 | 1.29E-08 | 5.44E-07 |
| GIMAP8 | 181.7903536 | 0.821672124 | 0.144507717 | 5.686008615 | 1.30E-08 | 5.47E-07 |
| SPOCD1 | 1394.305359 | 0.471916328 | 0.083007391 | 5.685232654 | 1.31E-08 | 5.48E-07 |
| ANXA11 | 4008.358647 | 0.389335455 | 0.068507947 | 5.683069956 | 1.32E-08 | 5.54E-07 |
| PHLDA2 | 449.0764474 | 0.542637546 | 0.095536666 | 5.679887822 | 1.35E-08 | 5.63E-07 |
| DBNDD2 | 1430.928321 | -0.319988398 | 0.056342332 | -5.67936024 | 1.35E-08 | 5.64E-07 |
| KLF9 | 488.3005999 | 0.34487031 | 0.060755732 | 5.676341924 | 1.38E-08 | 5.73E-07 |
| SLC37A2 | 611.888622 | -0.421378652 | 0.074273853 | -5.673310804 | 1.40E-08 | 5.82E-07 |
| DUSP4 | 46.82833946 | 0.95129257 | 0.167936435 | 5.664599044 | 1.47E-08 | 6.11E-07 |

| | | | | | | |
|---|---|---|---|---|---|---|
| RPGR | 126.8063105 | 0.524476258 | 0.092607297 | 5.663444154 | 1.48E-08 | 6.13E-07 |
| HEG1 | 494.5825755 | 0.47542592 | 0.084256333 | 5.642613465 | 1.67E-08 | 6.91E-07 |
| NTAN1 | 783.6012603 | -0.28514888 | 0.050591445 | -5.636306342 | 1.74E-08 | 7.15E-07 |
| MPP7 | 28.90421226 | -0.970638502 | 0.172294632 | -5.633596887 | 1.76E-08 | 7.25E-07 |
| RAPGEF3 | 40.15430517 | -0.999258873 | 0.177396246 | -5.632920055 | 1.77E-08 | 7.25E-07 |
| WBP1 | 270.1767489 | -0.386441065 | 0.068605058 | -5.632836363 | 1.77E-08 | 7.25E-07 |
| GBP5 | 1000.368692 | 1.660166837 | 0.294759643 | 5.632273198 | 1.78E-08 | 7.26E-07 |
| RRAD | 163.0125154 | 0.966498598 | 0.171641874 | 5.630902143 | 1.79E-08 | 7.30E-07 |
| IQGAP2 | 2028.815398 | 0.354414494 | 0.062974537 | 5.627901577 | 1.82E-08 | 7.41E-07 |
| USP12 | 1121.331903 | 0.359998058 | 0.063987995 | 5.626025018 | 1.84E-08 | 7.48E-07 |
| TNFRSF4 | 114.4549642 | 0.971531805 | 0.172948526 | 5.617462193 | 1.94E-08 | 7.84E-07 |
| SMPDL3A | 40.09868041 | -0.991401911 | 0.176521216 | -5.616332883 | 1.95E-08 | 7.88E-07 |
| PNPLA3 | 120.1211707 | -0.597221117 | 0.106416744 | -5.61209727 | 2.00E-08 | 8.06E-07 |
| KREMEN1 | 30.74954315 | 1.391647902 | 0.248426898 | 5.601840672 | 2.12E-08 | 8.53E-07 |
| IMPDH1 | 468.0700583 | -0.476812592 | 0.085254752 | -5.592797816 | 2.23E-08 | 8.97E-07 |
| TNFAIP8 | 1064.599739 | 0.466222404 | 0.08342047 | 5.588824934 | 2.29E-08 | 9.16E-07 |
| LTA4H | 5397.793789 | -0.525706495 | 0.094077555 | -5.588011902 | 2.30E-08 | 9.18E-07 |
| HDAC5 | 282.273404 | -0.39911553 | 0.071522206 | -5.580302296 | 2.40E-08 | 9.58E-07 |
| AICDA | 29.62689358 | -1.252791733 | 0.224536934 | -5.579446155 | 2.41E-08 | 9.60E-07 |
| FABP5 | 4614.615079 | -0.405373263 | 0.072758106 | -5.571520263 | 2.53E-08 | 1.00E-06 |
| JUNB | 3376.545444 | 0.373363793 | 0.067014564 | 5.571382869 | 2.53E-08 | 1.00E-06 |
| CYFIP1 | 1684.416591 | -0.273725195 | 0.049233902 | -5.559689199 | 2.70E-08 | 1.07E-06 |
| PGAP1 | 53.88604662 | 0.997780719 | 0.179663746 | 5.553600786 | 2.80E-08 | 1.10E-06 |
| LGMN | 1588.503697 | -0.455469481 | 0.082051515 | -5.551018534 | 2.84E-08 | 1.12E-06 |
| HS3ST3B1 | 55.85761899 | 1.057182568 | 0.190487319 | 5.549884224 | 2.86E-08 | 1.12E-06 |
| MST4 | 38.38670955 | 1.196598177 | 0.215605671 | 5.54993832 | 2.86E-08 | 1.12E-06 |
| SIK3 | 744.9755172 | 0.441013549 | 0.079606214 | 5.539938712 | 3.03E-08 | 1.18E-06 |
| BTG1 | 1954.688656 | 0.470442698 | 0.084992765 | 5.535091136 | 3.11E-08 | 1.22E-06 |
| PARP1 | 967.4342465 | -0.307278548 | 0.055527124 | -5.533845948 | 3.13E-08 | 1.22E-06 |
| PDSS1 | 175.8593555 | 0.570532552 | 0.10332204 | 5.521886265 | 3.35E-08 | 1.31E-06 |
| FCN1 | 42.51372094 | -0.898750715 | 0.162945893 | -5.515638955 | 3.48E-08 | 1.35E-06 |
| ELMO2 | 587.0225193 | 0.40832425 | 0.074047391 | 5.514363744 | 3.50E-08 | 1.36E-06 |
| MDGA1 | 12.67023441 | 1.394440969 | 0.252991244 | 5.511815134 | 3.55E-08 | 1.37E-06 |
| CXCL9 | 597.6961654 | 1.770043691 | 0.321781234 | 5.500767299 | 3.78E-08 | 1.46E-06 |
| PDK2 | 117.0524076 | -0.896802677 | 0.163118401 | -5.497863332 | 3.84E-08 | 1.48E-06 |
| PLK3 | 579.6857793 | 0.590900783 | 0.107521891 | 5.495632352 | 3.89E-08 | 1.50E-06 |
| FKBP9 | 136.3580694 | -0.480965496 | 0.087789168 | -5.478642829 | 4.29E-08 | 1.64E-06 |
| SNX8 | 1307.604316 | 0.299737802 | 0.054712123 | 5.478453168 | 4.29E-08 | 1.64E-06 |
| WSB1 | 879.7851647 | 0.284550813 | 0.051973004 | 5.47497334 | 4.38E-08 | 1.67E-06 |
| OR6N1 | 61.95414116 | -0.940852021 | 0.171946966 | -5.471757024 | 4.46E-08 | 1.70E-06 |
| PMP22 | 589.9259846 | 0.555970965 | 0.101613736 | 5.471415454 | 4.46E-08 | 1.70E-06 |
| CIITA | 1200.784206 | 0.553846757 | 0.101233485 | 5.470983826 | 4.48E-08 | 1.70E-06 |
| CYP2U1 | 27.06739532 | -1.013630424 | 0.185410285 | -5.466958983 | 4.58E-08 | 1.73E-06 |
| GATSL3 | 17.8751491 | -0.985629154 | 0.18027923 | -5.467236305 | 4.57E-08 | 1.73E-06 |
| HES2 | 47.85058451 | -1.074212264 | 0.196604553 | -5.4638219 | 4.66E-08 | 1.76E-06 |
| PLA2G16 | 2768.53468 | -0.309366506 | 0.056626386 | -5.46329243 | 4.67E-08 | 1.76E-06 |
| PARP9 | 277.8720816 | 0.868576819 | 0.159131544 | 5.458231581 | 4.81E-08 | 1.81E-06 |

| | | | | | | |
|---|---|---|---|---|---|---|
| NIPSNAP1 | 57.3297248 | -0.728581021 | 0.133531757 | -5.456237793 | 4.86E-08 | 1.83E-06 |
| AFMID | 13.61298987 | -1.467593263 | 0.269833518 | -5.438884211 | 5.36E-08 | 2.00E-06 |
| JAK3 | 86.2108141 | 0.962842178 | 0.177036198 | 5.438674061 | 5.37E-08 | 2.00E-06 |
| PCYOX1 | 215.9030429 | -0.476639518 | 0.087630535 | -5.439194433 | 5.35E-08 | 2.00E-06 |
| SORD | 98.413807 | -0.54171588 | 0.099588396 | -5.439548189 | 5.34E-08 | 2.00E-06 |
| AUH | 91.90883189 | -0.755065314 | 0.138853189 | -5.437868012 | 5.39E-08 | 2.01E-06 |
| OTUD7B | 69.67942755 | 0.731174718 | 0.134474742 | 5.437264346 | 5.41E-08 | 2.01E-06 |
| MFAP5 | 68.59914337 | -0.628030654 | 0.115520663 | -5.436522227 | 5.43E-08 | 2.01E-06 |
| PDGFB | 53.9438127 | 0.752897989 | 0.138709284 | 5.42788463 | 5.70E-08 | 2.11E-06 |
| CCDC28A | 272.2106726 | -0.381308333 | 0.070254573 | -5.427523327 | 5.71E-08 | 2.11E-06 |
| FMO5 | 75.02751769 | -0.597265014 | 0.110119376 | -5.423795855 | 5.83E-08 | 2.15E-06 |
| PPAP2B | 1839.090467 | 0.459334585 | 0.084687145 | 5.42389975 | 5.83E-08 | 2.15E-06 |
| RASGRP1 | 204.8939852 | 0.816842393 | 0.150658778 | 5.421804194 | 5.90E-08 | 2.17E-06 |
| SERPINB9 | 138.728803 | 1.114632248 | 0.205690115 | 5.418987927 | 5.99E-08 | 2.20E-06 |
| ANKRD12 | 2735.091531 | 0.39667685 | 0.073280693 | 5.413115416 | 6.19E-08 | 2.26E-06 |
| EPB41L3 | 3372.274746 | -0.21829404 | 0.040341962 | -5.411091283 | 6.26E-08 | 2.29E-06 |
| ALDH1B1 | 15.40143628 | -0.887265279 | 0.164069631 | -5.407858085 | 6.38E-08 | 2.32E-06 |
| LDHB | 1043.524569 | -0.422163999 | 0.078119584 | -5.40407381 | 6.51E-08 | 2.37E-06 |
| OR6K3 | 88.24812287 | -0.972291784 | 0.179990233 | -5.401914132 | 6.59E-08 | 2.39E-06 |
| MT2A | 11714.87947 | 0.47990035 | 0.088896369 | 5.39842465 | 6.72E-08 | 2.43E-06 |
| SHPK | 163.6939667 | 0.511558885 | 0.094805874 | 5.395856444 | 6.82E-08 | 2.47E-06 |
| TMCC3 | 321.190089 | 0.46882115 | 0.086906005 | 5.394577152 | 6.87E-08 | 2.48E-06 |
| PCNX | 889.9316832 | 0.339024226 | 0.062960635 | 5.384701465 | 7.26E-08 | 2.61E-06 |
| DCP1A | 764.5606532 | 0.400331852 | 0.074464481 | 5.376145084 | 7.61E-08 | 2.74E-06 |
| KCNAB1 | 84.05604413 | -1.227296328 | 0.228392214 | -5.373634713 | 7.72E-08 | 2.77E-06 |
| OSM | 261.3695544 | 0.723007805 | 0.134867486 | 5.360875546 | 8.28E-08 | 2.97E-06 |
| FARP1 | 538.3137037 | -0.534776531 | 0.099769015 | -5.360146466 | 8.32E-08 | 2.97E-06 |
| ANKRD29 | 96.5653672 | 0.704298854 | 0.131435077 | 5.358530395 | 8.39E-08 | 2.99E-06 |
| CCDC102B | 39.76604747 | 1.240076683 | 0.23162573 | 5.353795032 | 8.61E-08 | 3.07E-06 |
| MAP1LC3A | 195.9289794 | -0.559715339 | 0.10461067 | -5.350461273 | 8.77E-08 | 3.12E-06 |
| ATP2B1 | 1134.70532 | 0.375029399 | 0.070174634 | 5.344230214 | 9.08E-08 | 3.22E-06 |
| ITGAX | 4767.707299 | 0.402925689 | 0.075449956 | 5.34030382 | 9.28E-08 | 3.29E-06 |
| FCGR2C | 33.51677458 | 1.310180614 | 0.245651365 | 5.333496167 | 9.63E-08 | 3.41E-06 |
| ADORA2B | 739.0799193 | -0.411051404 | 0.077132521 | -5.329158135 | 9.87E-08 | 3.48E-06 |
| FGF10 | 15.23365506 | 1.839127517 | 0.3452903 | 5.326322575 | 1.00E-07 | 3.53E-06 |
| ABI3 | 107.1722171 | -0.707989081 | 0.133071881 | -5.320350741 | 1.04E-07 | 3.64E-06 |
| PTAFR | 567.5754725 | 0.511651699 | 0.096172754 | 5.320131532 | 1.04E-07 | 3.64E-06 |
| TOR3A | 675.0337073 | -0.554910425 | 0.104312178 | -5.319708933 | 1.04E-07 | 3.64E-06 |
| LINC00515 | 103.3176476 | 0.948716627 | 0.178414304 | 5.317491949 | 1.05E-07 | 3.68E-06 |
| IRF9 | 560.8163594 | 0.5034835 | 0.094743923 | 5.314150853 | 1.07E-07 | 3.74E-06 |
| BTG2 | 1357.418909 | 0.354329863 | 0.066725793 | 5.310238293 | 1.09E-07 | 3.81E-06 |
| MEGF9 | 466.2441769 | -0.394245378 | 0.074299517 | -5.306163384 | 1.12E-07 | 3.89E-06 |
| UGP2 | 1568.905552 | 0.320163504 | 0.060343131 | 5.305715761 | 1.12E-07 | 3.90E-06 |
| ESPL1 | 41.95525051 | 0.847802024 | 0.160019529 | 5.298115987 | 1.17E-07 | 4.05E-06 |
| MXD4 | 299.1886773 | -0.346727458 | 0.065547243 | -5.289733625 | 1.22E-07 | 4.24E-06 |
| LOC10012[illegible] | 33.09872068 | -1.240392716 | 0.234606382 | -5.287122654 | 1.24E-07 | 4.29E-06 |
| ITPR1 | 596.5043368 | -0.489863229 | 0.092716628 | -5.283445273 | 1.27E-07 | 4.37E-06 |

| | | | | | | |
|---|---|---|---|---|---|---|
| LRRC6 | 42.70240988 | -0.896896489 | 0.169956375 | -5.277215937 | 1.31E-07 | 4.51E-06 |
| RNF213 | 1654.445856 | 0.45570448 | 0.086372004 | 5.276066966 | 1.32E-07 | 4.53E-06 |
| CLDN12 | 305.3590903 | 0.443142196 | 0.084093586 | 5.269631349 | 1.37E-07 | 4.69E-06 |
| SLC25A22 | 127.5500043 | 0.599118986 | 0.113707913 | 5.268929567 | 1.37E-07 | 4.70E-06 |
| IRF4 | 42.81451045 | 0.885494956 | 0.168169873 | 5.265479116 | 1.40E-07 | 4.78E-06 |
| ATF3 | 2144.421326 | 0.46365134 | 0.088089352 | 5.263420952 | 1.41E-07 | 4.82E-06 |
| IGF1 | 54.90148528 | 1.126344319 | 0.214077607 | 5.261383168 | 1.43E-07 | 4.87E-06 |
| SPRR3 | 33.73225328 | -0.911526325 | 0.17329853 | -5.259861828 | 1.44E-07 | 4.90E-06 |
| TCP11L2 | 92.79051515 | -0.550065145 | 0.104616691 | -5.257909968 | 1.46E-07 | 4.94E-06 |
| IL6R | 976.3601357 | 0.583454082 | 0.111038373 | 5.254526576 | 1.48E-07 | 5.02E-06 |
| HDDC2 | 616.7412978 | -0.339854957 | 0.064738161 | -5.2496851 | 1.52E-07 | 5.15E-06 |
| VPS9D1 | 108.1104749 | 0.601804713 | 0.114842521 | 5.240260359 | 1.60E-07 | 5.41E-06 |
| MET | 178.7984272 | 0.957317854 | 0.182860608 | 5.235232799 | 1.65E-07 | 5.55E-06 |
| RBM8A | 1945.295223 | 0.379462603 | 0.072499474 | 5.234004913 | 1.66E-07 | 5.58E-06 |
| MUCL1 | 24.52409706 | 1.182507813 | 0.226006207 | 5.232191747 | 1.68E-07 | 5.62E-06 |
| IL2RG | 2867.771354 | 0.421389627 | 0.080598225 | 5.228274254 | 1.71E-07 | 5.73E-06 |
| FAM210B | 205.615992 | -0.407856858 | 0.078063968 | -5.224649345 | 1.74E-07 | 5.84E-06 |
| ENDOG | 127.2907468 | -0.344474612 | 0.065965093 | -5.222074229 | 1.77E-07 | 5.91E-06 |
| AQP9 | 5892.742647 | 0.366292894 | 0.070153364 | 5.221316148 | 1.78E-07 | 5.92E-06 |
| MMP10 | 969.4653755 | 0.664154779 | 0.127256838 | 5.219010537 | 1.80E-07 | 5.99E-06 |
| CALM1 | 2893.957653 | 0.212773909 | 0.040844262 | 5.209395327 | 1.89E-07 | 6.29E-06 |
| CLEC4A | 617.9034663 | 0.715311567 | 0.137425267 | 5.205094964 | 1.94E-07 | 6.43E-06 |
| ACSL5 | 4300.81584 | 0.472811325 | 0.090848374 | 5.204400546 | 1.95E-07 | 6.44E-06 |
| ZBTB4 | 831.5343372 | -0.41690328 | 0.080132892 | -5.20264863 | 1.96E-07 | 6.49E-06 |
| LXN | 61.86722594 | -0.588216554 | 0.113092295 | -5.201208049 | 1.98E-07 | 6.53E-06 |
| PIK3R6 | 14.84090163 | 1.27448452 | 0.245043487 | 5.201054454 | 1.98E-07 | 6.53E-06 |
| CLCF1 | 196.7157869 | 0.632912183 | 0.121749639 | 5.198472767 | 2.01E-07 | 6.61E-06 |
| KDM2B | 163.89388 | -0.553105663 | 0.106431757 | -5.196810392 | 2.03E-07 | 6.65E-06 |
| GPD2 | 2364.431467 | 0.30641277 | 0.058965749 | 5.196453424 | 2.03E-07 | 6.66E-06 |
| P2RY6 | 12.49661922 | 1.295762895 | 0.249492086 | 5.193603187 | 2.06E-07 | 6.75E-06 |
| CD84 | 1989.313741 | -0.609710212 | 0.117477616 | -5.190011793 | 2.10E-07 | 6.85E-06 |
| COQ2 | 297.1743655 | -0.304956196 | 0.058756681 | -5.19015353 | 2.10E-07 | 6.85E-06 |
| DOCK4 | 1088.522447 | 0.40520807 | 0.078089726 | 5.189006157 | 2.11E-07 | 6.88E-06 |
| PSME2 | 1305.767258 | 0.568092221 | 0.109617631 | 5.182489509 | 2.19E-07 | 7.11E-06 |
| CD4 | 996.7947728 | -0.404675822 | 0.078172576 | -5.176698033 | 2.26E-07 | 7.33E-06 |
| RUSC2 | 390.0150607 | 0.435625857 | 0.084174978 | 5.175241703 | 2.28E-07 | 7.37E-06 |
| GPR132 | 40.71177139 | 0.902794615 | 0.174458826 | 5.174829141 | 2.28E-07 | 7.37E-06 |
| NOTCH2 | 2520.000583 | -0.189116125 | 0.036646771 | -5.160512586 | 2.46E-07 | 7.95E-06 |
| PFKFB2 | 139.6631412 | -0.472774103 | 0.091642126 | -5.158916806 | 2.48E-07 | 8.00E-06 |
| PRSS21 | 191.064096 | -0.459063333 | 0.088989236 | -5.158638875 | 2.49E-07 | 8.00E-06 |
| CD300LF | 774.9405068 | 0.342572614 | 0.066414365 | 5.158110215 | 2.49E-07 | 8.01E-06 |
| ADCY3 | 1540.081265 | -0.250372523 | 0.048556658 | -5.156296399 | 2.52E-07 | 8.07E-06 |
| BRCA2 | 99.69799786 | 0.538020259 | 0.104388104 | 5.154038037 | 2.55E-07 | 8.16E-06 |
| LTBP3 | 149.5834937 | -0.537487274 | 0.104360904 | -5.150274214 | 2.60E-07 | 8.31E-06 |
| KIAA1147 | 566.6646842 | 0.57888615 | 0.112411842 | 5.14969009 | 2.61E-07 | 8.32E-06 |
| NUP107 | 387.1373386 | 0.341021925 | 0.06623052 | 5.149014786 | 2.62E-07 | 8.33E-06 |
| TMEM236 | 27.313469 | 1.1181004 | 0.2171384 | 5.149252271 | 2.62E-07 | 8.33E-06 |

| | | | | | | |
|---|---|---|---|---|---|---|
| ZNF697 | 254.5015984 | 0.444560639 | 0.086433468 | 5.14338544 | 2.70E-07 | 8.56E-06 |
| IFI16 | 2482.584338 | 0.325219722 | 0.063260555 | 5.140955873 | 2.73E-07 | 8.65E-06 |
| MXD1 | 421.091343 | 0.617836727 | 0.120174585 | 5.141159641 | 2.73E-07 | 8.65E-06 |
| CFB | 1232.569729 | 0.655815182 | 0.127627447 | 5.1385121 | 2.77E-07 | 8.75E-06 |
| ALOX5 | 1171.242556 | -0.52157412 | 0.101519035 | -5.137697762 | 2.78E-07 | 8.77E-06 |
| C15orf38 | 101.3446469 | -0.564215551 | 0.109835191 | -5.136928778 | 2.79E-07 | 8.79E-06 |
| SERPINE2 | 478.9458248 | 0.974395307 | 0.1896987 | 5.136541818 | 2.80E-07 | 8.80E-06 |
| EPB41L1 | 1654.927268 | -0.388838086 | 0.075715536 | -5.135512592 | 2.81E-07 | 8.83E-06 |
| PML | 245.2435382 | 0.614227428 | 0.119656102 | 5.133272918 | 2.85E-07 | 8.92E-06 |
| ELK3 | 263.1307276 | -0.382060954 | 0.07448396 | -5.129439322 | 2.91E-07 | 9.09E-06 |
| NFKBID | 215.2114218 | 0.443586554 | 0.086736032 | 5.114213148 | 3.15E-07 | 9.84E-06 |
| CAMK1D | 165.1231175 | -0.360385509 | 0.070526089 | -5.109960176 | 3.22E-07 | 1.00E-05 |
| CNNM3 | 387.635786 | -0.447981986 | 0.087706251 | -5.107754361 | 3.26E-07 | 1.01E-05 |
| STX11 | 1826.245144 | 0.294338763 | 0.057677838 | 5.103151815 | 3.34E-07 | 1.04E-05 |
| CASP4 | 2897.767741 | 0.435191748 | 0.085493794 | 5.090331415 | 3.57E-07 | 1.11E-05 |
| GADD45G | 351.6952299 | 0.483568925 | 0.095074238 | 5.086224544 | 3.65E-07 | 1.13E-05 |
| MARCO | 4083.971195 | -0.331830235 | 0.065256218 | -5.085036231 | 3.68E-07 | 1.14E-05 |
| H2AFY2 | 46.3910342 | -0.72064182 | 0.141795612 | -5.082257539 | 3.73E-07 | 1.15E-05 |
| SH2B2 | 225.2413388 | 0.400536874 | 0.07884149 | 5.080280395 | 3.77E-07 | 1.16E-05 |
| TRIM65 | 210.8887499 | -0.380722884 | 0.074943352 | -5.080142193 | 3.77E-07 | 1.16E-05 |
| DHRSX | 544.6111666 | -0.234512573 | 0.046227891 | -5.072967167 | 3.92E-07 | 1.20E-05 |
| ULK1 | 111.2108179 | -0.535041145 | 0.105674871 | -5.063087743 | 4.13E-07 | 1.27E-05 |
| RAI14 | 41.77960666 | 0.885618728 | 0.175113046 | 5.057411462 | 4.25E-07 | 1.30E-05 |
| PANX1 | 179.9890684 | 0.496644622 | 0.098277134 | 5.053511456 | 4.34E-07 | 1.33E-05 |
| NMT2 | 171.4437237 | -0.489219552 | 0.096869011 | -5.050320507 | 4.41E-07 | 1.34E-05 |
| ZDHHC7 | 548.8111359 | -0.312012881 | 0.061777716 | -5.050573309 | 4.40E-07 | 1.34E-05 |
| FCAR | 335.551716 | 0.739080093 | 0.14646131 | 5.046248008 | 4.51E-07 | 1.37E-05 |
| RAP2B | 563.0247364 | -0.42236995 | 0.083726527 | -5.044637146 | 4.54E-07 | 1.38E-05 |
| ZAK | 476.530189 | 0.464464794 | 0.092094461 | 5.043352112 | 4.57E-07 | 1.39E-05 |
| GTF3A | 1722.838164 | -0.278791761 | 0.055291241 | -5.042240982 | 4.60E-07 | 1.39E-05 |
| MRPS26 | 271.0666018 | -0.407401274 | 0.080880689 | -5.037064822 | 4.73E-07 | 1.43E-05 |
| ACOX3 | 154.738774 | 0.422001565 | 0.083802864 | 5.035646069 | 4.76E-07 | 1.44E-05 |
| CYBRD1 | 32.40006941 | -0.743479146 | 0.14775285 | -5.031910682 | 4.86E-07 | 1.46E-05 |
| TARP | 76.84692531 | 0.885672077 | 0.176069062 | 5.030253845 | 4.90E-07 | 1.47E-05 |
| CD48 | 125.9295998 | -0.647795872 | 0.128876285 | -5.026493982 | 5.00E-07 | 1.50E-05 |
| IFI30 | 77076.11511 | -0.2549052 | 0.050709197 | -5.026804128 | 4.99E-07 | 1.50E-05 |
| TSPAN15 | 207.8787835 | -0.42464202 | 0.084481499 | -5.026449861 | 5.00E-07 | 1.50E-05 |
| AKAP12 | 41.84021847 | 1.105392719 | 0.219997908 | 5.024560146 | 5.05E-07 | 1.51E-05 |
| KCTD17 | 314.5016128 | -0.47517837 | 0.094584839 | -5.023832335 | 5.07E-07 | 1.51E-05 |
| RAC2 | 7212.780328 | 0.36654479 | 0.072981495 | 5.022434634 | 5.10E-07 | 1.52E-05 |
| CECR1 | 2980.020053 | -0.295677483 | 0.058896779 | -5.020265772 | 5.16E-07 | 1.54E-05 |
| FCGR3A | 3174.608382 | -0.397640219 | 0.079352764 | -5.011044327 | 5.41E-07 | 1.61E-05 |
| AOC3 | 481.4724227 | -0.514612764 | 0.102704862 | -5.01059789 | 5.43E-07 | 1.61E-05 |
| HELZ2 | 78.30910201 | 0.921235118 | 0.183870636 | 5.010235131 | 5.44E-07 | 1.61E-05 |
| GFPT1 | 281.7911202 | 0.342278752 | 0.068329796 | 5.009216625 | 5.47E-07 | 1.62E-05 |
| ACSL1 | 4133.283368 | -0.485349717 | 0.097039929 | -5.001546497 | 5.69E-07 | 1.68E-05 |
| ADA | 119.3660912 | -0.834595258 | 0.167458317 | -4.983898507 | 6.23E-07 | 1.84E-05 |

| | | | | | | |
|---|---|---|---|---|---|---|
| SLC7A5 | 350.8088739 | 0.38440025 | 0.077229061 | 4.977404144 | 6.44E-07 | 1.90E-05 |
| HEXB | 3329.189183 | -0.339740357 | 0.068344807 | -4.970975449 | 6.66E-07 | 1.96E-05 |
| PNP | 1197.557229 | 0.405731679 | 0.081663254 | 4.968350611 | 6.75E-07 | 1.98E-05 |
| ADD3 | 79.98967939 | -0.458089115 | 0.092445004 | -4.955260886 | 7.22E-07 | 2.12E-05 |
| KCNK13 | 36.05918329 | 0.708620777 | 0.143098352 | 4.951984182 | 7.35E-07 | 2.15E-05 |
| CABLES1 | 205.4604406 | -0.584442834 | 0.118098154 | -4.948788905 | 7.47E-07 | 2.18E-05 |
| RFTN1 | 287.209683 | 0.35540323 | 0.07182432 | 4.948229638 | 7.49E-07 | 2.19E-05 |
| PSMB9 | 1403.814993 | 0.728954379 | 0.147358812 | 4.946798693 | 7.54E-07 | 2.20E-05 |
| MLLT6 | 1085.979488 | 0.291734226 | 0.059094333 | 4.936754726 | 7.94E-07 | 2.31E-05 |
| IRF5 | 2241.903136 | 0.307269094 | 0.062270621 | 4.934415091 | 8.04E-07 | 2.34E-05 |
| NLRP12 | 22.55840237 | -0.898678633 | 0.182428918 | -4.926185184 | 8.39E-07 | 2.43E-05 |
| TSPAN33 | 59.51910694 | -0.579047534 | 0.117628444 | -4.922682913 | 8.54E-07 | 2.47E-05 |
| MBNL2 | 800.7498483 | -0.400551364 | 0.08152021 | -4.913522234 | 8.95E-07 | 2.59E-05 |
| DPH3 | 651.2895312 | 0.248395706 | 0.050562982 | 4.912599966 | 8.99E-07 | 2.60E-05 |
| LOC64383[illegible] | 156.6024116 | 0.393513157 | 0.080117176 | 4.911720246 | 9.03E-07 | 2.60E-05 |
| CSF3R | 155.6499593 | -0.555323042 | 0.113076024 | -4.911059146 | 9.06E-07 | 2.61E-05 |
| TEX30 | 186.1223676 | 0.514177735 | 0.104727369 | 4.909678743 | 9.12E-07 | 2.62E-05 |
| CD44 | 595.3180082 | 0.468115679 | 0.095351259 | 4.909381196 | 9.14E-07 | 2.62E-05 |
| FXR2 | 1004.375592 | 0.239784814 | 0.048845417 | 4.909054509 | 9.15E-07 | 2.62E-05 |
| CSF2RB | 2669.151328 | 0.391143688 | 0.079721425 | 4.906381033 | 9.28E-07 | 2.66E-05 |
| CXADR | 105.613092 | -0.972529153 | 0.198346728 | -4.903177206 | 9.43E-07 | 2.70E-05 |
| TPCN1 | 43.33202687 | -1.033943415 | 0.211052943 | -4.898976526 | 9.63E-07 | 2.75E-05 |
| ATP2A2 | 1756.578916 | 0.502631253 | 0.102608221 | 4.898547607 | 9.65E-07 | 2.75E-05 |
| TNIP1 | 6284.699141 | 0.36906947 | 0.075428269 | 4.89298609 | 9.93E-07 | 2.83E-05 |
| AFF4 | 1044.136453 | 0.289075847 | 0.059125457 | 4.88919426 | 1.01E-06 | 2.87E-05 |
| RAP2C | 778.8758177 | 0.432765825 | 0.088521951 | 4.888796703 | 1.01E-06 | 2.87E-05 |
| TIMP2 | 5291.759892 | -0.441016882 | 0.090204804 | -4.889062048 | 1.01E-06 | 2.87E-05 |
| KLHDC2 | 286.366289 | -0.338603471 | 0.069289756 | -4.886775345 | 1.03E-06 | 2.90E-05 |
| FAM162A | 354.6295597 | -0.254773967 | 0.052146552 | -4.885729845 | 1.03E-06 | 2.91E-05 |
| NME3 | 164.7748754 | -0.536475179 | 0.10993785 | -4.87980416 | 1.06E-06 | 3.00E-05 |
| SLC50A1 | 671.4420881 | 0.300130059 | 0.061520794 | 4.878514074 | 1.07E-06 | 3.01E-05 |
| TRAPPC6A | 73.2817117 | -0.577546766 | 0.118393618 | -4.878191718 | 1.07E-06 | 3.01E-05 |
| DNAJB9 | 398.1817454 | 0.450154877 | 0.092359619 | 4.873936064 | 1.09E-06 | 3.07E-05 |
| SORT1 | 2968.227292 | 0.508711796 | 0.104405126 | 4.872479137 | 1.10E-06 | 3.09E-05 |
| MSRB2 | 228.1041061 | -0.483135099 | 0.099186861 | -4.870958632 | 1.11E-06 | 3.11E-05 |
| PVR | 1023.195105 | 0.464544957 | 0.095403263 | 4.869277431 | 1.12E-06 | 3.13E-05 |
| PARP14 | 1692.57793 | 0.41960048 | 0.086267214 | 4.863962321 | 1.15E-06 | 3.21E-05 |
| GFPT2 | 46.93570779 | 1.073308595 | 0.22083849 | 4.860151845 | 1.17E-06 | 3.27E-05 |
| ADAMTS1[illegible] | 44.64490709 | -0.958115674 | 0.197519493 | -4.850739836 | 1.23E-06 | 3.42E-05 |
| DDX60 | 331.1640984 | 0.582690451 | 0.12014975 | 4.849701749 | 1.24E-06 | 3.44E-05 |
| ZNF652 | 109.6457945 | -0.622591621 | 0.128476511 | -4.84595678 | 1.26E-06 | 3.50E-05 |
| MX1 | 1934.447958 | 0.794123067 | 0.163982849 | 4.842720273 | 1.28E-06 | 3.55E-05 |
| A2M | 476.2690543 | -0.462283459 | 0.095499401 | -4.84069488 | 1.29E-06 | 3.58E-05 |
| C7orf57 | 6.628351815 | 2.17305441 | 0.448923645 | 4.840588005 | 1.29E-06 | 3.58E-05 |
| EYA3 | 685.9706968 | 0.492576413 | 0.101883926 | 4.834682274 | 1.33E-06 | 3.68E-05 |
| NSUN3 | 335.7069125 | 0.288795486 | 0.059889346 | 4.822151307 | 1.42E-06 | 3.91E-05 |
| HPS5 | 1232.650397 | 0.255172693 | 0.052925907 | 4.821319277 | 1.43E-06 | 3.93E-05 |

| | | | | | | |
|---|---|---|---|---|---|---|
| C5orf15 | 1029.356603 | 0.385348344 | 0.079962524 | 4.819111784 | 1.44E-06 | 3.96E-05 |
| SCCPDH | 1427.133405 | -0.381762479 | 0.079222782 | -4.818847141 | 1.44E-06 | 3.96E-05 |
| #N/A | 1131.204124 | 0.472785516 | 0.098213818 | 4.813839086 | 1.48E-06 | 4.06E-05 |
| TGFBI | 2216.808555 | -0.526782305 | 0.109490718 | -4.811205123 | 1.50E-06 | 4.11E-05 |
| POLR3D | 275.1218008 | 0.378732468 | 0.078727987 | 4.81064591 | 1.50E-06 | 4.11E-05 |
| FAM110A | 108.5369112 | -0.43316353 | 0.090050088 | -4.810251073 | 1.51E-06 | 4.11E-05 |
| LNPEP | 53.0883327 | 0.86649185 | 0.180271578 | 4.806591596 | 1.54E-06 | 4.18E-05 |
| CLSTN3 | 74.82286959 | 0.529248483 | 0.110141764 | 4.805157131 | 1.55E-06 | 4.20E-05 |
| RPS6KA3 | 1718.14273 | 0.382586907 | 0.079619078 | 4.805216494 | 1.55E-06 | 4.20E-05 |
| IFI27 | 1792.601199 | 0.492947975 | 0.102732125 | 4.798381961 | 1.60E-06 | 4.34E-05 |
| TK2 | 272.4907558 | -0.337727207 | 0.070411966 | -4.79644618 | 1.62E-06 | 4.38E-05 |
| OAS2 | 631.7737248 | 0.881944802 | 0.183986735 | 4.793523838 | 1.64E-06 | 4.43E-05 |
| NOV | 12.51545495 | -1.409971635 | 0.29447817 | -4.788034499 | 1.68E-06 | 4.55E-05 |
| ATL1 | 98.34285566 | 0.528915358 | 0.110526849 | 4.785401579 | 1.71E-06 | 4.60E-05 |
| PHLDA3 | 390.5677799 | -0.46389134 | 0.097055985 | -4.779626342 | 1.76E-06 | 4.73E-05 |
| NDRG1 | 1314.47178 | 0.387723344 | 0.081137678 | 4.778585637 | 1.77E-06 | 4.75E-05 |
| ITGAM | 568.4355328 | 0.543595689 | 0.113786202 | 4.777342755 | 1.78E-06 | 4.76E-05 |
| NPC1 | 2717.269325 | 0.387795994 | 0.081171986 | 4.777460935 | 1.78E-06 | 4.76E-05 |
| PGM5 | 101.571914 | -0.708262604 | 0.148252795 | -4.777397971 | 1.78E-06 | 4.76E-05 |
| HAMP | 17.68375148 | -1.211596259 | 0.253639375 | -4.776846097 | 1.78E-06 | 4.76E-05 |
| FAM214A | 125.1634609 | -0.414703681 | 0.086834008 | -4.775821031 | 1.79E-06 | 4.78E-05 |
| TMTC3 | 194.5040885 | 0.433344044 | 0.090771095 | 4.774031234 | 1.81E-06 | 4.82E-05 |
| BRD3 | 337.9185263 | -0.33951315 | 0.071174405 | -4.770157928 | 1.84E-06 | 4.91E-05 |
| CERS6 | 897.3877508 | 0.244913382 | 0.051357485 | 4.768796252 | 1.85E-06 | 4.93E-05 |
| CXCL11 | 149.7027114 | 2.209001708 | 0.4633564 | 4.767392247 | 1.87E-06 | 4.96E-05 |
| OXER1 | 52.53258244 | -0.753714503 | 0.158144452 | -4.765987626 | 1.88E-06 | 4.99E-05 |
| GNA15 | 1671.940942 | 0.391814954 | 0.082232432 | 4.764725336 | 1.89E-06 | 5.01E-05 |
| STAU1 | 243.80211 | 0.487542523 | 0.102332579 | 4.764294276 | 1.90E-06 | 5.02E-05 |
| TCEB2 | 3250.961023 | -0.347874374 | 0.073025582 | -4.763733002 | 1.90E-06 | 5.02E-05 |
| MFNG | 367.5225441 | -0.374469402 | 0.078673197 | -4.759809147 | 1.94E-06 | 5.11E-05 |
| BATF3 | 95.41726773 | 0.738083735 | 0.155093447 | 4.758961431 | 1.95E-06 | 5.13E-05 |
| ICK | 142.2306772 | -0.318028269 | 0.066830615 | -4.758721255 | 1.95E-06 | 5.13E-05 |
| SFPQ | 1345.479006 | 0.272111985 | 0.057215786 | 4.755889976 | 1.98E-06 | 5.19E-05 |
| ELL | 173.9854743 | 0.432622729 | 0.091018664 | 4.753121051 | 2.00E-06 | 5.25E-05 |
| VAMP5 | 146.2152356 | 1.029684799 | 0.216628288 | 4.753233332 | 2.00E-06 | 5.25E-05 |
| SERTAD1 | 478.6481213 | 0.381172429 | 0.080213551 | 4.751970499 | 2.01E-06 | 5.26E-05 |
| TMEM223 | 94.08480905 | -0.497049235 | 0.104590286 | -4.752346075 | 2.01E-06 | 5.26E-05 |
| ZNF552 | 59.70831547 | -0.622984407 | 0.131100622 | -4.751956156 | 2.01E-06 | 5.26E-05 |
| BTN2A2 | 335.5581369 | 0.485110336 | 0.102094306 | 4.751590519 | 2.02E-06 | 5.26E-05 |
| SPTSSA | 354.3493038 | -0.28085427 | 0.059140959 | -4.748896138 | 2.05E-06 | 5.32E-05 |
| RPL23 | 6896.028424 | -0.250079446 | 0.052751602 | -4.74069863 | 2.13E-06 | 5.54E-05 |
| CDC37 | 1552.448952 | 0.339695052 | 0.071704062 | 4.737459002 | 2.16E-06 | 5.62E-05 |
| LIMK1 | 5259.325024 | 0.254181749 | 0.053663204 | 4.73661145 | 2.17E-06 | 5.64E-05 |
| RAP2A | 1921.810986 | 0.34618031 | 0.073174563 | 4.73088318 | 2.24E-06 | 5.79E-05 |
| C1orf233 | 53.51675429 | -0.842737298 | 0.178214702 | -4.728775387 | 2.26E-06 | 5.84E-05 |
| DDX21 | 496.9603015 | 0.408266969 | 0.086339644 | 4.728615406 | 2.26E-06 | 5.84E-05 |
| EPB41 | 279.2172158 | -0.392373334 | 0.083012045 | -4.726703644 | 2.28E-06 | 5.89E-05 |

| | | | | | | |
|---|---|---|---|---|---|---|
| SOWAHD | 170.8361787 | -0.449306318 | 0.095103362 | -4.724399945 | 2.31E-06 | 5.94E-05 |
| ARAP3 | 88.12429198 | 0.620810896 | 0.131421671 | 4.723809174 | 2.31E-06 | 5.95E-05 |
| FLVCR2 | 2071.076343 | 0.617071647 | 0.130664447 | 4.722567333 | 2.33E-06 | 5.98E-05 |
| HADH | 168.406961 | -0.516643667 | 0.109501124 | -4.718158583 | 2.38E-06 | 6.11E-05 |
| PCED1B-AS | 167.5749752 | -0.394796243 | 0.083688882 | -4.717427632 | 2.39E-06 | 6.12E-05 |
| BZW1 | 3858.040014 | 0.262113392 | 0.05557661 | 4.71625368 | 2.40E-06 | 6.15E-05 |
| DUSP1 | 3386.56126 | 0.4180785 | 0.088658852 | 4.71558669 | 2.41E-06 | 6.16E-05 |
| LPP | 372.1820126 | 0.424745206 | 0.090082521 | 4.715067936 | 2.42E-06 | 6.17E-05 |
| ERGIC2 | 1125.247885 | 0.293073444 | 0.062199833 | 4.711804378 | 2.46E-06 | 6.25E-05 |
| GTF2IRD2E | 80.91413138 | -0.437516725 | 0.092854072 | -4.711874403 | 2.45E-06 | 6.25E-05 |
| S100A8 | 143.3934268 | -0.889732616 | 0.188845577 | -4.711429463 | 2.46E-06 | 6.25E-05 |
| FRAT1 | 99.19170236 | -0.606583648 | 0.128777027 | -4.710340525 | 2.47E-06 | 6.28E-05 |
| ECH1 | 1204.489599 | -0.37296686 | 0.079197738 | -4.709312055 | 2.49E-06 | 6.29E-05 |
| METTL1 | 464.7654202 | 0.457680916 | 0.097183154 | 4.709467635 | 2.48E-06 | 6.29E-05 |
| SPHK1 | 1090.807783 | 0.357811489 | 0.076028973 | 4.706251765 | 2.52E-06 | 6.38E-05 |
| RBP4 | 3057.319981 | -0.362910177 | 0.077121965 | -4.705665581 | 2.53E-06 | 6.39E-05 |
| MRI1 | 122.6604284 | -0.511955994 | 0.108854877 | -4.70310573 | 2.56E-06 | 6.46E-05 |
| BBS2 | 668.9773197 | -0.39753921 | 0.084544898 | -4.702107622 | 2.57E-06 | 6.48E-05 |
| CTSK | 23.8890295 | -0.857130066 | 0.182426614 | -4.698492438 | 2.62E-06 | 6.59E-05 |
| SLC35B4 | 268.8925482 | 0.516369764 | 0.109968955 | 4.695595818 | 2.66E-06 | 6.68E-05 |
| LOC28597 | 24.85057112 | 1.434648652 | 0.305555105 | 4.695220698 | 2.66E-06 | 6.68E-05 |
| SLC35E3 | 614.6724203 | -0.336780257 | 0.071738808 | -4.694533771 | 2.67E-06 | 6.69E-05 |
| RNF141 | 111.416284 | -0.458174375 | 0.097817525 | -4.683970239 | 2.81E-06 | 7.04E-05 |
| IVNS1ABP | 1535.089382 | 0.406868849 | 0.086869759 | 4.683664997 | 2.82E-06 | 7.04E-05 |
| PLA1A | 34.73254377 | 0.972788532 | 0.207722747 | 4.68311029 | 2.83E-06 | 7.05E-05 |
| STAT2 | 1274.770242 | 0.467582839 | 0.099906451 | 4.68020669 | 2.87E-06 | 7.14E-05 |
| MYO1G | 160.1808302 | 0.538952768 | 0.115233504 | 4.677049206 | 2.91E-06 | 7.24E-05 |
| IGFBP3 | 19.17631277 | 1.375556804 | 0.294187448 | 4.675783464 | 2.93E-06 | 7.28E-05 |
| PINK1 | 462.1317752 | -0.390819453 | 0.083594721 | -4.675169065 | 2.94E-06 | 7.29E-05 |
| OTUD4 | 634.9188233 | 0.344208878 | 0.073630385 | 4.674821092 | 2.94E-06 | 7.30E-05 |
| B3GNT2 | 235.5970275 | -0.289590297 | 0.062013616 | -4.669785693 | 3.02E-06 | 7.47E-05 |
| SGMS1 | 1225.959535 | 0.292852514 | 0.06275191 | 4.666830261 | 3.06E-06 | 7.56E-05 |
| SLC39A14 | 48.46953894 | 0.865046119 | 0.185407977 | 4.665635926 | 3.08E-06 | 7.60E-05 |
| DCAF4 | 97.38862924 | -0.463683436 | 0.099448548 | -4.662546051 | 3.12E-06 | 7.70E-05 |
| ZNF763 | 33.66395726 | -0.606839939 | 0.130180712 | -4.661519583 | 3.14E-06 | 7.73E-05 |
| LOXHD1 | 7.073772321 | 2.020308116 | 0.43366435 | 4.658690792 | 3.18E-06 | 7.83E-05 |
| CTNNBIP1 | 110.9451617 | -0.520895204 | 0.11191068 | -4.654562045 | 3.25E-06 | 7.97E-05 |
| XYLT2 | 187.1249365 | -0.44960957 | 0.096591952 | -4.654731193 | 3.24E-06 | 7.97E-05 |
| DNM2 | 1477.598997 | 0.325988795 | 0.070083039 | 4.65146492 | 3.30E-06 | 8.08E-05 |
| ADAM19 | 21.88408528 | 1.104673148 | 0.237532722 | 4.650614611 | 3.31E-06 | 8.10E-05 |
| MDM4 | 363.5208388 | 0.348345192 | 0.074941689 | 4.648216457 | 3.35E-06 | 8.19E-05 |
| THYN1 | 81.40438903 | -0.505428672 | 0.108775005 | -4.646551587 | 3.38E-06 | 8.24E-05 |
| SLC24A4 | 27.78064021 | -1.027594146 | 0.221175983 | -4.64604761 | 3.38E-06 | 8.25E-05 |
| STOX2 | 16.49843946 | -1.121873846 | 0.241711006 | -4.641385037 | 3.46E-06 | 8.43E-05 |
| IFITM1 | 377.5448165 | 0.758343633 | 0.163426464 | 4.640274382 | 3.48E-06 | 8.46E-05 |
| CBX7 | 66.60411163 | -0.838476902 | 0.180757335 | -4.638688124 | 3.51E-06 | 8.52E-05 |
| DTX3L | 1009.305957 | 0.396070532 | 0.085430731 | 4.636159912 | 3.55E-06 | 8.61E-05 |

| | | | | | | |
|---|---|---|---|---|---|---|
| PIP5K1C | 1143.230444 | 0.353421941 | 0.076258428 | 4.63452959 | 3.58E-06 | 8.67E-05 |
| TUBB6 | 2003.660607 | 0.321167116 | 0.069370856 | 4.629712466 | 3.66E-06 | 8.86E-05 |
| PTPN22 | 468.8537879 | -0.383903863 | 0.083029182 | -4.623722085 | 3.77E-06 | 9.11E-05 |
| FAM168A | 275.753335 | -0.453003345 | 0.097993419 | -4.622793569 | 3.79E-06 | 9.14E-05 |
| FOSL1 | 145.5620925 | 0.473535256 | 0.10265225 | 4.613004179 | 3.97E-06 | 9.57E-05 |
| PLEK | 3241.240683 | 0.460186422 | 0.099972985 | 4.603107745 | 4.16E-06 | 0.000100239 |
| GDF15 | 597.1930823 | 0.594277615 | 0.129140206 | 4.601801657 | 4.19E-06 | 0.000100744 |
| MB21D1 | 424.3960701 | 0.474445439 | 0.103218399 | 4.596520047 | 4.30E-06 | 0.000103202 |
| MNDA | 766.6641386 | 0.534884145 | 0.116390023 | 4.595618505 | 4.31E-06 | 0.00010352 |
| RASAL1 | 18.10169786 | -1.020405057 | 0.222109827 | -4.594146374 | 4.35E-06 | 0.000104124 |
| RNPEPL1 | 1834.790934 | -0.324407063 | 0.070628935 | -4.593118408 | 4.37E-06 | 0.000104508 |
| EPSTI1 | 415.3518199 | 0.86832185 | 0.189076482 | 4.592437108 | 4.38E-06 | 0.00010472 |
| HARS | 300.1110359 | 0.297472282 | 0.064814115 | 4.589621926 | 4.44E-06 | 0.000106011 |
| SPIN4 | 290.5769648 | 0.599779331 | 0.130799882 | 4.58547303 | 4.53E-06 | 0.000108004 |
| CTSD | 53732.99392 | -0.292306192 | 0.063791081 | -4.582242338 | 4.60E-06 | 0.000109551 |
| TP53INP1 | 31.46138814 | -0.74782928 | 0.163213916 | -4.581896564 | 4.61E-06 | 0.000109597 |
| GALNT12 | 1189.392322 | -0.306691781 | 0.066993271 | -4.577949021 | 4.70E-06 | 0.000111547 |
| MYO1E | 1131.723483 | 0.283839745 | 0.06201006 | 4.577317711 | 4.71E-06 | 0.000111747 |
| CXCR7 | 17.00324674 | -1.386304579 | 0.303241237 | -4.571622887 | 4.84E-06 | 0.000114686 |
| WARS | 10213.07273 | 0.604723804 | 0.132374158 | 4.568291977 | 4.92E-06 | 0.000116381 |
| TET2 | 325.005386 | 0.329953858 | 0.07225484 | 4.566529487 | 4.96E-06 | 0.000117219 |
| ANPEP | 8882.763053 | 0.359083235 | 0.078649636 | 4.565605802 | 4.98E-06 | 0.000117592 |
| LOC10012[illegible] | 3322.862722 | 0.485927989 | 0.106445502 | 4.565040145 | 4.99E-06 | 0.000117765 |
| ISG15 | 774.2453415 | 0.742990387 | 0.162822814 | 4.563183556 | 5.04E-06 | 0.000118667 |
| C21orf91 | 181.3144083 | 0.46527941 | 0.101995951 | 4.561743915 | 5.07E-06 | 0.000119337 |
| ABHD12 | 1690.992386 | -0.335130568 | 0.073513633 | -4.558754005 | 5.15E-06 | 0.000120901 |
| RAB39A | 158.4191835 | 0.424172065 | 0.09313321 | 4.554466306 | 5.25E-06 | 0.000123243 |
| CMSS1 | 134.4961629 | -0.414763678 | 0.091098502 | -4.552914352 | 5.29E-06 | 0.000124005 |
| GRAMD1A | 1907.888088 | 0.455302641 | 0.100016811 | 4.552261127 | 5.31E-06 | 0.00012424 |
| TSPAN13 | 102.860968 | -0.696088146 | 0.152957964 | -4.550846045 | 5.34E-06 | 0.000124927 |
| CLCN4 | 127.463055 | -0.568473104 | 0.125085254 | -4.544685202 | 5.50E-06 | 0.000128481 |
| GRN | 12358.46678 | -0.446175468 | 0.098187415 | -4.544120752 | 5.52E-06 | 0.000128669 |
| TANC2 | 383.9950374 | -0.40413954 | 0.088985807 | -4.541617981 | 5.58E-06 | 0.000130049 |
| C17orf96 | 23.75433384 | 0.973056459 | 0.21428814 | 4.540878746 | 5.60E-06 | 0.000130349 |
| PLEKHM3 | 1290.616251 | 0.24451204 | 0.053887545 | 4.537449941 | 5.69E-06 | 0.000132325 |
| MYO5A | 1735.792087 | -0.257362122 | 0.056726248 | -4.536914257 | 5.71E-06 | 0.000132343 |
| TNFSF10 | 93.01260559 | 1.329133519 | 0.292953425 | 4.537013067 | 5.71E-06 | 0.000132343 |
| ZYG11B | 213.555178 | -0.347910119 | 0.076714083 | -4.535153181 | 5.76E-06 | 0.000133291 |
| CLEC2D | 147.5314436 | 0.825075859 | 0.182341637 | 4.524890054 | 6.04E-06 | 0.000139758 |
| DESI1 | 809.175795 | 0.484326471 | 0.107104534 | 4.521997829 | 6.13E-06 | 0.000141512 |
| ABCC5 | 112.6521512 | -0.431946199 | 0.095648205 | -4.515988536 | 6.30E-06 | 0.000145411 |
| RNF130 | 6080.357408 | -0.378387058 | 0.083800111 | -4.515352685 | 6.32E-06 | 0.000145674 |
| NDUFV2 | 800.1824546 | 0.601709124 | 0.133300697 | 4.51392332 | 6.36E-06 | 0.000146484 |
| CASP1 | 946.327503 | 0.536443266 | 0.118905371 | 4.511514157 | 6.44E-06 | 0.000147805 |
| STIP1 | 4252.333154 | 0.241764916 | 0.053587331 | 4.511605858 | 6.43E-06 | 0.000147805 |
| MS4A6E | 17.68525588 | -0.963836669 | 0.21374789 | -4.509221919 | 6.51E-06 | 0.000149233 |
| FKBP5 | 1214.323911 | 0.473313595 | 0.104994893 | 4.507967796 | 6.55E-06 | 0.000149904 |

| | | | | | | |
|---|---|---|---|---|---|---|
| UXS1 | 1842.050131 | 0.273561906 | 0.06068681 | 4.507765449 | 6.55E-06 | 0.000149904 |
| ME3 | 38.04566864 | -0.636176953 | 0.141146959 | -4.507195593 | 6.57E-06 | 0.000150129 |
| S1PR1 | 9.30155066 | -1.550574319 | 0.344404565 | -4.502188642 | 6.73E-06 | 0.000153528 |
| BCS1L | 99.145986 | -0.384732583 | 0.085516521 | -4.498926997 | 6.83E-06 | 0.000155718 |
| CLDN1 | 40.87231792 | 0.805909538 | 0.179190892 | 4.497491635 | 6.88E-06 | 0.000156587 |
| SECTM1 | 301.3366837 | 0.899924944 | 0.200153362 | 4.496176999 | 6.92E-06 | 0.000157186 |
| XAF1 | 143.3650222 | 0.878568126 | 0.195398079 | 4.496298689 | 6.91E-06 | 0.000157186 |
| NAMPT | 8044.64053 | 0.361722152 | 0.080492952 | 4.493836324 | 7.00E-06 | 0.000158738 |
| GSDMD | 1027.88199 | 0.411637323 | 0.091653496 | 4.49123426 | 7.08E-06 | 0.000160478 |
| TNS1 | 153.6528039 | -0.730662804 | 0.162694411 | -4.491013552 | 7.09E-06 | 0.000160478 |
| CLEC1A | 31.72050067 | 0.935482714 | 0.208351364 | 4.489928438 | 7.12E-06 | 0.000161109 |
| HIST1H1D | 192.5716761 | -0.386444434 | 0.086113403 | -4.487622361 | 7.20E-06 | 0.000162671 |
| CD22 | 81.35018381 | -0.539608649 | 0.120261449 | -4.486962791 | 7.22E-06 | 0.000162984 |
| AIF1 | 869.0312957 | -0.493826301 | 0.110125356 | -4.484219805 | 7.32E-06 | 0.000164902 |
| ZNF276 | 144.4160793 | 0.39843854 | 0.088962305 | 4.47873444 | 7.51E-06 | 0.000168998 |
| SERPINF2 | 37.46665712 | -0.577224878 | 0.12902735 | -4.473662953 | 7.69E-06 | 0.000172857 |
| SAMD8 | 198.0236792 | 0.391287616 | 0.087564137 | 4.46858302 | 7.87E-06 | 0.000176806 |
| GPX1 | 8796.179202 | -0.363652417 | 0.081446716 | -4.464911969 | 8.01E-06 | 0.000179655 |
| WIPI1 | 231.0091751 | -0.326167591 | 0.07308683 | -4.462740998 | 8.09E-06 | 0.000181275 |
| SLC30A7 | 938.8878328 | 0.483188969 | 0.108441977 | 4.455737378 | 8.36E-06 | 0.000187078 |
| IRF1 | 1319.116199 | 0.662942035 | 0.148836996 | 4.454148174 | 8.42E-06 | 0.00018825 |
| TSTD3 | 9.086655264 | -1.094486931 | 0.245938428 | -4.450247721 | 8.58E-06 | 0.00019148 |
| DYNLT1 | 875.411469 | 0.450536789 | 0.10128272 | 4.448308565 | 8.65E-06 | 0.000192993 |
| MMP14 | 8255.932036 | 0.221300297 | 0.049758768 | 4.447463375 | 8.69E-06 | 0.000193529 |
| IL1RAP | 79.25260858 | 0.884063573 | 0.198813108 | 4.446706663 | 8.72E-06 | 0.000193988 |
| KLF11 | 633.7455927 | -0.256706835 | 0.05774732 | -4.445346281 | 8.78E-06 | 0.000194994 |
| CPT1B | 170.5105202 | 0.411524747 | 0.092581822 | 4.444984336 | 8.79E-06 | 0.000195098 |
| DOCK10 | 579.6929988 | 0.317548799 | 0.071459544 | 4.443756284 | 8.84E-06 | 0.000195989 |
| DNAJC18 | 36.61502749 | 0.746467152 | 0.168067383 | 4.441475438 | 8.93E-06 | 0.000197851 |
| LPCAT1 | 617.0198323 | -0.310192186 | 0.069954907 | -4.434173381 | 9.24E-06 | 0.000204441 |
| LRG1 | 32.15614857 | 0.851001484 | 0.192038093 | 4.431420207 | 9.36E-06 | 0.000206832 |
| TOR1B | 788.8507747 | 0.397928526 | 0.089880275 | 4.427317633 | 9.54E-06 | 0.000210563 |
| MCL1 | 3511.618081 | 0.270579399 | 0.061123321 | 4.426778396 | 9.57E-06 | 0.000210848 |
| CLLU1OS | 96.34426902 | 1.19773288 | 0.271106498 | 4.41794235 | 9.96E-06 | 0.000219401 |
| ARFGAP3 | 1045.63191 | 0.327554703 | 0.074149468 | 4.41749229 | 9.99E-06 | 0.000219607 |
| HMGB2 | 337.5102778 | -0.382428882 | 0.086602402 | -4.415915422 | 1.01E-05 | 0.000220962 |
| C10orf54 | 141.2365568 | -0.484674133 | 0.109784143 | -4.414791795 | 1.01E-05 | 0.00022186 |
| KIF3C | 25.92559011 | -0.727752239 | 0.164910499 | -4.413013376 | 1.02E-05 | 0.000223429 |
| SLC18B1 | 26.2885797 | -0.599551461 | 0.135867215 | -4.412775097 | 1.02E-05 | 0.000223429 |
| AMICA1 | 40.17970967 | -0.925493158 | 0.210044378 | -4.406179139 | 1.05E-05 | 0.000230077 |
| LRRC45 | 83.36242162 | -0.499218386 | 0.113324456 | -4.405213187 | 1.06E-05 | 0.000230842 |
| FAM216A | 7.185021212 | -1.0971275 | 0.249081907 | -4.404685651 | 1.06E-05 | 0.000231143 |
| PWWP2B | 88.70477462 | -0.494421572 | 0.112266545 | -4.403997375 | 1.06E-05 | 0.000231615 |
| HADHA | 2483.003464 | -0.189444464 | 0.043048737 | -4.400697349 | 1.08E-05 | 0.000234901 |
| TNFRSF10A | 204.0741069 | 0.364149531 | 0.082770061 | 4.399531967 | 1.08E-05 | 0.000235899 |
| LGR4 | 12.6341925 | -0.901062256 | 0.20490487 | -4.397466283 | 1.10E-05 | 0.000237886 |
| RASSF8 | 257.5504622 | -0.269050255 | 0.061227014 | -4.394306294 | 1.11E-05 | 0.000241101 |

| | | | | | | |
|---|---|---|---|---|---|---|
| VWA7 | 30.41994525 | -0.604033735 | 0.137474053 | -4.393801764 | 1.11E-05 | 0.000241389 |
| CAMP | 93.06657405 | -0.576299988 | 0.131177287 | -4.393290937 | 1.12E-05 | 0.000241686 |
| SLAMF7 | 5431.856251 | 0.592460402 | 0.134863576 | 4.393034934 | 1.12E-05 | 0.000241699 |
| PKD2 | 330.1275487 | -0.401863089 | 0.091501025 | -4.391897148 | 1.12E-05 | 0.000242695 |
| PNPLA1 | 81.23318981 | 0.866249765 | 0.197250185 | 4.391629679 | 1.13E-05 | 0.000242722 |
| OSGIN2 | 973.5311731 | 0.323407062 | 0.073749806 | 4.38519202 | 1.16E-05 | 0.000249733 |
| KIAA0247 | 2029.976356 | 0.235463687 | 0.053748009 | 4.38088202 | 1.18E-05 | 0.00025444 |
| FCRLA | 16.84028029 | 1.232499734 | 0.281371992 | 4.380321318 | 1.19E-05 | 0.000254811 |
| SPARC | 2264.546515 | -0.461967543 | 0.105571843 | -4.375859415 | 1.21E-05 | 0.00025979 |
| STAT3 | 1772.353987 | 0.302029478 | 0.069049437 | 4.374104845 | 1.22E-05 | 0.000261597 |
| MAPRE3 | 30.74918206 | 1.029611699 | 0.235531064 | 4.371447585 | 1.23E-05 | 0.000264507 |
| HIVEP2 | 104.2676799 | 0.603462309 | 0.138142296 | 4.368410872 | 1.25E-05 | 0.000267913 |
| MFSD3 | 75.12722744 | -0.521335279 | 0.119357908 | -4.367831894 | 1.25E-05 | 0.000268326 |
| CCRL2 | 2645.468964 | 0.37246908 | 0.085354072 | 4.363811492 | 1.28E-05 | 0.000273004 |
| TMEM217 | 14.41863269 | 1.06597794 | 0.244312552 | 4.363173044 | 1.28E-05 | 0.000273499 |
| ACBD4 | 52.10906905 | -0.500268008 | 0.114665209 | -4.362857835 | 1.28E-05 | 0.000273591 |
| PKD2L1 | 17.41570401 | -1.274447117 | 0.292195948 | -4.361618031 | 1.29E-05 | 0.000274842 |
| H2AFZ | 3002.593197 | -0.218960585 | 0.050209968 | -4.360898723 | 1.30E-05 | 0.000275443 |
| LOC10050! | 52.88216066 | -0.564668679 | 0.129521362 | -4.359656758 | 1.30E-05 | 0.000276705 |
| MTSS1 | 62.95068121 | 0.556355081 | 0.127676016 | 4.35755357 | 1.32E-05 | 0.000279069 |
| TMEM167I | 768.697298 | 0.210093493 | 0.048231592 | 4.355931162 | 1.33E-05 | 0.000280835 |
| PTPRE | 2970.997911 | 0.387967587 | 0.089076026 | 4.355465851 | 1.33E-05 | 0.000281123 |
| HTR7 | 3.691382812 | 2.099950323 | 0.482358501 | 4.353505367 | 1.34E-05 | 0.000283339 |
| CRIP1 | 372.6910359 | -0.401530052 | 0.092265867 | -4.351880757 | 1.35E-05 | 0.000285134 |
| CTSL1 | 37553.32514 | 0.266657695 | 0.06131002 | 4.34933307 | 1.37E-05 | 0.000288151 |
| DDX60L | 2150.373112 | 0.440937692 | 0.101451544 | 4.346288615 | 1.38E-05 | 0.000291857 |
| EIF1AY | 168.2860919 | 0.384812195 | 0.088554554 | 4.345481722 | 1.39E-05 | 0.000292612 |
| SOWAHC | 540.9053893 | 0.398595152 | 0.09181463 | 4.341303261 | 1.42E-05 | 0.000297908 |
| CRTAP | 833.4674057 | -0.402215328 | 0.092724286 | -4.337756018 | 1.44E-05 | 0.000301767 |
| PIP5K1A | 204.995635 | 0.300539648 | 0.069282652 | 4.337877369 | 1.44E-05 | 0.000301767 |
| TRIB1 | 717.0514742 | 0.433132283 | 0.099850733 | 4.337797736 | 1.44E-05 | 0.000301767 |
| C3orf52 | 21.72206545 | 0.752616848 | 0.173521843 | 4.337303221 | 1.44E-05 | 0.000302061 |
| INPP5D | 236.634875 | -0.42859103 | 0.098823112 | -4.336951366 | 1.44E-05 | 0.000302216 |
| STRN3 | 495.2042534 | 0.301262096 | 0.069467886 | 4.336710276 | 1.45E-05 | 0.000302219 |
| TFCP2L1 | 173.5493367 | -0.366063173 | 0.084418058 | -4.336313581 | 1.45E-05 | 0.000302437 |
| FDXR | 32.90533927 | -0.513010288 | 0.118459794 | -4.330670105 | 1.49E-05 | 0.000309958 |
| ARHGAP9 | 274.1416435 | 0.401764776 | 0.092817944 | 4.328524843 | 1.50E-05 | 0.000312316 |
| IL24 | 9.851646527 | 1.977969344 | 0.456946313 | 4.328669005 | 1.50E-05 | 0.000312316 |
| SERPINB6 | 1099.864157 | -0.20546339 | 0.047488403 | -4.326601334 | 1.51E-05 | 0.000314715 |
| SEMA4B | 196.6569697 | 0.38723405 | 0.089544826 | 4.324471515 | 1.53E-05 | 0.000317428 |
| HSPA5 | 6765.582214 | 0.2990253 | 0.069152977 | 4.324113163 | 1.53E-05 | 0.000317602 |
| FAM102B | 420.8265638 | -0.335643873 | 0.077638113 | -4.323184315 | 1.54E-05 | 0.0003186 |
| FAM26F | 73.37588688 | 1.405575916 | 0.325334351 | 4.320404262 | 1.56E-05 | 0.000322294 |
| IL27 | 34.39352108 | 1.010674595 | 0.23398891 | 4.319326908 | 1.57E-05 | 0.000323524 |
| MSI2 | 40.75860992 | -0.50991444 | 0.118071991 | -4.318674002 | 1.57E-05 | 0.000324134 |
| ANTXR2 | 190.3979487 | 0.463495502 | 0.107425887 | 4.314560641 | 1.60E-05 | 0.000329875 |
| SLC11A2 | 1178.009261 | 0.412635336 | 0.095705379 | 4.311516649 | 1.62E-05 | 0.000334092 |

| | | | | | | |
|---|---|---|---|---|---|---|
| TBC1D8 | 317.4634065 | -0.475229786 | 0.110264598 | -4.309903586 | 1.63E-05 | 0.000336179 |
| ASTL | 62.843944 | 0.726212658 | 0.168544016 | 4.308741861 | 1.64E-05 | 0.000337229 |
| LRMP | 39.98255133 | -0.887460292 | 0.205963459 | -4.30882398 | 1.64E-05 | 0.000337229 |
| FEM1C | 151.2279987 | 0.319532769 | 0.074199435 | 4.306404364 | 1.66E-05 | 0.000340447 |
| SEMA6B | 2366.822054 | -0.577585396 | 0.134225655 | -4.303092415 | 1.68E-05 | 0.000345211 |
| CD300A | 317.027019 | 0.415136185 | 0.09651746 | 4.301151163 | 1.70E-05 | 0.000347879 |
| KIF16B | 399.9112615 | -0.305378384 | 0.071015977 | -4.300136357 | 1.71E-05 | 0.000349105 |
| PSAP | 77407.70055 | -0.266411072 | 0.061997964 | -4.297093898 | 1.73E-05 | 0.000353528 |
| STOM | 4206.911811 | 0.289611195 | 0.067400414 | 4.296875628 | 1.73E-05 | 0.000353528 |
| ZFAND5 | 4379.023501 | 0.324797973 | 0.075604301 | 4.296025084 | 1.74E-05 | 0.000354511 |
| SQRDL | 2092.955301 | 0.282167874 | 0.065703561 | 4.294559869 | 1.75E-05 | 0.000356483 |
| NUS1 | 1772.467491 | 0.391180808 | 0.091100317 | 4.293956617 | 1.76E-05 | 0.000357076 |
| LOC40139[illegible] | 794.4169302 | 0.31137221 | 0.072557011 | 4.291414538 | 1.78E-05 | 0.000360808 |
| GPR85 | 76.08422728 | 0.667730636 | 0.155638395 | 4.290269364 | 1.78E-05 | 0.000362291 |
| GNGT2 | 30.83306178 | -0.99115656 | 0.231106071 | -4.288751729 | 1.80E-05 | 0.000364391 |
| HIF1A | 2210.59553 | -0.249928317 | 0.058298878 | -4.28701761 | 1.81E-05 | 0.00036686 |
| ADM | 1454.872909 | 0.547160902 | 0.127663584 | 4.285959132 | 1.82E-05 | 0.000368224 |
| SSPN | 22.1569046 | -1.106469501 | 0.258322021 | -4.283295313 | 1.84E-05 | 0.000372271 |
| NRP1 | 4303.183349 | 0.278455966 | 0.065023253 | 4.282405978 | 1.85E-05 | 0.000373371 |
| ACP6 | 95.34546798 | -0.580804226 | 0.135712397 | -4.279669654 | 1.87E-05 | 0.000377595 |
| RPS6KA2 | 475.2754318 | -0.382154459 | 0.089308177 | -4.279053394 | 1.88E-05 | 0.000378246 |
| TTC4 | 1052.940885 | 0.328973943 | 0.076930514 | 4.276247845 | 1.90E-05 | 0.000382642 |
| VCAN | 71.80134858 | 0.987159889 | 0.230972583 | 4.273926706 | 1.92E-05 | 0.000386245 |
| PLBD2 | 1610.248573 | -0.348519496 | 0.081557228 | -4.273312166 | 1.93E-05 | 0.000386908 |
| SCN1B | 29.90494398 | 1.035588528 | 0.242401595 | 4.272201794 | 1.94E-05 | 0.000388435 |
| ITGB8 | 4079.486615 | 0.488569759 | 0.114418387 | 4.270028376 | 1.95E-05 | 0.000391832 |
| CTBS | 504.3465843 | -0.234115462 | 0.054868389 | -4.266855013 | 1.98E-05 | 0.000397032 |
| FAM19A4 | 13.61623197 | -0.857519224 | 0.200996884 | -4.266330926 | 1.99E-05 | 0.000397552 |
| NTPCR | 126.6402398 | -0.276756818 | 0.064901956 | -4.264229232 | 2.01E-05 | 0.000400896 |
| MCOLN1 | 2246.847273 | 0.267684773 | 0.062852229 | 4.258954358 | 2.05E-05 | 0.000410047 |
| CORO6 | 10.24172468 | -1.211844727 | 0.284727674 | -4.25615364 | 2.08E-05 | 0.000414785 |
| ACOT13 | 1538.957492 | -0.318331213 | 0.074818099 | -4.25473535 | 2.09E-05 | 0.000416991 |
| TOMM34 | 379.6180255 | 0.285477588 | 0.067182785 | 4.249266974 | 2.14E-05 | 0.00042686 |
| CUTA | 442.0547365 | -0.239742128 | 0.056431042 | -4.248408708 | 2.15E-05 | 0.000428056 |
| GPRC5B | 28.97182991 | 0.885270345 | 0.208630962 | 4.243235697 | 2.20E-05 | 0.000437596 |
| TTYH3 | 1141.320896 | -0.29510912 | 0.069562061 | -4.242386079 | 2.21E-05 | 0.000438804 |
| MICA | 255.9578088 | 0.3203199 | 0.075562752 | 4.239124341 | 2.24E-05 | 0.00044477 |
| HMGN4 | 188.2577324 | -0.336505322 | 0.079494287 | -4.233075519 | 2.31E-05 | 0.000456438 |
| LOC28307[illegible] | 169.3099999 | -0.45034682 | 0.106440488 | -4.230972898 | 2.33E-05 | 0.000459781 |
| TRIM56 | 139.7162745 | 0.350881032 | 0.082929367 | 4.231082986 | 2.33E-05 | 0.000459781 |
| CD99 | 3059.361282 | -0.143894742 | 0.034018725 | -4.229868718 | 2.34E-05 | 0.00046157 |
| CES1 | 23342.11392 | -0.252480692 | 0.059733852 | -4.226760602 | 2.37E-05 | 0.000467032 |
| LPCAT2 | 1163.494105 | -0.210721465 | 0.049853717 | -4.226795466 | 2.37E-05 | 0.000467032 |
| CPE | 466.0978349 | -0.396758878 | 0.093957296 | -4.222757516 | 2.41E-05 | 0.000474922 |
| DST | 767.788455 | -0.195182524 | 0.046224224 | -4.222515965 | 2.42E-05 | 0.000474947 |
| ELMSAN1 | 261.3562136 | -0.246837325 | 0.058485396 | -4.22049508 | 2.44E-05 | 0.000478551 |
| RTN4R | 27.56986658 | -0.930570053 | 0.220495798 | -4.220352775 | 2.44E-05 | 0.000478551 |

| | | | | | | |
|---|---|---|---|---|---|---|
| FPR1 | 115.4887873 | -0.793472162 | 0.188173879 | -4.216696632 | 2.48E-05 | 0.000485667 |
| SNTA1 | 52.09768379 | -0.4972319 | 0.117923442 | -4.216565361 | 2.48E-05 | 0.000485667 |
| NTHL1 | 19.73768206 | -0.800408738 | 0.19003015 | -4.212009193 | 2.53E-05 | 0.000495066 |
| BLVRB | 1737.825008 | -0.281602144 | 0.066906651 | -4.208881166 | 2.57E-05 | 0.00050146 |
| APBB1IP | 174.3315688 | -0.4356293 | 0.103723968 | -4.199890427 | 2.67E-05 | 0.000521264 |
| C2orf71 | 10.25972748 | 1.523796105 | 0.362994487 | 4.197849163 | 2.69E-05 | 0.000525451 |
| NPL | 1413.056031 | -0.313608129 | 0.074722025 | -4.196997187 | 2.70E-05 | 0.000526897 |
| MACF1 | 983.4592382 | 0.228512881 | 0.054471325 | 4.195104123 | 2.73E-05 | 0.000530781 |
| SETD7 | 473.2457986 | -0.246454894 | 0.058770207 | -4.193534526 | 2.75E-05 | 0.000533929 |
| CDA | 52.7870993 | -0.601148142 | 0.143393181 | -4.192306344 | 2.76E-05 | 0.000536288 |
| NBEAL2 | 193.0017536 | -0.533910619 | 0.127381108 | -4.191442728 | 2.77E-05 | 0.000537791 |
| RHOC | 3597.640899 | 0.295221403 | 0.070445201 | 4.190795095 | 2.78E-05 | 0.000538786 |
| MTMR9 | 263.7503533 | 0.291876618 | 0.069656202 | 4.190245922 | 2.79E-05 | 0.000539549 |
| MRAS | 1309.333959 | 0.287924833 | 0.068716967 | 4.190010798 | 2.79E-05 | 0.000539566 |
| NDUFAF3 | 1190.529021 | -0.186272958 | 0.044467132 | -4.18900321 | 2.80E-05 | 0.000541424 |
| OR6K6 | 43.89441193 | -0.888096378 | 0.212060783 | -4.187933121 | 2.82E-05 | 0.000543437 |
| ALDH7A1 | 29.86174732 | -0.752116725 | 0.179644257 | -4.186700641 | 2.83E-05 | 0.000545848 |
| ZNF207 | 1421.773558 | 0.273958525 | 0.065452846 | 4.185586137 | 2.84E-05 | 0.000547985 |
| NT5C3L | 316.7547073 | -0.255618328 | 0.061143157 | -4.180653103 | 2.91E-05 | 0.000559451 |
| CALHM2 | 150.4620612 | -0.490473891 | 0.117377597 | -4.178598834 | 2.93E-05 | 0.000563964 |
| EPHX1 | 2292.184946 | -0.33207185 | 0.079474262 | -4.178357158 | 2.94E-05 | 0.000564001 |
| HECW2 | 6.351545872 | 1.489425763 | 0.356591199 | 4.176843865 | 2.96E-05 | 0.000567199 |
| IFIH1 | 456.613956 | 0.606394393 | 0.145204384 | 4.176143831 | 2.96E-05 | 0.00056838 |
| FMNL3 | 3174.249917 | 0.279723138 | 0.067000295 | 4.174953806 | 2.98E-05 | 0.000570792 |
| IRF2BPL | 797.1642331 | 0.291977736 | 0.070003977 | 4.170873564 | 3.03E-05 | 0.000580533 |
| STARD10 | 268.6752807 | -0.291538954 | 0.069995491 | -4.165110499 | 3.11E-05 | 0.000594803 |
| NCK1 | 468.7182594 | -0.379240079 | 0.091134293 | -4.161332323 | 3.16E-05 | 0.000604132 |
| SAC3D1 | 16.96040343 | -0.942080615 | 0.226410096 | -4.160947902 | 3.17E-05 | 0.000604551 |
| MPZL2 | 47.13010966 | -0.930669979 | 0.223747809 | -4.15945964 | 3.19E-05 | 0.000607901 |
| HIST1H4E | 585.5568304 | -0.356555392 | 0.085771582 | -4.157034095 | 3.22E-05 | 0.000613782 |
| DUSP23 | 644.9963291 | -0.347587721 | 0.08362546 | -4.156482023 | 3.23E-05 | 0.000614659 |
| PDE6G | 116.520702 | -0.553178619 | 0.13316765 | -4.154001507 | 3.27E-05 | 0.000620751 |
| ATPAF1 | 147.804691 | -0.272657775 | 0.065731181 | -4.148073603 | 3.35E-05 | 0.000636413 |
| CTNNBL1 | 965.2875882 | 0.388991407 | 0.093790997 | 4.147427986 | 3.36E-05 | 0.000636955 |
| SLC46A2 | 37.60240993 | -0.885310375 | 0.213458562 | -4.147457783 | 3.36E-05 | 0.000636955 |
| TNFSF13 | 293.5258559 | 0.791787935 | 0.191045806 | 4.144492628 | 3.41E-05 | 0.000644535 |
| FAM50B | 19.14101433 | -0.73168579 | 0.176564995 | -4.144002546 | 3.41E-05 | 0.00064528 |
| FNDC3A | 1205.353513 | 0.344952305 | 0.083283107 | 4.141924055 | 3.44E-05 | 0.000650519 |
| MARCKS | 531.3187144 | 0.50734949 | 0.122621445 | 4.137526582 | 3.51E-05 | 0.000662459 |
| NCR3LG1 | 63.03768361 | 0.483878985 | 0.117004003 | 4.135576326 | 3.54E-05 | 0.000667457 |
| SLC43A3 | 4973.351608 | 0.372757854 | 0.090161624 | 4.134329391 | 3.56E-05 | 0.000670435 |
| PSMA3 | 204.3672659 | 0.453451624 | 0.109783764 | 4.130406989 | 3.62E-05 | 0.000681312 |
| GOLT1B | 840.8014681 | 0.276162544 | 0.066888798 | 4.128681502 | 3.65E-05 | 0.000685774 |
| H2AFY | 2330.626323 | -0.375606092 | 0.090980154 | -4.12843982 | 3.65E-05 | 0.000685826 |
| OAS3 | 1015.577093 | 0.616340314 | 0.149495691 | 4.122796516 | 3.74E-05 | 0.000702168 |
| CXCL6 | 12.21635332 | 2.106798884 | 0.511199286 | 4.121286825 | 3.77E-05 | 0.000706099 |
| H2AFV | 1353.017307 | -0.27588512 | 0.066950732 | -4.120718473 | 3.78E-05 | 0.000707155 |

| | | | | | | |
|---|---|---|---|---|---|---|
| B3GNT1 | 18.63958512 | -0.726738285 | 0.176443539 | -4.118814933 | 3.81E-05 | 0.000712329 |
| CD163 | 2239.654622 | -0.255068045 | 0.06197495 | -4.115663615 | 3.86E-05 | 0.000721432 |
| PPP2R3A | 257.1783535 | 0.288507396 | 0.070124201 | 4.114234355 | 3.88E-05 | 0.000725213 |
| LAMP3 | 82.48546006 | 0.721123731 | 0.175400139 | 4.1113065 | 3.93E-05 | 0.000733761 |
| ALCAM | 2597.735879 | 0.26683379 | 0.064912715 | 4.11065522 | 3.95E-05 | 0.000735123 |
| GBP4 | 714.1789824 | 0.786818636 | 0.19147972 | 4.109148663 | 3.97E-05 | 0.000739219 |
| NAB2 | 269.7528134 | 0.465085322 | 0.113279173 | 4.105656054 | 4.03E-05 | 0.000749754 |
| NUDT6 | 34.46777325 | -0.717360648 | 0.174752654 | -4.105005744 | 4.04E-05 | 0.000751141 |
| CHST14 | 54.98078971 | -0.61068766 | 0.148861771 | -4.102380722 | 4.09E-05 | 0.000758985 |
| PEPD | 2798.683946 | -0.2992452 | 0.072961176 | -4.101430574 | 4.11E-05 | 0.000761375 |
| RAB43 | 495.081231 | 0.285700821 | 0.069698606 | 4.099089443 | 4.15E-05 | 0.000768378 |
| MYO1B | 13.348382 | 1.306498392 | 0.319072306 | 4.094678128 | 4.23E-05 | 0.000782402 |
| SPRY2 | 185.2202282 | 0.469450568 | 0.114740257 | 4.091419869 | 4.29E-05 | 0.000792717 |
| RNF125 | 26.72786278 | -0.770725351 | 0.188420336 | -4.090457372 | 4.31E-05 | 0.00079525 |
| ARHGEF16 | 11.68702428 | -0.970556736 | 0.237361001 | -4.088947776 | 4.33E-05 | 0.000798911 |
| GPBP1 | 699.9604591 | 0.18263424 | 0.044664232 | 4.089049113 | 4.33E-05 | 0.000798911 |
| YWHAZ | 334.7568555 | 0.306372601 | 0.074954534 | 4.087445854 | 4.36E-05 | 0.00080333 |
| BAK1 | 729.8834563 | 0.356531934 | 0.087236608 | 4.086953214 | 4.37E-05 | 0.000804268 |
| CMPK2 | 464.1374418 | 0.779939833 | 0.190866902 | 4.086302147 | 4.38E-05 | 0.000805758 |
| TMEM194/ | 130.2291701 | 0.956953509 | 0.234259047 | 4.085022629 | 4.41E-05 | 0.000809439 |
| IRF8 | 1651.031709 | 0.310302053 | 0.076097597 | 4.07768529 | 4.55E-05 | 0.000834608 |
| FSCN1 | 248.2402893 | 0.39500576 | 0.096915163 | 4.075789045 | 4.59E-05 | 0.000840639 |
| RASSF2 | 42.9167357 | -0.781124168 | 0.191832998 | -4.071896787 | 4.66E-05 | 0.000854001 |
| ST14 | 779.2351456 | -0.289444816 | 0.071179077 | -4.066431148 | 4.77E-05 | 0.000873439 |
| CKAP2 | 51.94066209 | -0.476899713 | 0.117377635 | -4.062952142 | 4.85E-05 | 0.000884884 |
| THBD | 1595.542256 | 0.595721698 | 0.146622545 | 4.062961095 | 4.85E-05 | 0.000884884 |
| FTO | 654.140407 | -0.229050034 | 0.056383843 | -4.062334532 | 4.86E-05 | 0.000885603 |
| MECR | 39.50214188 | -0.488492735 | 0.120256209 | -4.062099896 | 4.86E-05 | 0.000885603 |
| SLC29A1 | 351.5845776 | -0.46841071 | 0.115307548 | -4.06227275 | 4.86E-05 | 0.000885603 |
| O3FAR1 | 148.5069262 | -0.612149739 | 0.150722981 | -4.061422713 | 4.88E-05 | 0.000887338 |
| SNTB1 | 986.276622 | -0.365732853 | 0.090093189 | -4.059495009 | 4.92E-05 | 0.000893851 |
| CCDC28B | 18.74379599 | -0.874061549 | 0.215428454 | -4.057317085 | 4.96E-05 | 0.000901374 |
| CAMKK2 | 152.4097277 | -0.477100849 | 0.117667557 | -4.05465077 | 5.02E-05 | 0.000910858 |
| STARD4 | 57.38179792 | 0.497684252 | 0.12287495 | 4.050331271 | 5.11E-05 | 0.000926964 |
| CASC4 | 521.9261623 | -0.286211957 | 0.070677513 | -4.049547639 | 5.13E-05 | 0.000928429 |
| DUSP7 | 558.8602546 | 0.398628096 | 0.098438308 | 4.049522024 | 5.13E-05 | 0.000928429 |
| AKR1C3 | 1230.809764 | -0.452963417 | 0.111869169 | -4.049046068 | 5.14E-05 | 0.000929446 |
| SLC25A19 | 625.7810515 | -0.254937421 | 0.063021488 | -4.045245992 | 5.23E-05 | 0.000943767 |
| PER1 | 319.6684856 | 0.372211122 | 0.092034459 | 4.044258267 | 5.25E-05 | 0.000946867 |
| NUP188 | 1660.701471 | 0.366135769 | 0.090545452 | 4.043668221 | 5.26E-05 | 0.000948366 |
| TCEAL4 | 401.9577282 | -0.331166001 | 0.081910657 | -4.043014829 | 5.28E-05 | 0.000950125 |
| SPAG9 | 1947.208075 | 0.211243891 | 0.05228506 | 4.040234246 | 5.34E-05 | 0.000960563 |
| CNIH4 | 1033.082467 | 0.217742609 | 0.053896558 | 4.040009555 | 5.34E-05 | 0.000960587 |
| UBE2L6 | 949.7464022 | 0.50421486 | 0.124857573 | 4.038320197 | 5.38E-05 | 0.000966628 |
| DEPDC1B | 10.76139636 | 1.078660595 | 0.267187128 | 4.037097912 | 5.41E-05 | 0.000970771 |
| MANF | 1282.386287 | 0.357176787 | 0.08853221 | 4.034427555 | 5.47E-05 | 0.000980963 |
| RPL30 | 7025.292765 | -0.202828829 | 0.05028432 | -4.0336397 | 5.49E-05 | 0.000982432 |

| | | | | | | |
|---|---|---|---|---|---|---|
| TMEM150I | 1994.700158 | 0.348723206 | 0.086451114 | 4.03376187 | 5.49E-05 | 0.000982432 |
| LDLRAD4 | 17.20209468 | -1.018344547 | 0.252531868 | -4.032538757 | 5.52E-05 | 0.000986132 |
| IFIT2 | 634.8932803 | 0.66758958 | 0.165616274 | 4.030941918 | 5.56E-05 | 0.000991936 |
| IFIT5 | 207.6108774 | 0.597458354 | 0.148258639 | 4.029838378 | 5.58E-05 | 0.000995681 |
| GDPD1 | 9.807915337 | -1.024926916 | 0.254352774 | -4.029548806 | 5.59E-05 | 0.000995986 |
| ANKZF1 | 101.6915201 | -0.323216962 | 0.080262784 | -4.026984197 | 5.65E-05 | 0.001005235 |
| SRCAP | 798.5924641 | 0.225899435 | 0.056097035 | 4.026940719 | 5.65E-05 | 0.001005235 |
| PLAGL1 | 214.4899393 | 0.434091137 | 0.107807895 | 4.026524576 | 5.66E-05 | 0.001006086 |
| HAGHL | 51.05557083 | -0.471447557 | 0.117129042 | -4.025027006 | 5.70E-05 | 0.001011576 |
| GPA33 | 42.59990477 | -0.711952506 | 0.176926501 | -4.024001505 | 5.72E-05 | 0.001015058 |
| FAM53B | 203.498413 | -0.404392113 | 0.100537462 | -4.022302769 | 5.76E-05 | 0.001020483 |
| FDX1 | 3604.601859 | -0.248242847 | 0.061719754 | -4.022097156 | 5.77E-05 | 0.001020483 |
| IFIT1 | 146.6493269 | 0.98254252 | 0.244265286 | 4.022440256 | 5.76E-05 | 0.001020483 |
| WIPI2 | 1075.096092 | -0.243163918 | 0.060460813 | -4.021843316 | 5.77E-05 | 0.001020647 |
| CHCHD10 | 431.9885781 | -0.294745415 | 0.073298445 | -4.021168742 | 5.79E-05 | 0.001022636 |
| ETV7 | 11.27622318 | 1.952644971 | 0.486082436 | 4.017106621 | 5.89E-05 | 0.001039465 |
| ZNF792 | 38.91008005 | -0.490190349 | 0.122036707 | -4.016745135 | 5.90E-05 | 0.001040108 |
| KLHDC3 | 652.5416986 | -0.355921665 | 0.088756979 | -4.010069634 | 6.07E-05 | 0.001068978 |
| IDUA | 134.9005344 | -0.415905409 | 0.103726668 | -4.009628546 | 6.08E-05 | 0.001069999 |
| IQGAP3 | 72.44187793 | -0.454481477 | 0.113365535 | -4.008991609 | 6.10E-05 | 0.00107191 |
| MFSD1 | 2058.834027 | -0.276553378 | 0.069005807 | -4.007682687 | 6.13E-05 | 0.001076884 |
| NLRP2 | 127.5640112 | 0.696700149 | 0.173857371 | 4.007308665 | 6.14E-05 | 0.001077608 |
| MRPL43 | 536.2321474 | -0.231109874 | 0.057681567 | -4.006650391 | 6.16E-05 | 0.001079631 |
| ARHGAP5 | 293.9473142 | 0.322773474 | 0.080563929 | 4.006426691 | 6.16E-05 | 0.001079671 |
| IFITM3 | 411.8585409 | 0.670076468 | 0.167290073 | 4.005476569 | 6.19E-05 | 0.001083036 |
| DAXX | 1040.058692 | 0.331992161 | 0.082892359 | 4.005099659 | 6.20E-05 | 0.00108378 |
| SIGLEC7 | 43.61106184 | -0.585923797 | 0.14648942 | -3.999768697 | 6.34E-05 | 0.001107479 |
| BAZ1A | 420.6924476 | 0.346267155 | 0.086630075 | 3.997077864 | 6.41E-05 | 0.001119124 |
| FICD | 359.471088 | 0.313812238 | 0.078558847 | 3.994613591 | 6.48E-05 | 0.001129804 |
| AIFM3 | 27.85822408 | -0.697992811 | 0.174823536 | -3.992556302 | 6.54E-05 | 0.001138624 |
| LY86 | 919.0040534 | -0.398087691 | 0.099921261 | -3.98401388 | 6.78E-05 | 0.001179297 |
| HLA-DOA | 586.2919111 | 0.474026869 | 0.119039014 | 3.982113516 | 6.83E-05 | 0.001187696 |
| HLTF | 144.8883437 | -0.481897744 | 0.121023596 | -3.981849486 | 6.84E-05 | 0.001187944 |
| FAM102A | 40.41973581 | -0.487112991 | 0.122504013 | -3.976302335 | 7.00E-05 | 0.001214888 |
| PLA2G4A | 166.6003537 | 0.429393875 | 0.10806428 | 3.973504225 | 7.08E-05 | 0.001227776 |
| ZCCHC24 | 78.68407365 | -0.602921601 | 0.1517409 | -3.973362497 | 7.09E-05 | 0.001227776 |
| ZNF385A | 914.3926461 | -0.333802755 | 0.084040564 | -3.971924239 | 7.13E-05 | 0.001234104 |
| SNRPA | 626.8182749 | 0.296408008 | 0.074682055 | 3.968932164 | 7.22E-05 | 0.001248579 |
| TLR8 | 325.5808942 | -0.543913242 | 0.137094094 | -3.967444738 | 7.26E-05 | 0.001255268 |
| ADCY1 | 7.78367136 | -1.545018894 | 0.389576728 | -3.965891144 | 7.31E-05 | 0.001260659 |
| SLC46A3 | 322.8124918 | -0.538303827 | 0.135737045 | -3.965784181 | 7.32E-05 | 0.001260659 |
| USP30 | 211.0159926 | -0.391094625 | 0.098617272 | -3.965782245 | 7.32E-05 | 0.001260659 |
| CDK2AP1 | 461.3663737 | -0.355313333 | 0.089609942 | -3.96511061 | 7.34E-05 | 0.001263084 |
| CD69 | 58.0982609 | 0.818001084 | 0.206416343 | 3.962869769 | 7.41E-05 | 0.001273864 |
| PSMA2 | 1164.243037 | 0.367516553 | 0.092760112 | 3.962010663 | 7.43E-05 | 0.001277317 |
| ACAA1 | 594.4269968 | -0.353455218 | 0.089288836 | -3.958560052 | 7.54E-05 | 0.001294541 |
| QSOX1 | 3119.056606 | 0.25924315 | 0.065492126 | 3.958386514 | 7.55E-05 | 0.001294541 |

| | | | | | | |
|---|---|---|---|---|---|---|
| TRIM36 | 121.286506 | 0.380207375 | 0.096079797 | 3.957204182 | 7.58E-05 | 0.001299806 |
| E2F7 | 2.87938941 | 2.145914426 | 0.542393542 | 3.956379015 | 7.61E-05 | 0.001303142 |
| HIVEP3 | 128.1528945 | 0.454045193 | 0.114855353 | 3.953191383 | 7.71E-05 | 0.001319459 |
| TMTC1 | 119.8627126 | 0.474899747 | 0.120163501 | 3.952113109 | 7.75E-05 | 0.001324244 |
| RARG | 89.80370301 | 0.507564902 | 0.128463872 | 3.951032259 | 7.78E-05 | 0.001329061 |
| OLFML2B | 19.36570263 | -0.88207027 | 0.223320422 | -3.949796726 | 7.82E-05 | 0.001334757 |
| NTN1 | 5.03784693 | 1.888378931 | 0.478283295 | 3.948243544 | 7.87E-05 | 0.001342255 |
| PROCR | 220.955792 | 0.403357592 | 0.102168835 | 3.94795138 | 7.88E-05 | 0.001342706 |
| GAB1 | 155.8318622 | -0.276716657 | 0.070101294 | -3.947382992 | 7.90E-05 | 0.001344707 |
| TERF2IP | 693.9212588 | 0.186320878 | 0.047228504 | 3.945093771 | 7.98E-05 | 0.001356421 |
| MAOB | 7.420048451 | 1.922223658 | 0.487815191 | 3.940475191 | 8.13E-05 | 0.001381586 |
| ARL6IP4 | 380.0583446 | -0.33067894 | 0.083941082 | -3.93941718 | 8.17E-05 | 0.001385249 |
| UACA | 298.6546993 | -0.274506044 | 0.06967906 | -3.93957731 | 8.16E-05 | 0.001385249 |
| ABCA7 | 15.07906674 | -0.971143426 | 0.24657395 | -3.938548353 | 8.20E-05 | 0.001389051 |
| COQ10B | 685.5143187 | 0.293797182 | 0.074613662 | 3.937578916 | 8.23E-05 | 0.001393448 |
| CTNNAL1 | 44.56289796 | 0.679396075 | 0.172701613 | 3.933930111 | 8.36E-05 | 0.001413541 |
| CCDC106 | 95.38699874 | -0.607942482 | 0.154566272 | -3.933215664 | 8.38E-05 | 0.001415265 |
| SEMA7A | 124.8388581 | 0.391025768 | 0.099413658 | 3.933320372 | 8.38E-05 | 0.001415265 |
| APMAP | 637.0410486 | -0.301419809 | 0.076652232 | -3.932303094 | 8.41E-05 | 0.001418165 |
| DDX58 | 869.6807848 | 0.506009778 | 0.128677867 | 3.93237618 | 8.41E-05 | 0.001418165 |
| PTPN12 | 3705.261445 | 0.400005167 | 0.101805437 | 3.929114003 | 8.53E-05 | 0.001435846 |
| C5orf4 | 6.747829316 | -1.193236849 | 0.30396041 | -3.925632447 | 8.65E-05 | 0.001454547 |
| SLCO4A1 | 14.03449825 | 0.965853031 | 0.246040826 | 3.925580343 | 8.65E-05 | 0.001454547 |
| TMEM106/ | 564.4767052 | 0.480797352 | 0.122493012 | 3.925100244 | 8.67E-05 | 0.001456181 |
| SNHG5 | 583.3094074 | -0.363159229 | 0.092538926 | -3.924394239 | 8.69E-05 | 0.001459185 |
| ADAR | 4006.941164 | 0.260876508 | 0.066483729 | 3.923915086 | 8.71E-05 | 0.00146082 |
| NLN | 569.1408838 | -0.331771559 | 0.084557689 | -3.923611954 | 8.72E-05 | 0.001461388 |
| TNFRSF12/ | 385.5799564 | 0.338294816 | 0.086236792 | 3.922859477 | 8.75E-05 | 0.001464688 |
| AVIL | 22.10070736 | 0.872186422 | 0.222347019 | 3.92263601 | 8.76E-05 | 0.001464775 |
| FBXW5 | 1020.016241 | -0.223198492 | 0.056913965 | -3.921682354 | 8.79E-05 | 0.00146931 |
| GUSB | 516.179774 | -0.361455562 | 0.092193288 | -3.920627746 | 8.83E-05 | 0.001474477 |
| FLJ39639 | 9.956720502 | -1.179147051 | 0.300930596 | -3.91833554 | 8.92E-05 | 0.001487277 |
| ATN1 | 1420.822144 | 0.270464509 | 0.069089188 | 3.914715417 | 9.05E-05 | 0.001508463 |
| NDEL1 | 314.0384189 | 0.391718932 | 0.100100223 | 3.913267329 | 9.11E-05 | 0.00151623 |
| PELO | 602.7016354 | 0.349186898 | 0.089288162 | 3.910786035 | 9.20E-05 | 0.001529249 |
| RNF113A | 151.3736819 | -0.283295497 | 0.072439093 | -3.910809537 | 9.20E-05 | 0.001529249 |
| CD52 | 8775.943444 | -0.337745035 | 0.086464707 | -3.906160658 | 9.38E-05 | 0.00155746 |
| RBP1 | 52.1155459 | -0.80110272 | 0.205206797 | -3.903880054 | 9.47E-05 | 0.001570863 |
| HN1 | 775.9675373 | -0.315340187 | 0.080801422 | -3.902656402 | 9.51E-05 | 0.001576366 |
| SMURF1 | 462.1464674 | 0.292461861 | 0.074939921 | 3.902617667 | 9.52E-05 | 0.001576366 |
| ZDHHC24 | 323.141759 | -0.250224274 | 0.064145206 | -3.900903756 | 9.58E-05 | 0.001586205 |
| TMEM88 | 16.10019146 | 0.89083921 | 0.228435902 | 3.899733804 | 9.63E-05 | 0.001592521 |
| ISCU | 1586.99441 | -0.177249491 | 0.045570169 | -3.889594816 | 0.000100412 | 0.001659119 |
| C1orf53 | 9.857473497 | -1.121834579 | 0.288456718 | -3.889091528 | 0.00010062 | 0.001659716 |
| RAB4B | 404.8393384 | 0.352431189 | 0.090619104 | 3.889148915 | 0.000100596 | 0.001659716 |
| LAMB2 | 70.05531207 | -0.473309931 | 0.121750737 | -3.88753238 | 0.000101268 | 0.001668981 |
| PRPF3 | 67.15816177 | 0.57433472 | 0.147750398 | 3.887195758 | 0.000101409 | 0.001669868 |

| | | | | | | |
|---|---|---|---|---|---|---|
| BAZ2B | 203.9339681 | 0.315329934 | 0.081137358 | 3.886371732 | 0.000101754 | 0.001674113 |
| ZNFX1 | 1023.135651 | 0.376397014 | 0.096899061 | 3.884423763 | 0.000102573 | 0.001686151 |
| CDR2 | 157.1743972 | -0.353033932 | 0.090896815 | -3.883897705 | 0.000102795 | 0.001688365 |
| C19orf24 | 646.5217059 | -0.202729093 | 0.052212783 | -3.882748306 | 0.000103282 | 0.001693481 |
| MLX | 2853.765393 | 0.218826766 | 0.056357121 | 3.882859233 | 0.000103235 | 0.001693481 |
| ATP13A2 | 454.5927613 | -0.493563004 | 0.127129352 | -3.882368597 | 0.000103444 | 0.001694686 |
| DEFB1 | 106.3747105 | -0.469980483 | 0.12118567 | -3.878185298 | 0.000105239 | 0.001721159 |
| IGFBP2 | 793.4410343 | -0.444651395 | 0.114652696 | -3.878246322 | 0.000105212 | 0.001721159 |
| ARL4C | 28.91858618 | -0.959915347 | 0.247579206 | -3.877205043 | 0.000105663 | 0.001726641 |
| UBASH3B | 1492.977266 | -0.407547225 | 0.105125683 | -3.876761736 | 0.000105856 | 0.001728322 |
| GLS | 3425.169499 | 0.255947587 | 0.066060746 | 3.874427735 | 0.000106876 | 0.00174318 |
| LRP12 | 151.2452675 | 0.756134765 | 0.195168557 | 3.874265275 | 0.000106947 | 0.00174318 |
| ACTN1 | 4644.555846 | 0.391188316 | 0.101014621 | 3.872591018 | 0.000107684 | 0.001753718 |
| IGFN1 | 6.764172492 | 1.727211864 | 0.446611993 | 3.867365614 | 0.000110017 | 0.001790199 |
| C14orf1 | 254.4481005 | -0.373791002 | 0.09668848 | -3.865931094 | 0.000110666 | 0.001799236 |
| PBX3 | 469.5137785 | -0.310649591 | 0.080429325 | -3.862392129 | 0.000112282 | 0.00182397 |
| PNRC2 | 3183.886708 | 0.182750283 | 0.047356818 | 3.859006799 | 0.000113849 | 0.001847861 |
| PCYOX1L | 67.29328866 | -0.432130296 | 0.112036149 | -3.85706131 | 0.000114758 | 0.001861056 |
| PFKP | 461.3624438 | 0.366650532 | 0.095066018 | 3.856799102 | 0.000114882 | 0.001861486 |
| CCR1 | 233.2798935 | 0.473358028 | 0.122758838 | 3.85599958 | 0.000115258 | 0.001866012 |
| GABARAPL | 849.6193203 | 0.263126758 | 0.068265682 | 3.85445145 | 0.000115989 | 0.001876282 |
| LOC54147[illegible] | 217.8099076 | 0.629964398 | 0.163649607 | 3.849470898 | 0.000118373 | 0.001913237 |
| VGLL4 | 167.1853063 | -0.345049075 | 0.089738391 | -3.845055267 | 0.000120525 | 0.001946387 |
| RPL21 | 8167.009117 | -0.210468272 | 0.054750313 | -3.844147413 | 0.000120972 | 0.00195197 |
| TRIM62 | 56.70704925 | -0.412936469 | 0.107484392 | -3.841827283 | 0.000122122 | 0.001968868 |
| PLK1S1 | 59.28208319 | -0.55058547 | 0.143345659 | -3.840963693 | 0.000122552 | 0.001974157 |
| RNF168 | 262.1901252 | -0.391548987 | 0.101974235 | -3.839685444 | 0.000123192 | 0.001982805 |
| ST3GAL5 | 62.84892214 | 0.415793322 | 0.108303644 | 3.839144329 | 0.000123464 | 0.001985521 |
| CENPN | 197.7185344 | 0.387565575 | 0.101014426 | 3.836734897 | 0.000124681 | 0.002003422 |
| HSPA13 | 719.727465 | 0.345375496 | 0.090070812 | 3.834488538 | 0.000125826 | 0.002020134 |
| GPX4 | 8236.659091 | -0.249000464 | 0.065061492 | -3.827155777 | 0.000129632 | 0.002079517 |
| DNMT3A | 106.2938893 | 0.484740039 | 0.126782851 | 3.823388068 | 0.00013163 | 0.002109809 |
| GALC | 461.9807995 | -0.357714484 | 0.09367349 | -3.818737647 | 0.000134136 | 0.002148188 |
| NIN | 250.3454908 | -0.337882979 | 0.08862349 | -3.81256684 | 0.000137531 | 0.002197075 |
| RWDD3 | 119.6092136 | -0.322549714 | 0.084600273 | -3.812632057 | 0.000137495 | 0.002197075 |
| USP15 | 440.854851 | 0.332530774 | 0.08721039 | 3.812972 | 0.000137306 | 0.002197075 |
| CPPED1 | 218.1942385 | -0.305780159 | 0.080233666 | -3.811120389 | 0.000138338 | 0.002208142 |
| LINC00032 | 8.812176678 | 1.208438687 | 0.317115578 | 3.810720037 | 0.000138563 | 0.00220989 |
| AK4 | 326.0961391 | 0.397120817 | 0.104267289 | 3.808680741 | 0.00013971 | 0.00222635 |
| TRAF3IP2 | 652.9157663 | 0.305083701 | 0.080131259 | 3.807299469 | 0.000140493 | 0.002236967 |
| LPAR1 | 884.3477387 | 0.384315241 | 0.100991247 | 3.805431194 | 0.000141557 | 0.00225206 |
| S100A4 | 3933.416464 | -0.360772562 | 0.094876283 | -3.802557906 | 0.00014321 | 0.002276468 |
| PSAT1 | 113.2909884 | -0.507270269 | 0.133481029 | -3.800317352 | 0.000144511 | 0.002295257 |
| OTUD1 | 890.6646572 | 0.260362317 | 0.068524488 | 3.799551442 | 0.000144958 | 0.002300465 |
| SEC24D | 803.9923273 | 0.195422488 | 0.05146862 | 3.796924975 | 0.000146502 | 0.002323053 |
| GALM | 76.17230564 | -0.548098132 | 0.144374856 | -3.796354474 | 0.000146839 | 0.002324577 |
| TNFRSF1B | 1267.414128 | 0.330863385 | 0.087150616 | 3.796454919 | 0.00014678 | 0.002324577 |

| | | | | | | |
|---|---|---|---|---|---|---|
| C2orf66 | 7.253831915 | 1.585388807 | 0.417717311 | 3.79536295 | 0.000147428 | 0.002331971 |
| XIRP1 | 6.711631799 | 2.319885819 | 0.611299325 | 3.795007983 | 0.000147639 | 0.002333394 |
| IBTK | 379.7547546 | 0.321609124 | 0.084766071 | 3.794078484 | 0.000148193 | 0.002340232 |
| BAIAP2L1 | 19.46506506 | 0.845081513 | 0.223200778 | 3.786194302 | 0.000152972 | 0.002409779 |
| C10orf11 | 241.687843 | -0.370347113 | 0.097810645 | -3.786368166 | 0.000152865 | 0.002409779 |
| DPH1 | 122.3149935 | -0.407413556 | 0.107596227 | -3.786504115 | 0.000152782 | 0.002409779 |
| ATP13A3 | 4122.438823 | 0.357415303 | 0.094405409 | 3.785962118 | 0.000153115 | 0.00241006 |
| NCOA3 | 830.1000307 | 0.213423674 | 0.056384753 | 3.785130958 | 0.000153628 | 0.002416155 |
| INPP1 | 145.8354897 | 0.389441151 | 0.10290644 | 3.784419622 | 0.000154068 | 0.0024211 |
| FAM173A | 187.1833028 | -0.364761403 | 0.096470553 | -3.781064686 | 0.000156159 | 0.002451965 |
| APRT | 1475.865972 | -0.318515456 | 0.084312319 | -3.777804496 | 0.000158217 | 0.002480434 |
| TAB2 | 1114.419032 | 0.295691939 | 0.078271257 | 3.777784481 | 0.00015823 | 0.002480434 |
| SAR1A | 1145.038148 | 0.178084844 | 0.047163185 | 3.775929118 | 0.000159412 | 0.002496941 |
| EXOSC5 | 138.7180704 | -0.341341604 | 0.090445132 | -3.774018516 | 0.000160639 | 0.002514109 |
| S100A9 | 227.8850875 | -0.453515858 | 0.120365574 | -3.767820347 | 0.000164679 | 0.00257525 |
| C16orf13 | 98.49081988 | -0.469731335 | 0.124719034 | -3.766316322 | 0.000165674 | 0.002588705 |
| NOTCH2NL | 883.4311712 | -0.2636959 | 0.070061917 | -3.763755149 | 0.000167381 | 0.002613256 |
| POPDC3 | 22.45589174 | 1.235288778 | 0.328376217 | 3.761809512 | 0.000168689 | 0.002631541 |
| PLAUR | 21274.85991 | 0.236633393 | 0.062978623 | 3.757360566 | 0.000171715 | 0.002676587 |
| CD55 | 95.67764884 | 0.378517197 | 0.100813056 | 3.754644616 | 0.000173588 | 0.002703588 |
| CCNDBP1 | 1412.913289 | -0.198125628 | 0.052786567 | -3.753334213 | 0.000174498 | 0.002715572 |
| BCOR | 257.9306016 | 0.358486115 | 0.095538195 | 3.752280614 | 0.000175233 | 0.002724812 |
| ATP2B4 | 282.7301759 | -0.360315135 | 0.096062634 | -3.750835474 | 0.000176246 | 0.002738356 |
| RPS23 | 7013.323956 | -0.267917453 | 0.071459741 | -3.749208262 | 0.000177394 | 0.002753963 |
| SDHA | 395.0150954 | -0.228995693 | 0.061098356 | -3.747984538 | 0.000178261 | 0.002765202 |
| MCTP1 | 406.7193304 | 0.298140965 | 0.079569801 | 3.746911033 | 0.000179026 | 0.002774824 |
| FAM126B | 309.6792717 | 0.397357563 | 0.106057445 | 3.746625846 | 0.000179229 | 0.002775746 |
| SLC2A4RG | 133.9468862 | -0.287486985 | 0.076738978 | -3.74629675 | 0.000179464 | 0.002777156 |
| GABARAPL | 1628.281693 | -0.157518557 | 0.042049574 | -3.746020278 | 0.000179662 | 0.002777985 |
| TRAPPC5 | 523.8536174 | -0.274690577 | 0.073420464 | -3.741335317 | 0.000183045 | 0.002828025 |
| PDE4A | 694.9150123 | 0.328728369 | 0.087901307 | 3.739743815 | 0.000184208 | 0.002843709 |
| ZNF134 | 40.30767055 | -0.41615279 | 0.111425081 | -3.734821505 | 0.000187848 | 0.002897587 |
| TMEM14A | 230.1871084 | -0.344173508 | 0.092161903 | -3.734444448 | 0.00018813 | 0.00289961 |
| ZNF808 | 319.5268057 | 0.256590281 | 0.068724166 | 3.733625265 | 0.000188743 | 0.002906737 |
| TWSG1 | 964.800701 | 0.236453751 | 0.06336987 | 3.731327722 | 0.000190473 | 0.002931036 |
| CD72 | 45.05880612 | -0.682961722 | 0.183110211 | -3.729785018 | 0.000191643 | 0.002946686 |
| ADAP1 | 505.8290598 | 0.504034832 | 0.13515283 | 3.729369411 | 0.00019196 | 0.002949197 |
| C16orf88 | 71.07398418 | -0.43911948 | 0.117844304 | -3.72626819 | 0.000194336 | 0.002980948 |
| MAPK6 | 1111.63873 | 0.422080572 | 0.113265699 | 3.726464188 | 0.000194185 | 0.002980948 |
| SHISA4 | 106.2464295 | -0.350932016 | 0.094192901 | -3.725673718 | 0.000194794 | 0.002985605 |
| USP18 | 88.04965744 | 0.793747549 | 0.213152685 | 3.723844941 | 0.000196211 | 0.003004936 |
| SNRK | 142.967135 | 0.326355744 | 0.08764608 | 3.723563498 | 0.00019643 | 0.003005754 |
| TUBE1 | 296.6022718 | 0.383381092 | 0.102966004 | 3.723375468 | 0.000196577 | 0.003005754 |
| HMGXB3 | 382.0557326 | 0.260578568 | 0.070033224 | 3.720784999 | 0.000198604 | 0.003034349 |
| SLMO2 | 1008.506761 | 0.220518729 | 0.05930449 | 3.718415415 | 0.000200476 | 0.003058098 |
| TDG | 580.7224405 | 0.297397747 | 0.079975543 | 3.718608677 | 0.000200323 | 0.003058098 |
| CTDSP1 | 1076.687082 | -0.177005658 | 0.047665664 | -3.713483482 | 0.000204426 | 0.003113413 |

| | | | | | | |
|---|---|---|---|---|---|---|
| TSPYL2 | 242.3759574 | 0.290014039 | 0.078095867 | 3.713564478 | 0.00020436 | 0.003113413 |
| IL1R2 | 5.972537666 | -1.39193332 | 0.374854239 | -3.713265515 | 0.000204602 | 0.003113636 |
| BPHL | 79.9347476 | -0.413017993 | 0.11128789 | -3.711257293 | 0.000206232 | 0.003135969 |
| TSC22D3 | 535.3453512 | -0.285029064 | 0.07681748 | -3.710471438 | 0.000206874 | 0.003143239 |
| STAB1 | 13.0473349 | -1.065933926 | 0.287432872 | -3.708462153 | 0.000208522 | 0.003165785 |
| CCNG2 | 175.1023826 | -0.342783502 | 0.092598086 | -3.7018422 | 0.00021404 | 0.003247 |
| MILR1 | 728.9177876 | -0.239431366 | 0.064704111 | -3.700404249 | 0.000215256 | 0.003262886 |
| JPH4 | 7.957128951 | -1.084377602 | 0.29305909 | -3.700201216 | 0.000215429 | 0.00326293 |
| CDKN2C | 27.79942808 | -0.469342513 | 0.126880606 | -3.699087897 | 0.000216376 | 0.0032747 |
| ZNF420 | 107.148534 | -0.302136411 | 0.081694165 | -3.698384201 | 0.000216976 | 0.003281213 |
| UTP6 | 800.6401333 | 0.18943173 | 0.051250543 | 3.696189707 | 0.000218859 | 0.003307095 |
| ICAM1 | 1840.561485 | 0.683407651 | 0.184915582 | 3.695781852 | 0.000219211 | 0.003309813 |
| SLC16A1 | 276.5121557 | -0.25199391 | 0.06819758 | -3.695056471 | 0.000219838 | 0.003315011 |
| TRERF1 | 70.26219936 | -0.642396119 | 0.173856205 | -3.694985282 | 0.0002199 | 0.003315011 |
| EBF1 | 3.238862711 | 2.166816381 | 0.586460763 | 3.694733762 | 0.000220117 | 0.0033157 |
| GCA | 305.2948882 | 0.443130853 | 0.119964638 | 3.693845617 | 0.000220888 | 0.003324709 |
| RPL38 | 4585.089232 | -0.217200988 | 0.058810432 | -3.693239114 | 0.000221416 | 0.00333005 |
| MGST2 | 504.6560886 | -0.358992471 | 0.097211782 | -3.692890547 | 0.000221719 | 0.003332018 |
| C6orf106 | 2468.519444 | 0.243676468 | 0.066003469 | 3.691873669 | 0.000222608 | 0.003340651 |
| MTA3 | 111.7911258 | -0.338518633 | 0.091693835 | -3.691836355 | 0.000222641 | 0.003340651 |
| SRXN1 | 2698.591394 | 0.279231342 | 0.075640366 | 3.691565173 | 0.000222878 | 0.003341613 |
| GALNT11 | 261.9860425 | -0.314007363 | 0.0850705 | -3.691142808 | 0.000223249 | 0.003344565 |
| C17orf58 | 161.2116346 | 0.423704681 | 0.114845598 | 3.689341949 | 0.000224835 | 0.00336571 |
| ADAM28 | 12.67019477 | -1.245064284 | 0.337641909 | -3.687528861 | 0.000226442 | 0.003387143 |
| SIGLEC10 | 92.14084003 | -0.432167196 | 0.117206086 | -3.687241926 | 0.000226698 | 0.003388332 |
| CXorf38 | 129.4011083 | 0.321565704 | 0.087227573 | 3.686514393 | 0.000227347 | 0.003395395 |
| NUDT9 | 342.0558484 | -0.215547807 | 0.058507746 | -3.684090081 | 0.000229521 | 0.003425215 |
| FBXL15 | 266.9463612 | -0.353213093 | 0.095950497 | -3.681201268 | 0.000232138 | 0.003461582 |
| ABCB1 | 21.97836403 | 0.604671826 | 0.164308123 | 3.680109136 | 0.000233134 | 0.003473753 |
| SEMA3A | 5.106222189 | 1.259409659 | 0.342384567 | 3.678348211 | 0.000234749 | 0.003495116 |
| KIAA0226L | 102.3240656 | 0.559489375 | 0.152161938 | 3.676933817 | 0.000236054 | 0.003511831 |
| GP1BA | 69.16793599 | 0.751936684 | 0.204646951 | 3.674311694 | 0.000238491 | 0.003545352 |
| SNX21 | 72.12524356 | -0.462080614 | 0.125860489 | -3.67137152 | 0.000241252 | 0.00358363 |
| CXCR2 | 11.79522699 | -1.178954031 | 0.321437866 | -3.667750931 | 0.000244693 | 0.003631943 |
| AR | 20.46835547 | 1.172974002 | 0.319962621 | 3.665971977 | 0.000246401 | 0.003654473 |
| PPP1R18 | 2226.524645 | 0.248551075 | 0.067810984 | 3.665351265 | 0.000246999 | 0.00366053 |
| PITPNB | 1150.874093 | 0.194152107 | 0.0529825 | 3.664457247 | 0.000247864 | 0.003670516 |
| CEP350 | 631.2521559 | 0.232196171 | 0.063409275 | 3.661864479 | 0.000250386 | 0.003705025 |
| C1orf122 | 644.881362 | 0.435066619 | 0.118836957 | 3.661038034 | 0.000251195 | 0.003714146 |
| SERPINH1 | 670.657678 | 0.239105925 | 0.06534947 | 3.658880873 | 0.000253319 | 0.003742672 |
| GOLGA2 | 334.4250307 | 0.415886379 | 0.113720116 | 3.657104791 | 0.00025508 | 0.00376292 |
| SESN1 | 172.592292 | -0.354742808 | 0.096996773 | -3.657264035 | 0.000254922 | 0.00376292 |
| AIM2 | 64.74641057 | 0.812478004 | 0.222305476 | 3.654781775 | 0.000257401 | 0.00379425 |
| PEBP1 | 587.8072992 | -0.246501102 | 0.067463202 | -3.653860096 | 0.000258327 | 0.003804992 |
| FBRS | 605.4665111 | 0.311941495 | 0.085439128 | 3.651037903 | 0.000261183 | 0.003844114 |
| PDE4DIP | 2554.186611 | 0.200436624 | 0.054950052 | 3.647614801 | 0.000264686 | 0.003892704 |
| RAPGEF2 | 170.1918866 | 0.48803977 | 0.133824647 | 3.646860134 | 0.000265464 | 0.003901171 |

| | | | | | | |
|---|---|---|---|---|---|---|
| TMEM160 | 214.3260155 | -0.339519509 | 0.093129224 | -3.645681724 | 0.000266684 | 0.003916106 |
| QPCTL | 287.4520862 | -0.21603033 | 0.059272266 | -3.644711851 | 0.000267692 | 0.00392791 |
| CPQ | 433.5092055 | -0.225599541 | 0.061902827 | -3.644414203 | 0.000268002 | 0.003929465 |
| DPP7 | 627.5206442 | -0.31423691 | 0.086248026 | -3.643409873 | 0.00026905 | 0.003941837 |
| LCN2 | 49.68068213 | 1.211477207 | 0.332572426 | 3.642747002 | 0.000269744 | 0.003949002 |
| GFRA2 | 12.88703292 | 1.091229845 | 0.299679379 | 3.64132444 | 0.000271239 | 0.003967875 |
| BTRC | 170.2280174 | 0.233196156 | 0.064070688 | 3.639669888 | 0.000272988 | 0.003990427 |
| VSIG10 | 25.0812842 | -0.556047304 | 0.152862101 | -3.637574651 | 0.000275217 | 0.004019969 |
| BMP6 | 20.13476112 | 0.725842534 | 0.199652056 | 3.635537478 | 0.000277402 | 0.004048803 |
| ZNF530 | 9.24536979 | -0.945773162 | 0.260431517 | -3.631561855 | 0.000281711 | 0.004108589 |
| GBP3 | 545.0684044 | 0.470072223 | 0.12946123 | 3.630988401 | 0.000282338 | 0.004114615 |
| XBP1 | 818.1058031 | 0.283480924 | 0.078110859 | 3.629212723 | 0.000284287 | 0.004139888 |
| ZNF616 | 78.9095605 | -0.309436581 | 0.085270477 | -3.628882952 | 0.00028465 | 0.004142048 |
| RPL26 | 4551.342325 | -0.25113368 | 0.069220535 | -3.628022792 | 0.0002856 | 0.004152733 |
| EIF4EBP3 | 282.9141934 | -0.241559518 | 0.066625805 | -3.62561502 | 0.000288275 | 0.00418846 |
| EPT1 | 323.6962861 | 0.402558267 | 0.111047723 | 3.6250925 | 0.000288858 | 0.004193775 |
| LOC72875 | 26.28806165 | -0.66671026 | 0.183972227 | -3.623972335 | 0.000290113 | 0.004208818 |
| CYCS | 1881.258296 | 0.24254413 | 0.066959254 | 3.622264511 | 0.000292035 | 0.004231447 |
| ZNF582-AS | 17.82568441 | -0.641531239 | 0.177111095 | -3.622196796 | 0.000292112 | 0.004231447 |
| SNAP29 | 571.951799 | -0.141620148 | 0.039102807 | -3.621738698 | 0.00029263 | 0.004235764 |
| LOC37444 | 150.6881627 | 0.513828309 | 0.142039149 | 3.617511883 | 0.000297449 | 0.004302287 |
| STX8 | 287.7585823 | -0.243994646 | 0.067465613 | -3.616577934 | 0.000298523 | 0.004314593 |
| PCED1A | 188.6671142 | -0.351819543 | 0.097307645 | -3.615538585 | 0.000299724 | 0.004328695 |
| HS3ST2 | 13.52111147 | -1.077447415 | 0.298176744 | -3.613452214 | 0.000302147 | 0.004360423 |
| AFF2 | 10.96779674 | 0.836739138 | 0.231596134 | 3.612923594 | 0.000302764 | 0.004366055 |
| FUZ | 58.99520627 | -0.399133498 | 0.110570924 | -3.609750945 | 0.000306491 | 0.004416498 |
| RGS16 | 69.92693479 | 0.527669122 | 0.146189867 | 3.609478091 | 0.000306814 | 0.004417842 |
| SOBP | 13.94607725 | 1.011712712 | 0.280321139 | 3.609120296 | 0.000307237 | 0.004420635 |
| ARHGAP4 | 107.3613238 | -0.331740366 | 0.091993383 | -3.606132904 | 0.000310794 | 0.004468476 |
| RCSD1 | 159.409471 | 0.410595222 | 0.113921731 | 3.604187007 | 0.000313132 | 0.004496374 |
| UST | 85.25047101 | 0.470525494 | 0.130551781 | 3.604129259 | 0.000313201 | 0.004496374 |
| NNT | 414.847014 | -0.241716415 | 0.067088045 | -3.602973018 | 0.000314598 | 0.004513067 |
| ERCC1 | 137.2383933 | 0.417558069 | 0.115911251 | 3.602394653 | 0.000315299 | 0.004519758 |
| SAMD4B | 436.4064145 | 0.287957793 | 0.079945876 | 3.601909262 | 0.000315889 | 0.00452484 |
| PIGK | 318.9341369 | -0.233367228 | 0.064820359 | -3.600214964 | 0.000317954 | 0.004551044 |
| TRHDE | 63.46393227 | -0.598998297 | 0.166415228 | -3.599419986 | 0.000318928 | 0.00456159 |
| ZNF568 | 41.77903942 | -0.498884448 | 0.138623965 | -3.598832626 | 0.000319649 | 0.004568512 |
| DHRS4-AS1 | 13.5064427 | -0.736261908 | 0.204718672 | -3.596457027 | 0.000322581 | 0.004607 |
| BTBD9 | 312.303307 | 0.219320353 | 0.060985675 | 3.596260163 | 0.000322825 | 0.004607071 |
| TAGAP | 270.0383544 | 0.436227506 | 0.121335405 | 3.595220274 | 0.000324117 | 0.004622089 |
| MAPK7 | 150.4307575 | 0.28351125 | 0.078866757 | 3.594813105 | 0.000324625 | 0.004623485 |
| TRAPPC2L | 1018.892199 | -0.249309211 | 0.069353573 | -3.594756551 | 0.000324695 | 0.004623485 |
| FHL3 | 198.7427048 | 0.342950927 | 0.095493073 | 3.591369659 | 0.000328945 | 0.004680538 |
| MLKL | 356.0474471 | 0.415151611 | 0.115617452 | 3.590734818 | 0.000329747 | 0.004688491 |
| NGFRAP1 | 55.74870584 | -0.948702688 | 0.264268223 | -3.589923439 | 0.000330775 | 0.004699641 |
| PAPD7 | 125.5461685 | 0.347903709 | 0.096933123 | 3.589110698 | 0.000331808 | 0.004710841 |
| PKI55 | 58.84784823 | -0.596816527 | 0.166456575 | -3.585418754 | 0.000336538 | 0.004774475 |

| | | | | | | |
|---|---|---|---|---|---|---|
| DEF6 | 80.47289833 | -0.477023954 | 0.133066499 | -3.584853872 | 0.000337267 | 0.004781299 |
| ANAPC13 | 513.1264451 | -0.172633744 | 0.048174912 | -3.58347811 | 0.000339049 | 0.004803031 |
| MICB | 240.3972092 | 0.244857324 | 0.068338009 | 3.583032736 | 0.000339628 | 0.00480722 |
| SLC39A8 | 2716.957365 | 0.412865814 | 0.115233371 | 3.58286675 | 0.000339844 | 0.00480722 |
| FAM3B | 119.645385 | -0.405119908 | 0.113107266 | -3.581731933 | 0.000341324 | 0.004818756 |
| MBOAT7 | 1049.668817 | 0.262643909 | 0.073330091 | 3.581666196 | 0.00034141 | 0.004818756 |
| OSBPL5 | 113.985986 | -0.392975183 | 0.109707924 | -3.582012753 | 0.000340957 | 0.004818756 |
| ABHD15 | 232.8479478 | -0.278677239 | 0.077843705 | -3.579958595 | 0.000343649 | 0.004843259 |
| NDRG2 | 46.53750121 | 0.579561115 | 0.161884398 | 3.580092469 | 0.000343473 | 0.004843259 |
| INPP5B | 361.9102205 | 0.294529186 | 0.082304885 | 3.578514017 | 0.000345553 | 0.004866543 |
| AGRN | 258.8782375 | 0.373701352 | 0.104519297 | 3.575429251 | 0.000349654 | 0.004920693 |
| DPF3 | 9.205968373 | 1.260649307 | 0.352699558 | 3.574286612 | 0.000351184 | 0.004938622 |
| H3F3B | 10852.16809 | 0.207683019 | 0.058136451 | 3.57233739 | 0.000353809 | 0.004971909 |
| RND3 | 1022.428136 | 0.352118683 | 0.098590606 | 3.571523678 | 0.00035491 | 0.00498375 |
| MED13 | 1218.192885 | 0.251590739 | 0.070460831 | 3.570646773 | 0.000356101 | 0.004993731 |
| TMEM63B | 550.4009537 | 0.413417018 | 0.115783035 | 3.570618249 | 0.00035614 | 0.004993731 |
| MBLAC2 | 21.08528021 | -0.683411173 | 0.191468521 | -3.569313473 | 0.000357918 | 0.005015016 |
| CHCHD6 | 19.10906903 | -0.606788705 | 0.17002876 | -3.568741578 | 0.0003587 | 0.005022322 |
| UBE2J1 | 1695.319355 | 0.209433239 | 0.058742099 | 3.565300583 | 0.000363439 | 0.005084985 |
| DERL1 | 949.2669101 | 0.283960832 | 0.079733435 | 3.561377128 | 0.000368915 | 0.005157846 |
| LOC10013[illegible] | 133.7481702 | 0.33877508 | 0.095136728 | 3.560928433 | 0.000369546 | 0.005162922 |
| PI4KB | 754.8242816 | 0.254170458 | 0.071415405 | 3.559042471 | 0.000372209 | 0.005196366 |
| ZNF443 | 21.82641086 | -0.644126628 | 0.181005724 | -3.558598119 | 0.00037284 | 0.005201395 |
| SYNJ1 | 125.2676578 | 0.32391496 | 0.09103003 | 3.55833083 | 0.000373219 | 0.005202922 |
| NDUFA3 | 371.3738752 | -0.242568153 | 0.068189829 | -3.55724829 | 0.00037476 | 0.005220625 |
| FST | 6.203164856 | 1.877638695 | 0.52828007 | 3.554248589 | 0.000379061 | 0.005276723 |
| GTPBP1 | 382.8274683 | 0.256618235 | 0.072217873 | 3.553389553 | 0.000380301 | 0.005290163 |
| C9orf89 | 399.4847809 | 0.355292642 | 0.100006686 | 3.552688886 | 0.000381315 | 0.005300446 |
| PELI2 | 48.99284568 | -0.495351427 | 0.139499162 | -3.550927614 | 0.000383876 | 0.005328592 |
| TMEM209 | 376.244413 | 0.293109206 | 0.082544678 | 3.550915848 | 0.000383893 | 0.005328592 |
| UBP1 | 580.5065531 | 0.167707615 | 0.047318459 | 3.544232426 | 0.000393758 | 0.005461584 |
| RAB11FIP1 | 1744.309432 | -0.185495272 | 0.052386973 | -3.540866368 | 0.000398816 | 0.005527755 |
| LOC10012[illegible] | 21.73651585 | -0.50186002 | 0.141744871 | -3.5405868 | 0.000399238 | 0.005529637 |
| EIF2AK2 | 805.6721609 | 0.466684102 | 0.131879825 | 3.538707328 | 0.000402091 | 0.005565153 |
| TBC1D10B | 600.5603181 | 0.256866749 | 0.072601699 | 3.53802668 | 0.000403129 | 0.005575513 |
| SLC30A1 | 2341.362374 | 0.231764557 | 0.065639038 | 3.530895101 | 0.000414156 | 0.005723909 |
| SIGLEC11 | 141.3540603 | -0.321485365 | 0.09107796 | -3.529782249 | 0.000415902 | 0.005743917 |
| SNX18 | 214.2381469 | -0.293130468 | 0.083052235 | -3.529471149 | 0.000416391 | 0.005746555 |
| DLGAP4 | 28.7150321 | 0.627192461 | 0.177879879 | 3.525932589 | 0.000421994 | 0.005815316 |
| SYNGR2 | 2935.447942 | -0.252495704 | 0.071610042 | -3.525981807 | 0.000421916 | 0.005815316 |
| TOP1MT | 282.0805586 | -0.26364224 | 0.074776129 | -3.525754031 | 0.000422279 | 0.005815316 |
| PPAPDC1B | 407.0990365 | 0.386085426 | 0.109550726 | 3.524261693 | 0.000424665 | 0.005843992 |
| AK8 | 137.1606661 | 0.311149661 | 0.088295211 | 3.523969852 | 0.000425133 | 0.005846256 |
| MANBA | 1079.745061 | -0.25201845 | 0.071528551 | -3.523326637 | 0.000426166 | 0.005856284 |
| TFAP4 | 39.45082959 | -0.817319028 | 0.232038968 | -3.522335209 | 0.000427763 | 0.005874041 |
| C5orf20 | 20.63098219 | -1.144777608 | 0.325024213 | -3.522130233 | 0.000428094 | 0.005874398 |
| ZC3H12C | 230.1178655 | 0.416854271 | 0.118457011 | 3.519034174 | 0.000433121 | 0.005939151 |

| | | | | | | |
|---|---|---|---|---|---|---|
| IL36B | 938.2701184 | 0.389178334 | 0.110605971 | 3.518601503 | 0.000433828 | 0.005940389 |
| PLCD1 | 137.3558387 | -0.383505508 | 0.108991375 | -3.518677582 | 0.000433703 | 0.005940389 |
| CD151 | 4408.941562 | -0.233520159 | 0.066380029 | -3.517927967 | 0.00043493 | 0.005951257 |
| VWA5A | 964.8523123 | -0.273394968 | 0.077771409 | -3.515366038 | 0.000439148 | 0.006004706 |
| CCDC124 | 1444.942183 | -0.177563392 | 0.050529785 | -3.514034153 | 0.000441356 | 0.006030614 |
| PTGES2 | 198.8877188 | -0.29005179 | 0.082589556 | -3.511966951 | 0.000444803 | 0.006073409 |
| UBE2V1 | 51.68378813 | -0.422060733 | 0.120211084 | -3.510996807 | 0.00044643 | 0.0060913 |
| OSBPL8 | 2337.839784 | 0.20972593 | 0.059750998 | 3.509998759 | 0.000448109 | 0.006109882 |
| SVIL | 1017.124667 | 0.349196886 | 0.099636076 | 3.504723377 | 0.000457082 | 0.006227822 |
| SLC9A1 | 800.7964461 | -0.29588591 | 0.084465471 | -3.503039849 | 0.000459981 | 0.006262888 |
| EEF2K | 247.6674758 | -0.32555705 | 0.092961843 | -3.502050307 | 0.000461692 | 0.006281756 |
| TFRC | 16849.4935 | 0.355268334 | 0.101452229 | 3.501828773 | 0.000462076 | 0.006282544 |
| EIF2AK4 | 436.961958 | -0.258584198 | 0.073854016 | -3.501288219 | 0.000463015 | 0.006290863 |
| GPX3 | 610.8584754 | -0.473446155 | 0.13528918 | -3.499512331 | 0.00046611 | 0.006328454 |
| CCDC23 | 147.3384587 | -0.375731744 | 0.107408695 | -3.498150176 | 0.000468497 | 0.006356386 |
| OMA1 | 152.6457922 | -0.313112204 | 0.089528251 | -3.49735642 | 0.000469894 | 0.006370844 |
| LFNG | 468.4741739 | -0.33519451 | 0.095929392 | -3.49417943 | 0.000475521 | 0.006438084 |
| TOP1 | 1234.677283 | 0.290360044 | 0.083095776 | 3.494281611 | 0.000475339 | 0.006438084 |
| CNPY3 | 2229.348239 | -0.227618198 | 0.065199749 | -3.491090103 | 0.000481054 | 0.006508417 |
| CPVL | 323.0382594 | -0.423739796 | 0.121428279 | -3.489630244 | 0.000483689 | 0.006539479 |
| MVD | 98.20330081 | 0.358906353 | 0.102861152 | 3.489231302 | 0.000484412 | 0.006544655 |
| KIAA1598 | 1669.043727 | -0.245464644 | 0.070380673 | -3.48767119 | 0.000487247 | 0.006573872 |
| PCGF3 | 364.0054297 | 0.250093683 | 0.071708042 | 3.487665775 | 0.000487257 | 0.006573872 |
| MMP19 | 2255.759208 | 0.276474174 | 0.079291857 | 3.486791532 | 0.000488852 | 0.006590782 |
| TDO2 | 2.836733052 | 1.857951204 | 0.533024246 | 3.485678592 | 0.00049089 | 0.006613633 |
| SMARCA2 | 420.867824 | -0.242282452 | 0.069534555 | -3.484346056 | 0.000493341 | 0.006642006 |
| PRRC2C | 1683.580076 | 0.295775797 | 0.084946756 | 3.481896318 | 0.000497876 | 0.006698385 |
| GPAM | 61.83475248 | -0.372037446 | 0.106860423 | -3.481526971 | 0.000498564 | 0.006702949 |
| ZNF318 | 339.7478077 | 0.30110715 | 0.086491822 | 3.481336671 | 0.000498918 | 0.006703036 |
| CTSL2 | 77.57956769 | -0.463189956 | 0.133075386 | -3.480658382 | 0.000500183 | 0.006710673 |
| DUSP22 | 434.71406 | 0.265170168 | 0.076183331 | 3.480684871 | 0.000500134 | 0.006710673 |
| GPAT2 | 56.32152201 | -0.41098279 | 0.11811805 | -3.479424104 | 0.000502493 | 0.006736969 |
| TNFRSF10I | 1115.904928 | -0.231903532 | 0.066664068 | -3.478688557 | 0.000503874 | 0.006750788 |
| CMBL | 10.58784065 | -1.614460888 | 0.46429539 | -3.477227907 | 0.000506627 | 0.006778361 |
| LOC28299 | 56.41797957 | -0.624061034 | 0.17947108 | -3.47722337 | 0.000506636 | 0.006778361 |
| SLIRP | 1102.866829 | -0.233949906 | 0.067317197 | -3.475336389 | 0.000510213 | 0.006821489 |
| NCKAP5L | 261.0596703 | -0.313409858 | 0.090186235 | -3.475140714 | 0.000510585 | 0.006821733 |
| SPTBN1 | 77.92832676 | -0.597480893 | 0.171977357 | -3.474183482 | 0.000512411 | 0.006841373 |
| CD109 | 865.3374144 | -0.296986931 | 0.08551107 | -3.473081706 | 0.000514519 | 0.006864763 |
| DKK2 | 4.123682356 | 1.829877243 | 0.527023116 | 3.472100535 | 0.000516403 | 0.006880372 |
| ENDOD1 | 139.9400518 | 0.504060059 | 0.145169252 | 3.472223303 | 0.000516167 | 0.006880372 |
| SP4 | 53.94086266 | 0.436081472 | 0.125673752 | 3.469948687 | 0.000520558 | 0.006930938 |
| DHX57 | 108.6313571 | -0.301682093 | 0.08695517 | -3.469397983 | 0.000521626 | 0.006935576 |
| DNAJC6 | 218.2452391 | 0.32344989 | 0.093228253 | 3.469440649 | 0.000521543 | 0.006935576 |
| TAPT1 | 264.6322865 | 0.326519668 | 0.094144908 | 3.468266903 | 0.000523827 | 0.006960032 |
| SLC30A5 | 389.2150041 | 0.498087567 | 0.143622329 | 3.468037106 | 0.000524275 | 0.006961185 |
| TBC1D10C | 74.95493976 | 0.3921807 | 0.113113804 | 3.46713386 | 0.00052604 | 0.00697981 |

| | | | | | | |
|---|---|---|---|---|---|---|
| SH3GLB2 | 631.331006 | -0.296093435 | 0.085431458 | -3.465859537 | 0.000528539 | 0.007008148 |
| KCNJ5 | 22.93738991 | -0.859560988 | 0.248081269 | -3.464836299 | 0.000530554 | 0.007030027 |
| ZNF823 | 54.29291309 | -0.375002913 | 0.108261732 | -3.463854735 | 0.000532494 | 0.007050879 |
| LITAF | 1927.633128 | 0.342928779 | 0.09900827 | 3.463637718 | 0.000532924 | 0.007051723 |
| APOBEC3G | 280.0709926 | 0.348639577 | 0.10069384 | 3.462372446 | 0.000535436 | 0.007080098 |
| ARID3B | 171.5104505 | 0.363004446 | 0.104857233 | 3.461892267 | 0.000536392 | 0.007087877 |
| CPEB4 | 746.4111715 | 0.168180852 | 0.048584208 | 3.461636166 | 0.000536902 | 0.007089762 |
| RASGRP2 | 42.0430812 | -0.575841477 | 0.16645008 | -3.459544616 | 0.00054109 | 0.007140159 |
| RCN2 | 211.4709247 | -0.3064801 | 0.088594796 | -3.459346516 | 0.000541488 | 0.007140523 |
| ZC3HAV1 | 980.8201347 | 0.347756855 | 0.100537432 | 3.458978886 | 0.000542227 | 0.007145385 |
| GALK1 | 148.185795 | -0.297539483 | 0.086031608 | -3.458490312 | 0.000543212 | 0.007153465 |
| ASTE1 | 68.17337966 | -0.379506273 | 0.109801741 | -3.456286484 | 0.000547673 | 0.007202366 |
| TRIP11 | 351.8619859 | 0.203648994 | 0.058920502 | 3.456334985 | 0.000547574 | 0.007202366 |
| GRAMD1B | 110.0649143 | -0.358714359 | 0.103861711 | -3.453768995 | 0.000552811 | 0.007260021 |
| PMAIP1 | 80.0099125 | 0.558114243 | 0.161588321 | 3.453926866 | 0.000552487 | 0.007260021 |
| FUT10 | 129.2711344 | -0.353441579 | 0.102348833 | -3.453303469 | 0.000553766 | 0.007267608 |
| RPLP2 | 19380.64828 | -0.194693878 | 0.056389101 | -3.452686329 | 0.000555034 | 0.007279295 |
| ZNF331 | 125.8190748 | -0.333601717 | 0.096655835 | -3.451438989 | 0.000557606 | 0.00730805 |
| DDX28 | 360.8926211 | -0.220161309 | 0.063795198 | -3.451063986 | 0.000558381 | 0.007313237 |
| LINC00656 | 75.60276598 | 0.489033105 | 0.141739676 | 3.450220278 | 0.000560129 | 0.007326172 |
| UBE2Z | 2361.288724 | 0.159425055 | 0.046206175 | 3.45029761 | 0.000559969 | 0.007326172 |
| CCL15 | 1686.993546 | 0.523864574 | 0.151870208 | 3.449422905 | 0.000561786 | 0.007342857 |
| HNMT | 1768.022906 | -0.196236701 | 0.056902619 | -3.448640942 | 0.000563415 | 0.007359159 |
| METTL7B | 23.96773466 | 0.814008113 | 0.236136256 | 3.447196656 | 0.000566436 | 0.007393603 |
| CPNE9 | 4.788781711 | -1.290491857 | 0.374476278 | -3.446124449 | 0.000568688 | 0.007417976 |
| GOSR1 | 337.5562033 | 0.254728513 | 0.074004215 | 3.442081139 | 0.000577257 | 0.007524651 |
| AIDA | 1077.427596 | 0.289564903 | 0.084139948 | 3.441467572 | 0.000578568 | 0.007531546 |
| MTHFD1 | 977.4288178 | 0.315986569 | 0.091813528 | 3.4416123 | 0.000578258 | 0.007531546 |
| FOLR3 | 93.62163537 | -0.801861495 | 0.233127418 | -3.439584675 | 0.000582608 | 0.007579012 |
| LOC10012[illegible] | 60.04809307 | 0.400029857 | 0.116323182 | 3.438952137 | 0.00058397 | 0.007591616 |
| INO80D | 273.3521078 | 0.346378474 | 0.100777174 | 3.437072691 | 0.000588038 | 0.007639336 |
| DGKQ | 164.3723033 | -0.289406322 | 0.084238876 | -3.435543495 | 0.000591367 | 0.007677404 |
| SNHG6 | 371.1513531 | -0.250661544 | 0.072970022 | -3.435130459 | 0.000592269 | 0.007683938 |
| SRSF5 | 3201.577779 | 0.211334448 | 0.06158121 | 3.431800822 | 0.000599588 | 0.007773659 |
| CCL8 | 35.43976865 | 2.032422444 | 0.592998748 | 3.4273638 | 0.000609472 | 0.007891187 |
| PPP3CC | 337.691481 | 0.522323439 | 0.15239299 | 3.427476815 | 0.000609218 | 0.007891187 |
| NR1H3 | 2557.082657 | -0.280715673 | 0.081910685 | -3.427094685 | 0.000610076 | 0.007893707 |
| MPPE1 | 172.0515192 | -0.25004111 | 0.072973652 | -3.426457384 | 0.00061151 | 0.007906944 |
| PTGS1 | 192.5074858 | -0.430421366 | 0.12565619 | -3.425389267 | 0.000613919 | 0.007932774 |
| NIPAL4 | 5.117776535 | 2.00347632 | 0.585014648 | 3.424660095 | 0.000615569 | 0.007948762 |
| DPYD | 509.1179153 | 0.240266344 | 0.07017556 | 3.423789467 | 0.000617544 | 0.007968929 |
| ZNF835 | 48.73882518 | -0.495142564 | 0.144731982 | -3.421099865 | 0.000623684 | 0.008042772 |
| PSME1 | 1166.326708 | 0.313520603 | 0.091651409 | 3.42079414 | 0.000624386 | 0.008046432 |
| HERC6 | 89.72414734 | 0.490813554 | 0.143527215 | 3.419654974 | 0.000627006 | 0.008074799 |
| TBC1D22B | 315.3780297 | 0.226640438 | 0.066298377 | 3.418491524 | 0.000629693 | 0.008103982 |
| TMEM141 | 186.5726863 | -0.403279275 | 0.118003075 | -3.417531924 | 0.000631917 | 0.008127176 |
| MCM6 | 230.9919642 | -0.314893733 | 0.092151899 | -3.417116047 | 0.000632883 | 0.008134172 |

| | | | | | | |
|---|---|---|---|---|---|---|
| GATSL2 | 16.54835072 | -0.696437076 | 0.203821724 | -3.416893271 | 0.000633401 | 0.008135403 |
| RBP7 | 31.54452711 | -0.565567306 | 0.165549908 | -3.416294894 | 0.000634795 | 0.00814787 |
| GMPPB | 102.200575 | 0.508726579 | 0.148920434 | 3.416096542 | 0.000635257 | 0.008148379 |
| STK17A | 582.531855 | 0.321071858 | 0.09401268 | 3.415197381 | 0.000637358 | 0.008169887 |
| TMSB4X | 30661.40921 | -0.337167055 | 0.098877074 | -3.409961913 | 0.00064972 | 0.008322803 |
| CAT | 1144.748218 | -0.370466962 | 0.108688888 | -3.408508159 | 0.000653191 | 0.008359797 |
| RPL7A | 3019.423207 | -0.213887979 | 0.062753384 | -3.408389587 | 0.000653475 | 0.008359797 |
| TSTD1 | 55.46951193 | -0.47771581 | 0.140361413 | -3.403469652 | 0.000665358 | 0.008506163 |
| MIB2 | 43.37592828 | -0.373472706 | 0.109850298 | -3.399833349 | 0.000674269 | 0.008614372 |
| SDSL | 334.5095205 | -0.403790623 | 0.118786996 | -3.399283049 | 0.000675628 | 0.008626004 |
| TIMM17A | 784.3911011 | 0.198477037 | 0.058414654 | 3.397726801 | 0.000679482 | 0.008669475 |
| KIAA1430 | 340.3898444 | -0.198633092 | 0.05847721 | -3.396760776 | 0.000681886 | 0.008694379 |
| ALOX5AP | 3927.796007 | 0.341268604 | 0.100534486 | 3.394542702 | 0.000687433 | 0.008755826 |
| SLC25A40 | 413.8316106 | 0.298064768 | 0.087808905 | 3.394470837 | 0.000687614 | 0.008755826 |
| PICALM | 338.1257584 | 0.249566982 | 0.073542604 | 3.393502129 | 0.00069005 | 0.008781049 |
| GBP2 | 4049.654246 | 0.408855464 | 0.120512574 | 3.392637385 | 0.000692232 | 0.008802999 |
| PAG1 | 677.2987972 | 0.30684687 | 0.090525226 | 3.389628316 | 0.000699874 | 0.008894315 |
| SLC29A3 | 1566.924721 | -0.367703771 | 0.108502361 | -3.388901111 | 0.000701733 | 0.008912057 |
| KARS | 386.2879539 | 0.228048721 | 0.067305421 | 3.388266738 | 0.000703358 | 0.008926812 |
| NCF1B | 935.9486988 | -0.359490631 | 0.106116689 | -3.387691732 | 0.000704834 | 0.008939657 |
| BAZ2A | 783.7540206 | 0.363208702 | 0.107245516 | 3.386702929 | 0.000707379 | 0.008964426 |
| FAM214B | 299.2437112 | 0.274720751 | 0.081122642 | 3.386486739 | 0.000707937 | 0.008964426 |
| FOXO3 | 570.3510639 | -0.205933449 | 0.060812064 | -3.386391359 | 0.000708183 | 0.008964426 |
| ATP1B1 | 2585.646759 | 0.240228503 | 0.070977925 | 3.384552347 | 0.000712944 | 0.009018768 |
| PLCG1 | 51.23855036 | -0.434279631 | 0.128413975 | -3.381872041 | 0.000719937 | 0.009101247 |
| C15orf26 | 14.06582436 | 0.847646419 | 0.250758684 | 3.380327278 | 0.000723996 | 0.009146557 |
| TSPYL5 | 205.8432988 | 0.312203925 | 0.092365572 | 3.380089753 | 0.000724622 | 0.009148467 |
| CERS5 | 381.6686331 | 0.299141065 | 0.088532964 | 3.378866493 | 0.000727853 | 0.009183251 |
| TCEB1 | 2239.918162 | 0.175514365 | 0.051958642 | 3.377962913 | 0.000730249 | 0.009207449 |
| MAP4K1 | 58.57332637 | -0.486017512 | 0.14390222 | -3.377414971 | 0.000731706 | 0.009219779 |
| COLEC12 | 632.1226494 | -0.372355958 | 0.110257816 | -3.377138884 | 0.000732441 | 0.009223007 |
| ITIH5 | 389.3786523 | -0.239520061 | 0.070940526 | -3.376350239 | 0.000734543 | 0.009237412 |
| LOC38879( | 207.3709471 | 0.324289358 | 0.09604396 | 3.376468001 | 0.000734229 | 0.009237412 |
| RAPH1 | 308.0706747 | 0.407989325 | 0.120937409 | 3.373557692 | 0.000742035 | 0.009325535 |
| DLC1 | 235.7738689 | 0.375793369 | 0.111443553 | 3.372051221 | 0.000746106 | 0.009370582 |
| SIRPB1 | 133.9674745 | -0.453158441 | 0.134405346 | -3.37158048 | 0.000747382 | 0.009380496 |
| ATP8B2 | 24.55480825 | 0.802716181 | 0.238178032 | 3.370236016 | 0.000751038 | 0.00942025 |
| SYT11 | 190.870704 | -0.315074956 | 0.093530899 | -3.368672396 | 0.000755311 | 0.009467684 |
| NUDT14 | 223.7953166 | -0.270942422 | 0.08046539 | -3.367192044 | 0.000759378 | 0.009512467 |
| KLF7 | 85.35723609 | -0.480060498 | 0.142765212 | -3.362587373 | 0.000772157 | 0.009659983 |
| LMTK2 | 127.9881281 | 0.303603168 | 0.090284831 | 3.362726204 | 0.000771769 | 0.009659983 |
| ARRB1 | 666.8400843 | -0.349997136 | 0.10409196 | -3.362383951 | 0.000772726 | 0.00966083 |
| NELF | 169.8687075 | -0.387647838 | 0.115351536 | -3.360578043 | 0.000777796 | 0.009717904 |
| ING4 | 106.3536678 | -0.291586838 | 0.086779238 | -3.360099097 | 0.000779145 | 0.009728458 |
| LY6G5B | 85.02401385 | 0.277852 | 0.082756516 | 3.357463719 | 0.000786611 | 0.009815312 |
| NDUFB1 | 1044.366511 | -0.26868445 | 0.080044794 | -3.356676154 | 0.000788855 | 0.009830576 |
| SH3BP4 | 107.5655442 | -0.4249107 | 0.126584674 | -3.356731007 | 0.000788698 | 0.009830576 |

| | | | | | | |
|---|---|---|---|---|---|---|
| ACE | 256.3904411 | 0.500811159 | 0.149275336 | 3.354949123 | 0.000793796 | 0.009879374 |
| GMPPA | 342.5621268 | 0.325773787 | 0.097099924 | 3.355036472 | 0.000793545 | 0.009879374 |
| MTMR1 | 509.3399227 | -0.297604944 | 0.088737541 | -3.353765942 | 0.000797198 | 0.009915307 |
| TLK2 | 1001.822829 | 0.200830561 | 0.059891328 | 3.353249446 | 0.000798687 | 0.009927421 |
| NHLRC3 | 219.7113343 | 0.377887988 | 0.112703833 | 3.352929349 | 0.000799611 | 0.009932501 |
| GHDC | 130.87084 | -0.439806375 | 0.131256684 | -3.350735069 | 0.000805974 | 0.010005083 |
| SURF4 | 1223.262281 | 0.312439319 | 0.093287708 | 3.349201361 | 0.000810449 | 0.010054157 |
| TMEM39A | 417.114373 | 0.393942738 | 0.117649238 | 3.348451259 | 0.000812646 | 0.010074925 |
| PBRM1 | 911.8577047 | 0.262525102 | 0.078410032 | 3.348106062 | 0.000813659 | 0.010080995 |
| HK3 | 1982.332241 | -0.252359463 | 0.075418681 | -3.346113438 | 0.000819529 | 0.010147197 |
| VNN1 | 200.3679754 | -0.339793417 | 0.101578007 | -3.345147519 | 0.000822388 | 0.010176063 |
| IDH2 | 285.4663492 | -0.287632957 | 0.086006864 | -3.344302335 | 0.000824898 | 0.010200567 |
| PLS3 | 55.82884928 | -0.435388968 | 0.130292233 | -3.341634108 | 0.000832868 | 0.010292514 |
| MALT1 | 91.77545239 | 0.308582165 | 0.092364425 | 3.34092012 | 0.000835012 | 0.010312404 |
| REV3L | 102.0821751 | -0.294170049 | 0.088072811 | -3.340077906 | 0.000837549 | 0.010337103 |
| ANKRD13A | 793.5771634 | 0.163992322 | 0.049131089 | 3.337852381 | 0.000844286 | 0.010409845 |
| NCOA2 | 237.3221536 | -0.325869399 | 0.097634874 | -3.337633231 | 0.000844952 | 0.010409845 |
| ZNF737 | 111.9609588 | -0.338455859 | 0.101407063 | -3.337596498 | 0.000845064 | 0.010409845 |
| PXMP2 | 30.46169359 | -0.480178323 | 0.143900135 | -3.336885841 | 0.000847227 | 0.010429829 |
| MEIS3P1 | 98.35149969 | -0.399550891 | 0.119771305 | -3.335948374 | 0.000850089 | 0.010458381 |
| DEXI | 45.02637887 | -0.6092954 | 0.1826919 | -3.335098047 | 0.000852693 | 0.01048372 |
| GORAB | 327.7692871 | 0.275441162 | 0.082604916 | 3.334440292 | 0.000854712 | 0.010501844 |
| YPEL4 | 166.6826819 | -0.352861416 | 0.105833623 | -3.334114496 | 0.000855714 | 0.010507451 |
| TOM1L2 | 272.1546792 | -0.264328836 | 0.079297364 | -3.333387411 | 0.000857954 | 0.010528242 |
| C21orf33 | 315.7585405 | -0.202225206 | 0.060701763 | -3.331455256 | 0.000863932 | 0.010594851 |
| ZNF133 | 5.775779581 | -1.104740383 | 0.331666444 | -3.330877765 | 0.000865726 | 0.010610101 |
| SLC39A12 | 4.974167761 | 1.310169241 | 0.393459674 | 3.329869174 | 0.000868868 | 0.010641838 |
| CHID1 | 376.1111665 | -0.228470155 | 0.068619017 | -3.32954572 | 0.000869878 | 0.010647437 |
| KSR1 | 12.90423274 | 0.893490389 | 0.268388341 | 3.329095385 | 0.000871286 | 0.010657897 |
| KANSL1-AS | 80.19380619 | -0.494621111 | 0.148841454 | -3.323140815 | 0.0008901 | 0.01088113 |
| GATM | 10.61778985 | -0.629783318 | 0.189559639 | -3.322349224 | 0.000892629 | 0.01090513 |
| MGST3 | 7005.972264 | -0.30579806 | 0.092086911 | -3.320754902 | 0.000897743 | 0.010954416 |
| PLEKHG2 | 78.27076449 | 0.375559442 | 0.113095202 | 3.320737167 | 0.0008978 | 0.010954416 |
| CD3EAP | 44.44467525 | -0.42916275 | 0.129305314 | -3.318987719 | 0.000903444 | 0.011016299 |
| DTNB | 173.327269 | -0.291208171 | 0.087819859 | -3.315971737 | 0.000913251 | 0.011128834 |
| VEGFA | 138.9032805 | -0.41399463 | 0.124937632 | -3.313610344 | 0.000920998 | 0.011216144 |
| ATHL1 | 26.08283135 | -0.511540058 | 0.154389434 | -3.313310014 | 0.000921987 | 0.011221103 |
| GAA | 1152.060313 | -0.340105172 | 0.102691947 | -3.311897198 | 0.000926656 | 0.011270803 |
| SUMF2 | 669.5144392 | -0.214946142 | 0.064963769 | -3.30870802 | 0.000937275 | 0.011392773 |
| TLE2 | 17.72853996 | -0.933070244 | 0.282256388 | -3.305754214 | 0.000947211 | 0.011506289 |
| RNF26 | 342.259516 | -0.218492973 | 0.066132389 | -3.303872355 | 0.000953592 | 0.011576504 |
| CRABP2 | 161.7837415 | -0.453213249 | 0.137186609 | -3.303625995 | 0.000954431 | 0.011579384 |
| LY6G5C | 42.03992991 | -0.436857574 | 0.132270977 | -3.30274701 | 0.000957427 | 0.011608429 |
| USP21 | 78.0403175 | -0.355375315 | 0.107663109 | -3.300808592 | 0.000964066 | 0.011681573 |
| LATS2 | 192.8107139 | 0.290875077 | 0.088191522 | 3.298220398 | 0.000972997 | 0.01178238 |
| TRAFD1 | 1140.832196 | 0.287380121 | 0.087136909 | 3.298029787 | 0.000973658 | 0.011782975 |
| PTP4A1 | 827.0716972 | 0.429534794 | 0.130247109 | 3.29784513 | 0.000974299 | 0.011783325 |

| | | | | | | |
|---|---|---|---|---|---|---|
| DYRK1B | 59.72912183 | -0.400362649 | 0.121426048 | -3.297172697 | 0.000976634 | 0.011804163 |
| ARHGAP35 | 402.4794967 | -0.25692239 | 0.077966837 | -3.295277854 | 0.000983244 | 0.011876602 |
| FAM213B | 104.808607 | -0.513179003 | 0.155917516 | -3.291349284 | 0.00099708 | 0.012036183 |
| ARMC7 | 125.2883152 | -0.315527331 | 0.095880221 | -3.29084902 | 0.000998855 | 0.012050057 |
| APOA1 | 2.594212309 | 1.919922688 | 0.584047301 | 3.287272599 | 0.001011629 | 0.012196518 |
| ARF1 | 9575.302341 | 0.135347373 | 0.041189068 | 3.286002314 | 0.001016202 | 0.012243992 |
| SLC35E4 | 41.14026921 | -0.447090752 | 0.136077472 | -3.285560391 | 0.001017797 | 0.012255551 |
| CD46 | 35.14742484 | -0.618325163 | 0.188244598 | -3.284690074 | 0.001020946 | 0.012284754 |
| PCNA | 309.7277278 | -0.217625365 | 0.06625753 | -3.284537861 | 0.001021498 | 0.012284754 |
| C10orf137 | 239.667041 | 0.253258594 | 0.077111216 | 3.28432888 | 0.001022256 | 0.012286198 |
| SOD2 | 83599.84538 | 0.211735144 | 0.064479228 | 3.28377292 | 0.001024274 | 0.012302784 |
| MEF2A | 263.2359685 | 0.348171814 | 0.1060661 | 3.282592779 | 0.001028571 | 0.012346701 |
| ALDH4A1 | 27.99719955 | -0.468021954 | 0.142601414 | -3.282028849 | 0.001030631 | 0.012363717 |
| RC3H2 | 433.0410094 | 0.202211803 | 0.06163294 | 3.280904709 | 0.001034747 | 0.012405373 |
| LYZ | 66510.70773 | -0.268415821 | 0.081850103 | -3.279358381 | 0.001040434 | 0.012465799 |
| NINJ2 | 79.3040586 | -0.485029515 | 0.147929712 | -3.278783598 | 0.001042556 | 0.012483452 |
| MBD2 | 4514.775245 | -0.178176243 | 0.054390578 | -3.275865972 | 0.001053385 | 0.012598615 |
| PRKACB | 216.5091034 | -0.402621628 | 0.122906367 | -3.275840277 | 0.001053481 | 0.012598615 |
| CCDC109B | 131.2508129 | 0.357604741 | 0.10920093 | 3.27474079 | 0.00105759 | 0.012639902 |
| EVL | 4098.455129 | 0.274726801 | 0.083913047 | 3.273946192 | 0.001060568 | 0.01266764 |
| DPY30 | 616.1183923 | -0.226248189 | 0.06913888 | -3.272372781 | 0.001066489 | 0.012730464 |
| AKIRIN2 | 432.6968603 | 0.302689564 | 0.092548075 | 3.270619771 | 0.001073121 | 0.012801701 |
| SSH3 | 69.97032207 | -0.374778994 | 0.114684506 | -3.267913047 | 0.001083436 | 0.012916762 |
| CYB5D1 | 21.19121876 | 0.606577354 | 0.18567077 | 3.266951248 | 0.001087124 | 0.012952709 |
| MICALL2 | 81.42849829 | 0.361411579 | 0.110738801 | 3.263639979 | 0.001099908 | 0.013096931 |
| ABCB6 | 70.76193747 | -0.293652302 | 0.090038354 | -3.261413471 | 0.001108583 | 0.013192065 |
| ALAD | 162.7117648 | -0.341535369 | 0.10479877 | -3.258963512 | 0.0011182 | 0.013298303 |
| IFNG | 13.80402886 | 1.888333406 | 0.579547956 | 3.258286716 | 0.001120871 | 0.013321838 |
| TOR1A | 1121.01333 | 0.360720828 | 0.110769628 | 3.256495801 | 0.001127966 | 0.013397899 |
| FAM156B | 214.0581031 | 0.367671663 | 0.112914707 | 3.25618931 | 0.001129184 | 0.013404107 |
| TMEFF1 | 108.3284297 | 0.373629296 | 0.114753065 | 3.255941749 | 0.001130169 | 0.013407539 |
| HYOU1 | 1429.611939 | 0.279859213 | 0.08597344 | 3.255182225 | 0.001133196 | 0.013426913 |
| SEC24C | 1090.999911 | 0.158375482 | 0.04865315 | 3.255194817 | 0.001133146 | 0.013426913 |
| NDUFB9 | 2105.55523 | -0.2533674 | 0.077848407 | -3.254625368 | 0.00113542 | 0.013431968 |
| PRDM1 | 290.9041478 | 0.297694764 | 0.091464455 | 3.254759055 | 0.001134886 | 0.013431968 |
| RPL11 | 18041.15814 | -0.230732869 | 0.070895437 | -3.254551784 | 0.001135715 | 0.013431968 |
| NAGLU | 302.8619134 | -0.280412742 | 0.086185726 | -3.253586833 | 0.001139579 | 0.013469403 |
| RUFY1 | 1106.958166 | -0.197360931 | 0.060693702 | -3.251753039 | 0.001146956 | 0.01354829 |
| CCDC107 | 237.3940598 | -0.239772039 | 0.073750638 | -3.251118183 | 0.001149521 | 0.01357026 |
| FAM149B1 | 176.3018636 | -0.29065746 | 0.089425463 | -3.250276277 | 0.001152929 | 0.013602167 |
| CACNB3 | 116.3460823 | 0.315045811 | 0.096964534 | 3.249082904 | 0.001157777 | 0.013651002 |
| ST18 | 8.902132586 | -1.006744812 | 0.310083041 | -3.246694206 | 0.001167538 | 0.013757663 |
| S100A16 | 228.1807966 | 0.343324296 | 0.105751525 | 3.246518627 | 0.001168258 | 0.013757738 |
| LILRB5 | 47.11978674 | -0.924636153 | 0.28494448 | -3.244969518 | 0.001174632 | 0.013824349 |
| MAGEH1 | 56.46385402 | -0.441862104 | 0.136213409 | -3.243895778 | 0.001179069 | 0.013868095 |
| CASD1 | 71.84358654 | -0.485764622 | 0.149843038 | -3.241823105 | 0.001187677 | 0.013960823 |
| TUG1 | 585.5882175 | 0.198037392 | 0.061130997 | 3.239557708 | 0.001197152 | 0.014063621 |

| | | | | | | |
|---|---|---|---|---|---|---|
| CHP1 | 1621.784412 | -0.210120781 | 0.064876844 | -3.2387639 | 0.001200489 | 0.014094224 |
| CLDN16 | 4.980159806 | 1.840994779 | 0.568618142 | 3.237664514 | 0.001205124 | 0.014131422 |
| NACC2 | 229.2884012 | -0.364973654 | 0.112725673 | -3.237715466 | 0.001204909 | 0.014131422 |
| TNNI2 | 88.9421797 | -0.437968236 | 0.135289152 | -3.237275335 | 0.001206769 | 0.014133505 |
| WDR43 | 230.6948551 | -0.24305171 | 0.075077251 | -3.237354948 | 0.001206433 | 0.014133505 |
| RNF19B | 1213.736568 | 0.495355764 | 0.153061269 | 3.236323382 | 0.001210801 | 0.014172113 |
| LIG4 | 135.4371755 | 0.255022272 | 0.078822408 | 3.235403207 | 0.001214711 | 0.014209238 |
| LOC10050( | 38.20366442 | -0.56402779 | 0.174349419 | -3.235042562 | 0.001216246 | 0.014210994 |
| TULP4 | 235.5148666 | -0.329931779 | 0.101987509 | -3.235021438 | 0.001216336 | 0.014210994 |
| DGCR6L | 205.8928824 | -0.315769103 | 0.097671355 | -3.232975557 | 0.00122508 | 0.014295821 |
| SERINC3 | 1704.42886 | -0.153961345 | 0.047621705 | -3.233007824 | 0.001224942 | 0.014295821 |
| PARP4 | 1454.651202 | -0.217217325 | 0.067245926 | -3.230193057 | 0.001237067 | 0.014426952 |
| FAM160A2 | 119.9867137 | 0.308581545 | 0.095543797 | 3.229739176 | 0.001239032 | 0.014441131 |
| AKNA | 100.3559541 | -0.381348172 | 0.11810606 | -3.228862036 | 0.001242838 | 0.014476738 |
| PSKH1 | 136.7629123 | -0.287513692 | 0.089092977 | -3.227119609 | 0.001250432 | 0.014556386 |
| GUSBP4 | 18.34382452 | -0.57464919 | 0.178170345 | -3.225279666 | 0.001258497 | 0.014641423 |
| IFT172 | 59.78941314 | -0.442184689 | 0.137128461 | -3.224601849 | 0.00126148 | 0.014666142 |
| MMAA | 187.0383035 | 0.318465436 | 0.098765781 | 3.224451153 | 0.001262144 | 0.014666142 |
| GGT8P | 15.18871256 | -0.696362148 | 0.216043506 | -3.223249613 | 0.001267451 | 0.014718928 |
| DLG5 | 65.72778097 | -0.450260888 | 0.139715487 | -3.222698489 | 0.001269891 | 0.014738391 |
| CSTF3 | 179.0411711 | 0.289306267 | 0.089801618 | 3.221615296 | 0.001274702 | 0.014785311 |
| CD38 | 174.320835 | 0.773899488 | 0.24028138 | 3.22080508 | 0.001278311 | 0.01481825 |
| EIF4A1 | 20017.80098 | 0.133388327 | 0.041430603 | 3.219560333 | 0.001283873 | 0.014867417 |
| RGL1 | 70.20257877 | 0.399786898 | 0.124182933 | 3.219338504 | 0.001284867 | 0.014867417 |
| ZNF586 | 299.5243504 | 0.278609947 | 0.086541844 | 3.219366888 | 0.00128474 | 0.014867417 |
| WFS1 | 174.5271927 | 0.52965216 | 0.164569602 | 3.21840821 | 0.001289042 | 0.014906775 |
| HFE | 183.7481037 | -0.353684813 | 0.109985498 | -3.215740429 | 0.001301085 | 0.01503701 |
| ATP6V0E2 | 19.98522684 | -0.615048355 | 0.191278831 | -3.215454378 | 0.001302382 | 0.01504298 |
| PLEKHO1 | 323.0316708 | 0.271348307 | 0.084423168 | 3.214145033 | 0.001308336 | 0.015102694 |
| ACOT2 | 446.8990816 | -0.350137284 | 0.109017221 | -3.211761221 | 0.00131924 | 0.015219445 |
| DGKG | 12.40660522 | -0.793101354 | 0.247116671 | -3.209420684 | 0.001330027 | 0.015334714 |
| SMG1 | 596.6541596 | 0.230991923 | 0.071990242 | 3.208656013 | 0.001333569 | 0.015366356 |
| KCNC4 | 30.32816104 | -0.526401695 | 0.164084524 | -3.208112996 | 0.00133609 | 0.015379816 |
| TMEM9 | 149.0763655 | -0.400839214 | 0.124947525 | -3.208060445 | 0.001336334 | 0.015379816 |
| JAK2 | 306.9884825 | 0.434955078 | 0.135596935 | 3.20770582 | 0.001337983 | 0.015389598 |
| RPL18A | 4249.833393 | -0.186310399 | 0.058132186 | -3.204943984 | 0.001350888 | 0.01552876 |
| NOB1 | 237.6752075 | -0.265709456 | 0.082931347 | -3.203968891 | 0.001355471 | 0.015572158 |
| MARVELD1 | 530.3430836 | -0.383837984 | 0.119808671 | -3.203757961 | 0.001356465 | 0.015574283 |
| CCND2 | 83.25178231 | 0.474891603 | 0.148277722 | 3.202717156 | 0.001361376 | 0.015618307 |
| NADKD1 | 239.2378474 | -0.276965145 | 0.086481286 | -3.202602068 | 0.00136192 | 0.015618307 |
| TBCA | 1043.939967 | 0.182804642 | 0.057085004 | 3.202323381 | 0.001363239 | 0.015624126 |
| TTLL1 | 36.34828072 | -0.416639717 | 0.130119103 | -3.201987312 | 0.00136483 | 0.015633066 |
| SIGLEC15 | 14.96988585 | 0.891085935 | 0.278360422 | 3.201194792 | 0.00136859 | 0.015666816 |
| SNORA70G | 511.2399656 | 0.346816081 | 0.108403333 | 3.199311969 | 0.00137756 | 0.015760139 |
| PER2 | 109.6199375 | 0.313913994 | 0.098209482 | 3.196371549 | 0.001391678 | 0.015912204 |
| COMMD3 | 494.0887721 | -0.18356112 | 0.05744477 | -3.195436621 | 0.001396194 | 0.015954379 |
| ASB13 | 62.96571697 | -0.520482349 | 0.162906151 | -3.194982796 | 0.001398392 | 0.015970015 |

| | | | | | | |
|---|---|---|---|---|---|---|
| SMURF2 | 138.2471918 | -0.308757088 | 0.09665755 | -3.194340109 | 0.001401509 | 0.015996133 |
| SRSF8 | 243.1366415 | -0.210117522 | 0.065794207 | -3.193556593 | 0.001405318 | 0.016030109 |
| LOC10013: | 22.45715299 | 0.584220447 | 0.183004972 | 3.192374728 | 0.001411081 | 0.016086328 |
| GRIPAP1 | 371.1988856 | 0.265096985 | 0.083091739 | 3.190413236 | 0.001420695 | 0.016186344 |
| TACC3 | 115.5348003 | -0.393973991 | 0.123500733 | -3.190053878 | 0.001422463 | 0.01618734 |
| ZNF143 | 126.2782203 | 0.44661627 | 0.139997252 | 3.190178842 | 0.001421848 | 0.01618734 |
| NAAA | 208.676425 | -0.349937435 | 0.109704074 | -3.189830817 | 0.001423561 | 0.016190276 |
| TIFAB | 30.04981761 | -0.842803001 | 0.264411335 | -3.187469256 | 0.001435237 | 0.016313439 |
| SLC45A3 | 41.48750569 | -0.546690213 | 0.171544291 | -3.186874997 | 0.001438189 | 0.016337354 |
| KLF3 | 937.2793325 | -0.244370059 | 0.076687308 | -3.186577594 | 0.001439669 | 0.016344523 |
| ARNTL2 | 314.9004911 | 0.590413528 | 0.185308762 | 3.186106916 | 0.001442013 | 0.016361497 |
| USO1 | 490.2996564 | 0.253850319 | 0.079694144 | 3.185307036 | 0.001446005 | 0.016397135 |
| ST6GAL1 | 62.71476142 | -0.578250285 | 0.181604002 | -3.184127444 | 0.001451911 | 0.016454419 |
| DBP | 22.81268378 | -0.713042552 | 0.223960729 | -3.183783851 | 0.001453635 | 0.016464277 |
| RARA | 217.7554428 | 0.4421948 | 0.138897285 | 3.183610116 | 0.001454508 | 0.016464481 |
| RTN2 | 64.55183837 | 0.39351432 | 0.12363885 | 3.182772401 | 0.001458722 | 0.016502492 |
| FAM195A | 188.6762274 | -0.342223969 | 0.107549656 | -3.182008954 | 0.001462573 | 0.016531694 |
| PPP1R10 | 748.6211947 | 0.292322115 | 0.09186971 | 3.181920502 | 0.00146302 | 0.016531694 |
| TBC1D19 | 20.93394275 | 0.561502434 | 0.176482304 | 3.1816359 | 0.001464458 | 0.016538246 |
| PPM1K | 198.8411627 | 0.30714495 | 0.096582161 | 3.18014163 | 0.001472031 | 0.016614029 |
| LOC72843: | 127.2093727 | 0.374297706 | 0.117782596 | 3.17786937 | 0.001483616 | 0.016734978 |
| SLC11A1 | 3293.287406 | 0.3742808 | 0.11781578 | 3.176830804 | 0.001488939 | 0.016785192 |
| SCP2 | 925.0726234 | -0.240682977 | 0.075814727 | -3.174620399 | 0.001500326 | 0.016903677 |
| ASPSCR1 | 288.6968706 | -0.244005642 | 0.076913726 | -3.172458984 | 0.001511539 | 0.017000182 |
| HNRNPF | 296.7881797 | 0.310284843 | 0.097801678 | 3.172592226 | 0.001510846 | 0.017000182 |
| RPL13 | 1065.163719 | -0.302833586 | 0.095449805 | -3.172699881 | 0.001510286 | 0.017000182 |
| CCDC125 | 268.7442445 | -0.270974087 | 0.085424695 | -3.172081409 | 0.001513506 | 0.01701237 |
| WAPAL | 603.2359367 | 0.211613532 | 0.066813628 | 3.167221078 | 0.001539033 | 0.017289215 |
| CLIP2 | 7.87181572 | -1.082522877 | 0.341939657 | -3.165830152 | 0.001546411 | 0.017361974 |
| TCHP | 267.0096355 | -0.217421723 | 0.068692872 | -3.165127868 | 0.001550148 | 0.017393799 |
| NEFH | 39.24505824 | -0.549167909 | 0.173558265 | -3.164170304 | 0.001555258 | 0.017440974 |
| HPS1 | 840.8705153 | -0.245688212 | 0.077712755 | -3.161491466 | 0.001569634 | 0.017591956 |
| CYLD | 1055.888499 | 0.236395853 | 0.074890899 | 3.156536459 | 0.001596549 | 0.017883208 |
| STAT4 | 10.4601501 | 0.967646333 | 0.306797676 | 3.154021067 | 0.001610375 | 0.018027587 |
| ELF4 | 904.4796918 | 0.176240441 | 0.05589096 | 3.153290672 | 0.00161441 | 0.018055805 |
| GLI3 | 29.15971942 | 0.53754156 | 0.170475797 | 3.153184019 | 0.001615 | 0.018055805 |
| MAPK10 | 2.885995317 | 1.730922367 | 0.548966539 | 3.153056232 | 0.001615707 | 0.018055805 |
| HOMER2 | 42.21452115 | -0.494743921 | 0.156925448 | -3.15273225 | 0.001617501 | 0.018065376 |
| CDK19 | 333.054131 | -0.254434521 | 0.080726339 | -3.151815449 | 0.001622588 | 0.018111691 |
| LOC10013: | 85.41463729 | -0.390235288 | 0.12382055 | -3.15161973 | 0.001623676 | 0.01811334 |
| C1RL | 813.8832309 | 0.357790969 | 0.113534988 | 3.151371891 | 0.001625054 | 0.018118227 |
| CBWD2 | 265.3102563 | 0.205586706 | 0.065255307 | 3.150497878 | 0.001629924 | 0.018162015 |
| ARC | 79.96075244 | 0.494350024 | 0.156998973 | 3.148746863 | 0.001639722 | 0.018260623 |
| ARID5A | 50.29851951 | 0.356637193 | 0.113304142 | 3.14760948 | 0.001646115 | 0.018321226 |
| CASZ1 | 34.21729753 | 0.403494381 | 0.12819947 | 3.147395077 | 0.001647322 | 0.018324081 |
| ACSL4 | 2578.839449 | 0.230397534 | 0.073254329 | 3.145172961 | 0.001659887 | 0.018453189 |
| CASP9 | 696.067015 | 0.173220452 | 0.055079897 | 3.144894236 | 0.001661469 | 0.018460126 |

| | | | | | | |
|---|---|---|---|---|---|---|
| CYP1B1 | 3094.613444 | 0.276024053 | 0.087802779 | 3.143682433 | 0.001668364 | 0.018526051 |
| POC1B | 185.9692895 | 0.270514664 | 0.08607423 | 3.142806678 | 0.001673363 | 0.018570861 |
| MICALL1 | 909.3412014 | 0.276251022 | 0.087925391 | 3.141879943 | 0.001678669 | 0.018619014 |
| ERI1 | 289.0977873 | 0.281886736 | 0.089736666 | 3.141265964 | 0.001682192 | 0.018647359 |
| SLC12A6 | 737.172949 | 0.291564912 | 0.092885047 | 3.138986532 | 0.001695332 | 0.018782216 |
| FGD5 | 957.6930055 | -0.310174839 | 0.098935869 | -3.135110062 | 0.001717896 | 0.019010335 |
| NPC2 | 5657.418389 | -0.183180226 | 0.058428457 | -3.135120036 | 0.001717838 | 0.019010335 |
| ACSF3 | 154.9589982 | -0.291259166 | 0.092937803 | -3.133914916 | 0.001724909 | 0.019076975 |
| USP13 | 292.1195935 | -0.261196048 | 0.083372148 | -3.132893356 | 0.001730923 | 0.019132511 |
| VPS26B | 241.7132774 | -0.21397636 | 0.068310394 | -3.132412905 | 0.001733759 | 0.019152863 |
| C8B | 86.36068617 | -0.398331303 | 0.127250563 | -3.130291085 | 0.001746332 | 0.019280702 |
| SURF1 | 118.6806704 | -0.255300313 | 0.081565915 | -3.129987727 | 0.001748136 | 0.01928957 |
| HLA-E | 14075.23027 | 0.34242147 | 0.109409149 | 3.129733424 | 0.00174965 | 0.019295225 |
| REXO1 | 429.2692347 | 0.240097866 | 0.076760544 | 3.127881238 | 0.001760713 | 0.019383942 |
| SENP2 | 259.1421168 | 0.243803476 | 0.077938385 | 3.128156635 | 0.001759064 | 0.019383942 |
| ZC3H12D | 22.59830356 | -0.574147429 | 0.183556836 | -3.127900017 | 0.001760601 | 0.019383942 |
| CTSZ | 11575.13261 | 0.204804418 | 0.065484854 | 3.127508181 | 0.001762949 | 0.019386403 |
| NEFM | 354.2001701 | 0.352688588 | 0.112765497 | 3.127628566 | 0.001762227 | 0.019386403 |
| GAPT | 16.60652006 | -0.75174621 | 0.240401694 | -3.127042058 | 0.001765747 | 0.019395026 |
| VAV2 | 191.8750927 | -0.251800971 | 0.0805203 | -3.127173775 | 0.001764956 | 0.019395026 |
| MZT2B | 173.4203979 | -0.316542815 | 0.101233918 | -3.126845431 | 0.001766928 | 0.019396943 |
| ERMAP | 60.83959832 | -0.330834908 | 0.105825611 | -3.12622724 | 0.001770647 | 0.019426697 |
| SLC12A2 | 59.91094051 | -0.414120289 | 0.132488265 | -3.125712968 | 0.001773746 | 0.019449623 |
| AREG | 41.39128596 | 0.538542322 | 0.172358428 | 3.12454882 | 0.00178078 | 0.019504869 |
| RPS18 | 1006.955326 | -0.245671086 | 0.078626221 | -3.124543987 | 0.001780809 | 0.019504869 |
| C19orf66 | 271.9238901 | 0.22879923 | 0.073291902 | 3.121753218 | 0.001797776 | 0.019679515 |
| CHRNA1 | 6.552004839 | -1.306290166 | 0.418559691 | -3.120917267 | 0.001802887 | 0.019713063 |
| PALD1 | 3.89626829 | -1.401945696 | 0.449197049 | -3.121003797 | 0.001802357 | 0.019713063 |
| TWIST1 | 2.987145649 | 1.985831269 | 0.636418047 | 3.120325197 | 0.001806515 | 0.019741528 |
| NUDT4 | 352.2872495 | 0.223390656 | 0.071602725 | 3.119862507 | 0.001809355 | 0.019755147 |
| SMARCD3 | 3.023573151 | -1.747069228 | 0.559996118 | -3.119788107 | 0.001809812 | 0.019755147 |
| METRNL | 824.8604462 | 0.27179439 | 0.087133122 | 3.119300504 | 0.00181281 | 0.019776666 |
| PIK3CG | 924.6306855 | 0.428391549 | 0.137380384 | 3.118287615 | 0.001819052 | 0.019833534 |
| ATP8A1 | 27.86099335 | -0.5805083 | 0.186193159 | -3.117774591 | 0.001822221 | 0.019856851 |
| TMEM66 | 5602.111485 | -0.206370415 | 0.06623955 | -3.115516553 | 0.00183623 | 0.0199982 |
| EFTUD1 | 575.6755289 | 0.171433714 | 0.055048505 | 3.114230149 | 0.001844256 | 0.020074254 |
| FAM108A1 | 817.5217994 | -0.202374113 | 0.064995875 | -3.113645489 | 0.001847914 | 0.02009137 |
| ZNF804A | 108.7010672 | 0.286631796 | 0.092055587 | 3.113681692 | 0.001847687 | 0.02009137 |
| TBXAS1 | 755.0997653 | -0.222429271 | 0.071445656 | -3.11326514 | 0.001850297 | 0.020105936 |
| APP | 662.0191151 | 0.177421328 | 0.056996245 | 3.112859966 | 0.001852839 | 0.020122209 |
| CCL13 | 14.43430413 | 1.123010082 | 0.36101608 | 3.110692691 | 0.001866491 | 0.020259048 |
| PIP4K2C | 592.6329649 | 0.16216814 | 0.052148009 | 3.109766679 | 0.001872352 | 0.020311222 |
| SAPCD1 | 10.91140983 | -0.973576343 | 0.313123329 | -3.109242439 | 0.001875677 | 0.020335847 |
| FPR3 | 3399.805888 | -0.293257078 | 0.094323886 | -3.109043628 | 0.00187694 | 0.020338091 |
| EYA4 | 9.4070752 | 0.85752965 | 0.275856278 | 3.108610238 | 0.001879695 | 0.02035474 |
| LONRF1 | 196.5560622 | 0.370904878 | 0.119320732 | 3.108469671 | 0.001880589 | 0.02035474 |
| KCNJ10 | 3.087738338 | -1.582593214 | 0.509191649 | -3.108050213 | 0.001883261 | 0.020372207 |

| | | | | | | |
|---|---|---|---|---|---|---|
| TBKBP1 | 163.2272454 | 0.329264637 | 0.106003973 | 3.106153741 | 0.001895381 | 0.020491817 |
| PLBD1 | 2609.671952 | -0.280497096 | 0.090311571 | -3.105882137 | 0.001897123 | 0.020499145 |
| AKIRIN1 | 1273.588759 | 0.219747283 | 0.070758779 | 3.105583317 | 0.001899041 | 0.020508367 |
| GNL1 | 426.551696 | 0.25162295 | 0.081039215 | 3.10495296 | 0.001903093 | 0.020540609 |
| PLEKHG6 | 11.5703773 | -0.776802442 | 0.250230905 | -3.104342533 | 0.001907024 | 0.020571515 |
| LAMC2 | 17.77500962 | 0.87069708 | 0.280557516 | 3.103453057 | 0.001912766 | 0.020621906 |
| DCSTAMP | 5038.136037 | 0.275073136 | 0.088649723 | 3.102921556 | 0.001916204 | 0.020647422 |
| HUWE1 | 1015.117063 | 0.259140314 | 0.083552102 | 3.101541513 | 0.001925159 | 0.020732313 |
| PLEKHB1 | 29.2898176 | -0.58675975 | 0.189290186 | -3.099789599 | 0.001936582 | 0.020843676 |
| GPR161 | 20.08741693 | 0.69311412 | 0.22363151 | 3.099358049 | 0.001939405 | 0.020861047 |
| MINK1 | 1395.250409 | -0.315023518 | 0.101646327 | -3.099212026 | 0.001940361 | 0.020861047 |
| RPS6KA4 | 332.7935289 | -0.270438795 | 0.087289091 | -3.098196954 | 0.00194702 | 0.020910396 |
| SOAT1 | 11085.38648 | 0.208439399 | 0.067277983 | 3.098181427 | 0.001947122 | 0.020910396 |
| AP2M1 | 6673.188112 | -0.210550684 | 0.068039435 | -3.094538972 | 0.00197119 | 0.021157077 |
| PXMP4 | 73.94162772 | -0.311438706 | 0.100750563 | -3.091185769 | 0.001993589 | 0.021373667 |
| RASAL2 | 812.0722322 | 0.197513414 | 0.06389427 | 3.091253932 | 0.001993131 | 0.021373667 |
| LY75 | 479.848243 | -0.368503716 | 0.119241267 | -3.090404235 | 0.001998843 | 0.021418075 |
| PTPMT1 | 1109.534726 | 0.174908088 | 0.056612184 | 3.089583853 | 0.002004371 | 0.021465378 |
| ACSS3 | 20.03584532 | -0.646951388 | 0.209410085 | -3.089399386 | 0.002005616 | 0.021466779 |
| SPATA7 | 38.53022866 | -0.435841251 | 0.141100407 | -3.088873088 | 0.002009172 | 0.021492902 |
| C4orf34 | 506.9944257 | -0.166578085 | 0.053940577 | -3.088177689 | 0.00201388 | 0.021531307 |
| CLMP | 5.009332083 | -1.417419408 | 0.459083437 | -3.087498465 | 0.002018488 | 0.021568604 |
| FLJ42393 | 44.34049112 | 0.641062967 | 0.207753948 | 3.085683677 | 0.002030847 | 0.02168864 |
| APEH | 471.5746994 | -0.216431551 | 0.070160731 | -3.084796133 | 0.002036917 | 0.021729373 |
| CISD2 | 823.1955259 | -0.275632625 | 0.089347385 | -3.084954586 | 0.002035832 | 0.021729373 |
| RPL36A | 1377.450449 | -0.254509254 | 0.082536904 | -3.083581307 | 0.002045252 | 0.021806213 |
| GTF2E2 | 185.0046943 | 0.244258599 | 0.079243716 | 3.082371859 | 0.002053581 | 0.021858727 |
| HTATIP2 | 423.2305573 | -0.262211596 | 0.085066619 | -3.082426447 | 0.002053205 | 0.021858727 |
| IL19 | 3.365531194 | 2.488887688 | 0.807415456 | 3.082536592 | 0.002052445 | 0.021858727 |
| ARHGAP22 | 74.89018538 | -0.378539335 | 0.122831662 | -3.081773282 | 0.002057715 | 0.021890633 |
| HELB | 67.96721974 | 0.398368163 | 0.129274297 | 3.081572852 | 0.002059101 | 0.021893287 |
| EGFL8 | 9.340713984 | -0.946325126 | 0.307226613 | -3.080218594 | 0.002068487 | 0.021968839 |
| PAK1 | 422.0328051 | -0.229960795 | 0.074656366 | -3.080257006 | 0.00206822 | 0.021968839 |
| MAML3 | 11.02062994 | -1.074723416 | 0.349084026 | -3.078695488 | 0.002079091 | 0.022069291 |
| GRAMD3 | 25.4833716 | 0.47158948 | 0.153229125 | 3.077675202 | 0.002086222 | 0.02213279 |
| SLC16A10 | 168.2797115 | -0.297180964 | 0.096578713 | -3.077085565 | 0.002090353 | 0.022164413 |
| TBC1D9B | 1416.638201 | -0.148631924 | 0.048326512 | -3.075577319 | 0.002100954 | 0.02226457 |
| EIF2S1 | 456.76123 | 0.195609352 | 0.063626118 | 3.074356215 | 0.002109574 | 0.022322474 |
| LOC10050( | 151.151749 | -0.43415409 | 0.141214966 | -3.074419826 | 0.002109124 | 0.022322474 |
| ZCCHC10 | 237.5351343 | 0.193087416 | 0.062806731 | 3.074310884 | 0.002109894 | 0.022322474 |
| CASP6 | 182.4324393 | -0.236900073 | 0.077099172 | -3.072666886 | 0.002121552 | 0.022421182 |
| SLC47A1 | 569.1100576 | -0.272721233 | 0.088754642 | -3.072754596 | 0.002120928 | 0.022421182 |
| LOC64551: | 45.62358063 | -0.38965677 | 0.126860338 | -3.071541331 | 0.002129567 | 0.022493549 |
| COX5B | 1612.494609 | -0.225077358 | 0.07329213 | -3.07096216 | 0.002133702 | 0.022524877 |
| MME | 42.87388687 | -0.302242402 | 0.098450951 | -3.069979524 | 0.002140734 | 0.022586742 |
| CCL23 | 1004.895712 | -0.445969314 | 0.145310333 | -3.069081919 | 0.002147177 | 0.022642317 |
| THBS1 | 765.8219943 | -0.393853967 | 0.128347047 | -3.068664039 | 0.002150182 | 0.022661606 |

| | | | | | | |
|---|---|---|---|---|---|---|
| ZNF629 | 47.12864361 | -0.425035639 | 0.138540506 | -3.067952125 | 0.002155311 | 0.022703242 |
| VAC14 | 3085.377442 | 0.178225669 | 0.058127846 | 3.066097966 | 0.002168722 | 0.022832026 |
| GPAA1 | 800.5195624 | -0.225564779 | 0.073574974 | -3.065781295 | 0.002171021 | 0.022843737 |
| PYCARD | 519.2212747 | -0.27419807 | 0.089467291 | -3.064785636 | 0.002178261 | 0.022907407 |
| C18orf8 | 315.9922495 | 0.325404033 | 0.106216741 | 3.06358519 | 0.002187019 | 0.022986968 |
| TBX20 | 64.06222741 | 0.455143121 | 0.148587572 | 3.063130473 | 0.002190345 | 0.023009375 |
| NR1H4 | 2.854739323 | 2.651393149 | 0.866047853 | 3.061485735 | 0.002202415 | 0.023123556 |
| CLDND1 | 761.5473505 | 0.420927695 | 0.137508777 | 3.061096936 | 0.002205277 | 0.023140993 |
| DGKE | 23.56356499 | 0.534843911 | 0.17488472 | 3.058265524 | 0.002226222 | 0.023322674 |
| GSPT2 | 59.20275904 | -0.317225125 | 0.103718808 | -3.058511088 | 0.002224398 | 0.023322674 |
| TUBB2A | 698.0383134 | -0.178293439 | 0.058295871 | -3.058423104 | 0.002225052 | 0.023322674 |
| GSG2 | 62.19313463 | -0.519505028 | 0.169924381 | -3.057271855 | 0.002233616 | 0.023387416 |
| TMC1 | 3.955715668 | 1.513909575 | 0.49539983 | 3.055934791 | 0.0022436 | 0.023479199 |
| LILRB4 | 3873.173105 | -0.238580116 | 0.078125533 | -3.053804635 | 0.002259592 | 0.023633711 |
| DOCK8 | 1714.751316 | 0.17779245 | 0.058245543 | 3.052464449 | 0.002269706 | 0.023726619 |
| ANP32A | 855.6965923 | -0.187398561 | 0.061409607 | -3.051616346 | 0.002276128 | 0.023780849 |
| FBL | 366.539739 | -0.260521414 | 0.085388693 | -3.051005998 | 0.00228076 | 0.023816328 |
| FNDC3B | 6113.521759 | 0.218286438 | 0.07155426 | 3.050642098 | 0.002283526 | 0.023832292 |
| NPLOC4 | 1347.650338 | 0.23605242 | 0.077409338 | 3.049404961 | 0.002292952 | 0.023917708 |
| ABHD14B | 296.130847 | -0.289165156 | 0.094932244 | -3.046016241 | 0.002318953 | 0.024175839 |
| NLK | 433.9387372 | 0.273529533 | 0.089842355 | 3.044549893 | 0.002330288 | 0.024280867 |
| CHI3L2 | 31.00963403 | 0.625963817 | 0.205621599 | 3.044251288 | 0.002332602 | 0.024291845 |
| GAS7 | 304.3498875 | -0.288545425 | 0.094841423 | -3.042398718 | 0.002347008 | 0.024415471 |
| POU2F2 | 10.72724171 | 0.90152687 | 0.296306242 | 3.042551057 | 0.00234582 | 0.024415471 |
| TPD52L1 | 5.858716072 | -1.532187984 | 0.503978958 | -3.040182449 | 0.002364349 | 0.024582592 |
| NCKAP5 | 2.981786799 | 2.063969621 | 0.679470994 | 3.037612552 | 0.002384603 | 0.024768924 |
| REXO2 | 163.7604825 | -0.250178653 | 0.082361106 | -3.037582472 | 0.002384841 | 0.024768924 |
| INTS6 | 234.8225466 | 0.471896304 | 0.155367049 | 3.037299779 | 0.002387079 | 0.024778809 |
| PPIL1 | 50.15808595 | 0.524063986 | 0.172555129 | 3.037081477 | 0.002388809 | 0.024783409 |
| TFEB | 282.528706 | -0.301232907 | 0.099260336 | -3.034776203 | 0.002407142 | 0.024960174 |
| NDUFA2 | 592.6237171 | -0.217209739 | 0.071579048 | -3.034543569 | 0.002409 | 0.024965996 |
| RAD51AP1 | 40.93108865 | -0.429914444 | 0.141723298 | -3.033477562 | 0.002417527 | 0.025040899 |
| SLC25A45 | 74.01542934 | -0.330256678 | 0.108892578 | -3.032866745 | 0.002422425 | 0.025078156 |
| IER5 | 2197.947276 | 0.266030832 | 0.087721292 | 3.032682547 | 0.002423904 | 0.025079991 |
| CLN6 | 939.8599853 | -0.259057227 | 0.085428378 | -3.032449325 | 0.002425778 | 0.025085906 |
| PCCB | 169.8231304 | -0.265964735 | 0.087716143 | -3.032107045 | 0.002428531 | 0.025100897 |
| GPR64 | 203.940869 | 0.299689331 | 0.09885124 | 3.03172051 | 0.002431642 | 0.025107347 |
| MRPL55 | 128.7693463 | -0.259760299 | 0.085681236 | -3.031705776 | 0.002431761 | 0.025107347 |
| CDC42SE1 | 2021.83736 | 0.213173467 | 0.070356169 | 3.029918606 | 0.002446197 | 0.025242861 |
| CHST15 | 100.9556961 | 0.450088102 | 0.148558026 | 3.02971245 | 0.002447867 | 0.025246574 |
| PRPF4B | 953.0307048 | 0.145690862 | 0.048100006 | 3.028915694 | 0.002454332 | 0.025299707 |
| SH3PXD2B | 432.7799164 | 0.403485067 | 0.133230732 | 3.028468442 | 0.002457967 | 0.025323637 |
| TPRG1L | 587.7414409 | -0.149834579 | 0.049479809 | -3.028196393 | 0.002460181 | 0.0253329 |
| PPP6R3 | 729.0337582 | 0.168180275 | 0.055542893 | 3.027935101 | 0.00246231 | 0.02534127 |
| LYL1 | 156.4526506 | -0.26312586 | 0.086926296 | -3.026999569 | 0.002469943 | 0.025406259 |
| PI4K2B | 121.5057643 | 0.387661954 | 0.128140346 | 3.025291923 | 0.002483933 | 0.025533107 |
| VGLL3 | 55.98785717 | 0.958211253 | 0.316746141 | 3.025171041 | 0.002484926 | 0.025533107 |

| | | | | | | |
|---|---|---|---|---|---|---|
| SMAGP | 6.221301757 | 1.453709167 | 0.48072206 | 3.024011771 | 0.002494467 | 0.025617488 |
| CREBZF | 230.6289806 | -0.25043929 | 0.082846228 | -3.022941369 | 0.002503307 | 0.025694576 |
| EPDR1 | 143.3476503 | -0.331861407 | 0.10988037 | -3.020206508 | 0.002526024 | 0.025913939 |
| TTLL11 | 33.83236967 | -0.500681167 | 0.165949153 | -3.017075753 | 0.00255226 | 0.026169157 |
| ICMT | 486.4869484 | -0.217932939 | 0.072271625 | -3.015470302 | 0.002565811 | 0.026256975 |
| KIAA2013 | 992.8889922 | -0.187772971 | 0.062262565 | -3.015824556 | 0.002562815 | 0.026256975 |
| PSMB8 | 465.4085519 | 0.466176314 | 0.154597718 | 3.015415237 | 0.002566277 | 0.026256975 |
| SSX1 | 636.2912718 | -0.222305038 | 0.073714999 | -3.015736847 | 0.002563556 | 0.026256975 |
| CCL7 | 113.0336391 | 0.608202768 | 0.201747755 | 3.01466932 | 0.002572596 | 0.026307659 |
| EBPL | 43.93377328 | -0.647243263 | 0.214796691 | -3.013283209 | 0.002584376 | 0.026386111 |
| KIAA0922 | 306.3018266 | -0.275953539 | 0.091575298 | -3.013405856 | 0.002583332 | 0.026386111 |
| NRAS | 1176.159717 | 0.225621482 | 0.074870968 | 3.013470888 | 0.002582778 | 0.026386111 |
| CHD6 | 69.49935227 | -0.431457228 | 0.143281949 | -3.011246223 | 0.002601778 | 0.02654971 |
| DAPK2 | 4.412871538 | -1.299592182 | 0.43167213 | -3.010600155 | 0.002607319 | 0.026592174 |
| SLC35E2B | 198.7950651 | -0.245696674 | 0.081622264 | -3.010167346 | 0.002611038 | 0.026616008 |
| ZNF438 | 275.6023062 | 0.216236851 | 0.071867791 | 3.008814501 | 0.002622692 | 0.026720668 |
| DSTN | 1896.015343 | -0.139507171 | 0.046393707 | -3.007027882 | 0.002638156 | 0.02686401 |
| SLX4 | 103.1478028 | 0.343653333 | 0.114375703 | 3.004600831 | 0.002659296 | 0.027064974 |
| C21orf7 | 55.06604539 | 0.46852731 | 0.155990412 | 3.003564789 | 0.002668367 | 0.027142959 |
| JHDM1D | 249.2179829 | 0.269305505 | 0.08971102 | 3.001922214 | 0.002682807 | 0.027275442 |
| C6orf48 | 340.0468033 | -0.228935206 | 0.076278253 | -3.001316852 | 0.002688147 | 0.027315316 |
| DDR1 | 23.75111146 | 0.621247263 | 0.207062406 | 3.000289985 | 0.002697227 | 0.027393132 |
| MGC1291( | 115.9252924 | 0.648168494 | 0.216074607 | 2.999743948 | 0.002702067 | 0.027427826 |
| KBTBD3 | 30.44907088 | -0.517117453 | 0.172417677 | -2.999213666 | 0.002706774 | 0.027455211 |
| RNF7 | 1051.53765 | -0.196505976 | 0.065521231 | -2.999119095 | 0.002707614 | 0.027455211 |
| NDUFB2 | 581.2045666 | -0.204204222 | 0.068100108 | -2.998588829 | 0.002712331 | 0.027488566 |
| DDB2 | 159.1206336 | -0.24221602 | 0.080788514 | -2.998149201 | 0.002716247 | 0.027513779 |
| DBF4 | 194.0563568 | 0.319497297 | 0.106693385 | 2.994537073 | 0.002748617 | 0.027827034 |
| IDI1 | 328.4468166 | 0.325992758 | 0.108888771 | 2.993814281 | 0.002755136 | 0.027878387 |
| TNIK | 1091.698781 | -0.202297107 | 0.06757951 | -2.993468073 | 0.002758264 | 0.027895385 |
| DDX59 | 250.4106587 | 0.22061921 | 0.073716707 | 2.992797962 | 0.002764327 | 0.027942036 |
| CCDC50 | 856.8138044 | 0.270212702 | 0.090300001 | 2.99238869 | 0.002768036 | 0.027964855 |
| SPG7 | 314.2680601 | -0.229646137 | 0.076779186 | -2.990994686 | 0.002780703 | 0.028078108 |
| MS4A6A | 57.26983782 | -0.392295372 | 0.131203583 | -2.989974534 | 0.002790007 | 0.028157295 |
| KBTBD7 | 75.0471175 | -0.374450292 | 0.125254525 | -2.989515076 | 0.002794207 | 0.028184913 |
| C7orf50 | 446.5505107 | -0.332284485 | 0.11116983 | -2.988980784 | 0.002799097 | 0.028204712 |
| PHTF2 | 362.0818703 | 0.333775322 | 0.111665212 | 2.989071661 | 0.002798265 | 0.028204712 |
| SLC22A5 | 138.3639163 | -0.2668003 | 0.089283478 | -2.988238205 | 0.002805908 | 0.028258556 |
| PAQR4 | 30.28971149 | -0.482695304 | 0.161554161 | -2.987823409 | 0.002809718 | 0.028282151 |
| DIS3L | 237.3739267 | -0.405334529 | 0.135692167 | -2.987162334 | 0.002815802 | 0.028313796 |
| NR1D1 | 81.077635 | 0.336301997 | 0.112578688 | 2.987261662 | 0.002814887 | 0.028313796 |
| PCYT1A | 839.2314562 | 0.212182462 | 0.071056261 | 2.986119154 | 0.002825425 | 0.028395745 |
| ABCA6 | 27.85413398 | 0.710724281 | 0.238134989 | 2.984543704 | 0.002840016 | 0.028527504 |
| GLB1 | 1150.104089 | -0.23266316 | 0.077969999 | -2.984008771 | 0.002844986 | 0.028562534 |
| RARRES3 | 305.3214643 | 0.643544169 | 0.215826132 | 2.981771315 | 0.00286586 | 0.028757112 |
| RPL29 | 5028.573228 | -0.189969767 | 0.063762228 | -2.979346439 | 0.00288864 | 0.028970606 |
| KCNAB2 | 2013.026761 | -0.317497002 | 0.106572563 | -2.979162675 | 0.002890373 | 0.028972905 |

| | | | | | | |
|---|---|---|---|---|---|---|
| RIN3 | 1239.696864 | -0.249159244 | 0.083643508 | -2.97882346 | 0.002893574 | 0.028989914 |
| MAPK3 | 158.8252194 | -0.31603667 | 0.106153785 | -2.977158742 | 0.002909333 | 0.029132648 |
| PPP1R15A | 3099.007601 | 0.300114559 | 0.100860776 | 2.975532923 | 0.0029248 | 0.029272305 |
| ARHGAP18 | 1218.786813 | 0.17043297 | 0.057288526 | 2.974993121 | 0.002929951 | 0.029308639 |
| CNKSR3 | 10.32313137 | 1.083687015 | 0.364581418 | 2.972414287 | 0.002954677 | 0.029540636 |
| RIT1 | 1125.417444 | 0.169128128 | 0.05693393 | 2.97060344 | 0.002972153 | 0.029699947 |
| C2CD2 | 127.194172 | -0.278054601 | 0.093692287 | -2.967742709 | 0.002999953 | 0.029962209 |
| TMEM107 | 145.5870817 | -0.24655759 | 0.08309534 | -2.967165047 | 0.003005596 | 0.030003009 |
| CLSTN1 | 477.2471168 | -0.21101185 | 0.07115121 | -2.96568182 | 0.003020128 | 0.030116863 |
| CMTM8 | 6.714459169 | -1.212738475 | 0.408910853 | -2.965777177 | 0.003019191 | 0.030116863 |
| TMEM199 | 369.5900243 | 0.158218169 | 0.053362153 | 2.964988475 | 0.003026943 | 0.030169207 |
| SZRD1 | 3201.704917 | 0.189326204 | 0.063865842 | 2.964436048 | 0.003032383 | 0.030207799 |
| LOC73010[illegible] | 25.90885715 | -0.51840984 | 0.174906678 | -2.963922508 | 0.003037448 | 0.030242618 |
| AKR1A1 | 1685.565885 | -0.19935958 | 0.067286527 | -2.962845428 | 0.003048096 | 0.030332963 |
| FOXRED2 | 14.04042813 | -0.821883275 | 0.277587844 | -2.960804275 | 0.003068369 | 0.030503194 |
| TALDO1 | 20205.19952 | -0.21723262 | 0.07336839 | -2.960847595 | 0.003067937 | 0.030503194 |
| PHLDB1 | 29.6254062 | 0.481664854 | 0.162699695 | 2.960453341 | 0.003071866 | 0.030522217 |
| CSRNP2 | 49.65997717 | 0.419952191 | 0.14193317 | 2.958802308 | 0.003088372 | 0.030670395 |
| LOC64563[illegible] | 211.5083755 | 0.280029241 | 0.094684081 | 2.957511333 | 0.003101333 | 0.030783251 |
| SLC9A7 | 905.2620286 | -0.201684339 | 0.068250969 | -2.955039926 | 0.003126286 | 0.031014945 |
| FN3K | 27.45962077 | -0.55254136 | 0.187026043 | -2.95435518 | 0.003133232 | 0.031067855 |
| RARRES1 | 91.4570736 | -0.428790458 | 0.145192157 | -2.953261853 | 0.003144351 | 0.031162073 |
| TMEM14E | 157.7994188 | 0.319930331 | 0.108398675 | 2.951422907 | 0.003163135 | 0.031332112 |
| AGAP3 | 1949.253637 | -0.245751079 | 0.083302372 | -2.950109012 | 0.003176618 | 0.0314495 |
| TRPC6 | 29.25365688 | -0.530848753 | 0.179982178 | -2.949451774 | 0.003183383 | 0.031500281 |
| LOC10013[illegible] | 25.29966385 | -0.530049133 | 0.179742212 | -2.948940751 | 0.003188651 | 0.031536217 |
| TRIM14 | 1643.110055 | 0.278331175 | 0.094398981 | 2.948455276 | 0.003193663 | 0.031569585 |
| SERTAD2 | 288.0650211 | 0.309449053 | 0.104960421 | 2.948245151 | 0.003195835 | 0.031574852 |
| ACOT11 | 43.46463614 | 0.420492971 | 0.142652605 | 2.94767117 | 0.003201774 | 0.031617317 |
| EIF4EBP2 | 2543.831924 | -0.220700568 | 0.074892632 | -2.946892943 | 0.003209843 | 0.03168045 |
| EIF5 | 1500.413639 | 0.22920018 | 0.077780995 | 2.946737559 | 0.003211456 | 0.03168045 |
| UNC119 | 239.3630384 | -0.191105387 | 0.06488071 | -2.945488545 | 0.003224451 | 0.03179236 |
| CD300E | 162.0316724 | 0.446366653 | 0.15159852 | 2.944399801 | 0.003235817 | 0.031888108 |
| HDAC9 | 18.53517548 | 0.623880585 | 0.212011011 | 2.942680101 | 0.003253845 | 0.032049371 |
| DUSP14 | 104.0662058 | -0.312446716 | 0.106206075 | -2.941891178 | 0.003262145 | 0.032114713 |
| BDH2 | 51.15241256 | -0.409141638 | 0.139084826 | -2.94166984 | 0.003264478 | 0.03212126 |
| DNAJC9 | 128.7003022 | -0.279523866 | 0.095059836 | -2.940504401 | 0.003276783 | 0.032225886 |
| RBM47 | 4356.007935 | 0.235630406 | 0.080144669 | 2.940063395 | 0.003281451 | 0.032255324 |
| UPB1 | 4.335791028 | 1.631296637 | 0.554885753 | 2.939878395 | 0.003283411 | 0.03225813 |
| SSH2 | 1083.614817 | -0.240612205 | 0.08185339 | -2.939550877 | 0.003286883 | 0.032275784 |
| TMEM206 | 624.2095061 | 0.216687088 | 0.073747628 | 2.938224494 | 0.003300979 | 0.032397688 |
| SH3BGRL | 3024.015281 | -0.266282266 | 0.090659236 | -2.937177484 | 0.003312145 | 0.032490724 |
| ZADH2 | 71.15916437 | -0.381513258 | 0.129928394 | -2.936334756 | 0.003321157 | 0.032562551 |
| AOAH | 8.251252446 | -0.876620361 | 0.298730367 | -2.934486939 | 0.003340996 | 0.032740403 |
| RPS12 | 5842.00042 | -0.171716992 | 0.058522834 | -2.934187924 | 0.003344217 | 0.032755301 |
| RTP4 | 150.7413594 | 0.469877869 | 0.160241818 | 2.932304913 | 0.003364562 | 0.032937834 |
| SMARCA1 | 6.210520644 | 1.063182406 | 0.362691492 | 2.931368476 | 0.003374722 | 0.033020518 |

| | | | | | | |
|---|---|---|---|---|---|---|
| FTL | 156066.0227 | -0.241311735 | 0.082339719 | -2.930684452 | 0.003382161 | 0.033076508 |
| ANGPTL4 | 14.52407821 | 0.93203588 | 0.318120279 | 2.929822276 | 0.003391559 | 0.033151588 |
| DDA1 | 1459.399735 | 0.155959909 | 0.053241766 | 2.92927753 | 0.003397509 | 0.033192907 |
| CREBRF | 1000.274103 | -0.244516594 | 0.083570583 | -2.925869208 | 0.003434953 | 0.033541717 |
| TTC33 | 173.8714764 | 0.208799971 | 0.071417967 | 2.923633624 | 0.003459717 | 0.033766417 |
| ZMYND8 | 148.2075102 | 0.228595998 | 0.078199542 | 2.923239601 | 0.003464098 | 0.033792061 |
| ANXA9 | 7.164247718 | -1.157427766 | 0.396021686 | -2.922637341 | 0.003470805 | 0.03384035 |
| GUCY1B2 | 9.054118041 | -0.908812457 | 0.310993117 | -2.922291227 | 0.003474665 | 0.033843727 |
| VPS37B | 673.8082618 | 0.243301954 | 0.083252861 | 2.922445568 | 0.003472943 | 0.033843727 |
| ATG4A | 181.406607 | 0.276614557 | 0.094717412 | 2.920419269 | 0.003495608 | 0.033996788 |
| EAPP | 249.6403025 | -0.266689121 | 0.091309571 | -2.920713754 | 0.003492305 | 0.033996788 |
| FES | 66.73670627 | -0.399549556 | 0.136812669 | -2.920413423 | 0.003495673 | 0.033996788 |
| IL18BP | 137.3543223 | 0.547199822 | 0.187403829 | 2.919896695 | 0.003501474 | 0.034018863 |
| MPST | 134.5803486 | -0.31891621 | 0.10921606 | -2.920048677 | 0.003499767 | 0.034018863 |
| ANKRD33B | 4.223258813 | 1.481672966 | 0.507563108 | 2.919189638 | 0.003509427 | 0.034078938 |
| TOP2A | 7.420910679 | -1.052891775 | 0.361071828 | -2.91601752 | 0.003545306 | 0.034410007 |
| GLIPR1 | 571.5422627 | -0.434112039 | 0.148947185 | -2.914536712 | 0.003562169 | 0.034556269 |
| GAL | 41.75257963 | -0.491447698 | 0.168649362 | -2.9140205 | 0.003568065 | 0.034596042 |
| MMP2 | 45.64819958 | -0.419212099 | 0.143869695 | -2.913831846 | 0.003570222 | 0.034599542 |
| FAM78A | 549.4762746 | -0.297334532 | 0.102066295 | -2.913151029 | 0.003578015 | 0.034657636 |
| ATP8B1 | 28.79335888 | -0.705277759 | 0.242144829 | -2.912627792 | 0.003584015 | 0.03469831 |
| CDIPT | 813.1228902 | -0.23298521 | 0.080010105 | -2.91194732 | 0.003591832 | 0.034756523 |
| CCND3 | 908.933466 | 0.264446093 | 0.090878315 | 2.909892133 | 0.003615535 | 0.034944251 |
| SHMT1 | 24.24400592 | -0.584384456 | 0.200840457 | -2.90969492 | 0.003617817 | 0.034944251 |
| TCEAL1 | 16.2355268 | -0.475169206 | 0.163295802 | -2.909867861 | 0.003615816 | 0.034944251 |
| ZNF93 | 50.71306362 | -0.315587211 | 0.108462743 | -2.909637003 | 0.003618488 | 0.034944251 |
| ABL2 | 1740.059641 | 0.395204237 | 0.135880083 | 2.908478034 | 0.003631927 | 0.03503891 |
| SLTM | 1069.330872 | 0.175335689 | 0.060283281 | 2.908529279 | 0.003631332 | 0.03503891 |
| FOXO1 | 130.118046 | -0.237831298 | 0.081779403 | -2.908205357 | 0.003635095 | 0.035051926 |
| FUK | 99.40496509 | -0.306340232 | 0.105366901 | -2.907366831 | 0.003644855 | 0.035128452 |
| DNAJC7 | 1062.478172 | 0.196781413 | 0.067710226 | 2.906228865 | 0.003658138 | 0.035238841 |
| PYURF | 695.0097168 | -0.145520979 | 0.050083566 | -2.905563476 | 0.003665925 | 0.035296206 |
| BAHCC1 | 14.25703261 | -0.627863564 | 0.216195055 | -2.904153215 | 0.003682479 | 0.035437882 |
| PDGFA | 31.03201191 | 0.495860874 | 0.170770945 | 2.903660647 | 0.003688277 | 0.035475958 |
| MAT2A | 1813.792215 | 0.20862858 | 0.071860033 | 2.90326309 | 0.003692963 | 0.035503302 |
| LRP5L | 16.62040062 | -0.592473817 | 0.20413841 | -2.902314254 | 0.003704168 | 0.035575519 |
| VEGFB | 582.4634727 | -0.292851614 | 0.100899362 | -2.902412941 | 0.003703001 | 0.035575519 |
| GNB5 | 40.79374393 | -0.569090224 | 0.196114558 | -2.901825487 | 0.003709951 | 0.035613316 |
| ARFIP2 | 176.259028 | 0.232997411 | 0.080310775 | 2.901197395 | 0.003717396 | 0.035660038 |
| PORCN | 74.50477862 | -0.296250881 | 0.102122138 | -2.900946705 | 0.003720371 | 0.035660038 |
| RAB31 | 4494.181159 | 0.256780737 | 0.088514685 | 2.900995882 | 0.003719788 | 0.035660038 |
| AOC2 | 65.80138508 | 0.398465597 | 0.137465772 | 2.898653187 | 0.003747692 | 0.035899913 |
| HSP90B1 | 11063.09856 | 0.194251269 | 0.067017089 | 2.898533374 | 0.003749124 | 0.035899913 |
| MSANTD3 | 844.9108578 | 0.274873083 | 0.09488729 | 2.896837738 | 0.003769447 | 0.036076582 |
| FCGR2B | 5.831613147 | 1.457611814 | 0.503265035 | 2.896310519 | 0.003775786 | 0.036109814 |
| USP7 | 1241.731081 | 0.140018488 | 0.048344965 | 2.896237257 | 0.003776667 | 0.036109814 |
| SUCO | 259.6422967 | 0.248582322 | 0.085900929 | 2.893825765 | 0.003805793 | 0.03637024 |

| | | | | | | |
|---|---|---|---|---|---|---|
| RPS6 | 4068.761649 | -0.164134751 | 0.056724324 | -2.893551463 | 0.003809119 | 0.036383976 |
| S1PR4 | 98.4008006 | -0.328664514 | 0.113601208 | -2.893142782 | 0.003814079 | 0.0364133 |
| FUBP1 | 759.2835703 | 0.234338273 | 0.081012112 | 2.892632565 | 0.003820279 | 0.036454433 |
| C6orf1 | 427.1660222 | -0.245945647 | 0.085036072 | -2.892250807 | 0.003824925 | 0.036480693 |
| FAM109B | 82.06722236 | -0.315369317 | 0.109144245 | -2.889472713 | 0.003858885 | 0.036786382 |
| MICALCL | 37.61067952 | 0.642241201 | 0.222641489 | 2.884642951 | 0.003918578 | 0.037336956 |
| MAP3K6 | 47.9235477 | -0.423407014 | 0.146876086 | -2.88274985 | 0.003942204 | 0.037514392 |
| PVRL2 | 404.5227705 | 0.298702424 | 0.103619927 | 2.882673552 | 0.003943159 | 0.037514392 |
| SNX20 | 146.2686531 | -0.298047682 | 0.103391768 | -2.882702261 | 0.003942799 | 0.037514392 |
| UBE2Q1 | 568.5382866 | 0.157703317 | 0.054710085 | 2.882527341 | 0.003944989 | 0.037514392 |
| ZBTB7A | 1091.953245 | 0.217830643 | 0.075610343 | 2.880963547 | 0.003964615 | 0.037682423 |
| SNX9 | 108.6155686 | 0.277242716 | 0.096251432 | 2.880400961 | 0.003971697 | 0.037731123 |
| ANAPC15 | 144.7649533 | -0.329817388 | 0.114565946 | -2.878843146 | 0.003991368 | 0.037898591 |
| SNRPG | 189.5637206 | 0.204086126 | 0.070895394 | 2.878693706 | 0.00399326 | 0.037898591 |
| AIFM2 | 89.80846513 | 0.317698657 | 0.110404602 | 2.877585286 | 0.004007316 | 0.038013267 |
| FAM190B | 550.8987502 | 0.201007595 | 0.0698674 | 2.876986911 | 0.004014922 | 0.038066681 |
| ABHD6 | 462.9903045 | -0.225286455 | 0.078328661 | -2.876168844 | 0.004025343 | 0.038146711 |
| SS18 | 744.6535977 | 0.267887921 | 0.093205287 | 2.874170879 | 0.004050897 | 0.038370003 |
| APOE | 29349.31247 | -0.244075853 | 0.084925412 | -2.874002565 | 0.004053057 | 0.038371593 |
| IQCC | 11.43856742 | -0.712721749 | 0.24826634 | -2.870794924 | 0.004094411 | 0.038744064 |
| NUDCD1 | 132.4652644 | 0.24221985 | 0.084394308 | 2.870096997 | 0.004103459 | 0.038810624 |
| UHRF1BP1 | 405.7884458 | 0.205437348 | 0.07159513 | 2.869431875 | 0.004112099 | 0.038873258 |
| CPNE2 | 390.9179583 | -0.241252558 | 0.084111162 | -2.86825852 | 0.004127381 | 0.038960377 |
| TLR5 | 34.69423839 | -0.679570011 | 0.23690842 | -2.868492429 | 0.004124331 | 0.038960377 |
| YIPF6 | 594.5703486 | 0.220447824 | 0.076856234 | 2.868314166 | 0.004126655 | 0.038960377 |
| LOC10028[illegible] | 316.1616998 | -0.182409435 | 0.063603216 | -2.867927849 | 0.004131697 | 0.038982019 |
| REEP6 | 8.294948073 | -0.965448541 | 0.336678888 | -2.867564835 | 0.00413644 | 0.039007666 |
| CUL1 | 845.3402552 | 0.190227132 | 0.066362442 | 2.866487819 | 0.004150541 | 0.039121492 |
| NOXA1 | 12.40484156 | -0.704269309 | 0.245766298 | -2.865605724 | 0.004162123 | 0.03921147 |
| CCDC88C | 15.56367945 | -0.661851303 | 0.230987848 | -2.865307882 | 0.00416604 | 0.03921169 |
| LSM6 | 161.6608534 | -0.296778668 | 0.103577023 | -2.865294443 | 0.004166217 | 0.03921169 |
| SAFB | 720.5416976 | 0.106598876 | 0.037205813 | 2.865113491 | 0.004168598 | 0.039214947 |
| MLLT11 | 11.95601532 | 0.614738584 | 0.214821073 | 2.861630733 | 0.004214676 | 0.039629066 |
| NUDT7 | 7.999292517 | -0.784928477 | 0.274385014 | -2.860682752 | 0.004227298 | 0.039728356 |
| MAZ | 2349.280647 | -0.231344003 | 0.080884424 | -2.860179889 | 0.004234008 | 0.039733264 |
| MED11 | 138.6185754 | -0.305995423 | 0.106981988 | -2.860251779 | 0.004233048 | 0.039733264 |
| ST3GAL2 | 866.5874216 | 0.302264139 | 0.105670084 | 2.860451397 | 0.004230384 | 0.039733264 |
| FBXL4 | 127.7875281 | -0.274000147 | 0.095854318 | -2.858506032 | 0.00425641 | 0.03992405 |
| PSIP1 | 48.40553329 | -0.381156798 | 0.133384805 | -2.857572855 | 0.004268946 | 0.04002215 |
| CDKL1 | 65.70073792 | 0.302190916 | 0.105812075 | 2.855920899 | 0.00429122 | 0.040211406 |
| ZC3H3 | 132.9343898 | 0.315093194 | 0.110347425 | 2.855464852 | 0.004297388 | 0.040249624 |
| CDK17 | 418.3786155 | 0.243612603 | 0.085336721 | 2.854721862 | 0.004307453 | 0.04030471 |
| CYBASC3 | 590.8537679 | -0.266058691 | 0.093195067 | -2.854858077 | 0.004305606 | 0.04030471 |
| ANKRD6 | 44.16818574 | -0.309886516 | 0.108562573 | -2.854450735 | 0.004311132 | 0.040319546 |
| C17orf100 | 21.39964062 | -0.68632583 | 0.240462683 | -2.854188527 | 0.004314692 | 0.040333262 |
| FAM127C | 61.2288777 | -0.368538653 | 0.129156687 | -2.853422932 | 0.004325102 | 0.040410965 |
| HK2 | 3088.559117 | 0.135500192 | 0.047520634 | 2.851396968 | 0.004352759 | 0.040649663 |

| | | | | | | |
|---|---|---|---|---|---|---|
| OR6N2 | 24.30796551 | -0.775253335 | 0.271950908 | -2.850710595 | 0.004362165 | 0.04071777 |
| UBE2M | 1124.162068 | 0.228755851 | 0.080253196 | 2.850426693 | 0.004366061 | 0.040734401 |
| EZH2 | 140.8992504 | 0.494513846 | 0.173542582 | 2.849524533 | 0.004378463 | 0.0408234 |
| HSP90AA1 | 13222.76596 | 0.220138438 | 0.077257155 | 2.849424611 | 0.004379838 | 0.0408234 |
| ST5 | 15.42628931 | -0.642566574 | 0.225525185 | -2.849200964 | 0.004382919 | 0.040832355 |
| COMMD9 | 417.4400664 | -0.252435936 | 0.088609229 | -2.848867322 | 0.004387517 | 0.040855442 |
| PNPLA7 | 6.070827235 | -1.017950935 | 0.357531378 | -2.847165304 | 0.004411045 | 0.041054683 |
| IFFO1 | 244.6129658 | -0.263515102 | 0.092580761 | -2.846326803 | 0.004422678 | 0.041143078 |
| STAP2 | 51.99119473 | -0.489878288 | 0.172119521 | -2.846151823 | 0.004425109 | 0.041145827 |
| PTGER3 | 48.87887079 | 0.556949043 | 0.195768248 | 2.844940624 | 0.00444197 | 0.041282681 |
| EIF2C1 | 200.3097459 | -0.288196628 | 0.101316513 | -2.844517827 | 0.00444787 | 0.041317578 |
| RPS11 | 12986.6518 | -0.191945308 | 0.067525053 | -2.842579139 | 0.004475012 | 0.041549677 |
| UBE2N | 1091.712731 | 0.203590064 | 0.071655851 | 2.841220383 | 0.004494124 | 0.041707033 |
| SEC61G | 2076.383301 | 0.203203705 | 0.071535249 | 2.840609462 | 0.004502742 | 0.041766886 |
| PDGFRL | 3.304313232 | 2.036556126 | 0.717184915 | 2.839652764 | 0.004516266 | 0.04187218 |
| CSF2RA | 1846.807072 | 0.183395064 | 0.064595554 | 2.839128266 | 0.004523697 | 0.041920897 |
| EIF2C3 | 393.2681021 | 0.22291403 | 0.078519592 | 2.838960621 | 0.004526074 | 0.041922762 |
| UBE3C | 346.0712414 | 0.276807937 | 0.097546212 | 2.837710768 | 0.004543834 | 0.042067034 |
| MFGE8 | 234.5326966 | -0.352682981 | 0.12431204 | -2.837078216 | 0.004552846 | 0.042130224 |
| PON2 | 34.29357336 | -0.717067626 | 0.252787391 | -2.836643172 | 0.004559053 | 0.042167413 |
| NUMB | 4181.315769 | 0.182428514 | 0.064330086 | 2.835819532 | 0.004570827 | 0.04225602 |
| CD14 | 3176.708984 | -0.373402726 | 0.131686133 | -2.835550853 | 0.004574673 | 0.042271296 |
| MCM4 | 201.9058716 | 0.337106475 | 0.118895923 | 2.835307263 | 0.004578163 | 0.042283264 |
| HAUS3 | 2.861678863 | 1.358144457 | 0.479039425 | 2.835141297 | 0.004580542 | 0.042284967 |
| SWAP70 | 1174.003799 | -0.232154648 | 0.081899962 | -2.834612421 | 0.004588131 | 0.042334739 |
| CENPF | 16.64893644 | -0.585907783 | 0.206809896 | -2.833074218 | 0.004610268 | 0.042518633 |
| ITFG3 | 315.1053785 | -0.202626638 | 0.07158329 | -2.830641586 | 0.004645475 | 0.042822829 |
| NECAP1 | 928.6753381 | 0.222822024 | 0.078724232 | 2.830412161 | 0.004648807 | 0.042833058 |
| TCFL5 | 19.28929331 | -0.756463815 | 0.267282961 | -2.83019843 | 0.004651914 | 0.042841195 |
| ETFB | 231.5930338 | -0.29367209 | 0.103770972 | -2.830002315 | 0.004654767 | 0.042846983 |
| CLN3 | 566.4023545 | -0.172531825 | 0.060984931 | -2.829089461 | 0.004668065 | 0.042948871 |
| DOCK3 | 101.9854877 | 0.375545593 | 0.132801928 | 2.82786251 | 0.004685993 | 0.043072679 |
| MCM5 | 261.8117686 | -0.306195785 | 0.108274989 | -2.827945644 | 0.004684776 | 0.043072679 |
| TMEM176I | 1117.828834 | -0.382827788 | 0.135453886 | -2.826259183 | 0.004709514 | 0.043268239 |
| SYPL1 | 707.8416372 | -0.13084548 | 0.046301482 | -2.825945862 | 0.004714123 | 0.04328994 |
| PTGIR | 7.079697021 | 1.000425 | 0.354056346 | 2.825609572 | 0.004719074 | 0.043314764 |
| GPR34 | 57.02716858 | -0.596931539 | 0.211296577 | -2.825088543 | 0.004726755 | 0.043364603 |
| DOCK5 | 363.1501367 | 0.279588345 | 0.099012602 | 2.823765242 | 0.004746314 | 0.0435049 |
| SACM1L | 1101.566364 | 0.147645219 | 0.052286961 | 2.823748336 | 0.004746564 | 0.0435049 |
| PDCD6IP | 3044.745128 | -0.246329934 | 0.087242916 | -2.823494978 | 0.004750317 | 0.043518596 |
| ZNF513 | 83.83449874 | 0.325854177 | 0.115438694 | 2.822746576 | 0.004761419 | 0.043599573 |
| FERMT1 | 1.502823436 | 2.219751561 | 0.786590095 | 2.821992771 | 0.004772625 | 0.043681425 |
| SLC22A23 | 4.584284651 | 0.961793145 | 0.34086951 | 2.821587493 | 0.00477866 | 0.04371589 |
| SLC25A29 | 134.1675909 | -0.381680695 | 0.135296866 | -2.821060863 | 0.004786512 | 0.04376694 |
| S100A3 | 4.48457931 | 1.291828596 | 0.457952655 | 2.820878059 | 0.00478924 | 0.043771113 |
| AMMECR1 | 183.7684237 | 0.244049857 | 0.086523228 | 2.820628212 | 0.004792972 | 0.043784443 |
| HMGCR | 311.4087169 | 0.184765634 | 0.065518256 | 2.820063371 | 0.004801417 | 0.043831735 |

| | | | | | | |
|---|---|---|---|---|---|---|
| RAB11FIP3 | 100.2323873 | -0.265454826 | 0.094133661 | -2.819977702 | 0.004802699 | 0.043831735 |
| PDZD7 | 26.99736513 | -0.630026229 | 0.223455084 | -2.81947592 | 0.004810214 | 0.043877347 |
| WHAMM | 341.4538952 | 0.244187912 | 0.086611729 | 2.819340015 | 0.004812251 | 0.043877347 |
| WLS | 388.6549857 | -0.248981864 | 0.088369189 | -2.817518932 | 0.004839626 | 0.044106072 |
| RRM2B | 122.7930714 | -0.233935729 | 0.08306727 | -2.816220272 | 0.004859234 | 0.044263829 |
| ZER1 | 250.02074 | -0.217994755 | 0.077418607 | -2.8157928 | 0.004865704 | 0.044301818 |
| LGALS8 | 888.1381081 | 0.184770348 | 0.065640973 | 2.814863016 | 0.004879803 | 0.044409203 |
| ZDHHC5 | 903.0267723 | 0.226220808 | 0.080373338 | 2.814625026 | 0.004883418 | 0.044421118 |
| ADH5 | 1563.98276 | -0.194535966 | 0.069127786 | -2.814150119 | 0.004890638 | 0.044465804 |
| AKAP11 | 312.8273846 | -0.174652545 | 0.062066059 | -2.813978326 | 0.004893253 | 0.044468588 |
| ANLN | 11.83502886 | -0.757108101 | 0.269105845 | -2.813421242 | 0.004901739 | 0.044488082 |
| GPR157 | 300.9206328 | -0.219844645 | 0.078142457 | -2.813382802 | 0.004902325 | 0.044488082 |
| XRN1 | 168.2596343 | 0.484948279 | 0.172368947 | 2.813431812 | 0.004901578 | 0.044488082 |
| CBX8 | 27.79868036 | -0.403222176 | 0.143334473 | -2.8131556 | 0.004905791 | 0.04449857 |
| COMMD7 | 20.11411896 | -0.550004329 | 0.195523093 | -2.812989099 | 0.004908332 | 0.044500666 |
| ZNF326 | 94.61619168 | 0.276179468 | 0.098243638 | 2.811168976 | 0.004936185 | 0.044732149 |
| TLE6 | 6.220834938 | -1.640364259 | 0.583558636 | -2.810967325 | 0.00493928 | 0.04473915 |
| PDCD6 | 919.1316586 | -0.154636841 | 0.055015181 | -2.810803081 | 0.004941802 | 0.044740958 |
| GLRX5 | 586.583274 | -0.161631154 | 0.057507097 | -2.810629704 | 0.004944466 | 0.044744046 |
| NMI | 279.7039791 | 0.305585837 | 0.108771954 | 2.809417549 | 0.004963123 | 0.044870731 |
| PVRL4 | 8.003138622 | -0.897379186 | 0.319414125 | -2.809453669 | 0.004962566 | 0.044870731 |
| LRP3 | 14.98467468 | -0.515512448 | 0.183517863 | -2.80905869 | 0.004968659 | 0.044899709 |
| SLC25A32 | 650.0826929 | 0.331452808 | 0.1180057 | 2.808786413 | 0.004972863 | 0.04491663 |
| TNPO2 | 116.3959605 | 0.254355548 | 0.090574387 | 2.808250271 | 0.00498115 | 0.0449704 |
| FGD2 | 232.0141237 | 0.321560773 | 0.114542736 | 2.807343227 | 0.004995199 | 0.045076112 |
| USP25 | 848.8947223 | 0.190892013 | 0.068056152 | 2.804919295 | 0.005032918 | 0.045395224 |
| MAPKAPK5 | 61.56236204 | -0.306108718 | 0.109221961 | -2.802629759 | 0.005068782 | 0.045697313 |
| ELL2 | 2899.50955 | 0.243281648 | 0.086851093 | 2.801135129 | 0.005092319 | 0.045888035 |
| AIG1 | 98.99376576 | -0.376793969 | 0.134551261 | -2.800374863 | 0.005104329 | 0.045931812 |
| ANKRD1 | 4.396790313 | 1.598341219 | 0.570748278 | 2.800431085 | 0.00510344 | 0.045931812 |
| SEC61B | 2257.283482 | 0.259419187 | 0.092631532 | 2.800549468 | 0.005101569 | 0.045931812 |
| APBA3 | 354.9945636 | 0.204263381 | 0.072959113 | 2.799696599 | 0.005115066 | 0.046006937 |
| AAMDC | 82.381955 | -0.305992434 | 0.109316174 | -2.799150612 | 0.005123723 | 0.0460633 |
| HLA-DQA1 | 7063.402962 | 0.368894831 | 0.131817466 | 2.798527698 | 0.005133617 | 0.046130717 |
| PTPRC | 546.6343534 | 0.22561596 | 0.080628429 | 2.798218481 | 0.005138534 | 0.046135419 |
| SIRPD | 19.64793153 | -0.626074448 | 0.223742358 | -2.798193656 | 0.005138929 | 0.046135419 |
| AACS | 99.77379703 | 0.260993714 | 0.093291677 | 2.797609854 | 0.005148226 | 0.046197353 |
| FBXW7 | 237.8891233 | 0.176187092 | 0.063007659 | 2.796280537 | 0.005169451 | 0.046366218 |
| C11orf96 | 26.43162105 | 0.891064547 | 0.318771557 | 2.79530757 | 0.005185036 | 0.046484366 |
| COA1 | 298.216718 | -0.225778344 | 0.080779424 | -2.794998189 | 0.005190001 | 0.046507233 |
| FAM104A | 356.7846086 | 0.156634745 | 0.056044939 | 2.794806239 | 0.005193083 | 0.04651322 |
| TTC19 | 181.3519808 | -0.174096283 | 0.062311041 | -2.793987724 | 0.005206246 | 0.046609445 |
| C7orf43 | 784.2630911 | 0.281848882 | 0.100887358 | 2.793698706 | 0.0052109 | 0.04662945 |
| DDX10 | 549.9213854 | 0.165563161 | 0.059282395 | 2.792788002 | 0.005225593 | 0.046739215 |
| AVPI1 | 923.3217256 | -0.206837111 | 0.074069933 | -2.792457133 | 0.00523094 | 0.04676533 |
| IL1R1 | 38.59793554 | 0.539536759 | 0.193294929 | 2.791261837 | 0.005250298 | 0.046916624 |
| MYEOV2 | 1733.661433 | -0.207833115 | 0.074463181 | -2.791085657 | 0.005253157 | 0.046920408 |

| | | | | | | |
|---|---|---|---|---|---|---|
| EIF4E | 1461.090631 | 0.231101515 | 0.082853447 | 2.789280644 | 0.005282527 | 0.047160871 |
| EHMT2 | 201.1780008 | -0.254829025 | 0.091383598 | -2.788564142 | 0.005294226 | 0.047243429 |
| RPL27 | 5839.699349 | -0.192726267 | 0.069138236 | -2.787549679 | 0.005310831 | 0.047369663 |
| EMR2 | 1066.012872 | 0.233604292 | 0.0838464 | 2.786098047 | 0.005334673 | 0.047560304 |
| NMB | 347.090275 | -0.379422173 | 0.136254547 | -2.784656966 | 0.005358438 | 0.047750076 |
| PTBP1 | 2647.563281 | 0.293269958 | 0.105450938 | 2.78110335 | 0.005417449 | 0.048253614 |
| ACYP2 | 223.8866005 | 0.344908837 | 0.124053657 | 2.780319782 | 0.005430539 | 0.048320693 |
| GSTP1 | 4929.658299 | -0.226833151 | 0.081588721 | -2.780202329 | 0.005432504 | 0.048320693 |
| PRKCB | 131.2645428 | 0.399633426 | 0.143739613 | 2.780259515 | 0.005431547 | 0.048320693 |
| XIAP | 510.3295503 | 0.177471835 | 0.063837818 | 2.780042321 | 0.005435181 | 0.048322199 |
| NCF1C | 683.019909 | -0.267670858 | 0.096363256 | -2.777727421 | 0.005474052 | 0.048645333 |
| TMEM245 | 618.7473063 | -0.208958246 | 0.075286055 | -2.77552393 | 0.005511284 | 0.048953621 |
| SRRT | 334.2692665 | 0.273241026 | 0.098452091 | 2.775370474 | 0.005513886 | 0.048954158 |
| CACFD1 | 27.62637796 | -0.40103893 | 0.144552905 | -2.774340164 | 0.005531381 | 0.049086864 |
| IFT122 | 169.226172 | -0.210029703 | 0.075716753 | -2.773886804 | 0.005539095 | 0.049132689 |
| SNURF | 99.23488897 | -0.365681929 | 0.131980585 | -2.77072518 | 0.005593161 | 0.049589435 |
| RAD9A | 89.09334453 | 0.2435239 | 0.087968031 | 2.768322731 | 0.005634563 | 0.049933527 |
| CGNL1 | 1.89049144 | 2.378515951 | 0.859270678 | 2.768063676 | 0.005639044 | 0.04995026 |
| SLC25A33 | 174.5825788 | 0.217825861 | 0.078698607 | 2.767849024 | 0.005642759 | 0.049958105 |
| VBP1 | 762.3100344 | 0.188968659 | 0.068276105 | 2.767712934 | 0.005645116 | 0.049958105 |

**Supplementary File 4. Differential expression of RNAs control versus M.tb infected AMs in all 28 donor AMs at 2h, 24h and 72h (Suppl. Tables 4A, B, and C), using separate non-infected control AMs for each time point**

**Suppl. File 4C: 72h**

| Gene | baseMean | log2FoldChange | lfcSE | stat | pvalue | padj |
|---|---|---|---|---|---|---|
| RSAD2 | 2766.166392 | 4.912946254 | 0.278596558 | 17.6346265 | 1.34E-69 | 2.49E-65 |
| INHBA | 1589.223749 | 4.704128144 | 0.298514944 | 15.75843434 | 6.01E-56 | 3.73E-52 |
| TNFAIP6 | 6758.379874 | 5.024082354 | 0.31833882 | 15.78218562 | 4.13E-56 | 3.73E-52 |
| CCL20 | 741.8013063 | 5.442185696 | 0.350716164 | 15.51735063 | 2.65E-54 | 9.87E-51 |
| GCH1 | 222.5524284 | 3.149521133 | 0.202884759 | 15.52369509 | 2.40E-54 | 9.87E-51 |
| CCL4 | 2974.633871 | 3.50536249 | 0.226630075 | 15.46733149 | 5.76E-54 | 1.79E-50 |
| AMICA1 | 301.6239116 | -2.201229689 | 0.154924876 | -14.20836824 | 8.13E-46 | 2.16E-42 |
| DUSP5 | 476.0351968 | 1.894539217 | 0.133440894 | 14.19759083 | 9.48E-46 | 2.21E-42 |
| ISG20 | 122.6101873 | 4.931518474 | 0.348843835 | 14.13675113 | 2.25E-45 | 4.67E-42 |
| HSD11B1 | 418.4518623 | 3.264902195 | 0.232085399 | 14.06767596 | 6.00E-45 | 1.12E-41 |
| SLC46A1 | 214.4879827 | -2.208056161 | 0.15705516 | -14.05911251 | 6.77E-45 | 1.15E-41 |
| PARP14 | 2221.977369 | 1.09076621 | 0.078136802 | 13.95969871 | 2.75E-44 | 4.26E-41 |
| GBP1 | 3987.926154 | 2.269253112 | 0.164831025 | 13.76714794 | 4.02E-43 | 5.76E-40 |
| ISG15 | 2322.407963 | 3.042976714 | 0.222291986 | 13.6890977 | 1.18E-42 | 1.57E-39 |
| GBP4 | 829.6809044 | 1.593093439 | 0.118418958 | 13.45302698 | 2.96E-41 | 3.67E-38 |
| DDX60 | 541.9428301 | 1.420289236 | 0.105611976 | 13.4481835 | 3.16E-41 | 3.67E-38 |
| MX1 | 5014.427796 | 2.317716106 | 0.178746178 | 12.96652123 | 1.89E-38 | 2.08E-35 |
| MT2A | 9296.041803 | 2.57891486 | 0.198986085 | 12.96027743 | 2.06E-38 | 2.13E-35 |
| IFI6 | 17405.81014 | 2.13935216 | 0.165889432 | 12.89625345 | 4.73E-38 | 4.63E-35 |
| DENND5A | 1840.833712 | 0.90270097 | 0.070175393 | 12.86349712 | 7.22E-38 | 6.73E-35 |
| LAP3 | 6161.425629 | 1.386212939 | 0.108939729 | 12.72458584 | 4.32E-37 | 3.83E-34 |
| DDX58 | 1316.752241 | 1.676124395 | 0.132996934 | 12.60272956 | 2.04E-36 | 1.66E-33 |
| EIF2AK2 | 1660.364012 | 1.423201061 | 0.112929968 | 12.60251008 | 2.05E-36 | 1.66E-33 |
| CCL15 | 447.0541085 | 3.561820127 | 0.283849539 | 12.54826813 | 4.06E-36 | 3.15E-33 |
| IRF7 | 1201.154915 | 2.144264023 | 0.171147333 | 12.52876095 | 5.20E-36 | 3.87E-33 |
| CXCL10 | 870.2579647 | 5.585627367 | 0.447468287 | 12.48273349 | 9.27E-36 | 6.65E-33 |
| DHX58 | 275.3946294 | 1.574567754 | 0.129377125 | 12.17037208 | 4.47E-34 | 3.09E-31 |
| PLSCR1 | 2242.680524 | 1.211268365 | 0.100193533 | 12.08928689 | 1.20E-33 | 8.01E-31 |
| TAP1 | 2681.04615 | 1.7444439 | 0.144647287 | 12.0599835 | 1.72E-33 | 1.10E-30 |
| CFB | 1057.334343 | 2.129105382 | 0.178067886 | 11.9567061 | 5.99E-33 | 3.72E-30 |
| PCED1B-AS | 250.6489094 | -1.722730348 | 0.144205036 | -11.94639519 | 6.78E-33 | 4.08E-30 |
| IL7R | 1092.261596 | 2.337986614 | 0.195863055 | 11.9368434 | 7.61E-33 | 4.43E-30 |
| USP18 | 300.4633859 | 2.522472499 | 0.212719279 | 11.85822231 | 1.95E-32 | 1.10E-29 |
| OAS3 | 1927.81696 | 2.192065824 | 0.185080395 | 11.84385751 | 2.32E-32 | 1.27E-29 |
| APMAP | 1007.919348 | -0.76067966 | 0.064340367 | -11.82274358 | 2.98E-32 | 1.59E-29 |
| IFITM3 | 1329.295818 | 2.404214647 | 0.204214841 | 11.77296733 | 5.38E-32 | 2.78E-29 |
| PARP9 | 439.2009568 | 1.622455721 | 0.137843289 | 11.77029168 | 5.55E-32 | 2.80E-29 |
| IFITM1 | 1202.969255 | 2.409818839 | 0.205110503 | 11.74888074 | 7.16E-32 | 3.51E-29 |
| CCL3 | 2341.89353 | 2.386573624 | 0.204115495 | 11.69227073 | 1.40E-31 | 6.67E-29 |
| ATF3 | 1732.626154 | 1.183190005 | 0.101266107 | 11.68396849 | 1.54E-31 | 7.17E-29 |
| TAP2 | 1042.801022 | 1.282061881 | 0.110455508 | 11.60704348 | 3.80E-31 | 1.72E-28 |
| IFIT1 | 737.6343114 | 2.990994854 | 0.258123489 | 11.58745712 | 4.77E-31 | 2.12E-28 |

| | | | | | | |
|---|---|---|---|---|---|---|
| HERC6 | 197.838323 | 1.938677731 | 0.16956914 | 11.43296316 | 2.86E-30 | 1.24E-27 |
| CYP27B1 | 238.3203023 | 2.945249934 | 0.258100488 | 11.41125286 | 3.67E-30 | 1.56E-27 |
| CCL5 | 1058.348222 | 2.931077686 | 0.257452285 | 11.38493559 | 4.97E-30 | 2.06E-27 |
| CD274 | 1261.138089 | 1.874900762 | 0.165263726 | 11.34490189 | 7.86E-30 | 3.18E-27 |
| PVALB | 324.2809752 | -1.744078735 | 0.153965048 | -11.32775755 | 9.56E-30 | 3.79E-27 |
| CXCL11 | 58.5470209 | 6.26355157 | 0.558115236 | 11.22268515 | 3.16E-29 | 1.22E-26 |
| APOC2 | 2469.901298 | -0.947206379 | 0.084511323 | -11.20804103 | 3.72E-29 | 1.39E-26 |
| HCAR3 | 118.1955462 | 1.952025123 | 0.174160356 | 11.20820586 | 3.72E-29 | 1.39E-26 |
| SLC27A1 | 409.9289624 | -0.959618396 | 0.085879494 | -11.17401084 | 5.47E-29 | 2.00E-26 |
| NUB1 | 1168.309329 | 0.989962293 | 0.088709551 | 11.15959085 | 6.43E-29 | 2.30E-26 |
| TBXAS1 | 1061.145358 | -0.729772197 | 0.065415415 | -11.15596686 | 6.70E-29 | 2.35E-26 |
| MMP2 | 289.6741024 | -1.291387165 | 0.115915986 | -11.14071675 | 7.95E-29 | 2.74E-26 |
| TNF | 404.1194724 | 1.769861229 | 0.159050901 | 11.12764036 | 9.20E-29 | 3.12E-26 |
| IL8 | 10902.81257 | 3.545304826 | 0.318915345 | 11.11675834 | 1.04E-28 | 3.46E-26 |
| SP110 | 124.3002386 | 1.145559185 | 0.10310926 | 11.11014844 | 1.12E-28 | 3.66E-26 |
| IDO1 | 1430.348017 | 5.278127417 | 0.476004459 | 11.0883991 | 1.43E-28 | 4.51E-26 |
| SLA | 1837.70283 | -1.164793013 | 0.105042199 | -11.08881022 | 1.42E-28 | 4.51E-26 |
| IFIT3 | 339.5508923 | 2.182106406 | 0.197046327 | 11.07407806 | 1.68E-28 | 5.20E-26 |
| SORL1 | 449.2567445 | -1.394893211 | 0.126205392 | -11.05256435 | 2.13E-28 | 6.51E-26 |
| HERC5 | 772.4780296 | 1.802035743 | 0.164637438 | 10.94547971 | 6.98E-28 | 2.10E-25 |
| SERPINE1 | 1289.02002 | 2.147360197 | 0.196457918 | 10.93038256 | 8.25E-28 | 2.44E-25 |
| CFP | 1022.458724 | -2.019951444 | 0.185098675 | -10.91283577 | 1.00E-27 | 2.87E-25 |
| EDN1 | 376.4675399 | 3.722651047 | 0.341106019 | 10.91347218 | 9.94E-28 | 2.87E-25 |
| CHMP5 | 2554.279105 | 0.832240721 | 0.076350989 | 10.90019564 | 1.15E-27 | 3.25E-25 |
| FAM129A | 1785.228785 | 0.945454443 | 0.086786811 | 10.89398755 | 1.23E-27 | 3.42E-25 |
| KIAA1462 | 115.3218534 | 3.306321166 | 0.304133075 | 10.87129759 | 1.58E-27 | 4.33E-25 |
| MT1E | 652.8391469 | 2.592876053 | 0.238733219 | 10.86097722 | 1.77E-27 | 4.78E-25 |
| IL15RA | 550.736194 | 1.599772569 | 0.148674208 | 10.76025621 | 5.30E-27 | 1.41E-24 |
| GPX1 | 17708.2123 | -0.472746566 | 0.043970907 | -10.75134894 | 5.84E-27 | 1.53E-24 |
| CA12 | 115.4887448 | 5.304719422 | 0.493594298 | 10.74712459 | 6.11E-27 | 1.58E-24 |
| CD180 | 107.4836498 | -2.797711456 | 0.260564778 | -10.73710531 | 6.81E-27 | 1.74E-24 |
| APOC1 | 11512.39423 | -1.165106105 | 0.10853132 | -10.73520621 | 6.96E-27 | 1.75E-24 |
| CD101 | 446.4578966 | -1.673527837 | 0.156220875 | -10.712575 | 8.89E-27 | 2.21E-24 |
| FCN1 | 113.0727082 | -2.239521806 | 0.209393539 | -10.69527656 | 1.07E-26 | 2.63E-24 |
| SOD2 | 40415.61856 | 1.494327972 | 0.14017155 | 10.66070808 | 1.55E-26 | 3.76E-24 |
| SLAMF7 | 3641.902894 | 1.678641859 | 0.158208718 | 10.61029937 | 2.67E-26 | 6.38E-24 |
| CP | 342.6400297 | 4.299839563 | 0.406090131 | 10.58838725 | 3.37E-26 | 7.96E-24 |
| CMPK2 | 899.2343634 | 2.67594995 | 0.25338719 | 10.5607152 | 4.53E-26 | 1.06E-23 |
| HCAR2 | 117.7607998 | 2.130306631 | 0.20185798 | 10.55349229 | 4.89E-26 | 1.13E-23 |
| TNS1 | 868.6747094 | -1.284127242 | 0.122078541 | -10.51886132 | 7.07E-26 | 1.61E-23 |
| MTF1 | 530.6519809 | 1.000704251 | 0.095148471 | 10.51729197 | 7.19E-26 | 1.61E-23 |
| PSME2 | 1534.389497 | 1.100478047 | 0.104768211 | 10.50393091 | 8.29E-26 | 1.84E-23 |
| MT1H | 1637.737026 | 3.988660452 | 0.379885967 | 10.49962566 | 8.67E-26 | 1.90E-23 |
| IFIT2 | 1584.107584 | 2.066882772 | 0.197514964 | 10.46443636 | 1.26E-25 | 2.73E-23 |
| C15orf48 | 3458.535489 | 1.893377838 | 0.181084795 | 10.45575271 | 1.38E-25 | 2.95E-23 |
| HAMP | 185.648606 | -1.964881907 | 0.188198962 | -10.44045028 | 1.62E-25 | 3.43E-23 |
| FNIP2 | 2034.43235 | 0.880503018 | 0.08441354 | 10.4308268 | 1.79E-25 | 3.75E-23 |

| | | | | | | |
|---|---|---|---|---|---|---|
| IFI44L | 468.4473681 | 3.918083374 | 0.376015792 | 10.41999686 | 2.01E-25 | 4.16E-23 |
| ABTB2 | 58.51100617 | 2.252638708 | 0.217109699 | 10.37557841 | 3.20E-25 | 6.56E-23 |
| OASL | 420.2742717 | 1.809657206 | 0.174474882 | 10.37202139 | 3.32E-25 | 6.72E-23 |
| SAMD9L | 508.4629199 | 1.489439716 | 0.143613818 | 10.37114487 | 3.35E-25 | 6.72E-23 |
| RNF213 | 2198.399036 | 0.827722097 | 0.080126283 | 10.33021953 | 5.14E-25 | 1.02E-22 |
| S100A16 | 216.3885508 | 1.861681979 | 0.180274568 | 10.32692518 | 5.32E-25 | 1.04E-22 |
| GBP5 | 1350.007194 | 2.354465415 | 0.228037795 | 10.32489116 | 5.44E-25 | 1.06E-22 |
| TMEM132/ | 131.106302 | 2.495729317 | 0.243042538 | 10.26869344 | 9.75E-25 | 1.87E-22 |
| ZNFX1 | 1083.590849 | 1.080947699 | 0.105563951 | 10.23974268 | 1.32E-24 | 2.50E-22 |
| PML | 355.8278722 | 1.454560492 | 0.142400979 | 10.21453995 | 1.71E-24 | 3.21E-22 |
| SLC39A8 | 1342.242482 | 1.992889426 | 0.195372779 | 10.20044573 | 1.97E-24 | 3.68E-22 |
| PLEKHM3 | 578.5265059 | 0.849872802 | 0.083458752 | 10.18314775 | 2.36E-24 | 4.35E-22 |
| GPR34 | 230.4374221 | -2.164033835 | 0.213364881 | -10.14240873 | 3.58E-24 | 6.54E-22 |
| DTX3L | 1300.4399 | 0.810374678 | 0.079921332 | 10.13965428 | 3.68E-24 | 6.60E-22 |
| RASGEF1B | 1386.449703 | 0.727803728 | 0.071771517 | 10.1405649 | 3.65E-24 | 6.60E-22 |
| TREM2 | 2341.249012 | -1.323681721 | 0.130593333 | -10.13590578 | 3.83E-24 | 6.79E-22 |
| ZNF697 | 158.1350193 | 1.028435855 | 0.10218939 | 10.06401794 | 7.97E-24 | 1.40E-21 |
| GP1BA | 38.99770464 | 3.58931006 | 0.358334513 | 10.01664626 | 1.29E-23 | 2.24E-21 |
| IFIT5 | 294.8087678 | 1.304902081 | 0.13028783 | 10.01553316 | 1.30E-23 | 2.25E-21 |
| MT1G | 301.4058655 | 3.877751981 | 0.38723744 | 10.01388703 | 1.32E-23 | 2.26E-21 |
| TSC22D1 | 1741.409104 | 1.387912981 | 0.138649335 | 10.01023904 | 1.37E-23 | 2.33E-21 |
| MPZL2 | 245.0265714 | -1.19550628 | 0.11952295 | -10.00231567 | 1.49E-23 | 2.50E-21 |
| C1S | 1082.185596 | 1.675232461 | 0.167508741 | 10.00086596 | 1.51E-23 | 2.51E-21 |
| IL1RN | 11877.21319 | 1.18378822 | 0.118420606 | 9.996471584 | 1.58E-23 | 2.60E-21 |
| AIF1 | 1914.490832 | -1.204311448 | 0.120662639 | -9.980814741 | 1.85E-23 | 3.02E-21 |
| NFKB2 | 1444.096116 | 1.176460617 | 0.11789751 | 9.978672255 | 1.89E-23 | 3.06E-21 |
| AMPD3 | 655.7629627 | 1.323543746 | 0.132772388 | 9.968516571 | 2.09E-23 | 3.36E-21 |
| SGPP2 | 30.91527904 | 4.823623143 | 0.484271113 | 9.960584096 | 2.27E-23 | 3.61E-21 |
| HADH | 391.1493115 | -1.033292891 | 0.103886434 | -9.946369784 | 2.62E-23 | 4.13E-21 |
| CIDEB | 63.98620249 | -2.09931081 | 0.211475956 | -9.926947965 | 3.18E-23 | 4.98E-21 |
| HAPLN3 | 79.41565647 | 3.193595517 | 0.322691397 | 9.896748234 | 4.30E-23 | 6.68E-21 |
| NEURL3 | 44.14800248 | 4.013884864 | 0.405728018 | 9.893043327 | 4.46E-23 | 6.87E-21 |
| NAIP | 250.4221679 | -1.261435718 | 0.128450644 | -9.820392359 | 9.20E-23 | 1.40E-20 |
| EMP1 | 1055.798164 | 1.02807715 | 0.104722065 | 9.817197044 | 9.49E-23 | 1.43E-20 |
| ETV7 | 16.18436872 | 3.420683831 | 0.348450982 | 9.816829367 | 9.53E-23 | 1.43E-20 |
| CTTN | 481.6368911 | 1.799225073 | 0.183469941 | 9.806647663 | 1.05E-22 | 1.57E-20 |
| HELZ2 | 164.1388679 | 2.758407713 | 0.281421391 | 9.801698804 | 1.11E-22 | 1.64E-20 |
| PPBP | 55.89162593 | 3.722882874 | 0.380539047 | 9.783182307 | 1.33E-22 | 1.95E-20 |
| PDLIM4 | 231.2708331 | 1.800260944 | 0.184659852 | 9.749065223 | 1.86E-22 | 2.71E-20 |
| AK4 | 251.3012323 | 1.480147192 | 0.151853806 | 9.747185324 | 1.90E-22 | 2.74E-20 |
| LDLR | 248.0545692 | 2.607715683 | 0.267593107 | 9.745077933 | 1.94E-22 | 2.78E-20 |
| PHLDA2 | 85.23705958 | 1.201206519 | 0.123333438 | 9.739504057 | 2.05E-22 | 2.91E-20 |
| TP53BP2 | 975.2876902 | 0.734546628 | 0.075487109 | 9.730755846 | 2.23E-22 | 3.15E-20 |
| G0S2 | 1765.142826 | 3.629002678 | 0.37308578 | 9.72699275 | 2.31E-22 | 3.24E-20 |
| SEMA4D | 838.2322886 | 1.3037253 | 0.134057881 | 9.72509258 | 2.36E-22 | 3.28E-20 |
| STAT1 | 7603.745133 | 1.682764921 | 0.173099554 | 9.72137062 | 2.44E-22 | 3.37E-20 |
| IFI27 | 3112.567635 | 1.58217176 | 0.162956932 | 9.709140579 | 2.76E-22 | 3.78E-20 |

| | | | | | | |
|---|---|---|---|---|---|---|
| IL3RA | 102.4733616 | 3.659943266 | 0.377348511 | 9.699106146 | 3.04E-22 | 4.14E-20 |
| BIRC3 | 394.4694124 | 1.056631001 | 0.109024638 | 9.691671681 | 3.27E-22 | 4.42E-20 |
| TSPAN3 | 2615.421522 | 0.959783028 | 0.099083742 | 9.686584397 | 3.44E-22 | 4.61E-20 |
| PTGS2 | 254.0264258 | 2.871438618 | 0.296505557 | 9.684265774 | 3.52E-22 | 4.68E-20 |
| TGFBI | 4883.406575 | -1.407917541 | 0.145599412 | -9.669802385 | 4.05E-22 | 5.34E-20 |
| ZC3H12A | 355.9599856 | 1.603544358 | 0.165838449 | 9.669315915 | 4.07E-22 | 5.34E-20 |
| STAB1 | 20.28398965 | -3.412805519 | 0.35312648 | -9.664541491 | 4.27E-22 | 5.56E-20 |
| TMEM86A | 584.9591892 | -0.96509504 | 0.099928403 | -9.657865118 | 4.55E-22 | 5.89E-20 |
| MICALL1 | 638.6245434 | 0.71078306 | 0.073686079 | 9.646096983 | 5.11E-22 | 6.56E-20 |
| STAT2 | 2097.829383 | 1.074877916 | 0.112002628 | 9.596899102 | 8.24E-22 | 1.05E-19 |
| KDM6B | 458.7156876 | 0.90738978 | 0.094570954 | 9.594804154 | 8.41E-22 | 1.07E-19 |
| MTHFD2 | 3052.802778 | 0.908179196 | 0.094877929 | 9.572080774 | 1.05E-21 | 1.32E-19 |
| KYNU | 300.5266468 | 1.329186962 | 0.138918644 | 9.568096317 | 1.09E-21 | 1.36E-19 |
| JAG1 | 174.8171111 | 1.49754911 | 0.156819634 | 9.549500093 | 1.30E-21 | 1.62E-19 |
| GRAMD1A | 1224.909816 | 1.699305205 | 0.178462202 | 9.52193342 | 1.70E-21 | 2.10E-19 |
| SLC7A11 | 1486.120197 | 1.629426791 | 0.171434874 | 9.504640196 | 2.01E-21 | 2.46E-19 |
| ELOVL7 | 173.9142272 | 2.133740279 | 0.224612561 | 9.499648045 | 2.11E-21 | 2.56E-19 |
| HIP1 | 479.5904614 | 1.286610544 | 0.135466946 | 9.497597637 | 2.15E-21 | 2.60E-19 |
| CTSF | 93.41506505 | -1.486355898 | 0.156582121 | -9.492500745 | 2.26E-21 | 2.71E-19 |
| TNFSF10 | 189.2661746 | 2.153299795 | 0.226874289 | 9.49115832 | 2.28E-21 | 2.73E-19 |
| AIG1 | 182.7069456 | -1.245925642 | 0.131367226 | -9.484295916 | 2.44E-21 | 2.90E-19 |
| CABLES1 | 558.105261 | -1.062295869 | 0.112020577 | -9.483042288 | 2.47E-21 | 2.91E-19 |
| IL6 | 110.8269696 | 5.53483502 | 0.584211736 | 9.474022309 | 2.69E-21 | 3.16E-19 |
| MMP12 | 255.0258503 | 1.876851757 | 0.198166514 | 9.471084265 | 2.77E-21 | 3.23E-19 |
| TNIP1 | 2949.643648 | 1.168473829 | 0.123508605 | 9.460667366 | 3.06E-21 | 3.54E-19 |
| XAF1 | 298.231981 | 1.88838232 | 0.199877045 | 9.447719805 | 3.46E-21 | 3.98E-19 |
| JUNB | 2184.750238 | 0.882946953 | 0.093536342 | 9.439613844 | 3.74E-21 | 4.28E-19 |
| OCSTAMP | 212.9800934 | 2.1524261 | 0.228180425 | 9.433000646 | 3.99E-21 | 4.53E-19 |
| CCL3L3 | 307.6899487 | 2.490609589 | 0.264289675 | 9.423786932 | 4.35E-21 | 4.91E-19 |
| GBP1P1 | 50.36034405 | 4.033499804 | 0.42827027 | 9.418117696 | 4.59E-21 | 5.12E-19 |
| NLRC5 | 368.4602274 | 0.781556655 | 0.082979411 | 9.418681666 | 4.57E-21 | 5.12E-19 |
| APOL1 | 1015.315051 | 0.841917358 | 0.089491081 | 9.40783539 | 5.06E-21 | 5.62E-19 |
| ALDH7A1 | 101.0741073 | -0.644303229 | 0.068522964 | -9.402734401 | 5.32E-21 | 5.86E-19 |
| IRAK2 | 78.29556686 | 1.721960302 | 0.183302024 | 9.394115032 | 5.77E-21 | 6.32E-19 |
| CCL22 | 13518.87543 | 1.907875845 | 0.203319026 | 9.38365621 | 6.37E-21 | 6.90E-19 |
| VNN2 | 256.2252571 | -0.896532759 | 0.09554208 | -9.383642869 | 6.37E-21 | 6.90E-19 |
| CERK | 750.3636364 | -0.837412597 | 0.089465633 | -9.360159543 | 7.96E-21 | 8.56E-19 |
| VEGFB | 994.1393424 | -0.681964375 | 0.072861129 | -9.359783271 | 7.99E-21 | 8.56E-19 |
| CD40 | 1347.850585 | 1.134959396 | 0.121278814 | 9.358265942 | 8.11E-21 | 8.63E-19 |
| CAT | 2343.331438 | -0.682117833 | 0.073048254 | -9.337907393 | 9.83E-21 | 1.04E-18 |
| ANGPT4 | 37.72767145 | -3.000038296 | 0.321568304 | -9.329396757 | 1.06E-20 | 1.12E-18 |
| IFI35 | 1371.554659 | 1.629164128 | 0.174672789 | 9.326948626 | 1.09E-20 | 1.14E-18 |
| RTP4 | 245.7044903 | 1.251144999 | 0.134155345 | 9.326091319 | 1.10E-20 | 1.14E-18 |
| HESX1 | 44.01114912 | 2.628347681 | 0.282666959 | 9.298390208 | 1.43E-20 | 1.48E-18 |
| TNIP3 | 412.9034906 | 2.070062231 | 0.222682009 | 9.296046127 | 1.46E-20 | 1.50E-18 |
| CYB5R3 | 62.55642274 | 3.054456131 | 0.330383138 | 9.245193789 | 2.35E-20 | 2.40E-18 |
| GPR64 | 85.22371964 | 1.73204785 | 0.187551456 | 9.235054138 | 2.58E-20 | 2.63E-18 |

| | | | | | | |
|---|---|---|---|---|---|---|
| TMEM170I | 370.4415151 | -0.924537806 | 0.10032575 | -9.215359008 | 3.10E-20 | 3.14E-18 |
| TNIK | 781.4658026 | 0.805490235 | 0.087481989 | 9.207497943 | 3.34E-20 | 3.36E-18 |
| P2RX1 | 120.7106185 | -1.221481918 | 0.132689446 | -9.205569492 | 3.40E-20 | 3.40E-18 |
| APOO | 275.6431047 | 1.333847343 | 0.144983465 | 9.199996298 | 3.58E-20 | 3.57E-18 |
| PSMB9 | 1580.897336 | 1.215948177 | 0.132192997 | 9.198279792 | 3.64E-20 | 3.61E-18 |
| DCUN1D3 | 270.4202723 | 1.301304929 | 0.141573843 | 9.191704506 | 3.87E-20 | 3.81E-18 |
| MFNG | 619.1876293 | -0.614022993 | 0.067058984 | -9.156461389 | 5.36E-20 | 5.26E-18 |
| PFKFB2 | 236.098616 | -0.886916757 | 0.097101502 | -9.133913912 | 6.61E-20 | 6.41E-18 |
| STX11 | 1503.485603 | 0.61961006 | 0.067836535 | 9.13386953 | 6.61E-20 | 6.41E-18 |
| CD4 | 2239.159719 | -0.842544504 | 0.092306502 | -9.127683199 | 7.00E-20 | 6.76E-18 |
| MPEG1 | 562.356056 | -1.477920534 | 0.162104392 | -9.117091245 | 7.72E-20 | 7.41E-18 |
| TCF7L2 | 603.7321274 | 0.687765664 | 0.075493285 | 9.110289326 | 8.22E-20 | 7.85E-18 |
| MSC | 634.2937579 | 1.709901897 | 0.187848383 | 9.102563809 | 8.82E-20 | 8.39E-18 |
| AGRN | 449.2613138 | 1.109486389 | 0.12194183 | 9.098488906 | 9.16E-20 | 8.66E-18 |
| HS3ST2 | 94.64166158 | -2.122827938 | 0.233437963 | -9.09375628 | 9.57E-20 | 9.00E-18 |
| COLEC12 | 844.5403452 | -1.194193125 | 0.131695921 | -9.067806504 | 1.21E-19 | 1.14E-17 |
| PSTPIP2 | 413.3866766 | 1.893455836 | 0.209091753 | 9.055621796 | 1.36E-19 | 1.27E-17 |
| ADORA3 | 16.63671804 | -3.352797455 | 0.37052906 | -9.048676111 | 1.45E-19 | 1.34E-17 |
| PNPT1 | 589.2981175 | 1.117686744 | 0.12352305 | 9.048406271 | 1.45E-19 | 1.34E-17 |
| INSIG1 | 373.8138838 | 1.451076872 | 0.160433469 | 9.044726632 | 1.50E-19 | 1.38E-17 |
| GAL3ST4 | 20.767042 | -3.523592909 | 0.390296939 | -9.02797988 | 1.75E-19 | 1.60E-17 |
| NRIP3 | 474.5866322 | 0.96628248 | 0.107071383 | 9.024656742 | 1.80E-19 | 1.64E-17 |
| KCNJ5 | 304.0937203 | -1.690605899 | 0.18766796 | -9.008495097 | 2.09E-19 | 1.89E-17 |
| FAM124A | 61.02931046 | 2.544006298 | 0.282599743 | 9.002153607 | 2.21E-19 | 1.99E-17 |
| WDR91 | 317.098171 | -0.969455757 | 0.107782545 | -8.994552477 | 2.37E-19 | 2.12E-17 |
| NAMPT | 5264.134648 | 0.815890208 | 0.090810116 | 8.98457404 | 2.60E-19 | 2.32E-17 |
| CCL8 | 103.5277561 | 4.972640343 | 0.554434811 | 8.968845823 | 3.00E-19 | 2.65E-17 |
| KIAA0040 | 98.25097545 | 0.962288547 | 0.107288841 | 8.969139154 | 2.99E-19 | 2.65E-17 |
| LAMP3 | 85.56816281 | 1.609957745 | 0.179555569 | 8.966348158 | 3.07E-19 | 2.68E-17 |
| RASGRP1 | 131.313989 | 2.132753801 | 0.237855618 | 8.966589967 | 3.06E-19 | 2.68E-17 |
| FBXO6 | 266.3884974 | 1.33495742 | 0.148910428 | 8.964835025 | 3.11E-19 | 2.71E-17 |
| TFPI2 | 18.90549937 | 2.915596308 | 0.325395268 | 8.960168135 | 3.24E-19 | 2.81E-17 |
| PDGFRL | 10.56282793 | 5.337547437 | 0.595856953 | 8.957766468 | 3.31E-19 | 2.86E-17 |
| TSPAN15 | 315.3515831 | -0.878407429 | 0.098075059 | -8.956481257 | 3.35E-19 | 2.88E-17 |
| CD80 | 444.4172936 | 1.223953982 | 0.137014402 | 8.933031589 | 4.14E-19 | 3.54E-17 |
| CYP51A1 | 983.4369427 | 0.851592091 | 0.095351606 | 8.931072369 | 4.22E-19 | 3.59E-17 |
| LY6E | 2095.277651 | 1.363195395 | 0.152668608 | 8.929113943 | 4.29E-19 | 3.64E-17 |
| LAMB3 | 66.85049286 | 2.401717925 | 0.269508413 | 8.911476632 | 5.04E-19 | 4.25E-17 |
| KCNN4 | 1090.463272 | 1.359193612 | 0.152566857 | 8.908839313 | 5.16E-19 | 4.31E-17 |
| STARD4 | 131.9760651 | 1.030361743 | 0.115649983 | 8.909311666 | 5.14E-19 | 4.31E-17 |
| ECE1 | 191.1033255 | 1.475167059 | 0.165647801 | 8.905443048 | 5.32E-19 | 4.42E-17 |
| ACSF2 | 475.6836662 | -0.542980904 | 0.061119087 | -8.883982605 | 6.45E-19 | 5.34E-17 |
| OSM | 725.8336165 | 2.765222952 | 0.311512078 | 8.876776057 | 6.88E-19 | 5.67E-17 |
| EHD1 | 1019.311678 | 1.060850011 | 0.119531092 | 8.875096807 | 6.99E-19 | 5.74E-17 |
| FARP1 | 688.5509539 | -0.678542962 | 0.076481955 | -8.871935369 | 7.19E-19 | 5.85E-17 |
| METTL7A | 178.2417263 | -1.439698295 | 0.162271328 | -8.872166852 | 7.17E-19 | 5.85E-17 |
| SEPP1 | 40.49693107 | -2.315130901 | 0.261346178 | -8.858483861 | 8.11E-19 | 6.57E-17 |

| | | | | | | |
|---|---|---|---|---|---|---|
| MMP9 | 11505.70524 | 1.161600084 | 0.131152193 | 8.856886455 | 8.23E-19 | 6.64E-17 |
| KIF1B | 528.5765929 | 0.545181058 | 0.061586313 | 8.852308743 | 8.57E-19 | 6.89E-17 |
| GAPT | 14.50557126 | -3.303389782 | 0.373404666 | -8.846675161 | 9.02E-19 | 7.21E-17 |
| TNFRSF4 | 46.67747045 | 3.847062578 | 0.435469699 | 8.834283047 | 1.01E-18 | 8.02E-17 |
| MASTL | 181.091577 | 0.774193856 | 0.087669897 | 8.830783266 | 1.04E-18 | 8.24E-17 |
| PRADC1 | 131.4385882 | -0.823663208 | 0.093414225 | -8.817320984 | 1.17E-18 | 9.26E-17 |
| RAB42 | 748.2076811 | -0.986282145 | 0.111892615 | -8.814541938 | 1.20E-18 | 9.45E-17 |
| B3GNT5 | 344.8983026 | 0.860481913 | 0.097685719 | 8.808676638 | 1.27E-18 | 9.91E-17 |
| ZNF589 | 814.8007175 | -0.657567896 | 0.074869444 | -8.782860632 | 1.59E-18 | 1.24E-16 |
| LY86 | 1875.032878 | -1.158530865 | 0.132118913 | -8.768849496 | 1.81E-18 | 1.40E-16 |
| DOCK4 | 712.8202955 | 0.923757923 | 0.105493315 | 8.756554116 | 2.01E-18 | 1.56E-16 |
| CISH | 366.454313 | 1.100971352 | 0.126090883 | 8.731569838 | 2.51E-18 | 1.93E-16 |
| TJP1 | 145.3272481 | 1.010349325 | 0.115962901 | 8.71269448 | 2.97E-18 | 2.28E-16 |
| BAALC | 44.00290595 | 2.908335927 | 0.333940468 | 8.709144896 | 3.06E-18 | 2.34E-16 |
| AQP3 | 269.7831115 | 1.398723197 | 0.16073502 | 8.702043898 | 3.26E-18 | 2.47E-16 |
| CCL17 | 19.84119682 | 3.888905295 | 0.446888829 | 8.702176118 | 3.26E-18 | 2.47E-16 |
| APOL3 | 774.8365656 | 1.388425509 | 0.159723485 | 8.692682285 | 3.54E-18 | 2.67E-16 |
| SPHK1 | 658.4874829 | 1.399917452 | 0.161089513 | 8.69030782 | 3.61E-18 | 2.72E-16 |
| CSRNP1 | 96.89764371 | 1.173332632 | 0.135223254 | 8.677003371 | 4.06E-18 | 3.03E-16 |
| ZSWIM4 | 172.5561241 | 1.736211462 | 0.200087451 | 8.677263148 | 4.05E-18 | 3.03E-16 |
| IL32 | 1694.010733 | 1.721940377 | 0.198544139 | 8.672834102 | 4.21E-18 | 3.13E-16 |
| TRPC6 | 77.18083208 | -2.307773378 | 0.266937319 | -8.645375588 | 5.36E-18 | 3.97E-16 |
| TLR2 | 364.5610856 | 1.222526906 | 0.141545282 | 8.637002172 | 5.77E-18 | 4.25E-16 |
| APOBEC3G | 328.5165165 | 0.775610941 | 0.08981354 | 8.635790832 | 5.83E-18 | 4.28E-16 |
| JAK3 | 87.74929099 | 1.760989987 | 0.204035468 | 8.630803263 | 6.09E-18 | 4.45E-16 |
| LOC72843[illegible] | 110.6166401 | 1.477386648 | 0.171353686 | 8.621855102 | 6.59E-18 | 4.80E-16 |
| TPCN1 | 230.938819 | -1.223975449 | 0.141978989 | -8.620820986 | 6.65E-18 | 4.82E-16 |
| IGFBP6 | 187.7593348 | 1.203985452 | 0.139702664 | 8.618199644 | 6.80E-18 | 4.91E-16 |
| LGALS17A | 8.380183364 | 4.729362036 | 0.549121535 | 8.612596188 | 7.14E-18 | 5.14E-16 |
| APBA1 | 112.3390499 | -1.176644159 | 0.13663972 | -8.611289326 | 7.22E-18 | 5.16E-16 |
| PELI3 | 309.39599 | -0.725208455 | 0.084212212 | -8.611678016 | 7.20E-18 | 5.16E-16 |
| PNPLA3 | 499.6667876 | -1.300478136 | 0.151223148 | -8.599729303 | 7.99E-18 | 5.68E-16 |
| H3F3B | 6636.597366 | 0.520597154 | 0.060699797 | 8.576588096 | 9.77E-18 | 6.92E-16 |
| CMBL | 62.47288303 | -1.905009958 | 0.222161103 | -8.574903223 | 9.92E-18 | 7.00E-16 |
| PDGFC | 306.5644927 | -0.793809985 | 0.092626216 | -8.570035783 | 1.03E-17 | 7.25E-16 |
| SERPINB2 | 82.61501584 | 3.46239586 | 0.404017302 | 8.569919769 | 1.04E-17 | 7.25E-16 |
| RELB | 859.4049109 | 0.885592818 | 0.103394128 | 8.565213853 | 1.08E-17 | 7.53E-16 |
| GALM | 279.6623032 | -0.972445092 | 0.113600842 | -8.560192676 | 1.13E-17 | 7.83E-16 |
| IL2RA | 86.42973264 | 4.266527463 | 0.498466303 | 8.559309708 | 1.14E-17 | 7.87E-16 |
| APP | 589.3496111 | 0.792636617 | 0.09267272 | 8.553073804 | 1.20E-17 | 8.27E-16 |
| GPC4 | 2545.187034 | 1.015371416 | 0.118787127 | 8.547823648 | 1.25E-17 | 8.62E-16 |
| DOK2 | 1449.942469 | -0.947446658 | 0.111015741 | -8.534345211 | 1.41E-17 | 9.66E-16 |
| ABHD12 | 2135.517782 | -0.543945692 | 0.063741749 | -8.533585868 | 1.42E-17 | 9.68E-16 |
| TBC1D9 | 2016.278936 | 0.704597018 | 0.0825738 | 8.532936766 | 1.43E-17 | 9.70E-16 |
| RBP1 | 482.5702505 | -1.244710705 | 0.14635154 | -8.504937525 | 1.82E-17 | 1.23E-15 |
| GAS7 | 797.2809132 | -0.705944232 | 0.083036421 | -8.501621565 | 1.87E-17 | 1.26E-15 |
| RPL10A | 2936.084275 | -0.541272399 | 0.063725239 | -8.493846541 | 2.00E-17 | 1.34E-15 |

| | | | | | | |
|---|---|---|---|---|---|---|
| RNF144B | 274.8177174 | 0.931884983 | 0.109775097 | 8.489038147 | 2.08E-17 | 1.40E-15 |
| DOCK2 | 1963.550585 | -0.647377802 | 0.076328953 | -8.481418634 | 2.22E-17 | 1.49E-15 |
| MICALL2 | 53.82000001 | 1.136134437 | 0.133994728 | 8.478948805 | 2.27E-17 | 1.51E-15 |
| PTGES | 742.9270412 | 3.03736528 | 0.359151431 | 8.457060217 | 2.74E-17 | 1.82E-15 |
| PLA2G15 | 742.7427184 | -0.759680015 | 0.08990435 | -8.449869376 | 2.92E-17 | 1.93E-15 |
| DSP | 165.6858491 | 1.404670883 | 0.166337464 | 8.44470542 | 3.05E-17 | 2.01E-15 |
| SQRDL | 1462.896018 | 0.60163803 | 0.071260215 | 8.442832143 | 3.10E-17 | 2.03E-15 |
| SP100 | 645.8188792 | 0.532029611 | 0.063050345 | 8.438171234 | 3.22E-17 | 2.11E-15 |
| PPTC7 | 1029.672582 | 0.448746111 | 0.053205966 | 8.434131383 | 3.34E-17 | 2.17E-15 |
| ITGB3 | 348.4412915 | 1.423280382 | 0.16878269 | 8.432620538 | 3.38E-17 | 2.19E-15 |
| DIRAS2 | 23.53047132 | 3.371928371 | 0.400409828 | 8.421192828 | 3.73E-17 | 2.41E-15 |
| RIPK2 | 1046.247793 | 0.924869 | 0.109940231 | 8.412470978 | 4.01E-17 | 2.59E-15 |
| SLC2A1 | 193.3175255 | 1.004033829 | 0.119456973 | 8.40498304 | 4.28E-17 | 2.75E-15 |
| ARHGAP9 | 395.5879922 | -0.881876941 | 0.104942403 | -8.403437649 | 4.34E-17 | 2.78E-15 |
| TM4SF1 | 102.5393713 | 2.953769062 | 0.351890116 | 8.394009748 | 4.70E-17 | 3.00E-15 |
| CCR7 | 110.8671194 | 2.132408069 | 0.254062997 | 8.393225672 | 4.73E-17 | 3.01E-15 |
| GEM | 222.7619689 | 1.585238159 | 0.189602891 | 8.360833256 | 6.23E-17 | 3.95E-15 |
| MT1M | 138.2959652 | 4.234603456 | 0.506519478 | 8.360198649 | 6.26E-17 | 3.96E-15 |
| IRF1 | 1229.424729 | 1.08288319 | 0.129679308 | 8.350470168 | 6.80E-17 | 4.27E-15 |
| RHOF | 313.6453943 | 0.962017635 | 0.11520059 | 8.350804734 | 6.78E-17 | 4.27E-15 |
| RHOD | 51.89880749 | 2.231342069 | 0.268428676 | 8.312606903 | 9.36E-17 | 5.85E-15 |
| PDE4B | 611.618811 | 1.252764899 | 0.150840572 | 8.305225075 | 9.96E-17 | 6.21E-15 |
| CSF1 | 8399.358594 | 1.511641271 | 0.182201955 | 8.296515114 | 1.07E-16 | 6.66E-15 |
| ARHGEF5 | 462.2414906 | 1.038765121 | 0.12524074 | 8.294147095 | 1.09E-16 | 6.77E-15 |
| FUCA1 | 3002.094675 | -2.038273486 | 0.245974719 | -8.286516172 | 1.17E-16 | 7.20E-15 |
| MERTK | 55.55372553 | -1.834835888 | 0.221702273 | -8.276125722 | 1.27E-16 | 7.83E-15 |
| CSF2RB | 1606.772118 | 0.746239 | 0.090210084 | 8.272234851 | 1.31E-16 | 8.06E-15 |
| OTUD4 | 529.4487694 | 0.514036834 | 0.062236278 | 8.259440503 | 1.46E-16 | 8.94E-15 |
| SOCS3 | 71.30916938 | 1.448044037 | 0.175328097 | 8.259052956 | 1.47E-16 | 8.94E-15 |
| EEPD1 | 191.7539117 | -1.054881417 | 0.127785575 | -8.255089945 | 1.52E-16 | 9.21E-15 |
| LOC28307( | 211.6667833 | -0.739587748 | 0.089653036 | -8.249444516 | 1.59E-16 | 9.63E-15 |
| MAFB | 1043.106435 | -0.940612932 | 0.114044267 | -8.247787976 | 1.61E-16 | 9.73E-15 |
| CFLAR | 1670.983479 | 0.523654307 | 0.063706584 | 8.219783131 | 2.04E-16 | 1.23E-14 |
| OAS2 | 1390.549315 | 1.941517078 | 0.236531068 | 8.20829625 | 2.24E-16 | 1.34E-14 |
| SLC2A3 | 784.1900698 | 0.905653216 | 0.110439998 | 8.200409542 | 2.40E-16 | 1.43E-14 |
| GCHFR | 307.9448311 | -1.14576068 | 0.139752266 | -8.198512353 | 2.43E-16 | 1.45E-14 |
| TDRD7 | 205.287478 | 0.720743476 | 0.087995124 | 8.19072058 | 2.60E-16 | 1.54E-14 |
| COTL1 | 5048.880368 | -0.636609081 | 0.077750291 | -8.187867554 | 2.66E-16 | 1.57E-14 |
| LOC10050( | 178.6749475 | 1.548697056 | 0.189365772 | 8.178336761 | 2.88E-16 | 1.70E-14 |
| PIK3AP1 | 5572.891108 | 0.5646967 | 0.06919891 | 8.160485511 | 3.34E-16 | 1.96E-14 |
| RCSD1 | 290.1383206 | -1.329621089 | 0.162981273 | -8.158121877 | 3.40E-16 | 1.99E-14 |
| GMPR | 213.9546611 | 1.243708229 | 0.152495185 | 8.155721289 | 3.47E-16 | 2.03E-14 |
| VDR | 789.6069129 | 1.17563248 | 0.144197696 | 8.152921397 | 3.55E-16 | 2.07E-14 |
| TMEM171 | 17.61505602 | 4.155907564 | 0.509841803 | 8.15136683 | 3.60E-16 | 2.09E-14 |
| JAK2 | 295.0367211 | 0.669873985 | 0.082491619 | 8.120509617 | 4.64E-16 | 2.69E-14 |
| HS3ST3A1 | 22.17396156 | 3.524896679 | 0.434318769 | 8.115920679 | 4.82E-16 | 2.78E-14 |
| IL1B | 3090.432093 | 1.834800433 | 0.22616051 | 8.112824082 | 4.95E-16 | 2.84E-14 |

| | | | | | | |
|---|---|---|---|---|---|---|
| DPEP2 | 719.7524396 | -1.071092715 | 0.132033378 | -8.112287419 | 4.97E-16 | 2.85E-14 |
| MS4A6A | 212.9299531 | -2.294921327 | 0.282949601 | -8.110707064 | 5.03E-16 | 2.88E-14 |
| ABI3BP | 8.984551896 | 3.275061154 | 0.403903696 | 8.1085199 | 5.12E-16 | 2.92E-14 |
| CD44 | 312.2080057 | 0.973343536 | 0.120056871 | 8.107353855 | 5.17E-16 | 2.94E-14 |
| LRRFIP2 | 1016.38479 | 0.497137856 | 0.061327937 | 8.1062217 | 5.22E-16 | 2.96E-14 |
| MARCKS | 182.9814838 | 1.68779094 | 0.208249335 | 8.104664247 | 5.29E-16 | 2.99E-14 |
| FAM53B | 271.1031988 | -0.537494352 | 0.06634064 | -8.10203754 | 5.40E-16 | 3.04E-14 |
| GPD2 | 1548.29429 | 0.741889071 | 0.091694553 | 8.090873988 | 5.92E-16 | 3.32E-14 |
| PAK1 | 590.4686309 | -0.617481867 | 0.076328783 | -8.089764329 | 5.98E-16 | 3.35E-14 |
| GBP2 | 3679.054436 | 1.021419737 | 0.126405412 | 8.08050635 | 6.45E-16 | 3.60E-14 |
| ITGB8 | 2344.678909 | 1.825665552 | 0.225980776 | 8.078853362 | 6.54E-16 | 3.63E-14 |
| SLC30A1 | 1975.58986 | 0.565126466 | 0.069948184 | 8.079215743 | 6.52E-16 | 3.63E-14 |
| LOC72837 | 110.401125 | 1.254589106 | 0.155383861 | 8.074127489 | 6.80E-16 | 3.76E-14 |
| METTL1 | 367.8167891 | 0.810480306 | 0.100468756 | 8.066988564 | 7.21E-16 | 3.97E-14 |
| TRANK1 | 172.7142577 | 0.707252043 | 0.087728361 | 8.061840379 | 7.52E-16 | 4.13E-14 |
| GFPT2 | 10.50508586 | 2.242623631 | 0.278609702 | 8.0493379 | 8.32E-16 | 4.55E-14 |
| SNTA1 | 121.1102738 | -0.947637611 | 0.11772593 | -8.049523253 | 8.31E-16 | 4.55E-14 |
| CRIM1 | 2516.885461 | 0.882335413 | 0.109723568 | 8.04143934 | 8.88E-16 | 4.84E-14 |
| ETS2 | 870.4990688 | 0.89687491 | 0.111610232 | 8.035776761 | 9.30E-16 | 5.05E-14 |
| PGPEP1 | 681.4253326 | -0.494396177 | 0.061661306 | -8.017932252 | 1.08E-15 | 5.83E-14 |
| CH25H | 21.46641784 | 1.5549856 | 0.193947436 | 8.017562019 | 1.08E-15 | 5.83E-14 |
| SLC7A8 | 1301.696992 | 1.078841824 | 0.134726661 | 8.007634237 | 1.17E-15 | 6.30E-14 |
| FRY | 183.2815482 | -0.873968187 | 0.1091611 | -8.006223721 | 1.18E-15 | 6.35E-14 |
| SLC2A6 | 1473.187914 | 1.079225572 | 0.134807977 | 8.005650666 | 1.19E-15 | 6.36E-14 |
| KLF9 | 282.7754962 | 0.944933168 | 0.118219467 | 7.993042015 | 1.32E-15 | 7.03E-14 |
| HIST1H4E | 735.3410197 | -0.922806199 | 0.115467253 | -7.991929954 | 1.33E-15 | 7.07E-14 |
| MFAP5 | 96.15165219 | -1.342722318 | 0.168104087 | -7.987446012 | 1.38E-15 | 7.31E-14 |
| HES4 | 12.89207466 | 3.047156509 | 0.382406334 | 7.968373526 | 1.61E-15 | 8.51E-14 |
| HS3ST3B1 | 30.24131316 | 2.874346057 | 0.361130708 | 7.959295609 | 1.73E-15 | 9.13E-14 |
| RPL23 | 7334.917665 | -0.608558878 | 0.076470777 | -7.958058031 | 1.75E-15 | 9.20E-14 |
| SH3BP5 | 1675.131529 | 0.800231068 | 0.100676311 | 7.948553777 | 1.89E-15 | 9.90E-14 |
| GAL | 188.4300787 | 1.244185348 | 0.156640076 | 7.94295676 | 1.97E-15 | 1.03E-13 |
| RICTOR | 295.678447 | 0.541891996 | 0.068220077 | 7.943292057 | 1.97E-15 | 1.03E-13 |
| KDM2B | 234.7311019 | -0.620082521 | 0.078157505 | -7.933755302 | 2.13E-15 | 1.11E-13 |
| CYBRD1 | 110.0477409 | -0.893832943 | 0.112745147 | -7.92790611 | 2.23E-15 | 1.16E-13 |
| CREBL2 | 1835.15249 | -0.545637154 | 0.068887298 | -7.920722282 | 2.36E-15 | 1.22E-13 |
| LILRA5 | 170.4409979 | -1.466038399 | 0.185144478 | -7.918347954 | 2.41E-15 | 1.24E-13 |
| B4GALT1 | 2237.145451 | 1.041618518 | 0.131851334 | 7.899946761 | 2.79E-15 | 1.44E-13 |
| DEFB1 | 93.00115669 | 2.04970503 | 0.26025836 | 7.875654908 | 3.39E-15 | 1.74E-13 |
| KCNAB1 | 414.2832614 | -1.267442002 | 0.160988687 | -7.872863786 | 3.47E-15 | 1.77E-13 |
| NBN | 963.0515701 | 1.092551307 | 0.138775934 | 7.872772115 | 3.47E-15 | 1.77E-13 |
| NCS1 | 1117.074122 | 1.076082291 | 0.136850549 | 7.863193116 | 3.74E-15 | 1.91E-13 |
| DYNLT1 | 1081.052856 | 0.789995388 | 0.100595258 | 7.853207023 | 4.06E-15 | 2.06E-13 |
| TAGAP | 452.5476629 | 1.012066258 | 0.128874749 | 7.853099712 | 4.06E-15 | 2.06E-13 |
| ACSL5 | 1201.996959 | 0.754777855 | 0.096119903 | 7.852461699 | 4.08E-15 | 2.06E-13 |
| FADS3 | 276.26096 | 1.010621188 | 0.12870218 | 7.852401493 | 4.08E-15 | 2.06E-13 |
| CKB | 104.8980404 | 1.635828984 | 0.208522503 | 7.844855879 | 4.33E-15 | 2.18E-13 |

| | | | | | | |
|---|---|---|---|---|---|---|
| PPP1R15B | 804.8212162 | 0.529203164 | 0.067510293 | 7.838851509 | 4.55E-15 | 2.28E-13 |
| PNPLA1 | 27.68644537 | 3.469732976 | 0.442793401 | 7.836008781 | 4.65E-15 | 2.32E-13 |
| SNAI2 | 18.17638385 | -2.728703248 | 0.348842116 | -7.822172613 | 5.19E-15 | 2.59E-13 |
| ZC3H12C | 257.5807517 | 0.93577995 | 0.11967287 | 7.819482815 | 5.30E-15 | 2.64E-13 |
| CXCL5 | 5778.527352 | 1.648223784 | 0.210801924 | 7.818827023 | 5.33E-15 | 2.64E-13 |
| SOWAHD | 220.433638 | -0.788866259 | 0.1008984 | -7.818421919 | 5.35E-15 | 2.64E-13 |
| ZBP1 | 30.64115028 | 2.368604068 | 0.303395845 | 7.806975971 | 5.86E-15 | 2.89E-13 |
| F3 | 217.4193051 | 1.490019148 | 0.190917885 | 7.804502682 | 5.97E-15 | 2.94E-13 |
| ZBTB4 | 1159.150176 | -0.575674575 | 0.073767235 | -7.803933171 | 6.00E-15 | 2.94E-13 |
| SLAMF8 | 5007.316661 | 0.501039947 | 0.064221123 | 7.801793655 | 6.10E-15 | 2.99E-13 |
| APOBEC3A | 50.66998298 | 3.122728941 | 0.400543769 | 7.796223989 | 6.38E-15 | 3.11E-13 |
| ADAMTS15 | 69.77860357 | -1.778847406 | 0.228204061 | -7.794985772 | 6.44E-15 | 3.13E-13 |
| EREG | 100.8094775 | 2.549051021 | 0.327046842 | 7.794146567 | 6.48E-15 | 3.15E-13 |
| UBE2L6 | 1162.099809 | 0.84697905 | 0.108682183 | 7.793172946 | 6.53E-15 | 3.16E-13 |
| TBX21 | 77.6287017 | 1.336561622 | 0.171619007 | 7.787958026 | 6.81E-15 | 3.29E-13 |
| ACSL4 | 2042.423169 | 0.508842777 | 0.065375845 | 7.783345289 | 7.06E-15 | 3.40E-13 |
| PLEK | 2932.749915 | 0.771661298 | 0.099274058 | 7.773040757 | 7.66E-15 | 3.68E-13 |
| RASGRP4 | 121.2621953 | -1.261556199 | 0.162957225 | -7.741640171 | 9.81E-15 | 4.70E-13 |
| OR2A7 | 89.97514511 | 1.046409875 | 0.135234315 | 7.737754099 | 1.01E-14 | 4.83E-13 |
| ECSCR | 18.76961013 | 3.353105117 | 0.433932165 | 7.727256433 | 1.10E-14 | 5.24E-13 |
| LRRC25 | 1216.391862 | -0.856204413 | 0.110886784 | -7.721428876 | 1.15E-14 | 5.47E-13 |
| NINJ2 | 194.8966116 | -1.105338442 | 0.143220961 | -7.717714173 | 1.18E-14 | 5.62E-13 |
| MAP7 | 498.5996862 | 0.946842711 | 0.122809589 | 7.709843498 | 1.26E-14 | 5.96E-13 |
| IFIH1 | 661.2281369 | 1.39153719 | 0.180932112 | 7.690935412 | 1.46E-14 | 6.89E-13 |
| WHAMMP | 1045.249358 | -0.660021713 | 0.085852402 | -7.687865426 | 1.50E-14 | 7.04E-13 |
| STAP2 | 68.41349214 | 1.516084625 | 0.197393141 | 7.680533458 | 1.58E-14 | 7.44E-13 |
| MAML3 | 19.27124599 | -2.318318733 | 0.301879853 | -7.679607331 | 1.60E-14 | 7.47E-13 |
| TRHDE | 111.1440934 | -1.173643798 | 0.15283389 | -7.679211702 | 1.60E-14 | 7.48E-13 |
| CYP3A7 | 16.67069487 | 5.229068392 | 0.681060471 | 7.67783275 | 1.62E-14 | 7.54E-13 |
| NUMB | 4003.267552 | 0.568657224 | 0.074136464 | 7.670411994 | 1.71E-14 | 7.97E-13 |
| CECR1 | 6198.500287 | -0.842032658 | 0.109829922 | -7.666696319 | 1.76E-14 | 8.18E-13 |
| PARP10 | 815.849602 | 0.882677595 | 0.115200206 | 7.662118182 | 1.83E-14 | 8.46E-13 |
| DNASE2 | 2689.041771 | -0.509384662 | 0.066631628 | -7.644787839 | 2.09E-14 | 9.65E-13 |
| DDX60L | 2263.417331 | 1.136983415 | 0.148759253 | 7.643110545 | 2.12E-14 | 9.76E-13 |
| USP12 | 764.2872825 | 0.667119861 | 0.087361 | 7.636357845 | 2.23E-14 | 1.03E-12 |
| MMP14 | 5914.251497 | 0.832412176 | 0.10915369 | 7.626056249 | 2.42E-14 | 1.11E-12 |
| HN1 | 1094.55085 | -0.664560519 | 0.087149987 | -7.625480395 | 2.43E-14 | 1.11E-12 |
| ME3 | 85.32598161 | -1.001601018 | 0.131370041 | -7.624272718 | 2.45E-14 | 1.12E-12 |
| GPR84 | 214.4750318 | 2.042238289 | 0.267914034 | 7.622737243 | 2.48E-14 | 1.13E-12 |
| CKAP4 | 265.6767271 | 1.127293775 | 0.147994213 | 7.617147682 | 2.59E-14 | 1.18E-12 |
| MXD4 | 431.8024954 | -0.819774337 | 0.107635488 | -7.616208661 | 2.61E-14 | 1.18E-12 |
| CADM1 | 203.2541146 | -1.276671636 | 0.167688484 | -7.613353073 | 2.67E-14 | 1.20E-12 |
| PNKD | 908.9846027 | 0.901824365 | 0.11845406 | 7.613283718 | 2.67E-14 | 1.20E-12 |
| HK3 | 2763.652474 | 0.801870708 | 0.105331236 | 7.61284821 | 2.68E-14 | 1.20E-12 |
| CYP2U1 | 69.95292597 | -1.260153889 | 0.165547635 | -7.612031972 | 2.70E-14 | 1.21E-12 |
| IFI44 | 865.1960987 | 1.934623757 | 0.254645266 | 7.597328564 | 3.02E-14 | 1.35E-12 |
| SASH3 | 1189.119045 | -0.77761091 | 0.102618291 | -7.577702821 | 3.52E-14 | 1.57E-12 |

| | | | | | | |
|---|---|---|---|---|---|---|
| BMP6 | 24.39980155 | 2.502209426 | 0.330221123 | 7.57737544 | 3.53E-14 | 1.57E-12 |
| EEF1B2 | 2309.364643 | -0.515131305 | 0.068141629 | -7.559715159 | 4.04E-14 | 1.79E-12 |
| AK8 | 92.22563718 | 0.995057436 | 0.131640529 | 7.558898817 | 4.06E-14 | 1.80E-12 |
| RPS3 | 5072.165844 | -0.478544325 | 0.063313313 | -7.558352267 | 4.08E-14 | 1.80E-12 |
| TLR8 | 732.5014515 | -0.698807439 | 0.092515427 | -7.553415275 | 4.24E-14 | 1.87E-12 |
| CLEC2D | 122.6830888 | 1.129728741 | 0.149591651 | 7.552084168 | 4.28E-14 | 1.88E-12 |
| CARS2 | 1094.318905 | -0.389001373 | 0.051523687 | -7.549952203 | 4.35E-14 | 1.91E-12 |
| SNX10 | 1057.54569 | 1.519127009 | 0.201275788 | 7.547490066 | 4.44E-14 | 1.94E-12 |
| RPL30 | 6312.012858 | -0.430462882 | 0.057161927 | -7.530587337 | 5.05E-14 | 2.20E-12 |
| SRPX2 | 85.70387263 | 1.184286129 | 0.157377128 | 7.525147684 | 5.27E-14 | 2.29E-12 |
| CHST3 | 55.39688646 | 1.977184836 | 0.262820837 | 7.52293791 | 5.36E-14 | 2.33E-12 |
| IER3 | 981.029595 | 1.345491151 | 0.17892727 | 7.519765739 | 5.49E-14 | 2.38E-12 |
| ST5 | 60.27372252 | -1.400649985 | 0.186319901 | -7.51744704 | 5.59E-14 | 2.41E-12 |
| OVOL1 | 104.783451 | 1.819871726 | 0.242396491 | 7.507830332 | 6.01E-14 | 2.59E-12 |
| CHPT1 | 610.5521351 | -0.783064366 | 0.104351796 | -7.504081331 | 6.19E-14 | 2.66E-12 |
| NLRP12 | 19.97445398 | -1.235599506 | 0.164893126 | -7.493335435 | 6.71E-14 | 2.88E-12 |
| DESI1 | 536.8677275 | 0.644874579 | 0.086209818 | 7.480291655 | 7.42E-14 | 3.18E-12 |
| PLA1A | 41.64300534 | 2.037134738 | 0.272568029 | 7.473857967 | 7.79E-14 | 3.33E-12 |
| SORD | 167.3720392 | -0.621490118 | 0.083196083 | -7.470184795 | 8.01E-14 | 3.41E-12 |
| CYP3A5 | 34.15091915 | 3.41722773 | 0.457637528 | 7.467105568 | 8.20E-14 | 3.49E-12 |
| DNAJB5 | 160.8784881 | 1.035855653 | 0.138786166 | 7.463680861 | 8.41E-14 | 3.57E-12 |
| IL27 | 37.68209581 | 1.89462678 | 0.253860966 | 7.463245759 | 8.44E-14 | 3.58E-12 |
| PXN | 646.4060262 | 1.324824743 | 0.17764694 | 7.457627731 | 8.81E-14 | 3.72E-12 |
| MSANTD3 | 554.9579933 | 1.131300556 | 0.151931818 | 7.44610687 | 9.61E-14 | 4.05E-12 |
| RASGRP2 | 117.3440212 | -1.088009921 | 0.146167061 | -7.44360538 | 9.80E-14 | 4.12E-12 |
| ZNF267 | 1371.363666 | 0.608469077 | 0.081967585 | 7.423289026 | 1.14E-13 | 4.79E-12 |
| SLC27A3 | 489.9267493 | 0.901158115 | 0.12142203 | 7.421701961 | 1.16E-13 | 4.84E-12 |
| MX2 | 426.9555556 | 1.805553895 | 0.243405664 | 7.417879547 | 1.19E-13 | 4.97E-12 |
| RPS18 | 1092.150846 | -0.448566872 | 0.060569336 | -7.405840976 | 1.30E-13 | 5.43E-12 |
| HSPA9 | 4024.355665 | 0.316449771 | 0.042739053 | 7.404229818 | 1.32E-13 | 5.49E-12 |
| EMR2 | 893.1922134 | 0.808327379 | 0.109185332 | 7.403259806 | 1.33E-13 | 5.51E-12 |
| PTGFRN | 111.8095134 | -1.006223836 | 0.13619146 | -7.388303447 | 1.49E-13 | 6.16E-12 |
| RAB32 | 655.5510941 | -0.620272741 | 0.083975357 | -7.386366189 | 1.51E-13 | 6.23E-12 |
| GZMB | 120.5470434 | 1.695959059 | 0.229621278 | 7.385896771 | 1.51E-13 | 6.24E-12 |
| SLED1 | 54.60054873 | 1.306050519 | 0.176853567 | 7.384926069 | 1.53E-13 | 6.27E-12 |
| ITGA5 | 3517.75822 | 1.070873876 | 0.145301519 | 7.370011552 | 1.71E-13 | 7.00E-12 |
| ZNF586 | 301.7032101 | 0.754674522 | 0.102451101 | 7.366192408 | 1.76E-13 | 7.19E-12 |
| TRPV4 | 268.0349381 | -0.710202633 | 0.096453752 | -7.363141594 | 1.80E-13 | 7.34E-12 |
| MGAT4A | 715.7853945 | -0.66472231 | 0.090288323 | -7.36221796 | 1.81E-13 | 7.38E-12 |
| HIPK2 | 4944.374995 | -0.773243536 | 0.10527865 | -7.344732621 | 2.06E-13 | 8.39E-12 |
| ZNF503 | 29.6885835 | -1.646426888 | 0.224175256 | -7.344373861 | 2.07E-13 | 8.39E-12 |
| CHN2 | 150.1624242 | -0.632346055 | 0.086142839 | -7.340668858 | 2.13E-13 | 8.61E-12 |
| UBA7 | 837.8837334 | 0.5612939 | 0.076467818 | 7.340263084 | 2.13E-13 | 8.62E-12 |
| SUSD1 | 2719.908641 | -0.522693589 | 0.071306884 | -7.330198173 | 2.30E-13 | 9.27E-12 |
| NPC1 | 2248.826811 | 0.365822684 | 0.049934637 | 7.326030692 | 2.37E-13 | 9.54E-12 |
| PRKACB | 583.1481043 | -0.645948652 | 0.088225055 | -7.321600969 | 2.45E-13 | 9.84E-12 |
| CD22 | 128.5596923 | -1.072079736 | 0.146453947 | -7.320251552 | 2.48E-13 | 9.92E-12 |

| | | | | | | |
|---|---|---|---|---|---|---|
| MCOLN2 | 128.2694049 | 1.589128693 | 0.21714388 | 7.318321336 | 2.51E-13 | 1.00E-11 |
| C17orf96 | 29.96144087 | 2.053348157 | 0.280600519 | 7.317691948 | 2.52E-13 | 1.01E-11 |
| HIST1H2Bk | 491.1586839 | 0.753449048 | 0.102985959 | 7.316036597 | 2.55E-13 | 1.02E-11 |
| TOR1B | 973.7637487 | 0.91225714 | 0.124770649 | 7.311472293 | 2.64E-13 | 1.05E-11 |
| EIF3F | 1516.519031 | -0.526154205 | 0.072069614 | -7.300638591 | 2.86E-13 | 1.14E-11 |
| CLEC6A | 437.0628092 | 1.570172877 | 0.215213801 | 7.295874466 | 2.97E-13 | 1.17E-11 |
| RPL29 | 5205.60302 | -0.423984034 | 0.058190379 | -7.286153495 | 3.19E-13 | 1.26E-11 |
| ABHD6 | 657.8493772 | -0.523958179 | 0.071964803 | -7.280756116 | 3.32E-13 | 1.31E-11 |
| GPR65 | 287.2792798 | -1.03951383 | 0.142784068 | -7.280320847 | 3.33E-13 | 1.31E-11 |
| TRIM5 | 402.0230563 | 0.827060316 | 0.113677034 | 7.275526849 | 3.45E-13 | 1.35E-11 |
| IQSEC3 | 110.651359 | -1.439760434 | 0.197949441 | -7.273374597 | 3.51E-13 | 1.37E-11 |
| C7orf41 | 42.48511864 | -1.490266665 | 0.204931244 | -7.272032476 | 3.54E-13 | 1.38E-11 |
| NNMT | 72.79388445 | 4.31577394 | 0.593919968 | 7.26659175 | 3.69E-13 | 1.44E-11 |
| EBI3 | 23.05978092 | 4.090451929 | 0.562969884 | 7.265845016 | 3.71E-13 | 1.44E-11 |
| NFKBIA | 3008.739479 | 0.794269106 | 0.109320734 | 7.265493743 | 3.72E-13 | 1.44E-11 |
| BLVRB | 2026.12202 | -0.583092115 | 0.080351346 | -7.256780959 | 3.96E-13 | 1.54E-11 |
| STBD1 | 50.2264039 | 1.000147431 | 0.137882905 | 7.253599961 | 4.06E-13 | 1.57E-11 |
| PSMA3 | 160.0515333 | 0.982295333 | 0.135560658 | 7.246168218 | 4.29E-13 | 1.65E-11 |
| SMPD2 | 236.9514466 | 0.642812535 | 0.088714729 | 7.245837793 | 4.30E-13 | 1.65E-11 |
| RPS12 | 4208.033635 | -0.521839397 | 0.072223593 | -7.22533144 | 5.00E-13 | 1.92E-11 |
| C3orf37 | 257.701358 | 0.524102683 | 0.072603675 | 7.218679813 | 5.25E-13 | 2.01E-11 |
| ABI3 | 293.9688254 | -1.175982432 | 0.162990933 | -7.21501747 | 5.39E-13 | 2.06E-11 |
| CLEC4A | 528.4290508 | -0.913370453 | 0.126692308 | -7.209359956 | 5.62E-13 | 2.15E-11 |
| C6orf106 | 1954.520108 | 0.377230792 | 0.05233528 | 7.207963552 | 5.68E-13 | 2.16E-11 |
| CLCF1 | 198.6046006 | 1.287497717 | 0.178624397 | 7.20784921 | 5.68E-13 | 2.16E-11 |
| EAF1 | 869.048365 | 0.680916679 | 0.09448008 | 7.206986695 | 5.72E-13 | 2.17E-11 |
| HSH2D | 20.40601476 | 2.995827018 | 0.416223129 | 7.197646668 | 6.13E-13 | 2.32E-11 |
| HSD3B7 | 2514.021838 | 0.619102767 | 0.086064334 | 7.193488177 | 6.32E-13 | 2.39E-11 |
| PDK4 | 750.5002433 | -1.173572 | 0.163263107 | -7.188225333 | 6.56E-13 | 2.47E-11 |
| USF2 | 2862.601944 | -0.502569424 | 0.069913911 | -7.188403828 | 6.56E-13 | 2.47E-11 |
| ARAP3 | 109.9001256 | 1.132991555 | 0.157625168 | 7.187884837 | 6.58E-13 | 2.47E-11 |
| LIMK1 | 5156.516915 | 0.459693421 | 0.064008018 | 7.181810017 | 6.88E-13 | 2.58E-11 |
| MBNL1 | 376.4311701 | 0.495604675 | 0.069122334 | 7.169964429 | 7.50E-13 | 2.81E-11 |
| C15orf26 | 7.095787696 | 3.193945137 | 0.445555122 | 7.168462397 | 7.58E-13 | 2.83E-11 |
| RBP7 | 55.72009154 | -1.241189735 | 0.173628969 | -7.148517571 | 8.77E-13 | 3.27E-11 |
| SSTR2 | 7.252278054 | 3.063163528 | 0.429149401 | 7.137755575 | 9.49E-13 | 3.53E-11 |
| JHDM1D | 144.4655126 | 0.679890594 | 0.095295623 | 7.134541689 | 9.71E-13 | 3.60E-11 |
| TIFA | 222.2667583 | 1.047970644 | 0.146985592 | 7.129750811 | 1.01E-12 | 3.72E-11 |
| SLC25A25 | 170.0283974 | 0.637611733 | 0.089482066 | 7.125581227 | 1.04E-12 | 3.83E-11 |
| KRT23 | 33.59019243 | 3.886725295 | 0.546011413 | 7.118395703 | 1.09E-12 | 4.03E-11 |
| CCDC115 | 737.5561857 | 0.736606077 | 0.103564221 | 7.112553669 | 1.14E-12 | 4.19E-11 |
| SLC43A3 | 4002.779541 | 0.476702262 | 0.067021589 | 7.112667288 | 1.14E-12 | 4.19E-11 |
| RPL18A | 4232.98397 | -0.396951039 | 0.055818348 | -7.111479579 | 1.15E-12 | 4.21E-11 |
| LDLRAP1 | 408.4673323 | 0.821080126 | 0.115705542 | 7.096290412 | 1.28E-12 | 4.69E-11 |
| NT5DC2 | 36.99995403 | -1.615691279 | 0.227947526 | -7.087996544 | 1.36E-12 | 4.97E-11 |
| TTC3 | 213.6097438 | -0.770935085 | 0.108805569 | -7.085437765 | 1.39E-12 | 5.05E-11 |
| C1RL | 1017.409618 | 0.851848572 | 0.12034873 | 7.078168337 | 1.46E-12 | 5.32E-11 |

| | | | | | | |
|---|---|---|---|---|---|---|
| RFTN1 | 367.3319212 | 0.922589959 | 0.130450869 | 7.072317482 | 1.52E-12 | 5.53E-11 |
| MSMO1 | 458.5295832 | 0.976842065 | 0.138130552 | 7.071875507 | 1.53E-12 | 5.54E-11 |
| RPS6 | 4434.007205 | -0.486226959 | 0.068894069 | -7.057602602 | 1.69E-12 | 6.13E-11 |
| MGC1291( | 50.28167004 | 2.189431746 | 0.310292517 | 7.056024963 | 1.71E-12 | 6.19E-11 |
| VWA8 | 717.6579118 | 1.261413849 | 0.178814742 | 7.054305665 | 1.73E-12 | 6.25E-11 |
| ANKRD22 | 91.94869677 | 2.398332121 | 0.340012845 | 7.053651525 | 1.74E-12 | 6.27E-11 |
| UBD | 1234.500547 | 2.598764283 | 0.368437095 | 7.053481626 | 1.74E-12 | 6.27E-11 |
| GPR82 | 26.37675128 | -2.490046866 | 0.353251455 | -7.048935919 | 1.80E-12 | 6.46E-11 |
| NFE2L3 | 209.8749397 | 0.899625291 | 0.127661252 | 7.046972176 | 1.83E-12 | 6.54E-11 |
| OTUD7B | 60.6401071 | 1.321600281 | 0.187602122 | 7.044697945 | 1.86E-12 | 6.63E-11 |
| RPLP0P2 | 31.07308323 | 1.767646677 | 0.251247642 | 7.035475674 | 1.99E-12 | 7.08E-11 |
| PPM1L | 74.50698275 | -0.615340673 | 0.087551151 | -7.028356183 | 2.09E-12 | 7.43E-11 |
| MMP10 | 12.65992575 | 4.199542305 | 0.597924972 | 7.023527206 | 2.16E-12 | 7.68E-11 |
| CPVL | 850.1631708 | -1.147193024 | 0.163537622 | -7.014856941 | 2.30E-12 | 8.15E-11 |
| MYO7A | 25.33902646 | -1.5832114 | 0.225829285 | -7.010655871 | 2.37E-12 | 8.39E-11 |
| GAB3 | 163.8284018 | -0.650432066 | 0.092784064 | -7.010170037 | 2.38E-12 | 8.40E-11 |
| CST3 | 13442.50268 | -0.51010279 | 0.072809403 | -7.006001574 | 2.45E-12 | 8.64E-11 |
| SLC45A3 | 82.96204549 | -1.005094717 | 0.143476481 | -7.005292527 | 2.46E-12 | 8.67E-11 |
| PLAUR | 12820.6138 | 0.806166844 | 0.115145856 | 7.001266647 | 2.54E-12 | 8.90E-11 |
| TRAFD1 | 862.9049422 | 0.49809127 | 0.071231703 | 6.992550375 | 2.70E-12 | 9.45E-11 |
| TCF7L1 | 14.06852983 | 1.964413432 | 0.280966631 | 6.991625398 | 2.72E-12 | 9.50E-11 |
| CXCR2 | 25.01367818 | -1.729780004 | 0.247471867 | -6.989804635 | 2.75E-12 | 9.61E-11 |
| SDC4 | 1254.788637 | 1.092522818 | 0.15642364 | 6.984384304 | 2.86E-12 | 9.97E-11 |
| PMAIP1 | 38.00075998 | 1.105443025 | 0.158283881 | 6.983926728 | 2.87E-12 | 9.98E-11 |
| LIF | 61.21341281 | 2.422284076 | 0.347506035 | 6.970480592 | 3.16E-12 | 1.10E-10 |
| NOV | 11.89700398 | -2.015675867 | 0.28939344 | -6.965174698 | 3.28E-12 | 1.14E-10 |
| PKD1L1 | 144.6514332 | 1.697488997 | 0.24375746 | 6.963844297 | 3.31E-12 | 1.14E-10 |
| ANKLE2 | 1191.122891 | 0.509734909 | 0.073200495 | 6.96354457 | 3.32E-12 | 1.15E-10 |
| HMGN2P4 | 17.08614817 | 2.203993443 | 0.317348802 | 6.94501894 | 3.78E-12 | 1.30E-10 |
| CD69 | 70.96032989 | 1.497068146 | 0.215670021 | 6.941475404 | 3.88E-12 | 1.33E-10 |
| INPP5D | 495.0884328 | -0.642564104 | 0.092592161 | -6.939724654 | 3.93E-12 | 1.35E-10 |
| PITPNB | 947.1527773 | 0.327537908 | 0.047223937 | 6.935844971 | 4.04E-12 | 1.38E-10 |
| PTPRF | 33.90826429 | 3.229351625 | 0.465683999 | 6.934641588 | 4.07E-12 | 1.39E-10 |
| NDP | 9.206915458 | 4.287018497 | 0.618452272 | 6.931850189 | 4.15E-12 | 1.42E-10 |
| SCARF1 | 195.4889453 | 0.994914229 | 0.143703474 | 6.923383296 | 4.41E-12 | 1.50E-10 |
| SPATA7 | 56.17795838 | -1.142322421 | 0.165080714 | -6.919781201 | 4.52E-12 | 1.54E-10 |
| LYZ | 63948.15382 | -0.51935031 | 0.075073822 | -6.917861632 | 4.59E-12 | 1.56E-10 |
| UXS1 | 954.905238 | 0.650060309 | 0.094047544 | 6.912039211 | 4.78E-12 | 1.62E-10 |
| LOC64563{ | 90.45258929 | 1.410827628 | 0.204150651 | 6.910718239 | 4.82E-12 | 1.63E-10 |
| TNFSF12 | 1220.577172 | -0.618675666 | 0.089728806 | -6.89495039 | 5.39E-12 | 1.82E-10 |
| CLEC4E | 1528.558242 | 1.41346579 | 0.205037084 | 6.893708016 | 5.44E-12 | 1.83E-10 |
| TRIM25 | 1168.433578 | 0.464299353 | 0.067456499 | 6.882944701 | 5.86E-12 | 1.97E-10 |
| ENTPD1 | 269.7721598 | 0.767987578 | 0.111666832 | 6.877490506 | 6.09E-12 | 2.05E-10 |
| ARL4C | 30.64511163 | -1.781993373 | 0.259347904 | -6.871053684 | 6.37E-12 | 2.14E-10 |
| C15orf38 | 164.5234791 | -0.739642241 | 0.107708752 | -6.867057941 | 6.55E-12 | 2.19E-10 |
| PLEKHB1 | 71.33009722 | -1.144920745 | 0.166744821 | -6.866304697 | 6.59E-12 | 2.20E-10 |
| UNC5B | 266.614222 | -0.705432031 | 0.102781325 | -6.863426094 | 6.72E-12 | 2.24E-10 |

| | | | | | | |
|---|---|---|---|---|---|---|
| FURIN | 1098.579198 | 0.444279043 | 0.064743271 | 6.862165514 | 6.78E-12 | 2.26E-10 |
| NFIA | 156.6630887 | -0.901214111 | 0.131373408 | -6.859943168 | 6.89E-12 | 2.29E-10 |
| RPL7 | 6109.838027 | -0.544556627 | 0.079397084 | -6.858647701 | 6.95E-12 | 2.30E-10 |
| SCD | 7689.667697 | -0.613812206 | 0.089492678 | -6.858798039 | 6.94E-12 | 2.30E-10 |
| ARHGAP15 | 242.6622434 | -0.585497145 | 0.085482335 | -6.849334913 | 7.42E-12 | 2.45E-10 |
| GTPBP2 | 815.3952998 | 0.608062361 | 0.088782383 | 6.848907833 | 7.44E-12 | 2.45E-10 |
| C19orf66 | 317.8150737 | 0.730335016 | 0.106848837 | 6.835217275 | 8.19E-12 | 2.69E-10 |
| MT1X | 311.6162287 | 1.839265516 | 0.269086845 | 6.835211567 | 8.19E-12 | 2.69E-10 |
| BATF3 | 76.27889229 | 1.082844274 | 0.158613853 | 6.826921201 | 8.68E-12 | 2.85E-10 |
| EIF3L | 2201.221816 | -0.507871711 | 0.074436257 | -6.822907675 | 8.92E-12 | 2.92E-10 |
| B4GALT5 | 1282.08585 | 0.625441793 | 0.091701681 | 6.820396136 | 9.08E-12 | 2.97E-10 |
| CLCN4 | 232.8695715 | -1.001836296 | 0.147031012 | -6.813775423 | 9.51E-12 | 3.10E-10 |
| CPT1B | 105.7172089 | 0.824136794 | 0.121028385 | 6.809450474 | 9.80E-12 | 3.19E-10 |
| TTC39B | 889.9048664 | 0.752716877 | 0.110562434 | 6.808070789 | 9.89E-12 | 3.22E-10 |
| YME1L1 | 902.0404469 | 0.389672194 | 0.057241165 | 6.80755179 | 9.93E-12 | 3.22E-10 |
| EPHX1 | 2283.138735 | -0.645376119 | 0.094837461 | -6.805075858 | 1.01E-11 | 3.27E-10 |
| BACE1 | 735.7236909 | -0.612250639 | 0.089975397 | -6.804645022 | 1.01E-11 | 3.28E-10 |
| TARS | 1449.116436 | 0.417928178 | 0.061605145 | 6.78398172 | 1.17E-11 | 3.78E-10 |
| SLIT3 | 107.6773486 | -1.065378809 | 0.157066156 | -6.782994117 | 1.18E-11 | 3.79E-10 |
| NPL | 2007.033077 | -0.77399851 | 0.114156772 | -6.780136641 | 1.20E-11 | 3.86E-10 |
| MMAA | 171.441098 | 0.522305703 | 0.077243815 | 6.761780251 | 1.36E-11 | 4.38E-10 |
| TRPS1 | 161.8939875 | -0.730211376 | 0.107998802 | -6.761291455 | 1.37E-11 | 4.39E-10 |
| SIRPB1 | 374.7454758 | -0.637102076 | 0.09424189 | -6.760285467 | 1.38E-11 | 4.41E-10 |
| TENM4 | 17.96382265 | -2.454774172 | 0.363166244 | -6.759367679 | 1.39E-11 | 4.43E-10 |
| GBP3 | 487.6779749 | 0.811853446 | 0.120117628 | 6.758820134 | 1.39E-11 | 4.44E-10 |
| CTNNBIP1 | 189.373322 | -0.626003479 | 0.092639643 | -6.757403862 | 1.40E-11 | 4.47E-10 |
| CSF2 | 13.61409724 | 4.070729307 | 0.602517681 | 6.756198918 | 1.42E-11 | 4.50E-10 |
| NCAM1 | 34.23201797 | 1.14878247 | 0.170037736 | 6.756044255 | 1.42E-11 | 4.50E-10 |
| APOL6 | 805.1980349 | 0.766640757 | 0.113479498 | 6.755764433 | 1.42E-11 | 4.50E-10 |
| SYNJ2 | 117.3443131 | 1.093574764 | 0.161891524 | 6.754984678 | 1.43E-11 | 4.52E-10 |
| PARP12 | 987.5231715 | 0.776502101 | 0.114976242 | 6.75358741 | 1.44E-11 | 4.56E-10 |
| GREM1 | 111.1157034 | 1.110859821 | 0.164783391 | 6.74133365 | 1.57E-11 | 4.95E-10 |
| RPLP2 | 13642.31461 | -0.330378964 | 0.049023214 | -6.739235135 | 1.59E-11 | 5.01E-10 |
| EGR2 | 4152.824676 | 0.795738402 | 0.118130963 | 6.73606972 | 1.63E-11 | 5.11E-10 |
| CDR2 | 247.1198718 | -0.544112547 | 0.080839785 | -6.730751986 | 1.69E-11 | 5.29E-10 |
| ALAS1 | 2221.06813 | 0.923855799 | 0.1375867 | 6.714717307 | 1.88E-11 | 5.90E-10 |
| DEF6 | 148.7787664 | -1.029988248 | 0.153520573 | -6.709121977 | 1.96E-11 | 6.11E-10 |
| ZNF552 | 116.6361275 | -0.861429884 | 0.128393057 | -6.709318278 | 1.96E-11 | 6.11E-10 |
| METTL9 | 459.3085671 | 0.546271299 | 0.081499648 | 6.702744284 | 2.05E-11 | 6.37E-10 |
| SEPN1 | 1707.019853 | 0.609702359 | 0.091026406 | 6.698082327 | 2.11E-11 | 6.57E-10 |
| C10orf54 | 422.9957702 | -0.577479544 | 0.086272443 | -6.693673224 | 2.18E-11 | 6.76E-10 |
| PIM3 | 1649.745914 | 0.599019443 | 0.089526502 | 6.690973358 | 2.22E-11 | 6.87E-10 |
| LOC65443[illegible] | 578.7697077 | 0.450994862 | 0.067410975 | 6.690229055 | 2.23E-11 | 6.89E-10 |
| MMP19 | 1199.715853 | 0.861527411 | 0.128776795 | 6.690082736 | 2.23E-11 | 6.89E-10 |
| H2AFV | 1381.289488 | -0.383135832 | 0.057355919 | -6.679970191 | 2.39E-11 | 7.37E-10 |
| APBB2 | 276.1251919 | 0.798343814 | 0.119621521 | 6.673914603 | 2.49E-11 | 7.67E-10 |
| USP30-AS1 | 148.9060556 | 1.047409023 | 0.157205797 | 6.662661577 | 2.69E-11 | 8.27E-10 |

| | | | | | | |
|---|---|---|---|---|---|---|
| LGR4 | 44.89121729 | -0.993848448 | 0.149190756 | -6.661595372 | 2.71E-11 | 8.32E-10 |
| NFKB1 | 653.068121 | 0.610584477 | 0.091664592 | 6.661072316 | 2.72E-11 | 8.33E-10 |
| CXCL3 | 222.1366557 | 1.005544681 | 0.151001026 | 6.659191066 | 2.75E-11 | 8.42E-10 |
| CASP8 | 110.5419929 | -0.69740701 | 0.104767383 | -6.656718845 | 2.80E-11 | 8.55E-10 |
| RIN3 | 1499.76146 | -0.382947152 | 0.05756949 | -6.651911459 | 2.89E-11 | 8.82E-10 |
| RPL36A | 1530.505101 | -0.612385826 | 0.092076007 | -6.650872991 | 2.91E-11 | 8.87E-10 |
| ATP1A1 | 15209.71044 | 0.449552405 | 0.067731735 | 6.637249193 | 3.20E-11 | 9.71E-10 |
| CUL1 | 817.5735053 | 0.447060142 | 0.06739998 | 6.632941784 | 3.29E-11 | 9.99E-10 |
| SLFN5 | 532.6365165 | 0.784092999 | 0.118257791 | 6.630370765 | 3.35E-11 | 1.01E-09 |
| WARS | 12493.94636 | 0.912626844 | 0.137822359 | 6.621761899 | 3.55E-11 | 1.07E-09 |
| PVRL2 | 290.7172049 | 0.787647882 | 0.118987551 | 6.61958225 | 3.60E-11 | 1.09E-09 |
| TLR7 | 117.8677195 | -1.234409377 | 0.186519278 | -6.618132919 | 3.64E-11 | 1.10E-09 |
| BZW1 | 3496.872183 | 0.327201522 | 0.049449279 | 6.616911916 | 3.67E-11 | 1.10E-09 |
| ZNF491 | 24.88870646 | -1.069997598 | 0.161751285 | -6.615079414 | 3.71E-11 | 1.12E-09 |
| DUSP1 | 1796.063335 | 0.563620249 | 0.085254447 | 6.611036349 | 3.82E-11 | 1.15E-09 |
| PAGR1 | 232.5675702 | -0.517128328 | 0.078267193 | -6.607216998 | 3.92E-11 | 1.17E-09 |
| TNFRSF10A | 178.2965648 | 0.766674505 | 0.116103324 | 6.603381166 | 4.02E-11 | 1.20E-09 |
| MYOF | 4202.427584 | 0.822260933 | 0.124876132 | 6.584612429 | 4.56E-11 | 1.36E-09 |
| DAG1 | 286.7507858 | 0.7238478 | 0.109938144 | 6.584136991 | 4.58E-11 | 1.36E-09 |
| BTBD2 | 507.7056289 | -0.597476698 | 0.090768095 | -6.58245276 | 4.63E-11 | 1.38E-09 |
| C1orf54 | 221.2150602 | 0.613913698 | 0.093273032 | 6.581899248 | 4.64E-11 | 1.38E-09 |
| GPNMB | 68939.51668 | -0.662961714 | 0.100757791 | -6.579756338 | 4.71E-11 | 1.40E-09 |
| HIST1H4B | 323.8998949 | -0.705508107 | 0.107231367 | -6.579307225 | 4.73E-11 | 1.40E-09 |
| GNB2L1 | 11471.51703 | -0.348800875 | 0.053104674 | -6.568176518 | 5.09E-11 | 1.51E-09 |
| QPCT | 644.0887018 | 0.995432915 | 0.151574144 | 6.567300269 | 5.12E-11 | 1.51E-09 |
| CHI3L1 | 26263.89662 | 0.838235545 | 0.127667758 | 6.565757544 | 5.18E-11 | 1.53E-09 |
| CFL1P1 | 6.411461731 | 2.974873266 | 0.453466962 | 6.560286668 | 5.37E-11 | 1.58E-09 |
| SERTAD1 | 201.6621855 | 0.743353557 | 0.113433095 | 6.553233485 | 5.63E-11 | 1.65E-09 |
| ADM | 238.2410163 | 1.299426883 | 0.198310276 | 6.552493964 | 5.66E-11 | 1.66E-09 |
| SLCO2B1 | 4427.070368 | -0.821003934 | 0.125392253 | -6.547485302 | 5.85E-11 | 1.71E-09 |
| MAP1LC3A | 336.1861252 | 1.469150955 | 0.22450972 | 6.54381893 | 6.00E-11 | 1.75E-09 |
| PFKP | 421.7181089 | 0.962696423 | 0.147115197 | 6.543827186 | 6.00E-11 | 1.75E-09 |
| TPT1 | 23055.34526 | -0.527890978 | 0.080673951 | -6.543512156 | 6.01E-11 | 1.75E-09 |
| RPL35 | 4832.109421 | -0.455080958 | 0.069596402 | -6.53885758 | 6.20E-11 | 1.80E-09 |
| CDCA7L | 101.036267 | -1.002250421 | 0.153304153 | -6.537659964 | 6.25E-11 | 1.82E-09 |
| SREBF2 | 816.2141987 | 0.708400097 | 0.108442356 | 6.532503746 | 6.47E-11 | 1.88E-09 |
| FKBP9 | 275.9698373 | -0.408142786 | 0.062542367 | -6.525860911 | 6.76E-11 | 1.96E-09 |
| SNCA | 41.60804551 | -1.42965035 | 0.219107997 | -6.52486614 | 6.81E-11 | 1.97E-09 |
| VAMP5 | 105.6867966 | 1.183234878 | 0.181389327 | 6.523178051 | 6.88E-11 | 1.99E-09 |
| CSF1R | 1703.199097 | -0.75432549 | 0.115764951 | -6.516009252 | 7.22E-11 | 2.08E-09 |
| PCNX | 849.096108 | 0.489371425 | 0.07511831 | 6.514675638 | 7.28E-11 | 2.10E-09 |
| HOMER3 | 449.0688289 | -0.874299552 | 0.134225132 | -6.513679979 | 7.33E-11 | 2.11E-09 |
| SNX29 | 586.8880554 | -0.404332054 | 0.062076767 | -6.513419935 | 7.35E-11 | 2.11E-09 |
| TBC1D14 | 197.9366784 | -0.497761089 | 0.076439963 | -6.511791304 | 7.43E-11 | 2.13E-09 |
| TBKBP1 | 232.6519589 | 0.807441627 | 0.124108899 | 6.505912399 | 7.72E-11 | 2.21E-09 |
| RGCC | 5067.042088 | 0.85532322 | 0.131556504 | 6.501565424 | 7.95E-11 | 2.27E-09 |
| GTPBP1 | 324.5862433 | 0.738014224 | 0.11354335 | 6.499845415 | 8.04E-11 | 2.29E-09 |

| | | | | | | |
|---|---|---|---|---|---|---|
| IL1A | 623.2437559 | 1.23010805 | 0.18931053 | 6.497832176 | 8.15E-11 | 2.32E-09 |
| HK2 | 2275.343531 | 0.462435855 | 0.071187441 | 6.496031467 | 8.25E-11 | 2.35E-09 |
| NFKBIZ | 672.1183643 | 0.602065255 | 0.092703526 | 6.494523816 | 8.33E-11 | 2.37E-09 |
| ENPP4 | 1034.289607 | 0.745989241 | 0.114943317 | 6.490061874 | 8.58E-11 | 2.43E-09 |
| TUBE1 | 217.8595316 | 0.555908628 | 0.085780708 | 6.480578693 | 9.14E-11 | 2.58E-09 |
| VILL | 53.44804036 | 1.136471906 | 0.175362867 | 6.480687306 | 9.13E-11 | 2.58E-09 |
| FAM129B | 2176.708204 | 0.876618772 | 0.135310021 | 6.478594599 | 9.26E-11 | 2.61E-09 |
| TK2 | 380.9834278 | -0.484220603 | 0.074767929 | -6.476314204 | 9.40E-11 | 2.65E-09 |
| CXCL9 | 493.5311724 | 3.000948716 | 0.463467814 | 6.474988395 | 9.48E-11 | 2.67E-09 |
| C2orf18 | 1306.720313 | -0.384223354 | 0.059342999 | -6.474619779 | 9.51E-11 | 2.67E-09 |
| RASSF2 | 113.5204871 | -0.998239262 | 0.154219928 | -6.472829262 | 9.62E-11 | 2.70E-09 |
| ZNF652 | 179.0287723 | -0.752041024 | 0.116230439 | -6.47025884 | 9.78E-11 | 2.74E-09 |
| MAZ | 2420.992133 | -0.39632433 | 0.06129598 | -6.465747529 | 1.01E-10 | 2.82E-09 |
| IRF9 | 589.3866833 | 0.489594994 | 0.075725709 | 6.465373516 | 1.01E-10 | 2.82E-09 |
| MT1F | 653.6257707 | 1.72286072 | 0.266486319 | 6.465100085 | 1.01E-10 | 2.82E-09 |
| S100A4 | 5779.02128 | -0.873081605 | 0.135120809 | -6.461488882 | 1.04E-10 | 2.89E-09 |
| PBXIP1 | 227.8385443 | -0.703489578 | 0.108886172 | -6.460779772 | 1.04E-10 | 2.89E-09 |
| PTGS1 | 395.0898392 | -0.845780276 | 0.130907753 | -6.460887584 | 1.04E-10 | 2.89E-09 |
| TMEFF1 | 62.41401475 | 1.019789616 | 0.158187844 | 6.446700279 | 1.14E-10 | 3.17E-09 |
| RPL37A | 4460.892965 | -0.378488516 | 0.058728895 | -6.444672867 | 1.16E-10 | 3.21E-09 |
| PLEKHG6 | 34.18612275 | 1.543424236 | 0.239607322 | 6.441473589 | 1.18E-10 | 3.27E-09 |
| TLE3 | 581.8054725 | -0.581254079 | 0.090250808 | -6.440430724 | 1.19E-10 | 3.29E-09 |
| FUT4 | 413.8010037 | 0.600788914 | 0.093295534 | 6.439632119 | 1.20E-10 | 3.30E-09 |
| BST2 | 3274.595144 | 0.605535293 | 0.09418674 | 6.429092815 | 1.28E-10 | 3.53E-09 |
| TMEM63B | 465.5638075 | 0.644168915 | 0.100211061 | 6.42812189 | 1.29E-10 | 3.55E-09 |
| BRK1 | 1460.780244 | -0.334015677 | 0.052025243 | -6.420261719 | 1.36E-10 | 3.73E-09 |
| TOP1MT | 507.4219573 | -0.842144635 | 0.131203527 | -6.418612781 | 1.38E-10 | 3.77E-09 |
| ERVFRD-1 | 7.737917641 | -2.565667191 | 0.399869658 | -6.416258746 | 1.40E-10 | 3.82E-09 |
| COL1A1 | 21.61506679 | 2.148957947 | 0.335221857 | 6.410554395 | 1.45E-10 | 3.96E-09 |
| ANXA6 | 289.2273381 | -0.680175606 | 0.106155398 | -6.407357681 | 1.48E-10 | 4.04E-09 |
| ZNF449 | 61.0065705 | -0.85533569 | 0.133494656 | -6.407265397 | 1.48E-10 | 4.04E-09 |
| PTRF | 118.110714 | 1.234036474 | 0.19286629 | 6.398404172 | 1.57E-10 | 4.27E-09 |
| MARCO | 2579.433179 | -0.7547516 | 0.117996842 | -6.396371184 | 1.59E-10 | 4.32E-09 |
| DRAM2 | 764.0305837 | -0.447980209 | 0.070075682 | -6.392805558 | 1.63E-10 | 4.41E-09 |
| ODC1 | 586.2355046 | 0.392883244 | 0.061456116 | 6.392907181 | 1.63E-10 | 4.41E-09 |
| CNNM4 | 702.0592585 | 0.567916841 | 0.08887491 | 6.390069351 | 1.66E-10 | 4.48E-09 |
| AKR1C2 | 1220.688673 | -0.740835641 | 0.116017906 | -6.385528485 | 1.71E-10 | 4.61E-09 |
| MATK | 1279.255925 | 0.879606908 | 0.137893991 | 6.378863231 | 1.78E-10 | 4.81E-09 |
| OAS1 | 2518.851683 | 0.877756579 | 0.137644571 | 6.376979287 | 1.81E-10 | 4.86E-09 |
| FOSB | 33.10562764 | 1.74060628 | 0.273176436 | 6.371729233 | 1.87E-10 | 5.03E-09 |
| SLC1A4 | 1140.848556 | 0.571493255 | 0.089699138 | 6.37122348 | 1.88E-10 | 5.04E-09 |
| NOTCH2 | 2527.648134 | -0.369815252 | 0.058048833 | -6.370761174 | 1.88E-10 | 5.04E-09 |
| GPR160 | 42.75057719 | -1.165804447 | 0.18333078 | -6.359021926 | 2.03E-10 | 5.44E-09 |
| PYGL | 821.3965513 | -0.466522459 | 0.073385049 | -6.35718674 | 2.05E-10 | 5.49E-09 |
| LRPAP1 | 2481.023 | -0.434561739 | 0.068360683 | -6.356895826 | 2.06E-10 | 5.50E-09 |
| UCP2 | 3358.243442 | -0.657688823 | 0.103563203 | -6.350603311 | 2.14E-10 | 5.72E-09 |
| ENPP3 | 62.97757835 | -1.167171126 | 0.183802468 | -6.350138474 | 2.15E-10 | 5.73E-09 |

| | | | | | | |
|---|---|---|---|---|---|---|
| CDKN1A | 3506.646485 | 0.934650166 | 0.147223675 | 6.348504508 | 2.17E-10 | 5.78E-09 |
| SNHG8 | 130.1158726 | -0.810547854 | 0.127685424 | -6.348006153 | 2.18E-10 | 5.79E-09 |
| PSMA5 | 899.8665151 | 0.532941522 | 0.083972251 | 6.346638487 | 2.20E-10 | 5.83E-09 |
| MOSPD3 | 80.68781548 | -0.635461845 | 0.100182803 | -6.343023195 | 2.25E-10 | 5.96E-09 |
| STOM | 4929.991363 | 0.767043968 | 0.121011606 | 6.338598353 | 2.32E-10 | 6.13E-09 |
| UBE2S | 420.9254457 | 0.658402355 | 0.103909441 | 6.336309275 | 2.35E-10 | 6.21E-09 |
| TMC8 | 287.9775788 | -0.981637317 | 0.154981919 | -6.333882853 | 2.39E-10 | 6.30E-09 |
| CR1 | 147.2272007 | -0.952293773 | 0.150365773 | -6.333181792 | 2.40E-10 | 6.32E-09 |
| RGS1 | 411.0347983 | 0.796073355 | 0.125700668 | 6.333087702 | 2.40E-10 | 6.32E-09 |
| IMPDH1 | 809.8662302 | -0.557098049 | 0.088128996 | -6.321393326 | 2.59E-10 | 6.80E-09 |
| PAX5 | 5.474027305 | 3.478076284 | 0.551006379 | 6.312225076 | 2.75E-10 | 7.21E-09 |
| FOSL1 | 43.43527858 | 0.963626716 | 0.152719941 | 6.309763536 | 2.79E-10 | 7.31E-09 |
| NUDT3 | 777.3311932 | -0.495574712 | 0.078545784 | -6.309373786 | 2.80E-10 | 7.32E-09 |
| CCL24 | 1640.217569 | 2.075668706 | 0.329000951 | 6.30900519 | 2.81E-10 | 7.33E-09 |
| VPS26B | 303.1308067 | -0.425351011 | 0.06746434 | -6.304827302 | 2.89E-10 | 7.52E-09 |
| CMTM7 | 726.0280841 | -0.506460898 | 0.080455776 | -6.294897969 | 3.08E-10 | 8.01E-09 |
| C11orf82 | 65.00104017 | -1.224956266 | 0.19462036 | -6.294080782 | 3.09E-10 | 8.04E-09 |
| AP1M2 | 13.98919166 | 2.275535863 | 0.361620935 | 6.292599919 | 3.12E-10 | 8.10E-09 |
| PRKCDBP | 166.1515687 | 0.616919799 | 0.098078088 | 6.29008795 | 3.17E-10 | 8.22E-09 |
| RPL13A | 3118.169433 | -0.474664481 | 0.075469356 | -6.289499543 | 3.18E-10 | 8.24E-09 |
| MEGF9 | 780.1245459 | -0.500510471 | 0.079631596 | -6.285325092 | 3.27E-10 | 8.46E-09 |
| ASAP2 | 68.24042694 | 1.377053323 | 0.219311563 | 6.278981851 | 3.41E-10 | 8.80E-09 |
| DOT1L | 323.0504092 | 0.702014449 | 0.11197178 | 6.269565857 | 3.62E-10 | 9.33E-09 |
| TUBA4A | 116.9252386 | 0.950559653 | 0.151642689 | 6.268417287 | 3.65E-10 | 9.39E-09 |
| RMI2 | 37.4019885 | 1.099781187 | 0.175765142 | 6.25710634 | 3.92E-10 | 1.01E-08 |
| MXI1 | 247.268576 | -0.621267581 | 0.099299863 | -6.256479757 | 3.94E-10 | 1.01E-08 |
| CNIH4 | 1004.67454 | 0.720866964 | 0.115254769 | 6.254552136 | 3.99E-10 | 1.02E-08 |
| CLDN12 | 256.4718353 | 0.800458357 | 0.128090666 | 6.249154444 | 4.13E-10 | 1.06E-08 |
| PKD2L1 | 53.5971698 | -1.062669271 | 0.170059951 | -6.248792056 | 4.14E-10 | 1.06E-08 |
| EPSTI1 | 770.4355599 | 1.564049905 | 0.250548402 | 6.242506003 | 4.31E-10 | 1.10E-08 |
| ATP13A3 | 2786.746141 | 0.4678857 | 0.074960787 | 6.241739456 | 4.33E-10 | 1.10E-08 |
| CD300E | 101.1503699 | 1.722687877 | 0.27601255 | 6.241338935 | 4.34E-10 | 1.10E-08 |
| GNGT2 | 94.13022648 | -0.946924058 | 0.151804745 | -6.237776405 | 4.44E-10 | 1.13E-08 |
| PPP2R2A | 536.8999461 | 0.328894251 | 0.052726798 | 6.237705768 | 4.44E-10 | 1.13E-08 |
| COL5A3 | 33.67938848 | 1.834478212 | 0.294199827 | 6.235483659 | 4.50E-10 | 1.14E-08 |
| CASC4 | 598.1225639 | -0.416553922 | 0.066953106 | -6.221577218 | 4.92E-10 | 1.25E-08 |
| MB21D2 | 111.8793403 | 0.569360136 | 0.091582398 | 6.216916659 | 5.07E-10 | 1.28E-08 |
| O3FAR1 | 281.1459028 | -1.050883363 | 0.169095095 | -6.214747781 | 5.14E-10 | 1.30E-08 |
| GRASP | 99.72982941 | 1.355628216 | 0.218194019 | 6.212948566 | 5.20E-10 | 1.31E-08 |
| ZCCHC24 | 177.8649181 | -0.755814763 | 0.121745934 | -6.208131435 | 5.36E-10 | 1.35E-08 |
| ZNF416 | 43.46722499 | -0.590425922 | 0.095133825 | -6.206267029 | 5.43E-10 | 1.36E-08 |
| ARHGDIB | 9796.404203 | -0.424301627 | 0.068436016 | -6.199975562 | 5.65E-10 | 1.42E-08 |
| B3GALNT1 | 243.2070229 | -0.739072401 | 0.119213159 | -6.199587398 | 5.66E-10 | 1.42E-08 |
| CXCL13 | 11.34335274 | 3.507890328 | 0.565815063 | 6.199711813 | 5.66E-10 | 1.42E-08 |
| VEGFA | 220.9701283 | 1.509615275 | 0.243686873 | 6.194897799 | 5.83E-10 | 1.46E-08 |
| RPL26 | 4447.553515 | -0.556228751 | 0.089793467 | -6.194534722 | 5.85E-10 | 1.46E-08 |
| STON1 | 17.0426383 | 2.27342841 | 0.367053905 | 6.19371808 | 5.88E-10 | 1.47E-08 |

| | | | | | | |
|---|---|---|---|---|---|---|
| LAG3 | 20.7690253 | 2.291441938 | 0.369986658 | 6.193309643 | 5.89E-10 | 1.47E-08 |
| FAM165B | 162.2270862 | -0.603876907 | 0.097520545 | -6.192304491 | 5.93E-10 | 1.48E-08 |
| FAM105A | 179.5357111 | -0.653869817 | 0.105648618 | -6.189099556 | 6.05E-10 | 1.50E-08 |
| IL18 | 1618.962136 | 0.617908566 | 0.099892361 | 6.185743951 | 6.18E-10 | 1.53E-08 |
| FPR1 | 303.7498791 | -1.031440557 | 0.16682092 | -6.182920924 | 6.29E-10 | 1.56E-08 |
| XIAP | 573.8306015 | 0.365234991 | 0.059091107 | 6.180879195 | 6.37E-10 | 1.58E-08 |
| SLC45A4 | 192.8267139 | -0.5288871 | 0.085594443 | -6.178988736 | 6.45E-10 | 1.59E-08 |
| HSPA1B | 567.3813708 | 0.562902871 | 0.091123918 | 6.177333902 | 6.52E-10 | 1.61E-08 |
| SPSB1 | 488.4368674 | 0.702351805 | 0.113780395 | 6.172871905 | 6.71E-10 | 1.65E-08 |
| CAMK1D | 215.0519833 | -0.4720043 | 0.076490562 | -6.170752145 | 6.80E-10 | 1.67E-08 |
| RPS15A | 2540.980567 | -0.505805129 | 0.081967642 | -6.170790247 | 6.79E-10 | 1.67E-08 |
| CXCR7 | 24.08721501 | -1.212405472 | 0.196486014 | -6.170441596 | 6.81E-10 | 1.67E-08 |
| HHEX | 132.4836932 | -0.75608519 | 0.12253783 | -6.17021854 | 6.82E-10 | 1.67E-08 |
| MFHAS1 | 507.9245434 | 0.609872062 | 0.098913169 | 6.165731684 | 7.02E-10 | 1.72E-08 |
| TGIF1 | 22.6957735 | 1.086340943 | 0.176560049 | 6.152812874 | 7.61E-10 | 1.86E-08 |
| CDKN2B | 48.64705253 | 0.923403357 | 0.150092227 | 6.152239681 | 7.64E-10 | 1.86E-08 |
| MCL1 | 2921.985881 | 0.589972192 | 0.095895923 | 6.152213509 | 7.64E-10 | 1.86E-08 |
| CDK14 | 1175.110656 | 0.675442776 | 0.109818795 | 6.150520729 | 7.72E-10 | 1.88E-08 |
| WBP1 | 384.3144018 | -0.530275116 | 0.086248687 | -6.148210895 | 7.84E-10 | 1.91E-08 |
| SLC25A28 | 116.3847672 | 0.891286507 | 0.145058727 | 6.144314955 | 8.03E-10 | 1.95E-08 |
| ACSL1 | 4091.009222 | 1.201415297 | 0.195573947 | 6.143023217 | 8.10E-10 | 1.96E-08 |
| AGPAT3 | 1435.894729 | 0.269426619 | 0.043876934 | 6.140506968 | 8.23E-10 | 1.99E-08 |
| RPS4X | 10267.73912 | -0.437028856 | 0.071175988 | -6.140116485 | 8.25E-10 | 2.00E-08 |
| RPL27A | 5236.594656 | -0.565913227 | 0.092287386 | -6.132075608 | 8.67E-10 | 2.10E-08 |
| MUM1 | 75.78203339 | -0.56361287 | 0.091942776 | -6.130039734 | 8.79E-10 | 2.12E-08 |
| NDRG1 | 1433.744986 | 0.833370375 | 0.135953462 | 6.129820918 | 8.80E-10 | 2.12E-08 |
| ELF4 | 746.0435439 | 0.479616458 | 0.078259767 | 6.128518841 | 8.87E-10 | 2.14E-08 |
| NPC2 | 8989.018187 | -0.421798243 | 0.068831227 | -6.128007072 | 8.90E-10 | 2.14E-08 |
| CNRIP1 | 92.52540876 | -0.801212251 | 0.130780572 | -6.126385891 | 8.99E-10 | 2.16E-08 |
| RRN3P1 | 130.1372555 | -0.669279753 | 0.109268764 | -6.125078482 | 9.06E-10 | 2.17E-08 |
| SLC39A4 | 131.8765708 | -0.738938898 | 0.120743913 | -6.119885313 | 9.36E-10 | 2.24E-08 |
| ENDOD1 | 78.93617085 | 0.757732577 | 0.123829777 | 6.119146761 | 9.41E-10 | 2.25E-08 |
| IRF4 | 24.09335441 | 1.757678987 | 0.287326252 | 6.117363015 | 9.51E-10 | 2.27E-08 |
| KIAA0247 | 1490.394693 | 0.453403519 | 0.074155462 | 6.114229593 | 9.70E-10 | 2.31E-08 |
| CXCR4 | 915.2070598 | -1.16967129 | 0.191502515 | -6.107863856 | 1.01E-09 | 2.41E-08 |
| MARVELD1 | 509.150339 | -0.50898754 | 0.083339263 | -6.107415897 | 1.01E-09 | 2.41E-08 |
| ELMO1 | 259.434537 | -0.666690546 | 0.109219167 | -6.10415336 | 1.03E-09 | 2.46E-08 |
| TSPAN13 | 99.38790734 | -0.882477288 | 0.144629867 | -6.101625528 | 1.05E-09 | 2.49E-08 |
| RAB13 | 1940.848698 | 0.488679888 | 0.080173733 | 6.095261753 | 1.09E-09 | 2.59E-08 |
| HMOX1 | 4079.107474 | -0.488374826 | 0.08017178 | -6.091605132 | 1.12E-09 | 2.65E-08 |
| CRTAP | 1472.210341 | -0.57034716 | 0.093733429 | -6.084778541 | 1.17E-09 | 2.75E-08 |
| ELMSAN1 | 267.4165858 | -0.466269726 | 0.076627213 | -6.08491039 | 1.17E-09 | 2.75E-08 |
| NR4A2 | 35.89997513 | 1.444082105 | 0.237324408 | 6.084844449 | 1.17E-09 | 2.75E-08 |
| SYT11 | 215.1954616 | 0.58544112 | 0.096238986 | 6.08320126 | 1.18E-09 | 2.78E-08 |
| ARHGAP4 | 193.7857438 | -0.568362963 | 0.093436382 | -6.082887135 | 1.18E-09 | 2.78E-08 |
| LILRB2 | 603.2587895 | -0.923764078 | 0.151966306 | -6.078742735 | 1.21E-09 | 2.85E-08 |
| CCND1 | 296.4333664 | 0.879038934 | 0.144620814 | 6.078232509 | 1.22E-09 | 2.85E-08 |

| | | | | | | |
|---|---|---|---|---|---|---|
| BHLHE40 | 2455.453728 | 0.85828289 | 0.141265556 | 6.07566994 | 1.23E-09 | 2.89E-08 |
| EXT1 | 95.80465205 | 0.698342735 | 0.115238601 | 6.059972346 | 1.36E-09 | 3.19E-08 |
| RBPJ | 869.0650655 | -0.298484145 | 0.049331252 | -6.050609608 | 1.44E-09 | 3.37E-08 |
| BAG3 | 1699.27894 | 0.645168214 | 0.106723048 | 6.045256639 | 1.49E-09 | 3.48E-08 |
| HLA-E | 11911.35907 | 0.82179664 | 0.136129677 | 6.036866167 | 1.57E-09 | 3.66E-08 |
| TSPAN4 | 573.7617719 | -0.60757973 | 0.10072829 | -6.031867806 | 1.62E-09 | 3.78E-08 |
| FXYD6 | 927.0714202 | -0.704457717 | 0.116878686 | -6.027255594 | 1.67E-09 | 3.88E-08 |
| FBXO10 | 256.4392196 | -0.714740731 | 0.118617253 | -6.025605164 | 1.68E-09 | 3.91E-08 |
| NR1H3 | 2371.293844 | 0.662968701 | 0.110038608 | 6.024873555 | 1.69E-09 | 3.93E-08 |
| CELF1 | 66.1125182 | 1.099042097 | 0.182517392 | 6.021574638 | 1.73E-09 | 4.00E-08 |
| ERCC2 | 100.9707428 | -0.706209987 | 0.117295409 | -6.02078113 | 1.74E-09 | 4.02E-08 |
| ALAD | 244.4845776 | -0.502034049 | 0.08339297 | -6.020100393 | 1.74E-09 | 4.03E-08 |
| RPS29 | 66.31770055 | -1.045259042 | 0.173849167 | -6.012447819 | 1.83E-09 | 4.22E-08 |
| RNPEPL1 | 1999.914404 | -0.385795568 | 0.064201594 | -6.009127605 | 1.87E-09 | 4.30E-08 |
| EID3 | 300.4431796 | 0.844024079 | 0.140656294 | 6.000613647 | 1.97E-09 | 4.53E-08 |
| XPNPEP1 | 360.8427527 | 0.393733198 | 0.065639717 | 5.998398781 | 1.99E-09 | 4.58E-08 |
| PDCD1LG2 | 1275.584232 | 0.866061665 | 0.144400025 | 5.997655905 | 2.00E-09 | 4.60E-08 |
| RNASE6 | 158.6543354 | -1.011466576 | 0.168725731 | -5.994738144 | 2.04E-09 | 4.68E-08 |
| BTD | 288.9427341 | -0.459718338 | 0.076731176 | -5.99128492 | 2.08E-09 | 4.77E-08 |
| LRRC16B | 20.15682396 | 1.766985711 | 0.294948282 | 5.990832358 | 2.09E-09 | 4.78E-08 |
| PLEKHG3 | 405.2128483 | 0.801519033 | 0.133912589 | 5.985389732 | 2.16E-09 | 4.94E-08 |
| PLK3 | 415.2994876 | 1.095077265 | 0.183082958 | 5.981317316 | 2.21E-09 | 5.05E-08 |
| LOC100500 | 300.1948078 | -0.615117242 | 0.102882884 | -5.978810244 | 2.25E-09 | 5.13E-08 |
| ST3GAL4 | 63.83232645 | 0.996287232 | 0.166725104 | 5.975628195 | 2.29E-09 | 5.22E-08 |
| NPR1 | 59.65604747 | 1.355452838 | 0.227008927 | 5.970923062 | 2.36E-09 | 5.37E-08 |
| RGS14 | 21.60191493 | -1.320181098 | 0.221201713 | -5.968222756 | 2.40E-09 | 5.45E-08 |
| PARP6 | 820.767395 | 0.568915439 | 0.095340669 | 5.967185294 | 2.41E-09 | 5.48E-08 |
| MIR155HG | 23.86342811 | 1.297593005 | 0.217478721 | 5.966528578 | 2.42E-09 | 5.49E-08 |
| GNPDA1 | 727.342357 | -0.289694713 | 0.0486454 | -5.955233499 | 2.60E-09 | 5.88E-08 |
| PIK3IP1 | 85.90590931 | -1.182121146 | 0.198505924 | -5.955092539 | 2.60E-09 | 5.88E-08 |
| KLHDC7B | 38.87496553 | 1.131181617 | 0.190333213 | 5.943164623 | 2.80E-09 | 6.31E-08 |
| ENO2 | 187.974382 | 0.907146131 | 0.15275379 | 5.938616195 | 2.87E-09 | 6.48E-08 |
| MDK | 544.8511944 | 0.867418977 | 0.146069019 | 5.938418579 | 2.88E-09 | 6.48E-08 |
| ATPAF1 | 244.1365264 | -0.391882522 | 0.066010051 | -5.93670986 | 2.91E-09 | 6.54E-08 |
| MFSD2A | 300.5448539 | 0.613485806 | 0.103423055 | 5.931808961 | 3.00E-09 | 6.73E-08 |
| MBNL3 | 139.981749 | -0.610806971 | 0.102996964 | -5.930339591 | 3.02E-09 | 6.78E-08 |
| PIK3R1 | 457.8080971 | -0.511241084 | 0.086206167 | -5.930446785 | 3.02E-09 | 6.78E-08 |
| TLR4 | 1907.832603 | -0.815901984 | 0.137686176 | -5.925809041 | 3.11E-09 | 6.96E-08 |
| DYRK4 | 318.9868607 | -1.125951722 | 0.190068664 | -5.923920837 | 3.14E-09 | 7.03E-08 |
| ABCG2 | 991.51359 | -0.533726245 | 0.090175187 | -5.918770582 | 3.24E-09 | 7.25E-08 |
| CSF3R | 967.9321499 | -0.821351282 | 0.138799418 | -5.917541232 | 3.27E-09 | 7.29E-08 |
| APPL2 | 195.6810004 | -0.563489729 | 0.095248773 | -5.91597886 | 3.30E-09 | 7.35E-08 |
| ATP2C1 | 1043.777228 | 0.819237015 | 0.138544177 | 5.913182602 | 3.36E-09 | 7.47E-08 |
| SYT7 | 15.32946501 | 2.792945582 | 0.472593707 | 5.909823899 | 3.42E-09 | 7.62E-08 |
| ARRB2 | 6134.939798 | -0.354664752 | 0.060021155 | -5.908995826 | 3.44E-09 | 7.64E-08 |
| ATP8B1 | 30.12847516 | -1.593594199 | 0.26987328 | -5.904972128 | 3.53E-09 | 7.82E-08 |
| IGSF10 | 20.78093734 | -1.565465611 | 0.265366526 | -5.899258048 | 3.65E-09 | 8.08E-08 |

| | | | | | | |
|---|---|---|---|---|---|---|
| PLEKHN1 | 16.27567293 | 1.599884508 | 0.27119243 | 5.899443832 | 3.65E-09 | 8.08E-08 |
| MET | 13.57086391 | 1.993754887 | 0.338595091 | 5.888315979 | 3.90E-09 | 8.62E-08 |
| C1R | 144.3703428 | 1.291100501 | 0.219348857 | 5.886059839 | 3.96E-09 | 8.73E-08 |
| LOC65378( | 20.24753381 | -1.086608663 | 0.184669724 | -5.884065015 | 4.00E-09 | 8.83E-08 |
| NDRG2 | 41.80496863 | 1.559341832 | 0.265080185 | 5.882528846 | 4.04E-09 | 8.90E-08 |
| MSRB2 | 428.9653044 | -0.497905702 | 0.08465743 | -5.881417641 | 4.07E-09 | 8.95E-08 |
| NFKBIE | 520.1791877 | 0.600120627 | 0.102048296 | 5.880751054 | 4.08E-09 | 8.97E-08 |
| TP53INP2 | 112.8508105 | 0.940556837 | 0.159944338 | 5.880525989 | 4.09E-09 | 8.98E-08 |
| GTF2I | 191.751597 | 0.567183475 | 0.096465306 | 5.879662844 | 4.11E-09 | 9.01E-08 |
| LILRB5 | 247.8557522 | -1.40764165 | 0.239555131 | -5.876065533 | 4.20E-09 | 9.20E-08 |
| ATP8B2 | 33.89272338 | 1.03987755 | 0.17710921 | 5.871391726 | 4.32E-09 | 9.45E-08 |
| FAM83G | 128.1727025 | 0.952682469 | 0.162483782 | 5.863246532 | 4.54E-09 | 9.91E-08 |
| PIM1 | 3311.803365 | 0.688008495 | 0.117343864 | 5.863182527 | 4.54E-09 | 9.91E-08 |
| PPA1 | 1266.216372 | 0.416990484 | 0.071123826 | 5.862880369 | 4.55E-09 | 9.91E-08 |
| EIF1B | 3330.203038 | 0.5227626 | 0.089291626 | 5.854553501 | 4.78E-09 | 1.04E-07 |
| ITGAE | 111.4007053 | -0.869941107 | 0.148729896 | -5.849134118 | 4.94E-09 | 1.07E-07 |
| ARHGEF35 | 51.06989543 | 1.31294772 | 0.224690966 | 5.843348936 | 5.12E-09 | 1.11E-07 |
| VASN | 20.5587021 | 1.582101994 | 0.270755591 | 5.843284678 | 5.12E-09 | 1.11E-07 |
| NGFRAP1 | 69.29004786 | -0.723473302 | 0.123897084 | -5.839308574 | 5.24E-09 | 1.14E-07 |
| MICALCL | 33.80652713 | 0.985326015 | 0.168836963 | 5.83596148 | 5.35E-09 | 1.16E-07 |
| NRSN2 | 47.3272992 | 1.301042234 | 0.222929931 | 5.836103883 | 5.34E-09 | 1.16E-07 |
| TMEM56 | 19.07831409 | -1.618777668 | 0.27747635 | -5.83393024 | 5.41E-09 | 1.17E-07 |
| H1F0 | 269.5928776 | 0.789156644 | 0.135338282 | 5.830993517 | 5.51E-09 | 1.19E-07 |
| LGALS3 | 9252.918758 | -0.458322918 | 0.078608909 | -5.830419565 | 5.53E-09 | 1.19E-07 |
| RNF125 | 52.88384622 | -1.032574723 | 0.177097496 | -5.830543894 | 5.52E-09 | 1.19E-07 |
| FABP5 | 4839.744092 | -0.653662296 | 0.112138413 | -5.829066742 | 5.57E-09 | 1.20E-07 |
| CD164 | 13290.66973 | 0.294200905 | 0.050473825 | 5.828781677 | 5.58E-09 | 1.20E-07 |
| LOC73010: | 19.42552487 | 1.533814234 | 0.263207502 | 5.827395576 | 5.63E-09 | 1.21E-07 |
| EPT1 | 268.5290685 | 0.607828945 | 0.104372558 | 5.823647096 | 5.76E-09 | 1.23E-07 |
| PCED1B | 115.3175561 | -0.698270853 | 0.119903261 | -5.82361854 | 5.76E-09 | 1.23E-07 |
| BRI3BP | 151.759872 | -0.675968845 | 0.116134861 | -5.820550707 | 5.87E-09 | 1.25E-07 |
| C11orf96 | 12.30173488 | 3.964985624 | 0.683600968 | 5.800146295 | 6.63E-09 | 1.41E-07 |
| CCL18 | 32426.69016 | 0.740129448 | 0.127634686 | 5.798811202 | 6.68E-09 | 1.42E-07 |
| MREG | 1731.019577 | 0.650238703 | 0.112182382 | 5.796264031 | 6.78E-09 | 1.44E-07 |
| ITGA4 | 218.7079474 | -0.738321445 | 0.127386337 | -5.795923333 | 6.79E-09 | 1.45E-07 |
| HPS5 | 1327.236167 | 0.343696185 | 0.059311223 | 5.794791781 | 6.84E-09 | 1.45E-07 |
| TPM1 | 74.5257404 | -0.453013672 | 0.078181398 | -5.794392061 | 6.86E-09 | 1.46E-07 |
| MOV10 | 255.6421961 | 0.612750408 | 0.105829347 | 5.78998572 | 7.04E-09 | 1.49E-07 |
| TM4SF19 | 6415.490551 | 0.608691799 | 0.105204491 | 5.785796748 | 7.22E-09 | 1.53E-07 |
| RHOBTB1 | 55.61745946 | -0.894760113 | 0.154802681 | -5.780003979 | 7.47E-09 | 1.58E-07 |
| BCAT1 | 1533.443007 | 0.759896233 | 0.131569538 | 5.775624377 | 7.67E-09 | 1.62E-07 |
| TYMP | 8218.49514 | 0.774076111 | 0.134052117 | 5.774441541 | 7.72E-09 | 1.63E-07 |
| MGST2 | 833.2352571 | -0.433812605 | 0.07520022 | -5.768767753 | 7.99E-09 | 1.68E-07 |
| FABP4 | 6653.34902 | -0.941323797 | 0.163212406 | -5.767476985 | 8.05E-09 | 1.69E-07 |
| GCLM | 336.9625198 | 0.597403138 | 0.103609209 | 5.765926998 | 8.12E-09 | 1.71E-07 |
| PECR | 83.56117218 | -0.658342807 | 0.114234362 | -5.763089096 | 8.26E-09 | 1.74E-07 |
| AP5B1 | 604.0981198 | 0.513721228 | 0.089156947 | 5.761987661 | 8.31E-09 | 1.74E-07 |

| | | | | | | |
|---|---|---|---|---|---|---|
| CCRL2 | 1878.936631 | 0.665819042 | 0.115603188 | 5.759521467 | 8.44E-09 | 1.77E-07 |
| LINC00263 | 63.24816366 | -0.734242415 | 0.127487671 | -5.759320938 | 8.45E-09 | 1.77E-07 |
| TNS3 | 2548.628956 | 0.591448783 | 0.102708858 | 5.758498303 | 8.49E-09 | 1.77E-07 |
| PRR11 | 42.25480946 | -0.871548666 | 0.151371268 | -5.75768886 | 8.53E-09 | 1.78E-07 |
| RPL21 | 7562.681858 | -0.43767423 | 0.076018616 | -5.757461147 | 8.54E-09 | 1.78E-07 |
| C5orf4 | 17.15993296 | -1.734516844 | 0.3015151 | -5.752669905 | 8.78E-09 | 1.83E-07 |
| PABPC1 | 11295.66516 | -0.472779302 | 0.082197265 | -5.751764382 | 8.83E-09 | 1.84E-07 |
| ARID5A | 41.75843467 | 0.702119754 | 0.122098102 | 5.750455917 | 8.90E-09 | 1.85E-07 |
| NRP1 | 2454.435142 | 0.534584487 | 0.0929779 | 5.749586608 | 8.95E-09 | 1.86E-07 |
| LOC28597[illegible] | 18.67261522 | 1.834234907 | 0.31909786 | 5.748189307 | 9.02E-09 | 1.87E-07 |
| HCK | 3461.339419 | 0.51009486 | 0.088789404 | 5.744996986 | 9.19E-09 | 1.91E-07 |
| SBNO2 | 558.0229018 | 0.548042269 | 0.09547717 | 5.740034709 | 9.47E-09 | 1.96E-07 |
| DERL1 | 886.945755 | 0.435423322 | 0.075910698 | 5.735994148 | 9.69E-09 | 2.00E-07 |
| NMB | 206.7490699 | -0.732516398 | 0.127737985 | -5.734522896 | 9.78E-09 | 2.02E-07 |
| GGA2 | 1730.629456 | -0.374386365 | 0.065308223 | -5.732606811 | 9.89E-09 | 2.04E-07 |
| CCDC106 | 233.4903339 | -0.5602669 | 0.097754387 | -5.731373494 | 9.96E-09 | 2.05E-07 |
| ZNF710 | 285.1402249 | -0.741643118 | 0.129530338 | -5.725632523 | 1.03E-08 | 2.12E-07 |
| HS3ST1 | 20.45249791 | -1.58605929 | 0.277045387 | -5.724907777 | 1.03E-08 | 2.13E-07 |
| C15orf52 | 91.28861739 | -0.777819816 | 0.135883192 | -5.724179766 | 1.04E-08 | 2.14E-07 |
| EYA4 | 7.265600571 | 2.96630243 | 0.518629876 | 5.719497791 | 1.07E-08 | 2.19E-07 |
| POPDC3 | 18.24589516 | 1.764375267 | 0.308538057 | 5.718501266 | 1.07E-08 | 2.20E-07 |
| ARID5B | 279.5066235 | 0.866447327 | 0.151587845 | 5.715810051 | 1.09E-08 | 2.24E-07 |
| SNX5 | 1473.128369 | -0.397871474 | 0.069612318 | -5.715532601 | 1.09E-08 | 2.24E-07 |
| UBE2J1 | 1616.683561 | 0.344045151 | 0.06019694 | 5.715326275 | 1.09E-08 | 2.24E-07 |
| RPL4 | 17690.55381 | -0.399097302 | 0.069870563 | -5.711952014 | 1.12E-08 | 2.28E-07 |
| TPD52L1 | 31.68273333 | -1.838125654 | 0.321842283 | -5.71126217 | 1.12E-08 | 2.29E-07 |
| FOSL2 | 597.5651183 | 1.134803808 | 0.198779431 | 5.708859311 | 1.14E-08 | 2.32E-07 |
| MIF4GD | 90.4597314 | -0.564899189 | 0.09895767 | -5.708493205 | 1.14E-08 | 2.32E-07 |
| METTL7B | 53.29602264 | 0.836579273 | 0.146580624 | 5.707297799 | 1.15E-08 | 2.33E-07 |
| ACAT1 | 849.828969 | -0.379101288 | 0.06642899 | -5.706865133 | 1.15E-08 | 2.34E-07 |
| ACOT7 | 619.4281681 | 0.719516637 | 0.126100423 | 5.705901846 | 1.16E-08 | 2.35E-07 |
| CYLD | 856.5264216 | 0.53870004 | 0.094425979 | 5.70499818 | 1.16E-08 | 2.36E-07 |
| DUSP14 | 190.2147511 | 0.745188373 | 0.130650581 | 5.703674373 | 1.17E-08 | 2.37E-07 |
| ASB6 | 221.7529207 | 0.81943731 | 0.143821398 | 5.697603567 | 1.22E-08 | 2.46E-07 |
| AFF4 | 911.220255 | 0.435795223 | 0.07651839 | 5.695300458 | 1.23E-08 | 2.49E-07 |
| PDXP | 105.9766708 | -0.732505259 | 0.128636154 | -5.69439644 | 1.24E-08 | 2.50E-07 |
| GLUL | 5422.001745 | -0.567537434 | 0.099690264 | -5.693007634 | 1.25E-08 | 2.51E-07 |
| C1orf21 | 49.12575002 | -1.000241657 | 0.175705008 | -5.692732755 | 1.25E-08 | 2.52E-07 |
| SNAI1 | 5.628004995 | 2.76542537 | 0.48594998 | 5.69076136 | 1.26E-08 | 2.54E-07 |
| RRAGD | 1797.254732 | -0.312992953 | 0.055006736 | -5.690084033 | 1.27E-08 | 2.55E-07 |
| PDE4A | 660.0712334 | 0.563837734 | 0.099122478 | 5.688293361 | 1.28E-08 | 2.57E-07 |
| CNP | 919.031252 | 0.656149723 | 0.115359633 | 5.687862441 | 1.29E-08 | 2.57E-07 |
| HMG20B | 1141.892346 | 0.587498215 | 0.103287189 | 5.688006637 | 1.29E-08 | 2.57E-07 |
| PPP1R18 | 1222.183221 | 0.59923264 | 0.105401178 | 5.685255642 | 1.31E-08 | 2.61E-07 |
| ZZZ3 | 316.5964554 | -0.419751106 | 0.073858881 | -5.683150061 | 1.32E-08 | 2.64E-07 |
| FICD | 350.2259884 | 0.443802521 | 0.078094422 | 5.682896544 | 1.32E-08 | 2.64E-07 |
| NFAT5 | 657.9144795 | 0.404041585 | 0.071105731 | 5.682264657 | 1.33E-08 | 2.65E-07 |

| | | | | | | |
|---|---|---|---|---|---|---|
| RPL12 | 2422.002879 | -0.402813247 | 0.070965775 | -5.676162139 | 1.38E-08 | 2.74E-07 |
| PDLIM5 | 2118.797452 | 0.399415257 | 0.070409137 | 5.672775957 | 1.41E-08 | 2.79E-07 |
| RNF31 | 777.6709771 | 0.333946966 | 0.058895839 | 5.670128333 | 1.43E-08 | 2.83E-07 |
| MYD88 | 1630.626407 | 0.549302495 | 0.096888642 | 5.669420907 | 1.43E-08 | 2.84E-07 |
| RGS16 | 17.25347752 | 1.87546156 | 0.330815801 | 5.669201877 | 1.43E-08 | 2.84E-07 |
| PDLIM7 | 1406.923906 | 0.739436441 | 0.13051337 | 5.665599159 | 1.47E-08 | 2.90E-07 |
| PKM | 39.51359901 | 0.74827779 | 0.132098496 | 5.664544373 | 1.47E-08 | 2.92E-07 |
| C17orf58 | 123.4353645 | 0.639600792 | 0.112928923 | 5.663746481 | 1.48E-08 | 2.93E-07 |
| ACOT9 | 1143.365508 | 0.479287277 | 0.084667462 | 5.66082015 | 1.51E-08 | 2.97E-07 |
| KARS | 348.222041 | 0.525352866 | 0.09283487 | 5.659003648 | 1.52E-08 | 3.00E-07 |
| SRXN1 | 1173.04578 | 0.661868816 | 0.116963675 | 5.658755303 | 1.52E-08 | 3.00E-07 |
| MEMO1 | 996.3074281 | -0.251528907 | 0.044501689 | -5.652120473 | 1.58E-08 | 3.12E-07 |
| SAMSN1 | 338.6653631 | -1.005415821 | 0.177945643 | -5.650128892 | 1.60E-08 | 3.15E-07 |
| LRP8 | 114.3537608 | 0.596205077 | 0.105534488 | 5.649386181 | 1.61E-08 | 3.16E-07 |
| ZNF33B | 41.37795953 | -1.023844982 | 0.181390055 | -5.644438354 | 1.66E-08 | 3.25E-07 |
| GPR68 | 48.43904696 | 1.089291346 | 0.193005572 | 5.643833677 | 1.66E-08 | 3.26E-07 |
| PDGFA | 67.36218458 | 1.223161623 | 0.2167355 | 5.643568415 | 1.67E-08 | 3.26E-07 |
| C10orf10 | 13.98502749 | 1.887056105 | 0.334964236 | 5.63360473 | 1.76E-08 | 3.45E-07 |
| FAIM3 | 256.5210545 | -0.754747167 | 0.133976358 | -5.633435463 | 1.77E-08 | 3.45E-07 |
| TRAF3IP2 | 528.2163923 | 0.477281408 | 0.084722494 | 5.633467435 | 1.77E-08 | 3.45E-07 |
| MAPK13 | 1877.548903 | 0.507989545 | 0.090225824 | 5.63020125 | 1.80E-08 | 3.51E-07 |
| ITGAX | 4145.446173 | 0.550816747 | 0.097855002 | 5.628907414 | 1.81E-08 | 3.53E-07 |
| NUP98 | 725.3583068 | 0.480097793 | 0.085290102 | 5.628997736 | 1.81E-08 | 3.53E-07 |
| SLC3A2 | 3612.322999 | 0.553684401 | 0.098436829 | 5.624768735 | 1.86E-08 | 3.61E-07 |
| DRAM1 | 4551.891546 | 0.433531563 | 0.077135254 | 5.620407567 | 1.91E-08 | 3.70E-07 |
| ZNF238 | 16.10023302 | -1.069043465 | 0.190329802 | -5.616794924 | 1.95E-08 | 3.77E-07 |
| C8B | 95.63127027 | -1.095915662 | 0.195158789 | -5.615507592 | 1.96E-08 | 3.79E-07 |
| CBLN3 | 23.91829796 | 1.155314429 | 0.20573966 | 5.615419177 | 1.96E-08 | 3.79E-07 |
| SNHG6 | 359.9025303 | -0.473710493 | 0.084392673 | -5.613170856 | 1.99E-08 | 3.84E-07 |
| CXCL1 | 62.57967252 | 1.870634936 | 0.333316227 | 5.612192816 | 2.00E-08 | 3.86E-07 |
| UBE2Z | 1739.158121 | 0.345156125 | 0.061508532 | 5.611516212 | 2.01E-08 | 3.87E-07 |
| CEMP1 | 46.49644314 | -0.829874389 | 0.147903991 | -5.610899209 | 2.01E-08 | 3.88E-07 |
| DUSP4 | 31.31831582 | 1.907263001 | 0.340014573 | 5.609356636 | 2.03E-08 | 3.91E-07 |
| IL23A | 64.81167508 | 2.674459949 | 0.47682879 | 5.608847465 | 2.04E-08 | 3.92E-07 |
| APOL4 | 142.5183879 | 1.263676711 | 0.225380281 | 5.606864557 | 2.06E-08 | 3.96E-07 |
| ARHGAP31 | 1096.32737 | 0.379343178 | 0.067672676 | 5.605558999 | 2.08E-08 | 3.98E-07 |
| OR6K3 | 76.66454864 | -0.796975986 | 0.142180687 | -5.605374432 | 2.08E-08 | 3.98E-07 |
| SERPINE2 | 56.24691384 | 2.293116925 | 0.409453374 | 5.600434796 | 2.14E-08 | 4.09E-07 |
| LOC10013[illegible] | 71.03711437 | 1.108035623 | 0.197874641 | 5.599684823 | 2.15E-08 | 4.11E-07 |
| FAM100B | 1774.365536 | 0.529636439 | 0.094622046 | 5.597389427 | 2.18E-08 | 4.16E-07 |
| SLC50A1 | 761.7602399 | 0.341037966 | 0.061019157 | 5.589031084 | 2.28E-08 | 4.36E-07 |
| LOC10012[illegible] | 36.98761349 | -0.878456066 | 0.157246617 | -5.586486268 | 2.32E-08 | 4.42E-07 |
| CMKLR1 | 26.02899383 | 2.240253979 | 0.401094064 | 5.585358097 | 2.33E-08 | 4.44E-07 |
| HCG4 | 21.75234317 | 1.098212871 | 0.196652826 | 5.58452625 | 2.34E-08 | 4.46E-07 |
| ABCC5 | 130.8007902 | -0.844306364 | 0.151192294 | -5.584321415 | 2.35E-08 | 4.46E-07 |
| TNFRSF9 | 148.5843405 | 1.415691394 | 0.253612311 | 5.582108332 | 2.38E-08 | 4.51E-07 |
| ZNF117 | 211.8759334 | -0.651947359 | 0.116789509 | -5.5822425 | 2.37E-08 | 4.51E-07 |

| | | | | | | |
|---|---|---|---|---|---|---|
| TBC1D7 | 379.7102285 | 0.515665824 | 0.092408358 | 5.580294193 | 2.40E-08 | 4.55E-07 |
| STK17A | 321.9714398 | 0.961586628 | 0.172346665 | 5.579374724 | 2.41E-08 | 4.57E-07 |
| PTGIR | 12.76906731 | 1.932449015 | 0.34647157 | 5.577511062 | 2.44E-08 | 4.62E-07 |
| TACC1 | 1748.607731 | -0.372827295 | 0.066900921 | -5.572827525 | 2.51E-08 | 4.74E-07 |
| APBA3 | 217.4148673 | 0.544695271 | 0.097748289 | 5.572427656 | 2.51E-08 | 4.74E-07 |
| ANP32A | 851.7777801 | -0.379977452 | 0.068200282 | -5.571493827 | 2.53E-08 | 4.76E-07 |
| IL4I1 | 833.9659885 | 0.926723166 | 0.166411498 | 5.568864986 | 2.56E-08 | 4.83E-07 |
| TFCP2L1 | 315.067797 | -0.47724835 | 0.085705003 | -5.568500483 | 2.57E-08 | 4.84E-07 |
| TNFRSF12A | 241.0977073 | 0.696086209 | 0.125009884 | 5.568249393 | 2.57E-08 | 4.84E-07 |
| LOC10028[illegible] | 78.56818108 | 1.199634485 | 0.215520678 | 5.566215261 | 2.60E-08 | 4.89E-07 |
| CFD | 1381.340594 | -0.918725252 | 0.165218493 | -5.560668407 | 2.69E-08 | 5.04E-07 |
| IDH2 | 633.7299326 | -0.429021841 | 0.077170386 | -5.559410355 | 2.71E-08 | 5.07E-07 |
| ADAMTS7 | 17.35307895 | -1.488755377 | 0.268068549 | -5.553636862 | 2.80E-08 | 5.24E-07 |
| EXOSC7 | 339.8344228 | 0.434286547 | 0.078235364 | 5.551026066 | 2.84E-08 | 5.31E-07 |
| RUFY3 | 903.9895088 | 0.710205936 | 0.12795412 | 5.550473354 | 2.85E-08 | 5.32E-07 |
| MYCL1 | 68.57439178 | 0.935636185 | 0.168657809 | 5.547541428 | 2.90E-08 | 5.40E-07 |
| SIGLEC1 | 169.9802881 | 1.773613774 | 0.319704048 | 5.547673815 | 2.89E-08 | 5.40E-07 |
| SYNGR2 | 4014.278665 | -0.347736404 | 0.062695877 | -5.54639988 | 2.92E-08 | 5.43E-07 |
| ZNF486 | 78.84266238 | -0.526594029 | 0.094978567 | -5.544345921 | 2.95E-08 | 5.49E-07 |
| SUSD3 | 52.57105815 | 0.64630022 | 0.116601407 | 5.542816643 | 2.98E-08 | 5.54E-07 |
| NAA50 | 1809.654543 | 0.426933021 | 0.077027293 | 5.542620069 | 2.98E-08 | 5.54E-07 |
| ENPP2 | 22.0282952 | 1.960026495 | 0.353905691 | 5.538273464 | 3.05E-08 | 5.67E-07 |
| UGDH | 359.8020412 | 0.598397259 | 0.108101298 | 5.535523334 | 3.10E-08 | 5.75E-07 |
| USE1 | 348.3007492 | -0.408193228 | 0.073738451 | -5.535690314 | 3.10E-08 | 5.75E-07 |
| ZNF581 | 112.0815188 | -0.749562192 | 0.135434924 | -5.534482332 | 3.12E-08 | 5.78E-07 |
| ABHD15 | 294.6102848 | -0.35194113 | 0.063610557 | -5.532747209 | 3.15E-08 | 5.83E-07 |
| RASAL2 | 732.1396258 | 0.387328427 | 0.070025448 | 5.531252366 | 3.18E-08 | 5.87E-07 |
| HMG20A | 370.6153198 | -0.455348217 | 0.082333582 | -5.530528436 | 3.19E-08 | 5.89E-07 |
| RCBTB1 | 153.9564403 | -0.461657168 | 0.083486637 | -5.529713335 | 3.21E-08 | 5.91E-07 |
| S100A9 | 420.1008451 | -1.405906305 | 0.254290528 | -5.52874035 | 3.23E-08 | 5.94E-07 |
| C7orf29 | 122.8669917 | 1.007879185 | 0.182534014 | 5.521596578 | 3.36E-08 | 6.18E-07 |
| CORO7 | 143.6846449 | -0.437724225 | 0.07929483 | -5.520211437 | 3.39E-08 | 6.22E-07 |
| DNTTIP1 | 390.0905237 | 0.364801655 | 0.066084704 | 5.520213214 | 3.39E-08 | 6.22E-07 |
| CCDC125 | 369.3746708 | -0.405085845 | 0.073384916 | -5.520015091 | 3.39E-08 | 6.22E-07 |
| TAGLN3 | 22.88625382 | 1.617992282 | 0.293339908 | 5.515759151 | 3.47E-08 | 6.36E-07 |
| SIGLEC15 | 39.17472141 | 1.131819356 | 0.205299792 | 5.513007804 | 3.53E-08 | 6.46E-07 |
| CALM2 | 3232.785435 | -0.584126054 | 0.105979753 | -5.511675948 | 3.55E-08 | 6.50E-07 |
| CUTA | 504.3156635 | -0.405756363 | 0.07368562 | -5.506588191 | 3.66E-08 | 6.68E-07 |
| TNFSF9 | 24.80378101 | 0.953675142 | 0.173210287 | 5.505880495 | 3.67E-08 | 6.70E-07 |
| TANC2 | 548.7061072 | -0.512958969 | 0.093202046 | -5.503730791 | 3.72E-08 | 6.78E-07 |
| YWHAZ | 308.0341729 | 0.562291237 | 0.1021707 | 5.503449013 | 3.72E-08 | 6.78E-07 |
| ZSWIM5 | 7.185776399 | 1.991977778 | 0.362098051 | 5.501210997 | 3.77E-08 | 6.86E-07 |
| SYT12 | 4.219885329 | 3.621814979 | 0.658462599 | 5.500411088 | 3.79E-08 | 6.89E-07 |
| SLC38A2 | 2972.175078 | 0.448560152 | 0.08161079 | 5.496333904 | 3.88E-08 | 7.04E-07 |
| CYSLTR2 | 54.90396104 | -1.043476043 | 0.190077058 | -5.489752704 | 4.02E-08 | 7.30E-07 |
| AIFM2 | 94.18425342 | 0.937517062 | 0.170791349 | 5.489253797 | 4.04E-08 | 7.32E-07 |
| C20orf160 | 6.839434511 | 2.848817898 | 0.519088968 | 5.488111044 | 4.06E-08 | 7.36E-07 |

| | | | | | | |
|---|---|---|---|---|---|---|
| RPS23 | 6931.047679 | -0.506579576 | 0.092340837 | -5.485975568 | 4.11E-08 | 7.44E-07 |
| RGS10 | 656.8219283 | -0.425523712 | 0.077581546 | -5.484857356 | 4.14E-08 | 7.48E-07 |
| RUSC2 | 314.8630963 | 0.662446625 | 0.120798793 | 5.483884468 | 4.16E-08 | 7.51E-07 |
| RPL11 | 18338.05697 | -0.376310877 | 0.068655205 | -5.481170415 | 4.23E-08 | 7.62E-07 |
| GAS2L3 | 309.8011401 | -0.485605106 | 0.088659469 | -5.477193927 | 4.32E-08 | 7.79E-07 |
| EIF2D | 93.58606728 | -0.465527924 | 0.08506433 | -5.47265725 | 4.43E-08 | 7.98E-07 |
| ZNF295 | 208.3536776 | 0.451705105 | 0.082613491 | 5.467691755 | 4.56E-08 | 8.20E-07 |
| PRSS36 | 20.8626572 | -1.146417186 | 0.209701344 | -5.466904328 | 4.58E-08 | 8.23E-07 |
| YPEL2 | 97.35813278 | -0.603905568 | 0.110476634 | -5.466364676 | 4.59E-08 | 8.25E-07 |
| GADD45G | 203.1439537 | 0.897418896 | 0.16420051 | 5.465384335 | 4.62E-08 | 8.28E-07 |
| FAM104A | 338.7338492 | 0.331088105 | 0.060626648 | 5.461098621 | 4.73E-08 | 8.48E-07 |
| ECM1 | 568.0609175 | 0.797791032 | 0.146096123 | 5.460726912 | 4.74E-08 | 8.49E-07 |
| MTHFD1 | 1214.812253 | 0.446561111 | 0.081804898 | 5.458855449 | 4.79E-08 | 8.57E-07 |
| STX6 | 1408.234696 | -0.358272541 | 0.065638399 | -5.458276674 | 4.81E-08 | 8.59E-07 |
| HEXB | 6747.614524 | -0.309534612 | 0.056734521 | -5.455842522 | 4.87E-08 | 8.70E-07 |
| RBMS2 | 469.3488715 | -0.555018498 | 0.101760745 | -5.45415129 | 4.92E-08 | 8.77E-07 |
| THBS1 | 467.6790905 | -1.414285954 | 0.259549985 | -5.448992634 | 5.07E-08 | 9.02E-07 |
| PRR5L | 36.33395593 | -1.431279591 | 0.262784541 | -5.446589774 | 5.13E-08 | 9.14E-07 |
| ATAD1 | 626.9420818 | -0.501348221 | 0.092192041 | -5.438085745 | 5.39E-08 | 9.58E-07 |
| LRMP | 104.270351 | -0.91452545 | 0.168212687 | -5.436721022 | 5.43E-08 | 9.64E-07 |
| NMD3 | 391.1371916 | -0.386986318 | 0.071185698 | -5.436293081 | 5.44E-08 | 9.65E-07 |
| ACVR1B | 812.9930105 | 0.520745929 | 0.095822545 | 5.43448234 | 5.50E-08 | 9.74E-07 |
| DHRS4 | 785.552167 | 0.463513354 | 0.085354689 | 5.430438076 | 5.62E-08 | 9.95E-07 |
| MAPK3 | 318.4651865 | -0.505267423 | 0.0930447 | -5.43037298 | 5.62E-08 | 9.95E-07 |
| AKIRIN2 | 447.2897627 | 0.387595419 | 0.071380897 | 5.429960084 | 5.64E-08 | 9.97E-07 |
| IPW | 18.79931431 | -1.020212317 | 0.18816249 | -5.421974997 | 5.89E-08 | 1.04E-06 |
| AFMID | 32.78829286 | -0.963506103 | 0.177738351 | -5.420924049 | 5.93E-08 | 1.05E-06 |
| ADORA2A | 23.87031554 | 2.425316866 | 0.447435638 | 5.420482098 | 5.94E-08 | 1.05E-06 |
| SLC7A2 | 45.05771328 | 4.383281162 | 0.808671284 | 5.420349715 | 5.95E-08 | 1.05E-06 |
| N4BP1 | 365.0154737 | 0.644597365 | 0.118926467 | 5.420133826 | 5.96E-08 | 1.05E-06 |
| POLR3D | 253.31694 | 0.482045982 | 0.08895741 | 5.41884012 | 6.00E-08 | 1.05E-06 |
| FMR1 | 208.2271643 | 0.32706688 | 0.060389363 | 5.415968328 | 6.10E-08 | 1.07E-06 |
| ARL3 | 172.1964839 | -0.449479564 | 0.082996826 | -5.415623507 | 6.11E-08 | 1.07E-06 |
| SLC2A9 | 214.9389045 | -0.587721817 | 0.108620063 | -5.410803525 | 6.27E-08 | 1.10E-06 |
| LOC64632! | 43.38519836 | 1.001707544 | 0.185260026 | 5.407035525 | 6.41E-08 | 1.12E-06 |
| ARRDC1 | 280.5962241 | -0.388092438 | 0.071823511 | -5.40341781 | 6.54E-08 | 1.14E-06 |
| CCNG2 | 260.7124132 | -0.57838381 | 0.10712481 | -5.399158329 | 6.70E-08 | 1.17E-06 |
| PROCR | 201.5389122 | 0.960884591 | 0.178004269 | 5.398098575 | 6.74E-08 | 1.18E-06 |
| UBTD2 | 530.8945972 | 0.363387996 | 0.067327902 | 5.397286795 | 6.77E-08 | 1.18E-06 |
| OSBPL11 | 1172.279549 | -0.415307463 | 0.076963214 | -5.396181356 | 6.81E-08 | 1.19E-06 |
| SPECC1 | 142.494686 | 0.552985033 | 0.102475652 | 5.396257776 | 6.80E-08 | 1.19E-06 |
| ITGAM | 1066.250719 | 0.552188283 | 0.102384457 | 5.393282329 | 6.92E-08 | 1.20E-06 |
| CD84 | 3412.992712 | -0.602607202 | 0.111784231 | -5.390806882 | 7.01E-08 | 1.22E-06 |
| SGMS1 | 1002.319619 | 0.425564904 | 0.078990119 | 5.387571369 | 7.14E-08 | 1.24E-06 |
| CDC42EP3 | 1045.522913 | -0.544968245 | 0.101162807 | -5.387041531 | 7.16E-08 | 1.24E-06 |
| H2AFY | 2884.382838 | -0.428987199 | 0.07964009 | -5.386573489 | 7.18E-08 | 1.24E-06 |
| RPS27 | 1643.690106 | -0.611618461 | 0.113651281 | -5.381536007 | 7.39E-08 | 1.28E-06 |

| | | | | | | |
|---|---|---|---|---|---|---|
| ADAR | 4938.686473 | 0.573979044 | 0.106699613 | 5.379392003 | 7.47E-08 | 1.29E-06 |
| PBX4 | 41.31608767 | 0.975572643 | 0.181423189 | 5.377331577 | 7.56E-08 | 1.31E-06 |
| RPL3 | 4957.370278 | -0.36534412 | 0.067960701 | -5.375814487 | 7.62E-08 | 1.32E-06 |
| NMI | 241.054215 | 0.632816007 | 0.117946089 | 5.365298751 | 8.08E-08 | 1.39E-06 |
| MTFP1 | 377.2661726 | 0.704421273 | 0.131320582 | 5.364134544 | 8.13E-08 | 1.40E-06 |
| FAM225A | 84.8615772 | 0.657710681 | 0.122637675 | 5.363039367 | 8.18E-08 | 1.41E-06 |
| GTF2A1 | 768.7084006 | 0.306416991 | 0.05716763 | 5.359973707 | 8.32E-08 | 1.43E-06 |
| RAI14 | 55.87795849 | 1.227655103 | 0.229059993 | 5.359535231 | 8.34E-08 | 1.43E-06 |
| GPR85 | 60.6612067 | -0.881770502 | 0.164628642 | -5.356118409 | 8.50E-08 | 1.46E-06 |
| NPPC | 6.118630574 | 2.089221333 | 0.390061215 | 5.356137072 | 8.50E-08 | 1.46E-06 |
| PLCB2 | 2198.188484 | -0.507209728 | 0.094727216 | -5.354424533 | 8.58E-08 | 1.47E-06 |
| SIRPD | 83.85684061 | -0.690313812 | 0.129001875 | -5.351192064 | 8.74E-08 | 1.50E-06 |
| RPLP0 | 4631.509061 | -0.377677858 | 0.070581506 | -5.350946398 | 8.75E-08 | 1.50E-06 |
| C1orf162 | 212.4172472 | -0.699628253 | 0.130842891 | -5.347086463 | 8.94E-08 | 1.53E-06 |
| ERV3-1 | 236.612832 | -0.754509349 | 0.141159152 | -5.345096913 | 9.04E-08 | 1.54E-06 |
| TNFSF15 | 84.41441699 | 1.057112588 | 0.197905932 | 5.341490144 | 9.22E-08 | 1.57E-06 |
| EGR3 | 17.26473764 | 1.692935525 | 0.317029751 | 5.33998945 | 9.30E-08 | 1.58E-06 |
| KIAA0141 | 268.5855098 | -0.424658448 | 0.079527347 | -5.339778853 | 9.31E-08 | 1.59E-06 |
| HLA-DMA | 3522.3214 | -0.433901878 | 0.081329574 | -5.335105744 | 9.55E-08 | 1.62E-06 |
| HBEGF | 3748.358605 | 0.638932531 | 0.119874019 | 5.330033467 | 9.82E-08 | 1.67E-06 |
| ACOT11 | 58.30032708 | -0.615404665 | 0.115470986 | -5.329517716 | 9.85E-08 | 1.67E-06 |
| RAP2B | 984.5545249 | -0.556731623 | 0.10446119 | -5.329554674 | 9.85E-08 | 1.67E-06 |
| LAD1 | 33.73231792 | 2.544660223 | 0.477533193 | 5.328760934 | 9.89E-08 | 1.67E-06 |
| RBM47 | 4167.022007 | 0.296896793 | 0.05571564 | 5.328787289 | 9.89E-08 | 1.67E-06 |
| SNX21 | 80.21179842 | -0.657563995 | 0.123402518 | -5.328610856 | 9.90E-08 | 1.67E-06 |
| SHC4 | 17.97557575 | 1.289106866 | 0.242047635 | 5.325839542 | 1.00E-07 | 1.70E-06 |
| NARS2 | 150.2185893 | -0.46412034 | 0.087243819 | -5.319807726 | 1.04E-07 | 1.75E-06 |
| SAMM50 | 263.7359246 | -0.488491543 | 0.091840586 | -5.318907052 | 1.04E-07 | 1.76E-06 |
| BAMBI | 51.46914803 | 0.930266655 | 0.174930765 | 5.317913389 | 1.05E-07 | 1.77E-06 |
| KLF6 | 1527.644219 | 0.49304952 | 0.092791472 | 5.313521901 | 1.08E-07 | 1.81E-06 |
| PPARD | 897.8943707 | 0.524344184 | 0.098700816 | 5.312460475 | 1.08E-07 | 1.82E-06 |
| PTPN22 | 685.2484261 | -0.607823703 | 0.114498446 | -5.308576018 | 1.10E-07 | 1.86E-06 |
| TEN1 | 144.1437716 | 0.45225384 | 0.085216832 | 5.307095192 | 1.11E-07 | 1.87E-06 |
| CKS2 | 86.62720267 | 0.446916666 | 0.084229662 | 5.305929739 | 1.12E-07 | 1.88E-06 |
| SARDH | 87.87904143 | 1.061070491 | 0.200037401 | 5.304360513 | 1.13E-07 | 1.90E-06 |
| SLC37A2 | 1220.993446 | -0.455944835 | 0.086059175 | -5.298038662 | 1.17E-07 | 1.96E-06 |
| TOMM34 | 275.1484616 | 0.458517029 | 0.086545575 | 5.297983487 | 1.17E-07 | 1.96E-06 |
| UPP1 | 51.33811855 | 0.838755815 | 0.158468002 | 5.292903316 | 1.20E-07 | 2.01E-06 |
| PCYOX1 | 397.6967742 | -0.564810197 | 0.10690132 | -5.283472603 | 1.27E-07 | 2.12E-06 |
| SLC44A2 | 582.894347 | 0.598412605 | 0.113292868 | 5.281997128 | 1.28E-07 | 2.13E-06 |
| FCRLA | 18.89494414 | 1.625734493 | 0.307818537 | 5.281470401 | 1.28E-07 | 2.14E-06 |
| SGSH | 224.549668 | -0.538584575 | 0.102025093 | -5.27894228 | 1.30E-07 | 2.17E-06 |
| XPO6 | 1321.457403 | 0.285999994 | 0.054196102 | 5.277132229 | 1.31E-07 | 2.19E-06 |
| ZFAND2A | 315.4510361 | 0.564657552 | 0.107083564 | 5.273055287 | 1.34E-07 | 2.23E-06 |
| IL27RA | 460.3432883 | 0.397716641 | 0.075447534 | 5.271433273 | 1.35E-07 | 2.25E-06 |
| TSPYL2 | 72.81276311 | -0.602386478 | 0.114279281 | -5.271178417 | 1.36E-07 | 2.25E-06 |
| CC2D2A | 5.887236086 | -2.248805031 | 0.426642301 | -5.270937801 | 1.36E-07 | 2.25E-06 |

| | | | | | | |
|---|---|---|---|---|---|---|
| PSME1 | 1210.54154 | 0.407644886 | 0.077356019 | 5.269724183 | 1.37E-07 | 2.27E-06 |
| PPARGC1B | 267.882652 | 0.614928981 | 0.116701132 | 5.269263212 | 1.37E-07 | 2.27E-06 |
| RRAD | 12.37020387 | 1.786406549 | 0.339087674 | 5.268273334 | 1.38E-07 | 2.28E-06 |
| IFT172 | 81.5681947 | -0.558052107 | 0.105949629 | -5.267145473 | 1.39E-07 | 2.29E-06 |
| TBX20 | 25.87754979 | 1.191372291 | 0.226188486 | 5.267165936 | 1.39E-07 | 2.29E-06 |
| AIM2 | 58.38087575 | 1.678817123 | 0.318865982 | 5.264961519 | 1.40E-07 | 2.31E-06 |
| RBM19 | 349.0975086 | 0.338502782 | 0.064317379 | 5.263006442 | 1.42E-07 | 2.34E-06 |
| WNT5A | 33.73420838 | 2.295653107 | 0.436269195 | 5.262010552 | 1.42E-07 | 2.35E-06 |
| SPP1 | 5206.372858 | 0.745986975 | 0.141868129 | 5.258312629 | 1.45E-07 | 2.39E-06 |
| TPD52 | 135.0022036 | 0.87219032 | 0.16589108 | 5.257608313 | 1.46E-07 | 2.40E-06 |
| LAMTOR1 | 1879.0226 | -0.285164221 | 0.054253277 | -5.256165856 | 1.47E-07 | 2.42E-06 |
| ZYG11B | 277.7553939 | -0.354507404 | 0.067453907 | -5.255550374 | 1.48E-07 | 2.42E-06 |
| PIP4K2C | 557.6431475 | 0.333494508 | 0.063513547 | 5.250761817 | 1.51E-07 | 2.48E-06 |
| DTNA | 104.3774419 | 0.930706622 | 0.177266626 | 5.250320619 | 1.52E-07 | 2.49E-06 |
| TRIAP1 | 306.8440359 | -0.349182227 | 0.066513024 | -5.249832398 | 1.52E-07 | 2.49E-06 |
| MYL12A | 8472.234916 | 0.248937127 | 0.047422743 | 5.249319415 | 1.53E-07 | 2.50E-06 |
| SLC25A19 | 672.9991222 | -0.323916298 | 0.061708268 | -5.249155599 | 1.53E-07 | 2.50E-06 |
| PPP1R15A | 1250.281097 | 0.651903365 | 0.124294538 | 5.244827144 | 1.56E-07 | 2.55E-06 |
| FAM20A | 88.25990199 | 0.906104251 | 0.172821215 | 5.243015164 | 1.58E-07 | 2.58E-06 |
| GTF2IRD2E | 83.36477082 | -0.774138579 | 0.14771985 | -5.240585992 | 1.60E-07 | 2.61E-06 |
| AGPAT6 | 517.914783 | 0.338443603 | 0.064616595 | 5.237719595 | 1.63E-07 | 2.65E-06 |
| A4GALT | 299.7828596 | 0.932768152 | 0.178126839 | 5.236539063 | 1.64E-07 | 2.66E-06 |
| CD9 | 8670.900841 | -0.451213103 | 0.08618914 | -5.235150319 | 1.65E-07 | 2.68E-06 |
| FCGR3A | 5154.728255 | -0.678525999 | 0.129608793 | -5.235184903 | 1.65E-07 | 2.68E-06 |
| DGAT2 | 179.8322778 | 0.783076066 | 0.150011796 | 5.220096596 | 1.79E-07 | 2.90E-06 |
| C19orf59 | 3001.105471 | 0.867128794 | 0.166183987 | 5.217884173 | 1.81E-07 | 2.94E-06 |
| C21orf91 | 139.9264329 | 0.566977017 | 0.108750339 | 5.213565521 | 1.85E-07 | 3.00E-06 |
| PSME4 | 1531.5516 | 0.502924666 | 0.096507392 | 5.211255359 | 1.88E-07 | 3.04E-06 |
| LAMA3 | 9.249102588 | 2.276452756 | 0.436971709 | 5.209611304 | 1.89E-07 | 3.06E-06 |
| DAXX | 657.6114404 | 0.658912019 | 0.126559197 | 5.206354307 | 1.93E-07 | 3.11E-06 |
| LYL1 | 198.1592471 | -0.513371102 | 0.098610881 | -5.206028959 | 1.93E-07 | 3.11E-06 |
| SLCO4A1 | 35.4709649 | 1.198579995 | 0.230223357 | 5.20616158 | 1.93E-07 | 3.11E-06 |
| CCNI | 4728.039521 | -0.510227616 | 0.09808775 | -5.201746572 | 1.97E-07 | 3.18E-06 |
| OSCAR | 373.6982696 | 0.603752499 | 0.116100041 | 5.200278066 | 1.99E-07 | 3.20E-06 |
| GSN | 6857.040159 | 0.612561041 | 0.117849865 | 5.197808572 | 2.02E-07 | 3.24E-06 |
| AGTRAP | 1773.454989 | 0.45861993 | 0.088252499 | 5.196679255 | 2.03E-07 | 3.26E-06 |
| STARD8 | 360.9429223 | 0.81546358 | 0.15706275 | 5.19196042 | 2.08E-07 | 3.34E-06 |
| RNF207 | 93.91886777 | 0.959469267 | 0.184884248 | 5.189567405 | 2.11E-07 | 3.38E-06 |
| CD55 | 69.13655728 | 0.932500886 | 0.179784116 | 5.186781262 | 2.14E-07 | 3.43E-06 |
| GAS5 | 102.8143676 | -0.62642094 | 0.120842956 | -5.183760497 | 2.17E-07 | 3.48E-06 |
| ZNF331 | 149.8263732 | -0.681891704 | 0.131601327 | -5.18149566 | 2.20E-07 | 3.52E-06 |
| FAM26F | 72.43142095 | 1.530860785 | 0.295466602 | 5.181163535 | 2.21E-07 | 3.53E-06 |
| SNHG12 | 170.3013072 | -0.560917645 | 0.108264354 | -5.181000263 | 2.21E-07 | 3.53E-06 |
| POLR2L | 2555.284869 | -0.386625141 | 0.07465 | -5.179171377 | 2.23E-07 | 3.56E-06 |
| BCL6 | 745.7111297 | 0.378417849 | 0.073074159 | 5.178545365 | 2.24E-07 | 3.57E-06 |
| GATM | 35.36285947 | -0.784905018 | 0.151637995 | -5.17617644 | 2.26E-07 | 3.61E-06 |
| CASP7 | 50.81553965 | 0.594550601 | 0.114906813 | 5.174198007 | 2.29E-07 | 3.65E-06 |

| | | | | | | |
|---|---|---|---|---|---|---|
| TRIM56 | 156.0801433 | 0.578554941 | 0.11191644 | 5.169525932 | 2.35E-07 | 3.73E-06 |
| PXK | 405.762142 | -0.501525758 | 0.097065355 | -5.166887376 | 2.38E-07 | 3.78E-06 |
| DNAJC5B | 267.6745268 | -0.659128731 | 0.127582117 | -5.16630973 | 2.39E-07 | 3.79E-06 |
| FSCN1 | 211.3905773 | 0.668151876 | 0.129394553 | 5.163678538 | 2.42E-07 | 3.84E-06 |
| HMGCS1 | 96.79570449 | 0.702503925 | 0.13605628 | 5.163333318 | 2.43E-07 | 3.85E-06 |
| PLXDC2 | 3918.618169 | -0.418545558 | 0.081113022 | -5.160029162 | 2.47E-07 | 3.91E-06 |
| CES1 | 25574.88286 | -0.435869031 | 0.084482091 | -5.159306837 | 2.48E-07 | 3.92E-06 |
| PLEKHO1 | 403.0621243 | 0.568161501 | 0.110170184 | 5.157125815 | 2.51E-07 | 3.97E-06 |
| RPS10 | 3159.046546 | -0.329707438 | 0.063939169 | -5.15657996 | 2.52E-07 | 3.97E-06 |
| SNURF | 229.0122654 | -0.54170386 | 0.105059158 | -5.156179335 | 2.52E-07 | 3.98E-06 |
| DIXDC1 | 108.0774035 | 0.615521666 | 0.119422273 | 5.154161369 | 2.55E-07 | 4.02E-06 |
| ABHD2 | 3836.646357 | -0.533180802 | 0.103498428 | -5.151583587 | 2.58E-07 | 4.07E-06 |
| RCN1 | 717.7762646 | 0.468826786 | 0.091057376 | 5.148696452 | 2.62E-07 | 4.13E-06 |
| SH2D4A | 41.17438625 | 0.819157078 | 0.159289778 | 5.14255897 | 2.71E-07 | 4.27E-06 |
| KCNE3 | 389.7396655 | -0.659642349 | 0.128286305 | -5.141954556 | 2.72E-07 | 4.28E-06 |
| MRO | 5.523295768 | -3.013156797 | 0.586066395 | -5.141323278 | 2.73E-07 | 4.29E-06 |
| CSNK1E | 387.2316975 | -0.331191242 | 0.064465553 | -5.137491668 | 2.78E-07 | 4.37E-06 |
| CD97 | 2450.402627 | 0.551897873 | 0.107474613 | 5.135146432 | 2.82E-07 | 4.42E-06 |
| IRF6 | 26.34484966 | 1.130614472 | 0.220172832 | 5.135122544 | 2.82E-07 | 4.42E-06 |
| PADI2 | 279.5798772 | -0.886328031 | 0.172593581 | -5.1353476 | 2.82E-07 | 4.42E-06 |
| TP53INP1 | 36.47606153 | -1.059712362 | 0.206415561 | -5.133878262 | 2.84E-07 | 4.44E-06 |
| TPST2 | 506.1689379 | -0.554953118 | 0.108095195 | -5.133929573 | 2.84E-07 | 4.44E-06 |
| MLLT4 | 279.3421945 | 0.548404369 | 0.106827408 | 5.133554925 | 2.84E-07 | 4.44E-06 |
| ATG2A | 593.3181109 | 0.569033094 | 0.110866406 | 5.132601594 | 2.86E-07 | 4.46E-06 |
| RPS6KA3 | 1215.205958 | 0.365952792 | 0.071334271 | 5.130111865 | 2.90E-07 | 4.52E-06 |
| HIST1H4F | 100.1034986 | -0.910132556 | 0.177470219 | -5.128367783 | 2.92E-07 | 4.55E-06 |
| ZNF483 | 25.17523726 | -1.004436344 | 0.195963234 | -5.125636704 | 2.97E-07 | 4.62E-06 |
| CD302 | 328.3863432 | -1.323721924 | 0.258504419 | -5.12069361 | 3.04E-07 | 4.73E-06 |
| ENDOG | 155.8761393 | -0.354531593 | 0.069238067 | -5.120472178 | 3.05E-07 | 4.74E-06 |
| SLC24A4 | 42.70073016 | -1.055433658 | 0.206195318 | -5.118611175 | 3.08E-07 | 4.78E-06 |
| SLC7A1 | 642.3737588 | 0.493963618 | 0.096524004 | 5.117521018 | 3.10E-07 | 4.80E-06 |
| SAMHD1 | 3264.254646 | -0.38954127 | 0.076142819 | -5.115929174 | 3.12E-07 | 4.84E-06 |
| LONRF1 | 189.2527475 | 0.797658128 | 0.155935508 | 5.115307845 | 3.13E-07 | 4.85E-06 |
| PTPMT1 | 1211.741879 | 0.472486335 | 0.092463678 | 5.109966931 | 3.22E-07 | 4.99E-06 |
| CCBL1 | 17.97468465 | -1.106544838 | 0.216681053 | -5.106790947 | 3.28E-07 | 5.07E-06 |
| CLLU1OS | 61.89922709 | 1.641663928 | 0.322083395 | 5.097015094 | 3.45E-07 | 5.33E-06 |
| HDDC2 | 743.9659961 | -0.370649396 | 0.072775202 | -5.09307274 | 3.52E-07 | 5.44E-06 |
| PCBP2 | 9331.489298 | -0.28578238 | 0.056130741 | -5.091370128 | 3.55E-07 | 5.48E-06 |
| CAMKK1 | 80.24558602 | -0.708786903 | 0.139261258 | -5.089620145 | 3.59E-07 | 5.53E-06 |
| OAT | 601.2052902 | -0.407208757 | 0.080010632 | -5.089433056 | 3.59E-07 | 5.53E-06 |
| DYRK3 | 92.13421563 | 0.805472169 | 0.158367721 | 5.086088016 | 3.66E-07 | 5.62E-06 |
| VCAM1 | 6.061703486 | 2.721082038 | 0.535052316 | 5.085637343 | 3.66E-07 | 5.63E-06 |
| GSTM2 | 31.42122416 | -1.298161265 | 0.255467747 | -5.081507469 | 3.74E-07 | 5.75E-06 |
| GNLY | 48.09580017 | 1.335056189 | 0.26293452 | 5.077523447 | 3.82E-07 | 5.87E-06 |
| ANAPC15 | 259.1362933 | -0.768760789 | 0.151517427 | -5.073745017 | 3.90E-07 | 5.98E-06 |
| APBB1IP | 283.8174165 | -0.620766633 | 0.122359937 | -5.073283349 | 3.91E-07 | 5.99E-06 |
| CR1L | 21.72930758 | 1.300622068 | 0.256379774 | 5.073029154 | 3.92E-07 | 5.99E-06 |

| | | | | | | |
|---|---|---|---|---|---|---|
| SRGN | 21937.7149 | 0.511434916 | 0.100852425 | 5.071121671 | 3.95E-07 | 6.05E-06 |
| SMPDL3A | 51.96313272 | -1.146181657 | 0.226235431 | -5.06632252 | 4.06E-07 | 6.20E-06 |
| ZNF528 | 41.74202712 | -0.671746123 | 0.132596603 | -5.066088472 | 4.06E-07 | 6.20E-06 |
| CD58 | 1135.471211 | 0.343210853 | 0.067760602 | 5.065050247 | 4.08E-07 | 6.23E-06 |
| RPL5 | 3413.434717 | -0.696289709 | 0.137594359 | -5.060452435 | 4.18E-07 | 6.38E-06 |
| NAAA | 382.8720831 | -0.587214468 | 0.116052833 | -5.05988915 | 4.20E-07 | 6.39E-06 |
| NUP85 | 184.8679782 | -0.418606406 | 0.082730091 | -5.059905084 | 4.19E-07 | 6.39E-06 |
| RFX7 | 149.4377541 | -0.433141847 | 0.085606524 | -5.059682688 | 4.20E-07 | 6.39E-06 |
| IGFBP4 | 14.8577171 | 2.068936281 | 0.408978076 | 5.058795088 | 4.22E-07 | 6.41E-06 |
| EIF4EBP1 | 754.0352683 | -0.372020861 | 0.073551001 | -5.057998622 | 4.24E-07 | 6.43E-06 |
| UBE4B | 387.1351509 | -0.374261062 | 0.074008923 | -5.056972121 | 4.26E-07 | 6.46E-06 |
| PDCD6IP | 2735.093898 | 0.495329601 | 0.097982984 | 5.055261427 | 4.30E-07 | 6.52E-06 |
| FFAR2 | 52.06165419 | 1.604469424 | 0.317689537 | 5.050432067 | 4.41E-07 | 6.68E-06 |
| CD38 | 687.8548159 | 0.903575756 | 0.178941557 | 5.04955792 | 4.43E-07 | 6.70E-06 |
| MBP | 311.8944687 | -0.655055669 | 0.12976457 | -5.04803174 | 4.46E-07 | 6.75E-06 |
| MMP1 | 11.14498608 | 3.102944603 | 0.614977498 | 5.045622989 | 4.52E-07 | 6.83E-06 |
| DLG4 | 166.3852881 | -0.512301126 | 0.101547141 | -5.044958646 | 4.54E-07 | 6.84E-06 |
| SLC2A5 | 10.36355016 | -1.544501392 | 0.306148146 | -5.044947715 | 4.54E-07 | 6.84E-06 |
| TBC1D22B | 285.6584131 | 0.379579343 | 0.075255219 | 5.043893914 | 4.56E-07 | 6.88E-06 |
| ARMC10 | 318.5749235 | -0.34787784 | 0.068990634 | -5.042392299 | 4.60E-07 | 6.92E-06 |
| CPE | 548.9929109 | -0.610201499 | 0.121011585 | -5.042504818 | 4.59E-07 | 6.92E-06 |
| SH3PXD2B | 430.1864644 | 0.741074716 | 0.147013933 | 5.040846828 | 4.63E-07 | 6.97E-06 |
| SERINC5 | 2111.28725 | -0.293224066 | 0.058243111 | -5.034484971 | 4.79E-07 | 7.20E-06 |
| ZNF318 | 277.2692723 | 0.374783661 | 0.0745181 | 5.029431247 | 4.92E-07 | 7.39E-06 |
| ZFYVE16 | 1436.730218 | 0.382733577 | 0.0761081 | 5.028815293 | 4.94E-07 | 7.40E-06 |
| DOCK10 | 588.4250293 | -0.387504575 | 0.077062489 | -5.028446155 | 4.94E-07 | 7.41E-06 |
| NPAS2 | 48.24262529 | 1.560106677 | 0.310713159 | 5.021051184 | 5.14E-07 | 7.70E-06 |
| LAIR1 | 2482.249951 | -0.420662007 | 0.083782439 | -5.020885169 | 5.14E-07 | 7.70E-06 |
| IL17RA | 2169.105386 | 0.389001556 | 0.077500164 | 5.019364295 | 5.18E-07 | 7.75E-06 |
| ANXA1 | 16204.0448 | 0.327140278 | 0.065215122 | 5.016325463 | 5.27E-07 | 7.87E-06 |
| LAMTOR4 | 250.3580222 | -0.476907777 | 0.095108034 | -5.0143795 | 5.32E-07 | 7.94E-06 |
| C22orf28 | 1126.904557 | 0.325135424 | 0.064849224 | 5.013713385 | 5.34E-07 | 7.97E-06 |
| LHFP | 6.492974453 | 2.192455983 | 0.437329868 | 5.013277493 | 5.35E-07 | 7.98E-06 |
| SLAMF1 | 12.95463325 | 2.151317191 | 0.429176634 | 5.012661502 | 5.37E-07 | 8.00E-06 |
| KDM2A | 1449.885931 | 0.333718076 | 0.066622659 | 5.00907772 | 5.47E-07 | 8.14E-06 |
| LILRB4 | 6684.697409 | 0.551185588 | 0.110044166 | 5.008766988 | 5.48E-07 | 8.15E-06 |
| TEAD3 | 4.171580038 | 2.152715822 | 0.430029683 | 5.005970301 | 5.56E-07 | 8.26E-06 |
| MAPKAPK5 | 96.98602138 | -0.517834256 | 0.103466649 | -5.004842252 | 5.59E-07 | 8.30E-06 |
| NANS | 2203.223351 | -0.229939322 | 0.045982246 | -5.000610875 | 5.71E-07 | 8.48E-06 |
| ZDHHC3 | 383.7814831 | 0.366252545 | 0.073287952 | 4.997445483 | 5.81E-07 | 8.61E-06 |
| CCR4 | 6.139489101 | 2.243935612 | 0.449570548 | 4.991286951 | 6.00E-07 | 8.88E-06 |
| ATN1 | 1238.009465 | 0.348273905 | 0.069779583 | 4.991057444 | 6.00E-07 | 8.89E-06 |
| SLC28A3 | 298.9446388 | 0.811171537 | 0.162676607 | 4.986405556 | 6.15E-07 | 9.10E-06 |
| UBL3 | 1481.972171 | -0.276441648 | 0.0555169 | -4.97941434 | 6.38E-07 | 9.42E-06 |
| KCTD10 | 421.2579343 | -0.343001139 | 0.068900319 | -4.978222779 | 6.42E-07 | 9.48E-06 |
| BATF | 121.8111453 | -0.856020227 | 0.172069387 | -4.974854868 | 6.53E-07 | 9.63E-06 |
| MT1JP | 28.3863593 | 2.1677965 | 0.435867004 | 4.973527425 | 6.57E-07 | 9.69E-06 |

| | | | | | | |
|---|---|---|---|---|---|---|
| TREML3P | 64.98249999 | 0.87939023 | 0.176859895 | 4.972242181 | 6.62E-07 | 9.75E-06 |
| FLI1 | 365.6198997 | -0.604263635 | 0.121551867 | -4.97124107 | 6.65E-07 | 9.79E-06 |
| GCNT2 | 145.7292287 | 0.490115278 | 0.098646394 | 4.968405438 | 6.75E-07 | 9.93E-06 |
| ITGAV | 1385.358147 | 0.518679947 | 0.104415498 | 4.967461328 | 6.78E-07 | 9.97E-06 |
| IGFN1 | 8.985255356 | 3.314752256 | 0.667455086 | 4.966255144 | 6.83E-07 | 1.00E-05 |
| TFEC | 836.4953978 | 0.336808563 | 0.067874417 | 4.962231414 | 6.97E-07 | 1.02E-05 |
| IKZF1 | 528.6202281 | -0.443555691 | 0.089504964 | -4.955654643 | 7.21E-07 | 1.06E-05 |
| SLC46A3 | 462.8943491 | -0.8405352 | 0.1697449 | -4.951755264 | 7.35E-07 | 1.08E-05 |
| POC1A | 27.75834476 | 0.750581519 | 0.15165926 | 4.949130838 | 7.45E-07 | 1.09E-05 |
| CA2 | 2226.922645 | 0.916809474 | 0.185292481 | 4.94790436 | 7.50E-07 | 1.10E-05 |
| PATZ1 | 22.50262906 | -1.027086402 | 0.207672438 | -4.945703964 | 7.59E-07 | 1.11E-05 |
| RPL15 | 11086.46866 | -0.442796119 | 0.089539399 | -4.945265712 | 7.60E-07 | 1.11E-05 |
| MLLT6 | 827.3932279 | 0.540397872 | 0.109308334 | 4.94379387 | 7.66E-07 | 1.12E-05 |
| ADO | 519.9242789 | 0.504956324 | 0.102145183 | 4.94351578 | 7.67E-07 | 1.12E-05 |
| SH3RF2 | 6.350180277 | 2.009349556 | 0.406533989 | 4.942636069 | 7.71E-07 | 1.12E-05 |
| PTPRC | 161.5123657 | 0.570459672 | 0.115425911 | 4.942215024 | 7.72E-07 | 1.12E-05 |
| NINJ1 | 2319.072122 | 0.672277208 | 0.136032737 | 4.942025154 | 7.73E-07 | 1.12E-05 |
| CLIC3 | 8.510036126 | 1.547042092 | 0.313060794 | 4.941666669 | 7.75E-07 | 1.13E-05 |
| ABL2 | 899.7504135 | 0.88636439 | 0.179438601 | 4.939652817 | 7.83E-07 | 1.14E-05 |
| C14orf159 | 52.10351451 | 0.890437217 | 0.180335605 | 4.937667298 | 7.91E-07 | 1.15E-05 |
| LINC00515 | 46.05743962 | 0.995171085 | 0.201546998 | 4.937662644 | 7.91E-07 | 1.15E-05 |
| AKIRIN1 | 1274.270317 | 0.45427852 | 0.092012603 | 4.937133684 | 7.93E-07 | 1.15E-05 |
| POLR1E | 164.6980964 | -0.373485253 | 0.075658067 | -4.936489498 | 7.95E-07 | 1.15E-05 |
| LRP10 | 1848.374091 | 0.402979193 | 0.081639955 | 4.93605358 | 7.97E-07 | 1.15E-05 |
| ABI1 | 2003.745487 | 0.368468428 | 0.074666826 | 4.934834476 | 8.02E-07 | 1.16E-05 |
| CLDN1 | 9.687171278 | 1.763223941 | 0.357349866 | 4.934167064 | 8.05E-07 | 1.16E-05 |
| SMAD7 | 570.2605052 | 0.466280664 | 0.094518721 | 4.933209603 | 8.09E-07 | 1.17E-05 |
| ANAPC13 | 497.3782255 | -0.384911617 | 0.078085492 | -4.929361499 | 8.25E-07 | 1.19E-05 |
| SESN3 | 42.99324508 | -1.087065885 | 0.22059021 | -4.927987907 | 8.31E-07 | 1.20E-05 |
| MLF2 | 1958.927331 | 0.302221498 | 0.061353432 | 4.925910255 | 8.40E-07 | 1.21E-05 |
| PNRC2 | 2133.274197 | 0.355588122 | 0.072205627 | 4.924659433 | 8.45E-07 | 1.22E-05 |
| ARG2 | 94.48660952 | -0.652866887 | 0.132593379 | -4.923827197 | 8.49E-07 | 1.22E-05 |
| UBR2 | 628.6859169 | 0.20919892 | 0.042489763 | 4.923513442 | 8.50E-07 | 1.22E-05 |
| DUSP2 | 216.2437629 | 0.838616556 | 0.170388358 | 4.921794935 | 8.58E-07 | 1.23E-05 |
| EIF2C1 | 313.0907455 | -0.501623703 | 0.101935118 | -4.921009678 | 8.61E-07 | 1.24E-05 |
| TBCK | 320.1819356 | -0.311032633 | 0.06321511 | -4.920226098 | 8.64E-07 | 1.24E-05 |
| ABAT | 184.5596028 | -0.859070077 | 0.174716425 | -4.916939418 | 8.79E-07 | 1.26E-05 |
| ANTXR2 | 188.9951521 | 0.662593592 | 0.134811883 | 4.914949464 | 8.88E-07 | 1.27E-05 |
| FAM40B | 54.48431652 | 0.898424319 | 0.182950719 | 4.910744936 | 9.07E-07 | 1.30E-05 |
| LOC10049 | 54.50529713 | 0.609898895 | 0.124200014 | 4.910618565 | 9.08E-07 | 1.30E-05 |
| NCF1 | 2924.403966 | 0.731182005 | 0.148897922 | 4.91062597 | 9.08E-07 | 1.30E-05 |
| RGS18 | 17.80712662 | -2.650401036 | 0.539937919 | -4.9087144 | 9.17E-07 | 1.31E-05 |
| LOC10028 | 342.3421757 | -0.411959242 | 0.083995837 | -4.904519751 | 9.37E-07 | 1.34E-05 |
| TTC4 | 650.8907242 | 0.332201838 | 0.067773701 | 4.901633431 | 9.50E-07 | 1.35E-05 |
| PPP1R1A | 17.71633251 | 1.42304675 | 0.290628625 | 4.896443878 | 9.76E-07 | 1.39E-05 |
| NR3C2 | 45.4575456 | -0.844355443 | 0.17246468 | -4.895816596 | 9.79E-07 | 1.39E-05 |
| SLC35A1 | 329.0527042 | -0.448247268 | 0.091559566 | -4.895690163 | 9.80E-07 | 1.39E-05 |

| | | | | | | |
|---|---|---|---|---|---|---|
| EIF3E | 2511.35524 | -0.368891115 | 0.075386429 | -4.89333584 | 9.91E-07 | 1.41E-05 |
| PTPRJ | 1426.637884 | 0.529008324 | 0.108118639 | 4.892850381 | 9.94E-07 | 1.41E-05 |
| C1orf122 | 250.3955663 | 0.742041685 | 0.151702383 | 4.891430652 | 1.00E-06 | 1.42E-05 |
| NYNRIN | 8.232180917 | -1.866590587 | 0.381910982 | -4.887501731 | 1.02E-06 | 1.45E-05 |
| AQP9 | 4732.35787 | 0.338114749 | 0.069223807 | 4.884370909 | 1.04E-06 | 1.47E-05 |
| ATRN | 351.9451741 | -0.485647302 | 0.09944206 | -4.883721255 | 1.04E-06 | 1.47E-05 |
| TANK | 700.293835 | 0.526999658 | 0.10791254 | 4.883581254 | 1.04E-06 | 1.47E-05 |
| A2M | 1927.374896 | -0.676094093 | 0.138454397 | -4.883153644 | 1.04E-06 | 1.47E-05 |
| SERPINB6 | 1605.874357 | -0.287748746 | 0.05892684 | -4.883152534 | 1.04E-06 | 1.47E-05 |
| MT1L | 196.3741111 | 0.935993474 | 0.191732722 | 4.881761767 | 1.05E-06 | 1.48E-05 |
| VAMP8 | 4028.328053 | -0.287634043 | 0.05892631 | -4.881249848 | 1.05E-06 | 1.49E-05 |
| IFNGR1 | 1589.809179 | -0.490799629 | 0.100592615 | -4.879082124 | 1.07E-06 | 1.50E-05 |
| VSIG4 | 697.1684689 | -0.674284395 | 0.13832018 | -4.874808542 | 1.09E-06 | 1.53E-05 |
| ESPL1 | 16.05534034 | 1.246844708 | 0.25582799 | 4.873761892 | 1.09E-06 | 1.54E-05 |
| EIF3H | 1542.933329 | -0.362526549 | 0.074402679 | -4.872493237 | 1.10E-06 | 1.55E-05 |
| C4B | 18.71323764 | 1.697211516 | 0.348554095 | 4.869291565 | 1.12E-06 | 1.57E-05 |
| SLC22A4 | 18.14044122 | 1.611921022 | 0.331018732 | 4.869576446 | 1.12E-06 | 1.57E-05 |
| SPPL2A | 2218.088267 | 0.348410566 | 0.071551835 | 4.869344924 | 1.12E-06 | 1.57E-05 |
| TBC1D24 | 212.737805 | -0.376107332 | 0.07724313 | -4.869136335 | 1.12E-06 | 1.57E-05 |
| SCCPDH | 2252.915704 | -0.311432211 | 0.063964279 | -4.868845779 | 1.12E-06 | 1.57E-05 |
| RAPGEF1 | 87.78636199 | 0.9329149 | 0.191844649 | 4.86286641 | 1.16E-06 | 1.62E-05 |
| RAP1B | 1959.04284 | 0.351517684 | 0.072316288 | 4.860836932 | 1.17E-06 | 1.63E-05 |
| BPHL | 134.5420798 | -0.421431769 | 0.086719348 | -4.859720227 | 1.18E-06 | 1.64E-05 |
| RPS8 | 6574.445873 | -0.325666726 | 0.067115113 | -4.852360519 | 1.22E-06 | 1.70E-05 |
| ACP5 | 24922.42612 | -0.638058233 | 0.131546701 | -4.850431272 | 1.23E-06 | 1.72E-05 |
| FMNL3 | 2206.002011 | 0.51868417 | 0.10704403 | 4.845521721 | 1.26E-06 | 1.76E-05 |
| CDYL | 514.2126264 | 0.360139076 | 0.074327238 | 4.845317642 | 1.26E-06 | 1.76E-05 |
| RELT | 138.5879927 | 0.48691092 | 0.100536074 | 4.843146334 | 1.28E-06 | 1.78E-05 |
| ATF5 | 1276.497898 | 0.423732178 | 0.087657001 | 4.833979859 | 1.34E-06 | 1.86E-05 |
| CTNNAL1 | 23.11495626 | 1.190274316 | 0.24623963 | 4.833804847 | 1.34E-06 | 1.86E-05 |
| HSDL2 | 476.4107288 | -0.376162788 | 0.077837743 | -4.832652848 | 1.35E-06 | 1.87E-05 |
| GPBP1 | 621.9262142 | 0.277411249 | 0.057405858 | 4.832455405 | 1.35E-06 | 1.87E-05 |
| TLE1 | 108.3414764 | 1.050367097 | 0.217369453 | 4.832174366 | 1.35E-06 | 1.87E-05 |
| TNFRSF18 | 14.8843674 | 2.12661711 | 0.440118857 | 4.831915464 | 1.35E-06 | 1.87E-05 |
| CASD1 | 104.053387 | -0.819925068 | 0.169751933 | -4.830136862 | 1.36E-06 | 1.89E-05 |
| IL36B | 82.22126213 | 0.734190634 | 0.152021229 | 4.829527014 | 1.37E-06 | 1.89E-05 |
| IFNAR2 | 671.8249449 | -0.333997598 | 0.069219091 | -4.825223687 | 1.40E-06 | 1.93E-05 |
| ST8SIA4 | 36.22661052 | -1.298441481 | 0.269175685 | -4.823769585 | 1.41E-06 | 1.95E-05 |
| WWC3 | 158.2001433 | -0.447047856 | 0.092685648 | -4.823269451 | 1.41E-06 | 1.95E-05 |
| EIF4EBP2 | 2878.692386 | -0.330419487 | 0.068544318 | -4.820523357 | 1.43E-06 | 1.97E-05 |
| GNA14 | 12.70785694 | 2.351039003 | 0.487841064 | 4.819272459 | 1.44E-06 | 1.99E-05 |
| NQO1 | 950.9269467 | 0.692882719 | 0.143866464 | 4.816151713 | 1.46E-06 | 2.02E-05 |
| TMEM38B | 1014.980893 | 0.618172179 | 0.128391286 | 4.814751819 | 1.47E-06 | 2.03E-05 |
| GAB1 | 137.4439728 | -0.439283034 | 0.091260468 | -4.813508442 | 1.48E-06 | 2.04E-05 |
| ADAT2 | 27.71832516 | -0.718120283 | 0.149330365 | -4.808936773 | 1.52E-06 | 2.08E-05 |
| BAK1 | 787.9494571 | 0.599346468 | 0.124632594 | 4.808906301 | 1.52E-06 | 2.08E-05 |
| SLC7A5 | 277.070375 | 0.903110493 | 0.187831051 | 4.808100077 | 1.52E-06 | 2.09E-05 |

| | | | | | | |
|---|---|---|---|---|---|---|
| RTN4RL2 | 101.2027315 | 1.270204548 | 0.264251015 | 4.806810485 | 1.53E-06 | 2.10E-05 |
| FBL | 349.0018738 | -0.355930224 | 0.074055952 | -4.80623387 | 1.54E-06 | 2.11E-05 |
| PLA2G6 | 99.61162736 | -0.756547405 | 0.157635975 | -4.799332155 | 1.59E-06 | 2.18E-05 |
| LONP1 | 1382.660306 | 0.43409072 | 0.090491218 | 4.797048061 | 1.61E-06 | 2.20E-05 |
| PARL | 322.1712585 | -0.364096871 | 0.075939226 | -4.794582359 | 1.63E-06 | 2.23E-05 |
| SLC12A9 | 200.9030091 | -0.383545625 | 0.080078962 | -4.789592875 | 1.67E-06 | 2.28E-05 |
| EVI2A | 1465.685999 | -0.49101931 | 0.102589163 | -4.786268819 | 1.70E-06 | 2.32E-05 |
| JUP | 169.65454 | 1.107393204 | 0.2315578 | 4.782361908 | 1.73E-06 | 2.36E-05 |
| CD47 | 1353.742016 | 0.406940335 | 0.085103784 | 4.78169493 | 1.74E-06 | 2.37E-05 |
| ACACB | 193.8379155 | -0.663656814 | 0.138818427 | -4.780754446 | 1.75E-06 | 2.38E-05 |
| MED13 | 865.4607048 | 0.296194734 | 0.061960706 | 4.780364071 | 1.75E-06 | 2.38E-05 |
| LNPEP | 176.6898354 | 0.868337447 | 0.181900297 | 4.773700004 | 1.81E-06 | 2.46E-05 |
| LAT | 157.5986459 | 0.678649645 | 0.142242735 | 4.771067184 | 1.83E-06 | 2.49E-05 |
| CARD11 | 39.12182689 | 0.915231619 | 0.191985563 | 4.76718981 | 1.87E-06 | 2.54E-05 |
| TCN2 | 691.7704443 | -0.619766242 | 0.130078152 | -4.764568333 | 1.89E-06 | 2.57E-05 |
| PCK2 | 706.2306951 | 0.390545637 | 0.081973335 | 4.764300954 | 1.90E-06 | 2.57E-05 |
| LRP1 | 7549.262783 | -0.696842082 | 0.146425254 | -4.759029339 | 1.95E-06 | 2.64E-05 |
| COMMD9 | 475.8503834 | -0.343367085 | 0.072178057 | -4.757222644 | 1.96E-06 | 2.66E-05 |
| KIAA0754 | 135.2399251 | -0.611835653 | 0.128686626 | -4.754461832 | 1.99E-06 | 2.69E-05 |
| LTA4H | 8527.550289 | -0.419358767 | 0.08821408 | -4.753875644 | 2.00E-06 | 2.70E-05 |
| TRHDE-AS1 | 19.38416899 | -1.107284193 | 0.23305486 | -4.751174015 | 2.02E-06 | 2.73E-05 |
| IL28RA | 176.9412631 | 0.66151967 | 0.139242152 | 4.750857854 | 2.03E-06 | 2.74E-05 |
| NCKAP5L | 321.4363598 | -0.397832542 | 0.083754102 | -4.750006636 | 2.03E-06 | 2.74E-05 |
| PIGF | 214.4958702 | -0.446103443 | 0.093915857 | -4.750033246 | 2.03E-06 | 2.74E-05 |
| TMBIM1 | 4513.960595 | 0.311957425 | 0.065779326 | 4.742484322 | 2.11E-06 | 2.84E-05 |
| PTPN1 | 951.4231636 | 0.439555999 | 0.092746877 | 4.739307796 | 2.14E-06 | 2.89E-05 |
| TAPBPL | 714.7008561 | 0.351700058 | 0.074216311 | 4.738851213 | 2.15E-06 | 2.89E-05 |
| C17orf49 | 662.6035687 | 0.414342312 | 0.087465312 | 4.737218698 | 2.17E-06 | 2.91E-05 |
| SOCS2 | 4.768618486 | 2.46791832 | 0.521035163 | 4.736567689 | 2.17E-06 | 2.92E-05 |
| ZNF671 | 59.47165787 | -0.532749351 | 0.112482924 | -4.736268685 | 2.18E-06 | 2.92E-05 |
| UBXN4 | 1406.835189 | 0.238060934 | 0.050274122 | 4.735257924 | 2.19E-06 | 2.93E-05 |
| ZMAT3 | 215.6000576 | -0.553372222 | 0.116963668 | -4.731146258 | 2.23E-06 | 2.99E-05 |
| BNC2 | 6.281067756 | -2.127583797 | 0.450076822 | -4.727156995 | 2.28E-06 | 3.05E-05 |
| DHRS1 | 117.951239 | -0.56876703 | 0.120316511 | -4.72725669 | 2.28E-06 | 3.05E-05 |
| RPL32 | 4616.477363 | -0.433648162 | 0.091819124 | -4.722852308 | 2.33E-06 | 3.11E-05 |
| PSMB7 | 1282.808191 | 0.538878009 | 0.114104572 | 4.722667995 | 2.33E-06 | 3.11E-05 |
| ITGA2 | 9.129615464 | 1.953451833 | 0.413979734 | 4.718713681 | 2.37E-06 | 3.17E-05 |
| RHBDF2 | 511.7496306 | 0.741020708 | 0.15717619 | 4.714586273 | 2.42E-06 | 3.23E-05 |
| MLPH | 494.3493027 | -0.514462028 | 0.109185813 | -4.711802884 | 2.46E-06 | 3.28E-05 |
| TNKS1BP1 | 43.23447373 | 0.849810229 | 0.180384579 | 4.711102434 | 2.46E-06 | 3.28E-05 |
| KIAA0196 | 492.7452439 | -0.266996122 | 0.056705934 | -4.708433567 | 2.50E-06 | 3.32E-05 |
| G6PD | 5809.005016 | 0.472617593 | 0.100382914 | 4.708147762 | 2.50E-06 | 3.33E-05 |
| SDF4 | 2754.860302 | 0.312230945 | 0.066328583 | 4.707336245 | 2.51E-06 | 3.34E-05 |
| DNAJA1 | 6891.58394 | 0.350121721 | 0.074416779 | 4.704876067 | 2.54E-06 | 3.38E-05 |
| RPLP1 | 6571.005089 | -0.275816586 | 0.058646664 | -4.703022603 | 2.56E-06 | 3.40E-05 |
| GALNT4 | 224.1337889 | 0.378483055 | 0.080495863 | 4.701894517 | 2.58E-06 | 3.42E-05 |
| DHRS9 | 457.9453982 | 1.042399607 | 0.221814845 | 4.699413176 | 2.61E-06 | 3.46E-05 |

| | | | | | | |
|---|---|---|---|---|---|---|
| TMEM194/ | 77.96903127 | 1.355397473 | 0.288530268 | 4.697591985 | 2.63E-06 | 3.49E-05 |
| CENPT | 3.691860109 | 1.934700505 | 0.411979129 | 4.696112902 | 2.65E-06 | 3.51E-05 |
| PCNXL2 | 75.94463446 | 0.627022315 | 0.133588821 | 4.693673558 | 2.68E-06 | 3.55E-05 |
| DHRS3 | 3117.580419 | -0.461153289 | 0.098269998 | -4.692716993 | 2.70E-06 | 3.57E-05 |
| SPTSSA | 452.0784749 | -0.432701608 | 0.092241945 | -4.690941909 | 2.72E-06 | 3.59E-05 |
| ZNF480 | 29.67320442 | -0.836143551 | 0.178257393 | -4.690652862 | 2.72E-06 | 3.60E-05 |
| OR2A4 | 106.4277769 | 0.967084741 | 0.20626794 | 4.688487908 | 2.75E-06 | 3.63E-05 |
| IFT20 | 239.360952 | -0.309891687 | 0.066107029 | -4.687726751 | 2.76E-06 | 3.64E-05 |
| PDGFB | 81.82243695 | 0.766381057 | 0.163599333 | 4.684499891 | 2.81E-06 | 3.70E-05 |
| SAT2 | 439.0356642 | -0.370230093 | 0.079036644 | -4.684284093 | 2.81E-06 | 3.70E-05 |
| FTO | 603.9342533 | -0.244650109 | 0.052229703 | -4.684118359 | 2.81E-06 | 3.70E-05 |
| PVR | 527.9437898 | 0.516233819 | 0.110230679 | 4.683213623 | 2.82E-06 | 3.71E-05 |
| PEA15 | 2119.148845 | 0.451079987 | 0.096344744 | 4.68193664 | 2.84E-06 | 3.73E-05 |
| CD244 | 8.103218318 | -2.281956754 | 0.487734109 | -4.678690118 | 2.89E-06 | 3.79E-05 |
| SERTAD2 | 253.4122351 | 0.536218085 | 0.114651955 | 4.67692054 | 2.91E-06 | 3.82E-05 |
| ACP2 | 1563.798196 | 0.598604466 | 0.128017228 | 4.675968042 | 2.93E-06 | 3.84E-05 |
| CYP27A1 | 24341.26651 | -0.371019218 | 0.079432731 | -4.670860672 | 3.00E-06 | 3.93E-05 |
| FAM161B | 50.10762361 | -0.615599262 | 0.131807343 | -4.67044739 | 3.01E-06 | 3.94E-05 |
| LSMD1 | 404.7895968 | -0.530040996 | 0.113499711 | -4.669976623 | 3.01E-06 | 3.94E-05 |
| ACAT2 | 178.0901321 | 0.52170211 | 0.111760397 | 4.668040935 | 3.04E-06 | 3.98E-05 |
| ZNF326 | 80.34964828 | 0.536945314 | 0.115051381 | 4.667004515 | 3.06E-06 | 3.99E-05 |
| MCTP1 | 347.3555612 | 0.690922657 | 0.148146501 | 4.663779777 | 3.10E-06 | 4.05E-05 |
| PLS1 | 24.8812727 | 0.814709556 | 0.174707084 | 4.663288627 | 3.11E-06 | 4.06E-05 |
| ID2 | 337.2204616 | -0.501068722 | 0.107459556 | -4.662858676 | 3.12E-06 | 4.07E-05 |
| DNAJB9 | 368.5667661 | 0.390396722 | 0.083729899 | 4.662572471 | 3.12E-06 | 4.07E-05 |
| SWI5 | 48.72559118 | -0.502371092 | 0.107761191 | -4.661892549 | 3.13E-06 | 4.08E-05 |
| CDC42EP1 | 673.6576702 | 0.797311544 | 0.171239993 | 4.656105919 | 3.22E-06 | 4.19E-05 |
| QSOX1 | 3035.387802 | 0.394183048 | 0.084693575 | 4.654226104 | 3.25E-06 | 4.23E-05 |
| DNAH1 | 112.2516091 | -0.969388787 | 0.208411661 | -4.651317412 | 3.30E-06 | 4.28E-05 |
| GRIPAP1 | 338.5203127 | 0.398655141 | 0.085707059 | 4.651368824 | 3.30E-06 | 4.28E-05 |
| JUND | 1275.968444 | 0.464478894 | 0.099904929 | 4.649209011 | 3.33E-06 | 4.32E-05 |
| PCCA | 510.3636565 | -0.414574167 | 0.089201441 | -4.647617384 | 3.36E-06 | 4.35E-05 |
| LMTK2 | 144.6323478 | 0.590693211 | 0.127111659 | 4.647041934 | 3.37E-06 | 4.36E-05 |
| SATB1 | 81.95566291 | -0.909628054 | 0.195914881 | -4.642975811 | 3.43E-06 | 4.45E-05 |
| FAS | 175.7647411 | 0.69593594 | 0.149897376 | 4.642749307 | 3.44E-06 | 4.45E-05 |
| UNC119 | 257.2355274 | -0.308312413 | 0.066433136 | -4.64094321 | 3.47E-06 | 4.48E-05 |
| PALD1 | 7.110565571 | -2.069874232 | 0.446081227 | -4.640128538 | 3.48E-06 | 4.50E-05 |
| CBX7 | 66.36558001 | -0.964880031 | 0.208003357 | -4.63877143 | 3.50E-06 | 4.53E-05 |
| ARMC2 | 11.926194 | -0.963772783 | 0.207776586 | -4.638505246 | 3.51E-06 | 4.53E-05 |
| TFAP4 | 52.68975464 | -0.831986951 | 0.179389858 | -4.637870602 | 3.52E-06 | 4.54E-05 |
| GUCY1B3 | 12.50143858 | 2.253245986 | 0.485852749 | 4.637713777 | 3.52E-06 | 4.54E-05 |
| EEF1G | 7651.158325 | -0.349675433 | 0.075421418 | -4.636288239 | 3.55E-06 | 4.57E-05 |
| PILRA | 2429.928353 | 0.419230503 | 0.090441898 | 4.635357186 | 3.56E-06 | 4.59E-05 |
| CLEC7A | 227.2161288 | -0.39396875 | 0.085011869 | -4.634279365 | 3.58E-06 | 4.61E-05 |
| HLX | 193.4224904 | 0.475122465 | 0.102526147 | 4.634158992 | 3.58E-06 | 4.61E-05 |
| AGTPBP1 | 148.9822232 | -0.49864667 | 0.107611934 | -4.633748801 | 3.59E-06 | 4.61E-05 |
| ACN9 | 93.33904608 | -0.416503703 | 0.089921749 | -4.631846126 | 3.62E-06 | 4.65E-05 |

| | | | | | | |
|---|---|---|---|---|---|---|
| RAPGEF2 | 145.2498641 | 0.608306393 | 0.131338525 | 4.631591468 | 3.63E-06 | 4.65E-05 |
| TIRAP | 43.69314438 | 0.591181141 | 0.127684568 | 4.630012454 | 3.66E-06 | 4.69E-05 |
| SCN1B | 29.27962804 | 1.818998827 | 0.393350287 | 4.624373965 | 3.76E-06 | 4.81E-05 |
| MMD | 651.8312036 | -0.548972615 | 0.118880415 | -4.61785581 | 3.88E-06 | 4.96E-05 |
| LILRA1 | 21.33630586 | -1.367166529 | 0.296124989 | -4.616856325 | 3.90E-06 | 4.98E-05 |
| C6orf228 | 151.3590977 | 0.402428727 | 0.08724668 | 4.612539134 | 3.98E-06 | 5.08E-05 |
| SLC31A2 | 93.89329248 | 0.419504436 | 0.090960453 | 4.611943125 | 3.99E-06 | 5.09E-05 |
| FAM135A | 141.6503818 | -0.42366962 | 0.091871214 | -4.611560055 | 4.00E-06 | 5.10E-05 |
| OTUD1 | 541.5400358 | 0.616427257 | 0.133678828 | 4.611255682 | 4.00E-06 | 5.10E-05 |
| RCOR1 | 530.1543396 | -0.312187368 | 0.067746232 | -4.608187942 | 4.06E-06 | 5.18E-05 |
| GLS | 2416.167976 | 0.364919937 | 0.079230147 | 4.605821768 | 4.11E-06 | 5.23E-05 |
| CD300A | 721.1459133 | -0.599578418 | 0.130240926 | -4.603609893 | 4.15E-06 | 5.28E-05 |
| CD83 | 4697.583219 | 0.525923364 | 0.114243747 | 4.6035199 | 4.15E-06 | 5.28E-05 |
| PDPN | 13.12130084 | 1.277981636 | 0.277599519 | 4.603688226 | 4.15E-06 | 5.28E-05 |
| DPY19L1 | 510.4446697 | -0.358422482 | 0.077875112 | -4.602529284 | 4.17E-06 | 5.30E-05 |
| CTSL1 | 17609.60167 | 0.34537967 | 0.075061202 | 4.601307459 | 4.20E-06 | 5.33E-05 |
| TMED5 | 2532.765092 | 0.315470671 | 0.068592062 | 4.599229996 | 4.24E-06 | 5.38E-05 |
| ANAPC1 | 215.5148764 | -0.327419952 | 0.071202 | -4.59846564 | 4.26E-06 | 5.39E-05 |
| CEACAM21 | 124.1224446 | -0.616685386 | 0.134104403 | -4.598546912 | 4.25E-06 | 5.39E-05 |
| SHKBP1 | 1237.59177 | 0.297725538 | 0.064752927 | 4.597869958 | 4.27E-06 | 5.40E-05 |
| NDRG4 | 10.0044457 | 1.479753758 | 0.321859375 | 4.597516403 | 4.28E-06 | 5.41E-05 |
| ADPRHL2 | 191.5154113 | 0.447522001 | 0.097367153 | 4.596231757 | 4.30E-06 | 5.44E-05 |
| DYX1C1 | 114.2627162 | -0.788918908 | 0.171751733 | -4.593367971 | 4.36E-06 | 5.51E-05 |
| PLCD1 | 183.2375469 | 0.586238179 | 0.12767327 | 4.591706475 | 4.40E-06 | 5.55E-05 |
| DAGLB | 144.6888857 | -0.45337055 | 0.098759781 | -4.590639498 | 4.42E-06 | 5.57E-05 |
| LGMN | 4892.703659 | -0.537609571 | 0.117138278 | -4.58952939 | 4.44E-06 | 5.60E-05 |
| TBK1 | 781.3245463 | 0.49369818 | 0.107640642 | 4.586540666 | 4.51E-06 | 5.68E-05 |
| HMBOX1 | 167.4596508 | -0.423373489 | 0.092321199 | -4.585875119 | 4.52E-06 | 5.69E-05 |
| SHMT1 | 77.46120703 | -0.672612999 | 0.146677067 | -4.585672539 | 4.53E-06 | 5.69E-05 |
| ASPHD1 | 43.86706971 | 0.764178559 | 0.166675446 | 4.584829839 | 4.54E-06 | 5.71E-05 |
| C5orf15 | 851.7754586 | 0.480017936 | 0.10469954 | 4.584718694 | 4.55E-06 | 5.71E-05 |
| NCF4 | 1399.722749 | -0.59998719 | 0.130981094 | -4.580715965 | 4.63E-06 | 5.82E-05 |
| TNIP2 | 372.3447838 | 0.607096486 | 0.13256555 | 4.579594685 | 4.66E-06 | 5.85E-05 |
| INO80D | 253.6027826 | 0.45311201 | 0.098980013 | 4.577813204 | 4.70E-06 | 5.89E-05 |
| RNF19B | 862.4051581 | 1.092274568 | 0.238644736 | 4.576989992 | 4.72E-06 | 5.91E-05 |
| TSFM | 222.6273426 | 0.525314316 | 0.114821501 | 4.575051799 | 4.76E-06 | 5.96E-05 |
| POC1B | 168.8725805 | 0.424489037 | 0.092814925 | 4.573499749 | 4.80E-06 | 6.00E-05 |
| ADRM1 | 1577.180817 | 0.442901773 | 0.096868838 | 4.572180098 | 4.83E-06 | 6.03E-05 |
| CKAP2 | 51.06617808 | -0.734570965 | 0.160660004 | -4.57220807 | 4.83E-06 | 6.03E-05 |
| ENPP5 | 19.82031784 | -1.461718484 | 0.319709923 | -4.572014756 | 4.83E-06 | 6.03E-05 |
| HAGHL | 116.7250126 | -0.342633992 | 0.074975362 | -4.569954479 | 4.88E-06 | 6.09E-05 |
| CNST | 120.3801834 | -0.28345184 | 0.062044572 | -4.568519522 | 4.91E-06 | 6.13E-05 |
| SLC16A6 | 472.0147792 | 0.723210701 | 0.158322697 | 4.567953383 | 4.93E-06 | 6.14E-05 |
| MS4A4A | 3457.851855 | -0.383331221 | 0.083943238 | -4.566552695 | 4.96E-06 | 6.18E-05 |
| PPHLN1 | 112.5738526 | -0.352420493 | 0.077227805 | -4.56338872 | 5.03E-06 | 6.27E-05 |
| SP140 | 45.26530284 | 0.940463248 | 0.206217077 | 4.560549784 | 5.10E-06 | 6.35E-05 |
| CHURC1 | 11.58337343 | 1.053068119 | 0.23091868 | 4.560341838 | 5.11E-06 | 6.35E-05 |

| | | | | | | |
|---|---|---|---|---|---|---|
| C17orf89 | 87.48225223 | -0.589260757 | 0.12922698 | -4.559889553 | 5.12E-06 | 6.36E-05 |
| RIN2 | 546.8484787 | 0.570080676 | 0.125038142 | 4.559254227 | 5.13E-06 | 6.37E-05 |
| BTN2A2 | 296.4211096 | 0.257730943 | 0.056540962 | 4.55830489 | 5.16E-06 | 6.38E-05 |
| CRYZ | 276.1465336 | -0.46305174 | 0.10158116 | -4.558441148 | 5.15E-06 | 6.38E-05 |
| IGSF6 | 2503.782937 | 0.553665756 | 0.121459701 | 4.558431752 | 5.15E-06 | 6.38E-05 |
| PCMTD2 | 221.1388279 | -0.416570293 | 0.09137911 | -4.558703784 | 5.15E-06 | 6.38E-05 |
| P2RY11 | 43.72333518 | -0.994917013 | 0.218323529 | -4.557076463 | 5.19E-06 | 6.42E-05 |
| CRISPLD2 | 30.79144744 | 2.32931069 | 0.511348998 | 4.555226858 | 5.23E-06 | 6.47E-05 |
| RILPL2 | 601.1748572 | 0.420273687 | 0.092273843 | 4.554635136 | 5.25E-06 | 6.48E-05 |
| SLC25A6 | 2228.512262 | -0.297928733 | 0.065416661 | -4.554324997 | 5.26E-06 | 6.49E-05 |
| AICDA | 27.16045542 | -1.150947529 | 0.252822423 | -4.552394975 | 5.30E-06 | 6.55E-05 |
| TCTEX1D4 | 68.89675351 | 1.16723285 | 0.256520296 | 4.550255353 | 5.36E-06 | 6.61E-05 |
| MANSC4 | 10.20399329 | -2.652593698 | 0.583202525 | -4.548323409 | 5.41E-06 | 6.66E-05 |
| RPSAP58 | 24.93514599 | -0.745905536 | 0.164138405 | -4.544369339 | 5.51E-06 | 6.79E-05 |
| HIATL1 | 2148.754979 | 0.280144897 | 0.061670793 | 4.542586299 | 5.56E-06 | 6.84E-05 |
| CYB5R1 | 1052.676224 | -0.259830266 | 0.057261474 | -4.537610474 | 5.69E-06 | 7.00E-05 |
| CSF2RA | 610.5701357 | 0.44989022 | 0.099157521 | 4.537126524 | 5.70E-06 | 7.01E-05 |
| INPP4A | 354.9103085 | -0.34859563 | 0.076881301 | -4.53420564 | 5.78E-06 | 7.10E-05 |
| ZBED3 | 25.27325387 | -0.888893456 | 0.196089973 | -4.533089804 | 5.81E-06 | 7.14E-05 |
| SEMA4A | 768.3067274 | 0.817476418 | 0.180348362 | 4.532763199 | 5.82E-06 | 7.14E-05 |
| RAC2 | 7787.713528 | 0.668628906 | 0.147559272 | 4.531256479 | 5.86E-06 | 7.19E-05 |
| TPM4 | 6347.77636 | 0.533324411 | 0.117703788 | 4.531072622 | 5.87E-06 | 7.19E-05 |
| ACPL2 | 15.1168817 | 1.171115073 | 0.258474215 | 4.530877766 | 5.87E-06 | 7.19E-05 |
| LPCAT2 | 1473.661586 | -0.357349571 | 0.078917045 | -4.528167136 | 5.95E-06 | 7.28E-05 |
| ANKRD5 | 7.463124656 | -1.59316284 | 0.352011273 | -4.525885856 | 6.01E-06 | 7.35E-05 |
| AKAP5 | 47.37777989 | 1.156687701 | 0.255631563 | 4.524823493 | 6.04E-06 | 7.39E-05 |
| SLC40A1 | 5.872546046 | -1.890119048 | 0.417808947 | -4.523883607 | 6.07E-06 | 7.41E-05 |
| DPYSL3 | 188.2277389 | 0.762531709 | 0.168658748 | 4.521151257 | 6.15E-06 | 7.51E-05 |
| TIMP3 | 478.5821816 | -0.680568934 | 0.150535779 | -4.52097794 | 6.16E-06 | 7.51E-05 |
| NFKBIB | 659.8030315 | 0.449841069 | 0.099529455 | 4.519677808 | 6.19E-06 | 7.55E-05 |
| TRIP4 | 379.5869735 | 0.360531117 | 0.079781357 | 4.518989509 | 6.21E-06 | 7.57E-05 |
| OGDH | 1776.975955 | 0.437588889 | 0.096860474 | 4.517724024 | 6.25E-06 | 7.61E-05 |
| ARHGEF2 | 1036.720614 | 0.328399502 | 0.07273505 | 4.515010333 | 6.33E-06 | 7.70E-05 |
| CD1D | 12.18541146 | -1.639743515 | 0.363204029 | -4.514662237 | 6.34E-06 | 7.71E-05 |
| TMEM217 | 5.497673022 | 2.224236183 | 0.493289815 | 4.508984611 | 6.51E-06 | 7.91E-05 |
| CHI3L2 | 18.51869959 | 1.357776118 | 0.301301865 | 4.506364802 | 6.59E-06 | 8.01E-05 |
| KCNQ1 | 255.4976857 | -0.904247982 | 0.200681203 | -4.505892779 | 6.61E-06 | 8.02E-05 |
| MDFIC | 1338.214492 | -0.29471989 | 0.065426584 | -4.504589283 | 6.65E-06 | 8.06E-05 |
| POLR2J | 511.7692941 | -0.275115822 | 0.061153124 | -4.498802446 | 6.83E-06 | 8.28E-05 |
| HIVEP2 | 68.39776808 | 0.529282246 | 0.117662785 | 4.498297805 | 6.85E-06 | 8.29E-05 |
| MRC1 | 5904.185002 | 0.628668177 | 0.139781673 | 4.497500726 | 6.88E-06 | 8.32E-05 |
| DGKH | 191.7525156 | 0.450251283 | 0.100202272 | 4.4934239 | 7.01E-06 | 8.48E-05 |
| TXNRD1 | 5.295821806 | 1.803490191 | 0.401442818 | 4.492520758 | 7.04E-06 | 8.50E-05 |
| YPEL3 | 297.0092714 | -0.492982369 | 0.109732789 | -4.492571228 | 7.04E-06 | 8.50E-05 |
| TIGIT | 7.810376338 | 1.654427984 | 0.368284749 | 4.492252228 | 7.05E-06 | 8.51E-05 |
| AKAP12 | 13.67761417 | 1.388981475 | 0.309308915 | 4.490596323 | 7.10E-06 | 8.57E-05 |
| GYPC | 1358.511294 | 0.427832398 | 0.095309375 | 4.488880555 | 7.16E-06 | 8.63E-05 |

| | | | | | | |
|---|---|---|---|---|---|---|
| FUNDC2 | 207.2393871 | 0.389336722 | 0.086753623 | 4.487843935 | 7.19E-06 | 8.67E-05 |
| ARRB1 | 712.3795833 | -0.42569621 | 0.094867641 | -4.487264622 | 7.21E-06 | 8.68E-05 |
| SYNDIG1 | 9.234223299 | -3.37127089 | 0.751613194 | -4.485380136 | 7.28E-06 | 8.76E-05 |
| MPV17L | 105.5361393 | -0.784145724 | 0.174865137 | -4.484288515 | 7.32E-06 | 8.79E-05 |
| RBP4 | 3117.792944 | -0.527752836 | 0.117893904 | -4.476506561 | 7.59E-06 | 9.11E-05 |
| TACC3 | 152.1999433 | -0.380468817 | 0.08499245 | -4.476501329 | 7.59E-06 | 9.11E-05 |
| ACBD6 | 133.1054197 | -0.527442565 | 0.117893314 | -4.473897183 | 7.68E-06 | 9.22E-05 |
| MYO1B | 11.15821587 | 1.313846837 | 0.293688251 | 4.473610474 | 7.69E-06 | 9.22E-05 |
| MOB3B | 233.8614117 | 0.741402465 | 0.165744611 | 4.473161812 | 7.71E-06 | 9.24E-05 |
| CTSA | 8392.921502 | -0.412761385 | 0.092288027 | -4.472534527 | 7.73E-06 | 9.26E-05 |
| LMNB1 | 60.68315101 | 0.903408817 | 0.202019093 | 4.471898185 | 7.75E-06 | 9.27E-05 |
| SRC | 693.7387687 | 0.355513329 | 0.079498883 | 4.471928594 | 7.75E-06 | 9.27E-05 |
| CCL13 | 4.008231947 | 1.840013505 | 0.411593583 | 4.470462081 | 7.81E-06 | 9.33E-05 |
| ZC3HAV1 | 1144.88529 | 0.483462574 | 0.108155356 | 4.47007519 | 7.82E-06 | 9.34E-05 |
| ADCY3 | 1720.538147 | -0.339950043 | 0.076061807 | -4.469392145 | 7.84E-06 | 9.36E-05 |
| INPP5K | 655.8156327 | 0.243365876 | 0.054450404 | 4.46949621 | 7.84E-06 | 9.36E-05 |
| ENO1 | 13911.8223 | 0.192223729 | 0.043040434 | 4.466119667 | 7.97E-06 | 9.50E-05 |
| FHIT | 87.13099302 | -0.599475689 | 0.134266177 | -4.464830237 | 8.01E-06 | 9.55E-05 |
| KIAA1586 | 86.65252505 | -0.565253461 | 0.126867177 | -4.455474419 | 8.37E-06 | 9.97E-05 |
| FTSJ2 | 330.0796178 | -0.377466821 | 0.084726929 | -4.455098578 | 8.39E-06 | 9.98E-05 |
| PSD3 | 25.6385483 | 1.081001441 | 0.242777207 | 4.452647993 | 8.48E-06 | 0.000100861 |
| SEMA7A | 363.7254834 | 0.610563937 | 0.137261534 | 4.44817947 | 8.66E-06 | 0.000102916 |
| MYLIP | 10.78130016 | -1.119485697 | 0.251734865 | -4.447082437 | 8.70E-06 | 0.000103377 |
| KLF5 | 35.30714229 | 0.786880894 | 0.176964138 | 4.446555688 | 8.73E-06 | 0.000103565 |
| LGALS2 | 94.6085907 | 1.087683759 | 0.244680639 | 4.445320094 | 8.78E-06 | 0.000104095 |
| LOC28299[illegible] | 61.80573052 | -0.658826783 | 0.148272372 | -4.443354978 | 8.86E-06 | 0.000104984 |
| JPH4 | 11.31263684 | -1.541713496 | 0.34709384 | -4.44177717 | 8.92E-06 | 0.00010569 |
| NDUFB3 | 50.68753101 | 0.825565521 | 0.185953533 | 4.439633416 | 9.01E-06 | 0.000106681 |
| SLC35E2B | 237.7140893 | -0.294487348 | 0.066362838 | -4.437533972 | 9.10E-06 | 0.000107658 |
| TIFAB | 42.1498536 | -1.016747432 | 0.229215614 | -4.435768631 | 9.17E-06 | 0.000108475 |
| RNF24 | 61.41467527 | 0.56512772 | 0.127409761 | 4.435513536 | 9.19E-06 | 0.000108535 |
| FN3K | 69.13938541 | -0.630503356 | 0.142156198 | -4.435285742 | 9.20E-06 | 0.000108581 |
| BRCA2 | 53.38722163 | 0.683210236 | 0.154064825 | 4.434563416 | 9.23E-06 | 0.000108876 |
| IL10 | 63.77972544 | -0.981555913 | 0.221530188 | -4.430799803 | 9.39E-06 | 0.000110724 |
| PGAM1 | 3517.713381 | 0.250872349 | 0.056628266 | 4.430161235 | 9.42E-06 | 0.000110982 |
| SMIM3 | 196.7669469 | 0.676717587 | 0.15279124 | 4.429033939 | 9.47E-06 | 0.000111493 |
| FLJ45340 | 568.8630695 | 0.308874468 | 0.069757937 | 4.427803951 | 9.52E-06 | 0.00011206 |
| FOLR3 | 94.51674392 | -1.02299549 | 0.231192433 | -4.424865798 | 9.65E-06 | 0.000113524 |
| FXN | 134.0324802 | -0.362020927 | 0.081818288 | -4.424694476 | 9.66E-06 | 0.000113543 |
| RPL10 | 5779.547635 | -0.419668465 | 0.094866721 | -4.423769068 | 9.70E-06 | 0.000113958 |
| DBNDD2 | 1346.826637 | -0.285107611 | 0.064479221 | -4.421697503 | 9.79E-06 | 0.000114984 |
| TOMM20 | 1183.679735 | -0.272995267 | 0.061748864 | -4.421057321 | 9.82E-06 | 0.000115253 |
| RRP1 | 368.3618837 | 0.298452356 | 0.067521776 | 4.420090427 | 9.87E-06 | 0.000115697 |
| P2RX7 | 1692.801522 | 0.470048225 | 0.106448784 | 4.415721901 | 1.01E-05 | 0.000117984 |
| CRELD1 | 246.50306 | 0.348575885 | 0.078942387 | 4.41557318 | 1.01E-05 | 0.000117991 |
| MYEOV2 | 1332.410854 | -0.268721164 | 0.060859518 | -4.41543365 | 1.01E-05 | 0.000117993 |
| ITGA1 | 36.54512628 | 0.764187745 | 0.173091919 | 4.414924463 | 1.01E-05 | 0.000118197 |

| | | | | | | |
|---|---|---|---|---|---|---|
| ARHGEF6 | 93.57271803 | -0.565885068 | 0.128181383 | -4.414721181 | 1.01E-05 | 0.000118234 |
| PRSS21 | 293.7092083 | -0.404005866 | 0.091528353 | -4.413996887 | 1.01E-05 | 0.000118556 |
| TMEM192 | 386.6658503 | -0.447385504 | 0.101360508 | -4.413804859 | 1.02E-05 | 0.000118587 |
| HADHA | 2949.273527 | -0.203539394 | 0.046134022 | -4.411915186 | 1.02E-05 | 0.000119552 |
| MAF | 89.49689612 | -0.406047347 | 0.09204223 | -4.411533127 | 1.03E-05 | 0.000119688 |
| ADAMTS6 | 36.25606369 | -0.652415717 | 0.147915432 | -4.410734634 | 1.03E-05 | 0.00011998 |
| TGIF2 | 102.2311093 | 0.599938936 | 0.136015903 | 4.410799919 | 1.03E-05 | 0.00011998 |
| NREP | 25.85517768 | -0.77927666 | 0.176695804 | -4.410272597 | 1.03E-05 | 0.000120161 |
| DNAJC1 | 462.5229212 | 0.40027361 | 0.090806655 | 4.407976611 | 1.04E-05 | 0.000121366 |
| NDUFS3 | 2025.080402 | 0.237043167 | 0.053783241 | 4.407379749 | 1.05E-05 | 0.000121625 |
| STX4 | 2858.68785 | 0.400671561 | 0.09093608 | 4.406079085 | 1.05E-05 | 0.000122281 |
| SGSM2 | 357.3236155 | -0.392784296 | 0.089205504 | -4.403139691 | 1.07E-05 | 0.000123873 |
| TNK2 | 625.2017723 | 0.468483087 | 0.106405605 | 4.402804606 | 1.07E-05 | 0.000123987 |
| TARBP1 | 100.3807476 | -0.52379139 | 0.118978223 | -4.402413952 | 1.07E-05 | 0.000124134 |
| CD72 | 124.3837117 | -0.743764268 | 0.168966112 | -4.401854661 | 1.07E-05 | 0.000124377 |
| IL6ST | 432.049589 | 0.388806889 | 0.088340363 | 4.40123715 | 1.08E-05 | 0.000124654 |
| ZFYVE26 | 3847.074477 | -0.213888569 | 0.048610376 | -4.400059944 | 1.08E-05 | 0.000125254 |
| ICOS | 53.10444988 | 0.867542944 | 0.197200764 | 4.399287956 | 1.09E-05 | 0.000125622 |
| NELF | 192.8924863 | -0.464749058 | 0.105724495 | -4.39585037 | 1.10E-05 | 0.000127548 |
| TDRKH | 46.1134518 | -0.628803513 | 0.143051969 | -4.395629896 | 1.10E-05 | 0.000127598 |
| SLC29A1 | 744.2242263 | -0.344137926 | 0.078321079 | -4.393937489 | 1.11E-05 | 0.000128437 |
| SLFN13 | 5.755124169 | 1.742129808 | 0.396477466 | 4.394019727 | 1.11E-05 | 0.000128437 |
| TDG | 420.7836308 | 0.301262065 | 0.06856838 | 4.393600451 | 1.11E-05 | 0.000128557 |
| SGK110 | 10.9827916 | 1.595884004 | 0.3632528 | 4.393315085 | 1.12E-05 | 0.000128646 |
| THOC5 | 433.9234437 | 0.343226947 | 0.078145908 | 4.392129507 | 1.12E-05 | 0.00012927 |
| CHST14 | 70.67513567 | -0.786520541 | 0.179206433 | -4.388907957 | 1.14E-05 | 0.000131038 |
| RNF130 | 10068.0002 | -0.276311976 | 0.062955972 | -4.388971625 | 1.14E-05 | 0.000131038 |
| SIGLEC9 | 909.9581326 | -0.360745485 | 0.082212613 | -4.3879579 | 1.14E-05 | 0.00013153 |
| PTTG1 | 18.53068954 | 0.874528649 | 0.199330536 | 4.387329033 | 1.15E-05 | 0.000131829 |
| PSMB2 | 6363.209153 | 0.503451517 | 0.11475997 | 4.386995894 | 1.15E-05 | 0.00013195 |
| ZC3HAV1L | 10.47094612 | -0.953182445 | 0.217309484 | -4.386290139 | 1.15E-05 | 0.000132297 |
| PHLDB1 | 15.49927025 | 1.690808529 | 0.385670777 | 4.384072199 | 1.16E-05 | 0.00013357 |
| LOC28314[illegible] | 13.95452297 | -1.146376756 | 0.261573152 | -4.382623935 | 1.17E-05 | 0.000134378 |
| GPR56 | 13.02908584 | 1.839843945 | 0.419833769 | 4.382315291 | 1.17E-05 | 0.000134486 |
| CCDC94 | 207.268796 | 0.412578083 | 0.094168219 | 4.381287944 | 1.18E-05 | 0.000135039 |
| PSRC1 | 3.835092832 | -2.004797205 | 0.457693559 | -4.380217209 | 1.19E-05 | 0.000135621 |
| PIP4K2A | 579.3965022 | -0.301031468 | 0.068756795 | -4.378206799 | 1.20E-05 | 0.000136794 |
| FAM222B | 418.6726357 | -0.306857668 | 0.070175047 | -4.372746158 | 1.23E-05 | 0.000140176 |
| RFFL | 785.2813794 | 0.39011667 | 0.089287235 | 4.369232275 | 1.25E-05 | 0.000142363 |
| NCOA7 | 308.3642932 | -0.478106382 | 0.109470268 | -4.367454187 | 1.26E-05 | 0.000143439 |
| BAZ2A | 741.4518493 | 0.37206124 | 0.085201932 | 4.366816956 | 1.26E-05 | 0.00014377 |
| LOC10012[illegible] | 15.92667503 | -0.776313641 | 0.177805327 | -4.36608765 | 1.26E-05 | 0.000144162 |
| PTPRE | 1662.645669 | 0.391643454 | 0.089721662 | 4.365093616 | 1.27E-05 | 0.000144721 |
| ZFYVE21 | 182.3565697 | -0.289279685 | 0.06627294 | -4.36497436 | 1.27E-05 | 0.000144721 |
| TMSB4X | 51521.7555 | -0.378717205 | 0.086832022 | -4.361492435 | 1.29E-05 | 0.000146954 |
| SEMA6B | 2451.014442 | -0.445660358 | 0.102240702 | -4.358932872 | 1.31E-05 | 0.000148592 |
| ALDH1A2 | 336.4386181 | 0.832855925 | 0.191162355 | 4.356798826 | 1.32E-05 | 0.000149865 |

| | | | | | | |
|---|---|---|---|---|---|---|
| AP1S2 | 713.8187611 | -0.503370253 | 0.115535696 | -4.356837519 | 1.32E-05 | 0.000149865 |
| UBP1 | 514.6493301 | 0.380445701 | 0.087390206 | 4.353413503 | 1.34E-05 | 0.000152107 |
| USO1 | 440.868944 | 0.336184174 | 0.077242942 | 4.352296363 | 1.35E-05 | 0.000152791 |
| SHROOM3 | 293.8139205 | 1.206688676 | 0.277390424 | 4.350145391 | 1.36E-05 | 0.000154203 |
| RPL13AP2( | 986.1570411 | -0.564953468 | 0.129945812 | -4.347608135 | 1.38E-05 | 0.000155903 |
| POLR2G | 471.259236 | -0.215149036 | 0.049498746 | -4.346555308 | 1.38E-05 | 0.000156473 |
| TCF4 | 11.13650351 | -1.050779923 | 0.241750849 | -4.346540783 | 1.38E-05 | 0.000156473 |
| RPL27 | 5800.363754 | -0.334800854 | 0.077070959 | -4.344059808 | 1.40E-05 | 0.000158155 |
| TMEM223 | 138.0503995 | -0.565353156 | 0.130150431 | -4.343843907 | 1.40E-05 | 0.000158215 |
| MCOLN1 | 2116.78703 | -0.327057318 | 0.075313849 | -4.342592 | 1.41E-05 | 0.000158927 |
| ZCCHC11 | 202.5962085 | -0.444162816 | 0.102278911 | -4.342662742 | 1.41E-05 | 0.000158927 |
| CD36 | 2253.016945 | -0.817862229 | 0.188342319 | -4.342424131 | 1.41E-05 | 0.000158952 |
| ATP6V1A | 1848.408576 | 0.320347313 | 0.073782924 | 4.341754106 | 1.41E-05 | 0.000159341 |
| FAM159A | 5.368678563 | -2.291228865 | 0.527845296 | -4.34072044 | 1.42E-05 | 0.000159996 |
| KRR1 | 391.596641 | 0.346586475 | 0.079904865 | 4.337489002 | 1.44E-05 | 0.000162072 |
| LOC14688( | 247.5815942 | -0.443102497 | 0.102151171 | -4.337713324 | 1.44E-05 | 0.000162072 |
| TMEM173 | 93.8523385 | 1.022000885 | 0.235613481 | 4.337616346 | 1.44E-05 | 0.000162072 |
| RASSF5 | 732.2720149 | 0.270294723 | 0.062348271 | 4.335240085 | 1.46E-05 | 0.000163639 |
| ARID3B | 151.0126963 | 0.503264506 | 0.116124201 | 4.333846894 | 1.47E-05 | 0.000164579 |
| TMEM220 | 126.7211771 | -0.405234171 | 0.093520488 | -4.333105848 | 1.47E-05 | 0.000165035 |
| ADRB2 | 24.05894857 | -1.119353595 | 0.258355851 | -4.332604001 | 1.47E-05 | 0.000165312 |
| PARP3 | 249.3573938 | 0.345165574 | 0.079718978 | 4.329779198 | 1.49E-05 | 0.000167346 |
| CCDC112 | 78.8568645 | -0.638747112 | 0.147639024 | -4.326411091 | 1.52E-05 | 0.000169747 |
| HSD17B4 | 4232.775895 | -0.280622043 | 0.064863077 | -4.326375777 | 1.52E-05 | 0.000169747 |
| SWAP70 | 1352.985912 | -0.420339866 | 0.097321682 | -4.319077272 | 1.57E-05 | 0.000175353 |
| TEAD4 | 3.662726822 | 1.983495761 | 0.459274541 | 4.318758359 | 1.57E-05 | 0.000175501 |
| YAP1 | 7.512175436 | 1.484066961 | 0.343763432 | 4.317117008 | 1.58E-05 | 0.000176705 |
| AKAP11 | 354.7832807 | -0.446480685 | 0.103498994 | -4.31386496 | 1.60E-05 | 0.000179218 |
| GGA3 | 512.6548972 | -0.307710831 | 0.071339591 | -4.313324839 | 1.61E-05 | 0.000179549 |
| KIAA1958 | 62.85234356 | 0.557813026 | 0.129340223 | 4.31275758 | 1.61E-05 | 0.000179814 |
| PHKA2 | 147.5490788 | -0.470425067 | 0.109078152 | -4.312734097 | 1.61E-05 | 0.000179814 |
| RNF146 | 227.009101 | -0.332788235 | 0.077170562 | -4.312372858 | 1.62E-05 | 0.000180001 |
| UBE3D | 47.52379897 | -0.541718706 | 0.125768387 | -4.307272433 | 1.65E-05 | 0.00018409 |
| CEACAM3 | 12.78926738 | -1.213450349 | 0.281741934 | -4.306956843 | 1.66E-05 | 0.000184243 |
| SNHG5 | 516.0267324 | -0.45270024 | 0.105214836 | -4.302627427 | 1.69E-05 | 0.000187769 |
| SPOCD1 | 1322.377528 | 0.570686296 | 0.132641109 | 4.302484366 | 1.69E-05 | 0.000187778 |
| KBTBD8 | 77.24816529 | 0.52756474 | 0.122671862 | 4.300617358 | 1.70E-05 | 0.000189254 |
| DBP | 57.25152809 | -1.038771757 | 0.241563573 | -4.300200329 | 1.71E-05 | 0.000189498 |
| ARL5B | 243.3733424 | 0.4760646 | 0.110829174 | 4.295480893 | 1.74E-05 | 0.000193229 |
| STAP1 | 21.63068946 | 1.813609251 | 0.422194189 | 4.295675546 | 1.74E-05 | 0.000193229 |
| SUMF1 | 519.2014573 | -0.293426875 | 0.068309415 | -4.295555348 | 1.74E-05 | 0.000193229 |
| SCARB1 | 1715.826207 | -0.600955892 | 0.139939449 | -4.294399436 | 1.75E-05 | 0.000194058 |
| CTNNBL1 | 796.8023641 | 0.525362231 | 0.122356556 | 4.29369908 | 1.76E-05 | 0.000194556 |
| DCAF11 | 699.106913 | 0.261940052 | 0.061019496 | 4.292727247 | 1.76E-05 | 0.000195294 |
| ACOX3 | 163.7570344 | 0.374336959 | 0.087252341 | 4.290279833 | 1.78E-05 | 0.000197341 |
| CA11 | 76.43763705 | 0.521893683 | 0.121660922 | 4.289739679 | 1.79E-05 | 0.000197609 |
| DCP1A | 654.5176034 | 0.359943631 | 0.083908504 | 4.289715729 | 1.79E-05 | 0.000197609 |

| | | | | | | |
|---|---|---|---|---|---|---|
| METTL21A | 139.4893383 | -0.467383963 | 0.108959258 | -4.289529612 | 1.79E-05 | 0.000197657 |
| IGBP1 | 567.7609004 | -0.422799236 | 0.098583013 | -4.288763565 | 1.80E-05 | 0.000198223 |
| ORAI3 | 48.3800975 | -0.544625388 | 0.127000928 | -4.288357557 | 1.80E-05 | 0.000198452 |
| SIK3 | 653.710079 | 0.602403213 | 0.140477822 | 4.288244257 | 1.80E-05 | 0.000198452 |
| GSTO1 | 660.3734022 | 0.302427091 | 0.070530487 | 4.287891701 | 1.80E-05 | 0.000198649 |
| TBC1D4 | 87.33489079 | -0.887986472 | 0.207117902 | -4.287347764 | 1.81E-05 | 0.000199018 |
| MFSD12 | 184.7061733 | 0.697907455 | 0.162788593 | 4.287201224 | 1.81E-05 | 0.000199032 |
| CTPS2 | 165.059182 | -0.484943118 | 0.113148379 | -4.28590424 | 1.82E-05 | 0.000200079 |
| EFTUD1 | 561.2344719 | 0.639431988 | 0.149205649 | 4.28557494 | 1.82E-05 | 0.000200257 |
| PPP2R4 | 1303.638213 | 0.459010899 | 0.107132291 | 4.284524252 | 1.83E-05 | 0.000201088 |
| LAMB2 | 197.0922907 | -0.386806545 | 0.090291617 | -4.283969653 | 1.84E-05 | 0.000201471 |
| PIAS2 | 550.1334326 | -0.5164021 | 0.120593477 | -4.28217273 | 1.85E-05 | 0.000202986 |
| TTBK2 | 96.63836081 | -0.45634357 | 0.106606372 | -4.280640668 | 1.86E-05 | 0.000204268 |
| CD82 | 1548.983977 | 0.713130921 | 0.166617251 | 4.280054546 | 1.87E-05 | 0.000204686 |
| SPTLC3 | 10.71311239 | -2.208348521 | 0.516027208 | -4.279519543 | 1.87E-05 | 0.000205059 |
| STXBP5-AS | 24.70655765 | 1.298547523 | 0.303571303 | 4.277570074 | 1.89E-05 | 0.000206741 |
| ASTE1 | 89.85437911 | -0.616672615 | 0.144302038 | -4.273485138 | 1.92E-05 | 0.000210443 |
| CDKN2A | 17.27412097 | 1.431006437 | 0.334878326 | 4.273213067 | 1.93E-05 | 0.000210576 |
| SLC37A3 | 16.35747096 | 0.824019275 | 0.192907383 | 4.271579777 | 1.94E-05 | 0.000212001 |
| PIK3CB | 1751.016361 | 0.324190812 | 0.075908848 | 4.270790818 | 1.95E-05 | 0.000212627 |
| GPR155 | 52.69999724 | -0.762466389 | 0.178564797 | -4.269970347 | 1.95E-05 | 0.000213286 |
| C12orf57 | 270.7179587 | 0.381344596 | 0.08931684 | 4.26957109 | 1.96E-05 | 0.000213543 |
| PET117 | 117.0606464 | -0.31399235 | 0.073552355 | -4.268963901 | 1.96E-05 | 0.000214 |
| ZNF514 | 120.0832404 | -0.528785492 | 0.1238879 | -4.268257753 | 1.97E-05 | 0.000214553 |
| IL2RB | 64.21636887 | 1.128486349 | 0.264463533 | 4.267077336 | 1.98E-05 | 0.000215565 |
| AOC2 | 16.20907959 | 0.826581992 | 0.193720548 | 4.266878238 | 1.98E-05 | 0.000215631 |
| ARHGAP35 | 479.1554114 | -0.361122629 | 0.084644632 | -4.266338214 | 1.99E-05 | 0.000216028 |
| ZNF280D | 107.0535983 | -0.584318799 | 0.13702763 | -4.264240707 | 2.01E-05 | 0.00021794 |
| MICAL1 | 2235.03912 | -0.412953383 | 0.096891419 | -4.262022259 | 2.03E-05 | 0.000219987 |
| AAMP | 552.8004558 | 0.442570666 | 0.103930212 | 4.258344669 | 2.06E-05 | 0.000223506 |
| SLC1A3 | 1703.103549 | 0.487320052 | 0.114446401 | 4.258063562 | 2.06E-05 | 0.000223657 |
| GPR132 | 25.46701289 | 1.176572192 | 0.276342629 | 4.257657224 | 2.07E-05 | 0.000223934 |
| CCNK | 1193.699206 | 0.370422845 | 0.087013907 | 4.257053355 | 2.07E-05 | 0.000224409 |
| KIDINS220 | 1137.012704 | -0.242411368 | 0.056948542 | -4.256673842 | 2.07E-05 | 0.000224659 |
| SAV1 | 524.1159948 | 0.451328193 | 0.106055707 | 4.255576674 | 2.09E-05 | 0.000225633 |
| RHOC | 2697.74864 | 0.571562287 | 0.13432885 | 4.254948111 | 2.09E-05 | 0.000226136 |
| LINC00622 | 114.1466068 | 0.781442877 | 0.183733284 | 4.253137259 | 2.11E-05 | 0.000227841 |
| LRG1 | 16.91715093 | -1.054083343 | 0.24793491 | -4.251451894 | 2.12E-05 | 0.00022943 |
| TAPBP | 64.36823896 | 0.723193009 | 0.170113589 | 4.251235981 | 2.13E-05 | 0.000229518 |
| PBLD | 19.25407473 | -0.793917858 | 0.186770646 | -4.250763572 | 2.13E-05 | 0.000229869 |
| FAM117B | 21.4925727 | -0.965914165 | 0.227247704 | -4.250490316 | 2.13E-05 | 0.000230017 |
| SLC26A11 | 513.9008153 | -0.374278569 | 0.088066137 | -4.249971478 | 2.14E-05 | 0.000230417 |
| SIGIRR | 67.47477397 | -0.585369441 | 0.137751672 | -4.249454342 | 2.14E-05 | 0.000230816 |
| ASCL1 | 3.327215829 | 2.947371257 | 0.693812259 | 4.248081841 | 2.16E-05 | 0.000232012 |
| ZDHHC2 | 401.2945055 | -0.605318042 | 0.142493565 | -4.248037758 | 2.16E-05 | 0.000232012 |
| C10orf11 | 402.7484521 | -0.42213489 | 0.099394372 | -4.247070329 | 2.17E-05 | 0.000232881 |
| BTN2A1 | 479.5220499 | 0.388916486 | 0.091602528 | 4.245696015 | 2.18E-05 | 0.000234043 |

| | | | | | | |
|---|---|---|---|---|---|---|
| DUSP6 | 964.9277916 | 0.279273809 | 0.065777059 | 4.245763073 | 2.18E-05 | 0.000234043 |
| NECAB3 | 100.6457381 | -0.464354549 | 0.109457827 | -4.242314697 | 2.21E-05 | 0.000237461 |
| PARVG | 2834.742606 | -0.363501303 | 0.085713569 | -4.240883974 | 2.23E-05 | 0.000238769 |
| PPP2R3A | 193.2665595 | 0.454125724 | 0.107084304 | 4.240824352 | 2.23E-05 | 0.000238769 |
| LGALS3BP | 5228.175511 | 0.52604895 | 0.124079421 | 4.239614782 | 2.24E-05 | 0.000239921 |
| NKD1 | 4.682873345 | 1.800363359 | 0.424689515 | 4.239246072 | 2.24E-05 | 0.000240177 |
| TRIM35 | 55.34454568 | 0.83817663 | 0.19784567 | 4.236517442 | 2.27E-05 | 0.000242973 |
| LGALS1 | 4141.906386 | -0.235101992 | 0.055526527 | -4.234048198 | 2.30E-05 | 0.000245517 |
| TUBB6 | 1097.285469 | 0.369095167 | 0.087179314 | 4.233747069 | 2.30E-05 | 0.000245705 |
| CASP6 | 229.2677003 | -0.320847372 | 0.075901179 | -4.227172423 | 2.37E-05 | 0.000252703 |
| YRDC | 146.6389533 | 0.426145431 | 0.100809086 | 4.2272522 | 2.37E-05 | 0.000252703 |
| MAP3K1 | 869.4447403 | -0.598891482 | 0.141718238 | -4.225930915 | 2.38E-05 | 0.000253955 |
| TRIM33 | 215.1145674 | 0.438624156 | 0.103797419 | 4.225771322 | 2.38E-05 | 0.00025399 |
| SIPA1L1 | 222.4707284 | 0.754079404 | 0.178469949 | 4.225245822 | 2.39E-05 | 0.000254437 |
| CXorf23 | 19.08404943 | -0.72280384 | 0.171115136 | -4.224078925 | 2.40E-05 | 0.000255613 |
| ACER3 | 622.3874407 | -0.375826779 | 0.088997161 | -4.222907517 | 2.41E-05 | 0.000256799 |
| SLC41A2 | 287.3986765 | 0.801775815 | 0.190035525 | 4.219083863 | 2.45E-05 | 0.000261043 |
| CYSLTR1 | 68.93288786 | -1.120263535 | 0.265782033 | -4.214970901 | 2.50E-05 | 0.000265694 |
| DNMBP | 65.21111842 | -0.571106882 | 0.135505151 | -4.214650704 | 2.50E-05 | 0.00026592 |
| MAVS | 215.5549082 | -0.355562448 | 0.084391742 | -4.213237456 | 2.52E-05 | 0.000267437 |
| FCHO2 | 2209.341358 | -0.295370656 | 0.070112472 | -4.21281188 | 2.52E-05 | 0.000267789 |
| FAM71A | 15.7155289 | 1.300124951 | 0.308687214 | 4.211787507 | 2.53E-05 | 0.000268854 |
| SNHG7 | 44.11965656 | -0.860576589 | 0.204476052 | -4.208691339 | 2.57E-05 | 0.000272408 |
| FBXL3 | 510.3288443 | 0.261828076 | 0.062253842 | 4.205813917 | 2.60E-05 | 0.000275741 |
| SHISA5 | 1830.151623 | 0.454154549 | 0.108011919 | 4.204670672 | 2.61E-05 | 0.000276981 |
| BCAS2 | 987.1931204 | 0.412983905 | 0.098279076 | 4.202154936 | 2.64E-05 | 0.00027976 |
| ZNF366 | 206.5924484 | 0.645350614 | 0.153572749 | 4.202246927 | 2.64E-05 | 0.00027976 |
| UBE2L3 | 3674.066225 | 0.236293715 | 0.056236028 | 4.201820884 | 2.65E-05 | 0.000280014 |
| SLC25A22 | 103.9689726 | 0.660096385 | 0.157124007 | 4.201117311 | 2.66E-05 | 0.000280726 |
| CCNG1 | 351.156982 | -0.552849142 | 0.131674588 | -4.198601645 | 2.69E-05 | 0.000283702 |
| PLAU | 786.54289 | 0.395195417 | 0.094142806 | 4.197829171 | 2.69E-05 | 0.00028451 |
| SNTB2 | 333.1777294 | 0.459029975 | 0.109368847 | 4.197081588 | 2.70E-05 | 0.000285289 |
| LRP12 | 106.4241221 | 1.015277004 | 0.24202486 | 4.194928592 | 2.73E-05 | 0.000287849 |
| GDPD1 | 15.21898839 | -0.932472543 | 0.222540408 | -4.190126876 | 2.79E-05 | 0.000293842 |
| CRABP2 | 3289.089184 | 0.77088229 | 0.184010216 | 4.189345052 | 2.80E-05 | 0.00029469 |
| KBTBD7 | 62.62990567 | -0.692670572 | 0.165377979 | -4.188408738 | 2.81E-05 | 0.000295707 |
| REPS2 | 15.61193175 | -1.393410494 | 0.332690684 | -4.188306314 | 2.81E-05 | 0.000295707 |
| ADAM19 | 19.89238889 | 1.351811777 | 0.322799157 | 4.187779763 | 2.82E-05 | 0.000296227 |
| RPS3A | 3105.515173 | -0.621928158 | 0.148515372 | -4.187634916 | 2.82E-05 | 0.000296249 |
| BRD3 | 367.867701 | -0.334434559 | 0.079886162 | -4.186389101 | 2.83E-05 | 0.000297711 |
| PSMB6 | 1063.436711 | 0.395347884 | 0.094496183 | 4.183744499 | 2.87E-05 | 0.000301027 |
| PIK3CD | 307.5815586 | -0.477027887 | 0.114031926 | -4.183283618 | 2.87E-05 | 0.000301468 |
| CEPT1 | 483.8240865 | -0.502273868 | 0.120075452 | -4.182985451 | 2.88E-05 | 0.000301694 |
| CCND2 | 211.7791507 | 0.513945806 | 0.122882267 | 4.182424513 | 2.88E-05 | 0.00030227 |
| RHOU | 427.0814765 | 0.531534828 | 0.127095191 | 4.182178923 | 2.89E-05 | 0.000302427 |
| FAM110A | 117.8940203 | -0.380844088 | 0.091090402 | -4.180946391 | 2.90E-05 | 0.0003039 |
| SPRED1 | 423.853983 | 0.39489424 | 0.094458687 | 4.180602663 | 2.91E-05 | 0.000304189 |

| | | | | | | |
|---|---|---|---|---|---|---|
| IPO5 | 446.5408064 | -0.392392358 | 0.093881911 | -4.179637541 | 2.92E-05 | 0.000305311 |
| SQLE | 35.34721803 | 0.992805776 | 0.237564727 | 4.1790959 | 2.93E-05 | 0.000305868 |
| DYRK1B | 70.34146324 | -0.568840984 | 0.136127228 | -4.178745066 | 2.93E-05 | 0.000306168 |
| PCSK7 | 552.8223026 | -0.189859297 | 0.045452531 | -4.177089653 | 2.95E-05 | 0.000308231 |
| TTF2 | 153.6933709 | -0.307507468 | 0.073642014 | -4.175706944 | 2.97E-05 | 0.000309936 |
| FOXRED2 | 50.1321919 | -0.793467123 | 0.19006461 | -4.174723133 | 2.98E-05 | 0.000311104 |
| C2orf49 | 245.1591314 | -0.284547138 | 0.068188245 | -4.172964676 | 3.01E-05 | 0.00031334 |
| BAIAP2L1 | 5.556091907 | 2.284357775 | 0.547581608 | 4.171721151 | 3.02E-05 | 0.00031488 |
| TP53TG1 | 20.56436485 | -0.772723415 | 0.185271561 | -4.170761078 | 3.04E-05 | 0.000316033 |
| ING2 | 170.1035118 | -0.441663537 | 0.105933538 | -4.169251253 | 3.06E-05 | 0.000317956 |
| FADS2 | 133.9382761 | 0.886295149 | 0.212624391 | 4.168360669 | 3.07E-05 | 0.000319022 |
| PPARG | 2768.29251 | -0.318973125 | 0.076527038 | -4.168110155 | 3.07E-05 | 0.000319195 |
| NDUFB1 | 1328.787683 | -0.464560782 | 0.1115008 | -4.166434521 | 3.09E-05 | 0.00032137 |
| ZNF573 | 33.38535237 | -0.85616334 | 0.205652268 | -4.163160209 | 3.14E-05 | 0.000325833 |
| AHNAK | 1279.129021 | -0.310997641 | 0.074769628 | -4.159411364 | 3.19E-05 | 0.000331043 |
| MFGE8 | 388.6645429 | -0.617714171 | 0.1485599 | -4.158014195 | 3.21E-05 | 0.000332888 |
| R3HDM1 | 387.7728661 | -0.371964175 | 0.089465143 | -4.157643556 | 3.22E-05 | 0.000333243 |
| ELMO2 | 426.8109273 | 0.327603132 | 0.078819845 | 4.156353411 | 3.23E-05 | 0.000334943 |
| PPIF | 2838.686151 | 0.505317929 | 0.121625774 | 4.154694444 | 3.26E-05 | 0.000337195 |
| MAPKAPK2 | 1443.466253 | 0.271832218 | 0.065430012 | 4.154549401 | 3.26E-05 | 0.000337222 |
| THYN1 | 91.47942482 | -0.474083241 | 0.114170354 | -4.152419832 | 3.29E-05 | 0.000340187 |
| TM6SF1 | 375.5303867 | -0.582382329 | 0.140284208 | -4.151446123 | 3.30E-05 | 0.000341449 |
| MCM4 | 235.2885462 | 0.760163453 | 0.183157576 | 4.150324936 | 3.32E-05 | 0.000342935 |
| MOSPD2 | 1041.663322 | -0.248381641 | 0.05986077 | -4.149322506 | 3.33E-05 | 0.00034425 |
| ANXA9 | 25.39967431 | -0.931711955 | 0.224578801 | -4.14870839 | 3.34E-05 | 0.000344983 |
| GOSR1 | 291.5019455 | 0.32098179 | 0.077393894 | 4.147378722 | 3.36E-05 | 0.0003468 |
| ARF1 | 8378.823737 | 0.192908094 | 0.04651847 | 4.146913963 | 3.37E-05 | 0.000347312 |
| TBC1D23 | 844.3629092 | 0.348287874 | 0.083995547 | 4.14650405 | 3.38E-05 | 0.000347742 |
| ICAM1 | 983.2686828 | 1.078416723 | 0.260089321 | 4.146332183 | 3.38E-05 | 0.000347811 |
| KIAA0922 | 443.9892692 | -0.413336914 | 0.099713899 | -4.145228696 | 3.39E-05 | 0.000349297 |
| KIF13A | 46.25981823 | 0.794867604 | 0.191771912 | 4.144859352 | 3.40E-05 | 0.000349668 |
| MRPL55 | 132.9331291 | -0.323036357 | 0.077953596 | -4.143957094 | 3.41E-05 | 0.000350853 |
| RP2 | 1104.427336 | 0.243750479 | 0.058826936 | 4.143518172 | 3.42E-05 | 0.000351332 |
| EIF5A | 5430.190748 | 0.226580996 | 0.054695556 | 4.142585157 | 3.43E-05 | 0.00035257 |
| GNG2 | 32.12972528 | -1.035575623 | 0.250078278 | -4.141005899 | 3.46E-05 | 0.000354616 |
| TUBG1 | 722.1176516 | 0.235684154 | 0.056913626 | 4.141084835 | 3.46E-05 | 0.000354616 |
| KIAA0146 | 457.8511571 | -0.288869332 | 0.069794163 | -4.138875205 | 3.49E-05 | 0.000357728 |
| RPL38 | 3741.131317 | -0.303424107 | 0.073372998 | -4.135364726 | 3.54E-05 | 0.000363042 |
| PSAP | 133711.6274 | -0.320394451 | 0.077535105 | -4.132250162 | 3.59E-05 | 0.000367795 |
| CCDC64 | 14.32798397 | 1.013248097 | 0.245263841 | 4.13125756 | 3.61E-05 | 0.000368985 |
| TMEM150A | 254.1564816 | -0.405748365 | 0.098214295 | -4.131255697 | 3.61E-05 | 0.000368985 |
| LDLRAD3 | 21.61123799 | -1.282331327 | 0.310793582 | -4.125990373 | 3.69E-05 | 0.000377323 |
| RPS20 | 4733.932337 | -0.549675677 | 0.133248763 | -4.125184086 | 3.70E-05 | 0.00037844 |
| RHBDD2 | 1261.027846 | 0.411249437 | 0.099727929 | 4.123713794 | 3.73E-05 | 0.000380656 |
| STARD13 | 46.90602504 | -0.894129321 | 0.216900784 | -4.122296405 | 3.75E-05 | 0.000382797 |
| UNC119B | 48.19938663 | -0.585547717 | 0.14205549 | -4.121964727 | 3.76E-05 | 0.000383139 |
| GALNT12 | 1662.00706 | 0.514922023 | 0.124939904 | 4.121357614 | 3.77E-05 | 0.00038394 |

| | | | | | | |
|---|---|---|---|---|---|---|
| TARP | 54.72786625 | 1.268864928 | 0.307907323 | 4.120931313 | 3.77E-05 | 0.000384441 |
| ADCK3 | 260.8966189 | -0.350369474 | 0.085039442 | -4.120082007 | 3.79E-05 | 0.00038565 |
| SESN2 | 868.6995498 | 0.369448603 | 0.089685239 | 4.119391409 | 3.80E-05 | 0.000386596 |
| CHST7 | 780.4006412 | 0.640269488 | 0.155559685 | 4.115908862 | 3.86E-05 | 0.000392267 |
| VGLL4 | 243.923645 | -0.300023746 | 0.072910644 | -4.114951236 | 3.87E-05 | 0.000393684 |
| CLSTN3 | 81.81388112 | 0.621520741 | 0.151077919 | 4.113908524 | 3.89E-05 | 0.000395252 |
| SCP2 | 1732.325279 | -0.311933993 | 0.075833671 | -4.113396967 | 3.90E-05 | 0.000395913 |
| ZNF510 | 102.0549826 | -0.481751052 | 0.11713279 | -4.112862432 | 3.91E-05 | 0.000396615 |
| PTRHD1 | 409.6839429 | -0.36409166 | 0.088545612 | -4.111910803 | 3.92E-05 | 0.000398037 |
| LOC38764[illegible] | 554.2415868 | 0.242907808 | 0.059089862 | 4.110820346 | 3.94E-05 | 0.000399704 |
| MAP4K3 | 298.8981572 | 0.481718345 | 0.117223545 | 4.109399221 | 3.97E-05 | 0.000401953 |
| AHRR | 5.990209949 | -1.628865177 | 0.396393937 | -4.109208101 | 3.97E-05 | 0.000402067 |
| CRYBB1 | 1.980959063 | -3.009325578 | 0.732445437 | -4.10860035 | 3.98E-05 | 0.000402908 |
| PHLDA3 | 379.9666965 | 0.347592188 | 0.08462004 | 4.107681671 | 4.00E-05 | 0.000404293 |
| ACSM3 | 27.53883731 | -0.788039287 | 0.191856416 | -4.107442977 | 4.00E-05 | 0.000404492 |
| SIGLEC11 | 303.26728 | -0.490703963 | 0.119530565 | -4.105259312 | 4.04E-05 | 0.00040811 |
| TLR5 | 54.57680357 | -1.194338873 | 0.290964864 | -4.104752907 | 4.05E-05 | 0.000408784 |
| ATP7A | 242.5238358 | -0.39139699 | 0.095383031 | -4.103423711 | 4.07E-05 | 0.000410918 |
| PSMB10 | 949.274957 | 0.373061403 | 0.090937207 | 4.102406663 | 4.09E-05 | 0.000412505 |
| FAM212B | 48.00893062 | -0.600703836 | 0.146573965 | -4.098298343 | 4.16E-05 | 0.000419667 |
| DANCR | 96.8810378 | -0.474863278 | 0.115930469 | -4.096104188 | 4.20E-05 | 0.000423433 |
| CDA | 155.3339023 | 0.513657445 | 0.125410605 | 4.095805497 | 4.21E-05 | 0.00042375 |
| DUSP16 | 232.1352488 | -0.469309431 | 0.11461428 | -4.094685491 | 4.23E-05 | 0.000425574 |
| AKR1C1 | 322.0549882 | 0.920682059 | 0.22486766 | 4.094328449 | 4.23E-05 | 0.000426 |
| IL18BP | 128.3820395 | -0.766229603 | 0.18717216 | -4.093715658 | 4.25E-05 | 0.000426897 |
| IFI16 | 2681.993291 | 0.292595164 | 0.071480143 | 4.093376862 | 4.25E-05 | 0.000427291 |
| FITM2 | 72.98705996 | -0.434265971 | 0.106179093 | -4.089938585 | 4.31E-05 | 0.000433441 |
| KCNMA1 | 18.47043461 | -0.846509675 | 0.207032172 | -4.08878323 | 4.34E-05 | 0.000435371 |
| DUSP8 | 74.53887007 | 0.615384963 | 0.150551322 | 4.087542731 | 4.36E-05 | 0.000437233 |
| N4BP2L1 | 162.6754005 | 0.372028286 | 0.091013453 | 4.087618619 | 4.36E-05 | 0.000437233 |
| ATP2B1 | 769.3748683 | 0.388689904 | 0.0951383 | 4.085525021 | 4.40E-05 | 0.000440813 |
| AHCYL1 | 2636.94148 | 0.322786762 | 0.079083454 | 4.081596667 | 4.47E-05 | 0.000448091 |
| BCL2A1 | 2243.525787 | 0.547151635 | 0.134058903 | 4.081427068 | 4.48E-05 | 0.000448177 |
| SOCS7 | 117.0004951 | -0.338007139 | 0.082819296 | -4.081260712 | 4.48E-05 | 0.000448258 |
| FAM214A | 140.8619509 | -0.35487579 | 0.086978723 | -4.080029897 | 4.50E-05 | 0.000450396 |
| RGL3 | 59.41771149 | -0.47540281 | 0.116530309 | -4.079649447 | 4.51E-05 | 0.000450891 |
| UTP6 | 769.6448958 | 0.240416011 | 0.058963374 | 4.077378789 | 4.55E-05 | 0.000455072 |
| ACE | 451.0377348 | 0.697491432 | 0.171140936 | 4.07553826 | 4.59E-05 | 0.000458441 |
| C8orf76 | 478.0717212 | 0.332183501 | 0.081512782 | 4.075232026 | 4.60E-05 | 0.000458799 |
| GTPBP4 | 299.6473831 | 0.423407394 | 0.103925247 | 4.074153366 | 4.62E-05 | 0.000460683 |
| HIAT1 | 329.0293639 | 0.287467885 | 0.070579312 | 4.07297655 | 4.64E-05 | 0.00046277 |
| AVPI1 | 873.9046107 | -0.240609333 | 0.059077807 | -4.072753312 | 4.65E-05 | 0.000462966 |
| KIAA1147 | 497.9483066 | 0.399345367 | 0.09814083 | 4.069105268 | 4.72E-05 | 0.000470023 |
| CUEDC1 | 286.8059761 | -0.392069154 | 0.096381046 | -4.067907215 | 4.74E-05 | 0.000472193 |
| AMIGO2 | 180.3137012 | -0.443405995 | 0.109026644 | -4.0669508 | 4.76E-05 | 0.000473882 |
| LOC10027[illegible] | 6.165688437 | -1.476316936 | 0.363036829 | -4.066576218 | 4.77E-05 | 0.00047439 |
| OSGEP | 232.1431544 | -0.255271595 | 0.062820802 | -4.063488331 | 4.83E-05 | 0.000480456 |

| | | | | | | |
|---|---|---|---|---|---|---|
| C11orf84 | 125.0229907 | -0.437936161 | 0.107777204 | -4.063346818 | 4.84E-05 | 0.000480492 |
| DIMT1 | 118.9731543 | -0.384645269 | 0.094685565 | -4.062343288 | 4.86E-05 | 0.000482306 |
| CXCR6 | 30.79412257 | 0.993171002 | 0.244523086 | 4.061665573 | 4.87E-05 | 0.000483451 |
| GLDN | 97.27632966 | 0.880924753 | 0.217023315 | 4.059124943 | 4.93E-05 | 0.00048848 |
| SNX7 | 161.9477372 | -0.318524404 | 0.078506524 | -4.057298538 | 4.96E-05 | 0.000492053 |
| ALG14 | 97.14717978 | -0.460917483 | 0.113612169 | -4.05693762 | 4.97E-05 | 0.000492551 |
| TKT | 3165.247377 | -0.559665173 | 0.138006638 | -4.055349659 | 5.01E-05 | 0.000495646 |
| TAGLN2 | 6043.911521 | 0.293072178 | 0.072305019 | 4.053275741 | 5.05E-05 | 0.000499797 |
| ZSCAN16 | 41.81018795 | 0.571971302 | 0.141199219 | 4.05081067 | 5.10E-05 | 0.000504824 |
| CSRP2 | 10.33504022 | 1.244225488 | 0.307200867 | 4.050201743 | 5.12E-05 | 0.000505659 |
| S1PR3 | 5.338158722 | 2.670643918 | 0.659389616 | 4.050175882 | 5.12E-05 | 0.000505659 |
| LOC73010: | 61.52712812 | -0.483713257 | 0.119455826 | -4.049306531 | 5.14E-05 | 0.000507272 |
| PNP | 590.1669836 | 0.526620968 | 0.130070313 | 4.048740696 | 5.15E-05 | 0.00050823 |
| CASS4 | 104.369885 | 0.584527143 | 0.144481136 | 4.045698704 | 5.22E-05 | 0.000514603 |
| DDX20 | 317.2970179 | -0.349573624 | 0.086415266 | -4.045276234 | 5.23E-05 | 0.000515259 |
| HMGCR | 295.1074902 | 0.399285636 | 0.098714615 | 4.044848227 | 5.24E-05 | 0.000515929 |
| RAP2C | 563.4425831 | 0.372251324 | 0.092047736 | 4.044111684 | 5.25E-05 | 0.000517279 |
| MPZL3 | 114.6966982 | 0.528036878 | 0.13057956 | 4.043794298 | 5.26E-05 | 0.000517707 |
| ERO1L | 958.7442353 | 0.243021927 | 0.060116166 | 4.042538671 | 5.29E-05 | 0.000519939 |
| LRRC37A3 | 13.59782511 | 0.966459813 | 0.239070301 | 4.042575797 | 5.29E-05 | 0.000519939 |
| PXMP4 | 69.20566073 | -0.485014896 | 0.120004349 | -4.041644331 | 5.31E-05 | 0.000521651 |
| NUDT14 | 308.6663617 | -0.375064468 | 0.092813153 | -4.0410702 | 5.32E-05 | 0.000522654 |
| FAM228B | 23.09703581 | -0.932453767 | 0.230976428 | -4.037008344 | 5.41E-05 | 0.000531502 |
| STAT4 | 17.21876092 | 1.227545209 | 0.304285294 | 4.034191699 | 5.48E-05 | 0.000537634 |
| IL4R | 1332.300334 | 0.363794022 | 0.090181291 | 4.034029848 | 5.48E-05 | 0.000537722 |
| ZDHHC11 | 15.21142862 | -1.095161856 | 0.271494761 | -4.033823163 | 5.49E-05 | 0.000537912 |
| GLTSCR2 | 890.3184787 | -0.667259541 | 0.165589943 | -4.029589776 | 5.59E-05 | 0.000547403 |
| SPAG9 | 1349.864486 | 0.303948197 | 0.075465959 | 4.027619906 | 5.63E-05 | 0.000551719 |
| TMCC3 | 458.9382271 | 0.423374553 | 0.105127613 | 4.027244054 | 5.64E-05 | 0.000552311 |
| FBXO15 | 10.6892357 | 0.994571122 | 0.246973631 | 4.02703364 | 5.65E-05 | 0.000552515 |
| MTA1 | 221.5011518 | -0.366794897 | 0.091087801 | -4.026827884 | 5.65E-05 | 0.000552708 |
| ZBTB17 | 175.4215126 | 0.343616086 | 0.085340515 | 4.026412162 | 5.66E-05 | 0.000553395 |
| TIMMDC1 | 447.9989253 | -0.332711988 | 0.082681278 | -4.024030538 | 5.72E-05 | 0.000558732 |
| GLCCI1 | 23.80106675 | -0.702877971 | 0.174804292 | -4.020942293 | 5.80E-05 | 0.000565516 |
| ST3GAL2 | 648.1325251 | 0.617773538 | 0.153635983 | 4.021021144 | 5.79E-05 | 0.000565516 |
| PKN1 | 1410.954278 | -0.250423646 | 0.062283171 | -4.020727286 | 5.80E-05 | 0.000565736 |
| PSMA2 | 965.4900687 | 0.335783124 | 0.083639134 | 4.014665245 | 5.95E-05 | 0.000580171 |
| CST7 | 22.36663754 | -1.117761017 | 0.278478296 | -4.013817346 | 5.97E-05 | 0.000581955 |
| MELK | 21.935441 | 0.695992782 | 0.173410305 | 4.013560678 | 5.98E-05 | 0.000582284 |
| SLC2A4RG | 182.0173239 | -0.461957432 | 0.115149505 | -4.011805639 | 6.03E-05 | 0.000586325 |
| LOC10050! | 109.5220269 | -0.640369641 | 0.159678762 | -4.010362003 | 6.06E-05 | 0.000589614 |
| ACOT13 | 1640.011613 | -0.336226827 | 0.083848599 | -4.00992777 | 6.07E-05 | 0.000590391 |
| NDST1 | 1486.026136 | 0.306357211 | 0.076412306 | 4.009265355 | 6.09E-05 | 0.000591741 |
| TNFAIP3 | 910.1634369 | 1.036016611 | 0.258466884 | 4.008314706 | 6.12E-05 | 0.000593818 |
| SAMD9 | 80.71200712 | 1.055318771 | 0.263294152 | 4.008136012 | 6.12E-05 | 0.000593958 |
| MYO7B | 87.85102833 | 0.760547985 | 0.189988733 | 4.003121516 | 6.25E-05 | 0.000606378 |
| RFXAP | 17.57371706 | -0.660012588 | 0.164888416 | -4.002783241 | 6.26E-05 | 0.00060693 |

| | | | | | | |
|---|---|---|---|---|---|---|
| CRYL1 | 406.2862886 | -0.431384139 | 0.107783192 | -4.002332182 | 6.27E-05 | 0.000607772 |
| NAV1 | 65.0271132 | -0.663852351 | 0.16587368 | -4.002156049 | 6.28E-05 | 0.000607909 |
| RPS27A | 2497.170233 | -0.523337124 | 0.130783842 | -4.00154267 | 6.29E-05 | 0.000609171 |
| ATP6V0D1 | 3149.568947 | -0.289772464 | 0.07244539 | -3.999874463 | 6.34E-05 | 0.000613162 |
| SDS | 154.711422 | -0.942430983 | 0.235791296 | -3.996886229 | 6.42E-05 | 0.000620629 |
| MLLT11 | 15.00985324 | 1.551285171 | 0.388190987 | 3.996190597 | 6.44E-05 | 0.000622131 |
| MPPE1 | 159.4603418 | -0.349264975 | 0.087429634 | -3.994812272 | 6.47E-05 | 0.000625437 |
| APOE | 65210.40951 | -0.578682721 | 0.14492024 | -3.993111805 | 6.52E-05 | 0.000629615 |
| CNKSR3 | 5.748564948 | 1.969797489 | 0.493377744 | 3.992473342 | 6.54E-05 | 0.000630986 |
| ZBTB2 | 222.7676749 | 0.37011863 | 0.092715003 | 3.992003671 | 6.55E-05 | 0.00063191 |
| ACBD4 | 60.14999095 | -0.548183041 | 0.137327147 | -3.991803912 | 6.56E-05 | 0.000632116 |
| H2AFZ | 2655.674567 | -0.402933156 | 0.100949177 | -3.991445667 | 6.57E-05 | 0.000632417 |
| WDFY2 | 200.0989163 | -0.465645841 | 0.116658976 | -3.991513178 | 6.57E-05 | 0.000632417 |
| SPATA12 | 2.958489965 | -2.211917773 | 0.554492608 | -3.989084326 | 6.63E-05 | 0.000638415 |
| AKR1A1 | 2253.280305 | -0.202399365 | 0.050750275 | -3.988143198 | 6.66E-05 | 0.000640622 |
| MAMLD1 | 25.667966 | 1.179741526 | 0.295872552 | 3.987330073 | 6.68E-05 | 0.000642489 |
| SYMPK | 1447.101205 | 0.352748881 | 0.088492756 | 3.986189354 | 6.71E-05 | 0.000645251 |
| NEXN | 4.290210372 | 1.68225103 | 0.422250893 | 3.984008221 | 6.78E-05 | 0.00065087 |
| KLHL36 | 262.6439132 | -0.526413205 | 0.132192414 | -3.98217408 | 6.83E-05 | 0.000655576 |
| GALC | 858.5333882 | -0.692222655 | 0.173854933 | -3.981610672 | 6.84E-05 | 0.000656794 |
| ASB2 | 11.92347203 | -1.021532208 | 0.256703941 | -3.979417703 | 6.91E-05 | 0.000662541 |
| RPS13 | 5020.478433 | -0.504402825 | 0.126763125 | -3.979097417 | 6.92E-05 | 0.000663092 |
| SLC31A1 | 3834.434259 | 0.436934923 | 0.109829698 | 3.97829488 | 6.94E-05 | 0.000664992 |
| PODXL | 44.62829275 | 1.037341207 | 0.260765156 | 3.978066784 | 6.95E-05 | 0.000665288 |
| BBS9 | 94.43097303 | -0.456813896 | 0.114859341 | -3.97715928 | 6.97E-05 | 0.000667365 |
| HMGA1 | 2110.466992 | 0.596965015 | 0.150101293 | 3.977081102 | 6.98E-05 | 0.000667365 |
| BCAR3 | 396.8135434 | 0.350683333 | 0.088226376 | 3.974812839 | 7.04E-05 | 0.000673411 |
| FERMT3 | 3393.658716 | 0.314950892 | 0.079243657 | 3.974461872 | 7.05E-05 | 0.000674059 |
| BIRC7 | 41.29695325 | -1.203247931 | 0.302850982 | -3.973069269 | 7.10E-05 | 0.000677666 |
| TYRO3 | 6.997500549 | 1.585052873 | 0.399001069 | 3.972552946 | 7.11E-05 | 0.00067879 |
| PNMA1 | 268.7535517 | 0.330724516 | 0.083266472 | 3.971880994 | 7.13E-05 | 0.000680359 |
| PLXNC1 | 195.6050249 | -0.774549617 | 0.195018606 | -3.971670351 | 7.14E-05 | 0.00068047 |
| ZNF101 | 80.76510635 | 0.358361479 | 0.090231047 | 3.971598361 | 7.14E-05 | 0.00068047 |
| ARNTL2 | 203.8222296 | 1.302473985 | 0.328128782 | 3.969398777 | 7.21E-05 | 0.000686431 |
| PRPF8 | 2842.930042 | -0.255385483 | 0.064430304 | -3.963747934 | 7.38E-05 | 0.000702533 |
| ISOC1 | 206.7436733 | 0.475436224 | 0.119955534 | 3.963437194 | 7.39E-05 | 0.000703089 |
| SLC9A9 | 104.3388409 | 0.719984313 | 0.181738301 | 3.961654258 | 7.44E-05 | 0.000707999 |
| ZDHHC7 | 651.3714007 | -0.349937439 | 0.088333925 | -3.961529386 | 7.45E-05 | 0.000708008 |
| ACCS | 14.71556513 | -0.791857733 | 0.199951903 | -3.960241036 | 7.49E-05 | 0.000711476 |
| TMEM176/ | 179.4583929 | -0.765518675 | 0.193310781 | -3.960041283 | 7.49E-05 | 0.000711709 |
| DNM1L | 968.1275197 | 0.291886562 | 0.073721679 | 3.959304316 | 7.52E-05 | 0.000713181 |
| REC8 | 38.5572171 | 0.538724661 | 0.136065106 | 3.95931533 | 7.52E-05 | 0.000713181 |
| TMEM88 | 4.665803718 | 2.161243948 | 0.545884044 | 3.959163072 | 7.52E-05 | 0.00071324 |
| IFFO1 | 294.0876204 | -0.481125944 | 0.121529645 | -3.958918358 | 7.53E-05 | 0.000713608 |
| GNPAT | 301.35256 | -0.314767647 | 0.079567764 | -3.955969503 | 7.62E-05 | 0.000721735 |
| ITK | 31.24341874 | 0.815372145 | 0.206109019 | 3.956023609 | 7.62E-05 | 0.000721735 |
| MAST3 | 204.5006555 | -0.401739842 | 0.10161032 | -3.953730718 | 7.69E-05 | 0.000728155 |

| | | | | | | |
|---|---|---|---|---|---|---|
| HK1 | 826.100611 | -0.313409172 | 0.079277133 | -3.953336344 | 7.71E-05 | 0.000728987 |
| CTAGE4 | 37.41270923 | 0.883256003 | 0.22350972 | 3.951756562 | 7.76E-05 | 0.000733444 |
| PPFIBP1 | 90.26205988 | 0.499834718 | 0.126566327 | 3.949191941 | 7.84E-05 | 0.00074097 |
| LTBP3 | 618.9724052 | 0.595671867 | 0.150871842 | 3.94819775 | 7.87E-05 | 0.000742923 |
| PARP1 | 1370.322439 | -0.30650149 | 0.077629619 | -3.948254453 | 7.87E-05 | 0.000742923 |
| SLC12A6 | 727.3580194 | -0.306990599 | 0.077751831 | -3.94833917 | 7.87E-05 | 0.000742923 |
| ACVR1 | 584.2853981 | 0.377360268 | 0.095605104 | 3.947072393 | 7.91E-05 | 0.000746045 |
| LOC64699! | 6.849867431 | 1.368588301 | 0.346757013 | 3.946822265 | 7.92E-05 | 0.000746447 |
| C1RL-AS1 | 9.412227501 | 0.98963248 | 0.250822427 | 3.945550215 | 7.96E-05 | 0.000750042 |
| ZNF248 | 44.12588028 | -0.519888333 | 0.131817317 | -3.944006341 | 8.01E-05 | 0.000754508 |
| DNTTIP2 | 1209.681291 | 0.246593689 | 0.06253696 | 3.943167215 | 8.04E-05 | 0.000756771 |
| BCS1L | 119.9777404 | -0.417912789 | 0.105994911 | -3.942762773 | 8.05E-05 | 0.000757667 |
| ATP5B | 5108.195437 | 0.337713247 | 0.085666758 | 3.94217378 | 8.07E-05 | 0.000759147 |
| ADAM28 | 32.0655292 | -0.910723886 | 0.231124064 | -3.940411354 | 8.13E-05 | 0.000764186 |
| DYNC2H1 | 15.9078159 | -0.848008928 | 0.215211836 | -3.940345211 | 8.14E-05 | 0.000764186 |
| BTG3 | 107.5215633 | 0.860006869 | 0.218284568 | 3.939842737 | 8.15E-05 | 0.000765402 |
| TRIM54 | 20.14462111 | 1.420739995 | 0.360900313 | 3.936654925 | 8.26E-05 | 0.000775247 |
| RAPGEF3 | 176.670056 | -0.460330249 | 0.117042048 | -3.933033084 | 8.39E-05 | 0.000786629 |
| FEM1B | 1893.403756 | 0.414654782 | 0.105492696 | 3.930649206 | 8.47E-05 | 0.000794071 |
| FLOT1 | 2212.148007 | 0.317825309 | 0.080866647 | 3.930239717 | 8.49E-05 | 0.000795025 |
| CASP10 | 699.3007376 | 0.351085909 | 0.089359938 | 3.928896073 | 8.53E-05 | 0.000799077 |
| INVS | 214.8442267 | -0.279941269 | 0.071265 | -3.92817329 | 8.56E-05 | 0.00080108 |
| HNRNPA1L | 46.32919948 | -0.55023782 | 0.140092736 | -3.927668449 | 8.58E-05 | 0.00080236 |
| RDX | 1202.75639 | 0.41920233 | 0.106758041 | 3.926658126 | 8.61E-05 | 0.000805332 |
| DNM2 | 1462.843027 | 0.253148844 | 0.064504203 | 3.924532559 | 8.69E-05 | 0.000812068 |
| ZNF337 | 76.25222926 | -0.418850826 | 0.106759552 | -3.923310081 | 8.73E-05 | 0.000815792 |
| CD226 | 66.24492934 | 0.67408628 | 0.171834238 | 3.922886899 | 8.75E-05 | 0.000816817 |
| COL8A2 | 62.41636368 | 0.765764831 | 0.195289568 | 3.921176332 | 8.81E-05 | 0.000822226 |
| S100A8 | 286.0532698 | -1.021849584 | 0.260839276 | -3.917544932 | 8.95E-05 | 0.00083429 |
| IL36G | 6.543970335 | 2.062412227 | 0.526877174 | 3.914408004 | 9.06E-05 | 0.000844787 |
| SKAP2 | 1265.045037 | -0.335935533 | 0.08585567 | -3.912793808 | 9.12E-05 | 0.000850031 |
| ZNF700 | 24.61271438 | -0.840073968 | 0.214738558 | -3.91207791 | 9.15E-05 | 0.00085213 |
| VPS35 | 679.2398496 | -0.426334812 | 0.108982951 | -3.911940444 | 9.16E-05 | 0.000852189 |
| GNAQ | 1253.664353 | -0.281811479 | 0.072049949 | -3.911334881 | 9.18E-05 | 0.000853902 |
| DPH1 | 155.0943109 | -0.30838909 | 0.078885717 | -3.909314645 | 9.26E-05 | 0.000860645 |
| GPR75 | 3.653896014 | 1.935828665 | 0.495213944 | 3.909075435 | 9.27E-05 | 0.000861068 |
| RNF141 | 154.0039714 | -0.419219445 | 0.107319146 | -3.906287572 | 9.37E-05 | 0.00087019 |
| TMEM55A | 118.1354827 | -0.398099124 | 0.101912264 | -3.906292635 | 9.37E-05 | 0.00087019 |
| TRIM39 | 177.073668 | 0.251593059 | 0.064525271 | 3.899139906 | 9.65E-05 | 0.000895833 |
| SYP | 31.20981066 | 0.829364432 | 0.21273289 | 3.898618744 | 9.67E-05 | 0.000897316 |
| SDC2 | 6061.763749 | 0.530131407 | 0.136174661 | 3.893025361 | 9.90E-05 | 0.000917811 |
| TNC | 8.538063863 | 1.653353069 | 0.424734496 | 3.892674327 | 9.91E-05 | 0.000918683 |
| CYTH4 | 358.3150604 | 0.374555481 | 0.096233809 | 3.892140231 | 9.94E-05 | 0.000920251 |
| ACO2 | 1312.225768 | 0.282931405 | 0.072744803 | 3.889369329 | 0.000100505 | 0.000930358 |
| MTMR10 | 1204.923572 | -0.249340554 | 0.064136359 | -3.887663068 | 0.000101214 | 0.000936456 |
| STRN3 | 357.0075421 | 0.322912865 | 0.083103854 | 3.885654519 | 0.000102055 | 0.000943764 |
| NME3 | 208.4349827 | -0.436913105 | 0.112451905 | -3.885333079 | 0.00010219 | 0.000944545 |

| | | | | | | |
|---|---|---|---|---|---|---|
| CARD9 | 166.867873 | -0.416181068 | 0.107143289 | -3.884340975 | 0.000102608 | 0.000947801 |
| FOS | 524.2115081 | -0.543121061 | 0.139826279 | -3.884255987 | 0.000102644 | 0.000947801 |
| NAPSB | 39.92221219 | -1.221832091 | 0.314622584 | -3.883485018 | 0.00010297 | 0.000950342 |
| HS2ST1 | 284.488442 | -0.321521639 | 0.08281811 | -3.882262434 | 0.000103489 | 0.000954661 |
| NDUFV2 | 603.5469567 | 0.310426366 | 0.079972768 | 3.881650895 | 0.00010375 | 0.000956592 |
| GZF1 | 162.7781359 | -0.298699678 | 0.077001626 | -3.879134668 | 0.000104829 | 0.000966062 |
| ZNF814 | 96.9385395 | -0.35508058 | 0.091541439 | -3.878905371 | 0.000104928 | 0.000966495 |
| IRF2BP2 | 1389.275293 | -0.400301468 | 0.103248027 | -3.87708589 | 0.000105715 | 0.000973268 |
| BIVM | 65.66169922 | -0.468045508 | 0.120848868 | -3.872982145 | 0.000107512 | 0.00098932 |
| CPNE6 | 13.99268016 | 1.413875513 | 0.365279997 | 3.870662296 | 0.00010854 | 0.00099829 |
| PRPF38B | 422.43568 | -0.253617043 | 0.065543254 | -3.869460677 | 0.000109076 | 0.001002728 |
| MRPS6 | 4596.880526 | 0.327643461 | 0.084733534 | 3.866750821 | 0.000110295 | 0.001013431 |
| MIER1 | 669.380954 | 0.259316564 | 0.067086884 | 3.865383942 | 0.000110915 | 0.001018621 |
| C4A | 17.6831386 | 1.104068079 | 0.285833001 | 3.862633338 | 0.000112171 | 0.001029284 |
| NME4 | 105.7707852 | -0.771365489 | 0.199701046 | -3.862601151 | 0.000112186 | 0.001029284 |
| ZBTB44 | 212.9770231 | -0.270990753 | 0.07017643 | -3.861563663 | 0.000112664 | 0.001032648 |
| ZNF69 | 17.38083949 | -1.013606243 | 0.262482009 | -3.861621777 | 0.000112637 | 0.001032648 |
| SLC11A1 | 3221.876338 | 0.367091315 | 0.095080101 | 3.860863742 | 0.000112987 | 0.001035102 |
| GMFG | 707.4590451 | -0.283866131 | 0.073563342 | -3.858798745 | 0.000113946 | 0.001043373 |
| DAPK1 | 3697.036235 | -0.264568633 | 0.068574218 | -3.858135601 | 0.000114255 | 0.001045694 |
| LOC72819( | 16.5178049 | -0.812023002 | 0.210479392 | -3.857969157 | 0.000114333 | 0.001045892 |
| CDKL1 | 53.19645954 | 0.526263347 | 0.136432509 | 3.857316341 | 0.000114639 | 0.001048174 |
| FRK | 9.161091358 | -1.152706011 | 0.298895024 | -3.856558048 | 0.000114995 | 0.001050399 |
| RASAL1 | 7.650005801 | -1.792774854 | 0.464851774 | -3.856659164 | 0.000114947 | 0.001050399 |
| SPTLC2 | 610.4612137 | 0.287677753 | 0.074606123 | 3.855953654 | 0.000115279 | 0.001052172 |
| TBXA2R | 10.3006993 | 0.938048825 | 0.243275871 | 3.855905733 | 0.000115302 | 0.001052172 |
| ITPK1 | 2958.267886 | -0.480328359 | 0.124580335 | -3.855571254 | 0.00011546 | 0.001053096 |
| AP2A2 | 668.0576004 | -0.388194563 | 0.100764101 | -3.852508569 | 0.000116914 | 0.001065838 |
| RASD1 | 9.294107318 | 1.235268294 | 0.320703009 | 3.851751489 | 0.000117276 | 0.001068617 |
| SLC43A1 | 11.34644791 | -1.268045159 | 0.329231913 | -3.851525657 | 0.000117384 | 0.00106908 |
| DHX8 | 297.0862366 | 0.306875274 | 0.079679601 | 3.851365619 | 0.000117461 | 0.001069256 |
| BRPF3 | 256.6806668 | 0.279736425 | 0.072639843 | 3.851005366 | 0.000117634 | 0.001070308 |
| PTPN7 | 288.2004753 | 0.359911669 | 0.093513914 | 3.848749923 | 0.000118722 | 0.001079682 |
| DCTN3 | 436.3246646 | -0.203720194 | 0.052948001 | -3.847552114 | 0.000119304 | 0.001084443 |
| CYFIP2 | 254.0586812 | -0.428676143 | 0.111419632 | -3.847402263 | 0.000119377 | 0.001084578 |
| TRIM22 | 2069.032489 | 0.461542539 | 0.119990741 | 3.846484605 | 0.000119825 | 0.001088115 |
| CCDC50 | 840.2795409 | 0.41342317 | 0.107508194 | 3.845503823 | 0.000120305 | 0.001091945 |
| LOC10013⁄ | 57.23634222 | -0.633911257 | 0.164921195 | -3.843722201 | 0.000121182 | 0.001098836 |
| SYNJ2BP | 151.6379456 | -0.326795659 | 0.085019419 | -3.84377669 | 0.000121155 | 0.001098836 |
| SLC43A2 | 1772.599706 | 0.415258874 | 0.108043131 | 3.843454658 | 0.000121314 | 0.0010995 |
| EHF | 63.96722959 | 0.903431033 | 0.235136671 | 3.84215286 | 0.00012196 | 0.001104813 |
| EIF4B | 938.5068882 | -0.872315255 | 0.227052852 | -3.841903981 | 0.000122084 | 0.001105396 |
| CYP1B1 | 1517.401013 | 0.561264467 | 0.146165648 | 3.839920498 | 0.000123074 | 0.001113824 |
| SLC19A1 | 45.28741517 | -0.746907738 | 0.194539631 | -3.839360308 | 0.000123355 | 0.001115826 |
| ARMC7 | 181.2043682 | -0.436673797 | 0.113773276 | -3.838105171 | 0.000123987 | 0.001121 |
| ARHGAP2( | 9.853691513 | 1.776592232 | 0.463090097 | 3.836385715 | 0.000124858 | 0.001127249 |
| NUP107 | 260.6526564 | 0.272737596 | 0.071092395 | 3.836382164 | 0.00012486 | 0.001127249 |

| | | | | | | |
|---|---|---|---|---|---|---|
| USP4 | 497.9440107 | 0.248465591 | 0.064762401 | 3.836571658 | 0.000124764 | 0.001127249 |
| CTNS | 583.3619949 | 0.471435625 | 0.123038246 | 3.831618537 | 0.000127303 | 0.001148748 |
| NDUFA7 | 638.3576266 | 0.215527545 | 0.056256387 | 3.831165812 | 0.000127538 | 0.001150307 |
| PLEC | 636.0259896 | 0.35176254 | 0.09188411 | 3.828328337 | 0.000129017 | 0.001162749 |
| PSMD11 | 2218.232131 | 0.397073437 | 0.103721104 | 3.828280097 | 0.000129042 | 0.001162749 |
| LAMA1 | 30.48488586 | -0.749656891 | 0.195851957 | -3.827671183 | 0.000129361 | 0.001165066 |
| SPATA13 | 390.3757121 | 0.495854131 | 0.129576642 | 3.826724663 | 0.00012986 | 0.001168988 |
| PDSS1 | 129.3068616 | 0.404168925 | 0.105627725 | 3.826352656 | 0.000130056 | 0.001170189 |
| EGLN1 | 494.8812351 | -0.349284621 | 0.09133618 | -3.824164961 | 0.000131216 | 0.001180045 |
| RBCK1 | 443.1595206 | 0.425998747 | 0.111399924 | 3.824048801 | 0.000131278 | 0.001180045 |
| KCTD12 | 1013.394941 | -0.396872072 | 0.10378726 | -3.823899688 | 0.000131357 | 0.00118019 |
| HEATR3 | 316.2642599 | -0.258213317 | 0.067535538 | -3.82336951 | 0.00013164 | 0.001181592 |
| TUBGCP3 | 537.1186055 | -0.183510938 | 0.047996697 | -3.823407608 | 0.00013162 | 0.001181592 |
| DOK3 | 440.7712546 | -0.430793563 | 0.112685855 | -3.822960426 | 0.000131859 | 0.001182985 |
| MFSD1 | 2512.972674 | -0.214431943 | 0.056093523 | -3.82275761 | 0.000131967 | 0.001183389 |
| ATP13A1 | 607.1612576 | 0.370628125 | 0.096995298 | 3.821093734 | 0.000132861 | 0.001190829 |
| SCIN | 84.71424115 | -0.568306295 | 0.14875016 | -3.820542413 | 0.000133159 | 0.001192921 |
| PRPF4B | 848.4309084 | 0.253335882 | 0.066318472 | 3.819989743 | 0.000133457 | 0.001195023 |
| ALPK3 | 139.3064269 | 0.570740864 | 0.149484617 | 3.818057494 | 0.000134507 | 0.00120384 |
| DGKG | 13.91814107 | -1.080103127 | 0.282912025 | -3.817805647 | 0.000134644 | 0.001204491 |
| TBC1D22A | 288.0970043 | -0.209885644 | 0.055000786 | -3.816048064 | 0.000135606 | 0.001212516 |
| AMDHD2 | 429.8325184 | -0.370102463 | 0.09701185 | -3.815023249 | 0.00013617 | 0.001216975 |
| ZNF516 | 101.430444 | -0.38293252 | 0.100423123 | -3.813190696 | 0.000137184 | 0.001225451 |
| SNX18 | 364.7418497 | -0.36555804 | 0.095884697 | -3.812475301 | 0.000137582 | 0.001228415 |
| LIPN | 15.17024582 | -1.440486306 | 0.377931316 | -3.811502902 | 0.000138124 | 0.001232423 |
| RRBP1 | 3249.885745 | 0.4540473 | 0.119127691 | 3.811433733 | 0.000138163 | 0.001232423 |
| RSPO3 | 104.5658674 | 1.267114787 | 0.332564039 | 3.810137715 | 0.000138889 | 0.001238309 |
| ABCC1 | 807.7064105 | 0.39539298 | 0.103779886 | 3.809919194 | 0.000139012 | 0.001238811 |
| FGF13 | 4.932957365 | 2.001426091 | 0.52543129 | 3.809110974 | 0.000139467 | 0.001242273 |
| SIRPB2 | 108.9342874 | -0.662282391 | 0.173922163 | -3.807924064 | 0.000140138 | 0.001247653 |
| EXOC3 | 377.9203467 | -0.348106183 | 0.091469159 | -3.805721913 | 0.000141391 | 0.001258206 |
| BTBD9 | 229.5389065 | 0.253954217 | 0.066744295 | 3.804882738 | 0.000141871 | 0.001261507 |
| ZNF780A | 313.4101803 | -0.38165629 | 0.100308181 | -3.804837115 | 0.000141898 | 0.001261507 |
| TMSB10 | 27375.4184 | 0.331392002 | 0.087101285 | 3.804674073 | 0.000141991 | 0.001261736 |
| HEY1 | 13.33321976 | 2.120995123 | 0.557635737 | 3.80354949 | 0.000142637 | 0.001266876 |
| NDEL1 | 281.9700708 | 0.30176907 | 0.079349142 | 3.803053951 | 0.000142923 | 0.00126881 |
| KIAA1467 | 90.14698973 | 0.523449267 | 0.137713761 | 3.800994624 | 0.000144116 | 0.001278793 |
| DPYD | 605.8292163 | -0.270765996 | 0.071279031 | -3.79867675 | 0.000145471 | 0.001290195 |
| CELSR2 | 4.560959442 | -1.582835337 | 0.41682705 | -3.797343134 | 0.000146255 | 0.001296537 |
| STIP1 | 3822.494394 | 0.210526623 | 0.055445791 | 3.796981166 | 0.000146469 | 0.001297813 |
| CHD3 | 134.6632991 | -0.375668278 | 0.09894424 | -3.79676754 | 0.000146595 | 0.001298272 |
| SLC46A2 | 83.15619879 | -0.749735471 | 0.197472486 | -3.796657876 | 0.00014666 | 0.001298272 |
| NOA1 | 309.2820541 | -0.325102677 | 0.085650056 | -3.795708846 | 0.000147222 | 0.00130263 |
| SLC38A6 | 1719.799972 | -0.379007414 | 0.099899207 | -3.793898119 | 0.000148301 | 0.001311548 |
| FZD4 | 16.91180384 | 1.204966263 | 0.317626452 | 3.793658414 | 0.000148444 | 0.001312193 |
| CBX6 | 101.5571033 | 0.51843225 | 0.136673798 | 3.793208782 | 0.000148713 | 0.001313949 |
| ACP1 | 1192.42428 | -0.139498314 | 0.036781487 | -3.792623043 | 0.000149064 | 0.001316429 |

| | | | | | | |
|---|---|---|---|---|---|---|
| GRHL1 | 58.47457871 | 0.617408877 | 0.162798744 | 3.792467077 | 0.000149158 | 0.001316632 |
| HAUS4 | 168.7387157 | -0.463144279 | 0.122129295 | -3.792245585 | 0.000149291 | 0.001317184 |
| ALCAM | 2033.058685 | 0.322292037 | 0.084991237 | 3.792061985 | 0.000149402 | 0.001317534 |
| C5orf20 | 15.0303213 | -1.477029447 | 0.389887901 | -3.788343891 | 0.000151655 | 0.001336772 |
| HIF1A | 3718.28755 | -0.476624447 | 0.125820773 | -3.788122083 | 0.00015179 | 0.001337333 |
| SPAG1 | 34.1690129 | 0.727967952 | 0.192236511 | 3.786835026 | 0.000152578 | 0.001343641 |
| DCUN1D4 | 82.49767448 | -0.622388088 | 0.164445269 | -3.784773452 | 0.000153849 | 0.001353549 |
| TPRG1 | 555.6857886 | 0.354639262 | 0.093700177 | 3.784830227 | 0.000153814 | 0.001353549 |
| SDHD | 1459.699773 | 0.317372786 | 0.083879199 | 3.783688779 | 0.000154521 | 0.001358822 |
| LOC72874: | 43.34412602 | 0.642951449 | 0.16998359 | 3.782432463 | 0.000155303 | 0.001365057 |
| ASB13 | 101.1378575 | -0.546511338 | 0.144500948 | -3.78206057 | 0.000155536 | 0.00136581 |
| IDI1 | 381.1776325 | 0.537231184 | 0.142044715 | 3.782127235 | 0.000155494 | 0.00136581 |
| CCNY | 1079.205612 | -0.238794737 | 0.063171159 | -3.780122807 | 0.000156751 | 0.001375835 |
| ARL5A | 222.611304 | -0.410895071 | 0.108724662 | -3.779226016 | 0.000157317 | 0.001380149 |
| CCND3 | 739.0072917 | 0.484550663 | 0.128303859 | 3.776586815 | 0.000158992 | 0.001394193 |
| PLAC8 | 92.44949121 | -0.968954735 | 0.256640521 | -3.775532912 | 0.000159666 | 0.001399443 |
| MAP3K7 | 204.0191033 | 0.293010665 | 0.077617913 | 3.775039222 | 0.000159982 | 0.001401558 |
| FKBP5 | 821.8852851 | 0.384611958 | 0.10189098 | 3.774739997 | 0.000160175 | 0.001402582 |
| OSGIN2 | 893.1500991 | 0.267258107 | 0.070811217 | 3.774234051 | 0.0001605 | 0.001404658 |
| SCN11A | 24.07488294 | -0.552751379 | 0.146457689 | -3.774136969 | 0.000160563 | 0.001404658 |
| DVL3 | 1004.268879 | -0.377587131 | 0.10016822 | -3.769530223 | 0.000163555 | 0.001430167 |
| TEX261 | 634.4999027 | -0.252006314 | 0.066883524 | -3.767838434 | 0.000164667 | 0.001439216 |
| HP1BP3 | 1285.193628 | -0.192641626 | 0.051147962 | -3.766359775 | 0.000165645 | 0.001447084 |
| MTIF3 | 779.3914371 | -0.214464938 | 0.056962655 | -3.765009511 | 0.000166543 | 0.001454244 |
| NFE2L2 | 1670.455607 | 0.234428693 | 0.062293509 | 3.763292481 | 0.000167691 | 0.001463584 |
| CSGALNAC | 757.1399454 | 0.248497398 | 0.066056355 | 3.761899928 | 0.000168628 | 0.00147107 |
| TMEM52 | 3.665794738 | 2.075979848 | 0.551884273 | 3.761621682 | 0.000168815 | 0.001472018 |
| RBMS1 | 1212.98381 | 0.323229154 | 0.08593288 | 3.761414199 | 0.000168955 | 0.001472551 |
| COPG1 | 2114.923311 | 0.232170603 | 0.061734375 | 3.760799438 | 0.000169371 | 0.001475485 |
| ABCA6 | 7.43472876 | 1.656242264 | 0.440495829 | 3.759949939 | 0.000169947 | 0.001479534 |
| AP1B1 | 2676.023455 | -0.291088512 | 0.077419623 | -3.759880241 | 0.000169995 | 0.001479534 |
| C20orf111 | 376.8919755 | 0.266957094 | 0.071032763 | 3.758224839 | 0.000171123 | 0.001488659 |
| USP11 | 653.0755688 | 0.38809589 | 0.1033031 | 3.756865861 | 0.000172055 | 0.001496065 |
| FAM211A | 72.16995197 | -0.559768004 | 0.149036325 | -3.755916578 | 0.000172708 | 0.001500347 |
| PXMP2 | 60.52399181 | -0.447363883 | 0.119105838 | -3.756019777 | 0.000172637 | 0.001500347 |
| ARHGAP6 | 115.7427505 | -0.506239587 | 0.134801821 | -3.755435821 | 0.00017304 | 0.00150253 |
| RELA | 816.418168 | 0.340716978 | 0.090735278 | 3.755066213 | 0.000173296 | 0.001504048 |
| FTSJD2 | 303.23658 | 0.450878614 | 0.120115728 | 3.753701716 | 0.000174242 | 0.001511559 |
| GSTK1 | 1726.16015 | -0.260166599 | 0.069322952 | -3.752964823 | 0.000174755 | 0.001515306 |
| DPP4 | 39.15603197 | 0.965296327 | 0.257235422 | 3.752579333 | 0.000175024 | 0.001516227 |
| DYNC1LI2 | 218.5725086 | 0.223155149 | 0.059466739 | 3.752604465 | 0.000175007 | 0.001516227 |
| PRSS22 | 25.1790385 | 1.0166587 | 0.270938877 | 3.752354446 | 0.000175182 | 0.001516883 |
| PTX3 | 8.69969312 | 1.359413405 | 0.362313308 | 3.752038297 | 0.000175403 | 0.001518093 |
| RBM43 | 32.50220001 | 0.659786009 | 0.175869483 | 3.751566201 | 0.000175733 | 0.001520248 |
| AMZ1 | 100.0234126 | 1.082971319 | 0.288748317 | 3.750571882 | 0.000176432 | 0.001525582 |
| CA13 | 46.56784335 | -0.553182 | 0.147541361 | -3.749335088 | 0.000177304 | 0.001532414 |
| GNA15 | 1048.142584 | 0.299457808 | 0.079976579 | 3.74431877 | 0.000180884 | 0.001562201 |

| | | | | | | |
|---|---|---|---|---|---|---|
| ZNF624 | 52.29532164 | -0.371964167 | 0.099342209 | -3.744271156 | 0.000180918 | 0.001562201 |
| FRG1 | 486.7168915 | 0.300396831 | 0.080270869 | 3.742289508 | 0.000182351 | 0.001573846 |
| NEIL2 | 40.02593293 | -0.685168436 | 0.183114856 | -3.741741384 | 0.00018275 | 0.001576553 |
| HOOK3 | 617.0774265 | -0.293950044 | 0.078628508 | -3.738466529 | 0.000185146 | 0.001596489 |
| SEC31B | 4.298983299 | -1.607315406 | 0.430134779 | -3.736771549 | 0.000186398 | 0.001606542 |
| CAV2 | 6.155324588 | 1.548511625 | 0.41445293 | 3.736278626 | 0.000186764 | 0.001608948 |
| CDK3 | 32.40972723 | 0.605929943 | 0.16219743 | 3.735755501 | 0.000187152 | 0.001611552 |
| DLST | 581.2754412 | 0.275544177 | 0.073828213 | 3.732234145 | 0.000189789 | 0.0016335 |
| MED20 | 62.07869182 | -0.329111517 | 0.088195989 | -3.731592785 | 0.000190273 | 0.001636909 |
| RPS21 | 3413.275033 | -0.352992775 | 0.094608168 | -3.731102548 | 0.000190644 | 0.001639341 |
| CAB39 | 565.5960859 | -0.255870339 | 0.06858185 | -3.730875409 | 0.000190816 | 0.001640063 |
| TNFRSF10( | 42.72411509 | -0.515833129 | 0.138281777 | -3.730304454 | 0.000191249 | 0.001643027 |
| RPS19 | 7293.075818 | -0.273462435 | 0.073312923 | -3.730071371 | 0.000191426 | 0.00164379 |
| GPR156 | 6.273160635 | 1.647458807 | 0.441759748 | 3.72930946 | 0.000192005 | 0.001648008 |
| USP25 | 940.1611838 | 0.249938433 | 0.067041017 | 3.728142022 | 0.000192897 | 0.001654897 |
| CDK19 | 437.717099 | -0.229093359 | 0.061456931 | -3.727705796 | 0.000193231 | 0.001656238 |
| TMEM194I | 33.02396031 | 0.565803753 | 0.151778728 | 3.727819835 | 0.000193143 | 0.001656238 |
| ZBTB10 | 54.96416015 | 0.405011806 | 0.108673082 | 3.726882477 | 0.000193863 | 0.001660892 |
| LINC00649 | 26.86388955 | 0.924905499 | 0.248332236 | 3.724468128 | 0.000195727 | 0.001676096 |
| FGD6 | 295.8392365 | 0.331774472 | 0.089110846 | 3.723166002 | 0.00019674 | 0.001683994 |
| HIVEP1 | 310.1248037 | 0.383057818 | 0.10292799 | 3.721609834 | 0.000197957 | 0.00169363 |
| FES | 97.54607005 | -0.549951427 | 0.147798744 | -3.720947895 | 0.000198476 | 0.001696518 |
| MUCL1 | 9.878488299 | 1.453710987 | 0.39067163 | 3.721055934 | 0.000198391 | 0.001696518 |
| ZKSCAN4 | 60.67713495 | -0.478567022 | 0.128769038 | -3.716475861 | 0.000202021 | 0.001726024 |
| WDFY1 | 746.9795611 | 0.398723503 | 0.107291517 | 3.716263095 | 0.000202191 | 0.001726686 |
| PABPC3 | 423.9844078 | -0.363188357 | 0.097746965 | -3.715597287 | 0.000202724 | 0.001730446 |
| ANAPC7 | 579.1083217 | -0.214059937 | 0.057632121 | -3.714247101 | 0.00020381 | 0.001738914 |
| LARP4 | 532.6978527 | 0.179257023 | 0.048264913 | 3.714023554 | 0.00020399 | 0.001739655 |
| HLA-DOA | 887.5277412 | -0.493050709 | 0.132764367 | -3.713727714 | 0.000204229 | 0.001740894 |
| XRN1 | 182.1319006 | 0.654361412 | 0.176264543 | 3.712382534 | 0.000205317 | 0.001749375 |
| JUN | 8.092521304 | 1.23440098 | 0.332525163 | 3.712203212 | 0.000205463 | 0.001749815 |
| SEMA4B | 135.5684835 | 0.386667729 | 0.104178034 | 3.711605146 | 0.000205949 | 0.001753155 |
| HSD3BP4 | 41.26219401 | 0.818765384 | 0.220689526 | 3.710032814 | 0.000207232 | 0.001763273 |
| RPS28 | 5576.219844 | -0.329569335 | 0.088917308 | -3.706470004 | 0.000210168 | 0.001787436 |
| ACPP | 13.73366092 | -0.896494422 | 0.241964612 | -3.705064205 | 0.000211337 | 0.001796559 |
| PRKDC | 1138.253999 | -0.208285461 | 0.056242392 | -3.703353547 | 0.000212768 | 0.001807898 |
| TNFAIP8 | 597.3531782 | 0.378805725 | 0.102314977 | 3.702348744 | 0.000213613 | 0.001814248 |
| LOC93622 | 28.24234491 | -0.636456662 | 0.171949471 | -3.70141681 | 0.000214399 | 0.001820096 |
| NIN | 321.5964907 | -0.312221298 | 0.084356154 | -3.701227283 | 0.000214559 | 0.001820626 |
| EIF4A1 | 20437.51699 | 0.285856247 | 0.077242486 | 3.700764473 | 0.000214951 | 0.00182312 |
| RABIF | 497.2664615 | 0.179579554 | 0.048537497 | 3.699810796 | 0.00021576 | 0.001829152 |
| WDHD1 | 33.62842175 | 0.673780585 | 0.182129858 | 3.699451554 | 0.000216066 | 0.00183091 |
| ZFP106 | 846.4821996 | -0.327800433 | 0.088675074 | -3.696646864 | 0.000218466 | 0.001850406 |
| CCDC176 | 8.319234144 | -1.355187341 | 0.366762954 | -3.694995166 | 0.000219891 | 0.00186163 |
| LST1 | 599.1576163 | -0.570085293 | 0.154297639 | -3.694711715 | 0.000220136 | 0.001862862 |
| GBGT1 | 289.2090538 | -0.286143264 | 0.077461035 | -3.694028413 | 0.000220729 | 0.00186703 |
| AZU1 | 3.238199418 | -2.96892443 | 0.804110196 | -3.692186025 | 0.000222335 | 0.001879757 |

| | | | | | | |
|---|---|---|---|---|---|---|
| CTAGE9 | 53.69506626 | 0.88770563 | 0.240449275 | 3.691862369 | 0.000222618 | 0.00188057 |
| FXYD2 | 37.10551051 | 0.915010391 | 0.247846337 | 3.691845525 | 0.000222633 | 0.00188057 |
| SIAH2 | 208.5205334 | 0.360217884 | 0.097587906 | 3.691214408 | 0.000223186 | 0.001884389 |
| ATP13A2 | 507.0060628 | -0.379150214 | 0.102730619 | -3.690722547 | 0.000223618 | 0.001887182 |
| EZH2 | 55.29467578 | 0.854971371 | 0.231687574 | 3.690190879 | 0.000224086 | 0.001890274 |
| R3HDM2 | 322.7883597 | -0.295561853 | 0.080106014 | -3.689633747 | 0.000224577 | 0.001893561 |
| ARHGAP32 | 58.7592258 | -0.485226494 | 0.131715052 | -3.683910721 | 0.000229683 | 0.001935734 |
| PKI55 | 51.2403265 | -0.842936509 | 0.228851127 | -3.683339993 | 0.000230198 | 0.001939198 |
| ABCG1 | 77.11110642 | -0.72773518 | 0.197676217 | -3.681450354 | 0.000231911 | 0.001952747 |
| LINC00239 | 5.863269566 | -1.342875295 | 0.364822391 | -3.680901524 | 0.000232411 | 0.001956071 |
| LDLRAD4 | 30.69041746 | -0.837504856 | 0.227550956 | -3.680515656 | 0.000232763 | 0.00195815 |
| ING4 | 114.2664757 | -0.323888587 | 0.088005634 | -3.680316486 | 0.000232945 | 0.001958795 |
| CNN2 | 1395.834087 | -0.41886721 | 0.113820346 | -3.680073232 | 0.000233167 | 0.001959781 |
| CCDC86 | 271.3473359 | -0.231552054 | 0.062923737 | -3.679884047 | 0.00023334 | 0.001960351 |
| RDH12 | 12.95462514 | -0.821797136 | 0.223351019 | -3.679397292 | 0.000233786 | 0.001963211 |
| CTNNB1 | 5782.002663 | -0.269706437 | 0.073356439 | -3.676656618 | 0.000236311 | 0.00198352 |
| GSG1 | 7.098996977 | 1.265075083 | 0.344279442 | 3.674558885 | 0.000238261 | 0.001998987 |
| HLA-H | 5.294106899 | 1.843263521 | 0.502007209 | 3.671786952 | 0.00024086 | 0.002019131 |
| RAD23B | 1273.044418 | 0.24991979 | 0.068065247 | 3.671767908 | 0.000240878 | 0.002019131 |
| LDHAL6B | 89.47603799 | 0.373809498 | 0.101817965 | 3.671351096 | 0.000241272 | 0.002021518 |
| NCF1B | 747.4708719 | 0.59100609 | 0.161049933 | 3.669707136 | 0.000242829 | 0.002033648 |
| PMEPA1 | 84.9289708 | 0.619818183 | 0.168921171 | 3.669274729 | 0.00024324 | 0.002036175 |
| EIF4G3 | 738.5576153 | -0.249274577 | 0.067955946 | -3.668179036 | 0.000244284 | 0.002044001 |
| FIG4 | 1286.949583 | -0.419420613 | 0.114353473 | -3.667755796 | 0.000244689 | 0.002046467 |
| PRKD2 | 192.1220209 | 0.318840389 | 0.086976938 | 3.665803784 | 0.000246563 | 0.002061217 |
| LAPTM5 | 43896.68671 | -0.236153877 | 0.064449107 | -3.664191608 | 0.000248121 | 0.002073312 |
| FAM125B | 204.6686965 | 0.406306521 | 0.110968406 | 3.661461289 | 0.000250781 | 0.002094598 |
| VMO1 | 779.5622681 | -0.315718498 | 0.086256394 | -3.66023298 | 0.000251986 | 0.002103722 |
| UAP1 | 229.9322203 | 0.337625704 | 0.092245851 | 3.660063838 | 0.000252152 | 0.002104168 |
| IL2RG | 2212.859267 | 0.539729721 | 0.147523898 | 3.658591768 | 0.000253605 | 0.002114396 |
| RPIA | 344.6493748 | 0.483394701 | 0.132124359 | 3.658634204 | 0.000253563 | 0.002114396 |
| ITGBL1 | 6.601776934 | 1.956705868 | 0.534957178 | 3.657686912 | 0.000254502 | 0.002119975 |
| MPST | 192.3714962 | -0.359165738 | 0.098194206 | -3.657708067 | 0.000254481 | 0.002119975 |
| ALDH2 | 13586.68698 | -0.303329803 | 0.082975508 | -3.655654687 | 0.000256526 | 0.002135887 |
| PSMC3 | 1703.089933 | 0.328458966 | 0.089925772 | 3.652556526 | 0.000259642 | 0.002160865 |
| LTB4R | 5.949832904 | -1.49655772 | 0.410079448 | -3.649433614 | 0.000262819 | 0.002186327 |
| SLC17A7 | 14.58135857 | -0.869089658 | 0.238295694 | -3.647106013 | 0.000265211 | 0.002205236 |
| CYCS | 1703.51774 | 0.372674268 | 0.102194091 | 3.646730085 | 0.000265599 | 0.002207478 |
| APOBEC3D | 11.71199355 | 1.010314585 | 0.277103891 | 3.64597762 | 0.000266377 | 0.002212961 |
| KIAA0226 | 889.7341446 | 0.291474784 | 0.079967488 | 3.644916099 | 0.000267479 | 0.002221125 |
| LRRC6 | 41.28217222 | -0.606588156 | 0.166432314 | -3.644653738 | 0.000267752 | 0.002222402 |
| TMEM150I | 1630.144728 | 0.370696809 | 0.10171992 | 3.644289228 | 0.000268132 | 0.002224562 |
| ATP1B1 | 2673.849112 | 0.225596725 | 0.061934124 | 3.642527111 | 0.000269975 | 0.002238854 |
| MPZL1 | 376.0616445 | 0.596026811 | 0.16382904 | 3.638102329 | 0.000274654 | 0.00227665 |
| DNAJC3 | 2159.639921 | 0.474682291 | 0.130512806 | 3.637055295 | 0.000275773 | 0.002284904 |
| RPL19 | 6278.929053 | -0.278161359 | 0.076493316 | -3.636413906 | 0.00027646 | 0.00228958 |
| ELL | 162.9756754 | 0.474923744 | 0.130609969 | 3.636198269 | 0.000276691 | 0.002289809 |

| | | | | | | |
|---|---|---|---|---|---|---|
| RASSF3 | 916.067083 | -0.309798578 | 0.0851994 | -3.636159134 | 0.000276733 | 0.002289809 |
| TNNI2 | 291.6092973 | -0.663623164 | 0.182520725 | -3.635878404 | 0.000277035 | 0.002291287 |
| MRI1 | 126.1351288 | -0.369083274 | 0.101585579 | -3.633225073 | 0.000279901 | 0.002313961 |
| CEP85L | 36.39628633 | -0.440736713 | 0.121407523 | -3.630225715 | 0.000283173 | 0.00233998 |
| RRP12 | 402.4094591 | 0.459424535 | 0.126585273 | 3.629367957 | 0.000284116 | 0.002346728 |
| CNTRL | 178.1448556 | -0.335769061 | 0.092541311 | -3.628315382 | 0.000285277 | 0.00235527 |
| ADK | 54.59974724 | -0.699915462 | 0.193079841 | -3.625005386 | 0.000288955 | 0.002384586 |
| SLC22A5 | 202.6876223 | -0.322810406 | 0.089073954 | -3.624071821 | 0.000290001 | 0.002392155 |
| LSM10 | 220.6013933 | -0.2784982 | 0.076872348 | -3.622865797 | 0.000291357 | 0.002401215 |
| MICAL2 | 181.3757863 | 0.715702241 | 0.197545748 | 3.622969607 | 0.00029124 | 0.002401215 |
| DNAJC4 | 381.3686457 | -0.279048049 | 0.077029417 | -3.622616675 | 0.000291638 | 0.002402467 |
| PRPF3 | 27.71859127 | 0.958161455 | 0.264539262 | 3.622000932 | 0.000292333 | 0.00240713 |
| PGM3 | 34.88256364 | 0.626080301 | 0.172900903 | 3.621035464 | 0.000293426 | 0.002415064 |
| ZNF33A | 171.8341755 | -0.402202777 | 0.111082318 | -3.620763277 | 0.000293735 | 0.002416539 |
| CCDC147 | 6.013212318 | 1.432785526 | 0.395749105 | 3.620439083 | 0.000294103 | 0.002418502 |
| CEBPD | 189.8353821 | -0.687713107 | 0.189993671 | -3.619663241 | 0.000294987 | 0.002424694 |
| PI3 | 4.059780876 | 2.814161201 | 0.777565758 | 3.619193842 | 0.000295522 | 0.002428025 |
| SLC9A7 | 1013.10094 | 0.33539277 | 0.09267632 | 3.618969451 | 0.000295779 | 0.00242906 |
| PPAP2A | 156.1801292 | -0.408393925 | 0.112865339 | -3.618417555 | 0.00029641 | 0.002433173 |
| SNHG15 | 24.88075388 | 0.577363682 | 0.159695814 | 3.615396465 | 0.000299888 | 0.002460642 |
| IQSEC2 | 77.45912486 | -0.486685764 | 0.134646491 | -3.614544722 | 0.000300876 | 0.002467659 |
| WDR17 | 6.356402468 | -1.38447474 | 0.383073507 | -3.614122915 | 0.000301366 | 0.002470592 |
| PPIP5K1 | 199.0581271 | -0.392129206 | 0.108530821 | -3.613067713 | 0.000302596 | 0.002479581 |
| OPTN | 1005.897793 | 0.862853659 | 0.238849594 | 3.612539775 | 0.000303213 | 0.002483544 |
| PEF1 | 712.9968843 | 0.270594721 | 0.074921562 | 3.611706887 | 0.000304188 | 0.002490441 |
| SMAD5 | 220.6796198 | -0.28054074 | 0.077717774 | -3.609737213 | 0.000306507 | 0.002508326 |
| SMOX | 53.49873954 | 0.753277074 | 0.208871855 | 3.606407741 | 0.000310465 | 0.0025396 |
| USP28 | 256.8667048 | 0.271357393 | 0.075267125 | 3.60525786 | 0.000311843 | 0.002549752 |
| EYA3 | 644.3538608 | 0.385802596 | 0.107019488 | 3.60497517 | 0.000312183 | 0.002550291 |
| SSBP3 | 532.9480321 | 0.360249908 | 0.099930864 | 3.604991421 | 0.000312163 | 0.002550291 |
| HPGDS | 384.4545785 | -0.564975402 | 0.156738524 | -3.604572689 | 0.000312667 | 0.002552827 |
| INF2 | 811.5368887 | 0.242577861 | 0.067298813 | 3.604489452 | 0.000312767 | 0.002552827 |
| MLEC | 738.0374588 | -0.485717473 | 0.134792128 | -3.603455778 | 0.000314014 | 0.002561883 |
| TLK2 | 862.989773 | 0.342982067 | 0.095196053 | 3.602902183 | 0.000314684 | 0.002566224 |
| FAM200B | 138.7256659 | -0.435849005 | 0.121002283 | -3.601989917 | 0.000315791 | 0.002573425 |
| KIAA0494 | 681.8783453 | -0.271662547 | 0.075421038 | -3.601946563 | 0.000315843 | 0.002573425 |
| NUP50 | 565.2161492 | 0.181201102 | 0.050309879 | 3.601700223 | 0.000316143 | 0.00257474 |
| MKL2 | 288.5387952 | -0.273134252 | 0.075854643 | -3.600758498 | 0.00031729 | 0.002582801 |
| WDR7 | 406.5587316 | -0.190907034 | 0.053020003 | -3.600660557 | 0.00031741 | 0.002582801 |
| PIWIL4 | 12.47060475 | 1.055067847 | 0.293079417 | 3.599938396 | 0.000318293 | 0.002588854 |
| RAPGEFL1 | 173.9754977 | -0.500098018 | 0.139031142 | -3.597021589 | 0.000321882 | 0.002616904 |
| PER2 | 68.53749511 | 0.391223157 | 0.108831242 | 3.594768829 | 0.00032468 | 0.002638501 |
| LOC55310[illegible] | 29.65454078 | -0.58036842 | 0.161490737 | -3.59381864 | 0.000325867 | 0.002646992 |
| CCDC149 | 53.19094572 | -0.64050466 | 0.178267781 | -3.592935618 | 0.000326973 | 0.002654825 |
| SIVA1 | 165.65844 | -0.375268168 | 0.104489728 | -3.591435984 | 0.000328861 | 0.002668987 |
| TYROBP | 42958.59535 | -0.258370658 | 0.071952998 | -3.590825488 | 0.000329632 | 0.002674083 |
| ABRACL | 986.2827456 | -0.246028862 | 0.06851847 | -3.590694032 | 0.000329799 | 0.002674268 |

| | | | | | | |
|---|---|---|---|---|---|---|
| UBE2D2 | 728.9351584 | 0.301767737 | 0.08406684 | 3.589616749 | 0.000331164 | 0.002684175 |
| FAM102B | 335.8728019 | -0.2345986 | 0.065379352 | -3.588267454 | 0.000332883 | 0.002696929 |
| PRRC2C | 1493.462576 | 0.305295056 | 0.085105178 | 3.587267719 | 0.000334161 | 0.00270611 |
| HSPB7 | 26.79316617 | 1.330753407 | 0.37106685 | 3.586290195 | 0.000335416 | 0.002715089 |
| EIF2S1 | 392.00864 | 0.241117578 | 0.067273024 | 3.584164416 | 0.000338159 | 0.002736107 |
| AMZ2P1 | 14.56171433 | -1.029756251 | 0.287318226 | -3.584026904 | 0.000338337 | 0.002736361 |
| COX6A1 | 5833.312987 | 0.178533624 | 0.049817753 | 3.583734962 | 0.000338716 | 0.002738234 |
| GGH | 272.3275099 | 0.347711311 | 0.097148394 | 3.579177123 | 0.000344678 | 0.002785224 |
| SGK223 | 182.5571157 | -0.480182232 | 0.134173807 | -3.578807546 | 0.000345166 | 0.002787956 |
| PRKD3 | 457.3847043 | 0.224754665 | 0.062835539 | 3.576871762 | 0.000347731 | 0.002807458 |
| TMEM159 | 162.5891301 | -0.464792471 | 0.129953584 | -3.576603706 | 0.000348087 | 0.00280912 |
| PSMD14 | 1208.162576 | 0.255342944 | 0.071408369 | 3.575812547 | 0.000349142 | 0.00281641 |
| HCG26 | 44.38404294 | 0.573833032 | 0.160495937 | 3.575374204 | 0.000349727 | 0.002819913 |
| GTF3A | 2323.323503 | -0.282022795 | 0.07888364 | -3.575174709 | 0.000349994 | 0.002820413 |
| QPRT | 38.02134687 | -0.600063211 | 0.167845085 | -3.575101476 | 0.000350092 | 0.002820413 |
| RALGAPA2 | 205.3675596 | -0.262088702 | 0.073350485 | -3.573101173 | 0.000352778 | 0.002840826 |
| MAP2K3 | 477.9501075 | 0.684450386 | 0.191571124 | 3.572826479 | 0.000353149 | 0.002842581 |
| RAB18 | 985.8103117 | 0.220311663 | 0.061719816 | 3.569545045 | 0.000357602 | 0.002877181 |
| STK39 | 191.0749663 | -0.697380878 | 0.195381657 | -3.569326258 | 0.0003579 | 0.002878342 |
| COX10 | 442.1880325 | 0.267501149 | 0.074958903 | 3.568637438 | 0.000358843 | 0.002884673 |
| BTN3A2 | 514.8698495 | 0.469491783 | 0.131698665 | 3.564894013 | 0.000364003 | 0.002924897 |
| AAMDC | 82.99071791 | -0.424832849 | 0.119218283 | -3.563487381 | 0.00036596 | 0.002938811 |
| SLC2A8 | 45.77099838 | -1.355221267 | 0.380314463 | -3.563422899 | 0.00036605 | 0.002938811 |
| CENPV | 205.2002156 | -0.316183117 | 0.088744103 | -3.562863393 | 0.000366832 | 0.002943816 |
| KLF7 | 132.4657327 | -0.66266711 | 0.186135369 | -3.560135373 | 0.000370664 | 0.002972688 |
| TMEM110 | 357.9996061 | 0.492010421 | 0.138202251 | 3.560075309 | 0.000370748 | 0.002972688 |
| MN1 | 13.1466213 | 3.810022428 | 1.070601986 | 3.55876645 | 0.000372601 | 0.002986254 |
| SEC61A1 | 3919.538702 | 0.359649228 | 0.101150315 | 3.555591788 | 0.000377129 | 0.00302125 |
| BCL2 | 184.5783224 | 0.541811264 | 0.15241437 | 3.554856818 | 0.000378185 | 0.003028405 |
| DCLK2 | 11.13405406 | -0.988069795 | 0.278061424 | -3.553422772 | 0.000380253 | 0.003043656 |
| CCNA1 | 10.10824767 | 1.291793585 | 0.363560355 | 3.553175059 | 0.000380611 | 0.003045216 |
| C3orf18 | 87.62427291 | 0.39526782 | 0.111279529 | 3.55202636 | 0.000382277 | 0.003057228 |
| KBTBD3 | 31.99018981 | -0.592838118 | 0.166910318 | -3.551836242 | 0.000382553 | 0.003058125 |
| LRP3 | 32.87033717 | -0.768171333 | 0.216395982 | -3.549841016 | 0.000385464 | 0.003078754 |
| TTC7A | 442.7289329 | -0.367610822 | 0.10355439 | -3.549929871 | 0.000385334 | 0.003078754 |
| EIF2AK4 | 579.3916796 | -0.234948131 | 0.066191223 | -3.549536003 | 0.000385911 | 0.003081002 |
| SOAT1 | 11712.10773 | 0.326053352 | 0.091866065 | 3.5492252 | 0.000386366 | 0.00308332 |
| ERLIN2 | 124.5463274 | -0.31017781 | 0.087396412 | -3.549090883 | 0.000386564 | 0.003083573 |
| DEPDC1 | 25.30567024 | 0.888625153 | 0.250405947 | 3.548738211 | 0.000387082 | 0.003085484 |
| DIS3L | 322.885343 | -0.349627067 | 0.09852251 | -3.54870239 | 0.000387134 | 0.003085484 |
| TPP1 | 11037.29127 | -0.233434446 | 0.065789008 | -3.548228711 | 0.000387831 | 0.003089717 |
| ARRDC4 | 654.6575063 | 0.542322034 | 0.152851934 | 3.54802205 | 0.000388136 | 0.003090821 |
| ACTA2 | 225.7034075 | 0.368774306 | 0.103984316 | 3.54644162 | 0.000390471 | 0.003106764 |
| XIRP1 | 6.337441979 | 1.943906302 | 0.548123131 | 3.546477411 | 0.000390418 | 0.003106764 |
| EBPL | 70.61066263 | -0.538289681 | 0.151790414 | -3.546269268 | 0.000390727 | 0.00310747 |
| LACC1 | 246.8767658 | -0.769676019 | 0.217048857 | -3.546095698 | 0.000390984 | 0.003108191 |
| GATSL2 | 18.82815504 | -0.701152064 | 0.197746896 | -3.545704529 | 0.000391565 | 0.003111481 |

| | | | | | | |
|---|---|---|---|---|---|---|
| KDM3A | 139.0276055 | -0.342410639 | 0.096577812 | -3.545437948 | 0.000391961 | 0.003113302 |
| USP30 | 276.7990181 | -0.431034152 | 0.121596234 | -3.544798531 | 0.000392913 | 0.003119534 |
| PAG1 | 714.9718447 | 0.359105506 | 0.101313847 | 3.544485933 | 0.000393379 | 0.003121905 |
| CD247 | 18.85136786 | 0.92004989 | 0.259650133 | 3.543421603 | 0.000394971 | 0.003133199 |
| FKBP14 | 77.00313821 | -0.373213159 | 0.105353048 | -3.542499863 | 0.000396354 | 0.003142831 |
| ARHGAP27 | 196.8938499 | -0.311444462 | 0.087942148 | -3.541469816 | 0.000397904 | 0.003153786 |
| NIPAL1 | 4.234878953 | 1.934154191 | 0.546227526 | 3.540931388 | 0.000398717 | 0.003158885 |
| SH3YL1 | 83.52738341 | -0.544496863 | 0.153830856 | -3.539581565 | 0.000400762 | 0.003173735 |
| SPATA6 | 21.95001978 | -0.715016389 | 0.202032997 | -3.539106977 | 0.000401483 | 0.003178095 |
| FANCC | 76.36088329 | 0.444103136 | 0.125502671 | 3.538595102 | 0.000402262 | 0.003182912 |
| PHACTR2 | 15.45523713 | 0.845503043 | 0.239119696 | 3.535898786 | 0.00040639 | 0.00321421 |
| SLC12A2 | 86.07612206 | -0.404937766 | 0.114569261 | -3.534436393 | 0.000408646 | 0.003230677 |
| TMPO | 70.26004009 | 0.60795644 | 0.172075106 | 3.533087702 | 0.000410736 | 0.003245826 |
| CCDC71L | 192.6107168 | 0.670023358 | 0.189666823 | 3.532633425 | 0.000411443 | 0.00325003 |
| PIR | 232.5088366 | 0.349753354 | 0.09901227 | 3.532424346 | 0.000411768 | 0.003251223 |
| HIST1H2BE | 91.48457135 | 0.627617101 | 0.177690538 | 3.532079461 | 0.000412305 | 0.003254087 |
| CBWD2 | 266.6391484 | 0.258157852 | 0.073129984 | 3.530123204 | 0.000415366 | 0.003276856 |
| FXR2 | 824.0223866 | 0.19260757 | 0.054571588 | 3.529447754 | 0.000416428 | 0.003283842 |
| TGFBR2 | 262.0076597 | -0.687496402 | 0.194922712 | -3.527020504 | 0.000420264 | 0.003312692 |
| DCSTAMP | 6344.46666 | 0.40612287 | 0.115187154 | 3.5257653 | 0.000422261 | 0.003327024 |
| LPAR1 | 644.5511867 | 0.400119367 | 0.113513058 | 3.524875232 | 0.000423682 | 0.003336811 |
| PSPC1 | 92.54027625 | 0.468739047 | 0.133104826 | 3.521578143 | 0.000428986 | 0.003377156 |
| ITGA7 | 10.64070528 | 1.219619609 | 0.34646734 | 3.520157511 | 0.000431291 | 0.003393864 |
| LILRA3 | 94.25855599 | 0.970614281 | 0.275830594 | 3.518878266 | 0.000433376 | 0.003407443 |
| LOC37444 | 135.873391 | 0.329037454 | 0.093506453 | 3.518874312 | 0.000433382 | 0.003407443 |
| TMEM189 | 1735.318863 | 0.284480334 | 0.080877407 | 3.517426488 | 0.000435753 | 0.00342464 |
| KCNA3 | 295.5282135 | 0.408487283 | 0.116245568 | 3.514003076 | 0.000441408 | 0.003467619 |
| ZNF124 | 529.702189 | -0.43879772 | 0.124912488 | -3.512841081 | 0.000443343 | 0.003481351 |
| TRIM61 | 31.89907038 | -0.695492235 | 0.198039352 | -3.511889066 | 0.000444934 | 0.003492373 |
| CREB3L2 | 612.3852215 | -0.209780088 | 0.059742712 | -3.511392132 | 0.000445766 | 0.003497436 |
| UBE2O | 189.8288831 | 0.274524816 | 0.07819896 | 3.510594212 | 0.000447106 | 0.003506473 |
| SNORA70G | 569.4661735 | 0.600804207 | 0.17129039 | 3.507518481 | 0.000452307 | 0.003545767 |
| RNF166 | 157.6326948 | -0.617926395 | 0.176271156 | -3.505544587 | 0.000455674 | 0.003570661 |
| NECAP1 | 834.0320829 | 0.163704455 | 0.046700354 | 3.505422144 | 0.000455884 | 0.003570802 |
| CLU | 60.80214161 | 0.56571478 | 0.161420887 | 3.504594677 | 0.000457303 | 0.003580413 |
| FLJ27354 | 32.83986204 | -0.617280307 | 0.176210147 | -3.503091726 | 0.000459891 | 0.003599164 |
| ERMAP | 97.02093769 | -0.485378189 | 0.138566417 | -3.502855889 | 0.000460298 | 0.00360084 |
| BAZ1A | 305.496684 | 0.387993148 | 0.110787487 | 3.502138718 | 0.000461539 | 0.003609032 |
| TIMM8A | 65.67058942 | -0.411874404 | 0.117619515 | -3.501752265 | 0.000462209 | 0.003612754 |
| CXADR | 143.718531 | -0.508572075 | 0.145268458 | -3.500911914 | 0.000463669 | 0.003622645 |
| GPD1L | 40.42597442 | -0.463815832 | 0.132511992 | -3.50018005 | 0.000464944 | 0.003631084 |
| FAM216A | 10.27445871 | -1.091709288 | 0.311962318 | -3.499490883 | 0.000466148 | 0.003638958 |
| ARL6IP4 | 412.9347774 | -0.282010052 | 0.080591694 | -3.499244638 | 0.000466578 | 0.003640796 |
| SUPT5H | 1330.993639 | 0.329215991 | 0.094086818 | 3.499066044 | 0.000466891 | 0.00364171 |
| CORO1A | 246.2775063 | -0.70046789 | 0.200236436 | -3.498203939 | 0.000468403 | 0.003651974 |
| TBC1D8 | 447.4684359 | -0.546253922 | 0.156214567 | -3.496818068 | 0.000470843 | 0.003669463 |
| PSEN1 | 2085.911331 | 0.309411764 | 0.088493768 | 3.496424336 | 0.000471538 | 0.003673346 |

| | | | | | | |
|---|---|---|---|---|---|---|
| CDK18 | 549.1089327 | 0.27874686 | 0.079734421 | 3.495941375 | 0.000472392 | 0.003678463 |
| SRCRB4D | 50.19014982 | 0.789294868 | 0.225803867 | 3.495488709 | 0.000473194 | 0.003683169 |
| NBPF1 | 286.1053605 | -0.837530444 | 0.239801204 | -3.492603167 | 0.000478337 | 0.00372164 |
| IPPK | 102.9461299 | 0.40179665 | 0.115127005 | 3.490029555 | 0.000482967 | 0.003756097 |
| MYOM1 | 25.29288252 | -1.007913825 | 0.288852295 | -3.489374472 | 0.000484152 | 0.003763745 |
| PCM1 | 340.1613012 | -0.257272196 | 0.073770169 | -3.487482791 | 0.00048759 | 0.00378889 |
| FLJ46906 | 25.0453884 | -0.643648583 | 0.184589664 | -3.486915619 | 0.000488625 | 0.003795117 |
| PSMB3 | 2339.317754 | 0.244135783 | 0.070016731 | 3.486820656 | 0.000488799 | 0.003795117 |
| ZFAND1 | 266.7498923 | -0.2513385 | 0.072092872 | -3.486315535 | 0.000489723 | 0.003800707 |
| COL6A2 | 992.6880093 | 0.631276562 | 0.1811555 | 3.484722042 | 0.000492649 | 0.00382182 |
| RAB21 | 699.2758212 | 0.258957869 | 0.074317529 | 3.484478986 | 0.000493096 | 0.003823701 |
| CLIC4 | 4563.289085 | 0.526447637 | 0.151089397 | 3.48434535 | 0.000493342 | 0.00382402 |
| SZRD1 | 2344.315219 | 0.41122793 | 0.118042513 | 3.483727332 | 0.000494483 | 0.003831266 |
| LTB4R2 | 9.427848384 | -1.251953798 | 0.359555031 | -3.481953224 | 0.000497771 | 0.003854164 |
| TRIM16L | 22.38777784 | 0.744931758 | 0.213943457 | 3.481909513 | 0.000497852 | 0.003854164 |
| CASP4 | 2333.194955 | 0.246534007 | 0.070832056 | 3.480542855 | 0.000500399 | 0.003872272 |
| RABGAP1L | 377.8790185 | -0.326555188 | 0.093869112 | -3.478835373 | 0.000503598 | 0.00389541 |
| CCL7 | 148.0370375 | 0.702598754 | 0.202064178 | 3.47710694 | 0.000506856 | 0.003916853 |
| IDH1 | 2015.346094 | -0.222032076 | 0.063856648 | -3.477039324 | 0.000506983 | 0.003916853 |
| RASA3 | 360.8896395 | -0.377522206 | 0.108576048 | -3.477030264 | 0.000507001 | 0.003916853 |
| PLTP | 1546.396998 | 0.47754712 | 0.137389692 | 3.475858444 | 0.000509221 | 0.003930747 |
| TMEM50B | 57.58644968 | -0.31508452 | 0.090648883 | -3.475878671 | 0.000509183 | 0.003930747 |
| DNAH17 | 22.99130775 | -1.210792218 | 0.348776962 | -3.471537259 | 0.000517488 | 0.003992904 |
| PIP4K2B | 648.2746081 | -0.371627806 | 0.107058647 | -3.471254463 | 0.000518033 | 0.003995457 |
| ERN1 | 125.5079259 | 0.408976845 | 0.117932764 | 3.46788145 | 0.000524579 | 0.004040925 |
| RAB35 | 1295.801046 | 0.165998601 | 0.047867337 | 3.467888759 | 0.000524564 | 0.004040925 |
| SLFN12L | 4.464229014 | 1.756853679 | 0.50657967 | 3.468069848 | 0.000524211 | 0.004040925 |
| JTB | 1364.975649 | -0.186324629 | 0.053734157 | -3.467526814 | 0.000525271 | 0.004043812 |
| TPPP | 13.93067397 | -1.470594407 | 0.424111962 | -3.467467413 | 0.000525387 | 0.004043812 |
| TMEM45B | 23.72994867 | 0.790778016 | 0.228114068 | 3.466590305 | 0.000527105 | 0.004055354 |
| GNAI1 | 19.96209393 | 0.814357632 | 0.234995322 | 3.465420615 | 0.000529403 | 0.004071354 |
| GPRC5B | 41.4119547 | -1.124988043 | 0.324741249 | -3.464259761 | 0.000531693 | 0.004087279 |
| TDP2 | 3894.491461 | 0.218909112 | 0.063202141 | 3.463634427 | 0.00053293 | 0.004095102 |
| RALGAPB | 645.9550173 | 0.204678402 | 0.059096887 | 3.463437981 | 0.00053332 | 0.004096404 |
| UQCRC2 | 1190.90835 | -0.243514453 | 0.070316829 | -3.463103435 | 0.000533983 | 0.004099811 |
| COG8 | 218.0745784 | -0.327993066 | 0.094721744 | -3.462700878 | 0.000534783 | 0.004103025 |
| ERAP2 | 496.9066522 | 0.626671416 | 0.180979202 | 3.462670889 | 0.000534842 | 0.004103025 |
| MYO15A | 29.179132 | 0.74795845 | 0.216052112 | 3.461935374 | 0.000536306 | 0.004112562 |
| NUBPL | 49.41363017 | -0.789696373 | 0.228124217 | -3.461694608 | 0.000536786 | 0.004114549 |
| LOC10013: | 54.91418113 | -0.548916435 | 0.158665786 | -3.459576571 | 0.000541025 | 0.004144436 |
| NPTXR | 7.192389782 | -1.154067802 | 0.333591442 | -3.459524608 | 0.00054113 | 0.004144436 |
| DRD5 | 29.51971371 | -0.665389306 | 0.192346836 | -3.459320263 | 0.000541541 | 0.004145878 |
| GAK | 1130.484033 | 0.213950642 | 0.061856692 | 3.458811585 | 0.000542564 | 0.00415201 |
| SLC25A46 | 668.3133239 | -0.202705713 | 0.058609862 | -3.458559781 | 0.000543072 | 0.004154187 |
| ITPR3 | 50.1269576 | 0.671050515 | 0.194095852 | 3.457315072 | 0.000545587 | 0.00417 |
| PKIA | 106.600551 | 0.487684399 | 0.141054305 | 3.457422996 | 0.000545368 | 0.00417 |
| LYRM1 | 89.45233204 | 0.43519595 | 0.125964232 | 3.454916852 | 0.000550463 | 0.004205543 |

| | | | | | | |
|---|---|---|---|---|---|---|
| PAQR5 | 1073.950507 | 0.338390946 | 0.097962217 | 3.454300604 | 0.000551722 | 0.004213438 |
| RPL36 | 2736.290728 | -0.241665661 | 0.069992894 | -3.452717086 | 0.000554971 | 0.004236511 |
| ZNF780B | 222.9273113 | -0.344243039 | 0.099717278 | -3.4521905 | 0.000556055 | 0.00424305 |
| ANGEL1 | 754.7347817 | -0.313970912 | 0.091010849 | -3.44981851 | 0.000560964 | 0.004278753 |
| RTN4R | 140.7482178 | -0.391853489 | 0.11359565 | -3.44954662 | 0.000561529 | 0.004281312 |
| CYGB | 378.7839614 | -0.496441786 | 0.143933094 | -3.449114957 | 0.000562427 | 0.004286408 |
| GALNT11 | 349.165986 | -0.243419418 | 0.070598171 | -3.447956451 | 0.000564845 | 0.004303076 |
| UBA52 | 3405.994981 | -0.218753847 | 0.063459899 | -3.447119379 | 0.000566598 | 0.004314667 |
| C16orf88 | 69.86235572 | -0.421462567 | 0.12228836 | -3.446465119 | 0.000567972 | 0.004323361 |
| DLGAP4 | 13.97004402 | 0.842519682 | 0.244528607 | 3.445485145 | 0.000570035 | 0.004333822 |
| DUSP22 | 445.9626032 | -0.31463526 | 0.091312406 | -3.445701117 | 0.00056958 | 0.004333822 |
| PROS1 | 350.6517369 | -0.345743138 | 0.10034684 | -3.445481058 | 0.000570044 | 0.004333822 |
| SMAD2 | 927.5262263 | -0.196112571 | 0.056920804 | -3.445358399 | 0.000570303 | 0.004334021 |
| RAB3D | 22.19029521 | -0.603112413 | 0.175163353 | -3.443142657 | 0.000574996 | 0.004367906 |
| HIP1R | 4.580865835 | 1.840105647 | 0.534911736 | 3.440017341 | 0.000581677 | 0.004413356 |
| SLC30A7 | 707.0864549 | 0.334796577 | 0.097323859 | 3.440025714 | 0.000581659 | 0.004413356 |
| VAV3 | 92.78539862 | -0.353649718 | 0.102804808 | -3.440011483 | 0.00058169 | 0.004413356 |
| PITPNA | 942.4409725 | 0.228832619 | 0.06652829 | 3.439628766 | 0.000582513 | 0.004417802 |
| GARNL3 | 5.74140258 | -1.772016639 | 0.515256286 | -3.439097566 | 0.000583657 | 0.004424679 |
| TRAPPC5 | 561.8576174 | -0.272882181 | 0.079367528 | -3.438209386 | 0.000585575 | 0.004437413 |
| RPL7A | 3130.384577 | -0.303927072 | 0.088399932 | -3.438091709 | 0.000585829 | 0.004437538 |
| LEPRE1 | 211.6517322 | -0.413786611 | 0.120450412 | -3.435327487 | 0.000591838 | 0.004481232 |
| PCGF2 | 7.123664977 | 1.159421004 | 0.33752899 | 3.435026442 | 0.000592496 | 0.004484391 |
| SCRN2 | 25.65090695 | -0.836803992 | 0.243644419 | -3.434529699 | 0.000593583 | 0.004490794 |
| CLEC4D | 30.97431141 | 1.091071974 | 0.317750613 | 3.433736804 | 0.000595322 | 0.004502122 |
| C3orf52 | 1.414820388 | 2.990080959 | 0.871020723 | 3.432847094 | 0.000597279 | 0.004515089 |
| PITPNC1 | 291.8869029 | -0.273484537 | 0.079675351 | -3.432486126 | 0.000598074 | 0.00451927 |
| PTP4A2 | 1271.02479 | -0.233017488 | 0.067891601 | -3.432199059 | 0.000598708 | 0.004522223 |
| CST2 | 48.62749531 | -1.509324804 | 0.439794415 | -3.431887154 | 0.000599397 | 0.004523865 |
| LIG3 | 121.5838096 | 0.24883225 | 0.072506088 | 3.431880811 | 0.000599411 | 0.004523865 |
| NR4A3 | 343.0124582 | 0.519624435 | 0.151472889 | 3.430478153 | 0.000602518 | 0.004545478 |
| CPPED1 | 270.7792331 | -0.306623759 | 0.089397574 | -3.42988904 | 0.000603828 | 0.004553514 |
| ANKRD33B | 3.367125919 | 2.231601687 | 0.650907299 | 3.428447786 | 0.000607043 | 0.004575908 |
| LY6G5C | 43.68615596 | -0.571580102 | 0.167009842 | -3.422433648 | 0.000620632 | 0.004676452 |
| ADIPOR2 | 667.7500559 | 0.343059743 | 0.100273768 | 3.421231206 | 0.000623383 | 0.004695279 |
| UQCR11 | 1999.034169 | -0.217116191 | 0.063469935 | -3.420772238 | 0.000624436 | 0.00470131 |
| SHPK | 231.9487035 | 0.555260318 | 0.162359188 | 3.419950076 | 0.000626326 | 0.004713637 |
| SLC47A1 | 1130.987173 | -0.557181239 | 0.162945264 | -3.419438076 | 0.000627506 | 0.004720609 |
| LAMB1 | 195.9827876 | 0.441600119 | 0.129150539 | 3.419266547 | 0.000627902 | 0.00472168 |
| SART3 | 418.101105 | 0.247579546 | 0.072457776 | 3.416880287 | 0.000633431 | 0.004761339 |
| PSMD12 | 486.8365368 | 0.259401963 | 0.07594065 | 3.415851218 | 0.00063583 | 0.00477744 |
| HERPUD2 | 655.6356549 | 0.233587388 | 0.068396828 | 3.415178651 | 0.000637402 | 0.004787323 |
| GMEB2 | 82.5999584 | 0.326251428 | 0.095564797 | 3.413928974 | 0.000640333 | 0.004807397 |
| MAP4K1 | 84.38696733 | -0.580115898 | 0.169972268 | -3.413003217 | 0.000642512 | 0.004821815 |
| TMEM229B | 114.1772494 | 0.503795286 | 0.147635764 | 3.412420345 | 0.000643887 | 0.004830192 |
| ABHD16A | 642.6783601 | 0.287001695 | 0.084116032 | 3.41197379 | 0.000644943 | 0.00483422 |
| JOSD1 | 625.4817633 | 0.20568795 | 0.060283323 | 3.412020764 | 0.000644832 | 0.00483422 |

| | | | | | | |
|---|---|---|---|---|---|---|
| SWT1 | 21.80482645 | 0.602111749 | 0.176477271 | 3.411837378 | 0.000645266 | 0.004834695 |
| SLC7A6 | 36.46535355 | -0.526894486 | 0.154538173 | -3.409477903 | 0.000650874 | 0.00487475 |
| MYL12B | 5714.739014 | 0.189607314 | 0.055646271 | 3.40736783 | 0.000655927 | 0.004909808 |
| TGFB1 | 4416.315199 | 0.180236572 | 0.05289713 | 3.407303428 | 0.000656081 | 0.004909808 |
| MED21 | 138.516686 | -0.260437591 | 0.076471063 | -3.405701216 | 0.000659944 | 0.004936728 |
| GDE1 | 875.8124408 | -0.305599965 | 0.089749619 | -3.405027978 | 0.000661573 | 0.004940981 |
| TIMP2 | 8406.170548 | -0.265018913 | 0.077829654 | -3.405114887 | 0.000661362 | 0.004940981 |
| TTC39A | 4.909149958 | 2.74201322 | 0.805219899 | 3.405297388 | 0.00066092 | 0.004940981 |
| TXNDC16 | 26.44827289 | -0.730152269 | 0.214424237 | -3.405176016 | 0.000661214 | 0.004940981 |
| CHID1 | 544.0527222 | -0.295118757 | 0.086688227 | -3.404369499 | 0.00066317 | 0.004950924 |
| HPS4 | 278.0632481 | -0.508240534 | 0.149300271 | -3.404150111 | 0.000663703 | 0.004950935 |
| SH3BGRL | 4042.795272 | -0.299201607 | 0.08789304 | -3.404155856 | 0.000663689 | 0.004950935 |
| MRPL39 | 284.1234563 | 0.195619704 | 0.057470685 | 3.403817197 | 0.000664512 | 0.004953007 |
| MXD1 | 263.1444048 | 0.676207722 | 0.198657748 | 3.403882956 | 0.000664352 | 0.004953007 |
| PTGER3 | 41.24923018 | 0.755355744 | 0.221977395 | 3.402849848 | 0.000666869 | 0.004968589 |
| NPLOC4 | 1129.447544 | 0.340077335 | 0.09996323 | 3.402024282 | 0.000668887 | 0.004979727 |
| SLC20A1 | 1667.063875 | 0.263619689 | 0.077489176 | 3.402019532 | 0.000668899 | 0.004979727 |
| PAPSS1 | 1014.816001 | 0.380552447 | 0.111901444 | 3.400782284 | 0.000671933 | 0.005000321 |
| RAD51B | 11.16754188 | 0.759665274 | 0.223413892 | 3.400259794 | 0.000673219 | 0.005007886 |
| RHOH | 30.48780775 | 1.050192901 | 0.308967763 | 3.399037135 | 0.000676235 | 0.00502832 |
| DDIT4L | 4.91965562 | -1.201807614 | 0.353600794 | -3.398769554 | 0.000676897 | 0.005031234 |
| OLFML2B | 36.79618775 | -0.728857405 | 0.21450619 | -3.397838566 | 0.000679205 | 0.005046373 |
| USP13 | 249.4557917 | 0.525940854 | 0.154859238 | 3.396251089 | 0.000683157 | 0.005073711 |
| TRIM9 | 8.704329625 | 1.281750219 | 0.37747192 | 3.395617399 | 0.00068474 | 0.005081642 |
| ZNF136 | 125.334512 | -0.222425541 | 0.065503941 | -3.395605456 | 0.00068477 | 0.005081642 |
| CCDC134 | 177.3476995 | 0.308091087 | 0.090826186 | 3.39209537 | 0.000693603 | 0.005145142 |
| PTOV1 | 674.8039155 | -0.263251091 | 0.077614365 | -3.391783096 | 0.000694394 | 0.00514896 |
| SEL1L3 | 370.663341 | -0.483638693 | 0.142622508 | -3.391040439 | 0.000696278 | 0.005160879 |
| CCDC102B | 23.48728943 | 0.748434458 | 0.220717774 | 3.390911589 | 0.000696606 | 0.005161253 |
| TNFAIP8L2 | 1130.322202 | -0.236767814 | 0.069829744 | -3.390644149 | 0.000697286 | 0.005164238 |
| MTMR11 | 6.331827137 | 1.82263786 | 0.537645908 | 3.390033909 | 0.00069884 | 0.005173691 |
| MEP1A | 38.97363566 | 0.559656013 | 0.165117843 | 3.38943389 | 0.000700371 | 0.005182967 |
| SH3KBP1 | 72.05719302 | -0.433381211 | 0.127890124 | -3.388699578 | 0.000702249 | 0.005194803 |
| WASH2P | 12.42937628 | -0.794681772 | 0.234611446 | -3.387225066 | 0.000706034 | 0.005220732 |
| LOC10050( | 27.71354734 | -0.628521585 | 0.185675336 | -3.385056945 | 0.000711635 | 0.005260057 |
| EXOC3L4 | 7.075671904 | 1.487147138 | 0.439398816 | 3.384504199 | 0.000713069 | 0.005266482 |
| GMPR2 | 399.4493567 | -0.373927092 | 0.110479723 | -3.384576658 | 0.000712881 | 0.005266482 |
| ITLN1 | 5.221856207 | -1.853812059 | 0.547975289 | -3.383021274 | 0.000716931 | 0.005292903 |
| MVK | 96.83510584 | 0.550482848 | 0.162760071 | 3.382173812 | 0.000719146 | 0.005307157 |
| KIF21A | 12.78385528 | 1.035126764 | 0.306102519 | 3.381634256 | 0.00072056 | 0.005315486 |
| MAN1C1 | 39.80609732 | -0.485168741 | 0.143480373 | -3.381429336 | 0.000721098 | 0.005317347 |
| IQCA1 | 5.428591495 | -1.79822286 | 0.531831307 | -3.381190306 | 0.000721725 | 0.00531987 |
| TAF10 | 865.8213484 | -0.247042993 | 0.073101566 | -3.379448721 | 0.000726314 | 0.005351572 |
| NDOR1 | 100.2841288 | 0.282523747 | 0.083604892 | 3.379272891 | 0.000726778 | 0.00535288 |
| TFPI | 395.5496155 | 0.404561093 | 0.11973465 | 3.378813842 | 0.000727993 | 0.005359707 |
| MGMT | 147.7627338 | -0.349061157 | 0.10331366 | -3.378654448 | 0.000728415 | 0.005360697 |
| KHK | 30.73282545 | -0.527834113 | 0.156344243 | -3.376102002 | 0.000735207 | 0.005408543 |

| | | | | | | |
|---|---|---|---|---|---|---|
| MAD1L1 | 254.1299494 | -0.285744849 | 0.084644369 | -3.375828228 | 0.000735939 | 0.005411791 |
| RECK | 39.20314606 | -0.589918712 | 0.174935065 | -3.372215358 | 0.000745661 | 0.005481124 |
| NOP9 | 127.8886398 | -0.23569585 | 0.069900533 | -3.371874841 | 0.000746584 | 0.005485741 |
| AOC3 | 699.1253999 | -0.491465036 | 0.145812379 | -3.370530264 | 0.000750237 | 0.00551041 |
| SAP130 | 576.4651571 | 0.237529814 | 0.070479195 | 3.370211802 | 0.000751104 | 0.005514609 |
| GRK5 | 52.11295145 | 0.730744916 | 0.216850637 | 3.369807552 | 0.000752207 | 0.005518315 |
| RNASE1 | 12.66430829 | -1.591343644 | 0.472247946 | -3.369720628 | 0.000752444 | 0.005518315 |
| SLC25A39 | 638.1474312 | -0.223073976 | 0.066199929 | -3.369701149 | 0.000752497 | 0.005518315 |
| C6orf48 | 312.5770723 | -0.390827581 | 0.116066458 | -3.367274131 | 0.000759152 | 0.005564923 |
| STOX2 | 24.16433852 | -0.766406288 | 0.227659611 | -3.366456984 | 0.000761405 | 0.005579242 |
| IFT27 | 22.24250496 | -0.650000145 | 0.193096431 | -3.366194498 | 0.00076213 | 0.005582359 |
| CBR3 | 11.95223617 | 0.771561824 | 0.229256634 | 3.365493991 | 0.000764067 | 0.005594354 |
| ZFP161 | 174.1649058 | -0.348311548 | 0.103515202 | -3.364834737 | 0.000765895 | 0.005605535 |
| APH1B | 386.5491599 | -0.232299719 | 0.069057042 | -3.363881693 | 0.000768545 | 0.005622718 |
| PAQR7 | 13.58062569 | -0.937284544 | 0.278652216 | -3.363635715 | 0.00076923 | 0.005625523 |
| KCNAB2 | 2779.114921 | -0.445834433 | 0.132621249 | -3.361711904 | 0.000774609 | 0.005662638 |
| FLJ39639 | 8.069346351 | -1.33278259 | 0.396516028 | -3.361232572 | 0.000775955 | 0.00567025 |
| TCHP | 288.8193298 | -0.3197016 | 0.095122717 | -3.360938468 | 0.000776781 | 0.005674066 |
| ANKRD29 | 180.9631494 | 0.354455316 | 0.105483573 | 3.360289253 | 0.000778609 | 0.005685189 |
| PTH2R | 21.60690467 | -0.633448247 | 0.188626761 | -3.358209846 | 0.00078449 | 0.005725888 |
| PWP2 | 174.6142214 | -0.243750863 | 0.07261245 | -3.356874226 | 0.00078829 | 0.005751366 |
| KIAA0664 | 575.0435913 | 0.245103678 | 0.073024713 | 3.356448354 | 0.000789505 | 0.005757976 |
| ATP1B4 | 11.8512781 | 1.317840194 | 0.392693128 | 3.355903372 | 0.000791062 | 0.005767077 |
| SLC36A1 | 229.6284235 | -0.782706469 | 0.233314788 | -3.354722927 | 0.000794445 | 0.005789476 |
| CDR2L | 9.985704433 | -0.833849955 | 0.248597488 | -3.35421714 | 0.000795899 | 0.005795537 |
| RPS11 | 12571.30796 | -0.324701862 | 0.096802829 | -3.354260037 | 0.000795775 | 0.005795537 |
| SNX24 | 157.3151666 | -0.385758951 | 0.115013601 | -3.354028966 | 0.00079644 | 0.005797214 |
| ASB8 | 217.7822642 | -0.250730417 | 0.074781516 | -3.352839443 | 0.000799871 | 0.005819912 |
| POP5 | 192.3327021 | -0.298352347 | 0.089018452 | -3.351578689 | 0.000803522 | 0.005844196 |
| C1orf233 | 94.83085566 | -0.413295352 | 0.123324673 | -3.351278728 | 0.000804393 | 0.005848247 |
| SLC5A3 | 2294.490225 | 0.457826733 | 0.136627384 | 3.350914876 | 0.000805451 | 0.005853653 |
| ERMP1 | 619.8032826 | -0.281083486 | 0.083930118 | -3.349018127 | 0.000810985 | 0.005888036 |
| HAAO | 173.5182693 | 0.605112227 | 0.18068614 | 3.348968704 | 0.00081113 | 0.005888036 |
| HIST1H4C | 766.4415189 | -0.319263298 | 0.0953315 | -3.348980126 | 0.000811096 | 0.005888036 |
| PLEKHA4 | 25.63891252 | 0.880700496 | 0.26308694 | 3.347564483 | 0.00081525 | 0.005915643 |
| NSUN2 | 780.2959714 | 0.186764762 | 0.055816951 | 3.346022314 | 0.000819798 | 0.005946328 |
| AKR1C3 | 1082.899054 | -0.324229452 | 0.096959126 | -3.343980758 | 0.000825855 | 0.005987928 |
| CNBP | 3453.931724 | 0.217553886 | 0.065124249 | 3.34059724 | 0.000835984 | 0.006059014 |
| INSR | 157.4711998 | -0.24838021 | 0.074368676 | -3.339849821 | 0.000838237 | 0.006072982 |
| LOC10024( | 28.77811509 | -0.634012341 | 0.189843057 | -3.339665666 | 0.000838793 | 0.006074648 |
| SORT1 | 3797.628156 | 0.265518403 | 0.07951155 | 3.339368974 | 0.00083969 | 0.006078778 |
| IGIP | 107.0116645 | -0.338839822 | 0.101498765 | -3.338363984 | 0.000842733 | 0.00609844 |
| DDA1 | 1121.019479 | 0.216567092 | 0.064883727 | 3.337772071 | 0.00084453 | 0.006109073 |
| FERMT2 | 15.40245642 | 1.034946961 | 0.310166671 | 3.336744584 | 0.000847658 | 0.006129321 |
| SUMO3 | 1290.760168 | -0.254053504 | 0.076147842 | -3.336319162 | 0.000848956 | 0.006136328 |
| IL17RB | 488.5284427 | 0.566378763 | 0.16994222 | 3.33277254 | 0.000859852 | 0.006212673 |
| CBY1 | 42.07489758 | -0.475700046 | 0.142801269 | -3.331203216 | 0.000864715 | 0.006245384 |

| | | | | | | |
|---|---|---|---|---|---|---|
| SERPINB9 | 112.1044773 | 0.849876747 | 0.255143815 | 3.330971391 | 0.000865435 | 0.006248165 |
| BAD | 348.8088564 | -0.308470483 | 0.092618754 | -3.3305402 | 0.000866776 | 0.006255427 |
| RHBDF1 | 16.2933076 | 0.681199693 | 0.204598291 | 3.329449581 | 0.000870178 | 0.006277546 |
| SLA2 | 12.62808065 | 1.246183224 | 0.374343514 | 3.328983077 | 0.000871637 | 0.006285636 |
| PRAM1 | 517.0430213 | -0.351556945 | 0.105641428 | -3.327832199 | 0.000875246 | 0.006309218 |
| B9D1 | 31.89766331 | -0.544962463 | 0.163764349 | -3.327723438 | 0.000875587 | 0.006309241 |
| NAA35 | 172.5459236 | 0.226613722 | 0.068115244 | 3.326916387 | 0.000878127 | 0.006325095 |
| RAP2A | 1901.290525 | 0.256506382 | 0.077120304 | 3.32605512 | 0.000880845 | 0.006342221 |
| PTPLAD2 | 489.2289183 | -0.347623059 | 0.104521263 | -3.325859736 | 0.000881463 | 0.006344217 |
| MID2 | 50.28895342 | 0.70587219 | 0.212330907 | 3.32439682 | 0.0008861 | 0.006375131 |
| HSF4 | 7.648219046 | -1.164270453 | 0.350267727 | -3.323944406 | 0.000887539 | 0.006383018 |
| TMEM99 | 34.90659919 | -0.549548436 | 0.165351221 | -3.323522102 | 0.000888884 | 0.006390224 |
| ITFG2 | 362.6654461 | -0.258748053 | 0.077873042 | -3.322690976 | 0.000891536 | 0.006406821 |
| FAM117A | 181.4264386 | -0.359758604 | 0.108291739 | -3.322124183 | 0.000893349 | 0.006417375 |
| FCGR2C | 28.40177336 | -0.661934198 | 0.199315294 | -3.321040676 | 0.000896825 | 0.006439858 |
| RWDD2A | 109.0958524 | 0.322009072 | 0.097018718 | 3.319040679 | 0.000903273 | 0.006483661 |
| ALDH18A1 | 135.7581901 | -0.39582838 | 0.119268695 | -3.318795255 | 0.000904067 | 0.006486863 |
| MAP3K12 | 93.17992817 | -0.441950217 | 0.133215781 | -3.317551521 | 0.000908102 | 0.006513305 |
| GOLT1B | 641.2202487 | 0.218639125 | 0.065916647 | 3.316903023 | 0.000910212 | 0.00652593 |
| PCYOX1L | 119.3065929 | -0.443711601 | 0.133789956 | -3.316479168 | 0.000911594 | 0.006533323 |
| UGGT1 | 449.7236355 | -0.226686684 | 0.068361974 | -3.315976272 | 0.000913236 | 0.006542575 |
| ASRGL1 | 47.12473201 | -1.132206887 | 0.341702491 | -3.313428835 | 0.000921596 | 0.006599759 |
| ZNF737 | 159.6992173 | -0.399148354 | 0.120467484 | -3.313328536 | 0.000921926 | 0.006599759 |
| PSMA4 | 1201.854181 | 0.403993049 | 0.121966585 | 3.31232566 | 0.000925238 | 0.006620922 |
| LMLN | 161.2372295 | 0.296500728 | 0.089522046 | 3.312041449 | 0.000926178 | 0.006625107 |
| PHF6 | 268.8802681 | -0.190813327 | 0.057620663 | -3.311543427 | 0.000927828 | 0.006634365 |
| NCDN | 86.2547939 | 0.290554634 | 0.087756319 | 3.310925503 | 0.00092988 | 0.006646481 |
| RNF6 | 842.7406519 | 0.214673423 | 0.064840839 | 3.310774904 | 0.00093038 | 0.006647509 |
| PRKAG2-A! | 24.35700197 | -0.604114401 | 0.182530627 | -3.309660478 | 0.000934092 | 0.006671472 |
| CIR1 | 1076.628752 | 0.204616135 | 0.06186296 | 3.307571034 | 0.000941088 | 0.006718866 |
| RPL6 | 3121.53918 | -0.514589044 | 0.155642406 | -3.306226492 | 0.000945616 | 0.006748607 |
| LBR | 590.9665384 | -0.194596282 | 0.058876616 | -3.305153983 | 0.000949242 | 0.006771892 |
| ALDH1A1 | 2277.313003 | -0.58254786 | 0.176475534 | -3.301012024 | 0.000963368 | 0.006870031 |
| LOC10050! | 44.12421207 | -0.414005214 | 0.125446113 | -3.300263386 | 0.000965941 | 0.00688348 |
| TCEAL1 | 23.82952853 | -0.61584094 | 0.18660441 | -3.300248588 | 0.000965992 | 0.00688348 |
| DIEXF | 421.7666201 | 0.24783164 | 0.075116757 | 3.299285658 | 0.000969312 | 0.006901858 |
| HSD17B12 | 1224.696961 | 0.331466948 | 0.100464226 | 3.29935304 | 0.00096908 | 0.006901858 |
| HIST2H2AE | 131.4179754 | -0.383898097 | 0.116370879 | -3.298918946 | 0.000970579 | 0.006908241 |
| FAM40A | 341.8782012 | 0.247826239 | 0.075131657 | 3.298559456 | 0.000971823 | 0.006914452 |
| C5orf42 | 95.33937789 | -0.390931492 | 0.118521915 | -3.298389937 | 0.00097241 | 0.006915987 |
| TOM1L2 | 374.5683594 | -0.28309651 | 0.085832788 | -3.298232716 | 0.000972955 | 0.006917221 |
| LIMK2 | 260.2508871 | 0.619796487 | 0.187936656 | 3.297901012 | 0.000974105 | 0.006922756 |
| KLF13 | 1243.734597 | -0.222729167 | 0.067550897 | -3.297205169 | 0.000976521 | 0.006937285 |
| CORO6 | 16.55866928 | -0.81926873 | 0.248611961 | -3.29537134 | 0.000982917 | 0.006980059 |
| ARL2BP | 1019.929337 | 0.341305337 | 0.103613381 | 3.294027598 | 0.000987628 | 0.007010842 |
| LOC10028: | 25.7685382 | -0.613436842 | 0.186238685 | -3.293820731 | 0.000988355 | 0.007013332 |
| PLAGL2 | 141.3247279 | 0.405126309 | 0.123117039 | 3.290578711 | 0.000999815 | 0.007091952 |

| | | | | | | |
|---|---|---|---|---|---|---|
| TDRD9 | 333.7820934 | -0.395136092 | 0.12011133 | -3.289748696 | 0.001002769 | 0.007110197 |
| RPS9 | 2592.569742 | -0.433230699 | 0.131745004 | -3.288403245 | 0.001007574 | 0.00714155 |
| MCOLN3 | 83.43392752 | 0.421388344 | 0.128178488 | 3.287512201 | 0.001010768 | 0.007159388 |
| TPM2 | 45.54669075 | -0.451231653 | 0.137257326 | -3.287486843 | 0.001010859 | 0.007159388 |
| ZNF781 | 2.117693343 | -2.352384293 | 0.715636681 | -3.287120905 | 0.001012174 | 0.007165974 |
| L3HYPDH | 49.95095428 | -0.382614069 | 0.116438656 | -3.285971193 | 0.001016314 | 0.007189824 |
| SAP30BP | 1155.894819 | 0.246583075 | 0.075041088 | 3.285974131 | 0.001016303 | 0.007189824 |
| GPBAR1 | 102.3712665 | 0.669597766 | 0.20385574 | 3.284664767 | 0.001021038 | 0.0072205 |
| CKS1B | 183.9381312 | 0.403764379 | 0.122942421 | 3.284174631 | 0.001022815 | 0.007230326 |
| DNAJC7 | 1006.174912 | 0.34631307 | 0.105459397 | 3.283852173 | 0.001023986 | 0.007235858 |
| CLDN23 | 343.4675206 | 0.576014488 | 0.175433619 | 3.283375733 | 0.001025719 | 0.00724453 |
| PIP5K1A | 175.0472476 | 0.295391216 | 0.089967757 | 3.283300883 | 0.001025991 | 0.00724453 |
| HSPA6 | 154.3946189 | 0.500324945 | 0.152441534 | 3.282077614 | 0.001030452 | 0.007273276 |
| TLE2 | 37.59677439 | -0.641305313 | 0.195462581 | -3.280962057 | 0.001034537 | 0.007299339 |
| PER1 | 232.0439054 | 0.250410851 | 0.076325131 | 3.280844023 | 0.00103497 | 0.007299631 |
| ZNF554 | 7.164458202 | -0.841816866 | 0.256595976 | -3.28070954 | 0.001035463 | 0.007300348 |
| PLEKHA7 | 22.91264895 | 0.841320777 | 0.256486512 | 3.280175523 | 0.001037425 | 0.007311415 |
| SRSF12 | 5.365880397 | 1.966849095 | 0.599690348 | 3.279774474 | 0.001038901 | 0.007319048 |
| RPS24 | 3.804867421 | -1.728351272 | 0.527310686 | -3.277671625 | 0.001046671 | 0.007370999 |
| CD300LB | 768.5989855 | -0.404909713 | 0.123706679 | -3.273143516 | 0.001063585 | 0.007487282 |
| SMAD6 | 124.7044496 | -0.324479798 | 0.099156954 | -3.272385703 | 0.00106644 | 0.007504547 |
| NRP2 | 3996.229055 | 0.466717459 | 0.142673338 | 3.271231086 | 0.001070804 | 0.007532411 |
| MRPL48 | 81.87226942 | -0.444060616 | 0.13576148 | -3.270888145 | 0.001072103 | 0.007538705 |
| FSTL1 | 5.607052498 | 1.351066053 | 0.413086491 | 3.270661433 | 0.001072963 | 0.007541905 |
| ATP8B3 | 19.99493888 | -0.700972697 | 0.214375511 | -3.269835697 | 0.0010761 | 0.007561102 |
| VCP | 8265.111184 | 0.360548926 | 0.110375275 | 3.266573306 | 0.001088576 | 0.007645883 |
| PAAF1 | 78.70212165 | -0.425134888 | 0.130168056 | -3.266046225 | 0.001090604 | 0.007654359 |
| ZC2HC1A | 113.6880789 | -0.358947734 | 0.109900361 | -3.266119686 | 0.001090322 | 0.007654359 |
| LSM14B | 260.044225 | -0.309864854 | 0.094929159 | -3.264169388 | 0.001097855 | 0.007702346 |
| IFITM2 | 273.1315257 | 0.821930064 | 0.251829276 | 3.263838413 | 0.001099138 | 0.007708447 |
| CCDC107 | 226.0709766 | -0.234066223 | 0.071763137 | -3.261649817 | 0.001107659 | 0.00776528 |
| ADAP1 | 377.2957369 | -0.372119039 | 0.114136415 | -3.260300755 | 0.001112941 | 0.007796447 |
| PDE4DIP | 1712.922134 | 0.261569882 | 0.080226876 | 3.260377268 | 0.001112641 | 0.007796447 |
| KBTBD2 | 327.464403 | 0.272024025 | 0.083443417 | 3.259981838 | 0.001114193 | 0.007802285 |
| PSMC4 | 1414.810553 | 0.32121022 | 0.098581156 | 3.258332874 | 0.001120689 | 0.007844821 |
| SNX30 | 702.8060161 | -0.315040714 | 0.096729437 | -3.256926996 | 0.001126254 | 0.007880817 |
| ATP8B4 | 14.1955159 | -0.995114779 | 0.305680625 | -3.255406783 | 0.001132301 | 0.007920154 |
| TES | 359.0530542 | 0.284448392 | 0.087386796 | 3.255050035 | 0.001133724 | 0.007927134 |
| MTRNR2L8 | 23.43475694 | 0.855158112 | 0.262732704 | 3.254859778 | 0.001134484 | 0.00792947 |
| ZDHHC9 | 121.6624531 | 0.447554588 | 0.137517526 | 3.254527641 | 0.001135811 | 0.007935772 |
| MED22 | 228.5399885 | -0.224996499 | 0.069148722 | -3.253805623 | 0.001138702 | 0.007952986 |
| PBX3 | 772.7011653 | -0.303719283 | 0.093369321 | -3.252880916 | 0.001142414 | 0.007975922 |
| EZR | 1470.072007 | 0.281477472 | 0.086538094 | 3.252642364 | 0.001143373 | 0.00797963 |
| ANXA7 | 2617.685109 | 0.229881702 | 0.070695226 | 3.251728793 | 0.001147054 | 0.00800135 |
| GPR124 | 20.78044252 | -1.014631315 | 0.312035161 | -3.251656999 | 0.001147344 | 0.00800135 |
| TMEM14B | 236.4753212 | -0.202280409 | 0.062280004 | -3.247919041 | 0.001162523 | 0.008104176 |
| ZNF688 | 122.8977714 | 0.361326627 | 0.111264074 | 3.247468972 | 0.001164364 | 0.008113968 |

| | | | | | | |
|---|---|---|---|---|---|---|
| SH2B1 | 232.2450379 | -0.400290074 | 0.123274115 | -3.247154312 | 0.001165652 | 0.008119908 |
| SEPW1 | 99.09547048 | -0.478644228 | 0.147410248 | -3.247021378 | 0.001166196 | 0.008120666 |
| SLC7A6OS | 373.499636 | -0.189627276 | 0.058442423 | -3.244685368 | 0.001175805 | 0.008184514 |
| NPRL3 | 260.2782487 | -0.309154224 | 0.095290961 | -3.244318448 | 0.00117732 | 0.008192005 |
| ZNF271 | 245.3110344 | -0.273816348 | 0.084406965 | -3.244001839 | 0.00117863 | 0.008198055 |
| VASP | 452.4573638 | 0.484033754 | 0.14924356 | 3.243247184 | 0.001181756 | 0.008216734 |
| WBP4 | 184.0577659 | 0.406888069 | 0.125475348 | 3.242772973 | 0.001183725 | 0.008227352 |
| CAMTA1 | 135.0669154 | 0.47218424 | 0.145637535 | 3.242187796 | 0.001186158 | 0.008238828 |
| MACF1 | 967.7865171 | 0.219985393 | 0.067853453 | 3.242066285 | 0.001186664 | 0.008238828 |
| TNFSF8 | 15.62250505 | -0.966031528 | 0.297968697 | -3.24205709 | 0.001186702 | 0.008238828 |
| LINC00113 | 3.950742348 | 2.86569419 | 0.884476884 | 3.239987662 | 0.001195349 | 0.008295765 |
| DHCR24 | 856.9393008 | 0.408687701 | 0.126174488 | 3.23906765 | 0.001199211 | 0.008317359 |
| PNPLA7 | 21.16568238 | -0.841204892 | 0.259714683 | -3.238957774 | 0.001199673 | 0.008317359 |
| SELK | 480.0399971 | 0.274237878 | 0.084669339 | 3.238927827 | 0.001199799 | 0.008317359 |
| CD79A | 33.6032393 | 0.714217764 | 0.220532156 | 3.238610533 | 0.001201135 | 0.008319733 |
| CRTAM | 7.145987288 | 1.965716518 | 0.606960212 | 3.238625003 | 0.001201074 | 0.008319733 |
| PDCD1 | 6.492619924 | 1.129065456 | 0.34863536 | 3.238528236 | 0.001201481 | 0.008319733 |
| CCDC9 | 218.6284342 | 0.352333746 | 0.108798015 | 3.238420713 | 0.001201934 | 0.008319778 |
| SLC35B4 | 270.3409587 | -0.244551404 | 0.075525915 | -3.237980042 | 0.001203792 | 0.008329545 |
| ESR1 | 34.35749764 | 0.558708896 | 0.172559204 | 3.237780914 | 0.001204633 | 0.008331718 |
| NUP153 | 762.0925477 | 0.218486132 | 0.067482024 | 3.237693813 | 0.001205001 | 0.008331718 |
| GNB5 | 56.01376054 | -0.450473292 | 0.139172642 | -3.236794867 | 0.001208803 | 0.008354907 |
| ZFP14 | 58.42016355 | -0.469101826 | 0.144998439 | -3.235219838 | 0.001215491 | 0.00839802 |
| CCDC103 | 26.75371716 | 0.508909236 | 0.157312452 | 3.235021953 | 0.001216334 | 0.008400728 |
| RDH10 | 353.0841466 | -0.518591048 | 0.160329868 | -3.234525516 | 0.00121845 | 0.008412228 |
| NEFM | 4.539560182 | 1.472168972 | 0.455190365 | 3.234183069 | 0.001219912 | 0.008419202 |
| ATL3 | 877.7164038 | -0.165711456 | 0.051244141 | -3.233763944 | 0.001221704 | 0.008428445 |
| VNN1 | 356.1785419 | 0.370875185 | 0.114695369 | 3.233567224 | 0.001222546 | 0.008431131 |
| BTAF1 | 473.200981 | 0.214503513 | 0.066352755 | 3.232774767 | 0.001225942 | 0.008451424 |
| KAT2A | 19.66597935 | -0.727853517 | 0.225220775 | -3.231733465 | 0.001230418 | 0.008479143 |
| LIN7A | 7.055541265 | -1.246571482 | 0.385742857 | -3.231612609 | 0.001230938 | 0.008479594 |
| GEMIN6 | 76.09474563 | -0.387638959 | 0.11996876 | -3.231165838 | 0.001232864 | 0.008489721 |
| GPR137B | 3937.704728 | -0.19717295 | 0.061028006 | -3.230860087 | 0.001234183 | 0.008495668 |
| ARID3A | 386.4826246 | 0.312716489 | 0.096827328 | 3.229630487 | 0.001239503 | 0.008525987 |
| MAPK6 | 643.6828325 | 0.296334108 | 0.091754731 | 3.229633016 | 0.001239492 | 0.008525987 |
| PUM2 | 947.900063 | -0.178250575 | 0.055215516 | -3.228269662 | 0.001245415 | 0.008563493 |
| ZNF544 | 155.1719229 | -0.31926675 | 0.098916504 | -3.227638822 | 0.001248165 | 0.008579233 |
| REEP4 | 90.46206385 | -0.507677112 | 0.157305907 | -3.227323886 | 0.001249539 | 0.008585515 |
| FBXW8 | 56.97259394 | -0.533342699 | 0.165325274 | -3.226020349 | 0.001255244 | 0.008618358 |
| KIF5B | 3064.491474 | 0.298104114 | 0.092404285 | 3.226085401 | 0.001254959 | 0.008618358 |
| MAN2A2 | 727.1425473 | -0.35384441 | 0.109697269 | -3.22564465 | 0.001256893 | 0.008626499 |
| ENC1 | 80.48807789 | -0.44107739 | 0.136763175 | -3.225118091 | 0.001259207 | 0.0086392 |
| #N/A | 76.32255067 | -0.458233935 | 0.142119927 | -3.224276469 | 0.001262914 | 0.008661443 |
| DGKD | 83.14201859 | -0.327621686 | 0.101620776 | -3.223963619 | 0.001264295 | 0.008667721 |
| CCDC91 | 654.8764782 | -0.176201476 | 0.054687594 | -3.221964326 | 0.00127315 | 0.00872522 |
| ZNF625 | 109.288579 | -0.667614416 | 0.20725208 | -3.221267622 | 0.001276249 | 0.008743245 |
| EIF5 | 831.7814777 | 0.268129655 | 0.083274902 | 3.219813527 | 0.00128274 | 0.008784483 |

| | | | | | | |
|---|---|---|---|---|---|---|
| TREML1 | 6.933048138 | -1.342957723 | 0.41715339 | -3.219337909 | 0.00128487 | 0.008795835 |
| LHFPL2 | 80.30914587 | 0.475353293 | 0.147673873 | 3.21893971 | 0.001286655 | 0.008804824 |
| ZNF618 | 52.5200869 | 0.501964869 | 0.156043093 | 3.216834922 | 0.001296132 | 0.008866415 |
| MTSS1 | 37.29026448 | 0.592363785 | 0.184198124 | 3.215905628 | 0.001300336 | 0.008891911 |
| ABHD14A | 37.27974726 | -0.565713771 | 0.175923194 | -3.215686105 | 0.001301331 | 0.008895451 |
| ATOX1 | 4638.584726 | 0.288555246 | 0.089738413 | 3.215515378 | 0.001302105 | 0.00889748 |
| DNAJB6 | 2306.44985 | 0.299201835 | 0.093052687 | 3.215402408 | 0.001302618 | 0.00889772 |
| HDAC7 | 222.5462751 | 0.315049428 | 0.098045377 | 3.213302237 | 0.001312181 | 0.008949922 |
| MDM4 | 274.4888686 | 0.318410591 | 0.099088364 | 3.213400433 | 0.001311733 | 0.008949922 |
| RAB39A | 96.64753887 | 0.370997874 | 0.11545573 | 3.213334441 | 0.001312034 | 0.008949922 |
| WAS | 1722.833319 | -0.380738988 | 0.11847981 | -3.213534769 | 0.001311119 | 0.008949922 |
| CLIC2 | 592.6634471 | 0.257141499 | 0.080067271 | 3.211568158 | 0.001320127 | 0.009000819 |
| LINC00094 | 35.96817633 | -0.471570537 | 0.146845857 | -3.211330202 | 0.00132122 | 0.009004981 |
| PSMC6 | 1342.308417 | 0.200667989 | 0.062500028 | 3.210686369 | 0.001324184 | 0.009021879 |
| LMO4 | 740.918401 | 0.94288251 | 0.294086821 | 3.206136567 | 0.001345301 | 0.009162406 |
| PPP1R12A | 107.8637078 | 0.284363542 | 0.088745312 | 3.20426549 | 0.001354076 | 0.009218795 |
| EPHX2 | 2.812977133 | -1.98894224 | 0.620875321 | -3.203448701 | 0.001357922 | 0.009237176 |
| RPL13AP6 | 48.36020303 | 0.452606054 | 0.141290305 | 3.203376587 | 0.001358262 | 0.009237176 |
| WTAP | 215.679645 | 0.535714435 | 0.167233457 | 3.203392702 | 0.001358186 | 0.009237176 |
| FANCE | 60.2196125 | 0.707729417 | 0.2211065 | 3.200853058 | 0.001370214 | 0.009315055 |
| ZMIZ2 | 1087.652478 | 0.277348695 | 0.086659314 | 3.20044877 | 0.001372138 | 0.009324731 |
| LCLAT1 | 122.9270891 | -0.208858471 | 0.065266036 | -3.200109617 | 0.001373753 | 0.009332307 |
| BCAS3 | 131.9673847 | -0.337727279 | 0.105572188 | -3.199017526 | 0.001378968 | 0.009364317 |
| SNX9 | 78.76751558 | 0.327462412 | 0.102382717 | 3.198414957 | 0.001381853 | 0.00938049 |
| ALG1 | 170.7203546 | -0.276239282 | 0.086410234 | -3.196835227 | 0.001389443 | 0.009425146 |
| STAC | 446.8428201 | 0.552672171 | 0.172878895 | 3.196874733 | 0.001389252 | 0.009425146 |
| SLC25A40 | 377.5721491 | 0.302218592 | 0.094564531 | 3.195897951 | 0.001393964 | 0.009452375 |
| CSRNP2 | 49.71405976 | 0.491600557 | 0.15385489 | 3.195222186 | 0.001397232 | 0.009467646 |
| FAIM2 | 42.28703457 | 0.955643495 | 0.299079264 | 3.195285029 | 0.001396928 | 0.009467646 |
| NFATC1 | 62.60688448 | -0.425367439 | 0.133156785 | -3.194485664 | 0.001400802 | 0.009488387 |
| EIF4A2 | 4489.102569 | -0.207163857 | 0.064945895 | -3.189791387 | 0.001423755 | 0.009638278 |
| SIGLEC7 | 132.7298992 | -0.470719659 | 0.147572649 | -3.189748645 | 0.001423966 | 0.009638278 |
| TIMM13 | 230.2929476 | -0.228247221 | 0.0715784 | -3.188772327 | 0.001428784 | 0.009667376 |
| SNRNP25 | 69.37822053 | -0.511054566 | 0.16031093 | -3.187895966 | 0.001433121 | 0.009693203 |
| EEF1A1 | 51947.92577 | -0.307389204 | 0.09649377 | -3.185586012 | 0.001444612 | 0.009767377 |
| XXYLT1 | 9.127383927 | -0.814300019 | 0.255658472 | -3.185108681 | 0.001446997 | 0.009779954 |
| PRX | 4.512595489 | -1.264996606 | 0.397312489 | -3.183883318 | 0.001453136 | 0.009817887 |
| CIB1 | 911.9659935 | -0.178619085 | 0.056104438 | -3.18368907 | 0.001454111 | 0.009820916 |
| P2RY6 | 17.2186804 | -0.901337138 | 0.283234443 | -3.182300602 | 0.001461101 | 0.009864548 |
| EPOR | 24.46076992 | -0.741721654 | 0.233140671 | -3.181433982 | 0.001465479 | 0.009890525 |
| LSM7 | 658.4788297 | -0.175200855 | 0.055099505 | -3.179717425 | 0.001474187 | 0.009945694 |
| GRB10 | 80.58685682 | -0.656462495 | 0.20650457 | -3.178924778 | 0.001478224 | 0.009969321 |
| MAN1A2 | 184.194824 | -0.27624616 | 0.086913898 | -3.178388804 | 0.00148096 | 0.009984157 |
| ZMYND10 | 6.985845561 | -1.179192992 | 0.37109303 | -3.177620965 | 0.001484887 | 0.010007013 |
| HDAC9 | 15.16059988 | 0.582584402 | 0.183455608 | 3.175615117 | 0.001495192 | 0.010072814 |
| DECR2 | 121.8662589 | -0.351989483 | 0.110917127 | -3.173445735 | 0.001506411 | 0.010144725 |
| SLC35B2 | 60.40551762 | -0.607224394 | 0.1913555 | -3.173279021 | 0.001507276 | 0.010146886 |

| | | | | | | |
|---|---|---|---|---|---|---|
| PDIA3P | 5828.311314 | 0.187401243 | 0.059060483 | 3.173039483 | 0.00150852 | 0.010151594 |
| PFAS | 59.10506425 | -0.453357117 | 0.142895147 | -3.172655802 | 0.001510515 | 0.010161348 |
| MCMBP | 715.0468692 | 0.245790506 | 0.077485846 | 3.172069732 | 0.001513567 | 0.010178203 |
| KIAA1429 | 236.9831206 | -0.239237373 | 0.07542409 | -3.171896043 | 0.001514472 | 0.010180618 |
| PEX13 | 524.3931271 | 0.205555595 | 0.064821953 | 3.171079948 | 0.001518733 | 0.010205582 |
| UNC93B1 | 2099.568891 | 0.325169428 | 0.102551637 | 3.170787311 | 0.001520264 | 0.010212185 |
| DBF4 | 134.5106515 | 0.433375716 | 0.136707552 | 3.170093454 | 0.001523899 | 0.010232914 |
| GSDMD | 981.8058514 | 0.370770901 | 0.116990237 | 3.169246523 | 0.001528347 | 0.010259084 |
| PPP1R3G | 21.24972721 | -0.804097976 | 0.253951703 | -3.166342126 | 0.001543691 | 0.010358351 |
| NIP7 | 458.8082975 | 0.261741059 | 0.082707569 | 3.164656673 | 0.00155266 | 0.010414786 |
| NCLN | 1749.455975 | 0.445177153 | 0.140739746 | 3.163123173 | 0.001560863 | 0.010466037 |
| LRRC59 | 3418.658821 | 0.171916097 | 0.054369218 | 3.162011586 | 0.001566833 | 0.010502292 |
| NT5C3L | 512.1131076 | -0.195525498 | 0.061841598 | -3.161714839 | 0.001568431 | 0.010508667 |
| XPC | 449.2272546 | 0.254811905 | 0.080595223 | 3.161625426 | 0.001568912 | 0.010508667 |
| RAB3IP | 47.71223743 | 0.397982948 | 0.12590895 | 3.160878928 | 0.001572939 | 0.01053185 |
| KLF3 | 716.7259118 | -0.438678974 | 0.138821644 | -3.16001857 | 0.001577591 | 0.010559205 |
| CPNE2 | 477.1283422 | -0.304743651 | 0.09646879 | -3.158986982 | 0.001583186 | 0.010589046 |
| PSMB8 | 473.8056281 | 0.676918235 | 0.214278759 | 3.159054305 | 0.00158282 | 0.010589046 |
| GLB1 | 2051.180716 | -0.193894426 | 0.061392471 | -3.158276963 | 0.001587047 | 0.010611063 |
| CCR5 | 200.0246814 | 0.478090675 | 0.151393298 | 3.157938175 | 0.001588893 | 0.010619592 |
| CTDSPL | 16.11741134 | -0.66972718 | 0.212268643 | -3.155092386 | 0.001604473 | 0.010719881 |
| LOC10063( | 31.94393811 | -0.483945285 | 0.153405509 | -3.154679964 | 0.001606743 | 0.010731197 |
| PARVA | 29.01877095 | 0.629379324 | 0.199520222 | 3.154463828 | 0.001607933 | 0.010735301 |
| NT5C3 | 205.3374515 | 0.255583199 | 0.081031444 | 3.154123707 | 0.001609808 | 0.010741953 |
| SPATS2L | 214.6440415 | 0.527676109 | 0.167299849 | 3.154074023 | 0.001610082 | 0.010741953 |
| C12orf52 | 42.24894455 | -0.43023272 | 0.13642335 | -3.153658951 | 0.001612374 | 0.010753392 |
| A2M-AS1 | 6.170685012 | -1.001202989 | 0.317520051 | -3.153196113 | 0.001614933 | 0.010766603 |
| CHPF | 95.52142091 | 0.405521609 | 0.128690194 | 3.151146146 | 0.001626311 | 0.010838583 |
| NMNAT1 | 191.3877544 | -0.266511531 | 0.084614128 | -3.149728517 | 0.001634222 | 0.010887416 |
| RASGRF1 | 5.298777982 | 1.54921508 | 0.492245403 | 3.147241338 | 0.001648189 | 0.010976536 |
| R3HDM4 | 289.3626082 | 0.328109095 | 0.104303084 | 3.145727645 | 0.001656742 | 0.011029558 |
| ATP7B | 2.304494542 | -1.53692025 | 0.488724846 | -3.144755711 | 0.001662256 | 0.011062312 |
| NRGN | 7.834050104 | -1.052995694 | 0.334861531 | -3.144570504 | 0.001663308 | 0.011065365 |
| EIF4G1 | 5490.374625 | 0.310719618 | 0.098826351 | 3.144096849 | 0.001666003 | 0.011079336 |
| CAMK2G | 45.37579155 | -0.452402855 | 0.143908634 | -3.143681123 | 0.001668371 | 0.011091128 |
| TIE1 | 142.9915282 | 0.594709581 | 0.189183766 | 3.143555033 | 0.00166909 | 0.01109195 |
| FKBP7 | 4.439848149 | -1.409963123 | 0.448552146 | -3.143365013 | 0.001670174 | 0.011095197 |
| KDM5B | 211.8328135 | 0.25191326 | 0.080149006 | 3.143061552 | 0.001671907 | 0.011101761 |
| TTLL4 | 610.3593322 | -0.246858567 | 0.078542756 | -3.142983266 | 0.001672354 | 0.011101761 |
| ABCB5 | 23.1747584 | 0.876447914 | 0.278914484 | 3.142353528 | 0.001675955 | 0.011121707 |
| PWWP2B | 116.2006863 | -0.290489661 | 0.092455127 | -3.141952979 | 0.00167825 | 0.011132968 |
| FAM58A | 119.6934739 | -0.243645506 | 0.077555159 | -3.141577029 | 0.001680406 | 0.011143304 |
| COX5B | 2298.550961 | 0.209373763 | 0.066658309 | 3.14100022 | 0.001683719 | 0.011161302 |
| B2M | 91742.52397 | 0.307808487 | 0.098042207 | 3.139550766 | 0.001692071 | 0.011212678 |
| FBXO18 | 6.983741726 | -0.80626917 | 0.256874545 | -3.138766315 | 0.001696607 | 0.011234745 |
| LIPA | 24559.56893 | -0.419713465 | 0.133716024 | -3.138841958 | 0.001696169 | 0.011234745 |
| OSBPL1A | 834.9354648 | -0.34944065 | 0.111350121 | -3.138215265 | 0.0016998 | 0.011251891 |

| | | | | | | |
|---|---|---|---|---|---|---|
| GPANK1 | 132.8001761 | 0.27304976 | 0.087039603 | 3.137074975 | 0.001706425 | 0.01128934 |
| HLTF | 188.4394649 | -0.423171735 | 0.134895531 | -3.137033029 | 0.001706669 | 0.01128934 |
| PIGM | 87.65192573 | -0.460131412 | 0.146761073 | -3.13524154 | 0.001717127 | 0.011354485 |
| GATAD2A | 1772.913977 | 0.163123263 | 0.052030775 | 3.13513037 | 0.001717778 | 0.011354759 |
| MOB3A | 403.6884731 | -0.35051133 | 0.111816935 | -3.13468912 | 0.001720363 | 0.011367818 |
| PSMC1 | 988.1412311 | 0.233354 | 0.074458956 | 3.133995061 | 0.001724438 | 0.011390702 |
| EFCAB11 | 24.43763349 | -0.555968235 | 0.177412902 | -3.133753116 | 0.00172586 | 0.011396058 |
| METTL2A | 123.6061101 | 0.218498297 | 0.069735096 | 3.133261561 | 0.001728753 | 0.011411118 |
| POFUT2 | 189.6774174 | -0.266898003 | 0.085190189 | -3.132966442 | 0.001730492 | 0.011418553 |
| TCEB1 | 1763.271049 | 0.201399583 | 0.064342676 | 3.130108915 | 0.001747415 | 0.011526135 |
| METTL20 | 6.884781635 | -0.819645036 | 0.261911469 | -3.129473628 | 0.001751198 | 0.011547 |
| RPP30 | 29.76182976 | 0.332286017 | 0.106217822 | 3.128345236 | 0.001757936 | 0.011587327 |
| ARL14EP | 131.2336591 | -0.429888976 | 0.137435434 | -3.127934068 | 0.001760397 | 0.011599445 |
| NMRK1 | 168.2765282 | -0.292116594 | 0.093440027 | -3.126246895 | 0.001770528 | 0.011662081 |
| YWHAB | 1961.050331 | -0.261806231 | 0.083748881 | -3.126086325 | 0.001771495 | 0.011664327 |
| SERPINH1 | 498.0027635 | 0.270631395 | 0.086576522 | 3.125921308 | 0.00177249 | 0.011666752 |
| BMF | 506.5837015 | -0.384537489 | 0.123047438 | -3.12511576 | 0.001777351 | 0.011694618 |
| FUT10 | 162.9540745 | -0.264162958 | 0.084549469 | -3.124359751 | 0.001781924 | 0.011720572 |
| ARHGEF12 | 321.8570081 | 0.351438412 | 0.112495952 | 3.124009416 | 0.001784047 | 0.011730395 |
| CACYBP | 395.5056863 | -0.378798677 | 0.121262344 | -3.123794783 | 0.001785349 | 0.011734814 |
| UAP1L1 | 364.8028273 | -0.260730159 | 0.083480226 | -3.12325649 | 0.001788618 | 0.011752154 |
| FAM171A2 | 4.922047081 | -1.582846739 | 0.507078374 | -3.121503147 | 0.001799303 | 0.011818194 |
| RPL35A | 46.34760945 | -0.799430618 | 0.256124617 | -3.121256478 | 0.001800811 | 0.011823931 |
| STK11 | 219.4246312 | -0.354822209 | 0.113694036 | -3.120851561 | 0.001803289 | 0.01183603 |
| CBLB | 115.7055095 | 0.356125422 | 0.114145013 | 3.119938524 | 0.001808888 | 0.011868597 |
| SPINT2 | 1158.936739 | -0.284127853 | 0.091101727 | -3.11879767 | 0.001815906 | 0.011910452 |
| C20orf96 | 85.06190818 | -0.400875727 | 0.128548174 | -3.118486351 | 0.001817826 | 0.011918847 |
| FER1L6 | 5.745733 | 1.322510164 | 0.424175347 | 3.117838346 | 0.001821827 | 0.011940881 |
| SLU7 | 1216.481856 | 0.175992329 | 0.056460225 | 3.117102872 | 0.001826378 | 0.011966503 |
| AP3S2 | 319.0502694 | 0.158828152 | 0.050985901 | 3.115138663 | 0.001838585 | 0.012042244 |
| CLEC10A | 13.04055858 | -1.104990952 | 0.35515994 | -3.111248841 | 0.001862979 | 0.012197732 |
| FAM213A | 962.7127501 | 0.33345135 | 0.107204026 | 3.110436831 | 0.001868108 | 0.012227023 |
| ARHGEF9 | 65.33461659 | -0.452990575 | 0.145641708 | -3.110308034 | 0.001868923 | 0.012228061 |
| SOBP | 42.03315843 | 0.577579534 | 0.185705878 | 3.110184457 | 0.001869705 | 0.012228884 |
| CEP135 | 97.50764486 | 0.31968344 | 0.102824726 | 3.109013283 | 0.001877133 | 0.012273155 |
| PSMB1 | 1806.523254 | 0.188364444 | 0.060590628 | 3.108804978 | 0.001878457 | 0.012275458 |
| TRIM21 | 373.0688159 | 0.378404396 | 0.12172234 | 3.108750586 | 0.001878802 | 0.012275458 |
| MFN1 | 287.4845198 | 0.406931026 | 0.130934073 | 3.107907798 | 0.001884168 | 0.012306202 |
| PPP1R16B | 26.94008548 | 0.944993751 | 0.304164614 | 3.106849741 | 0.001890925 | 0.012346003 |
| HOMER1 | 317.9751302 | 0.396159815 | 0.127526883 | 3.106480797 | 0.001893286 | 0.01235709 |
| TMEM187 | 52.76764989 | -0.344139317 | 0.110810532 | -3.105655297 | 0.001898579 | 0.012387297 |
| MLKL | 427.9323939 | 0.322749339 | 0.103928121 | 3.1055054 | 0.001899542 | 0.012389239 |
| XBP1 | 1075.736607 | 0.326369679 | 0.105107954 | 3.105090224 | 0.00190221 | 0.012402302 |
| CAST | 3413.295003 | 0.18240945 | 0.05877865 | 3.103328334 | 0.001913572 | 0.012467658 |
| ZNHIT1 | 418.2844113 | -0.2485699 | 0.080097433 | -3.103344144 | 0.00191347 | 0.012467658 |
| CMTM6 | 1284.996899 | 0.285784126 | 0.092116185 | 3.102431183 | 0.001919382 | 0.012501139 |
| PCED1A | 326.6999937 | -0.280378691 | 0.090482817 | -3.098695437 | 0.001943747 | 0.012655411 |

| | | | | | | |
|---|---|---|---|---|---|---|
| SCARB2 | 2793.844922 | 0.247835875 | 0.080042945 | 3.096286293 | 0.001959611 | 0.012749785 |
| SCML1 | 4.577812203 | 1.390426612 | 0.449060711 | 3.096299851 | 0.001959521 | 0.012749785 |
| CHAMP1 | 126.4193552 | -0.270911827 | 0.087502959 | -3.096030488 | 0.001961302 | 0.012756335 |
| ARHGAP5 | 296.3633157 | 0.302531312 | 0.097786526 | 3.093793421 | 0.00197615 | 0.012848425 |
| CHD1L | 148.3233096 | -0.315983296 | 0.102165536 | -3.092856046 | 0.001982403 | 0.01288458 |
| PNPLA2 | 1397.141188 | -0.221875139 | 0.071748593 | -3.092397079 | 0.001985471 | 0.012900021 |
| APLN | 2.381659442 | -1.953634206 | 0.631924869 | -3.091560884 | 0.001991071 | 0.012931902 |
| DYNLL2 | 558.8303794 | -0.22160397 | 0.071691681 | -3.091069536 | 0.001994369 | 0.012944297 |
| LOC28483[illegible] | 10.32167142 | -0.972516333 | 0.31461179 | -3.091163021 | 0.001993741 | 0.012944297 |
| ZNF684 | 77.65740876 | 0.418924157 | 0.13554101 | 3.090755748 | 0.001996478 | 0.012953471 |
| RTN2 | 43.83211629 | 0.51973119 | 0.168259446 | 3.088867827 | 0.002009208 | 0.013031529 |
| CD37 | 2678.487686 | -0.498735954 | 0.161471962 | -3.08868454 | 0.002010448 | 0.013035033 |
| FCGRT | 2044.0235 | -0.433922971 | 0.140516048 | -3.088067005 | 0.00201463 | 0.013057608 |
| SLC35C2 | 899.9722605 | 0.301152809 | 0.09752989 | 3.087800157 | 0.00201644 | 0.013064794 |
| C17orf53 | 8.550639573 | 0.973657718 | 0.315337657 | 3.087667125 | 0.002017343 | 0.0130661 |
| RND3 | 666.7701643 | 0.526395189 | 0.170490347 | 3.087536622 | 0.002018229 | 0.013067297 |
| KCTD6 | 119.4495569 | -0.400463815 | 0.129751208 | -3.086397572 | 0.002025977 | 0.013112908 |
| C17orf61 | 149.5112656 | -0.53363851 | 0.172954966 | -3.085418834 | 0.002032657 | 0.013151573 |
| S100A1 | 97.36324628 | 0.685464109 | 0.222223004 | 3.084577631 | 0.002038414 | 0.013184244 |
| CBX8 | 30.68508212 | -0.389043376 | 0.126181079 | -3.083214848 | 0.002047773 | 0.013240178 |
| ETV5 | 56.1517822 | -0.523939034 | 0.169978691 | -3.082380691 | 0.00205352 | 0.013272736 |
| CHTOP | 266.8957303 | 0.22354991 | 0.072536862 | 3.081880069 | 0.002056977 | 0.013290468 |
| LRRC61 | 32.81771637 | 0.922089221 | 0.299254247 | 3.081290336 | 0.002061056 | 0.013312205 |
| COL7A1 | 9.166648155 | 1.036572731 | 0.336430697 | 3.08108844 | 0.002062454 | 0.01331662 |
| PER3 | 187.2076894 | -0.555242066 | 0.180231319 | -3.080719099 | 0.002065014 | 0.013328529 |
| ALPK1 | 338.9642354 | 0.235586647 | 0.0764896 | 3.079982712 | 0.002070126 | 0.013356901 |
| AGFG2 | 114.1802649 | 0.354191833 | 0.115010048 | 3.07965989 | 0.002072371 | 0.013366757 |
| CYP20A1 | 437.4063655 | -0.186544538 | 0.060578168 | -3.079402089 | 0.002074165 | 0.013373701 |
| RNASEH2B | 183.1586752 | -0.423654674 | 0.137595522 | -3.078985916 | 0.002077065 | 0.013387765 |
| MCM6 | 325.4364921 | -0.294640322 | 0.095697437 | -3.078873686 | 0.002077848 | 0.013388178 |
| FAM72A | 23.06538926 | 0.70326603 | 0.228450829 | 3.078413125 | 0.002081062 | 0.013399623 |
| GABRR2 | 10.02908569 | 1.001465333 | 0.325311838 | 3.078477993 | 0.002080609 | 0.013399623 |
| PPDPF | 1687.000681 | 0.27152247 | 0.088237606 | 3.077174022 | 0.002089733 | 0.013450804 |
| MGST3 | 7483.109368 | -0.184908373 | 0.06011728 | -3.075794064 | 0.002099428 | 0.013508543 |
| SULT1C2 | 116.6191906 | 1.001729004 | 0.325829278 | 3.074398384 | 0.002109276 | 0.013567222 |
| NCKAP1L | 4335.890184 | -0.216749811 | 0.070575379 | -3.071181664 | 0.002132134 | 0.013709518 |
| FAM172A | 178.7848048 | -0.250157487 | 0.08146914 | -3.070579713 | 0.002136436 | 0.013727729 |
| MIOS | 212.7495037 | -0.27331711 | 0.089011579 | -3.070579287 | 0.00213644 | 0.013727729 |
| SPR | 727.7664418 | 0.391477207 | 0.127508154 | 3.070213111 | 0.002139061 | 0.013739834 |
| PSTPIP1 | 79.70445147 | 0.498600797 | 0.162439768 | 3.069450315 | 0.002144531 | 0.013770223 |
| AGPAT9 | 1086.683378 | 0.41519004 | 0.135326667 | 3.06805782 | 0.002154549 | 0.013829568 |
| MAPK1 | 987.9074292 | 0.229466881 | 0.074794622 | 3.067959622 | 0.002155257 | 0.013829568 |
| ASAP1 | 2411.066535 | 0.337191093 | 0.10992466 | 3.067474522 | 0.002158759 | 0.013847266 |
| FAM122B | 471.3556555 | 0.321229256 | 0.104759582 | 3.066347251 | 0.002166915 | 0.013894801 |
| FAM210B | 299.670349 | -0.211700458 | 0.069060477 | -3.065435777 | 0.002173531 | 0.013932428 |
| ICT1 | 274.0111777 | 0.15434768 | 0.050352735 | 3.065328612 | 0.00217431 | 0.013932629 |
| THAP11 | 400.4653724 | -0.213718616 | 0.069731548 | -3.064876967 | 0.002177596 | 0.013948888 |

| | | | | | | |
|---|---|---|---|---|---|---|
| ZNF362 | 93.72794188 | -0.556714924 | 0.181718648 | -3.063609212 | 0.002186844 | 0.014003314 |
| CLEC12A | 204.6787188 | -0.472532151 | 0.154247422 | -3.063468706 | 0.002187871 | 0.014005079 |
| RFX5 | 664.8439057 | 0.20662151 | 0.067476944 | 3.062105315 | 0.002197861 | 0.014064198 |
| NFKBID | 107.0834909 | 0.253689057 | 0.082855 | 3.061843688 | 0.002199783 | 0.014066834 |
| TXNIP | 4768.671732 | -0.364896285 | 0.119174091 | -3.061875973 | 0.002199546 | 0.014066834 |
| COMMD1 | 790.951949 | 0.120346103 | 0.039309896 | 3.061470915 | 0.002202524 | 0.01407953 |
| ACTRT3 | 14.33250132 | 0.825542924 | 0.269733111 | 3.060591714 | 0.002209001 | 0.014116092 |
| FUZ | 73.78702832 | -0.330304623 | 0.107944822 | -3.059939482 | 0.002213817 | 0.014140776 |
| KLF12 | 16.08243766 | -0.5436895 | 0.177684253 | -3.05986316 | 0.002214381 | 0.014140776 |
| CCNB2 | 81.11695789 | 0.391674598 | 0.128091916 | 3.057762035 | 0.002229966 | 0.014235417 |
| LYRM5 | 33.94477993 | -0.581121972 | 0.190137191 | -3.056329857 | 0.002240646 | 0.014298698 |
| CLIC1 | 19142.79987 | 0.217339649 | 0.071130502 | 3.055505619 | 0.002246814 | 0.01433315 |
| MGC5734€ | 28.92585302 | -0.388672588 | 0.127211407 | -3.055328109 | 0.002248144 | 0.014336728 |
| C1GALT1 | 185.0713197 | 0.257111243 | 0.084173964 | 3.054522207 | 0.002254193 | 0.014370386 |
| CDC20 | 5.996617461 | 0.948136182 | 0.310474039 | 3.053834017 | 0.00225937 | 0.014398464 |
| CENPN | 173.165469 | 0.359500461 | 0.117743092 | 3.053261605 | 0.002263685 | 0.014418651 |
| PLCL1 | 56.31990623 | 0.504205802 | 0.165139656 | 3.053208503 | 0.002264086 | 0.014418651 |
| ATP6V0D2 | 708.9643495 | 0.244431967 | 0.08007427 | 3.052565661 | 0.002268941 | 0.014444634 |
| MYO9B | 1812.755903 | -0.326877716 | 0.107112346 | -3.051727719 | 0.002275284 | 0.014480068 |
| TMEM199 | 325.1672549 | 0.213571066 | 0.070000073 | 3.051012048 | 0.002280714 | 0.014509672 |
| PDLIM2 | 49.85252006 | -0.501169386 | 0.164326116 | -3.049846227 | 0.002289586 | 0.014561139 |
| DUT | 159.4611087 | -0.56094498 | 0.183962796 | -3.049230561 | 0.002294283 | 0.014576091 |
| SLMO2 | 791.0387835 | 0.276627962 | 0.090718642 | 3.04929565 | 0.002293786 | 0.014576091 |
| TSPAN14 | 1119.406184 | -0.225802269 | 0.074051215 | -3.049271647 | 0.00229397 | 0.014576091 |
| PPP1R3B | 105.1453876 | -0.295031427 | 0.096765554 | -3.048930269 | 0.002296578 | 0.014585696 |
| NEDD9 | 43.2067207 | 0.547995753 | 0.179745686 | 3.048728266 | 0.002298122 | 0.014590533 |
| YES1 | 43.94940369 | 0.607397233 | 0.19927707 | 3.048003631 | 0.002303671 | 0.014620781 |
| SH2D3A | 7.701381835 | 1.129159251 | 0.370477102 | 3.047851664 | 0.002304837 | 0.01462225 |
| TTC39C | 46.58919178 | -0.531400068 | 0.174357079 | -3.047768806 | 0.002305472 | 0.01462225 |
| GK | 7.650494043 | 1.128928683 | 0.370570391 | 3.04646218 | 0.002315516 | 0.014680955 |
| C7orf57 | 10.66785673 | 1.093902262 | 0.359088286 | 3.046332351 | 0.002316516 | 0.0146823 |
| PION | 1583.860974 | -0.207605184 | 0.068153824 | -3.046126727 | 0.002318101 | 0.014682358 |
| TCEAL4 | 406.0828091 | -0.332013499 | 0.108992999 | -3.046191062 | 0.002317605 | 0.014682358 |
| CD163 | 2979.225706 | -0.334608825 | 0.109911151 | -3.044357384 | 0.00233178 | 0.014763976 |
| GLT8D1 | 295.0163475 | -0.218771146 | 0.071875091 | -3.043768632 | 0.002336347 | 0.014783354 |
| PSMA6 | 3168.61351 | 0.501741818 | 0.164842852 | 3.043758407 | 0.002336427 | 0.014783354 |
| MLLT3 | 8.887001522 | -0.728709556 | 0.239429281 | -3.043527309 | 0.002338222 | 0.014789692 |
| ZDHHC5 | 836.6449565 | 0.221800558 | 0.072894886 | 3.042745091 | 0.002344308 | 0.014823156 |
| NXF1 | 541.5848101 | 0.274903019 | 0.090415965 | 3.040425651 | 0.00236244 | 0.014932547 |
| PI4KB | 575.1121421 | 0.258127468 | 0.084901209 | 3.040327351 | 0.002363211 | 0.014932547 |
| GPR157 | 372.8857654 | 0.390258546 | 0.128372154 | 3.040056077 | 0.002365341 | 0.014937757 |
| PVRL4 | 17.01511309 | 0.896265619 | 0.29482246 | 3.040018116 | 0.002365639 | 0.014937757 |
| ZMYM6 | 263.3285561 | -0.224851233 | 0.073975059 | -3.03955463 | 0.002369282 | 0.014955693 |
| TMEM241 | 65.41699992 | 0.396097784 | 0.130332577 | 3.039131067 | 0.002372616 | 0.014971665 |
| C1orf63 | 135.5555841 | 0.261624651 | 0.0861022 | 3.038536189 | 0.002377305 | 0.014986032 |
| GALNT6 | 226.7926835 | 0.30088957 | 0.09902361 | 3.038563936 | 0.002377087 | 0.014986032 |
| SIX5 | 6.087914807 | 1.445797205 | 0.475808952 | 3.038608667 | 0.002376734 | 0.014986032 |

| | | | | | | |
|---|---|---|---|---|---|---|
| SAT1 | 11231.837 | 0.46797194 | 0.154050301 | 3.037786601 | 0.002383227 | 0.015018277 |
| MAT2B | 1335.933858 | 0.272076498 | 0.089590545 | 3.036888518 | 0.002390338 | 0.015052912 |
| PPM1K | 286.6415563 | 0.276024141 | 0.090887625 | 3.036982668 | 0.002389592 | 0.015052912 |
| EXOC6B | 135.4751207 | -0.246822402 | 0.081282175 | -3.036611683 | 0.002392535 | 0.015061652 |
| SF3B4 | 40.66831712 | 0.612383416 | 0.201703863 | 3.036051993 | 0.00239698 | 0.015084542 |
| GPRIN3 | 1169.398086 | -0.314888146 | 0.103722258 | -3.035878235 | 0.002398362 | 0.015088142 |
| APLP2 | 9983.554579 | -0.376720217 | 0.124151505 | -3.034358842 | 0.002410475 | 0.015159229 |
| HTATIP2 | 360.1380224 | 0.274961438 | 0.090647757 | 3.033295538 | 0.002418986 | 0.015207618 |
| MSL3 | 422.5764203 | 0.327125851 | 0.107859695 | 3.032883153 | 0.002422294 | 0.015223279 |
| STAC2 | 10.68488804 | 0.767802082 | 0.253202007 | 3.032369648 | 0.002426419 | 0.015244061 |
| BLOC1S5 | 319.0109588 | 0.172327656 | 0.056834044 | 3.03212029 | 0.002428424 | 0.015251518 |
| PPAPDC1B | 209.3371214 | 0.217522532 | 0.071764107 | 3.031076982 | 0.002436831 | 0.015299161 |
| STX8 | 308.8813071 | -0.210532103 | 0.069479331 | -3.030140044 | 0.002444404 | 0.015341535 |
| TMEM42 | 64.90224643 | -0.301540601 | 0.09956699 | -3.028519806 | 0.00245755 | 0.015418849 |
| HYOU1 | 1050.004333 | 0.168054497 | 0.055494953 | 3.028284332 | 0.002459466 | 0.015423236 |
| THBD | 361.8829732 | -0.476590566 | 0.15738253 | -3.028230423 | 0.002459904 | 0.015423236 |
| KCNIP2 | 6.019727718 | -1.012772487 | 0.334498066 | -3.027737946 | 0.002463917 | 0.015443195 |
| CPT1A | 506.4855142 | -0.255409319 | 0.084361282 | -3.027565656 | 0.002465322 | 0.015446806 |
| CSPG4 | 66.32239783 | 0.615372794 | 0.203307419 | 3.02680934 | 0.002471498 | 0.015476095 |
| VPS37C | 828.9149528 | 0.218569128 | 0.07221153 | 3.02678987 | 0.002471657 | 0.015476095 |
| ABCB6 | 84.51982537 | 0.557247885 | 0.184119214 | 3.026560199 | 0.002473536 | 0.015482655 |
| CHP1 | 1883.948955 | -0.255725961 | 0.084581219 | -3.023436685 | 0.002499213 | 0.015638126 |
| LOC10012( | 2.764168682 | 2.04917132 | 0.677820933 | 3.023175032 | 0.002501375 | 0.01564115 |
| SETD5 | 442.7445268 | 0.248152317 | 0.082081968 | 3.023225706 | 0.002500956 | 0.01564115 |
| UBE2V1 | 52.60605397 | -0.35253919 | 0.116619651 | -3.022982725 | 0.002502965 | 0.015645842 |
| VDAC1 | 1695.914164 | 0.193787634 | 0.064161395 | 3.020315153 | 0.002525118 | 0.015779023 |
| SNRPA1 | 1024.549959 | 0.3453143 | 0.114363081 | 3.019456065 | 0.00253229 | 0.015818537 |
| NFE2L1 | 1394.893278 | 0.257366242 | 0.085255605 | 3.018760377 | 0.002538112 | 0.01584959 |
| PTK6 | 19.76202606 | 0.700942703 | 0.232246565 | 3.018097184 | 0.002543673 | 0.015878997 |
| PSMB4 | 2932.533854 | 0.205735753 | 0.068194916 | 3.016878151 | 0.002553924 | 0.015937652 |
| SUGP2 | 319.183075 | -0.278232237 | 0.092253553 | -3.015951461 | 0.002561743 | 0.015981089 |
| MRPL30 | 157.2946906 | -0.358778007 | 0.118970188 | -3.015696721 | 0.002563896 | 0.015989167 |
| FAM46C | 18.00919166 | 0.88024742 | 0.291937764 | 3.015188609 | 0.002568195 | 0.016010621 |
| CYFIP1 | 1993.242374 | -0.192052852 | 0.063721218 | -3.013954517 | 0.002578665 | 0.016070514 |
| DTWD2 | 210.0717051 | -0.369997531 | 0.122790146 | -3.013250999 | 0.00258465 | 0.016102433 |
| TMX2 | 1591.497881 | 0.124919784 | 0.041462981 | 3.012802775 | 0.002588471 | 0.016120844 |
| FAM35A | 235.1921739 | -0.227535869 | 0.07553103 | -3.012482017 | 0.002591208 | 0.016132498 |
| SLC16A13 | 58.04909117 | 0.458070727 | 0.152078125 | 3.012075061 | 0.002594684 | 0.016148747 |
| KCNJ10 | 10.34692782 | -0.831140189 | 0.275945585 | -3.011971319 | 0.002595571 | 0.016148873 |
| EIF2S3 | 2248.284835 | -0.271015668 | 0.089998036 | -3.011350923 | 0.002600881 | 0.016165889 |
| FOXP1 | 534.3926896 | -0.377579061 | 0.12538541 | -3.011347651 | 0.002600909 | 0.016165889 |
| TNFSF13B | 399.226596 | 0.512064061 | 0.170036886 | 3.011488108 | 0.002599706 | 0.016165889 |
| ADCK2 | 419.8449047 | 0.216194372 | 0.071813427 | 3.010500693 | 0.002608174 | 0.016205637 |
| MFSD7 | 610.7619249 | 0.386985189 | 0.128559703 | 3.010159323 | 0.002611107 | 0.016218455 |
| SRPR | 1112.328276 | 0.177207026 | 0.058878786 | 3.009692259 | 0.002615125 | 0.016238001 |
| ZNF549 | 24.41379673 | -0.487488623 | 0.162037813 | -3.00848681 | 0.002625522 | 0.016297127 |
| EVI5 | 249.2274306 | -0.289062387 | 0.096119136 | -3.007334412 | 0.002635497 | 0.016353594 |

| | | | | | | |
|---|---|---|---|---|---|---|
| EDF1 | 1211.87866 | 0.380836838 | 0.126663283 | 3.006686923 | 0.002641116 | 0.016383009 |
| FAM70A | 395.9274809 | -0.342221399 | 0.11383009 | -3.006422988 | 0.00264341 | 0.016386598 |
| ZNF192 | 273.363057 | -0.293632738 | 0.097668634 | -3.006417975 | 0.002643454 | 0.016386598 |
| SLC39A13 | 104.4757301 | 0.509312928 | 0.169472055 | 3.005291516 | 0.002653264 | 0.016441944 |
| SGPP1 | 1661.293558 | 0.233311974 | 0.077689123 | 3.003148523 | 0.00267202 | 0.016552666 |
| PLAGL1 | 215.6643463 | 0.291634004 | 0.097217748 | 2.999802095 | 0.002701551 | 0.016724484 |
| WDR77 | 251.4307619 | -0.239590439 | 0.079868693 | -2.99980419 | 0.002701532 | 0.016724484 |
| ATP5G2 | 2981.348376 | -0.312988397 | 0.10434399 | -2.999582407 | 0.0027035 | 0.016725437 |
| LPIN2 | 917.5203405 | 0.221964224 | 0.073997082 | 2.999634818 | 0.002703035 | 0.016725437 |
| GSK3A | 1374.413847 | 0.174140807 | 0.058069447 | 2.998837039 | 0.002710122 | 0.016760842 |
| SULT1B1 | 2.998259921 | -2.290102417 | 0.76407151 | -2.997235713 | 0.0027244 | 0.016843551 |
| MECR | 59.9650716 | -0.37487735 | 0.125091735 | -2.996819491 | 0.002728122 | 0.01686097 |
| FAAH | 3.847854408 | -1.252390377 | 0.417986037 | -2.996249314 | 0.002733229 | 0.016886931 |
| CHAF1B | 17.79743188 | 0.575332488 | 0.192044995 | 2.995821302 | 0.002737068 | 0.016905045 |
| PPM1H | 51.09071568 | -0.637972984 | 0.212991559 | -2.995297034 | 0.002741777 | 0.01692852 |
| RGS19 | 141.0636184 | -0.326582724 | 0.10903705 | -2.995153711 | 0.002743066 | 0.016930867 |
| AIFM3 | 34.74162815 | -0.496802197 | 0.165886796 | -2.994826647 | 0.002746009 | 0.01693781 |
| SMARCC1 | 616.1174155 | -0.233443599 | 0.077947227 | -2.994892908 | 0.002745412 | 0.01693781 |
| SPTAN1 | 2344.087271 | 0.382629411 | 0.127776931 | 2.994510878 | 0.002748853 | 0.016949741 |
| DDRGK1 | 1013.091751 | 0.200618518 | 0.067023327 | 2.993264085 | 0.002760108 | 0.017013514 |
| NOTCH1 | 494.9484348 | -0.401898467 | 0.134377379 | -2.99081937 | 0.0027823 | 0.017144637 |
| EIF2C4 | 581.3826222 | -0.230598765 | 0.07713276 | -2.989634569 | 0.002793114 | 0.017205581 |
| KLHL28 | 207.4162062 | 0.339864612 | 0.113699543 | 2.989146681 | 0.002797578 | 0.017227385 |
| TWF1 | 482.8518498 | 0.243727754 | 0.081542716 | 2.988958009 | 0.002799306 | 0.017232332 |
| C3orf58 | 9.268918075 | -0.910977178 | 0.304799382 | -2.988776327 | 0.002800971 | 0.017236887 |
| TUT1 | 233.791975 | 0.257878643 | 0.086287556 | 2.988595971 | 0.002802625 | 0.017241369 |
| FNBP1 | 1574.507022 | -0.319773729 | 0.107107727 | -2.98553371 | 0.002830839 | 0.017409195 |
| CLINT1 | 543.7027358 | 0.293815834 | 0.09842709 | 2.985111446 | 0.00283475 | 0.017427495 |
| SEPT7P2 | 71.39480855 | -0.384525149 | 0.12883558 | -2.984619227 | 0.002839315 | 0.017449802 |
| BICD2 | 678.586998 | -0.1807511 | 0.060627154 | -2.98135552 | 0.002869754 | 0.017631058 |
| LCN2 | 21.84452345 | 1.49323518 | 0.501467483 | 2.977730822 | 0.002903909 | 0.01782914 |
| RABGAP1 | 496.7986492 | -0.176949796 | 0.059423914 | -2.977753966 | 0.00290369 | 0.01782914 |
| DTWD1 | 88.70549653 | -0.488518883 | 0.164067715 | -2.977544255 | 0.002905677 | 0.017834119 |
| REV3L | 104.0331336 | -0.518606745 | 0.174288359 | -2.97556732 | 0.002924472 | 0.017943564 |
| SNRPD3 | 580.1792208 | -0.162811825 | 0.054732894 | -2.974661337 | 0.002933122 | 0.017990714 |
| KLF4 | 159.7449052 | -0.2748166 | 0.092407277 | -2.973971424 | 0.002939725 | 0.01802528 |
| COL11A2 | 77.11149688 | 1.001370071 | 0.336762692 | 2.97351843 | 0.002944067 | 0.01804597 |
| SLC48A1 | 281.2763934 | -0.395536026 | 0.133058899 | -2.972638657 | 0.002952518 | 0.01809182 |
| SAMD8 | 141.0979442 | 0.350152038 | 0.117864774 | 2.970794644 | 0.002970303 | 0.018194816 |
| FKBP1A | 5687.526787 | 0.30259989 | 0.101874058 | 2.970333148 | 0.00297477 | 0.018216186 |
| CMC2 | 570.7972807 | 0.244791827 | 0.08242335 | 2.969933007 | 0.002978647 | 0.018230897 |
| MYL5 | 13.28255683 | 0.777800487 | 0.261895972 | 2.969883348 | 0.002979129 | 0.018230897 |
| MAPK1IP1 | 2479.28022 | 0.239308949 | 0.080591869 | 2.969393205 | 0.002983885 | 0.01825401 |
| RRAGB | 178.9588537 | -0.269827727 | 0.090879532 | -2.969070372 | 0.002987022 | 0.018267202 |
| GRAMD4 | 12.04000392 | -1.02115593 | 0.344097894 | -2.967632024 | 0.003001034 | 0.018346871 |
| ZNF430 | 487.6216724 | 0.354805271 | 0.119568208 | 2.967388053 | 0.003003416 | 0.018355415 |
| C21orf7 | 63.91971226 | 0.560963905 | 0.189074499 | 2.96689351 | 0.003008251 | 0.018363463 |

| | | | | | | |
|---|---|---|---|---|---|---|
| NXPE3 | 261.4679231 | -0.345053205 | 0.116302874 | -2.966850201 | 0.003008675 | 0.018363463 |
| PRKCB | 194.8894398 | -0.323216313 | 0.108941177 | -2.966888381 | 0.003008301 | 0.018363463 |
| SETMAR | 25.74618664 | 0.594723723 | 0.20045137 | 2.966922715 | 0.003007966 | 0.018363463 |
| ELOVL5 | 2745.610389 | 0.301467631 | 0.101652983 | 2.965654537 | 0.003020396 | 0.018428962 |
| IL6R | 437.7226791 | 0.467368459 | 0.157603388 | 2.965472155 | 0.003022187 | 0.018433857 |
| GNB4 | 275.6512343 | -0.362546911 | 0.122269777 | -2.965139231 | 0.00302546 | 0.01844778 |
| SMTN | 11.24184177 | 0.896283066 | 0.302343467 | 2.964453228 | 0.003032213 | 0.018482912 |
| GCNT4 | 2.537944661 | 2.770206276 | 0.934781435 | 2.963480202 | 0.003041816 | 0.018535384 |
| SKP2 | 87.73319988 | -0.351414179 | 0.118588426 | -2.963309252 | 0.003043506 | 0.01853962 |
| TXNDC5 | 582.4624769 | -0.284568662 | 0.096053849 | -2.962595109 | 0.003050575 | 0.018576609 |
| KIF16B | 577.2018404 | -0.302438351 | 0.102113797 | -2.961777553 | 0.003058687 | 0.018619918 |
| PCTP | 146.9852294 | -0.273807086 | 0.092450741 | -2.961653765 | 0.003059917 | 0.018621321 |
| ASMTL | 67.79125928 | -0.364208817 | 0.123011134 | -2.960779272 | 0.003068618 | 0.018668176 |
| DENND5B | 39.52542545 | 0.417094385 | 0.140926701 | 2.959654791 | 0.003079839 | 0.018729946 |
| HIST1H3I | 264.9907948 | -0.420468139 | 0.14207114 | -2.959560544 | 0.003080782 | 0.018729946 |
| GCSHP3 | 214.0004676 | -0.347836959 | 0.117561063 | -2.958776904 | 0.003088626 | 0.018771513 |
| RP9 | 114.8772132 | -0.308489887 | 0.10427901 | -2.958312389 | 0.003093285 | 0.018793697 |
| MYCBP2 | 397.9711637 | -0.235254006 | 0.079527793 | -2.958135733 | 0.003095058 | 0.018798342 |
| ADRA2B | 60.87185935 | 0.67772089 | 0.229155938 | 2.957465978 | 0.00310179 | 0.018833089 |
| RBBP9 | 212.6222792 | 0.269624117 | 0.091186556 | 2.956840652 | 0.003108087 | 0.018865177 |
| PDCD4 | 1150.078834 | -0.26392055 | 0.089278391 | -2.956152623 | 0.003115029 | 0.018901156 |
| ZNF329 | 75.01889339 | -0.308539272 | 0.104394725 | -2.955506355 | 0.003121563 | 0.018934635 |
| ITM2B | 7908.983431 | -0.189278947 | 0.06408633 | -2.953499552 | 0.003141931 | 0.01905198 |
| CECR2 | 24.95720704 | 0.555546722 | 0.188111396 | 2.953285844 | 0.003144107 | 0.019058974 |
| CD7 | 36.38193142 | 0.941954488 | 0.319017104 | 2.95267707 | 0.003150313 | 0.01908418 |
| DPH3 | 567.6448453 | 0.183257271 | 0.062063017 | 2.952761248 | 0.003149454 | 0.01908418 |
| RYBP | 174.9309873 | 0.29766121 | 0.10081864 | 2.952442228 | 0.003152711 | 0.019092496 |
| AMD1 | 1540.110121 | 0.159403363 | 0.054032444 | 2.950141648 | 0.003176283 | 0.019228997 |
| CHCHD6 | 41.20189351 | -0.437742487 | 0.148416007 | -2.94942909 | 0.003183616 | 0.019267134 |
| GIMAP4 | 115.0730721 | 0.50347679 | 0.170760237 | 2.9484428 | 0.003193792 | 0.019322444 |
| NANOS1 | 41.39224889 | 0.645455417 | 0.218924991 | 2.948294821 | 0.003195322 | 0.019325422 |
| LOC15038: | 64.26811599 | -0.522654334 | 0.177537111 | -2.943915966 | 0.00324088 | 0.019594599 |
| OSBPL8 | 1825.881673 | 0.225203783 | 0.076515462 | 2.943245429 | 0.003247908 | 0.019630724 |
| HRASLS | 14.6142336 | -0.843546206 | 0.286622428 | -2.943057218 | 0.003249883 | 0.019636293 |
| CXCL16 | 6937.211303 | 0.303994772 | 0.103310363 | 2.942539011 | 0.003255328 | 0.019662812 |
| GPR108 | 839.8588954 | 0.268004268 | 0.091128707 | 2.940942278 | 0.003272155 | 0.01974689 |
| KL | 67.48706475 | -0.533682202 | 0.181459419 | -2.941055393 | 0.00327096 | 0.01974689 |
| ORC6 | 30.51064487 | -0.619026808 | 0.210487715 | -2.940916571 | 0.003272427 | 0.01974689 |
| WDR19 | 36.91633366 | -0.398879699 | 0.135642648 | -2.94066582 | 0.003275077 | 0.019756483 |
| LSS | 14.38685257 | 0.867930728 | 0.295206339 | 2.94008161 | 0.003281258 | 0.019787367 |
| KIFC2 | 37.84326285 | -0.42548725 | 0.144724482 | -2.939981155 | 0.003282322 | 0.019787379 |
| PPAP2B | 2213.355664 | 0.226983946 | 0.07722182 | 2.939375775 | 0.003288741 | 0.019819662 |
| PYROXD1 | 558.0645578 | 0.32602153 | 0.110942526 | 2.938652484 | 0.003296425 | 0.019859546 |
| RPS6KA2 | 631.4004431 | -0.450361579 | 0.153276644 | -2.938227029 | 0.003300952 | 0.019880394 |
| RC3H2 | 420.0106502 | -0.193646898 | 0.065909324 | -2.938080477 | 0.003302513 | 0.019883368 |
| GPER | 11.81112302 | 1.117962735 | 0.380580181 | 2.937522209 | 0.003308465 | 0.019912769 |
| ACADVL | 633.934933 | 0.26369795 | 0.089799674 | 2.936513443 | 0.003319244 | 0.019964751 |

| | | | | | | |
|---|---|---|---|---|---|---|
| LRPPRC | 703.4646423 | -0.227979004 | 0.077635587 | -2.936527075 | 0.003319098 | 0.019964751 |
| SLC22A16 | 27.76117747 | -0.837830456 | 0.285463481 | -2.934982973 | 0.00333566 | 0.020057014 |
| TMIE | 3.352468301 | -1.686094763 | 0.574800705 | -2.933355421 | 0.003353198 | 0.020155964 |
| BTN3A1 | 151.7418936 | 0.295476608 | 0.100743653 | 2.932955076 | 0.003357525 | 0.020175464 |
| MTOR | 612.0006627 | -0.201869629 | 0.06889877 | -2.929945329 | 0.003390216 | 0.020365343 |
| FDPS | 346.1061846 | 0.306676575 | 0.104684002 | 2.929545769 | 0.003394578 | 0.020384973 |
| CASP9 | 521.8048322 | 0.186847474 | 0.063787283 | 2.929227682 | 0.003398054 | 0.020399272 |
| HAUS7 | 59.99709714 | -0.426818512 | 0.145728066 | -2.928869668 | 0.00340197 | 0.020416205 |
| MAP3K3 | 438.9042608 | -0.306714241 | 0.10472873 | -2.928654268 | 0.003404329 | 0.020423779 |
| PTDSS1 | 1010.072027 | -0.176994251 | 0.060449444 | -2.927971526 | 0.003411813 | 0.020456785 |
| TMEM176I | 2695.465186 | -0.362666049 | 0.12386338 | -2.927952135 | 0.003412026 | 0.020456785 |
| JMJD6 | 140.1788744 | 0.51012659 | 0.174271135 | 2.927200703 | 0.003420281 | 0.020499683 |
| P2RY2 | 44.53409947 | 0.559319368 | 0.191099655 | 2.926846559 | 0.003424178 | 0.02051644 |
| PLCG2 | 534.0821283 | -0.27383764 | 0.093565293 | -2.926701031 | 0.00342578 | 0.020519444 |
| CASP3 | 534.3655481 | 0.401844186 | 0.137342401 | 2.925856717 | 0.003435091 | 0.020561994 |
| CD74 | 21552.33768 | -0.228794633 | 0.078195395 | -2.925934881 | 0.003434228 | 0.020561994 |
| SIK1 | 21.44049858 | 0.73250613 | 0.250547226 | 2.923624985 | 0.003459813 | 0.020703325 |
| MTFMT | 349.2267722 | -0.205111638 | 0.070199922 | -2.921821463 | 0.00347991 | 0.020816897 |
| SDE2 | 332.4482222 | 0.200660795 | 0.068706618 | 2.920545377 | 0.003494193 | 0.020895633 |
| GLCE | 123.744646 | -0.377716834 | 0.12934717 | -2.920178575 | 0.003498309 | 0.020906826 |
| RBBP7 | 592.4951839 | -0.155017828 | 0.053083982 | -2.920237365 | 0.003497649 | 0.020906826 |
| RGS20 | 16.79297879 | 0.714025457 | 0.244527604 | 2.920019845 | 0.003500091 | 0.020910771 |
| C17orf51 | 79.26190416 | -0.404490904 | 0.138659794 | -2.917146299 | 0.0035325 | 0.021097632 |
| PRKAR2B | 293.312646 | 0.362904362 | 0.124417712 | 2.916822353 | 0.003536171 | 0.021112788 |
| GTF2E2 | 176.2022327 | 0.280791292 | 0.096326565 | 2.914993302 | 0.003556962 | 0.021216931 |
| OXER1 | 82.48552776 | 0.475770692 | 0.163215355 | 2.914987338 | 0.00355703 | 0.021216931 |
| RGAG4 | 25.56163659 | -0.522725833 | 0.179317428 | -2.915086611 | 0.003555898 | 0.021216931 |
| SENP7 | 86.75331094 | -0.271039365 | 0.093002388 | -2.914326923 | 0.003564564 | 0.021255068 |
| LINC00630 | 23.42068928 | -0.516955331 | 0.177445093 | -2.913325589 | 0.003576015 | 0.021316529 |
| SLAIN2 | 1989.904165 | 0.209528042 | 0.071957697 | 2.911822511 | 0.003593267 | 0.021412519 |
| AUH | 166.9724948 | -0.174151802 | 0.059814703 | -2.911521628 | 0.00359673 | 0.0214263 |
| PYCARD | 1035.317164 | -0.350053307 | 0.120293619 | -2.909990649 | 0.003614396 | 0.021524656 |
| RPS16 | 9024.455298 | -0.387568182 | 0.133216126 | -2.909318828 | 0.003622173 | 0.021564078 |
| COQ10A | 46.12945586 | -0.371855164 | 0.127823508 | -2.909129693 | 0.003624365 | 0.021570237 |
| ASS1 | 41.91398355 | 0.636238607 | 0.218744866 | 2.908587606 | 0.003630654 | 0.021600771 |
| RPS15 | 6480.785083 | -0.209654374 | 0.072098028 | -2.907907189 | 0.003638563 | 0.021640914 |
| SCAF4 | 545.8199582 | 0.181546399 | 0.062469321 | 2.906168916 | 0.003658839 | 0.021754563 |
| PARD3B | 8.514282347 | -0.943575985 | 0.324696763 | -2.906022148 | 0.003660555 | 0.021757827 |
| HSPA2 | 16.66278784 | 0.865940404 | 0.298022355 | 2.905622305 | 0.003665236 | 0.021765704 |
| NPM1 | 5743.282902 | -0.171343102 | 0.058966135 | -2.905788246 | 0.003663293 | 0.021765704 |
| TCTEX1D2 | 58.41798203 | 0.608581736 | 0.209450622 | 2.905609587 | 0.003665385 | 0.021765704 |
| TCF20 | 56.05390507 | 0.304526688 | 0.1048108 | 2.905489594 | 0.00366679 | 0.021767115 |
| PPP1R10 | 653.9236751 | 0.18001565 | 0.061980535 | 2.90439006 | 0.003679694 | 0.021836758 |
| ZP3 | 7.629980162 | -1.032725818 | 0.355764681 | -2.902834014 | 0.003698026 | 0.021938559 |
| IDUA | 165.685076 | -0.335772505 | 0.115681999 | -2.90254756 | 0.00370141 | 0.021951644 |
| TSR1 | 180.4448274 | 0.243007807 | 0.083754511 | 2.901429481 | 0.003714644 | 0.022023122 |
| CRHBP | 2.924090468 | -2.192137558 | 0.755756413 | -2.900587436 | 0.003724639 | 0.022075358 |

| | | | | | | |
|---|---|---|---|---|---|---|
| IFNA1 | 69.95491945 | 0.512976833 | 0.176884841 | 2.900061032 | 0.0037309 | 0.022098408 |
| MBOAT7 | 1084.783445 | 0.286467569 | 0.098779706 | 2.900064998 | 0.003730853 | 0.022098408 |
| PRKCD | 634.1231659 | 0.319933334 | 0.11032726 | 2.899857504 | 0.003733323 | 0.022105735 |
| FNDC3B | 4506.620646 | 0.1845209 | 0.063658315 | 2.898614262 | 0.003748157 | 0.022186518 |
| MBOAT1 | 200.4248133 | -0.357374457 | 0.123299761 | -2.898419699 | 0.003750483 | 0.022193238 |
| PPP5C | 272.6751203 | 0.24617465 | 0.084941715 | 2.898159628 | 0.003753595 | 0.022204599 |
| SGK1 | 1.841049853 | -2.951678917 | 1.018761044 | -2.897322128 | 0.003763631 | 0.022256902 |
| SLC11A2 | 1043.676844 | 0.214743761 | 0.074145883 | 2.896233113 | 0.003776717 | 0.022327205 |
| ARHGAP3( | 962.6336294 | -0.308275888 | 0.106463382 | -2.895604873 | 0.003784286 | 0.022364852 |
| BAX | 859.0719899 | -0.189764586 | 0.065571745 | -2.893999346 | 0.00380369 | 0.022465278 |
| PDHA1 | 1304.590272 | 0.164883333 | 0.056972835 | 2.894069297 | 0.003802842 | 0.022465278 |
| H3F3A | 5374.595994 | -0.207601827 | 0.071757109 | -2.893118597 | 0.003814372 | 0.022521234 |
| CYP19A1 | 5.262911011 | 2.232308505 | 0.771921552 | 2.891885191 | 0.003829379 | 0.022602673 |
| KLHL7 | 106.3439572 | -0.307446896 | 0.106320399 | -2.891701861 | 0.003831614 | 0.022608704 |
| EMILIN1 | 14.69494292 | 0.825800577 | 0.285624234 | 2.891213272 | 0.003837576 | 0.022636717 |
| ARFGAP3 | 832.4883777 | 0.381301028 | 0.131894801 | 2.890948127 | 0.003840815 | 0.022648655 |
| ABCA5 | 75.56826822 | 0.588941576 | 0.203745687 | 2.89057199 | 0.003845414 | 0.022668603 |
| FGD5 | 1107.293405 | -0.183690223 | 0.063558959 | -2.890076001 | 0.003851487 | 0.02269722 |
| MCEE | 39.67626843 | -0.462278224 | 0.160009603 | -2.889065502 | 0.003863886 | 0.022763088 |
| NSUN7 | 31.5327669 | 0.498054239 | 0.172415367 | 2.888688214 | 0.003868524 | 0.022783212 |
| MANEA | 229.269517 | -0.262636189 | 0.090923473 | -2.888541103 | 0.003870334 | 0.02278667 |
| RTF1 | 1724.071875 | 0.187765657 | 0.065014299 | 2.888067098 | 0.003876172 | 0.02281383 |
| CD276 | 448.6379768 | 0.460348326 | 0.159476432 | 2.88662293 | 0.003894006 | 0.022905188 |
| HSF2 | 28.86458251 | -0.606526975 | 0.210117315 | -2.8866111 | 0.003894152 | 0.022905188 |
| ANKFY1 | 649.4744091 | 0.228002716 | 0.078992296 | 2.886391808 | 0.003896867 | 0.022913922 |
| KIAA0513 | 1581.620524 | -0.263100457 | 0.091167902 | -2.885889138 | 0.003903096 | 0.02294331 |
| MRAS | 1288.732213 | 0.265103468 | 0.091878727 | 2.88536287 | 0.003909627 | 0.022974455 |
| CACFD1 | 33.84590736 | -0.456304631 | 0.158174437 | -2.884819067 | 0.003916387 | 0.022999669 |
| NOXA1 | 27.20278696 | -0.560192913 | 0.194181221 | -2.884897464 | 0.003915412 | 0.022999669 |
| MAP3K10 | 99.13707752 | 0.368637446 | 0.127794083 | 2.884620617 | 0.003918856 | 0.02300692 |
| TREX1 | 45.69513084 | 0.493573993 | 0.171142963 | 2.88398649 | 0.003926756 | 0.023046038 |
| PPM1N | 75.05138768 | 0.313322724 | 0.108667202 | 2.88332375 | 0.003935028 | 0.023087314 |
| ALKBH8 | 142.7694587 | 0.205319144 | 0.071238636 | 2.882131902 | 0.003949943 | 0.023160241 |
| MTERFD3 | 31.06433911 | -0.490624825 | 0.170225064 | -2.882212614 | 0.003948932 | 0.023160241 |
| EXOC5 | 816.1499466 | 0.30079365 | 0.104384529 | 2.881592264 | 0.003956714 | 0.02319264 |
| ST6GALNA | 574.7519936 | 0.269770734 | 0.093656543 | 2.880425923 | 0.003971383 | 0.023271303 |
| HMGXB3 | 386.1480504 | 0.210096675 | 0.072950489 | 2.879989931 | 0.003976879 | 0.023296183 |
| KIFC1 | 9.054734441 | 0.915299903 | 0.317849951 | 2.879660354 | 0.003981038 | 0.023313218 |
| DNAJC9 | 129.1283204 | -0.236776287 | 0.082262337 | -2.878307317 | 0.003998154 | 0.023406098 |
| ARF4 | 3851.951136 | 0.201874244 | 0.070155089 | 2.877542435 | 0.00400786 | 0.023455548 |
| UBXN1 | 1238.651514 | -0.214034398 | 0.074384435 | -2.877408413 | 0.004009563 | 0.023458146 |
| LUZP1 | 622.5324076 | 0.131391636 | 0.045666118 | 2.877223679 | 0.004011911 | 0.023464517 |
| VENTX | 10.53735761 | -0.962292871 | 0.334465949 | -2.877102661 | 0.00401345 | 0.023466152 |
| LINC00032 | 5.735516099 | 1.143124308 | 0.39734721 | 2.876890234 | 0.004016153 | 0.023474589 |
| ATP6V0E2 | 43.49951637 | -0.488204482 | 0.169782209 | -2.875474902 | 0.004034202 | 0.023572695 |
| MYO1C | 5.843888949 | 1.415383615 | 0.492287758 | 2.875114387 | 0.004038811 | 0.02359223 |
| SLC25A42 | 4.671929611 | -1.157746984 | 0.40284387 | -2.873934718 | 0.004053927 | 0.023673107 |

| | | | | | | |
|---|---|---|---|---|---|---|
| ZNF620 | 10.19520418 | 0.564271964 | 0.196370519 | 2.873506499 | 0.004059427 | 0.023697797 |
| PAK6 | 7.573703962 | -1.04950686 | 0.365295181 | -2.873037792 | 0.004065455 | 0.023718123 |
| RRAS2 | 8.553445168 | 1.331112902 | 0.463299335 | 2.873116368 | 0.004064444 | 0.023718123 |
| PMS2 | 45.31270364 | -0.772169141 | 0.268808979 | -2.872557098 | 0.004071645 | 0.023746802 |
| DDB1 | 3288.183355 | -0.13659545 | 0.047580575 | -2.870823866 | 0.004094036 | 0.023869919 |
| RNASET2 | 2816.12393 | -0.219103634 | 0.076340532 | -2.87008262 | 0.004103646 | 0.023918465 |
| SAYSD1 | 111.0040572 | -0.290031934 | 0.101066023 | -2.869727376 | 0.004108258 | 0.023937863 |
| CYP4V2 | 568.1624155 | -0.203365922 | 0.070876132 | -2.869314622 | 0.004113624 | 0.023939184 |
| DCTN6 | 786.6345168 | 0.223774023 | 0.077986553 | 2.86939241 | 0.004112612 | 0.023939184 |
| DUSP7 | 790.8544161 | 0.294276974 | 0.102553216 | 2.869505084 | 0.004111147 | 0.023939184 |
| IL15 | 34.06358903 | 0.49932144 | 0.174012213 | 2.869462044 | 0.004111707 | 0.023939184 |
| SIKE1 | 149.1043304 | 0.299438294 | 0.104399221 | 2.868204293 | 0.004128089 | 0.024015863 |
| TRIM4 | 177.3919199 | -0.251142077 | 0.087586023 | -2.867376177 | 0.004138907 | 0.024071285 |
| NKG7 | 47.35364629 | 0.623806144 | 0.217667713 | 2.865864372 | 0.004158724 | 0.024178989 |
| DNAJC17 | 43.07978991 | -0.428661377 | 0.149668144 | -2.864078919 | 0.004182238 | 0.024308117 |
| SAMD4A | 221.9124092 | -0.351638933 | 0.122853376 | -2.862265129 | 0.004206249 | 0.02444005 |
| VLDLR | 40.086147 | -0.627308076 | 0.219180837 | -2.86205713 | 0.00420901 | 0.024448472 |
| RGS2 | 24.42319844 | -0.753641082 | 0.263406322 | -2.861135133 | 0.004221271 | 0.024512048 |
| GIMAP7 | 24.33316778 | -0.949534655 | 0.332113399 | -2.859067591 | 0.004248882 | 0.024664696 |
| CDK1 | 11.54510461 | 1.078784272 | 0.377412662 | 2.858367992 | 0.004258262 | 0.024711449 |
| HMGN4 | 194.5566297 | -0.175280652 | 0.061326822 | -2.858140134 | 0.004261322 | 0.024713808 |
| MORC3 | 932.7588285 | 0.23663766 | 0.082791861 | 2.858223703 | 0.004260199 | 0.024713808 |
| SLC26A2 | 76.36014218 | -0.240103196 | 0.084022168 | -2.857617239 | 0.004268349 | 0.024746864 |
| RHOQ | 1393.272652 | 0.283047809 | 0.099080427 | 2.856747972 | 0.004280055 | 0.024799301 |
| UBE2Q2 | 150.2907753 | -0.356346592 | 0.124734558 | -2.856839326 | 0.004278824 | 0.024799301 |
| EIF2AK3 | 310.2798114 | 0.315074987 | 0.110299001 | 2.856553399 | 0.00428268 | 0.024806792 |
| KIAA1522 | 382.8041423 | 0.318843404 | 0.111626659 | 2.85633742 | 0.004285594 | 0.024815961 |
| ACTL6A | 72.55335242 | -0.451783197 | 0.15818703 | -2.856006575 | 0.004290063 | 0.024834118 |
| SLC16A10 | 191.9357504 | 0.403379356 | 0.141245594 | 2.855872138 | 0.004291879 | 0.024836919 |
| TET3 | 140.7725686 | -0.411678986 | 0.144172897 | -2.855453385 | 0.004297543 | 0.024861974 |
| TSEN15 | 64.49280582 | -0.348713197 | 0.12220072 | -2.853609996 | 0.004322556 | 0.024998917 |
| PTPLAD1 | 338.4295079 | -0.21607177 | 0.075744252 | -2.852649069 | 0.004335647 | 0.025066848 |
| CCNB1IP1 | 76.81043616 | -0.353188333 | 0.123817682 | -2.852487038 | 0.004337858 | 0.025071851 |
| RNF20 | 464.3847664 | -0.147348218 | 0.051662162 | -2.852149645 | 0.004342465 | 0.025090697 |
| HLA-DPB1 | 11013.53389 | -0.242140429 | 0.085027663 | -2.847784125 | 0.004402478 | 0.025429562 |
| CCDC36 | 2.466913198 | -1.744986527 | 0.61280707 | -2.847530016 | 0.004405994 | 0.025441986 |
| FKBP1AP1 | 9.501396867 | 1.048584871 | 0.368305527 | 2.847051683 | 0.00441262 | 0.025472353 |
| SPOPL | 213.8241093 | -0.403571017 | 0.141794396 | -2.846170422 | 0.004424851 | 0.025535047 |
| RPS25 | 4301.435436 | -0.4238922 | 0.1489902 | -2.845101214 | 0.004439731 | 0.025612989 |
| HOPX | 9.855197189 | 1.119788751 | 0.393662329 | 2.844541294 | 0.004447542 | 0.025650108 |
| KLHDC8B | 1038.308057 | -0.314783182 | 0.11068547 | -2.84394312 | 0.0044559 | 0.02569036 |
| TREML4 | 21.36365012 | 1.304752189 | 0.45887508 | 2.843371204 | 0.004463905 | 0.025728549 |
| FAM83H | 78.72128618 | 0.348535909 | 0.122599719 | 2.842876888 | 0.004470834 | 0.025760518 |
| MAFF | 24.83734459 | 0.923482882 | 0.324854183 | 2.842761242 | 0.004472456 | 0.0257619 |
| IER5L | 63.40501704 | -0.448813236 | 0.157891088 | -2.842549519 | 0.004475428 | 0.025771051 |
| FBN2 | 8.445875424 | -1.352758867 | 0.476056176 | -2.841595039 | 0.004488847 | 0.025840338 |
| ERCC6 | 44.14338096 | -0.37867678 | 0.133284861 | -2.841108717 | 0.004495698 | 0.025867057 |

| | | | | | | |
|---|---|---|---|---|---|---|
| KIAA0825 | 11.00160212 | -0.820081032 | 0.288652322 | -2.841068539 | 0.004496265 | 0.025867057 |
| H2AFX | 236.7206503 | 0.341788095 | 0.12030911 | 2.84091616 | 0.004498414 | 0.025871432 |
| ZNF850 | 20.28939043 | -0.556665858 | 0.196003595 | -2.840079831 | 0.004510225 | 0.025931356 |
| FAM126B | 262.3496785 | 0.606222718 | 0.213463342 | 2.839938284 | 0.004512226 | 0.025934863 |
| FAU | 2268.645416 | -0.214917394 | 0.075707914 | -2.838770519 | 0.004528771 | 0.026021932 |
| TTC7B | 911.6379466 | 0.209800845 | 0.073911112 | 2.838556189 | 0.004531814 | 0.026031388 |
| SLC39A12 | 5.755623495 | 1.219139472 | 0.42953049 | 2.838307173 | 0.004535351 | 0.026043678 |
| P4HA2 | 282.6392338 | 0.237809515 | 0.083817552 | 2.837228112 | 0.004550709 | 0.026123816 |
| QARS | 752.7053812 | -0.338869918 | 0.119469034 | -2.836466533 | 0.004561576 | 0.026178135 |
| XPR1 | 280.3750814 | 0.236386546 | 0.083358289 | 2.835789322 | 0.004571259 | 0.026225628 |
| ERCC8 | 68.46647147 | -0.381779045 | 0.134667843 | -2.834968142 | 0.004583026 | 0.026285041 |
| ACAP3 | 93.22294266 | -0.267005124 | 0.094200243 | -2.834441982 | 0.004590579 | 0.026312167 |
| TMEM164 | 422.9622112 | 0.290000074 | 0.10230965 | 2.834532951 | 0.004589272 | 0.026312167 |
| ACOT8 | 168.4909326 | -0.291257776 | 0.102778362 | -2.833843347 | 0.004599187 | 0.026353399 |
| ZC3H3 | 134.9089106 | 0.284730033 | 0.100548704 | 2.831762337 | 0.004629224 | 0.026517358 |
| MRPS18A | 389.9006364 | 0.203887128 | 0.072118157 | 2.827126162 | 0.004696782 | 0.026896077 |
| CSNK1A1 | 2978.075069 | 0.293195048 | 0.103727721 | 2.826583327 | 0.00470475 | 0.02693343 |
| DHFR | 287.4330165 | -0.259331562 | 0.091780031 | -2.825577178 | 0.004719552 | 0.027009867 |
| SRCIN1 | 44.00656923 | -0.428904981 | 0.151799387 | -2.825472426 | 0.004721095 | 0.027010404 |
| TAF1D | 518.7833093 | -0.239468836 | 0.084759867 | -2.825262066 | 0.004724196 | 0.027019849 |
| ATG7 | 476.3917758 | -0.206475989 | 0.073090044 | -2.824953664 | 0.004728745 | 0.027035453 |
| MBNL2 | 999.1020326 | -0.239596197 | 0.084816402 | -2.824880456 | 0.004729826 | 0.027035453 |
| SGSM3 | 352.4104148 | -0.311190762 | 0.1102292 | -2.823124562 | 0.004755809 | 0.027175635 |
| TMC6 | 546.7356007 | -0.195850859 | 0.069395646 | -2.82223554 | 0.004769014 | 0.027242735 |
| GINS3 | 15.32156542 | 0.584787769 | 0.20726456 | 2.82145568 | 0.004780624 | 0.02730069 |
| SLC15A2 | 7.422877681 | -1.076801065 | 0.381717487 | -2.820937219 | 0.004788357 | 0.027336474 |
| PABPC4 | 2331.205334 | -0.349588602 | 0.123990158 | -2.8194867 | 0.004810052 | 0.027451919 |
| EVI2B | 1168.708846 | -0.323697405 | 0.114818665 | -2.819205433 | 0.00481427 | 0.027467575 |
| FIGNL1 | 57.20155464 | 0.386563969 | 0.137133645 | 2.818884957 | 0.004819079 | 0.027486597 |
| EEF1E1 | 349.725394 | 0.210565805 | 0.074719257 | 2.818092868 | 0.004830983 | 0.027539483 |
| LSAMP | 171.2585552 | -0.408389227 | 0.144917985 | -2.818071387 | 0.004831307 | 0.027539483 |
| NET1 | 36.27335961 | -0.527738269 | 0.187457218 | -2.81524645 | 0.004873984 | 0.027774257 |
| TIGD1 | 62.15742748 | -0.358404898 | 0.12734633 | -2.814410886 | 0.004886672 | 0.027838048 |
| ITPKB | 384.6837616 | -0.345199371 | 0.122718567 | -2.812935159 | 0.004909155 | 0.027957577 |
| LOC22072 | 139.5418008 | -0.246959105 | 0.087827656 | -2.811860356 | 0.004925588 | 0.028042594 |
| NRBP2 | 90.35730669 | -0.298435021 | 0.106183817 | -2.810550889 | 0.004945677 | 0.028147128 |
| SLC25A51 | 215.0762103 | -0.279627624 | 0.099495084 | -2.810466736 | 0.00494697 | 0.028147128 |
| KRTCAP2 | 1186.191312 | -0.134817494 | 0.047973011 | -2.810277947 | 0.004949873 | 0.028155048 |
| DFNB31 | 10.65057031 | 0.817507524 | 0.29098345 | 2.809463984 | 0.004962407 | 0.028217728 |
| SCN8A | 4.542005399 | 1.529805806 | 0.54455578 | 2.809272917 | 0.004965353 | 0.028225868 |
| ARHGAP2 | 15.56100435 | -0.822008259 | 0.292637188 | -2.808967186 | 0.004970071 | 0.028239933 |
| C1orf85 | 1632.702046 | -0.257997363 | 0.091855852 | -2.808719922 | 0.00497389 | 0.028239933 |
| KIAA1755 | 3.62123196 | 2.018713966 | 0.718706583 | 2.808815189 | 0.004972418 | 0.028239933 |
| PITPNA-AS | 16.30628239 | -0.572213807 | 0.203723884 | -2.808771348 | 0.004973095 | 0.028239933 |
| ZAK | 275.0402931 | -0.310622561 | 0.110607686 | -2.808327085 | 0.004979962 | 0.028265796 |
| ITFG3 | 329.6315122 | -0.206908407 | 0.073695845 | -2.807599378 | 0.004991228 | 0.028321114 |
| LOC38964 | 39.99867152 | 0.431665226 | 0.153806455 | 2.806548178 | 0.005007542 | 0.028405036 |

| | | | | | | |
|---|---|---|---|---|---|---|
| BNIP3L | 1670.323107 | -0.286537028 | 0.10211607 | -2.805993506 | 0.00501617 | 0.028445319 |
| EHMT2 | 236.1635596 | -0.209973617 | 0.074858992 | -2.804921773 | 0.005032879 | 0.028531388 |
| CASP2 | 337.1898623 | -0.16720685 | 0.059615879 | -2.804736787 | 0.005035768 | 0.028539084 |
| C14orf166 | 923.1456007 | -0.225293003 | 0.080333487 | -2.804471848 | 0.005039909 | 0.028553865 |
| TAS1R3 | 3.404898841 | -1.233180247 | 0.43998807 | -2.802758373 | 0.005066761 | 0.028697274 |
| CDCA2 | 1.989017313 | 1.937971794 | 0.691932273 | 2.800811395 | 0.00509743 | 0.028853437 |
| TMEM67 | 28.03547837 | 0.324525905 | 0.115866881 | 2.800851303 | 0.0050968 | 0.028853437 |
| CTSD | 127278.683 | -0.302075738 | 0.107892796 | -2.799776718 | 0.005113797 | 0.028937286 |
| C7orf69 | 4.251450077 | 1.55003481 | 0.553688076 | 2.799472984 | 0.00511861 | 0.028955731 |
| SFXN4 | 218.1587254 | 0.210663343 | 0.075255605 | 2.799304381 | 0.005121284 | 0.028962064 |
| PQLC2 | 856.2624233 | 0.225669359 | 0.080643344 | 2.798363092 | 0.005136234 | 0.029037798 |
| DISC1 | 58.82793006 | -0.395444728 | 0.141321148 | -2.798199243 | 0.00513884 | 0.029043721 |
| DPY30 | 610.0081968 | -0.236387142 | 0.084495512 | -2.797629576 | 0.005147911 | 0.029086168 |
| PTBP1 | 2554.562039 | 0.301005522 | 0.107603567 | 2.797356367 | 0.005152267 | 0.029086857 |
| RPS6KA4 | 457.9882426 | -0.332337719 | 0.118800469 | -2.797444507 | 0.005150861 | 0.029086857 |
| TRPM7 | 508.3252684 | -0.212868727 | 0.076097158 | -2.797328199 | 0.005152716 | 0.029086857 |
| DOCK3 | 154.981667 | 0.436853336 | 0.156176784 | 2.797172061 | 0.005155207 | 0.029092105 |
| TSPAN32 | 21.60834281 | -0.683645411 | 0.244430442 | -2.796891437 | 0.005159687 | 0.029108569 |
| ARPC2 | 20318.47171 | 0.185833488 | 0.066461067 | 2.796125572 | 0.00517193 | 0.029168809 |
| HSPE1 | 1190.401449 | -0.188249395 | 0.06733163 | -2.795853825 | 0.005176281 | 0.029184512 |
| CPQ | 464.7047979 | -0.225662351 | 0.080730091 | -2.795269376 | 0.005185649 | 0.029228486 |
| FBP1 | 3755.108325 | 0.356196324 | 0.127534845 | 2.792933373 | 0.005223245 | 0.029430394 |
| MTHFD1L | 131.6778193 | -0.479468088 | 0.171677137 | -2.792847642 | 0.005224629 | 0.029430394 |
| ATHL1 | 36.79178611 | -0.457791311 | 0.163957289 | -2.792137598 | 0.005236108 | 0.029486142 |
| HNMT | 1851.478218 | -0.225606802 | 0.080810557 | -2.791798623 | 0.005241597 | 0.02950813 |
| PLAA | 382.5060446 | 0.25420389 | 0.091068006 | 2.791363287 | 0.005248653 | 0.029538928 |
| GUSBP11 | 83.12021363 | -0.307997161 | 0.110346234 | -2.791188685 | 0.005251485 | 0.029545945 |
| SRD5A1 | 102.4110497 | 0.317561308 | 0.11378242 | 2.790952321 | 0.005255321 | 0.029558605 |
| PON2 | 74.6136575 | -0.612839823 | 0.219681869 | -2.789669565 | 0.005276186 | 0.029667004 |
| POP1 | 25.03706143 | 0.480081505 | 0.172107185 | 2.789433247 | 0.005280038 | 0.029679707 |
| TMED1 | 349.5985926 | 0.376546403 | 0.135034635 | 2.788517209 | 0.005294993 | 0.029754797 |
| ALDH5A1 | 76.65384975 | -0.44308088 | 0.15894087 | -2.787708908 | 0.005308222 | 0.02982014 |
| ARL8B | 6657.36751 | 0.22333572 | 0.080132354 | 2.787085489 | 0.005318445 | 0.029868565 |
| KIAA0930 | 3960.715491 | -0.20940252 | 0.075140148 | -2.786826039 | 0.005322704 | 0.029883481 |
| TSHZ1 | 22.59136059 | -0.492399102 | 0.17676283 | -2.785648437 | 0.005342077 | 0.029983214 |
| POLR2A | 1936.552199 | 0.250595723 | 0.089973893 | 2.785204867 | 0.005349391 | 0.030015224 |
| NAA40 | 130.7167352 | -0.325346326 | 0.116832376 | -2.784727477 | 0.005357273 | 0.030050398 |
| ADI1 | 684.9619572 | -0.272821497 | 0.098010274 | -2.783600988 | 0.005375912 | 0.030145878 |
| ELOVL6 | 8.901181462 | 0.759593718 | 0.272906615 | 2.78334667 | 0.005380129 | 0.030160444 |
| LOC92249 | 63.67434299 | -0.466005567 | 0.167439168 | -2.783133559 | 0.005383664 | 0.030171186 |
| ATP9B | 624.5248038 | 0.137267067 | 0.049376085 | 2.780031385 | 0.005435365 | 0.030445809 |
| SYNM | 12.41601227 | 0.830935579 | 0.298897979 | 2.77999732 | 0.005435935 | 0.030445809 |
| ENTPD7 | 259.7430587 | 0.363579265 | 0.130891062 | 2.77772416 | 0.005474107 | 0.030650392 |
| TRIB2 | 93.43159265 | 0.384511436 | 0.138445347 | 2.777351823 | 0.005480382 | 0.030676312 |
| RARA | 153.6724178 | -0.547585283 | 0.19718061 | -2.777074701 | 0.005485057 | 0.03069326 |
| TBL2 | 473.7058605 | -0.186857026 | 0.067324315 | -2.775476064 | 0.005512096 | 0.030835302 |
| MTCH2 | 977.5714264 | -0.161730262 | 0.058275005 | -2.775293839 | 0.005515185 | 0.030843326 |

| | | | | | | |
|---|---|---|---|---|---|---|
| KAT5 | 580.3743502 | 0.204577784 | 0.07373909 | 2.774346456 | 0.005531274 | 0.030924019 |
| ANKRD24 | 11.11664547 | 0.772142905 | 0.278329254 | 2.774206786 | 0.005533649 | 0.03092802 |
| GTF3C5 | 197.7607088 | -0.288756806 | 0.104161947 | -2.772190936 | 0.005568037 | 0.031109633 |
| TMEM140 | 640.249201 | 0.515922734 | 0.186112167 | 2.772106421 | 0.005569482 | 0.031109633 |
| HDAC10 | 330.590529 | -0.235307067 | 0.084887879 | -2.771974869 | 0.005571734 | 0.031112882 |
| MICU1 | 1023.014828 | 0.144875646 | 0.052270959 | 2.771627873 | 0.005577676 | 0.031136733 |
| DAAM1 | 181.9058822 | 0.256067647 | 0.092450018 | 2.769795541 | 0.005609149 | 0.03130305 |
| NEK6 | 119.9738741 | 0.385558756 | 0.139288147 | 2.768065806 | 0.005639007 | 0.031441428 |
| TECPR2 | 179.974372 | 0.331206608 | 0.119651338 | 2.768097834 | 0.005638453 | 0.031441428 |
| ZBED2 | 6.495766511 | 2.584633743 | 0.93373128 | 2.768070213 | 0.005638931 | 0.031441428 |
| FAM186B | 4.738271546 | 1.171908112 | 0.423391307 | 2.767907828 | 0.005641741 | 0.031447263 |
| NUS1 | 1716.259062 | 0.356844992 | 0.128929083 | 2.767761795 | 0.005644269 | 0.031451948 |
| GTDC1 | 247.8840128 | -0.28572886 | 0.103299731 | -2.76601747 | 0.005674549 | 0.031599613 |
| NETO2 | 314.9421263 | 0.252489549 | 0.09128518 | 2.765942386 | 0.005675856 | 0.031599613 |
| PSMB5 | 476.391923 | 0.248716941 | 0.089917674 | 2.766051775 | 0.005673953 | 0.031599613 |
| TSC2 | 530.9814258 | -0.220402822 | 0.079687744 | -2.765830874 | 0.005677797 | 0.031600978 |
| TAOK2 | 385.4742902 | -0.245756589 | 0.088859841 | -2.765665404 | 0.005680679 | 0.031607575 |
| AMOTL1 | 275.909851 | -0.264902127 | 0.095797129 | -2.765240783 | 0.005688079 | 0.031639304 |
| ENTPD5 | 188.6508432 | 0.263993417 | 0.095515566 | 2.763878474 | 0.005711881 | 0.031761783 |
| TRIP10 | 308.9189729 | 0.436609156 | 0.157975047 | 2.763785567 | 0.005713507 | 0.031761783 |
| KRBA1 | 21.52667036 | -0.476411824 | 0.1723959 | -2.763475367 | 0.005718941 | 0.031782507 |
| HERC1 | 401.9159211 | -0.272227577 | 0.098521647 | -2.763124494 | 0.005725093 | 0.031805602 |
| TRAP1 | 344.023372 | 0.173292074 | 0.062717819 | 2.763043673 | 0.00572651 | 0.031805602 |
| NCF2 | 11779.96189 | 0.214959407 | 0.077816926 | 2.762373405 | 0.005738281 | 0.03186148 |
| NISCH | 205.5343461 | -0.326757572 | 0.118297467 | -2.762168794 | 0.005741879 | 0.031871958 |
| CCSAP | 783.0014692 | -0.178558967 | 0.064649879 | -2.761938123 | 0.005745937 | 0.031884986 |
| FAM102A | 55.9397287 | -0.454877638 | 0.164719215 | -2.761533539 | 0.005753061 | 0.031905516 |
| TERF1 | 26.52297316 | 0.56836385 | 0.205812501 | 2.761561362 | 0.005752571 | 0.031905516 |
| GOLGA5 | 626.3976952 | 0.19129707 | 0.069338881 | 2.758871599 | 0.005800132 | 0.032156995 |
| PORCN | 118.5473869 | -0.418217266 | 0.151680923 | -2.757217316 | 0.005829559 | 0.032310532 |
| ACAD8 | 224.0064765 | -0.245802455 | 0.089152121 | -2.757112818 | 0.005831423 | 0.03231125 |
| LATS2 | 152.4560112 | 0.23738679 | 0.086107044 | 2.756880021 | 0.005835576 | 0.03232465 |
| PYCR1 | 5.479720525 | 0.966697871 | 0.350677065 | 2.756661233 | 0.005839482 | 0.032336671 |
| DLX4 | 6.217790029 | -1.093492195 | 0.3968851 | -2.755185812 | 0.005865881 | 0.032473212 |
| LRWD1 | 107.0368749 | 0.315511184 | 0.114581194 | 2.753603571 | 0.005894312 | 0.03262091 |
| GALNT14 | 7.1453559 | 1.589067732 | 0.577533263 | 2.751473956 | 0.005932774 | 0.032815079 |
| LOC72985: | 252.4155934 | -0.269325231 | 0.097884268 | -2.751465965 | 0.005932918 | 0.032815079 |
| ZNF702P | 98.42276 | 0.493063372 | 0.179248254 | 2.750729014 | 0.005946281 | 0.032879229 |
| TTYH2 | 461.479088 | 0.304965255 | 0.110873338 | 2.750573395 | 0.005949106 | 0.032885092 |
| BDH1 | 123.9647373 | 0.288864695 | 0.105040359 | 2.750035296 | 0.005958885 | 0.032929376 |
| LOC38963 | 6.49420314 | 1.282131677 | 0.46628436 | 2.749677639 | 0.005965392 | 0.032955563 |
| LIPT1 | 9.758227376 | -0.747985776 | 0.272060454 | -2.749336649 | 0.005971602 | 0.032980092 |
| KLHDC2 | 338.536439 | -0.244465029 | 0.088928825 | -2.748996508 | 0.005977802 | 0.033004554 |
| ELP4 | 80.49714998 | -0.286652843 | 0.104295143 | -2.748477394 | 0.005987276 | 0.03304707 |
| C1QB | 26069.03887 | 0.270843618 | 0.098547277 | 2.748362275 | 0.005989379 | 0.033048887 |
| HP | 26.45930642 | -1.291867525 | 0.470139481 | -2.747838838 | 0.005998949 | 0.033091892 |
| NCAPD3 | 135.765087 | -0.237882188 | 0.086584507 | -2.747399 | 0.006007001 | 0.033119335 |

| | | | | | | |
|---|---|---|---|---|---|---|
| PRPS1 | 378.2709755 | -0.199101941 | 0.072469937 | -2.747372922 | 0.006007478 | 0.033119335 |
| DESI2 | 140.6139113 | -0.351951435 | 0.128143079 | -2.746550469 | 0.006022562 | 0.03319267 |
| COG2 | 226.0968551 | -0.188748071 | 0.06872684 | -2.746351643 | 0.006026213 | 0.033202975 |
| LOC64285Z | 27.45944083 | -0.343219616 | 0.124990269 | -2.745970687 | 0.006033215 | 0.033231728 |
| MFAP1 | 622.4849843 | 0.190599322 | 0.069413796 | 2.745842074 | 0.006035581 | 0.033234934 |
| LAYN | 7.647061493 | 0.854243864 | 0.311186519 | 2.745118479 | 0.006048906 | 0.033298465 |
| ASAH1 | 16399.57393 | -0.262172514 | 0.095521175 | -2.744653367 | 0.006057484 | 0.033335842 |
| SF3B14 | 315.2477657 | -0.137929604 | 0.050269253 | -2.743816469 | 0.006072948 | 0.033411076 |
| ZNF417 | 81.59087421 | -0.295323827 | 0.107639304 | -2.743643047 | 0.006076157 | 0.033418863 |
| CYB5A | 1130.214399 | 0.248016193 | 0.090421108 | 2.742901495 | 0.006089895 | 0.03348454 |
| MAD2L1BF | 316.9476813 | 0.294995225 | 0.107565866 | 2.742461304 | 0.006098063 | 0.033519562 |
| NAGA | 1441.567481 | -0.218989404 | 0.079857062 | -2.742267227 | 0.006101668 | 0.033529484 |
| ARPC4 | 2272.247903 | -0.155938665 | 0.05687476 | -2.741790297 | 0.006110534 | 0.033556604 |
| MOCOS | 44.11792787 | 0.388320138 | 0.141639224 | 2.741614409 | 0.006113807 | 0.033556604 |
| NANOG | 125.4926879 | -0.460926764 | 0.168122196 | -2.741617555 | 0.006113748 | 0.033556604 |
| ZNF93 | 67.37117866 | -0.400742505 | 0.146155126 | -2.741898391 | 0.006108523 | 0.033556604 |
| ABCA7 | 21.46198487 | -0.661424288 | 0.241275262 | -2.741368023 | 0.006118393 | 0.033571892 |
| PTER | 87.94324671 | -0.27091304 | 0.098845579 | -2.740770432 | 0.006129532 | 0.033623107 |
| CMTM8 | 11.07288267 | -0.711756015 | 0.259786022 | -2.739777956 | 0.00614807 | 0.033704956 |
| EEFSEC | 111.3514952 | -0.22152387 | 0.080854385 | -2.739787952 | 0.006147883 | 0.033704956 |
| RGMB | 38.62120945 | -0.41349178 | 0.150961186 | -2.739060227 | 0.006161509 | 0.033762578 |
| SH3PXD2A | 164.0595326 | -0.568106904 | 0.207412238 | -2.739023068 | 0.006162205 | 0.033762578 |
| CEBPE | 12.88105079 | -0.73968414 | 0.270103591 | -2.738520199 | 0.006171637 | 0.033804316 |
| KIAA0391 | 299.5046507 | -0.235446392 | 0.085984141 | -2.738253702 | 0.006176641 | 0.033821781 |
| TRIM37 | 87.40730243 | -0.337997905 | 0.123466224 | -2.737573849 | 0.006189422 | 0.033881813 |
| AK1 | 398.6440279 | 0.265426177 | 0.097025767 | 2.735625643 | 0.006226181 | 0.034063024 |
| NAF1 | 218.4057458 | 0.400281154 | 0.14632082 | 2.735640446 | 0.006225901 | 0.034063024 |
| CXorf26 | 809.3346194 | 0.187798208 | 0.068652403 | 2.735493593 | 0.00622868 | 0.034066692 |
| LOC40132: | 35.43114217 | -0.604507 | 0.221027386 | -2.734986876 | 0.006238276 | 0.034109167 |
| RCL1 | 110.8181706 | -0.309582529 | 0.113201703 | -2.734786868 | 0.006242068 | 0.034119886 |
| PMP22 | 417.433758 | -0.3115002 | 0.113907429 | -2.734678523 | 0.006244123 | 0.034121108 |
| KDM4B | 207.6082679 | -0.388788221 | 0.142211608 | -2.733871217 | 0.006259452 | 0.034184827 |
| STAU2 | 267.4281166 | -0.192077997 | 0.070258524 | -2.73387464 | 0.006259387 | 0.034184827 |
| STOML1 | 25.25348417 | 0.537365982 | 0.196602768 | 2.733257464 | 0.006271129 | 0.034238564 |
| TAMM41 | 114.186219 | -0.260063822 | 0.095157265 | -2.732989663 | 0.00627623 | 0.034256378 |
| ANXA11 | 3944.809434 | 0.217789682 | 0.079817007 | 2.728612493 | 0.006360139 | 0.034704198 |
| ARSK | 52.4973803 | -0.418763174 | 0.153511742 | -2.727890181 | 0.006374082 | 0.034759922 |
| RPL23AP82 | 247.0286714 | -0.45058454 | 0.165173917 | -2.727940035 | 0.006373119 | 0.034759922 |
| ACTN1 | 6522.073951 | 0.37728808 | 0.138387862 | 2.726309049 | 0.0064047 | 0.034906457 |
| SMYD3 | 36.56679912 | -0.537825913 | 0.19727186 | -2.726318453 | 0.006404517 | 0.034906457 |
| EFR3B | 20.81338094 | 0.783732004 | 0.287500026 | 2.726024111 | 0.006410232 | 0.03492639 |
| ALDH4A1 | 44.0122839 | -0.425851464 | 0.156251597 | -2.725421507 | 0.006421944 | 0.034969758 |
| TMBIM4 | 1544.304784 | -0.120193736 | 0.044100247 | -2.725466274 | 0.006421074 | 0.034969758 |
| TMEM218 | 3.773736637 | -1.161619501 | 0.426396597 | -2.724270103 | 0.006444378 | 0.035081664 |
| HELB | 45.41435495 | 0.430834371 | 0.15819787 | 2.723389209 | 0.006461588 | 0.035154814 |
| NEDD4L | 108.8886692 | 0.425081446 | 0.156083691 | 2.723419996 | 0.006460986 | 0.035154814 |
| UBB | 25367.19625 | 0.205863629 | 0.075609552 | 2.722719851 | 0.006474694 | 0.035215833 |

| | | | | | | |
|---|---|---|---|---|---|---|
| LGALS8 | 842.90194 | 0.157310758 | 0.057812137 | 2.721068031 | 0.006507137 | 0.035381963 |
| RPL36AL | 1182.816773 | -0.34305643 | 0.126107032 | -2.720359241 | 0.006521103 | 0.035447558 |
| EED | 405.3282261 | 0.187044451 | 0.068772529 | 2.719755303 | 0.006533024 | 0.035502004 |
| AVIL | 16.05000425 | 0.632760677 | 0.232691265 | 2.719314264 | 0.006541742 | 0.035539016 |
| AP4E1 | 140.1802087 | 0.316309597 | 0.116350311 | 2.718596923 | 0.006555945 | 0.035605791 |
| EPAS1 | 1363.866595 | -0.28207871 | 0.10377353 | -2.718214463 | 0.006563528 | 0.035636591 |
| FAM54B | 434.8586772 | -0.215698528 | 0.079374845 | -2.717467078 | 0.00657837 | 0.035706771 |
| MAX | 357.0981662 | 0.353676335 | 0.130158017 | 2.71728428 | 0.006582005 | 0.035716096 |
| VSIG10 | 39.26907227 | -0.468315858 | 0.172389526 | -2.716614338 | 0.006595341 | 0.035778045 |
| SH3BP1 | 1489.781215 | 0.231913912 | 0.085417004 | 2.715078962 | 0.006625997 | 0.035928116 |
| TRIM65 | 296.418249 | -0.182390503 | 0.067177938 | -2.715035762 | 0.006626861 | 0.035928116 |
| POLE3 | 380.7219099 | -0.279370099 | 0.10290323 | -2.714881712 | 0.006629945 | 0.035934377 |
| STIM1 | 670.0488113 | -0.289396103 | 0.106632352 | -2.713961551 | 0.006648388 | 0.036023863 |
| HIST1H2BH | 87.02442853 | -0.554302452 | 0.204253833 | -2.713792172 | 0.006651788 | 0.036031809 |
| GK3P | 6.072383555 | 0.859606974 | 0.317108429 | 2.710766714 | 0.006712784 | 0.036351645 |
| KIF1C | 1270.634169 | -0.245031303 | 0.090413391 | -2.710121812 | 0.00672585 | 0.036411822 |
| STAM | 235.1863663 | -0.330759937 | 0.122062991 | -2.709747917 | 0.006733436 | 0.036442303 |
| TOB1-AS1 | 4.492676844 | -1.027736633 | 0.379432878 | -2.70861249 | 0.00675652 | 0.03655662 |
| CTSL2 | 12.03952572 | 0.720204359 | 0.265975931 | 2.7077802 | 0.006773487 | 0.036633026 |
| TRAPPC9 | 473.3227681 | -0.147012215 | 0.05429359 | -2.707726931 | 0.006774574 | 0.036633026 |
| DUSP28 | 28.53440174 | -0.460475666 | 0.170092763 | -2.707203157 | 0.006785272 | 0.036680232 |
| HSPA1A | 6876.624627 | 0.267779457 | 0.098926258 | 2.70685926 | 0.006792305 | 0.036699847 |
| WDR35 | 54.18799765 | -0.35466217 | 0.131024764 | -2.706833114 | 0.00679284 | 0.036699847 |
| TRAPPC6A | 161.2505676 | -0.385589295 | 0.142489781 | -2.706083851 | 0.006808185 | 0.036772094 |
| RUNX1 | 124.6442909 | 0.52505081 | 0.19407616 | 2.705385394 | 0.006822519 | 0.036838832 |
| OR2A20P | 614.6824572 | 0.474735375 | 0.175509488 | 2.704898633 | 0.006832524 | 0.036882168 |
| LAMC1 | 304.532574 | 0.296590986 | 0.10971218 | 2.703355135 | 0.006864336 | 0.037032438 |
| TLN1 | 3360.384771 | -0.205191086 | 0.075900515 | -2.703421516 | 0.006862965 | 0.037032438 |
| ATP11B | 859.1335306 | 0.185095997 | 0.068503749 | 2.701983466 | 0.006892719 | 0.037174798 |
| CIITA | 1114.073906 | -0.332813196 | 0.123188916 | -2.701648872 | 0.006899658 | 0.037201457 |
| RNF144A | 2.191552005 | 1.611213053 | 0.596456037 | 2.701310665 | 0.006906679 | 0.037228539 |
| HPCAL1 | 1695.940881 | -0.30032288 | 0.11119168 | -2.700947419 | 0.006914227 | 0.037258445 |
| ANP32AP1 | 8.264060591 | -1.003858737 | 0.371726378 | -2.700531347 | 0.006922881 | 0.037293028 |
| POR | 1939.473457 | -0.186854622 | 0.069193973 | -2.700446501 | 0.006924647 | 0.037293028 |
| SNX8 | 1103.638677 | 0.176525761 | 0.065417213 | 2.698460431 | 0.006966102 | 0.037494611 |
| TMCO6 | 59.21118838 | -0.290609702 | 0.107693126 | -2.698498159 | 0.006965312 | 0.037494611 |
| CES2 | 330.3520102 | -0.147106191 | 0.054540176 | -2.697207839 | 0.006992361 | 0.037621635 |
| NPTX2 | 4.211647252 | 2.330506085 | 0.864064963 | 2.69714221 | 0.00699374 | 0.037621635 |
| ARSB | 922.5941955 | 0.184805243 | 0.068555458 | 2.695704305 | 0.007023999 | 0.037762607 |
| CHST13 | 2.396510618 | -1.556734732 | 0.577485458 | -2.695712437 | 0.007023827 | 0.037762607 |
| CERS5 | 261.6505061 | 0.252434796 | 0.093659573 | 2.695237529 | 0.007033847 | 0.037793744 |
| PLCG1 | 112.3514541 | -0.38329479 | 0.142211789 | -2.695239202 | 0.007033812 | 0.037793744 |
| ZNF438 | 267.5182874 | 0.155820487 | 0.057822489 | 2.694807683 | 0.007042927 | 0.037831622 |
| SYNC | 57.25787058 | 0.549673907 | 0.204058291 | 2.693710238 | 0.007066157 | 0.037945464 |
| CD300LF | 1104.497555 | 0.352838088 | 0.131014496 | 2.693122504 | 0.007078625 | 0.037994333 |
| PLK1S1 | 54.66076044 | -0.353400499 | 0.131224957 | -2.693089095 | 0.007079335 | 0.037994333 |
| CAMLG | 379.8886597 | -0.234349554 | 0.087029834 | -2.692749642 | 0.007086546 | 0.038022084 |

| | | | | | | |
|---|---|---|---|---|---|---|
| CENPP | 65.8702047 | -0.328858492 | 0.122156059 | -2.692117719 | 0.007099988 | 0.03808324 |
| KLRB1 | 30.44407802 | -0.962451333 | 0.357558411 | -2.691731764 | 0.007108209 | 0.038116365 |
| C10orf12 | 371.3599204 | 0.252035934 | 0.093682038 | 2.690333588 | 0.007138063 | 0.038265437 |
| RBM22 | 430.5284301 | 0.17326106 | 0.064424163 | 2.689380095 | 0.007158486 | 0.038363884 |
| CSPP1 | 6.401843578 | -1.012783891 | 0.376694467 | -2.688608356 | 0.007175055 | 0.038441623 |
| TBRG4 | 368.2414028 | 0.180286757 | 0.067063583 | 2.688295907 | 0.007181772 | 0.038466555 |
| MDN1 | 164.517417 | -0.230783484 | 0.085874669 | -2.687445389 | 0.007200087 | 0.038553572 |
| DRAP1 | 1822.39264 | 0.298568407 | 0.111106503 | 2.687227103 | 0.007204795 | 0.038567695 |
| AHCY | 563.5751355 | -0.274927362 | 0.102325611 | -2.686789353 | 0.007214243 | 0.038607183 |
| TFAP2E | 30.42254011 | -0.328818722 | 0.122417381 | -2.686046037 | 0.007230313 | 0.038682069 |
| ENG | 4385.574813 | -0.183555661 | 0.068356722 | -2.68526129 | 0.007247312 | 0.038754901 |
| LIMD1-AS1 | 4.990186406 | -1.004746295 | 0.374175743 | -2.685225626 | 0.007248086 | 0.038754901 |
| PRR3 | 75.31296333 | 0.274975288 | 0.10242249 | 2.684715903 | 0.007259148 | 0.038802916 |
| DHX34 | 215.6097088 | 0.370462992 | 0.138060318 | 2.68334159 | 0.00728905 | 0.038951578 |
| RAB8A | 1963.399135 | 0.326237217 | 0.121588403 | 2.683127728 | 0.007293713 | 0.038965322 |
| GSTP1 | 7329.607052 | -0.289708427 | 0.108020287 | -2.681981643 | 0.007318748 | 0.03908786 |
| BRD2 | 1027.926504 | 0.173599541 | 0.064737363 | 2.681597359 | 0.007327159 | 0.039116269 |
| ERVK13-1 | 86.85302599 | -0.28957686 | 0.107992597 | -2.681451024 | 0.007330364 | 0.039116269 |
| TRERF1 | 118.1906013 | -0.311773459 | 0.116267718 | -2.681513525 | 0.007328995 | 0.039116269 |
| PTPN13 | 20.37222165 | -0.704104911 | 0.262649178 | -2.680780945 | 0.007345058 | 0.039183456 |
| NBPF14 | 217.0533598 | -0.23468451 | 0.087547056 | -2.680667075 | 0.007347558 | 0.039185572 |
| HIST1H1D | 160.0932259 | -0.406891398 | 0.151837686 | -2.679778697 | 0.007367085 | 0.039278236 |
| ME1 | 489.0926661 | -0.47792957 | 0.178352896 | -2.679684934 | 0.007369148 | 0.039278236 |
| ADAMTSL4 | 1147.146205 | 0.302689469 | 0.112979943 | 2.6791434 | 0.007381078 | 0.039329813 |
| IMPDH2 | 305.5799527 | -0.266393808 | 0.09943577 | -2.679054105 | 0.007383046 | 0.039329813 |
| HERC2P2 | 285.4158547 | -0.242176977 | 0.090426177 | -2.678173341 | 0.00740249 | 0.039422119 |
| LOC65365 | 21.48893425 | 0.682902257 | 0.25511497 | 2.67684118 | 0.007431985 | 0.039567891 |
| F11R | 4530.064976 | -0.126690648 | 0.047347035 | -2.675788442 | 0.007455369 | 0.039681047 |
| FAM98C | 95.41450427 | -0.352497304 | 0.131759658 | -2.675305243 | 0.007466124 | 0.039726942 |
| PDK2 | 226.5691331 | -0.24170581 | 0.09036895 | -2.674655527 | 0.007480607 | 0.039792644 |
| AGGF1 | 280.6582415 | -0.216669278 | 0.081035001 | -2.67377399 | 0.007500298 | 0.039886003 |
| LOXHD1 | 3.181727086 | 1.957492423 | 0.732203905 | 2.673425272 | 0.0075081 | 0.039916102 |
| C7orf49 | 105.787104 | -0.2853383 | 0.106737328 | -2.673275644 | 0.00751145 | 0.039922522 |
| TMEM158 | 98.04257101 | 0.517993333 | 0.193810997 | 2.672672555 | 0.007524966 | 0.039982954 |
| SLC4A1AP | 311.0228868 | 0.237569653 | 0.088892931 | 2.672537055 | 0.007528006 | 0.039987703 |
| FAM210A | 118.8202402 | 0.247863072 | 0.092751133 | 2.672345489 | 0.007532305 | 0.039987743 |
| HIST1H3D | 71.91977935 | 0.449905533 | 0.168355501 | 2.672354216 | 0.007532109 | 0.039987743 |
| POLH | 84.80708929 | -0.300053026 | 0.11228484 | -2.672248774 | 0.007534477 | 0.039987878 |
| SMCHD1 | 544.584341 | 0.255422223 | 0.095590018 | 2.672059583 | 0.007538726 | 0.039999038 |
| NAT9 | 141.135181 | -0.215251056 | 0.080564181 | -2.671796007 | 0.00754465 | 0.040019073 |
| KIAA0319 | 18.04445598 | 0.666496776 | 0.249508391 | 2.671239925 | 0.007557161 | 0.040074029 |
| TFDP2 | 88.27167123 | -0.326764746 | 0.12238993 | -2.669866279 | 0.007588146 | 0.040226888 |
| PLP2 | 740.1468983 | 0.191450483 | 0.071713498 | 2.66965758 | 0.007592864 | 0.040240449 |
| FBXO3 | 260.7245989 | -0.288728647 | 0.108203391 | -2.668388152 | 0.007621615 | 0.040369862 |
| MARVELD2 | 7.072040789 | 0.984484037 | 0.36893057 | 2.668480514 | 0.00761952 | 0.040369862 |
| CTAGE15P | 8.528578429 | 0.896430254 | 0.3360498 | 2.667551816 | 0.007640611 | 0.040458977 |
| FPR2 | 172.3932303 | 0.588095373 | 0.220585356 | 2.666067155 | 0.007674436 | 0.040615008 |

| | | | | | | |
|---|---|---|---|---|---|---|
| PARP11 | 180.4073078 | 0.377621951 | 0.141637553 | 2.666114632 | 0.007673352 | 0.040615008 |
| APOC1P1 | 3.57705668 | -1.251759229 | 0.469842777 | -2.664208729 | 0.007716966 | 0.040828492 |
| FBXO48 | 41.63671069 | 0.374709625 | 0.140682213 | 2.663518143 | 0.007732824 | 0.040900779 |
| CPNE9 | 8.572924084 | -0.854940673 | 0.321065128 | -2.662826323 | 0.00774874 | 0.04097333 |
| FAM168B | 664.155933 | -0.221818448 | 0.083324511 | -2.662103205 | 0.007765407 | 0.041049811 |
| LOC10012ł | 16.87842404 | -0.721264452 | 0.27097155 | -2.661771885 | 0.007773054 | 0.041078583 |
| C11orf74 | 43.80669697 | -0.306448583 | 0.115168016 | -2.660882704 | 0.00779361 | 0.041175541 |
| FOXD2 | 16.59956201 | 0.738486968 | 0.277652974 | 2.659748094 | 0.007819911 | 0.041302786 |
| MSL1 | 286.2807068 | -0.18019589 | 0.067779502 | -2.658560239 | 0.007847532 | 0.041436926 |
| ZNF383 | 77.43128824 | -0.386761155 | 0.145542209 | -2.657381383 | 0.00787503 | 0.041570341 |
| DDHD2 | 277.2486269 | -0.451670101 | 0.170056218 | -2.656004632 | 0.007907253 | 0.041727064 |
| MB21D1 | 299.8553508 | 0.476573859 | 0.179438219 | 2.655921695 | 0.007909198 | 0.041727064 |
| HYPK | 571.1720306 | 0.144623129 | 0.054464318 | 2.655373884 | 0.007922055 | 0.041783066 |
| HLA-DQA2 | 6.766583656 | 1.947854257 | 0.73367539 | 2.654926529 | 0.007932568 | 0.041826678 |
| EIF4EBP3 | 294.6063946 | -0.246512689 | 0.092863697 | -2.654564682 | 0.007941081 | 0.041847882 |
| PLXNA1 | 907.9232533 | 0.261240201 | 0.098410968 | 2.654584202 | 0.007940622 | 0.041847882 |
| CREB3 | 192.9124942 | 0.292964976 | 0.110370538 | 2.65437663 | 0.007945509 | 0.041859375 |
| KLHL24 | 251.0225773 | -0.259304271 | 0.097717878 | -2.653601114 | 0.007963791 | 0.041943831 |
| FUT8 | 150.8256449 | -0.239447079 | 0.09024334 | -2.653349025 | 0.007969741 | 0.041963312 |
| C2orf71 | 20.48701818 | 0.905314177 | 0.341298999 | 2.652554448 | 0.007988524 | 0.042050328 |
| LOC10050( | 2.683816478 | 2.087029169 | 0.787202764 | 2.651196444 | 0.008020718 | 0.042207865 |
| CD28 | 17.83112113 | 0.578985244 | 0.218413484 | 2.650867678 | 0.008028529 | 0.042237043 |
| DUSP23 | 667.8740679 | -0.264254458 | 0.099695308 | -2.650620801 | 0.008034399 | 0.042255995 |
| PGM5P2 | 43.83670344 | -0.461751181 | 0.174235312 | -2.650158434 | 0.008045403 | 0.042301931 |
| PMFBP1 | 2.128179875 | -1.715784028 | 0.647518063 | -2.649785583 | 0.008054287 | 0.042334252 |
| UQCRFS1 | 1901.74392 | 0.251932978 | 0.095079461 | 2.649709781 | 0.008056094 | 0.042334252 |
| DPP7 | 1159.273448 | -0.261630446 | 0.098755877 | -2.649264577 | 0.008066715 | 0.042378114 |
| TRAF4 | 53.8953267 | 0.454331554 | 0.171551196 | 2.648372993 | 0.008088023 | 0.04247808 |
| C3orf38 | 481.1169477 | 0.213599292 | 0.080667117 | 2.647910333 | 0.0080991 | 0.042512292 |
| CDK20 | 18.34661297 | -0.545676943 | 0.206072445 | -2.64798597 | 0.008097289 | 0.042512292 |
| ARSG | 4.365077031 | -1.241142106 | 0.468857545 | -2.647162486 | 0.008117034 | 0.042594428 |
| PRKACA | 474.610411 | -0.244430808 | 0.09236439 | -2.646374963 | 0.008135958 | 0.042681711 |
| SIPA1 | 599.7344575 | -0.185425334 | 0.070126017 | -2.644173209 | 0.008189074 | 0.04294827 |
| DOCK6 | 38.5896507 | 0.489119594 | 0.185009505 | 2.643753863 | 0.008199226 | 0.042989411 |
| LOC55064: | 545.2581528 | -0.315741479 | 0.119438502 | -2.643548559 | 0.0082042 | 0.043003391 |
| MDGA1 | 8.795341442 | 1.108210398 | 0.419252846 | 2.64329845 | 0.008210263 | 0.043023071 |
| ZNF140 | 213.4875652 | 0.164123806 | 0.062095312 | 2.643094972 | 0.008215199 | 0.043036833 |
| AKT3 | 1264.267959 | 0.219860333 | 0.083195688 | 2.642689023 | 0.008225054 | 0.043076351 |
| DEPDC7 | 13.35655418 | -0.52823511 | 0.199903856 | -2.642445825 | 0.008230963 | 0.043095186 |
| KHNYN | 565.0223068 | 0.164528619 | 0.062284107 | 2.641582718 | 0.008251965 | 0.043193012 |
| HIST1H2A( | 333.5193153 | -0.411086922 | 0.155677564 | -2.640630487 | 0.008275192 | 0.043302422 |
| CA5B | 29.02387546 | -0.42886306 | 0.162420136 | -2.640455 | 0.008279479 | 0.043312691 |
| GPR162 | 17.88769982 | -0.784195516 | 0.297020419 | -2.640207423 | 0.00828553 | 0.043332182 |
| NIPAL4 | 3.986216433 | 2.56063636 | 0.969905186 | 2.640089357 | 0.008288417 | 0.043335119 |
| COL6A1 | 424.9642878 | 0.345353175 | 0.130826327 | 2.639783466 | 0.008295901 | 0.043362083 |
| ZBTB33 | 269.3067265 | -0.27369947 | 0.103781664 | -2.637262302 | 0.008357818 | 0.043673465 |
| ATP6V1E1 | 3444.857296 | 0.136036018 | 0.051587391 | 2.637001309 | 0.008364251 | 0.043694828 |

| | | | | | | |
|---|---|---|---|---|---|---|
| TMA7 | 455.0117798 | -0.426826835 | 0.161927069 | -2.635920217 | 0.008390946 | 0.043821997 |
| MNS1 | 1.991222928 | -1.481691969 | 0.562180797 | -2.635614693 | 0.008398504 | 0.043849179 |
| LIPC | 19.86851992 | -0.69415073 | 0.263486396 | -2.634484139 | 0.008426524 | 0.043983151 |
| GNG12 | 563.7208997 | -0.156395466 | 0.059368553 | -2.634314942 | 0.008430725 | 0.043992754 |
| UTP20 | 85.00758212 | -0.255054301 | 0.096881034 | -2.632654609 | 0.008472045 | 0.044195995 |
| TNFRSF1B | 1621.065227 | 0.226884601 | 0.086232355 | 2.631084365 | 0.00851129 | 0.044388297 |
| CRAMP1L | 95.07040845 | 0.271549419 | 0.103219545 | 2.630794573 | 0.008518551 | 0.044413732 |
| KITLG | 12.17495279 | -0.63091368 | 0.240044821 | -2.628316146 | 0.008580873 | 0.044726148 |
| PFDN5 | 4264.888046 | -0.241530863 | 0.091904416 | -2.628065915 | 0.008587187 | 0.044746547 |
| MSI2 | 54.22094235 | -0.337774208 | 0.128545306 | -2.627666607 | 0.008597273 | 0.044786577 |
| SUV420H1 | 337.6871316 | -0.231669374 | 0.088178333 | -2.627282318 | 0.008606989 | 0.044824661 |
| TMEM184I | 1000.897207 | -0.291340821 | 0.11089917 | -2.627078449 | 0.008612148 | 0.044838994 |
| SRF | 486.3217199 | 0.122504007 | 0.046636438 | 2.626787404 | 0.008619517 | 0.044864826 |
| ATP6V1F | 143.3691889 | 0.216432008 | 0.082415613 | 2.626104454 | 0.008636831 | 0.044907692 |
| NT5E | 4.33534953 | 2.05381266 | 0.782082358 | 2.626082331 | 0.008637392 | 0.044907692 |
| SHFM1 | 1826.996127 | 0.24765207 | 0.094302777 | 2.626137628 | 0.008635989 | 0.044907692 |
| TWISTNB | 178.5331282 | 0.27042816 | 0.102971239 | 2.626249469 | 0.008633152 | 0.044907692 |
| USP27X | 56.57115288 | -0.366116899 | 0.139550549 | -2.623543231 | 0.00870204 | 0.04523119 |
| TPK1 | 416.7807434 | 0.246790576 | 0.094087329 | 2.622994817 | 0.00871606 | 0.045291428 |
| LANCL2 | 247.813769 | 0.219171309 | 0.08356244 | 2.622844783 | 0.008719899 | 0.045298744 |
| ZNF84 | 88.57387592 | -0.285322202 | 0.108806907 | -2.622280231 | 0.008734358 | 0.045361211 |
| KLHL5 | 358.7613339 | 0.182342431 | 0.069556194 | 2.621512492 | 0.008754055 | 0.045450841 |
| CLIC6 | 11.32993807 | 1.139844441 | 0.434846083 | 2.621259536 | 0.008760554 | 0.045471912 |
| GAMT | 5.438794186 | -1.149827027 | 0.438757354 | -2.620644454 | 0.008776374 | 0.04554134 |
| BLMH | 179.5436677 | -0.309545039 | 0.118125618 | -2.620473388 | 0.008780778 | 0.04555151 |
| VPS36 | 696.9925862 | -0.141957652 | 0.054181776 | -2.620025826 | 0.008792311 | 0.045598642 |
| C19orf73 | 11.81540671 | -0.774156842 | 0.29553157 | -2.619540243 | 0.008804838 | 0.045650906 |
| NDUFA9 | 1021.856706 | 0.272378178 | 0.104009901 | 2.618771646 | 0.0088247 | 0.045741157 |
| ELP2 | 111.4670873 | -0.41690915 | 0.159268883 | -2.617643465 | 0.008853926 | 0.045879883 |
| PCDHGB8P | 12.49544263 | -0.771143746 | 0.294652115 | -2.617132908 | 0.008867181 | 0.045935793 |
| PRKCA | 127.8256908 | -0.28334314 | 0.108325303 | -2.615669034 | 0.008905283 | 0.046120357 |
| FAM176C | 8.355916927 | -0.906611667 | 0.346663537 | -2.615249573 | 0.008916228 | 0.046164209 |
| SLX4 | 70.92050955 | 0.31409079 | 0.120185631 | 2.613380544 | 0.008965142 | 0.04640457 |
| RPA1 | 560.883477 | -0.200043348 | 0.076565335 | -2.612714334 | 0.008982635 | 0.046482204 |
| PDK3 | 54.30703823 | -0.640540636 | 0.245211135 | -2.612200446 | 0.008996149 | 0.046522042 |
| PRDX6 | 2871.58164 | 0.212618525 | 0.081393216 | 2.612238886 | 0.008995138 | 0.046522042 |
| PURB | 323.3538196 | 0.269492826 | 0.10316949 | 2.612136833 | 0.008997823 | 0.046522042 |
| DNPEP | 237.8019126 | 0.330483206 | 0.126544261 | 2.611601689 | 0.009011918 | 0.046581993 |
| NOTCH3 | 105.0992176 | 0.452685834 | 0.173343115 | 2.611501673 | 0.009014555 | 0.046582699 |
| SMG1P1 | 1571.584472 | -0.225356628 | 0.08633491 | -2.610260777 | 0.009047323 | 0.046739065 |
| KCTD2 | 252.7898364 | -0.188475115 | 0.072214542 | -2.609932985 | 0.009055996 | 0.046770907 |
| SPIN4 | 182.3369259 | 0.404840027 | 0.155129642 | 2.609688395 | 0.009062473 | 0.046791389 |
| MED8 | 365.523499 | 0.187005863 | 0.07166259 | 2.609532567 | 0.009066602 | 0.046799738 |
| HCST | 1029.21713 | 0.210411018 | 0.080641254 | 2.609223039 | 0.009074807 | 0.046803749 |
| SCGB1A1 | 13.7188764 | 1.366163496 | 0.523567791 | 2.609334491 | 0.009071852 | 0.046803749 |
| TMEM41B | 249.3695886 | 0.355720967 | 0.136333595 | 2.609195238 | 0.009075545 | 0.046803749 |
| ZNF569 | 54.78366871 | -0.347561495 | 0.133210017 | -2.609124323 | 0.009077426 | 0.046803749 |

| | | | | | | |
|---|---|---|---|---|---|---|
| PLEKHG4 | 12.05997841 | -0.738072277 | 0.282967779 | -2.608326218 | 0.00909862 | 0.04690005 |
| KCNS3 | 1.692789282 | 1.917863043 | 0.735383078 | 2.607978211 | 0.009107875 | 0.046934775 |
| PCDHGA12 | 16.41483521 | 0.668835101 | 0.256544693 | 2.607089988 | 0.009131536 | 0.047043693 |
| ASF1A | 298.3661736 | -0.255953317 | 0.098266941 | -2.604673694 | 0.00919618 | 0.047337461 |
| PARK2 | 8.934603848 | -0.875024992 | 0.33593406 | -2.604752231 | 0.009194072 | 0.047337461 |
| RPL23A | 274.789786 | 0.257558983 | 0.098882636 | 2.604693747 | 0.009195641 | 0.047337461 |
| SDC3 | 334.0353912 | -0.356057497 | 0.1367115 | -2.604444375 | 0.009202336 | 0.047356068 |
| CYBB | 13787.73318 | 0.272181479 | 0.104540042 | 2.603609817 | 0.009224771 | 0.047458416 |
| SUZ12P1 | 89.51473354 | -0.249517239 | 0.095864775 | -2.60280419 | 0.009246475 | 0.047556945 |
| NUCB2 | 119.0262535 | -0.27471767 | 0.105572468 | -2.602171518 | 0.009263551 | 0.047621073 |
| TJP2 | 107.6918991 | 0.300780681 | 0.115589166 | 2.602152879 | 0.009264054 | 0.047621073 |
| HMGN3 | 57.56803642 | -0.524476607 | 0.201599301 | -2.601579498 | 0.009279556 | 0.047680786 |
| PSENEN | 453.9558806 | -0.172792196 | 0.066419351 | -2.601533933 | 0.009280788 | 0.047680786 |
| MARK4 | 199.805678 | -0.218565421 | 0.0840325 | -2.600962965 | 0.009296249 | 0.047747051 |
| SRP68 | 397.1278744 | 0.167633251 | 0.064480969 | 2.599732189 | 0.009329654 | 0.047902192 |
| ZNF35 | 57.63645064 | -0.334732913 | 0.128760228 | -2.599660758 | 0.009331596 | 0.047902192 |
| AKTIP | 303.691323 | -0.246805442 | 0.094958986 | -2.599074121 | 0.009347559 | 0.047967405 |
| TMEM30A | 2465.219728 | -0.149547816 | 0.057540417 | -2.599004743 | 0.009349448 | 0.047967405 |
| ESYT1 | 3482.603226 | -0.179412885 | 0.0690388 | -2.598725423 | 0.009357059 | 0.047993237 |
| GTF2A2 | 668.2700267 | -0.160538269 | 0.061812082 | -2.597198885 | 0.009398749 | 0.048193804 |
| FOLR2 | 4.472509353 | -1.378687536 | 0.531036058 | -2.596222076 | 0.009425512 | 0.048317743 |
| RNASE2 | 4.728410205 | -1.286548705 | 0.495820657 | -2.594786415 | 0.009464972 | 0.048506678 |
| PMF1 | 342.7884326 | -0.164741932 | 0.063498154 | -2.594436561 | 0.00947461 | 0.048542721 |
| HS6ST1 | 166.5107956 | -0.201488767 | 0.077697145 | -2.593258307 | 0.009507134 | 0.048695967 |
| CPNE3 | 544.265898 | -0.280517972 | 0.108188252 | -2.592869067 | 0.0095179 | 0.048710944 |
| LOC15409: | 18.76252027 | 0.609252048 | 0.234962728 | 2.592973165 | 0.009515019 | 0.048710944 |
| SUPT3H | 43.70958661 | -0.324169732 | 0.125021399 | -2.592913971 | 0.009516657 | 0.048710944 |
| SIRT1 | 116.3634959 | -0.302954357 | 0.116848042 | -2.592720861 | 0.009522002 | 0.048718557 |
| MIB2 | 45.20613212 | -0.353724682 | 0.136435222 | -2.592619978 | 0.009524795 | 0.048719471 |
| SESN1 | 218.5632943 | -0.49501298 | 0.191081651 | -2.590583535 | 0.009581336 | 0.048995232 |
| KCNK6 | 30.27967546 | -0.729173915 | 0.281565353 | -2.589714634 | 0.009605552 | 0.049105586 |
| EDA | 8.953027126 | -1.013390928 | 0.391484942 | -2.58858214 | 0.009637196 | 0.049253842 |
| TYW5 | 23.33008403 | 0.507600675 | 0.196109962 | 2.588347225 | 0.009643772 | 0.049273934 |
| MIIP | 639.6656225 | 0.157796265 | 0.060966996 | 2.588224381 | 0.009647212 | 0.049277999 |
| LOH12CR1 | 262.5710835 | 0.187143452 | 0.072327443 | 2.587447372 | 0.009668996 | 0.049375741 |
| GPAM | 63.64249775 | -0.333984392 | 0.12908777 | -2.587265948 | 0.009674089 | 0.049388214 |
| CYTH2 | 270.696884 | 0.21174893 | 0.081857345 | 2.586804288 | 0.00968706 | 0.049439009 |
| TRMT6 | 173.8004601 | 0.23326331 | 0.090177151 | 2.586722987 | 0.009689345 | 0.049439009 |
| FAH | 123.3301882 | -0.324979127 | 0.125640319 | -2.586583117 | 0.009693279 | 0.04944554 |
| TRAPPC12 | 772.4940643 | -0.199701437 | 0.077243796 | -2.585339492 | 0.009728316 | 0.049610682 |
| ENGASE | 67.38876631 | -0.445914485 | 0.172487744 | -2.585195177 | 0.009732389 | 0.049617875 |
| PARP4 | 1520.948808 | -0.147090767 | 0.05691381 | -2.58444771 | 0.009753509 | 0.049711951 |
| PFKFB3 | 615.1946951 | -0.383001118 | 0.148252569 | -2.583436635 | 0.009782143 | 0.049844261 |
| ATP1B3 | 2869.933835 | 0.189528294 | 0.073366745 | 2.583299766 | 0.009786025 | 0.049850409 |
| UBR5 | 383.4489487 | -0.227122379 | 0.087931557 | -2.582945049 | 0.009796092 | 0.049888053 |
| LHPP | 334.9664673 | -0.274444495 | 0.106284296 | -2.582173532 | 0.00981802 | 0.049958763 |
| MOGAT1 | 25.06014812 | -0.504108725 | 0.195212352 | -2.58236079 | 0.009812694 | 0.049958763 |

| | | | | | | |
|---|---|---|---|---|---|---|
| ZNF174 | 128.832971 | -0.263871624 | 0.102189267 | -2.58218531 | 0.009817685 | 0.049958763 |
