## Supplemental Table 5 for "Human alveolar macrophage response to *Mycobacterium tuberculosis*: immune characteristics underlying large inter-individual variability"

**Supplementary File 5: A. Variably expressed genes (VE genes)**

| Symbol | Average RPM across all samples | DE Expr Log2 Ratio | DE Expr p-value | Time point (hrs) | FDR Levene | tau ratio | FDR entropy | Cor to generation time - 2hr | Cor to generation time - 24hr | Cor to generation time - 72hr |
|---|---|---|---|---|---|---|---|---|---|---|
| IFI6 | 1,165.2 | 7.1 | 1.73E-10 | 72 | 0.03 | 0.88 | 0.00 | -0.14 | 0.16 | 0.27 |
| IL1B | 780.3 | 6.9 | 7.26E-12 | 24 | 0.00 | 1.57 | 0.00 | -0.20 | -0.15 | -0.11 |
| TNFAIP6 | 611.9 | 4.8 | 1E-10 | 72 | 0.11 | 0.91 | 0.00 | -0.07 | 0.08 | 0.03 |
| TYMP | 552.2 | 6.6 | 0.00022 | 72 | 0.05 | 0.86 | 0.00 | 0.14 | 0.30 | 0.26 |
| CCL4 | 474.2 | 4.9 | 1.16E-10 | 72 | 0.02 | 1.33 | 0.00 | -0.19 | 0.15 | 0.01 |
| LAP3 | 462.5 | 6.3 | 1.13E-10 | 72 | 0.00 | 0.55 | 0.00 | -0.07 | 0.17 | -0.02 |
| CCL20 | 380.7 | 2.8 | 3.73E-11 | 72 | 0.02 | 1.80 | 0.00 | -0.36 | -0.06 | -0.07 |
| CCL3 | 355.0 | 6.0 | 7.74E-12 | 24 | 0.01 | 1.41 | 0.00 | -0.11 | 0.08 | -0.08 |
| IL7R | 333.3 | 4.2 | 1.4E-09 | 72 | 0.04 | 1.01 | 0.00 | -0.06 | -0.03 | -0.08 |
| SLAMF7 | 309.5 | 5.6 | 2.37E-08 | 72 | 0.00 | 0.95 | 0.00 | -0.06 | 0.05 | -0.05 |
| MX1 | 307.5 | 5.8 | 5.64E-11 | 72 | 0.00 | 0.69 | 0.00 | | | |
| GBP1 | 298.7 | 5.5 | 4.12E-11 | 72 | 0.00 | 0.40 | 0.00 | -0.22 | -0.03 | -0.09 |
| TAP1 | 194.1 | 5.3 | 4.39E-09 | 72 | 0.00 | 0.56 | 0.00 | -0.17 | -0.03 | 0.05 |
| PARP14 | 158.4 | 5.3 | 1.07E-11 | 72 | 0.00 | 0.67 | 0.00 | 0.03 | -0.05 | 0.17 |
| NPL | 151.5 | 5.2 | 2.78E-06 | 72 | 0.00 | 1.51 | 0.00 | -0.11 | -0.25 | -0.26 |
| STAT2 | 140.4 | 5.2 | 3.4E-09 | 72 | 0.00 | 0.55 | 0.00 | -0.34 | -0.03 | -0.08 |
| ISG15 | 136.3 | 4.8 | 4.69E-11 | 72 | 0.00 | 0.65 | 0.00 | -0.29 | 0.20 | 0.28 |
| PSMB9 | 134.1 | 4.9 | 9.61E-08 | 72 | 0.00 | 0.56 | 0.00 | -0.24 | 0.00 | -0.07 |
| CD274 | 118.9 | 4.9 | 6.47E-10 | 24 | 0.00 | 0.91 | 0.00 | 0.05 | 0.11 | 0.09 |
| OAS3 | 118.4 | 4.8 | 5.71E-11 | 72 | 0.00 | 0.65 | 0.00 | -0.01 | 0.13 | 0.36 |
| RIPK2 | 110.2 | 4.5 | 1.62E-07 | 72 | 0.01 | 0.89 | 0.00 | -0.06 | 0.08 | 0.05 |
| IFI35 | 106.3 | 4.7 | 6.35E-08 | 72 | 0.00 | 0.58 | 0.00 | -0.20 | -0.05 | -0.05 |
| APOL1 | 106.0 | 4.5 | 8.44E-08 | 72 | 0.05 | 0.68 | 0.00 | -0.18 | -0.07 | -0.05 |
| ZNFX1 | 98.4 | 4.6 | 2.04E-09 | 72 | 0.04 | 0.87 | 0.00 | | | |
| GBP5 | 93.2 | 3.9 | 1.55E-07 | 72 | 0.07 | 0.92 | 0.00 | -0.12 | 0.03 | 0.02 |
| IDO1 | 89.4 | 1.9 | 1.46E-08 | 72 | 0.14 | 1.83 | 0.00 | | -0.06 | -0.12 |
| IFITM3 | 84.0 | 4.4 | 8.52E-10 | 72 | 0.00 | 0.78 | 0.00 | -0.05 | 0.18 | 0.28 |
| VDR | 81.1 | 4.4 | 3.04E-08 | 24 | 0.08 | 1.09 | 0.00 | 0.36 | -0.05 | 0.23 |
| DYNLT1 | 79.3 | 4.6 | 6.38E-06 | 72 | 0.00 | 0.70 | 0.00 | -0.24 | 0.30 | 0.21 |
| C1RL | 78.0 | 4.5 | 0.0000065 | 72 | 0.03 | 0.88 | 0.00 | 0.20 | 0.30 | 0.24 |
| IFITM1 | 75.7 | 4.3 | 8.5E-10 | 72 | 0.00 | 0.79 | 0.00 | -0.03 | 0.15 | 0.29 |
| EPSTI1 | 55.5 | 4.0 | 2.41E-06 | 72 | 0.00 | 0.81 | 0.00 | -0.30 | -0.15 | -0.24 |
| IFIH1 | 45.2 | 4.0 | 2.61E-08 | 72 | 0.00 | 0.73 | 0.00 | -0.39 | -0.25 | -0.22 |
| SAMD9L | 43.7 | 3.8 | 3.5E-10 | 72 | 0.00 | 0.80 | 0.00 | -0.35 | -0.19 | -0.23 |
| DDX60 | 38.6 | 3.8 | 1.3E-11 | 72 | 0.00 | 0.62 | 0.00 | | | |
| TAGAP | 34.1 | 3.7 | 4.93E-08 | 72 | 0.00 | 0.62 | 0.00 | -0.21 | -0.01 | -0.05 |
| DCUN1D3 | 33.5 | 3.6 | 2.42E-12 | 24 | 0.00 | 1.03 | 0.00 | -0.25 | -0.33 | -0.25 |
| CXCL1 | 31.5 | 3.4 | 2.59E-11 | 24 | 0.00 | 1.44 | 0.00 | | | |
| PARP9 | 29.5 | 3.5 | 1.21E-11 | 72 | 0.00 | 0.66 | 0.00 | 0.00 | 0.24 | 0.38 |
| HSD11B1 | 29.3 | 3.4 | 7.91E-11 | 24 | 0.00 | 1.49 | 0.00 | | 0.33 | 0.24 |
| GCH1 | 26.6 | 2.4 | 5.84E-11 | 72 | 0.02 | 0.89 | 0.00 | 0.14 | 0.15 | 0.04 |
| IFIT3 | 26.3 | 3.1 | 2.81E-11 | 72 | 0.00 | 0.99 | 0.00 | -0.49 | 0.02 | -0.10 |
| PML | 26.0 | 3.4 | 1.92E-09 | 72 | 0.00 | 0.55 | 0.00 | -0.18 | -0.09 | 0.18 |
| WDR91 | 23.6 | 3.3 | 3.77E-08 | 72 | 0.00 | 1.31 | 0.00 | 0.03 | 0.03 | 0.06 |
| IFIT5 | 21.5 | 3.2 | 8.5E-09 | 72 | 0.00 | 0.61 | 0.00 | -0.26 | 0.11 | 0.25 |
| DHX58 | 20.7 | 3.1 | 1.93E-10 | 72 | 0.01 | 0.61 | 0.00 | | | |
| FBXO6 | 19.4 | 3.1 | 1.94E-08 | 72 | 0.00 | 0.74 | 0.00 | | | |
| F3 | 18.9 | 2.5 | 2.02E-08 | 24 | 0.03 | 1.49 | 0.00 | -0.05 | 0.07 | -0.04 |
| CYP27B1 | 18.3 | 2.6 | 3.77E-10 | 24 | 0.00 | 1.60 | 0.00 | | -0.23 | -0.16 |
| SCARF1 | 17.5 | 3.0 | 2.06E-12 | 24 | 0.00 | 0.71 | 0.00 | 0.04 | -0.18 | 0.02 |

| | | | | | | | | | | |
|---|---|---|---|---|---|---|---|---|---|---|
| RTP4 | 15.9 | 3.0 | 6.14E-09 | 72 | 0.00 | 0.60 | 0.00 | -0.39 | 0.21 | 0.32 |
| HERC6 | 12.5 | 2.7 | 9.76E-11 | 72 | 0.00 | 0.72 | 0.00 | -0.15 | 0.18 | 0.20 |
| PDLIM4 | 10.8 | 2.8 | 1.13E-08 | 72 | 0.01 | 1.12 | 0.00 | | 0.26 | 0.17 |
| HELZ2 | 10.1 | 2.2 | 1.79E-08 | 72 | 0.01 | 0.77 | 0.00 | -0.23 | -0.12 | -0.10 |
| APBA1 | 8.0 | 2.3 | 3.52E-08 | 72 | 0.02 | 1.29 | 0.00 | 0.10 | 0.21 | -0.09 |
| ISG20 | 7.9 | 1.1 | 5.91E-11 | 72 | 0.01 | 0.53 | 0.00 | | -0.14 | 0.06 |
| CCL8 | 4.7 | 0.4 | 2.85E-08 | 72 | 0.01 | 1.01 | 0.00 | | | -0.23 |
| CHMP5 | 223.2 | 5.5 | 3.96E-09 | 72 | 0.01 | 0.75 | 0.00 | -0.06 | 0.34 | 0.27 |
| DDX58 | 117.1 | 4.6 | 1.12E-10 | 72 | 0.12 | 1.11 | 0.00 | | | |
| EMP1 | 106.8 | 5.0 | 1.5E-10 | 24 | 0.00 | 0.91 | 0.00 | 0.13 | -0.16 | 0.00 |
| G0S2 | 106.1 | 3.7 | 8.36E-08 | 72 | 0.04 | 1.09 | 0.00 | -0.04 | 0.16 | 0.12 |
| DTX3L | 102.2 | 4.8 | 1.67E-08 | 72 | 0.00 | 0.63 | 0.00 | | | |
| OAS2 | 90.3 | 4.6 | 4.32E-07 | 72 | 0.00 | 0.76 | 0.00 | -0.27 | -0.16 | -0.11 |
| KCNN4 | 86.8 | 4.8 | 4.27E-08 | 24 | 0.00 | 1.18 | 0.00 | -0.26 | -0.09 | -0.02 |
| TLR2 | 62.9 | 4.6 | 7.93E-11 | 24 | 0.01 | 1.12 | 0.00 | 0.09 | -0.14 | 0.00 |
| CMPK2 | 57.5 | 3.9 | 2.63E-09 | 72 | 0.00 | 0.82 | 0.00 | -0.25 | -0.19 | -0.04 |
| EYA3 | 56.9 | 4.1 | 8.54E-08 | 24 | 0.15 | 0.77 | 0.00 | 0.20 | 0.19 | 0.34 |
| HERC5 | 49.0 | 4.1 | 3.99E-10 | 72 | 0.00 | 0.62 | 0.00 | -0.20 | 0.20 | 0.19 |
| N4BP1 | 40.3 | 3.9 | 1.85E-08 | 24 | 0.02 | 0.91 | 0.00 | -0.26 | -0.30 | -0.14 |
| IL15RA | 36.1 | 3.7 | 3.61E-09 | 72 | 0.00 | 0.60 | 0.00 | -0.46 | -0.14 | -0.04 |
| NLRC5 | 29.5 | 3.5 | 2.93E-08 | 72 | 0.00 | 0.71 | 0.00 | -0.12 | -0.12 | 0.01 |
| EEPD1 | 12.6 | 2.5 | 6.53E-09 | 24 | 0.00 | 1.00 | 0.00 | 0.13 | -0.10 | -0.10 |
| CSF1 | 1,031.9 | 7.3 | 4.21E-11 | 24 | 0.00 | 1.00 | 0.00 | 0.08 | -0.14 | -0.22 |
| ALAS1 | 191.1 | 5.3 | 4.41E-06 | 72 | 0.00 | 0.75 | 0.00 | -0.17 | -0.29 | -0.18 |
| SLC30A1 | 172.6 | 5.2 | 0.000012 | 72 | 0.03 | 0.80 | 0.00 | 0.35 | 0.22 | 0.30 |
| PARP12 | 82.1 | 4.6 | 2.45E-08 | 72 | 0.00 | 0.71 | 0.00 | -0.27 | -0.19 | -0.29 |
| CERK | 51.9 | 4.2 | 8.24E-10 | 72 | 0.02 | 1.38 | 0.00 | -0.20 | -0.04 | -0.01 |
| PPBP | 25.4 | 3.5 | 3.02E-11 | 24 | 0.00 | 1.16 | 0.00 | 0.05 | -0.33 | -0.03 |
| USP18 | 17.4 | 2.9 | 9.73E-10 | 72 | 0.00 | 0.84 | 0.00 | -0.27 | 0.12 | 0.15 |
| STAT1 | 491.6 | 6.4 | 5.64E-09 | 72 | 0.00 | 0.73 | 0.00 | -0.35 | -0.22 | -0.33 |
| PIK3AP1 | 451.0 | 6.3 | 2.98E-07 | 72 | 0.01 | 0.89 | 0.00 | -0.15 | 0.09 | -0.05 |
| PLSCR1 | 197.5 | 5.3 | 1.37E-10 | 72 | 0.01 | 0.87 | 0.00 | -0.30 | -0.03 | -0.02 |
| TAP2 | 93.1 | 4.5 | 1.34E-09 | 72 | 0.00 | 0.63 | 0.00 | -0.25 | -0.10 | 0.04 |
| SEMA4D | 74.2 | 4.2 | 1.48E-08 | 72 | 0.00 | 0.68 | 0.00 | -0.20 | -0.01 | -0.06 |
| SPRED1 | 65.4 | 4.4 | 1.55E-11 | 24 | 0.00 | 0.80 | 0.00 | 0.16 | -0.13 | -0.14 |
| IFIT1 | 41.4 | 3.6 | 2.68E-10 | 72 | 0.00 | 0.80 | 0.00 | -0.39 | 0.07 | 0.07 |
| CA12 | 8.1 | 0.8 | 3.96E-09 | 72 | 0.00 | 1.37 | 0.00 | | -0.39 | -0.23 |
| CHST3 | 5.3 | 1.3 | 1.84E-07 | 72 | 0.01 | 1.19 | 0.00 | | -0.06 | 0.00 |
| NBN | 98.9 | 4.5 | 6.69E-08 | 72 | 0.01 | 0.87 | 0.00 | -0.31 | -0.16 | -0.02 |
| CHPT1 | 52.6 | 4.0 | 1.27E-06 | 72 | 0.00 | 1.18 | 0.00 | -0.17 | -0.16 | -0.23 |
| HIP1 | 40.8 | 3.6 | 5.24E-09 | 72 | 0.00 | 0.78 | 0.00 | -0.08 | 0.05 | 0.21 |
| C19orf66 | 28.5 | 3.4 | 1.73E-06 | 72 | 0.07 | 0.83 | 0.00 | -0.28 | 0.27 | 0.25 |
| STOM | 437.1 | 6.2 | 8.08E-06 | 72 | 0.02 | 0.73 | 0.00 | -0.01 | -0.13 | -0.15 |
| SORL1 | 26.9 | 2.1 | 1.27E-10 | 24 | 0.06 | 0.98 | 0.00 | -0.06 | -0.06 | 0.06 |
| YPEL3 | 22.1 | 2.9 | 3.02E-11 | 24 | 0.11 | 1.04 | 0.00 | 0.00 | 0.09 | 0.07 |
| GPR84 | 19.1 | 3.0 | 1.47E-07 | 24 | 0.00 | 0.97 | 0.00 | -0.28 | -0.23 | -0.14 |
| SLAMF8 | 383.1 | 6.2 | 8.28E-08 | 24 | 0.00 | 0.86 | 0.00 | 0.07 | 0.02 | -0.03 |
| TMEM86A | 39.6 | 4.0 | 7.35E-10 | 72 | 0.01 | 1.66 | 0.00 | -0.15 | -0.01 | -0.11 |
| IL1A | 156.1 | 4.8 | 4.39E-10 | 24 | 0.06 | 1.32 | 0.00 | -0.01 | -0.08 | -0.05 |
| TPST2 | 45.1 | 3.9 | 0.00041 | 72 | 0.15 | 0.99 | 0.00 | -0.21 | -0.04 | -0.18 |
| MB21D2 | 11.6 | 2.8 | 6.56E-09 | 24 | 0.00 | 0.77 | 0.00 | -0.07 | -0.07 | -0.03 |
| WARS | 852.5 | 7.0 | 0.000017 | 72 | 0.00 | 0.69 | 0.00 | 0.03 | 0.06 | -0.05 |
| DAPK1 | 327.0 | 6.0 | 1.47E-07 | 24 | 0.03 | 1.15 | 0.00 | 0.15 | 0.11 | 0.40 |
| GBP4 | 64.7 | 4.1 | 9.76E-11 | 72 | 0.03 | 0.41 | 0.00 | -0.13 | 0.00 | 0.02 |
| CFP | 58.8 | 4.1 | 2.68E-10 | 72 | 0.00 | 1.47 | 0.00 | 0.07 | 0.22 | 0.31 |

| | | | | | | | | | | |
|---|---|---|---|---|---|---|---|---|---|---|
| PSTPIP2 | 43.5 | 3.1 | 9.86E-07 | 72 | 0.00 | 0.94 | 0.00 | -0.04 | 0.08 | 0.03 |
| IFI44L | 30.5 | 2.7 | 1.96E-10 | 72 | 0.00 | 1.23 | 0.00 | -0.27 | -0.14 | -0.17 |
| LIMK2 | 26.9 | 3.4 | 7.88E-07 | 24 | 0.07 | 0.87 | 0.00 | -0.30 | -0.24 | -0.14 |
| VWA8 | 96.5 | 4.9 | 3.09E-09 | 24 | 0.01 | 1.04 | 0.00 | -0.20 | -0.13 | -0.14 |
| CCL5 | 58.8 | 3.4 | 3.81E-09 | 72 | 0.14 | 0.55 | 0.00 | -0.30 | -0.13 | -0.07 |
| TNFSF10 | 16.8 | 2.5 | 1.58E-09 | 72 | 0.06 | 1.13 | 0.00 | -0.32 | 0.12 | 0.04 |
| FAM110A | 10.0 | 2.4 | 0.00116 | 24 | 0.01 | 1.26 | 0.00 | -0.25 | -0.36 | -0.10 |
| MIR155HG | 9.4 | 2.6 | 7.79E-09 | 24 | 0.02 | 0.94 | 0.00 | -0.08 | -0.29 | |
| PVALB | 17.0 | 3.1 | 6.71E-10 | 72 | 0.03 | 1.39 | 0.00 | -0.43 | 0.08 | -0.13 |
| SLC27A1 | 23.6 | 3.6 | 4.69E-11 | 72 | 0.00 | 1.45 | 0.00 | 0.03 | -0.33 | -0.13 |
| LSM6 | 15.5 | 2.7 | 0.00531 | 24 | 0.09 | 1.22 | 0.00 | -0.05 | 0.07 | -0.03 |
| RUNX1 | 13.4 | 2.6 | 2.55E-09 | 24 | 0.02 | 0.99 | 0.00 | | | |
| TMEM150A | 23.2 | 3.2 | 0.0017 | 72 | 0.15 | 1.25 | 0.00 | 0.01 | 0.31 | 0.24 |
| FUCA1 | 141.9 | 4.0 | 4.27E-08 | 24 | 0.05 | 1.50 | 0.00 | -0.31 | -0.20 | -0.14 |
| RASGRP1 | 13.0 | 2.1 | 7.05E-08 | 72 | 0.00 | 1.12 | 0.00 | -0.13 | 0.03 | 0.05 |
| SLC46A1 | 10.4 | 2.6 | 4.12E-11 | 72 | 0.00 | 1.34 | 0.00 | -0.15 | | 0.02 |
| CD80 | 48.6 | 3.7 | 3.1E-08 | 72 | 0.00 | 0.99 | 0.00 | -0.04 | 0.01 | 0.02 |
| RAB42 | 41.9 | 4.1 | 5.39E-08 | 72 | 0.00 | 1.08 | 0.00 | -0.14 | 0.24 | 0.06 |
| CASS4 | 11.5 | 2.7 | 1.19E-09 | 24 | 0.00 | 0.92 | 0.00 | | -0.43 | -0.12 |
| SP110 | 9.4 | 2.4 | 2.57E-10 | 72 | 0.01 | 0.70 | 0.00 | -0.15 | 0.14 | 0.38 |
| MT2A | 899.2 | 6.3 | 1.93E-10 | 72 | 0.06 | 0.85 | 0.00 | -0.36 | -0.32 | -0.33 |
| XAF1 | 27.6 | 3.0 | 2.51E-09 | 72 | 0.03 | 1.23 | 0.00 | | | |
| TRAFD1 | 86.2 | 4.4 | 6.99E-08 | 72 | 0.10 | 0.88 | 0.00 | -0.23 | -0.01 | -0.09 |
| UBD | 62.7 | 2.5 | 3.08E-06 | 24 | 0.14 | 0.95 | 0.00 | 0.00 | -0.12 | -0.20 |
| GMPR | 12.1 | 2.9 | 7.96E-08 | 72 | 0.01 | 0.62 | 0.00 | 0.03 | 0.20 | 0.21 |
| EIF2AK2 | 118.4 | 4.9 | 3.73E-11 | 72 | 0.01 | 0.78 | 0.00 | -0.02 | 0.25 | 0.31 |
| PIM1 | 269.7 | 5.6 | 2.61E-08 | 24 | 0.03 | 0.72 | 0.00 | 0.06 | 0.10 | 0.20 |
| EPHX1 | 185.1 | 5.4 | 8.87E-06 | 72 | 0.00 | 1.15 | 0.00 | | | |
| AGRN | 29.9 | 3.6 | 2.17E-08 | 72 | 0.00 | 0.67 | 0.00 | 0.05 | 0.23 | 0.45 |
| NEK6 | 10.3 | 2.5 | 7.43E-06 | 24 | 0.05 | 0.78 | 0.00 | 0.16 | -0.04 | 0.01 |
| CCL24 | 283.2 | 5.9 | 6.27E-09 | 24 | 0.00 | 0.98 | 0.00 | 0.14 | -0.15 | -0.03 |
| IRF1 | 111.5 | 4.5 | 1.47E-06 | 72 | 0.11 | 0.45 | 0.00 | -0.28 | -0.13 | -0.05 |
| LDLRAP1 | 32.4 | 3.5 | 1.85E-08 | 24 | 0.00 | 0.90 | 0.00 | -0.01 | -0.04 | -0.11 |
| MICAL2 | 15.8 | 2.9 | 1.56E-07 | 24 | 0.00 | 0.88 | 0.00 | -0.41 | -0.69 | -0.47 |
| CFB | 79.0 | 4.1 | 1.22E-09 | 72 | 0.01 | 0.96 | 0.00 | 0.14 | 0.08 | 0.13 |
| NUB1 | 102.0 | 4.7 | 1.01E-07 | 72 | 0.01 | 0.58 | 0.00 | -0.06 | 0.13 | 0.07 |
| BATF3 | 7.3 | 1.8 | 1.59E-06 | 72 | 0.16 | 1.05 | 0.00 | -0.04 | 0.26 | 0.12 |
| RUSC2 | 29.8 | 3.3 | 0.000102 | 72 | 0.06 | 0.84 | 0.00 | 0.27 | -0.06 | 0.21 |
| ACSF2 | 33.5 | 3.8 | 8.72E-08 | 72 | 0.08 | 1.19 | 0.00 | -0.06 | 0.14 | -0.04 |
| MX2 | 27.5 | 3.3 | 2.62E-06 | 72 | 0.02 | 0.72 | 0.00 | | | |
| PGPEP1 | 45.5 | 4.2 | 2.23E-07 | 72 | 0.02 | 1.25 | 0.00 | 0.00 | 0.16 | 0.11 |
| APOBEC3G | 26.5 | 3.4 | 6.73E-08 | 72 | 0.00 | 0.54 | 0.00 | -0.31 | 0.01 | -0.18 |
| PBXIP1 | 21.0 | 3.0 | 0.000002 | 72 | 0.15 | 1.09 | 0.00 | | | |
| PTPRF | 6.5 | 2.2 | 2.61E-08 | 24 | 0.01 | 1.17 | 0.00 | | -0.35 | -0.39 |
| CD40 | 138.8 | 4.7 | 1.79E-08 | 72 | 0.00 | 0.81 | 0.00 | -0.15 | -0.06 | -0.10 |
| PSME1 | 108.7 | 4.8 | 0.0000834 | 72 | 0.01 | 0.72 | 0.00 | -0.48 | -0.03 | -0.12 |
| ITGB3 | 34.2 | 3.9 | 2.93E-08 | 24 | 0.00 | 1.25 | 0.00 | | -0.01 | 0.12 |
| TBXAS1 | 82.9 | 4.6 | 1.99E-10 | 72 | 0.01 | 1.57 | 0.00 | | | |
| XRN1 | 16.3 | 2.7 | 0.000454 | 24 | 0.01 | 0.84 | 0.00 | | | |
| NUP85 | 15.0 | 2.9 | 0.0000468 | 72 | 0.00 | 1.30 | 0.00 | -0.38 | -0.50 | -0.23 |
| FARP1 | 51.6 | 4.2 | 5.14E-08 | 72 | 0.00 | 1.28 | 0.00 | 0.06 | -0.02 | -0.01 |
| ASB6 | 18.9 | 3.0 | 0.0000737 | 72 | 0.13 | 0.77 | 0.00 | 0.09 | 0.07 | 0.17 |
| SLC46A3 | 38.0 | 3.7 | 0.000634 | 72 | 0.04 | 1.10 | 0.00 | -0.22 | -0.21 | -0.15 |
| OSM | 37.4 | 3.4 | 6.3E-07 | 72 | 0.11 | 1.05 | 0.00 | -0.07 | 0.03 | 0.00 |

| | | | | | | | | | | |
|---|---|---|---|---|---|---|---|---|---|---|
| MTF1 | 65.7 | 3.9 | 3.65E-09 | 72 | 0.06 | 1.05 | 0.00 | 0.01 | -0.06 | -0.04 |
| CXCL5 | 1,254.9 | 7.7 | 3.34E-09 | 24 | 0.02 | 1.12 | 0.00 | | | |
| BCS1L | 9.9 | 2.2 | 0.000455 | 24 | 0.01 | 1.35 | 0.00 | -0.04 | -0.01 | 0.11 |
| PARP10 | 66.7 | 4.3 | 3.1E-07 | 72 | 0.01 | 0.81 | 0.00 | -0.03 | 0.02 | 0.37 |
| CD180 | 6.3 | 1.7 | 1.04E-09 | 72 | 0.05 | 1.57 | 0.00 | -0.51 | | -0.37 |
| FAM129B | 197.1 | 5.7 | 2.13E-08 | 24 | 0.02 | 0.95 | 0.00 | 0.15 | -0.09 | 0.15 |
| APBB2 | 32.3 | 3.5 | 7.87E-10 | 24 | 0.07 | 0.88 | 0.00 | 0.00 | 0.05 | 0.04 |
| DRAM2 | 62.0 | 4.3 | 0.0000149 | 72 | 0.06 | 1.31 | 0.00 | -0.19 | -0.06 | -0.06 |
| IFI44 | 57.9 | 4.1 | 6.81E-07 | 72 | 0.01 | 0.88 | 0.00 | -0.34 | -0.07 | -0.09 |
| PNPLA1 | 4.8 | 0.0 | 7.32E-07 | 72 | 0.15 | 1.07 | 0.00 | -0.18 | -0.37 | |
| TBC1D8 | 30.9 | 3.4 | 0.0000136 | 24 | 0.06 | 1.11 | 0.00 | 0.22 | 0.07 | 0.18 |
| KIAA0040 | 7.9 | 2.2 | 4.27E-08 | 72 | 0.07 | 0.69 | 0.00 | -0.24 | -0.15 | -0.05 |
| H2AFY | 234.0 | 5.6 | 0.0013 | 72 | 0.06 | 1.26 | 0.00 | -0.28 | -0.29 | -0.30 |
| CLEC2D | 10.8 | 2.1 | 3.06E-06 | 72 | 0.01 | 0.97 | 0.00 | -0.28 | 0.33 | 0.17 |
| IGSF6 | 201.4 | 5.4 | 3.32E-08 | 24 | 0.16 | 1.10 | 0.00 | -0.35 | -0.32 | -0.43 |
| BAK1 | 63.6 | 4.2 | 0.0000566 | 24 | 0.03 | 0.74 | 0.00 | 0.22 | 0.09 | 0.23 |
| VAMP5 | 10.8 | 2.0 | 4.65E-06 | 72 | 0.06 | 0.48 | 0.00 | -0.21 | -0.15 | -0.19 |
| LAMB3 | 6.0 | 2.1 | 2.13E-08 | 24 | 0.06 | 0.86 | 0.00 | | 0.04 | 0.10 |
| PSME2 | 115.0 | 4.9 | 1.72E-08 | 72 | 0.00 | 0.58 | 0.00 | -0.32 | -0.06 | -0.21 |
| ELMO2 | 43.5 | 4.0 | 0.0000164 | 24 | 0.04 | 0.86 | 0.00 | -0.18 | -0.07 | -0.02 |
| TREM2 | 160.4 | 5.2 | 4.36E-09 | 72 | 0.14 | 0.95 | 0.00 | 0.01 | 0.13 | 0.07 |
| CASC4 | 49.5 | 4.1 | 0.0000151 | 72 | 0.05 | 1.19 | 0.00 | 0.04 | 0.26 | 0.12 |
| ARHGAP10 | 64.4 | 4.4 | 4.59E-08 | 24 | 0.07 | 0.76 | 0.00 | -0.18 | -0.42 | -0.19 |
| TOR1B | 83.8 | 4.5 | 6.92E-06 | 72 | 0.05 | 0.78 | 0.00 | -0.16 | 0.21 | 0.07 |
| GBP2 | 462.3 | 5.6 | 1.46E-06 | 72 | 0.08 | 0.78 | 0.00 | 0.14 | -0.09 | 0.01 |
| CASP8 | 10.2 | 2.3 | 0.0000143 | 72 | 0.06 | 1.26 | 0.00 | -0.09 | 0.19 | 0.01 |
| RNF213 | 167.9 | 5.3 | 8.5E-10 | 72 | 0.00 | 0.74 | 0.00 | -0.09 | -0.17 | -0.12 |
| DOCK4 | 67.4 | 4.2 | 2.92E-07 | 72 | 0.04 | 0.95 | 0.00 | 0.11 | -0.18 | 0.05 |
| CBX6 | 8.6 | 2.2 | 2.93E-07 | 24 | 0.01 | 0.80 | 0.00 | | | |
| ERCC2 | 9.4 | 2.3 | 4.96E-07 | 72 | 0.06 | 1.25 | 0.00 | -0.20 | -0.07 | -0.11 |
| OAS1 | 172.4 | 5.4 | 1.47E-06 | 72 | 0.03 | 0.72 | 0.00 | -0.23 | -0.03 | -0.02 |
| SPOCD1 | 98.4 | 4.9 | 0.0000285 | 24 | 0.01 | 0.86 | 0.00 | 0.13 | 0.06 | -0.01 |
| AMDHD2 | 31.6 | 3.5 | 2.76E-09 | 24 | 0.03 | 1.09 | 0.00 | 0.30 | 0.37 | 0.24 |
| TMEM223 | 11.4 | 2.2 | 0.0000137 | 24 | 0.15 | 1.19 | 0.00 | -0.37 | 0.13 | 0.07 |
| HSD3B7 | 184.4 | 5.4 | 0.0000143 | 72 | 0.09 | 0.90 | 0.00 | 0.10 | 0.33 | 0.24 |
| DPP4 | 6.7 | 1.8 | 2.35E-10 | 24 | 0.02 | 1.11 | 0.00 | | -0.22 | -0.34 |
| SLC25A22 | 11.4 | 2.5 | 0.000169 | 24 | 0.08 | 0.78 | 0.01 | -0.04 | 0.18 | 0.29 |
| MRPS6 | 348.0 | 5.7 | 1.79E-08 | 24 | 0.03 | 0.87 | 0.01 | 0.30 | 0.12 | 0.20 |
| NLRP3 | 8.3 | 2.1 | 1.81E-07 | 24 | 0.11 | 0.97 | 0.01 | -0.10 | -0.31 | -0.07 |
| PPARD | 100.7 | 4.8 | 8.12E-09 | 24 | 0.08 | 0.93 | 0.01 | -0.18 | -0.24 | -0.09 |
| AMPD3 | 51.3 | 4.0 | 3.19E-08 | 72 | 0.05 | 1.15 | 0.01 | -0.15 | -0.12 | -0.01 |
| CLCN4 | 15.4 | 3.0 | 2.65E-08 | 72 | 0.04 | 1.24 | 0.01 | -0.05 | -0.04 | -0.05 |
| MCTP1 | 33.4 | 3.5 | 0.0000332 | 72 | 0.04 | 0.73 | 0.01 | -0.29 | -0.20 | -0.13 |
| HAPLN3 | 8.7 | 0.8 | 4.63E-07 | 72 | 0.08 | 0.91 | 0.01 | -0.10 | -0.12 | -0.10 |
| UBE2L6 | 94.3 | 4.6 | 2.4E-08 | 72 | 0.01 | 0.77 | 0.01 | -0.37 | -0.22 | -0.21 |
| TSFM | 20.6 | 3.4 | 3.76E-08 | 24 | 0.01 | 0.87 | 0.01 | -0.44 | -0.16 | -0.15 |
| BST2 | 256.5 | 5.7 | 0.0000109 | 72 | 0.03 | 0.73 | 0.01 | -0.10 | 0.16 | 0.10 |
| ZDHHC7 | 53.2 | 4.0 | 0.000144 | 24 | 0.01 | 1.28 | 0.01 | -0.19 | -0.30 | -0.28 |
| ICK | 11.8 | 2.6 | 0.00417 | 24 | 0.07 | 1.26 | 0.01 | -0.23 | -0.13 | -0.10 |
| HAMP | 8.0 | 2.3 | 1.34E-09 | 72 | 0.04 | 0.95 | 0.01 | | | 0.01 |
| MMP12 | 17.6 | 2.5 | 1.09E-09 | 72 | 0.01 | 0.93 | 0.01 | | -0.38 | -0.41 |
| MATK | 65.8 | 4.1 | 6.66E-08 | 24 | 0.04 | 0.88 | 0.01 | | 0.33 | 0.23 |
| LAMP3 | 7.0 | 1.6 | 1.65E-06 | 72 | 0.04 | 0.90 | 0.01 | | 0.12 | -0.03 |
| CYB5R3 | 5.3 | 0.9 | 8.36E-08 | 72 | 0.03 | 1.29 | 0.01 | | -0.23 | -0.16 |
| PDGFB | 6.3 | 1.5 | 0.000159 | 24 | 0.15 | 0.71 | 0.01 | 0.12 | 0.18 | 0.13 |

| | | | | | | | | | | |
|---|---|---|---|---|---|---|---|---|---|---|
| SLC45A3 | 4.8 | 2.0 | 2.17E-06 | 72 | 0.11 | 1.04 | 0.01 | | -0.13 | -0.03 |
| GYPC | 119.8 | 4.9 | 0.00144 | 72 | 0.04 | 0.67 | 0.01 | 0.12 | 0.16 | 0.06 |
| ANGEL1 | 51.7 | 3.8 | 6.66E-09 | 24 | 0.13 | 1.10 | 0.01 | 0.05 | 0.28 | 0.26 |
| ITGA5 | 298.8 | 5.9 | 4.52E-08 | 24 | 0.02 | 0.73 | 0.01 | 0.17 | 0.04 | 0.23 |
| APOO | 16.4 | 3.1 | 2.75E-07 | 72 | 0.06 | 0.64 | 0.01 | 0.19 | 0.13 | 0.14 |
| ITGB8 | 230.3 | 5.1 | 0.0000014 | 72 | 0.15 | 1.24 | 0.01 | -0.19 | -0.46 | -0.15 |
| ARHGAP35 | 38.6 | 3.8 | 0.0000947 | 72 | 0.02 | 1.43 | 0.01 | 0.30 | 0.16 | 0.20 |
| MT1H | 86.0 | 3.3 | 1.12E-07 | 72 | 0.13 | 0.62 | 0.01 | -0.25 | -0.05 | -0.22 |
| MMP2 | 11.8 | 3.1 | 2.1E-08 | 72 | 0.04 | 1.19 | 0.01 | | -0.17 | -0.22 |
| CCL22 | 547.4 | 6.5 | 1.76E-09 | 72 | 0.10 | 0.77 | 0.01 | -0.16 | -0.15 | -0.06 |
| SLIT3 | 6.4 | 1.9 | 0.0000322 | 72 | 0.06 | 1.16 | 0.01 | | 0.21 | 0.27 |
| PSMA2 | 92.6 | 4.6 | 0.000525 | 72 | 0.02 | 0.73 | 0.01 | -0.24 | -0.15 | -0.16 |
| ITGAE | 8.5 | 2.2 | 0.000433 | 72 | 0.13 | 1.56 | 0.01 | 0.06 | -0.09 | -0.09 |
| CCDC71L | 24.6 | 3.6 | 6.28E-09 | 24 | 0.00 | 0.83 | 0.01 | -0.29 | -0.14 | -0.17 |
| SWAP70 | 106.1 | 4.8 | 0.00187 | 72 | 0.13 | 1.25 | 0.01 | -0.08 | -0.25 | -0.02 |
| RCSD1 | 19.3 | 3.1 | 6.23E-08 | 72 | 0.04 | 1.89 | 0.01 | -0.16 | -0.31 | -0.11 |
| CNP | 76.2 | 4.4 | 0.0000168 | 72 | 0.04 | 0.60 | 0.01 | 0.49 | 0.26 | 0.40 |
| SLC46A2 | 5.2 | 1.0 | 5.7E-07 | 24 | 0.04 | 1.34 | 0.01 | 0.17 | 0.13 | 0.07 |
| RHBDF2 | 39.8 | 3.8 | 0.000919 | 72 | 0.06 | 0.92 | 0.02 | -0.50 | -0.47 | -0.18 |
| TNFRSF4 | 5.3 | 0.0 | 4.52E-07 | 72 | 0.01 | 1.17 | 0.02 | | 0.01 | 0.12 |
| SLC28A3 | 18.8 | 2.7 | 2.66E-08 | 24 | 0.06 | 1.15 | 0.02 | -0.19 | -0.05 | 0.02 |
| PSMB8 | 44.0 | 3.7 | 0.0107 | 72 | 0.08 | 0.81 | 0.02 | -0.37 | -0.42 | -0.27 |
| PNPT1 | 39.5 | 3.9 | 3.96E-08 | 72 | 0.01 | 0.46 | 0.02 | 0.08 | 0.44 | 0.21 |
| NAIP | 17.5 | 3.0 | 5.24E-09 | 72 | 0.03 | 1.33 | 0.02 | -0.01 | 0.02 | -0.03 |
| RFTN1 | 24.8 | 3.4 | 7.67E-07 | 72 | 0.00 | 0.76 | 0.02 | -0.24 | -0.18 | 0.00 |
| PPARGC1B | 20.9 | 3.2 | 0.0000157 | 72 | 0.08 | 0.99 | 0.02 | -0.12 | -0.04 | 0.07 |
| LILRA5 | 9.9 | 2.5 | 1.02E-07 | 72 | 0.16 | 1.22 | 0.02 | -0.36 | | -0.26 |
| PTGS1 | 23.0 | 3.5 | 6.68E-06 | 72 | 0.00 | 1.28 | 0.02 | 0.14 | 0.05 | 0.10 |
| TRIM56 | 12.9 | 2.7 | 1.07E-06 | 72 | 0.11 | 0.91 | 0.02 | -0.05 | -0.42 | -0.08 |
| IL1RN | 1,556.9 | 6.8 | 3.45E-08 | 72 | 0.07 | 1.11 | 0.02 | 0.11 | -0.01 | -0.08 |
| SGSH | 16.1 | 3.0 | 0.000236 | 72 | 0.01 | 1.27 | 0.02 | -0.23 | -0.27 | -0.20 |
| CTNS | 48.4 | 4.1 | 4.59E-08 | 24 | 0.10 | 0.86 | 0.02 | -0.25 | -0.49 | -0.41 |
| LACC1 | 20.0 | 2.9 | 0.00669 | 72 | 0.08 | 1.23 | 0.02 | -0.25 | -0.27 | -0.20 |
| ABCA5 | 7.1 | 2.2 | 0.0202 | 24 | 0.13 | 0.96 | 0.02 | -0.06 | -0.06 | -0.07 |
| CD84 | 209.2 | 5.2 | 6.51E-07 | 24 | 0.05 | 0.95 | 0.02 | -0.12 | -0.03 | -0.07 |
| KCNJ5 | 12.9 | 3.0 | 6.4E-09 | 72 | 0.00 | 1.13 | 0.02 | -0.08 | | 0.17 |
| MFAP5 | 6.6 | 1.9 | 5.52E-07 | 72 | 0.04 | 1.21 | 0.02 | | -0.22 | -0.30 |
| LMNB1 | 6.0 | 1.6 | 5.57E-07 | 24 | 0.05 | 0.66 | 0.02 | -0.59 | -0.40 | -0.28 |
| TGFA | 6.8 | 1.9 | 1.37E-10 | 24 | 0.03 | 0.80 | 0.02 | | -0.23 | -0.21 |
| ABHD12 | 155.5 | 5.3 | 1.66E-09 | 72 | 0.02 | 1.23 | 0.03 | -0.03 | 0.38 | 0.13 |
| TRIB2 | 6.9 | 1.9 | 6.73E-11 | 24 | 0.02 | 0.95 | 0.03 | | 0.06 | -0.03 |
| NUP188 | 120.2 | 5.1 | 0.00163 | 24 | 0.01 | 0.80 | 0.03 | -0.03 | 0.09 | 0.12 |
| ITGAX | 326.7 | 5.9 | 0.0000359 | 72 | 0.14 | 0.93 | 0.03 | | | |
| GALM | 13.9 | 3.1 | 2.83E-07 | 72 | 0.13 | 1.25 | 0.03 | -0.13 | -0.09 | -0.17 |
| HIPK2 | 341.0 | 6.1 | 5.38E-07 | 72 | 0.12 | 1.48 | 0.03 | 0.19 | -0.06 | 0.06 |
| TNFSF15 | 7.1 | 1.9 | 5.19E-08 | 24 | 0.15 | 1.04 | 0.03 | -0.10 | -0.13 | -0.12 |
| RASSF2 | 6.1 | 2.3 | 7.88E-06 | 72 | 0.09 | 1.13 | 0.03 | | -0.73 | -0.34 |
| C1S | 53.4 | 4.4 | 5E-08 | 72 | 0.01 | 0.83 | 0.03 | | -0.16 | -0.19 |
| ZNF641 | 8.1 | 2.2 | 0.042 | 24 | 0.15 | 1.39 | 0.03 | 0.03 | -0.18 | 0.17 |
| NPC2 | 587.8 | 6.7 | 0.000176 | 72 | 0.16 | 1.09 | 0.03 | -0.36 | -0.07 | -0.18 |
| TRAPPC5 | 40.6 | 3.9 | 0.00295 | 24 | 0.16 | 1.04 | 0.03 | -0.18 | 0.11 | 0.03 |
| LPCAT1 | 47.7 | 4.1 | 0.0000412 | 24 | 0.16 | 1.12 | 0.03 | 0.03 | 0.04 | 0.04 |
| JAK3 | 6.3 | 1.8 | 2.98E-07 | 72 | 0.06 | 1.11 | 0.03 | | -0.12 | 0.03 |
| ZNF710 | 27.0 | 3.3 | 0.000012 | 72 | 0.14 | 1.18 | 0.03 | -0.22 | -0.28 | -0.20 |
| RCN1 | 68.0 | 4.3 | 0.000156 | 72 | 0.11 | 0.79 | 0.03 | -0.20 | -0.20 | -0.10 |

| | | | | | | | | | | |
|---|---|---|---|---|---|---|---|---|---|---|
| RNF130 | 693.2 | 6.4 | 0.000235 | 24 | 0.15 | 1.15 | 0.04 | -0.09 | 0.30 | 0.10 |
| GAS7 | 49.0 | 4.2 | 2.95E-08 | 72 | 0.08 | 1.23 | 0.04 | -0.02 | 0.02 | -0.08 |
| MCOLN2 | 11.0 | 2.2 | 0.0000013 | 72 | 0.16 | 0.97 | 0.04 | -0.34 | -0.19 | -0.26 |
| RSPO3 | 14.2 | 2.2 | 1.65E-09 | 24 | 0.06 | 0.88 | 0.04 | -0.24 | -0.34 | -0.31 |
| GNA15 | 114.6 | 5.1 | 0.0000263 | 24 | 0.10 | 0.83 | 0.04 | -0.08 | -0.47 | -0.13 |
| ARNTL2 | 18.5 | 3.3 | 0.000489 | 24 | 0.03 | 1.14 | 0.04 | | -0.42 | -0.21 |
| MOV10 | 18.3 | 3.2 | 0.0000126 | 72 | 0.03 | 0.81 | 0.04 | -0.24 | -0.19 | 0.08 |
| GPNMB | 4,056.6 | 8.0 | 1.09E-07 | 24 | 0.04 | 1.28 | 0.04 | 0.01 | 0.39 | 0.09 |
| VGLL4 | 16.9 | 2.8 | 0.00157 | 24 | 0.06 | 1.35 | 0.04 | -0.15 | -0.31 | -0.07 |
| CD226 | 10.5 | 2.5 | 3.78E-08 | 24 | 0.07 | 0.90 | 0.05 | -0.18 | -0.05 | -0.27 |
| ADAR | 412.2 | 6.2 | 0.0000332 | 72 | 0.03 | 0.75 | 0.05 | -0.24 | -0.14 | -0.15 |
| CYBRD1 | 6.3 | 2.2 | 6.54E-08 | 72 | 0.16 | 1.12 | 0.05 | | | 0.01 |
| TRIM5 | 28.8 | 3.5 | 4.52E-06 | 72 | 0.07 | 0.75 | 0.05 | 0.17 | 0.16 | 0.24 |
| NOTCH2 | 203.9 | 5.5 | 1.71E-06 | 72 | 0.11 | 1.10 | 0.05 | 0.24 | -0.09 | 0.05 |
| PXN | 30.6 | 3.0 | 3.83E-07 | 24 | 0.15 | 0.73 | 0.05 | | | |
| SBNO2 | 56.0 | 4.3 | 2.4E-07 | 24 | 0.06 | 0.86 | 0.05 | 0.11 | -0.19 | 0.12 |
| RBP1 | 19.1 | 3.6 | 2.44E-08 | 72 | 0.08 | 1.12 | 0.05 | | 0.06 | 0.03 |
| CADM1 | 9.9 | 2.6 | 2.76E-09 | 72 | 0.12 | 1.27 | 0.05 | | 0.03 | 0.05 |
| MTHFD1 | 97.8 | 4.7 | 0.000523 | 72 | 0.07 | 0.78 | 0.06 | -0.10 | 0.14 | -0.10 |
| PDE4A | 59.1 | 4.1 | 6.74E-06 | 72 | 0.08 | 0.81 | 0.06 | 0.03 | 0.17 | 0.32 |
| LOC100129034 | 270.9 | 5.7 | 0.000237 | 24 | 0.14 | 0.76 | 0.06 | 0.19 | 0.20 | 0.17 |
| LPAR1 | 56.3 | 4.4 | 0.00499 | 24 | 0.03 | 0.89 | 0.06 | -0.01 | -0.08 | -0.12 |
| TPD52 | 12.0 | 2.5 | 4.66E-06 | 72 | 0.13 | 0.86 | 0.06 | -0.20 | 0.05 | -0.32 |
| TMEM132A | 6.1 | 1.9 | 1.46E-08 | 72 | 0.13 | 1.11 | 0.06 | | -0.33 | 0.03 |
| AIM2 | 5.3 | 1.1 | 3.95E-06 | 72 | 0.02 | 0.93 | 0.06 | | -0.18 | -0.20 |
| HLX | 23.6 | 3.4 | 4.65E-08 | 24 | 0.03 | 0.85 | 0.06 | | | |
| PFKP | 35.5 | 3.5 | 0.0000025 | 72 | 0.11 | 0.92 | 0.07 | -0.07 | -0.12 | 0.02 |
| LRP12 | 9.6 | 2.6 | 0.0000232 | 24 | 0.07 | 1.06 | 0.07 | | -0.44 | -0.26 |
| SGSM2 | 26.0 | 3.5 | 3.65E-06 | 72 | 0.10 | 1.29 | 0.08 | -0.05 | 0.22 | 0.19 |
| TPCN1 | 10.0 | 2.9 | 1.03E-08 | 72 | 0.01 | 1.11 | 0.08 | | 0.01 | -0.18 |
| PRDM1 | 20.6 | 3.3 | 0.0139 | 24 | 0.16 | 0.84 | 0.08 | -0.07 | 0.09 | 0.16 |
| PLEKHO1 | 26.5 | 3.7 | 0.0000657 | 72 | 0.01 | 0.87 | 0.08 | -0.20 | -0.37 | -0.32 |
| TNS1 | 41.0 | 4.2 | 1.04E-09 | 72 | 0.03 | 1.24 | 0.08 | -0.03 | -0.16 | 0.03 |
| GBP3 | 54.5 | 3.5 | 0.0000341 | 72 | 0.07 | 0.79 | 0.10 | -0.11 | -0.06 | -0.03 |
| CLLU1OS | 5.7 | 1.7 | 0.0000525 | 24 | 0.06 | 0.98 | 0.10 | | -0.17 | -0.18 |
| CSF2RB | 191.4 | 5.0 | 1.19E-07 | 72 | 0.14 | 0.76 | 0.10 | -0.05 | -0.13 | 0.00 |
| ARHGAP9 | 35.5 | 3.5 | 3.01E-08 | 72 | 0.07 | 1.31 | 0.11 | -0.27 | -0.28 | -0.09 |
| OCSTAMP | 10.1 | 1.6 | 2.52E-09 | 24 | 0.01 | 1.09 | 0.11 | | 0.08 | 0.11 |
| ACVR1B | 67.9 | 4.3 | 0.0000198 | 72 | 0.07 | 0.78 | 0.12 | -0.25 | 0.07 | -0.03 |
| PCYOX1 | 25.7 | 3.0 | 1.03E-07 | 24 | 0.14 | 1.10 | 0.12 | | | |
| NMI | 22.1 | 3.1 | 1.85E-06 | 72 | 0.14 | 0.88 | 0.13 | -0.36 | -0.16 | -0.18 |
| RARRES3 | 40.0 | 2.8 | 0.085 | 24 | 0.14 | 0.54 | 0.14 | -0.26 | -0.08 | -0.13 |
| SAMHD1 | 235.3 | 5.7 | 0.000117 | 72 | 0.10 | 1.27 | 0.14 | -0.55 | -0.10 | -0.12 |
| SDS | 6.5 | 0.7 | 0.000019 | 24 | 0.07 | 1.26 | 0.14 | | | -0.20 |
| ANTXR2 | 14.8 | 2.8 | 0.000219 | 24 | 0.12 | 0.92 | 0.15 | | | |
| LILRB2 | 43.6 | 3.9 | 7.74E-06 | 72 | 0.13 | 1.53 | 0.16 | -0.02 | -0.20 | -0.14 |
| MAP7 | 42.9 | 3.6 | 1.47E-06 | 72 | 0.13 | 0.99 | 0.18 | 0.31 | 0.24 | 0.07 |
| APOL3 | 49.3 | 3.9 | 1.09E-06 | 72 | 0.07 | 0.52 | 0.18 | -0.06 | 0.00 | -0.06 |
| PDCD1LG2 | 86.0 | 4.7 | 1.54E-06 | 72 | 0.15 | 0.67 | 0.19 | -0.36 | -0.20 | -0.15 |
| CCL2 | 120.7 | 4.6 | 6.06E-09 | 24 | 0.16 | 0.95 | 0.23 | -0.31 | -0.01 | -0.33 |
| LCP2 | 123.7 | 4.9 | 3.39E-06 | 24 | 0.08 | 0.78 | 0.23 | -0.36 | -0.35 | -0.17 |
| C1R | 8.7 | 2.4 | 0.0000203 | 72 | 0.04 | 0.77 | 0.34 | | -0.24 | -0.16 |
| IFITM2 | 23.7 | 2.9 | 0.000525 | 72 | 0.13 | 0.78 | 0.35 | -0.20 | -0.12 | -0.11 |

| | | | | | | | | | | |
|---|---|---|---|---|---|---|---|---|---|---|
| INPP5F | 15.3 | 2.9 | 0.00261 | 72 | 0.14 | 0.71 | 0.36 | -0.32 | -0.40 | -0.32 |
| CD38 | 31.5 | 4.0 | 0.000548 | 72 | 0.08 | 0.72 | 0.43 | -0.25 | 0.06 | 0.06 |

**Supplementary File 5: B. Reactome Pathways significantly overrepresented among VE genes**

| geneSet | description | link | size | overlap | expect | enrichmentRatio | pValue | FDR | overlapId | userId |
|---|---|---|---|---|---|---|---|---|---|---|
| R-HSA-168256 | Immune System | http://reactome.org/PathwayBrowser/#/R-HSA-168256 | 1997 | 109 | 44.84451393 | 2.430620614 | 0 | 0 | 10125;103;10410;10577;10581;10666;11274;114548;115361;115362;118788;140459;1435;1439;151636;1727;2040;23511;23586;2517;2537;25939;26270;2633;2634;2635;272;29121;29126;2919;3430;3434;3437;353514;3552;3553;3557;3575;3601;3659;3669;3687;3718;3937;4001;4064;4313;4502;4599;4600;4938;4939;4940;5008;51191;5199;5292;5371;53831;54209;5473;54739;55008;55819;5610;5683;5696;5698;5720;5721;57674;57823;629;6347;6348;6351;6352;6364;6367;63916;64135;64333;6772;6773;684;6890;6891;6993;7097;7130;715;716;7293;79132;79888;79902;80380;81844;841;84166;8519;85363;8767;9246;941;9447;958;9636;9966 | IFI6;IL1B;TNFAIP6;CCL4;CCL20;CCL3;IL7R;SLAMF7;MX1;GBP1;TAP1;STAT2;ISG15;PSMB9;CD274;OAS3;RIPK2;IFI35;GBP5;IFITM3;DYNLT1;IFITM1;IFIH1;CXCL1;IFIT3;PML;DHX58;FBXO6;HERC6;ISG20;DDX58;DTX3L;OAS2;TLR2;HERC5;IL15RA;NLRC5;CSF1;PPBP;USP18;STAT1;PIK3AP1;TAP2;IFIT1;STOM;GPR84;IL1A;GBP4;CFP;CCL5;FUCA1;RASGRP1;CD80;MT2A;XAF1;EIF2AK2;PIM1;IRF1;CFB;MX2;CD40;PSME1;NUP85;ASB6;OSM;CD180;CLEC2D;PSME2;ELMO2;TREM2;GBP2;CASP8;RNF213;OAS1;NLRP3;AMPD3;UBE2L6;BST2;CYB5R3;MMP2;CCL22;PSMA2;TNFRSF4;PSMB8;LILRA5;TRIM56;IL1RN;LMNB1;NUP188;ITGAX;TNFSF15;C1S;NPC2;LPCAT1;JAK3;RNF130;CD226;ADAR;TRIM5;AIM2;GBP3;CSF2RB;ARHGAP9;SAMHD1;PDCD1LG2;CCL2;LCP2;C1R;IFITM2 |
| R-HSA-1280215 | Cytokine Signaling in Immune system | http://reactome.org/PathwayBrowser/#/R-HSA-1280215 | 688 | 66 | 15.44968732 | 4.271931116 | 0 | 0 | 103;10410;10581;11274;115361;115362;1435;1439;23511;23586;2537;25939;2633;2634;2635;2919;3430;3434;3437;3552;3553;3557;3575;3601;3659;3669;3687;3718;4001;4313;4502;4599;4600;4938;4939;4940;5008;51191;5292;5371;54739;5610;5683;5696;5698;5720;5721;6347;6348;6351;6352;6364;6367;6772;6773;684;7293;79902;8519;85363;8767;9246;941;958;9636;9966 | IFI6;IL1B;CCL4;CCL20;CCL3;IL7R;MX1;GBP1;STAT2;ISG15;PSMB9;OAS3;RIPK2;IFI35;GBP5;IFITM3;IFITM1;CXCL1;IFIT3;PML;ISG20;DDX58;OAS2;HERC5;IL15RA;CSF1;USP18;STAT1;IFIT1;IL1A;GBP4;CCL5;CD80;MT2A;XAF1;EIF2AK2;PIM1;IRF1;MX2;CD40;PSME1;NUP85;OSM;PSME2;GBP2;OAS1;UBE2L6;BST2;MMP2;CCL22;PSMA2;TNFRSF4;PSMB8;IL1RN;LMNB1;NUP188;ITGAX;TNFSF15;JAK3;ADAR;TRIM5;GBP3;CSF2RB;SAMHD1;CCL2;IFITM2 |
| R-HSA-913531 | Interferon Signaling | http://reactome.org/PathwayBrowser/#/R-HSA-913531 | 197 | 37 | 4.423820352 | 8.363811604 | 0 | 0 | 103;10410;10581;11274;115361;115362;23511;23586;2537;25939;2633;2634;2635;3430;3434;3437;3659;3669;4502;4599;4600;4938;4939;4940;51191;5371;54739;5610;5696;6772;6773;684;79902;8519;85363;9246;9636 | IFI6;MX1;GBP1;STAT2;ISG15;OAS3;IFI35;GBP5;IFITM3;IFITM1;IFIT3;PML;ISG20;DDX58;OAS2;HERC5;USP18;STAT1;IFIT1;GBP4;MT2A;XAF1;EIF2AK2;IRF1;MX2;NUP85;GBP2;OAS1;UBE2L6;BST2;PSMB8;NUP188;ADAR;TRIM5;GBP3;SAMHD1;IFITM2 |
| R-HSA-909733 | Interferon alpha/beta signaling | http://reactome.org/PathwayBrowser/#/R-HSA-909733 | 69 | 24 | 1.54945992 | 15.48926802 | 0 | 0 | 103;10410;10581;11274;2537;25939;2634;3430;3434;3437;3659;3669;4599;4600;4938;4939;4940;54739;5696;6772;6773;684;8519;9636 | IFI6;MX1;STAT2;ISG15;OAS3;IFI35;IFITM3;IFITM1;IFIT3;ISG20;OAS2;USP18;STAT1;IFIT1;XAF1;IRF1;MX2;GBP2;OAS1;BST2;PSMB8;ADAR;SAMHD1;IFITM2 |
| R-HSA-1169410 | Antiviral mechanism by IFN-stimulated genes | http://reactome.org/PathwayBrowser/#/R-HSA-1169410 | 78 | 15 | 1.751563388 | 8.563777994 | 1.49E-10 | 5.14E-08 | 11274;23511;23586;3434;4599;4600;4938;4939;4940;51191;5610;6772;79902;9246;9636 | MX1;ISG15;OAS3;DDX58;OAS2;HERC5;USP18;STAT1;IFIT1;EIF2AK2;MX2;NUP85;OAS1;UBE2L6;NUP188 |
| R-HSA-6783783 | Interleukin-10 signaling | http://reactome.org/PathwayBrowser/#/R-HSA-6783783 | 47 | 12 | 1.055429221 | 11.36978185 | 3.25E-10 | 9.35E-08 | 1435;2919;3552;3553;3557;6347;6348;6351;6352;6364;6367;941 | IL1B;CCL4;CCL20;CCL3;CXCL1;CSF1;IL1A;CCL5;CD80;CCL22;IL1RN;CCL2 |
| R-HSA-1169408 | ISG15 antiviral mechanism | http://reactome.org/PathwayBrowser/#/R-HSA-1169408 | 71 | 12 | 1.594371802 | 7.526475308 | 4.94E-08 | 1.22E-05 | 11274;23511;23586;3434;4599;4600;51191;5610;6772;79902;9246;9636 | MX1;ISG15;DDX58;HERC5;USP18;STAT1;IFIT1;EIF2AK2;MX2;NUP85;UBE2L6;NUP188 |

| | | | | | | | | | | |
|---|---|---|---|---|---|---|---|---|---|---|
| R-HSA-877300 | Interferon gamma signaling | http://reactome.org/PathwayBrowser/#/R-HSA-877300 | 92 | 13 | 2.065946561 | 6.292515135 | 1.23E-07 | 2.66E-05 | 115361;115362;2633;2634;2635;3659;4502;4938;4939;4940;5371;6772;85363 | GBP1;OAS3;GBP5;PML;OAS2;STAT1;GBP4;MT2A;IRF1;GBP2;OAS1;TRIM5;GBP3 |
| R-HSA-449147 | Signaling by Interleukins | http://reactome.org/PathwayBrowser/#/R-HSA-449147 | 462 | 29 | 10.37464468 | 2.795276454 | 4.88E-07 | 9.37E-05 | 1435;1439;2919;3552;3553;3557;3575;3601;3687;3718;4001;4313;5008;5292;5683;5696;5698;5720;5721;6347;6348;6351;6352;6364;6367;6772;6773;8767;941 | IL1B;CCL4;CCL20;CCL3;IL7R;STAT2;PSMB9;RIPK2;CXCL1;IL15RA;CSF1;STAT1;IL1A;CCL5;CD80;PIM1;PSME1;OSM;PSME2;MMP2;CCL22;PSMA2;PSMB8;IL1RN;LMNB1;ITGAX;JAK3;CSF2RB;CCL2 |
| R-HSA-8983711 | OAS antiviral response | http://reactome.org/PathwayBrowser/#/R-HSA-8983711 | 8 | 4 | 0.179647527 | 22.26582278 | 1.62E-05 | 0.002794 | 23586;4938;4939;4940 | OAS3;DDX58;OAS2;OAS1 |
