## Supplemental Table 6 for "Human alveolar macrophage response to *Mycobacterium tuberculosis*: immune characteristics underlying large inter-individual variability"

**Table 6. Gene module membership and correlation with growth rate (48/72h), with adjusted p values for each module**

**Supplementary File 6: 6A. Network module gene membership and correlations between RNA expression (log2 RPMs) at 2, 24 and 72h in infected cells, and M.tb generation times**

| Network Module Membership - 2hr | | | | Network Module Membership - 24hr | | | | Network Module Membership - 72hr | | | |
|---|---|---|---|---|---|---|---|---|---|---|---|
| GeneID | Module | COR | Pval | GeneID | Module | COR | Pval | GeneID | Module | COR | Pval |
| PCCB | black | -0.577896 | 0.001989 | RASSF2 | black | -0.729181847 | 1.59971E-05 | DGKA | black | -0.370933 | 0.062103 |
| ABHD16B | black | 0.53643 | 0.004727 | PDCD6IP | black | -0.685350601 | 7.98935E-05 | SHISA4 | black | -0.359351 | 0.071389 |
| PEX19 | black | -0.533967 | 0.00496 | PHF21A | black | -0.675323488 | 0.000111161 | LOC550643 | black | -0.338612 | 0.090631 |
| HAUS7 | black | -0.494853 | 0.010167 | MARK4 | black | -0.611263324 | 0.000705987 | NMT2 | black | -0.299195 | 0.137589 |
| HECA | black | 0.465064 | 0.016669 | C17orf80 | black | -0.603885183 | 0.000851902 | ACTR6 | black | -0.289842 | 0.150926 |
| ATP5J2 | black | -0.427637 | 0.029318 | STARD7 | black | -0.601455907 | 0.000905362 | TEFM | black | -0.285995 | 0.156671 |
| SNORA70G | black | 0.426341 | 0.029865 | CLEC5A | black | -0.593389197 | 0.001104342 | PAFAH1B1 | black | -0.257774 | 0.203598 |
| IMMT | black | -0.42047 | 0.032451 | CAMTA2 | black | -0.585320655 | 0.001340223 | DISP1 | black | -0.248281 | 0.221333 |
| SRI | black | -0.418974 | 0.033137 | TMUB1 | black | -0.568501855 | 0.001975229 | FGFR1OP | black | -0.247832 | 0.222198 |
| C17orf75 | black | -0.414612 | 0.035206 | MCRS1 | black | -0.556015551 | 0.002600385 | YARS2 | black | -0.244568 | 0.228544 |
| COX18 | black | -0.411357 | 0.036815 | PSMC5 | black | -0.551980055 | 0.002835674 | PSKH1 | black | -0.242041 | 0.233541 |
| NLRP2 | black | -0.402986 | 0.041225 | GOT2 | black | -0.542645797 | 0.003450692 | CASD1 | black | -0.241273 | 0.235072 |
| PLGRKT | black | -0.396891 | 0.044692 | MRPS7 | black | -0.541055872 | 0.003566043 | CORO1A | black | -0.240057 | 0.237513 |
| TANC2 | black | 0.394104 | 0.046352 | UTP14C | black | -0.527922804 | 0.004651101 | FAM20A | black | -0.232732 | 0.252567 |
| IFIH1 | black | -0.389068 | 0.049474 | ATN1 | black | -0.527223272 | 0.004716001 | GPX4 | black | -0.226533 | 0.26578 |
| FLYWCH2 | black | -0.379022 | 0.056193 | TTBK2 | black | -0.523684853 | 0.005056189 | SUB1 | black | -0.226458 | 0.265943 |
| PANK3 | black | 0.378379 | 0.056646 | HSD17B10 | black | -0.514147822 | 0.00607832 | EBNA1BP2 | black | -0.218019 | 0.284644 |
| EXOSC10 | black | -0.377706 | 0.057124 | PSME3 | black | -0.511263523 | 0.00641984 | PSME2 | black | -0.211324 | 0.300057 |
| PDXDC2P | black | 0.374228 | 0.05964 | MSANTD4 | black | -0.511109338 | 0.006438543 | FBXO46 | black | -0.209454 | 0.304453 |
| INIP | black | -0.36617 | 0.065802 | NDUFA11 | black | -0.510555605 | 0.006506092 | ACRBP | black | -0.203905 | 0.317731 |
| REV3L | black | -0.359855 | 0.070964 | PLEKHM2 | black | -0.509926555 | 0.006583553 | GTF3C3 | black | -0.196831 | 0.335168 |
| NAA50 | black | 0.359553 | 0.071218 | VTI1B | black | -0.504540428 | 0.007279213 | GLO1 | black | -0.196196 | 0.336762 |
| ZMPSTE24 | black | -0.356515 | 0.073818 | SLC39A13 | black | -0.500363388 | 0.007860386 | HIST1H3I | black | -0.195793 | 0.337774 |
| GPN3 | black | -0.355563 | 0.074647 | CCZ1B | black | -0.498711623 | 0.008100716 | TMEM192 | black | -0.194008 | 0.342284 |
| NDUFS2 | black | -0.353797 | 0.076203 | USO1 | black | -0.495910299 | 0.008522401 | CD36 | black | -0.193468 | 0.343655 |
| IGIP | black | 0.352256 | 0.077582 | ACTL6A | black | -0.495529398 | 0.008581135 | GNL2 | black | -0.191783 | 0.347954 |
| NECAP2 | black | -0.352191 | 0.077641 | NUP85 | black | -0.495380778 | 0.008604143 | HIST1H2BE | black | -0.191746 | 0.348049 |
| CRK | black | 0.349685 | 0.079924 | FLYWCH1 | black | -0.494508224 | 0.008740272 | HIST1H4J | black | -0.191378 | 0.348992 |
| STAT1 | black | -0.348262 | 0.081244 | ORAI2 | black | -0.493577347 | 0.008887486 | ATP6V1C1 | black | -0.191285 | 0.349231 |
| MB21D1 | black | -0.347394 | 0.082057 | ARAP1 | black | -0.489322634 | 0.009587057 | CRYZL1 | black | -0.190598 | 0.350998 |
| IMPDH2 | black | 0.346712 | 0.0827 | HLA-F | black | -0.489143919 | 0.009617421 | ACAA2 | black | -0.175613 | 0.390829 |
| SIGLEC7 | black | -0.343989 | 0.085305 | SH2B3 | black | -0.484679869 | 0.010402377 | MRPS23 | black | -0.173923 | 0.395478 |
| ASTE1 | black | 0.343234 | 0.086038 | CAPN2 | black | -0.484360645 | 0.010460499 | ASUN | black | -0.173416 | 0.396878 |
| FNTA | black | -0.342352 | 0.086901 | METRNL | black | -0.481887876 | 0.010919963 | SMC6 | black | -0.172285 | 0.400012 |
| STAT2 | black | -0.341693 | 0.087549 | CTPS1 | black | -0.480671889 | 0.011152001 | EVI5 | black | -0.171098 | 0.403316 |
| AP1S1 | black | -0.339932 | 0.089301 | RPUSD3 | black | -0.479853135 | 0.01131054 | VDAC3 | black | -0.170737 | 0.404325 |
| SLC25A38 | black | -0.338094 | 0.091157 | RTFDC1 | black | -0.476848939 | 0.011908421 | DYM | black | -0.168518 | 0.41055 |
| RAN | black | -0.333088 | 0.09636 | RRAS | black | -0.47595693 | 0.012090922 | TRIAP1 | black | -0.168181 | 0.411499 |
| C18orf32 | black | -0.331883 | 0.097645 | NFKBIE | black | -0.474708999 | 0.012350138 | ANAPC1 | black | -0.167218 | 0.414223 |
| CCDC84 | black | -0.329198 | 0.100554 | LATS2 | black | -0.471586694 | 0.01301898 | HEATR1 | black | -0.167036 | 0.414738 |
| FOXO3 | black | 0.327727 | 0.102175 | MIDN | black | -0.470940996 | 0.01316098 | ZNF830 | black | -0.166355 | 0.416671 |
| CDK14 | black | 0.326406 | 0.103647 | SRSF2 | black | -0.470565781 | 0.013244085 | FAM175B | black | -0.162585 | 0.427455 |
| RRN3P2 | black | -0.3263 | 0.103765 | METTL2B | black | -0.464564738 | 0.014633469 | DTWD2 | black | -0.162572 | 0.427494 |
| SSR2 | black | -0.326008 | 0.104094 | HIST2H2AC | black | -0.464390607 | 0.014675517 | ENOPH1 | black | -0.162227 | 0.428488 |
| UBE2Q2 | black | -0.325869 | 0.10425 | ARMCX3 | black | -0.462455238 | 0.01514961 | HINT3 | black | -0.161822 | 0.429658 |
| FAM136A | black | -0.325145 | 0.105067 | COPS5 | black | -0.461137331 | 0.015479632 | H2AFZ | black | -0.156198 | 0.44607 |
| NR4A3 | black | 0.325033 | 0.105193 | KIAA0240 | black | -0.457582141 | 0.016399606 | HIST1H2AD | black | -0.154749 | 0.450353 |
| ZMAT2 | black | -0.324984 | 0.105249 | TIMM13 | black | -0.454316289 | 0.017283958 | LYST | black | -0.152735 | 0.456342 |
| GGCT | black | -0.324978 | 0.105256 | MAP3K11 | black | -0.453765155 | 0.017436996 | BCCIP | black | -0.151168 | 0.46103 |
| UPRT | black | 0.324527 | 0.105767 | ARRB1 | black | -0.452241262 | 0.017865958 | KIAA0922 | black | -0.150617 | 0.462685 |
| PAPD5 | black | -0.323503 | 0.106938 | GLUL | black | -0.451741576 | 0.018008489 | PPIA | black | -0.150062 | 0.464354 |
| TAF1C | black | -0.3211 | 0.109719 | TRIM27 | black | -0.451150626 | 0.018178255 | CLCN3 | black | -0.148559 | 0.468891 |

**Table 6. Gene module membership and correlation with growth rate (48/72h), with adjusted p values for each module**

| | | | | | | | | | | | |
|---|---|---|---|---|---|---|---|---|---|---|---|
| ZNF264 | black | 0.320641 | 0.110256 | SRSF4 | black | -0.450512962 | 0.018362914 | C2orf47 | black | -0.146672 | 0.474619 |
| BET1L | black | -0.32023 | 0.11074 | RNPS1 | black | -0.450400877 | 0.018395531 | ZNF654 | black | -0.142907 | 0.486156 |
| HADHB | black | 0.319672 | 0.111398 | NACC1 | black | -0.447082573 | 0.019382921 | DYNC2LI1 | black | -0.140092 | 0.494873 |
| HIST1H4D | black | -0.319589 | 0.111496 | CDKN1B | black | -0.445044337 | 0.020010657 | RILPL2 | black | -0.139729 | 0.496002 |
| USP30 | black | -0.316815 | 0.114811 | ABR | black | -0.440311358 | 0.021532678 | WDR75 | black | -0.138406 | 0.500132 |
| RLIM | black | 0.31567 | 0.116201 | RIC8A | black | -0.440241372 | 0.021555874 | PPM1F | black | -0.137582 | 0.502713 |
| DAXX | black | -0.31476 | 0.117315 | APLP2 | black | -0.440032459 | 0.021625235 | ABCE1 | black | -0.13637 | 0.506517 |
| SYNCRIP | black | 0.313559 | 0.118796 | TUBB | black | -0.438776721 | 0.022046004 | ITGB2 | black | -0.136034 | 0.507576 |
| BUD31 | black | -0.312087 | 0.12063 | RAC1 | black | -0.438294063 | 0.022209498 | RHEB | black | -0.134683 | 0.511841 |
| ARHGEF3 | black | -0.310157 | 0.123067 | PLEKHJ1 | black | -0.435678296 | 0.023112846 | SLC25A24 | black | -0.129692 | 0.527745 |
| FHIT | black | -0.308527 | 0.125152 | GTF2H4 | black | -0.429926539 | 0.025204653 | GNA15 | black | -0.129582 | 0.528097 |
| POM121 | black | -0.304914 | 0.129866 | CTTNBP2NL | black | -0.428705259 | 0.025668007 | MRPL40 | black | -0.127177 | 0.535844 |
| RAB8A | black | -0.302669 | 0.132859 | COPA | black | -0.427991594 | 0.025941955 | MGC72080 | black | -0.126352 | 0.538515 |
| MRPL17 | black | -0.297805 | 0.139516 | C12orf10 | black | -0.4272586 | 0.026225785 | VPS8 | black | -0.125274 | 0.542016 |
| ALKBH7 | black | -0.297336 | 0.140171 | FGD3 | black | -0.42710149 | 0.026286948 | KIAA0368 | black | -0.125163 | 0.542374 |
| COMMD4 | black | -0.295451 | 0.142822 | NUP155 | black | -0.426626733 | 0.026472473 | FBXO3 | black | -0.123194 | 0.548797 |
| CBX5 | black | -0.295352 | 0.142962 | SBF1 | black | -0.425516827 | 0.026910343 | DBI | black | -0.122695 | 0.550428 |
| OXA1L | black | -0.294137 | 0.144691 | BRF1 | black | -0.423372485 | 0.027772917 | TMED4 | black | -0.121203 | 0.555322 |
| PPP2R5B | black | -0.290832 | 0.149473 | RELA | black | -0.423184625 | 0.027849538 | AGPS | black | -0.120134 | 0.558843 |
| LETMD1 | black | -0.286754 | 0.155525 | RNH1 | black | -0.421564307 | 0.028517523 | PSME1 | black | -0.119019 | 0.562524 |
| GLT25D1 | black | -0.286483 | 0.155933 | ERGIC1 | black | -0.420127688 | 0.029120549 | ERMP1 | black | -0.115974 | 0.572633 |
| XAB2 | black | -0.286069 | 0.156559 | RELT | black | -0.418676007 | 0.029740311 | TIMM21 | black | -0.112506 | 0.584242 |
| RASA3 | black | 0.285444 | 0.157506 | FBXO7 | black | -0.417280531 | 0.030346063 | PTCD3 | black | -0.104869 | 0.610162 |
| HNRNPA2B1 | black | -0.284504 | 0.158937 | RNF185 | black | -0.415723454 | 0.031033659 | UBR4 | black | -0.104679 | 0.610814 |
| PRPF18 | black | -0.283315 | 0.160763 | HNRNPUL2 | black | -0.415081505 | 0.031320766 | CYTH2 | black | -0.104539 | 0.611294 |
| TRAPPC12 | black | -0.28316 | 0.161001 | SDF2L1 | black | -0.414075671 | 0.031774913 | FAM110A | black | -0.104225 | 0.612371 |
| TLR1 | black | -0.282023 | 0.162763 | LRFN4 | black | -0.413847392 | 0.031878717 | MED6 | black | -0.099139 | 0.62992 |
| CAPRIN2 | black | 0.279279 | 0.167068 | PPP1R35 | black | -0.412385579 | 0.032549926 | CCNK | black | -0.097745 | 0.634765 |
| TRAF7 | black | -0.279265 | 0.16709 | FCGRT | black | -0.412307403 | 0.032586139 | KCTD17 | black | -0.097083 | 0.637071 |
| IKZF1 | black | -0.278821 | 0.167795 | CCDC127 | black | -0.411877624 | 0.0327858 | CD40 | black | -0.096219 | 0.640085 |
| VIM | black | -0.277357 | 0.170131 | COX8A | black | -0.411384781 | 0.033015966 | HIST1H2BD | black | -0.09255 | 0.652947 |
| ZDHHC6 | black | -0.275138 | 0.173716 | SARS | black | -0.411101713 | 0.03314875 | COG6 | black | -0.090759 | 0.65926 |
| FKRP | black | -0.274775 | 0.174307 | EZR | black | -0.40734161 | 0.034953576 | NRAS | black | -0.090524 | 0.660089 |
| CC2D1B | black | -0.273865 | 0.175795 | USB1 | black | -0.406152258 | 0.035540557 | MAP1S | black | -0.089216 | 0.66472 |
| KDSR | black | 0.273607 | 0.176219 | MAP1S | black | -0.405642323 | 0.03579463 | INTS7 | black | -0.087217 | 0.671814 |
| SPOP | black | -0.269755 | 0.18263 | TRADD | black | -0.398503088 | 0.039506686 | EPRS | black | -0.086486 | 0.674416 |
| DMAP1 | black | -0.269671 | 0.182771 | AHNAK | black | -0.397632702 | 0.03997946 | ANO10 | black | -0.085359 | 0.678434 |
| RBM42 | black | -0.269259 | 0.183466 | GNPTG | black | -0.396456539 | 0.040625464 | ACE | black | -0.080139 | 0.697158 |
| TMEM116 | black | -0.268654 | 0.184491 | ENTPD6 | black | -0.395424051 | 0.041199371 | IER3IP1 | black | -0.077831 | 0.70549 |
| S100A11 | black | -0.266894 | 0.187494 | ZC3H3 | black | -0.394383992 | 0.041783982 | UTP3 | black | -0.075473 | 0.714041 |
| PPIL4 | black | -0.265879 | 0.189243 | CDC37 | black | -0.394023348 | 0.041988229 | PSMB9 | black | -0.072878 | 0.723486 |
| USP18 | black | -0.265056 | 0.190666 | C7orf26 | black | -0.392950691 | 0.042600403 | CXCL16 | black | -0.070884 | 0.730774 |
| C8orf73 | black | -0.263066 | 0.194144 | MED15 | black | -0.392920064 | 0.042617986 | HSD17B4 | black | -0.070576 | 0.7319 |
| ZNF713 | black | 0.262047 | 0.19594 | CALHM2 | black | -0.392806502 | 0.04268323 | ZFP91 | black | -0.07016 | 0.733425 |
| LOH12CR1 | black | -0.261706 | 0.196543 | GOSR2 | black | -0.388591455 | 0.045161468 | ZNF747 | black | -0.068409 | 0.739849 |
| MTCH2 | black | -0.260656 | 0.19841 | IRAK1 | black | -0.386200075 | 0.046617258 | ST6GALNAC | black | -0.066927 | 0.7453 |
| USP42 | black | -0.257186 | 0.204667 | CYTH2 | black | -0.384721639 | 0.047535648 | DHX29 | black | -0.066608 | 0.746475 |
| EMC4 | black | -0.255159 | 0.208384 | TMSB4X | black | -0.384333776 | 0.047778933 | NADKD1 | black | -0.066291 | 0.747641 |
| VPS13B | black | 0.252734 | 0.212889 | ST3GAL5 | black | -0.38122945 | 0.049761651 | CCDC50 | black | -0.065936 | 0.74895 |
| CTNS | black | -0.252305 | 0.213694 | NDUFB10 | black | -0.381205929 | 0.049776917 | NIP7 | black | -0.062409 | 0.761989 |
| PQLC2 | black | 0.252236 | 0.213822 | FAM50A | black | -0.380600134 | 0.05017137 | UBQLN2 | black | -0.060339 | 0.769672 |
| NF2 | black | 0.250417 | 0.217256 | VAMP3 | black | -0.380332249 | 0.050346581 | ZCCHC4 | black | -0.058588 | 0.776187 |
| LILRB3 | black | -0.249115 | 0.219735 | KAT8 | black | -0.378119681 | 0.051812176 | NUDT22 | black | -0.058254 | 0.777429 |
| PANK4 | black | 0.248635 | 0.220655 | ZC3H15 | black | -0.377585142 | 0.052171226 | SKIV2L2 | black | -0.055939 | 0.786069 |
| EIF2C4 | black | 0.246889 | 0.224018 | SIVA1 | black | -0.375777569 | 0.053399849 | CYB5D2 | black | -0.054611 | 0.791037 |
| HEBP1 | black | 0.246883 | 0.22403 | ISY1 | black | -0.375608297 | 0.053516056 | KLF3 | black | -0.051447 | 0.802902 |

**Table 6. Gene module membership and correlation with growth rate (48/72h), with adjusted p values for each module**

| | | | | | | | | | | | |
|---|---|---|---|---|---|---|---|---|---|---|---|
| VIPAS39 | black | -0.244933 | 0.227828 | WDR74 | black | -0.375317313 | 0.053716281 | IRF1 | black | -0.050861 | 0.805106 |
| FAM102A | black | -0.244859 | 0.227973 | IFT57 | black | -0.374618883 | 0.054199264 | CHRM3 | black | -0.046569 | 0.82128 |
| SAAL1 | black | 0.244166 | 0.229334 | SGK223 | black | -0.372987902 | 0.055340371 | LMLN | black | -0.046536 | 0.821403 |
| RHOG | black | -0.24395 | 0.229761 | CDK11B | black | -0.372972789 | 0.055351031 | CAHM | black | -0.044728 | 0.82824 |
| MBTPS1 | black | 0.242618 | 0.232394 | PNPLA4 | black | -0.370443551 | 0.057157921 | SMARCE1 | black | -0.042067 | 0.838321 |
| CTIF | black | -0.239456 | 0.238726 | SCARNA2 | black | -0.369409033 | 0.057910107 | CD4 | black | -0.04132 | 0.841158 |
| CMAS | black | -0.238647 | 0.240364 | ATP6V1E1 | black | -0.36900267 | 0.058207672 | SH3TC1 | black | -0.03797 | 0.853894 |
| CLCN5 | black | 0.237975 | 0.241729 | UPF2 | black | -0.368081303 | 0.058886769 | HIST1H3A | black | -0.037644 | 0.855133 |
| BHLHE40 | black | -0.23692 | 0.243885 | RPL13AP20 | black | -0.367084224 | 0.059628605 | PATL1 | black | -0.037556 | 0.85547 |
| BTK | black | -0.236216 | 0.24533 | FMNL1 | black | -0.362325492 | 0.063269833 | HSPA14 | black | -0.037208 | 0.856795 |
| SPAG7 | black | -0.233455 | 0.251054 | RPS9 | black | -0.361066284 | 0.064261593 | MAGT1 | black | -0.037007 | 0.85756 |
| CTSS | black | -0.232406 | 0.253249 | IPO9 | black | -0.36023839 | 0.06492018 | LLGL1 | black | -0.035439 | 0.863538 |
| NUP133 | black | -0.230927 | 0.256368 | AGPAT2 | black | -0.357100855 | 0.067463593 | HHEX | black | -0.034432 | 0.867384 |
| RIMKLB | black | 0.23092 | 0.256384 | PGP | black | -0.355552815 | 0.0687465 | JUP | black | -0.034375 | 0.867599 |
| SSBP4 | black | -0.230013 | 0.258309 | DOK3 | black | -0.340143482 | 0.082564529 | DHRS1 | black | -0.032937 | 0.873092 |
| PRKAR1A | black | -0.22942 | 0.259573 | CNTROB | black | -0.339681659 | 0.083008997 | CSNK1G2 | black | -0.032496 | 0.874781 |
| PEPD | black | 0.229295 | 0.25984 | RPS15A | black | -0.33919918 | 0.083475284 | GBP3 | black | -0.032199 | 0.875915 |
| FCAR | black | -0.228854 | 0.260782 | ANKRD27 | black | -0.337339012 | 0.085291675 | RALB | black | -0.026506 | 0.897731 |
| ARHGAP15 | black | -0.228653 | 0.261213 | UBL4A | black | -0.33681457 | 0.085809152 | QSOX1 | black | -0.025447 | 0.901796 |
| SGSH | black | -0.227985 | 0.262647 | TCEA1 | black | -0.336798505 | 0.085825041 | HNRNPU | black | -0.024818 | 0.904212 |
| USP34 | black | 0.227811 | 0.263021 | RNASEH2B | black | -0.3359456 | 0.086671817 | MCM4 | black | -0.022736 | 0.912217 |
| ALG12 | black | -0.226412 | 0.266043 | CPT1A | black | -0.333214655 | 0.089425739 | SIPA1L3 | black | -0.019941 | 0.922974 |
| SLMAP | black | -0.226213 | 0.266474 | TSC22D4 | black | -0.331516733 | 0.091170967 | WAS | black | -0.01975 | 0.923708 |
| VMP1 | black | -0.225747 | 0.267487 | MAVS | black | -0.329482912 | 0.093295129 | PIM2 | black | -0.015885 | 0.938608 |
| SRSF9 | black | -0.223967 | 0.271379 | TUFM | black | -0.329395788 | 0.093386948 | TCF25 | black | -0.011458 | 0.955698 |
| TFIP11 | black | -0.221927 | 0.275882 | COMMD9 | black | -0.324622373 | 0.098522287 | SPATS2 | black | -0.010598 | 0.959019 |
| BRD9 | black | 0.219968 | 0.280252 | CORO7 | black | -0.322518524 | 0.100851647 | CD72 | black | -0.009981 | 0.961404 |
| GBGT1 | black | -0.219264 | 0.281834 | TMEM214 | black | -0.31933131 | 0.104458591 | TEX264 | black | -0.008914 | 0.965529 |
| CD2BP2 | black | -0.21887 | 0.28272 | ZBTB6 | black | -0.318221832 | 0.105736462 | ZFYVE27 | black | -0.008039 | 0.96891 |
| SYNC | black | -0.218163 | 0.284317 | PPP1R11 | black | -0.317871558 | 0.106142305 | CCSAP | black | -0.005808 | 0.977536 |
| TIMM23 | black | 0.215844 | 0.289594 | TP53I11 | black | -0.315662494 | 0.108728575 | C3orf17 | black | -0.004485 | 0.982651 |
| INTS4 | black | 0.212915 | 0.296347 | PHF13 | black | -0.314850015 | 0.109691457 | PTAFR | black | 0.003018 | 0.988326 |
| SREBF1 | black | -0.212382 | 0.297586 | MRPL28 | black | -0.314446558 | 0.110171943 | SMG6 | black | 0.003751 | 0.98549 |
| BANF1 | black | 0.210975 | 0.300874 | ECI1 | black | -0.313500568 | 0.111304661 | CCDC130 | black | 0.01014 | 0.960789 |
| USP10 | black | -0.210551 | 0.301869 | RABEP2 | black | -0.312259733 | 0.112803466 | CD99L2 | black | 0.015325 | 0.940768 |
| CRKL | black | -0.206061 | 0.312531 | TUBGCP2 | black | -0.310243655 | 0.115270436 | RNF6 | black | 0.017833 | 0.931097 |
| TBRG1 | black | 0.205667 | 0.313478 | PEX16 | black | -0.307767193 | 0.118354962 | ARMC1 | black | 0.020734 | 0.919921 |
| PITPNB | black | 0.205095 | 0.314854 | LAMP1 | black | -0.307655416 | 0.118495602 | SETDB2 | black | 0.021955 | 0.91522 |
| KIAA1468 | black | -0.204982 | 0.315127 | NAGPA | black | -0.304906204 | 0.121993552 | TPCN2 | black | 0.022031 | 0.914927 |
| BBS7 | black | -0.204952 | 0.3152 | PEX11B | black | -0.30002971 | 0.128383495 | KIAA0020 | black | 0.024006 | 0.907334 |
| LARP4 | black | 0.202836 | 0.320331 | CMC4 | black | -0.297092455 | 0.132348236 | TULP3 | black | 0.025418 | 0.901906 |
| RBPJ | black | -0.201916 | 0.322577 | EI24 | black | -0.296749768 | 0.132816531 | EXOC8 | black | 0.026005 | 0.899652 |
| PA2G4 | black | 0.200372 | 0.326369 | CDKN2D | black | -0.296648599 | 0.132955012 | JUNB | black | 0.027221 | 0.894985 |
| STAC | black | 0.20012 | 0.326991 | CHCHD10 | black | -0.29664641 | 0.132958009 | DCP2 | black | 0.030644 | 0.881867 |
| SNTB2 | black | 0.199612 | 0.328245 | CCDC86 | black | -0.296414344 | 0.133276065 | TBL1X | black | 0.03113 | 0.880005 |
| VEZT | black | 0.19923 | 0.329191 | OR2A1 | black | -0.296393524 | 0.133304626 | PFKFB3 | black | 0.032913 | 0.873186 |
| SEMA4D | black | -0.19908 | 0.329564 | TARDBP | black | -0.292178679 | 0.139178652 | NDUFAF2 | black | 0.036961 | 0.857735 |
| RNF113A | black | 0.198752 | 0.330376 | TRMT112 | black | -0.291696123 | 0.139862906 | RAB5B | black | 0.040724 | 0.84342 |
| BCCIP | black | -0.198239 | 0.331653 | MRPL27 | black | -0.287024701 | 0.146612838 | HCK | black | 0.042349 | 0.83725 |
| PMPCA | black | -0.197583 | 0.333288 | TRIO | black | -0.286407882 | 0.147521275 | EPHA2 | black | 0.043452 | 0.833072 |
| PKN2 | black | -0.197362 | 0.333838 | MAP3K3 | black | -0.286355126 | 0.14759916 | TGIF2 | black | 0.044039 | 0.830848 |
| BTBD7 | black | 0.193984 | 0.342344 | CRAMP1L | black | -0.28601502 | 0.148101974 | FXN | black | 0.046561 | 0.821309 |
| TRIM36 | black | 0.193094 | 0.344607 | PCGF5 | black | -0.28208238 | 0.15400542 | FOXO4 | black | 0.052134 | 0.800323 |
| IQGAP1 | black | 0.191776 | 0.347973 | MYO9B | black | -0.281264778 | 0.155253527 | AGXT2L2 | black | 0.054219 | 0.792503 |
| GAB2 | black | -0.191341 | 0.349089 | POMP | black | -0.279895068 | 0.157360573 | C16orf13 | black | 0.055016 | 0.789521 |
| ARIH2 | black | -0.19128 | 0.349246 | LRR1 | black | -0.274620471 | 0.165664427 | AHI1 | black | 0.055193 | 0.78886 |

**Table 6. Gene module membership and correlation with growth rate (48/72h), with adjusted p values for each module**

| | | | | | | | | | | | |
|---|---|---|---|---|---|---|---|---|---|---|---|
| USP39 | black | -0.191073 | 0.349777 | HK1 | black | -0.273656938 | 0.167214116 | WASL | black | 0.055454 | 0.787884 |
| SRRT | black | -0.190408 | 0.351487 | AP1S1 | black | -0.272703961 | 0.168756851 | GYPC | black | 0.056596 | 0.783615 |
| UXT | black | -0.189245 | 0.354491 | ZFP3 | black | -0.271982987 | 0.169930646 | CALU | black | 0.057071 | 0.781842 |
| MDH2 | black | -0.189145 | 0.354751 | ZRANB2 | black | -0.271356453 | 0.170955339 | POLRMT | black | 0.058761 | 0.775541 |
| TROVE2 | black | -0.189005 | 0.355113 | ZGPAT | black | -0.27004669 | 0.173111461 | CYTH1 | black | 0.062234 | 0.762638 |
| C6orf47 | black | 0.188445 | 0.356568 | CDK10 | black | -0.269499151 | 0.174018444 | UMPS | black | 0.064645 | 0.753715 |
| UFM1 | black | 0.187615 | 0.358729 | CRY1 | black | -0.266808876 | 0.178523202 | MBNL2 | black | 0.066356 | 0.747402 |
| ZDHHC7 | black | -0.187146 | 0.359954 | LRPAP1 | black | -0.265755277 | 0.180309399 | VAV1 | black | 0.070535 | 0.73205 |
| BBX | black | -0.186862 | 0.360696 | SERINC1 | black | -0.264105516 | 0.183131247 | OCEL1 | black | 0.070976 | 0.730438 |
| PPT1 | black | -0.186553 | 0.361506 | POLR3F | black | -0.26115941 | 0.188246533 | CCDC159 | black | 0.073687 | 0.720538 |
| NAPG | black | 0.186349 | 0.362041 | S100A6 | black | -0.258298537 | 0.193307684 | ALDH2 | black | 0.074691 | 0.716883 |
| PML | black | -0.184854 | 0.365973 | NIT1 | black | -0.257261242 | 0.195165704 | WBP1L | black | 0.079071 | 0.701011 |
| HPS3 | black | 0.184634 | 0.366554 | ZNF655 | black | -0.24639372 | 0.215372741 | MARS | black | 0.080025 | 0.69757 |
| MED17 | black | 0.184365 | 0.367264 | POP7 | black | -0.246136428 | 0.215867653 | NBR1 | black | 0.08027 | 0.696687 |
| PIP4K2B | black | 0.183767 | 0.368847 | ACO2 | black | -0.24482025 | 0.2184114 | H6PD | black | 0.082637 | 0.688175 |
| IRF8 | black | -0.183619 | 0.369241 | SKP2 | black | -0.243671672 | 0.22064768 | TNIP2 | black | 0.087555 | 0.670615 |
| RAB5C | black | 0.183108 | 0.370598 | DVL2 | black | -0.242937843 | 0.222084482 | CA13 | black | 0.088686 | 0.666598 |
| RPS25 | black | -0.181302 | 0.375416 | SNRPB | black | -0.240631001 | 0.226642063 | DSTYK | black | 0.092509 | 0.653092 |
| ZNF597 | black | -0.180535 | 0.377472 | PRKAG1 | black | -0.240517122 | 0.226868661 | HIST1H4E | black | 0.092542 | 0.652975 |
| CSNK1G1 | black | -0.180479 | 0.377624 | JMJD7 | black | -0.239804477 | 0.228290132 | PTGS1 | black | 0.096054 | 0.640662 |
| NAAA | black | 0.179043 | 0.381495 | NOM1 | black | -0.238535859 | 0.230835274 | MDC1 | black | 0.097071 | 0.637111 |
| PLEKHB2 | black | 0.178795 | 0.382165 | PLEKHM1P | black | -0.23819994 | 0.231512361 | CLSTN1 | black | 0.099451 | 0.628836 |
| SNHG1 | black | 0.17858 | 0.382748 | RPGR | black | -0.238095561 | 0.231723019 | ENGASE | black | 0.101276 | 0.622522 |
| RNF187 | black | -0.177874 | 0.384661 | GHITM | black | -0.23669482 | 0.234562361 | COL9A2 | black | 0.104207 | 0.612432 |
| TIMMDC1 | black | -0.176335 | 0.388853 | PRDM4 | black | -0.233776151 | 0.240552615 | TH1L | black | 0.104464 | 0.611552 |
| SSH1 | black | -0.175756 | 0.390438 | PTPN18 | black | -0.23323044 | 0.241683746 | RRP12 | black | 0.109018 | 0.596023 |
| ZNF141 | black | 0.175434 | 0.391322 | TNPO1 | black | -0.232904511 | 0.242360992 | C11orf58 | black | 0.109482 | 0.59445 |
| ULK4 | black | -0.174924 | 0.392721 | POLG | black | -0.230291443 | 0.247835947 | BCL2L11 | black | 0.111068 | 0.589088 |
| MPZL2 | black | 0.174329 | 0.39436 | DDX23 | black | -0.226860713 | 0.255146494 | GOLGA3 | black | 0.112061 | 0.585739 |
| EIF2AK1 | black | 0.171551 | 0.402054 | DNAJC17 | black | -0.224584015 | 0.260074775 | RXRB | black | 0.112329 | 0.584837 |
| FAM192A | black | -0.171389 | 0.402504 | TXNDC5 | black | -0.222095163 | 0.265532567 | PHF12 | black | 0.115754 | 0.573366 |
| ZNF586 | black | 0.169943 | 0.406546 | C1orf122 | black | -0.222040869 | 0.265652445 | WBP2 | black | 0.117409 | 0.567858 |
| EXOSC8 | black | 0.169834 | 0.406851 | RIPK1 | black | -0.21431468 | 0.283068499 | TNF | black | 0.118008 | 0.56587 |
| TSPYL4 | black | -0.168962 | 0.409301 | SON | black | -0.213169228 | 0.285710891 | ABCD4 | black | 0.122374 | 0.55148 |
| APOOL | black | 0.168626 | 0.410247 | RNF141 | black | -0.213087401 | 0.28590025 | HIST1H4K | black | 0.130641 | 0.5247 |
| RBM10 | black | -0.167006 | 0.414822 | ELAVL1 | black | -0.209266814 | 0.294830233 | ZHX2 | black | 0.134195 | 0.513383 |
| STYXL1 | black | -0.165569 | 0.418906 | PARP2 | black | -0.208544384 | 0.296538287 | PLSCR3 | black | 0.137896 | 0.501728 |
| MMAB | black | -0.163663 | 0.424357 | AKT1S1 | black | -0.206311106 | 0.301857669 | TTC28 | black | 0.143973 | 0.482875 |
| SPRED1 | black | 0.162762 | 0.426946 | PAGR1 | black | -0.205877499 | 0.30289733 | RAB43 | black | 0.147045 | 0.473484 |
| LNPEP | black | 0.162623 | 0.427347 | PIR | black | -0.203518557 | 0.308592457 | SLC26A6 | black | 0.147392 | 0.472431 |
| ZNF224 | black | 0.160704 | 0.432893 | FAM86A | black | -0.203376488 | 0.308937556 | BBC3 | black | 0.156882 | 0.444057 |
| RAE1 | black | -0.160473 | 0.433563 | SNORA14B | black | -0.198390582 | 0.321200288 | DLG4 | black | 0.15813 | 0.440396 |
| PABPC4 | black | -0.160226 | 0.434282 | ARL6IP4 | black | -0.198291603 | 0.321446705 | TBC1D17 | black | 0.158708 | 0.438705 |
| UBA2 | black | 0.159773 | 0.435599 | ZBTB44 | black | -0.195186722 | 0.329235273 | MFGE8 | black | 0.159338 | 0.436867 |
| RALGDS | black | -0.159666 | 0.435909 | PCIF1 | black | -0.19381022 | 0.332724608 | ANKRD13D | black | 0.160618 | 0.433143 |
| TTLL5 | black | -0.158032 | 0.440683 | YLPM1 | black | -0.192832073 | 0.335217706 | HNRNPF | black | 0.161521 | 0.430528 |
| NBR1 | black | -0.155875 | 0.447024 | TFE3 | black | -0.185264885 | 0.354884499 | CTSH | black | 0.167798 | 0.412582 |
| THOC2 | black | 0.15579 | 0.447276 | MIA3 | black | -0.185102524 | 0.35531381 | ASB6 | black | 0.172973 | 0.398104 |
| ATP8B3 | black | -0.155716 | 0.447493 | NAA10 | black | -0.184171142 | 0.357782485 | PLEKHM1 | black | 0.173215 | 0.397434 |
| RCN3 | black | -0.15556 | 0.447955 | CSNK1E | black | -0.183538208 | 0.359465872 | SDC3 | black | 0.174845 | 0.392939 |
| SLC35B1 | black | 0.155142 | 0.449191 | CCRN4L | black | -0.179920177 | 0.369177941 | ADA | black | 0.176058 | 0.389611 |
| BAG4 | black | 0.154928 | 0.449823 | AMOTL1 | black | -0.174909716 | 0.382877661 | FAM220A | black | 0.176718 | 0.387809 |
| RPN1 | black | 0.154911 | 0.449874 | FIS1 | black | -0.171820942 | 0.391466646 | WDR48 | black | 0.180215 | 0.378334 |
| PRMT5 | black | 0.151804 | 0.459124 | FAM89A | black | -0.167957075 | 0.402363914 | TYK2 | black | 0.184287 | 0.36747 |
| EIF2S1 | black | 0.151169 | 0.461026 | ETFA | black | -0.162590617 | 0.417778403 | RUNX2 | black | 0.18611 | 0.362667 |
| ZNF304 | black | -0.150976 | 0.461606 | SLC39A3 | black | -0.161871801 | 0.419867594 | GRN | black | 0.186861 | 0.360697 |

**Table 6. Gene module membership and correlation with growth rate (48/72h), with adjusted p values for each module**

| | | | | | | | | | | | |
|---|---|---|---|---|---|---|---|---|---|---|---|
| INPPL1 | black | -0.150465 | 0.463142 | TRIM44 | black | -0.158284266 | 0.430380255 | MLLT6 | black | 0.188567 | 0.35625 |
| TMEM9B | black | 0.149061 | 0.467375 | CEMP1 | black | -0.157756767 | 0.431937991 | AMPD2 | black | 0.190696 | 0.350746 |
| TRMT6 | black | 0.146481 | 0.475201 | CISD3 | black | -0.154352544 | 0.442064269 | CPNE1 | black | 0.197409 | 0.333722 |
| FAM134C | black | -0.146157 | 0.47619 | RANBP17 | black | -0.150866569 | 0.45256444 | TNFAIP2 | black | 0.204977 | 0.315138 |
| MYO5A | black | 0.145976 | 0.476742 | SOX13 | black | -0.148924343 | 0.458471486 | STAT5B | black | 0.205325 | 0.3143 |
| ARHGEF1 | black | -0.144783 | 0.480389 | KIAA0040 | black | -0.148640623 | 0.459337767 | RARRES1 | black | 0.205372 | 0.314187 |
| SPG11 | black | -0.144073 | 0.482568 | ZFPL1 | black | -0.145750648 | 0.468210549 | DIP2A | black | 0.207029 | 0.310213 |
| TMEM87A | black | 0.14263 | 0.48701 | PPCS | black | -0.143475737 | 0.475257097 | ENTPD4 | black | 0.208046 | 0.307789 |
| CA5BP1 | black | -0.140131 | 0.494751 | USF2 | black | -0.139220074 | 0.488584233 | OSCAR | black | 0.211172 | 0.300411 |
| ANKH | black | 0.137712 | 0.502303 | MECR | black | -0.133288448 | 0.507470219 | ARRDC2 | black | 0.212151 | 0.298124 |
| LRRC28 | black | 0.137515 | 0.502923 | SBNO1 | black | -0.129939568 | 0.518289357 | LOC1001310[illegible] | black | 0.21239 | 0.297567 |
| CALR | black | -0.137114 | 0.504181 | POLE4 | black | -0.128168793 | 0.524055018 | PPP1R15A | black | 0.212754 | 0.296721 |
| DHX35 | black | 0.137029 | 0.504447 | SAR1A | black | -0.126981422 | 0.527938307 | FANCC | black | 0.215553 | 0.290261 |
| ZNF266 | black | 0.135289 | 0.509927 | VPS36 | black | -0.11763208 | 0.558987406 | MTMR3 | black | 0.219681 | 0.280896 |
| CIRBP | black | -0.134305 | 0.513035 | HYOU1 | black | -0.106770513 | 0.596068303 | PCED1A | black | 0.22652 | 0.26581 |
| GAS2L3 | black | 0.134239 | 0.513247 | DNAJC14 | black | -0.105252874 | 0.601331977 | TMED8 | black | 0.22743 | 0.263841 |
| PDPK1 | black | -0.132532 | 0.518666 | ANXA5 | black | -0.102660323 | 0.610369 | ZNF414 | black | 0.227956 | 0.262709 |
| PDCD10 | black | 0.132386 | 0.519131 | PYGL | black | -0.100984203 | 0.61624148 | FAM78A | black | 0.228255 | 0.262066 |
| SLC30A9 | black | 0.132155 | 0.519867 | LAMP2 | black | -0.100204112 | 0.618982531 | UNKL | black | 0.242469 | 0.232689 |
| AGGF1 | black | -0.132117 | 0.519986 | CBLB | black | -0.098204256 | 0.626032222 | EZH1 | black | 0.248334 | 0.221232 |
| USP9X | black | 0.131977 | 0.520434 | ZCCHC16 | black | -0.095016702 | 0.637335072 | MLLT1 | black | 0.249777 | 0.218472 |
| HNRNPUL1 | black | -0.131815 | 0.520949 | NENF | black | -0.090027211 | 0.655186722 | AMBRA1 | black | 0.251848 | 0.214551 |
| ZNF782 | black | -0.129114 | 0.5296 | FBL | black | -0.078592852 | 0.696787915 | GPR108 | black | 0.252964 | 0.212459 |
| WDR33 | black | 0.127042 | 0.536281 | VPS13C | black | -0.066507638 | 0.741704913 | ZNF41 | black | 0.255937 | 0.206953 |
| TMEM123 | black | -0.126744 | 0.537246 | NAA16 | black | -0.058456726 | 0.772103413 | ANGEL1 | black | 0.2599 | 0.199762 |
| DHPS | black | 0.126614 | 0.537666 | DNM2 | black | -0.050840784 | 0.801165796 | ARHGEF2 | black | 0.26555 | 0.189811 |
| NSMCE1 | black | 0.126324 | 0.538606 | ACSL4 | black | -0.043012996 | 0.831307827 | LGALS3BP | black | 0.266386 | 0.188367 |
| ZNF44 | black | 0.125516 | 0.541229 | MRPL51 | black | -0.037597437 | 0.852301078 | SEC16A | black | 0.267131 | 0.187089 |
| NDOR1 | black | -0.125218 | 0.542198 | ALG6 | black | -0.036629278 | 0.856064913 | MED29 | black | 0.270634 | 0.181153 |
| IMPACT | black | 0.124588 | 0.544248 | FAM195A | black | -0.015437336 | 0.93908223 | DHX8 | black | 0.273816 | 0.175876 |
| TERF2IP | black | -0.12441 | 0.544828 | NDFIP2 | black | -0.012744067 | 0.949695599 | C19orf38 | black | 0.277023 | 0.170667 |
| NLRC5 | black | -0.123697 | 0.547152 | MIB1 | black | -0.008315897 | 0.967163091 | SMAD4 | black | 0.281366 | 0.163787 |
| POLD2 | black | 0.122155 | 0.552199 | ATR | black | 0.001984192 | 0.992163081 | PHF8 | black | 0.284368 | 0.159147 |
| ASNA1 | black | 0.121311 | 0.554969 | NUDT3 | black | 0.007195489 | 0.971585346 | SLC25A22 | black | 0.285182 | 0.157904 |
| YWHAH | black | -0.119801 | 0.559939 | CAAP1 | black | 0.008240586 | 0.967460317 | RAB11B | black | 0.28556 | 0.157329 |
| CD33 | black | -0.118894 | 0.562937 | MGAM | black | 0.02666546 | 0.894964851 | HDAC7 | black | 0.306397 | 0.127915 |
| SP1 | black | 0.117991 | 0.565927 | FBXO48 | black | 0.026744048 | 0.894656987 | ZBTB8A | black | 0.326346 | 0.103714 |
| DOCK2 | black | -0.117548 | 0.567397 | PCMTD1 | black | 0.029287405 | 0.884701756 | PLEKHO2 | black | 0.332444 | 0.097045 |
| SEC62 | black | 0.116767 | 0.569991 | MTERFD2 | black | 0.033887522 | 0.866740245 | MAU2 | black | 0.33616 | 0.093142 |
| SCARB2 | black | 0.114903 | 0.576208 | TAOK1 | black | 0.035187685 | 0.861674957 | SUPT6H | black | 0.33648 | 0.092811 |
| PITPNM1 | black | -0.114702 | 0.576879 | CARD16 | black | 0.035530348 | 0.86034086 | PFKL | black | 0.341898 | 0.087347 |
| FAM76B | black | -0.112789 | 0.58329 | ATAT1 | black | 0.036054683 | 0.858300181 | CRTC2 | black | 0.343406 | 0.085871 |
| SUOX | black | 0.112779 | 0.583325 | GALNT4 | black | 0.041859624 | 0.835769881 | SASH1 | black | 0.345152 | 0.084184 |
| GIMAP2 | black | -0.112062 | 0.585737 | REST | black | 0.051744719 | 0.797702035 | SETD1B | black | 0.355028 | 0.075116 |
| NDUFS6 | black | -0.110652 | 0.590491 | CCNT2 | black | 0.062383081 | 0.757234259 | CALCOCO1 | black | 0.366992 | 0.065151 |
| CCT4 | black | 0.109968 | 0.592803 | WDR82 | black | 0.066179292 | 0.742937644 | PARP10 | black | 0.367274 | 0.064929 |
| IPPK | black | -0.108047 | 0.59932 | UBE2G1 | black | 0.067710576 | 0.737193969 | TMEM63B | black | 0.372031 | 0.061274 |
| C2CD3 | black | -0.106188 | 0.605653 | TMEM70 | black | 0.072055804 | 0.720971269 | PPP1R16A | black | 0.376537 | 0.05796 |
| MYCBP2 | black | 0.106013 | 0.60625 | KIAA1432 | black | 0.078883819 | 0.695717991 | IRF7 | black | 0.380333 | 0.055278 |
| ZZZ3 | black | -0.105929 | 0.606536 | DCK | black | 0.079249597 | 0.694373789 | FARP2 | black | 0.422208 | 0.031667 |
| ZCCHC11 | black | -0.102952 | 0.616746 | TMEM30A | black | 0.085741619 | 0.670669468 | PKD1 | black | 0.442213 | 0.023697 |
| ZNF680 | black | 0.10267 | 0.617714 | EIF3D | black | 0.089877928 | 0.655723759 | SLC3A2 | black | 0.445951 | 0.022406 |
| HSPH1 | black | -0.102424 | 0.618563 | PRDM1 | black | 0.093648561 | 0.642210988 | AGRN | black | 0.449945 | 0.02109 |
| SACM1L | black | -0.101792 | 0.620741 | SRFBP1 | black | 0.101330761 | 0.615025367 | MARK2 | black | 0.454018 | 0.019815 |
| CTCF | black | 0.101666 | 0.621174 | ABCF2 | black | 0.103332745 | 0.608019667 | NBPF24 | black | 0.49667 | 0.009851 |
| PPP2R4 | black | 0.101117 | 0.62307 | CNOT7 | black | 0.104337011 | 0.604517985 | TTC39C | blue | -0.509192 | 0.00789 |

**Table 6. Gene module membership and correlation with growth rate (48/72h), with adjusted p values for each module**

| | | | | | | | | | | | |
|---|---|---|---|---|---|---|---|---|---|---|---|
| SEC63 | black | -0.099632 | 0.62821 | RBM7 | black | 0.115541294 | 0.56604286 | TBCE | blue | -0.474829 | 0.014241 |
| EEF1A1 | black | -0.099456 | 0.628821 | MTR | black | 0.116181505 | 0.563878182 | SLC35D2 | blue | -0.404069 | 0.040632 |
| PPFIBP1 | black | -0.099435 | 0.628893 | TMEM59 | black | 0.122398188 | 0.543055598 | CDK6 | blue | -0.393984 | 0.046425 |
| KLHL6 | black | -0.097299 | 0.636316 | GUSBP11 | black | 0.125991223 | 0.531187237 | C10orf76 | blue | -0.377052 | 0.05759 |
| PRR14 | black | -0.095693 | 0.641922 | BUD13 | black | 0.12667146 | 0.528954297 | G2E3 | blue | -0.371928 | 0.061351 |
| LITAF | black | -0.095334 | 0.643179 | DCP2 | black | 0.129142061 | 0.520882233 | DDX41 | blue | -0.365466 | 0.066362 |
| C10orf12 | black | 0.09442 | 0.64638 | GOLGA7 | black | 0.132338638 | 0.510527392 | TMSB4X | blue | -0.361969 | 0.069203 |
| NEMF | black | -0.092928 | 0.651617 | SFMBT2 | black | 0.136795348 | 0.496261173 | ARHGEF6 | blue | -0.359869 | 0.070952 |
| PPM1D | black | 0.092837 | 0.651938 | ENPP4 | black | 0.137585288 | 0.493753529 | RAB9A | blue | -0.358619 | 0.07201 |
| GPBP1 | black | -0.092618 | 0.652708 | PIK3CG | black | 0.144053805 | 0.473461372 | TXNL4A | blue | -0.355978 | 0.074284 |
| SCAF4 | black | -0.091055 | 0.658216 | DLAT | black | 0.144216773 | 0.472955755 | LEPR | blue | -0.353519 | 0.07645 |
| LUC7L2 | black | -0.090288 | 0.660925 | TMLHE | black | 0.145146625 | 0.470076199 | ZNF25 | blue | -0.351876 | 0.077925 |
| FAM20B | black | 0.089406 | 0.664046 | CD164 | black | 0.145660453 | 0.468488889 | NPAT | blue | -0.351146 | 0.078587 |
| TFEC | black | -0.088559 | 0.667047 | UBASH3B | black | 0.146822249 | 0.464910179 | RRM1 | blue | -0.34923 | 0.080345 |
| HMGB3 | black | -0.087831 | 0.669634 | PRMT7 | black | 0.147532723 | 0.462728743 | LIN52 | blue | -0.335033 | 0.094313 |
| ZCCHC17 | black | 0.087546 | 0.670646 | PDE12 | black | 0.14902013 | 0.458179214 | TPM3 | blue | -0.333157 | 0.096287 |
| SFT2D1 | black | -0.084805 | 0.680415 | EIF5B | black | 0.152078767 | 0.448898247 | PPP5C | blue | -0.330087 | 0.099584 |
| PLIN3 | black | -0.083411 | 0.685399 | TRIM5 | black | 0.158912776 | 0.428528228 | CYFIP2 | blue | -0.324871 | 0.105377 |
| DGKH | black | -0.082176 | 0.689831 | NOD1 | black | 0.159170506 | 0.427770037 | SMARCC1 | blue | -0.3245 | 0.105798 |
| TMEM138 | black | -0.081812 | 0.691137 | SNW1 | black | 0.159375678 | 0.427166986 | SNHG4 | blue | -0.322084 | 0.108573 |
| RNASEH1 | black | -0.080515 | 0.695802 | ZBTB11 | black | 0.161087703 | 0.422153079 | RPS20 | blue | -0.319081 | 0.112097 |
| COA1 | black | 0.079834 | 0.698257 | RAB12 | black | 0.161904095 | 0.41977361 | DPP8 | blue | -0.318012 | 0.113372 |
| BACE1 | black | 0.078089 | 0.704559 | PDS5A | black | 0.165305799 | 0.409938974 | CUTA | blue | -0.315485 | 0.116427 |
| ZNF789 | black | 0.076971 | 0.708604 | MSTO1 | black | 0.165890878 | 0.408260517 | MTX3 | blue | -0.314614 | 0.117494 |
| POLR1C | black | -0.075983 | 0.712187 | FANCL | black | 0.169331889 | 0.398467116 | C16orf62 | blue | -0.313684 | 0.118641 |
| ATG14 | black | 0.075962 | 0.712266 | HARS2 | black | 0.169990466 | 0.39660802 | TMEM138 | blue | -0.312783 | 0.11976 |
| SZRD1 | black | -0.074174 | 0.718765 | BBS4 | black | 0.171856678 | 0.391366654 | BIVM | blue | -0.311718 | 0.121094 |
| SCAMP3 | black | 0.074105 | 0.719016 | SEC23A | black | 0.171948766 | 0.391109045 | OBFC1 | blue | -0.31035 | 0.122821 |
| SBNO1 | black | -0.074045 | 0.719236 | PRKDC | black | 0.174677422 | 0.383519808 | KPNA5 | blue | -0.310223 | 0.122983 |
| THUMPD1 | black | -0.072611 | 0.724461 | TUBGCP4 | black | 0.175122025 | 0.382291299 | HNRNPH1 | blue | -0.309989 | 0.12328 |
| MAML2 | black | 0.072129 | 0.726221 | C1D | black | 0.177491784 | 0.375781592 | HELQ | blue | -0.309161 | 0.124338 |
| C1orf112 | black | -0.072097 | 0.72634 | STK3 | black | 0.178065803 | 0.374214492 | SNX1 | blue | -0.307225 | 0.126836 |
| CIITA | black | 0.068904 | 0.738033 | POP5 | black | 0.181055983 | 0.366112687 | HNRNPA3 | blue | -0.30707 | 0.127038 |
| ADPGK | black | 0.068451 | 0.739696 | ZNF562 | black | 0.182298263 | 0.362777208 | DROSHA | blue | -0.303218 | 0.132123 |
| DRG1 | black | 0.068261 | 0.740394 | RRP1B | black | 0.183610404 | 0.359273622 | MOB1A | blue | -0.300621 | 0.135633 |
| PVRL2 | black | -0.067743 | 0.742296 | NIPBL | black | 0.18496969 | 0.355665276 | IDH2 | blue | -0.300451 | 0.135865 |
| TMEM127 | black | -0.065321 | 0.75122 | POLH | black | 0.187865348 | 0.348050367 | RSU1 | blue | -0.292174 | 0.147518 |
| MRPS31 | black | -0.064664 | 0.753648 | MTIF2 | black | 0.188291767 | 0.346937263 | B2M | blue | -0.29191 | 0.147901 |
| INO80 | black | 0.062671 | 0.761019 | ANAPC7 | black | 0.192100914 | 0.337088636 | WHAMMP1 | blue | -0.289912 | 0.150823 |
| NAB1 | black | -0.061261 | 0.766246 | FLJ23867 | black | 0.198802461 | 0.320176131 | ASCC1 | blue | -0.289894 | 0.15085 |
| GMEB2 | black | -0.059977 | 0.771017 | MBTD1 | black | 0.200163634 | 0.316805781 | SUCLA2 | blue | -0.288609 | 0.152751 |
| SCAF8 | black | -0.059274 | 0.773632 | RARS2 | black | 0.201727679 | 0.312960187 | NOL9 | blue | -0.287923 | 0.153772 |
| TSPAN3 | black | -0.058024 | 0.778289 | ABHD3 | black | 0.20373682 | 0.308062738 | PSMC2 | blue | -0.285997 | 0.156667 |
| CEP97 | black | 0.05693 | 0.782368 | MOSPD2 | black | 0.206588355 | 0.301194079 | CNOT8 | blue | -0.284473 | 0.158986 |
| DCLRE1C | black | -0.056905 | 0.782463 | FRY | black | 0.206647495 | 0.301052646 | TDRD3 | blue | -0.282011 | 0.162781 |
| SIAE | black | 0.056439 | 0.784202 | PION | black | 0.20716674 | 0.299812661 | ARL14EP | blue | -0.280415 | 0.165276 |
| ENO1 | black | 0.056194 | 0.785117 | DSN1 | black | 0.208359569 | 0.296976245 | GIMAP8 | blue | -0.279665 | 0.166458 |
| PIGN | black | -0.056052 | 0.785648 | XPO7 | black | 0.208369525 | 0.296952642 | IAH1 | blue | -0.279181 | 0.167223 |
| CSTF2T | black | 0.054996 | 0.789595 | ADO | black | 0.208966325 | 0.295539931 | LRRC8B | blue | -0.278369 | 0.168513 |
| USP24 | black | -0.054876 | 0.790044 | CREBRF | black | 0.209748887 | 0.293693906 | WDR5 | blue | -0.278145 | 0.168869 |
| DGAT1 | black | -0.05452 | 0.791376 | WWOX | black | 0.211767351 | 0.288966034 | PECAM1 | blue | -0.274431 | 0.174869 |
| CCDC85B | black | -0.053934 | 0.793572 | BTBD10 | black | 0.21219291 | 0.287975424 | RAB7A | blue | -0.274418 | 0.174889 |
| ARHGEF7 | black | 0.053719 | 0.794377 | PTGES3 | black | 0.213052374 | 0.285981331 | DCTN5 | blue | -0.274195 | 0.175254 |
| FYCO1 | black | -0.053633 | 0.794701 | DNAJA2 | black | 0.214292434 | 0.283119669 | SERAC1 | blue | -0.272548 | 0.177966 |
| KLHDC10 | black | -0.049894 | 0.808743 | CACUL1 | black | 0.214564759 | 0.282493678 | C21orf7 | blue | -0.271174 | 0.180249 |
| PRPF4B | black | -0.049796 | 0.80911 | PCF11 | black | 0.216805949 | 0.277375335 | ATG4C | blue | -0.270508 | 0.181363 |

**Table 6. Gene module membership and correlation with growth rate (48/72h), with adjusted p values for each module**

| | | | | | | | | | | | |
|---|---|---|---|---|---|---|---|---|---|---|---|
| IRF9 | black | 0.048648 | 0.813436 | ATP11B | black | 0.217971493 | 0.274737098 | CD302 | blue | -0.268669 | 0.184467 |
| RAD21 | black | -0.047818 | 0.816565 | MCPH1 | black | 0.221634789 | 0.266550168 | FPR3 | blue | -0.267818 | 0.185915 |
| HN1 | black | -0.047303 | 0.818507 | DPY19L1 | black | 0.222758302 | 0.264071195 | RBMX2 | blue | -0.267544 | 0.186381 |
| RPL23A | black | -0.047156 | 0.819064 | CDC42BPA | black | 0.226680933 | 0.255533424 | AURKAIP1 | blue | -0.267533 | 0.1864 |
| PDE3B | black | 0.045174 | 0.826551 | PLEKHA1 | black | 0.22695045 | 0.2549535 | TMEM242 | blue | -0.267452 | 0.186539 |
| PIGT | black | 0.042832 | 0.835418 | CHM | black | 0.233916177 | 0.240262937 | RBM23 | blue | -0.266153 | 0.188769 |
| UTP14C | black | -0.04105 | 0.842183 | ATXN3 | black | 0.235670453 | 0.23665336 | GRB2 | blue | -0.264707 | 0.191274 |
| KANSL1-AS1 | black | 0.04057 | 0.844003 | LEMD3 | black | 0.235868838 | 0.236247444 | LOC10013108 | blue | -0.264685 | 0.191312 |
| AK3 | black | 0.040034 | 0.84604 | FUT4 | black | 0.241312438 | 0.225289298 | HEATR5B | blue | -0.263399 | 0.193558 |
| TNPO3 | black | -0.037361 | 0.856213 | POLA2 | black | 0.24301752 | 0.221928175 | METTL10 | blue | -0.263257 | 0.193808 |
| EIF4EBP1 | black | 0.034119 | 0.868577 | FBXW2 | black | 0.24395001 | 0.220104347 | DERA | blue | -0.26315 | 0.193995 |
| EFR3A | black | -0.031638 | 0.878062 | STXBP3 | black | 0.244059901 | 0.219890082 | STK17A | blue | -0.262944 | 0.194358 |
| LOXL3 | black | -0.031151 | 0.879926 | PTCD3 | black | 0.244907509 | 0.218242134 | IRAK3 | blue | -0.262169 | 0.195724 |
| APOA1BP | black | -0.02901 | 0.888126 | FNDC3A | black | 0.247697862 | 0.212875963 | ARRDC3 | blue | -0.261605 | 0.196722 |
| SPIN1 | black | -0.028506 | 0.890057 | CASK | black | 0.252970187 | 0.202982357 | MORF4L1 | blue | -0.261338 | 0.197196 |
| C5orf43 | black | 0.028029 | 0.891887 | NDUFV2 | black | 0.254192125 | 0.200735005 | ERCC6L2 | blue | -0.260227 | 0.199178 |
| DARS | black | 0.026141 | 0.89913 | DHX15 | black | 0.257418586 | 0.194883079 | ARMC10 | blue | -0.260128 | 0.199354 |
| SRP72 | black | 0.025952 | 0.899855 | PDLIM5 | black | 0.257466987 | 0.194796196 | URM1 | blue | -0.259348 | 0.200754 |
| MKLN1 | black | 0.025842 | 0.90028 | FUT8 | black | 0.258788062 | 0.192435086 | PSMG3 | blue | -0.257173 | 0.204691 |
| CDC16 | black | 0.025624 | 0.901117 | FGD4 | black | 0.258834737 | 0.192352028 | MORC4 | blue | -0.256733 | 0.205495 |
| MKI67IP | black | 0.02479 | 0.904318 | SMG1 | black | 0.259578856 | 0.191031212 | REV3L | blue | -0.25571 | 0.20737 |
| USP16 | black | 0.024599 | 0.905055 | UBR3 | black | 0.259620335 | 0.190957771 | PCMT1 | blue | -0.255309 | 0.208107 |
| OSTC | black | -0.021655 | 0.916374 | RRAGB | black | 0.259703571 | 0.190810457 | ADRBK2 | blue | -0.255096 | 0.2085 |
| ZNF583 | black | 0.019952 | 0.922932 | ZNF277 | black | 0.263297967 | 0.184523661 | RDH10 | blue | -0.252725 | 0.212906 |
| BRD4 | black | 0.01975 | 0.923711 | SPTLC1 | black | 0.263339607 | 0.184451685 | CCDC43 | blue | -0.25209 | 0.214097 |
| DKC1 | black | 0.018994 | 0.92662 | CASP10 | black | 0.263868256 | 0.183539582 | CTSS | blue | -0.251253 | 0.215674 |
| MED14 | black | -0.01675 | 0.935272 | ORC5 | black | 0.264131993 | 0.183085718 | GPN3 | blue | -0.25104 | 0.216076 |
| KCMF1 | black | 0.015262 | 0.941012 | REXO1 | black | 0.265870122 | 0.180114097 | ZNHIT1 | blue | -0.250591 | 0.216926 |
| MTFMT | black | 0.015176 | 0.941342 | C12orf57 | black | 0.266162468 | 0.179617606 | ZBTB37 | blue | -0.245727 | 0.226277 |
| COX19 | black | -0.015086 | 0.941692 | SLC25A32 | black | 0.268130886 | 0.176299486 | C17orf89 | blue | -0.245531 | 0.22666 |
| SYPL1 | black | 0.013583 | 0.947492 | KCTD10 | black | 0.27102633 | 0.171496998 | ZNF131 | blue | -0.244973 | 0.227751 |
| USP14 | black | 0.009729 | 0.962379 | UBE2D2 | black | 0.271550925 | 0.170636818 | POLK | blue | -0.244692 | 0.228302 |
| EXOSC1 | black | -0.009428 | 0.963542 | SLC50A1 | black | 0.271621191 | 0.170521833 | TCP11L1 | blue | -0.243237 | 0.231166 |
| CCDC71 | black | -0.008199 | 0.968291 | KNTC1 | black | 0.273902338 | 0.166818463 | SNHG16 | blue | -0.24184 | 0.233941 |
| ABCB10 | black | -0.006498 | 0.974869 | TBC1D5 | black | 0.276491465 | 0.162684253 | HIST1H4D | blue | -0.241604 | 0.234412 |
| PHB2 | black | 0.006351 | 0.975438 | HSD17B7P2 | black | 0.276962786 | 0.161939541 | ERICH1 | blue | -0.240588 | 0.236445 |
| ZNF81 | black | -0.005738 | 0.977808 | CEP57L1 | black | 0.277053347 | 0.161796727 | EIF2B1 | blue | -0.240497 | 0.236628 |
| SMAP1 | black | -0.005626 | 0.978239 | SPPL2A | black | 0.27706758 | 0.161774289 | CHCHD6 | blue | -0.240125 | 0.237376 |
| ANO5 | black | -0.005474 | 0.978828 | PIK3CB | black | 0.277266595 | 0.161460788 | GLTSCR2 | blue | -0.239869 | 0.237892 |
| SMAD2 | black | 0.005349 | 0.979311 | LYPLA1 | black | 0.27843754 | 0.159624957 | SOCS4 | blue | -0.239001 | 0.239645 |
| GMEB1 | black | -0.003923 | 0.984827 | GMCL1 | black | 0.280453135 | 0.156499649 | MPHOSPH9 | blue | -0.238443 | 0.240777 |
| ZNF273 | black | -0.003371 | 0.986959 | C2orf43 | black | 0.281889665 | 0.154298962 | NDUFS8 | blue | -0.236574 | 0.244594 |
| EID1 | black | -0.002449 | 0.990527 | GPR89C | black | 0.282776167 | 0.152951948 | MITD1 | blue | -0.236461 | 0.244828 |
| AMBRA1 | black | 0.001558 | 0.993975 | ZNF408 | black | 0.283779291 | 0.151437876 | METTL23 | blue | -0.236113 | 0.245544 |
| NFAT5 | black | -0.001094 | 0.995766 | PTPLB | black | 0.28687263 | 0.146836431 | ERCC5 | blue | -0.235679 | 0.246436 |
| TMCO7 | black | 0.001087 | 0.995794 | GTF3A | black | 0.287041714 | 0.146587839 | RPP38 | blue | -0.234447 | 0.248987 |
| RPUSD3 | blue | -0.604371 | 0.001076 | MAP3K2 | black | 0.288273534 | 0.144785893 | CPM | blue | -0.233759 | 0.250419 |
| RAB9A | blue | -0.577678 | 0.001999 | POLA1 | black | 0.28928519 | 0.143317963 | RSL1D1 | blue | -0.233724 | 0.250492 |
| PAICS | blue | -0.540614 | 0.004353 | RPL4 | black | 0.297970795 | 0.131153435 | ALDH5A1 | blue | -0.233107 | 0.25178 |
| TPD52L2 | blue | 0.515194 | 0.007072 | YPEL5 | black | 0.299111337 | 0.129613698 | ZNF559 | blue | -0.232801 | 0.252421 |
| TULP4 | blue | 0.500264 | 0.009251 | HSPA4 | black | 0.299875743 | 0.128589145 | RPL24 | blue | -0.232559 | 0.25293 |
| IFIT3 | blue | -0.492271 | 0.01063 | RRNAD1 | black | 0.303779552 | 0.123448707 | RECQL | blue | -0.232512 | 0.253028 |
| NCKAP5L | blue | 0.480551 | 0.012959 | LOC374443 | black | 0.304582044 | 0.122410934 | NUP37 | blue | -0.232419 | 0.253222 |
| C16orf88 | blue | -0.467556 | 0.016019 | SMPD2 | black | 0.30483749 | 0.122081939 | DNAJC24 | blue | -0.232167 | 0.253753 |
| KL | blue | -0.465598 | 0.016528 | ITSN2 | black | 0.306881832 | 0.119472316 | WDR3 | blue | -0.23204 | 0.254019 |
| TSTA3 | blue | -0.464054 | 0.016938 | DENND4A | black | 0.307849299 | 0.118251733 | SELPLG | blue | -0.231504 | 0.25515 |

**Table 6. Gene module membership and correlation with growth rate (48/72h), with adjusted p values for each module**

| | | | | | | | | | | | |
|---|---|---|---|---|---|---|---|---|---|---|---|
| BUB1 | blue | -0.45608 | 0.019193 | PIGK | black | 0.308197101 | 0.117815185 | ARHGEF9 | blue | -0.230645 | 0.256966 |
| HIST2H2AC | blue | -0.450472 | 0.020922 | FXN | black | 0.308785008 | 0.117079968 | CCDC75 | blue | -0.229269 | 0.259894 |
| TFG | blue | -0.450422 | 0.020938 | WDPCP | black | 0.313590215 | 0.11119695 | PIGB | blue | -0.229243 | 0.259951 |
| TBCB | blue | -0.429052 | 0.028729 | GNB2L1 | black | 0.31731048 | 0.106794809 | TMA7 | blue | -0.229071 | 0.260318 |
| SNRNP25 | blue | -0.428866 | 0.028806 | LOC202181 | black | 0.319382084 | 0.104400388 | RAP1GDS1 | blue | -0.228659 | 0.2612 |
| C19orf60 | blue | -0.425415 | 0.030262 | ATL2 | black | 0.319789023 | 0.103934774 | CEP57 | blue | -0.228562 | 0.261407 |
| SLC22A5 | blue | -0.422727 | 0.031436 | ATIC | black | 0.32197581 | 0.10145916 | PPP1CC | blue | -0.226885 | 0.265018 |
| TCTN2 | blue | -0.418522 | 0.033347 | ATP5G2 | black | 0.324578309 | 0.098570657 | DUT | blue | -0.225923 | 0.267104 |
| TPM3 | blue | -0.415782 | 0.034641 | TPP2 | black | 0.326558082 | 0.096414911 | BBS9 | blue | -0.225816 | 0.267338 |
| IKZF2 | blue | 0.414097 | 0.035456 | WRB | black | 0.326716669 | 0.096243771 | DPY19L4 | blue | -0.225305 | 0.26845 |
| SLC1A5 | blue | 0.413698 | 0.035652 | CLEC2D | black | 0.328169969 | 0.094686045 | MMAB | blue | -0.225283 | 0.268499 |
| CDK12 | blue | 0.411161 | 0.036914 | SUV39H1 | black | 0.33223248 | 0.090432173 | ATE1 | blue | -0.224665 | 0.269849 |
| KLHL18 | blue | -0.410104 | 0.037451 | MEMO1 | black | 0.334772209 | 0.087847097 | PIK3C2A | blue | -0.224635 | 0.269914 |
| A2M | blue | 0.409871 | 0.037569 | C16orf52 | black | 0.338094665 | 0.084550224 | WDR92 | blue | -0.224352 | 0.270533 |
| NKAP | blue | -0.408847 | 0.038096 | LIMD1 | black | 0.339535591 | 0.083149953 | RAB11FIP4 | blue | -0.223887 | 0.271555 |
| C9orf85 | blue | -0.408105 | 0.038481 | RABIF | black | 0.340941596 | 0.081800678 | ZNF714 | blue | -0.223867 | 0.271599 |
| IPO9 | blue | -0.40775 | 0.038667 | SRSF7 | black | 0.343103184 | 0.079758892 | POC5 | blue | -0.223353 | 0.272729 |
| SRSF4 | blue | -0.404146 | 0.040591 | PSMC6 | black | 0.344198987 | 0.078738783 | PRDX2 | blue | -0.222495 | 0.274625 |
| ERCC5 | blue | -0.401917 | 0.041818 | DCSTAMP | black | 0.346136319 | 0.076959682 | PHKB | blue | -0.222461 | 0.274699 |
| COMMD9 | blue | -0.399475 | 0.043195 | SNAP23 | black | 0.348434383 | 0.074889355 | HLA-DRA | blue | -0.222054 | 0.275602 |
| ARPC3 | blue | -0.399414 | 0.04323 | PLEKHA8 | black | 0.349049831 | 0.07434221 | SP4 | blue | -0.222022 | 0.275672 |
| RP9 | blue | -0.396825 | 0.044731 | FBXO5 | black | 0.349178328 | 0.074228361 | WDR47 | blue | -0.222004 | 0.275711 |
| TRAF6 | blue | -0.39668 | 0.044816 | SCP2 | black | 0.351590877 | 0.072115562 | TIAL1 | blue | -0.221931 | 0.275874 |
| C16orf62 | blue | -0.394529 | 0.046096 | RAB22A | black | 0.354810949 | 0.069367933 | FADS1 | blue | -0.221765 | 0.276242 |
| MDM4 | blue | -0.392526 | 0.047313 | ZNF267 | black | 0.355297799 | 0.068959631 | BIRC6 | blue | -0.221691 | 0.276408 |
| GORAB | blue | -0.391156 | 0.04816 | PRPF39 | black | 0.360338331 | 0.064840401 | KIF3B | blue | -0.220867 | 0.278242 |
| CKS1B | blue | -0.389342 | 0.0493 | MGST2 | black | 0.362313543 | 0.063279187 | RPS3A | blue | -0.220754 | 0.278495 |
| PRPSAP1 | blue | -0.387987 | 0.050165 | MAP4K3 | black | 0.362868809 | 0.062845595 | DCUN1D4 | blue | -0.21941 | 0.281505 |
| ASCC1 | blue | -0.386772 | 0.050951 | MGA | black | 0.364795211 | 0.061359164 | MS4A14 | blue | -0.218429 | 0.283716 |
| PLEKHO2 | blue | 0.386737 | 0.050974 | GEN1 | black | 0.364953151 | 0.061238517 | FAM188A | blue | -0.218353 | 0.283888 |
| RNF14 | blue | -0.38662 | 0.05105 | PPWD1 | black | 0.36784943 | 0.05905864 | BCLAF1 | blue | -0.218288 | 0.284034 |
| C14orf80 | blue | 0.385545 | 0.051754 | HIBADH | black | 0.369900087 | 0.057552111 | FAM126B | blue | -0.21741 | 0.286024 |
| GUSBP3 | blue | -0.385118 | 0.052036 | SCARNA9 | black | 0.371514098 | 0.056387585 | LOC645513 | blue | -0.217132 | 0.286657 |
| C11orf96 | blue | -0.384007 | 0.052775 | SRSF6 | black | 0.371823369 | 0.056166559 | VCPIP1 | blue | -0.216642 | 0.287771 |
| ARL6IP5 | blue | -0.383263 | 0.053274 | ABHD12 | black | 0.377519806 | 0.052215245 | JAGN1 | blue | -0.215643 | 0.290054 |
| MRPL51 | blue | -0.383004 | 0.053449 | TPK1 | black | 0.378696475 | 0.051426924 | P4HA2 | blue | -0.21557 | 0.290222 |
| FRMD8 | blue | 0.382699 | 0.053656 | C1QBP | black | 0.382419242 | 0.048994226 | LYAR | blue | -0.215368 | 0.290685 |
| BSG | blue | 0.382067 | 0.054085 | RASGEF1B | black | 0.384012874 | 0.047980957 | SNRNP27 | blue | -0.214808 | 0.291973 |
| VPS51 | blue | -0.379261 | 0.056025 | PPFIA1 | black | 0.384297056 | 0.047802016 | DDX26B | blue | -0.214647 | 0.292343 |
| KPNA1 | blue | -0.378393 | 0.056636 | ZFAND4 | black | 0.386468508 | 0.046452023 | NOL10 | blue | -0.214492 | 0.2927 |
| NUP85 | blue | -0.378322 | 0.056687 | SPCS2 | black | 0.387975053 | 0.045533226 | UBA6 | blue | -0.212246 | 0.297903 |
| MRPS33 | blue | -0.377396 | 0.057345 | MAGOHB | black | 0.388572986 | 0.045172572 | CXorf21 | blue | -0.211105 | 0.300568 |
| RAPGEF6 | blue | 0.376347 | 0.058097 | PCYT1A | black | 0.388769863 | 0.045054318 | TMEM181 | blue | -0.211104 | 0.30057 |
| ECH1 | blue | -0.372789 | 0.060706 | NSMCE2 | black | 0.390857776 | 0.043815222 | ZNF615 | blue | -0.210164 | 0.302779 |
| LRRC37BP1 | blue | -0.372415 | 0.060986 | PCNXL4 | black | 0.391272415 | 0.043572393 | RPS27A | blue | -0.209926 | 0.303339 |
| TCEANC2 | blue | -0.36949 | 0.063206 | TMEM48 | black | 0.393636467 | 0.042208215 | LOC1005056 | blue | -0.209735 | 0.30379 |
| TLR5 | blue | -0.366214 | 0.065766 | TMX2 | black | 0.396902373 | 0.040379621 | ZRSR2 | blue | -0.209698 | 0.303876 |
| RSF1 | blue | -0.365803 | 0.066093 | UBE2D4 | black | 0.398345821 | 0.039591779 | RPS25 | blue | -0.208729 | 0.306168 |
| YPEL4 | blue | 0.364899 | 0.066817 | PSMD14 | black | 0.402684295 | 0.037297253 | OSBP | blue | -0.207969 | 0.307972 |
| ALG1 | blue | -0.360885 | 0.070102 | PSMB2 | black | 0.404246319 | 0.036497621 | CYSLTR1 | blue | -0.207656 | 0.308717 |
| VDR | blue | 0.356786 | 0.073583 | INPP5B | black | 0.405381358 | 0.035925215 | PRKCB | blue | -0.206842 | 0.31066 |
| BAG3 | blue | 0.353868 | 0.076141 | SAP30BP | black | 0.413557184 | 0.032011077 | CCNL2 | blue | -0.20666 | 0.311094 |
| STX2 | blue | -0.353232 | 0.076707 | PSMB6 | black | 0.418359681 | 0.02987676 | ZNF570 | blue | -0.206459 | 0.311577 |
| UTRN | blue | -0.353184 | 0.076749 | KIAA1430 | black | 0.420146726 | 0.029112491 | NOL8 | blue | -0.206202 | 0.312192 |
| S100A9 | blue | -0.352901 | 0.077002 | AKIRIN1 | black | 0.427986426 | 0.025943947 | SNRPD3 | blue | -0.205925 | 0.312858 |
| SF3B4 | blue | 0.35226 | 0.077578 | ZNF37A | black | 0.428401376 | 0.025784367 | B3GALNT2 | blue | -0.205048 | 0.314968 |

**Table 6. Gene module membership and correlation with growth rate (48/72h), with adjusted p values for each module**

| | | | | | | | | | | | |
|---|---|---|---|---|---|---|---|---|---|---|---|
| TMA7 | blue | -0.351239 | 0.078502 | COPZ1 | black | 0.435353713 | 0.023226998 | C4orf33 | blue | -0.20483 | 0.315494 |
| LAIR1 | blue | -0.349537 | 0.080061 | SNX5 | black | 0.446950618 | 0.019423066 | TMTC3 | blue | -0.204509 | 0.316269 |
| MRPL3 | blue | -0.349179 | 0.080391 | POLR2H | black | 0.447562016 | 0.019237632 | WDR61 | blue | -0.204501 | 0.31629 |
| EXD2 | blue | -0.348119 | 0.081377 | MRPS18B | black | 0.4581508 | 0.0162495 | TBC1D15 | blue | -0.204065 | 0.317345 |
| LIMA1 | blue | -0.347538 | 0.081921 | IGIP | black | 0.464826396 | 0.014570474 | CLEC4A | blue | -0.203316 | 0.319163 |
| PPP1CC | blue | -0.347506 | 0.081952 | NDUFAF4 | black | 0.472305539 | 0.012862391 | ZNF568 | blue | -0.202728 | 0.320595 |
| PLXNA3 | blue | 0.344796 | 0.084526 | ZBED6 | black | 0.477209855 | 0.011835234 | DNAJC5 | blue | -0.201256 | 0.324196 |
| PIGS | blue | -0.344177 | 0.085123 | MRPS15 | black | 0.49628375 | 0.008465144 | GGPS1 | blue | -0.200454 | 0.326168 |
| S1PR4 | blue | -0.34244 | 0.086814 | ARL1 | black | 0.499943701 | 0.007920875 | LACC1 | blue | -0.200273 | 0.326614 |
| BTG3 | blue | -0.341587 | 0.087654 | ZNF138 | black | 0.521347377 | 0.005292141 | ZW10 | blue | -0.200133 | 0.326958 |
| PHLDA3 | blue | -0.340619 | 0.088614 | GEMIN2 | black | 0.54280014 | 0.003439666 | ZZZ3 | blue | -0.199222 | 0.329211 |
| NME6 | blue | 0.340004 | 0.089229 | NOP10 | black | 0.569852023 | 0.001916123 | C20orf111 | blue | -0.197015 | 0.334707 |
| TMEM106E | blue | -0.339837 | 0.089397 | MICAL2 | blue | -0.687274515 | 7.4878E-05 | CDC5L | blue | -0.195797 | 0.337764 |
| HEATR5B | blue | -0.339449 | 0.089786 | WDR47 | blue | -0.647687929 | 0.000259842 | PES1 | blue | -0.19521 | 0.339241 |
| DUS3L | blue | 0.338552 | 0.090692 | ZCCHC6 | blue | -0.643056872 | 0.000297157 | CRBN | blue | -0.194458 | 0.341143 |
| SNUPN | blue | -0.337559 | 0.091703 | IFNGR2 | blue | -0.597475501 | 0.000999264 | PFDN6 | blue | -0.194223 | 0.341739 |
| ULBP2 | blue | -0.335842 | 0.09347 | ZNF786 | blue | -0.593501594 | 0.001101329 | SHQ1 | blue | -0.193623 | 0.34326 |
| PWP2 | blue | -0.335666 | 0.093654 | GMPPB | blue | -0.592760856 | 0.001121321 | RETN | blue | -0.193365 | 0.343916 |
| PMS1 | blue | -0.334009 | 0.095387 | MAPKAP1 | blue | -0.585284999 | 0.001341355 | RUFY1 | blue | -0.193036 | 0.344754 |
| WDR34 | blue | -0.332313 | 0.097185 | ZNF354B | blue | -0.574711054 | 0.001715829 | RAB11FIP2 | blue | -0.192792 | 0.345375 |
| ZNF131 | blue | -0.332173 | 0.097334 | LOC1005067 | blue | -0.574378361 | 0.001728942 | ARHGAP10 | blue | -0.19245 | 0.346249 |
| RDH11 | blue | -0.331797 | 0.097738 | DLG1 | blue | -0.573583264 | 0.001760632 | GTPBP3 | blue | -0.192245 | 0.346773 |
| UBASH3B | blue | 0.331725 | 0.097814 | CNOT8 | blue | -0.570815415 | 0.001874886 | ELF1 | blue | -0.192033 | 0.347316 |
| JPH4 | blue | 0.331283 | 0.098289 | MRPL45 | blue | -0.56473042 | 0.002148713 | THEM4 | blue | -0.1914 | 0.348937 |
| REL | blue | -0.330525 | 0.099108 | C3orf62 | blue | -0.561540208 | 0.00230553 | EPAS1 | blue | -0.191326 | 0.349126 |
| SSBP1 | blue | -0.328405 | 0.101425 | SLC36A1 | blue | -0.561373066 | 0.00231401 | REEP3 | blue | -0.190852 | 0.350345 |
| CENPL | blue | -0.326163 | 0.103919 | SECISBP2 | blue | -0.557236657 | 0.002532573 | TCAIM | blue | -0.190711 | 0.350708 |
| ADRBK2 | blue | -0.326074 | 0.10402 | ARL6IP1 | blue | -0.556627003 | 0.002566237 | VPS26B | blue | -0.19009 | 0.352308 |
| ETNK1 | blue | 0.325811 | 0.104315 | FBXO38 | blue | -0.556313311 | 0.002583707 | PAK2 | blue | -0.190085 | 0.35232 |
| SPPL3 | blue | 0.324884 | 0.105363 | LACTB2 | blue | -0.551765832 | 0.002848657 | ERCC3 | blue | -0.189865 | 0.35289 |
| GGA3 | blue | -0.323813 | 0.106583 | APEX2 | blue | -0.549791587 | 0.002970726 | MAP3K3 | blue | -0.189652 | 0.353439 |
| CDK2AP1 | blue | -0.322842 | 0.107697 | ARFRP1 | blue | -0.548460201 | 0.003055549 | TAPT1 | blue | -0.188764 | 0.35574 |
| NLE1 | blue | -0.320136 | 0.11085 | F8A1 | blue | -0.547579682 | 0.003112778 | CLPX | blue | -0.188216 | 0.357164 |
| LOC493754 | blue | 0.318345 | 0.112974 | RPL35A | blue | -0.545445655 | 0.003255291 | MPEG1 | blue | -0.188111 | 0.357437 |
| TMEM97 | blue | -0.318145 | 0.113213 | PPP2CA | blue | -0.54476752 | 0.003301728 | PMP22 | blue | -0.187772 | 0.358319 |
| COX5A | blue | -0.318008 | 0.113377 | RPL8 | blue | -0.544115359 | 0.00334692 | FIGNL1 | blue | -0.187009 | 0.360312 |
| CLDND1 | blue | -0.317878 | 0.113533 | TPRKB | blue | -0.544098244 | 0.003348113 | ATRX | blue | -0.186885 | 0.360635 |
| ZBTB10 | blue | -0.316656 | 0.115004 | TRIT1 | blue | -0.540867364 | 0.003579935 | DDHD2 | blue | -0.186317 | 0.362123 |
| SRD5A3 | blue | -0.315305 | 0.116647 | MRPL38 | blue | -0.540218206 | 0.003628129 | SRP68 | blue | -0.186105 | 0.362679 |
| CFDP1 | blue | -0.314053 | 0.118185 | PPP2R3C | blue | -0.53949058 | 0.003682804 | TAF3 | blue | -0.185903 | 0.363211 |
| RHOBTB2 | blue | 0.31365 | 0.118683 | KLHL18 | blue | -0.538970911 | 0.003722281 | BCOR | blue | -0.18563 | 0.363927 |
| ADPRM | blue | -0.312181 | 0.120513 | SDCBP | blue | -0.538307125 | 0.003773231 | SPATA2L | blue | -0.1854 | 0.364533 |
| TSSC1 | blue | -0.311713 | 0.1211 | SKI | blue | -0.536568068 | 0.00390954 | C11orf24 | blue | -0.185217 | 0.365015 |
| FAM50A | blue | -0.311682 | 0.121138 | MTAP | blue | -0.53437095 | 0.004087729 | LIX1L | blue | -0.185098 | 0.365329 |
| SRPK1 | blue | -0.311007 | 0.121989 | LMO4 | blue | -0.534248047 | 0.004097897 | LRIG1 | blue | -0.185075 | 0.365388 |
| RINT1 | blue | -0.310416 | 0.122737 | ATRX | blue | -0.532721462 | 0.004226003 | BLOC1S1 | blue | -0.184968 | 0.365671 |
| PDLIM7 | blue | 0.310223 | 0.122983 | CSNK1G1 | blue | -0.532133355 | 0.004276256 | PRRG1 | blue | -0.184296 | 0.367448 |
| TCEAL8 | blue | -0.309864 | 0.12344 | GOLGA2 | blue | -0.531464273 | 0.004334045 | DTYMK | blue | -0.184092 | 0.367987 |
| DAZAP2 | blue | -0.308689 | 0.124944 | C22orf25 | blue | -0.531071899 | 0.004368242 | PLCL2 | blue | -0.183296 | 0.370098 |
| DR1 | blue | -0.30867 | 0.124968 | DDX19A | blue | -0.527958607 | 0.004647799 | TMX4 | blue | -0.182897 | 0.371158 |
| CCDC91 | blue | -0.307586 | 0.126368 | MXD1 | blue | -0.525531473 | 0.004876147 | TRAF6 | blue | -0.182551 | 0.37208 |
| ARV1 | blue | -0.307048 | 0.127066 | MORF4L2 | blue | -0.521703782 | 0.005255573 | C3orf62 | blue | -0.182125 | 0.373215 |
| CLIP4 | blue | -0.306878 | 0.127288 | HPRT1 | blue | -0.520382915 | 0.005392179 | CCDC149 | blue | -0.18155 | 0.374751 |
| FAM174A | blue | -0.30655 | 0.127716 | PSMD1 | blue | -0.519360684 | 0.005499952 | LTN1 | blue | -0.181546 | 0.374763 |
| FBXO11 | blue | -0.304876 | 0.129916 | TFCP2 | blue | -0.517901488 | 0.005656948 | DIMT1 | blue | -0.181307 | 0.375403 |
| ETF1 | blue | -0.3048 | 0.130017 | SERPINB9 | blue | -0.516089499 | 0.005857162 | SEC31A | blue | -0.181139 | 0.375852 |

**Table 6. Gene module membership and correlation with growth rate (48/72h), with adjusted p values for each module**

| | | | | | | | | | | | |
|---|---|---|---|---|---|---|---|---|---|---|---|
| TBC1D13 | blue | -0.304489 | 0.130429 | PPIL2 | blue | -0.51394946 | 0.006101306 | PDRG1 | blue | -0.181104 | 0.375945 |
| PRRG1 | blue | -0.304105 | 0.130938 | MTCH1 | blue | -0.512402223 | 0.006283125 | HNRNPK | blue | -0.180031 | 0.37883 |
| LZTS2 | blue | 0.304076 | 0.130977 | RER1 | blue | -0.512315127 | 0.006293495 | UQCRQ | blue | -0.179984 | 0.378956 |
| XRCC5 | blue | -0.303975 | 0.131112 | OSTM1 | blue | -0.511046814 | 0.00644614 | VRK2 | blue | -0.179973 | 0.378984 |
| ATP1B3 | blue | 0.303761 | 0.131397 | ATM | blue | -0.510546525 | 0.006507205 | ITCH | blue | -0.179971 | 0.378991 |
| MKS1 | blue | -0.303574 | 0.131647 | AKAP13 | blue | -0.509649168 | 0.006617957 | CHKB | blue | -0.179927 | 0.379108 |
| SQSTM1 | blue | 0.301966 | 0.133806 | MB21D1 | blue | -0.509593442 | 0.006624887 | TLR1 | blue | -0.179706 | 0.379704 |
| SDC4 | blue | 0.301956 | 0.133821 | MLYCD | blue | -0.508590578 | 0.006750648 | TUBD1 | blue | -0.177541 | 0.385566 |
| POLR2M | blue | -0.301243 | 0.134787 | SLC39A6 | blue | -0.507574045 | 0.00688017 | SNRPD1 | blue | -0.17754 | 0.38557 |
| ZNF783 | blue | 0.30104 | 0.135063 | STX3 | blue | -0.50716038 | 0.006933473 | PBRM1 | blue | -0.177056 | 0.386888 |
| CCDC6 | blue | -0.298765 | 0.138183 | DENND1B | blue | -0.505920407 | 0.00709534 | FTL | blue | -0.175425 | 0.391345 |
| SP140L | blue | -0.29815 | 0.139036 | SLC35D2 | blue | -0.505016112 | 0.007215384 | EEA1 | blue | -0.175312 | 0.391657 |
| LYRM4 | blue | -0.297603 | 0.139797 | USP1 | blue | -0.504408244 | 0.007297034 | ESYT2 | blue | -0.175281 | 0.391742 |
| RUFY2 | blue | -0.297446 | 0.140016 | IP6K2 | blue | -0.503037032 | 0.007484076 | ATL1 | blue | -0.174386 | 0.394203 |
| FMNL3 | blue | 0.297211 | 0.140344 | LAMC1 | blue | -0.502851334 | 0.007509713 | ZNF143 | blue | -0.174128 | 0.394914 |
| C6orf70 | blue | -0.296698 | 0.141064 | SNX19 | blue | -0.502730391 | 0.007526451 | AP1G2 | blue | -0.174 | 0.395266 |
| ICMT | blue | -0.294677 | 0.143921 | GLTP | blue | -0.502574875 | 0.007548018 | PTAR1 | blue | -0.173462 | 0.396751 |
| THUMPD3 | blue | -0.294514 | 0.144154 | FEM1A | blue | -0.500793875 | 0.007798744 | ZUFSP | blue | -0.172293 | 0.399991 |
| AKT1S1 | blue | -0.293368 | 0.145794 | BRWD3 | blue | -0.500747429 | 0.007805375 | GOLGB1 | blue | -0.172007 | 0.400783 |
| RAB14 | blue | -0.293224 | 0.146002 | VPS37A | blue | -0.49899394 | 0.008059208 | ODF2 | blue | -0.171867 | 0.401174 |
| FAM102B | blue | -0.293211 | 0.146019 | GLIPR1 | blue | -0.496272355 | 0.008466887 | PCBD2 | blue | -0.171668 | 0.401726 |
| MAP3K5 | blue | -0.291974 | 0.147808 | IL1RAP | blue | -0.495298218 | 0.008616947 | ZNF320 | blue | -0.171618 | 0.401866 |
| SUPT6H | blue | 0.291925 | 0.147879 | PIGX | blue | -0.493520033 | 0.008896617 | AP3M1 | blue | -0.171213 | 0.402996 |
| ANKRD54 | blue | 0.290834 | 0.14947 | CCNL2 | blue | -0.492042886 | 0.009134684 | HSBP1 | blue | -0.170942 | 0.40375 |
| PANX1 | blue | -0.290828 | 0.149478 | GK5 | blue | -0.491733318 | 0.009185246 | GSTM4 | blue | -0.170881 | 0.403922 |
| PHKB | blue | -0.290809 | 0.149505 | TTPAL | blue | -0.488691928 | 0.009694575 | SFMBT1 | blue | -0.169505 | 0.407775 |
| PPP2R5A | blue | -0.289868 | 0.150888 | HSPH1 | blue | -0.486639412 | 0.010051473 | WDR44 | blue | -0.169206 | 0.408615 |
| ATPIF1 | blue | -0.289197 | 0.151878 | SAE1 | blue | -0.485648688 | 0.01022763 | CNOT4 | blue | -0.169117 | 0.408864 |
| NDUFV1 | blue | 0.287647 | 0.154185 | DERL2 | blue | -0.484444558 | 0.010445194 | MAP3K1 | blue | -0.168904 | 0.409464 |
| PLA2G12A | blue | -0.28673 | 0.155562 | SMUG1 | blue | -0.482950799 | 0.010720443 | LOC150381 | blue | -0.168859 | 0.40959 |
| PGM1 | blue | -0.285088 | 0.158047 | IL2RG | blue | -0.482469097 | 0.010810483 | RFC1 | blue | -0.16758 | 0.413198 |
| CPT2 | blue | -0.284393 | 0.159108 | PRDX3 | blue | -0.482232173 | 0.010854999 | SPEN | blue | -0.167304 | 0.413979 |
| ARHGEF6 | blue | -0.283257 | 0.160852 | RERE | blue | -0.480237417 | 0.011235898 | ITGB5 | blue | -0.165519 | 0.419049 |
| APTX | blue | -0.282706 | 0.161703 | ANKRD32 | blue | -0.480088249 | 0.011264823 | LETMD1 | blue | -0.165474 | 0.419178 |
| XPO1 | blue | -0.282439 | 0.162117 | PDHB | blue | -0.479293946 | 0.011419893 | NET1 | blue | -0.165443 | 0.419265 |
| LILRB5 | blue | -0.282359 | 0.16224 | TMEM87B | blue | -0.479061977 | 0.011465514 | ACLY | blue | -0.16502 | 0.420471 |
| LOC729852 | blue | -0.280992 | 0.164371 | P2RY1 | blue | -0.478867667 | 0.011503845 | MCM8 | blue | -0.164769 | 0.421188 |
| IRAK3 | blue | -0.28072 | 0.164797 | CHPF2 | blue | -0.478216915 | 0.011632994 | ATPIF1 | blue | -0.164341 | 0.422413 |
| NCBP1 | blue | -0.280238 | 0.165554 | TMEM159 | blue | -0.477536708 | 0.011769277 | HMGN4 | blue | -0.164165 | 0.422916 |
| VDAC3 | blue | -0.279357 | 0.166944 | RPL39 | blue | -0.477193265 | 0.01183859 | INPP5A | blue | -0.163303 | 0.425391 |
| USP53 | blue | -0.278112 | 0.168923 | LRRC41 | blue | -0.476230982 | 0.012034607 | GPATCH8 | blue | -0.162768 | 0.426928 |
| FXC1 | blue | 0.276634 | 0.171292 | ANXA2 | blue | -0.476097769 | 0.012061953 | FLYWCH2 | blue | -0.162114 | 0.428813 |
| TEP1 | blue | -0.274937 | 0.174042 | MRPS30 | blue | -0.475886725 | 0.012105384 | TRAP1 | blue | -0.161975 | 0.429214 |
| EMC7 | blue | -0.27488 | 0.174136 | DENND1A | blue | -0.475850652 | 0.01211282 | CYB5R4 | blue | -0.161838 | 0.429612 |
| IL13RA1 | blue | -0.274722 | 0.174392 | NBPF1 | blue | -0.475671184 | 0.012149873 | CDCA7L | blue | -0.161522 | 0.430525 |
| TMEM14B | blue | -0.274663 | 0.17449 | TIMM44 | blue | -0.475575036 | 0.012169763 | FBXO8 | blue | -0.160873 | 0.432403 |
| NIT2 | blue | -0.274299 | 0.175085 | VRK1 | blue | -0.47556919 | 0.012170973 | SNRPB2 | blue | -0.160746 | 0.432771 |
| RILP | blue | -0.27363 | 0.176181 | CLDND1 | blue | -0.475122746 | 0.012263689 | RTN4 | blue | -0.160371 | 0.43386 |
| RUSC2 | blue | 0.273251 | 0.176804 | PAPD4 | blue | -0.473482188 | 0.012609451 | PINX1 | blue | -0.159107 | 0.43754 |
| SEC61G | blue | -0.273188 | 0.176909 | PAIP1 | blue | -0.472683675 | 0.012780649 | MLH3 | blue | -0.158736 | 0.438622 |
| UBAP1 | blue | -0.272861 | 0.177448 | FEZ2 | blue | -0.471755788 | 0.012982004 | SETD7 | blue | -0.158009 | 0.440751 |
| KSR1 | blue | 0.271592 | 0.179552 | ETF1 | blue | -0.468607831 | 0.013684833 | GEMIN4 | blue | -0.157805 | 0.441348 |
| AMN1 | blue | -0.271458 | 0.179777 | RALA | blue | -0.468408117 | 0.013730464 | LPPR2 | blue | -0.157625 | 0.441876 |
| DNAJC16 | blue | -0.271047 | 0.180462 | COPB2 | blue | -0.468147598 | 0.013790179 | ZNF561 | blue | -0.15754 | 0.442124 |
| SNORA14B | blue | -0.270672 | 0.181089 | IMMT | blue | -0.468046683 | 0.013813367 | UBE3A | blue | -0.157236 | 0.443016 |
| FASTKD1 | blue | -0.270538 | 0.181314 | GRINA | blue | -0.467118674 | 0.014028127 | PLCL1 | blue | -0.156604 | 0.444876 |

**Table 6. Gene module membership and correlation with growth rate (48/72h), with adjusted p values for each module**

| | | | |
|---|---|---|---|
| NUDT22 | blue | -0.269641 | 0.182821 |
| ST3GAL2 | blue | 0.268443 | 0.18485 |
| EPS15 | blue | -0.268344 | 0.185018 |
| CREBBP | blue | 0.266932 | 0.18743 |
| GEMIN8 | blue | -0.266078 | 0.188899 |
| SNRPD1 | blue | -0.265921 | 0.18917 |
| HDHD3 | blue | -0.265706 | 0.189541 |
| FAM73B | blue | 0.265688 | 0.189572 |
| C11orf21 | blue | 0.265515 | 0.189872 |
| AIDA | blue | -0.264957 | 0.190839 |
| ATP6V1H | blue | -0.264727 | 0.191239 |
| CEP41 | blue | -0.26415 | 0.192245 |
| GCSH | blue | -0.263727 | 0.192985 |
| VAMP3 | blue | -0.263672 | 0.19308 |
| UBR7 | blue | -0.263026 | 0.194214 |
| NDUFA9 | blue | -0.262338 | 0.195425 |
| S100A8 | blue | -0.26134 | 0.197192 |
| ECT2 | blue | -0.260267 | 0.199106 |
| CHFR | blue | 0.259885 | 0.199789 |
| NSL1 | blue | -0.259738 | 0.200053 |
| RBM45 | blue | -0.258643 | 0.202024 |
| PTPN23 | blue | 0.258244 | 0.202745 |
| SUMO1P3 | blue | -0.257291 | 0.204476 |
| YLPM1 | blue | -0.257215 | 0.204614 |
| POLG | blue | 0.256562 | 0.205806 |
| GALT | blue | -0.255622 | 0.20753 |
| HAUS8 | blue | -0.255555 | 0.207654 |
| ALDH3B1 | blue | 0.255512 | 0.207734 |
| PAQR5 | blue | 0.255429 | 0.207886 |
| RNASEL | blue | -0.255112 | 0.208471 |
| GGPS1 | blue | -0.254345 | 0.20989 |
| JKAMP | blue | -0.252944 | 0.212497 |
| RNF5 | blue | -0.251905 | 0.214444 |
| CIC | blue | 0.25137 | 0.215452 |
| LCP1 | blue | -0.251321 | 0.215546 |
| RMND5B | blue | -0.249216 | 0.219543 |
| CNOT10 | blue | -0.248794 | 0.22035 |
| ACVR1B | blue | -0.248686 | 0.220557 |
| SAP18 | blue | -0.24813 | 0.221624 |
| ATP8A1 | blue | -0.247986 | 0.221901 |
| ACOT13 | blue | -0.247578 | 0.222688 |
| CDK7 | blue | -0.247472 | 0.222892 |
| RING1 | blue | 0.247231 | 0.223357 |
| RB1CC1 | blue | -0.247222 | 0.223375 |
| ZNF101 | blue | -0.246965 | 0.223872 |
| KRTCAP2 | blue | -0.246034 | 0.225678 |
| EEFSEC | blue | -0.245813 | 0.226109 |
| BPGM | blue | -0.245351 | 0.227012 |
| CCDC149 | blue | -0.245175 | 0.227356 |
| TRMT61B | blue | -0.244582 | 0.228517 |
| SLC36A4 | blue | -0.244545 | 0.22859 |
| RACGAP1 | blue | -0.244355 | 0.228963 |
| ITGB3BP | blue | -0.244221 | 0.229226 |
| IRF2 | blue | -0.244153 | 0.229359 |
| INHBA | blue | 0.24297 | 0.231696 |
| PSMB9 | blue | -0.242228 | 0.233168 |
| TRAPPC3 | blue | -0.466947555 | 0.014068027 |
| RHBDF2 | blue | -0.466850233 | 0.014090762 |
| ARL4A | blue | -0.466008505 | 0.014288665 |
| OCRL | blue | -0.465831713 | 0.014330522 |
| MDM4 | blue | -0.465328229 | 0.014450283 |
| SLC35D1 | blue | -0.465081848 | 0.01450919 |
| HAVCR2 | blue | -0.464157805 | 0.014731887 |
| FAM96A | blue | -0.462336942 | 0.015178994 |
| DPP8 | blue | -0.46068384 | 0.015594556 |
| ACOX1 | blue | -0.460632627 | 0.015607578 |
| SECISBP2L | blue | -0.460584971 | 0.015619704 |
| PITRM1 | blue | -0.458852065 | 0.016065952 |
| ZW10 | blue | -0.457155132 | 0.016513071 |
| KDM5C | blue | -0.45602569 | 0.016816306 |
| COL4A3BP | blue | -0.454811734 | 0.017147328 |
| SNX2 | blue | -0.454808942 | 0.017148096 |
| PDCD2 | blue | -0.454535796 | 0.017223314 |
| ATP6V0A2 | blue | -0.454530085 | 0.01722489 |
| FECH | blue | -0.453454944 | 0.017523624 |
| ST6GALNAC | blue | -0.452644481 | 0.017751621 |
| CUL9 | blue | -0.451410256 | 0.018103508 |
| PET112 | blue | -0.450862232 | 0.01826158 |
| STX2 | blue | -0.450642147 | 0.01832538 |
| HSP90AA1 | blue | -0.450420128 | 0.018389925 |
| NLK | blue | -0.447923604 | 0.019128649 |
| OXR1 | blue | -0.447731827 | 0.019186388 |
| VTA1 | blue | -0.447305689 | 0.019315197 |
| TMEM194A | blue | -0.446798767 | 0.019469348 |
| GMPPA | blue | -0.446209671 | 0.019649752 |
| RFC1 | blue | -0.442350531 | 0.02086573 |
| PSMG2 | blue | -0.44131276 | 0.021203009 |
| TMEM248 | blue | -0.441163567 | 0.021251862 |
| LRP12 | blue | -0.440818753 | 0.021365121 |
| GCLC | blue | -0.440140761 | 0.021589255 |
| TMEM5 | blue | -0.439008089 | 0.02196798 |
| ATG4C | blue | -0.438620085 | 0.022098954 |
| TLR1 | blue | -0.437821519 | 0.022370523 |
| SELT | blue | -0.437109548 | 0.022614932 |
| UCHL3 | blue | -0.437016878 | 0.022646904 |
| RBM26 | blue | -0.436825133 | 0.022713174 |
| TRAF5 | blue | -0.43297302 | 0.024078359 |
| ZFP90 | blue | -0.431532032 | 0.024605885 |
| C12orf5 | blue | -0.43099101 | 0.024806353 |
| TPI1 | blue | -0.430195495 | 0.025103529 |
| TNFRSF1A | blue | -0.428634526 | 0.025695053 |
| RAN | blue | -0.428155924 | 0.025878665 |
| AP1G2 | blue | -0.427770958 | 0.026027126 |
| SLC9B2 | blue | -0.427397046 | 0.026171984 |
| SERPINB8 | blue | -0.427295852 | 0.0262113 |
| ZNF354A | blue | -0.426938906 | 0.026350363 |
| TTC37 | blue | -0.426347397 | 0.026582126 |
| TAF4 | blue | -0.426233146 | 0.026627082 |
| SCLY | blue | -0.42577687 | 0.02680723 |
| MSANTD2 | blue | -0.425523698 | 0.026907614 |
| RSL24D1 | blue | -0.425417274 | 0.026949902 |
| CCNG2 | blue | -0.425225683 | 0.027026168 |
| PPIL4 | blue | -0.156065 | 0.446464 |
| WWP1 | blue | -0.156021 | 0.446592 |
| CXCR4 | blue | -0.155921 | 0.446888 |
| UBE2E3 | blue | -0.155481 | 0.448186 |
| ERAL1 | blue | -0.155358 | 0.44855 |
| CNOT1 | blue | -0.154353 | 0.451528 |
| KDM4A | blue | -0.153895 | 0.452889 |
| GNG5 | blue | -0.153475 | 0.454137 |
| CMTM3 | blue | -0.153314 | 0.454615 |
| LOC654433 | blue | -0.152749 | 0.456301 |
| MLEC | blue | -0.152619 | 0.456687 |
| CPNE8 | blue | -0.152228 | 0.457855 |
| ANKRD44 | blue | -0.152064 | 0.458347 |
| LARP7 | blue | -0.150858 | 0.461962 |
| KIAA0317 | blue | -0.150616 | 0.462688 |
| CHP1 | blue | -0.150464 | 0.463145 |
| AAMDC | blue | -0.149587 | 0.465785 |
| PPP2R3A | blue | -0.14904 | 0.467435 |
| ASH2L | blue | -0.148064 | 0.470391 |
| WHSC1L1 | blue | -0.14766 | 0.471615 |
| C1orf43 | blue | -0.147435 | 0.472298 |
| CHMP3 | blue | -0.146675 | 0.474609 |
| MERTK | blue | -0.146404 | 0.475437 |
| PI4K2B | blue | -0.146171 | 0.476145 |
| RBM39 | blue | -0.146111 | 0.47633 |
| ACY1 | blue | -0.146054 | 0.476503 |
| FAM126A | blue | -0.14603 | 0.476578 |
| DOM3Z | blue | -0.145577 | 0.47796 |
| PTGER2 | blue | -0.145443 | 0.478369 |
| WDSUB1 | blue | -0.145358 | 0.47863 |
| METTL14 | blue | -0.14517 | 0.479206 |
| SASH3 | blue | -0.144966 | 0.479829 |
| NSUN3 | blue | -0.144563 | 0.481063 |
| GIGYF2 | blue | -0.144241 | 0.482053 |
| KIAA1009 | blue | -0.144051 | 0.482635 |
| ZNF788 | blue | -0.143819 | 0.483349 |
| PGGT1B | blue | -0.142786 | 0.486528 |
| MKS1 | blue | -0.141504 | 0.490492 |
| TBL1XR1 | blue | -0.141285 | 0.491169 |
| GLOD4 | blue | -0.139594 | 0.496424 |
| TMEM106B | blue | -0.139456 | 0.496855 |
| FAM190B | blue | -0.139307 | 0.497318 |
| SMARCC2 | blue | -0.138806 | 0.49888 |
| SLC2A13 | blue | -0.138741 | 0.499083 |
| PPID | blue | -0.138494 | 0.499857 |
| ARL13B | blue | -0.137939 | 0.501593 |
| USP42 | blue | -0.135884 | 0.508048 |
| SPATA20 | blue | -0.135555 | 0.509087 |
| SMA5 | blue | -0.135087 | 0.510562 |
| USP33 | blue | -0.135004 | 0.510824 |
| NEK9 | blue | -0.133736 | 0.514838 |
| MIER3 | blue | -0.13283 | 0.517717 |
| CLIP4 | blue | -0.13262 | 0.518386 |
| RHOB | blue | -0.132418 | 0.51903 |
| NDNL2 | blue | -0.132406 | 0.519066 |
| C16orf80 | blue | -0.131883 | 0.520734 |

**Table 6. Gene module membership and correlation with growth rate (48/72h), with adjusted p values for each module**

| | | | | | | | | | | | |
|---|---|---|---|---|---|---|---|---|---|---|---|
| PYGB | blue | 0.242154 | 0.233316 | STK38 | blue | -0.424645452 | 0.027258203 | ZNF800 | blue | -0.131607 | 0.521615 |
| ACTR1B | blue | -0.242141 | 0.233341 | C2orf69 | blue | -0.424231737 | 0.027424631 | EFTUD1 | blue | -0.131413 | 0.522234 |
| SNX6 | blue | -0.241559 | 0.234503 | GID4 | blue | -0.42415403 | 0.027455983 | LOC1005057( | blue | -0.13138 | 0.522339 |
| P4HB | blue | -0.24139 | 0.234839 | PSMB8 | blue | -0.422713296 | 0.028042527 | PARP11 | blue | -0.130645 | 0.52469 |
| ZNF32 | blue | -0.240639 | 0.236343 | SLC24A6 | blue | -0.422395707 | 0.028173175 | ZNF160 | blue | -0.130608 | 0.524806 |
| FTSJD2 | blue | -0.239466 | 0.238705 | SSR2 | blue | -0.420892036 | 0.028798443 | SUCLG1 | blue | -0.130057 | 0.526572 |
| KIAA0040 | blue | -0.23654 | 0.244665 | TIMELESS | blue | -0.420886788 | 0.028800645 | CPED1 | blue | -0.129992 | 0.526779 |
| PAF1 | blue | 0.236438 | 0.244875 | PTP4A1 | blue | -0.42028486 | 0.029054078 | TERF2 | blue | -0.12938 | 0.528747 |
| TMX4 | blue | -0.235489 | 0.246828 | PTCD1 | blue | -0.420208623 | 0.029086305 | OLR1 | blue | -0.127761 | 0.533959 |
| SLCO3A1 | blue | 0.23527 | 0.247283 | FAM82B | blue | -0.420180611 | 0.029098153 | FRS2 | blue | -0.127648 | 0.534324 |
| SPCS1 | blue | -0.235211 | 0.247403 | PTPN12 | blue | -0.419715774 | 0.029295337 | TIMM9 | blue | -0.127596 | 0.534492 |
| PCOLCE2 | blue | -0.234405 | 0.249074 | BTG3 | blue | -0.418880574 | 0.029652337 | RAP2C | blue | -0.12664 | 0.537582 |
| RBCK1 | blue | 0.234017 | 0.249881 | ANKRD42 | blue | -0.418825961 | 0.029675803 | ZNF24 | blue | -0.126048 | 0.5395 |
| ATRX | blue | -0.233994 | 0.24993 | UVRAG | blue | -0.418149787 | 0.029967578 | RSPRY1 | blue | -0.125959 | 0.539792 |
| SCARNA3 | blue | -0.233565 | 0.250825 | WDR36 | blue | -0.418063612 | 0.030004929 | CCP110 | blue | -0.125683 | 0.540687 |
| HRSP12 | blue | -0.233385 | 0.251201 | ESD | blue | -0.417861092 | 0.030092855 | FAM114A2 | blue | -0.125405 | 0.541589 |
| TARS2 | blue | -0.233331 | 0.251312 | SPG20 | blue | -0.417408278 | 0.0302902 | FAM70A | blue | -0.1247 | 0.543881 |
| UBE2E3 | blue | -0.232916 | 0.25218 | CAND1 | blue | -0.417158984 | 0.030399292 | KCTD11 | blue | -0.124491 | 0.544564 |
| TRIM37 | blue | -0.232375 | 0.253316 | RIOK3 | blue | -0.417043926 | 0.030449748 | TNNT1 | blue | -0.124443 | 0.544718 |
| RAB33B | blue | 0.23155 | 0.255053 | WDYHV1 | blue | -0.416993863 | 0.030471723 | PPP3CB | blue | -0.123832 | 0.546712 |
| NEK8 | blue | -0.231042 | 0.256125 | EMR3 | blue | -0.416627206 | 0.030633057 | ABCC5 | blue | -0.123172 | 0.548868 |
| RPS3A | blue | -0.230571 | 0.257123 | KMO | blue | -0.416494784 | 0.030691493 | U2AF1 | blue | -0.122929 | 0.549661 |
| FAM54A | blue | -0.228958 | 0.260558 | KLF7 | blue | -0.415041037 | 0.031338936 | CCL7 | blue | -0.122598 | 0.550745 |
| ACTR10 | blue | -0.228749 | 0.261006 | RBM23 | blue | -0.415009012 | 0.031353322 | BTG1 | blue | -0.12168 | 0.553757 |
| FOCAD | blue | -0.228657 | 0.261204 | RAB1A | blue | -0.414720391 | 0.031483208 | COX16 | blue | -0.121474 | 0.554432 |
| SVIL | blue | -0.227447 | 0.263804 | NHEJ1 | blue | -0.413338698 | 0.032111016 | CRNKL1 | blue | -0.120871 | 0.556415 |
| ERAL1 | blue | -0.226457 | 0.265945 | VPS53 | blue | -0.413309861 | 0.032124225 | ZDHHC21 | blue | -0.120866 | 0.55643 |
| NAA15 | blue | -0.226344 | 0.266191 | CNOT4 | blue | -0.412232406 | 0.032620909 | MBNL3 | blue | -0.120835 | 0.556534 |
| PTK2B | blue | -0.226141 | 0.266632 | FCAR | blue | -0.412004354 | 0.032726823 | LARGE | blue | -0.120517 | 0.55758 |
| CENPC1 | blue | -0.225759 | 0.267462 | EXTL2 | blue | -0.411735727 | 0.032851935 | LY96 | blue | -0.119644 | 0.560457 |
| HSPA5 | blue | 0.225715 | 0.267557 | TMEM8A | blue | -0.410937289 | 0.033226075 | ZNF397 | blue | -0.119519 | 0.560871 |
| ZNF17 | blue | 0.225503 | 0.268019 | PIK3R2 | blue | -0.410889779 | 0.033248445 | SPOP | blue | -0.118891 | 0.562947 |
| RECK | blue | -0.224973 | 0.269175 | RECQL | blue | -0.410707698 | 0.03333429 | CLIC4 | blue | -0.118668 | 0.563684 |
| HDAC5 | blue | 0.223785 | 0.271779 | PNPLA8 | blue | -0.410372795 | 0.033492649 | TRAK2 | blue | -0.118385 | 0.564621 |
| PLP2 | blue | -0.223721 | 0.27192 | CREG1 | blue | -0.410241083 | 0.033555095 | RMRP | blue | -0.117882 | 0.566289 |
| METTL8 | blue | -0.221732 | 0.276315 | TNFSF13 | blue | -0.409847036 | 0.033742472 | ADSS | blue | -0.117647 | 0.567068 |
| UQCRQ | blue | -0.221322 | 0.277227 | CREB3 | blue | -0.409546326 | 0.03388603 | CCDC66 | blue | -0.116997 | 0.569228 |
| COL4A3BP | blue | -0.22018 | 0.279778 | TFRC | blue | -0.409274366 | 0.034016284 | YWHAQ | blue | -0.116732 | 0.570108 |
| MARK2 | blue | 0.219811 | 0.280606 | STOML2 | blue | -0.408854126 | 0.034218345 | NUP43 | blue | -0.1167 | 0.570214 |
| LRRC8D | blue | -0.219632 | 0.281007 | TBC1D2 | blue | -0.408635744 | 0.034323728 | CBX5 | blue | -0.116409 | 0.571181 |
| TFE3 | blue | 0.219568 | 0.281149 | SLC38A9 | blue | -0.407873847 | 0.034693428 | SRP54 | blue | -0.116328 | 0.571452 |
| SEC24D | blue | 0.218964 | 0.282508 | FRMD4B | blue | -0.407640185 | 0.034807446 | C17orf58 | blue | -0.115689 | 0.573582 |
| FCHSD1 | blue | 0.21852 | 0.28351 | PAQR3 | blue | -0.40759039 | 0.034831782 | LUZP1 | blue | -0.115503 | 0.574203 |
| SEC61B | blue | 0.218374 | 0.28384 | TMEM181 | blue | -0.407441363 | 0.0349047 | CREB1 | blue | -0.11547 | 0.574313 |
| TOMM6 | blue | -0.21785 | 0.285026 | FAM126B | blue | -0.407319827 | 0.034964256 | PDIA3 | blue | -0.115334 | 0.574768 |
| ARAF | blue | -0.217616 | 0.285557 | DOCK11 | blue | -0.407041904 | 0.035100753 | TSEN2 | blue | -0.114248 | 0.578398 |
| RSL1D1 | blue | -0.217435 | 0.285967 | SNX3 | blue | -0.406544252 | 0.035346231 | KL | blue | -0.113817 | 0.57984 |
| OR2A4 | blue | -0.217414 | 0.286014 | ZNF622 | blue | -0.406087076 | 0.035572953 | TUBA1A | blue | -0.113412 | 0.5812 |
| GATA3 | blue | -0.217008 | 0.286938 | UFL1 | blue | -0.403223297 | 0.037019761 | PPP1R14B | blue | -0.113126 | 0.582159 |
| MOB3A | blue | -0.216834 | 0.287333 | TGFBR2 | blue | -0.403188445 | 0.037037654 | TNFRSF10C | blue | -0.112757 | 0.583397 |
| ANKRD27 | blue | -0.215682 | 0.289966 | C9orf156 | blue | -0.403018636 | 0.037124931 | UGGT1 | blue | -0.112505 | 0.584246 |
| EIF4H | blue | 0.215614 | 0.290122 | LIN52 | blue | -0.401879013 | 0.037714923 | ISOC2 | blue | -0.112398 | 0.584604 |
| UQCRC1 | blue | -0.214871 | 0.291828 | SNUPN | blue | -0.400656436 | 0.038356155 | GSS | blue | -0.111955 | 0.586097 |
| NCOR1 | blue | -0.214389 | 0.292936 | C12orf4 | blue | -0.400385205 | 0.038499585 | TSPYL1 | blue | -0.111472 | 0.587722 |
| ARL14EP | blue | -0.214047 | 0.293727 | C2orf42 | blue | -0.399837803 | 0.03879036 | XRCC6BP1 | blue | -0.111227 | 0.588551 |
| TMCO4 | blue | 0.214001 | 0.293832 | FARSB | blue | -0.399387862 | 0.039030672 | CLCC1 | blue | -0.11058 | 0.590735 |

**Table 6. Gene module membership and correlation with growth rate (48/72h), with adjusted p values for each module**

| | | | | | | | | | | | |
|---|---|---|---|---|---|---|---|---|---|---|---|
| PTCD2 | blue | -0.213231 | 0.295615 | ATP6V0A1 | blue | -0.39828533 | 0.039624548 | FAM173B | blue | -0.110409 | 0.591312 |
| ZUFSP | blue | -0.212887 | 0.296412 | MGST1 | blue | -0.396563458 | 0.040566398 | FAM199X | blue | -0.110395 | 0.59136 |
| DCBLD2 | blue | 0.212748 | 0.296735 | TRAPPC4 | blue | -0.396447452 | 0.040630487 | SPRTN | blue | -0.108913 | 0.596376 |
| LOC728407 | blue | -0.212161 | 0.298101 | CHSY1 | blue | -0.395156137 | 0.041349339 | ECSIT | blue | -0.108811 | 0.596723 |
| NAP1L1 | blue | -0.212135 | 0.298163 | COMMD2 | blue | -0.392382496 | 0.042927534 | LCMT2 | blue | -0.108756 | 0.596909 |
| ZNF449 | blue | -0.211917 | 0.29867 | GANC | blue | -0.392171951 | 0.043049257 | COA3 | blue | -0.108735 | 0.596981 |
| RABAC1 | blue | -0.211725 | 0.299119 | MRPL35 | blue | -0.388431384 | 0.045257777 | MDM4 | blue | -0.1081 | 0.599137 |
| MAN2A2 | blue | 0.211336 | 0.300027 | POU2F1 | blue | -0.388255199 | 0.045363969 | NBAS | blue | -0.107818 | 0.600096 |
| TRIM22 | blue | -0.20968 | 0.30392 | MRPL44 | blue | -0.387919065 | 0.045567113 | CDC42BPB | blue | -0.107329 | 0.601761 |
| DCXR | blue | 0.209341 | 0.304719 | MED13L | blue | -0.384458879 | 0.047700356 | ACER2 | blue | -0.107146 | 0.602386 |
| ZDHHC24 | blue | 0.209338 | 0.304728 | RRM1 | blue | -0.383925399 | 0.048036144 | CR1 | blue | -0.107132 | 0.602433 |
| TMEM161E | blue | -0.209307 | 0.304799 | EHMT1 | blue | -0.38326976 | 0.048451371 | KIAA0753 | blue | -0.105764 | 0.607101 |
| EREG | blue | -0.208955 | 0.305632 | ZMAT3 | blue | -0.383169827 | 0.048514908 | PRPF4 | blue | -0.105526 | 0.607914 |
| PTDSS1 | blue | -0.208933 | 0.305685 | SMARCA5 | blue | -0.382689452 | 0.048821245 | FAM179B | blue | -0.105343 | 0.608541 |
| EBNA1BP2 | blue | -0.208121 | 0.307612 | EDEM1 | blue | -0.382605222 | 0.048875115 | GLCE | blue | -0.105234 | 0.608913 |
| TXN | blue | -0.20812 | 0.307613 | RPS27 | blue | -0.382492851 | 0.048947056 | C8orf59 | blue | -0.105215 | 0.60898 |
| MCL1 | blue | -0.206972 | 0.310348 | CCDC75 | blue | -0.379243604 | 0.051063567 | RND3 | blue | -0.104747 | 0.610583 |
| IRAK4 | blue | -0.206777 | 0.310814 | PREX1 | blue | -0.379151502 | 0.051124591 | EXOSC4 | blue | -0.102868 | 0.617035 |
| BCL2A1 | blue | -0.205864 | 0.313004 | TCP11L1 | blue | -0.378245451 | 0.05172798 | CDC26 | blue | -0.102124 | 0.619596 |
| PHLPP1 | blue | 0.205685 | 0.313435 | PEAK1 | blue | -0.377208964 | 0.052425072 | FMN1 | blue | -0.101308 | 0.622412 |
| KRT79 | blue | 0.205154 | 0.314713 | TAF1C | blue | -0.377143587 | 0.052469286 | DCUN1D1 | blue | -0.101155 | 0.622941 |
| NOC3L | blue | -0.2051 | 0.314844 | PINX1 | blue | -0.377092925 | 0.05250357 | TYW1 | blue | -0.100682 | 0.624576 |
| PCGF3 | blue | 0.205095 | 0.314856 | DNAJC25 | blue | -0.374682895 | 0.054154857 | HIRA | blue | -0.100384 | 0.625606 |
| METTL14 | blue | -0.204709 | 0.315787 | MOB1A | blue | -0.374540989 | 0.05425334 | NXF1 | blue | -0.100359 | 0.625693 |
| CCDC43 | blue | -0.204703 | 0.3158 | RRP36 | blue | -0.374305169 | 0.054417309 | TTC33 | blue | -0.100078 | 0.626664 |
| ASB13 | blue | -0.20449 | 0.316317 | PNPLA1 | blue | -0.374224268 | 0.05447365 | TCTN3 | blue | -0.098835 | 0.630976 |
| HNRNPR | blue | -0.204391 | 0.316556 | HIST1H4D | blue | -0.373926875 | 0.054681152 | GLT1D1 | blue | -0.098745 | 0.631288 |
| ZFP1 | blue | -0.204078 | 0.317313 | JAK1 | blue | -0.372688049 | 0.05555219 | GIPC1 | blue | -0.098699 | 0.631446 |
| VAC14 | blue | 0.204028 | 0.317433 | VMP1 | blue | -0.372021427 | 0.056025369 | LOC1001299 | blue | -0.097712 | 0.63488 |
| NOMO1 | blue | -0.203621 | 0.318422 | ALOX5 | blue | -0.370553203 | 0.057078643 | DDOST | blue | -0.097369 | 0.636073 |
| TAF2 | blue | -0.203061 | 0.319784 | HNRNPK | blue | -0.370105474 | 0.057402889 | NDUFAF3 | blue | -0.096188 | 0.640192 |
| EARS2 | blue | -0.202444 | 0.321287 | ATF6 | blue | -0.368545478 | 0.05854388 | POLR2M | blue | -0.095527 | 0.642503 |
| NNMT | blue | -0.201869 | 0.322692 | SLC36A4 | blue | -0.367324212 | 0.059449391 | AGPAT9 | blue | -0.095374 | 0.643039 |
| KLF8 | blue | -0.201078 | 0.324632 | RNF19B | blue | -0.367239974 | 0.059512249 | MTHFD1 | blue | -0.095087 | 0.644043 |
| NOL12 | blue | 0.200773 | 0.325382 | TTI2 | blue | -0.366074734 | 0.060387064 | TMEM101 | blue | -0.094911 | 0.64466 |
| MICB | blue | -0.200494 | 0.32607 | ARHGEF11 | blue | -0.365155333 | 0.061084345 | ANKRD13A | blue | -0.094791 | 0.645079 |
| IL1B | blue | -0.200453 | 0.32617 | PRPF18 | blue | -0.364363861 | 0.0616896 | FAM200B | blue | -0.094653 | 0.645564 |
| ISCU | blue | -0.200205 | 0.326781 | PELO | blue | -0.36396726 | 0.061994636 | ATG16L1 | blue | -0.094153 | 0.647315 |
| MLH3 | blue | -0.199353 | 0.328888 | TAPBP | blue | -0.362650243 | 0.063015992 | DOCK7 | blue | -0.093825 | 0.648465 |
| GTF3C1 | blue | -0.199004 | 0.329751 | G2E3 | blue | -0.362494304 | 0.063137782 | SNORA8 | blue | -0.093299 | 0.650314 |
| ZRANB1 | blue | 0.198559 | 0.330855 | INSIG2 | blue | -0.355025496 | 0.069187772 | NME6 | blue | -0.09317 | 0.650767 |
| HCCS | blue | 0.198547 | 0.330886 | HMGXB3 | blue | -0.353985478 | 0.070064477 | SIRT1 | blue | -0.093033 | 0.651246 |
| PNO1 | blue | -0.198425 | 0.331188 | COQ10B | blue | -0.353164454 | 0.070762597 | SNRPD2 | blue | -0.090316 | 0.660826 |
| FBXL20 | blue | 0.198247 | 0.331633 | TPMT | blue | -0.352817709 | 0.071059038 | PWWP2B | blue | -0.090293 | 0.660909 |
| COX15 | blue | -0.19824 | 0.331649 | TMEM50A | blue | -0.352435273 | 0.071387099 | SWSAP1 | blue | -0.089847 | 0.662484 |
| TMEM62 | blue | -0.197623 | 0.333189 | PGM2 | blue | -0.352351793 | 0.071458865 | LITAF | blue | -0.089148 | 0.664961 |
| NUPL2 | blue | 0.197424 | 0.333685 | MED9 | blue | -0.351080513 | 0.072558615 | PIGP | blue | -0.087892 | 0.669415 |
| TEC | blue | -0.197114 | 0.33446 | MTA2 | blue | -0.350399391 | 0.073153158 | NANOS1 | blue | -0.087727 | 0.67 |
| ARRDC1 | blue | -0.197012 | 0.334715 | EPB41 | blue | -0.349806574 | 0.073673658 | KIAA1919 | blue | -0.087123 | 0.672151 |
| RANBP2 | blue | -0.196664 | 0.335585 | KDM1A | blue | -0.348530448 | 0.074803748 | CARD8 | blue | -0.086473 | 0.674464 |
| VPS36 | blue | -0.196365 | 0.336335 | ZBTB24 | blue | -0.348169098 | 0.075126147 | RAB31 | blue | -0.086195 | 0.675453 |
| WBP4 | blue | -0.196144 | 0.336892 | SMCR8 | blue | -0.347047912 | 0.076133253 | REEP5 | blue | -0.085375 | 0.678378 |
| RPL6 | blue | -0.195432 | 0.338684 | DIMT1 | blue | -0.346894359 | 0.076271983 | CCDC90B | blue | -0.085375 | 0.67838 |
| ERCC2 | blue | -0.195282 | 0.339061 | PTPRF | blue | -0.346794166 | 0.076362609 | RALGAPA1 | blue | -0.085301 | 0.678643 |
| MPST | blue | -0.195201 | 0.339264 | IRAK3 | blue | -0.346479884 | 0.076647412 | FKBP4 | blue | -0.08499 | 0.679752 |
| LTN1 | blue | -0.194917 | 0.339981 | SERAC1 | blue | -0.346286184 | 0.076823348 | GARS | blue | -0.084934 | 0.679954 |

**Table 6. Gene module membership and correlation with growth rate (48/72h), with adjusted p values for each module**

| | | | | | | | | | | | |
|---|---|---|---|---|---|---|---|---|---|---|---|
| FAM188A | blue | -0.194872 | 0.340096 | AP3B1 | blue | -0.3455657 | 0.077480471 | CPSF4 | blue | -0.084633 | 0.681028 |
| ZNF616 | blue | -0.194232 | 0.341715 | FGD2 | blue | -0.344508797 | 0.078452188 | RFNG | blue | -0.084059 | 0.683079 |
| OTUD1 | blue | 0.194007 | 0.342285 | RSPO3 | blue | -0.343914314 | 0.079002829 | TSPAN14 | blue | -0.083931 | 0.68354 |
| TOR1A | blue | -0.193918 | 0.342512 | TTC23 | blue | -0.342738657 | 0.080100463 | HIF1A | blue | -0.082653 | 0.688118 |
| PEF1 | blue | -0.19342 | 0.343776 | SEC23B | blue | -0.342613722 | 0.080217786 | RNF138 | blue | -0.082338 | 0.689248 |
| PTPN22 | blue | -0.193264 | 0.344175 | SEC11A | blue | -0.338216145 | 0.084431485 | LAPTM5 | blue | -0.082336 | 0.689256 |
| BAG6 | blue | 0.192261 | 0.346733 | C3AR1 | blue | -0.336554024 | 0.086067121 | CD300A | blue | -0.082005 | 0.690444 |
| SNORA8 | blue | -0.191532 | 0.348599 | FBXO8 | blue | -0.335843291 | 0.086773813 | FTH1 | blue | -0.080834 | 0.694656 |
| ZNF516 | blue | 0.191318 | 0.349146 | FER | blue | -0.334875172 | 0.087743488 | EPB41L3 | blue | -0.079715 | 0.698685 |
| GTF2IRD2B | blue | -0.191098 | 0.349713 | TGDS | blue | -0.334653401 | 0.087966766 | ATRN | blue | -0.079143 | 0.70075 |
| SLC11A1 | blue | -0.191035 | 0.349874 | INTS3 | blue | -0.334343279 | 0.088279714 | FAM105A | blue | -0.078996 | 0.70128 |
| ATXN7 | blue | -0.190004 | 0.352529 | ALKBH7 | blue | -0.33319691 | 0.089443847 | REL | blue | -0.078293 | 0.703822 |
| POMT1 | blue | 0.189896 | 0.35281 | NUS1 | blue | -0.332999134 | 0.089645855 | OSBPL1A | blue | -0.078073 | 0.704615 |
| KLF13 | blue | -0.189769 | 0.353136 | PROSER1 | blue | -0.331385465 | 0.091306955 | ATP11C | blue | -0.077077 | 0.708222 |
| GPR153 | blue | 0.188774 | 0.355714 | SRRM1 | blue | -0.33108547 | 0.09161831 | TCP11L2 | blue | -0.076675 | 0.70968 |
| MRE11A | blue | -0.188512 | 0.356394 | DCUN1D5 | blue | -0.329460611 | 0.093318624 | PPP3CA | blue | -0.074912 | 0.716081 |
| POR | blue | -0.188455 | 0.356542 | ERLEC1 | blue | -0.32785145 | 0.095025818 | LANCL2 | blue | -0.074488 | 0.717624 |
| SLC4A2 | blue | 0.188369 | 0.356766 | RFC2 | blue | -0.327336077 | 0.095577521 | BCL2L13 | blue | -0.073951 | 0.719578 |
| DCTN6 | blue | -0.18749 | 0.359056 | ATF7IP2 | blue | -0.326987587 | 0.095951937 | XAF1 | blue | -0.073271 | 0.722055 |
| GLO1 | blue | -0.187274 | 0.35962 | FAM177A1 | blue | -0.32659188 | 0.096378419 | ANKRD12 | blue | -0.072723 | 0.724053 |
| GSTM4 | blue | -0.187247 | 0.359689 | SNX10 | blue | -0.32565508 | 0.097393744 | METTL21A | blue | -0.072199 | 0.725965 |
| TK1 | blue | -0.187089 | 0.360102 | CNOT10 | blue | -0.325491652 | 0.097571689 | CBWD2 | blue | -0.071549 | 0.72834 |
| TYW1 | blue | -0.185968 | 0.36304 | VWA9 | blue | -0.325047759 | 0.098056245 | PSMD13 | blue | -0.071462 | 0.728659 |
| METTL4 | blue | -0.185794 | 0.363496 | TIPRL | blue | -0.324971514 | 0.098139654 | TXN | blue | -0.070645 | 0.731651 |
| ITGB8 | blue | -0.185601 | 0.364003 | ALG5 | blue | -0.3249615 | 0.098150613 | CENPQ | blue | -0.070334 | 0.732786 |
| RBBP6 | blue | -0.185491 | 0.364293 | LOC1001310 | blue | -0.324905457 | 0.098211962 | CIRBP | blue | -0.069761 | 0.734889 |
| MLLT6 | blue | 0.184991 | 0.365611 | STK39 | blue | -0.324565222 | 0.098585026 | CD300LF | blue | -0.069332 | 0.736462 |
| SPAG5 | blue | 0.184666 | 0.366468 | PIGA | blue | -0.322813926 | 0.10052212 | POLR3GL | blue | -0.069145 | 0.737146 |
| RPL34 | blue | -0.18418 | 0.367753 | RAB39A | blue | -0.322625664 | 0.100732037 | USP53 | blue | -0.069023 | 0.737595 |
| NUDT16 | blue | -0.183289 | 0.370116 | RAB21 | blue | -0.321600443 | 0.101880941 | KLF11 | blue | -0.069003 | 0.737667 |
| BACH1 | blue | -0.183074 | 0.370689 | NLRP2 | blue | -0.320645897 | 0.102959417 | RABL3 | blue | -0.068279 | 0.740328 |
| CDCA7L | blue | -0.18295 | 0.371018 | CHKB | blue | -0.319073251 | 0.104754785 | LOC728613 | blue | -0.06824 | 0.74047 |
| RSRC2 | blue | -0.18263 | 0.37187 | MTHFD1L | blue | -0.318991888 | 0.104848301 | TCEB1 | blue | -0.067647 | 0.74265 |
| TMEM241 | blue | 0.182432 | 0.372396 | FADS1 | blue | -0.317948849 | 0.106052653 | PARN | blue | -0.06758 | 0.742897 |
| ZNF559 | blue | -0.182388 | 0.372515 | FIG4 | blue | -0.317475332 | 0.106602786 | TTC17 | blue | -0.067223 | 0.744211 |
| AP4S1 | blue | -0.182055 | 0.373401 | GEMIN4 | blue | -0.316559502 | 0.107672819 | DIEXF | blue | -0.067176 | 0.744382 |
| DUSP11 | blue | -0.181757 | 0.3742 | SIPA1L1 | blue | -0.315141445 | 0.109345354 | EXOSC2 | blue | -0.065782 | 0.74952 |
| PFDN1 | blue | -0.181634 | 0.374528 | SAMD8 | blue | -0.314759824 | 0.109798732 | PPFIBP2 | blue | -0.065775 | 0.749544 |
| MIOS | blue | -0.181572 | 0.374694 | PRMT2 | blue | -0.31352715 | 0.111272714 | IPO8 | blue | -0.065472 | 0.750664 |
| NHLRC3 | blue | -0.181507 | 0.374868 | TRMT61B | blue | -0.31179247 | 0.113371725 | TSPAN31 | blue | -0.065471 | 0.750666 |
| C12orf44 | blue | -0.181493 | 0.374903 | GART | blue | -0.310532068 | 0.114915102 | ATG12 | blue | -0.063196 | 0.759076 |
| PLCL2 | blue | -0.181309 | 0.375397 | REPIN1 | blue | -0.309827437 | 0.11578466 | KLHL8 | blue | -0.063113 | 0.759381 |
| AGSK1 | blue | 0.18112 | 0.375904 | C17orf51 | blue | -0.3096592 | 0.11599299 | RYBP | blue | -0.063084 | 0.759489 |
| TAF9B | blue | -0.181049 | 0.376095 | ZNF362 | blue | -0.309572749 | 0.116100151 | TBC1D12 | blue | -0.062522 | 0.76157 |
| DBI | blue | -0.180743 | 0.376915 | CAPN3 | blue | -0.309461209 | 0.116238519 | SNAPIN | blue | -0.06157 | 0.765099 |
| GEMIN7 | blue | -0.180459 | 0.377677 | PLA2G4B | blue | -0.309011981 | 0.11679703 | NOMO1 | blue | -0.061166 | 0.766601 |
| RNF214 | blue | -0.180238 | 0.378271 | IL1R1 | blue | -0.30876887 | 0.117100105 | IRF2BP2 | blue | -0.060859 | 0.76774 |
| HSF1 | blue | 0.180011 | 0.378882 | RWDD2B | blue | -0.308325768 | 0.117653989 | RASAL3 | blue | -0.059533 | 0.772666 |
| RALY | blue | -0.179981 | 0.378965 | TRIM38 | blue | -0.307536727 | 0.118645074 | RBM45 | blue | -0.058951 | 0.774834 |
| YAE1D1 | blue | -0.179513 | 0.380224 | ZBTB37 | blue | -0.306826364 | 0.119542576 | CCAR1 | blue | -0.058552 | 0.776321 |
| CCM2 | blue | 0.179486 | 0.380296 | ANKIB1 | blue | -0.306581921 | 0.119852567 | UAP1 | blue | -0.058279 | 0.777338 |
| ICOS | blue | -0.179415 | 0.38049 | RMND1 | blue | -0.303451202 | 0.123875178 | CUL3 | blue | -0.057599 | 0.779873 |
| GUSB | blue | -0.179303 | 0.380791 | SNRNP27 | blue | -0.300704247 | 0.127485362 | C19orf59 | blue | -0.057442 | 0.780457 |
| ANXA2 | blue | -0.178963 | 0.38171 | CASP1 | blue | -0.300450498 | 0.127822684 | ARIH1 | blue | -0.057366 | 0.780743 |
| TXLNG | blue | -0.178746 | 0.382298 | NUBPL | blue | -0.299750667 | 0.128756383 | DNAJC6 | blue | -0.057131 | 0.781618 |
| SSR1 | blue | -0.178294 | 0.383521 | SLC41A2 | blue | -0.299159806 | 0.129548558 | ZNF736 | blue | -0.056265 | 0.78485 |

**Table 6. Gene module membership and correlation with growth rate (48/72h), with adjusted p values for each module**

| | | | | | | | | | | | |
|---|---|---|---|---|---|---|---|---|---|---|---|
| ZNF445 | blue | -0.177938 | 0.384488 | N4BP1 | blue | -0.29872262 | 0.130136982 | ISG20L2 | blue | -0.05621 | 0.785055 |
| DHX34 | blue | 0.17793 | 0.38451 | IGSF8 | blue | -0.29711061 | 0.132323461 | OTUB2 | blue | -0.055538 | 0.787567 |
| UBE3A | blue | -0.177318 | 0.386173 | PMS2 | blue | -0.296851139 | 0.132677879 | PWP1 | blue | -0.055444 | 0.78792 |
| RPN2 | blue | -0.176796 | 0.387595 | ARFGAP3 | blue | -0.293962726 | 0.136669911 | CCDC69 | blue | -0.054575 | 0.791171 |
| PGF | blue | -0.176397 | 0.388685 | GGCT | blue | -0.292568703 | 0.138627378 | WARS | blue | -0.053968 | 0.793444 |
| UNC45A | blue | 0.176248 | 0.389094 | ATXN10 | blue | -0.292551429 | 0.13865176 | SHPRH | blue | -0.05301 | 0.797034 |
| RSPH3 | blue | -0.175826 | 0.390246 | OCIAD2 | blue | -0.291115942 | 0.140688802 | SAFB | blue | -0.052011 | 0.800784 |
| SNORA19 | blue | -0.17578 | 0.390374 | ZNF207 | blue | -0.290125824 | 0.142106363 | MRPL36 | blue | -0.051915 | 0.801145 |
| RTN1 | blue | 0.175372 | 0.391491 | HINT1 | blue | -0.290117674 | 0.142118075 | RPP21 | blue | -0.05173 | 0.80184 |
| FBXO28 | blue | -0.175061 | 0.392345 | PAPD7 | blue | -0.289346026 | 0.143230032 | SPECC1 | blue | -0.050552 | 0.806266 |
| YPEL2 | blue | -0.174707 | 0.393317 | TBC1D1 | blue | -0.289154519 | 0.143506964 | TMEM135 | blue | -0.049917 | 0.808657 |
| BAZ1A | blue | -0.174362 | 0.394268 | CNN2 | blue | -0.288837133 | 0.143966774 | ZNF670 | blue | -0.048602 | 0.813608 |
| ANKRD12 | blue | -0.174089 | 0.39502 | SESN1 | blue | -0.286316946 | 0.147655544 | ANXA11 | blue | -0.048574 | 0.813714 |
| MLX | blue | -0.173748 | 0.39596 | FAM160A2 | blue | -0.286070595 | 0.148019728 | SLC4A8 | blue | -0.048535 | 0.813861 |
| ANKRD36B | blue | 0.17367 | 0.396177 | ZNF143 | blue | -0.285392928 | 0.149024851 | COX20 | blue | -0.048533 | 0.813869 |
| NDFIP2 | blue | -0.173519 | 0.396595 | EREG | blue | -0.28503495 | 0.149557778 | RNPEP | blue | -0.047764 | 0.816769 |
| PPP6R2 | blue | -0.173385 | 0.396964 | PAICS | blue | -0.282130231 | 0.153932595 | ZBTB40 | blue | -0.04683 | 0.820293 |
| CHMP2B | blue | 0.173184 | 0.39752 | BPNT1 | blue | -0.280647563 | 0.156200496 | PANK2 | blue | -0.046713 | 0.820735 |
| ZNHIT6 | blue | -0.173127 | 0.397677 | PRKACA | blue | -0.280200733 | 0.15688861 | SBF2 | blue | -0.046382 | 0.821987 |
| OGDH | blue | 0.172423 | 0.399628 | RAB8A | blue | -0.27825492 | 0.159910291 | LANCL1 | blue | -0.046125 | 0.822957 |
| VWA5A | blue | 0.172319 | 0.399917 | SCOC | blue | -0.273240743 | 0.16788665 | DSTN | blue | -0.045828 | 0.82408 |
| SLC25A45 | blue | 0.172094 | 0.400543 | PRDM2 | blue | -0.272192329 | 0.169589232 | CAPNS1 | blue | -0.045555 | 0.825113 |
| DHRS3 | blue | -0.171648 | 0.401782 | TFEB | blue | -0.271405597 | 0.170874809 | CPVL | blue | -0.045041 | 0.827055 |
| RNF219 | blue | -0.171624 | 0.401851 | C1orf159 | blue | -0.269466867 | 0.174072026 | USP30-AS1 | blue | -0.043641 | 0.832355 |
| PPP1R14B | blue | -0.1714 | 0.402474 | SLC22A15 | blue | -0.268042314 | 0.176447861 | PRKX | blue | -0.043406 | 0.833245 |
| AGFG1 | blue | 0.170419 | 0.405213 | SVIL | blue | -0.266804558 | 0.178530498 | PTCD2 | blue | -0.043396 | 0.833282 |
| TAP1 | blue | -0.170241 | 0.405712 | ARAP2 | blue | -0.26539531 | 0.180922506 | JMJD4 | blue | -0.042996 | 0.8348 |
| MAP3K7 | blue | -0.169981 | 0.406439 | URM1 | blue | -0.263271982 | 0.184568588 | NAPRT1 | blue | -0.042607 | 0.836274 |
| LINC00493 | blue | -0.169596 | 0.407519 | GALK2 | blue | -0.262570236 | 0.185784732 | CDK2AP1 | blue | -0.041673 | 0.839818 |
| CGRRF1 | blue | -0.169526 | 0.407714 | TDRD3 | blue | -0.257789186 | 0.194218512 | ZNF333 | blue | -0.041151 | 0.841797 |
| CKAP5 | blue | -0.169436 | 0.407968 | RBL2 | blue | -0.25716056 | 0.195346698 | FGD6 | blue | -0.039986 | 0.846222 |
| PDSS2 | blue | -0.169402 | 0.408064 | FOXK2 | blue | -0.256209932 | 0.19706133 | RBM41 | blue | -0.039909 | 0.846518 |
| TM2D2 | blue | -0.168818 | 0.409704 | TTC39C | blue | -0.253912732 | 0.201247347 | CDK2AP2 | blue | -0.039241 | 0.849057 |
| OSGEP | blue | -0.168642 | 0.4102 | MTERFD1 | blue | -0.252698443 | 0.203484468 | ZMYND19 | blue | -0.039217 | 0.849149 |
| MTMR3 | blue | 0.168424 | 0.410815 | NPL | blue | -0.252691568 | 0.203497181 | HPGD | blue | -0.039184 | 0.849272 |
| SREBF2 | blue | -0.168281 | 0.411219 | PNO1 | blue | -0.250788725 | 0.207037058 | EIF3C | blue | -0.039143 | 0.84943 |
| SC5DL | blue | -0.167862 | 0.4124 | ALG12 | blue | -0.249206049 | 0.210013087 | MICB | blue | -0.038892 | 0.850385 |
| MR1 | blue | 0.167738 | 0.412751 | CBX5 | blue | -0.244006035 | 0.219995092 | CYB5R1 | blue | -0.038533 | 0.85175 |
| PSMC4 | blue | 0.167593 | 0.413162 | TMEM167E | blue | -0.243906641 | 0.220188947 | GM2A | blue | -0.038185 | 0.853075 |
| SUCLG1 | blue | -0.166938 | 0.415016 | LYRM1 | blue | -0.242692888 | 0.22256549 | LRRC57 | blue | -0.037667 | 0.855048 |
| GPS2 | blue | -0.166879 | 0.415183 | TMA7 | blue | -0.239608695 | 0.228681687 | CTSD | blue | -0.037512 | 0.855636 |
| CPSF3L | blue | 0.166627 | 0.415897 | NAA60 | blue | -0.238744041 | 0.23041632 | G3BP2 | blue | -0.037299 | 0.856447 |
| MPV17L | blue | 0.165515 | 0.419059 | SIGLEC1 | blue | -0.238143545 | 0.231626162 | SENP2 | blue | -0.037032 | 0.857465 |
| LOC150381 | blue | -0.16538 | 0.419445 | LIMK2 | blue | -0.235887107 | 0.236210086 | RNF25 | blue | -0.036035 | 0.861266 |
| FAM78A | blue | -0.164385 | 0.422287 | GON4L | blue | -0.234699222 | 0.238647282 | ERF | blue | -0.035177 | 0.86454 |
| VEGFB | blue | -0.164265 | 0.422631 | YEATS4 | blue | -0.232639583 | 0.242912407 | PMPCA | blue | -0.033618 | 0.870493 |
| CTSZ | blue | -0.163873 | 0.423753 | CHMP3 | blue | -0.231311321 | 0.245689494 | MFNG | blue | -0.033595 | 0.870579 |
| CLASRP | blue | -0.163802 | 0.423958 | TBC1D12 | blue | -0.2294549 | 0.249605713 | FAM3C | blue | -0.033123 | 0.872382 |
| COPE | blue | 0.163765 | 0.424063 | SEC61G | blue | -0.228916257 | 0.250749627 | CSRNP2 | blue | -0.032841 | 0.873461 |
| PABPN1 | blue | 0.163193 | 0.425705 | RPS27A | blue | -0.228578172 | 0.251469367 | ZFAND3 | blue | -0.031173 | 0.87984 |
| CRADD | blue | -0.162954 | 0.426392 | HCFC2 | blue | -0.226866301 | 0.255134473 | TNFRSF10B | blue | -0.030868 | 0.881009 |
| DAD1 | blue | -0.162466 | 0.427798 | CXCR4 | blue | -0.226727823 | 0.255432468 | GPR183 | blue | -0.030678 | 0.881737 |
| METTL10 | blue | -0.162309 | 0.428252 | UBN2 | blue | -0.22589468 | 0.257230133 | SLAMF8 | blue | -0.02894 | 0.888392 |
| RAB43 | blue | 0.161732 | 0.429916 | HSD17B12 | blue | -0.216319123 | 0.278482055 | CEP192 | blue | -0.027923 | 0.892293 |
| ZEB2 | blue | -0.161666 | 0.430109 | TTL | blue | -0.214426625 | 0.282811096 | NFATC3 | blue | -0.027467 | 0.894043 |
| ATP13A2 | blue | -0.160664 | 0.433008 | LGALSL | blue | -0.213290105 | 0.285431309 | UPP1 | blue | -0.026478 | 0.897838 |

**Table 6. Gene module membership and correlation with growth rate (48/72h), with adjusted p values for each module**

| | | | | | | | | | | | |
|---|---|---|---|---|---|---|---|---|---|---|---|
| GSK3B | blue | 0.16063 | 0.433107 | SMAD1 | blue | -0.212363922 | 0.287577949 | CYBA | blue | -0.026385 | 0.898196 |
| GPBP1L1 | blue | 0.159595 | 0.436117 | SLC46A3 | blue | -0.211907806 | 0.288638847 | LSM6 | blue | -0.025616 | 0.901145 |
| SLC9A6 | blue | -0.159461 | 0.436507 | PAFAH1B1 | blue | -0.210198232 | 0.292637211 | RFX2 | blue | -0.025555 | 0.901379 |
| ST6GALNAC | blue | -0.159374 | 0.43676 | MRPL22 | blue | -0.208183014 | 0.297395008 | SMYD4 | blue | -0.025387 | 0.902025 |
| PATL2 | blue | -0.158821 | 0.438375 | LEPR | blue | -0.207661036 | 0.29863523 | NUDC | blue | -0.025005 | 0.903492 |
| ZNF85 | blue | -0.158493 | 0.439333 | LUZP1 | blue | -0.205447891 | 0.303929604 | AMZ2 | blue | -0.024803 | 0.904271 |
| DEPDC5 | blue | 0.158258 | 0.440022 | ERMAP | blue | -0.202558928 | 0.310928146 | HOMER2 | blue | -0.023655 | 0.908683 |
| SUPT7L | blue | -0.158188 | 0.440226 | FDFT1 | blue | -0.202531547 | 0.310994949 | ZNF12 | blue | -0.023606 | 0.908871 |
| CCDC130 | blue | 0.157929 | 0.440984 | NDUFB2 | blue | -0.201264715 | 0.314095479 | PI4KB | blue | -0.023312 | 0.91 |
| SAFB2 | blue | 0.157438 | 0.442424 | PSD3 | blue | -0.199004262 | 0.319675075 | TMOD2 | blue | -0.021356 | 0.917524 |
| ARMC6 | blue | -0.157169 | 0.443213 | ACO1 | blue | -0.19699639 | 0.324681897 | DDX31 | blue | -0.021193 | 0.918154 |
| RPS13 | blue | -0.157096 | 0.443428 | SYNGAP1 | blue | -0.196769754 | 0.325250026 | USP7 | blue | -0.019762 | 0.923665 |
| NRP1 | blue | 0.156667 | 0.444689 | MLH3 | blue | -0.194981732 | 0.329753493 | GPAT2 | blue | -0.018844 | 0.9272 |
| ATF4 | blue | 0.156481 | 0.445236 | ITGA6 | blue | -0.194865258 | 0.330048162 | FTSJD2 | blue | -0.018819 | 0.927297 |
| BCAT2 | blue | -0.156295 | 0.445785 | ASH2L | blue | -0.194658303 | 0.330572137 | CPNE2 | blue | -0.018732 | 0.92763 |
| ADRBK1 | blue | 0.156135 | 0.446257 | CMPK2 | blue | -0.192634101 | 0.335723667 | STX8 | blue | -0.018334 | 0.929164 |
| BTF3 | blue | -0.156017 | 0.446605 | ERCC3 | blue | -0.191373748 | 0.338955583 | RB1 | blue | -0.015411 | 0.940437 |
| FKBP9 | blue | -0.155776 | 0.447317 | HDAC1 | blue | -0.188817457 | 0.345567958 | NCR3LG1 | blue | -0.014985 | 0.94208 |
| NBAS | blue | -0.155367 | 0.448523 | SLC39A14 | blue | -0.187123441 | 0.349992073 | PSTPIP1 | blue | -0.014578 | 0.94365 |
| EFTUD2 | blue | -0.154437 | 0.45128 | CRY2 | blue | -0.187104538 | 0.35004163 | SPTBN1 | blue | -0.014282 | 0.944792 |
| DTNB | blue | -0.154128 | 0.452195 | NEK8 | blue | -0.185092101 | 0.355341382 | NR3C2 | blue | -0.013447 | 0.948018 |
| TMTC4 | blue | -0.153838 | 0.453056 | STAT3 | blue | -0.18093393 | 0.366441362 | GPR137B | blue | -0.012857 | 0.950297 |
| PARP15 | blue | 0.153442 | 0.454236 | NSUN3 | blue | -0.179514966 | 0.370275117 | BAG2 | blue | -0.011657 | 0.95493 |
| AMOTL1 | blue | -0.15283 | 0.456059 | GNL1 | blue | -0.179367931 | 0.370673708 | HSPA8 | blue | -0.011383 | 0.955986 |
| NUDT9 | blue | -0.151365 | 0.460438 | PINK1 | blue | -0.179101525 | 0.371396533 | ANXA4 | blue | -0.011286 | 0.956364 |
| FYB | blue | -0.151188 | 0.460969 | URB1 | blue | -0.176676478 | 0.378013944 | OPA1 | blue | -0.010202 | 0.960549 |
| ABHD10 | blue | -0.150409 | 0.46331 | HIATL1 | blue | -0.174331574 | 0.384477003 | NBR2 | blue | -0.010122 | 0.96086 |
| RNF181 | blue | -0.149559 | 0.465869 | AZI2 | blue | -0.172491877 | 0.389591706 | THBS1 | blue | -0.009481 | 0.963336 |
| WDR75 | blue | -0.148764 | 0.468271 | CBFA2T2 | blue | -0.165259911 | 0.410070777 | TSPAN33 | blue | -0.009453 | 0.963445 |
| GPR157 | blue | 0.148599 | 0.468771 | ULK4 | blue | -0.165228108 | 0.410162139 | UBE2Q1 | blue | -0.008862 | 0.965728 |
| UBE4B | blue | -0.148496 | 0.469083 | OAS2 | blue | -0.164508314 | 0.412232958 | NAA50 | blue | -0.008719 | 0.966283 |
| LINC00265 | blue | -0.147824 | 0.471117 | RAPGEF2 | blue | -0.160666407 | 0.423383892 | TRMT13 | blue | -0.008693 | 0.966384 |
| COBLL1 | blue | -0.147682 | 0.471548 | XBP1 | blue | -0.159666199 | 0.426313866 | PRAM1 | blue | -0.008572 | 0.966851 |
| FAM129B | blue | 0.147367 | 0.472506 | KLHL2 | blue | -0.157416328 | 0.432944955 | FAM53C | blue | -0.007138 | 0.972395 |
| SIPA1 | blue | -0.146691 | 0.474562 | CYB5R2 | blue | -0.155876578 | 0.437515168 | HSPA5 | blue | -0.004786 | 0.981489 |
| TMEM209 | blue | -0.146376 | 0.475521 | TTC38 | blue | -0.155180968 | 0.439588354 | DGUOK | blue | -0.004102 | 0.984135 |
| SLC9A7 | blue | 0.14635 | 0.475601 | CEP192 | blue | -0.153295042 | 0.445235681 | CCDC111 | blue | -0.002684 | 0.989618 |
| CAPN1 | blue | -0.146114 | 0.47632 | MDH1 | blue | -0.147647551 | 0.462376678 | BOD1L1 | blue | -0.00087 | 0.996635 |
| RAVER1 | blue | 0.145491 | 0.478224 | SLC30A7 | blue | -0.147467488 | 0.462928817 | PAOX | blue | 0.000446 | 0.998276 |
| SNORA10 | blue | -0.145223 | 0.479043 | LIG4 | blue | -0.143733603 | 0.47445562 | LOC550112 | blue | 0.000722 | 0.997207 |
| CFB | blue | 0.144986 | 0.479767 | HMOX1 | blue | -0.138105647 | 0.492105146 | SLC31A2 | blue | 0.000729 | 0.997182 |
| SPATA5L1 | blue | -0.144635 | 0.480843 | SEC22B | blue | -0.136917253 | 0.495873775 | TMEM116 | blue | 0.000926 | 0.996417 |
| CACFD1 | blue | 0.144064 | 0.482596 | EMR4P | blue | -0.134028591 | 0.505094174 | FAM63B | blue | 0.001389 | 0.994627 |
| GBP2 | blue | 0.14302 | 0.485808 | STOM | blue | -0.128458559 | 0.523109431 | PPIC | blue | 0.001591 | 0.993844 |
| TIMM9 | blue | -0.143019 | 0.485812 | ZNF141 | blue | -0.126214372 | 0.530454236 | RAPH1 | blue | 0.002882 | 0.98885 |
| RCC2 | blue | 0.142798 | 0.486492 | ABCC5 | blue | -0.123249276 | 0.540233159 | MRPS24 | blue | 0.003672 | 0.985798 |
| CCND3 | blue | -0.14269 | 0.486826 | FAM20C | blue | -0.123243541 | 0.540252153 | WDR13 | blue | 0.003809 | 0.985266 |
| TMEM65 | blue | -0.141798 | 0.48958 | MRPS36 | blue | -0.121802697 | 0.545034499 | SNTB2 | blue | 0.004033 | 0.9844 |
| CCL24 | blue | 0.141521 | 0.490439 | SPATS2L | blue | -0.117184189 | 0.560495457 | C10orf128 | blue | 0.004209 | 0.983718 |
| EP400 | blue | 0.140948 | 0.492214 | ZNF841 | blue | -0.112665915 | 0.575810931 | GMPR2 | blue | 0.004412 | 0.982935 |
| ZNF7 | blue | -0.14061 | 0.493264 | SYNE1 | blue | -0.112037537 | 0.57795552 | SSFA2 | blue | 0.005463 | 0.978869 |
| BBC3 | blue | -0.139929 | 0.495381 | PIGS | blue | -0.104887229 | 0.602603092 | ATAD3A | blue | 0.006034 | 0.976661 |
| DGAT2 | blue | 0.139257 | 0.497474 | PPP2R2D | blue | -0.097717167 | 0.627754158 | RBP4 | blue | 0.006293 | 0.975662 |
| SYNGR2 | blue | -0.138998 | 0.498282 | KIAA0586 | blue | -0.091733391 | 0.649060809 | RBM15B | blue | 0.008037 | 0.968918 |
| CYFIP2 | blue | -0.138855 | 0.49873 | MRPL9 | blue | -0.071345383 | 0.723615759 | DOLPP1 | blue | 0.009097 | 0.96482 |
| GORASP1 | blue | 0.138228 | 0.50069 | NAPB | blue | -0.067966149 | 0.736236677 | NUFIP2 | blue | 0.010258 | 0.960336 |

**Table 6. Gene module membership and correlation with growth rate (48/72h), with adjusted p values for each module**

| | | | | | | | | | | | |
|---|---|---|---|---|---|---|---|---|---|---|---|
| FNTB | blue | -0.137519 | 0.502909 | NRP2 | blue | -0.063008807 | 0.754872251 | TEX261 | blue | 0.011035 | 0.95733 |
| ARHGDIB | blue | -0.137438 | 0.503162 | SLC25A45 | blue | -0.062735015 | 0.755905507 | PLEKHG2 | blue | 0.012144 | 0.953049 |
| TWISTNB | blue | -0.13733 | 0.503501 | ZFP36L1 | blue | -0.059714756 | 0.767330406 | DGCR6 | blue | 0.012961 | 0.949892 |
| IL15 | blue | -0.137319 | 0.503537 | AP3M2 | blue | -0.05723313 | 0.776753555 | C7orf25 | blue | 0.013695 | 0.947059 |
| NPRL2 | blue | 0.137212 | 0.503871 | ZC3H12A | blue | -0.05312335 | 0.792426514 | VSIG10 | blue | 0.01464 | 0.943412 |
| OGFR | blue | -0.13721 | 0.503878 | GAA | blue | -0.044353631 | 0.826127721 | SFXN1 | blue | 0.015033 | 0.941895 |
| MRPL15 | blue | -0.137182 | 0.503965 | ARHGAP6 | blue | -0.038056023 | 0.850519366 | PSPH | blue | 0.016152 | 0.937578 |
| TIAL1 | blue | -0.136779 | 0.505233 | SPRTN | blue | -0.037177284 | 0.853934091 | FEM1C | blue | 0.016216 | 0.937333 |
| TOP2B | blue | -0.136707 | 0.505457 | FAM120A | blue | -0.029523724 | 0.883777595 | SPR | blue | 0.016231 | 0.937273 |
| CRTAP | blue | 0.136587 | 0.505834 | FAM211A | blue | -0.024500734 | 0.903450781 | TPRN | blue | 0.0164 | 0.936621 |
| SPAG1 | blue | -0.13611 | 0.507337 | DOK2 | blue | -0.02368674 | 0.906644446 | TARP | blue | 0.016468 | 0.936361 |
| FAM32A | blue | 0.135787 | 0.508353 | TMEM245 | blue | -0.009208243 | 0.963641672 | THAP8 | blue | 0.016714 | 0.93541 |
| GLIPR2 | blue | -0.135631 | 0.508845 | TH1L | blue | -0.005877312 | 0.976789231 | MRPS9 | blue | 0.017634 | 0.931865 |
| GLS | blue | 0.135602 | 0.508939 | LOC1001282! | blue | -0.004848168 | 0.980852733 | PFN1 | blue | 0.017644 | 0.931824 |
| ZBTB7C | blue | 0.135536 | 0.509146 | GRK6 | blue | 0.001220132 | 0.995180827 | ZDHHC16 | blue | 0.017776 | 0.931316 |
| ARHGAP17 | blue | 0.134922 | 0.511086 | DPM1 | blue | 0.001865166 | 0.992633182 | ZNF22 | blue | 0.020255 | 0.921763 |
| CLTB | blue | -0.134082 | 0.513744 | HDAC7 | blue | 0.003572488 | 0.9858903 | RAD23B | blue | 0.020949 | 0.919095 |
| MYO1G | blue | -0.133438 | 0.515786 | UBA2 | blue | 0.004448501 | 0.982430918 | DNAJB11 | blue | 0.02112 | 0.918436 |
| EIF4E | blue | -0.13299 | 0.51721 | ANKZF1 | blue | 0.008550142 | 0.966238643 | INCENP | blue | 0.021168 | 0.918248 |
| UFD1L | blue | 0.132081 | 0.520102 | BCL2L13 | blue | 0.011402765 | 0.954984505 | NONO | blue | 0.02137 | 0.917474 |
| DSC2 | blue | 0.132009 | 0.520331 | ZNF582 | blue | 0.018153981 | 0.928387023 | FAM173A | blue | 0.022017 | 0.914982 |
| TRIM28 | blue | 0.13193 | 0.520583 | STARD10 | blue | 0.02326016 | 0.908318677 | BOP1 | blue | 0.023273 | 0.910151 |
| NASP | blue | -0.131674 | 0.521401 | LOC1005061! | blue | 0.027959259 | 0.889898362 | ZNF136 | blue | 0.024356 | 0.905989 |
| MFSD6 | blue | 0.13156 | 0.521765 | SS18L2 | blue | 0.030669651 | 0.879298416 | CEP85 | blue | 0.024365 | 0.905952 |
| TMTC2 | blue | -0.131453 | 0.522106 | BRF2 | blue | 0.030799426 | 0.878791377 | ANO6 | blue | 0.025124 | 0.903037 |
| RETSAT | blue | -0.130992 | 0.523579 | WWP2 | blue | 0.033934714 | 0.8665563 | TYROBP | blue | 0.025479 | 0.901672 |
| MMS19 | blue | -0.1308 | 0.524192 | FKBP9 | blue | 0.038076585 | 0.850439497 | FAM108A1 | blue | 0.025562 | 0.901355 |
| N6AMT1 | blue | -0.130426 | 0.525389 | TOMM40L | blue | 0.040842279 | 0.839709817 | C12orf49 | blue | 0.025716 | 0.900762 |
| AQR | blue | -0.130409 | 0.525445 | PELI3 | blue | 0.045653722 | 0.821111051 | MALSU1 | blue | 0.026409 | 0.898102 |
| TRPV2 | blue | -0.130257 | 0.525933 | POLD4 | blue | 0.049576476 | 0.806016606 | FAM210A | blue | 0.026519 | 0.897678 |
| PTAFR | blue | -0.129868 | 0.527179 | CTPS2 | blue | 0.050120634 | 0.803927946 | ADK | blue | 0.026765 | 0.896737 |
| ADIPOR1 | blue | 0.129834 | 0.527289 | ZNF767 | blue | 0.050877935 | 0.801023364 | FAH | blue | 0.026826 | 0.8965 |
| METTL6 | blue | -0.129621 | 0.527972 | LOC146880 | blue | 0.061214281 | 0.761651973 | RAPGEF3 | blue | 0.027135 | 0.895316 |
| WDR26 | blue | -0.128974 | 0.53005 | PARVB | blue | 0.06244778 | 0.756989934 | HNRNPH3 | blue | 0.027612 | 0.893485 |
| ERGIC3 | blue | 0.128062 | 0.532987 | LSM6 | blue | 0.068144242 | 0.735569822 | RPS2 | blue | 0.027734 | 0.893017 |
| HK1 | blue | -0.127573 | 0.534567 | DCTD | blue | 0.069459525 | 0.730650687 | PIN1 | blue | 0.0279 | 0.892383 |
| HUS1 | blue | -0.127254 | 0.535596 | SLCO3A1 | blue | 0.071770484 | 0.722032978 | RFK | blue | 0.028163 | 0.891374 |
| MBOAT1 | blue | -0.127051 | 0.536252 | LOC729013 | blue | 0.072147296 | 0.720630927 | MAP2K1 | blue | 0.028825 | 0.888833 |
| PRR13 | blue | -0.127032 | 0.536315 | ACACB | blue | 0.073701877 | 0.714855956 | MORC3 | blue | 0.028841 | 0.888775 |
| SEPHS1 | blue | 0.127016 | 0.536367 | ZNF783 | blue | 0.073702516 | 0.714853586 | AKT1 | blue | 0.028983 | 0.888229 |
| FIZ1 | blue | 0.126862 | 0.536864 | ZNF544 | blue | 0.075463711 | 0.708329567 | ATG2B | blue | 0.029081 | 0.887853 |
| NAA40 | blue | -0.126801 | 0.537063 | BLVRB | blue | 0.079273629 | 0.694285504 | MDFIC | blue | 0.029246 | 0.887222 |
| FHL3 | blue | 0.126236 | 0.538891 | C1orf112 | blue | 0.083067955 | 0.68039614 | CRY1 | blue | 0.030216 | 0.883507 |
| FAM96B | blue | 0.125767 | 0.540414 | DDOST | blue | 0.084085575 | 0.676688097 | MED21 | blue | 0.030456 | 0.882585 |
| TXNIP | blue | -0.125717 | 0.540575 | MPLKIP | blue | 0.084258541 | 0.676058562 | PAPD7 | blue | 0.030775 | 0.881365 |
| ABCC1 | blue | 0.124878 | 0.543301 | TRA2B | blue | 0.086437615 | 0.668145863 | IFRD2 | blue | 0.031287 | 0.879407 |
| WASL | blue | 0.124496 | 0.544546 | C12orf73 | blue | 0.086899737 | 0.666472197 | HCST | blue | 0.03146 | 0.878743 |
| LONRF1 | blue | -0.123905 | 0.546475 | BAK1 | blue | 0.086900186 | 0.666470572 | CFL1 | blue | 0.031688 | 0.877869 |
| METTL16 | blue | -0.123632 | 0.547364 | C18orf25 | blue | 0.088136225 | 0.662001695 | GIMAP5 | blue | 0.032938 | 0.873092 |
| INTU | blue | -0.123137 | 0.548983 | WDR13 | blue | 0.089759018 | 0.656151647 | CCDC142 | blue | 0.034013 | 0.868983 |
| RABL2A | blue | -0.123099 | 0.549105 | DDX20 | blue | 0.09120616 | 0.650951438 | UBTD1 | blue | 0.034077 | 0.868737 |
| ZMYM6 | blue | -0.122306 | 0.551704 | DNAJC7 | blue | 0.095690498 | 0.634939118 | GANAB | blue | 0.034612 | 0.866695 |
| PTBP2 | blue | -0.122114 | 0.55233 | LYSMD1 | blue | 0.099865142 | 0.620175142 | ACTB | blue | 0.035439 | 0.863538 |
| BTBD2 | blue | 0.121603 | 0.554009 | RUVBL1 | blue | 0.102903939 | 0.609517408 | SUMO3 | blue | 0.036214 | 0.860585 |
| TMEM104 | blue | 0.12151 | 0.554315 | ZNF737 | blue | 0.104900382 | 0.602557348 | FAIM2 | blue | 0.036402 | 0.859867 |
| TRIM41 | blue | 0.121368 | 0.55478 | NFIA | blue | 0.107567874 | 0.593310718 | PVR | blue | 0.036974 | 0.857687 |

**Table 6. Gene module membership and correlation with growth rate (48/72h), with adjusted p values for each module**

| | | | | | | | | | | | |
|---|---|---|---|---|---|---|---|---|---|---|---|
| BMP1 | blue | 0.12084 | 0.556517 | ZC3HC1 | blue | 0.109250232 | 0.587510537 | FKBP1A | blue | 0.037877 | 0.854248 |
| GCN1L1 | blue | 0.120217 | 0.558567 | PPDPF | blue | 0.110784642 | 0.582242011 | ULK3 | blue | 0.038561 | 0.851645 |
| HNRNPF | blue | -0.119894 | 0.559632 | ZCCHC17 | blue | 0.11084703 | 0.582028237 | GLG1 | blue | 0.038795 | 0.850754 |
| EPT1 | blue | 0.119866 | 0.559727 | C7orf25 | blue | 0.111720202 | 0.579039892 | DCTN6 | blue | 0.039013 | 0.849925 |
| CYTIP | blue | -0.119681 | 0.560335 | KIAA1191 | blue | 0.117285222 | 0.560155116 | PCNT | blue | 0.039057 | 0.849755 |
| HOXB7 | blue | 0.119675 | 0.560356 | SORBS3 | blue | 0.119258923 | 0.553525439 | KANK1 | blue | 0.039411 | 0.848408 |
| CCDC50 | blue | -0.119615 | 0.560554 | PDCD6 | blue | 0.120076705 | 0.550789119 | RPS6KB2 | blue | 0.039982 | 0.846241 |
| GTF3C2 | blue | -0.119455 | 0.561083 | MRI1 | blue | 0.120790247 | 0.548406704 | XKR8 | blue | 0.040988 | 0.842417 |
| BLNK | blue | -0.119051 | 0.562416 | KXD1 | blue | 0.124031651 | 0.537644679 | AMIGO3 | blue | 0.041054 | 0.842167 |
| OC1001310[illegible] | blue | 0.11861 | 0.563876 | PDGFC | blue | 0.124845849 | 0.534957138 | CAPG | blue | 0.041258 | 0.841392 |
| CRELD1 | blue | 0.118511 | 0.564203 | ZNF14 | blue | 0.124870503 | 0.534875861 | TOB2 | blue | 0.041342 | 0.841072 |
| ATPAF2 | blue | 0.118285 | 0.564953 | RDH14 | blue | 0.125937449 | 0.531363946 | SNORA23 | blue | 0.041386 | 0.840904 |
| MNDA | blue | -0.118267 | 0.565011 | ANKRD36 | blue | 0.126760473 | 0.528662437 | DGKH | blue | 0.041934 | 0.838827 |
| ORMDL2 | blue | 0.117765 | 0.566678 | ABHD6 | blue | 0.127601565 | 0.525908418 | STXBP3 | blue | 0.043176 | 0.834116 |
| NME4 | blue | -0.117556 | 0.567372 | ATP5B | blue | 0.132267636 | 0.51075629 | C11orf63 | blue | 0.04369 | 0.832169 |
| PTTG1IP | blue | -0.117172 | 0.568644 | APIP | blue | 0.132743004 | 0.509224746 | CSDA | blue | 0.044235 | 0.830107 |
| DIRC2 | blue | -0.116649 | 0.570385 | FIZ1 | blue | 0.132878829 | 0.508787558 | C8orf82 | blue | 0.044325 | 0.829763 |
| OSTF1 | blue | -0.116361 | 0.571341 | LOC1005060[illegible] | blue | 0.137030157 | 0.495515113 | BCL2 | blue | 0.044474 | 0.829199 |
| ACY1 | blue | -0.115749 | 0.573381 | RASA3 | blue | 0.13708635 | 0.495336654 | CELF1 | blue | 0.04575 | 0.824375 |
| DPF2 | blue | 0.115647 | 0.573721 | POGLUT1 | blue | 0.139636262 | 0.487272607 | RHOU | blue | 0.045916 | 0.823747 |
| ZNF142 | blue | 0.115516 | 0.574161 | CHID1 | blue | 0.140409136 | 0.484841623 | TAP1 | blue | 0.046108 | 0.823022 |
| GPR89A | blue | -0.115515 | 0.574162 | SH3BP4 | blue | 0.140589343 | 0.484275689 | FAM222B | blue | 0.048051 | 0.815685 |
| NLRP1 | blue | 0.115181 | 0.575277 | ALS2 | blue | 0.140929193 | 0.483209321 | SMARCAL1 | blue | 0.048192 | 0.815153 |
| UHMK1 | blue | -0.115102 | 0.57554 | KRTCAP2 | blue | 0.141308006 | 0.482022107 | FBXW4 | blue | 0.048284 | 0.814807 |
| NIPA1 | blue | 0.114941 | 0.57608 | LRRC47 | blue | 0.141576193 | 0.4811825 | MID2 | blue | 0.049056 | 0.811899 |
| RCOR3 | blue | 0.114934 | 0.576104 | RTN1 | blue | 0.14296105 | 0.476858881 | SDC2 | blue | 0.049133 | 0.811609 |
| S100A10 | blue | -0.114738 | 0.576758 | ZNF223 | blue | 0.144598976 | 0.471771052 | CDADC1 | blue | 0.050446 | 0.806665 |
| UBAP2L | blue | 0.114506 | 0.577533 | TXNL4A | blue | 0.144899839 | 0.470839556 | TRNAU1AP | blue | 0.05067 | 0.805825 |
| GIT2 | blue | -0.114306 | 0.578203 | C14orf80 | blue | 0.145087866 | 0.470257893 | L3MBTL2 | blue | 0.050734 | 0.805582 |
| SETD1A | blue | 0.114041 | 0.57909 | SPICE1 | blue | 0.145757743 | 0.468188656 | CD300LB | blue | 0.051312 | 0.803411 |
| RAB5B | blue | -0.1138 | 0.579899 | SYPL1 | blue | 0.145836388 | 0.467946037 | BTBD7 | blue | 0.052088 | 0.800494 |
| UBQLN2 | blue | -0.113055 | 0.582397 | ARID3A | blue | 0.147221002 | 0.463685189 | PLIN2 | blue | 0.052793 | 0.797849 |
| CDK13 | blue | -0.112838 | 0.583125 | VBP1 | blue | 0.147364253 | 0.463245526 | NUBP1 | blue | 0.053336 | 0.795811 |
| LOC440354 | blue | -0.112698 | 0.583595 | HDAC6 | blue | 0.147553 | 0.462666562 | LGALS9 | blue | 0.053373 | 0.795673 |
| VPS28 | blue | -0.112492 | 0.58429 | RAPGEF3 | blue | 0.149395385 | 0.457035152 | RAP1GAP2 | blue | 0.053413 | 0.795524 |
| ACER2 | blue | -0.112227 | 0.585182 | EXOSC5 | blue | 0.15019682 | 0.45459683 | PSMF1 | blue | 0.054057 | 0.79311 |
| TCIRG1 | blue | 0.112004 | 0.585931 | THAP11 | blue | 0.152386587 | 0.447969796 | ZFP1 | blue | 0.054342 | 0.792042 |
| PYGO2 | blue | -0.111671 | 0.587052 | ZNF526 | blue | 0.155879051 | 0.437507808 | CTSL1 | blue | 0.054664 | 0.790839 |
| PAQR4 | blue | 0.111612 | 0.587252 | MRRF | blue | 0.155923903 | 0.437374313 | PPP4C | blue | 0.055373 | 0.788184 |
| NAA25 | blue | -0.111146 | 0.588822 | PGM5 | blue | 0.156649375 | 0.435218138 | LHFPL2 | blue | 0.056334 | 0.784595 |
| SLC43A3 | blue | -0.111076 | 0.589059 | MSL3 | blue | 0.162078796 | 0.419265388 | DYNLRB1 | blue | 0.056462 | 0.784116 |
| PTPRM | blue | -0.110958 | 0.589459 | NUDCD2 | blue | 0.164427082 | 0.412467024 | MYH9 | blue | 0.05673 | 0.783113 |
| DNAJC5B | blue | -0.11061 | 0.590633 | TMEM150B | blue | 0.166048872 | 0.407807929 | CASP9 | blue | 0.057174 | 0.781457 |
| TSR2 | blue | -0.110381 | 0.591406 | RNF26 | blue | 0.169147815 | 0.398987618 | ELAC2 | blue | 0.057538 | 0.780098 |
| ZFAND2A | blue | 0.110168 | 0.592128 | EMR2 | blue | 0.169801439 | 0.39714112 | ZNF282 | blue | 0.058191 | 0.777665 |
| PSMD9 | blue | 0.109807 | 0.593349 | FKBP15 | blue | 0.170291489 | 0.395759901 | CYP27A1 | blue | 0.058423 | 0.776801 |
| DMTF1 | blue | -0.109028 | 0.595988 | NUPL2 | blue | 0.172757346 | 0.388851264 | ARHGEF17 | blue | 0.059219 | 0.773837 |
| JUND | blue | 0.108978 | 0.596158 | PYROXD1 | blue | 0.174124677 | 0.385050283 | ASNSD1 | blue | 0.059762 | 0.771817 |
| TSPYL1 | blue | -0.108715 | 0.59705 | SUPT6H | blue | 0.176386081 | 0.378810916 | DHRSX | blue | 0.05982 | 0.771601 |
| NSUN7 | blue | -0.10851 | 0.597746 | SLC25A30 | blue | 0.176402229 | 0.378766572 | TRAF4 | blue | 0.060188 | 0.770231 |
| EHMT2 | blue | -0.108322 | 0.598383 | HPS3 | blue | 0.178152951 | 0.373976904 | NUP210 | blue | 0.060551 | 0.768882 |
| HYAL2 | blue | -0.108036 | 0.599357 | EMC3 | blue | 0.178388404 | 0.373335441 | ARHGAP5 | blue | 0.060742 | 0.768175 |
| NPL | blue | -0.107772 | 0.600254 | TNFRSF21 | blue | 0.179483037 | 0.37036165 | COG3 | blue | 0.060861 | 0.767734 |
| METAP2 | blue | -0.10741 | 0.601485 | CD81 | blue | 0.181160303 | 0.365831904 | NRDE2 | blue | 0.061092 | 0.766876 |
| SORBS3 | blue | 0.107246 | 0.602043 | NACC2 | blue | 0.18361234 | 0.359268465 | TUBA1C | blue | 0.061335 | 0.765972 |
| SBF2 | blue | -0.106881 | 0.603288 | TMED3 | blue | 0.184600955 | 0.356641988 | FDX1L | blue | 0.062139 | 0.762992 |

**Table 6. Gene module membership and correlation with growth rate (48/72h), with adjusted p values for each module**

| | | | |
|---|---|---|---|
| C2orf29 | blue | 0.106719 | 0.603842 |
| CARD8 | blue | -0.106541 | 0.604448 |
| RIOK2 | blue | -0.106441 | 0.604788 |
| ATP7A | blue | -0.106119 | 0.605889 |
| BCAS3 | blue | -0.106069 | 0.60606 |
| RABIF | blue | -0.106008 | 0.606269 |
| ZCCHC8 | blue | 0.105681 | 0.607384 |
| LPCAT2 | blue | -0.105243 | 0.608884 |
| CCNG1 | blue | -0.104715 | 0.610691 |
| VEZF1 | blue | -0.104565 | 0.611205 |
| FAM219B | blue | -0.104404 | 0.611756 |
| BECN1 | blue | -0.104349 | 0.611947 |
| E2F4 | blue | -0.104054 | 0.612958 |
| PAK4 | blue | 0.103801 | 0.613827 |
| ZNF844 | blue | -0.103749 | 0.614005 |
| MCTS1 | blue | -0.103675 | 0.614259 |
| RUNDC1 | blue | -0.103511 | 0.614822 |
| RALGAPA1 | blue | -0.103473 | 0.614954 |
| FAM214A | blue | -0.102801 | 0.617265 |
| SMURF1 | blue | 0.102552 | 0.618122 |
| FLOT2 | blue | -0.102384 | 0.618699 |
| G6PC3 | blue | -0.102382 | 0.618706 |
| BST2 | blue | -0.102182 | 0.619397 |
| FLCN | blue | 0.102176 | 0.619418 |
| MFSD7 | blue | 0.102024 | 0.619939 |
| RPS6KA1 | blue | 0.10178 | 0.620781 |
| ZNF830 | blue | -0.100401 | 0.625547 |
| CCT6A | blue | -0.099433 | 0.628901 |
| TSPAN31 | blue | -0.099281 | 0.629426 |
| FKBP14 | blue | 0.098851 | 0.630918 |
| CDKN2AIPN | blue | -0.097876 | 0.634308 |
| SUMO2 | blue | 0.097661 | 0.635058 |
| BRI3 | blue | 0.097543 | 0.635468 |
| SH3BP5L | blue | -0.097231 | 0.636554 |
| BBS5 | blue | -0.097148 | 0.636845 |
| RPS12 | blue | -0.097141 | 0.636867 |
| AMMECR1L | blue | -0.096898 | 0.637714 |
| RBBP9 | blue | -0.096843 | 0.637906 |
| ACTN4 | blue | 0.096781 | 0.638124 |
| RYK | blue | -0.096492 | 0.639132 |
| GBA | blue | 0.096338 | 0.639669 |
| MTDH | blue | -0.095777 | 0.64163 |
| TTC32 | blue | -0.095356 | 0.643102 |
| SLC44A1 | blue | -0.094962 | 0.64448 |
| ILK | blue | -0.094765 | 0.64517 |
| ARHGAP5 | blue | -0.094677 | 0.645478 |
| ACBD6 | blue | -0.094609 | 0.645718 |
| TRAF3IP2 | blue | 0.094198 | 0.647156 |
| RNF213 | blue | -0.09408 | 0.647573 |
| SF3B1 | blue | 0.093431 | 0.64985 |
| ST13 | blue | -0.093082 | 0.651076 |
| RPA2 | blue | -0.093069 | 0.651122 |
| PPP1CB | blue | 0.092374 | 0.653566 |
| LILRA6 | blue | -0.092076 | 0.654616 |
| CCDC97 | blue | -0.09194 | 0.655096 |
| ANKRD37 | blue | -0.091815 | 0.655534 |
| RFX3 | blue | 0.185031456 | 0.355501823 |
| GPBP1L1 | blue | 0.185741115 | 0.353627039 |
| DNASE1L1 | blue | 0.188791792 | 0.345634736 |
| POMGNT1 | blue | 0.189321934 | 0.344256953 |
| LOC283070 | blue | 0.189904953 | 0.342745554 |
| EYA3 | blue | 0.192062774 | 0.337186405 |
| DOLPP1 | blue | 0.194107608 | 0.331968859 |
| SRPRB | blue | 0.194768649 | 0.330292697 |
| KPNA2 | blue | 0.195240452 | 0.329099524 |
| SDC4 | blue | 0.196701061 | 0.325422347 |
| BRAT1 | blue | 0.198580799 | 0.320727057 |
| DNASE2 | blue | 0.202114309 | 0.312014026 |
| KLHL22 | blue | 0.204344129 | 0.306591792 |
| UBR2 | blue | 0.204818936 | 0.305444823 |
| AIF1 | blue | 0.205453436 | 0.303916267 |
| ZNF117 | blue | 0.207615871 | 0.298742694 |
| NCAPD2 | blue | 0.211380778 | 0.289867761 |
| CDK12 | blue | 0.213334254 | 0.285329238 |
| CD151 | blue | 0.214713771 | 0.282151517 |
| MRPS26 | blue | 0.216608021 | 0.277824952 |
| ADCK3 | blue | 0.216842877 | 0.2772915 |
| STAT6 | blue | 0.219276038 | 0.271803366 |
| SNORA70G | blue | 0.22008205 | 0.270000861 |
| WHSC1 | blue | 0.222684073 | 0.264234515 |
| EMC8 | blue | 0.22432749 | 0.260633913 |
| KIAA1704 | blue | 0.224493344 | 0.260272319 |
| SWSAP1 | blue | 0.224753993 | 0.259704709 |
| AP2M1 | blue | 0.228126363 | 0.252433321 |
| FDPS | blue | 0.229243822 | 0.25005357 |
| C6orf120 | blue | 0.230886478 | 0.246582133 |
| RPL23A | blue | 0.233699978 | 0.240710291 |
| DHX35 | blue | 0.234181212 | 0.239715283 |
| ZNF506 | blue | 0.236769941 | 0.234409506 |
| PXMP4 | blue | 0.237274279 | 0.233384991 |
| KCTD6 | blue | 0.240737589 | 0.226430111 |
| TNIK | blue | 0.241561462 | 0.224796297 |
| RAB42 | blue | 0.24185915 | 0.224207906 |
| CLK3 | blue | 0.242059344 | 0.223812795 |
| USP28 | blue | 0.244539319 | 0.218956955 |
| MCM5 | blue | 0.244860795 | 0.218332739 |
| PDCD4 | blue | 0.246365863 | 0.215426288 |
| DNAJC19 | blue | 0.247426917 | 0.213393066 |
| CRYL1 | blue | 0.248382149 | 0.211573776 |
| ZNF680 | blue | 0.248767542 | 0.210842761 |
| PRKRIP1 | blue | 0.25049095 | 0.207594782 |
| TCF7L2 | blue | 0.250853555 | 0.206915767 |
| LTA4H | blue | 0.254044503 | 0.201005598 |
| AKAP8L | blue | 0.255057292 | 0.199154158 |
| TMEM66 | blue | 0.257832403 | 0.194141116 |
| MFN2 | blue | 0.25935308 | 0.191431302 |
| CARD9 | blue | 0.259998613 | 0.190288911 |
| HECTD1 | blue | 0.260035345 | 0.190224048 |
| LPL | blue | 0.261016229 | 0.188497631 |
| POLR3K | blue | 0.261387767 | 0.187846541 |
| POLR2L | blue | 0.263983196 | 0.183341686 |
| SLC10A7 | blue | 0.265137939 | 0.181361758 |
| FAM125B | blue | 0.062348 | 0.762216 |
| PSMD4 | blue | 0.062736 | 0.760778 |
| WDR1 | blue | 0.063082 | 0.759498 |
| UXS1 | blue | 0.063673 | 0.757309 |
| RUNX3 | blue | 0.06372 | 0.757137 |
| TRIM11 | blue | 0.065249 | 0.751487 |
| YPEL3 | blue | 0.065357 | 0.751088 |
| TOMM40 | blue | 0.06537 | 0.75104 |
| TNPO3 | blue | 0.065765 | 0.749582 |
| MEN1 | blue | 0.065918 | 0.749018 |
| SNTB1 | blue | 0.067025 | 0.744939 |
| ARMC7 | blue | 0.069082 | 0.737379 |
| WDR60 | blue | 0.069377 | 0.736297 |
| JTB | blue | 0.070549 | 0.731999 |
| ZFP36 | blue | 0.070809 | 0.731048 |
| CTBP1 | blue | 0.070818 | 0.731016 |
| ZDHHC14 | blue | 0.070899 | 0.730718 |
| HERC2P2 | blue | 0.070965 | 0.730479 |
| BCL7C | blue | 0.071354 | 0.729055 |
| CCDC57 | blue | 0.071626 | 0.72806 |
| ZNF445 | blue | 0.071808 | 0.727394 |
| MMS19 | blue | 0.071859 | 0.727209 |
| WDR19 | blue | 0.07227 | 0.725706 |
| ALYREF | blue | 0.07245 | 0.725051 |
| PIGV | blue | 0.072546 | 0.724699 |
| VPS52 | blue | 0.072635 | 0.724375 |
| SEC61B | blue | 0.074106 | 0.719013 |
| FAM174A | blue | 0.074931 | 0.71601 |
| SRXN1 | blue | 0.075388 | 0.714351 |
| DOCK5 | blue | 0.075719 | 0.713149 |
| KLF13 | blue | 0.075938 | 0.712351 |
| ADIPOR1 | blue | 0.076729 | 0.709481 |
| MRPL21 | blue | 0.076949 | 0.708684 |
| CHD6 | blue | 0.077097 | 0.708151 |
| PRKRIR | blue | 0.077591 | 0.70636 |
| PPP1CB | blue | 0.077699 | 0.705968 |
| RPLP1 | blue | 0.077746 | 0.705799 |
| AGPAT6 | blue | 0.078083 | 0.70458 |
| DNAJC4 | blue | 0.078114 | 0.704467 |
| MCOLN1 | blue | 0.078144 | 0.704361 |
| PSPC1 | blue | 0.07821 | 0.70412 |
| TNNI2 | blue | 0.078887 | 0.701675 |
| HMOX2 | blue | 0.079561 | 0.699241 |
| MLKL | blue | 0.079709 | 0.698706 |
| CDK4 | blue | 0.080665 | 0.695261 |
| RAC2 | blue | 0.082065 | 0.69023 |
| THRA | blue | 0.083596 | 0.684737 |
| ZNF514 | blue | 0.083659 | 0.684511 |
| ZDHHC12 | blue | 0.083799 | 0.684013 |
| SLC35A2 | blue | 0.084169 | 0.682687 |
| FZR1 | blue | 0.085328 | 0.678546 |
| FOXN3-AS1 | blue | 0.086299 | 0.675082 |
| COPS7A | blue | 0.086647 | 0.673844 |
| GPNMB | blue | 0.088399 | 0.667617 |
| SLED1 | blue | 0.088516 | 0.667202 |
| ZNF841 | blue | 0.088732 | 0.666435 |

**Table 6. Gene module membership and correlation with growth rate (48/72h), with adjusted p values for each module**

| | | | | | | | | | | | |
|---|---|---|---|---|---|---|---|---|---|---|---|
| POGLUT1 | blue | -0.091438 | 0.656866 | COX4I1 | blue | 0.267073372 | 0.178076742 | DOHH | blue | 0.089037 | 0.665355 |
| SSSCA1 | blue | -0.090767 | 0.659233 | NPEPL1 | blue | 0.267998102 | 0.176521959 | ATOX1 | blue | 0.0893 | 0.664422 |
| CD22 | blue | -0.090709 | 0.659439 | SMAD7 | blue | 0.271420755 | 0.170849974 | PXMP4 | blue | 0.089669 | 0.663114 |
| ZBTB7A | blue | 0.09058 | 0.659892 | TALDO1 | blue | 0.271729808 | 0.170344197 | C1QB | blue | 0.090724 | 0.659386 |
| HIST2H2BE | blue | -0.09054 | 0.660035 | RAC2 | blue | 0.272083975 | 0.169765886 | TBC1D30 | blue | 0.091317 | 0.657291 |
| KIAA0754 | blue | -0.090288 | 0.660924 | C19orf66 | blue | 0.273632062 | 0.16725426 | VASP | blue | 0.092361 | 0.653612 |
| AP4B1 | blue | -0.08989 | 0.662332 | ARMCX5 | blue | 0.27526693 | 0.164630403 | NFIX | blue | 0.093038 | 0.651231 |
| MYO1F | blue | -0.089727 | 0.662909 | SENP3 | blue | 0.275465762 | 0.164313287 | GDF15 | blue | 0.093498 | 0.649613 |
| HSPA1B | blue | 0.088971 | 0.665586 | NFS1 | blue | 0.275470893 | 0.164305108 | ANKRD17 | blue | 0.093761 | 0.648692 |
| VAV1 | blue | -0.088714 | 0.666497 | CREBZF | blue | 0.275929913 | 0.163574694 | ZNF511 | blue | 0.094183 | 0.64721 |
| SH3BP1 | blue | 0.088528 | 0.667158 | GREM1 | blue | 0.277382975 | 0.161277659 | RTEL1 | blue | 0.094184 | 0.647207 |
| HIAT1 | blue | -0.087729 | 0.669995 | ATXN7L3 | blue | 0.277919164 | 0.160435841 | SHB | blue | 0.094326 | 0.646708 |
| PHC3 | blue | -0.087163 | 0.672007 | POLD2 | blue | 0.27893595 | 0.158848051 | DFFA | blue | 0.095009 | 0.644316 |
| USP3 | blue | 0.086921 | 0.672867 | CNIH4 | blue | 0.279498942 | 0.157973712 | TWF2 | blue | 0.095129 | 0.643895 |
| CAMK2G | blue | -0.086841 | 0.673152 | WIPI2 | blue | 0.280008494 | 0.15718532 | STAC | blue | 0.095351 | 0.643117 |
| ARHGAP1 | blue | 0.086808 | 0.67327 | PHB2 | blue | 0.282593059 | 0.153229484 | DAZAP1 | blue | 0.096142 | 0.640353 |
| ZNF609 | blue | -0.086771 | 0.673403 | ANGEL1 | blue | 0.282922144 | 0.152730948 | CDKAL1 | blue | 0.096902 | 0.637701 |
| CTBP1 | blue | -0.086408 | 0.674696 | PLAGL1 | blue | 0.284780444 | 0.149937492 | DCAF15 | blue | 0.097451 | 0.635788 |
| TDG | blue | 0.086339 | 0.674942 | CYP27A1 | blue | 0.285072639 | 0.149501605 | SPSB1 | blue | 0.097537 | 0.635489 |
| MYD88 | blue | -0.086304 | 0.675067 | MEF2B | blue | 0.285937315 | 0.148217025 | XRCC1 | blue | 0.097858 | 0.634373 |
| GCA | blue | -0.0862 | 0.675435 | PI4KAP2 | blue | 0.286539278 | 0.14732742 | MED14 | blue | 0.09871 | 0.631408 |
| PIDD | blue | -0.086183 | 0.675496 | CSTF2T | blue | 0.288691218 | 0.144178521 | GOLM1 | blue | 0.098872 | 0.630846 |
| AGAP6 | blue | -0.085982 | 0.676214 | DCXR | blue | 0.288839598 | 0.143963199 | UBE2J2 | blue | 0.098953 | 0.630567 |
| AQP3 | blue | -0.085278 | 0.678726 | PARK7 | blue | 0.289821981 | 0.142543436 | P2RX4 | blue | 0.099412 | 0.628973 |
| RAB6A | blue | 0.085207 | 0.67898 | TST | blue | 0.289987874 | 0.142304681 | EMC8 | blue | 0.099636 | 0.628196 |
| IL17RC | blue | -0.085145 | 0.679199 | PYGB | blue | 0.290325037 | 0.141820324 | SLC9A6 | blue | 0.099756 | 0.62778 |
| HDAC10 | blue | -0.085049 | 0.679541 | NAGK | blue | 0.292409064 | 0.138852826 | LENG1 | blue | 0.100018 | 0.626872 |
| TPR | blue | -0.084904 | 0.68006 | PDCD10 | blue | 0.29270446 | 0.138435863 | OTUD1 | blue | 0.10044 | 0.625411 |
| SUN2 | blue | 0.084748 | 0.680617 | INPP5K | blue | 0.293465441 | 0.137365881 | PRDX6 | blue | 0.100808 | 0.62414 |
| FBXO22 | blue | -0.08469 | 0.680826 | NCOA7 | blue | 0.293962585 | 0.136670108 | QTRT1 | blue | 0.10114 | 0.622992 |
| WSB1 | blue | -0.08462 | 0.681075 | DTYMK | blue | 0.296309622 | 0.13341977 | THOP1 | blue | 0.10127 | 0.622541 |
| GNA15 | blue | -0.084016 | 0.683236 | THOC7 | blue | 0.296651183 | 0.132951473 | TRIB3 | blue | 0.101273 | 0.622533 |
| POLDIP3 | blue | 0.083715 | 0.684312 | MYD88 | blue | 0.296978543 | 0.132503769 | SEPHS2 | blue | 0.10141 | 0.622058 |
| SMCR8 | blue | -0.083366 | 0.685562 | OSGEP | blue | 0.297683188 | 0.131543798 | FAM120A | blue | 0.101947 | 0.620206 |
| UBE2H | blue | -0.082966 | 0.686995 | SF3B5 | blue | 0.298474741 | 0.130471474 | PNPLA6 | blue | 0.102312 | 0.618947 |
| DNASE1L1 | blue | 0.08284 | 0.687446 | URGCP | blue | 0.300655617 | 0.127549958 | ADH5 | blue | 0.102568 | 0.618066 |
| LMAN2L | blue | -0.082646 | 0.688142 | ACTG1 | blue | 0.301256395 | 0.126753607 | ARHGEF10L | blue | 0.102596 | 0.61797 |
| SLCO2B1 | blue | -0.082365 | 0.689153 | NMRAL1 | blue | 0.301837325 | 0.125987025 | EGR2 | blue | 0.102967 | 0.616694 |
| LSAMP | blue | -0.082323 | 0.689303 | EIF2C3 | blue | 0.303048757 | 0.124399356 | SMIM3 | blue | 0.103369 | 0.61531 |
| ROCK1 | blue | -0.082227 | 0.689647 | CIR1 | blue | 0.307237088 | 0.119023044 | RBBP4 | blue | 0.104275 | 0.6122 |
| CABLES2 | blue | 0.082147 | 0.689934 | ATP5D | blue | 0.308203336 | 0.117807371 | RPL13 | blue | 0.10453 | 0.611325 |
| NLGN3 | blue | 0.082088 | 0.690147 | MED4 | blue | 0.308803927 | 0.117056365 | FSD1L | blue | 0.104591 | 0.611117 |
| RPS26 | blue | 0.082046 | 0.690298 | UBAP2L | blue | 0.310829131 | 0.114549956 | ISOC1 | blue | 0.104943 | 0.60991 |
| NUDT18 | blue | -0.081958 | 0.690612 | NFE2L2 | blue | 0.311614974 | 0.113588138 | HSP90AB1 | blue | 0.10512 | 0.609304 |
| RPL23AP82 | blue | -0.081636 | 0.69177 | DNM1L | blue | 0.313444636 | 0.111371902 | GPX3 | blue | 0.106081 | 0.606017 |
| CPT1A | blue | -0.081498 | 0.692264 | OGG1 | blue | 0.314992267 | 0.109522417 | WDR37 | blue | 0.106187 | 0.605656 |
| RRM2B | blue | -0.081315 | 0.692925 | DPP7 | blue | 0.316606527 | 0.107617683 | TMED9 | blue | 0.106249 | 0.605444 |
| WDFY1 | blue | 0.081277 | 0.693062 | ZKSCAN4 | blue | 0.317010443 | 0.107144958 | DNTTIP1 | blue | 0.106849 | 0.603397 |
| PTPN6 | blue | 0.081199 | 0.69334 | C6orf47 | blue | 0.319675772 | 0.104064198 | SRC | blue | 0.108257 | 0.598605 |
| PREP | blue | -0.080862 | 0.694553 | NOLC1 | blue | 0.324290346 | 0.0988872 | NSUN5 | blue | 0.108899 | 0.596424 |
| CCDC115 | blue | -0.080719 | 0.695068 | MEA1 | blue | 0.331426151 | 0.091264789 | PKD2 | blue | 0.109375 | 0.594811 |
| INTS9 | blue | 0.079289 | 0.700222 | HEBP1 | blue | 0.334049775 | 0.088576669 | UQCRFS1 | blue | 0.109391 | 0.594756 |
| C6orf228 | blue | 0.07906 | 0.701049 | GTDC2 | blue | 0.334624553 | 0.087995842 | AGTRAP | blue | 0.109596 | 0.594063 |
| ATF7IP | blue | 0.078747 | 0.70218 | CHST11 | blue | 0.338873345 | 0.083791309 | IGF2R | blue | 0.10965 | 0.593881 |
| ADIPOR2 | blue | 0.078682 | 0.702414 | ZSCAN16 | blue | 0.338875543 | 0.083789174 | BCS1L | blue | 0.109786 | 0.593419 |
| UROD | blue | -0.078581 | 0.70278 | ARF5 | blue | 0.34050454 | 0.082218302 | GTF3A | blue | 0.110351 | 0.591507 |

**Table 6. Gene module membership and correlation with growth rate (48/72h), with adjusted p values for each module**

| | | | | | | | | | | | |
|---|---|---|---|---|---|---|---|---|---|---|---|
| RHBDD2 | blue | -0.078473 | 0.70317 | ASNA1 | blue | 0.341567651 | 0.081205273 | PGD | blue | 0.11066 | 0.590462 |
| CD99L2 | blue | 0.077889 | 0.705282 | PEPD | blue | 0.343016992 | 0.079839556 | SEL1L | blue | 0.110799 | 0.589995 |
| POLRMT | blue | -0.077731 | 0.705852 | PPM1M | blue | 0.343252509 | 0.079619292 | SFI1 | blue | 0.111071 | 0.589075 |
| AMICA1 | blue | 0.077708 | 0.705936 | COA5 | blue | 0.34572268 | 0.077336931 | GALNT12 | blue | 0.113661 | 0.580363 |
| MAT2A | blue | -0.077463 | 0.706825 | DCTPP1 | blue | 0.348428978 | 0.074894173 | GLYCTK | blue | 0.113807 | 0.579874 |
| NANOS1 | blue | -0.07699 | 0.708536 | FAM120C | blue | 0.349503707 | 0.073940673 | TUBB2A | blue | 0.113892 | 0.57959 |
| PPP2R1A | blue | 0.076513 | 0.710265 | C19orf53 | blue | 0.352509589 | 0.071323259 | OGFOD3 | blue | 0.113982 | 0.579288 |
| CCDC88B | blue | -0.075977 | 0.712211 | ABAT | blue | 0.352740851 | 0.071124875 | SPATA5L1 | blue | 0.114035 | 0.57911 |
| SSFA2 | blue | -0.075837 | 0.712718 | NSFL1C | blue | 0.353227149 | 0.070709099 | RRM2B | blue | 0.114342 | 0.578083 |
| CCT2 | blue | -0.075805 | 0.712834 | PLCD1 | blue | 0.354105875 | 0.06996255 | MED10 | blue | 0.11448 | 0.577619 |
| PSMD4 | blue | -0.075762 | 0.71299 | PDXDC2P | blue | 0.355005254 | 0.069204754 | CSAD | blue | 0.114755 | 0.576702 |
| PATL1 | blue | -0.075663 | 0.713352 | VPS16 | blue | 0.359492563 | 0.065517943 | RUVBL2 | blue | 0.115186 | 0.575261 |
| ZNF268 | blue | 0.075639 | 0.713437 | SNX25 | blue | 0.361824646 | 0.063662871 | AGPAT1 | blue | 0.115223 | 0.575136 |
| TMEM101 | blue | -0.075229 | 0.714928 | TRMT12 | blue | 0.362058086 | 0.063479444 | HEBP2 | blue | 0.116536 | 0.570759 |
| GTDC2 | blue | -0.075151 | 0.715209 | POGK | blue | 0.373164283 | 0.055216069 | KDELR2 | blue | 0.116964 | 0.569337 |
| PAK1IP1 | blue | -0.074935 | 0.715994 | PAF1 | blue | 0.373870827 | 0.054720328 | CACUL1 | blue | 0.117335 | 0.568103 |
| RANBP3 | blue | -0.074847 | 0.716316 | BOLA2B | blue | 0.375269879 | 0.053748976 | WDR18 | blue | 0.117785 | 0.56661 |
| MDM2 | blue | -0.07466 | 0.716995 | PPAN | blue | 0.375470369 | 0.053610891 | PIGQ | blue | 0.118179 | 0.565302 |
| B3GALTL | blue | 0.074295 | 0.718323 | MRPS2 | blue | 0.376561321 | 0.052864372 | BLOC1S4 | blue | 0.118304 | 0.564889 |
| DNAJC3 | blue | -0.07411 | 0.718998 | POMT1 | blue | 0.380167657 | 0.050454472 | SLC35A4 | blue | 0.118456 | 0.564386 |
| SIN3A | blue | 0.073512 | 0.721177 | SLBP | blue | 0.380699042 | 0.050106801 | RNASEH1 | blue | 0.119325 | 0.561512 |
| JMJD8 | blue | -0.073346 | 0.721782 | HEMK1 | blue | 0.385344304 | 0.047147131 | DHX34 | blue | 0.119536 | 0.560815 |
| FMNL1 | blue | -0.073044 | 0.722882 | SCO2 | blue | 0.388931682 | 0.044957305 | TRIM4 | blue | 0.120197 | 0.558633 |
| PCSK7 | blue | 0.073013 | 0.722995 | PTPN6 | blue | 0.391124525 | 0.043658881 | LOC728431 | blue | 0.120921 | 0.55625 |
| ATF7 | blue | 0.07288 | 0.72348 | SLC1A5 | blue | 0.392389022 | 0.042923766 | RASSF8 | blue | 0.122752 | 0.550242 |
| POU2F2 | blue | 0.072535 | 0.724739 | HMGCL | blue | 0.393534069 | 0.042266593 | VKORC1L1 | blue | 0.12494 | 0.543102 |
| FMN1 | blue | -0.072293 | 0.725623 | TRAPPC2L | blue | 0.395256938 | 0.041292864 | KIAA1467 | blue | 0.125027 | 0.542819 |
| LAP3 | blue | -0.072158 | 0.726115 | TMEM160 | blue | 0.39821764 | 0.039661242 | PPP2R1A | blue | 0.125641 | 0.540824 |
| SCFD2 | blue | -0.071881 | 0.72713 | SH3GLB2 | blue | 0.399659165 | 0.038885628 | LOC285359 | blue | 0.125657 | 0.54077 |
| DUSP23 | blue | 0.071697 | 0.727802 | NFU1 | blue | 0.403127313 | 0.037069055 | MYO1D | blue | 0.125734 | 0.540522 |
| C6orf120 | blue | 0.071431 | 0.728772 | MRPL53 | blue | 0.403964568 | 0.036640833 | MAP4K2 | blue | 0.126433 | 0.538254 |
| UNC5B | blue | 0.071428 | 0.728784 | BET1 | blue | 0.406331475 | 0.035451607 | TTLL4 | blue | 0.126469 | 0.538138 |
| ENPP3 | blue | -0.071 | 0.730349 | RHOBTB2 | blue | 0.406993736 | 0.035124453 | C19orf25 | blue | 0.126531 | 0.537937 |
| GALNT11 | blue | -0.07048 | 0.732253 | HEXDC | blue | 0.407394714 | 0.034927549 | ZRANB1 | blue | 0.126866 | 0.536852 |
| ADAP2 | blue | -0.070452 | 0.732355 | AKR7A2 | blue | 0.411643856 | 0.032894812 | EIF4H | blue | 0.127513 | 0.534761 |
| HSPB1 | blue | -0.070325 | 0.73282 | SMU1 | blue | 0.412546227 | 0.032475612 | ZNF330 | blue | 0.128221 | 0.532475 |
| DCAF12 | blue | -0.070175 | 0.73337 | SNX4 | blue | 0.414843971 | 0.031427541 | MBOAT7 | blue | 0.130061 | 0.52656 |
| DUSP18 | blue | -0.07017 | 0.733389 | AP1M1 | blue | 0.419562283 | 0.029360684 | CMC2 | blue | 0.130073 | 0.526522 |
| PFKP | blue | -0.070043 | 0.733853 | SNRPA | blue | 0.419665536 | 0.029316712 | SLC25A44 | blue | 0.132378 | 0.519157 |
| GDAP1 | blue | -0.069945 | 0.734213 | WDR33 | blue | 0.42143756 | 0.028570317 | WDR45L | blue | 0.133474 | 0.515672 |
| TTLL4 | blue | 0.069847 | 0.734572 | PSMG4 | blue | 0.424089264 | 0.027482135 | TPRG1L | blue | 0.133628 | 0.515183 |
| SND1 | blue | -0.068576 | 0.739238 | RBM28 | blue | 0.428613921 | 0.025702936 | SLC41A3 | blue | 0.133739 | 0.514829 |
| MMP10 | blue | -0.067512 | 0.743149 | ZBTB7C | blue | 0.440624104 | 0.021429273 | MIOS | blue | 0.133888 | 0.514359 |
| FOXO4 | blue | 0.067032 | 0.744912 | BCL2L12 | blue | 0.443668899 | 0.020443594 | CHD8 | blue | 0.134212 | 0.513331 |
| SLC27A3 | blue | 0.06667 | 0.746248 | KEAP1 | blue | 0.443681871 | 0.020439475 | DHX16 | blue | 0.135458 | 0.50939 |
| KIAA0664 | blue | 0.066458 | 0.747027 | HSF2BP | blue | 0.44603466 | 0.01970361 | MAOA | blue | 0.135793 | 0.508336 |
| CCDC94 | blue | 0.066287 | 0.747657 | SNHG9 | blue | 0.446966799 | 0.01941814 | MAPK13 | blue | 0.136364 | 0.506538 |
| CMTM3 | blue | 0.066269 | 0.747723 | EPN2 | blue | 0.44876874 | 0.018875897 | MTFP1 | blue | 0.136401 | 0.506422 |
| SNX17 | blue | -0.066162 | 0.748117 | TIMM10 | blue | 0.449719599 | 0.018594807 | NEU1 | blue | 0.13689 | 0.504883 |
| KIAA2026 | blue | -0.064967 | 0.752526 | NGDN | blue | 0.465492835 | 0.014411038 | NSUN5P1 | blue | 0.137383 | 0.503335 |
| SMARCE1 | blue | -0.064902 | 0.752766 | UBA52 | blue | 0.472683494 | 0.012780688 | NOTCH2NL | blue | 0.138678 | 0.499282 |
| EVI5L | blue | 0.06476 | 0.753292 | ZDHHC24 | blue | 0.474919533 | 0.012306086 | VAT1 | blue | 0.140492 | 0.493631 |
| RABEP1 | blue | -0.064294 | 0.755014 | NDUFV1 | blue | 0.475309356 | 0.012224864 | SQSTM1 | blue | 0.142152 | 0.488487 |
| MIER2 | blue | 0.064289 | 0.755032 | EXOSC8 | blue | 0.486476093 | 0.010080336 | VAC14 | blue | 0.142583 | 0.487154 |
| ZNF764 | blue | 0.06416 | 0.755509 | TFDP1 | blue | 0.488295651 | 0.009762643 | TMEM18 | blue | 0.142982 | 0.485924 |
| HSPBP1 | blue | -0.063969 | 0.756214 | TMEM9B | blue | 0.488696812 | 0.009693738 | TMEM184B | blue | 0.143069 | 0.485656 |

**Table 6. Gene module membership and correlation with growth rate (48/72h), with adjusted p values for each module**

| | | | | | | | | | | | |
|---|---|---|---|---|---|---|---|---|---|---|---|
| AHNAK | blue | -0.063864 | 0.756604 | RPS21 | blue | 0.489556812 | 0.009547391 | RPN1 | blue | 0.143288 | 0.484982 |
| CHI3L2 | blue | -0.063576 | 0.757667 | CYB5B | blue | 0.498024226 | 0.008202533 | RABGGTA | blue | 0.143369 | 0.484734 |
| CCDC146 | blue | 0.063366 | 0.758446 | ACAD9 | blue | 0.501959543 | 0.007633866 | TCIRG1 | blue | 0.143951 | 0.482944 |
| AHCYL1 | blue | -0.063365 | 0.75845 | THOC5 | blue | 0.509122463 | 0.0066837 | C15orf23 | blue | 0.144753 | 0.480482 |
| NISCH | blue | -0.063229 | 0.758951 | SAAL1 | blue | 0.51530978 | 0.005945141 | PPP6R2 | blue | 0.145192 | 0.479139 |
| FUT11 | blue | 0.062889 | 0.760212 | CXorf56 | blue | 0.519084831 | 0.005529345 | TCTN1 | blue | 0.145202 | 0.479106 |
| DIDO1 | blue | 0.06252 | 0.761578 | RNASEH2C | blue | 0.532503666 | 0.004244555 | LTBR | blue | 0.145639 | 0.477771 |
| NBR2 | blue | -0.062339 | 0.762248 | SHFM1 | blue | 0.577925054 | 0.001593497 | NEDD8 | blue | 0.145666 | 0.47769 |
| CHID1 | blue | -0.061979 | 0.763582 | NFKBIA | blue | 0.612626493 | 0.000681557 | NT5C | blue | 0.146482 | 0.475197 |
| TNRC6A | blue | -0.061786 | 0.7643 | NOP9 | brown | -0.472128055 | 0.012900907 | SDSL | blue | 0.146621 | 0.474774 |
| ZYX | blue | 0.061774 | 0.764342 | MFSD9 | brown | -0.431602233 | 0.024579969 | SGK3 | blue | 0.146826 | 0.47415 |
| SCO2 | blue | -0.061754 | 0.764419 | HIST1H3E | brown | -0.425962468 | 0.026733833 | BBS2 | blue | 0.147519 | 0.472045 |
| TMED5 | blue | 0.06161 | 0.764952 | GHRLOS2 | brown | -0.411572447 | 0.03292817 | ABHD16B | blue | 0.147652 | 0.471639 |
| SLAMF7 | blue | -0.061445 | 0.765564 | ASB3 | brown | -0.394891069 | 0.04149814 | TIGD2 | blue | 0.147686 | 0.471538 |
| DCAF8 | blue | 0.06135 | 0.765917 | BEST1 | brown | -0.383005769 | 0.048619358 | MOGS | blue | 0.147973 | 0.470666 |
| AP2A2 | blue | -0.060969 | 0.76733 | TUSC1 | brown | -0.377789197 | 0.052033932 | ANKRD10 | blue | 0.149477 | 0.466117 |
| BCR | blue | 0.060931 | 0.767473 | PRRC2A | brown | -0.369304349 | 0.05798665 | XIAP | blue | 0.149952 | 0.464685 |
| C4orf52 | blue | -0.060385 | 0.769501 | TBP | brown | -0.352974225 | 0.07092511 | HPCAL1 | blue | 0.150236 | 0.463831 |
| ITGAL | blue | -0.060219 | 0.770118 | SPECC1L | brown | -0.334845899 | 0.087772935 | SNRNP200 | blue | 0.15113 | 0.461145 |
| UAP1L1 | blue | -0.059989 | 0.770974 | RPP40 | brown | -0.334250202 | 0.088373804 | CHCHD2 | blue | 0.152639 | 0.456628 |
| GOLGA3 | blue | 0.05997 | 0.771042 | MTVR2 | brown | -0.326190724 | 0.096812223 | RASSF4 | blue | 0.153308 | 0.454634 |
| CSF1R | blue | 0.059595 | 0.772438 | POU5F1P4 | brown | -0.325154935 | 0.097939086 | EFHD2 | blue | 0.15358 | 0.453824 |
| NFIA | blue | 0.059548 | 0.772613 | SLC25A38 | brown | -0.323176337 | 0.100118944 | PA2G4 | blue | 0.153717 | 0.453416 |
| ANKRD39 | blue | -0.059394 | 0.773185 | NASP | brown | -0.321218446 | 0.102311517 | ZNF835 | blue | 0.155551 | 0.447981 |
| DSTN | blue | -0.059251 | 0.773718 | LOC1001299[illegible] | brown | -0.313583799 | 0.111204656 | BRMS1 | blue | 0.156226 | 0.44599 |
| NCLN | blue | 0.059096 | 0.774295 | PCGF1 | brown | -0.311710232 | 0.113471956 | PTRH1 | blue | 0.15628 | 0.445829 |
| UTP6 | blue | -0.058862 | 0.775167 | CEP290 | brown | -0.308392829 | 0.117570038 | NARFL | blue | 0.156536 | 0.445076 |
| AGPAT9 | blue | -0.058705 | 0.775749 | ZFAND5 | brown | -0.304313647 | 0.122757307 | RNF26 | blue | 0.156587 | 0.444924 |
| IMPA2 | blue | 0.058313 | 0.777211 | PI4KA | brown | -0.302001943 | 0.125770417 | SLC35B1 | blue | 0.156994 | 0.443728 |
| PKD1L1 | blue | 0.0574 | 0.780616 | ZNF584 | brown | -0.299748809 | 0.128758868 | ACTR1B | blue | 0.15819 | 0.440219 |
| NDUFAF2 | blue | 0.057348 | 0.780809 | CYCSP52 | brown | -0.295904062 | 0.133977368 | SEPHS1 | blue | 0.158249 | 0.440047 |
| CAMK1D | blue | 0.056946 | 0.78231 | CECR6 | brown | -0.295433183 | 0.13462689 | NEFH | blue | 0.160088 | 0.434683 |
| FTSJ3 | blue | 0.056918 | 0.782415 | CCDC23 | brown | -0.294857014 | 0.13542475 | GPS1 | blue | 0.160355 | 0.433907 |
| CCNY | blue | -0.056903 | 0.782469 | ERF | brown | -0.291319933 | 0.140398018 | ACTG1 | blue | 0.16039 | 0.433806 |
| SLC15A3 | blue | -0.056803 | 0.782842 | DND1 | brown | -0.279091592 | 0.158605993 | FAM76A | blue | 0.160423 | 0.43371 |
| LOC729013 | blue | 0.056416 | 0.784286 | CELF6 | brown | -0.276744423 | 0.162284265 | ZBTB9 | blue | 0.161337 | 0.431059 |
| SLC25A1 | blue | 0.056151 | 0.785277 | ZNF558 | brown | -0.272297577 | 0.169417767 | LARP1 | blue | 0.161509 | 0.430563 |
| ABCA5 | blue | -0.055725 | 0.786869 | FRAT2 | brown | -0.272172506 | 0.169621542 | MAP2K2 | blue | 0.161521 | 0.430526 |
| PIP4K2C | blue | -0.055707 | 0.786937 | STK36 | brown | -0.266883073 | 0.178397882 | ERCC4 | blue | 0.16153 | 0.430502 |
| IFRD2 | blue | -0.055487 | 0.78776 | CARD6 | brown | -0.263698353 | 0.18383238 | MRRF | blue | 0.161902 | 0.429427 |
| LGALS9 | blue | 0.05532 | 0.788382 | RIPK3 | brown | -0.263454018 | 0.18425402 | RRS1 | blue | 0.163494 | 0.42484 |
| PDCL | blue | -0.054677 | 0.790791 | CCPG1 | brown | -0.262174862 | 0.18647237 | PRKD2 | blue | 0.163919 | 0.423622 |
| SMG6 | blue | -0.054513 | 0.791402 | PEX14 | brown | -0.260945324 | 0.188622062 | CD83 | blue | 0.164009 | 0.423364 |
| ESYT2 | blue | 0.054375 | 0.791921 | RSPRY1 | brown | -0.26006981 | 0.190163203 | DPF2 | blue | 0.164172 | 0.422896 |
| NR3C2 | blue | -0.05418 | 0.792652 | PCGF6 | brown | -0.25742049 | 0.19487966 | SCFD2 | blue | 0.164643 | 0.42155 |
| EIF3K | blue | 0.053719 | 0.794377 | PXDC1 | brown | -0.256218092 | 0.197046568 | HERC2 | blue | 0.16482 | 0.421043 |
| CREBRF | blue | -0.053472 | 0.795305 | NKAPL | brown | -0.250236025 | 0.208073062 | TRIM8 | blue | 0.164924 | 0.420746 |
| AP3D1 | blue | -0.05239 | 0.799361 | LOC1002165[illegible] | brown | -0.249425019 | 0.209599619 | TMEM206 | blue | 0.164942 | 0.420693 |
| EIF3C | blue | -0.051879 | 0.801281 | MCOLN3 | brown | -0.248764038 | 0.210849401 | ARFGAP1 | blue | 0.165614 | 0.418779 |
| GLE1 | blue | -0.051334 | 0.803326 | ITFG3 | brown | -0.248498752 | 0.211352422 | DDX28 | blue | 0.165878 | 0.418027 |
| CARKD | blue | -0.051089 | 0.80425 | ZNF398 | brown | -0.246055872 | 0.216022764 | SLC25A29 | blue | 0.166101 | 0.417392 |
| TRUB2 | blue | -0.050346 | 0.807043 | UNK | brown | -0.244734575 | 0.218577679 | ATPAF2 | blue | 0.166434 | 0.416445 |
| PLD2 | blue | -0.050297 | 0.807226 | SIRPA | brown | -0.238698873 | 0.230507174 | STRN4 | blue | 0.166604 | 0.415964 |
| KIF5B | blue | -0.049973 | 0.808444 | EIF4G3 | brown | -0.2381329 | 0.231647645 | VWA5A | blue | 0.167059 | 0.414672 |
| MSN | blue | -0.049453 | 0.810404 | FOXJ3 | brown | -0.237479544 | 0.232968868 | TIMP1 | blue | 0.167206 | 0.414256 |
| ATP11C | blue | 0.048977 | 0.812195 | SLC38A10 | brown | -0.237170545 | 0.233595472 | ZNF707 | blue | 0.167602 | 0.413137 |

**Table 6. Gene module membership and correlation with growth rate (48/72h), with adjusted p values for each module**

| | | | | | | | | | | | |
|---|---|---|---|---|---|---|---|---|---|---|---|
| FTX | blue | 0.048899 | 0.812489 | TTC30B | brown | -0.23270344 | 0.242779422 | PHF19 | blue | 0.167729 | 0.412777 |
| SSR3 | blue | -0.048876 | 0.812576 | TSPAN15 | brown | -0.232294702 | 0.243631475 | POMGNT1 | blue | 0.167852 | 0.412429 |
| LMLN | blue | -0.04887 | 0.812599 | ATMIN | brown | -0.229980787 | 0.248492197 | PRDX4 | blue | 0.168352 | 0.411019 |
| SPRYD3 | blue | 0.048585 | 0.813674 | C11orf83 | brown | -0.229795129 | 0.248884937 | MCM7 | blue | 0.168475 | 0.410672 |
| IL28RA | blue | -0.047997 | 0.81589 | GGA3 | brown | -0.228305526 | 0.252050781 | MESDC1 | blue | 0.168692 | 0.410059 |
| R3HDM2 | blue | -0.047669 | 0.817125 | CAPNS2 | brown | -0.224520547 | 0.260213041 | EIF2B5 | blue | 0.168874 | 0.409548 |
| RBX1 | blue | -0.046175 | 0.822769 | IFFO2 | brown | -0.224438911 | 0.260390957 | SNORA70 | blue | 0.169283 | 0.408397 |
| ZDHHC18 | blue | -0.045487 | 0.825369 | HBA2 | brown | -0.2229343 | 0.263684221 | PEX10 | blue | 0.169864 | 0.406766 |
| MAEA | blue | -0.045478 | 0.825401 | CCDC85B | brown | -0.221544774 | 0.26674943 | PMS2L2 | blue | 0.169936 | 0.406564 |
| TNFRSF10B | blue | 0.044943 | 0.827425 | CREB3L2 | brown | -0.214737052 | 0.282098084 | PDCL3 | blue | 0.170002 | 0.406381 |
| PLOD1 | blue | 0.044491 | 0.829137 | FYN | brown | -0.213659864 | 0.284577157 | MAPKAPK3 | blue | 0.170652 | 0.404561 |
| ALDH2 | blue | 0.043706 | 0.832109 | EIF2A | brown | -0.213265997 | 0.285487055 | PPP1R9B | blue | 0.170831 | 0.40406 |
| ACOT2 | blue | 0.042833 | 0.835417 | SET | brown | -0.21295974 | 0.286195836 | FANCE | blue | 0.172534 | 0.399321 |
| YWHAG | blue | 0.042586 | 0.836354 | CLCN7 | brown | -0.210309021 | 0.292377045 | FBXW5 | blue | 0.174414 | 0.394124 |
| NPTN | blue | 0.042319 | 0.837367 | AGGF1 | brown | -0.209012881 | 0.295429904 | COX7A2L | blue | 0.175462 | 0.391244 |
| GOLPH3L | blue | 0.042284 | 0.837498 | PGAP1 | brown | -0.203909555 | 0.307643917 | RALGAPB | blue | 0.175769 | 0.390402 |
| ZNF223 | blue | -0.042094 | 0.83822 | NSUN5 | brown | -0.20337169 | 0.308949216 | RCE1 | blue | 0.179551 | 0.380123 |
| FAM120C | blue | -0.04208 | 0.838273 | REXO2 | brown | -0.202554194 | 0.310939696 | WSB2 | blue | 0.179598 | 0.379997 |
| ABHD14B | blue | 0.041514 | 0.840418 | ZNF16 | brown | -0.199523538 | 0.318387969 | DNAJC21 | blue | 0.179678 | 0.379779 |
| RPS5 | blue | 0.041321 | 0.841151 | LILRB2 | brown | -0.197980506 | 0.322221956 | MKL1 | blue | 0.179811 | 0.379423 |
| WBP1L | blue | 0.041264 | 0.841368 | ZNF101 | brown | -0.192953143 | 0.334908513 | IFITM10 | blue | 0.179828 | 0.379376 |
| ALCAM | blue | 0.041104 | 0.841978 | MACF1 | brown | -0.192084429 | 0.337130892 | UNC45A | blue | 0.179887 | 0.379215 |
| SLC35F5 | blue | -0.040182 | 0.84548 | LPCAT4 | brown | -0.192020418 | 0.337294998 | CDCP1 | blue | 0.180059 | 0.378753 |
| ANKZF1 | blue | 0.040108 | 0.845762 | JRKL | brown | -0.190018281 | 0.342452231 | C12orf23 | blue | 0.180768 | 0.376848 |
| RECQL5 | blue | 0.040103 | 0.845781 | IQSEC2 | brown | -0.187628641 | 0.348669173 | ABCB6 | blue | 0.181481 | 0.374936 |
| CHTF8 | blue | -0.04007 | 0.845906 | TAS2R43 | brown | -0.186097071 | 0.352688883 | TALDO1 | blue | 0.182051 | 0.373413 |
| SPINT2 | blue | -0.040007 | 0.846142 | ZNF785 | brown | -0.185905691 | 0.353193099 | DUSP18 | blue | 0.182065 | 0.373375 |
| ZADH2 | blue | 0.039973 | 0.846275 | CHD3 | brown | -0.185207004 | 0.355037513 | BAHD1 | blue | 0.182139 | 0.373177 |
| MAD2L2 | blue | 0.039898 | 0.846559 | EML3 | brown | -0.184369889 | 0.357254848 | GBA | blue | 0.183235 | 0.370259 |
| ZNF43 | blue | -0.039205 | 0.849193 | PHC1 | brown | -0.183054507 | 0.360755495 | RNF123 | blue | 0.183268 | 0.370171 |
| MTMR10 | blue | 0.038844 | 0.850567 | LOC148696 | brown | -0.181418785 | 0.365136727 | MSRA | blue | 0.183388 | 0.369852 |
| ADCK1 | blue | 0.038818 | 0.850666 | TMEM14E | brown | -0.180605617 | 0.367326323 | KIAA1967 | blue | 0.184111 | 0.367937 |
| SF3A2 | blue | -0.038781 | 0.850805 | SLC10A3 | brown | -0.179576595 | 0.370108124 | RAB20 | blue | 0.184316 | 0.367395 |
| ST6GALNAC | blue | -0.038462 | 0.852019 | POM121C | brown | -0.175442932 | 0.381405994 | EIF2AK3 | blue | 0.185001 | 0.365585 |
| SLC25A22 | blue | -0.038267 | 0.852764 | LOC1002892 | brown | -0.170435041 | 0.395355811 | RAB34 | blue | 0.185082 | 0.365372 |
| STAT5B | blue | -0.038027 | 0.853675 | CD74 | brown | -0.169444222 | 0.398149663 | TPD52L2 | blue | 0.185656 | 0.36386 |
| BCS1L | blue | -0.038002 | 0.85377 | LOC1002165 | brown | -0.167598723 | 0.403383142 | PIP4K2C | blue | 0.186035 | 0.362863 |
| UBTD1 | blue | 0.03699 | 0.857625 | SH3RF3-AS1 | brown | -0.16720542 | 0.404503451 | NDN | blue | 0.187335 | 0.35946 |
| SLC25A11 | blue | -0.036904 | 0.857954 | LRRC45 | brown | -0.166830925 | 0.405571806 | LMF2 | blue | 0.188191 | 0.357228 |
| AIFM1 | blue | -0.036606 | 0.859088 | PABPC1 | brown | -0.165756682 | 0.408645155 | RFX1 | blue | 0.188945 | 0.35527 |
| ANXA4 | blue | -0.036566 | 0.85924 | WBP1 | brown | -0.16517923 | 0.410302577 | RUSC1 | blue | 0.190486 | 0.351287 |
| PVR | blue | -0.036513 | 0.859442 | KBTBD4 | brown | -0.16436487 | 0.412646336 | SLC25A35 | blue | 0.19106 | 0.34981 |
| KIAA0141 | blue | -0.035631 | 0.862807 | MYO19 | brown | -0.163672139 | 0.414645888 | CUL4A | blue | 0.191176 | 0.349513 |
| VPS11 | blue | -0.035631 | 0.862807 | MED23 | brown | -0.16253235 | 0.417947539 | C6orf1 | blue | 0.191202 | 0.349444 |
| PECR | blue | -0.035513 | 0.863257 | RPL29P2 | brown | -0.160941177 | 0.42258093 | OSGEP | blue | 0.192508 | 0.346102 |
| SLC25A3 | blue | -0.034978 | 0.8653 | ZNF563 | brown | -0.159876478 | 0.425696964 | FBXL12 | blue | 0.194097 | 0.342057 |
| TBK1 | blue | -0.034539 | 0.866975 | FLJ42393 | brown | -0.159247757 | 0.427542921 | EMC10 | blue | 0.194154 | 0.341914 |
| R3HDM4 | blue | 0.034535 | 0.86699 | APPL2 | brown | -0.157374195 | 0.433069666 | SLC39A1 | blue | 0.194627 | 0.340714 |
| ADCY9 | blue | 0.034374 | 0.867606 | LOC653653 | brown | -0.157186679 | 0.433624938 | FAM83G | blue | 0.195915 | 0.337467 |
| SERINC2 | blue | 0.033945 | 0.869242 | FRAT1 | brown | -0.155939383 | 0.437328244 | HIPK1 | blue | 0.196083 | 0.337044 |
| TNFAIP1 | blue | 0.033786 | 0.869851 | ACLY | brown | -0.154590526 | 0.441352245 | WDFY1 | blue | 0.196499 | 0.336 |
| C16orf5 | blue | 0.033708 | 0.870147 | BLOC1S5 | brown | -0.154185338 | 0.442564906 | ATP5D | blue | 0.196908 | 0.334974 |
| USP25 | blue | 0.033612 | 0.870514 | PVRL2 | brown | -0.153311223 | 0.445187064 | DUS1L | blue | 0.199157 | 0.329373 |
| NUP188 | blue | -0.033465 | 0.871076 | LOC90834 | brown | -0.152092467 | 0.448856903 | PCSK7 | blue | 0.199611 | 0.328248 |
| PRCC | blue | -0.033464 | 0.871081 | ATP5L2 | brown | -0.151959672 | 0.449257739 | GIMAP4 | blue | 0.199672 | 0.328097 |
| MAU2 | blue | -0.033361 | 0.871473 | MBOAT7 | brown | -0.151951801 | 0.449281502 | RPS6KA1 | blue | 0.199922 | 0.327479 |

### Table 6. Gene module membership and correlation with growth rate (48/72h), with adjusted p values for each module

| | | | | | | | | | | | |
|---|---|---|---|---|---|---|---|---|---|---|---|
| TAOK1 | blue | 0.033194 | 0.872111 | LINC00341 | brown | -0.150756071 | 0.45289942 | PLCB3 | blue | 0.200566 | 0.325891 |
| ARL14EPL | blue | -0.032732 | 0.873878 | POLD1 | brown | -0.150617321 | 0.453320233 | NUBP2 | blue | 0.200629 | 0.325737 |
| ARMCX1 | blue | 0.032331 | 0.875411 | GUSB | brown | -0.14871816 | 0.459100939 | WDR46 | blue | 0.200842 | 0.325212 |
| CSPG4 | blue | 0.032253 | 0.875711 | NR4A2 | brown | -0.1478285 | 0.461822171 | ENOSF1 | blue | 0.202891 | 0.320198 |
| TOPORS | blue | 0.031842 | 0.87728 | FAM115A | brown | -0.147619597 | 0.462462372 | PIM1 | blue | 0.202911 | 0.320147 |
| RPS6KB2 | blue | -0.030658 | 0.881814 | CD300C | brown | -0.146283614 | 0.466567574 | MFSD6 | blue | 0.206523 | 0.311424 |
| MOB1B | blue | 0.030368 | 0.882924 | ZNF260 | brown | -0.145948378 | 0.467600657 | HIP1 | blue | 0.20693 | 0.310449 |
| GANAB | blue | -0.030221 | 0.883484 | SF3B2 | brown | -0.144444789 | 0.472248797 | DUSP3 | blue | 0.207175 | 0.309863 |
| PDIK1L | blue | 0.030142 | 0.883789 | PSMB5 | brown | -0.143771623 | 0.474337512 | CLN3 | blue | 0.207591 | 0.308872 |
| NARFL | blue | 0.029684 | 0.885542 | S100A2 | brown | -0.14349549 | 0.475195676 | SP100 | blue | 0.208563 | 0.306562 |
| TAX1BP3 | blue | -0.029484 | 0.886309 | ZFHX3 | brown | -0.142704531 | 0.477658239 | CD27-AS1 | blue | 0.209495 | 0.304356 |
| ATP5G1 | blue | 0.028754 | 0.889107 | ALG10 | brown | -0.140316672 | 0.485132131 | BAMBI | blue | 0.212483 | 0.297352 |
| MAP4K2 | blue | -0.028736 | 0.889175 | HIST1H2BM | brown | -0.140249906 | 0.485341956 | NCLN | blue | 0.213473 | 0.295054 |
| CTNND1 | blue | -0.027885 | 0.892439 | SIGMAR1 | brown | -0.140054804 | 0.485955366 | RHOBTB2 | blue | 0.215034 | 0.291453 |
| ATP6V0D1 | blue | 0.027847 | 0.892586 | DAG1 | brown | -0.138313623 | 0.4914471 | PLEKHB2 | blue | 0.215479 | 0.290431 |
| IL17RA | blue | 0.027672 | 0.893256 | SAP30 | brown | -0.136550075 | 0.497041082 | TESK2 | blue | 0.216191 | 0.288802 |
| ARHGEF35 | blue | 0.027114 | 0.895395 | UPK3B | brown | -0.136534714 | 0.497089946 | CRK | blue | 0.216334 | 0.288474 |
| FAM212B | blue | 0.026867 | 0.896344 | RASSF5 | brown | -0.135936315 | 0.498995369 | SPATA2 | blue | 0.217559 | 0.285686 |
| UBE2R2 | blue | 0.026741 | 0.896828 | NAA35 | brown | -0.134460468 | 0.503710285 | JMJD6 | blue | 0.217732 | 0.285293 |
| PDE4A | blue | 0.026529 | 0.897641 | SUB1 | brown | -0.134357574 | 0.504039824 | OGDH | blue | 0.218294 | 0.28402 |
| GFOD1 | blue | -0.026375 | 0.898234 | PRKAB2 | brown | -0.133327396 | 0.507345047 | IPO13 | blue | 0.220212 | 0.279705 |
| OCEL1 | blue | -0.02634 | 0.898367 | ZNFX1 | brown | -0.132442408 | 0.510192946 | LOC1005064! | blue | 0.220931 | 0.278098 |
| RPIA | blue | 0.026309 | 0.898484 | TES | brown | -0.127715044 | 0.525537378 | PYGB | blue | 0.221689 | 0.276411 |
| USP5 | blue | -0.02605 | 0.899482 | AMIGO3 | brown | -0.126882031 | 0.528263987 | C7orf43 | blue | 0.222427 | 0.274775 |
| TBC1D25 | blue | -0.025997 | 0.899682 | THEM4 | brown | -0.126516656 | 0.529462059 | TEC | blue | 0.222764 | 0.27403 |
| C1orf85 | blue | -0.025666 | 0.900956 | MKL2 | brown | -0.125100422 | 0.534118139 | H3F3B | blue | 0.223889 | 0.271549 |
| B3GAT3 | blue | -0.02566 | 0.900978 | MFSD8 | brown | -0.124905632 | 0.534760055 | GBF1 | blue | 0.224584 | 0.270027 |
| SUN1 | blue | -0.025625 | 0.901111 | SRSF8 | brown | -0.123727429 | 0.538650499 | FMNL3 | blue | 0.22835 | 0.261863 |
| ZNF33A | blue | 0.025081 | 0.903202 | SPP1 | brown | -0.123468669 | 0.539506709 | RDBP | blue | 0.22882 | 0.260854 |
| IMP4 | blue | 0.024928 | 0.903789 | AAAS | brown | -0.123297235 | 0.540074316 | NIPAL3 | blue | 0.229327 | 0.259772 |
| CYC1 | blue | -0.02462 | 0.904974 | ZNF304 | brown | -0.123164468 | 0.540514093 | BAK1 | blue | 0.229399 | 0.259618 |
| ELAVL1 | blue | 0.024517 | 0.90537 | KIAA0317 | brown | -0.121997766 | 0.544385885 | PSMC3 | blue | 0.23008 | 0.258167 |
| QKI | blue | -0.02439 | 0.905859 | CTAGE9 | brown | -0.121596121 | 0.545721762 | CHUK | blue | 0.230109 | 0.258104 |
| RPS16 | blue | 0.02381 | 0.908086 | RGS16 | brown | -0.120006311 | 0.551024414 | CERS2 | blue | 0.230112 | 0.258099 |
| EGLN2 | blue | 0.023727 | 0.908406 | ACER2 | brown | -0.117550849 | 0.559260773 | MFSD1 | blue | 0.230212 | 0.257885 |
| GRIK5 | blue | 0.023644 | 0.908725 | TMEM69 | brown | -0.116624297 | 0.56238321 | IRF2BPL | blue | 0.230572 | 0.25712 |
| FAM82A2 | blue | -0.02361 | 0.908854 | GOLGA6B | brown | -0.115939951 | 0.564694485 | DDX54 | blue | 0.231188 | 0.255816 |
| NUPL1 | blue | 0.023433 | 0.909535 | TIAF1 | brown | -0.115073872 | 0.567625667 | FAM203A | blue | 0.231357 | 0.255459 |
| KLHDC5 | blue | -0.023275 | 0.910142 | TACSTD2 | brown | -0.114551823 | 0.569395797 | SUPT16H | blue | 0.232498 | 0.253057 |
| TGFBI | blue | -0.022761 | 0.912118 | SH3GL1P1 | brown | -0.113146863 | 0.574171893 | ST3GAL2 | blue | 0.23276 | 0.252508 |
| FOXP1 | blue | 0.022396 | 0.913524 | CMTM7 | brown | -0.112724443 | 0.575611359 | GYS1 | blue | 0.23326 | 0.251462 |
| NQO1 | blue | 0.02237 | 0.913624 | XRCC3 | brown | -0.111921694 | 0.578351266 | ZADH2 | blue | 0.2337 | 0.250542 |
| POLD4 | blue | -0.022324 | 0.913801 | MAF | brown | -0.111140836 | 0.581021961 | DCTPP1 | blue | 0.234516 | 0.248844 |
| GNB2 | blue | -0.022257 | 0.914058 | ASXL1 | brown | -0.110113503 | 0.584543876 | RPS6KA5 | blue | 0.235921 | 0.245939 |
| AKT3 | blue | -0.021744 | 0.916032 | LOC1003350: | brown | -0.10991296 | 0.585232465 | SMC3 | blue | 0.236356 | 0.245043 |
| RBP7 | blue | 0.021661 | 0.916353 | TNFRSF10D | brown | -0.109910112 | 0.585242247 | MRPS2 | blue | 0.236998 | 0.243726 |
| TRIM23 | blue | -0.021557 | 0.916752 | PRRC1 | brown | -0.109038622 | 0.588238737 | AXIN1 | blue | 0.237339 | 0.243027 |
| DNAJB4 | blue | 0.021486 | 0.917026 | LOC1002722: | brown | -0.108067519 | 0.591585547 | C11orf21 | blue | 0.237425 | 0.242853 |
| MCOLN1 | blue | 0.020423 | 0.921117 | ZNF347 | brown | -0.106671568 | 0.596410877 | C6orf228 | blue | 0.237844 | 0.241997 |
| WWP2 | blue | -0.019382 | 0.925127 | TRAF4 | brown | -0.105721713 | 0.599703788 | BLOC1S3 | blue | 0.237889 | 0.241905 |
| NIPSNAP3A | blue | 0.019177 | 0.925917 | SNHG11 | brown | -0.104320873 | 0.604574188 | C1RL | blue | 0.238701 | 0.240254 |
| ZNF800 | blue | -0.019177 | 0.925918 | TSC2 | brown | -0.103970422 | 0.605795225 | TRAF3 | blue | 0.238843 | 0.239967 |
| LRWD1 | blue | -0.018748 | 0.927569 | GABPB1 | brown | -0.10077662 | 0.616970388 | BCL2L1 | blue | 0.239438 | 0.238762 |
| PCNT | blue | -0.018717 | 0.927689 | HS6ST1 | brown | -0.100422941 | 0.618213113 | HEXA | blue | 0.239708 | 0.238218 |
| HRAS | blue | 0.018331 | 0.929175 | PRDM15 | brown | -0.09967258 | 0.620853059 | KRT79 | blue | 0.240262 | 0.237101 |
| INF2 | blue | -0.018218 | 0.929611 | FOXD2-AS1 | brown | -0.097601514 | 0.628163289 | RCOR3 | blue | 0.24066 | 0.236302 |

**Table 6. Gene module membership and correlation with growth rate (48/72h), with adjusted p values for each module**

| | | | | | | | | | | | |
|---|---|---|---|---|---|---|---|---|---|---|---|
| THEM4 | blue | 0.018059 | 0.930226 | GSG2 | brown | -0.096869681 | 0.630754693 | SIRT6 | blue | 0.24075 | 0.236122 |
| WDR82 | blue | -0.017779 | 0.931306 | C7orf29 | brown | -0.096084946 | 0.63353816 | RSAD1 | blue | 0.240873 | 0.235874 |
| MCM5 | blue | 0.016987 | 0.934357 | LINC00515 | brown | -0.09540725 | 0.635945885 | TMEM150A | blue | 0.240959 | 0.235702 |
| ARSB | blue | 0.016978 | 0.934392 | LOC1002870 | brown | -0.094815639 | 0.638050722 | ITPK1 | blue | 0.241176 | 0.235268 |
| NADSYN1 | blue | -0.016482 | 0.936305 | DNASE1 | brown | -0.094548143 | 0.639003323 | MFSD7 | blue | 0.241642 | 0.234337 |
| ANKRD36BP | blue | 0.016437 | 0.936479 | SCAMP1 | brown | -0.094426711 | 0.639435948 | NOL12 | blue | 0.243923 | 0.229813 |
| TAF6L | blue | -0.015854 | 0.938728 | FIP1L1 | brown | -0.094412517 | 0.639486526 | UBAP2L | blue | 0.244752 | 0.228184 |
| UBE2E2 | blue | -0.01551 | 0.940056 | MGC16025 | brown | -0.093503379 | 0.642729257 | PNKP | blue | 0.24534 | 0.227032 |
| LENG8 | blue | -0.015493 | 0.940122 | ARFGAP2 | brown | -0.092520848 | 0.646240966 | AP5S1 | blue | 0.245822 | 0.226092 |
| FASTKD3 | blue | -0.015079 | 0.941719 | DUSP8 | brown | -0.091725706 | 0.649088351 | SPARC | blue | 0.246495 | 0.224782 |
| PHF10 | blue | -0.015015 | 0.941966 | FLNA | brown | -0.08906575 | 0.658648431 | KXD1 | blue | 0.246917 | 0.223966 |
| ACAD8 | blue | 0.014859 | 0.942568 | PI4K2A | brown | -0.087582305 | 0.664002998 | PDIK1L | blue | 0.247551 | 0.222739 |
| RUVBL1 | blue | -0.014321 | 0.944642 | FTX | brown | -0.086957153 | 0.666264363 | STK19 | blue | 0.247918 | 0.222032 |
| ARRDC2 | blue | -0.014205 | 0.945092 | ERO1LB | brown | -0.085294235 | 0.672293478 | GNB1 | blue | 0.248937 | 0.220077 |
| GRIN2D | blue | -0.013687 | 0.947089 | MRPL55 | brown | -0.083793347 | 0.677752181 | DFNA5 | blue | 0.249841 | 0.21835 |
| TATDN2 | blue | 0.013162 | 0.949117 | MYBBP1A | brown | -0.082196982 | 0.683575601 | ARCN1 | blue | 0.250304 | 0.217471 |
| UCP2 | blue | 0.013148 | 0.949171 | TUBB6 | brown | -0.081932293 | 0.684542887 | MFSD3 | blue | 0.250943 | 0.216259 |
| ATR | blue | -0.012729 | 0.950789 | HIST1H1A | brown | -0.079524085 | 0.693365661 | GTPBP2 | blue | 0.252509 | 0.213311 |
| POLDIP2 | blue | 0.012344 | 0.952276 | MST1 | brown | -0.078457303 | 0.697286538 | TM2D2 | blue | 0.252829 | 0.212711 |
| C19orf59 | blue | -0.012286 | 0.952501 | TMEM167A | brown | -0.076914555 | 0.702970213 | ZSCAN21 | blue | 0.252868 | 0.21264 |
| PHTF2 | blue | -0.012179 | 0.952915 | AES | brown | -0.075596186 | 0.707839646 | TIAM1 | blue | 0.253029 | 0.212338 |
| DNAJB6 | blue | -0.012139 | 0.953067 | RPL23AP64 | brown | -0.075203871 | 0.709290846 | PRSS21 | blue | 0.25315 | 0.212112 |
| PCBP2 | blue | 0.012028 | 0.953497 | FFAR2 | brown | -0.075026484 | 0.70994734 | SH2D4A | blue | 0.253746 | 0.211001 |
| ADD1 | blue | 0.011949 | 0.953801 | RGS2 | brown | -0.073403706 | 0.715962429 | PADI2 | blue | 0.255833 | 0.207143 |
| CD151 | blue | 0.011808 | 0.954345 | SNX15 | brown | -0.073240839 | 0.716567042 | TECPR2 | blue | 0.256328 | 0.206235 |
| PLSCR3 | blue | 0.011631 | 0.955032 | ZNF678 | brown | -0.07253029 | 0.719206771 | GPR157 | blue | 0.257702 | 0.203729 |
| CAPN10 | blue | -0.011613 | 0.955101 | C8orf58 | brown | -0.069950857 | 0.728815765 | SLC16A6 | blue | 0.258051 | 0.203096 |
| BIN2 | blue | -0.011607 | 0.955123 | SIGLEC10 | brown | -0.069874471 | 0.729100942 | TTLL12 | blue | 0.258579 | 0.20214 |
| CCS | blue | -0.011272 | 0.956416 | SLC26A11 | brown | -0.068001967 | 0.736102543 | DUSP22 | blue | 0.260365 | 0.19893 |
| CSRP1 | blue | -0.010174 | 0.960657 | GNG10 | brown | -0.067704473 | 0.737216835 | LMF1 | blue | 0.260611 | 0.198491 |
| FAM3A | blue | 0.010165 | 0.960693 | KRT23 | brown | -0.067403083 | 0.738346249 | CHST11 | blue | 0.260725 | 0.198287 |
| CPSF7 | blue | -0.009949 | 0.961529 | OR6K6 | brown | -0.066102502 | 0.743226031 | MAP3K8 | blue | 0.262909 | 0.194419 |
| SPR | blue | 0.009917 | 0.961653 | PDHX | brown | -0.06537292 | 0.745967684 | TYMP | blue | 0.263388 | 0.193578 |
| BCKDK | blue | -0.009894 | 0.961742 | VAV1 | brown | -0.065194982 | 0.746636808 | PTPN6 | blue | 0.264855 | 0.191017 |
| NR1H3 | blue | -0.008825 | 0.965873 | RPS10P7 | brown | -0.064590775 | 0.748910217 | TMEM189 | blue | 0.267334 | 0.186741 |
| RBMS2 | blue | 0.008471 | 0.96724 | NUMA1 | brown | -0.064101147 | 0.750754014 | MGAT3 | blue | 0.267796 | 0.185952 |
| SH3BP5 | blue | -0.008247 | 0.968108 | MOCS3 | brown | -0.063734921 | 0.752133986 | C2orf29 | blue | 0.269734 | 0.182665 |
| LRRC8C | blue | 0.007793 | 0.969861 | VNN2 | brown | -0.063283799 | 0.75383488 | ZNF408 | blue | 0.270281 | 0.181745 |
| NCOA2 | blue | 0.007128 | 0.972433 | PUM1 | brown | -0.063107812 | 0.754498718 | C14orf169 | blue | 0.271033 | 0.180486 |
| GPATCH1 | blue | 0.006821 | 0.97362 | C9orf3 | brown | -0.062030189 | 0.758567307 | ENTHD2 | blue | 0.271423 | 0.179834 |
| CUL3 | blue | 0.006722 | 0.974003 | TTC28 | brown | -0.061071024 | 0.762193946 | MDK | blue | 0.27208 | 0.178741 |
| DCP2 | blue | -0.005996 | 0.976809 | NUCB1 | brown | -0.06077552 | 0.763312248 | ZMIZ1 | blue | 0.272554 | 0.177956 |
| RDBP | blue | 0.005792 | 0.977596 | RGL1 | brown | -0.060570436 | 0.764088641 | ZNF675 | blue | 0.272825 | 0.177508 |
| SON | blue | 0.005692 | 0.977984 | FLJ10038 | brown | -0.06056201 | 0.764120546 | FRMD8 | blue | 0.273417 | 0.176532 |
| LRRC58 | blue | -0.005201 | 0.979884 | ZNF527 | brown | -0.060184627 | 0.76554981 | ARMCX5 | blue | 0.274499 | 0.174758 |
| AAMP | blue | 0.004953 | 0.980841 | LOC1002893 | brown | -0.059484188 | 0.768204572 | ELOVL5 | blue | 0.275228 | 0.173569 |
| ARID1B | blue | 0.00478 | 0.981509 | SPATA2 | brown | -0.059244705 | 0.769112834 | TMEM66 | blue | 0.276894 | 0.170874 |
| ATAD3A | blue | -0.004607 | 0.982178 | ZNF586 | brown | -0.057752648 | 0.774778258 | HEBP1 | blue | 0.277452 | 0.169978 |
| ZFP64 | blue | 0.004191 | 0.983789 | TSTD2 | brown | -0.057428899 | 0.776009048 | C5orf24 | blue | 0.277924 | 0.169223 |
| CTDSPL2 | blue | -0.004099 | 0.984146 | ZNF254 | brown | -0.056940677 | 0.777866109 | ZBTB7C | blue | 0.278138 | 0.168881 |
| PCBP1 | blue | -0.003839 | 0.985151 | UQCRBP1 | brown | -0.052223537 | 0.795868775 | TIA1 | blue | 0.278687 | 0.168007 |
| ACTN1 | blue | -0.003688 | 0.985734 | XPO6 | brown | -0.050888829 | 0.800981602 | HSF2BP | blue | 0.279842 | 0.166179 |
| ZBTB9 | blue | 0.00362 | 0.985997 | RAD51AP1 | brown | -0.050487063 | 0.802522211 | TMUB2 | blue | 0.280076 | 0.165809 |
| MINK1 | blue | -0.003514 | 0.986406 | WDR5B | brown | -0.049347968 | 0.806894083 | ZNF839 | blue | 0.282927 | 0.161361 |
| ZNF461 | blue | 0.003446 | 0.986669 | MAGEF1 | brown | -0.048668457 | 0.809504768 | MFSD10 | blue | 0.283299 | 0.160787 |
| ZNF638 | blue | -0.003406 | 0.986824 | CTC1 | brown | -0.047336841 | 0.814626596 | GADD45B | blue | 0.283731 | 0.160123 |

**Table 6. Gene module membership and correlation with growth rate (48/72h), with adjusted p values for each module**

| | | | | | | | | | | | |
|---|---|---|---|---|---|---|---|---|---|---|---|
| ZNF280C | blue | 0.00285 | 0.988977 | MT1H | brown | -0.046998754 | 0.81592818 | CTDNEP1 | blue | 0.286083 | 0.156537 |
| FBXO6 | blue | 0.002285 | 0.99116 | VPS18 | brown | -0.045239631 | 0.822708171 | GPR153 | blue | 0.286967 | 0.155205 |
| TMEM167B | blue | 0.002236 | 0.991352 | SLED1 | brown | -0.044850845 | 0.824208322 | ATP6AP1 | blue | 0.28884 | 0.152408 |
| C5orf4 | blue | 0.001966 | 0.992396 | ERI3 | brown | -0.043137422 | 0.830826764 | ZNF232 | blue | 0.289949 | 0.150769 |
| PDK2 | blue | 0.001628 | 0.993703 | FOXO3B | brown | -0.042796121 | 0.832146467 | BRD9 | blue | 0.290575 | 0.149848 |
| MYSM1 | blue | -0.001495 | 0.994219 | LOC646471 | brown | -0.042616065 | 0.832842869 | FAM109B | blue | 0.290997 | 0.149231 |
| SAT2 | blue | 0.001389 | 0.994627 | SRRM5 | brown | -0.042109208 | 0.834803892 | SH3BP1 | blue | 0.291421 | 0.148612 |
| POGZ | blue | -0.001026 | 0.996031 | NFX1 | brown | -0.041038196 | 0.838950779 | SLC7A1 | blue | 0.292092 | 0.147636 |
| PMEPA1 | blue | 0.000937 | 0.996377 | POU5F1P3 | brown | -0.040103542 | 0.842573138 | RABL6 | blue | 0.292532 | 0.147 |
| EIF6 | blue | -0.000836 | 0.996767 | CLCN4 | brown | -0.038172986 | 0.850065055 | TBC1D20 | blue | 0.294343 | 0.144398 |
| TMED7 | blue | 0.000693 | 0.99732 | CAHM | brown | -0.037870182 | 0.851241316 | SLC39A7 | blue | 0.294486 | 0.144194 |
| ZNF256 | blue | -0.000619 | 0.997604 | OPLAH | brown | -0.037853298 | 0.851306915 | BRF2 | blue | 0.295122 | 0.143289 |
| DCAF10 | blue | 0.000499 | 0.998071 | ZNF324B | brown | -0.037278 | 0.853542585 | TAF5L | blue | 0.296348 | 0.141556 |
| SCAF1 | blue | 0.000112 | 0.999567 | SPDYE7P | brown | -0.035794762 | 0.859311667 | ARHGEF18 | blue | 0.299986 | 0.136502 |
| LMNB1 | brown | -0.593894 | 0.00138 | HMGB3 | brown | -0.035503596 | 0.860445001 | GET4 | blue | 0.300164 | 0.136258 |
| MZT2B | brown | -0.560958 | 0.002871 | NAPRT1 | brown | -0.035288589 | 0.861282068 | ZFP64 | blue | 0.302693 | 0.132827 |
| GIMAP8 | brown | -0.54612 | 0.003899 | POLR1E | brown | -0.034447493 | 0.86455802 | ARL11 | blue | 0.302951 | 0.132481 |
| SLFN5 | brown | 0.535964 | 0.00477 | SAP25 | brown | -0.032023626 | 0.874010696 | HAUS5 | blue | 0.304016 | 0.131057 |
| GRAP | brown | 0.526127 | 0.005766 | PGBD4 | brown | -0.031893195 | 0.874519845 | DPP7 | blue | 0.30478 | 0.130043 |
| EIF4A1 | brown | 0.526118 | 0.005767 | MCM4 | brown | -0.030814714 | 0.878731651 | EEF1D | blue | 0.304922 | 0.129855 |
| MYBL2 | brown | -0.523489 | 0.006061 | PFN1P2 | brown | -0.030747743 | 0.878993302 | NCAPH2 | blue | 0.305044 | 0.129694 |
| GNG11 | brown | -0.517198 | 0.006816 | DCAF12 | brown | -0.030714318 | 0.879123892 | SETD1A | blue | 0.305347 | 0.129295 |
| CCZ1B | brown | -0.515952 | 0.006974 | LOC440173 | brown | -0.030425739 | 0.880251516 | LOC283070 | blue | 0.305699 | 0.128831 |
| FZD5 | brown | 0.513789 | 0.007257 | ZNHIT2 | brown | -0.02936614 | 0.884393835 | XYLT2 | blue | 0.306524 | 0.12775 |
| ITSN2 | brown | 0.510021 | 0.007772 | ASS1 | brown | -0.02909762 | 0.885444042 | CCT4 | blue | 0.307938 | 0.125912 |
| RHBDF2 | brown | -0.495811 | 0.009999 | RBBP9 | brown | -0.029041994 | 0.885661624 | RAB11FIP3 | blue | 0.308799 | 0.124803 |
| STX4 | brown | 0.494516 | 0.010226 | GTF2IRD2B | brown | -0.028817278 | 0.886540687 | PSMC4 | blue | 0.308976 | 0.124575 |
| NDUFA11 | brown | -0.48184 | 0.012684 | ZNF485 | brown | -0.027240784 | 0.892711386 | MUS81 | blue | 0.312525 | 0.120082 |
| CDK5RAP2 | brown | 0.479964 | 0.013086 | TREX1 | brown | -0.02616108 | 0.896941097 | GORASP1 | blue | 0.315003 | 0.117016 |
| SOX4 | brown | -0.475795 | 0.014017 | LOC339240 | brown | -0.025934201 | 0.897830241 | PTPN23 | blue | 0.315268 | 0.116692 |
| DHX57 | brown | -0.473958 | 0.014445 | SSBP3 | brown | -0.025080656 | 0.901176379 | ZDHHC24 | blue | 0.315935 | 0.115879 |
| C3AR1 | brown | -0.473179 | 0.014629 | SLC25A3 | brown | -0.025016213 | 0.901429084 | SLC1A5 | blue | 0.316521 | 0.115167 |
| MAPKAP1 | brown | -0.471577 | 0.015014 | MAPK3 | brown | -0.024698337 | 0.902675718 | TSNARE1 | blue | 0.317117 | 0.114447 |
| APEX2 | brown | -0.467334 | 0.016076 | HIST1H2AE | brown | -0.022387398 | 0.911745265 | LOXL3 | blue | 0.317251 | 0.114285 |
| TAB3 | brown | 0.466974 | 0.016169 | ULK3 | brown | -0.020566005 | 0.918901199 | ARHGAP1 | blue | 0.318153 | 0.113204 |
| C21orf59 | brown | -0.464978 | 0.016692 | DHCR24 | brown | -0.020263347 | 0.9200909 | RRBP1 | blue | 0.320054 | 0.110946 |
| PHYH | brown | -0.462952 | 0.017236 | NANOS1 | brown | -0.020239427 | 0.920184934 | SNORA71C | blue | 0.322613 | 0.107961 |
| SLC35B2 | brown | -0.461723 | 0.017574 | RBM15B | brown | -0.020196992 | 0.920351757 | DDX27 | blue | 0.324039 | 0.106324 |
| AKNA | brown | -0.459659 | 0.018152 | TAS2R39 | brown | -0.019700814 | 0.922302577 | BSG | blue | 0.325808 | 0.104318 |
| ECI1 | brown | -0.458809 | 0.018395 | RAB3GAP1 | brown | -0.01957917 | 0.922780911 | TMEM150B | blue | 0.326518 | 0.103521 |
| HIST1H2BC | brown | -0.456668 | 0.019019 | IBA57 | brown | -0.01904822 | 0.924869045 | HSF1 | blue | 0.327397 | 0.102541 |
| EIF2C3 | brown | 0.456643 | 0.019026 | ZSCAN5A | brown | -0.018437984 | 0.927269598 | TELO2 | blue | 0.327808 | 0.102085 |
| ARMCX5 | brown | 0.454202 | 0.019759 | GPR32 | brown | -0.01841626 | 0.92735507 | COPE | blue | 0.329486 | 0.100239 |
| IL4R | brown | -0.452272 | 0.020354 | GANAB | brown | -0.018357167 | 0.927587566 | C6orf89 | blue | 0.329625 | 0.100087 |
| STOML2 | brown | -0.44866 | 0.021506 | IFNA13 | brown | -0.018208109 | 0.928174045 | PLD3 | blue | 0.329781 | 0.099916 |
| C10orf2 | brown | 0.448502 | 0.021558 | RHEB | brown | -0.017523336 | 0.930868808 | MTHFR | blue | 0.330495 | 0.099141 |
| PDK3 | brown | -0.44499 | 0.022732 | ENTPD4 | brown | -0.013314902 | 0.947445349 | PAF1 | blue | 0.334566 | 0.094802 |
| TUBB | brown | -0.442609 | 0.023557 | PHF12 | brown | -0.012675337 | 0.949966559 | FXYD5 | blue | 0.334654 | 0.094709 |
| TIMM8B | brown | -0.438222 | 0.02514 | MRM1 | brown | -0.01172064 | 0.953730913 | PELP1 | blue | 0.336361 | 0.092934 |
| PRPS2 | brown | -0.433506 | 0.026938 | PILRA | brown | -0.011566881 | 0.954337273 | UBE2D4 | blue | 0.337298 | 0.09197 |
| RWDD2B | brown | -0.431347 | 0.027795 | STX6 | brown | -0.010948235 | 0.956777202 | TMEM79 | blue | 0.337699 | 0.09156 |
| BLVRA | brown | -0.431094 | 0.027896 | ACTN4 | brown | -0.010330943 | 0.959212172 | CORO1B | blue | 0.340252 | 0.088981 |
| GOT2 | brown | -0.430574 | 0.028107 | IL1RN | brown | -0.010011544 | 0.960472213 | HSPBP1 | blue | 0.340942 | 0.088293 |
| MRPL52 | brown | -0.430525 | 0.028126 | LOC349196 | brown | -0.009778458 | 0.961391807 | FZD5 | blue | 0.342838 | 0.086425 |
| PHF21A | brown | -0.429805 | 0.028419 | RPS15AP10 | brown | -0.009500398 | 0.962488895 | SF3B4 | blue | 0.343131 | 0.086139 |
| TNFSF13 | brown | -0.427828 | 0.029238 | RASGRP2 | brown | -0.009483047 | 0.962557359 | HMGCL | blue | 0.34385 | 0.085439 |

**Table 6. Gene module membership and correlation with growth rate (48/72h), with adjusted p values for each module**

| | | | | | | | | | | | |
|---|---|---|---|---|---|---|---|---|---|---|---|
| KLF10 | brown | -0.427661 | 0.029308 | MRPL43 | brown | -0.008735056 | 0.965508907 | ASB1 | blue | 0.347142 | 0.082293 |
| USP4 | brown | 0.42611 | 0.029964 | CCL2 | brown | -0.008225907 | 0.967518248 | C19orf6 | blue | 0.34766 | 0.081807 |
| SKP2 | brown | -0.425103 | 0.030396 | SUMO1P1 | brown | -0.007561803 | 0.970139406 | SFXN5 | blue | 0.349343 | 0.08024 |
| CDYL2 | brown | -0.425012 | 0.030436 | HIST1H2AK | brown | -0.00507732 | 0.979947897 | ABCD1 | blue | 0.350791 | 0.07891 |
| C4orf48 | brown | -0.423606 | 0.031048 | ANXA2P2 | brown | -0.00506096 | 0.980012497 | SEC61A1 | blue | 0.351308 | 0.07844 |
| STAC3 | brown | -0.42218 | 0.03168 | HIST1H3B | brown | -0.004281235 | 0.98309143 | PKD1P1 | blue | 0.353126 | 0.076802 |
| ARFRP1 | brown | -0.421409 | 0.032026 | FKSG29 | brown | -0.004275825 | 0.983112797 | C2CD2L | blue | 0.35547 | 0.074728 |
| EMC1 | brown | -0.419167 | 0.033048 | NDE1 | brown | -0.003974531 | 0.984302595 | CLK3 | blue | 0.355889 | 0.074362 |
| OSTM1 | brown | -0.41889 | 0.033176 | FZD7 | brown | -0.003410636 | 0.986529483 | DAP | blue | 0.35691 | 0.073475 |
| CYBB | brown | -0.41855 | 0.033334 | LOC644961 | brown | -0.001029395 | 0.995934177 | OXER1 | blue | 0.357326 | 0.073117 |
| NSF | brown | -0.417331 | 0.033904 | CEBPA-AS1 | brown | 0.002603797 | 0.98971595 | B9D2 | blue | 0.360505 | 0.070419 |
| CDKN1B | brown | -0.415403 | 0.034823 | TMEM81 | brown | 0.00380967 | 0.984953638 | ZNF335 | blue | 0.362117 | 0.06908 |
| TBC1D2B | brown | -0.415048 | 0.034994 | ZNF33A | brown | 0.004258596 | 0.983180832 | TRIM23 | blue | 0.363352 | 0.068068 |
| MLYCD | brown | -0.414847 | 0.035092 | FAM133B | brown | 0.004881036 | 0.980722946 | FAM131A | blue | 0.363945 | 0.067586 |
| SMG1 | brown | 0.4129 | 0.036045 | TFPI2 | brown | 0.005052935 | 0.980044185 | SF3A1 | blue | 0.364268 | 0.067325 |
| ZBED6 | brown | 0.411286 | 0.036851 | CDC27 | brown | 0.007211531 | 0.97152202 | CCM2 | blue | 0.364311 | 0.067291 |
| MICAL2 | brown | -0.410581 | 0.037208 | OR4F21 | brown | 0.007715499 | 0.969532756 | CCDC9 | blue | 0.365182 | 0.06659 |
| SRSF3 | brown | -0.410216 | 0.037393 | GSPT2 | brown | 0.008434735 | 0.966694091 | SEC24C | blue | 0.366433 | 0.065593 |
| NLRC4 | brown | -0.409765 | 0.037624 | DUX4L4 | brown | 0.009651237 | 0.961893751 | YPEL4 | blue | 0.367139 | 0.065035 |
| COBRA1 | brown | -0.407145 | 0.038985 | SMG1P1 | brown | 0.009721509 | 0.961616496 | WDR81 | blue | 0.367264 | 0.064937 |
| ICAM1 | brown | -0.404383 | 0.040462 | LOC84989 | brown | 0.011032572 | 0.956444557 | MMACHC | blue | 0.36831 | 0.064119 |
| FBXO7 | brown | -0.403857 | 0.040748 | LOC440944 | brown | 0.011538219 | 0.954450308 | MAPK1IP1L | blue | 0.369378 | 0.063293 |
| ZNF354C | brown | -0.403393 | 0.041002 | CST3 | brown | 0.013039419 | 0.948531262 | SUV39H1 | blue | 0.371118 | 0.061963 |
| SAPCD1 | brown | -0.402815 | 0.041319 | NDST2 | brown | 0.013047778 | 0.948498312 | SLCO3A1 | blue | 0.372452 | 0.060958 |
| DCTN3 | brown | -0.401675 | 0.041952 | DGCR11 | brown | 0.013427008 | 0.947003467 | TRMT2A | blue | 0.373452 | 0.060213 |
| STRA13 | brown | -0.401624 | 0.041981 | HIST1H3J | brown | 0.01470164 | 0.941980454 | IPPK | blue | 0.373928 | 0.059861 |
| IK | brown | -0.401139 | 0.042252 | DNMT3A | brown | 0.014764518 | 0.94173272 | NCAPD2 | blue | 0.374058 | 0.059766 |
| ATP6V1G1 | brown | -0.400155 | 0.042808 | LOC644656 | brown | 0.01626264 | 0.935831903 | KIAA0226 | blue | 0.374888 | 0.059156 |
| NDUFA2 | brown | -0.398791 | 0.043587 | AP4S1 | brown | 0.01798965 | 0.929033656 | FAM122B | blue | 0.375564 | 0.058663 |
| LACTB2 | brown | -0.398435 | 0.043793 | SLC35A4 | brown | 0.020361865 | 0.919703623 | SLC16A3 | blue | 0.375773 | 0.058512 |
| CXXC5 | brown | -0.396765 | 0.044766 | CHCHD4 | brown | 0.021011694 | 0.917149571 | EDEM2 | blue | 0.383517 | 0.053104 |
| APOL6 | brown | -0.396397 | 0.044982 | ARF1 | brown | 0.021725848 | 0.91434363 | ZNF324 | blue | 0.389535 | 0.049178 |
| TNFAIP3 | brown | -0.394831 | 0.045914 | C1orf85 | brown | 0.02218998 | 0.912520569 | PRICKLE3 | blue | 0.392322 | 0.047438 |
| SAMM50 | brown | -0.393996 | 0.046417 | OR51T1 | brown | 0.022987379 | 0.909389485 | TSSC4 | blue | 0.39335 | 0.046809 |
| SRSF2 | brown | -0.393914 | 0.046467 | ARF3 | brown | 0.023844354 | 0.906025941 | TOR2A | blue | 0.394014 | 0.046406 |
| OC1002718 | brown | 0.393883 | 0.046485 | FZD2 | brown | 0.024749724 | 0.902474175 | C22orf13 | blue | 0.396493 | 0.044926 |
| EID3 | brown | 0.393617 | 0.046647 | ZNF830 | brown | 0.025255586 | 0.900490468 | CHFR | blue | 0.396746 | 0.044777 |
| HSPD1 | brown | 0.392251 | 0.047482 | L3MBTL3 | brown | 0.025538293 | 0.899382106 | CTDSP1 | blue | 0.398703 | 0.043638 |
| TBC1D10A | brown | -0.391434 | 0.047987 | OR11H2 | brown | 0.02734767 | 0.892292817 | ARID3A | blue | 0.399234 | 0.043333 |
| DUT | brown | -0.391187 | 0.048141 | ACTR3BP2 | brown | 0.028393655 | 0.888198209 | PTBP1 | blue | 0.403456 | 0.040967 |
| CENPJ | brown | -0.391146 | 0.048166 | DNAJC1 | brown | 0.028721006 | 0.886917333 | MEPCE | blue | 0.407147 | 0.038984 |
| KAT8 | brown | -0.38805 | 0.050124 | C5orf55 | brown | 0.028923494 | 0.886125168 | C19orf10 | blue | 0.407477 | 0.03881 |
| KLF7 | brown | -0.38757 | 0.050434 | SLC19A2 | brown | 0.028966191 | 0.885958142 | CECR5 | blue | 0.408043 | 0.038514 |
| F8A1 | brown | -0.387405 | 0.05054 | MPPE1 | brown | 0.028995404 | 0.885843867 | ACADS | blue | 0.412613 | 0.036187 |
| HLA-F | brown | -0.386758 | 0.05096 | ANKRD37 | brown | 0.029203714 | 0.885029074 | BAG3 | blue | 0.419856 | 0.032731 |
| NDUFS4 | brown | -0.385988 | 0.051463 | HSDL1 | brown | 0.029377781 | 0.88434831 | FAM211A | blue | 0.429364 | 0.0286 |
| DAGLB | brown | -0.385421 | 0.051836 | DNAJB7 | brown | 0.029493004 | 0.883897722 | CHST7 | blue | 0.432151 | 0.027473 |
| LOC730102 | brown | -0.384729 | 0.052294 | COPG1 | brown | 0.029619061 | 0.883404808 | FAM195B | blue | 0.449958 | 0.021086 |
| RTFDC1 | brown | -0.384608 | 0.052375 | PIGO | brown | 0.030069155 | 0.881645181 | ABHD4 | blue | 0.453595 | 0.019944 |
| RPL8 | brown | -0.383351 | 0.053216 | KIAA0754 | brown | 0.031136668 | 0.877473975 | CNNM4 | blue | 0.455452 | 0.019381 |
| WDR36 | brown | -0.383108 | 0.05338 | KLHL15 | brown | 0.031509921 | 0.876016274 | PPP2CB | blue | 0.457521 | 0.018769 |
| ANAPC15 | brown | -0.382664 | 0.053679 | TMEM203 | brown | 0.03182919 | 0.874769715 | MBD1 | blue | 0.475299 | 0.014132 |
| MED30 | brown | -0.382424 | 0.053843 | RPRD1A | brown | 0.033245543 | 0.869243233 | NDUFV1 | blue | 0.487512 | 0.01153 |
| SCARNA5 | brown | -0.381653 | 0.054369 | LINC00115 | brown | 0.033984584 | 0.866361924 | BTN2A1 | blue | 0.51607 | 0.006959 |
| FKBP8 | brown | -0.37992 | 0.055565 | RPS10 | brown | 0.034151175 | 0.86571266 | ABHD16A | blue | 0.53013 | 0.005342 |
| HSD17B10 | brown | -0.379806 | 0.055644 | MFSD3 | brown | 0.034543122 | 0.864185446 | FOXK2 | brown | -0.531768 | 0.005176 |

**Table 6. Gene module membership and correlation with growth rate (48/72h), with adjusted p values for each module**

| | | | | | | | | | | | |
|---|---|---|---|---|---|---|---|---|---|---|---|
| MICU1 | brown | -0.379611 | 0.055781 | CXADRP2 | brown | 0.034687323 | 0.863623691 | FAM102A | brown | -0.503835 | 0.008684 |
| C1QC | brown | -0.379473 | 0.055877 | TAS2R20 | brown | 0.034790395 | 0.863222195 | MICAL2 | brown | -0.467165 | 0.016119 |
| DDIT3 | brown | -0.379389 | 0.055935 | ANKRD20A9 | brown | 0.034920221 | 0.862716536 | TP53INP1 | brown | -0.42844 | 0.028982 |
| STAP1 | brown | -0.377933 | 0.056962 | INTS1 | brown | 0.035040402 | 0.862248491 | SLC39A6 | brown | -0.415508 | 0.034773 |
| ZBTB49 | brown | -0.377247 | 0.057451 | PIDD | brown | 0.036273011 | 0.857450722 | CTNS | brown | -0.4125 | 0.036243 |
| ID3 | brown | -0.376796 | 0.057774 | SCGB3A1 | brown | 0.036458066 | 0.856730845 | C12orf5 | brown | -0.410742 | 0.037126 |
| RPL13AP20 | brown | -0.376667 | 0.057867 | ZNF259 | brown | 0.036994224 | 0.854645777 | MAPKAP1 | brown | -0.406837 | 0.039147 |
| ARL15 | brown | -0.376404 | 0.058056 | KCNQ1OT1 | brown | 0.037174411 | 0.853945261 | ADD3 | brown | -0.404326 | 0.040493 |
| ALG9 | brown | -0.376401 | 0.058058 | USP17L5 | brown | 0.037693747 | 0.851926835 | TRAPPC11 | brown | -0.404196 | 0.040563 |
| ACTR5 | brown | -0.376401 | 0.058058 | REXO1L2P | brown | 0.038517316 | 0.848727861 | MTCH1 | brown | -0.397207 | 0.044506 |
| DTD1 | brown | -0.376185 | 0.058214 | SDHAF1 | brown | 0.038989933 | 0.846893136 | ARL15 | brown | -0.396719 | 0.044793 |
| TP53I3 | brown | -0.37484 | 0.059191 | RBM4B | brown | 0.039174945 | 0.84617512 | DNAJB2 | brown | -0.395858 | 0.045301 |
| CEBPG | brown | -0.372658 | 0.060804 | PAFAH1B2 | brown | 0.039721523 | 0.844054599 | PDK3 | brown | -0.392943 | 0.047057 |
| ARL6IP1 | brown | -0.372513 | 0.060912 | SCGB1A1 | brown | 0.040877257 | 0.839574292 | PTPRF | brown | -0.386214 | 0.051315 |
| SLC2A4RG | brown | 0.372457 | 0.060955 | EBPL | brown | 0.041014911 | 0.839040987 | MTAP | brown | -0.377451 | 0.057305 |
| NFATC1 | brown | -0.371311 | 0.061817 | OR1N2 | brown | 0.041319231 | 0.83786222 | LY6G5C | brown | -0.376435 | 0.058033 |
| BST1 | brown | -0.370766 | 0.06223 | FAM173A | brown | 0.042495676 | 0.833308562 | SNX19 | brown | -0.375453 | 0.058745 |
| DNAH1 | brown | -0.370338 | 0.062556 | AHCY | brown | 0.042735516 | 0.832380856 | ORAI3 | brown | -0.373 | 0.060549 |
| RRAS | brown | -0.37006 | 0.062769 | FSD1L | brown | 0.043244114 | 0.830414309 | CASP6 | brown | -0.372308 | 0.061066 |
| UBE2F | brown | -0.369458 | 0.063231 | LOC1001909 | brown | 0.043702548 | 0.828642583 | CHSY1 | brown | -0.37133 | 0.061802 |
| PGM2 | brown | -0.368612 | 0.063885 | RPL13AP6 | brown | 0.044027403 | 0.827387597 | COL6A1 | brown | -0.371003 | 0.06205 |
| INSIG2 | brown | -0.366244 | 0.065743 | SIRPB1 | brown | 0.044973417 | 0.823735304 | LAMC1 | brown | -0.365267 | 0.066521 |
| CXCL2 | brown | -0.365915 | 0.066004 | NCSTN | brown | 0.046522425 | 0.817762788 | PDCD2 | brown | -0.362876 | 0.068457 |
| BTAF1 | brown | 0.365141 | 0.066622 | TSN | brown | 0.048498383 | 0.810158507 | UTP11L | brown | -0.362705 | 0.068597 |
| COPS5 | brown | -0.364821 | 0.066879 | LINC00622 | brown | 0.049812205 | 0.805111638 | FRMD4B | brown | -0.362656 | 0.068637 |
| TRADD | brown | -0.364579 | 0.067074 | PSIMCT-1 | brown | 0.049908659 | 0.80474142 | ZNF564 | brown | -0.362034 | 0.069149 |
| SART1 | brown | -0.363527 | 0.067925 | ANXA2P3 | brown | 0.050226544 | 0.803521584 | APEX2 | brown | -0.361377 | 0.069692 |
| ALDH1A1 | brown | -0.363405 | 0.068025 | INPP5E | brown | 0.050815471 | 0.801262845 | TMEM216 | brown | -0.359987 | 0.070854 |
| NBPF14 | brown | 0.362102 | 0.069093 | OCLM | brown | 0.051567084 | 0.798382417 | VPS53 | brown | -0.355997 | 0.074267 |
| LMO4 | brown | -0.361714 | 0.069414 | KRTAP5-7 | brown | 0.053537697 | 0.790842688 | LACTB2 | brown | -0.355655 | 0.074566 |
| CA5B | brown | -0.361704 | 0.069422 | DPM3 | brown | 0.055419695 | 0.783659104 | ALOX5 | brown | -0.355313 | 0.074866 |
| SSU72 | brown | -0.36147 | 0.069615 | TMOD2 | brown | 0.055565629 | 0.783102791 | FYTTD1 | brown | -0.354103 | 0.075932 |
| UQCRC2 | brown | -0.360897 | 0.070091 | AMBRA1 | brown | 0.056246618 | 0.78050817 | CSNK1G1 | brown | -0.353239 | 0.0767 |
| DDX19A | brown | -0.360268 | 0.070618 | KRTAP5-4 | brown | 0.05628919 | 0.780346044 | PRPS2 | brown | -0.352677 | 0.077204 |
| SETD8 | brown | -0.359822 | 0.070992 | NFE2L1 | brown | 0.057953 | 0.774016848 | QDPR | brown | -0.351645 | 0.078134 |
| WDYHV1 | brown | -0.359698 | 0.071097 | ADAMTSL4 | brown | 0.059031738 | 0.76992078 | PET112 | brown | -0.348914 | 0.080638 |
| SNX19 | brown | -0.358566 | 0.072055 | HIST1H4I | brown | 0.05970077 | 0.767383423 | MRPL45 | brown | -0.348257 | 0.081249 |
| LCMT1 | brown | -0.358331 | 0.072255 | OR2T27 | brown | 0.060767272 | 0.76334347 | YOD1 | brown | -0.34819 | 0.081311 |
| SCRIB | brown | -0.358275 | 0.072303 | ZFAND2A | brown | 0.060949882 | 0.762652337 | PDIA4 | brown | -0.346576 | 0.082828 |
| MS4A14 | brown | -0.35782 | 0.072692 | LRRC37A4P | brown | 0.061252926 | 0.76150579 | GUK1 | brown | -0.345527 | 0.083826 |
| POLR1D | brown | -0.357596 | 0.072885 | LOC389791 | brown | 0.064317185 | 0.74994031 | SULT1C2 | brown | -0.34532 | 0.084023 |
| AXL | brown | -0.357087 | 0.073322 | SYF2 | brown | 0.065291643 | 0.746273297 | PRKCSH | brown | -0.345021 | 0.08431 |
| ATP5F1 | brown | -0.356683 | 0.073672 | NQO2 | brown | 0.066160521 | 0.743008138 | SLC36A1 | brown | -0.342754 | 0.086507 |
| NCL | brown | -0.356518 | 0.073815 | ADAM1A | brown | 0.066298748 | 0.742489091 | CPOX | brown | -0.341627 | 0.087614 |
| STUB1 | brown | -0.356471 | 0.073855 | NR2C2AP | brown | 0.066559595 | 0.741509904 | HEATR6 | brown | -0.341612 | 0.087629 |
| LCP2 | brown | -0.355344 | 0.074838 | LDHAL6B | brown | 0.066844672 | 0.740440207 | FEM1A | brown | -0.340157 | 0.089075 |
| COX8A | brown | -0.354809 | 0.075308 | GAS5 | brown | 0.066918633 | 0.740162762 | NHEJ1 | brown | -0.33955 | 0.089684 |
| NSRP1 | brown | -0.353834 | 0.076171 | SETD1B | brown | 0.067456985 | 0.73814422 | RPL39 | brown | -0.336692 | 0.092592 |
| CUL9 | brown | -0.353808 | 0.076194 | HSD3BP4 | brown | 0.067458175 | 0.73813976 | WFS1 | brown | -0.336086 | 0.093218 |
| LOC338758 | brown | -0.35339 | 0.076565 | TMEM110 | brown | 0.067555244 | 0.737775983 | ARRDC4 | brown | -0.335879 | 0.093432 |
| ZNF362 | brown | -0.35293 | 0.076977 | PIGP | brown | 0.068405228 | 0.734592922 | ALG9 | brown | -0.335455 | 0.093873 |
| LY86 | brown | -0.35263 | 0.077246 | AURKAPS1 | brown | 0.069025888 | 0.732271346 | USP21 | brown | -0.335404 | 0.093926 |
| IGSF6 | brown | -0.352603 | 0.07727 | HTR7P1 | brown | 0.069557333 | 0.730285298 | PMS2 | brown | -0.335249 | 0.094088 |
| PPFIA1 | brown | 0.351399 | 0.078357 | NGFRAP1 | brown | 0.069904809 | 0.728987676 | PHYH | brown | -0.334885 | 0.094467 |
| PDIA4 | brown | -0.351247 | 0.078495 | DALRD3 | brown | 0.07050212 | 0.726758765 | TFRC | brown | -0.333964 | 0.095434 |
| GSTZ1 | brown | -0.350971 | 0.078746 | EIF4EBP3 | brown | 0.070768911 | 0.725763913 | NKAP | brown | -0.33262 | 0.096858 |

**Table 6. Gene module membership and correlation with growth rate (48/72h), with adjusted p values for each module**

| | | | | | | | | | | | |
|---|---|---|---|---|---|---|---|---|---|---|---|
| FARSB | brown | -0.350552 | 0.079128 | SNORA41 | brown | 0.071008463 | 0.724871003 | ZFYVE20 | brown | -0.332361 | 0.097134 |
| STARD7 | brown | -0.350484 | 0.079191 | MGAT3 | brown | 0.072148347 | 0.720627016 | PTBP3 | brown | -0.331092 | 0.098496 |
| DHRS7B | brown | -0.349619 | 0.079986 | TIGD1 | brown | 0.072339849 | 0.71991481 | ZNF362 | brown | -0.330962 | 0.098635 |
| WIPI1 | brown | 0.349077 | 0.080486 | PPP1R3D | brown | 0.072457968 | 0.719475632 | STRA13 | brown | -0.330865 | 0.09874 |
| AK2 | brown | -0.348566 | 0.080961 | BMPR1A | brown | 0.074793415 | 0.710810217 | TBC1D2B | brown | -0.330766 | 0.098847 |
| EML4 | brown | -0.348212 | 0.081291 | EP400 | brown | 0.07540749 | 0.708537523 | KIAA0240 | brown | -0.329821 | 0.099873 |
| SNORA49 | brown | -0.348005 | 0.081483 | FADS2 | brown | 0.075537265 | 0.708057535 | PPP1R8 | brown | -0.329645 | 0.100066 |
| PEX14 | brown | -0.347177 | 0.08226 | NHLRC1 | brown | 0.076277441 | 0.705321996 | NLRC4 | brown | -0.329171 | 0.100583 |
| ACTL6A | brown | -0.347117 | 0.082318 | SNORA33 | brown | 0.07845709 | 0.69728732 | RWDD2B | brown | -0.328896 | 0.100885 |
| HDGFRP2 | brown | -0.347055 | 0.082376 | ACTR1A | brown | 0.079184152 | 0.694614225 | DENND1B | brown | -0.327565 | 0.102354 |
| RNF19B | brown | -0.347038 | 0.082392 | CPD | brown | 0.079839634 | 0.692207353 | VPS37A | brown | -0.327405 | 0.102532 |
| PRKCSH | brown | -0.346616 | 0.08279 | OR52K2 | brown | 0.080110251 | 0.691214519 | HAVCR2 | brown | -0.325498 | 0.104668 |
| NUP205 | brown | -0.346449 | 0.082948 | RBPMS2 | brown | 0.080559343 | 0.689567998 | SOS2 | brown | -0.323808 | 0.106588 |
| IDH3B | brown | -0.346349 | 0.083043 | REXO1L1 | brown | 0.081276904 | 0.686940049 | DHRS9 | brown | -0.323777 | 0.106624 |
| TINF2 | brown | -0.344867 | 0.084458 | RPL30 | brown | 0.081616701 | 0.685696829 | PSMA1 | brown | -0.323659 | 0.106759 |
| C2orf69 | brown | -0.344726 | 0.084593 | HLA-DPA1 | brown | 0.083320197 | 0.679476332 | NUP205 | brown | -0.323511 | 0.106928 |
| PET112 | brown | -0.34465 | 0.084667 | PNPLA2 | brown | 0.083757719 | 0.677881951 | FARSB | brown | -0.322258 | 0.108372 |
| DDX39B | brown | -0.344635 | 0.084681 | CH25H | brown | 0.085433947 | 0.671786167 | MAP3K4 | brown | -0.321849 | 0.108846 |
| TTC5 | brown | -0.343332 | 0.085943 | SMAD6 | brown | 0.086561182 | 0.667698189 | CDYL2 | brown | -0.321538 | 0.109208 |
| ALG8 | brown | -0.342895 | 0.086369 | OR2B11 | brown | 0.086881641 | 0.666537708 | COBRA1 | brown | -0.321448 | 0.109313 |
| DOPEY2 | brown | -0.342191 | 0.087058 | DNAJC30 | brown | 0.087206204 | 0.665363127 | DPP9 | brown | -0.321324 | 0.109457 |
| SHISA4 | brown | -0.341346 | 0.087893 | BOP1 | brown | 0.087463408 | 0.664432864 | CXXC5 | brown | -0.319392 | 0.111728 |
| TMEM222 | brown | -0.340719 | 0.088515 | BLOC1S4 | brown | 0.087491706 | 0.664330544 | TMEM175 | brown | -0.318953 | 0.11225 |
| KCTD20 | brown | 0.340361 | 0.088872 | TARS2 | brown | 0.088540911 | 0.660541002 | SCARNA5 | brown | -0.318619 | 0.112647 |
| SDHC | brown | -0.338195 | 0.091055 | EPM2AIP1 | brown | 0.090088319 | 0.654966941 | ARFRP1 | brown | -0.318578 | 0.112696 |
| RALBP1 | brown | -0.3376 | 0.091661 | KIAA1033 | brown | 0.0909006 | 0.652048131 | CDC42SE2 | brown | -0.318175 | 0.113177 |
| TNFRSF1A | brown | -0.337473 | 0.091791 | ASH1L-AS1 | brown | 0.09110531 | 0.651313321 | SIRT2 | brown | -0.31804 | 0.113338 |
| YWHAB | brown | -0.337105 | 0.092168 | OR5B21 | brown | 0.091438461 | 0.650118154 | C10orf32 | brown | -0.317363 | 0.11415 |
| RPL39 | brown | -0.336751 | 0.092532 | LIN37 | brown | 0.093439362 | 0.642957836 | HIST1H2AC | brown | -0.316563 | 0.115116 |
| C14orf178 | brown | 0.336646 | 0.09264 | ARID3B | brown | 0.093778151 | 0.641748517 | SECISBP2 | brown | -0.315614 | 0.11627 |
| C5orf22 | brown | -0.336124 | 0.093179 | STRBP | brown | 0.094363768 | 0.639660241 | ZNF354B | brown | -0.314948 | 0.117085 |
| RNF166 | brown | -0.336006 | 0.093301 | LOC254100 | brown | 0.094424032 | 0.639445493 | ZCCHC6 | brown | -0.314782 | 0.117288 |
| COX6C | brown | -0.335988 | 0.093319 | RPL38 | brown | 0.096062281 | 0.633618627 | LMAN1 | brown | -0.310958 | 0.122051 |
| TRAPPC3 | brown | -0.335765 | 0.093551 | TOP3A | brown | 0.096101208 | 0.633480429 | TNFSF13 | brown | -0.310545 | 0.122573 |
| PFKFB2 | brown | 0.335679 | 0.09364 | LOC642361 | brown | 0.096167521 | 0.633245035 | CCDC174 | brown | -0.310142 | 0.123086 |
| ISY1 | brown | -0.335472 | 0.093855 | FAM122A | brown | 0.096589791 | 0.631746905 | PAPD4 | brown | -0.309551 | 0.123839 |
| FECH | brown | -0.335467 | 0.09386 | LOC1001323! | brown | 0.098100323 | 0.626399479 | XRCC5 | brown | -0.309216 | 0.124268 |
| PLEKHJ1 | brown | -0.334906 | 0.094445 | CTSF | brown | 0.098723129 | 0.624200025 | FBXO38 | brown | -0.308215 | 0.125554 |
| CISD3 | brown | -0.334667 | 0.094696 | STK40 | brown | 0.099415414 | 0.621758885 | RPL8 | brown | -0.308146 | 0.125644 |
| PHLDA1 | brown | 0.334451 | 0.094922 | E2F6 | brown | 0.099499261 | 0.621463489 | ZNF493 | brown | -0.30805 | 0.125767 |
| PMS2 | brown | -0.334105 | 0.095285 | OR8G2 | brown | 0.099691579 | 0.620786159 | GPN2 | brown | -0.30784 | 0.126038 |
| CASP1 | brown | -0.333601 | 0.095816 | KIAA0664 | brown | 0.099775418 | 0.620490978 | UIMC1 | brown | -0.307685 | 0.126239 |
| MYOF | brown | 0.333295 | 0.096141 | CLDN24 | brown | 0.102764019 | 0.610006456 | ARFGEF2 | brown | -0.306803 | 0.127386 |
| GOLGA2 | brown | -0.333261 | 0.096176 | OR6K3 | brown | 0.102917898 | 0.609468629 | TMF1 | brown | -0.30653 | 0.127742 |
| LMNB2 | brown | -0.333137 | 0.096308 | SNORA66 | brown | 0.103160502 | 0.608621094 | SETD8 | brown | -0.306498 | 0.127784 |
| ZFR | brown | -0.332383 | 0.09711 | HERC4 | brown | 0.104112282 | 0.605300833 | RALBP1 | brown | -0.305783 | 0.128721 |
| STK25 | brown | -0.331539 | 0.098014 | LOC648987 | brown | 0.104469083 | 0.60405811 | ARHGAP12 | brown | -0.305604 | 0.128956 |
| SZT2 | brown | -0.331513 | 0.098042 | OR56A5 | brown | 0.104539712 | 0.603812238 | TMEM176A | brown | -0.305358 | 0.12928 |
| VTI1B | brown | -0.331436 | 0.098125 | C6orf141 | brown | 0.105911994 | 0.599043513 | RAB1A | brown | -0.304453 | 0.130477 |
| TAF11 | brown | -0.330783 | 0.098829 | HMGB2 | brown | 0.106423652 | 0.597269593 | HSPB11 | brown | -0.303013 | 0.132397 |
| UBAC1 | brown | -0.330705 | 0.098913 | LOC606724 | brown | 0.106433557 | 0.597235276 | RALA | brown | -0.302469 | 0.133128 |
| MRPL45 | brown | -0.330014 | 0.099663 | C1orf233 | brown | 0.107543537 | 0.593394803 | C21orf59 | brown | -0.302102 | 0.133624 |
| TUSC2 | brown | -0.329947 | 0.099736 | LOC282997 | brown | 0.107824673 | 0.592423772 | YBX1 | brown | -0.302097 | 0.13363 |
| C22orf25 | brown | -0.329753 | 0.099948 | IFIT5 | brown | 0.107824991 | 0.592422676 | ARL6IP1 | brown | -0.301084 | 0.135003 |
| C14orf28 | brown | 0.329346 | 0.100392 | MON1A | brown | 0.108255342 | 0.590937594 | STK25 | brown | -0.300561 | 0.135716 |
| PLOD3 | brown | -0.328825 | 0.100963 | UTP3 | brown | 0.108276415 | 0.590864914 | TMEM87B | brown | -0.300126 | 0.13631 |

**Table 6. Gene module membership and correlation with growth rate (48/72h), with adjusted p values for each module**

| | | | | | | | | | | | |
|---|---|---|---|---|---|---|---|---|---|---|---|
| EMR3 | brown | -0.328705 | 0.101095 | MMP1 | brown | 0.108661633 | 0.589537016 | FKBP8 | brown | -0.299215 | 0.137561 |
| ATP5SL | brown | -0.328087 | 0.101776 | OR6N1 | brown | 0.109563202 | 0.586434248 | ARL4A | brown | -0.299008 | 0.137848 |
| JMY | brown | -0.327633 | 0.102279 | OR6X1 | brown | 0.109581333 | 0.586371924 | NVL | brown | -0.298389 | 0.138704 |
| QDPR | brown | -0.327577 | 0.102341 | TAS2R14 | brown | 0.109859898 | 0.585414718 | SF1 | brown | -0.298382 | 0.138714 |
| PARP16 | brown | -0.327335 | 0.10261 | MLL2 | brown | 0.110479153 | 0.58328928 | PPP2CA | brown | -0.298315 | 0.138807 |
| BMI1 | brown | -0.326463 | 0.103582 | KRTAP21-2 | brown | 0.111686857 | 0.579153887 | RIOK1 | brown | -0.298001 | 0.139243 |
| ZNF707 | brown | 0.326265 | 0.103805 | RBM12B-AS | brown | 0.112650085 | 0.575864912 | SLC39A11 | brown | -0.297502 | 0.139938 |
| SLC5A3 | brown | 0.326205 | 0.103873 | OR51A4 | brown | 0.114571742 | 0.569328211 | BMI1 | brown | -0.296931 | 0.140737 |
| CXCL1 | brown | -0.325924 | 0.104187 | TBC1D3F | brown | 0.116376837 | 0.563218474 | TIMM8B | brown | -0.296849 | 0.140852 |
| NAGPA | brown | -0.325858 | 0.104262 | LOC1002708( | brown | 0.11651741 | 0.562743923 | MLYCD | brown | -0.296649 | 0.141133 |
| FZD1 | brown | 0.325582 | 0.104573 | TIGD5 | brown | 0.116712739 | 0.562084825 | AK2 | brown | -0.296285 | 0.141645 |
| SIRT2 | brown | -0.325437 | 0.104736 | EXOSC4 | brown | 0.116922158 | 0.561378572 | OSTM1 | brown | -0.296111 | 0.14189 |
| UBN1 | brown | -0.324685 | 0.105588 | OR4M2 | brown | 0.117078399 | 0.560851922 | IP6K2 | brown | -0.295356 | 0.142957 |
| DNAJB2 | brown | -0.324652 | 0.105626 | KRTAP19-2 | brown | 0.117125436 | 0.560693415 | SELT | brown | -0.295017 | 0.143438 |
| GPN1 | brown | -0.324204 | 0.106136 | C15orf62 | brown | 0.118186086 | 0.557124631 | CEBPB | brown | -0.294933 | 0.143557 |
| INPP5F | brown | -0.324038 | 0.106325 | FAM106A | brown | 0.121678336 | 0.545448189 | PRRC2C | brown | -0.294636 | 0.143979 |
| TRIP10 | brown | -0.323316 | 0.107152 | DKFZP434I07 | brown | 0.121835087 | 0.544926774 | OPTN | brown | -0.294524 | 0.14414 |
| ACP5 | brown | 0.323183 | 0.107305 | C21orf33 | brown | 0.122547056 | 0.542561412 | BLVRA | brown | -0.294006 | 0.144879 |
| C16orf52 | brown | 0.32259 | 0.107987 | CDRT15P1 | brown | 0.122667771 | 0.54216084 | STX12 | brown | -0.293812 | 0.145157 |
| RANBP9 | brown | -0.322183 | 0.108458 | MRGPRD | brown | 0.124514019 | 0.53605169 | PARP12 | brown | -0.293765 | 0.145225 |
| RUNX1 | brown | -0.322046 | 0.108618 | CDK3 | brown | 0.124724067 | 0.535358717 | DLG1 | brown | -0.292973 | 0.146363 |
| TSG101 | brown | -0.322029 | 0.108638 | ARGFXP2 | brown | 0.125465042 | 0.532917544 | ZBTB24 | brown | -0.292336 | 0.147283 |
| MRPL32 | brown | -0.321995 | 0.108677 | HABP4 | brown | 0.125962338 | 0.531282155 | SAE1 | brown | -0.291851 | 0.147987 |
| TMEM50A | brown | -0.321644 | 0.109085 | NEAT1 | brown | 0.128205055 | 0.52393664 | ZCCHC9 | brown | -0.291207 | 0.148924 |
| RBM18 | brown | -0.321604 | 0.109132 | SDC3 | brown | 0.129126228 | 0.520933774 | NSRP1 | brown | -0.291206 | 0.148926 |
| SNX3 | brown | -0.321326 | 0.109455 | SHMT2 | brown | 0.130976204 | 0.514928411 | COPB2 | brown | -0.291071 | 0.149123 |
| RAB1B | brown | 0.321111 | 0.109706 | HUS1B | brown | 0.131243446 | 0.514063694 | EMR3 | brown | -0.290497 | 0.149963 |
| C12orf5 | brown | -0.321074 | 0.10975 | GMPR2 | brown | 0.132687002 | 0.509405056 | PAIP1 | brown | -0.288532 | 0.152865 |
| MAP2K5 | brown | -0.320154 | 0.110828 | PRR24 | brown | 0.133662173 | 0.50626978 | RNF10 | brown | -0.288159 | 0.153421 |
| HPRT1 | brown | -0.319415 | 0.111701 | FAM106CP | brown | 0.134715882 | 0.502892734 | SDCBP | brown | -0.287278 | 0.154738 |
| POLR3H | brown | -0.318716 | 0.112531 | SNORA84 | brown | 0.137277683 | 0.494729257 | TMEM131 | brown | -0.286801 | 0.155454 |
| HDHD2 | brown | 0.318647 | 0.112614 | HSPA7 | brown | 0.137676631 | 0.493463972 | F8A1 | brown | -0.286388 | 0.156077 |
| MED24 | brown | -0.318586 | 0.112686 | OR9A2 | brown | 0.138713029 | 0.490184606 | ZFR | brown | -0.284878 | 0.158367 |
| APH1A | brown | -0.317164 | 0.11439 | OR3A1 | brown | 0.140657915 | 0.484060431 | ERCC1 | brown | -0.284541 | 0.158882 |
| ACYP2 | brown | -0.316844 | 0.114776 | TAS2R19 | brown | 0.141059473 | 0.48280085 | UBN1 | brown | -0.284145 | 0.159488 |
| ZNF92 | brown | -0.316164 | 0.1156 | TIGD2 | brown | 0.141166081 | 0.48246673 | OR2A4 | brown | -0.284069 | 0.159605 |
| HDDC2 | brown | -0.315851 | 0.11598 | BRICD5 | brown | 0.141642633 | 0.480974614 | ORMDL1 | brown | -0.283938 | 0.159805 |
| TRPM4 | brown | -0.313759 | 0.118549 | MAS1 | brown | 0.141670005 | 0.480888981 | LMO4 | brown | -0.282636 | 0.161811 |
| GRINA | brown | -0.313448 | 0.118933 | LRRC16A | brown | 0.143186907 | 0.476155636 | KIAA0100 | brown | -0.28157 | 0.163468 |
| TMEM43 | brown | -0.313234 | 0.119199 | OR2T2 | brown | 0.14445741 | 0.472209683 | INIP | brown | -0.281546 | 0.163505 |
| CEBPB | brown | -0.312479 | 0.12014 | RABGAP1L | brown | 0.145365769 | 0.469398883 | SMAD1 | brown | -0.281398 | 0.163736 |
| USP31 | brown | 0.312451 | 0.120174 | IQCB1 | brown | 0.146191443 | 0.466851495 | PSMG2 | brown | -0.280993 | 0.16437 |
| RPS9 | brown | -0.312415 | 0.12022 | LENG9 | brown | 0.146931092 | 0.46457564 | SPG20 | brown | -0.280575 | 0.165025 |
| PTPRA | brown | -0.312179 | 0.120514 | LOC401321 | brown | 0.147052865 | 0.46420151 | MTG1 | brown | -0.280534 | 0.165089 |
| UBE4A | brown | -0.31174 | 0.121066 | MRPL36 | brown | 0.147267278 | 0.463543138 | EIF2A | brown | -0.280278 | 0.165492 |
| ARPC1A | brown | -0.311191 | 0.121757 | FLJ42627 | brown | 0.14750181 | 0.462823547 | C22orf25 | brown | -0.279362 | 0.166936 |
| C10orf32 | brown | -0.310938 | 0.122077 | PEX5 | brown | 0.147947161 | 0.461458731 | PEAK1 | brown | -0.279246 | 0.16712 |
| WHSC1 | brown | 0.310654 | 0.122435 | LOC441454 | brown | 0.149861338 | 0.455616676 | RBM26 | brown | -0.279095 | 0.167359 |
| XPA | brown | -0.310622 | 0.122476 | HIRIP3 | brown | 0.150229341 | 0.45449803 | USP1 | brown | -0.27769 | 0.169597 |
| KDM1A | brown | -0.310567 | 0.122546 | TOP1P2 | brown | 0.150628549 | 0.453286171 | SMYD2 | brown | -0.277677 | 0.169618 |
| BAD | brown | -0.310564 | 0.12255 | CAMSAP3 | brown | 0.151574842 | 0.450420402 | CSTF1 | brown | -0.277072 | 0.170587 |
| CHDH | brown | 0.310428 | 0.122722 | SMAP2 | brown | 0.151965647 | 0.449239699 | C12orf4 | brown | -0.276804 | 0.171018 |
| GNPTG | brown | -0.310191 | 0.123023 | PRORSD1P | brown | 0.154112152 | 0.442784129 | DENND2D | brown | -0.274308 | 0.17507 |
| GPX4 | brown | -0.310097 | 0.123143 | NDUFAF3 | brown | 0.154681117 | 0.441081366 | YTHDC2 | brown | -0.274083 | 0.175437 |
| YBX1 | brown | -0.310004 | 0.123261 | ENG | brown | 0.154984452 | 0.440175006 | MXD1 | brown | -0.273948 | 0.175659 |
| MRPS7 | brown | -0.309705 | 0.123643 | OXER1 | brown | 0.158139347 | 0.430807903 | SYK | brown | -0.272957 | 0.177289 |

**Table 6. Gene module membership and correlation with growth rate (48/72h), with adjusted p values for each module**

| | | | | | | | | | | | |
|---|---|---|---|---|---|---|---|---|---|---|---|
| RRP36 | brown | -0.309179 | 0.124314 | ARFIP1 | brown | 0.159286131 | 0.427430129 | MSANTD3 | brown | -0.272491 | 0.17806 |
| SLC36A1 | brown | -0.308638 | 0.125009 | PPP1R3G | brown | 0.159382979 | 0.427145533 | VCAN | brown | -0.271949 | 0.178958 |
| WWC2 | brown | 0.308181 | 0.125598 | SNORA30 | brown | 0.161220206 | 0.42176638 | GUSB | brown | -0.271641 | 0.179471 |
| ATXN2 | brown | -0.308125 | 0.12567 | CECR2 | brown | 0.161341444 | 0.421412728 | PHIP | brown | -0.270987 | 0.180563 |
| MCRS1 | brown | -0.308121 | 0.125675 | OR11H12 | brown | 0.161480851 | 0.421006278 | STAG2 | brown | -0.270851 | 0.18079 |
| FAH | brown | 0.307843 | 0.126035 | LOC1002876 | brown | 0.16194934 | 0.419641955 | WDTC1 | brown | -0.270596 | 0.181217 |
| TRAPPC4 | brown | -0.307806 | 0.126082 | LOC1001283 | brown | 0.163020634 | 0.416531342 | DDX52 | brown | -0.270401 | 0.181544 |
| ZNF45 | brown | -0.307671 | 0.126257 | TMEM104 | brown | 0.163165791 | 0.416110848 | GLTP | brown | -0.270098 | 0.182052 |
| ZNF491 | brown | 0.307641 | 0.126297 | SMC3 | brown | 0.163371121 | 0.415516446 | TAF11 | brown | -0.269746 | 0.182645 |
| INTS6 | brown | -0.307506 | 0.126471 | TRIM14 | brown | 0.165796198 | 0.408531873 | EML4 | brown | -0.269567 | 0.182947 |
| PPP1R8 | brown | -0.307298 | 0.126742 | PCID2 | brown | 0.165942605 | 0.408112311 | COX14 | brown | -0.268683 | 0.184443 |
| TBC1D2 | brown | -0.306563 | 0.127699 | NANOG | brown | 0.16691303 | 0.405337442 | TRAPPC3 | brown | -0.267706 | 0.186105 |
| MT1M | brown | -0.306532 | 0.12774 | GHDC | brown | 0.16790051 | 0.402524699 | CCR1 | brown | -0.267601 | 0.186284 |
| AFG3L2 | brown | -0.306441 | 0.127858 | PNRC2 | brown | 0.16794077 | 0.402410257 | SMS | brown | -0.266672 | 0.187875 |
| XRN2 | brown | -0.305786 | 0.128717 | TTC30A | brown | 0.170326852 | 0.395660334 | DDX19A | brown | -0.265712 | 0.189531 |
| PCNP | brown | -0.305611 | 0.128947 | SHKBP1 | brown | 0.175191072 | 0.382100718 | HGS | brown | -0.265619 | 0.189692 |
| HSPA4 | brown | 0.305187 | 0.129505 | IFNA1 | brown | 0.176223521 | 0.379257469 | PEX7 | brown | -0.265539 | 0.189829 |
| LMAN1 | brown | -0.304976 | 0.129784 | ARHGAP32 | brown | 0.178436209 | 0.37320528 | ETF1 | brown | -0.265162 | 0.190483 |
| VRK1 | brown | -0.304813 | 0.13 | PDGFB | brown | 0.178485781 | 0.373070336 | ZNF354A | brown | -0.2643 | 0.191983 |
| PIK3CD | brown | -0.304587 | 0.130299 | PTDSS2 | brown | 0.178657015 | 0.372604425 | N4BP2 | brown | -0.264271 | 0.192033 |
| ADCY7 | brown | 0.304479 | 0.130442 | DCAF13P3 | brown | 0.179325029 | 0.370790057 | TMEM248 | brown | -0.264166 | 0.192216 |
| PIK3R2 | brown | -0.30432 | 0.130653 | OR4C46 | brown | 0.180910798 | 0.366503672 | SLC25A13 | brown | -0.264046 | 0.192427 |
| GLTSCR2 | brown | -0.304019 | 0.131054 | BNIP3 | brown | 0.181244334 | 0.365605821 | ATP6V0A2 | brown | -0.263998 | 0.192511 |
| UQCR10 | brown | -0.303916 | 0.13119 | CCDC58 | brown | 0.18180912 | 0.364088403 | POLR2J | brown | -0.263823 | 0.192816 |
| CHMP1A | brown | -0.303843 | 0.131288 | LOC1002718 | brown | 0.183900636 | 0.358501367 | TRIP10 | brown | -0.263301 | 0.193731 |
| MTG1 | brown | -0.30377 | 0.131385 | ABL1 | brown | 0.186551354 | 0.351493725 | FDXACB1 | brown | -0.263018 | 0.194228 |
| GPN2 | brown | -0.303651 | 0.131544 | TSPYL6 | brown | 0.18727476 | 0.34959552 | XPNPEP3 | brown | -0.262477 | 0.19518 |
| LIMK2 | brown | -0.303447 | 0.131817 | SNORA58 | brown | 0.188790614 | 0.345637799 | VSIG10L | brown | -0.262228 | 0.19562 |
| SAE1 | brown | -0.302736 | 0.132769 | TYW1B | brown | 0.189391157 | 0.344077292 | LRP12 | brown | -0.261274 | 0.197311 |
| IMPAD1 | brown | 0.302687 | 0.132835 | PKI55 | brown | 0.192318761 | 0.336530539 | PLGRKT | brown | -0.261251 | 0.197352 |
| WDPCP | brown | 0.302586 | 0.132971 | C14orf178 | brown | 0.193045681 | 0.334672303 | GLTPD1 | brown | -0.260739 | 0.198262 |
| CAPRIN1 | brown | -0.302254 | 0.133418 | LPAR6 | brown | 0.194722219 | 0.330410258 | UCHL3 | brown | -0.260354 | 0.19895 |
| ANKRD32 | brown | -0.30136 | 0.134628 | LOC1004991 | brown | 0.195220646 | 0.329149559 | CBFB | brown | -0.260352 | 0.198954 |
| MLL | brown | -0.300724 | 0.135493 | GPR21 | brown | 0.196928423 | 0.324852212 | EIF4B | brown | -0.260098 | 0.199408 |
| TFCP2 | brown | -0.300471 | 0.135838 | ZNF532 | brown | 0.198544096 | 0.320818334 | WDYHV1 | brown | -0.259713 | 0.200098 |
| ZNF276 | brown | -0.300355 | 0.135997 | UFSP1 | brown | 0.204594217 | 0.305987332 | SLC24A6 | brown | -0.259496 | 0.200487 |
| EIF4B | brown | -0.299565 | 0.13708 | COX17 | brown | 0.206164034 | 0.302210055 | KDM5C | brown | -0.259218 | 0.200988 |
| SNRNP48 | brown | 0.299279 | 0.137474 | ZNF571 | brown | 0.206981821 | 0.300253891 | SEC24A | brown | -0.258585 | 0.202128 |
| CORO2A | brown | -0.298828 | 0.138096 | SEC14L1P1 | brown | 0.210867241 | 0.291068392 | RMND1 | brown | -0.258223 | 0.202783 |
| NIT1 | brown | -0.298557 | 0.138471 | KIAA1109 | brown | 0.214665955 | 0.282261283 | TIPRL | brown | -0.257733 | 0.203673 |
| QARS | brown | -0.298287 | 0.138846 | MEIS3P1 | brown | 0.215862426 | 0.279522842 | FCAR | brown | -0.257504 | 0.204089 |
| CDK8 | brown | 0.298005 | 0.139237 | FANCF | brown | 0.217087776 | 0.276735932 | MTHFD1L | brown | -0.257111 | 0.204804 |
| ORMDL1 | brown | -0.297724 | 0.139629 | GAS5-AS1 | brown | 0.217321109 | 0.276207268 | ZCCHC14 | brown | -0.256957 | 0.205084 |
| ARHGEF9 | brown | -0.29703 | 0.140598 | FAM104B | brown | 0.21811784 | 0.274406979 | CCDC59 | brown | -0.256914 | 0.205164 |
| NCF4 | brown | -0.296967 | 0.140687 | MPDU1 | brown | 0.218817849 | 0.272831463 | CCNH | brown | -0.256255 | 0.206368 |
| MSANTD3 | brown | -0.296816 | 0.140898 | MARS2 | brown | 0.220296366 | 0.269522879 | TRIM38 | brown | -0.255808 | 0.207189 |
| MPHOSPH8 | brown | -0.296765 | 0.140969 | SNORA70B | brown | 0.224062556 | 0.261212199 | ACVRL1 | brown | -0.255073 | 0.208543 |
| CREG1 | brown | -0.296709 | 0.141048 | C17orf59 | brown | 0.225411041 | 0.258277445 | FAM82B | brown | -0.255038 | 0.208607 |
| RER1 | brown | -0.296698 | 0.141064 | FZD1 | brown | 0.225623971 | 0.257816008 | C7orf49 | brown | -0.254491 | 0.209618 |
| NAMPT | brown | 0.295913 | 0.142169 | ARHGAP1 | brown | 0.226170281 | 0.256634562 | CUL9 | brown | -0.253688 | 0.21111 |
| GLTP | brown | -0.295696 | 0.142475 | NDN | brown | 0.227163137 | 0.254496464 | ACTR5 | brown | -0.253581 | 0.211308 |
| SMYD2 | brown | -0.295666 | 0.142518 | RIMKLB | brown | 0.22938806 | 0.249747475 | TMEM5 | brown | -0.253196 | 0.212027 |
| PSMA7 | brown | 0.295347 | 0.14297 | YBEY | brown | 0.229942858 | 0.248572399 | TBC1D10C | brown | -0.252755 | 0.212851 |
| MNT | brown | -0.295185 | 0.143199 | EBP | brown | 0.230613564 | 0.247156678 | PAPSS2 | brown | -0.252542 | 0.213248 |
| HENMT1 | brown | -0.294841 | 0.143688 | SCARNA1 | brown | 0.231174295 | 0.245977168 | NLRP2 | brown | -0.25254 | 0.213252 |
| REEP4 | brown | -0.294808 | 0.143735 | LOC152217 | brown | 0.235342508 | 0.237325387 | TMEM194A | brown | -0.25227 | 0.213759 |

**Table 6. Gene module membership and correlation with growth rate (48/72h), with adjusted p values for each module**

| | | | |
|---|---|---|---|
| FCGR2C | brown | -0.294706 | 0.14388 |
| PPIL2 | brown | -0.294327 | 0.144421 |
| RNF170 | brown | -0.294297 | 0.144463 |
| GAB3 | brown | -0.294113 | 0.144727 |
| C10orf11 | brown | 0.293476 | 0.145639 |
| KCNE3 | brown | -0.293168 | 0.146082 |
| SPIN4 | brown | 0.292807 | 0.146602 |
| PINX1 | brown | -0.2927 | 0.146757 |
| TMC8 | brown | -0.292374 | 0.147228 |
| ZMAT5 | brown | -0.291647 | 0.148284 |
| COMMD2 | brown | -0.290428 | 0.150065 |
| NAGA | brown | 0.289801 | 0.150986 |
| KPNA6 | brown | 0.289723 | 0.151101 |
| UBAC2 | brown | -0.289359 | 0.151639 |
| EFCAB4A | brown | 0.28868 | 0.152645 |
| LOC541471 | brown | -0.288526 | 0.152874 |
| RARA | brown | -0.288038 | 0.1536 |
| PMM1 | brown | 0.287849 | 0.153883 |
| PSME3 | brown | -0.287778 | 0.153989 |
| ZNF354B | brown | -0.287225 | 0.154817 |
| NENF | brown | -0.286291 | 0.156223 |
| SARS | brown | -0.28624 | 0.156299 |
| PTPDC1 | brown | 0.286238 | 0.156303 |
| KIAA0100 | brown | -0.285954 | 0.156733 |
| ZNF354A | brown | -0.285596 | 0.157275 |
| SLC39A13 | brown | -0.28528 | 0.157755 |
| RAF1 | brown | -0.285209 | 0.157862 |
| MGRN1 | brown | -0.283962 | 0.159768 |
| PRUNE | brown | -0.283831 | 0.15997 |
| SUPV3L1 | brown | -0.283154 | 0.161012 |
| FNDC3A | brown | 0.283142 | 0.161029 |
| IL2RG | brown | -0.282584 | 0.161892 |
| SH2B3 | brown | -0.282246 | 0.162416 |
| MRPL38 | brown | -0.281854 | 0.163025 |
| HECTD1 | brown | 0.28161 | 0.163405 |
| GTF3C5 | brown | -0.2814 | 0.163733 |
| RAB38 | brown | 0.279294 | 0.167045 |
| CDK11B | brown | -0.279031 | 0.167461 |
| KCTD6 | brown | 0.278658 | 0.168053 |
| CHD1L | brown | -0.278404 | 0.168458 |
| TMEM248 | brown | -0.278116 | 0.168916 |
| GPR84 | brown | -0.277856 | 0.169331 |
| SCMH1 | brown | 0.277757 | 0.16949 |
| PCNXL3 | brown | 0.277684 | 0.169607 |
| TMEM33 | brown | 0.277333 | 0.170169 |
| CCDC59 | brown | -0.277113 | 0.170521 |
| CUEDC1 | brown | -0.27674 | 0.171121 |
| NBPF1 | brown | -0.276507 | 0.171498 |
| RNF185 | brown | -0.276476 | 0.171547 |
| CMIP | brown | -0.27636 | 0.171735 |
| ZNF330 | brown | 0.274966 | 0.173996 |
| SLC39A3 | brown | -0.274849 | 0.174187 |
| MAX | brown | 0.274836 | 0.174208 |
| TRIM33 | brown | -0.274176 | 0.175285 |
| RCL1 | brown | -0.27177 | 0.179256 |
| RPGR | brown | -0.271763 | 0.179268 |
| YY2 | brown | 0.240238789 | 0.227423128 |
| GOLGA8B | brown | 0.24045533 | 0.226991679 |
| EID2 | brown | 0.244128131 | 0.219757118 |
| USP17L2 | brown | 0.247794459 | 0.212691813 |
| LCMT2 | brown | 0.249918306 | 0.2086702 |
| FAM108A1 | brown | 0.253772736 | 0.201504404 |
| TATDN1 | brown | 0.254716683 | 0.199775502 |
| WBP11 | brown | 0.255844245 | 0.197723659 |
| HNRNPA1 | brown | 0.259953875 | 0.19036793 |
| STX4 | brown | 0.266897498 | 0.178373525 |
| LOC392196 | brown | 0.279351398 | 0.158202519 |
| HYI | brown | 0.279644968 | 0.15774749 |
| CPT2 | brown | 0.281658451 | 0.154651671 |
| ABHD14B | brown | 0.282542788 | 0.153305743 |
| TMED10P1 | brown | 0.290256487 | 0.141918706 |
| TOR1AIP1 | brown | 0.293422403 | 0.137426236 |
| ADPGK | brown | 0.299635788 | 0.128910125 |
| GIMAP5 | brown | 0.313198947 | 0.111667624 |
| RPL7A | brown | 0.319440822 | 0.104333084 |
| RPS8 | brown | 0.332970458 | 0.089675174 |
| LOC285359 | brown | 0.343621031 | 0.079275568 |
| C11orf58 | brown | 0.349738927 | 0.073733233 |
| NPM1 | brown | 0.360967987 | 0.064339516 |
| RIN2 | brown | 0.365139877 | 0.061096121 |
| HYPK | brown | 0.37609258 | 0.053184118 |
| TRIM23 | brown | 0.382287754 | 0.049078577 |
| C14orf169 | brown | 0.425963346 | 0.026733486 |
| NME3 | brown | 0.433820636 | 0.023772381 |
| IL4R | cyan | -0.489327078 | 0.009586303 |
| WDR77 | cyan | -0.481617417 | 0.010971222 |
| TBC1D10C | cyan | -0.444032368 | 0.020328449 |
| CYBB | cyan | -0.408120842 | 0.034573229 |
| PVRL1 | cyan | -0.352731019 | 0.0711333 |
| ZNF329 | cyan | -0.351693734 | 0.072026522 |
| NEURL4 | cyan | -0.34115121 | 0.081600957 |
| COPS6 | cyan | -0.339745026 | 0.082947903 |
| RAB40C | cyan | -0.314495224 | 0.110113903 |
| IKZF1 | cyan | -0.313656812 | 0.111116983 |
| HAUS7 | cyan | -0.305561667 | 0.121152783 |
| NCOA3 | cyan | -0.30308807 | 0.124348081 |
| PDS5B | cyan | -0.292492776 | 0.138734572 |
| RPL32P3 | cyan | -0.28679737 | 0.146947178 |
| C1orf162 | cyan | -0.283511965 | 0.151840315 |
| STXBP2 | cyan | -0.268472241 | 0.175728463 |
| C19orf60 | cyan | -0.257946798 | 0.193936353 |
| MPV17L2 | cyan | -0.256491593 | 0.196552229 |
| EIF3G | cyan | -0.253364027 | 0.202256149 |
| PDIA5 | cyan | -0.232314597 | 0.243589955 |
| IKBKB | cyan | -0.223061066 | 0.263405722 |
| SMG7 | cyan | -0.220606191 | 0.268832848 |
| NDUFS7 | cyan | -0.214838121 | 0.281866189 |
| TACC1 | cyan | -0.209690085 | 0.293832365 |
| LSM12 | cyan | -0.203619282 | 0.308347929 |
| RHOT2 | cyan | -0.203422149 | 0.308826616 |
| TLE3 | cyan | -0.200183494 | 0.31675677 |
| GOLPH3 | cyan | -0.195947615 | 0.327316051 |
| TFCP2 | brown | -0.252252 | 0.213794 |
| SNX3 | brown | -0.252052 | 0.214169 |
| SPNS1 | brown | -0.252025 | 0.214219 |
| ZC3H12D | brown | -0.251251 | 0.215677 |
| SECISBP2L | brown | -0.251026 | 0.216103 |
| PHF11 | brown | -0.25082 | 0.216493 |
| GCLC | brown | -0.250647 | 0.216819 |
| SLC23A2 | brown | -0.250017 | 0.218016 |
| GOLGA2 | brown | -0.249847 | 0.218339 |
| ACOX1 | brown | -0.249156 | 0.219657 |
| TIPIN | brown | -0.248594 | 0.220734 |
| DERL2 | brown | -0.248377 | 0.22115 |
| MS4A7 | brown | -0.247775 | 0.222308 |
| PRCP | brown | -0.247077 | 0.223655 |
| MRPS25 | brown | -0.246879 | 0.224039 |
| FHIT | brown | -0.246808 | 0.224176 |
| HLA-F | brown | -0.246466 | 0.22484 |
| DAXX | brown | -0.24564 | 0.226447 |
| IFNGR2 | brown | -0.245588 | 0.226549 |
| CYB5A | brown | -0.245003 | 0.227693 |
| CHST14 | brown | -0.244903 | 0.227887 |
| VTI1A | brown | -0.24466 | 0.228364 |
| PLOD3 | brown | -0.244485 | 0.228707 |
| TIMELESS | brown | -0.243961 | 0.229739 |
| TPRKB | brown | -0.243813 | 0.23003 |
| TUSC2 | brown | -0.24268 | 0.23227 |
| NCL | brown | -0.241969 | 0.233683 |
| RER1 | brown | -0.241839 | 0.233943 |
| PRDM2 | brown | -0.241606 | 0.234409 |
| SIPA1L1 | brown | -0.241357 | 0.234906 |
| C3AR1 | brown | -0.241251 | 0.235118 |
| CANX | brown | -0.240928 | 0.235765 |
| ICAM1 | brown | -0.240775 | 0.23607 |
| UBAC2 | brown | -0.240695 | 0.236231 |
| MRPL44 | brown | -0.240617 | 0.236388 |
| CREG1 | brown | -0.240284 | 0.237057 |
| MGRN1 | brown | -0.240267 | 0.23709 |
| MYO7B | brown | -0.239723 | 0.238186 |
| GAB2 | brown | -0.239379 | 0.238881 |
| MAP2K3 | brown | -0.2393 | 0.239041 |
| NUS1 | brown | -0.239228 | 0.239186 |
| ARNTL | brown | -0.239121 | 0.239403 |
| LRCH3 | brown | -0.239013 | 0.239622 |
| TTC23 | brown | -0.238769 | 0.240115 |
| BET1L | brown | -0.238636 | 0.240387 |
| USP16 | brown | -0.238133 | 0.241408 |
| TBC1D2 | brown | -0.238039 | 0.241599 |
| C3 | brown | -0.238029 | 0.241621 |
| LOC731275 | brown | -0.237869 | 0.241946 |
| ME2 | brown | -0.2377 | 0.242291 |
| EPSTI1 | brown | -0.237152 | 0.24341 |
| C17orf75 | brown | -0.237056 | 0.243607 |
| AEN | brown | -0.236698 | 0.24434 |
| ARL16 | brown | -0.236598 | 0.244546 |
| UBAC1 | brown | -0.23551 | 0.246785 |
| SERPINB8 | brown | -0.235265 | 0.247293 |

**Table 6. Gene module membership and correlation with growth rate (48/72h), with adjusted p values for each module**

| | | | |
|---|---|---|---|
| ERLEC1 | brown | -0.271491 | 0.179721 |
| RIC8A | brown | -0.270986 | 0.180564 |
| AGPAT2 | brown | -0.270844 | 0.180802 |
| SMAD7 | brown | 0.270812 | 0.180856 |
| CDK5RAP3 | brown | -0.270619 | 0.181177 |
| AP5Z1 | brown | -0.270508 | 0.181363 |
| PARP12 | brown | -0.270494 | 0.181387 |
| BCAP31 | brown | -0.269911 | 0.182367 |
| HACL1 | brown | -0.269816 | 0.182527 |
| ISPD | brown | 0.269594 | 0.182901 |
| TPMT | brown | -0.269463 | 0.183122 |
| SNX27 | brown | -0.26943 | 0.183179 |
| OAS2 | brown | -0.269374 | 0.183272 |
| ABCG1 | brown | -0.269288 | 0.183419 |
| GLTPD1 | brown | -0.269207 | 0.183555 |
| IP6K2 | brown | -0.268515 | 0.184727 |
| THOC3 | brown | -0.268204 | 0.185257 |
| PSMD6 | brown | -0.267779 | 0.185981 |
| WRAP73 | brown | -0.267622 | 0.186249 |
| POLE3 | brown | -0.267615 | 0.18626 |
| YOD1 | brown | -0.267105 | 0.187132 |
| RAB2B | brown | 0.267102 | 0.187138 |
| VTI1A | brown | -0.266871 | 0.187534 |
| CCDC144B | brown | 0.266379 | 0.18838 |
| TMF1 | brown | -0.265454 | 0.189977 |
| TAB1 | brown | 0.264965 | 0.190825 |
| UTP11L | brown | -0.264581 | 0.191492 |
| KIAA0513 | brown | -0.264004 | 0.192499 |
| NRBP1 | brown | -0.263979 | 0.192544 |
| STRN | brown | -0.263888 | 0.192702 |
| ERMAP | brown | -0.263774 | 0.192901 |
| DVL2 | brown | -0.263557 | 0.193281 |
| DOK3 | brown | -0.263552 | 0.19329 |
| CDC37 | brown | -0.263434 | 0.193497 |
| RPS15AP10 | brown | 0.263213 | 0.193885 |
| CD46 | brown | -0.262993 | 0.194271 |
| TMBIM6 | brown | -0.26166 | 0.196626 |
| SERPINB9 | brown | -0.261602 | 0.196728 |
| TAOK2 | brown | -0.261434 | 0.197026 |
| ING1 | brown | -0.261089 | 0.19764 |
| CLEC2B | brown | -0.260731 | 0.198276 |
| PDK1 | brown | -0.260347 | 0.198963 |
| PITHD1 | brown | -0.260212 | 0.199204 |
| ACSL1 | brown | -0.259404 | 0.200652 |
| SMARCD1 | brown | -0.259258 | 0.200915 |
| CD68 | brown | -0.258986 | 0.201406 |
| METTL23 | brown | -0.25881 | 0.201723 |
| CCDC174 | brown | -0.258734 | 0.20186 |
| IDH3G | brown | 0.258132 | 0.202948 |
| PSMB2 | brown | 0.257739 | 0.203661 |
| SDCBP | brown | -0.257618 | 0.203881 |
| IL17RB | brown | -0.256169 | 0.206526 |
| ATXN10 | brown | -0.256065 | 0.206718 |
| RIOK1 | brown | -0.255769 | 0.20726 |
| WDR77 | brown | -0.255433 | 0.207879 |
| CS | brown | -0.254504 | 0.209594 |

| | | | |
|---|---|---|---|
| RSBN1 | cyan | -0.192107762 | 0.337071083 |
| VPS41 | cyan | -0.184814014 | 0.35607744 |
| REXO4 | cyan | -0.184409111 | 0.357150773 |
| LONP2 | cyan | -0.183680038 | 0.359088247 |
| ZFAND3 | cyan | -0.183676364 | 0.359098027 |
| PTPRE | cyan | -0.181539347 | 0.364812744 |
| TPCN2 | cyan | -0.179864578 | 0.369328373 |
| CISD2 | cyan | -0.172191413 | 0.390430723 |
| C9orf89 | cyan | -0.170638552 | 0.394783343 |
| PSENEN | cyan | -0.168615023 | 0.400496346 |
| COG5 | cyan | -0.16479892 | 0.411396196 |
| ZBTB33 | cyan | -0.160370559 | 0.424249385 |
| TP53I13 | cyan | -0.157831914 | 0.43171589 |
| HM13 | cyan | -0.156914431 | 0.434431804 |
| CAPZA1 | cyan | -0.152066358 | 0.448935698 |
| PLBD1 | cyan | -0.151845931 | 0.449601211 |
| PLD2 | cyan | -0.147225852 | 0.463670299 |
| BZW2 | cyan | -0.147099425 | 0.464058501 |
| MTHFD2 | cyan | -0.146350416 | 0.466361856 |
| PHACTR4 | cyan | -0.146068137 | 0.467231464 |
| TAOK3 | cyan | -0.142157892 | 0.479363959 |
| AKAP8 | cyan | -0.140815771 | 0.483565078 |
| HNRNPUL1 | cyan | -0.140464464 | 0.484667829 |
| SDE2 | cyan | -0.138251356 | 0.491644069 |
| OVCA2 | cyan | -0.131562752 | 0.513031443 |
| KPNA3 | cyan | -0.129900998 | 0.518414614 |
| KIAA0947 | cyan | -0.127116727 | 0.5274951 |
| OAZ1 | cyan | -0.124055059 | 0.537567325 |
| SERINC3 | cyan | -0.122332405 | 0.54327404 |
| TRAPPC8 | cyan | -0.121737441 | 0.545251557 |
| PRKAB1 | cyan | -0.114043198 | 0.571122789 |
| PPP1R18 | cyan | -0.111143898 | 0.581011475 |
| TERF2 | cyan | -0.105531214 | 0.600365128 |
| MILR1 | cyan | -0.083799911 | 0.677728271 |
| ARPC5 | cyan | -0.07564875 | 0.707645281 |
| ENOPH1 | cyan | -0.075163855 | 0.709438926 |
| RBM4 | cyan | -0.073508543 | 0.715573328 |
| BIRC6 | cyan | -0.072546191 | 0.719147667 |
| LSM14A | cyan | -0.061673457 | 0.759915549 |
| WSB1 | cyan | -0.060311324 | 0.765069884 |
| GLRX2 | cyan | -0.054515798 | 0.787107159 |
| ZDHHC12 | cyan | -0.053271908 | 0.791858561 |
| ANKRD36BP | cyan | -0.04737773 | 0.814469212 |
| WDR11 | cyan | -0.047288897 | 0.814811144 |
| YTHDF2 | cyan | -0.04106035 | 0.838864959 |
| SMARCAL1 | cyan | -0.037943699 | 0.850955707 |
| MAST2 | cyan | -0.03730739 | 0.853428344 |
| SENP5 | cyan | -0.034254648 | 0.865309433 |
| C11orf84 | cyan | -0.033602338 | 0.867851985 |
| PPP3R1 | cyan | -0.029212723 | 0.884993838 |
| SSR3 | cyan | -0.028461023 | 0.887934587 |
| PICK1 | cyan | -0.026149165 | 0.896987788 |
| MED27 | cyan | -0.022127085 | 0.912767588 |
| C9orf64 | cyan | -0.018707528 | 0.926209186 |
| CNPY2 | cyan | -0.018566527 | 0.926763882 |
| MCAT | cyan | -0.011121563 | 0.95609356 |

| | | | |
|---|---|---|---|
| CMIP | brown | -0.235088 | 0.247659 |
| SEC11A | brown | -0.235061 | 0.247713 |
| TEP1 | brown | -0.234931 | 0.247983 |
| PLA2G4C | brown | -0.234927 | 0.247992 |
| ATM | brown | -0.234414 | 0.249056 |
| STK39 | brown | -0.234299 | 0.249295 |
| VIM | brown | -0.233566 | 0.250822 |
| RBM42 | brown | -0.232083 | 0.25393 |
| SCRIB | brown | -0.231991 | 0.254124 |
| WTAP | brown | -0.231785 | 0.254556 |
| RAPGEF2 | brown | -0.230722 | 0.256804 |
| TROVE2 | brown | -0.230452 | 0.257376 |
| SAMD9L | brown | -0.23006 | 0.25821 |
| PARG | brown | -0.229834 | 0.25869 |
| TGFBR2 | brown | -0.229833 | 0.258691 |
| TTC31 | brown | -0.229331 | 0.259761 |
| PTP4A1 | brown | -0.22921 | 0.260021 |
| IL10RA | brown | -0.228697 | 0.261118 |
| RSL24D1 | brown | -0.228588 | 0.261351 |
| DNLZ | brown | -0.228288 | 0.261996 |
| NKTR | brown | -0.227501 | 0.263688 |
| MAP2K5 | brown | -0.226766 | 0.265275 |
| SMARCA5 | brown | -0.226606 | 0.265623 |
| AP1S1 | brown | -0.226546 | 0.265753 |
| GART | brown | -0.226098 | 0.266725 |
| MLL | brown | -0.226007 | 0.266922 |
| UFL1 | brown | -0.225688 | 0.267616 |
| CUTC | brown | -0.224925 | 0.26928 |
| RRP36 | brown | -0.224468 | 0.270281 |
| KDM1A | brown | -0.224381 | 0.27047 |
| BLZF1 | brown | -0.222956 | 0.273605 |
| FAM160B2 | brown | -0.222528 | 0.27455 |
| SUDS3 | brown | -0.222391 | 0.274855 |
| SCLY | brown | -0.222178 | 0.275325 |
| CDK11A | brown | -0.221798 | 0.276169 |
| MED24 | brown | -0.221403 | 0.277047 |
| PTPRA | brown | -0.221173 | 0.27756 |
| HDDC2 | brown | -0.220686 | 0.278647 |
| TTC37 | brown | -0.220674 | 0.278672 |
| ESD | brown | -0.220427 | 0.279225 |
| MB21D1 | brown | -0.220113 | 0.279928 |
| ATRAID | brown | -0.220111 | 0.279931 |
| TNFAIP3 | brown | -0.219882 | 0.280445 |
| MPRIP | brown | -0.21985 | 0.280517 |
| HMGA1 | brown | -0.219643 | 0.280982 |
| HTT | brown | -0.219612 | 0.281051 |
| SNX6 | brown | -0.219284 | 0.281788 |
| USP13 | brown | -0.217844 | 0.285039 |
| ANKRD42 | brown | -0.2176 | 0.285594 |
| SRSF9 | brown | -0.217437 | 0.285963 |
| RAPGEF1 | brown | -0.217418 | 0.286005 |
| FECH | brown | -0.216678 | 0.28769 |
| PQBP1 | brown | -0.216374 | 0.288382 |
| C9orf156 | brown | -0.216184 | 0.288817 |
| CLDND1 | brown | -0.215869 | 0.289538 |
| UBXN2A | brown | -0.215861 | 0.289555 |

**Table 6. Gene module membership and correlation with growth rate (48/72h), with adjusted p values for each module**

| | | | | | | | | | | | |
|---|---|---|---|---|---|---|---|---|---|---|---|
| ZC3H7A | brown | -0.25414 | 0.210269 | RBM25 | cyan | -0.009796428 | 0.961320906 | ARPC3 | brown | -0.215659 | 0.290017 |
| TMEM173 | brown | -0.25346 | 0.211533 | PLCG2 | cyan | -0.007976297 | 0.968503399 | SZT2 | brown | -0.215151 | 0.291184 |
| RASSF3 | brown | -0.253455 | 0.211542 | MRTO4 | cyan | -0.003879342 | 0.9846785 | PCYOX1L | brown | -0.21506 | 0.291393 |
| CMPK2 | brown | -0.25345 | 0.211553 | UST | cyan | -0.003763004 | 0.985137927 | PLK3 | brown | -0.213633 | 0.294682 |
| ITGB1BP1 | brown | 0.252961 | 0.212465 | ALDH9A1 | cyan | 0.005600431 | 0.977882426 | URB1 | brown | -0.21355 | 0.294876 |
| DNMT1 | brown | -0.252952 | 0.212482 | EIF3L | cyan | 0.011460954 | 0.954755021 | UTP14A | brown | -0.213411 | 0.295197 |
| VPS9D1 | brown | -0.251719 | 0.214794 | ZNF706 | cyan | 0.024071169 | 0.90513598 | AAED1 | brown | -0.213264 | 0.295537 |
| NDUFAF4 | brown | 0.251042 | 0.216072 | PUM2 | cyan | 0.025224747 | 0.900611386 | FASTKD5 | brown | -0.213083 | 0.295956 |
| FCGRT | brown | -0.251031 | 0.216092 | KTN1 | cyan | 0.026578526 | 0.89530543 | MCL1 | brown | -0.212913 | 0.296352 |
| PKP4 | brown | 0.250703 | 0.216714 | C19orf10 | cyan | 0.026791872 | 0.894469645 | CNDP2 | brown | -0.212782 | 0.296655 |
| LSM14B | brown | -0.250661 | 0.216794 | IPO7 | cyan | 0.028516898 | 0.887715948 | NUCKS1 | brown | -0.212073 | 0.298307 |
| NAB2 | brown | -0.250647 | 0.21682 | RFK | cyan | 0.035008075 | 0.862374385 | GABARAP | brown | -0.211972 | 0.298543 |
| MPI | brown | 0.250325 | 0.21743 | YWHAE | cyan | 0.043925884 | 0.827779742 | TSPAN17 | brown | -0.21188 | 0.298756 |
| SLC15A4 | brown | -0.250195 | 0.217678 | AAMP | cyan | 0.045423383 | 0.821999366 | GCFC1 | brown | -0.211754 | 0.299051 |
| ATP6V0B | brown | -0.24984 | 0.218352 | NCEH1 | cyan | 0.049329827 | 0.806963755 | IDE | brown | -0.210553 | 0.301865 |
| VPS37A | brown | -0.249822 | 0.218387 | NEK4 | cyan | 0.051211029 | 0.799746618 | SESN1 | brown | -0.210372 | 0.302289 |
| VRK3 | brown | -0.249674 | 0.218668 | ADTRP | cyan | 0.052050786 | 0.796530068 | FOSL2 | brown | -0.209923 | 0.303346 |
| MAP1LC3B2 | brown | 0.24953 | 0.218943 | NNT | cyan | 0.052283538 | 0.795639123 | HIATL1 | brown | -0.20902 | 0.305479 |
| NHP2L1 | brown | -0.249238 | 0.219501 | NGRN | cyan | 0.065760297 | 0.744511604 | PTPN11 | brown | -0.208739 | 0.306145 |
| BRPF1 | brown | -0.248738 | 0.220457 | KLHL28 | cyan | 0.069221349 | 0.731540696 | BCL3 | brown | -0.208038 | 0.307807 |
| CLASP2 | brown | -0.248679 | 0.220571 | HSPA8 | cyan | 0.070963918 | 0.725037014 | PARVB | brown | -0.207199 | 0.309806 |
| CBFB | brown | -0.248638 | 0.220648 | ZBTB1 | cyan | 0.074004921 | 0.713731976 | HNRNPH2 | brown | -0.207116 | 0.310004 |
| NHEJ1 | brown | -0.248557 | 0.220804 | FANCC | cyan | 0.074741154 | 0.711003745 | TNFRSF1A | brown | -0.20687 | 0.310592 |
| MRPL44 | brown | -0.24848 | 0.220952 | RNF6 | cyan | 0.081248043 | 0.687045677 | PPM1A | brown | -0.206189 | 0.312224 |
| POLR3C | brown | -0.247999 | 0.221876 | PRPS1 | cyan | 0.090267516 | 0.654322595 | RNF19B | brown | -0.204677 | 0.315862 |
| TFRC | brown | -0.247711 | 0.222431 | LRRC57 | cyan | 0.095243885 | 0.636526833 | EHMT1 | brown | -0.203851 | 0.317863 |
| ZNF410 | brown | 0.247254 | 0.223312 | RLIM | cyan | 0.101591333 | 0.614111642 | SRRM1 | brown | -0.203225 | 0.319382 |
| APIP | brown | -0.24662 | 0.224541 | MKRN1 | cyan | 0.109994596 | 0.584952116 | PPM1H | brown | -0.202717 | 0.320621 |
| ACOX1 | brown | -0.246434 | 0.224901 | RANGAP1 | cyan | 0.111737012 | 0.578982427 | TTPAL | brown | -0.202508 | 0.32113 |
| MBD6 | brown | -0.245769 | 0.226196 | FBXL17 | cyan | 0.115162481 | 0.567325461 | SERPINB9 | brown | -0.202344 | 0.321531 |
| C12orf45 | brown | -0.245518 | 0.226686 | PKNOX1 | cyan | 0.122926879 | 0.541301502 | TBC1D10B | brown | -0.202094 | 0.322141 |
| UBL4A | brown | -0.245147 | 0.22741 | DHX29 | cyan | 0.130555288 | 0.516291809 | MTO1 | brown | -0.202022 | 0.322317 |
| PSD4 | brown | -0.245131 | 0.227442 | KLC4 | cyan | 0.145799208 | 0.46806073 | OCRL | brown | -0.200858 | 0.325174 |
| SYS1 | brown | 0.245041 | 0.227618 | SNAPC3 | cyan | 0.154671552 | 0.441109962 | TTI2 | brown | -0.200849 | 0.325194 |
| ZNF684 | brown | -0.244858 | 0.227976 | TMBIM1 | cyan | 0.156679525 | 0.435128656 | SHROOM3 | brown | -0.200631 | 0.325732 |
| ACP2 | brown | -0.244848 | 0.227996 | C16orf70 | cyan | 0.156954449 | 0.434313151 | JAK1 | brown | -0.200448 | 0.326181 |
| PPP1R11 | brown | -0.244161 | 0.229343 | NCAPG2 | cyan | 0.157745055 | 0.431972611 | MTA2 | brown | -0.200014 | 0.327252 |
| PRKACA | brown | -0.243889 | 0.229879 | AAGAB | cyan | 0.167482578 | 0.403713794 | WDR59 | brown | -0.199821 | 0.32773 |
| RPL10 | brown | -0.243611 | 0.230427 | CNIH | cyan | 0.181746754 | 0.364255782 | SYNC | brown | -0.199614 | 0.328242 |
| TFEB | brown | -0.243142 | 0.231356 | KANSL1 | cyan | 0.196537173 | 0.325833691 | CORO2A | brown | -0.19863 | 0.330681 |
| MRPL22 | brown | -0.243082 | 0.231474 | MED19 | cyan | 0.209085912 | 0.295257361 | PSD4 | brown | -0.198538 | 0.33091 |
| TMEM106A | brown | -0.243074 | 0.231489 | SLC38A2 | cyan | 0.212872779 | 0.286397294 | ERP29 | brown | -0.198228 | 0.331679 |
| ULK2 | brown | 0.242961 | 0.231714 | YTHDC1 | cyan | 0.214394265 | 0.282885489 | NOMO2 | brown | -0.19765 | 0.333121 |
| HNRNPH2 | brown | -0.242934 | 0.231767 | GTPBP5 | cyan | 0.218765827 | 0.27294835 | JMY | brown | -0.196128 | 0.336931 |
| GTF2A1 | brown | 0.242865 | 0.231904 | PPTC7 | cyan | 0.243012524 | 0.221937974 | GID4 | brown | -0.19611 | 0.336976 |
| MIEN1 | brown | 0.24286 | 0.231914 | SMAD5 | cyan | 0.252068463 | 0.204651764 | SDS | brown | -0.196038 | 0.337157 |
| DENND1C | brown | -0.242794 | 0.232044 | GOLGA5 | cyan | 0.288747642 | 0.144096614 | TBCD | brown | -0.195222 | 0.339212 |
| STIM1 | brown | -0.242655 | 0.23232 | STYX | cyan | 0.290356368 | 0.141775376 | DISC1 | brown | -0.194479 | 0.341089 |
| ATXN1 | brown | -0.242514 | 0.232599 | BFAR | cyan | 0.296855816 | 0.132671484 | SGPL1 | brown | -0.194362 | 0.341386 |
| IRAK1 | brown | -0.242277 | 0.23307 | SPAG9 | cyan | 0.307210733 | 0.119056331 | LRRC41 | brown | -0.19403 | 0.342227 |
| NPM1 | brown | 0.241977 | 0.233667 | PIGH | cyan | 0.318032135 | 0.10595611 | SKI | brown | -0.193839 | 0.342712 |
| H1FX | brown | -0.241942 | 0.233737 | PPP2R2A | cyan | 0.344949069 | 0.078046277 | FUS | brown | -0.193275 | 0.344145 |
| RSPO3 | brown | -0.241856 | 0.233909 | EPC2 | cyan | 0.479405228 | 0.011398061 | BRWD3 | brown | -0.193007 | 0.344828 |
| MGAT5 | brown | -0.241428 | 0.234763 | CDC42SE2 | darkgreen | -0.495532586 | 0.008580642 | ARAP2 | brown | -0.192644 | 0.345753 |
| MAP3K4 | brown | -0.241344 | 0.234932 | TUBB4B | darkgreen | -0.490848671 | 0.009331029 | DR1 | brown | -0.191404 | 0.348928 |
| TCF3 | brown | -0.240674 | 0.236273 | C10orf76 | darkgreen | -0.438572719 | 0.022114987 | SEC22B | brown | -0.191181 | 0.349499 |

**Table 6. Gene module membership and correlation with growth rate (48/72h), with adjusted p values for each module**

| | | | | | | | | | | | |
|---|---|---|---|---|---|---|---|---|---|---|---|
| ADAR | brown | -0.240672 | 0.236277 | ZNF28 | darkgreen | -0.362259868 | 0.063321224 | SART1 | brown | -0.190885 | 0.350259 |
| TP53I11 | brown | -0.240446 | 0.236731 | SATB1 | darkgreen | -0.351106489 | 0.072536015 | ARHGEF28 | brown | -0.18901 | 0.355102 |
| HCLS1 | brown | 0.239566 | 0.238504 | LOC1001709 | darkgreen | -0.34156873 | 0.08120425 | STOML2 | brown | -0.188205 | 0.357191 |
| DHX37 | brown | -0.239315 | 0.23901 | ZNF100 | darkgreen | -0.327715825 | 0.095170771 | LOC1005067 | brown | -0.188161 | 0.357305 |
| PCF11 | brown | 0.238972 | 0.239704 | UGP2 | darkgreen | -0.326364395 | 0.096624238 | ZNF45 | brown | -0.188099 | 0.357467 |
| DENND2D | brown | -0.238825 | 0.240002 | PRKCD | darkgreen | -0.320176751 | 0.103492584 | NR1D2 | brown | -0.188086 | 0.357502 |
| DOK1 | brown | 0.238701 | 0.240255 | LOC653501 | darkgreen | -0.313941203 | 0.110775981 | FAM3B | brown | -0.18785 | 0.358116 |
| USP54 | brown | -0.23851 | 0.240642 | ZNF350 | darkgreen | -0.308310426 | 0.117673201 | ZCCHC2 | brown | -0.187509 | 0.359005 |
| SAV1 | brown | -0.238355 | 0.240957 | DDT | darkgreen | -0.298256079 | 0.130767058 | NDUFB9 | brown | -0.186821 | 0.360803 |
| INTS3 | brown | -0.238312 | 0.241045 | NBPF10 | darkgreen | -0.294884952 | 0.135385984 | PIK3R2 | brown | -0.185902 | 0.363213 |
| PLRG1 | brown | -0.238165 | 0.241344 | BEX2 | darkgreen | -0.290962436 | 0.140907907 | ZHX1 | brown | -0.18581 | 0.363454 |
| PPP2CA | brown | -0.237992 | 0.241695 | ABCB7 | darkgreen | -0.28536573 | 0.149065293 | UVRAG | brown | -0.185657 | 0.363856 |
| SMYD4 | brown | -0.236186 | 0.245393 | LOC1001322 | darkgreen | -0.28048374 | 0.156452533 | HDHD3 | brown | -0.185267 | 0.364883 |
| PSMA2 | brown | -0.235645 | 0.246508 | MEF2D | darkgreen | -0.280253477 | 0.156807274 | METTL21B | brown | -0.184178 | 0.367758 |
| CTNNA1 | brown | -0.235566 | 0.246671 | RRP9 | darkgreen | -0.274012185 | 0.166641573 | ZMAT5 | brown | -0.182635 | 0.371855 |
| TBX19 | brown | -0.235258 | 0.247306 | OTUB1 | darkgreen | -0.27328881 | 0.167808879 | GANC | brown | -0.182546 | 0.372092 |
| CANX | brown | -0.234592 | 0.248687 | PHACTR1 | darkgreen | -0.270323026 | 0.17265498 | KLC2 | brown | -0.182492 | 0.372236 |
| GTF2H4 | brown | -0.234386 | 0.249114 | HS3ST3B1 | darkgreen | -0.270230542 | 0.17280766 | SERBP1 | brown | -0.182231 | 0.372933 |
| CCDC137 | brown | -0.234371 | 0.249145 | FAM127A | darkgreen | -0.267127524 | 0.177985432 | STAM | brown | -0.181931 | 0.373733 |
| C2orf42 | brown | -0.233736 | 0.250468 | TM9SF2 | darkgreen | -0.265409467 | 0.180898367 | AP3B1 | brown | -0.181753 | 0.374208 |
| SNX12 | brown | -0.233236 | 0.25151 | TECPR1 | darkgreen | -0.263841744 | 0.18358525 | SERPINB2 | brown | -0.181638 | 0.374518 |
| TSC22D2 | brown | -0.231838 | 0.254445 | ZNF598 | darkgreen | -0.263029358 | 0.184988434 | CLLU1OS | brown | -0.181305 | 0.375409 |
| TTC31 | brown | -0.231616 | 0.254913 | LSM4 | darkgreen | -0.261392965 | 0.187837443 | KCTD9 | brown | -0.180443 | 0.377722 |
| KDM5C | brown | -0.230805 | 0.256628 | PYCR2 | darkgreen | -0.25616946 | 0.197134558 | PLAA | brown | -0.179858 | 0.379295 |
| HTT | brown | -0.229563 | 0.259267 | PPP1R12C | darkgreen | -0.254642746 | 0.199910555 | PPIL2 | brown | -0.179404 | 0.380519 |
| DIMT1 | brown | -0.229508 | 0.259384 | ARMC9 | darkgreen | -0.253567678 | 0.201881331 | TBX19 | brown | -0.17915 | 0.381206 |
| KLHL11 | brown | -0.229351 | 0.25972 | MKRN2 | darkgreen | -0.253185543 | 0.202585037 | SH3GL1 | brown | -0.178975 | 0.381678 |
| SLC24A1 | brown | 0.229053 | 0.260357 | HERC3 | darkgreen | -0.252996608 | 0.202933583 | DDX21 | brown | -0.178916 | 0.381837 |
| TTL | brown | -0.228885 | 0.260716 | TPRA1 | darkgreen | -0.252089558 | 0.204612605 | MRGBP | brown | -0.178794 | 0.382168 |
| KIAA1432 | brown | 0.22865 | 0.261218 | DCTN3 | darkgreen | -0.2500438 | 0.2084342 | INTS3 | brown | -0.178486 | 0.383 |
| GALNT4 | brown | 0.228149 | 0.262294 | KIAA0494 | darkgreen | -0.243927121 | 0.220148994 | INSIG2 | brown | -0.177391 | 0.385973 |
| TMUB1 | brown | -0.227796 | 0.263054 | ZNF625 | darkgreen | -0.242901339 | 0.22215612 | DCXR | brown | -0.177216 | 0.386451 |
| VCAN | brown | -0.227746 | 0.263161 | SLC11A2 | darkgreen | -0.229414806 | 0.249690742 | TMEM185E | brown | -0.176901 | 0.38731 |
| DPP9 | brown | -0.227557 | 0.263568 | RPL23AP7 | darkgreen | -0.229109143 | 0.250339602 | RABGEF1 | brown | -0.176844 | 0.387466 |
| PNPLA4 | brown | -0.227446 | 0.263807 | FAM21B | darkgreen | -0.225957389 | 0.257094539 | OXR1 | brown | -0.17682 | 0.387531 |
| CPNE3 | brown | -0.227294 | 0.264135 | HS3ST3A1 | darkgreen | -0.214693325 | 0.28219845 | ARID4B | brown | -0.176799 | 0.387589 |
| SLC39A11 | brown | -0.226965 | 0.264847 | TOMM7 | darkgreen | -0.198810129 | 0.320157084 | LOC1002884 | brown | -0.176342 | 0.388836 |
| HELZ2 | brown | -0.226606 | 0.265622 | FCGR2A | darkgreen | -0.197826438 | 0.322606314 | MED9 | brown | -0.175847 | 0.390189 |
| ST6GALNAC | brown | -0.226463 | 0.265933 | NOMO3 | darkgreen | -0.196847468 | 0.325055146 | AKAP13 | brown | -0.175613 | 0.39083 |
| PI4KA | brown | -0.226176 | 0.266556 | ZNF440 | darkgreen | -0.195810834 | 0.327660556 | ZNF786 | brown | -0.174227 | 0.39464 |
| CHAF1A | brown | -0.225316 | 0.268426 | LYPLA2 | darkgreen | -0.194812304 | 0.330182186 | PAQR3 | brown | -0.174143 | 0.394871 |
| HAVCR2 | brown | -0.224973 | 0.269174 | TUBA1B | darkgreen | -0.193485342 | 0.333551406 | AKNA | brown | -0.174068 | 0.395079 |
| PSMD2 | brown | -0.224448 | 0.270324 | SLC35E3 | darkgreen | -0.186198806 | 0.352421021 | CXCL1 | brown | -0.173624 | 0.396303 |
| TSPAN17 | brown | -0.22444 | 0.270341 | FAM100A | darkgreen | -0.180388557 | 0.367912092 | IFI44L | brown | -0.173594 | 0.396386 |
| FNBP1 | brown | -0.22429 | 0.27067 | ZNF182 | darkgreen | -0.165906423 | 0.408215976 | RPS6KC1 | brown | -0.172976 | 0.398097 |
| WDR35 | brown | 0.223322 | 0.272797 | LRRC37B | darkgreen | -0.161704185 | 0.420355587 | TSTA3 | brown | -0.172564 | 0.399239 |
| LEO1 | brown | -0.223306 | 0.272834 | PCSK5 | darkgreen | -0.161428074 | 0.421160128 | DNMT1 | brown | -0.172286 | 0.40001 |
| RPS6KA3 | brown | 0.222988 | 0.273533 | TMEM205 | darkgreen | -0.145575593 | 0.468750847 | GRINA | brown | -0.172262 | 0.400077 |
| KLHL28 | brown | 0.222931 | 0.27366 | YIF1A | darkgreen | -0.133922432 | 0.505434632 | ST6GALNAC | brown | -0.170093 | 0.406126 |
| ATG9A | brown | -0.222532 | 0.274541 | BSDC1 | darkgreen | -0.131551895 | 0.513066523 | SCRN3 | brown | -0.169335 | 0.40825 |
| MFN2 | brown | 0.221674 | 0.276446 | IFITM2 | darkgreen | -0.11757262 | 0.559187503 | MCM6 | brown | -0.168719 | 0.409983 |
| ARNTL | brown | -0.221254 | 0.277379 | ZNF765 | darkgreen | -0.092697243 | 0.645609958 | NINJ2 | brown | -0.168454 | 0.410729 |
| EPN2 | brown | 0.221155 | 0.277599 | ARMC10 | darkgreen | -0.064179245 | 0.750459828 | CHPF | brown | -0.168261 | 0.411276 |
| MIDN | brown | -0.221075 | 0.277778 | DHX9 | darkgreen | -0.05640073 | 0.779921309 | DDX60L | brown | -0.167743 | 0.412738 |
| SHFM1 | brown | 0.22103 | 0.277878 | LGALS1 | darkgreen | -0.044173895 | 0.826821801 | ZNF200 | brown | -0.167627 | 0.413066 |
| KHDRBS1 | brown | 0.220973 | 0.278006 | SUPT3H | darkgreen | -0.043741466 | 0.828492213 | SSBP4 | brown | -0.167546 | 0.413294 |

**Table 6. Gene module membership and correlation with growth rate (48/72h), with adjusted p values for each module**

| | | | | | | | | | | | |
|---|---|---|---|---|---|---|---|---|---|---|---|
| RPL12 | brown | -0.220504 | 0.279053 | WASL | darkgreen | -0.040246563 | 0.842018641 | NT5C3L | brown | -0.165634 | 0.418722 |
| TIMM44 | brown | -0.219562 | 0.281165 | ANP32A | darkgreen | -0.038045872 | 0.850558798 | ABHD15 | brown | -0.165536 | 0.418999 |
| PNRC2 | brown | 0.218783 | 0.282918 | CTAGE5 | darkgreen | 0.005730665 | 0.977368223 | CXCL5 | brown | -0.165267 | 0.419766 |
| OTUB1 | brown | -0.217983 | 0.284725 | SEC24D | darkgreen | 0.0640634 | 0.750896213 | CHERP | brown | -0.164834 | 0.421002 |
| CDS2 | brown | 0.217372 | 0.286112 | PMF1 | darkgreen | 0.077826891 | 0.699607149 | SLC41A2 | brown | -0.163984 | 0.423434 |
| ZNF710 | brown | -0.217147 | 0.286621 | MNF1 | darkgreen | 0.091103706 | 0.651319078 | SERTAD2 | brown | -0.163956 | 0.423516 |
| FLJ23867 | brown | 0.215397 | 0.290618 | CLEC4E | darkgreen | 0.099538613 | 0.621324871 | PREX1 | brown | -0.16376 | 0.424077 |
| CDK3 | brown | 0.215 | 0.29153 | POLR3D | darkgreen | 0.118744869 | 0.55524867 | SCARNA2 | brown | -0.163649 | 0.424397 |
| DAAM1 | brown | 0.214172 | 0.293436 | C17orf85 | darkgreen | 0.151484036 | 0.450694981 | ADSL | brown | -0.162538 | 0.42759 |
| OC1005067[illegible] | brown | -0.214087 | 0.293633 | FAM165B | darkgreen | 0.161764135 | 0.420181012 | MARK3 | brown | -0.1624 | 0.42799 |
| UHRF1BP1 | brown | -0.213307 | 0.295439 | LUC7L | darkgreen | 0.1700652 | 0.396397363 | ATP6V0A1 | brown | -0.162085 | 0.428899 |
| KAT7 | brown | 0.213255 | 0.29556 | ZMYM6NB | darkgreen | 0.218839464 | 0.272782907 | CDK17 | brown | -0.161945 | 0.429302 |
| LOC401093 | brown | 0.21323 | 0.295616 | TBC1D3 | darkgreen | 0.241306917 | 0.225300238 | NUP93 | brown | -0.161424 | 0.430808 |
| RPS28 | brown | -0.212408 | 0.297527 | RPS6 | darkgreen | 0.345889408 | 0.077184701 | MBTPS2 | brown | -0.160402 | 0.433769 |
| M6PR | brown | -0.211656 | 0.299279 | ITGB7 | darkgrey | -0.398869251 | 0.039309131 | PLXNC1 | brown | -0.160401 | 0.433774 |
| TTI1 | brown | -0.211593 | 0.299426 | PARVA | darkgrey | -0.398675917 | 0.039413342 | DENND1A | brown | -0.159635 | 0.436002 |
| JAK1 | brown | -0.211407 | 0.299861 | PLXNC1 | darkgrey | -0.365652369 | 0.060706617 | PPAPDC2 | brown | -0.159484 | 0.436441 |
| NACC1 | brown | -0.211074 | 0.300641 | COL6A1 | darkgrey | -0.332022363 | 0.090648586 | TDRD7 | brown | -0.157048 | 0.443569 |
| RBFA | brown | -0.21101 | 0.300792 | GIMAP8 | darkgrey | -0.32553909 | 0.097520012 | MRPS18C | brown | -0.156626 | 0.444812 |
| ATP6AP2 | brown | -0.21096 | 0.30091 | CD5 | darkgrey | -0.317014671 | 0.107140018 | FDFT1 | brown | -0.155717 | 0.447489 |
| CD37 | brown | -0.210608 | 0.301736 | CD3G | darkgrey | -0.293408885 | 0.137445196 | AP2A1 | brown | -0.155285 | 0.448768 |
| ASCC3 | brown | 0.2104 | 0.302224 | CD96 | darkgrey | -0.268045277 | 0.176442896 | ITGB8 | brown | -0.154105 | 0.452263 |
| ELK1 | brown | -0.210311 | 0.302433 | IL2RB | darkgrey | -0.24624877 | 0.215651464 | ACOT13 | brown | -0.152495 | 0.457059 |
| RPUSD4 | brown | -0.209539 | 0.304252 | CORO1A | darkgrey | -0.238314094 | 0.231282121 | YKT6 | brown | -0.152369 | 0.457436 |
| RAB1A | brown | -0.209403 | 0.304572 | EIF4E3 | darkgrey | -0.231750895 | 0.244768145 | CCDC53 | brown | -0.151754 | 0.459273 |
| SRSF6 | brown | 0.208378 | 0.307 | STAT1 | darkgrey | -0.222532276 | 0.264568707 | MED13L | brown | -0.151543 | 0.459905 |
| RNPS1 | brown | -0.208377 | 0.307002 | ZNF626 | darkgrey | -0.210447027 | 0.292053168 | BRD7 | brown | -0.151454 | 0.460173 |
| MCU | brown | -0.207735 | 0.308529 | CD3E | darkgrey | -0.205055944 | 0.304873295 | DECR2 | brown | -0.150762 | 0.462248 |
| MAP2K3 | brown | -0.207298 | 0.309571 | FAM26F | darkgrey | -0.204822547 | 0.30543611 | STOM | brown | -0.150038 | 0.464426 |
| ZFP92 | brown | -0.206422 | 0.311665 | ANXA6 | darkgrey | -0.199752075 | 0.317822516 | CBFA2T2 | brown | -0.148356 | 0.469506 |
| TMEM8A | brown | -0.205959 | 0.312776 | CCR5 | darkgrey | -0.188005579 | 0.347684078 | ZMYND15 | brown | -0.147945 | 0.470752 |
| ANAPC2 | brown | -0.205871 | 0.312987 | NCR3LG1 | darkgrey | -0.16556265 | 0.409201653 | ZNF32 | brown | -0.147898 | 0.470895 |
| MED25 | brown | -0.205758 | 0.313258 | CD3D | darkgrey | -0.145502147 | 0.468977631 | SLC46A3 | brown | -0.147512 | 0.472066 |
| VPS37C | brown | -0.205676 | 0.313456 | IL15RA | darkgrey | -0.138960406 | 0.489403485 | SERTAD3 | brown | -0.147323 | 0.472638 |
| KIAA1143 | brown | -0.205637 | 0.31355 | NNMT | darkgrey | -0.134342267 | 0.504088858 | LSM10 | brown | -0.146988 | 0.473657 |
| DENND1B | brown | -0.205336 | 0.314273 | CCL5 | darkgrey | -0.129899904 | 0.518418168 | USP54 | brown | -0.146787 | 0.474268 |
| CEP192 | brown | -0.205081 | 0.31489 | NKG7 | darkgrey | -0.128283558 | 0.523680407 | SAT1 | brown | -0.146733 | 0.474433 |
| SIRPD | brown | 0.204707 | 0.315791 | HAPLN3 | darkgrey | -0.122418282 | 0.542988881 | DENND1C | brown | -0.145741 | 0.47746 |
| CUL1 | brown | 0.204225 | 0.316958 | JAK3 | darkgrey | -0.120645938 | 0.548888145 | SPTAN1 | brown | -0.1438 | 0.483408 |
| PORCN | brown | 0.204167 | 0.317098 | UBD | darkgrey | -0.118854492 | 0.554880982 | PRMT10 | brown | -0.143694 | 0.483732 |
| HELQ | brown | -0.203419 | 0.318912 | NLRC5 | darkgrey | -0.116676459 | 0.562207216 | ELP2 | brown | -0.142981 | 0.485929 |
| CNIH2 | brown | 0.202299 | 0.321642 | ST6GAL1 | darkgrey | -0.115933353 | 0.564716788 | TMEM140 | brown | -0.141651 | 0.490035 |
| LOC642361 | brown | 0.202289 | 0.321664 | KLHL21 | darkgrey | -0.107141895 | 0.594783238 | PHF2 | brown | -0.141424 | 0.490738 |
| GNPNAT1 | brown | 0.201825 | 0.3228 | CLSTN3 | darkgrey | -0.104438248 | 0.604165465 | N4BP1 | brown | -0.141398 | 0.490819 |
| RPS6KC1 | brown | -0.201465 | 0.323682 | CD2 | darkgrey | -0.1017715 | 0.613480194 | KIAA0586 | brown | -0.140889 | 0.492399 |
| MRPS22 | brown | 0.20144 | 0.323743 | ACAP1 | darkgrey | -0.10138251 | 0.61484386 | ANGEL2 | brown | -0.140653 | 0.493131 |
| MRPL48 | brown | -0.201394 | 0.323856 | IQGAP3 | darkgrey | -0.09600964 | 0.633805529 | FUCA1 | brown | -0.140624 | 0.493219 |
| ZMYND15 | brown | -0.200656 | 0.32567 | JAKMIP2 | darkgrey | -0.092223024 | 0.647306901 | POU2F1 | brown | -0.139832 | 0.495682 |
| MRPL53 | brown | -0.200403 | 0.326293 | MAP4K1 | darkgrey | -0.091486537 | 0.649945752 | SSH1 | brown | -0.139629 | 0.496315 |
| WHAMM | brown | -0.200329 | 0.326476 | RARRES3 | darkgrey | -0.084351639 | 0.675719809 | IL4I1 | brown | -0.139118 | 0.497907 |
| WTAP | brown | -0.200267 | 0.326627 | ZFYVE9 | darkgrey | -0.080642656 | 0.689262696 | CDC37L1 | brown | -0.138426 | 0.50007 |
| CABLES1 | brown | 0.19968 | 0.328077 | GIMAP6 | darkgrey | -0.079862035 | 0.69212515 | GRPEL2 | brown | -0.138322 | 0.500394 |
| NFATC2IP | brown | 0.199349 | 0.328896 | HAGHL | darkgrey | -0.07870902 | 0.696360682 | PLEKHA3 | brown | -0.136092 | 0.507393 |
| CADPS2 | brown | 0.199224 | 0.329207 | PARP11 | darkgrey | -0.076891607 | 0.703054878 | SVIL | brown | -0.135489 | 0.509294 |
| KIAA1430 | brown | 0.19792 | 0.332448 | IL32 | darkgrey | -0.075522366 | 0.708112633 | PPP1R2 | brown | -0.135087 | 0.510562 |
| GCDH | brown | -0.1977 | 0.332995 | CD69 | darkgrey | -0.059650746 | 0.767573062 | TRAF7 | brown | -0.135022 | 0.510769 |

**Table 6. Gene module membership and correlation with growth rate (48/72h), with adjusted p values for each module**

| | | | | | | | | | | | |
|---|---|---|---|---|---|---|---|---|---|---|---|
| ELMO1 | brown | -0.1968 | 0.335244 | IDO1 | darkgrey | -0.05665581 | 0.778950217 | WIPF1 | brown | -0.133845 | 0.514493 |
| CHPF2 | brown | -0.196111 | 0.336973 | LAMB1 | darkgrey | -0.048615659 | 0.809707702 | ARHGEF11 | brown | -0.133704 | 0.51494 |
| SH2B1 | brown | -0.195802 | 0.33775 | HK3 | darkgrey | -0.046071109 | 0.819501921 | RFC2 | brown | -0.133178 | 0.516611 |
| GSE1 | brown | -0.19577 | 0.337832 | NR1H3 | darkgrey | -0.04461713 | 0.825110411 | STARD13 | brown | -0.13303 | 0.517081 |
| CIAO1 | brown | -0.19547 | 0.338586 | SERINC2 | darkgrey | -0.03932933 | 0.845576058 | CPEB3 | brown | -0.131926 | 0.520596 |
| FAM203A | brown | 0.19478 | 0.340328 | LYSMD3 | darkgrey | -0.036295977 | 0.857361377 | TCEB3 | brown | -0.131663 | 0.521436 |
| FAM213B | brown | -0.194502 | 0.341032 | ABI3 | darkgrey | -0.031504352 | 0.87603802 | CD37 | brown | -0.130965 | 0.523666 |
| RIPK3 | brown | -0.193844 | 0.342699 | ITGA4 | darkgrey | -0.024876294 | 0.901977782 | PYGL | brown | -0.129195 | 0.52934 |
| PRPF38B | brown | 0.193649 | 0.343194 | SEMA4D | darkgrey | -0.012153288 | 0.952024862 | PSIP1 | brown | -0.128964 | 0.530082 |
| ARID3B | brown | 0.193359 | 0.343931 | DCBLD2 | darkgrey | 0.004977307 | 0.98034281 | MAN1B1 | brown | -0.128539 | 0.531449 |
| CHCHD7 | brown | -0.19303 | 0.34477 | MMP14 | darkgrey | 0.008669677 | 0.965766911 | DPM1 | brown | -0.128375 | 0.531979 |
| SEC14L1 | brown | 0.192834 | 0.34527 | APOBEC3G | darkgrey | 0.011647723 | 0.954018461 | PMS2CL | brown | -0.128187 | 0.532586 |
| SLC25A13 | brown | -0.192444 | 0.346263 | TUBB2A | darkgrey | 0.029419314 | 0.884185888 | UHRF1BP1 | brown | -0.128175 | 0.532623 |
| SLC38A9 | brown | -0.191885 | 0.347695 | GBP5 | darkgrey | 0.034297514 | 0.865142398 | SPAST | brown | -0.127069 | 0.536196 |
| RPS15 | brown | 0.191445 | 0.348822 | ABCG2 | darkgrey | 0.047438638 | 0.814234787 | ELK1 | brown | -0.126455 | 0.538184 |
| SCAMP2 | brown | 0.191378 | 0.348993 | DNAJC5B | darkgrey | 0.057774099 | 0.774696725 | LSM7 | brown | -0.126066 | 0.539445 |
| RPL5 | brown | -0.190176 | 0.352087 | RYK | darkgrey | 0.101399429 | 0.614784522 | ENPP3 | brown | -0.12386 | 0.546621 |
| RPL36 | brown | -0.189957 | 0.352651 | TRPM2 | darkgrey | 0.107540716 | 0.593404549 | KRTCAP2 | brown | -0.122927 | 0.549669 |
| QRSL1 | brown | -0.189865 | 0.352888 | ACSS1 | darkgrey | 0.114224335 | 0.570507487 | CYP3A5 | brown | -0.122848 | 0.549929 |
| TMEM81 | brown | -0.189845 | 0.352941 | TRIM36 | darkgrey | 0.155264568 | 0.439338914 | UBE2Q2 | brown | -0.122056 | 0.552524 |
| IER5 | brown | 0.188883 | 0.355429 | SIGLEC5 | darkgrey | 0.169929589 | 0.396779662 | SAMHD1 | brown | -0.121261 | 0.555133 |
| IRF3 | brown | -0.187815 | 0.358207 | TUG1 | darkgrey | 0.260506687 | 0.189393098 | CHPF2 | brown | -0.120855 | 0.556468 |
| ANKRD49 | brown | -0.187796 | 0.358257 | PTPN11 | darkorange | -0.468913686 | 0.013615193 | RNF166 | brown | -0.120673 | 0.557067 |
| PLA2G4C | brown | -0.187474 | 0.359098 | SLC25A13 | darkorange | -0.465255489 | 0.014467654 | STAT3 | brown | -0.120167 | 0.558733 |
| ANXA5 | brown | -0.187356 | 0.359404 | STK35 | darkorange | -0.423369304 | 0.027774213 | GLT25D1 | brown | -0.119499 | 0.560937 |
| MAP3K11 | brown | -0.186458 | 0.361754 | PRR14L | darkorange | -0.414590654 | 0.031541734 | PITPNM1 | brown | -0.119162 | 0.562052 |
| TMEM140 | brown | -0.186429 | 0.361831 | ASPH | darkorange | -0.382117661 | 0.049187861 | WDR49 | brown | -0.118008 | 0.56587 |
| WDTC1 | brown | -0.186282 | 0.362216 | TCEB3 | darkorange | -0.365947432 | 0.06048324 | C20orf72 | brown | -0.117685 | 0.566941 |
| SLC16A5 | brown | -0.186104 | 0.362683 | ESCO1 | darkorange | -0.354754706 | 0.069415221 | RELL2 | brown | -0.117155 | 0.568702 |
| ENPP4 | brown | 0.185789 | 0.363509 | UQCRHL | darkorange | -0.324247454 | 0.098934414 | USP30 | brown | -0.117037 | 0.569093 |
| MPP1 | brown | -0.185309 | 0.364773 | ZNF621 | darkorange | -0.316580081 | 0.107648688 | C1orf159 | brown | -0.116953 | 0.569374 |
| EHMT1 | brown | -0.185303 | 0.36479 | DLD | darkorange | -0.296832643 | 0.132703169 | C15orf57 | brown | -0.116848 | 0.569721 |
| POLR2L | brown | 0.184963 | 0.365686 | MLL3 | darkorange | -0.285154092 | 0.149380257 | ITFG3 | brown | -0.116497 | 0.57089 |
| DFFB | brown | -0.184943 | 0.365738 | NEDD1 | darkorange | -0.281990724 | 0.154144979 | UPF1 | brown | -0.115862 | 0.573005 |
| LAMTOR2 | brown | 0.184721 | 0.366324 | HOOK3 | darkorange | -0.278526408 | 0.159486234 | TMEFF1 | brown | -0.115648 | 0.57372 |
| STRADA | brown | 0.184594 | 0.366659 | GIGYF2 | darkorange | -0.268785278 | 0.17520595 | FAM8A1 | brown | -0.114949 | 0.576054 |
| MXD4 | brown | 0.184081 | 0.368014 | MACC1 | darkorange | -0.24575539 | 0.216602009 | MYADM | brown | -0.114645 | 0.577069 |
| TPM3P9 | brown | 0.183725 | 0.368958 | TMX3 | darkorange | -0.241239813 | 0.225433213 | ZC3H7A | brown | -0.11462 | 0.577152 |
| FMO5 | brown | -0.183317 | 0.370042 | THUMPD3 | darkorange | -0.236562914 | 0.234830925 | OAS2 | brown | -0.114232 | 0.57845 |
| PAPOLG | brown | -0.182459 | 0.372325 | KLHL12 | darkorange | -0.232941391 | 0.242284297 | WHAMM | brown | -0.114153 | 0.578716 |
| MYL12A | brown | -0.180883 | 0.376538 | VPS13D | darkorange | -0.232927427 | 0.242313334 | GBGT1 | brown | -0.113986 | 0.579274 |
| TBC1D10B | brown | -0.180773 | 0.376833 | KIAA1598 | darkorange | -0.230204626 | 0.248019228 | PIGS | brown | -0.112867 | 0.583028 |
| TCEB3 | brown | -0.179694 | 0.379736 | RAB11A | darkorange | -0.219520466 | 0.27125593 | FAHD2A | brown | -0.112787 | 0.583299 |
| TMED10 | brown | -0.179647 | 0.379862 | QTRTD1 | darkorange | -0.216706505 | 0.277601176 | MTRF1L | brown | -0.110798 | 0.589998 |
| HNRNPA1 | brown | 0.179492 | 0.380282 | SKIL | darkorange | -0.20547687 | 0.303859902 | CHCHD3 | brown | -0.109817 | 0.593316 |
| MRPS30 | brown | -0.179406 | 0.380513 | MBD2 | darkorange | -0.20282702 | 0.310274525 | RCBTB1 | brown | -0.108041 | 0.599341 |
| PPP2R5D | brown | -0.179089 | 0.381371 | CDYL | darkorange | -0.192317955 | 0.336532605 | THEMIS2 | brown | -0.107465 | 0.601298 |
| YWHAQ | brown | -0.178107 | 0.384029 | FAM190B | darkorange | -0.191713988 | 0.338081269 | ZNF428 | brown | -0.106239 | 0.605477 |
| PCYOX1L | brown | -0.17807 | 0.384131 | EIF3A | darkorange | -0.190826846 | 0.340363808 | E2F4 | brown | -0.105315 | 0.608636 |
| ZNF341 | brown | -0.177978 | 0.384378 | MED16 | darkorange | -0.19004921 | 0.342372204 | TRAPPC2L | brown | -0.104984 | 0.609769 |
| GNL3 | brown | -0.177937 | 0.384492 | MCM7 | darkorange | -0.182769281 | 0.361517227 | PRKAR2B | brown | -0.104893 | 0.610082 |
| MYLIP | brown | -0.17777 | 0.384945 | FUBP1 | darkorange | -0.182033934 | 0.363485425 | SIGLEC1 | brown | -0.104597 | 0.611094 |
| ACAD10 | brown | 0.177212 | 0.386461 | IARS2 | darkorange | -0.169405712 | 0.398258476 | MRPL23 | brown | -0.103964 | 0.613266 |
| CASP10 | brown | -0.176669 | 0.387943 | YARS | darkorange | -0.13287012 | 0.508815585 | DAD1 | brown | -0.103927 | 0.613392 |
| GCLC | brown | -0.176354 | 0.388803 | SNRNP40 | darkorange | -0.128231001 | 0.523851945 | YRDC | brown | -0.103078 | 0.616312 |
| MGAT4B | brown | -0.175674 | 0.390664 | USP47 | darkorange | -0.119713222 | 0.55200457 | RRAGC | brown | -0.102409 | 0.618615 |

**Table 6. Gene module membership and correlation with growth rate (48/72h), with adjusted p values for each module**

| | | | | | | | | | | | |
|---|---|---|---|---|---|---|---|---|---|---|---|
| MED15 | brown | -0.175655 | 0.390716 | ZNF160 | darkorange | -0.097793459 | 0.627484328 | STK35 | brown | -0.101814 | 0.620666 |
| MRAS | brown | -0.175191 | 0.391987 | DEPDC5 | darkorange | -0.077301966 | 0.701541462 | RERE | brown | -0.101623 | 0.621325 |
| SLC35C2 | brown | 0.17509 | 0.392265 | LARP1 | darkorange | -0.067655884 | 0.73739888 | SKIV2L | brown | -0.101425 | 0.622007 |
| GPHN | brown | -0.173786 | 0.395857 | PPP3CA | darkorange | -0.051175818 | 0.799881557 | PRMT2 | brown | -0.100751 | 0.624334 |
| WDR49 | brown | -0.173751 | 0.395954 | GALK1 | darkorange | -0.014280113 | 0.943641357 | LIPJ | brown | -0.099909 | 0.627251 |
| MLXIP | brown | -0.172575 | 0.399206 | GYS1 | darkorange | -0.010594701 | 0.958171707 | SRPRB | brown | -0.099303 | 0.62935 |
| SSH2 | brown | 0.172422 | 0.399632 | ZNF318 | darkorange | 0.007973679 | 0.968513734 | ATF7IP2 | brown | -0.098703 | 0.631434 |
| POLR3D | brown | 0.172319 | 0.399918 | BBX | darkorange | 0.013351791 | 0.947299944 | DNTTIP2 | brown | -0.098051 | 0.633699 |
| RNF10 | brown | -0.172184 | 0.400294 | ZBTB2 | darkorange | 0.041336201 | 0.837796498 | ATXN1 | brown | -0.097534 | 0.6355 |
| GGT7 | brown | -0.1719 | 0.401081 | IER3IP1 | darkorange | 0.058217468 | 0.773012086 | MBD6 | brown | -0.097449 | 0.635794 |
| YIF1B | brown | 0.171548 | 0.402063 | UROS | darkorange | 0.065574319 | 0.745210556 | IFIT3 | brown | -0.096885 | 0.63776 |
| PACS1 | brown | -0.171303 | 0.402744 | AGTRAP | darkorange | 0.132754573 | 0.509187501 | LIG4 | brown | -0.096522 | 0.639027 |
| DPY19L1 | brown | 0.170972 | 0.403667 | AKIP1 | darkorange | 0.138178926 | 0.491873236 | ZNF276 | brown | -0.096134 | 0.640382 |
| LPIN2 | brown | 0.170793 | 0.404168 | PRPF19 | darkorange | 0.157999296 | 0.431221408 | C1orf198 | brown | -0.09606 | 0.640641 |
| IGSF8 | brown | -0.170748 | 0.404294 | TMEM63B | darkorange | 0.163458407 | 0.415263905 | DERL1 | brown | -0.09597 | 0.640955 |
| TRIM5 | brown | 0.170384 | 0.405312 | TAF5L | darkorange | 0.169006553 | 0.399387322 | MMP7 | brown | -0.094867 | 0.644811 |
| ZNF384 | brown | -0.170167 | 0.405918 | SNTB1 | darkorange | 0.170261124 | 0.395845407 | PTCD1 | brown | -0.094345 | 0.646641 |
| CLN6 | brown | 0.169287 | 0.408386 | RAB20 | darkorange | 0.191666853 | 0.33820231 | COA5 | brown | -0.094023 | 0.647771 |
| SDHD | brown | -0.169131 | 0.408825 | VPS25 | darkorange | 0.273336561 | 0.167731648 | BCRP3 | brown | -0.093294 | 0.65033 |
| OLA1 | brown | 0.167895 | 0.412308 | YIPF3 | darkorange | 0.285213165 | 0.149292295 | APBA1 | brown | -0.093223 | 0.650581 |
| AAED1 | brown | -0.167771 | 0.412657 | FBXO25 | darkorange | 0.32881076 | 0.094005264 | ARHGEF3 | brown | -0.093147 | 0.650846 |
| FAM151B | brown | 0.167066 | 0.414651 | EIF4A2 | darkorange | 0.432057454 | 0.024412458 | NAB2 | brown | -0.092726 | 0.652329 |
| TTC7A | brown | -0.165988 | 0.417713 | FIBP | darkorange | 0.440430017 | 0.021493398 | IL23A | brown | -0.092463 | 0.653255 |
| FLYWCH1 | brown | -0.165905 | 0.417949 | SCAMP2 | darkorange | 0.446224521 | 0.019645188 | MKKS | brown | -0.091993 | 0.654909 |
| ARFGEF2 | brown | -0.165902 | 0.417959 | FKRP | darkred | -0.518131565 | 0.005631945 | PIGN | brown | -0.09142 | 0.656926 |
| SNX24 | brown | -0.165719 | 0.418478 | EXOSC10 | darkred | -0.491878866 | 0.009161445 | ITGAE | brown | -0.090911 | 0.658723 |
| AZI2 | brown | -0.165689 | 0.418564 | TFG | darkred | -0.450137348 | 0.018472405 | SP140 | brown | -0.090513 | 0.66013 |
| IFT172 | brown | 0.164928 | 0.420734 | ZNF561 | darkred | -0.415291392 | 0.031226661 | IFI44 | brown | -0.090405 | 0.660512 |
| ACAD11 | brown | 0.164477 | 0.422025 | ZNF639 | darkred | -0.389274734 | 0.044752186 | MAPKAPK5-A | brown | -0.089092 | 0.665159 |
| NPRL3 | brown | -0.163254 | 0.42553 | ZNF559 | darkred | -0.387868848 | 0.045597523 | GUF1 | brown | -0.089089 | 0.66517 |
| SRRM2 | brown | -0.162138 | 0.428746 | GALNT6 | darkred | -0.374925783 | 0.053986618 | EPM2A | brown | -0.089025 | 0.665397 |
| MCMBP | brown | -0.162053 | 0.428991 | KPNA1 | darkred | -0.359479736 | 0.065528261 | SRRM2 | brown | -0.088403 | 0.667603 |
| POMZP3 | brown | 0.16121 | 0.431426 | UBAP1 | darkred | -0.357125371 | 0.067443426 | CDK5RAP3 | brown | -0.087457 | 0.670963 |
| PEX26 | brown | -0.160878 | 0.43239 | ACBD5 | darkred | -0.355418786 | 0.068858452 | H3F3A | brown | -0.087318 | 0.671454 |
| CCNI | brown | -0.160089 | 0.434679 | FBXO11 | darkred | -0.341236482 | 0.081519816 | MX2 | brown | -0.086441 | 0.674577 |
| NFIC | brown | -0.160042 | 0.434816 | MFSD11 | darkred | -0.32333343 | 0.099944556 | ZNF101 | brown | -0.083059 | 0.686661 |
| TMEM64 | brown | -0.15949 | 0.436423 | EIF1AD | darkred | -0.31918633 | 0.104624919 | KCMF1 | brown | -0.08294 | 0.687091 |
| SLC7A5P1 | brown | 0.159245 | 0.437138 | ATP1B1 | darkred | -0.313282223 | 0.111567324 | CHMP4B | brown | -0.081263 | 0.693112 |
| CDC42BPA | brown | 0.158571 | 0.439105 | GPATCH8 | darkred | -0.293745181 | 0.136974059 | ATP6V1F | brown | -0.080709 | 0.695106 |
| ATP6V0A1 | brown | -0.15827 | 0.439985 | TLR5 | darkred | -0.260601831 | 0.189225669 | STAT2 | brown | -0.080663 | 0.69527 |
| EMG1 | brown | 0.158136 | 0.440378 | S100A9 | darkred | -0.255560154 | 0.198239254 | SMCR8 | brown | -0.079896 | 0.698033 |
| PTGES2 | brown | -0.157682 | 0.441709 | ATE1 | darkred | -0.240900851 | 0.226105717 | MFSD9 | brown | -0.079305 | 0.700164 |
| RPL28 | brown | -0.15614 | 0.446242 | AHSA2 | darkred | -0.234592555 | 0.238866943 | C7orf31 | brown | -0.07819 | 0.704194 |
| TRIM26 | brown | -0.155972 | 0.446738 | C4orf29 | darkred | -0.229078577 | 0.250404547 | MTX1 | brown | -0.077706 | 0.705945 |
| PHF2 | brown | -0.154884 | 0.449952 | BTG1 | darkred | -0.221385762 | 0.267101661 | NUP50 | brown | -0.077567 | 0.706447 |
| SNTA1 | brown | -0.154831 | 0.450112 | HMGN3 | darkred | -0.217915827 | 0.274862732 | GVINP1 | brown | -0.076713 | 0.70954 |
| DAPK1 | brown | 0.154717 | 0.450448 | SEH1L | darkred | -0.199039445 | 0.319587768 | DZIP1L | brown | -0.076665 | 0.709715 |
| CTDP1 | brown | -0.153547 | 0.453924 | COG2 | darkred | -0.197028368 | 0.324601782 | ABCF1 | brown | -0.076164 | 0.711531 |
| SETD6 | brown | -0.153482 | 0.454116 | EXD2 | darkred | -0.192997819 | 0.334794462 | NLRP3 | brown | -0.073253 | 0.722121 |
| ENY2 | brown | 0.153343 | 0.454529 | SAMD9 | darkred | -0.191955773 | 0.33746078 | PPRC1 | brown | -0.07313 | 0.722569 |
| PPRC1 | brown | -0.152899 | 0.455854 | IRF2BP2 | darkred | -0.189927139 | 0.34268812 | IGSF8 | brown | -0.07008 | 0.733718 |
| TRIP6 | brown | 0.152751 | 0.456293 | HP1BP3 | darkred | -0.189285972 | 0.344350312 | ARAP1 | brown | -0.069194 | 0.736967 |
| URB2 | brown | 0.15229 | 0.45767 | PAPLN | darkred | -0.18920146 | 0.344569762 | INO80B | brown | -0.06915 | 0.73713 |
| RPAP1 | brown | -0.152185 | 0.457986 | TMEM102 | darkred | -0.186944209 | 0.350462123 | RABEP2 | brown | -0.068802 | 0.738405 |
| B3GNT2 | brown | 0.151206 | 0.460916 | RFX7 | darkred | -0.186621412 | 0.351309628 | GDE1 | brown | -0.06734 | 0.743781 |
| PTCD1 | brown | -0.150654 | 0.462575 | FBXO28 | darkred | -0.18349298 | 0.359586342 | ERN1 | brown | -0.066411 | 0.747199 |

**Table 6. Gene module membership and correlation with growth rate (48/72h), with adjusted p values for each module**

| | | | | | | | | | | | |
|---|---|---|---|---|---|---|---|---|---|---|---|
| PROS1 | brown | 0.150074 | 0.464317 | SCAF11 | darkred | -0.165306046 | 0.409938263 | CLPTM1L | brown | -0.066049 | 0.748536 |
| MAPKAPK5 | brown | 0.149965 | 0.464645 | GNAQ | darkred | -0.162058803 | 0.419323531 | PYGO2 | brown | -0.064268 | 0.755109 |
| FBXL17 | brown | 0.149549 | 0.4659 | NBPF11 | darkred | -0.160850567 | 0.422845628 | AHSA2 | brown | -0.063648 | 0.757403 |
| SHMT1 | brown | 0.148009 | 0.470557 | GEMIN7 | darkred | -0.144225392 | 0.472929024 | PLEKHG3 | brown | -0.063586 | 0.75763 |
| SNRPF | brown | -0.14787 | 0.47098 | DHTKD1 | darkred | -0.143625671 | 0.474791001 | ULK4 | brown | -0.063295 | 0.758709 |
| MTR | brown | 0.147636 | 0.47169 | HNRNPC | darkred | -0.140949764 | 0.483144811 | ASRGL1 | brown | -0.063162 | 0.759199 |
| PHF20 | brown | 0.147034 | 0.473518 | AMMECR1L | darkred | -0.136007996 | 0.49876693 | SLC19A3 | brown | -0.06044 | 0.769298 |
| KLHL36 | brown | -0.14675 | 0.474381 | ZNF518A | darkred | -0.128310494 | 0.523592505 | SLC25A46 | brown | -0.056853 | 0.782656 |
| RNASEH2C | brown | 0.146213 | 0.476019 | RNASEL | darkred | -0.110246326 | 0.584088005 | KIF13A | brown | -0.055604 | 0.787322 |
| PEA15 | brown | -0.145454 | 0.478337 | TMEM183A | darkred | -0.083389769 | 0.679222714 | NDUFA8 | brown | -0.055267 | 0.788584 |
| EMC9 | brown | 0.145163 | 0.479225 | VPS13B | darkred | -0.083000303 | 0.680642913 | PLIN3 | brown | -0.054688 | 0.790749 |
| TRMT61A | brown | -0.144477 | 0.481329 | HBP1 | darkred | -0.074917201 | 0.710351889 | FLCN | brown | -0.054383 | 0.791892 |
| NFU1 | brown | 0.144099 | 0.482487 | SH3PXD2A | darkred | -0.074729559 | 0.711046685 | FBXO32 | brown | -0.054026 | 0.793229 |
| ZNF420 | brown | -0.143884 | 0.483148 | SLFN11 | darkred | -0.066970007 | 0.739970063 | APH1B | brown | -0.053656 | 0.794613 |
| CHMP4B | brown | -0.143773 | 0.48349 | SRRD | darkred | -0.062556691 | 0.756578696 | MKLN1 | brown | -0.052828 | 0.797718 |
| SLC25A46 | brown | -0.143351 | 0.484788 | BTBD8 | darkred | -0.050771859 | 0.80143006 | EPB41L2 | brown | -0.052812 | 0.797779 |
| C14orf119 | brown | -0.143083 | 0.485614 | BEX4 | darkred | -0.037941886 | 0.85096275 | IFI35 | brown | -0.052746 | 0.798024 |
| DPP3 | brown | -0.142666 | 0.486898 | PSMB4 | darkred | -0.033965194 | 0.8664375 | ZC3H7B | brown | -0.052178 | 0.800158 |
| SLC24A6 | brown | -0.141917 | 0.489212 | DDX39A | darkred | -0.031455093 | 0.876230374 | MBD2 | brown | -0.050548 | 0.806283 |
| OC1005061 | brown | 0.141893 | 0.489288 | MGEA5 | darkred | -0.026419033 | 0.895930319 | C4orf34 | brown | -0.050226 | 0.807494 |
| C16orf72 | brown | -0.141878 | 0.489335 | RNMT | darkred | -0.019414397 | 0.923428883 | SLC16A10 | brown | -0.049681 | 0.809543 |
| GNAQ | brown | -0.141778 | 0.489643 | TBC1D9B | darkred | -0.005677353 | 0.977578715 | ZNF496 | brown | -0.049316 | 0.81092 |
| ARID1A | brown | -0.141022 | 0.491985 | C1orf50 | darkred | -0.005177701 | 0.979551541 | NDUFB8 | brown | -0.049285 | 0.811034 |
| LOC93622 | brown | 0.140679 | 0.49305 | TCTEX1D4 | darkred | 0.005529102 | 0.978164056 | PINK1 | brown | -0.048438 | 0.814225 |
| AIF1 | brown | -0.139278 | 0.497408 | UFSP2 | darkred | 0.005736897 | 0.97734362 | ATP10A | brown | -0.047932 | 0.816133 |
| PDE1B | brown | 0.138533 | 0.499733 | GPKOW | darkred | 0.011693198 | 0.953839131 | ZNF90 | brown | -0.046374 | 0.822014 |
| DDX24 | brown | -0.1385 | 0.499838 | RWDD3 | darkred | 0.020019783 | 0.921048437 | LENG8 | brown | -0.045702 | 0.824556 |
| LZTFL1 | brown | 0.137518 | 0.502912 | MIOS | darkred | 0.030362128 | 0.880500106 | TMEM129 | brown | -0.044392 | 0.829511 |
| RAP2A | brown | 0.137484 | 0.503018 | ZC3H14 | darkred | 0.040490302 | 0.84107382 | SRP72 | brown | -0.04366 | 0.832283 |
| ABHD3 | brown | 0.136983 | 0.50459 | ARHGAP18 | darkred | 0.046883802 | 0.816370839 | KLC1 | brown | -0.043068 | 0.834526 |
| C21orf91 | brown | 0.136407 | 0.506403 | PHF3 | darkred | 0.052409064 | 0.795158729 | OST4 | brown | -0.042058 | 0.838356 |
| HYPK | brown | -0.136355 | 0.506564 | PRPF4B | darkred | 0.059885488 | 0.766683275 | CMPK2 | brown | -0.040229 | 0.845298 |
| CPSF6 | brown | -0.136282 | 0.506794 | MTOR | darkred | 0.065880996 | 0.744058095 | PRKACA | brown | -0.038323 | 0.852549 |
| ZNF18 | brown | -0.134471 | 0.512512 | BMP1 | darkred | 0.068507887 | 0.734208771 | UBE2J1 | brown | -0.038035 | 0.853644 |
| FUT8 | brown | 0.13441 | 0.512703 | KAT6A | darkred | 0.069350232 | 0.731059047 | UBQLN4 | brown | -0.037946 | 0.853986 |
| LRR1 | brown | -0.134361 | 0.51286 | DCAF11 | darkred | 0.109359553 | 0.58713449 | TXNDC5 | brown | -0.037556 | 0.855469 |
| COG1 | brown | 0.134236 | 0.513256 | CARS2 | darkred | 0.10973801 | 0.585833468 | PAPOLG | brown | -0.036681 | 0.858805 |
| LRRC41 | brown | -0.134007 | 0.513979 | TXLNA | darkred | 0.126915594 | 0.528154 | C6orf203 | brown | -0.035496 | 0.863324 |
| S100P | brown | 0.133641 | 0.515142 | SLC9A7 | darkred | 0.161389364 | 0.421272989 | NDUFA3 | brown | -0.034448 | 0.867323 |
| PNPLA8 | brown | -0.133206 | 0.516521 | SEC24C | darkred | 0.241066956 | 0.225775995 | ACBD3 | brown | -0.02974 | 0.885327 |
| FLJ31306 | brown | -0.132891 | 0.517523 | ZNF438 | darkred | 0.290680017 | 0.141311657 | PIAS4 | brown | -0.028607 | 0.88967 |
| NOL7 | brown | -0.132399 | 0.51909 | GPD1 | darkred | 0.300816372 | 0.127336516 | KIFC3 | brown | -0.028091 | 0.891649 |
| FASTKD5 | brown | -0.131251 | 0.52275 | A2M | darkred | 0.37511105 | 0.053858565 | ZNF212 | brown | -0.026899 | 0.896223 |
| ARHGEF11 | brown | -0.130915 | 0.523826 | BRI3 | darkred | 0.419467731 | 0.029400997 | FOXO1 | brown | -0.026567 | 0.897495 |
| HIBADH | brown | -0.130914 | 0.523829 | HIST1H1D | darkturquoise | -0.335027376 | 0.087590499 | POLR2D | brown | -0.02626 | 0.898673 |
| PSMA3 | brown | -0.130449 | 0.525318 | B2M | darkturquoise | -0.271237447 | 0.171150463 | AEBP2 | brown | -0.024526 | 0.905336 |
| SH3BP4 | brown | 0.130416 | 0.525422 | SLC25A40 | darkturquoise | -0.229541566 | 0.249421981 | RSBN1 | brown | -0.024189 | 0.906631 |
| ADCK4 | brown | -0.130302 | 0.525788 | KRT8 | darkturquoise | -0.205102744 | 0.304760518 | ZNF777 | brown | -0.024153 | 0.906766 |
| SIPA1L1 | brown | -0.130252 | 0.525948 | SURF1 | darkturquoise | -0.199676728 | 0.318008874 | SRP9 | brown | -0.023924 | 0.907648 |
| TNPO2 | brown | -0.130136 | 0.526319 | HNRNPR | darkturquoise | -0.18148567 | 0.364956968 | ZNF85 | brown | -0.023168 | 0.910555 |
| NDUFA13 | brown | 0.129755 | 0.527541 | SNORA10 | darkturquoise | -0.168470546 | 0.40090602 | LRRC23 | brown | -0.021146 | 0.918333 |
| PRKDC | brown | 0.129071 | 0.52974 | GPR65 | darkturquoise | -0.163090041 | 0.416330251 | BRAP | brown | -0.01848 | 0.928602 |
| CBFA2T2 | brown | -0.128788 | 0.530649 | CTBS | darkturquoise | -0.160033449 | 0.42523677 | EVI5L | brown | -0.017234 | 0.933407 |
| KARS | brown | 0.127268 | 0.53555 | AP1G1 | darkturquoise | -0.157964845 | 0.431323159 | LNPEP | brown | -0.016531 | 0.936118 |
| PPWD1 | brown | 0.127126 | 0.53601 | LGMN | darkturquoise | -0.149345738 | 0.457186428 | BTG2 | brown | -0.016058 | 0.93794 |
| GPATCH3 | brown | 0.125867 | 0.540088 | TERC | darkturquoise | -0.147397208 | 0.463144412 | XAB2 | brown | -0.015371 | 0.940591 |

**Table 6. Gene module membership and correlation with growth rate (48/72h), with adjusted p values for each module**

| | | | | | | | | | | | |
|---|---|---|---|---|---|---|---|---|---|---|---|
| TBPL1 | brown | -0.125586 | 0.541003 | SCARNA16 | darkturquoise | -0.14432522 | 0.472619451 | POLR1A | brown | -0.01526 | 0.941021 |
| PABPC1 | brown | -0.124795 | 0.543573 | RPPH1 | darkturquoise | -0.139616825 | 0.487333824 | ADCK4 | brown | -0.014894 | 0.942431 |
| MXI1 | brown | -0.124646 | 0.544058 | LMBR1 | darkturquoise | -0.136992306 | 0.495635339 | RPAP1 | brown | -0.014295 | 0.944743 |
| FAM89A | brown | -0.124316 | 0.545134 | SNORA63 | darkturquoise | -0.134695161 | 0.502959035 | ZNF429 | brown | -0.013635 | 0.947291 |
| CHUK | brown | 0.124189 | 0.545548 | RFXANK | darkturquoise | -0.127378086 | 0.526639492 | PRR14 | brown | -0.013217 | 0.948904 |
| ZMYM6NB | brown | -0.123598 | 0.547476 | SNORA65 | darkturquoise | -0.124421193 | 0.536358069 | P4HA1 | brown | -0.011082 | 0.95715 |
| KLC2 | brown | -0.122605 | 0.550724 | DEGS1 | darkturquoise | -0.099750114 | 0.620580064 | TTC13 | brown | -0.010622 | 0.958927 |
| TCF25 | brown | -0.12176 | 0.553493 | SCARNA6 | darkturquoise | -0.097179432 | 0.629657348 | NRP2 | brown | -0.009378 | 0.963734 |
| PUSL1 | brown | -0.121138 | 0.555537 | SNORA9 | darkturquoise | -0.092904088 | 0.644870326 | NEK8 | brown | -0.007151 | 0.972344 |
| CASK | brown | 0.121114 | 0.555616 | SNORA16A | darkturquoise | -0.092113831 | 0.647697878 | COX6B1 | brown | -0.006906 | 0.973288 |
| KDM4B | brown | -0.12093 | 0.556219 | SNORA31 | darkturquoise | -0.07616374 | 0.705741982 | RANBP6 | brown | -0.006658 | 0.974249 |
| SETDB1 | brown | 0.120532 | 0.557531 | SNORA5C | darkturquoise | -0.071944414 | 0.721385707 | TMEM14A | brown | -0.005323 | 0.979412 |
| C7orf26 | brown | -0.120127 | 0.558866 | SNORA55 | darkturquoise | -0.068719726 | 0.733416256 | PEX6 | brown | -0.004513 | 0.982543 |
| SLC39A4 | brown | -0.119392 | 0.561289 | ZNF514 | darkturquoise | -0.067423227 | 0.738270745 | MPV17 | brown | -0.003548 | 0.986276 |
| COX10 | brown | 0.119236 | 0.561804 | GGA1 | darkturquoise | -0.060461194 | 0.764502292 | GCC1 | brown | -0.003013 | 0.988344 |
| NRBF2 | brown | -0.119156 | 0.562069 | BPIFB1 | darkturquoise | -0.057546996 | 0.775560018 | NAA38 | brown | -0.002964 | 0.988535 |
| SMG9 | brown | -0.118919 | 0.562853 | TRABD | darkturquoise | -0.051092644 | 0.800200329 | AP2S1 | brown | -0.002388 | 0.990762 |
| FUCA2 | brown | -0.118556 | 0.564053 | SNORA62 | darkturquoise | -0.049829343 | 0.805045855 | TMEM184C | brown | -0.002278 | 0.991187 |
| SLC23A2 | brown | -0.118122 | 0.565492 | SNORA43 | darkturquoise | -0.047217375 | 0.815086469 | BACE2 | brown | -0.001451 | 0.994388 |
| RRNAD1 | brown | 0.118107 | 0.565543 | SNORA71A | darkturquoise | -0.047183009 | 0.815218768 | INTS12 | brown | 0.000816 | 0.996842 |
| FZD7 | brown | 0.118058 | 0.565706 | SNORA6 | darkturquoise | -0.04381921 | 0.828191845 | BARD1 | brown | 0.004858 | 0.981208 |
| MINPP1 | brown | -0.1177 | 0.566894 | CBLL1 | darkturquoise | -0.032560354 | 0.871916046 | C16orf5 | brown | 0.005912 | 0.977132 |
| PLK3 | brown | -0.1171 | 0.568884 | NUP88 | darkturquoise | -0.029457275 | 0.884037441 | PRNP | brown | 0.006734 | 0.973954 |
| PYROXD1 | brown | 0.116881 | 0.569614 | LBR | darkturquoise | -0.024262205 | 0.90438649 | YWHAG | brown | 0.007964 | 0.969202 |
| PFDN5 | brown | 0.116392 | 0.571239 | KRT19 | darkturquoise | -0.021485347 | 0.915288458 | FAM122A | brown | 0.008422 | 0.96743 |
| DHX15 | brown | 0.115346 | 0.574725 | SNORA5A | darkturquoise | -0.01373054 | 0.945807135 | C1orf21 | brown | 0.00892 | 0.965505 |
| WFS1 | brown | -0.115343 | 0.574737 | GTDC1 | darkturquoise | -0.009545382 | 0.962311408 | SYNRG | brown | 0.009609 | 0.962843 |
| POM121C | brown | 0.114936 | 0.576097 | CLK4 | darkturquoise | -0.006880003 | 0.972830723 | SREBF1 | brown | 0.009781 | 0.962176 |
| UTP14A | brown | -0.113355 | 0.581392 | LOC399753 | darkturquoise | -0.003096962 | 0.987768262 | CASP8 | brown | 0.010338 | 0.960024 |
| MIB1 | brown | 0.113003 | 0.582572 | CLCN5 | darkturquoise | -0.002649687 | 0.989534712 | FAM160A2 | brown | 0.010509 | 0.959363 |
| ZNF496 | brown | -0.109942 | 0.592892 | SCARNA12 | darkturquoise | 0.002668474 | 0.989460515 | ITIH5 | brown | 0.012134 | 0.953087 |
| PLCB2 | brown | -0.109794 | 0.593393 | HLX | darkturquoise | 0.005349295 | 0.978874005 | NCAPG2 | brown | 0.013076 | 0.94945 |
| INPP1 | brown | -0.108668 | 0.597208 | NFKB1 | darkturquoise | 0.011397868 | 0.955003818 | GATSL3 | brown | 0.014359 | 0.944498 |
| SLC48A1 | brown | -0.108558 | 0.597584 | TSSC4 | darkturquoise | 0.014781598 | 0.94166543 | CD80 | brown | 0.015362 | 0.940627 |
| PDGFD | brown | 0.108125 | 0.599053 | SNORD17 | darkturquoise | 0.015623151 | 0.938350341 | PEMT | brown | 0.016458 | 0.936399 |
| ATP5G2 | brown | 0.107334 | 0.601744 | AQP9 | darkturquoise | 0.021737549 | 0.914297664 | RNLS | brown | 0.017172 | 0.933645 |
| LSM14A | brown | -0.106739 | 0.603772 | SNORA74A | darkturquoise | 0.023414293 | 0.907713696 | TMEM208 | brown | 0.017937 | 0.930697 |
| SNAP23 | brown | 0.106722 | 0.603829 | SNORD15A | darkturquoise | 0.024984198 | 0.901554626 | POC1B | brown | 0.021436 | 0.917218 |
| ZNF75A | brown | 0.10291 | 0.61689 | SCARNA13 | darkturquoise | 0.02921975 | 0.884966355 | CDC123 | brown | 0.021917 | 0.915366 |
| MANBAL | brown | 0.102655 | 0.617766 | SNORA38 | darkturquoise | 0.03684148 | 0.855239688 | LIN9 | brown | 0.022166 | 0.91441 |
| FCGR1C | brown | -0.102646 | 0.617798 | ADAM10 | darkturquoise | 0.041832799 | 0.835873719 | VAMP4 | brown | 0.023255 | 0.910218 |
| NUDT5 | brown | 0.102354 | 0.618803 | SNORA21 | darkturquoise | 0.060511704 | 0.764311026 | PDE4DIP | brown | 0.02675 | 0.896793 |
| NLRP3 | brown | -0.102304 | 0.618976 | SNORA52 | darkturquoise | 0.074849472 | 0.710602647 | DTNB | brown | 0.026825 | 0.896505 |
| SMARCB1 | brown | -0.101786 | 0.620762 | SNORA7B | darkturquoise | 0.078588484 | 0.696803981 | SIN3B | brown | 0.028793 | 0.888956 |
| ZNF669 | brown | 0.101013 | 0.623429 | KRT7 | darkturquoise | 0.084139925 | 0.67649026 | ATP6V1D | brown | 0.028915 | 0.88849 |
| ANKRD13A | brown | -0.100663 | 0.624639 | SNORA50 | darkturquoise | 0.090834547 | 0.652285295 | BZW1 | brown | 0.029003 | 0.888152 |
| ZNF658 | brown | -0.10049 | 0.625239 | SNORA71B | darkturquoise | 0.095614248 | 0.635210078 | METTL3 | brown | 0.030129 | 0.883836 |
| EIF2C1 | brown | 0.100331 | 0.625789 | SNORA42 | darkturquoise | 0.097594721 | 0.628187325 | ZCCHC17 | brown | 0.033833 | 0.869672 |
| ANKFY1 | brown | -0.098084 | 0.633584 | SNORA28 | darkturquoise | 0.104370078 | 0.604402831 | KIAA0226L | brown | 0.034312 | 0.867841 |
| PINK1 | brown | -0.098084 | 0.633584 | KRT17 | darkturquoise | 0.105174068 | 0.601605839 | CCDC124 | brown | 0.03538 | 0.863766 |
| NF1 | brown | -0.097655 | 0.635079 | SNORD15B | darkturquoise | 0.105623763 | 0.600043796 | SLC25A33 | brown | 0.035619 | 0.862853 |
| HMGA1 | brown | -0.096943 | 0.637557 | SNORA34 | darkturquoise | 0.107001269 | 0.595269695 | GTF2IRD1 | brown | 0.036296 | 0.860269 |
| CHERP | brown | -0.096303 | 0.639792 | MFAP3 | darkturquoise | 0.154977348 | 0.440196221 | FITM2 | brown | 0.036808 | 0.85832 |
| NGLY1 | brown | -0.095913 | 0.641152 | LCN2 | darkturquoise | 0.162059605 | 0.4193212 | LTA4H | brown | 0.038092 | 0.853427 |
| CSTF2 | brown | 0.095759 | 0.641691 | SNORA74B | darkturquoise | 0.16559731 | 0.409102215 | AFTPH | brown | 0.038722 | 0.85103 |
| GOPC | brown | -0.095466 | 0.642718 | CES1 | darkturquoise | 0.16979993 | 0.397145378 | STRADA | brown | 0.038898 | 0.85036 |

**Table 6. Gene module membership and correlation with growth rate (48/72h), with adjusted p values for each module**

| | | | | | | | | | | | |
|---|---|---|---|---|---|---|---|---|---|---|---|
| CRY2 | brown | 0.09459 | 0.645784 | SNORA2A | darkturquoise | 0.182217301 | 0.362994043 | MTMR4 | brown | 0.039364 | 0.848589 |
| CNN2 | brown | -0.094445 | 0.646291 | SNORA46 | darkturquoise | 0.189350902 | 0.344181765 | TMEM63A | brown | 0.039877 | 0.846638 |
| CRTC3 | brown | -0.094072 | 0.6476 | NPM3 | darkturquoise | 0.189759523 | 0.343122189 | IFIT2 | brown | 0.040982 | 0.84244 |
| CDC25B | brown | -0.093924 | 0.648118 | PSMD13 | darkturquoise | 0.21593435 | 0.279358767 | ITGB1BP1 | brown | 0.041873 | 0.839058 |
| NUP214 | brown | -0.093545 | 0.649448 | COASY | darkturquoise | 0.284081344 | 0.150984076 | FDPS | brown | 0.045564 | 0.825078 |
| ANKRD13C | brown | 0.093201 | 0.650659 | OR2A20P | green | -0.53287558 | 0.004212918 | HMGXB3 | brown | 0.046819 | 0.820334 |
| KDM4C | brown | 0.092652 | 0.652588 | RIN1 | green | -0.491631817 | 0.009201875 | KANSL1L | brown | 0.047277 | 0.818605 |
| MRPL49 | brown | -0.092625 | 0.652684 | OR2A9P | green | -0.455045452 | 0.017083185 | SYPL1 | brown | 0.047863 | 0.816396 |
| DCLRE1B | brown | -0.092518 | 0.653059 | FAM219B | green | -0.441848604 | 0.021028306 | GAN | brown | 0.047959 | 0.816033 |
| AEN | brown | -0.091653 | 0.656108 | FASN | green | -0.419945168 | 0.029197894 | ZNF780A | brown | 0.049689 | 0.809513 |
| MESDC2 | brown | 0.091303 | 0.65734 | SNORA57 | green | -0.413022628 | 0.032256035 | BRICD5 | brown | 0.050384 | 0.806898 |
| SLC7A6OS | brown | 0.091301 | 0.657348 | CTBP2 | green | -0.412542185 | 0.03247748 | KANSL2 | brown | 0.050584 | 0.806148 |
| WDFY4 | brown | -0.090842 | 0.658967 | MT1A | green | -0.412421365 | 0.03253336 | CRY2 | brown | 0.050744 | 0.805543 |
| SLC35D1 | brown | -0.090157 | 0.661389 | RTN4IP1 | green | -0.410816851 | 0.033282807 | GPX7 | brown | 0.05097 | 0.804693 |
| TIMM17A | brown | -0.090037 | 0.661815 | RRN3 | green | -0.410759054 | 0.03331006 | CREBBP | brown | 0.051916 | 0.80114 |
| SLC7A6 | brown | -0.088439 | 0.667473 | SPINK1 | green | -0.402154284 | 0.037571732 | DCTD | brown | 0.05307 | 0.79681 |
| RRP8 | brown | -0.086289 | 0.675117 | CTAGE4 | green | -0.398563086 | 0.039474262 | DOCK3 | brown | 0.056498 | 0.783982 |
| DNMBP | brown | -0.085842 | 0.676711 | LOC1002887 | green | -0.385180141 | 0.047249318 | NAA35 | brown | 0.056499 | 0.783977 |
| LTBP3 | brown | 0.085713 | 0.677172 | CDKN1C | green | -0.380851735 | 0.050007247 | AIMP2 | brown | 0.056952 | 0.782286 |
| AP1B1 | brown | 0.085646 | 0.677413 | NUP50 | green | -0.373324235 | 0.055103534 | SLBP | brown | 0.058349 | 0.777075 |
| ARL3 | brown | -0.08525 | 0.678826 | NCOR2 | green | -0.371661242 | 0.056282342 | LIPA | brown | 0.058503 | 0.776501 |
| FAM162A | brown | 0.084841 | 0.680285 | MIR22HG | green | -0.36902704 | 0.058189793 | ZNF544 | brown | 0.059525 | 0.772696 |
| FAM20C | brown | -0.083447 | 0.685273 | OR2A4 | green | -0.35928793 | 0.065682694 | COMMD1 | brown | 0.062535 | 0.761521 |
| FAM109A | brown | -0.082695 | 0.687966 | LINC00338 | green | -0.357866484 | 0.066835964 | ACOT9 | brown | 0.0633 | 0.758691 |
| HMGXB3 | brown | -0.082011 | 0.690423 | KIAA0319 | green | -0.356176772 | 0.068227165 | IFFO1 | brown | 0.063463 | 0.758085 |
| EDF1 | brown | 0.081749 | 0.691363 | MLL4 | green | -0.356052833 | 0.06833008 | CD33 | brown | 0.064087 | 0.755778 |
| WDR24 | brown | 0.080993 | 0.694081 | LCP2 | green | -0.354835802 | 0.069347045 | MAGEH1 | brown | 0.064712 | 0.75347 |
| PSMB3 | brown | 0.080681 | 0.695207 | IGF2BP2 | green | -0.353487268 | 0.070487471 | CDC16 | brown | 0.064853 | 0.752946 |
| CTC1 | brown | -0.080665 | 0.695261 | SCARNA17 | green | -0.352138861 | 0.071642167 | HNRNPD | brown | 0.066378 | 0.747322 |
| H2AFV | brown | 0.080446 | 0.696051 | LIMS2 | green | -0.348848786 | 0.074520604 | SLC35E2 | brown | 0.06696 | 0.745178 |
| RAD54L2 | brown | -0.079135 | 0.70078 | GSTM3 | green | -0.345105385 | 0.077902547 | SULT1A4 | brown | 0.068004 | 0.741338 |
| RPF1 | brown | 0.079082 | 0.70097 | CRISPLD2 | green | -0.342607828 | 0.080223325 | TOR1B | brown | 0.068661 | 0.738926 |
| TSC22D4 | brown | -0.078902 | 0.701621 | LOC439994 | green | -0.332255742 | 0.090408238 | ANAPC16 | brown | 0.068672 | 0.738884 |
| IDS | brown | -0.077879 | 0.705318 | ACTN2 | green | -0.320098367 | 0.103581864 | HTRA4 | brown | 0.069467 | 0.735967 |
| CARD16 | brown | -0.077151 | 0.707955 | RPL13P5 | green | -0.313594169 | 0.1111922 | IFIT1 | brown | 0.069842 | 0.734592 |
| FAM210B | brown | 0.077011 | 0.708462 | ARC | green | -0.312505216 | 0.112505769 | AP1B1 | brown | 0.070009 | 0.733978 |
| CYTH3 | brown | -0.076749 | 0.70941 | FOS | green | -0.311480004 | 0.113752906 | GGA1 | brown | 0.070271 | 0.73302 |
| GNAS | brown | 0.076701 | 0.709583 | HIST1H4J | green | -0.305203203 | 0.121612059 | WDR24 | brown | 0.070549 | 0.732 |
| FAHD2A | brown | -0.076277 | 0.71112 | LOC338758 | green | -0.304888708 | 0.122016052 | GORAB | brown | 0.070725 | 0.731357 |
| TBC1D23 | brown | 0.076215 | 0.711347 | DYRK1B | green | -0.303863452 | 0.123339908 | C5orf43 | brown | 0.071432 | 0.728768 |
| HIVEP3 | brown | -0.075898 | 0.712498 | NEFM | green | -0.299315962 | 0.129338853 | WDR12 | brown | 0.072249 | 0.725784 |
| MED9 | brown | -0.075466 | 0.714065 | AREG | green | -0.298254119 | 0.130769711 | EMR4P | brown | 0.073133 | 0.722557 |
| PTTG1 | brown | -0.075224 | 0.714945 | CSRP2BP | green | -0.296547846 | 0.133093028 | PRKACB | brown | 0.075184 | 0.71509 |
| C11orf49 | brown | -0.074226 | 0.718574 | NUDT12 | green | -0.296397534 | 0.133299125 | GPKOW | brown | 0.075636 | 0.71345 |
| PLEKHA1 | brown | 0.073825 | 0.720036 | LAT | green | -0.295532162 | 0.13449017 | COQ7 | brown | 0.075877 | 0.712573 |
| CFP | brown | 0.072018 | 0.726628 | PER1 | green | -0.287894026 | 0.145339343 | MRPS16 | brown | 0.07627 | 0.711147 |
| BTBD9 | brown | 0.07169 | 0.727827 | LOC1001331 | green | -0.287301196 | 0.146206928 | ABCB10 | brown | 0.077233 | 0.707658 |
| CDC40 | brown | -0.070022 | 0.733929 | RRAD | green | -0.286616575 | 0.147213466 | BUB3 | brown | 0.078159 | 0.704305 |
| ACTR8 | brown | 0.069893 | 0.734405 | USP35 | green | -0.28451164 | 0.150339288 | PPIL1 | brown | 0.079309 | 0.700152 |
| GPBAR1 | brown | 0.069738 | 0.73497 | PRKCB | green | -0.283088829 | 0.152478875 | RRP7B | brown | 0.080146 | 0.697132 |
| TMEM147 | brown | 0.068632 | 0.739029 | WASH1 | green | -0.278245574 | 0.159924904 | SH3BP4 | brown | 0.08365 | 0.684545 |
| PLXNB2 | brown | 0.068612 | 0.739105 | STON2 | green | -0.277348542 | 0.161331824 | ULK1 | brown | 0.083704 | 0.684352 |
| MRPL24 | brown | 0.068493 | 0.739539 | HIST1H2AL | green | -0.277058246 | 0.161789003 | BRD1 | brown | 0.084936 | 0.679944 |
| C15orf57 | brown | -0.067835 | 0.741959 | BTK | green | -0.276473904 | 0.162712047 | SUOX | brown | 0.085733 | 0.677102 |
| STIL | brown | 0.06722 | 0.74422 | DNAJC9 | green | -0.275765166 | 0.163836583 | PHPT1 | brown | 0.087405 | 0.671145 |
| BOLA2B | brown | 0.066377 | 0.747328 | SLC1A3 | green | -0.275080597 | 0.164927979 | SBF2-AS1 | brown | 0.08886 | 0.66598 |

**Table 6. Gene module membership and correlation with growth rate (48/72h), with adjusted p values for each module**

| | | | | | | | | | | | |
|---|---|---|---|---|---|---|---|---|---|---|---|
| POLR1A | brown | -0.065819 | 0.749384 | MYLIP | green | -0.270190662 | 0.172873526 | LPIN2 | brown | 0.088915 | 0.665786 |
| DOK2 | brown | 0.064845 | 0.752977 | ADM | green | -0.268315 | 0.175991336 | ANKRD36 | brown | 0.089916 | 0.662241 |
| HLA-C | brown | 0.064003 | 0.756088 | BIRC3 | green | -0.267452157 | 0.177438724 | IER5L | brown | 0.092923 | 0.651634 |
| ELMOD3 | brown | 0.063329 | 0.758582 | HSPA2 | green | -0.266934727 | 0.178310673 | AP2M1 | brown | 0.09325 | 0.650487 |
| C8orf33 | brown | -0.062822 | 0.760458 | ADAMTS15 | green | -0.265985313 | 0.179918354 | DARS | brown | 0.093327 | 0.650215 |
| CCDC142 | brown | -0.06278 | 0.760615 | BTBD6 | green | -0.265111178 | 0.181407473 | MRPL53 | brown | 0.095283 | 0.643355 |
| SPRED2 | brown | 0.062583 | 0.761344 | FLJ45445 | green | -0.264681423 | 0.182142715 | GLUD1 | brown | 0.09643 | 0.639349 |
| TCEB2 | brown | 0.062419 | 0.761953 | HIST1H2AM | green | -0.263592143 | 0.18401558 | IKBKE | brown | 0.097859 | 0.634368 |
| CAST | brown | 0.06215 | 0.762949 | LILRB5 | green | -0.261750477 | 0.187212426 | SCARNA1 | brown | 0.098673 | 0.631539 |
| RPA1 | brown | -0.061793 | 0.764272 | MT1G | green | -0.259836098 | 0.190576067 | SLC18B1 | brown | 0.102583 | 0.618014 |
| ANKRD36 | brown | 0.061189 | 0.766515 | LOC1004994[illegible] | green | -0.259748563 | 0.190730861 | AP5M1 | brown | 0.105095 | 0.609391 |
| UHRF2 | brown | -0.060187 | 0.770238 | CD46 | green | -0.258552245 | 0.192855099 | DDX56 | brown | 0.10749 | 0.601213 |
| EIF4ENIF1 | brown | 0.060062 | 0.770702 | TNFSF9 | green | -0.256790833 | 0.196012345 | NDOR1 | brown | 0.109804 | 0.593358 |
| PRDM2 | brown | -0.05983 | 0.771564 | IKBKG | green | -0.253551257 | 0.201911536 | MRFAP1 | brown | 0.109904 | 0.59302 |
| FOXK2 | brown | -0.058278 | 0.77734 | TRIM16L | green | -0.247528718 | 0.213198678 | EID2 | brown | 0.11153 | 0.587529 |
| LRRFIP1 | brown | -0.057777 | 0.779207 | NOMO2 | green | -0.247367597 | 0.213506392 | MYEOV2 | brown | 0.111777 | 0.586696 |
| ABL2 | brown | 0.057493 | 0.780268 | DUSP12 | green | -0.246858966 | 0.214479768 | C6orf108 | brown | 0.112191 | 0.585301 |
| C16orf87 | brown | -0.057385 | 0.780671 | H2AFY2 | green | -0.245832721 | 0.216452835 | SPECC1L | brown | 0.112214 | 0.585226 |
| PPP6R3 | brown | 0.056304 | 0.784705 | KDM6B | green | -0.245572838 | 0.216954428 | OLA1 | brown | 0.117261 | 0.568348 |
| MLH1 | brown | 0.055554 | 0.787509 | C1R | green | -0.243745296 | 0.220503874 | ZNF141 | brown | 0.119354 | 0.561417 |
| RNF217 | brown | 0.055254 | 0.788632 | SPEN | green | -0.241789971 | 0.224344547 | TMEM14C | brown | 0.121355 | 0.554823 |
| HERPUD1 | brown | 0.054119 | 0.792878 | ASTL | green | -0.24147496 | 0.224967466 | C7orf73 | brown | 0.122133 | 0.552268 |
| PRR3 | brown | -0.053814 | 0.794022 | KCTD11 | green | -0.241467642 | 0.224981951 | ZNF582 | brown | 0.122926 | 0.549673 |
| LRRK1 | brown | 0.053442 | 0.795416 | FOSL1 | green | -0.239998905 | 0.227901729 | ZNF571 | brown | 0.123453 | 0.547949 |
| MFSD9 | brown | -0.053118 | 0.796629 | PLEKHM1 | green | -0.239261833 | 0.229376496 | TMCO1 | brown | 0.124328 | 0.545095 |
| MORC2 | brown | 0.051405 | 0.803061 | MAP3K5 | green | -0.238049823 | 0.231815367 | EIF3K | brown | 0.124478 | 0.544606 |
| GADD45GIP | brown | -0.050705 | 0.805691 | HIST1H2AG | green | -0.234140202 | 0.23979997 | C6orf47 | brown | 0.125224 | 0.542178 |
| ZNF542 | brown | -0.050681 | 0.805781 | ATP1B2 | green | -0.233178997 | 0.241790556 | FUT11 | brown | 0.125768 | 0.540411 |
| POLR3K | brown | -0.050632 | 0.805966 | CYB5R4 | green | -0.231817076 | 0.244629627 | RASGRP2 | brown | 0.126568 | 0.537816 |
| PHF14 | brown | -0.049003 | 0.812099 | HIST1H2BN | green | -0.229032363 | 0.250502763 | HCLS1 | brown | 0.127644 | 0.534336 |
| HIST1H3D | brown | 0.046154 | 0.822847 | CYB5R3 | green | -0.227763224 | 0.253209844 | KIAA1191 | brown | 0.129911 | 0.527042 |
| PEMT | brown | -0.045471 | 0.825427 | WDR26 | green | -0.227190466 | 0.254437777 | RBM28 | brown | 0.133463 | 0.515707 |
| TPK1 | brown | -0.044161 | 0.830386 | TRIM47 | green | -0.225764509 | 0.257511743 | ZCCHC24 | brown | 0.134847 | 0.511322 |
| MRFAP1L1 | brown | -0.044158 | 0.830396 | GATAD2A | green | -0.22576328 | 0.257514402 | ARL3 | brown | 0.135087 | 0.510564 |
| CASC4 | brown | 0.044106 | 0.830594 | DNAJB1 | green | -0.225443872 | 0.258206261 | WRB | brown | 0.137268 | 0.503697 |
| SNAPC2 | brown | -0.043485 | 0.832945 | GPR84 | green | -0.225090048 | 0.25897408 | MTMR6 | brown | 0.137295 | 0.503613 |
| LEMD3 | brown | 0.041559 | 0.84025 | LOC646329 | green | -0.224606785 | 0.260025182 | DNASE2 | brown | 0.137813 | 0.501989 |
| METRNL | brown | -0.041407 | 0.840824 | GFPT2 | green | -0.223790514 | 0.261806865 | LPXN | brown | 0.142042 | 0.488828 |
| DDX5 | brown | -0.041039 | 0.842223 | HIST1H3D | green | -0.222795056 | 0.263990351 | KDSR | brown | 0.142501 | 0.487408 |
| FAM72D | brown | -0.040389 | 0.844691 | SNORA48 | green | -0.220726646 | 0.268564885 | SFT2D2 | brown | 0.143403 | 0.484629 |
| KIAA1109 | brown | -0.03961 | 0.847653 | UAP1 | green | -0.217433244 | 0.275953432 | DDA1 | brown | 0.143654 | 0.483857 |
| CBR4 | brown | -0.038895 | 0.850372 | FBXO32 | green | -0.216644394 | 0.277742292 | NTAN1 | brown | 0.144724 | 0.48057 |
| RDH14 | brown | -0.038625 | 0.851401 | HSPA1B | green | -0.216579925 | 0.277888815 | PRKRA | brown | 0.145009 | 0.479699 |
| ZNF571 | brown | 0.038114 | 0.853344 | AKR1C1 | green | -0.216125103 | 0.278923913 | KIF1C | brown | 0.146259 | 0.475877 |
| C6orf203 | brown | 0.038023 | 0.853693 | TRPV4 | green | -0.215997211 | 0.279215418 | GALE | brown | 0.146541 | 0.475018 |
| CD101 | brown | 0.037196 | 0.856839 | NPTN | green | -0.215976961 | 0.279261591 | ZBTB17 | brown | 0.146626 | 0.474759 |
| COX6A1 | brown | 0.036902 | 0.857961 | DYNC1H1 | green | -0.215249173 | 0.280924309 | PCCA | brown | 0.147538 | 0.471986 |
| FAM21A | brown | 0.036355 | 0.860047 | PHC2 | green | -0.213639876 | 0.284623286 | POLD2 | brown | 0.147694 | 0.471512 |
| PILRB | brown | -0.036065 | 0.861151 | BTG2 | green | -0.211589289 | 0.289381166 | USP18 | brown | 0.151737 | 0.459325 |
| PRKCDBP | brown | 0.035969 | 0.861517 | EPS8 | green | -0.208544839 | 0.296537209 | SLX4 | brown | 0.153815 | 0.453127 |
| UBE2E1 | brown | 0.035053 | 0.86501 | PDCD1LG2 | green | -0.20474083 | 0.305633318 | BOLA2B | brown | 0.155127 | 0.449234 |
| UNG | brown | 0.034523 | 0.867034 | TMEM241 | green | -0.202748527 | 0.310465807 | HPS3 | brown | 0.155859 | 0.447071 |
| JARID2 | brown | 0.032965 | 0.872986 | CTSZ | green | -0.202091896 | 0.312068828 | MYNN | brown | 0.158209 | 0.440164 |
| UPF1 | brown | -0.032397 | 0.87516 | HIST1H2AD | green | -0.200997724 | 0.314751358 | ZNF737 | brown | 0.160096 | 0.434661 |
| NME3 | brown | -0.03227 | 0.875646 | LINC00656 | green | -0.199753124 | 0.317819922 | AK3 | brown | 0.161098 | 0.431752 |
| METTL5 | brown | -0.03223 | 0.875796 | HES4 | green | -0.199751588 | 0.31782372 | IL7 | brown | 0.163993 | 0.423411 |

**Table 6. Gene module membership and correlation with growth rate (48/72h), with adjusted p values for each module**

| | | | | | | | | | | | |
|---|---|---|---|---|---|---|---|---|---|---|---|
| FABP4 | brown | -0.030708 | 0.881623 | FAM20A | green | -0.199448126 | 0.318574692 | LOC283922 | brown | 0.167217 | 0.414224 |
| ERN1 | brown | 0.029948 | 0.88453 | MCTP1 | green | -0.196754174 | 0.325289105 | OGG1 | brown | 0.168918 | 0.409424 |
| STK11IP | brown | 0.029175 | 0.887491 | FZD8 | green | -0.196019149 | 0.32713597 | MATR3 | brown | 0.170663 | 0.404532 |
| FAM207A | brown | 0.029035 | 0.888028 | GRASP | green | -0.194274069 | 0.331546289 | EMC3 | brown | 0.17087 | 0.403952 |
| C17orf51 | brown | -0.02893 | 0.888433 | SH3D21 | green | -0.192022028 | 0.337290871 | ZC3H12C | brown | 0.17239 | 0.399722 |
| TLE3 | brown | 0.028267 | 0.890974 | CLDN7 | green | -0.190432997 | 0.341380113 | NBPF9 | brown | 0.172946 | 0.39818 |
| ELP3 | brown | -0.028082 | 0.891685 | ZNF844 | green | -0.189130615 | 0.344753792 | RPL26L1 | brown | 0.17304 | 0.397919 |
| SNORA66 | brown | -0.028003 | 0.891984 | SYNM | green | -0.188754693 | 0.345731275 | PMS2P3 | brown | 0.175332 | 0.3916 |
| IFFO1 | brown | -0.027639 | 0.893381 | THBD | green | -0.188499373 | 0.346396108 | POLR2L | brown | 0.177105 | 0.386754 |
| MSL3 | brown | -0.027226 | 0.894967 | FBXO44 | green | -0.188015489 | 0.347658202 | NANS | brown | 0.17881 | 0.382123 |
| RRAGA | brown | -0.026516 | 0.897693 | ENTPD1 | green | -0.185299047 | 0.354794207 | TRMT6 | brown | 0.18049 | 0.377595 |
| SRP9 | brown | -0.026508 | 0.897722 | BIVM | green | -0.184654286 | 0.356500626 | GTF2A1 | brown | 0.182999 | 0.370887 |
| NANP | brown | 0.026309 | 0.898487 | TMEM57 | green | -0.18462709 | 0.356572709 | DNAJC19 | brown | 0.183204 | 0.370343 |
| WDR91 | brown | 0.025039 | 0.903362 | C6orf136 | green | -0.184424078 | 0.357111064 | MEA1 | brown | 0.186212 | 0.362399 |
| RPS21 | brown | 0.024706 | 0.904643 | DUSP6 | green | -0.184057557 | 0.358084237 | THOC2 | brown | 0.186505 | 0.361632 |
| LYPLA1 | brown | 0.024456 | 0.905603 | PLAA | green | -0.18380493 | 0.358755914 | FUBP3 | brown | 0.186965 | 0.360427 |
| EXOSC3 | brown | -0.023381 | 0.909736 | FAM53C | green | -0.182121953 | 0.363249506 | RNF113A | brown | 0.187652 | 0.358633 |
| AAR2 | brown | -0.02295 | 0.911392 | CHRNE | green | -0.181390005 | 0.36521409 | UBIAD1 | brown | 0.188802 | 0.355641 |
| HGS | brown | -0.022507 | 0.913099 | DANCR | green | -0.174760984 | 0.383288738 | SOD1 | brown | 0.189525 | 0.353767 |
| CHCHD5 | brown | 0.022183 | 0.914344 | TULP3 | green | -0.172780084 | 0.38878788 | C9orf142 | brown | 0.1913 | 0.349194 |
| ABI2 | brown | 0.022159 | 0.914435 | EMP3 | green | -0.172421957 | 0.389786858 | GATC | brown | 0.192831 | 0.345276 |
| MTMR6 | brown | -0.022104 | 0.914648 | SCAF4 | green | -0.17153306 | 0.392272713 | EXOSC8 | brown | 0.193538 | 0.343476 |
| L3HYPDH | brown | 0.021959 | 0.915207 | DUSP2 | green | -0.170648413 | 0.394755616 | C14orf1 | brown | 0.196052 | 0.337121 |
| TLK2 | brown | 0.021455 | 0.917146 | KLHL8 | green | -0.169519427 | 0.397937214 | CLN6 | brown | 0.196989 | 0.334772 |
| PET100 | brown | -0.021417 | 0.917291 | NFAM1 | green | -0.168922777 | 0.399624474 | TRMU | brown | 0.198323 | 0.331442 |
| ALPK1 | brown | -0.021364 | 0.917494 | PDK4 | green | -0.167393428 | 0.403967699 | PLEKHF2 | brown | 0.199211 | 0.329239 |
| CDC23 | brown | -0.021327 | 0.917637 | TAF4B | green | -0.167297816 | 0.404240107 | SNAI3 | brown | 0.199427 | 0.328702 |
| EIF4G2 | brown | 0.0203 | 0.921593 | GNS | green | -0.167227366 | 0.404440893 | DIAPH2 | brown | 0.200302 | 0.326541 |
| MAPK9 | brown | 0.019336 | 0.925306 | PKN1 | green | -0.166680189 | 0.406002269 | KLHL20 | brown | 0.202022 | 0.322319 |
| DEAF1 | brown | 0.019089 | 0.926255 | B3GNT5 | green | -0.165052391 | 0.410667135 | LEPROT | brown | 0.202046 | 0.322259 |
| CCNB1 | brown | 0.018598 | 0.928147 | EMP1 | green | -0.163453458 | 0.415278222 | LOC1001348( | brown | 0.202644 | 0.320798 |
| ZNF174 | brown | -0.018462 | 0.928673 | FAM46C | green | -0.162750603 | 0.417314198 | PDE7A | brown | 0.205448 | 0.314005 |
| GLTSCR1 | brown | -0.017647 | 0.931815 | ZNF302 | green | -0.162529517 | 0.417955764 | SGMS1 | brown | 0.208626 | 0.306411 |
| TBC1D7 | brown | -0.017573 | 0.9321 | BTN3A3 | green | -0.161963569 | 0.419600558 | NDUFA13 | brown | 0.208857 | 0.305865 |
| PRKD3 | brown | 0.017396 | 0.932782 | PRKAR2B | green | -0.161076855 | 0.422184746 | MEIS3P1 | brown | 0.20988 | 0.303447 |
| SPTAN1 | brown | -0.016655 | 0.935637 | TMEM99 | green | -0.160843279 | 0.42286692 | B3GNT2 | brown | 0.210066 | 0.303009 |
| UST | brown | -0.015096 | 0.941654 | COQ10A | green | -0.160169291 | 0.424838741 | MKNK2 | brown | 0.210159 | 0.30279 |
| XBP1 | brown | -0.014452 | 0.944137 | SNAI1 | green | -0.159185159 | 0.427726952 | PANK4 | brown | 0.21564 | 0.290062 |
| MANBA | brown | 0.0129 | 0.950131 | IQSEC1 | green | -0.156580154 | 0.435423621 | FUT10 | brown | 0.215899 | 0.289469 |
| GALNT2 | brown | -0.011771 | 0.95449 | FLNB | green | -0.152460096 | 0.447748229 | NOLC1 | brown | 0.216294 | 0.288565 |
| LRMP | brown | -0.011171 | 0.956805 | UBE2S | green | -0.152281074 | 0.448287929 | PSMA7 | brown | 0.21699 | 0.286979 |
| TMEM184C | brown | -0.010984 | 0.957528 | MPEG1 | green | -0.151712739 | 0.450003599 | LSM2 | brown | 0.217521 | 0.285772 |
| EMC3 | brown | -0.010243 | 0.960391 | ILVBL | green | -0.150617546 | 0.453319552 | LILRA3 | brown | 0.217527 | 0.285759 |
| LYRM2 | brown | 0.009611 | 0.962834 | ABCB6 | green | -0.150417511 | 0.453926598 | ASPSCR1 | brown | 0.219242 | 0.281884 |
| ZWINT | brown | -0.009508 | 0.96323 | HIST1H2BB | green | -0.149456809 | 0.456848031 | PIGT | brown | 0.219626 | 0.281021 |
| C14orf1 | brown | 0.009264 | 0.964174 | NFE2L3 | green | -0.148947255 | 0.458401565 | MTF2 | brown | 0.22108 | 0.277766 |
| C14orf166 | brown | 0.008642 | 0.96658 | ZNF646 | green | -0.148811852 | 0.45881485 | SAAL1 | brown | 0.221962 | 0.275805 |
| KBTBD2 | brown | 0.008475 | 0.967226 | HLA-DOA | green | -0.147548971 | 0.462678918 | RPS21 | brown | 0.222863 | 0.273811 |
| ATP6V1D | brown | -0.007842 | 0.969671 | HIST3H2BB | green | -0.146922526 | 0.464601965 | RAB38 | brown | 0.22302 | 0.273463 |
| FBRS | brown | -0.006876 | 0.973406 | ZNF268 | green | -0.146919183 | 0.46461224 | UNG | brown | 0.224152 | 0.270973 |
| ALDOC | brown | 0.00671 | 0.974048 | FAM117A | green | -0.146307993 | 0.466492494 | EMC9 | brown | 0.224161 | 0.270953 |
| TMEM245 | brown | -0.006405 | 0.975226 | FSCN1 | green | -0.145543081 | 0.46885123 | PPP2R4 | brown | 0.22468 | 0.269816 |
| PARK7 | brown | 0.005097 | 0.980285 | R3HCC1 | green | -0.145277442 | 0.469671819 | NSFL1C | brown | 0.224762 | 0.269636 |
| LRRC16A | brown | 0.004507 | 0.982568 | IFI44L | green | -0.14293833 | 0.476929652 | ZNF512 | brown | 0.225538 | 0.267942 |
| CUL2 | brown | 0.004128 | 0.984033 | LYSMD2 | green | -0.140898772 | 0.483304725 | TDP2 | brown | 0.227059 | 0.264642 |
| MYO18A | brown | -0.002098 | 0.991882 | NXF1 | green | -0.139949955 | 0.486285179 | ABHD3 | brown | 0.229187 | 0.26007 |

Table 6. Gene module membership and correlation with growth rate (48/72h), with adjusted p values for each module

| | | | | | | | | | | | |
|---|---|---|---|---|---|---|---|---|---|---|---|
| LCORL | brown | -0.001631 | 0.993693 | HOMER2 | green | -0.13956537 | 0.4874959 | DDX55 | brown | 0.229769 | 0.258828 |
| EHBP1L1 | brown | 0.001177 | 0.995446 | SLC6A8 | green | -0.137337385 | 0.494539805 | DDX20 | brown | 0.230483 | 0.25731 |
| TADA1 | brown | 0.000752 | 0.997091 | ST6GALNAC | green | -0.135786006 | 0.499474557 | GSPT2 | brown | 0.230942 | 0.256337 |
| DPY19L3 | brown | -0.000541 | 0.997908 | CTSC | green | -0.134805821 | 0.502605006 | POLR2F | brown | 0.232393 | 0.253277 |
| SIVA1 | cyan | -0.489782 | 0.011093 | C20orf111 | green | -0.13341249 | 0.507071626 | CYB5B | brown | 0.232454 | 0.25315 |
| CRYZ | cyan | -0.471532 | 0.015025 | FPR2 | green | -0.132586755 | 0.509727901 | EED | brown | 0.234526 | 0.248823 |
| METTL17 | cyan | -0.467783 | 0.015961 | LY86 | green | -0.131251552 | 0.514037476 | ROMO1 | brown | 0.234726 | 0.248408 |
| PDCD2L | cyan | -0.466268 | 0.016352 | ZFP36 | green | -0.130223079 | 0.517369112 | ZNF526 | brown | 0.235757 | 0.246276 |
| C6orf57 | cyan | -0.434206 | 0.026664 | AKR1C2 | green | -0.129632582 | 0.519286697 | ECI2 | brown | 0.236631 | 0.244478 |
| WDR5 | cyan | -0.432654 | 0.027273 | ABHD16B | green | -0.128438031 | 0.52317639 | NANP | brown | 0.241897 | 0.233828 |
| RIT1 | cyan | 0.422535 | 0.031522 | AKAP7 | green | -0.127254703 | 0.527043325 | PSMB7 | brown | 0.242532 | 0.232565 |
| NAPSB | cyan | -0.419046 | 0.033104 | USP33 | green | -0.124616637 | 0.535713088 | PEPD | brown | 0.242757 | 0.232119 |
| HIST1H4F | cyan | -0.410306 | 0.037347 | HIST1H2BG | green | -0.124343526 | 0.536614477 | PGM5 | brown | 0.242944 | 0.231748 |
| DCSTAMP | cyan | 0.399415 | 0.043229 | DDIT4 | green | -0.120635866 | 0.548921754 | YIF1B | brown | 0.24857 | 0.22078 |
| R3HCC1 | cyan | -0.389661 | 0.049098 | RHOB | green | -0.120370556 | 0.549807412 | WDR33 | brown | 0.249912 | 0.218216 |
| SHARPIN | cyan | -0.384041 | 0.052753 | COQ2 | green | -0.119843326 | 0.551569375 | VHL | brown | 0.251268 | 0.215645 |
| CUTA | cyan | -0.383188 | 0.053325 | HIST1H4C | green | -0.11909063 | 0.554089326 | ZNF223 | brown | 0.252768 | 0.212826 |
| GLUL | cyan | -0.380105 | 0.055437 | KHDRBS3 | green | -0.118142732 | 0.557270299 | NUPL2 | brown | 0.252998 | 0.212396 |
| POLH | cyan | 0.375001 | 0.059074 | PLK2 | green | -0.118132341 | 0.557305217 | TPM3P9 | brown | 0.253005 | 0.212383 |
| REXO1 | cyan | 0.374292 | 0.059593 | C4orf33 | green | -0.117886489 | 0.558131636 | KPNA2 | brown | 0.254023 | 0.210486 |
| NLRX1 | cyan | -0.360342 | 0.070556 | STARD5 | green | -0.116796221 | 0.561803239 | COG4 | brown | 0.254187 | 0.210182 |
| SH2B2 | cyan | -0.353174 | 0.076759 | HIPK4 | green | -0.114726021 | 0.568804863 | MANEA | brown | 0.254464 | 0.209669 |
| LY6E | cyan | -0.347827 | 0.08165 | DHRS4L2 | green | -0.11438171 | 0.56997314 | MRFAP1L1 | brown | 0.255154 | 0.208393 |
| CDC123 | cyan | -0.343591 | 0.085691 | NEFH | green | -0.111051361 | 0.581328328 | CNIH4 | brown | 0.256233 | 0.20641 |
| ATMIN | cyan | -0.34171 | 0.087533 | C5orf15 | green | -0.110541431 | 0.583075715 | MAPRE1 | brown | 0.256935 | 0.205125 |
| ODF3L1 | cyan | -0.334849 | 0.094505 | RNU12 | green | -0.109682236 | 0.586025121 | SGTA | brown | 0.257819 | 0.203516 |
| ALDH16A1 | cyan | -0.330302 | 0.09935 | SLC7A5 | green | -0.108748602 | 0.589237404 | L3MBTL4 | brown | 0.25795 | 0.203279 |
| PARVB | cyan | -0.328173 | 0.101682 | SERPING1 | green | -0.105742985 | 0.599629959 | UBTF | brown | 0.258557 | 0.20218 |
| FAM160B2 | cyan | -0.326658 | 0.103364 | ZNF187 | green | -0.103957105 | 0.605841646 | CLPP | brown | 0.259798 | 0.199946 |
| ARPC1B | cyan | -0.322157 | 0.108489 | COX20 | green | -0.103258371 | 0.60827933 | NFU1 | brown | 0.260128 | 0.199354 |
| ZNF426 | cyan | 0.308597 | 0.125061 | KLF4 | green | -0.103082634 | 0.608893074 | SS18L2 | brown | 0.260761 | 0.198223 |
| DRG2 | cyan | -0.304716 | 0.130128 | NRARP | green | -0.10221524 | 0.611926127 | EIF4EBP1 | brown | 0.264341 | 0.191911 |
| PGAM1 | cyan | -0.293229 | 0.145994 | NUDT4 | green | -0.101388186 | 0.614823952 | CRELD2 | brown | 0.264975 | 0.190808 |
| CSNK1G2 | cyan | -0.292343 | 0.147273 | RAB31 | green | -0.098354444 | 0.625501669 | RDX | brown | 0.268696 | 0.18442 |
| TRABD | cyan | -0.290724 | 0.149631 | EEPD1 | green | -0.098166879 | 0.626164286 | UBE2G2 | brown | 0.269314 | 0.183374 |
| PACS2 | cyan | -0.289341 | 0.151666 | NFKBID | green | -0.097870929 | 0.627210378 | DDX49 | brown | 0.269426 | 0.183186 |
| CSGALNACT | cyan | 0.288268 | 0.153258 | HIST2H2AB | green | -0.094920227 | 0.637678418 | INPP5B | brown | 0.270681 | 0.181074 |
| ATP13A3 | cyan | 0.287425 | 0.154517 | HIST3H2A | green | -0.093803669 | 0.641657466 | GSK3B | brown | 0.271463 | 0.179768 |
| ARPC4 | cyan | -0.285926 | 0.156774 | ANXA1 | green | -0.093601789 | 0.642377937 | RPAP3 | brown | 0.272604 | 0.177873 |
| MED13 | cyan | 0.284241 | 0.159341 | CTSO | green | -0.092152142 | 0.647560693 | VBP1 | brown | 0.273354 | 0.176635 |
| ZNF804A | cyan | 0.281756 | 0.163179 | HIST1H3C | green | -0.091011379 | 0.651650448 | TARSL2 | brown | 0.274798 | 0.174269 |
| KIAA1919 | cyan | 0.281712 | 0.163246 | SLC7A7 | green | -0.087637048 | 0.663805111 | PPM1M | brown | 0.276608 | 0.171334 |
| P4HTM | cyan | -0.280201 | 0.165613 | CRELD2 | green | -0.086291291 | 0.668676127 | ACAD9 | brown | 0.277861 | 0.169324 |
| TMEM179B | cyan | -0.27661 | 0.171331 | ITIH5 | green | -0.086010023 | 0.66969585 | NAGK | brown | 0.279394 | 0.166886 |
| ATP5G3 | cyan | -0.275038 | 0.173878 | CPED1 | green | -0.084235784 | 0.676141379 | HIF1AN | brown | 0.283856 | 0.159931 |
| NAA10 | cyan | -0.265322 | 0.190206 | GADD45A | green | -0.083988877 | 0.677040132 | MKI67IP | brown | 0.284833 | 0.158436 |
| SSR4 | cyan | -0.263269 | 0.193787 | SLC7A5P1 | green | -0.080420765 | 0.690075926 | EFNB1 | brown | 0.284972 | 0.158224 |
| B3GALT4 | cyan | -0.257398 | 0.204281 | TFPI | green | -0.078742026 | 0.696239313 | CDK8 | brown | 0.286512 | 0.15589 |
| CROT | cyan | 0.256192 | 0.206485 | C11orf82 | green | -0.078640227 | 0.69661367 | BNIP3L | brown | 0.287675 | 0.154144 |
| UXS1 | cyan | -0.253783 | 0.210932 | FABP4 | green | -0.078538572 | 0.696987572 | NMRAL1 | brown | 0.28877 | 0.152512 |
| LRSAM1 | cyan | -0.251514 | 0.215182 | ADRB2 | green | -0.078117579 | 0.698536769 | PRDX5 | brown | 0.291637 | 0.148298 |
| RAC1 | cyan | -0.25146 | 0.215282 | CLEC7A | green | -0.078058361 | 0.698754775 | BUD13 | brown | 0.292946 | 0.146402 |
| USP36 | cyan | 0.249904 | 0.21823 | C1orf63 | green | -0.076462246 | 0.704639552 | KEAP1 | brown | 0.293108 | 0.146169 |
| FAM211A | cyan | -0.244791 | 0.228106 | HIST1H3A | green | -0.07641231 | 0.704823935 | NF2 | brown | 0.293265 | 0.145942 |
| C16orf54 | cyan | -0.241553 | 0.234513 | EGR1 | green | -0.075795598 | 0.707102384 | ADO | brown | 0.294076 | 0.144779 |
| ATP6V1E1 | cyan | -0.237046 | 0.243627 | TRMT6 | green | -0.075741901 | 0.707300886 | LOC642361 | brown | 0.297021 | 0.14061 |

**Table 6. Gene module membership and correlation with growth rate (48/72h), with adjusted p values for each module**

| | | | | | | | | | | | |
|---|---|---|---|---|---|---|---|---|---|---|---|
| CAMKK1 | cyan | 0.235584 | 0.246634 | HEXIM1 | green | -0.075257691 | 0.709091703 | HUWE1 | brown | 0.298896 | 0.138002 |
| TNRC18 | cyan | -0.234523 | 0.24883 | APOL1 | green | -0.074881392 | 0.710484463 | PARVG | brown | 0.301094 | 0.134989 |
| C11orf24 | cyan | -0.23284 | 0.252341 | FOSB | green | -0.073393697 | 0.715999581 | POGLUT1 | brown | 0.304528 | 0.130377 |
| TIMM22 | cyan | -0.232276 | 0.253524 | TUBA4A | green | -0.070421459 | 0.727059631 | JOSD2 | brown | 0.306904 | 0.127254 |
| SURF1 | cyan | -0.229905 | 0.258539 | KLF5 | green | -0.07016701 | 0.728008985 | UBE2V2 | brown | 0.307841 | 0.126037 |
| PLCXD1 | cyan | 0.228443 | 0.261663 | SNORA12 | green | -0.070040511 | 0.728481103 | C15orf62 | brown | 0.310044 | 0.123211 |
| MYO1E | cyan | 0.225658 | 0.267682 | MRPL18 | green | -0.068971118 | 0.732476118 | EIF2AK2 | brown | 0.311286 | 0.121637 |
| ZNF791 | cyan | 0.220556 | 0.278936 | IFI44 | green | -0.068130561 | 0.735621042 | ARL8B | brown | 0.312566 | 0.120031 |
| KIF9 | cyan | 0.218528 | 0.283493 | P2RY13 | green | -0.06799755 | 0.736119084 | PRMT5 | brown | 0.312812 | 0.119724 |
| HECTD3 | cyan | -0.216152 | 0.28889 | SLC7A5P2 | green | -0.067831945 | 0.736739313 | ZNF134 | brown | 0.313542 | 0.118817 |
| PHF20L1 | cyan | 0.213613 | 0.29473 | HIRA | green | -0.066054659 | 0.743405725 | FUCA2 | brown | 0.324619 | 0.105663 |
| CLTC | cyan | 0.2079 | 0.308137 | VASN | green | -0.065229922 | 0.746505406 | PCYT1A | brown | 0.325045 | 0.105179 |
| GAB1 | cyan | 0.207188 | 0.309833 | AGAP5 | green | -0.063733046 | 0.752141056 | SART3 | brown | 0.3341 | 0.095291 |
| B4GALT7 | cyan | -0.207083 | 0.310083 | NUP160 | green | -0.062754694 | 0.755831224 | PSMD11 | brown | 0.334668 | 0.094694 |
| FAM18B2 | cyan | 0.205626 | 0.313577 | UTP18 | green | -0.062107353 | 0.758275763 | BAG4 | brown | 0.33527 | 0.094065 |
| TPP2 | cyan | 0.203606 | 0.318458 | CLEC4A | green | -0.061623881 | 0.760102971 | DDX58 | brown | 0.336656 | 0.09263 |
| CPEB4 | cyan | 0.202741 | 0.320562 | HIST1H2AH | green | -0.061457691 | 0.760731351 | RBM17 | brown | 0.339163 | 0.090074 |
| ZFYVE16 | cyan | 0.200295 | 0.32656 | ADORA2B | green | -0.060623997 | 0.763885854 | RRAGA | brown | 0.340948 | 0.088288 |
| EXT1 | cyan | 0.199857 | 0.32764 | TIAM1 | green | -0.059672838 | 0.767489312 | MPLKIP | brown | 0.342421 | 0.086833 |
| ITPR1 | cyan | 0.188778 | 0.355703 | MAFB | green | -0.059256156 | 0.769069399 | CPT2 | brown | 0.349076 | 0.080487 |
| ADNP2 | cyan | 0.188315 | 0.356906 | ZSCAN22 | green | -0.058897105 | 0.770431668 | ANKS3 | brown | 0.353004 | 0.076911 |
| HDLBP | cyan | 0.182701 | 0.371679 | ARHGAP31 | green | -0.058281871 | 0.772767461 | SNHG9 | brown | 0.356425 | 0.073896 |
| NDST2 | cyan | 0.179446 | 0.380406 | GOLIM4 | green | -0.056908268 | 0.777989428 | GDI1 | brown | 0.363042 | 0.068321 |
| SAP30BP | cyan | 0.178856 | 0.381999 | GTF2E1 | green | -0.056113921 | 0.781013578 | POMT1 | brown | 0.364345 | 0.067263 |
| SLC44A2 | cyan | -0.177484 | 0.385722 | PMAIP1 | green | -0.055410571 | 0.783693889 | CLPTM1 | brown | 0.365272 | 0.066517 |
| TBC1D5 | cyan | 0.176449 | 0.388544 | STK11IP | green | -0.053745036 | 0.790050444 | SH3GLB2 | brown | 0.373664 | 0.060057 |
| TRIM65 | cyan | -0.175587 | 0.390902 | SNORA64 | green | -0.052838515 | 0.79351575 | CDK14 | brown | 0.381086 | 0.054757 |
| CLEC7A | cyan | 0.17327 | 0.397281 | SNORA8 | green | -0.052784726 | 0.793721489 | DPEP2 | brown | 0.383436 | 0.053158 |
| LIMK1 | cyan | -0.167902 | 0.412288 | FAM213A | green | -0.052725509 | 0.793948002 | CHRAC1 | brown | 0.384201 | 0.052646 |
| SERINC5 | cyan | 0.164538 | 0.42185 | TUFT1 | green | -0.052613894 | 0.794374991 | HDHD2 | brown | 0.388389 | 0.049907 |
| FOXK1 | cyan | -0.163061 | 0.426086 | CTSW | green | -0.050674126 | 0.801804808 | PPAN | brown | 0.396717 | 0.044794 |
| DST | cyan | 0.161855 | 0.429561 | GPR85 | green | -0.049948911 | 0.804586935 | ACSF3 | brown | 0.397016 | 0.044619 |
| ZNF326 | cyan | 0.160204 | 0.434346 | USE1 | green | -0.048719775 | 0.809307535 | YIPF5 | brown | 0.402298 | 0.041606 |
| EWSR1 | cyan | 0.156689 | 0.444624 | FAM83H | green | -0.047210288 | 0.815113749 | RBMS1 | brown | 0.404833 | 0.040218 |
| ABCF3 | cyan | 0.15588 | 0.447009 | FBP1 | green | -0.047141243 | 0.815379562 | FOXJ2 | brown | 0.415622 | 0.034718 |
| NIPBL | cyan | 0.155089 | 0.449348 | IFI35 | green | -0.04669846 | 0.817084669 | NME3 | brown | 0.419818 | 0.032748 |
| SLU7 | cyan | 0.153086 | 0.455296 | CD4 | green | -0.046581612 | 0.817534775 | AARS | brown | 0.425418 | 0.030261 |
| KBTBD8 | cyan | 0.150763 | 0.462246 | DTNB | green | -0.046322667 | 0.818532446 | KTI12 | brown | 0.430975 | 0.027944 |
| PWWP2B | cyan | -0.147793 | 0.471213 | MPC1 | green | -0.046260512 | 0.81877196 | STAG1 | brown | 0.434731 | 0.026461 |
| ATP6V1C1 | cyan | -0.146712 | 0.474497 | ELAC2 | green | -0.04523816 | 0.822713846 | ARHGEF12 | brown | 0.443031 | 0.023409 |
| SCD | cyan | 0.144328 | 0.481784 | NCKAP1L | green | -0.044998374 | 0.823639 | WHSC1 | brown | 0.460908 | 0.0178 |
| PSMD14 | cyan | 0.143262 | 0.485062 | CKS2 | green | -0.044604886 | 0.825157677 | TAB3 | brown | 0.472523 | 0.014786 |
| AHI1 | cyan | 0.139901 | 0.495469 | RASD1 | green | -0.044193012 | 0.826747971 | SSRP1 | brown | 0.483134 | 0.012413 |
| UHRF1BP1L | cyan | 0.139642 | 0.496274 | APOL4 | green | -0.043705381 | 0.828631633 | EXOG | brown | 0.543064 | 0.004146 |
| FBXO30 | cyan | 0.138866 | 0.498694 | SNORA47 | green | -0.042605556 | 0.832883518 | SMG5 | brown | 0.584614 | 0.00171 |
| NSFP1 | cyan | -0.137819 | 0.50197 | BAIAP2 | green | -0.042592385 | 0.832934464 | HSP90AA1 | cyan | -0.38534 | 0.05189 |
| C11orf54 | cyan | -0.134557 | 0.512239 | MMAB | green | -0.039528582 | 0.844803019 | FUNDC2 | cyan | -0.344669 | 0.084648 |
| TEX264 | cyan | -0.133438 | 0.515785 | EPHA2 | green | -0.037491764 | 0.852711745 | RCL1 | cyan | -0.343552 | 0.085729 |
| ROCK2 | cyan | 0.131282 | 0.522652 | JUND | green | -0.035417481 | 0.860780246 | PIGX | cyan | -0.318065 | 0.113308 |
| SPAG9 | cyan | 0.12925 | 0.529162 | FBXL5 | green | -0.035156685 | 0.861795668 | HPRT1 | cyan | -0.317604 | 0.113861 |
| FAM46A | cyan | 0.124883 | 0.543288 | LRRK2 | green | -0.03395628 | 0.866472244 | ATP5H | cyan | -0.315654 | 0.116221 |
| DMXL2 | cyan | 0.122375 | 0.551476 | GBP1 | green | -0.032280983 | 0.873006224 | RAN | cyan | -0.311885 | 0.120884 |
| HPS1 | cyan | -0.121605 | 0.554001 | CECR1 | green | -0.031843511 | 0.874713807 | MTRR | cyan | -0.304647 | 0.130219 |
| SASH1 | cyan | 0.119348 | 0.561437 | SLC47A1 | green | -0.031475876 | 0.876149218 | SSR2 | cyan | -0.288846 | 0.152399 |
| ACSL4 | cyan | 0.118026 | 0.565812 | CTSL2 | green | -0.030904126 | 0.878382343 | LINC00674 | cyan | -0.285556 | 0.157335 |
| PAK1 | cyan | -0.116259 | 0.571683 | HSPA6 | green | -0.028764651 | 0.886746579 | SRSF3 | cyan | -0.284677 | 0.158675 |

**Table 6. Gene module membership and correlation with growth rate (48/72h), with adjusted p values for each module**

| | | | | | | | | | | | |
|---|---|---|---|---|---|---|---|---|---|---|---|
| ZFC3H1 | cyan | 0.11531 | 0.574847 | ZNF808 | green | -0.027915933 | 0.890067956 | STARD7 | cyan | -0.276295 | 0.171839 |
| SPCS2 | cyan | 0.110804 | 0.589977 | TCTN3 | green | -0.026894385 | 0.89406809 | GMPPA | cyan | -0.270896 | 0.180714 |
| C6orf136 | cyan | 0.103739 | 0.614039 | CD84 | green | -0.026538212 | 0.895463375 | DNAJC25 | cyan | -0.267245 | 0.186894 |
| KAT6A | cyan | 0.103537 | 0.614733 | PKDCC | green | -0.026048052 | 0.897384039 | DDX39B | cyan | -0.265187 | 0.19044 |
| ABHD5 | cyan | 0.099647 | 0.628158 | OAS1 | green | -0.02544963 | 0.89972969 | PRDX3 | cyan | -0.263886 | 0.192705 |
| CDKAL1 | cyan | 0.099078 | 0.630131 | SERTAD1 | green | -0.024512663 | 0.90340399 | MRPL28 | cyan | -0.263697 | 0.193037 |
| DNAJC8 | cyan | -0.093885 | 0.648255 | TLR7 | green | -0.02365088 | 0.906785173 | RBM18 | cyan | -0.254006 | 0.210518 |
| ADPRH | cyan | 0.089898 | 0.662306 | PCED1B | green | -0.023056465 | 0.909118273 | GGCT | cyan | -0.250589 | 0.21693 |
| NRAS | cyan | -0.083768 | 0.684123 | PCYOX1 | green | -0.022202448 | 0.912471604 | ATP5J2 | cyan | -0.245729 | 0.226273 |
| IDH3A | cyan | -0.082713 | 0.687903 | IL18 | green | -0.019580155 | 0.922777041 | CNOT6 | cyan | -0.239345 | 0.238949 |
| RPL27A | cyan | -0.082627 | 0.68821 | HIST1H3I | green | -0.013533424 | 0.946584029 | MRPL3 | cyan | -0.23776 | 0.242168 |
| ASNS | cyan | -0.08092 | 0.694344 | TAGAP | green | -0.013183251 | 0.947964287 | MRPS14 | cyan | -0.236926 | 0.243873 |
| KRT8 | cyan | -0.080766 | 0.6949 | PLIN2 | green | -0.013155863 | 0.948072244 | PSMD1 | cyan | -0.233045 | 0.25191 |
| ELL2 | cyan | -0.080569 | 0.695609 | C7orf41 | green | -0.013066727 | 0.948423613 | TBC1D1 | cyan | -0.229385 | 0.259648 |
| MAP2K4 | cyan | -0.080015 | 0.697603 | GADD45B | green | -0.010287481 | 0.959383624 | ARL6IP5 | cyan | -0.225963 | 0.267017 |
| TERF2 | cyan | -0.077647 | 0.706156 | ZDHHC18 | green | -0.009960184 | 0.96067484 | XPO5 | cyan | -0.22434 | 0.27056 |
| DNAH17 | cyan | 0.074901 | 0.716118 | THAP9-AS1 | green | -0.00746772 | 0.970510767 | ATP2A2 | cyan | -0.223643 | 0.272091 |
| SRSF1 | cyan | 0.073206 | 0.722293 | TTC19 | green | -0.006785282 | 0.973204643 | WDR73 | cyan | -0.222793 | 0.273965 |
| FBXO46 | cyan | -0.072402 | 0.725225 | FBN1 | green | -0.005401356 | 0.978668447 | HMGN1 | cyan | -0.2216 | 0.27661 |
| MON2 | cyan | 0.067427 | 0.743461 | LOC1001320 | green | -0.003811754 | 0.984945411 | CENPW | cyan | -0.220652 | 0.278722 |
| PAN3 | cyan | 0.066239 | 0.747833 | ENO2 | green | -0.002474131 | 0.990228062 | ACO1 | cyan | -0.217865 | 0.284992 |
| STAU2 | cyan | 0.064051 | 0.755911 | LY75 | green | -0.002000379 | 0.992099149 | RNF34 | cyan | -0.21554 | 0.290292 |
| HMBOX1 | cyan | 0.06227 | 0.762505 | PPT1 | green | -0.001687156 | 0.993336249 | HNRNPA2B | cyan | -0.214756 | 0.292093 |
| AP1G1 | cyan | 0.059154 | 0.774077 | EID3 | green | -0.000445542 | 0.998240227 | PABPC3 | cyan | -0.214528 | 0.292616 |
| GMCL1 | cyan | 0.058689 | 0.775808 | TYSND1 | green | -0.000235441 | 0.999070072 | TFIP11 | cyan | -0.213086 | 0.29595 |
| RIPK2 | cyan | -0.058189 | 0.777672 | GBP4 | green | 0.001518909 | 0.994000762 | ZNF782 | cyan | -0.212919 | 0.296338 |
| ZNF565 | cyan | 0.056259 | 0.784874 | RASL11A | green | 0.002388533 | 0.990566128 | TIMM44 | cyan | -0.208953 | 0.305638 |
| RSAD2 | cyan | 0.04648 | 0.821616 | SPATA13 | green | 0.002478491 | 0.99021084 | NDUFB2 | cyan | -0.207518 | 0.309046 |
| SIGLEC11 | cyan | -0.044451 | 0.829288 | GLDN | green | 0.004299874 | 0.983017829 | VMP1 | cyan | -0.191152 | 0.349574 |
| LGMN | cyan | -0.041512 | 0.840429 | H2AFX | green | 0.004438815 | 0.98246917 | DPP3 | cyan | -0.188878 | 0.355445 |
| NCOR2 | cyan | -0.041185 | 0.841671 | BLNK | green | 0.004551294 | 0.98202501 | MGAT4B | cyan | -0.188259 | 0.357051 |
| PDS5A | cyan | 0.037975 | 0.853874 | HIST1H2BI | green | 0.006427949 | 0.974615307 | DCUN1D5 | cyan | -0.18487 | 0.365931 |
| SCML1 | cyan | 0.03599 | 0.861439 | ST6GALNAC | green | 0.00698878 | 0.972401318 | MRPS30 | cyan | -0.181459 | 0.374996 |
| SNAPC1 | cyan | 0.035899 | 0.861785 | HIST1H2BD | green | 0.007458889 | 0.970545627 | COMMD4 | cyan | -0.181065 | 0.376051 |
| ZNF140 | cyan | 0.027527 | 0.893812 | UBL3 | green | 0.009706162 | 0.961677043 | SLC36A4 | cyan | -0.176391 | 0.388702 |
| CALM1 | cyan | -0.025893 | 0.900084 | PEBP1 | green | 0.009874644 | 0.961012317 | IKZF1 | cyan | -0.16998 | 0.406442 |
| NIP7 | cyan | 0.024488 | 0.905481 | PPP1R15A | green | 0.010447113 | 0.958753901 | MRPL32 | cyan | -0.169533 | 0.407696 |
| NECAP1 | cyan | -0.023934 | 0.907609 | TPCN1 | green | 0.010918772 | 0.956893414 | RWDD1 | cyan | -0.168864 | 0.409577 |
| MOSPD1 | cyan | -0.020884 | 0.919342 | PIM2 | green | 0.01198743 | 0.952678862 | AKAP5 | cyan | -0.158237 | 0.440081 |
| PHKA2 | cyan | -0.019625 | 0.92419 | PEX12 | green | 0.012941391 | 0.948917697 | TMEM8A | cyan | -0.154533 | 0.450995 |
| OPA1 | cyan | -0.019301 | 0.925438 | TCEA3 | green | 0.014320441 | 0.943482449 | PLRG1 | cyan | -0.135982 | 0.507739 |
| MANEA | cyan | 0.017891 | 0.930871 | SFXN4 | green | 0.014784813 | 0.941652762 | EMC7 | cyan | -0.133954 | 0.514148 |
| TERC | cyan | -0.016939 | 0.934542 | EGR3 | green | 0.014998723 | 0.940810028 | NOP2 | cyan | -0.133835 | 0.514527 |
| SRPK2 | cyan | 0.01585 | 0.938742 | WDR83OS | green | 0.01695954 | 0.93308805 | ARFGAP3 | cyan | -0.132353 | 0.519235 |
| TBCEL | cyan | -0.01352 | 0.947733 | GSS | green | 0.018640693 | 0.926472111 | CMAS | cyan | -0.129785 | 0.527444 |
| RNF4 | cyan | -0.009428 | 0.963542 | WDR49 | green | 0.019095961 | 0.924681269 | MRPL22 | cyan | -0.121131 | 0.555561 |
| RBM25 | cyan | 0.008044 | 0.968892 | IFIT3 | green | 0.020170351 | 0.920456487 | BRF1 | cyan | -0.120322 | 0.558223 |
| CREBL2 | cyan | 0.008042 | 0.968897 | HIST1H4K | green | 0.0202932 | 0.919973544 | MRPL15 | cyan | -0.118276 | 0.564983 |
| KAT6B | cyan | 0.007726 | 0.97012 | TARP | green | 0.020820568 | 0.917900681 | ZNF616 | cyan | -0.109396 | 0.594741 |
| PDCD6IP | darkgreen | -0.5472 | 0.003815 | TEX14 | green | 0.021386653 | 0.915676219 | GPATCH4 | cyan | -0.103384 | 0.61526 |
| MT1F | darkgreen | -0.435898 | 0.026013 | NUFIP2 | green | 0.022096811 | 0.912886491 | CLDN23 | cyan | -0.098045 | 0.63372 |
| IL10RA | darkgreen | -0.416794 | 0.034159 | NAIP | green | 0.02331127 | 0.908118062 | DEXI | cyan | -0.096404 | 0.639438 |
| ARSA | darkgreen | -0.406325 | 0.039419 | TRIB3 | green | 0.024266261 | 0.904370577 | SZRD1 | cyan | -0.090212 | 0.661196 |
| UNC13D | darkgreen | -0.399026 | 0.043452 | DLC1 | green | 0.025650625 | 0.898941755 | ATL3 | cyan | -0.090137 | 0.66146 |
| C12orf10 | darkgreen | -0.374871 | 0.059168 | FLJ43663 | green | 0.026709224 | 0.894793409 | EIF1AX | cyan | -0.08069 | 0.695172 |
| POLE4 | darkgreen | -0.371561 | 0.061628 | PEX3 | green | 0.027094154 | 0.893285639 | RAPGEF6 | cyan | -0.064714 | 0.753461 |

**Table 6. Gene module membership and correlation with growth rate (48/72h), with adjusted p values for each module**

| | | | | | | | | | | | |
|---|---|---|---|---|---|---|---|---|---|---|---|
| ENTPD6 | darkgreen | -0.370997 | 0.062055 | FCGR1C | green | 0.027219421 | 0.892795047 | CWC25 | cyan | -0.060306 | 0.769795 |
| BID | darkgreen | -0.369909 | 0.062885 | IFT122 | green | 0.029197016 | 0.885055271 | HIAT1 | cyan | -0.056206 | 0.78507 |
| SETD3 | darkgreen | -0.353808 | 0.076194 | RSBN1L-AS1 | green | 0.029399955 | 0.884261593 | TACO1 | cyan | -0.053659 | 0.794602 |
| NR4A1 | darkgreen | -0.347684 | 0.081784 | ARSD | green | 0.029432384 | 0.884134777 | VPS13B | cyan | -0.053516 | 0.795138 |
| FAM156B | darkgreen | 0.340926 | 0.08831 | HSPA1A | green | 0.030240242 | 0.880976464 | SLC25A11 | cyan | -0.053024 | 0.796984 |
| MT1X | darkgreen | -0.340319 | 0.088914 | DARS | green | 0.03133725 | 0.876690577 | TXNDC11 | cyan | -0.047473 | 0.817865 |
| MTIF2 | darkgreen | 0.339521 | 0.089713 | NR1D1 | green | 0.031345252 | 0.876659323 | KLF8 | cyan | -0.034774 | 0.866076 |
| RNASEH2B | darkgreen | -0.326335 | 0.103727 | HIST2H3D | green | 0.032756874 | 0.871149311 | HDGF | cyan | -0.032636 | 0.874246 |
| NEAT1 | darkgreen | 0.321426 | 0.109338 | HLA-C | green | 0.033275587 | 0.869126068 | ALG13 | cyan | -0.032148 | 0.876111 |
| PCNXL4 | darkgreen | 0.304262 | 0.13073 | ARHGAP24 | green | 0.033983529 | 0.866366035 | MB21D2 | cyan | -0.029591 | 0.885897 |
| VPS53 | darkgreen | -0.297614 | 0.139782 | EHHADH | green | 0.035954001 | 0.858691962 | FBXL5 | cyan | -0.020652 | 0.920235 |
| SMS | darkgreen | -0.284288 | 0.159268 | RWDD2A | green | 0.036690718 | 0.855825967 | TFAP4 | cyan | -0.016939 | 0.934543 |
| FGD2 | darkgreen | -0.283651 | 0.160246 | CCL15 | green | 0.037320555 | 0.853377174 | WDR55 | cyan | -0.015583 | 0.939774 |
| MRPL27 | darkgreen | -0.278264 | 0.16868 | STAP2 | green | 0.038121313 | 0.850265757 | USP9X | cyan | -0.010848 | 0.958056 |
| MT1E | darkgreen | -0.274718 | 0.174399 | C3orf18 | green | 0.038318752 | 0.849498927 | CRYL1 | cyan | -0.005446 | 0.978934 |
| POGK | darkgreen | 0.253743 | 0.211007 | CREBL2 | green | 0.039343186 | 0.845522294 | PREB | cyan | -0.003139 | 0.987856 |
| UBE2A | darkgreen | -0.253508 | 0.211444 | NMRK1 | green | 0.03982086 | 0.843669323 | TAF15 | cyan | 0.00055 | 0.997871 |
| RPL35A | darkgreen | -0.253087 | 0.212229 | RARRES1 | green | 0.040834735 | 0.839739047 | MED17 | cyan | 0.004585 | 0.982266 |
| SF1 | darkgreen | -0.2436 | 0.230451 | TLR8 | green | 0.040954536 | 0.839274885 | NGRN | cyan | 0.009811 | 0.962061 |
| POP7 | darkgreen | -0.241216 | 0.235188 | TMX4 | green | 0.04194541 | 0.835437831 | GPATCH2 | cyan | 0.01027 | 0.960286 |
| SCARNA2 | darkgreen | -0.239939 | 0.237751 | KCNAB1 | green | 0.04245609 | 0.833461703 | NAA30 | cyan | 0.027153 | 0.895247 |
| CAPN2 | darkgreen | -0.235111 | 0.247611 | KLLN | green | 0.042770938 | 0.832243862 | ZNF609 | cyan | 0.028056 | 0.891784 |
| CTTNBP2NL | darkgreen | -0.233838 | 0.250255 | RBM17 | green | 0.042939094 | 0.831593579 | ATF2 | cyan | 0.029306 | 0.886991 |
| SIGLEC1 | darkgreen | -0.230259 | 0.257785 | ENOSF1 | green | 0.044312023 | 0.826288389 | NDUFS6 | cyan | 0.033524 | 0.87085 |
| SERBP1 | darkgreen | -0.228192 | 0.2622 | ACOT1 | green | 0.044581295 | 0.825248747 | LOC728407 | cyan | 0.03439 | 0.867543 |
| TCEA1 | darkgreen | -0.209133 | 0.305212 | PACRGL | green | 0.04638436 | 0.818294729 | ZNF780B | cyan | 0.048683 | 0.813303 |
| CCDC92 | darkgreen | -0.208919 | 0.305718 | RFFL | green | 0.049580704 | 0.80600037 | RCN3 | cyan | 0.063887 | 0.756517 |
| SNORA14A | darkgreen | -0.203297 | 0.319208 | SNORA68 | green | 0.0508371 | 0.801179921 | SLC46A2 | cyan | 0.065546 | 0.75039 |
| LAMP1 | darkgreen | -0.199747 | 0.327913 | TPD52 | green | 0.051536277 | 0.798500428 | ZNF266 | cyan | 0.069248 | 0.736769 |
| TMEM175 | darkgreen | -0.196688 | 0.335525 | CXCL11 | green | 0.053681332 | 0.790293838 | ZFP36L1 | cyan | 0.080036 | 0.697528 |
| USO1 | darkgreen | -0.18094 | 0.376387 | SUCNR1 | green | 0.055343215 | 0.783950697 | ACACB | cyan | 0.093538 | 0.649473 |
| RNF8 | darkgreen | -0.179576 | 0.380055 | DST | green | 0.055614399 | 0.782916897 | NR3C1 | cyan | 0.095446 | 0.642788 |
| PAPSS2 | darkgreen | -0.171506 | 0.402178 | CBX6 | green | 0.057653127 | 0.775156546 | FKBP9 | cyan | 0.096233 | 0.640036 |
| SFT2D2 | darkgreen | 0.169491 | 0.407814 | WDHD1 | green | 0.058021673 | 0.773755913 | CDK2 | cyan | 0.100016 | 0.626878 |
| MCCC2 | darkgreen | -0.163143 | 0.42585 | ERAP2 | green | 0.059511247 | 0.768101966 | PDK2 | cyan | 0.107189 | 0.602239 |
| SLC2A9 | darkgreen | -0.155943 | 0.446823 | MTHFS | green | 0.061669717 | 0.759929684 | BDH2 | cyan | 0.107772 | 0.600254 |
| PTPN4 | darkgreen | 0.154237 | 0.451871 | PHLDA1 | green | 0.064511302 | 0.749209396 | ASH1L | cyan | 0.113781 | 0.579961 |
| DOPEY1 | darkgreen | 0.153962 | 0.452688 | SIAE | green | 0.064770876 | 0.748232348 | ABCA11P | cyan | 0.115781 | 0.573276 |
| SRGAP2 | darkgreen | -0.149584 | 0.465796 | DNAH17 | green | 0.066015313 | 0.743553515 | PTK2B | cyan | 0.12264 | 0.550609 |
| NEIL2 | darkgreen | -0.142604 | 0.48709 | STEAP3 | green | 0.066060048 | 0.743385484 | NLRP1 | cyan | 0.123195 | 0.548791 |
| GALNT7 | darkgreen | -0.131752 | 0.521152 | FZD5 | green | 0.067139809 | 0.739333261 | INO80 | cyan | 0.124426 | 0.544775 |
| PTGR1 | darkgreen | -0.13151 | 0.521924 | DUSP1 | green | 0.070996593 | 0.72491524 | RASA3 | cyan | 0.126207 | 0.538986 |
| LINC00641 | darkgreen | 0.125519 | 0.54122 | GLB1 | green | 0.072864783 | 0.717963719 | LRRC28 | cyan | 0.130574 | 0.524916 |
| MBTD1 | darkgreen | 0.124226 | 0.545428 | ACVR1B | green | 0.07314246 | 0.716932337 | FAM120C | cyan | 0.139247 | 0.497507 |
| TMEM57 | darkgreen | -0.122462 | 0.55119 | IL36B | green | 0.076921171 | 0.702945808 | ZNF823 | cyan | 0.140861 | 0.492485 |
| RTF1 | darkgreen | 0.113463 | 0.581026 | IL17RB | green | 0.077326768 | 0.70145003 | SIN3A | cyan | 0.148401 | 0.469369 |
| EPS15L1 | darkgreen | 0.110812 | 0.58995 | NIPSNAP3A | green | 0.077526906 | 0.700712351 | BCAS3 | cyan | 0.149272 | 0.466736 |
| INPP5B | darkgreen | 0.109768 | 0.593482 | IER5L | green | 0.07782788 | 0.699603507 | ORMDL3 | cyan | 0.157331 | 0.442737 |
| PDCD4 | darkgreen | 0.086762 | 0.673436 | CFB | green | 0.077976002 | 0.699058017 | CTPS2 | cyan | 0.15739 | 0.442565 |
| DDX55 | darkgreen | 0.08115 | 0.693517 | PDE4B | green | 0.0779891 | 0.699009789 | ZNF142 | cyan | 0.158065 | 0.440587 |
| FAM204A | darkgreen | 0.077687 | 0.706011 | CYP4V2 | green | 0.078597344 | 0.696771392 | SUN2 | cyan | 0.160542 | 0.433365 |
| CWC27 | darkgreen | -0.06612 | 0.748273 | SNHG12 | green | 0.078993803 | 0.695313714 | LOC493754 | cyan | 0.169954 | 0.406516 |
| CCT6P3 | darkgreen | -0.062221 | 0.762686 | AKR1A1 | green | 0.079135362 | 0.694793495 | SPG11 | cyan | 0.175107 | 0.392218 |
| FLVCR2 | darkgreen | -0.057911 | 0.77871 | SCFD2 | green | 0.07946533 | 0.693581411 | WDR70 | cyan | 0.175995 | 0.389784 |
| TRIM56 | darkgreen | -0.052521 | 0.798871 | ZNF296 | green | 0.080371243 | 0.69025747 | DCAF6 | cyan | 0.176807 | 0.387565 |
| BRWD1 | darkgreen | 0.047044 | 0.819487 | PHLDA2 | green | 0.081322538 | 0.68677304 | SNORA66 | cyan | 0.18299 | 0.37091 |

**Table 6. Gene module membership and correlation with growth rate (48/72h), with adjusted p values for each module**

| | | | |
|---|---|---|---|
| MGST3 | darkgreen | 0.044432 | 0.829358 |
| CCDC15 | darkgreen | 0.041394 | 0.840874 |
| ZNF562 | darkgreen | 0.025013 | 0.903462 |
| MGA | darkgreen | -0.018331 | 0.929177 |
| IREB2 | darkgreen | 0.014867 | 0.942538 |
| NXPE3 | darkgreen | -0.012944 | 0.94996 |
| NT5C3 | darkgreen | -0.009812 | 0.962059 |
| YBEY | darkgreen | 0.009731 | 0.962372 |
| MYNN | darkgreen | -0.002621 | 0.98986 |
| RPE | darkgreen | 0.002487 | 0.99038 |
| RBBP8 | darkgrey | -0.377483 | 0.057283 |
| GMIP | darkgrey | -0.335082 | 0.094262 |
| FAM40A | darkgrey | 0.277917 | 0.169235 |
| ARHGAP9 | darkgrey | -0.272461 | 0.178109 |
| PCTP | darkgrey | -0.26897 | 0.183956 |
| TMEM38B | darkgrey | -0.240907 | 0.235806 |
| SDCCAG8 | darkgrey | -0.214542 | 0.292585 |
| HIVEP2 | darkgrey | -0.207266 | 0.309646 |
| SREK1 | darkgrey | -0.206039 | 0.312583 |
| POMP | darkgrey | -0.201221 | 0.32428 |
| PBX2 | darkgrey | -0.200394 | 0.326314 |
| ELP6 | darkgrey | -0.198044 | 0.332139 |
| AKAP1 | darkgrey | -0.186886 | 0.360634 |
| TRIP12 | darkgrey | -0.177166 | 0.386586 |
| BPHL | darkgrey | -0.175894 | 0.390061 |
| SRRD | darkgrey | -0.174789 | 0.393093 |
| SDF4 | darkgrey | -0.173535 | 0.396549 |
| NDUFAF7 | darkgrey | -0.173155 | 0.397599 |
| PPP2R5E | darkgrey | -0.170454 | 0.405116 |
| SNRK | darkgrey | -0.159807 | 0.4355 |
| ACADS | darkgrey | 0.157842 | 0.44124 |
| VCPIP1 | darkgrey | -0.155214 | 0.448977 |
| HTRA2 | darkgrey | -0.150233 | 0.46384 |
| ACBD4 | darkgrey | -0.142114 | 0.488604 |
| TWSG1 | darkgrey | -0.140771 | 0.492764 |
| GLRX2 | darkgrey | -0.140731 | 0.492889 |
| CBLB | darkgrey | -0.139684 | 0.496145 |
| ERBB2IP | darkgrey | -0.136924 | 0.504777 |
| TSEN34 | darkgrey | -0.132014 | 0.520317 |
| FRG1 | darkgrey | -0.131201 | 0.52291 |
| IFFO2 | darkgrey | -0.129745 | 0.527574 |
| ATP6V1A | darkgrey | -0.127458 | 0.534937 |
| MX1 | darkgrey | -0.124922 | 0.543159 |
| LAPTM5 | darkgrey | -0.121718 | 0.55363 |
| RNF41 | darkgrey | -0.121431 | 0.554574 |
| DEDD | darkgrey | -0.115396 | 0.574559 |
| ECD | darkgrey | -0.115321 | 0.57481 |
| EIF3A | darkgrey | -0.112989 | 0.582618 |
| TMPPE | darkgrey | -0.096623 | 0.638674 |
| NDST1 | darkgrey | 0.092637 | 0.652641 |
| RABL6 | darkgrey | 0.088868 | 0.665952 |
| PRMT7 | darkgrey | -0.078837 | 0.701856 |
| UGCG | darkgrey | -0.07452 | 0.717506 |
| PRKD2 | darkgrey | 0.043793 | 0.83178 |
| GNB2L1 | darkgrey | 0.027358 | 0.894461 |
| RC3H2 | darkgrey | 0.026627 | 0.897263 |
| LHPP | green | 0.082205521 | 0.683544406 |
| CSGALNACT | green | 0.082268806 | 0.683313211 |
| DVL1 | green | 0.083526455 | 0.678724536 |
| PLTP | green | 0.084845731 | 0.673922994 |
| TSPAN4 | green | 0.085078313 | 0.673077791 |
| CD72 | green | 0.085095934 | 0.673013771 |
| JARID2 | green | 0.089114886 | 0.65847135 |
| RASSF4 | green | 0.092533474 | 0.64619579 |
| GBAP1 | green | 0.09300117 | 0.644523294 |
| FAM18B2 | green | 0.095110309 | 0.637001999 |
| EXOC6B | green | 0.10087716 | 0.616617307 |
| NCAPH2 | green | 0.101033734 | 0.616067609 |
| SLC25A51 | green | 0.101531427 | 0.61432166 |
| CR1 | green | 0.10481592 | 0.602851121 |
| NFKB2 | green | 0.104898627 | 0.602563451 |
| HIST1H2AB | green | 0.105153056 | 0.60167887 |
| SLC2A4RG | green | 0.105703815 | 0.599765911 |
| RNF122 | green | 0.106242419 | 0.597897672 |
| MYL12A | green | 0.106452135 | 0.597170909 |
| C7orf55 | green | 0.106840466 | 0.595826158 |
| MAP2K4 | green | 0.106847582 | 0.595801532 |
| SYNJ2 | green | 0.109245288 | 0.587527546 |
| JAG1 | green | 0.110575302 | 0.582959579 |
| TBCEL | green | 0.111216792 | 0.580761937 |
| RRP7B | green | 0.111560919 | 0.579584518 |
| TPD52L2 | green | 0.112144724 | 0.577589451 |
| MNDA | green | 0.116281908 | 0.563539039 |
| IDH1 | green | 0.119121993 | 0.553984221 |
| ABRACL | green | 0.123917543 | 0.538021841 |
| PHLPP1 | green | 0.124151943 | 0.537247213 |
| SLAMF6 | green | 0.124766781 | 0.535217852 |
| NINJ2 | green | 0.125784122 | 0.531867953 |
| APOO | green | 0.126371327 | 0.529938959 |
| TNFSF12 | green | 0.129607767 | 0.519367356 |
| CTSH | green | 0.133791679 | 0.505854121 |
| LNPEP | green | 0.136343334 | 0.497698943 |
| PRMT3 | green | 0.136771821 | 0.496335956 |
| HAAO | green | 0.137611065 | 0.493671807 |
| PDE6G | green | 0.138565549 | 0.490650588 |
| C2 | green | 0.139196236 | 0.488659413 |
| PNKD | green | 0.139412098 | 0.487978844 |
| TNFAIP8L2 | green | 0.140245639 | 0.485355366 |
| GAN | green | 0.140774259 | 0.483695317 |
| CXCL10 | green | 0.141646899 | 0.480961268 |
| MTHFD1 | green | 0.143696139 | 0.47457202 |
| CLEC12A | green | 0.145751685 | 0.46820735 |
| ACY1 | green | 0.146299206 | 0.466519556 |
| FAM131A | green | 0.146702052 | 0.465279764 |
| GCH1 | green | 0.147102551 | 0.464048902 |
| CSNK2A2 | green | 0.147963925 | 0.461407398 |
| HIST1H4A | green | 0.149398792 | 0.457024772 |
| BDH1 | green | 0.152990155 | 0.446152282 |
| PRKACB | green | 0.160516779 | 0.423821502 |
| SLC31A1 | green | 0.161336952 | 0.421425829 |
| ENTPD5 | green | 0.164241641 | 0.41300164 |
| BRAF | green | 0.164702126 | 0.411674798 |
| ATF4 | cyan | 0.183776 | 0.368824 |
| KLHL21 | cyan | 0.184414 | 0.367135 |
| NCOA7 | cyan | 0.186823 | 0.360797 |
| ZNF586 | cyan | 0.189683 | 0.35336 |
| ARHGAP35 | cyan | 0.203674 | 0.318293 |
| MAP3K12 | cyan | 0.21938 | 0.281573 |
| DGKQ | cyan | 0.227486 | 0.26372 |
| TLE4 | cyan | 0.243966 | 0.229729 |
| USP3 | cyan | 0.252572 | 0.213192 |
| LURAP1 | cyan | 0.253216 | 0.211989 |
| DCAF12 | cyan | 0.259408 | 0.200645 |
| FOXO3 | cyan | 0.284949 | 0.158259 |
| IDUA | cyan | 0.285838 | 0.156908 |
| HINFP | cyan | 0.293847 | 0.145106 |
| EPHB2 | cyan | 0.29715 | 0.140431 |
| ZNF506 | cyan | 0.310303 | 0.122881 |
| SLC7A5P1 | cyan | 0.314645 | 0.117456 |
| LOC146880 | cyan | 0.362098 | 0.069097 |
| SPPL2B | cyan | 0.380929 | 0.054866 |
| RBCK1 | cyan | 0.388369 | 0.04992 |
| ASB7 | cyan | 0.4051 | 0.040074 |
| HECA | cyan | 0.408317 | 0.038371 |
| ADCK3 | cyan | 0.411469 | 0.036759 |
| TBRG1 | cyan | 0.417032 | 0.034046 |
| ADCY7 | cyan | 0.41904 | 0.033107 |
| CNNM3 | cyan | 0.448094 | 0.021692 |
| RNF216 | cyan | 0.44919 | 0.021334 |
| NPEPL1 | cyan | 0.475303 | 0.014131 |
| LOC1001328 | darkgreen | -0.388354 | 0.049929 |
| ZWILCH | darkgreen | -0.334189 | 0.095197 |
| ZNF639 | darkgreen | -0.326416 | 0.103636 |
| CLIP1 | darkgreen | -0.308314 | 0.125426 |
| C4orf29 | darkgreen | -0.295285 | 0.143057 |
| UTRN | darkgreen | -0.288782 | 0.152494 |
| KPNA1 | darkgreen | -0.273481 | 0.176427 |
| CKS1B | darkgreen | -0.272662 | 0.177778 |
| STX3 | darkgreen | -0.264139 | 0.192263 |
| MFSD11 | darkgreen | -0.259725 | 0.200076 |
| UBAP1 | darkgreen | -0.257493 | 0.204109 |
| PAICS | darkgreen | -0.255237 | 0.20824 |
| AMMECR1L | darkgreen | -0.254734 | 0.209169 |
| UBP1 | darkgreen | -0.230617 | 0.257026 |
| COQ10B | darkgreen | -0.212286 | 0.29781 |
| ARNTL2 | darkgreen | -0.209903 | 0.303393 |
| CRADD | darkgreen | -0.208518 | 0.306667 |
| SERPINB1 | darkgreen | -0.208143 | 0.307557 |
| DAZAP2 | darkgreen | -0.205292 | 0.314379 |
| SPCS3 | darkgreen | -0.203165 | 0.319529 |
| FBXO11 | darkgreen | -0.196153 | 0.336869 |
| SH3PXD2A | darkgreen | -0.194342 | 0.341437 |
| UBE3C | darkgreen | -0.191063 | 0.349802 |
| MGST1 | darkgreen | -0.185029 | 0.36551 |
| RIOK2 | darkgreen | -0.179667 | 0.379809 |
| ZNF792 | darkgreen | -0.178083 | 0.384093 |
| CTSA | darkgreen | -0.177166 | 0.386586 |
| GLRX3 | darkgreen | -0.161845 | 0.429589 |

**Table 6. Gene module membership and correlation with growth rate (48/72h), with adjusted p values for each module**

| | | | |
|---|---|---|---|
| TSR3 | darkgrey | 0.01967 | 0.924019 |
| STX17 | darkgrey | -0.019605 | 0.924269 |
| SLC12A9 | darkgrey | 0.017282 | 0.93322 |
| LEPROTL1 | darkgrey | -0.009603 | 0.962866 |
| RIC8B | darkgrey | 0.005816 | 0.977503 |
| ELP4 | darkorange | -0.295163 | 0.14323 |
| RMI1 | darkorange | -0.27981 | 0.166228 |
| AKR1B1 | darkorange | -0.254519 | 0.209566 |
| FAM60A | darkorange | -0.249287 | 0.219407 |
| ZFP161 | darkorange | -0.246533 | 0.22471 |
| SNHG6 | darkorange | -0.226658 | 0.26551 |
| TUBGCP3 | darkorange | -0.217828 | 0.285077 |
| NUDT3 | darkorange | -0.207375 | 0.309387 |
| ID2 | darkorange | -0.184467 | 0.366995 |
| MTMR12 | darkorange | -0.180234 | 0.378283 |
| FAM43A | darkorange | -0.1508 | 0.462135 |
| C19orf25 | darkorange | 0.149063 | 0.467367 |
| TSNAX | darkorange | -0.14059 | 0.493325 |
| NPHS1 | darkorange | 0.134984 | 0.510887 |
| NIPAL2 | darkorange | -0.131671 | 0.52141 |
| FPGS | darkorange | -0.125758 | 0.540442 |
| CCNE2 | darkorange | -0.123788 | 0.546856 |
| ALMS1 | darkorange | -0.123749 | 0.546982 |
| TMEM165 | darkorange | -0.114659 | 0.577022 |
| SUGT1 | darkorange | -0.113239 | 0.581779 |
| TXNDC16 | darkorange | -0.096925 | 0.63762 |
| LOC644656 | darkorange | -0.096423 | 0.639371 |
| ADRA2B | darkorange | 0.085251 | 0.678819 |
| RPF2 | darkorange | -0.071145 | 0.729819 |
| VPS18 | darkorange | -0.070066 | 0.733768 |
| PLXNA1 | darkorange | -0.064415 | 0.754567 |
| COPG1 | darkorange | -0.060639 | 0.768556 |
| EFNB2 | darkorange | -0.059943 | 0.771142 |
| PIR | darkorange | -0.057108 | 0.781703 |
| HSD17B7 | darkorange | 0.056412 | 0.784303 |
| RAC3 | darkorange | -0.053438 | 0.795432 |
| TMEM236 | darkorange | -0.051854 | 0.801373 |
| AKR7A2P1 | darkorange | -0.04207 | 0.838312 |
| TMEM115 | darkorange | 0.038725 | 0.851019 |
| FTSJ1 | darkorange | 0.037909 | 0.854126 |
| PIAS2 | darkorange | -0.033853 | 0.869594 |
| POLR2C | darkorange | -0.03264 | 0.87423 |
| PRDM4 | darkorange | -0.025215 | 0.902686 |
| POU3F1 | darkorange | -0.015376 | 0.940571 |
| GAS5-AS1 | darkorange | -0.007098 | 0.97255 |
| SNX5 | darkorange | 0.001088 | 0.99579 |
| CD180 | darkred | -0.50955 | 0.007839 |
| PPIG | darkred | -0.437604 | 0.02537 |
| ERCC6 | darkred | -0.423439 | 0.031122 |
| LIMD1 | darkred | 0.380528 | 0.055143 |
| ZNF747 | darkred | -0.364219 | 0.067365 |
| OC1001709 | darkred | -0.351222 | 0.078517 |
| MT1JP | darkred | -0.34739 | 0.082061 |
| BEX2 | darkred | -0.344644 | 0.084672 |
| DGKA | darkred | -0.343123 | 0.086146 |
| ZNF320 | darkred | -0.333578 | 0.095841 |
| GYPC | green | 0.164876913 | 0.411171787 |
| LAP3 | green | 0.165378308 | 0.409730754 |
| FABP5 | green | 0.16625617 | 0.407214532 |
| SASH1 | green | 0.166442232 | 0.406682334 |
| WDR7 | green | 0.173795777 | 0.385962626 |
| LRMP | green | 0.1782691 | 0.373660391 |
| ALDH18A1 | green | 0.180085835 | 0.368729942 |
| HERC6 | green | 0.180841034 | 0.366691633 |
| PLEK | green | 0.181379415 | 0.36524256 |
| MSR1 | green | 0.182438996 | 0.362400475 |
| CCL23 | green | 0.188677805 | 0.345931404 |
| FKBP5 | green | 0.191095865 | 0.339670669 |
| UBE3D | green | 0.191340135 | 0.339042034 |
| RASA4 | green | 0.194757559 | 0.330320775 |
| NAAA | green | 0.195455944 | 0.328555426 |
| LRRC59 | green | 0.204113639 | 0.307149542 |
| HPS5 | green | 0.205214779 | 0.304490647 |
| BAG3 | green | 0.20532506 | 0.304225148 |
| RTP4 | green | 0.206860872 | 0.300542702 |
| RGMB | green | 0.207911967 | 0.298038614 |
| TOR1B | green | 0.20930619 | 0.294737313 |
| DYRK4 | green | 0.211394022 | 0.289836838 |
| B3GNTL1 | green | 0.216980968 | 0.276978146 |
| FDX1 | green | 0.217169364 | 0.276551004 |
| KAL1 | green | 0.219501405 | 0.271298596 |
| ITPR1 | green | 0.220917775 | 0.268140051 |
| FRMD4A | green | 0.221644264 | 0.266529199 |
| PKD2 | green | 0.227326426 | 0.254145943 |
| MAP1LC3A | green | 0.234330029 | 0.23940814 |
| WDR67 | green | 0.236214419 | 0.235541451 |
| MPZL2 | green | 0.241125294 | 0.22566027 |
| BDH2 | green | 0.245049885 | 0.21796614 |
| TNF | green | 0.245463998 | 0.217164731 |
| GALNT1 | green | 0.269583863 | 0.173877904 |
| DDX51 | green | 0.277610631 | 0.160919857 |
| DOK1 | green | 0.281555056 | 0.154809582 |
| ARHGAP25 | green | 0.286164981 | 0.147880119 |
| SORT1 | green | 0.290619768 | 0.141397899 |
| B3GNT2 | green | 0.292235575 | 0.139098135 |
| BLOC1S6 | green | 0.309138323 | 0.116639754 |
| SPARC | green | 0.311588098 | 0.113620934 |
| TBC1D9 | green | 0.319860891 | 0.103852705 |
| SUCLG2 | green | 0.328220826 | 0.094631879 |
| CROT | green | 0.331048166 | 0.091657082 |
| ZNF248 | green | 0.333790307 | 0.088839816 |
| HSD11B1 | green | 0.334733003 | 0.087886574 |
| C17orf49 | green | 0.336842403 | 0.085781628 |
| PPM1K | green | 0.360875641 | 0.064412788 |
| TARS | green | 0.368638193 | 0.058475578 |
| CRELD1 | green | 0.394123193 | 0.041931604 |
| DENND5A | green | 0.425428116 | 0.026945592 |
| CSTF1 | greenyellow | -0.453641655 | 0.017471442 |
| WDR3 | greenyellow | -0.440108685 | 0.021599906 |
| TGS1 | greenyellow | -0.354600551 | 0.069544961 |
| TRDMT1 | greenyellow | -0.341595928 | 0.081178459 |
| TMEM242 | greenyellow | -0.335659369 | 0.086957404 |
| CGGBP1 | darkgreen | -0.159907 | 0.43521 |
| TPM4 | darkgreen | -0.155867 | 0.447047 |
| CDK13 | darkgreen | -0.145188 | 0.479149 |
| VWA8 | darkgreen | -0.144294 | 0.481891 |
| RNMT | darkgreen | -0.144092 | 0.48251 |
| SRPK1 | darkgreen | -0.142208 | 0.488314 |
| MPZL1 | darkgreen | -0.135096 | 0.510535 |
| MYCBP | darkgreen | -0.128779 | 0.530677 |
| BAZ1A | darkgreen | -0.125577 | 0.541031 |
| VWA9 | darkgreen | -0.120687 | 0.55702 |
| PPP4R2 | darkgreen | -0.117661 | 0.56702 |
| CLMN | darkgreen | -0.115948 | 0.572717 |
| PECR | darkgreen | -0.114172 | 0.578653 |
| ITFG1 | darkgreen | -0.110916 | 0.589598 |
| BACH1 | darkgreen | -0.101939 | 0.620234 |
| KLHL2 | darkgreen | -0.096736 | 0.638281 |
| CCDC14 | darkgreen | -0.092636 | 0.652645 |
| BEX4 | darkgreen | -0.081929 | 0.690718 |
| PAPSS1 | darkgreen | -0.071185 | 0.729674 |
| BAZ1B | darkgreen | -0.068746 | 0.738612 |
| ABCA5 | darkgreen | -0.067705 | 0.742436 |
| PAPLN | darkgreen | -0.06033 | 0.769705 |
| NCBP2 | darkgreen | -0.055813 | 0.78654 |
| RGPD6 | darkgreen | -0.04653 | 0.821426 |
| ZC3H14 | darkgreen | -0.044636 | 0.828586 |
| DIAPH1 | darkgreen | -0.04295 | 0.834974 |
| FAM98C | darkgreen | -0.042542 | 0.83652 |
| SAMD9 | darkgreen | -0.039347 | 0.848652 |
| ZBED5 | darkgreen | -0.035156 | 0.864619 |
| ITGAV | darkgreen | -0.033913 | 0.869366 |
| FBXO28 | darkgreen | -0.028555 | 0.88987 |
| YY1 | darkgreen | -0.027567 | 0.893658 |
| CLASRP | darkgreen | -0.025026 | 0.903413 |
| SWAP70 | darkgreen | -0.021655 | 0.916374 |
| TMCO3 | darkgreen | -0.005141 | 0.980116 |
| CHST3 | darkgreen | -0.002197 | 0.9915 |
| GNAI2 | darkgreen | 0.000571 | 0.997792 |
| ASAP1 | darkgreen | 0.011688 | 0.954809 |
| SFSWAP | darkgreen | 0.013244 | 0.948803 |
| SLC9A8 | darkgreen | 0.026409 | 0.8981 |
| LRWD1 | darkgreen | 0.044031 | 0.830879 |
| RUNDC1 | darkgreen | 0.051343 | 0.803292 |
| ZNF574 | darkgreen | 0.056507 | 0.783947 |
| TCTEX1D4 | darkgreen | 0.057577 | 0.779955 |
| USP32 | darkgreen | 0.063521 | 0.757871 |
| PTPLAD1 | darkgreen | 0.06811 | 0.740949 |
| BTBD3 | darkgreen | 0.072403 | 0.725221 |
| BRD2 | darkgreen | 0.096928 | 0.637612 |
| RASL11A | darkgreen | 0.107464 | 0.601302 |
| CCDC101 | darkgreen | 0.10872 | 0.597032 |
| GPR77 | darkgreen | 0.109673 | 0.593801 |
| PIK3CG | darkgreen | 0.130942 | 0.523739 |
| ALDH3B1 | darkgreen | 0.134607 | 0.512081 |
| TCERG1 | darkgreen | 0.170664 | 0.404528 |
| DNASE1L1 | darkgreen | 0.189283 | 0.354394 |
| NUDT18 | darkgreen | 0.207926 | 0.308073 |

**Table 6. Gene module membership and correlation with growth rate (48/72h), with adjusted p values for each module**

| | | | | | | | | | | | |
|---|---|---|---|---|---|---|---|---|---|---|---|
| EXOC2 | darkred | -0.33248 | 0.097007 | OXA1L | greenyellow | -0.324481606 | 0.098676873 | VAMP2 | darkgreen | 0.240014 | 0.2376 |
| NUP155 | darkred | -0.32552 | 0.104644 | FOXN2 | greenyellow | -0.316558333 | 0.10767419 | MOB3C | darkgreen | 0.268164 | 0.185325 |
| ZNF28 | darkred | -0.305588 | 0.128977 | BCRP3 | greenyellow | -0.299326376 | 0.129324877 | RCC1 | darkgreen | 0.292038 | 0.147716 |
| LSM4 | darkred | -0.300968 | 0.13516 | FBXO3 | greenyellow | -0.295484672 | 0.134555755 | ZMYM3 | darkgreen | 0.303884 | 0.131233 |
| HIST2H2BC | darkred | -0.295224 | 0.143143 | HDAC8 | greenyellow | -0.293577585 | 0.137208708 | HDAC5 | darkgreen | 0.353449 | 0.076513 |
| LOC595101 | darkred | -0.291827 | 0.148021 | OXSR1 | greenyellow | -0.289141758 | 0.143525431 | RAB33B | darkgreen | 0.411088 | 0.036951 |
| LAS1L | darkred | -0.285941 | 0.156753 | VCPIP1 | greenyellow | -0.276534462 | 0.162616214 | IRF5 | darkgreen | 0.418647 | 0.033289 |
| SATB1 | darkred | -0.285179 | 0.157908 | RAD23B | greenyellow | -0.272251386 | 0.169493005 | KIAA0913 | darkgreen | 0.482236 | 0.0126 |
| NOMO3 | darkred | -0.281886 | 0.162976 | SMARCC1 | greenyellow | -0.272220277 | 0.169543689 | PIK3R5 | darkgreen | 0.501184 | 0.009102 |
| SUCLG2 | darkred | 0.27958 | 0.166591 | MAN1A1 | greenyellow | -0.268705911 | 0.175338325 | RANBP9 | darkgrey | -0.394952 | 0.045842 |
| TM9SF2 | darkred | -0.274198 | 0.17525 | DNAJC5 | greenyellow | -0.261509148 | 0.187634168 | PDZD11 | darkgrey | -0.354908 | 0.075221 |
| MKRN2 | darkred | -0.267454 | 0.186536 | TMTC3 | greenyellow | -0.252482994 | 0.203883162 | SETD3 | darkgrey | -0.350497 | 0.079179 |
| ABCB7 | darkred | -0.267176 | 0.187011 | GOT1 | greenyellow | -0.252297274 | 0.20422727 | UBAP2 | darkgrey | -0.320379 | 0.110565 |
| TOMM7 | darkred | -0.266369 | 0.188397 | NRF1 | greenyellow | -0.248987325 | 0.210426646 | INPP5F | darkgrey | -0.318609 | 0.112658 |
| RRP9 | darkred | -0.265841 | 0.189307 | UNC119 | greenyellow | -0.245144758 | 0.217782363 | RNF2 | darkgrey | -0.303361 | 0.131932 |
| CARD6 | darkred | -0.264913 | 0.190915 | WHSC1L1 | greenyellow | -0.235235626 | 0.237544683 | CTAGE4 | darkgrey | -0.300203 | 0.136205 |
| PYCR2 | darkred | -0.264195 | 0.192166 | MAP2K2 | greenyellow | -0.235083864 | 0.237856293 | GNL3 | darkgrey | -0.298629 | 0.138371 |
| MTHFS | darkred | -0.263538 | 0.193314 | CCDC57 | greenyellow | -0.233035049 | 0.242089597 | SSU72 | darkgrey | -0.289769 | 0.151034 |
| PEX11A | darkred | -0.262857 | 0.194511 | EFTUD1 | greenyellow | -0.232918282 | 0.242332353 | SEC11C | darkgrey | -0.288536 | 0.15286 |
| FTSJD1 | darkred | -0.26065 | 0.198422 | ANAPC5 | greenyellow | -0.230904864 | 0.246543458 | NAPB | darkgrey | -0.28201 | 0.162782 |
| FAM100A | darkred | -0.254997 | 0.208682 | SH3BGRL3 | greenyellow | -0.230878675 | 0.246598549 | LRRC42 | darkgrey | -0.254865 | 0.208926 |
| TMX2 | darkred | 0.247578 | 0.222688 | SETD7 | greenyellow | -0.221657429 | 0.266500066 | RRAGD | darkgrey | -0.248749 | 0.220436 |
| SIDT2 | darkred | 0.245478 | 0.226763 | DIXDC1 | greenyellow | -0.220185316 | 0.269770481 | GNG10 | darkgrey | -0.248554 | 0.220811 |
| MBTPS2 | darkred | -0.244718 | 0.228249 | UCK1 | greenyellow | -0.21699841 | 0.276938582 | TMEM251 | darkgrey | -0.247852 | 0.22216 |
| FAM127A | darkred | -0.242641 | 0.232348 | TGFBR1 | greenyellow | -0.209650352 | 0.293925945 | TTC39B | darkgrey | -0.232506 | 0.253041 |
| HERC3 | darkred | -0.238254 | 0.241163 | RNF167 | greenyellow | -0.19204159 | 0.337240713 | GSPT1 | darkgrey | -0.226867 | 0.265058 |
| TPRA1 | darkred | -0.238004 | 0.241672 | ATP6V1C1 | greenyellow | -0.190182333 | 0.342027883 | NDUFA9 | darkgrey | -0.224063 | 0.271168 |
| LGALS1 | darkred | -0.237358 | 0.242989 | ZNF124 | greenyellow | -0.183374306 | 0.359902556 | TIMM17A | darkgrey | -0.220354 | 0.279387 |
| ISCA1 | darkred | -0.236536 | 0.244673 | ANAPC1 | greenyellow | -0.179942904 | 0.369116459 | CSF1 | darkgrey | -0.218557 | 0.283427 |
| MEF2D | darkred | -0.232162 | 0.253764 | SMA5 | greenyellow | -0.177159766 | 0.376689751 | SRM | darkgrey | -0.214978 | 0.291581 |
| TUBB4B | darkred | -0.229752 | 0.258864 | TXNDC9 | greenyellow | -0.171973044 | 0.391041145 | MPP1 | darkgrey | -0.197615 | 0.333208 |
| PPP1R12C | darkred | -0.228362 | 0.261837 | SEL1L3 | greenyellow | -0.168323993 | 0.401321823 | SCARB1 | darkgrey | -0.193957 | 0.342414 |
| ZNF440 | darkred | -0.219128 | 0.282139 | TTC39B | greenyellow | -0.168115891 | 0.401912669 | SET | darkgrey | -0.186884 | 0.36064 |
| OC1001322[illegible] | darkred | -0.218945 | 0.282551 | DDX1 | greenyellow | -0.165108836 | 0.410504878 | SLC48A1 | darkgrey | -0.18103 | 0.376145 |
| KIAA0494 | darkred | -0.215989 | 0.289262 | PJA1 | greenyellow | -0.160984618 | 0.422454059 | AFAP1L1 | darkgrey | -0.177743 | 0.385019 |
| ZNF598 | darkred | -0.213713 | 0.294499 | XBP1 | greenyellow | -0.159666199 | 0.426313866 | LDHB | darkgrey | -0.172622 | 0.399077 |
| FCGR2A | darkred | -0.21267 | 0.296916 | MAP3K1 | greenyellow | -0.156213792 | 0.436512043 | RNF125 | darkgrey | -0.163447 | 0.424977 |
| ZDHHC9 | darkred | -0.208012 | 0.307869 | SNRPD3 | greenyellow | -0.153511406 | 0.444585825 | PIPOX | darkgrey | -0.151483 | 0.460085 |
| PPARG | darkred | -0.206941 | 0.310423 | OR2A7 | greenyellow | -0.153283686 | 0.445269804 | KCNK13 | darkgrey | -0.149327 | 0.46657 |
| LYPLA2 | darkred | -0.204359 | 0.316633 | FAM114A2 | greenyellow | -0.152721949 | 0.44695944 | TEX10 | darkgrey | -0.145236 | 0.479003 |
| FAM21B | darkred | -0.200363 | 0.326391 | RCAN3 | greenyellow | -0.150050546 | 0.455041347 | OPN3 | darkgrey | -0.127095 | 0.536112 |
| RPL23AP7 | darkred | -0.190859 | 0.350327 | IGFBP2 | greenyellow | -0.145689832 | 0.468398217 | TSPAN13 | darkgrey | -0.126353 | 0.538511 |
| YIF1A | darkred | -0.189576 | 0.353635 | ARL13B | greenyellow | -0.134463624 | 0.503700181 | TSEN54 | darkgrey | -0.12557 | 0.541055 |
| BASP1 | darkred | -0.178358 | 0.383349 | YTHDF3 | greenyellow | -0.109028925 | 0.588272118 | FAM127B | darkgrey | -0.115016 | 0.575829 |
| ZNF625 | darkred | -0.177332 | 0.386134 | MLEC | greenyellow | -0.106184034 | 0.598100071 | TLR8 | darkgrey | -0.108547 | 0.597621 |
| LRIG2 | darkred | -0.17711 | 0.38674 | NDFIP1 | greenyellow | -0.102210067 | 0.611944236 | LIMK1 | darkgrey | -0.069613 | 0.735431 |
| R3HCC1L | darkred | 0.17557 | 0.390948 | HNRNPH3 | greenyellow | -0.102073531 | 0.612422243 | ZNF3 | darkgrey | -0.067938 | 0.741579 |
| ZNF595 | darkred | -0.172118 | 0.400476 | B3GALNT2 | greenyellow | -0.098085389 | 0.626452259 | ADTRP | darkgrey | -0.062992 | 0.759831 |
| NBPF10 | darkred | -0.159912 | 0.435195 | PDIA3P | greenyellow | -0.097849939 | 0.627284599 | SOCS7 | darkgrey | -0.050155 | 0.807759 |
| TLR8 | darkred | -0.154734 | 0.450398 | ZNF740 | greenyellow | -0.089565837 | 0.65684702 | IQCE | darkgrey | -0.044002 | 0.830987 |
| ZNF765 | darkred | -0.148864 | 0.467968 | CASD1 | greenyellow | -0.070631683 | 0.726275573 | RTCA | darkgrey | -0.043957 | 0.831158 |
| SYNJ2BP | darkred | -0.129074 | 0.529728 | FTH1 | greenyellow | -0.065665311 | 0.744868562 | CIITA | darkgrey | -0.024699 | 0.90467 |
| TP53BP1 | darkred | 0.114718 | 0.576826 | FTL | greenyellow | -0.063411981 | 0.753351475 | NCOA5 | darkgrey | -0.020842 | 0.919504 |
| ATF1 | darkred | 0.089254 | 0.664586 | ECSIT | greenyellow | -0.061859517 | 0.759212263 | ZNF79 | darkgrey | -0.018143 | 0.929902 |
| FLJ43663 | darkred | 0.081875 | 0.690913 | TAPBPL | greenyellow | -0.058403487 | 0.772305582 | SORD | darkgrey | -0.008437 | 0.967372 |
| SELO | darkred | -0.076525 | 0.710221 | QRICH1 | greenyellow | -0.039319344 | 0.845614802 | SLC24A4 | darkgrey | -0.005948 | 0.976995 |

**Table 6. Gene module membership and correlation with growth rate (48/72h), with adjusted p values for each module**

| | | | | | | | | | | | |
|---|---|---|---|---|---|---|---|---|---|---|---|
| IRGQ | darkred | -0.059026 | 0.774556 | TWSG1 | greenyellow | -0.029079584 | 0.885514591 | FMO5 | darkgrey | 0.008358 | 0.967676 |
| LRRC37B | darkred | -0.052678 | 0.798282 | GEMIN5 | greenyellow | -0.026468117 | 0.895738001 | FAM127C | darkgrey | 0.008716 | 0.966292 |
| NAT10 | darkred | -0.041741 | 0.839557 | SIGLEC7 | greenyellow | -0.025687163 | 0.898798527 | TBC1D7 | darkgrey | 0.022454 | 0.9133 |
| LARS | darkred | -0.039616 | 0.84763 | SIPA1L3 | greenyellow | -0.021856135 | 0.913831834 | SH3D21 | darkgrey | 0.030812 | 0.881224 |
| TOE1 | darkred | 0.039339 | 0.848684 | LENG1 | greenyellow | -0.020558139 | 0.918932116 | FAM49A | darkgrey | 0.045956 | 0.823594 |
| CD164 | darkred | -0.033666 | 0.870309 | C11orf63 | greenyellow | -0.018422376 | 0.927331004 | RPS29 | darkgrey | 0.046943 | 0.819867 |
| MAPK8IP3 | darkred | 0.002692 | 0.989587 | NCAM1 | greenyellow | -0.013081949 | 0.948363608 | ARID5B | darkgrey | 0.053177 | 0.79641 |
| SNAI1 | darkturquoise | -0.384157 | 0.052675 | VSIG4 | greenyellow | -0.007167587 | 0.971695482 | GIMAP2 | darkgrey | 0.05409 | 0.792989 |
| PCDH17 | darkturquoise | -0.335029 | 0.094317 | RTEL1 | greenyellow | 0.00056502 | 0.99776832 | ARF3 | darkgrey | 0.067863 | 0.741855 |
| FSCN1 | darkturquoise | -0.271992 | 0.178887 | PRAM1 | greenyellow | 0.000632526 | 0.997501692 | METTL7A | darkgrey | 0.068225 | 0.740525 |
| ARL4D | darkturquoise | -0.269365 | 0.183288 | NSUN5P1 | greenyellow | 0.002564195 | 0.989872358 | MXD4 | darkgrey | 0.084915 | 0.68002 |
| PPP1R36 | darkturquoise | -0.265453 | 0.189978 | SNRPD2 | greenyellow | 0.016000506 | 0.936864166 | LRRC8C | darkgrey | 0.099553 | 0.628484 |
| TPM2 | darkturquoise | -0.259133 | 0.201141 | PPP4C | greenyellow | 0.023232329 | 0.908427922 | OSBPL2 | darkgrey | 0.101818 | 0.620649 |
| RGS16 | darkturquoise | -0.255428 | 0.207888 | HIST1H4E | greenyellow | 0.026107607 | 0.897150649 | SH3KBP1 | darkgrey | 0.124671 | 0.543975 |
| ID1 | darkturquoise | -0.245581 | 0.226562 | CCDC124 | greenyellow | 0.031017431 | 0.877939728 | FBXO42 | darkgrey | 0.124863 | 0.543353 |
| RND1 | darkturquoise | -0.21894 | 0.282564 | SEC13 | greenyellow | 0.033509153 | 0.868215304 | SLC52A2 | darkgrey | 0.127825 | 0.533752 |
| GSPT1 | darkturquoise | -0.216344 | 0.288451 | HARS | greenyellow | 0.03721756 | 0.853777528 | ZNF674 | darkgrey | 0.130246 | 0.525966 |
| ITGA6 | darkturquoise | -0.208115 | 0.307625 | ZNF22 | greenyellow | 0.040625511 | 0.840549798 | TMEM185A | darkgrey | 0.152288 | 0.457675 |
| PLEKHG2 | darkturquoise | -0.196623 | 0.33569 | AKT1 | greenyellow | 0.050667343 | 0.80183082 | B3GNTL1 | darkgrey | 0.159081 | 0.437614 |
| TNFSF9 | darkturquoise | -0.191497 | 0.348689 | TWF2 | greenyellow | 0.050760001 | 0.801475524 | PRDM1 | darkgrey | 0.159364 | 0.436791 |
| MAFB | darkturquoise | 0.188665 | 0.355996 | CHD8 | greenyellow | 0.053905799 | 0.78943631 | CAMP | darkgrey | 0.180549 | 0.377435 |
| GADD45G | darkturquoise | -0.180698 | 0.377035 | C12orf23 | greenyellow | 0.054248995 | 0.788125671 | PPIL3 | darkgrey | 0.188978 | 0.355183 |
| FRAT1 | darkturquoise | -0.174881 | 0.392839 | PNKP | greenyellow | 0.055619741 | 0.782896535 | PIM3 | darkgrey | 0.190063 | 0.352378 |
| ZFP36L1 | darkturquoise | 0.169315 | 0.408309 | ZDHHC3 | greenyellow | 0.056071601 | 0.781174778 | TARS2 | darkgrey | 0.210146 | 0.302821 |
| PDK4 | darkturquoise | -0.164843 | 0.420977 | SNORA23 | greenyellow | 0.056869637 | 0.778136428 | NAB1 | darkgrey | 0.221558 | 0.276703 |
| PCDH10 | darkturquoise | -0.16078 | 0.432672 | MESDC1 | greenyellow | 0.060815209 | 0.763162024 | CCDC144A | darkgrey | 0.225259 | 0.26855 |
| CDC42BPG | darkturquoise | -0.156236 | 0.445961 | PHF1 | greenyellow | 0.062806756 | 0.755634724 | KIAA0319L | darkgrey | 0.238457 | 0.24075 |
| SLC15A2 | darkturquoise | 0.151733 | 0.459336 | CTSD | greenyellow | 0.068764139 | 0.733250136 | RFWD3 | darkgrey | 0.252284 | 0.213732 |
| TRNAU1AP | darkturquoise | 0.145322 | 0.478741 | COQ5 | greenyellow | 0.074083317 | 0.713441302 | TSC22D3 | darkgrey | 0.25988 | 0.199799 |
| ABAT | darkturquoise | 0.14193 | 0.489172 | TOPBP1 | greenyellow | 0.074768568 | 0.710902226 | EDEM3 | darkgrey | 0.303322 | 0.131984 |
| RASD1 | darkturquoise | -0.134055 | 0.513828 | MMD | greenyellow | 0.082862321 | 0.681146325 | PCNXL3 | darkgrey | 0.320512 | 0.110407 |
| MRPS18A | darkturquoise | -0.133189 | 0.516576 | DFFA | greenyellow | 0.088310146 | 0.661373786 | ACVR2B | darkgrey | 0.325549 | 0.10461 |
| TAF4B | darkturquoise | -0.126449 | 0.538203 | SLC25A33 | greenyellow | 0.089950968 | 0.655460982 | PHF20 | darkgrey | 0.382211 | 0.053988 |
| SLC20A1 | darkturquoise | 0.124387 | 0.544901 | PRKD2 | greenyellow | 0.094215085 | 0.640190184 | RIN2 | darkgrey | 0.384936 | 0.052157 |
| KLF5 | darkturquoise | -0.121338 | 0.55488 | HADHA | greenyellow | 0.094472981 | 0.639271089 | DONSON | darkolivegreen | -0.485764 | 0.011876 |
| CCNE1 | darkturquoise | -0.119256 | 0.561739 | EPRS | greenyellow | 0.096736725 | 0.631225946 | PRPF38A | darkolivegreen | -0.307175 | 0.126901 |
| C12orf65 | darkturquoise | -0.118609 | 0.563878 | FBXW5 | greenyellow | 0.099746913 | 0.620591333 | ATP6V0B | darkolivegreen | -0.299033 | 0.137813 |
| RASL11B | darkturquoise | 0.117915 | 0.566178 | CSAD | greenyellow | 0.10428285 | 0.604706619 | HNRNPM | darkolivegreen | -0.282165 | 0.162542 |
| NEFM | darkturquoise | -0.117848 | 0.566402 | RNF123 | greenyellow | 0.10965966 | 0.586102705 | HSPE1 | darkolivegreen | -0.26472 | 0.191252 |
| TNFRSF10D | darkturquoise | -0.113312 | 0.581533 | TRNAU1AP | greenyellow | 0.110746499 | 0.582372728 | LONP2 | darkolivegreen | -0.237723 | 0.242244 |
| NEU1 | darkturquoise | 0.107727 | 0.600405 | CHMP2A | greenyellow | 0.111043824 | 0.581354137 | CHPT1 | darkolivegreen | -0.231174 | 0.255847 |
| ENO2 | darkturquoise | 0.105743 | 0.607173 | NOTCH2NL | greenyellow | 0.113236142 | 0.573867863 | LSM12 | darkolivegreen | -0.220295 | 0.27952 |
| MPO | darkturquoise | 0.097717 | 0.634863 | IFT20 | greenyellow | 0.120508762 | 0.549345971 | LSM14B | darkolivegreen | -0.215175 | 0.291129 |
| ACTN2 | darkturquoise | -0.086755 | 0.67346 | TOB2 | greenyellow | 0.125265544 | 0.533574279 | TBCCD1 | darkolivegreen | -0.212873 | 0.296446 |
| FOSB | darkturquoise | -0.082693 | 0.687975 | WDR1 | greenyellow | 0.126824158 | 0.528453669 | FAM195A | darkolivegreen | -0.192019 | 0.34735 |
| HEXIM1 | darkturquoise | 0.071887 | 0.727105 | IER3 | greenyellow | 0.137890428 | 0.492786577 | SNRPF | darkolivegreen | -0.188982 | 0.355173 |
| STARD5 | darkturquoise | -0.065739 | 0.749677 | ARHGEF10L | greenyellow | 0.139131946 | 0.4888622 | CAPZA1 | darkolivegreen | -0.162766 | 0.426936 |
| LOC729966 | darkturquoise | 0.062895 | 0.760189 | ARAP3 | greenyellow | 0.143566861 | 0.474973794 | EIF3A | darkolivegreen | -0.149274 | 0.466729 |
| HIST2H3D | darkturquoise | 0.059878 | 0.771385 | IFI30 | greenyellow | 0.146432446 | 0.466109307 | VPS29 | darkolivegreen | -0.146986 | 0.473663 |
| PPP1R10 | darkturquoise | 0.059418 | 0.773097 | LONRF3 | greenyellow | 0.147448604 | 0.46298674 | SRRD | darkolivegreen | -0.146461 | 0.475262 |
| SEMA6B | darkturquoise | 0.05926 | 0.773683 | DCAF15 | greenyellow | 0.15161453 | 0.450300421 | LYN | darkolivegreen | -0.145714 | 0.477542 |
| SYNM | darkturquoise | -0.059252 | 0.773715 | ACTB | greenyellow | 0.167526939 | 0.403587486 | KIAA1598 | darkolivegreen | -0.145169 | 0.479207 |
| EGR3 | darkturquoise | 0.053786 | 0.794125 | MYH9 | greenyellow | 0.16753527 | 0.403563768 | TMEM161B | darkolivegreen | -0.123992 | 0.546189 |
| CDC14A | darkturquoise | -0.049697 | 0.809483 | MED10 | greenyellow | 0.168973361 | 0.399481271 | TCF12 | darkolivegreen | -0.11865 | 0.563742 |
| RAB3A | darkturquoise | 0.047705 | 0.81699 | SRGN | greenyellow | 0.175020019 | 0.382572958 | ZNF706 | darkolivegreen | -0.108246 | 0.598642 |
| SEMA3B | darkturquoise | 0.04249 | 0.836715 | SPHK1 | greenyellow | 0.176269335 | 0.379131589 | LRRFIP1 | darkolivegreen | -0.097942 | 0.634079 |

**Table 6. Gene module membership and correlation with growth rate (48/72h), with adjusted p values for each module**

| | | | | | | | | | | | |
|---|---|---|---|---|---|---|---|---|---|---|---|
| SLC7A5P2 | darkturquoise | 0.033848 | 0.869614 | WDR45L | greenyellow | 0.176881278 | 0.377452471 | UBE2N | darkolivegreen | -0.057554 | 0.780041 |
| SLC12A2 | darkturquoise | 0.028392 | 0.890496 | RPS2 | greenyellow | 0.184570049 | 0.356723924 | BTN3A2 | darkolivegreen | -0.057073 | 0.781835 |
| ZFP36 | darkturquoise | -0.027112 | 0.895404 | GLG1 | greenyellow | 0.188411637 | 0.346624743 | TMEM220 | darkolivegreen | -0.052451 | 0.799131 |
| GAN | darkturquoise | 0.023261 | 0.910196 | HIST1H4H | greenyellow | 0.192071242 | 0.337164695 | ZNF778 | darkolivegreen | 0.020489 | 0.920866 |
| RPP14 | darkturquoise | -0.022628 | 0.912633 | WDR46 | greenyellow | 0.193606742 | 0.333242303 | ZNF490 | darkolivegreen | 0.021667 | 0.91633 |
| GRASP | darkturquoise | 0.020346 | 0.921415 | VASP | greenyellow | 0.202859645 | 0.310195042 | NDST2 | darkolivegreen | 0.051299 | 0.803459 |
| NOL9 | darkturquoise | -0.011047 | 0.957285 | LY6G5B | greenyellow | 0.208204606 | 0.297343776 | GPRC5B | darkolivegreen | 0.072411 | 0.725193 |
| HIPK4 | darkturquoise | 0.009372 | 0.963759 | NT5C | greenyellow | 0.212508811 | 0.287241462 | HLA-E | darkolivegreen | 0.117254 | 0.568374 |
| SAMHD1 | green | -0.551367 | 0.003505 | DYNLRB1 | greenyellow | 0.213353652 | 0.285284398 | ACTA2 | darkolivegreen | 0.171873 | 0.401157 |
| COQ2 | green | -0.497353 | 0.009735 | GALE | greenyellow | 0.213670883 | 0.284551726 | RNF215 | darkolivegreen | 0.183301 | 0.370083 |
| POLR3GL | green | -0.467772 | 0.015963 | MAPK13 | greenyellow | 0.221149203 | 0.267626224 | RAB40C | darkolivegreen | 0.201287 | 0.324119 |
| MELK | green | -0.465751 | 0.016487 | TESK2 | greenyellow | 0.224927311 | 0.259327726 | EFCAB4A | darkolivegreen | 0.224611 | 0.269967 |
| DNAJA1 | green | 0.464562 | 0.016802 | UBFD1 | greenyellow | 0.229195517 | 0.250156136 | CCDC144B | darkolivegreen | 0.271321 | 0.180004 |
| CEP85 | green | -0.462841 | 0.017266 | RPL13 | greenyellow | 0.230466095 | 0.2474675 | GPATCH3 | darkolivegreen | 0.288328 | 0.153169 |
| MAP7D3 | green | -0.451168 | 0.020701 | DUS1L | greenyellow | 0.234824776 | 0.238388898 | FAM120B | darkolivegreen | 0.295503 | 0.142748 |
| C5orf44 | green | -0.422321 | 0.031617 | CEP76 | greenyellow | 0.238584799 | 0.230736739 | ZC3HAV1 | darkolivegreen | 0.31811 | 0.113254 |
| GPC4 | green | 0.413981 | 0.035513 | MKL1 | greenyellow | 0.245164368 | 0.21774439 | PGS1 | darkolivegreen | 0.356134 | 0.074148 |
| POLI | green | -0.401813 | 0.041876 | NEDD8 | greenyellow | 0.25684995 | 0.195905807 | ATG2A | darkolivegreen | 0.373547 | 0.060143 |
| EXOSC4 | green | -0.377865 | 0.057011 | GNB1 | greenyellow | 0.260068729 | 0.190165112 | WDFY2 | darkolivegreen | 0.43724 | 0.025506 |
| VAT1 | green | 0.376656 | 0.057875 | ADRM1 | greenyellow | 0.263421458 | 0.184310257 | NCOA6 | darkolivegreen | 0.445969 | 0.0224 |
| GSKIP | green | -0.357604 | 0.072878 | CRTC1 | greenyellow | 0.264323549 | 0.18275656 | SIKE1 | darkolivegreen | 0.50669 | 0.008253 |
| ULBP3 | green | 0.342656 | 0.086603 | MAP3K8 | greenyellow | 0.274168776 | 0.16638964 | ABCG1 | darkorange | -0.418092 | 0.033547 |
| CREB3 | green | -0.339898 | 0.089335 | MFSD10 | greenyellow | 0.276783554 | 0.162222451 | FLAD1 | darkorange | -0.395501 | 0.045514 |
| COPS6 | green | -0.337828 | 0.091429 | KIAA1967 | greenyellow | 0.277780155 | 0.160653785 | SH2B3 | darkorange | -0.371599 | 0.061599 |
| EPB41 | green | -0.336273 | 0.093025 | SH3BP2 | greenyellow | 0.279366365 | 0.158179298 | TMEM43 | darkorange | -0.317843 | 0.113574 |
| BIVM | green | -0.336246 | 0.093053 | RAB34 | greenyellow | 0.282671608 | 0.153110383 | VPS9D1 | darkorange | -0.312337 | 0.120317 |
| RANBP1 | green | -0.335104 | 0.094239 | HIPK1 | greenyellow | 0.28351764 | 0.151831764 | FLYWCH1 | darkorange | -0.311906 | 0.120857 |
| MCCC1 | green | -0.334196 | 0.095189 | ZNF511 | greenyellow | 0.288216411 | 0.1448691 | RPL38 | darkorange | -0.297186 | 0.14038 |
| NDUFS8 | green | -0.333325 | 0.096109 | WDR37 | greenyellow | 0.288450837 | 0.144527844 | TP53I3 | darkorange | -0.293996 | 0.144894 |
| KDM6A | green | -0.331238 | 0.098338 | OGFOD3 | greenyellow | 0.298095007 | 0.130985105 | APH1A | darkorange | -0.271856 | 0.179113 |
| TRAK1 | green | 0.32891 | 0.100869 | HEXA | greenyellow | 0.330168643 | 0.092574816 | SMARCD1 | darkorange | -0.27081 | 0.180858 |
| APEX1 | green | -0.325018 | 0.105211 | KIAA1467 | greenyellow | 0.338288055 | 0.084361257 | EIF3E | darkorange | -0.252596 | 0.213149 |
| USP33 | green | -0.321872 | 0.108819 | MFSD1 | greenyellow | 0.345248088 | 0.077771511 | TRPM4 | darkorange | -0.250856 | 0.216424 |
| FBXO44 | green | 0.320292 | 0.110666 | CCDC9 | greenyellow | 0.346202286 | 0.076899649 | FAM105B | darkorange | -0.249111 | 0.219743 |
| OC1002884 | green | -0.31992 | 0.111105 | DHX16 | greenyellow | 0.347385131 | 0.075829266 | EHD4 | darkorange | -0.238911 | 0.239828 |
| HIST1H2AL | green | -0.319132 | 0.112037 | ZNF335 | greenyellow | 0.368789788 | 0.058364033 | RUNX1 | darkorange | -0.234394 | 0.249098 |
| LBX2-AS1 | green | -0.31827 | 0.113063 | NUBP2 | greenyellow | 0.372098844 | 0.055970257 | C10orf88 | darkorange | -0.23172 | 0.254694 |
| EIF2D | green | -0.317993 | 0.113395 | FXYD5 | greenyellow | 0.381235928 | 0.049757447 | AAR2 | darkorange | -0.228288 | 0.261996 |
| AGXT2L2 | green | -0.316863 | 0.114753 | CSE1L | greenyellow | 0.383759287 | 0.048141079 | TMPPE | darkorange | -0.207674 | 0.308674 |
| LRP11 | green | -0.313196 | 0.119246 | JMJD6 | greenyellow | 0.389135025 | 0.044835632 | PI4KA | darkorange | -0.204294 | 0.316789 |
| SNORA30 | green | 0.311294 | 0.121628 | CTDSP2 | greenyellow | 0.425137936 | 0.027061155 | ZNF384 | darkorange | -0.201458 | 0.3237 |
| RBM23 | green | -0.311147 | 0.121812 | HEBP2 | greenyellow | 0.426576035 | 0.026492347 | ZNF420 | darkorange | -0.191191 | 0.349473 |
| AURKAIP1 | green | -0.308756 | 0.124858 | GAPDH | greenyellow | 0.44319305 | 0.020595146 | DGAT1 | darkorange | -0.188143 | 0.357353 |
| ARHGAP35 | green | 0.304846 | 0.129956 | ZNF672 | grey60 | -0.552111691 | 0.002827721 | METRNL | darkorange | -0.185894 | 0.363235 |
| MOB1A | green | -0.304569 | 0.130323 | C7orf49 | grey60 | -0.530261628 | 0.004439587 | LRRC33 | darkorange | -0.175208 | 0.391941 |
| RPS18 | green | -0.303841 | 0.13129 | VIM | grey60 | -0.454879597 | 0.017128683 | ST3GAL1 | darkorange | -0.167551 | 0.413279 |
| UBN2 | green | -0.302657 | 0.132875 | ARHGAP10 | grey60 | -0.418230314 | 0.029932709 | ZNF18 | darkorange | -0.148603 | 0.468757 |
| KLHL8 | green | -0.29981 | 0.136744 | GCFC1 | grey60 | -0.411301686 | 0.0330549 | ABCC10 | darkorange | -0.147414 | 0.472363 |
| FASTK | green | -0.296529 | 0.141301 | ARHGAP27 | grey60 | -0.403237029 | 0.037012713 | ADAR | darkorange | -0.145839 | 0.477159 |
| COA5 | green | -0.295699 | 0.142471 | DZIP1L | grey60 | -0.397678511 | 0.039954465 | USP20 | darkorange | -0.141749 | 0.489732 |
| TOMM22 | green | 0.291255 | 0.148855 | FBXL6 | grey60 | -0.383426098 | 0.048352104 | FIP1L1 | darkorange | -0.114238 | 0.578433 |
| ISG15 | green | -0.289193 | 0.151885 | SUDS3 | grey60 | -0.375624649 | 0.053504821 | CTC1 | darkorange | -0.101349 | 0.622271 |
| RPS27A | green | -0.286044 | 0.156596 | NOB1 | grey60 | -0.349957165 | 0.073541169 | CPSF6 | darkorange | -0.096463 | 0.639233 |
| DDX52 | green | -0.285683 | 0.157142 | DEXI | grey60 | -0.338869026 | 0.083795504 | SMG9 | darkorange | -0.091894 | 0.655258 |
| CDC26 | green | -0.284557 | 0.158856 | CCDC96 | grey60 | -0.307459785 | 0.118742045 | ANAPC2 | darkorange | -0.090921 | 0.658688 |
| PSMG3 | green | -0.282802 | 0.161555 | GNG11 | grey60 | -0.280184043 | 0.156914355 | CCNB1IP1 | darkorange | -0.07186 | 0.727206 |

**Table 6. Gene module membership and correlation with growth rate (48/72h), with adjusted p values for each module**

| | | | | | | | | | | | |
|---|---|---|---|---|---|---|---|---|---|---|---|
| HP1BP3 | green | -0.282708 | 0.161701 | S100A4 | grey60 | -0.280173239 | 0.156931021 | CCT6P3 | darkorange | -0.043228 | 0.83392 |
| HSPA8 | green | 0.282339 | 0.162273 | ORAI3 | grey60 | -0.275036143 | 0.164999027 | KANSL3 | darkorange | -0.024796 | 0.904296 |
| PIAS3 | green | 0.280665 | 0.164884 | SNORA24 | grey60 | -0.25685337 | 0.195899645 | SLC43A2 | darkorange | -0.021771 | 0.915931 |
| PLAA | green | -0.27951 | 0.166703 | SNORA2B | grey60 | -0.215820536 | 0.279618432 | C22orf32 | darkorange | -0.017185 | 0.933595 |
| SMA5 | green | -0.278467 | 0.168356 | CHST14 | grey60 | -0.186936817 | 0.350481517 | RPL22 | darkorange | -0.016681 | 0.935537 |
| TMEM181 | green | -0.274751 | 0.174345 | SNORA1 | grey60 | -0.157234813 | 0.433482367 | VPS72 | darkorange | -0.000919 | 0.996443 |
| SCARNA7 | green | 0.271994 | 0.178884 | CPOX | grey60 | -0.112851887 | 0.575176904 | ITPRIPL2 | darkorange | 0.013021 | 0.949663 |
| VIMP | green | -0.271495 | 0.179715 | HCG26 | grey60 | -0.067205108 | 0.739088417 | RPL29 | darkorange | 0.015922 | 0.938467 |
| IFNA1 | green | 0.270833 | 0.18082 | ACBD4 | grey60 | -0.057507614 | 0.775709747 | FAIM3 | darkorange | 0.03935 | 0.848641 |
| HSP90AB1 | green | 0.270297 | 0.181718 | CORO1C | grey60 | -0.051597802 | 0.798264748 | SCAMP2 | darkorange | 0.045955 | 0.823597 |
| SF3A3 | green | -0.268719 | 0.184382 | FLII | grey60 | -0.018742076 | 0.926073279 | MRPS18B | darkorange | 0.091376 | 0.657082 |
| MFSD10 | green | 0.26601 | 0.189016 | ALG2 | grey60 | -0.013098585 | 0.948298031 | RPL11 | darkorange | 0.101923 | 0.620289 |
| ZDHHC12 | green | -0.265508 | 0.189884 | PLA2G15 | grey60 | -0.010978192 | 0.956659043 | SEMA6B | darkorange | 0.103252 | 0.615713 |
| FAM104A | green | 0.263227 | 0.193861 | DOCK2 | grey60 | -0.00158455 | 0.993741504 | SRP14 | darkorange | 0.104393 | 0.611796 |
| SLC1A3 | green | -0.261975 | 0.196066 | FUBP3 | grey60 | 0.020163335 | 0.920484069 | RPA1 | darkorange | 0.116037 | 0.572422 |
| CTSO | green | -0.258728 | 0.201871 | SLC12A6 | grey60 | 0.027077808 | 0.893349662 | ANKFY1 | darkorange | 0.121056 | 0.555806 |
| CEP76 | green | 0.258716 | 0.201892 | SLC35A2 | grey60 | 0.027959593 | 0.889897055 | RPL27 | darkorange | 0.125062 | 0.542705 |
| THAP8 | green | -0.255115 | 0.208464 | UBIAD1 | grey60 | 0.034277771 | 0.865219331 | GNPNAT1 | darkorange | 0.131105 | 0.523219 |
| HIST1H2BM | green | -0.253855 | 0.210799 | AP3M1 | grey60 | 0.034284891 | 0.865191587 | EMG1 | darkorange | 0.133569 | 0.515369 |
| CCDC66 | green | -0.253189 | 0.21204 | ABHD2 | grey60 | 0.036676916 | 0.855879643 | NDUFB5 | darkorange | 0.133952 | 0.514154 |
| NACC2 | green | 0.253177 | 0.212062 | SYNRG | grey60 | 0.048558717 | 0.809926578 | PEX13 | darkorange | 0.134923 | 0.511083 |
| PDE6D | green | -0.251794 | 0.214653 | PPP1CA | grey60 | 0.053143987 | 0.79234761 | PFDN5 | darkorange | 0.135419 | 0.509515 |
| GNG10 | green | -0.251326 | 0.215536 | RAB4B | grey60 | 0.053918617 | 0.789387349 | ALKBH1 | darkorange | 0.136533 | 0.506006 |
| ZNF702P | green | -0.250367 | 0.21735 | LEPROT | grey60 | 0.061783407 | 0.759499925 | UBE2Z | darkorange | 0.139295 | 0.497354 |
| HDAC1 | green | -0.249178 | 0.219615 | CDIPT | grey60 | 0.064983124 | 0.747433718 | RPLP2 | darkorange | 0.140385 | 0.493963 |
| TYW1B | green | 0.24847 | 0.220971 | DOCK5 | grey60 | 0.065937193 | 0.743846973 | MRPL24 | darkorange | 0.141238 | 0.491314 |
| SNORA67 | green | 0.248448 | 0.221014 | ARHGEF2 | grey60 | 0.069209629 | 0.731584501 | CBR4 | darkorange | 0.151271 | 0.46072 |
| HNRNPH1 | green | -0.248094 | 0.221694 | AKT2 | grey60 | 0.074542086 | 0.711741082 | ACTR8 | darkorange | 0.155908 | 0.446926 |
| IFNA13 | green | 0.247972 | 0.221928 | DCAF16 | grey60 | 0.08310937 | 0.680245087 | KIAA0664 | darkorange | 0.160603 | 0.433188 |
| FXYD5 | green | 0.247703 | 0.222447 | CTCF | grey60 | 0.090947231 | 0.651880721 | TRIP6 | darkorange | 0.173175 | 0.397545 |
| FAM110A | green | -0.247685 | 0.22248 | SIPA1 | grey60 | 0.097852849 | 0.627274311 | RPF1 | darkorange | 0.179777 | 0.379514 |
| ZBTB2 | green | -0.247566 | 0.222711 | SEC16A | grey60 | 0.101220285 | 0.615412934 | ZNF174 | darkorange | 0.199692 | 0.328047 |
| SMARCA5 | green | -0.246537 | 0.224701 | RECQL5 | grey60 | 0.101475956 | 0.614516154 | KPNA6 | darkorange | 0.249052 | 0.219856 |
| CNOT6 | green | -0.245855 | 0.226028 | UBE2J1 | grey60 | 0.119704649 | 0.552033253 | USP49 | darkorange | 0.254548 | 0.209512 |
| PARVG | green | 0.243943 | 0.229773 | KIAA0319L | grey60 | 0.136002288 | 0.498785119 | SLC7A6OS | darkorange | 0.254862 | 0.208932 |
| ZNF490 | green | 0.242348 | 0.23293 | UMPS | grey60 | 0.136549471 | 0.497043003 | TMEM147 | darkorange | 0.283152 | 0.161014 |
| SOCS4 | green | -0.242167 | 0.23329 | DSCR3 | grey60 | 0.14317307 | 0.476198706 | RABL5 | darkorange | 0.28334 | 0.160724 |
| HIST1H2AE | green | -0.240173 | 0.237279 | CHD4 | grey60 | 0.147308654 | 0.463416145 | CASK | darkorange | 0.326556 | 0.103478 |
| LOC146880 | green | 0.240166 | 0.237293 | HMGXB4 | grey60 | 0.148879508 | 0.458608323 | IFT43 | darkorange | 0.357333 | 0.073111 |
| FLJ42627 | green | 0.240106 | 0.237415 | RPRD2 | grey60 | 0.151628136 | 0.450259294 | MIIP | darkorange | 0.371726 | 0.061503 |
| SNORA55 | green | 0.239783 | 0.238065 | SLC35E1 | grey60 | 0.156679944 | 0.435127411 | SHFM1 | darkorange | 0.380268 | 0.055323 |
| AES | green | -0.238653 | 0.240351 | SPAST | grey60 | 0.157446498 | 0.432855663 | CD3D | darkred | -0.423735 | 0.030992 |
| MAPK14 | green | -0.237316 | 0.243074 | TXNDC11 | grey60 | 0.164335615 | 0.412730672 | ANXA6 | darkred | -0.395475 | 0.045529 |
| ANAPC10 | green | -0.236688 | 0.244361 | CNOT6L | grey60 | 0.165145009 | 0.410400915 | HIST1H2AM | darkred | -0.380059 | 0.055468 |
| OXSM | green | -0.236563 | 0.244619 | SCAMP3 | grey60 | 0.16645305 | 0.406651404 | TBX21 | darkred | -0.357008 | 0.073391 |
| SPG20 | green | -0.235621 | 0.246557 | DNAJC11 | grey60 | 0.169858647 | 0.396979738 | HIST1H2AG | darkred | -0.351219 | 0.07852 |
| NOL10 | green | -0.233544 | 0.250867 | AIMP2 | grey60 | 0.170351632 | 0.395590575 | DPP4 | darkred | -0.339595 | 0.08964 |
| DDOST | green | -0.230974 | 0.256269 | GTPBP2 | grey60 | 0.171579076 | 0.392143806 | HIST1H2AJ | darkred | -0.334123 | 0.095266 |
| OAS1 | green | -0.22763 | 0.26341 | FAM54B | grey60 | 0.177579991 | 0.375540537 | HIST1H2AL | darkred | -0.328633 | 0.101174 |
| STX8 | green | -0.227262 | 0.264204 | FOXJ2 | grey60 | 0.178933174 | 0.371853733 | STAT1 | darkred | -0.326395 | 0.103659 |
| CERS2 | green | 0.226388 | 0.266095 | HCK | grey60 | 0.180938416 | 0.366429279 | ITGA1 | darkred | -0.322319 | 0.108301 |
| RECQL | green | -0.224371 | 0.270493 | ZNF281 | grey60 | 0.18408834 | 0.358002443 | SNHG6 | darkred | -0.311306 | 0.121612 |
| GBP1 | green | -0.223927 | 0.271467 | SIN3A | grey60 | 0.184290105 | 0.357466605 | HIST1H3G | darkred | -0.307658 | 0.126274 |
| PSMD1 | green | -0.223904 | 0.271517 | ZNF836 | grey60 | 0.186746531 | 0.350980981 | CD5 | darkred | -0.30328 | 0.13204 |
| SNORA63 | green | -0.223679 | 0.272012 | ZC3H12C | grey60 | 0.190901274 | 0.340171958 | HIST1H1D | darkred | -0.302971 | 0.132455 |
| ABHD15 | green | -0.22133 | 0.27721 | RFWD3 | grey60 | 0.193442112 | 0.333661519 | GNLY | darkred | -0.295366 | 0.142943 |

**Table 6. Gene module membership and correlation with growth rate (48/72h), with adjusted p values for each module**

| | | | | | | | | | | | |
|---|---|---|---|---|---|---|---|---|---|---|---|
| HCAR3 | green | -0.220341 | 0.279418 | CSNK1D | grey60 | 0.196906691 | 0.32490668 | HIST1H3B | darkred | -0.292593 | 0.146911 |
| ZNF655 | green | -0.219316 | 0.281717 | CNPPD1 | grey60 | 0.197877598 | 0.322478654 | CBWD1 | darkred | -0.284226 | 0.159363 |
| TAF3 | green | -0.217514 | 0.285787 | C22orf13 | grey60 | 0.199592211 | 0.318217991 | LMNB1 | darkred | -0.275437 | 0.17323 |
| RIN1 | green | -0.217348 | 0.286166 | HECA | grey60 | 0.203826958 | 0.307844141 | HIST1H4C | darkred | -0.274209 | 0.175231 |
| ATL1 | green | -0.217282 | 0.286315 | PRPF8 | grey60 | 0.204734981 | 0.305647434 | HIST2H2AB | darkred | -0.266897 | 0.18749 |
| PDE9A | green | -0.217059 | 0.286821 | IQGAP1 | grey60 | 0.206823869 | 0.300631096 | CD226 | darkred | -0.265067 | 0.190648 |
| GLOD4 | green | -0.216626 | 0.287808 | CDC16 | grey60 | 0.20764247 | 0.298679402 | ACAP1 | darkred | -0.263715 | 0.193004 |
| LRRC47 | green | -0.214239 | 0.293284 | MTMR6 | grey60 | 0.208217391 | 0.297313442 | DYSF | darkred | -0.246117 | 0.225518 |
| C5orf25 | green | 0.212822 | 0.296562 | WDR20 | grey60 | 0.208272761 | 0.297182097 | GPR141 | darkred | -0.236912 | 0.243903 |
| C7orf43 | green | -0.210962 | 0.300904 | ACAD8 | grey60 | 0.209152139 | 0.295100948 | HIST1H2AH | darkred | -0.233698 | 0.250546 |
| ACO1 | green | -0.210703 | 0.301512 | STK32C | grey60 | 0.213418522 | 0.285134478 | CD96 | darkred | -0.229601 | 0.259186 |
| CCNL2 | green | -0.209298 | 0.304821 | SUOX | grey60 | 0.215709799 | 0.279871222 | PCNXL2 | darkred | -0.227304 | 0.264114 |
| SNAPIN | green | -0.208409 | 0.306928 | UBXN6 | grey60 | 0.217798838 | 0.275126888 | HIST1H3F | darkred | -0.224127 | 0.271027 |
| HIST1H2BG | green | -0.208386 | 0.306982 | VEZT | grey60 | 0.218670756 | 0.273162042 | HIST1H2BI | darkred | -0.222503 | 0.274606 |
| APEH | green | -0.208081 | 0.307705 | BAP1 | grey60 | 0.221669664 | 0.266472994 | CD3G | darkred | -0.218568 | 0.283401 |
| PPIC | green | -0.20808 | 0.307708 | SGSM2 | grey60 | 0.22201244 | 0.26571523 | GIMAP6 | darkred | -0.216018 | 0.289196 |
| MTA2 | green | -0.206554 | 0.311348 | CEP78 | grey60 | 0.225679914 | 0.257694863 | RRM2 | darkred | -0.211709 | 0.299156 |
| DPEP2 | green | 0.205397 | 0.314128 | F11R | grey60 | 0.226830865 | 0.255210708 | IL2RB | darkred | -0.211446 | 0.299771 |
| CNOT2 | green | -0.204398 | 0.316537 | NDUFA12 | grey60 | 0.231778253 | 0.244710878 | HIST1H3C | darkred | -0.210408 | 0.302204 |
| TOP1MT | green | -0.20404 | 0.317404 | SEC23IP | grey60 | 0.232498125 | 0.243207173 | HIST1H2BH | darkred | -0.209054 | 0.305398 |
| EIF5A | green | -0.202217 | 0.321841 | TMEM87A | grey60 | 0.23335446 | 0.241426372 | MAP4K1 | darkred | -0.20547 | 0.313951 |
| ZW10 | green | -0.202172 | 0.321952 | THOC2 | grey60 | 0.235327438 | 0.237356298 | CA11 | darkred | -0.203047 | 0.319817 |
| TTC33 | green | -0.202008 | 0.322353 | SGPP1 | grey60 | 0.236117937 | 0.235738415 | HIST1H3J | darkred | -0.201401 | 0.323839 |
| HN1L | green | -0.201735 | 0.32302 | SRSF1 | grey60 | 0.240586128 | 0.226731334 | UBD | darkred | -0.195976 | 0.337313 |
| DNAJC21 | green | 0.200939 | 0.324974 | SLC30A9 | grey60 | 0.241488359 | 0.224940947 | CD2 | darkred | -0.193598 | 0.343325 |
| BAZ1B | green | -0.198873 | 0.330076 | FAM20B | grey60 | 0.249050277 | 0.210307561 | FYN | darkred | -0.193044 | 0.344734 |
| SNRPB2 | green | -0.198662 | 0.330601 | SESN2 | grey60 | 0.256621834 | 0.196317125 | VAMP5 | darkred | -0.192553 | 0.345987 |
| HERPUD2 | green | -0.198012 | 0.332218 | TIMM23 | grey60 | 0.265370713 | 0.180964454 | NUSAP1 | darkred | -0.190609 | 0.350971 |
| MITD1 | green | -0.197668 | 0.333076 | THG1L | grey60 | 0.26957469 | 0.173893119 | APOBEC3G | darkred | -0.17916 | 0.381178 |
| XRCC6BP1 | green | -0.197659 | 0.333097 | PTBP1 | grey60 | 0.272997253 | 0.16828099 | HIST1H2AE | darkred | -0.173732 | 0.396006 |
| RPL29P2 | green | 0.196855 | 0.335107 | UBE2G2 | grey60 | 0.27329258 | 0.167802782 | P2RY13 | darkred | -0.168495 | 0.410616 |
| TMEM67 | green | -0.196778 | 0.3353 | PCNXL3 | grey60 | 0.274586027 | 0.16571965 | ANKRD22 | darkred | -0.163988 | 0.423425 |
| CIB1 | green | -0.196015 | 0.337214 | PDE8A | grey60 | 0.280585418 | 0.15629607 | HLA-DOA | darkred | -0.163683 | 0.424298 |
| SLC38A2 | green | 0.195564 | 0.338351 | TADA1 | grey60 | 0.291796897 | 0.139719811 | GBP1P1 | darkred | -0.155249 | 0.448872 |
| HNRNPK | green | -0.193505 | 0.343561 | FLVCR2 | grey60 | 0.291888111 | 0.139590382 | SYNE1 | darkred | -0.155029 | 0.449524 |
| SARS2 | green | 0.192863 | 0.345195 | COG4 | grey60 | 0.315167179 | 0.109314832 | CD69 | darkred | -0.151561 | 0.459853 |
| ZNF780A | green | -0.192804 | 0.345346 | VPS72 | grey60 | 0.316400029 | 0.107859956 | CD7 | darkred | -0.149167 | 0.467052 |
| CDC73 | green | -0.192547 | 0.346 | CLPTM1 | grey60 | 0.318926793 | 0.104923164 | CD3E | darkred | -0.147055 | 0.473455 |
| P2RX4 | green | -0.192334 | 0.346545 | MUL1 | grey60 | 0.339669067 | 0.083021141 | APOL4 | darkred | -0.14635 | 0.475602 |
| AFF1 | green | -0.191961 | 0.347498 | LMAN2 | grey60 | 0.34285193 | 0.079994204 | FAM26F | darkred | -0.139048 | 0.498127 |
| CHMP3 | green | -0.191712 | 0.348138 | NFRKB | grey60 | 0.352209634 | 0.071581203 | CXCL9 | darkred | -0.136774 | 0.505247 |
| RABGGTA | green | -0.191365 | 0.349028 | AK3 | grey60 | 0.353039954 | 0.070868925 | HIST1H2BM | darkred | -0.134903 | 0.511146 |
| RPL36AL | green | -0.189812 | 0.353027 | C3orf38 | grey60 | 0.355583057 | 0.068721258 | NKG7 | darkred | -0.130889 | 0.523909 |
| NUCKS1 | green | -0.189702 | 0.35331 | C19orf43 | grey60 | 0.360510134 | 0.064703435 | RARRES3 | darkred | -0.130214 | 0.526071 |
| HIST1H4J | green | -0.18845 | 0.356555 | ROCK2 | grey60 | 0.373357189 | 0.055080371 | STK17B | darkred | -0.120778 | 0.556721 |
| SMPD1 | green | -0.188333 | 0.356858 | C6orf203 | grey60 | 0.38234159 | 0.049044027 | HIST1H4A | darkred | -0.10414 | 0.612662 |
| ESCO1 | green | -0.187419 | 0.359241 | KAT5 | grey60 | 0.388875677 | 0.044990862 | LTB | darkred | -0.103655 | 0.614329 |
| LOC283070 | green | 0.184226 | 0.367633 | CCDC88A | grey60 | 0.391748287 | 0.04329502 | GALNT11 | darkred | -0.103405 | 0.615185 |
| DTYMK | green | -0.182646 | 0.371826 | RNF139 | grey60 | 0.41100285 | 0.033195226 | SERPING1 | darkred | -0.098956 | 0.630554 |
| KLHL2 | green | -0.182259 | 0.372858 | STAG1 | grey60 | 0.413768964 | 0.031914444 | HAPLN3 | darkred | -0.096137 | 0.64037 |
| RNF167 | green | -0.182132 | 0.373196 | MBTPS1 | grey60 | 0.420854891 | 0.02881403 | GBP1 | darkred | -0.094203 | 0.647139 |
| CNOT4 | green | -0.181944 | 0.373698 | KLHL20 | grey60 | 0.426376496 | 0.026570686 | IL32 | darkred | -0.088913 | 0.665793 |
| PSME4 | green | 0.181488 | 0.374919 | GTF2A1 | grey60 | 0.432420275 | 0.024279613 | CXCL10 | darkred | -0.086928 | 0.672842 |
| ZNF230 | green | -0.181476 | 0.374949 | SF3B3 | grey60 | 0.441733985 | 0.021065577 | CEP170 | darkred | -0.083793 | 0.684034 |
| ATP5E | green | -0.181042 | 0.376112 | ASCC2 | grey60 | 0.460229775 | 0.015710329 | CCL5 | darkred | -0.074588 | 0.717258 |
| HIST1H2BN | green | -0.180187 | 0.37841 | RBMS1 | grey60 | 0.482172714 | 0.010866195 | DLC1 | darkred | -0.06975 | 0.734926 |

**Table 6. Gene module membership and correlation with growth rate (48/72h), with adjusted p values for each module**

| | | | | | | | | | | | |
|---|---|---|---|---|---|---|---|---|---|---|---|
| FAR2 | green | -0.180078 | 0.378704 | TMEM33 | grey60 | 0.48853946 | 0.009720717 | RFX5 | darkred | -0.052405 | 0.799306 |
| IPO13 | green | 0.180059 | 0.378753 | PSMD8 | grey60 | 0.532818224 | 0.004217784 | HIST1H2BG | darkred | -0.050529 | 0.806355 |
| FLJ11235 | green | 0.179594 | 0.380005 | NSMCE1 | grey60 | 0.618926012 | 0.000577973 | APOL1 | darkred | -0.048185 | 0.815179 |
| INO80B | green | -0.179424 | 0.380466 | ANAPC15 | lightcyan | -0.499158337 | 0.008035119 | MACF1 | darkred | -0.045897 | 0.823818 |
| ACTB | green | 0.178537 | 0.382862 | ITCH | lightcyan | -0.460095741 | 0.015744639 | PPA1 | darkred | -0.040243 | 0.845246 |
| DCTN1 | green | -0.177538 | 0.385576 | ZNF320 | lightcyan | -0.444487395 | 0.020185048 | LAP3 | darkred | -0.021348 | 0.917556 |
| GOLGB1 | green | -0.177476 | 0.385743 | SEC61A2 | lightcyan | -0.39633979 | 0.040690038 | PLA1A | darkred | -0.013579 | 0.947508 |
| CDK9 | green | -0.176794 | 0.387601 | MRPL11 | lightcyan | -0.384871178 | 0.047442113 | IL17RB | darkred | 0.001975 | 0.992358 |
| TCF7L2 | green | -0.176312 | 0.388917 | C1QC | lightcyan | -0.363989679 | 0.061977362 | NLRC5 | darkred | 0.007887 | 0.969497 |
| CD27-AS1 | green | 0.176191 | 0.389248 | MT1F | lightcyan | -0.363716283 | 0.062188272 | SERINC2 | darkred | 0.00878 | 0.966046 |
| C2CD2L | green | 0.17604 | 0.389662 | EHD4 | lightcyan | -0.347519572 | 0.075708333 | TPM2 | darkred | 0.009057 | 0.964974 |
| ARL13B | green | -0.175914 | 0.390005 | IKBKAP | lightcyan | -0.346377011 | 0.076740812 | IL27 | darkred | 0.013407 | 0.94817 |
| ITM2B | green | -0.17572 | 0.390538 | ZNF493 | lightcyan | -0.343642642 | 0.079255446 | GBP4 | darkred | 0.016372 | 0.936731 |
| RHOC | green | 0.1757 | 0.390593 | AHSA1 | lightcyan | -0.338199955 | 0.084447303 | GBP5 | darkred | 0.016427 | 0.936517 |
| ZC3H18 | green | -0.175568 | 0.390955 | RPS20 | lightcyan | -0.324085917 | 0.099112379 | RAB11FIP1 | darkred | 0.021742 | 0.916041 |
| FIP1L1 | green | -0.174575 | 0.393682 | HACL1 | lightcyan | -0.31968048 | 0.104058815 | CXCL11 | darkred | 0.028196 | 0.891247 |
| DNAJC4 | green | -0.173685 | 0.396136 | TMEM185E | lightcyan | -0.308807698 | 0.117051662 | SEC23IP | darkred | 0.029294 | 0.887037 |
| PPM1B | green | -0.173527 | 0.396571 | ALDH1A1 | lightcyan | -0.303916293 | 0.123271422 | KLHDC7B | darkred | 0.031536 | 0.878452 |
| ZNF16 | green | -0.172818 | 0.398534 | IST1 | lightcyan | -0.295032759 | 0.135181022 | USE1 | darkred | 0.03738 | 0.85614 |
| PPID | green | -0.172485 | 0.399456 | CTSS | lightcyan | -0.279897768 | 0.157356399 | TNFSF10 | darkred | 0.038801 | 0.850731 |
| ARMC7 | green | -0.171944 | 0.40096 | ZMAT5 | lightcyan | -0.279024081 | 0.158710955 | SLC19A2 | darkred | 0.051727 | 0.801851 |
| DRAP1 | green | -0.171113 | 0.403275 | SREK1IP1 | lightcyan | -0.276949768 | 0.161960077 | CHM | darkred | 0.076296 | 0.711051 |
| SPN | green | -0.170088 | 0.406139 | MZT2B | lightcyan | -0.275979291 | 0.163496258 | TFCP2L1 | darkred | 0.079429 | 0.699718 |
| OC1002165 | green | 0.169253 | 0.408483 | ELP2 | lightcyan | -0.259086922 | 0.191903695 | CDK19 | darkred | 0.083098 | 0.686522 |
| CPM | green | -0.169223 | 0.408566 | ZNF92 | lightcyan | -0.254941259 | 0.199365678 | TUBGCP5 | darkred | 0.083677 | 0.684449 |
| KIAA1279 | green | -0.168394 | 0.410901 | ATPIF1 | lightcyan | -0.251332079 | 0.206022002 | HLA-C | darkred | 0.089532 | 0.663599 |
| ATG16L1 | green | -0.167622 | 0.413079 | COX6C | lightcyan | -0.251078389 | 0.206495505 | MYO6 | darkred | 0.092791 | 0.652101 |
| BRWD3 | green | -0.167067 | 0.414648 | IFIH1 | lightcyan | -0.250817444 | 0.206983322 | ERAP2 | darkred | 0.095484 | 0.642652 |
| ZNF780B | green | -0.167 | 0.414841 | RPL24 | lightcyan | -0.232587105 | 0.243021732 | SESTD1 | darkred | 0.122719 | 0.550351 |
| XPO5 | green | -0.166686 | 0.415729 | BAG1 | lightcyan | -0.230176249 | 0.248079157 | INO80D | darkred | 0.137201 | 0.503907 |
| SUPT3H | green | 0.166389 | 0.416574 | CCDC43 | lightcyan | -0.229389669 | 0.249744062 | SMG1P1 | darkred | 0.147688 | 0.47153 |
| RASL11A | green | 0.166087 | 0.417432 | PDCL | lightcyan | -0.227047371 | 0.254745165 | RMI2 | darkred | 0.158461 | 0.439428 |
| ZNF850 | green | -0.165007 | 0.42051 | C4orf48 | lightcyan | -0.22265804 | 0.264291809 | PNPT1 | darkred | 0.2131 | 0.295919 |
| FAM222B | green | -0.164804 | 0.421088 | APOL6 | lightcyan | -0.218225682 | 0.274163878 | GTF3C4 | darkred | 0.339042 | 0.090197 |
| TRAPPC2P1 | green | -0.163862 | 0.423787 | ZNF496 | lightcyan | -0.214103056 | 0.283555511 | PSME3 | darkturquoise | -0.294852 | 0.143672 |
| MXD1 | green | -0.16326 | 0.425514 | TMEM251 | lightcyan | -0.209832428 | 0.293497267 | COPS6 | darkturquoise | -0.256785 | 0.2054 |
| IKBKB | green | -0.162927 | 0.426472 | CHMP4B | lightcyan | -0.20543934 | 0.303950171 | ZNF655 | darkturquoise | -0.25363 | 0.211217 |
| GNL1 | green | -0.162765 | 0.426937 | STXBP5 | lightcyan | -0.203014954 | 0.30981684 | C17orf79 | darkturquoise | -0.227802 | 0.263039 |
| MAN1B1 | green | -0.162387 | 0.428028 | PARP12 | lightcyan | -0.194750137 | 0.330339567 | COX18 | darkturquoise | -0.221886 | 0.275973 |
| CPSF4 | green | -0.162154 | 0.428699 | ERAP1 | lightcyan | -0.187634456 | 0.348653962 | KDM4B | darkturquoise | -0.214551 | 0.292563 |
| TSPO | green | -0.162092 | 0.428879 | ARHGEF28 | lightcyan | -0.17552269 | 0.381186145 | SDE2 | darkturquoise | -0.206588 | 0.311267 |
| ZNF720 | green | -0.161824 | 0.429652 | ATRAID | lightcyan | -0.156870541 | 0.434561959 | KLHL11 | darkturquoise | -0.199979 | 0.327338 |
| DCTPP1 | green | 0.161704 | 0.429997 | EPSTI1 | lightcyan | -0.154782327 | 0.440778842 | NDUFS7 | darkturquoise | -0.192887 | 0.345133 |
| CAPZB | green | -0.161652 | 0.430148 | ZRSR2 | lightcyan | -0.151333265 | 0.451151082 | POLI | darkturquoise | -0.184717 | 0.366333 |
| HIST1H2BE | green | -0.160811 | 0.432584 | CDK5 | lightcyan | -0.150365527 | 0.454084423 | TMEM183A | darkturquoise | -0.180858 | 0.376606 |
| YARS | green | -0.160107 | 0.434627 | ZMYND15 | lightcyan | -0.149331289 | 0.45723046 | CCT6A | darkturquoise | -0.176708 | 0.387836 |
| HIST1H2AD | green | -0.159441 | 0.436565 | PRPS2 | lightcyan | -0.144312542 | 0.472658759 | APEX1 | darkturquoise | -0.176566 | 0.388224 |
| PPP1R7 | green | -0.158896 | 0.438155 | ALKBH3 | lightcyan | -0.134719386 | 0.502881524 | NECAP2 | darkturquoise | -0.169291 | 0.408375 |
| FAM190B | green | -0.158478 | 0.439377 | C15orf57 | lightcyan | -0.123057914 | 0.540867167 | HAUS7 | darkturquoise | -0.152243 | 0.45781 |
| HSDL2 | green | -0.157838 | 0.441251 | CCDC93 | lightcyan | -0.114456761 | 0.569718394 | NPC1 | darkturquoise | -0.131639 | 0.521512 |
| XYLT2 | green | 0.157792 | 0.441387 | MARK3 | lightcyan | -0.114270003 | 0.570352402 | GALNT6 | darkturquoise | -0.123863 | 0.546611 |
| ACADSB | green | -0.157095 | 0.44343 | LDHB | lightcyan | -0.109103265 | 0.588016242 | TMEM167A | darkturquoise | -0.122319 | 0.55166 |
| CPED1 | green | -0.156851 | 0.444148 | TMEM127 | lightcyan | -0.107171108 | 0.594682205 | SIRPB2 | darkturquoise | -0.107819 | 0.600092 |
| TSEN2 | green | -0.15633 | 0.445683 | PPFIBP2 | lightcyan | -0.099964017 | 0.619827168 | SERPINH1 | darkturquoise | -0.103097 | 0.616244 |
| IQSEC1 | green | 0.156299 | 0.445774 | EPS15 | lightcyan | -0.097476646 | 0.628605141 | PDSS1 | darkturquoise | -0.094054 | 0.647661 |
| MRPL10 | green | -0.15584 | 0.447127 | C11orf75 | lightcyan | -0.0943941 | 0.639552152 | OXLD1 | darkturquoise | -0.091323 | 0.65727 |

**Table 6. Gene module membership and correlation with growth rate (48/72h), with adjusted p values for each module**

| | | | | | | | | | | | |
|---|---|---|---|---|---|---|---|---|---|---|---|
| CXCR4 | green | -0.155418 | 0.448374 | NR2C2 | lightcyan | -0.092754174 | 0.64540635 | PPP2R5D | darkturquoise | -0.088861 | 0.665976 |
| RABL3 | green | -0.154903 | 0.449899 | DKC1 | lightcyan | -0.073287608 | 0.716393404 | SYAP1 | darkturquoise | -0.088528 | 0.667157 |
| IL1R1 | green | -0.154788 | 0.450239 | LOC1005056 | lightcyan | -0.071633872 | 0.722541505 | UBA1 | darkturquoise | -0.086978 | 0.672666 |
| TMEM48 | green | -0.153399 | 0.454362 | GCHFR | lightcyan | -0.056571741 | 0.779270234 | UBA5 | darkturquoise | -0.085509 | 0.677902 |
| KDM4A | green | -0.153316 | 0.454611 | TMEM184E | lightcyan | -0.033553734 | 0.868041483 | PRPS1 | darkturquoise | -0.083226 | 0.686062 |
| FKBP1A | green | -0.153127 | 0.455175 | ABHD4 | lightcyan | -0.03343554 | 0.868502331 | RSBN1L | darkturquoise | -0.079105 | 0.700888 |
| WASH1 | green | -0.152726 | 0.45637 | PPP1R12A | lightcyan | -0.031246585 | 0.877044666 | GTF2H2 | darkturquoise | -0.07302 | 0.722968 |
| PAK2 | green | -0.152486 | 0.457085 | KDM2A | lightcyan | -0.026316997 | 0.896330122 | UBE2R2 | darkturquoise | -0.063962 | 0.756241 |
| ZNF252P | green | 0.152406 | 0.457323 | MED22 | lightcyan | -0.024178143 | 0.904716277 | GPAA1 | darkturquoise | -0.061524 | 0.765272 |
| PAFAH1B2 | green | -0.149986 | 0.464583 | PMS2P3 | lightcyan | -0.016956547 | 0.933099835 | SNX15 | darkturquoise | -0.057407 | 0.780588 |
| ANO6 | green | 0.149559 | 0.465869 | NDUFB3 | lightcyan | -0.015406892 | 0.939202146 | RPS10P7 | darkturquoise | -0.056628 | 0.783496 |
| ARID3A | green | -0.149013 | 0.467517 | RPS6KA3 | lightcyan | -0.006000958 | 0.976301061 | FAM136A | darkturquoise | -0.055593 | 0.787363 |
| UBE2D4 | green | -0.148887 | 0.4679 | RALGAPB | lightcyan | -0.001992976 | 0.992128388 | RPGR | darkturquoise | -0.054982 | 0.789649 |
| C7orf73 | green | -0.148781 | 0.46822 | XIAP | lightcyan | 0.005918022 | 0.9766285 | NGLY1 | darkturquoise | -0.052211 | 0.800033 |
| SNORA81 | green | 0.14865 | 0.468615 | KLF9 | lightcyan | 0.012743687 | 0.949697094 | LUC7L2 | darkturquoise | -0.051246 | 0.803659 |
| MFNG | green | -0.148343 | 0.469544 | ZCCHC8 | lightcyan | 0.030849415 | 0.878596079 | PFDN2 | darkturquoise | -0.05054 | 0.806311 |
| TMEM86A | green | -0.147921 | 0.470823 | PIGL | lightcyan | 0.04403709 | 0.827350179 | SMC1A | darkturquoise | -0.047712 | 0.816963 |
| UEVLD | green | -0.147919 | 0.470829 | SLC16A6 | lightcyan | 0.044424218 | 0.825855173 | IKBKB | darkturquoise | -0.032436 | 0.875008 |
| SNORA38 | green | 0.147171 | 0.4731 | CERS2 | lightcyan | 0.049684953 | 0.805600133 | OAS1 | darkturquoise | -0.021049 | 0.918708 |
| TBC1D22B | green | -0.147136 | 0.473209 | PEMT | lightcyan | 0.056825453 | 0.778304566 | ELOVL1 | darkturquoise | -0.010679 | 0.958708 |
| ATP13A1 | green | -0.146804 | 0.474218 | LATS1 | lightcyan | 0.056887152 | 0.778069779 | SUCNR1 | darkturquoise | -0.010449 | 0.959595 |
| TIMM10 | green | 0.146217 | 0.476008 | NCF2 | lightcyan | 0.056949479 | 0.777832617 | KLHL28 | darkturquoise | 0.00946 | 0.963416 |
| HIST1H3I | green | -0.145698 | 0.47759 | SPOCD1 | lightcyan | 0.061443611 | 0.760784593 | HFE | darkturquoise | 0.013381 | 0.948273 |
| CARM1 | green | 0.143564 | 0.484133 | TADA3 | lightcyan | 0.06255381 | 0.756589574 | ZBTB33 | darkturquoise | 0.025014 | 0.903458 |
| RASSF8 | green | 0.14276 | 0.486609 | SLC25A29 | lightcyan | 0.068761418 | 0.733260314 | CKAP5 | darkturquoise | 0.026641 | 0.897213 |
| MTF2 | green | -0.141157 | 0.491565 | RHOC | lightcyan | 0.069118787 | 0.731924052 | GEMIN7 | darkturquoise | 0.029393 | 0.886659 |
| ANAPC13 | green | -0.140974 | 0.492134 | KBTBD2 | lightcyan | 0.074988356 | 0.710088476 | RPL29P2 | darkturquoise | 0.031671 | 0.877935 |
| EDEM1 | green | -0.140854 | 0.492507 | MPZL3 | lightcyan | 0.090619637 | 0.653057166 | GDAP1 | darkturquoise | 0.050189 | 0.807631 |
| RBM8A | green | -0.140282 | 0.494283 | PYCARD | lightcyan | 0.102695444 | 0.610246201 | RAP2A | darkturquoise | 0.055527 | 0.787608 |
| C17orf49 | green | -0.14024 | 0.494414 | ELOF1 | lightcyan | 0.103633234 | 0.606971029 | ASTE1 | darkturquoise | 0.055771 | 0.786699 |
| LY6G5C | green | 0.140114 | 0.494805 | TM2D2 | lightcyan | 0.105086118 | 0.601911542 | PRPF4B | darkturquoise | 0.056732 | 0.783107 |
| SNHG15 | green | -0.139987 | 0.495201 | FNIP1 | lightcyan | 0.105460689 | 0.600610045 | UBA2 | darkturquoise | 0.069995 | 0.734028 |
| TYMP | green | 0.139972 | 0.495247 | CPNE2 | lightcyan | 0.108492346 | 0.590120413 | RLIM | darkturquoise | 0.083475 | 0.68517 |
| KIAA1737 | green | 0.139465 | 0.496826 | PSMB7 | lightcyan | 0.112764259 | 0.575475613 | IPO7 | darkturquoise | 0.100467 | 0.62532 |
| DDX59 | green | -0.139323 | 0.49727 | SNIP1 | lightcyan | 0.127328594 | 0.526801462 | GOLGA5 | darkturquoise | 0.116089 | 0.57225 |
| RSBN1L | green | -0.138674 | 0.499293 | TRIB1 | lightcyan | 0.129128165 | 0.520927468 | C8B | darkturquoise | 0.117973 | 0.565987 |
| ARNT | green | 0.138185 | 0.500824 | DEDD | lightcyan | 0.131127034 | 0.514440282 | IVNS1ABP | darkturquoise | 0.118318 | 0.564842 |
| PRPF4 | green | -0.138179 | 0.500842 | USP4 | lightcyan | 0.131454815 | 0.513380267 | EML2 | darkturquoise | 0.120856 | 0.556465 |
| IFI6 | green | -0.137497 | 0.502976 | USP31 | lightcyan | 0.135988095 | 0.498830347 | FABP3 | darkturquoise | 0.124876 | 0.543311 |
| CREB1 | green | -0.137468 | 0.50307 | NANP | lightcyan | 0.144090104 | 0.473348728 | C7orf13 | darkturquoise | 0.150732 | 0.46234 |
| COQ10B | green | -0.136508 | 0.506084 | AGPAT1 | lightcyan | 0.147011114 | 0.464329766 | SNAP29 | darkturquoise | 0.151715 | 0.459391 |
| FAM126A | green | -0.13638 | 0.506486 | MAT2B | lightcyan | 0.153153453 | 0.445661223 | NDUFS1 | darkturquoise | 0.158327 | 0.43982 |
| HDAC7 | green | 0.136131 | 0.507271 | PCBP1 | lightcyan | 0.156133631 | 0.436750388 | AIMP1 | darkturquoise | 0.161356 | 0.431003 |
| H2AFX | green | 0.135804 | 0.508301 | SLC16A3 | lightcyan | 0.163656409 | 0.414691357 | SPDYE7P | darkturquoise | 0.163135 | 0.425872 |
| CARD9 | green | -0.13575 | 0.508472 | SHB | lightcyan | 0.170336087 | 0.395634334 | BRI3 | darkturquoise | 0.171543 | 0.402077 |
| TMED1 | green | 0.135506 | 0.50924 | CLASP1 | lightcyan | 0.172054927 | 0.390812189 | SEC62 | darkturquoise | 0.180166 | 0.378466 |
| FUT10 | green | -0.134337 | 0.512934 | PDLIM7 | lightcyan | 0.184370679 | 0.357252751 | MR1 | darkturquoise | 0.190751 | 0.350606 |
| NEK7 | green | -0.134136 | 0.51357 | FAM122B | lightcyan | 0.189620917 | 0.343481382 | STRADB | darkturquoise | 0.200665 | 0.325647 |
| USP27X | green | 0.133784 | 0.514686 | CEP68 | lightcyan | 0.190700997 | 0.340688356 | EIF4A1 | darkturquoise | 0.20713 | 0.309971 |
| DHRSX | green | -0.133725 | 0.514874 | RBM47 | lightcyan | 0.193339168 | 0.333923817 | CRCP | darkturquoise | 0.208969 | 0.3056 |
| MED6 | green | -0.132553 | 0.5186 | ARMC8 | lightcyan | 0.197733267 | 0.322838889 | DTX3L | darkturquoise | 0.215796 | 0.289705 |
| ATP2A2 | green | -0.132185 | 0.519772 | EIF2C2 | lightcyan | 0.202263363 | 0.311649734 | EIF2AK1 | darkturquoise | 0.223681 | 0.272008 |
| IRF2BP1 | green | -0.131659 | 0.521449 | STX5 | lightcyan | 0.204229166 | 0.306869905 | IFNA1 | darkturquoise | 0.276081 | 0.172185 |
| RAD23A | green | 0.130221 | 0.526047 | CCDC144B | lightcyan | 0.204735144 | 0.305647041 | PIP4K2B | darkturquoise | 0.281045 | 0.164288 |
| ZMAT3 | green | -0.12999 | 0.526787 | GSK3A | lightcyan | 0.204964484 | 0.305093765 | BFAR | darkturquoise | 0.29245 | 0.147119 |
| TMTC3 | green | -0.129024 | 0.52989 | ZNF410 | lightcyan | 0.209008564 | 0.295440105 | TRAK1 | darkturquoise | 0.298715 | 0.138252 |

**Table 6. Gene module membership and correlation with growth rate (48/72h), with adjusted p values for each module**

| | | | | | | | | | | | |
|---|---|---|---|---|---|---|---|---|---|---|---|
| BSDC1 | green | -0.128794 | 0.53063 | FEM1B | lightcyan | 0.211600135 | 0.289355868 | ATP1B3 | darkturquoise | 0.313807 | 0.118489 |
| BCL9L | green | 0.128777 | 0.530685 | POMZP3 | lightcyan | 0.217626255 | 0.27551687 | CADPS2 | darkturquoise | 0.319484 | 0.11162 |
| GMPR2 | green | -0.128635 | 0.53114 | SLC3A2 | lightcyan | 0.218135209 | 0.274367815 | MTMR1 | darkturquoise | 0.326825 | 0.103179 |
| GALM | green | -0.127522 | 0.534732 | PNISR | lightcyan | 0.218648731 | 0.273211564 | HADHB | darkturquoise | 0.35981 | 0.071002 |
| MRPS5 | green | 0.126104 | 0.539321 | RPRD1B | lightcyan | 0.221317237 | 0.267253545 | DNAJA1 | darkturquoise | 0.416911 | 0.034103 |
| TMED3 | green | -0.125901 | 0.539979 | CSNK1A1 | lightcyan | 0.226874016 | 0.255117878 | HEG1 | darkturquoise | 0.418418 | 0.033395 |
| AGPAT1 | green | 0.125891 | 0.54001 | SEC61A1 | lightcyan | 0.240937009 | 0.226033916 | MRPL11 | green | -0.492763 | 0.010541 |
| ATP5D | green | 0.12558 | 0.541021 | CLIC1 | lightcyan | 0.259335351 | 0.191462743 | CCZ1B | green | -0.440245 | 0.0244 |
| SEC23IP | green | 0.125502 | 0.541274 | ARNT | lightcyan | 0.261248698 | 0.188090065 | IGSF6 | green | -0.427133 | 0.02953 |
| PFKFB4 | green | -0.125213 | 0.542214 | YIPF6 | lightcyan | 0.286422523 | 0.147499665 | UQCRC2 | green | -0.410283 | 0.037359 |
| EFTUD1 | green | -0.124296 | 0.545197 | LOC1005067: | lightcyan | 0.288932456 | 0.143828564 | CYBB | green | -0.401682 | 0.041949 |
| GDE1 | green | -0.124257 | 0.545324 | TIMP1 | lightcyan | 0.293907103 | 0.13674763 | GOT2 | green | -0.400163 | 0.042803 |
| GMPPB | green | -0.124007 | 0.546142 | PLEKHF2 | lightcyan | 0.294845007 | 0.135441414 | RTFDC1 | green | -0.38437 | 0.052533 |
| DUSP3 | green | 0.12377 | 0.546915 | PAQR5 | lightcyan | 0.305632614 | 0.121062034 | NDUFA11 | green | -0.381445 | 0.054511 |
| PRDX2 | green | -0.123111 | 0.549066 | MAPK9 | lightcyan | 0.306656534 | 0.119757884 | MCCC1 | green | -0.369899 | 0.062892 |
| SMARCC2 | green | -0.122767 | 0.550191 | SRP9 | lightcyan | 0.311323746 | 0.113943882 | ETFB | green | -0.36717 | 0.065011 |
| CYP27A1 | green | 0.122045 | 0.55256 | MAPKAPK5 | lightcyan | 0.319789788 | 0.1039339 | UQCR11 | green | -0.36624 | 0.065746 |
| MPND | green | -0.118153 | 0.565388 | TMEM189 | lightcyan | 0.33908978 | 0.08358129 | MRPS7 | green | -0.3648 | 0.066896 |
| GYPC | green | 0.118102 | 0.56556 | CHST7 | lightcyan | 0.385584208 | 0.046998111 | LYRM1 | green | -0.364182 | 0.067395 |
| AMMECR1 | green | 0.117554 | 0.567376 | TMEM52B | lightcyan | 0.459208766 | 0.015973248 | EIF2B2 | green | -0.363994 | 0.067547 |
| TMC6 | green | -0.117345 | 0.56807 | CCDC92 | lightgreen | -0.517498982 | 0.005700915 | TERC | green | -0.360674 | 0.070278 |
| HECTD4 | green | -0.117003 | 0.569208 | GRSF1 | lightgreen | -0.465403197 | 0.014432399 | R3HCC1 | green | -0.354787 | 0.075328 |
| DYRK3 | green | -0.116513 | 0.570837 | GCSH | lightgreen | -0.411775693 | 0.032833297 | C12orf10 | green | -0.353353 | 0.076599 |
| AMACR | green | -0.115987 | 0.572588 | RIC8B | lightgreen | -0.381706963 | 0.049452524 | ATP6V1G1 | green | -0.349591 | 0.080011 |
| SUMO3 | green | 0.115136 | 0.575427 | NRD1 | lightgreen | -0.379770701 | 0.050715424 | HIST1H1C | green | -0.343064 | 0.086204 |
| GLRX5 | green | -0.113706 | 0.580212 | GID8 | lightgreen | -0.368259383 | 0.058755035 | ATP5F1 | green | -0.342763 | 0.086498 |
| GAS2L1 | green | 0.113056 | 0.582393 | APBB1IP | lightgreen | -0.354024021 | 0.070031834 | TUSC1 | green | -0.342127 | 0.087122 |
| PTOV1 | green | -0.112956 | 0.582731 | ZNF789 | lightgreen | -0.353502828 | 0.070474231 | IK | green | -0.342044 | 0.087203 |
| POLR3A | green | -0.11288 | 0.582987 | MRPL12 | lightgreen | -0.309036349 | 0.116766684 | TRIM21 | green | -0.340746 | 0.088489 |
| CSDE1 | green | -0.11265 | 0.583759 | HNRNPM | lightgreen | -0.307410382 | 0.11880434 | GMPPB | green | -0.338234 | 0.091015 |
| TOM1 | green | -0.112547 | 0.584104 | WDR43 | lightgreen | -0.293026039 | 0.137982972 | IDH3B | green | -0.337188 | 0.092083 |
| CLUAP1 | green | -0.111526 | 0.587541 | CDV3 | lightgreen | -0.282680453 | 0.153096976 | MRPL12 | green | -0.331119 | 0.098466 |
| MLEC | green | -0.111111 | 0.588943 | B3GALT4 | lightgreen | -0.279113251 | 0.158572328 | CCDC22 | green | -0.324966 | 0.105269 |
| BOD1L1 | green | -0.110678 | 0.590404 | AFF1 | lightgreen | -0.278280597 | 0.159870151 | HDGFRP2 | green | -0.324643 | 0.105636 |
| RBM15B | green | 0.110202 | 0.592011 | IQCE | lightgreen | -0.271276981 | 0.171085626 | ECI1 | green | -0.324535 | 0.105759 |
| MMACHC | green | -0.109805 | 0.593355 | RPL5 | lightgreen | -0.268138046 | 0.176287495 | TINF2 | green | -0.324073 | 0.106285 |
| JTB | green | -0.109691 | 0.59374 | BRD7 | lightgreen | -0.254750099 | 0.199714487 | SNORA27 | green | -0.321888 | 0.108802 |
| STK11 | green | -0.10931 | 0.595033 | MPP1 | lightgreen | -0.240637622 | 0.226628893 | ZNF622 | green | -0.320166 | 0.110815 |
| STX3 | green | -0.108302 | 0.598453 | USP22 | lightgreen | -0.23322021 | 0.241704983 | TMEM11 | green | -0.319051 | 0.112133 |
| NDUFB9 | green | -0.108168 | 0.598908 | RNF169 | lightgreen | -0.230646738 | 0.247086793 | FBXO7 | green | -0.318353 | 0.112964 |
| KIAA0226 | green | 0.107784 | 0.600214 | CHMP4C | lightgreen | -0.221752957 | 0.266288733 | QPCT | green | -0.317778 | 0.113652 |
| ALYREF | green | 0.107098 | 0.602549 | NUP214 | lightgreen | -0.214632904 | 0.282337171 | RPL13A | green | -0.315951 | 0.115859 |
| IDUA | green | 0.106884 | 0.603279 | PIAS4 | lightgreen | -0.206668948 | 0.301001352 | ADAL | green | -0.310709 | 0.122366 |
| GOLGA6B | green | 0.106807 | 0.603539 | THOC3 | lightgreen | -0.20490854 | 0.305228671 | TRAPPC4 | green | -0.309213 | 0.124271 |
| IGF2R | green | 0.106523 | 0.604508 | LOC728554 | lightgreen | -0.18403091 | 0.358155051 | COA4 | green | -0.308979 | 0.124571 |
| SMC1A | green | -0.106443 | 0.604784 | HNRNPL | lightgreen | -0.162828595 | 0.417088005 | VTA1 | green | -0.308517 | 0.125165 |
| CSE1L | green | -0.106433 | 0.604817 | SNORA14A | lightgreen | -0.160010328 | 0.425304538 | SLC25A38 | green | -0.30761 | 0.126336 |
| ZNF414 | green | -0.10638 | 0.604997 | TAF1D | lightgreen | -0.149508838 | 0.456689562 | CYB561D2 | green | -0.307515 | 0.126459 |
| FRS2 | green | -0.106154 | 0.60577 | CXorf26 | lightgreen | -0.133791156 | 0.505855799 | TSTD1 | green | -0.305404 | 0.12922 |
| CDC42BPB | green | -0.106109 | 0.605922 | TMEM179E | lightgreen | -0.126352957 | 0.529999254 | HIST1H1E | green | -0.301889 | 0.133912 |
| TMEM107 | green | -0.105441 | 0.608204 | UBE3A | lightgreen | -0.124641648 | 0.535630577 | PSMA3 | green | -0.30026 | 0.136127 |
| PELO | green | -0.105034 | 0.6096 | NUDT14 | lightgreen | -0.111763786 | 0.578890907 | XPA | green | -0.29667 | 0.141102 |
| MED26 | green | -0.104799 | 0.610402 | PUS1 | lightgreen | -0.109345912 | 0.587181407 | TBCC | green | -0.295604 | 0.142605 |
| RBBP4 | green | -0.10444 | 0.611633 | ZHX2 | lightgreen | -0.099509878 | 0.62142609 | UGP2 | green | -0.294559 | 0.144089 |
| CD81 | green | 0.104406 | 0.61175 | PDZD8 | lightgreen | -0.09024165 | 0.654415589 | BAD | green | -0.294065 | 0.144796 |
| PCOLCE | green | -0.102896 | 0.616937 | LCP1 | lightgreen | -0.088912676 | 0.659200204 | C19orf60 | green | -0.293536 | 0.145554 |

**Table 6. Gene module membership and correlation with growth rate (48/72h), with adjusted p values for each module**

| | | | | | | | | | | | |
|---|---|---|---|---|---|---|---|---|---|---|---|
| EXOSC2 | green | -0.102807 | 0.617243 | ZNF823 | lightgreen | -0.082615706 | 0.682046402 | CRAT | green | -0.292308 | 0.147323 |
| ATP10A | green | -0.10273 | 0.617507 | NSFP1 | lightgreen | -0.067800655 | 0.736856518 | SRI | green | -0.28624 | 0.1563 |
| ZNF585A | green | 0.102157 | 0.619481 | SUPT7L | lightgreen | -0.058397738 | 0.772327414 | TRMT61A | green | -0.280757 | 0.164739 |
| ZSCAN5A | green | 0.10196 | 0.62016 | FNDC3B | lightgreen | -0.052446273 | 0.795016341 | ITGB7 | green | -0.277086 | 0.170566 |
| MTA1 | green | 0.101751 | 0.620883 | PAK1IP1 | lightgreen | -0.035011705 | 0.862360247 | NDUFS4 | green | -0.275162 | 0.173677 |
| HNRNPH3 | green | -0.101541 | 0.621606 | CDC23 | lightgreen | -0.026659086 | 0.894989822 | SMUG1 | green | -0.274106 | 0.1754 |
| NAPRT1 | green | 0.101512 | 0.621707 | CD44 | lightgreen | -0.019385081 | 0.923544172 | SIGLEC7 | green | -0.271978 | 0.178911 |
| KIAA0317 | green | -0.101379 | 0.622168 | ZNF3 | lightgreen | -0.01126901 | 0.955512021 | APOL6 | green | -0.270408 | 0.181532 |
| HIST1H2BD | green | -0.100626 | 0.624769 | IFFO1 | lightgreen | -0.011110159 | 0.95613854 | CD209 | green | -0.268766 | 0.184302 |
| IPO8 | green | -0.100044 | 0.626782 | C2CD3 | lightgreen | -0.008659155 | 0.965808437 | TSG101 | green | -0.268735 | 0.184354 |
| CECR2 | green | 0.099765 | 0.627749 | NDST1 | lightgreen | -0.003810623 | 0.984949877 | PCCB | green | -0.26708 | 0.187176 |
| CRYL1 | green | -0.099389 | 0.629052 | COX7B | lightgreen | 0.002961572 | 0.988302961 | PSMB8 | green | -0.266371 | 0.188394 |
| RAD9A | green | -0.098538 | 0.632005 | TMED5 | lightgreen | 0.003606988 | 0.985754055 | PANX1 | green | -0.266121 | 0.188825 |
| MYADM | green | -0.097999 | 0.63388 | AFG3L1P | lightgreen | 0.005804706 | 0.977075893 | NDUFB3 | green | -0.26534 | 0.190174 |
| C5 | green | 0.097776 | 0.634656 | TMEM185A | lightgreen | 0.030350876 | 0.880544079 | SGTB | green | -0.260935 | 0.197914 |
| MRPS26 | green | -0.097756 | 0.634724 | MED14 | lightgreen | 0.03520262 | 0.861616801 | CTPS1 | green | -0.25918 | 0.201056 |
| STIP1 | green | 0.097726 | 0.634831 | NEK11 | lightgreen | 0.038307853 | 0.849541254 | CD48 | green | -0.257821 | 0.203513 |
| NAV1 | green | -0.097485 | 0.635668 | NMNAT1 | lightgreen | 0.060740969 | 0.763443035 | CHMP1A | green | -0.257122 | 0.204784 |
| MIER3 | green | -0.097303 | 0.636303 | ZCCHC7 | lightgreen | 0.061867301 | 0.759182846 | TRIM26 | green | -0.256694 | 0.205565 |
| EFHD2 | green | 0.096663 | 0.638533 | LPCAT3 | lightgreen | 0.062250753 | 0.757734048 | SDHC | green | -0.256179 | 0.206509 |
| FAM160A2 | green | -0.096516 | 0.639046 | ALKBH1 | lightgreen | 0.062839934 | 0.755509506 | COX6C | green | -0.255926 | 0.206973 |
| GTDC1 | green | -0.096268 | 0.639912 | KIF1C | lightgreen | 0.066077499 | 0.743319941 | PSMD6 | green | -0.25446 | 0.209676 |
| REEP5 | green | -0.096209 | 0.64012 | TM9SF4 | lightgreen | 0.084565082 | 0.674943389 | C15orf39 | green | -0.252964 | 0.21246 |
| NRDE2 | green | -0.094046 | 0.647689 | MANEA | lightgreen | 0.098888448 | 0.623616723 | SNORA53 | green | -0.252633 | 0.21308 |
| PHLDA2 | green | 0.093064 | 0.651138 | FBXO42 | lightgreen | 0.103701719 | 0.606732137 | SLC1A3 | green | -0.252594 | 0.213152 |
| AUH | green | -0.092728 | 0.652321 | DEAF1 | lightgreen | 0.113320655 | 0.57358013 | HDGFRP3 | green | -0.250684 | 0.21675 |
| ALOX5AP | green | -0.092347 | 0.65366 | RNF217 | lightgreen | 0.121335465 | 0.546589525 | TLE1 | green | -0.248019 | 0.221838 |
| MLL5 | green | -0.091655 | 0.656099 | RAB7L1 | lightgreen | 0.122018695 | 0.544316319 | LDB1 | green | -0.247309 | 0.223207 |
| SRP54 | green | -0.091108 | 0.658028 | DNAJC8 | lightgreen | 0.126141409 | 0.530693852 | EIF2B4 | green | -0.246333 | 0.225097 |
| RAB3GAP1 | green | -0.090509 | 0.660143 | UACA | lightgreen | 0.126791726 | 0.528559981 | WWOX | green | -0.24594 | 0.225863 |
| CLIC4 | green | -0.089803 | 0.662641 | SUMF2 | lightgreen | 0.1275121 | 0.526201029 | UBB | green | -0.243967 | 0.229727 |
| SCARNA1 | green | 0.089668 | 0.66312 | NCOA6 | lightgreen | 0.130507515 | 0.51644666 | ALKBH7 | green | -0.24352 | 0.230607 |
| FAM76A | green | -0.089079 | 0.665204 | CD82 | lightgreen | 0.130899509 | 0.515176703 | ACSL1 | green | -0.241475 | 0.234669 |
| PCK2 | green | 0.088838 | 0.666059 | WARS2 | lightgreen | 0.133999534 | 0.50518735 | SUPV3L1 | green | -0.240683 | 0.236254 |
| PRR24 | green | 0.087928 | 0.669287 | TTC4 | lightgreen | 0.134913481 | 0.502260692 | MPC2 | green | -0.239997 | 0.237635 |
| SLTM | green | -0.087833 | 0.669627 | ZNF684 | lightgreen | 0.141170396 | 0.48245321 | CIDECP | green | -0.238869 | 0.239914 |
| BTN2A2 | green | -0.087664 | 0.670225 | LYRM2 | lightgreen | 0.142915986 | 0.476999256 | PDCD5 | green | -0.226847 | 0.2651 |
| GSTP1 | green | 0.087337 | 0.671388 | TMA16 | lightgreen | 0.145794266 | 0.468075974 | PIK3CD | green | -0.225606 | 0.267794 |
| BRF2 | green | -0.087132 | 0.672118 | PSMA5 | lightgreen | 0.146238715 | 0.46670587 | CD2BP2 | green | -0.2253 | 0.26846 |
| KIAA2013 | green | 0.086875 | 0.673031 | SLC35B4 | lightgreen | 0.146910533 | 0.46463882 | AP3S2 | green | -0.224489 | 0.270233 |
| NFS1 | green | -0.086555 | 0.674172 | TMEM60 | lightgreen | 0.161080597 | 0.422173824 | ABCD3 | green | -0.222696 | 0.274179 |
| KRTAP19-2 | green | -0.086549 | 0.674192 | PAN3 | lightgreen | 0.162417786 | 0.4182802 | PSMD2 | green | -0.222236 | 0.275197 |
| YTHDC2 | green | -0.086366 | 0.674845 | USP36 | lightgreen | 0.162895354 | 0.416894445 | TMEM158 | green | -0.221889 | 0.275967 |
| WDR70 | green | 0.086213 | 0.675388 | RPL10A | lightgreen | 0.163898026 | 0.41399328 | M6PR | green | -0.221764 | 0.276245 |
| RBM39 | green | -0.085457 | 0.678085 | PHKA2 | lightgreen | 0.16669344 | 0.405964418 | TBC1D10A | green | -0.221539 | 0.276745 |
| SEC14L1P1 | green | 0.08488 | 0.680146 | NDUFS6 | lightgreen | 0.170595039 | 0.394905705 | AGAP5 | green | -0.221085 | 0.277756 |
| RCAN3 | green | -0.084875 | 0.680164 | NRIP1 | lightgreen | 0.181403589 | 0.365177574 | PARP3 | green | -0.22098 | 0.27799 |
| CTSB | green | -0.084524 | 0.681419 | FARSA | lightgreen | 0.183579087 | 0.359357008 | ATP6V1E1 | green | -0.213408 | 0.295203 |
| ZGPAT | green | 0.08375 | 0.684189 | MOAP1 | lightgreen | 0.183644899 | 0.359181785 | SURF4 | green | -0.211854 | 0.298818 |
| LARP1 | green | 0.082829 | 0.687487 | VPS52 | lightgreen | 0.186923522 | 0.350516399 | SNORA49 | green | -0.211802 | 0.298939 |
| NUDT17 | green | 0.08217 | 0.68985 | ILF3 | lightgreen | 0.187494159 | 0.349021032 | TMEM205 | green | -0.211478 | 0.299696 |
| PRNP | green | -0.081645 | 0.691738 | PSMD3 | lightgreen | 0.206310291 | 0.301859621 | MTCH2 | green | -0.210283 | 0.302499 |
| TPRG1L | green | 0.080131 | 0.697188 | EEF2 | lightgreen | 0.222381664 | 0.26490056 | LRBA | green | -0.206938 | 0.310431 |
| CECR1 | green | -0.08 | 0.697657 | NARS | lightgreen | 0.224758273 | 0.259695396 | SARS | green | -0.206555 | 0.311348 |
| GRB10 | green | -0.078965 | 0.701393 | UROD | lightgreen | 0.232207877 | 0.243812723 | TMEM51 | green | -0.204069 | 0.317334 |
| TBC1D30 | green | -0.078849 | 0.701811 | ELP3 | lightgreen | 0.233328544 | 0.241480141 | ZNF789 | green | -0.200711 | 0.325536 |

**Table 6. Gene module membership and correlation with growth rate (48/72h), with adjusted p values for each module**

| | | | | | | | | | | | |
|---|---|---|---|---|---|---|---|---|---|---|---|
| RNF26 | green | -0.07879 | 0.702024 | FAM120B | lightgreen | 0.240004095 | 0.227891367 | S100A6 | green | -0.197528 | 0.333426 |
| ZKSCAN4 | green | -0.078413 | 0.703387 | CASC3 | lightgreen | 0.258983857 | 0.192086836 | QRSL1 | green | -0.197095 | 0.334507 |
| TTLL12 | green | 0.078268 | 0.703911 | PSMB3 | lightgreen | 0.275070869 | 0.164943524 | DYRK3 | green | -0.196268 | 0.33658 |
| STK17A | green | -0.07813 | 0.70441 | NIF3L1 | lightgreen | 0.276788064 | 0.162215328 | SSX1 | green | -0.196037 | 0.337161 |
| RUVBL2 | green | 0.078059 | 0.704666 | C11orf30 | lightgreen | 0.277148802 | 0.161646291 | NDUFA2 | green | -0.195354 | 0.33888 |
| H3F3A | green | -0.077947 | 0.705073 | GRPEL1 | lightgreen | 0.283773885 | 0.151446008 | RAE1 | green | -0.193416 | 0.343787 |
| LIN54 | green | -0.077501 | 0.706685 | TECR | lightgreen | 0.285230798 | 0.149266047 | SCO1 | green | -0.192768 | 0.345437 |
| LCMT2 | green | 0.077352 | 0.707227 | MESDC2 | lightgreen | 0.305817282 | 0.120826058 | DFFB | green | -0.192522 | 0.346065 |
| SLC10A7 | green | -0.076689 | 0.709629 | EWSR1 | lightgreen | 0.308852705 | 0.116995526 | PTMA | green | -0.189541 | 0.353727 |
| SNORA70 | green | 0.076225 | 0.711311 | COQ9 | lightgreen | 0.32954425 | 0.093230526 | RTN4IP1 | green | -0.184266 | 0.367525 |
| CARS2 | green | 0.075745 | 0.713054 | SATB2 | lightgreen | 0.334774443 | 0.087844849 | MRPL38 | green | -0.184266 | 0.367527 |
| PDCD6 | green | -0.075644 | 0.713419 | STT3A | lightgreen | 0.340012892 | 0.082690027 | ZNF672 | green | -0.183376 | 0.369884 |
| ADSS | green | -0.07465 | 0.717034 | EIF4ENIF1 | lightgreen | 0.346998671 | 0.07617772 | UCHL1 | green | -0.182883 | 0.371196 |
| CXorf56 | green | 0.074332 | 0.718188 | RAB1B | lightgreen | 0.370190699 | 0.057341058 | C19orf24 | green | -0.18126 | 0.375528 |
| PLCL1 | green | 0.07417 | 0.71878 | NOP16 | lightgreen | 0.495761474 | 0.008545309 | TPST2 | green | -0.178765 | 0.382247 |
| OC1005062[illegible] | green | -0.072644 | 0.724342 | KLHL11 | lightyellow | -0.574863828 | 0.001709836 | RHBDF2 | green | -0.178358 | 0.383347 |
| HIF1A | green | -0.072392 | 0.725263 | RAB9A | lightyellow | -0.542968599 | 0.003427665 | PGK1 | green | -0.178149 | 0.383914 |
| CD109 | green | -0.071339 | 0.729111 | TFIP11 | lightyellow | -0.540180559 | 0.00363094 | KMO | green | -0.177915 | 0.384549 |
| SUV39H1 | green | 0.069514 | 0.735794 | XPO5 | lightyellow | -0.485406339 | 0.010271111 | NACC1 | green | -0.177414 | 0.385912 |
| EPHX1 | green | -0.069259 | 0.73673 | ATP2A2 | lightyellow | -0.473860947 | 0.012528914 | MAGED2 | green | -0.177163 | 0.386596 |
| NUBP1 | green | -0.068784 | 0.738472 | KDM4B | lightyellow | -0.467029598 | 0.014048885 | CHST15 | green | -0.17648 | 0.388458 |
| KANSL2 | green | -0.068764 | 0.738546 | PANX1 | lightyellow | -0.432973291 | 0.024078261 | MLX | green | -0.172759 | 0.398696 |
| CIRH1A | green | -0.068554 | 0.739318 | GAB2 | lightyellow | -0.427659886 | 0.026070089 | TUBB6 | green | -0.172577 | 0.399203 |
| MTRNR2L8 | green | 0.068291 | 0.740282 | LETMD1 | lightyellow | -0.411424864 | 0.032997198 | PSMB5 | green | -0.169219 | 0.408578 |
| STARD10 | green | 0.067262 | 0.744069 | COX18 | lightyellow | -0.394821245 | 0.041537408 | PTPMT1 | green | -0.16858 | 0.410375 |
| TMEM14C | green | -0.067041 | 0.74488 | OXNAD1 | lightyellow | -0.389029809 | 0.044898557 | PAK1 | green | -0.167844 | 0.412453 |
| FDX1L | green | 0.06684 | 0.745621 | SRSF3 | lightyellow | -0.373900138 | 0.054699838 | ZNF775 | green | -0.16518 | 0.420017 |
| MBLAC2 | green | -0.066353 | 0.747413 | NECAP2 | lightyellow | -0.365728651 | 0.060648807 | C14orf93 | green | -0.164552 | 0.421811 |
| ZNF621 | green | -0.066313 | 0.747563 | DAZAP2 | lightyellow | -0.365352203 | 0.060934513 | NDUFAB1 | green | -0.164375 | 0.422316 |
| PAPD7 | green | 0.066297 | 0.74762 | AIDA | lightyellow | -0.359881614 | 0.065205598 | IL2RG | green | -0.163536 | 0.424719 |
| NDUFAF3 | green | -0.065821 | 0.749374 | STK17A | lightyellow | -0.353939626 | 0.070103324 | C1R | green | -0.162886 | 0.42659 |
| IFT52 | green | 0.065556 | 0.750354 | PGM2L1 | lightyellow | -0.343056402 | 0.079802665 | LOC729737 | green | -0.15707 | 0.443504 |
| PES1 | green | -0.06475 | 0.75333 | POLI | lightyellow | -0.327150709 | 0.095776543 | SSR1 | green | -0.156948 | 0.443863 |
| PPP1R21 | green | -0.064628 | 0.753778 | RPL36AL | lightyellow | -0.326057683 | 0.096956414 | SLMAP | green | -0.150776 | 0.462208 |
| VHL | green | 0.064605 | 0.753866 | RABL3 | lightyellow | -0.324188034 | 0.098999849 | PNMA1 | green | -0.149759 | 0.465266 |
| NUB1 | green | -0.06435 | 0.754804 | SOCS4 | lightyellow | -0.32392633 | 0.099288432 | XBP1 | green | -0.14939 | 0.466379 |
| SENP2 | green | -0.063686 | 0.757263 | MRPL17 | lightyellow | -0.316328815 | 0.107943601 | MRPL49 | green | -0.141285 | 0.491169 |
| FMO4 | green | 0.06317 | 0.759171 | SRRT | lightyellow | -0.308659794 | 0.117236273 | LIMK2 | green | -0.141073 | 0.491827 |
| WDR59 | green | -0.062944 | 0.760006 | SF3A3 | lightyellow | -0.295347912 | 0.134744754 | ATHL1 | green | -0.139851 | 0.495623 |
| CALU | green | 0.062476 | 0.76174 | UQCRC2 | lightyellow | -0.294292405 | 0.136209922 | NOD2 | green | -0.13648 | 0.506172 |
| EPC1 | green | -0.062407 | 0.761995 | PDE6D | lightyellow | -0.293740248 | 0.136980962 | GSTZ1 | green | -0.135115 | 0.510475 |
| AKR7A2 | green | 0.062074 | 0.763232 | GPN3 | lightyellow | -0.265851863 | 0.180145138 | DDX24 | green | -0.133443 | 0.51577 |
| PFN1 | green | 0.061306 | 0.76608 | CCDC91 | lightyellow | -0.261437026 | 0.187760335 | ATP5SL | green | -0.129589 | 0.528074 |
| PIH1D1 | green | 0.06047 | 0.769184 | APEX1 | lightyellow | -0.253216553 | 0.202527869 | ZNF394 | green | -0.129062 | 0.529769 |
| CHMP5 | green | -0.060392 | 0.769474 | FAM136A | lightyellow | -0.250424532 | 0.207719321 | TSEN34 | green | -0.127105 | 0.536078 |
| MRPL21 | green | -0.060375 | 0.769537 | PCMT1 | lightyellow | -0.244332669 | 0.219358846 | SLC25A26 | green | -0.122755 | 0.550233 |
| TUBA1A | green | 0.059965 | 0.771062 | WDR92 | lightyellow | -0.236808611 | 0.234330844 | SACM1L | green | -0.119482 | 0.560994 |
| BCAT1 | green | 0.05988 | 0.771376 | YME1L1 | lightyellow | -0.233574154 | 0.240970897 | TYSND1 | green | -0.118941 | 0.56278 |
| ANXA2P2 | green | -0.059719 | 0.771977 | CDK9 | lightyellow | -0.229102408 | 0.250353911 | KLHDC8B | green | -0.114199 | 0.578561 |
| SIK2 | green | -0.059245 | 0.773741 | UBL7 | lightyellow | -0.211171309 | 0.290357113 | NEK4 | green | -0.107375 | 0.601606 |
| HSDL1 | green | -0.058968 | 0.774771 | FAM3C | lightyellow | -0.186834438 | 0.350750189 | SNORA64 | green | -0.106319 | 0.605204 |
| SEC13 | green | -0.058669 | 0.775884 | BOD1L1 | lightyellow | -0.186399633 | 0.351892616 | TRIP4 | green | -0.106202 | 0.605605 |
| ITGAE | green | 0.058377 | 0.776973 | GCNT2 | lightyellow | -0.171631523 | 0.391996911 | ZEB2 | green | -0.105952 | 0.606457 |
| HIST1H4K | green | -0.05837 | 0.776997 | CHP1 | lightyellow | -0.134701008 | 0.502940325 | GHRLOS2 | green | -0.101595 | 0.621421 |
| CEP164 | green | -0.058229 | 0.777523 | METTL10 | lightyellow | -0.112589397 | 0.576071889 | RPS26 | green | -0.099579 | 0.628393 |
| AIMP1 | green | 0.058054 | 0.778176 | HSP90B1 | lightyellow | -0.098513051 | 0.624941571 | MOCS3 | green | -0.099426 | 0.628923 |

**Table 6. Gene module membership and correlation with growth rate (48/72h), with adjusted p values for each module**

| | | | | | | | | | | | |
|---|---|---|---|---|---|---|---|---|---|---|---|
| DGKQ | green | -0.058047 | 0.778201 | PARP8 | lightyellow | -0.093299307 | 0.643458028 | MME | green | -0.099101 | 0.630053 |
| PEX10 | green | -0.057492 | 0.780273 | GLOD4 | lightyellow | -0.038021099 | 0.85065503 | ZNF304 | green | -0.098988 | 0.630443 |
| ZMYM3 | green | 0.057455 | 0.780409 | HINFP | lightyellow | 0.001104712 | 0.995636697 | ACADVL | green | -0.098202 | 0.633175 |
| NFIX | green | -0.057338 | 0.780847 | TSEN2 | lightyellow | 0.012397524 | 0.951061866 | HMGB1 | green | -0.098001 | 0.633872 |
| LSM2 | green | 0.056966 | 0.782234 | SIGLEC11 | lightyellow | 0.027149832 | 0.893067581 | ACTR2 | green | -0.091474 | 0.656739 |
| ANKRD10 | green | -0.05693 | 0.782369 | FAM49A | lightyellow | 0.035931457 | 0.858779692 | DNAJB5 | green | -0.086803 | 0.673287 |
| UBR2 | green | 0.056735 | 0.783096 | SHQ1 | lightyellow | 0.06330492 | 0.753755222 | CHAF1A | green | -0.084883 | 0.680134 |
| TLR4 | green | 0.056405 | 0.784328 | LASP1 | lightyellow | 0.10744605 | 0.593731677 | SNX21 | green | -0.083881 | 0.683719 |
| PNPLA6 | green | -0.056259 | 0.784873 | ATP1A1 | lightyellow | 0.107484686 | 0.593598155 | MANF | green | -0.08304 | 0.686731 |
| C11orf35 | green | -0.055127 | 0.789105 | TSC22D3 | lightyellow | 0.132687122 | 0.509404672 | TMEM14E | green | -0.081583 | 0.69196 |
| LUZP1 | green | -0.05505 | 0.789395 | KHNYN | lightyellow | 0.135403439 | 0.500695217 | PPFIBP1 | green | -0.081174 | 0.693431 |
| UNC119 | green | 0.055049 | 0.789396 | ARHGAP35 | lightyellow | 0.155148927 | 0.439683976 | NOP9 | green | -0.080174 | 0.697031 |
| USP30-AS1 | green | 0.05449 | 0.791491 | SUMO3 | lightyellow | 0.172147033 | 0.390554738 | MILR1 | green | -0.080091 | 0.697329 |
| GPR77 | green | 0.054154 | 0.792747 | IRF2BPL | lightyellow | 0.190822623 | 0.340374696 | TMEM203 | green | -0.075308 | 0.714642 |
| UQCRFS1 | green | 0.053887 | 0.793746 | HSP90AB1 | lightyellow | 0.193242886 | 0.334169257 | MPG | green | -0.074807 | 0.71646 |
| MECP2 | green | -0.053845 | 0.793906 | PSPH | lightyellow | 0.198410522 | 0.32115066 | RBPJ | green | -0.071578 | 0.728234 |
| TMEM150E | green | 0.053621 | 0.794745 | CARM1 | lightyellow | 0.207855162 | 0.298173607 | DRAP1 | green | -0.071328 | 0.729149 |
| CLN3 | green | 0.05361 | 0.794785 | LRRFIP2 | lightyellow | 0.208125369 | 0.297531814 | PSAT1 | green | -0.067182 | 0.744361 |
| LSM6 | green | -0.05336 | 0.795724 | ZNF264 | lightyellow | 0.220327402 | 0.269453706 | UNC119B | green | -0.064775 | 0.753235 |
| MTRNR2L1 | green | 0.053062 | 0.79684 | TMEM79 | lightyellow | 0.233550679 | 0.24101954 | FAM214A | green | -0.064521 | 0.754176 |
| DOLK | green | -0.052953 | 0.797249 | MYO1F | lightyellow | 0.240146779 | 0.22760662 | ANXA2P2 | green | -0.063892 | 0.756499 |
| NCAPD2 | green | -0.052823 | 0.797738 | TBRG1 | lightyellow | 0.24906453 | 0.210280605 | BBX | green | -0.062748 | 0.760735 |
| CEACAM3 | green | 0.052315 | 0.799643 | YPEL4 | lightyellow | 0.266029685 | 0.179842992 | CLDN7 | green | -0.061453 | 0.765533 |
| PELI3 | green | 0.052019 | 0.800756 | EEF1D | lightyellow | 0.274068319 | 0.16655123 | ACOT11 | green | -0.050828 | 0.805229 |
| ZNF121 | green | -0.051949 | 0.801015 | BRD9 | lightyellow | 0.280822982 | 0.15593094 | STPG1 | green | -0.040818 | 0.843061 |
| AGRN | green | 0.051171 | 0.803939 | SMARCD2 | lightyellow | 0.28271478 | 0.153044952 | ADAM1A | green | -0.039148 | 0.849408 |
| CLP1 | green | -0.050637 | 0.805946 | SP1 | lightyellow | 0.284366085 | 0.150557178 | PROS1 | green | -0.037598 | 0.855309 |
| IVNS1ABP | green | 0.05056 | 0.806238 | SMIM7 | lightyellow | 0.308997927 | 0.116814535 | ACSF2 | green | -0.037066 | 0.857335 |
| WDR55 | green | -0.050152 | 0.807772 | JTB | lightyellow | 0.319280841 | 0.104516469 | PRKD3 | green | -0.036313 | 0.860205 |
| ERF | green | 0.049955 | 0.808514 | LAMTOR1 | lightyellow | 0.325620293 | 0.0974316 | CAPNS2 | green | -0.035778 | 0.862245 |
| SIGLEC10 | green | -0.049387 | 0.810653 | NGLY1 | lightyellow | 0.336338192 | 0.086281263 | RNF13 | green | -0.033555 | 0.870731 |
| TCTN1 | green | 0.04935 | 0.81079 | TRAK1 | lightyellow | 0.345560767 | 0.077484984 | HNRNPUL1 | green | -0.032324 | 0.875437 |
| C8orf82 | green | 0.049223 | 0.811269 | KIAA2013 | lightyellow | 0.360236299 | 0.064921849 | SRRT | green | -0.026687 | 0.897034 |
| TBC1D9B | green | -0.049147 | 0.811555 | EPS15L1 | lightyellow | 0.363234197 | 0.062561533 | FAM70B | green | -0.026239 | 0.898754 |
| DNAJB12 | green | 0.047983 | 0.815944 | CCT4 | lightyellow | 0.372339508 | 0.055799199 | THAP3 | green | -0.02437 | 0.905933 |
| CHD7 | green | -0.047142 | 0.819116 | PLD3 | lightyellow | 0.380800206 | 0.050040826 | ANO5 | green | -0.022964 | 0.911341 |
| MAPKAPK3 | green | -0.046508 | 0.821508 | EIF4A1 | lightyellow | 0.382913377 | 0.048678258 | PPP1R7 | green | -0.015728 | 0.939212 |
| CHP1 | green | -0.046429 | 0.821808 | SMG5 | lightyellow | 0.387742969 | 0.045673823 | RICTOR | green | -0.015396 | 0.940493 |
| HSP90B1 | green | 0.04601 | 0.823393 | PA2G4 | lightyellow | 0.393459838 | 0.042308952 | MAD1L1 | green | -0.015345 | 0.940692 |
| KIAA0368 | green | -0.045757 | 0.824348 | HADHB | lightyellow | 0.550931376 | 0.002899718 | AP3M2 | green | -0.015076 | 0.941731 |
| SLC39A1 | green | 0.045584 | 0.825003 | XPA | magenta | -0.476684029 | 0.011941988 | SNX8 | green | -0.010629 | 0.958902 |
| FAM3C | green | 0.04547 | 0.825431 | NRBF2 | magenta | -0.413686823 | 0.031951896 | GSDMD | green | -0.009526 | 0.963164 |
| MAP3K3 | green | -0.045123 | 0.826745 | PLA2G12A | magenta | -0.412487725 | 0.032502658 | PPP2R5E | green | -0.007688 | 0.970265 |
| ARF5 | green | 0.044962 | 0.827354 | TET2 | magenta | -0.383654518 | 0.048207356 | ITPKC | green | -0.005765 | 0.977704 |
| TADA3 | green | 0.044944 | 0.827423 | MGLL | magenta | -0.380282816 | 0.050378966 | MED26 | green | -0.004948 | 0.980861 |
| NUS1 | green | -0.044694 | 0.828367 | BCL10 | magenta | -0.376897253 | 0.052636148 | HN1 | green | -0.004311 | 0.983326 |
| ZNF24 | green | -0.044671 | 0.828454 | RPL22L1 | magenta | -0.376390261 | 0.052980883 | MAP3K6 | green | 0.000638 | 0.997534 |
| SURF6 | green | -0.044578 | 0.828807 | SKA2 | magenta | -0.369007911 | 0.058203827 | DOCK8 | green | 0.007966 | 0.969193 |
| FANCD2 | green | -0.043913 | 0.831323 | ZNF720 | magenta | -0.358372567 | 0.066423581 | USP27X | green | 0.008349 | 0.96771 |
| CNIH4 | green | 0.043683 | 0.832197 | BACH1 | magenta | -0.340627355 | 0.082100784 | RBM4B | green | 0.008381 | 0.967588 |
| RPLP1 | green | -0.042915 | 0.835106 | ZNF32 | magenta | -0.339010776 | 0.083657905 | MAML2 | green | 0.009139 | 0.96466 |
| RAB11FIP3 | green | 0.042737 | 0.835779 | TWF1 | magenta | -0.331793541 | 0.090884708 | EFR3A | green | 0.013937 | 0.946124 |
| ATXN7L3B | green | 0.041539 | 0.840325 | ZMAT2 | magenta | -0.331178058 | 0.09152213 | PRKDC | green | 0.015179 | 0.941333 |
| C5orf42 | green | -0.041336 | 0.841095 | PLEKHA3 | magenta | -0.331075994 | 0.091628157 | HIPK3 | green | 0.015383 | 0.940545 |
| DEGS1 | green | -0.040026 | 0.846072 | IGHMBP2 | magenta | -0.329860777 | 0.092897688 | DAAM1 | green | 0.018325 | 0.929202 |
| )C1001299 | green | 0.038537 | 0.851735 | ANKRD52 | magenta | -0.322916524 | 0.100407859 | ZNF17 | green | 0.020072 | 0.922471 |

**Table 6. Gene module membership and correlation with growth rate (48/72h), with adjusted p values for each module**

| | | | | | | | | | | | |
|---|---|---|---|---|---|---|---|---|---|---|---|
| RNLS | green | 0.038371 | 0.852368 | RSPH3 | magenta | -0.319787735 | 0.103936245 | FAM35A | green | 0.0208 | 0.919665 |
| TRMT12 | green | 0.038341 | 0.852482 | ANP32B | magenta | -0.31483769 | 0.109706112 | RSC1A1 | green | 0.021456 | 0.91714 |
| HCST | green | -0.038234 | 0.852889 | TYW1 | magenta | -0.303043028 | 0.12440683 | BHLHE41 | green | 0.024947 | 0.903715 |
| RABGEF1 | green | -0.037358 | 0.856223 | TYW3 | magenta | -0.297575442 | 0.131690257 | RNF217 | green | 0.026812 | 0.896556 |
| RENBP | green | -0.036867 | 0.858095 | RB1CC1 | magenta | -0.297447319 | 0.131864569 | TRAM2 | green | 0.028137 | 0.891472 |
| SNORA43 | green | 0.03656 | 0.859264 | GFM2 | magenta | -0.296900448 | 0.132610473 | GZF1 | green | 0.031669 | 0.877942 |
| BLOC1S4 | green | -0.036484 | 0.859556 | FNTA | magenta | -0.292170496 | 0.139190236 | INPPL1 | green | 0.032573 | 0.874485 |
| EIF1AX | green | -0.036279 | 0.860335 | HIGD1A | magenta | -0.291262389 | 0.14048 | TBRG4 | green | 0.034094 | 0.868672 |
| CAMSAP1 | green | -0.035035 | 0.86508 | HGSNAT | magenta | -0.29118808 | 0.140585921 | ZNF595 | green | 0.035157 | 0.864617 |
| FUBP1 | green | -0.034932 | 0.865474 | PLAC8 | magenta | -0.286553397 | 0.147306601 | ZCCHC11 | green | 0.041015 | 0.842313 |
| HSPA13 | green | 0.034753 | 0.866157 | PPM1B | magenta | -0.286045373 | 0.148057049 | ADPRHL2 | green | 0.044144 | 0.830448 |
| KCTD11 | green | 0.034425 | 0.86741 | MTRF1 | magenta | -0.285750221 | 0.148494303 | DCBLD2 | green | 0.049324 | 0.810887 |
| PNN | green | 0.034395 | 0.867525 | SUMO1P3 | magenta | -0.274308606 | 0.166164899 | MYO5A | green | 0.049542 | 0.810069 |
| RGMB | green | 0.033679 | 0.870257 | RIOK2 | magenta | -0.272848214 | 0.168522685 | PSIMCT-1 | green | 0.050565 | 0.806217 |
| SLMO2 | green | -0.033131 | 0.872354 | EEA1 | magenta | -0.270944001 | 0.171632268 | PIDD | green | 0.05533 | 0.788347 |
| ZNF514 | green | -0.032601 | 0.874378 | ZCRB1 | magenta | -0.26645091 | 0.179128681 | MYO9A | green | 0.057448 | 0.780436 |
| POLR2F | green | 0.03161 | 0.878169 | REL | magenta | -0.265891341 | 0.180078029 | ACAP2 | green | 0.057944 | 0.778585 |
| NGRN | green | -0.030751 | 0.881458 | CMC1 | magenta | -0.265197344 | 0.181260306 | NAAA | green | 0.058473 | 0.776614 |
| CR1 | green | -0.030729 | 0.881539 | SKIV2L | magenta | -0.25710618 | 0.195444505 | CTNNB1 | green | 0.058778 | 0.775478 |
| PPM1F | green | 0.030593 | 0.882061 | CFL2 | magenta | -0.255175366 | 0.198939076 | NFIA | green | 0.064547 | 0.754078 |
| SF3B2 | green | 0.030357 | 0.882966 | ARHGEF6 | magenta | -0.252335006 | 0.204157327 | CNEP1R1 | green | 0.064739 | 0.753368 |
| MRPS16 | green | -0.030265 | 0.883316 | CEP57 | magenta | -0.248601523 | 0.211157456 | AGAP6 | green | 0.065387 | 0.750976 |
| CHD6 | green | -0.030216 | 0.883506 | FERMT3 | magenta | -0.246580285 | 0.215014353 | SC5DL | green | 0.066852 | 0.745575 |
| TSPYL5 | green | -0.029191 | 0.887431 | ANKRD26 | magenta | -0.237877603 | 0.232163315 | TRA2A | green | 0.069867 | 0.734499 |
| SMTN | green | 0.029143 | 0.887616 | CLNS1A | magenta | -0.23780625 | 0.232307575 | KAL1 | green | 0.070817 | 0.731018 |
| OSCP1 | green | 0.028964 | 0.888303 | RPS3A | magenta | -0.237628145 | 0.232667927 | DDRGK1 | green | 0.075853 | 0.712661 |
| NOP56 | green | 0.02858 | 0.889775 | TTC1 | magenta | -0.233196057 | 0.241755131 | TBC1D22A | green | 0.078037 | 0.704747 |
| STRADB | green | 0.028203 | 0.891219 | ZNF93 | magenta | -0.232107743 | 0.244021865 | SLC38A10 | green | 0.078249 | 0.703978 |
| PRPS1 | green | -0.028039 | 0.891848 | RSL1D1 | magenta | -0.23171089 | 0.244851902 | ZNF252P | green | 0.078506 | 0.703052 |
| PGGT1B | green | -0.027911 | 0.892338 | ME1 | magenta | -0.225351315 | 0.258406973 | QSER1 | green | 0.080762 | 0.694915 |
| WRB | green | 0.027221 | 0.894987 | C11orf74 | magenta | -0.222142044 | 0.265429083 | KANSL1-AS1 | green | 0.080902 | 0.694411 |
| ULK3 | green | -0.026961 | 0.895983 | WDR60 | magenta | -0.220528222 | 0.269006391 | NUMBL | green | 0.081664 | 0.691671 |
| XPO7 | green | 0.026782 | 0.896669 | RNF149 | magenta | -0.220014634 | 0.27015133 | OCLM | green | 0.083457 | 0.685236 |
| SUCO | green | -0.026681 | 0.897056 | B4GALT7 | magenta | -0.217719053 | 0.275307134 | PTPDC1 | green | 0.094843 | 0.644895 |
| DNAJB5 | green | 0.026389 | 0.898178 | NISCH | magenta | -0.21422168 | 0.283282457 | ATF7 | green | 0.098252 | 0.633 |
| FEM1C | green | -0.025848 | 0.900257 | PRCC | magenta | -0.21160265 | 0.289350002 | ADAM10 | green | 0.098563 | 0.631919 |
| TRHDE-AS1 | green | -0.025613 | 0.901157 | TRRAP | magenta | -0.211486877 | 0.289620096 | ENTPD5 | green | 0.100191 | 0.626273 |
| STK24 | green | 0.025459 | 0.90175 | VEGFB | magenta | -0.203552442 | 0.308510182 | PCMTD2 | green | 0.102276 | 0.61907 |
| CTPS2 | green | -0.025059 | 0.903286 | CAMSAP2 | magenta | -0.199892972 | 0.317474213 | ETNK1 | green | 0.103472 | 0.614955 |
| GNPDA2 | green | 0.024705 | 0.904648 | N6AMT1 | magenta | -0.199581672 | 0.318244075 | RCC2 | green | 0.105375 | 0.608433 |
| CACNB3 | green | -0.02456 | 0.905204 | SNX6 | magenta | -0.198631882 | 0.320600042 | WDR7 | green | 0.105912 | 0.606596 |
| TRIM4 | green | -0.023599 | 0.908899 | ANP32E | magenta | -0.197934753 | 0.322336068 | GPR155 | green | 0.109477 | 0.594466 |
| CBL | green | 0.0235 | 0.909276 | LGALS8 | magenta | -0.193901245 | 0.332493176 | SCP2 | green | 0.111231 | 0.588538 |
| VPS37B | green | -0.023405 | 0.909642 | ABHD8 | magenta | -0.189338612 | 0.344213663 | ABAT | green | 0.112206 | 0.585252 |
| C6orf1 | green | 0.023119 | 0.910745 | NUDT19 | magenta | -0.186995989 | 0.350326287 | TTLL5 | green | 0.112829 | 0.583157 |
| HIST1H4E | green | -0.022829 | 0.911858 | XRN1 | magenta | -0.186977285 | 0.35037535 | F11R | green | 0.113379 | 0.58131 |
| SS18L2 | green | -0.022661 | 0.912505 | ISCA1 | magenta | -0.185322184 | 0.354733064 | AMMECR1 | green | 0.113831 | 0.579795 |
| SLC41A2 | green | -0.022623 | 0.912653 | IQCG | magenta | -0.185183401 | 0.35509992 | DPY19L3 | green | 0.11435 | 0.578056 |
| SLC2A13 | green | -0.0217 | 0.916202 | ZNF76 | magenta | -0.180130608 | 0.368608914 | NOP56 | green | 0.11453 | 0.577452 |
| RALB | green | -0.021393 | 0.917382 | KIF1B | magenta | -0.178764179 | 0.372313012 | SIRPB1 | green | 0.116281 | 0.571608 |
| NT5DC3 | green | -0.020782 | 0.919734 | ZNF436 | magenta | -0.178513861 | 0.37299391 | BTRC | green | 0.118882 | 0.562978 |
| LSM3 | green | 0.019895 | 0.923152 | NCOA2 | magenta | -0.178287708 | 0.373609697 | S100A1 | green | 0.119381 | 0.561325 |
| PRELID1 | green | 0.01936 | 0.925212 | SH3TC1 | magenta | -0.16818348 | 0.401720715 | PFKFB2 | green | 0.122539 | 0.550939 |
| RND3 | green | 0.019288 | 0.925491 | CYTH1 | magenta | -0.167438897 | 0.40383819 | DPY19L1 | green | 0.122798 | 0.550091 |
| RPS19 | green | -0.018868 | 0.927106 | CNTLN | magenta | -0.167333966 | 0.404137101 | LZTFL1 | green | 0.124272 | 0.545275 |
| EMC10 | green | -0.018246 | 0.929505 | NFYB | magenta | -0.163374295 | 0.41550726 | TPT1-AS1 | green | 0.124344 | 0.545043 |

**Table 6. Gene module membership and correlation with growth rate (48/72h), with adjusted p values for each module**

| | | | | | | | | | | | |
|---|---|---|---|---|---|---|---|---|---|---|---|
| NTAN1 | green | 0.018202 | 0.929675 | ZNF609 | magenta | -0.160619763 | 0.423520281 | PVRL2 | green | 0.127285 | 0.535497 |
| BCAR3 | green | -0.018158 | 0.929845 | RAD50 | magenta | -0.160419459 | 0.424106261 | CCNDBP1 | green | 0.127871 | 0.533604 |
| GTF3A | green | 0.017676 | 0.9317 | KDM3A | magenta | -0.158104284 | 0.430911407 | SRCAP | green | 0.13065 | 0.524673 |
| NSMAF | green | -0.017344 | 0.932981 | ITGB3BP | magenta | -0.157458721 | 0.432819492 | IMPDH2 | green | 0.131433 | 0.52217 |
| ECI2 | green | 0.017294 | 0.933175 | GLYR1 | magenta | -0.152249944 | 0.448381814 | IQGAP1 | green | 0.1326 | 0.518449 |
| TBL1X | green | 0.017213 | 0.933486 | CCDC117 | magenta | -0.151923454 | 0.449367093 | DDX60 | green | 0.134488 | 0.512457 |
| ZNF563 | green | 0.016723 | 0.935377 | PYURF | magenta | -0.150374032 | 0.454058602 | MSH2 | green | 0.137182 | 0.503966 |
| MARS | green | -0.015724 | 0.939229 | RABEP1 | magenta | -0.150008455 | 0.455169301 | SLC10A7 | green | 0.137781 | 0.502086 |
| HSPA7 | green | -0.014645 | 0.943394 | EAPP | magenta | -0.148822327 | 0.458782872 | S100A13 | green | 0.141767 | 0.489677 |
| CEP170 | green | 0.013351 | 0.948386 | DNAJB4 | magenta | -0.145931354 | 0.467653152 | RYK | green | 0.142333 | 0.487927 |
| ABRACL | green | 0.012786 | 0.950568 | ZC3H6 | magenta | -0.145809872 | 0.468027833 | PRMT3 | green | 0.153735 | 0.453363 |
| C9orf89 | green | -0.012621 | 0.951207 | ARMC1 | magenta | -0.144346809 | 0.472552514 | LOC148696 | green | 0.156291 | 0.445798 |
| SNORA7B | green | -0.012461 | 0.951826 | PTCD2 | magenta | -0.14402556 | 0.473549031 | PANK3 | green | 0.156402 | 0.445469 |
| SLC41A3 | green | 0.012403 | 0.952049 | DNAJC16 | magenta | -0.138017495 | 0.492384197 | FBXW7 | green | 0.156569 | 0.444977 |
| GRAMD1B | green | 0.012307 | 0.95242 | RUFY1 | magenta | -0.135501929 | 0.500380822 | NPEPPS | green | 0.156695 | 0.444607 |
| TBL1XR1 | green | -0.012289 | 0.952489 | COMMD8 | magenta | -0.1352345 | 0.501234721 | H2AFV | green | 0.157561 | 0.442063 |
| PIGA | green | -0.012273 | 0.95255 | SYNJ2BP | magenta | -0.133228617 | 0.507662531 | PPFIA1 | green | 0.157732 | 0.441561 |
| EXOC6B | green | 0.012153 | 0.953013 | INO80C | magenta | -0.132844035 | 0.508899534 | PHLDA1 | green | 0.158263 | 0.440006 |
| GOLT1B | green | 0.012136 | 0.953079 | ARRDC1 | magenta | -0.132676361 | 0.509439323 | R3HDM1 | green | 0.159057 | 0.437685 |
| PI4K2A | green | 0.011981 | 0.953677 | TEX261 | magenta | -0.132193294 | 0.510996011 | HEXIM1 | green | 0.160426 | 0.4337 |
| PREB | green | 0.011447 | 0.955741 | SSFA2 | magenta | -0.131372268 | 0.513647117 | KIAA1704 | green | 0.160483 | 0.433536 |
| RFX2 | green | 0.011213 | 0.956643 | CEP152 | magenta | -0.131274805 | 0.513962272 | EIF2C4 | green | 0.161418 | 0.430824 |
| PSPH | green | -0.010989 | 0.957509 | ZNF574 | magenta | -0.130028149 | 0.518001749 | ALDH18A1 | green | 0.161862 | 0.429542 |
| ZBTB40 | green | -0.010517 | 0.959333 | METAP2 | magenta | -0.125512187 | 0.532762402 | FAM20B | green | 0.163549 | 0.424682 |
| WDR18 | green | 0.010517 | 0.959334 | ATP7A | magenta | -0.122632858 | 0.542276678 | ZNF641 | green | 0.165684 | 0.418579 |
| STOM | green | -0.009594 | 0.962898 | MBIP | magenta | -0.122331354 | 0.543277529 | SYNCRIP | green | 0.16597 | 0.417765 |
| UBA1 | green | 0.009068 | 0.964934 | RBBP4 | magenta | -0.118864988 | 0.554845782 | AFF4 | green | 0.170237 | 0.405721 |
| ABHD2 | green | -0.00901 | 0.965155 | PSMA4 | magenta | -0.118521213 | 0.555999186 | LOC401093 | green | 0.170411 | 0.405236 |
| C20orf112 | green | 0.008784 | 0.966031 | RSRC1 | magenta | -0.115873653 | 0.564918625 | LOC1001290 | green | 0.170803 | 0.404141 |
| ZNF670 | green | 0.00878 | 0.966044 | ASUN | magenta | -0.115300844 | 0.566856834 | HECTD1 | green | 0.173716 | 0.396049 |
| PXMP4 | green | 0.008478 | 0.967211 | PRPF40A | magenta | -0.113844842 | 0.571796921 | FRRS1 | green | 0.178866 | 0.381971 |
| PPP1R13B | green | 0.007844 | 0.969666 | CACYBP | magenta | -0.109736534 | 0.585838538 | LY6G5B | green | 0.182138 | 0.373179 |
| FBXW5 | green | 0.007544 | 0.970824 | ETHE1 | magenta | -0.107755207 | 0.59266364 | RPS15AP10 | green | 0.184281 | 0.367487 |
| TARP | green | 0.006876 | 0.973406 | MIER1 | magenta | -0.105045268 | 0.60205355 | CPEB4 | green | 0.184281 | 0.367486 |
| SPICE1 | green | 0.006712 | 0.97404 | RPF2 | magenta | -0.10171326 | 0.613684284 | ZNF567 | green | 0.18454 | 0.366802 |
| KLLN | green | 0.006449 | 0.975058 | WAPAL | magenta | -0.101310352 | 0.615096956 | DCAF16 | green | 0.185068 | 0.365408 |
| TIFA | green | -0.00643 | 0.975131 | SLC2A6 | magenta | -0.099646293 | 0.620945628 | NAGA | green | 0.185297 | 0.364805 |
| TRIM8 | green | -0.005769 | 0.977686 | FXR1 | magenta | -0.098400702 | 0.625338295 | SGSM2 | green | 0.185539 | 0.364167 |
| RLF | green | 0.004763 | 0.981576 | DNAJB14 | magenta | -0.09554994 | 0.635438632 | GNA12 | green | 0.188131 | 0.357384 |
| RASGRP2 | green | 0.004568 | 0.982331 | SLC25A44 | magenta | -0.094933096 | 0.637632616 | GBE1 | green | 0.188853 | 0.355509 |
| HSD17B14 | green | 0.004132 | 0.984017 | RNPEP | magenta | -0.094430514 | 0.639422397 | CDR2 | green | 0.189754 | 0.353177 |
| MRI1 | green | -0.003923 | 0.984824 | RPL6 | magenta | -0.092089495 | 0.647785031 | CDC40 | green | 0.192014 | 0.347363 |
| PARP9 | green | -0.003538 | 0.986316 | SNORA75 | magenta | -0.091658114 | 0.649330617 | ZNF542 | green | 0.192319 | 0.346583 |
| PGPEP1 | green | 0.003515 | 0.986403 | RAPH1 | magenta | -0.090753842 | 0.652575117 | RBAK | green | 0.196854 | 0.33511 |
| CNNM3 | green | 0.002581 | 0.990017 | TMEM170A | magenta | -0.085775823 | 0.670545365 | SPICE1 | green | 0.196951 | 0.334868 |
| PEAK1 | green | -0.001435 | 0.994449 | NAP1L4 | magenta | -0.081028085 | 0.687850908 | LATS1 | green | 0.197 | 0.334744 |
| DDX28 | green | -0.001016 | 0.99607 | SREBF2 | magenta | -0.079411184 | 0.693780256 | DHPS | green | 0.199568 | 0.328353 |
| GDI2 | green | 0.000916 | 0.996457 | C1orf131 | magenta | -0.078618132 | 0.696694931 | RNF111 | green | 0.201687 | 0.323138 |
| VPS13A | green | -0.000821 | 0.996825 | TATDN3 | magenta | -0.077633324 | 0.700320217 | STK32C | green | 0.202493 | 0.321167 |
| SAP25 | green | -0.0005 | 0.998064 | DAZAP1 | magenta | -0.072669154 | 0.718690636 | MED31 | green | 0.211735 | 0.299096 |
| IRF5 | green | -0.000481 | 0.998138 | DHX32 | magenta | -0.071949712 | 0.721365991 | DCLRE1C | green | 0.215491 | 0.290404 |
| HPCAL1 | green | 0.000343 | 0.998675 | CSK | magenta | -0.071351374 | 0.723593444 | BACE1 | green | 0.217323 | 0.286222 |
| PLA2G4B | greenyellow | -0.501579 | 0.009039 | ATF7 | magenta | -0.069952338 | 0.728810239 | LOC1004991 | green | 0.217721 | 0.285318 |
| HIST1H1D | greenyellow | -0.487172 | 0.011597 | KATNBL1 | magenta | -0.064376297 | 0.749717715 | RBFOX2 | green | 0.218311 | 0.283983 |
| EIF2B2 | greenyellow | -0.452526 | 0.020274 | CSDA | magenta | -0.061742616 | 0.759654113 | DYNC1LI2 | green | 0.218658 | 0.283199 |
| OCLM | greenyellow | 0.436131 | 0.025925 | CEBPZ | magenta | -0.061430572 | 0.760833903 | SLC41A1 | green | 0.218996 | 0.282436 |

**Table 6. Gene module membership and correlation with growth rate (48/72h), with adjusted p values for each module**

| | | | | | | | | | | | |
|---|---|---|---|---|---|---|---|---|---|---|---|
| ARL6IP4 | greenyellow | -0.381139 | 0.054721 | HECTD3 | magenta | -0.058427477 | 0.772214481 | LYPLA1 | green | 0.222433 | 0.274762 |
| SNORA70B | greenyellow | 0.36704 | 0.065113 | CRYZL1 | magenta | -0.05667053 | 0.778894187 | TRMT12 | green | 0.226528 | 0.265791 |
| EPM2AIP1 | greenyellow | 0.329641 | 0.100069 | MRPS28 | magenta | -0.05400689 | 0.789050193 | CCDC88A | green | 0.227743 | 0.263167 |
| ADM | greenyellow | -0.321464 | 0.109294 | ADD1 | magenta | -0.052557789 | 0.794589646 | ARGFXP2 | green | 0.229218 | 0.260004 |
| PRORSD1P | greenyellow | 0.312853 | 0.119673 | EXOC5 | magenta | -0.051915468 | 0.797048163 | IGIP | green | 0.229813 | 0.258733 |
| EZR | greenyellow | -0.306451 | 0.127846 | ASH1L | magenta | -0.051661364 | 0.798021286 | ZNF808 | green | 0.232691 | 0.252651 |
| POLR3F | greenyellow | -0.304261 | 0.130731 | SLC38A7 | magenta | -0.049197597 | 0.807471637 | ZNF680 | green | 0.23402 | 0.249876 |
| IFIT2 | greenyellow | -0.299655 | 0.136956 | MPHOSPH6 | magenta | -0.046490193 | 0.817886965 | PAQR5 | green | 0.237551 | 0.242595 |
| TIAF1 | greenyellow | 0.296964 | 0.14069 | CBX3 | magenta | -0.046463285 | 0.817990634 | INSR | green | 0.23778 | 0.242127 |
| PRDX4 | greenyellow | -0.293356 | 0.145812 | DDX50 | magenta | -0.045000809 | 0.823629604 | C10orf12 | green | 0.244049 | 0.229566 |
| CDKN1C | greenyellow | -0.291445 | 0.148578 | CTDSPL2 | magenta | -0.044439476 | 0.825796264 | SETD2 | green | 0.246208 | 0.225341 |
| ORC5 | greenyellow | -0.276344 | 0.17176 | RAB8B | magenta | -0.044119266 | 0.827032785 | GALNT1 | green | 0.247491 | 0.222855 |
| APLP2 | greenyellow | -0.274597 | 0.174598 | GFM1 | magenta | -0.041759855 | 0.836156095 | TUBG2 | green | 0.248304 | 0.221291 |
| COPA | greenyellow | -0.271205 | 0.180198 | PALB2 | magenta | -0.041643767 | 0.836605526 | FMR1 | green | 0.249427 | 0.219141 |
| KTI12 | greenyellow | 0.266244 | 0.188612 | CSRP1 | magenta | -0.040763349 | 0.840015653 | RIMKLB | green | 0.249833 | 0.218366 |
| NSUN5 | greenyellow | -0.249871 | 0.218294 | SNRPD1 | magenta | -0.037902097 | 0.851117327 | LOC202181 | green | 0.250707 | 0.216705 |
| LOC90834 | greenyellow | 0.244192 | 0.229284 | OTUD1 | magenta | -0.036817831 | 0.855331649 | TADA1 | green | 0.251373 | 0.215447 |
| LOC148696 | greenyellow | 0.243898 | 0.229863 | SKAP2 | magenta | -0.036629862 | 0.856062642 | MLL2 | green | 0.252281 | 0.213739 |
| PLEKHB1 | greenyellow | -0.239714 | 0.238205 | BRIX1 | magenta | -0.029503985 | 0.883854784 | YY2 | green | 0.252445 | 0.213432 |
| ELOVL1 | greenyellow | -0.237195 | 0.243322 | PAAF1 | magenta | -0.02736055 | 0.892242381 | KCTD6 | green | 0.252815 | 0.212738 |
| FANCL | greenyellow | -0.222626 | 0.274335 | AVL9 | magenta | -0.018135756 | 0.928458735 | ZSWIM6 | green | 0.252839 | 0.212694 |
| RGPD2 | greenyellow | -0.218196 | 0.284242 | DHFR | magenta | -0.017088416 | 0.93258072 | RUVBL1 | green | 0.253777 | 0.210944 |
| SLC19A1 | greenyellow | -0.213105 | 0.295906 | GOLGA4 | magenta | -0.015170706 | 0.940132514 | PRORSD1P | green | 0.254284 | 0.210001 |
| NDN | greenyellow | 0.208792 | 0.306019 | ZNF567 | magenta | -0.014759056 | 0.941754241 | RFX3 | green | 0.255606 | 0.207561 |
| KCNA10 | greenyellow | 0.206691 | 0.311021 | TNRC6A | magenta | -0.012884238 | 0.949143003 | BAGE2 | green | 0.256023 | 0.206794 |
| CAT | greenyellow | -0.205866 | 0.313 | SMURF1 | magenta | -0.00901908 | 0.964388103 | C1orf112 | green | 0.257005 | 0.204998 |
| NXF1 | greenyellow | -0.20233 | 0.321564 | BTBD7 | magenta | -0.007923866 | 0.968710338 | SLC29A1 | green | 0.258789 | 0.201761 |
| TMED10P1 | greenyellow | 0.201665 | 0.323191 | C1orf174 | magenta | -0.00712205 | 0.971875236 | TANC2 | green | 0.259571 | 0.200353 |
| STX6 | greenyellow | -0.199855 | 0.327644 | WIPF2 | magenta | -0.005203471 | 0.979449787 | APOOL | green | 0.26094 | 0.197905 |
| OC1004991 | greenyellow | 0.197818 | 0.3327 | CCSAP | magenta | 0.001781047 | 0.992965416 | TPRG1 | green | 0.263027 | 0.194211 |
| FH | greenyellow | -0.197769 | 0.332823 | PIGQ | magenta | 0.00614237 | 0.975742758 | MTA1 | green | 0.264131 | 0.192278 |
| IFI16 | greenyellow | -0.197689 | 0.333023 | NADKD1 | magenta | 0.00619067 | 0.975552067 | ZFP36L2 | green | 0.265815 | 0.189353 |
| GCSHP3 | greenyellow | -0.196014 | 0.337218 | DCP1A | magenta | 0.011342715 | 0.955221335 | TBC1D3 | green | 0.266966 | 0.187371 |
| LMTK2 | greenyellow | -0.19369 | 0.34309 | SLC16A13 | magenta | 0.011491965 | 0.954632719 | EXOC6B | green | 0.269779 | 0.182589 |
| ATF3 | greenyellow | -0.191493 | 0.348699 | ARHGAP17 | magenta | 0.013084853 | 0.948352161 | ZNF805 | green | 0.270111 | 0.182031 |
| AREG | greenyellow | -0.190562 | 0.351091 | STX18 | magenta | 0.013397102 | 0.947121345 | ZNF432 | green | 0.27104 | 0.180474 |
| TM9SF1 | greenyellow | -0.190388 | 0.351539 | KLHDC5 | magenta | 0.015689259 | 0.938089968 | IFT52 | green | 0.271156 | 0.18028 |
| GVINP1 | greenyellow | 0.188722 | 0.355849 | SNX30 | magenta | 0.016056755 | 0.936642655 | C16orf52 | green | 0.271454 | 0.179782 |
| TYSND1 | greenyellow | -0.185266 | 0.364885 | GALNS | magenta | 0.020012937 | 0.921075352 | ENG | green | 0.272091 | 0.178723 |
| PIK3CB | greenyellow | 0.184795 | 0.366129 | BCAR3 | magenta | 0.020946054 | 0.917407524 | SEC14L1P1 | green | 0.274201 | 0.175244 |
| PRRC2C | greenyellow | -0.1827 | 0.371683 | ZNF274 | magenta | 0.024573951 | 0.903163591 | ARL17A | green | 0.274445 | 0.174845 |
| OC1002722 | greenyellow | 0.180875 | 0.376561 | GCSHP3 | magenta | 0.024617393 | 0.902993196 | MPZL2 | green | 0.276945 | 0.170793 |
| EVL | greenyellow | -0.176696 | 0.387869 | FCF1 | magenta | 0.028093686 | 0.889372188 | NR4A3 | green | 0.279442 | 0.16681 |
| AURKAPS1 | greenyellow | 0.176459 | 0.388516 | NDNL2 | magenta | 0.028381938 | 0.888244062 | ZNF783 | green | 0.280439 | 0.165239 |
| LOC150776 | greenyellow | -0.176066 | 0.389591 | PFDN4 | magenta | 0.028535313 | 0.887643892 | PCNXL4 | green | 0.281206 | 0.164036 |
| LOC84989 | greenyellow | 0.174526 | 0.393817 | LLPH | magenta | 0.030493565 | 0.879986465 | WDPCP | green | 0.281653 | 0.163339 |
| REXO1L2P | greenyellow | -0.171651 | 0.401774 | ZC3H8 | magenta | 0.032290116 | 0.872970581 | BRWD1 | green | 0.285744 | 0.157051 |
| MYO19 | greenyellow | -0.164799 | 0.421104 | CENPN | magenta | 0.041508727 | 0.837128392 | CACNB3 | green | 0.286161 | 0.15642 |
| TSPYL6 | greenyellow | 0.161628 | 0.430219 | PPM1G | magenta | 0.042251447 | 0.83425347 | ANKRD13C | green | 0.286791 | 0.15547 |
| LOC349196 | greenyellow | -0.159816 | 0.435473 | CLCN6 | magenta | 0.04454781 | 0.825378014 | EDC3 | green | 0.288417 | 0.153036 |
| DCAF4L1 | greenyellow | -0.15973 | 0.435725 | PLSCR3 | magenta | 0.046363513 | 0.818375056 | PI4KAP1 | green | 0.289146 | 0.151954 |
| SEPP1 | greenyellow | -0.159007 | 0.43783 | MPHOSPH1 | magenta | 0.048707079 | 0.809356327 | ACAD11 | green | 0.291337 | 0.148734 |
| NDUFA10 | greenyellow | -0.157772 | 0.441445 | DAGLA | magenta | 0.04948181 | 0.806380097 | INPP4A | green | 0.292042 | 0.147709 |
| HPS6 | greenyellow | 0.15594 | 0.446832 | EIF2S2 | magenta | 0.052283262 | 0.79564018 | C18orf25 | green | 0.294607 | 0.144021 |
| FOXJ3 | greenyellow | -0.155112 | 0.44928 | POLR2E | magenta | 0.056654178 | 0.778956428 | VASH1 | green | 0.295915 | 0.142166 |
| NOP58 | greenyellow | -0.152413 | 0.457303 | ATP5A1 | magenta | 0.063053731 | 0.754702753 | ZNF295 | green | 0.29646 | 0.141398 |

**Table 6. Gene module membership and correlation with growth rate (48/72h), with adjusted p values for each module**

| | | | | | | | | | | | |
|---|---|---|---|---|---|---|---|---|---|---|---|
| PFN1P2 | greenyellow | 0.150881 | 0.46189 | RPS6KA2 | magenta | 0.065078383 | 0.747075368 | EIF2C2 | green | 0.298751 | 0.138202 |
| DDX60 | greenyellow | -0.150842 | 0.462007 | NAA20 | magenta | 0.068231661 | 0.735242559 | TMED10P1 | green | 0.300993 | 0.135126 |
| DAG1 | greenyellow | -0.15019 | 0.46397 | C6orf89 | magenta | 0.072216379 | 0.720373976 | TMEM123 | green | 0.302183 | 0.133514 |
| BMPR1A | greenyellow | -0.148326 | 0.469596 | MLLT1 | magenta | 0.072391832 | 0.719721518 | PNISR | green | 0.316098 | 0.11568 |
| NUFIP1 | greenyellow | 0.147649 | 0.47165 | DERL1 | magenta | 0.075894863 | 0.706735483 | ARL10 | green | 0.326283 | 0.103785 |
| ZMYM1 | greenyellow | -0.147292 | 0.472733 | EGLN2 | magenta | 0.08259546 | 0.682120313 | KIAA2013 | green | 0.327272 | 0.10268 |
| VPS33B | greenyellow | -0.144773 | 0.480421 | C19orf59 | magenta | 0.084181574 | 0.676338669 | EPS15L1 | green | 0.327685 | 0.102222 |
| CLDN24 | greenyellow | 0.143971 | 0.482883 | SNRPC | magenta | 0.087594889 | 0.663957505 | TBC1D3F | green | 0.330549 | 0.099083 |
| SFR1 | greenyellow | -0.143508 | 0.484305 | TBCA | magenta | 0.089094385 | 0.658545231 | SPTLC2 | green | 0.337297 | 0.091971 |
| OR2T27 | greenyellow | -0.142434 | 0.487616 | DGKH | magenta | 0.089695506 | 0.656380233 | SAP25 | green | 0.337869 | 0.091387 |
| KCNQ1OT1 | greenyellow | 0.142415 | 0.487674 | ST3GAL2 | magenta | 0.093181887 | 0.643877497 | DHX35 | green | 0.338212 | 0.091037 |
| UPF3A | greenyellow | -0.142219 | 0.48828 | SAFB2 | magenta | 0.094801364 | 0.638101546 | TMEM33 | green | 0.341588 | 0.087653 |
| LONP2 | greenyellow | -0.141925 | 0.48919 | SLC41A3 | magenta | 0.097240186 | 0.62944221 | EPN2 | green | 0.345184 | 0.084154 |
| RARS2 | greenyellow | 0.141814 | 0.48953 | RRP15 | magenta | 0.098935521 | 0.623450675 | KIF13B | green | 0.347689 | 0.081779 |
| EBPL | greenyellow | -0.141579 | 0.490258 | ARGLU1 | magenta | 0.107419674 | 0.593822833 | SNORA70G | green | 0.348331 | 0.081179 |
| E2F6 | greenyellow | 0.140502 | 0.493598 | G3BP2 | magenta | 0.107942984 | 0.592015337 | PIKFYVE | green | 0.348753 | 0.080787 |
| FLJ10038 | greenyellow | 0.13898 | 0.49834 | MFSD6 | magenta | 0.111149175 | 0.58099341 | ADAM15 | green | 0.351748 | 0.078041 |
| FANCF | greenyellow | -0.135468 | 0.50936 | TMEM53 | magenta | 0.120162329 | 0.550502979 | MAP3K2 | green | 0.354344 | 0.075719 |
| MYL12B | greenyellow | -0.135265 | 0.51 | GPATCH1 | magenta | 0.121683807 | 0.545429987 | PPM1D | green | 0.357095 | 0.073316 |
| FAM122A | greenyellow | -0.134296 | 0.513066 | MRPL14 | magenta | 0.127206812 | 0.527200115 | ZMYM5 | green | 0.357847 | 0.07267 |
| ANKRD20A9 | greenyellow | 0.132373 | 0.519171 | VAMP2 | magenta | 0.128429985 | 0.523202638 | PPP1R12A | green | 0.35855 | 0.072069 |
| DUX4L4 | greenyellow | -0.131909 | 0.52065 | SIK3 | magenta | 0.129227274 | 0.520604887 | RGL2 | green | 0.360516 | 0.07041 |
| RPL13AP6 | greenyellow | 0.130145 | 0.526289 | FMNL2 | magenta | 0.132219944 | 0.510910069 | STARD9 | green | 0.365159 | 0.066608 |
| LOC644961 | greenyellow | 0.129918 | 0.527018 | SHISA5 | magenta | 0.132902133 | 0.50871257 | CDK5RAP2 | green | 0.366963 | 0.065175 |
| FAM58A | greenyellow | -0.129515 | 0.528311 | GABARAPL2 | magenta | 0.139352534 | 0.488166587 | SNX25 | green | 0.371281 | 0.061839 |
| MARK4 | greenyellow | -0.128075 | 0.532947 | OSBPL9 | magenta | 0.141082738 | 0.482727925 | NFYA | green | 0.376595 | 0.057919 |
| JOSD2 | greenyellow | -0.127715 | 0.534107 | TMCO7 | magenta | 0.141278154 | 0.48211561 | MTFMT | green | 0.377367 | 0.057365 |
| PRRC2A | greenyellow | -0.12765 | 0.534319 | NDUFB1 | magenta | 0.144655876 | 0.471594814 | SPIN4 | green | 0.378721 | 0.056405 |
| UTP15 | greenyellow | 0.126337 | 0.538563 | FAM78A | magenta | 0.147893423 | 0.461623302 | MTIF2 | green | 0.380131 | 0.055418 |
| OTUD6B | greenyellow | -0.120502 | 0.557631 | NCK1 | magenta | 0.150956339 | 0.452292395 | MBTPS1 | green | 0.389614 | 0.049127 |
| SAP30 | greenyellow | 0.120338 | 0.55817 | SHC1 | magenta | 0.155026732 | 0.440048754 | GAS2L1 | green | 0.393587 | 0.046665 |
| USP17L5 | greenyellow | -0.118478 | 0.564313 | GRIN2D | magenta | 0.155857796 | 0.437571077 | DAPK1 | green | 0.397216 | 0.044501 |
| LOC441454 | greenyellow | 0.117027 | 0.569127 | MAP4K2 | magenta | 0.157670448 | 0.432193187 | LOC646471 | green | 0.402799 | 0.041328 |
| AK4 | greenyellow | 0.114968 | 0.575989 | SMC6 | magenta | 0.158873503 | 0.428643825 | PACRGL | green | 0.406027 | 0.039577 |
| ZNF394 | greenyellow | 0.112212 | 0.58523 | TRAPPC10 | magenta | 0.163920064 | 0.413929641 | ZFYVE1 | green | 0.40746 | 0.038819 |
| TATDN1 | greenyellow | 0.109955 | 0.592846 | HIGD2A | magenta | 0.16512561 | 0.410456668 | FLJ45340 | green | 0.409127 | 0.037951 |
| REC8 | greenyellow | 0.109204 | 0.595392 | WBSCR22 | magenta | 0.16601731 | 0.407898321 | DEM1 | green | 0.410827 | 0.037083 |
| LOC401321 | greenyellow | 0.103999 | 0.613145 | EIF4H | magenta | 0.171423604 | 0.392579437 | RAPGEFL1 | green | 0.41153 | 0.036728 |
| TRIO | greenyellow | 0.103888 | 0.613529 | MRPL23 | magenta | 0.171433976 | 0.392550366 | PIGL | green | 0.413494 | 0.035752 |
| SUPT16H | greenyellow | 0.102987 | 0.616624 | GTF2H5 | magenta | 0.177145171 | 0.376729702 | ITSN2 | green | 0.420996 | 0.032212 |
| ITFG2 | greenyellow | -0.102712 | 0.617569 | GET4 | magenta | 0.190146561 | 0.342120387 | PLAGL1 | green | 0.42278 | 0.031413 |
| BRICD5 | greenyellow | -0.102115 | 0.619626 | POLDIP2 | magenta | 0.201230214 | 0.314180188 | SCMH1 | green | 0.433846 | 0.026805 |
| FUT4 | greenyellow | 0.101249 | 0.622615 | SNRNP200 | magenta | 0.230536011 | 0.247320106 | PRRC2B | green | 0.439585 | 0.02464 |
| SPDYE7P | greenyellow | 0.099303 | 0.629351 | GSK3B | magenta | 0.252729107 | 0.203427766 | ZBED6 | green | 0.442609 | 0.023557 |
| DNAJC30 | greenyellow | -0.098607 | 0.631767 | GPI | magenta | 0.272354575 | 0.169324961 | C10orf118 | green | 0.444585 | 0.022871 |
| CYCSP52 | greenyellow | 0.098372 | 0.632584 | CPSF3L | magenta | 0.284645329 | 0.150139359 | SETDB1 | green | 0.449451 | 0.02125 |
| TAS2R14 | greenyellow | 0.097931 | 0.634116 | GSTP1 | magenta | 0.303812369 | 0.123406142 | DAPK3 | green | 0.45078 | 0.020824 |
| DEPDC1 | greenyellow | 0.096345 | 0.639646 | MSRA | magenta | 0.32652709 | 0.096448382 | SP1 | green | 0.454593 | 0.01964 |
| MGC16025 | greenyellow | -0.095238 | 0.643512 | RBX1 | magenta | 0.328201359 | 0.094652609 | WWC2 | green | 0.466072 | 0.016403 |
| VNN2 | greenyellow | -0.094135 | 0.647378 | TMCO1 | magenta | 0.353371876 | 0.070585722 | PDXDC2P | green | 0.469074 | 0.015633 |
| ATP5L2 | greenyellow | 0.092479 | 0.653197 | ERGIC3 | magenta | 0.386659492 | 0.046334744 | ZNF192 | green | 0.477771 | 0.013569 |
| TMEM203 | greenyellow | -0.092396 | 0.65349 | STK10 | midnightblue | -0.460930244 | 0.015532025 | ZNF767 | green | 0.477834 | 0.013555 |
| DDX58 | greenyellow | -0.092272 | 0.653925 | EXOC8 | midnightblue | -0.384358816 | 0.047763197 | EPC2 | green | 0.478203 | 0.013473 |
| SBDSP1 | greenyellow | -0.089106 | 0.665108 | CCNK | midnightblue | -0.359186717 | 0.065764299 | CDK12 | green | 0.480791 | 0.012907 |
| PPP1R3D | greenyellow | 0.088671 | 0.666651 | SOGA1 | midnightblue | -0.358704089 | 0.066154508 | EIF2C3 | green | 0.484505 | 0.01213 |
| INSIG1 | greenyellow | -0.087349 | 0.671344 | CD276 | midnightblue | -0.357995615 | 0.066730553 | SEC14L1 | green | 0.497881 | 0.009645 |

**Table 6. Gene module membership and correlation with growth rate (48/72h), with adjusted p values for each module**

| | | | | | | | | | | | |
|---|---|---|---|---|---|---|---|---|---|---|---|
| SERTAD1 | greenyellow | -0.086619 | 0.673943 | PLEC | midnightblue | -0.343710594 | 0.079192203 | CCDC127 | greenyellow | -0.42584 | 0.030079 |
| UTP3 | greenyellow | -0.086331 | 0.674969 | TCTA | midnightblue | -0.342474691 | 0.080348501 | APBB1IP | greenyellow | -0.331558 | 0.097994 |
| CHIT1 | greenyellow | 0.081756 | 0.691339 | RAPGEF6 | midnightblue | -0.314889864 | 0.109644084 | PITRM1 | greenyellow | -0.331485 | 0.098072 |
| DSN1 | greenyellow | -0.081483 | 0.692321 | TADA2B | midnightblue | -0.303727455 | 0.123516301 | POP7 | greenyellow | -0.303315 | 0.131993 |
| ZNF829 | greenyellow | 0.080342 | 0.696426 | SHARPIN | midnightblue | -0.283244133 | 0.152244281 | IFT122 | greenyellow | -0.292293 | 0.147345 |
| DNM2 | greenyellow | -0.07941 | 0.699786 | VEGFA | midnightblue | -0.281545845 | 0.154823656 | CEPT1 | greenyellow | -0.271462 | 0.179769 |
| LOC145474 | greenyellow | 0.078304 | 0.703782 | BAHD1 | midnightblue | -0.259979855 | 0.19032204 | JRKL | greenyellow | -0.265089 | 0.190609 |
| SSBP3 | greenyellow | -0.077183 | 0.707837 | RNF115 | midnightblue | -0.25815819 | 0.193558359 | PLA2G12A | greenyellow | -0.262552 | 0.195048 |
| EID2 | greenyellow | 0.0771 | 0.708137 | DDX42 | midnightblue | -0.254840189 | 0.199550048 | SLC9A3R1 | greenyellow | -0.25144 | 0.21532 |
| ACTR3BP2 | greenyellow | -0.076288 | 0.711081 | THUMPD2 | midnightblue | -0.252724211 | 0.203436819 | CHMP7 | greenyellow | -0.246398 | 0.224972 |
| ATG3 | greenyellow | -0.070912 | 0.730672 | SWAP70 | midnightblue | -0.249361946 | 0.209718661 | SLK | greenyellow | -0.230998 | 0.256219 |
| H1F0 | greenyellow | -0.069254 | 0.736748 | CHMP7 | midnightblue | -0.240297173 | 0.227306746 | RAB39A | greenyellow | -0.216705 | 0.287628 |
| LOC254100 | greenyellow | 0.069169 | 0.737059 | DDX6 | midnightblue | -0.240099551 | 0.227700845 | CCDC88C | greenyellow | -0.209717 | 0.303831 |
| OR52K2 | greenyellow | 0.064928 | 0.75267 | MON1B | midnightblue | -0.233215141 | 0.241715507 | RPL18A | greenyellow | -0.204835 | 0.315483 |
| STAG3 | greenyellow | -0.06479 | 0.753181 | PTPRJ | midnightblue | -0.211895675 | 0.288667097 | TTC21B | greenyellow | -0.202063 | 0.322219 |
| CDC27 | greenyellow | -0.063635 | 0.75745 | DDX59 | midnightblue | -0.203011461 | 0.309825343 | MAP4K5 | greenyellow | -0.200223 | 0.326736 |
| ASS1 | greenyellow | -0.063488 | 0.757994 | DHX33 | midnightblue | -0.201544967 | 0.313407935 | RPL30 | greenyellow | -0.196826 | 0.335179 |
| RBM12B-AS | greenyellow | 0.063061 | 0.759574 | CCDC130 | midnightblue | -0.198961985 | 0.319780007 | MMGT1 | greenyellow | -0.196094 | 0.337017 |
| C7orf13 | greenyellow | 0.062539 | 0.761507 | ZNF317 | midnightblue | -0.197792902 | 0.322690017 | ZYG11B | greenyellow | -0.181613 | 0.374584 |
| RRP1B | greenyellow | -0.061185 | 0.766529 | MAML1 | midnightblue | -0.184109276 | 0.357946821 | METTL16 | greenyellow | -0.179531 | 0.380177 |
| TIGD1 | greenyellow | -0.0611 | 0.766846 | CCS | midnightblue | -0.177420081 | 0.375977613 | TRAPPC8 | greenyellow | -0.179385 | 0.380569 |
| HARBI1 | greenyellow | -0.061071 | 0.766953 | TNIP2 | midnightblue | -0.176452243 | 0.378629253 | LOC1001323 | greenyellow | -0.17849 | 0.382991 |
| GHRLOS2 | greenyellow | 0.059641 | 0.772266 | GATAD2B | midnightblue | -0.176131189 | 0.379511244 | PPIP5K2 | greenyellow | -0.177191 | 0.386519 |
| MTMR1 | greenyellow | -0.057511 | 0.780199 | SETD5 | midnightblue | -0.174196797 | 0.384850395 | ZFHX2 | greenyellow | -0.176148 | 0.389367 |
| WDR11 | greenyellow | 0.056134 | 0.785341 | ZBTB7A | midnightblue | -0.172093644 | 0.390703957 | BTBD1 | greenyellow | -0.171751 | 0.401495 |
| CNOT7 | greenyellow | -0.054412 | 0.79178 | NR3C1 | midnightblue | -0.168320545 | 0.401331607 | LEO1 | greenyellow | -0.170399 | 0.405269 |
| UFSP1 | greenyellow | 0.053974 | 0.793422 | YWHAG | midnightblue | -0.163221248 | 0.41595026 | YIPF1 | greenyellow | -0.169987 | 0.406422 |
| ZNF277 | greenyellow | -0.049318 | 0.81091 | SS18L1 | midnightblue | -0.161174595 | 0.42189947 | CHORDC1 | greenyellow | -0.169381 | 0.408122 |
| MST1 | greenyellow | 0.048353 | 0.814549 | CHD2 | midnightblue | -0.145294511 | 0.469619067 | IARS2 | greenyellow | -0.166835 | 0.415308 |
| STAM2 | greenyellow | 0.047918 | 0.816186 | SIPA1L2 | midnightblue | -0.137547715 | 0.493872658 | OVCA2 | greenyellow | -0.165943 | 0.417842 |
| DRD5 | greenyellow | -0.047865 | 0.816387 | ZNF81 | midnightblue | -0.131159419 | 0.514335503 | RPS18 | greenyellow | -0.159543 | 0.436269 |
| TOMM20 | greenyellow | 0.04759 | 0.817425 | GOSR1 | midnightblue | -0.127899847 | 0.524933399 | RSPH3 | greenyellow | -0.155359 | 0.448548 |
| DDX20 | greenyellow | -0.046456 | 0.821706 | XRCC5 | midnightblue | -0.12765525 | 0.525732868 | ANKRD36BP | greenyellow | -0.153555 | 0.4539 |
| NBEA | greenyellow | 0.04567 | 0.824678 | GNE | midnightblue | -0.124480952 | 0.53616082 | EP300 | greenyellow | -0.152998 | 0.455559 |
| FKSG29 | greenyellow | 0.044641 | 0.828569 | CNOT3 | midnightblue | -0.123156879 | 0.540539236 | SLC35F5 | greenyellow | -0.152948 | 0.455707 |
| ZFAND4 | greenyellow | -0.043626 | 0.832413 | TMEM115 | midnightblue | -0.118242351 | 0.556935606 | DNAJC3 | greenyellow | -0.152941 | 0.455728 |
| STYX | greenyellow | 0.04312 | 0.834329 | RCE1 | midnightblue | -0.116535248 | 0.562683717 | LOC729852 | greenyellow | -0.151293 | 0.460654 |
| OC1001283 | greenyellow | 0.041515 | 0.840415 | MAFK | midnightblue | -0.115393277 | 0.566543869 | RUFY2 | greenyellow | -0.148465 | 0.469176 |
| LOC220729 | greenyellow | -0.040893 | 0.842779 | ADAP2 | midnightblue | -0.114150114 | 0.570759571 | ALDH9A1 | greenyellow | -0.146252 | 0.475898 |
| LINC00628 | greenyellow | 0.039585 | 0.847749 | RCC2 | midnightblue | -0.113606221 | 0.572608369 | NUP214 | greenyellow | -0.142149 | 0.488496 |
| LOC606724 | greenyellow | -0.038135 | 0.853265 | SUGP1 | midnightblue | -0.110449028 | 0.583392599 | GDAP2 | greenyellow | -0.142032 | 0.488859 |
| WDR5B | greenyellow | -0.036493 | 0.859521 | HS2ST1 | midnightblue | -0.107854777 | 0.592319836 | RPL21 | greenyellow | -0.139743 | 0.495961 |
| ATP5S | greenyellow | -0.035024 | 0.865124 | MAP4 | midnightblue | -0.098674723 | 0.62437086 | NCOA3 | greenyellow | -0.136711 | 0.505447 |
| ADAL | greenyellow | 0.033125 | 0.872374 | BICD2 | midnightblue | -0.09651415 | 0.632015161 | SGMS2 | greenyellow | -0.13387 | 0.514414 |
| BET1 | greenyellow | 0.032776 | 0.873708 | SNX11 | midnightblue | -0.095758169 | 0.634698685 | SUCO | greenyellow | -0.131472 | 0.522044 |
| NANOG | greenyellow | -0.0324 | 0.875145 | IRGQ | midnightblue | -0.095532287 | 0.635501381 | SKIL | greenyellow | -0.131307 | 0.522571 |
| OR8G2 | greenyellow | 0.030211 | 0.883523 | SRXN1 | midnightblue | -0.09553022 | 0.635508726 | PROSER1 | greenyellow | -0.131007 | 0.523532 |
| DNAAF2 | greenyellow | -0.022487 | 0.913173 | AFF4 | midnightblue | -0.094987004 | 0.637440756 | NSMAF | greenyellow | -0.128839 | 0.530486 |
| JAG1 | greenyellow | -0.020199 | 0.921982 | DTNBP1 | midnightblue | -0.086379719 | 0.668355653 | MTPN | greenyellow | -0.126707 | 0.537368 |
| OR4F21 | greenyellow | 0.018626 | 0.928042 | FADD | midnightblue | -0.084010678 | 0.676960761 | MPHOSPH1 | greenyellow | -0.12618 | 0.539073 |
| OC1005061 | greenyellow | -0.017708 | 0.931579 | HTATSF1 | midnightblue | -0.080974216 | 0.688048162 | TCN2 | greenyellow | -0.12497 | 0.543003 |
| ZNF532 | greenyellow | 0.017296 | 0.933166 | DOCK1 | midnightblue | -0.080966051 | 0.688078063 | STX7 | greenyellow | -0.121759 | 0.553497 |
| RBM22 | greenyellow | 0.015887 | 0.938601 | DUSP18 | midnightblue | -0.08011607 | 0.691193175 | HLA-DPB1 | greenyellow | -0.120844 | 0.556502 |
| KCTD3 | greenyellow | 0.014462 | 0.944098 | NDUFA9 | midnightblue | -0.077757346 | 0.699863315 | TMCO6 | greenyellow | -0.117118 | 0.568826 |
| LOC282997 | greenyellow | 0.014053 | 0.945676 | GTF2F1 | midnightblue | -0.068403963 | 0.734597655 | PLBD1 | greenyellow | -0.116398 | 0.571221 |
| RAB32 | greenyellow | 0.012975 | 0.94984 | NDUFB6 | midnightblue | -0.063410806 | 0.753355905 | TRIP11 | greenyellow | -0.115261 | 0.575009 |

**Table 6. Gene module membership and correlation with growth rate (48/72h), with adjusted p values for each module**

| | | | |
|---|---|---|---|
| FGL2 | greenyellow | -0.012547 | 0.951492 |
| CEMP1 | greenyellow | 0.012221 | 0.95275 |
| PKI55 | greenyellow | 0.010858 | 0.958015 |
| AKTIP | greenyellow | 0.010158 | 0.960719 |
| HUS1B | greenyellow | 0.006756 | 0.97387 |
| LOC727896 | greenyellow | 0.00523 | 0.979771 |
| ZNF442 | greenyellow | 0.002737 | 0.989411 |
| SLC50A1 | greenyellow | -0.000808 | 0.996873 |
| KRT8P41 | greenyellow | 0.000568 | 0.997801 |
| ARL2 | grey60 | -0.546422 | 0.003875 |
| AKIRIN1 | grey60 | 0.531427 | 0.00521 |
| ZFYVE1 | grey60 | 0.517822 | 0.006738 |
| SGMS1 | grey60 | 0.505723 | 0.008397 |
| TBC1D9 | grey60 | 0.500737 | 0.009174 |
| HADH | grey60 | -0.467743 | 0.015971 |
| ABCD1 | grey60 | 0.452835 | 0.020179 |
| RASGEF1B | grey60 | 0.44515 | 0.022677 |
| CHAMP1 | grey60 | -0.440633 | 0.02426 |
| LUC7L | grey60 | 0.433539 | 0.026925 |
| TTC12 | grey60 | -0.420439 | 0.032465 |
| LOC401397 | grey60 | -0.418762 | 0.033236 |
| AKAP8L | grey60 | 0.416098 | 0.03449 |
| ACAD9 | grey60 | 0.4135 | 0.035749 |
| VCP | grey60 | 0.406373 | 0.039393 |
| YARS2 | grey60 | -0.404545 | 0.040374 |
| NFYA | grey60 | 0.403416 | 0.040989 |
| EEF2 | grey60 | 0.402717 | 0.041374 |
| SAMD4A | grey60 | 0.397822 | 0.044148 |
| FLJ45340 | grey60 | 0.395437 | 0.045552 |
| C4orf46 | grey60 | -0.38886 | 0.049606 |
| CDK5 | grey60 | -0.383384 | 0.053193 |
| CD82 | grey60 | 0.38014 | 0.055412 |
| NUMB | grey60 | 0.376822 | 0.057755 |
| MRPS18C | grey60 | -0.371181 | 0.061915 |
| LPIN1 | grey60 | 0.369961 | 0.062845 |
| DDX49 | grey60 | 0.363488 | 0.067957 |
| NPEPL1 | grey60 | 0.36069 | 0.070264 |
| TRMT2A | grey60 | 0.358001 | 0.072538 |
| SUPT5H | grey60 | 0.354626 | 0.07547 |
| BRCC3 | grey60 | -0.351342 | 0.078409 |
| SNORA53 | grey60 | -0.349737 | 0.079877 |
| EIF4G1 | grey60 | 0.348247 | 0.081257 |
| GNA12 | grey60 | 0.345266 | 0.084075 |
| CIZ1 | grey60 | 0.345 | 0.084331 |
| CHST7 | grey60 | 0.340057 | 0.089176 |
| CNNM4 | grey60 | 0.331911 | 0.097615 |
| RRP7B | grey60 | 0.331496 | 0.09806 |
| MICA | grey60 | -0.329835 | 0.099858 |
| GLRX | grey60 | -0.329257 | 0.100489 |
| AKAP7 | grey60 | -0.324663 | 0.105613 |
| NUDT14 | grey60 | -0.324311 | 0.106014 |
| MRPL11 | grey60 | -0.322339 | 0.108278 |
| FAM100B | grey60 | 0.321757 | 0.108953 |
| STX5 | grey60 | 0.318036 | 0.113343 |
| SNHG8 | grey60 | -0.308268 | 0.125486 |
| PDHB | grey60 | -0.303244 | 0.132088 |
| KCNAB2 | midnightblue | -0.061635963 | 0.760057291 |
| PSMD4 | midnightblue | -0.059090667 | 0.769697196 |
| ICAM2 | midnightblue | -0.055838691 | 0.782062129 |
| PABPN1 | midnightblue | -0.045647379 | 0.821135507 |
| COIL | midnightblue | -0.040875171 | 0.839582373 |
| TSPYL1 | midnightblue | -0.037555298 | 0.852464837 |
| THAP6 | midnightblue | -0.036399337 | 0.856959294 |
| NPHP4 | midnightblue | -0.035244389 | 0.861454162 |
| TMUB2 | midnightblue | -0.034461654 | 0.864502845 |
| NAPEPLD | midnightblue | -0.025780003 | 0.898434616 |
| RPS13 | midnightblue | -0.025492912 | 0.899560012 |
| ZFAND2B | midnightblue | -0.024908917 | 0.901849847 |
| CCDC142 | midnightblue | -0.014446794 | 0.942984572 |
| ITM2B | midnightblue | -0.012846752 | 0.949290784 |
| CEP41 | midnightblue | -0.011699486 | 0.953814334 |
| NDUFAB1 | midnightblue | -0.010227551 | 0.959620047 |
| GRHPR | midnightblue | -0.009819299 | 0.961230673 |
| ZNF41 | midnightblue | -0.005221821 | 0.979377331 |
| RPL23AP82 | midnightblue | -0.002461489 | 0.990277989 |
| CLASRP | midnightblue | 0.002869907 | 0.988664979 |
| SKP1 | midnightblue | 0.004734411 | 0.981301924 |
| GPRIN3 | midnightblue | 0.007676442 | 0.969686914 |
| STAMBP | midnightblue | 0.010433779 | 0.958806499 |
| ZNF407 | midnightblue | 0.012028628 | 0.952516412 |
| FUT11 | midnightblue | 0.014208293 | 0.943924369 |
| SMG6 | midnightblue | 0.021155343 | 0.91658509 |
| HDGFRP3 | midnightblue | 0.02216828 | 0.912605796 |
| CANT1 | midnightblue | 0.027212741 | 0.89282121 |
| MPST | midnightblue | 0.035279111 | 0.86131897 |
| NIPA1 | midnightblue | 0.042353055 | 0.833860327 |
| EIF3E | midnightblue | 0.044143065 | 0.826940869 |
| ZFP91 | midnightblue | 0.0453024 | 0.82246603 |
| FAM174A | midnightblue | 0.049074456 | 0.807944677 |
| IWS1 | midnightblue | 0.053547174 | 0.790806471 |
| ZFAND1 | midnightblue | 0.062691177 | 0.756070983 |
| TIMM21 | midnightblue | 0.064784758 | 0.748180105 |
| AKIRIN2 | midnightblue | 0.065295715 | 0.746257985 |
| RPS11 | midnightblue | 0.066503512 | 0.741720398 |
| FASTKD3 | midnightblue | 0.08634651 | 0.668476 |
| PHF14 | midnightblue | 0.090092327 | 0.654952527 |
| LSAMP | midnightblue | 0.104337943 | 0.604514741 |
| PPP1R10 | midnightblue | 0.112676006 | 0.57577652 |
| EXOC3 | midnightblue | 0.11766963 | 0.558861061 |
| SCARNA18 | midnightblue | 0.12082016 | 0.548306934 |
| PDHA1 | midnightblue | 0.121661247 | 0.54550505 |
| VKORC1L1 | midnightblue | 0.123467731 | 0.539509814 |
| ZNHIT3 | midnightblue | 0.123571749 | 0.539165551 |
| ATG3 | midnightblue | 0.128623064 | 0.522572969 |
| SMN1 | midnightblue | 0.135562505 | 0.500187504 |
| HERPUD2 | midnightblue | 0.138370552 | 0.49126705 |
| MARK2 | midnightblue | 0.14043786 | 0.484751393 |
| HEATR3 | midnightblue | 0.145944579 | 0.46761237 |
| DDX18 | midnightblue | 0.148917357 | 0.458492804 |
| ACBD3 | midnightblue | 0.177924995 | 0.374598554 |
| RBM8A | midnightblue | 0.188387834 | 0.346686787 |
| WDR75 | midnightblue | 0.195894455 | 0.327449918 |
| EIF3G | greenyellow | -0.113698 | 0.580241 |
| PPP4R1L | greenyellow | -0.106144 | 0.605803 |
| ZMYND11 | greenyellow | -0.105159 | 0.609172 |
| UBR5 | greenyellow | -0.10439 | 0.611805 |
| RASGEF1A | greenyellow | -0.103206 | 0.615871 |
| RUFY3 | greenyellow | -0.102158 | 0.619479 |
| EIF3M | greenyellow | -0.100528 | 0.625108 |
| MLST8 | greenyellow | -0.098726 | 0.631352 |
| ETHE1 | greenyellow | -0.095749 | 0.641726 |
| IPO5 | greenyellow | -0.095637 | 0.642117 |
| USP47 | greenyellow | -0.094249 | 0.64698 |
| UBR1 | greenyellow | -0.093954 | 0.648013 |
| ALOX15B | greenyellow | -0.092257 | 0.653979 |
| ARHGAP9 | greenyellow | -0.090577 | 0.659905 |
| EMP3 | greenyellow | -0.090096 | 0.661604 |
| UBE2Q2P1 | greenyellow | -0.089785 | 0.662704 |
| SSR3 | greenyellow | -0.089414 | 0.664018 |
| SNORA12 | greenyellow | -0.089022 | 0.665407 |
| TGFBRAP1 | greenyellow | -0.084807 | 0.680406 |
| TRMT11 | greenyellow | -0.084101 | 0.682931 |
| HCFC2 | greenyellow | -0.074391 | 0.717975 |
| CCNJ | greenyellow | -0.07234 | 0.72545 |
| TBXAS1 | greenyellow | -0.07077 | 0.731192 |
| DENND5B | greenyellow | -0.070605 | 0.731794 |
| SLC17A5 | greenyellow | -0.068514 | 0.739464 |
| EEF1G | greenyellow | -0.067257 | 0.744085 |
| ORC3 | greenyellow | -0.06517 | 0.751779 |
| RPS16 | greenyellow | -0.06414 | 0.755581 |
| MTA3 | greenyellow | -0.063716 | 0.75715 |
| AATF | greenyellow | -0.061072 | 0.766948 |
| ANKRD9 | greenyellow | -0.060958 | 0.767374 |
| ZNF581 | greenyellow | -0.06026 | 0.769963 |
| EIF2D | greenyellow | -0.059045 | 0.774485 |
| RREB1 | greenyellow | -0.058771 | 0.775504 |
| RNASET2 | greenyellow | -0.057089 | 0.781773 |
| NEURL4 | greenyellow | -0.055151 | 0.789014 |
| IFT20 | greenyellow | -0.054519 | 0.79138 |
| AAGAB | greenyellow | -0.054418 | 0.79176 |
| GAPVD1 | greenyellow | -0.053013 | 0.797023 |
| GHITM | greenyellow | -0.051524 | 0.802612 |
| SEMA3C | greenyellow | -0.05107 | 0.804319 |
| NSD1 | greenyellow | -0.045736 | 0.824427 |
| ZBTB41 | greenyellow | -0.044211 | 0.830196 |
| CAPN3 | greenyellow | -0.042812 | 0.835495 |
| RWDD4 | greenyellow | -0.042222 | 0.837733 |
| MOB1B | greenyellow | -0.04067 | 0.843623 |
| MLLT10 | greenyellow | -0.03778 | 0.854616 |
| VPS13C | greenyellow | -0.037337 | 0.856303 |
| NCF2 | greenyellow | -0.034238 | 0.868122 |
| SOWAHC | greenyellow | -0.034077 | 0.868737 |
| C11orf84 | greenyellow | -0.029991 | 0.884366 |
| SNAPC3 | greenyellow | -0.029527 | 0.886146 |
| ZMYM1 | greenyellow | -0.02626 | 0.898674 |
| SPRYD7 | greenyellow | -0.025927 | 0.899952 |
| SERTAD1 | greenyellow | -0.023758 | 0.908286 |
| C16orf70 | greenyellow | -0.021487 | 0.917023 |

**Table 6. Gene module membership and correlation with growth rate (48/72h), with adjusted p values for each module**

| | | | |
|---|---|---|---|
| U2AF2 | grey60 | 0.302653 | 0.132881 |
| UGP2 | grey60 | -0.301149 | 0.134914 |
| NRD1 | grey60 | -0.296315 | 0.141603 |
| NPAT | grey60 | -0.294634 | 0.143983 |
| RNF216 | grey60 | 0.291764 | 0.148113 |
| GRSF1 | grey60 | -0.291233 | 0.148886 |
| SOD1 | grey60 | 0.288044 | 0.153591 |
| ZNF324 | grey60 | 0.286203 | 0.156355 |
| CLOCK | grey60 | -0.28565 | 0.157192 |
| TPM1 | grey60 | -0.284446 | 0.159026 |
| RAPGEFL1 | grey60 | 0.2793 | 0.167034 |
| RAP2C | grey60 | -0.27711 | 0.170527 |
| GRPEL1 | grey60 | 0.269919 | 0.182353 |
| OC1005067[illegible] | grey60 | 0.267271 | 0.186849 |
| KHNYN | grey60 | 0.264705 | 0.191276 |
| ATP1B1 | grey60 | -0.264096 | 0.192338 |
| ZNF776 | grey60 | 0.254458 | 0.209679 |
| RPL3 | grey60 | 0.252297 | 0.213709 |
| TTC30B | grey60 | -0.250701 | 0.216717 |
| NREP | grey60 | -0.249758 | 0.218509 |
| HPS4 | grey60 | -0.248471 | 0.220969 |
| NCOA6 | grey60 | 0.247063 | 0.223683 |
| DAPK3 | grey60 | 0.243715 | 0.230222 |
| CASC3 | grey60 | 0.239744 | 0.238144 |
| PDE12 | grey60 | 0.237236 | 0.243238 |
| SLC3A2 | grey60 | 0.233962 | 0.249997 |
| GTF2H2 | grey60 | -0.233746 | 0.250446 |
| RPRD1B | grey60 | 0.23196 | 0.254188 |
| MOAP1 | grey60 | 0.231212 | 0.255765 |
| CACTIN | grey60 | 0.230571 | 0.257123 |
| TTC4 | grey60 | 0.224148 | 0.27098 |
| PTBP1 | grey60 | 0.221442 | 0.27696 |
| SCOC | grey60 | -0.217738 | 0.28528 |
| NFE2L1 | grey60 | 0.215221 | 0.291022 |
| SPA17 | grey60 | -0.212081 | 0.298288 |
| VPS4A | grey60 | 0.208944 | 0.305659 |
| SGPL1 | grey60 | -0.208283 | 0.307226 |
| HLA-DMB | grey60 | 0.205851 | 0.313035 |
| RBM47 | grey60 | 0.201333 | 0.324006 |
| PARP6 | grey60 | 0.199223 | 0.329208 |
| IFT122 | grey60 | -0.196826 | 0.335179 |
| TNK2 | grey60 | 0.194838 | 0.340182 |
| POFUT2 | grey60 | 0.19099 | 0.349989 |
| HAUS1 | grey60 | -0.184567 | 0.366731 |
| ZKSCAN1 | grey60 | 0.182035 | 0.373456 |
| NR2C2 | grey60 | 0.179014 | 0.381573 |
| DNAJC15 | grey60 | -0.178484 | 0.383006 |
| C11orf30 | grey60 | 0.162046 | 0.429011 |
| MTHFD2 | grey60 | 0.155271 | 0.448809 |
| TMEM220 | grey60 | -0.149977 | 0.46461 |
| UPF3B | grey60 | 0.137276 | 0.503672 |
| RALGAPB | grey60 | 0.135069 | 0.510619 |
| PDS5B | grey60 | -0.133018 | 0.517119 |
| NNT | grey60 | -0.131389 | 0.522311 |
| C2 | grey60 | 0.118102 | 0.565559 |
| HSPE1 | grey60 | -0.117499 | 0.56756 |
| RCN2 | midnightblue | 0.204541965 | 0.306113564 |
| PPIA | midnightblue | 0.222191191 | 0.265320625 |
| UQCRQ | midnightblue | 0.231200066 | 0.245923048 |
| KCTD3 | midnightblue | 0.248002494 | 0.212295586 |
| IFI16 | midnightblue | 0.261579084 | 0.187511881 |
| PIGB | midnightblue | 0.261947565 | 0.186868486 |
| HSPA14 | midnightblue | 0.270888123 | 0.171724121 |
| FAM46A | midnightblue | 0.274054103 | 0.166574107 |
| CCT8 | midnightblue | 0.291014584 | 0.140833446 |
| DDX58 | midnightblue | 0.291720999 | 0.139827573 |
| COPS3 | midnightblue | 0.362996123 | 0.062746505 |
| RPL27 | midnightblue | 0.402215383 | 0.037540008 |
| NDUFB4 | midnightblue | 0.412778697 | 0.032368315 |
| ING3 | midnightblue | 0.445190548 | 0.01996508 |
| FBXL19 | NA | -0.659386967 | 0.000183285 |
| NAB2 | NA | -0.608886784 | 0.000750406 |
| ZHX1 | NA | -0.59813596 | 0.000983122 |
| TGIF2 | NA | -0.578623171 | 0.001567944 |
| C12orf45 | NA | -0.566424564 | 0.002069228 |
| CSRNP2 | NA | -0.535351857 | 0.004007341 |
| C1orf109 | NA | -0.52055305 | 0.005374416 |
| DIP2B | NA | -0.516412431 | 0.005821047 |
| PIP5K1A | NA | -0.513499659 | 0.006153699 |
| MSC | NA | -0.512176771 | 0.006309997 |
| UBAP2 | NA | -0.5100676 | 0.006566118 |
| WDR5 | NA | -0.507564852 | 0.006881351 |
| LARS2 | NA | -0.506484617 | 0.007021298 |
| DHX57 | NA | -0.503031605 | 0.007484824 |
| MICALCL | NA | -0.498631322 | 0.008112555 |
| FAM102A | NA | -0.494780242 | 0.008697642 |
| CTNS | NA | -0.494390667 | 0.00875875 |
| PUF60 | NA | -0.486628137 | 0.010053463 |
| DGKA | NA | -0.480119955 | 0.01125867 |
| HIST1H2BH | NA | -0.474427363 | 0.012409273 |
| SLC44A1 | NA | -0.472341724 | 0.01285455 |
| GNA15 | NA | -0.469863494 | 0.013400801 |
| FPR3 | NA | -0.463969623 | 0.014777584 |
| CUEDC1 | NA | -0.463296996 | 0.014941879 |
| STK17B | NA | -0.462853276 | 0.015051086 |
| GMIP | NA | -0.461915537 | 0.015284049 |
| ARHGEF40 | NA | -0.461437726 | 0.015403892 |
| C6orf57 | NA | -0.460830199 | 0.015557389 |
| ZNF792 | NA | -0.459977999 | 0.015774829 |
| KHSRP | NA | -0.452186687 | 0.01788148 |
| XAB2 | NA | -0.446098538 | 0.019683938 |
| IAH1 | NA | -0.443958675 | 0.020351752 |
| ULBP2 | NA | -0.443174526 | 0.020601064 |
| METTL21B | NA | -0.435553025 | 0.023156848 |
| ZEB2 | NA | -0.433551177 | 0.023869307 |
| POM121 | NA | -0.433510988 | 0.023883791 |
| CLU | NA | -0.432718875 | 0.024170724 |
| C16orf88 | NA | -0.432188547 | 0.024364391 |
| RASGRP4 | NA | -0.427610481 | 0.026089217 |
| CASS4 | NA | -0.427359564 | 0.026186541 |
| CYFIP2 | NA | -0.426705021 | 0.026441807 |
| CTDP1 | NA | -0.425809297 | 0.026794395 |
| MAST2 | greenyellow | -0.019561 | 0.924438 |
| RPL3 | greenyellow | -0.017638 | 0.931847 |
| MAPK1 | greenyellow | -0.013233 | 0.948845 |
| RNF20 | greenyellow | -0.012878 | 0.950214 |
| HEATR5A | greenyellow | -0.011181 | 0.95677 |
| UTP20 | greenyellow | -0.010052 | 0.961131 |
| ARGLU1 | greenyellow | -0.00767 | 0.970336 |
| ZFP92 | greenyellow | -0.006822 | 0.973616 |
| LOC220729 | greenyellow | -0.003209 | 0.987587 |
| PALB2 | greenyellow | -0.002913 | 0.98873 |
| PIK3IP1 | greenyellow | -0.002183 | 0.991556 |
| PHF10 | greenyellow | -0.001097 | 0.995755 |
| ZC3HC1 | greenyellow | -0.000489 | 0.998108 |
| PPP1R18 | greenyellow | 0.005516 | 0.978663 |
| YTHDC1 | greenyellow | 0.006975 | 0.973023 |
| CD2AP | greenyellow | 0.009765 | 0.962237 |
| RBM4 | greenyellow | 0.010192 | 0.960587 |
| DIDO1 | greenyellow | 0.011955 | 0.953779 |
| EXOSC1 | greenyellow | 0.012902 | 0.950121 |
| FBXO44 | greenyellow | 0.013935 | 0.946133 |
| SNX30 | greenyellow | 0.016101 | 0.937773 |
| VMO1 | greenyellow | 0.019476 | 0.924766 |
| C17orf59 | greenyellow | 0.02194 | 0.91528 |
| PHLPP2 | greenyellow | 0.024115 | 0.906915 |
| C11orf10 | greenyellow | 0.032115 | 0.876238 |
| AIP | greenyellow | 0.033046 | 0.872676 |
| MRPL2 | greenyellow | 0.040084 | 0.845851 |
| DHODH | greenyellow | 0.041105 | 0.841972 |
| IRS2 | greenyellow | 0.041389 | 0.840893 |
| HVCN1 | greenyellow | 0.041781 | 0.839406 |
| C17orf62 | greenyellow | 0.047932 | 0.816135 |
| DDB2 | greenyellow | 0.048512 | 0.813946 |
| HIST1H4H | greenyellow | 0.048796 | 0.812876 |
| FAM208A | greenyellow | 0.052769 | 0.797937 |
| FAU | greenyellow | 0.056055 | 0.785634 |
| CCDC94 | greenyellow | 0.059417 | 0.773099 |
| MRPL10 | greenyellow | 0.060143 | 0.7704 |
| SUGP1 | greenyellow | 0.061199 | 0.766477 |
| PCBD1 | greenyellow | 0.062626 | 0.761185 |
| THAP7 | greenyellow | 0.063487 | 0.757996 |
| RPL18 | greenyellow | 0.064614 | 0.753829 |
| DGCR6L | greenyellow | 0.0723 | 0.725598 |
| TMBIM1 | greenyellow | 0.072573 | 0.724601 |
| PGAP3 | greenyellow | 0.081682 | 0.691604 |
| CASP7 | greenyellow | 0.087841 | 0.669597 |
| TMEM60 | greenyellow | 0.092471 | 0.653225 |
| DCAF10 | greenyellow | 0.098513 | 0.632092 |
| PSMB10 | greenyellow | 0.100962 | 0.623606 |
| TMEM117 | greenyellow | 0.101725 | 0.62097 |
| PYCARD | greenyellow | 0.103096 | 0.616248 |
| CNPY2 | greenyellow | 0.106127 | 0.605859 |
| C17orf85 | greenyellow | 0.112875 | 0.583 |
| SHMT2 | greenyellow | 0.113624 | 0.580489 |
| PMS2P1 | greenyellow | 0.119227 | 0.561835 |
| NDUFB11 | greenyellow | 0.119332 | 0.56149 |
| SLC25A19 | greenyellow | 0.119719 | 0.560211 |

**Table 6. Gene module membership and correlation with growth rate (48/72h), with adjusted p values for each module**

| | | | | | | | | | | | |
|---|---|---|---|---|---|---|---|---|---|---|---|
| BTN3A2 | grey60 | -0.117021 | 0.569147 | RPL7L1 | NA | -0.425703975 | 0.026836102 | EGLN2 | greenyellow | 0.120128 | 0.558862 |
| FLNA | grey60 | 0.113391 | 0.581268 | HLA-A | NA | -0.424200692 | 0.027437153 | TJP1 | greenyellow | 0.120494 | 0.557655 |
| PLXNA2 | grey60 | 0.110819 | 0.589928 | DUSP16 | NA | -0.422542537 | 0.028112712 | FAM3A | greenyellow | 0.121033 | 0.555882 |
| HLA-E | grey60 | 0.107905 | 0.599801 | STYXL1 | NA | -0.421627668 | 0.028491161 | EXOSC5 | greenyellow | 0.129121 | 0.529578 |
| ABHD4 | grey60 | 0.101561 | 0.621536 | CEP72 | NA | -0.420047828 | 0.02915437 | DUS3L | greenyellow | 0.135386 | 0.509619 |
| AKAP17A | grey60 | 0.097628 | 0.63517 | INPP1 | NA | -0.41951016 | 0.029382902 | C2 | greenyellow | 0.13671 | 0.505449 |
| PRKAB1 | grey60 | -0.096385 | 0.639504 | ATP5F1 | NA | -0.418129884 | 0.029976201 | SPINT2 | greenyellow | 0.148731 | 0.46837 |
| FXR2 | grey60 | 0.091606 | 0.656273 | TRIM56 | NA | -0.417463723 | 0.03026598 | CARHSP1 | greenyellow | 0.153769 | 0.453264 |
| TRPC6 | grey60 | -0.086188 | 0.675479 | SEMA4B | NA | -0.416234564 | 0.030806586 | SPTY2D1 | greenyellow | 0.153868 | 0.452967 |
| TOMM40L | grey60 | 0.085823 | 0.676779 | ARHGEF9 | NA | -0.414422131 | 0.031617888 | KCNA3 | greenyellow | 0.154347 | 0.451546 |
| CTNNBL1 | grey60 | 0.076783 | 0.709288 | JMJD4 | NA | -0.411452948 | 0.032984054 | LAGE3 | greenyellow | 0.168301 | 0.411163 |
| CD300A | grey60 | 0.075718 | 0.713151 | SRRM2 | NA | -0.409706594 | 0.033809458 | CCZ1 | greenyellow | 0.17192 | 0.401025 |
| DDB1 | grey60 | 0.06863 | 0.739038 | PRPSAP1 | NA | -0.409376923 | 0.033967118 | LILRA2 | greenyellow | 0.1779 | 0.384592 |
| PRPF31 | grey60 | 0.068583 | 0.739208 | CCDC104 | NA | -0.407008187 | 0.035117341 | MUL1 | greenyellow | 0.180833 | 0.376673 |
| ANXA1 | grey60 | 0.056227 | 0.784993 | PPAPDC2 | NA | -0.406999542 | 0.035121595 | GSTT1 | greenyellow | 0.18375 | 0.368892 |
| TFDP1 | grey60 | 0.052197 | 0.800085 | PTX3 | NA | -0.406493185 | 0.035371499 | POLD4 | greenyellow | 0.193002 | 0.344841 |
| NSFL1C | grey60 | 0.049917 | 0.808656 | UBP1 | NA | -0.406085714 | 0.03557363 | FARSA | greenyellow | 0.194243 | 0.341689 |
| ARF1 | grey60 | 0.041406 | 0.840829 | TBCE | NA | -0.404420239 | 0.036409443 | R3HDM4 | greenyellow | 0.195802 | 0.337751 |
| SMPD2 | grey60 | 0.034968 | 0.865338 | CLOCK | NA | -0.402740242 | 0.037268373 | TRAPPC6A | greenyellow | 0.199683 | 0.328069 |
| MLF2 | grey60 | 0.031742 | 0.877665 | SOCS3 | NA | -0.399716908 | 0.038854813 | CTSF | greenyellow | 0.204229 | 0.316948 |
| GTPBP1 | grey60 | 0.031235 | 0.879602 | MTSS1 | NA | -0.395524636 | 0.04114318 | CDIPT | greenyellow | 0.252127 | 0.214028 |
| SRPR | grey60 | 0.027398 | 0.894305 | LMNB1 | NA | -0.395421279 | 0.041200921 | BCL2L12 | greenyellow | 0.260272 | 0.199096 |
| ANKLE2 | grey60 | -0.024947 | 0.903717 | LOC150381 | NA | -0.39295686 | 0.042596863 | NFKBIB | greenyellow | 0.267488 | 0.186477 |
| LOC648987 | grey60 | 0.020943 | 0.919116 | RALY | NA | -0.389414403 | 0.044668887 | GHDC | greenyellow | 0.279646 | 0.166487 |
| MAF1 | grey60 | 0.016599 | 0.935854 | ADORA2A | NA | -0.388090302 | 0.045463535 | CDC34 | greenyellow | 0.282484 | 0.162047 |
| AKAP8 | grey60 | 0.010367 | 0.959911 | CA12 | NA | -0.385806692 | 0.046860245 | APBA3 | greenyellow | 0.332409 | 0.097082 |
| NFKBIB | grey60 | 0.003515 | 0.986402 | CBX7 | NA | -0.383668897 | 0.048198255 | METTL13 | greenyellow | 0.338476 | 0.090769 |
| ATP6V0E2 | lightcyan | -0.477192 | 0.013699 | CKAP2 | NA | -0.379895851 | 0.050633038 | GTF2F2 | grey60 | -0.342056 | 0.087191 |
| UBE2T | lightcyan | -0.381849 | 0.054235 | DYM | NA | -0.379552005 | 0.050859644 | MRPS11 | grey60 | -0.274641 | 0.174526 |
| MIA3 | lightcyan | -0.342204 | 0.087046 | GMEB2 | NA | -0.379255357 | 0.051055784 | HIGD1A | grey60 | -0.260615 | 0.198484 |
| IL27 | lightcyan | -0.325152 | 0.105059 | CALML4 | NA | -0.378483957 | 0.051568606 | BUD31 | grey60 | -0.258773 | 0.20179 |
| C10orf76 | lightcyan | -0.323074 | 0.107429 | SLC35E4 | NA | -0.376546857 | 0.052874216 | UQCRB | grey60 | -0.238689 | 0.240278 |
| KIAA0240 | lightcyan | -0.316099 | 0.11568 | TRPS1 | NA | -0.376086672 | 0.053188157 | ZNF583 | grey60 | -0.230764 | 0.256715 |
| MRGBP | lightcyan | -0.314201 | 0.118003 | PLEKHO1 | NA | -0.374234325 | 0.054466644 | DDX1 | grey60 | -0.224251 | 0.270755 |
| PTH2R | lightcyan | -0.313329 | 0.119082 | PEX7 | NA | -0.372235276 | 0.055873234 | CCDC104 | grey60 | -0.212808 | 0.296597 |
| ABHD13 | lightcyan | -0.312511 | 0.1201 | SNHG7 | NA | -0.369950991 | 0.057515099 | NT5DC1 | grey60 | -0.206789 | 0.310787 |
| ACN9 | lightcyan | -0.295939 | 0.142133 | MRPS14 | NA | -0.369317134 | 0.057977298 | LAPTM4A | grey60 | -0.205053 | 0.314956 |
| DONSON | lightcyan | -0.284594 | 0.158801 | C7orf31 | NA | -0.367920697 | 0.059005774 | ZNF124 | grey60 | -0.203582 | 0.318515 |
| GTF2F2 | lightcyan | -0.283347 | 0.160713 | FUOM | NA | -0.3675382 | 0.059289946 | C20orf96 | grey60 | -0.199973 | 0.327352 |
| TRDMT1 | lightcyan | -0.281479 | 0.163609 | NAIF1 | NA | -0.36522745 | 0.061029426 | POLR2G | grey60 | -0.193238 | 0.34424 |
| SNX4 | lightcyan | 0.281276 | 0.163927 | GCLM | NA | -0.363225907 | 0.062567967 | RSBN1L-AS1 | grey60 | -0.187345 | 0.359433 |
| ICT1 | lightcyan | -0.277989 | 0.16912 | FAM110A | NA | -0.363010618 | 0.062735231 | VSIG4 | grey60 | -0.187283 | 0.359595 |
| CTSF | lightcyan | 0.273862 | 0.1758 | PPM1H | NA | -0.362446688 | 0.063175008 | CTNNA1 | grey60 | -0.177222 | 0.386435 |
| P2RX7 | lightcyan | -0.272603 | 0.177875 | MALSU1 | NA | -0.360723844 | 0.064533372 | FAF1 | grey60 | -0.175443 | 0.391297 |
| FLI1 | lightcyan | -0.272064 | 0.178768 | MCCC1 | NA | -0.360596905 | 0.064634344 | FRG1 | grey60 | -0.167615 | 0.413098 |
| TSPAN15 | lightcyan | -0.268965 | 0.183964 | ZNF845 | NA | -0.36043048 | 0.064766909 | ENO1 | grey60 | -0.167573 | 0.413218 |
| HADHA | lightcyan | -0.265823 | 0.189338 | PFKFB3 | NA | -0.359864933 | 0.065218967 | COQ5 | grey60 | -0.164976 | 0.420598 |
| MT1H | lightcyan | -0.254648 | 0.209329 | MMP7 | NA | -0.358777712 | 0.066094868 | C1orf174 | grey60 | -0.163123 | 0.425906 |
| NOP10 | lightcyan | 0.251667 | 0.214893 | SESTD1 | NA | -0.357945212 | 0.066771683 | SMN2 | grey60 | -0.161635 | 0.430197 |
| HSD17B11 | lightcyan | -0.243809 | 0.230037 | IFT80 | NA | -0.35552972 | 0.06876578 | SEC63 | grey60 | -0.157756 | 0.44149 |
| CEP120 | lightcyan | -0.243325 | 0.230992 | INPPL1 | NA | -0.355296598 | 0.068960636 | SUV420H1 | grey60 | -0.15767 | 0.441744 |
| LDHA | lightcyan | -0.234086 | 0.249738 | PPT2 | NA | -0.35255568 | 0.071283686 | ALDH4A1 | grey60 | -0.153093 | 0.455276 |
| DCPS | lightcyan | 0.227835 | 0.26297 | MIB2 | NA | -0.352228448 | 0.071565002 | CAPRIN2 | grey60 | -0.149743 | 0.465314 |
| PRDM10 | lightcyan | -0.227264 | 0.2642 | SAMD1 | NA | -0.352098663 | 0.071676813 | CALR | grey60 | -0.145797 | 0.477288 |
| TET3 | lightcyan | -0.209294 | 0.304831 | IFT140 | NA | -0.349661139 | 0.073801785 | CTAGE9 | grey60 | -0.145183 | 0.479165 |
| SERINC1 | lightcyan | -0.2062 | 0.312196 | ZC3H18 | NA | -0.349387013 | 0.074043751 | TSPYL4 | grey60 | -0.144182 | 0.482233 |

**Table 6. Gene module membership and correlation with growth rate (48/72h), with adjusted p values for each module**

| | | | | | | | | | | | |
|---|---|---|---|---|---|---|---|---|---|---|---|
| KAT2B | lightcyan | -0.206139 | 0.312345 | SPA17 | NA | -0.348553101 | 0.074783572 | UTP6 | grey60 | -0.144139 | 0.482365 |
| THOC1 | lightcyan | -0.20011 | 0.327015 | CDKN2B | NA | -0.348138202 | 0.075153762 | NDUFS2 | grey60 | -0.14131 | 0.491091 |
| TKT | lightcyan | -0.199089 | 0.32954 | CBY1 | NA | -0.347956864 | 0.075315999 | NEMF | grey60 | -0.140902 | 0.492358 |
| NOP14 | lightcyan | -0.197155 | 0.334357 | OTUD7B | NA | -0.34783235 | 0.075427552 | SMNDC1 | grey60 | -0.139893 | 0.495494 |
| RPL38 | lightcyan | -0.189802 | 0.35305 | MICALL2 | NA | -0.347291764 | 0.075913339 | CAB39 | grey60 | -0.134874 | 0.511235 |
| RPL7A | lightcyan | 0.183704 | 0.369016 | AKAP11 | NA | -0.347255684 | 0.075945846 | ZNF597 | grey60 | -0.124221 | 0.545444 |
| C22orf28 | lightcyan | -0.180523 | 0.377506 | C16orf54 | NA | -0.345203947 | 0.077812025 | CC2D1B | grey60 | -0.12152 | 0.554282 |
| C1orf54 | lightcyan | -0.178607 | 0.382674 | PAK2 | NA | -0.344988895 | 0.078009638 | RBM10 | grey60 | -0.11711 | 0.568853 |
| APOC1 | lightcyan | -0.17751 | 0.385652 | TMC8 | NA | -0.344682169 | 0.078292155 | MDH2 | grey60 | -0.115804 | 0.573199 |
| SAP30L | lightcyan | -0.169871 | 0.406747 | LOC1001327 | NA | -0.342948835 | 0.079903385 | EEF1A1 | grey60 | -0.114576 | 0.577299 |
| PIGP | lightcyan | -0.163036 | 0.426158 | STRN3 | NA | -0.342038798 | 0.080759373 | SCAF4 | grey60 | -0.114393 | 0.577911 |
| SNORA75 | lightcyan | -0.151106 | 0.461218 | ZNF470 | NA | -0.341416225 | 0.081348982 | RFT1 | grey60 | -0.110186 | 0.592066 |
| PSMD13 | lightcyan | -0.150374 | 0.463416 | ZNF212 | NA | -0.341300615 | 0.08145883 | DMAP1 | grey60 | -0.104934 | 0.609942 |
| CCL18 | lightcyan | -0.148098 | 0.470288 | USP24 | NA | -0.340558957 | 0.082166217 | ANKHD1 | grey60 | -0.074358 | 0.718093 |
| IL3RA | lightcyan | 0.147995 | 0.470599 | OSGIN1 | NA | -0.338791498 | 0.083870834 | COMMD3 | grey60 | -0.071216 | 0.72956 |
| ARHGAP4 | lightcyan | -0.145477 | 0.478265 | FAM161B | NA | -0.338372339 | 0.084279002 | RALGDS | grey60 | -0.069835 | 0.734616 |
| OC1005055 | lightcyan | 0.140966 | 0.49216 | TMEM161E | NA | -0.336664223 | 0.08595794 | ISCA1 | grey60 | -0.068391 | 0.739915 |
| FAM108A1 | lightcyan | -0.139058 | 0.498096 | SERGEF | NA | -0.336253068 | 0.086365831 | DBNDD2 | grey60 | -0.066074 | 0.748442 |
| ANGPT4 | lightcyan | -0.137961 | 0.501523 | BBS9 | NA | -0.335948508 | 0.086668918 | CEP120 | grey60 | -0.063684 | 0.757267 |
| RPL30 | lightcyan | -0.131763 | 0.521117 | ACD | NA | -0.335611297 | 0.087005438 | MAN1C1 | grey60 | -0.060271 | 0.769926 |
| ACLY | lightcyan | -0.130922 | 0.523801 | CHST15 | NA | -0.335410197 | 0.087206595 | REPS1 | grey60 | -0.058054 | 0.778176 |
| DHX38 | lightcyan | 0.130914 | 0.523827 | C10orf88 | NA | -0.335200926 | 0.0874163 | XRCC6 | grey60 | -0.057538 | 0.780101 |
| AGFG2 | lightcyan | -0.130384 | 0.525524 | DCUN1D3 | NA | -0.333249887 | 0.089389795 | NARF | grey60 | -0.055917 | 0.786153 |
| TBC1D4 | lightcyan | -0.130236 | 0.525997 | CC2D1B | NA | -0.331163897 | 0.091536835 | VPS18 | grey60 | -0.053029 | 0.796964 |
| SCNM1 | lightcyan | -0.130148 | 0.52628 | NAT9 | NA | -0.330666367 | 0.092054622 | SLC12A6 | grey60 | -0.051258 | 0.803613 |
| UPF2 | lightcyan | -0.128634 | 0.531145 | ERCC1 | NA | -0.330311853 | 0.092424915 | FAM192A | grey60 | -0.051071 | 0.804317 |
| DVL3 | lightcyan | -0.124537 | 0.544413 | STARD4 | NA | -0.329743221 | 0.093021197 | ZNF175 | grey60 | -0.050378 | 0.806922 |
| SRP14 | lightcyan | 0.122894 | 0.549776 | UBXN2A | NA | -0.32945104 | 0.09332871 | VEZT | grey60 | -0.047381 | 0.818213 |
| AAAS | lightcyan | -0.122683 | 0.550469 | UPF1 | NA | -0.329316509 | 0.093470558 | CABIN1 | grey60 | -0.044614 | 0.82867 |
| FOSL2 | lightcyan | -0.120903 | 0.556308 | S100A8 | NA | -0.328751457 | 0.094068112 | PLXDC2 | grey60 | -0.04368 | 0.832205 |
| FAM222A | lightcyan | -0.120851 | 0.556479 | RC3H2 | NA | -0.328597841 | 0.09423106 | DHCR24 | grey60 | -0.042154 | 0.837992 |
| NGFRAP1 | lightcyan | -0.117909 | 0.566198 | MED25 | NA | -0.328572168 | 0.094258314 | SIAE | grey60 | -0.04149 | 0.840511 |
| DBF4B | lightcyan | -0.117618 | 0.567164 | PPBP | NA | -0.328252219 | 0.094598456 | RNF187 | grey60 | -0.041342 | 0.841071 |
| P2RY1 | lightcyan | -0.112509 | 0.584231 | TDP1 | NA | -0.32779402 | 0.095087177 | TIMMDC1 | grey60 | -0.038815 | 0.850675 |
| ZFYVE19 | lightcyan | -0.107075 | 0.602627 | ITPRIPL2 | NA | -0.326067757 | 0.096945491 | ENOX2 | grey60 | -0.036158 | 0.860798 |
| GATAD2B | lightcyan | -0.105953 | 0.606456 | MFHAS1 | NA | -0.325297898 | 0.09778297 | ABHD2 | grey60 | -0.032935 | 0.873103 |
| HIRIP3 | lightcyan | -0.105327 | 0.608596 | MICA | NA | -0.325221276 | 0.097866618 | AGL | grey60 | -0.031863 | 0.877199 |
| DIS3L2 | lightcyan | -0.103784 | 0.613885 | MMP9 | NA | -0.324899225 | 0.098218787 | COX19 | grey60 | -0.031658 | 0.877984 |
| RAB13 | lightcyan | 0.1021 | 0.619679 | DOM3Z | NA | -0.324076107 | 0.099123195 | ZNF664 | grey60 | -0.026574 | 0.897467 |
| ATP11A | lightcyan | -0.099887 | 0.627325 | CCND2 | NA | -0.323741744 | 0.099492355 | COX17 | grey60 | -0.019569 | 0.924407 |
| ATP5I | lightcyan | 0.096974 | 0.637449 | MAN1B1 | NA | -0.320979447 | 0.102581598 | SENP7 | grey60 | -0.018933 | 0.926857 |
| NCAPD3 | lightcyan | 0.096165 | 0.640273 | GGT1 | NA | -0.320921895 | 0.102646714 | DNAJA3 | grey60 | -0.017517 | 0.932314 |
| MAML3 | lightcyan | -0.091785 | 0.655643 | WDR83 | NA | -0.320693352 | 0.102905601 | ELMO2 | grey60 | -0.017145 | 0.933747 |
| CHMP4A | lightcyan | -0.090924 | 0.658679 | ZNF568 | NA | -0.320265958 | 0.103391045 | SLC30A9 | grey60 | -0.016557 | 0.936015 |
| ADPRHL2 | lightcyan | -0.090557 | 0.659975 | POC5 | NA | -0.320069462 | 0.103614802 | BAZ2B | grey60 | -0.016339 | 0.936856 |
| MMP14 | lightcyan | -0.090462 | 0.66031 | LOC1001328 | NA | -0.317535981 | 0.106532206 | MAN2B2 | grey60 | -0.004891 | 0.981082 |
| CCL23 | lightcyan | -0.089181 | 0.664844 | IL2RA | NA | -0.315388088 | 0.109053073 | PKN2 | grey60 | 0.002271 | 0.991215 |
| NCOA1 | lightcyan | -0.088255 | 0.668128 | UBTD2 | NA | -0.315294387 | 0.109164044 | ASS1 | grey60 | 0.003268 | 0.987358 |
| C12orf43 | lightcyan | -0.086578 | 0.67409 | NFATC1 | NA | -0.314396107 | 0.110232136 | PDZD8 | grey60 | 0.004548 | 0.982407 |
| FONG | lightcyan | -0.08586 | 0.676648 | TBX21 | NA | -0.314136772 | 0.110541932 | AGGF1 | grey60 | 0.004628 | 0.982097 |
| RNF130 | lightcyan | -0.085209 | 0.67897 | ARL6IP5 | NA | -0.312764471 | 0.112192002 | CTIF | grey60 | 0.009945 | 0.961542 |
| FAM214B | lightcyan | -0.084157 | 0.68273 | ATL1 | NA | -0.311995245 | 0.113124862 | NUP160 | grey60 | 0.01042 | 0.959707 |
| TOM1L1 | lightcyan | 0.082546 | 0.688502 | PRMT10 | NA | -0.311882519 | 0.113262048 | MED8 | grey60 | 0.016259 | 0.937164 |
| RNF7 | lightcyan | 0.078174 | 0.704253 | RNF114 | NA | -0.311859466 | 0.113290118 | BYSL | grey60 | 0.01651 | 0.936199 |
| RPS10 | lightcyan | -0.077226 | 0.707682 | RASAL3 | NA | -0.311516166 | 0.113708744 | BTBD2 | grey60 | 0.016678 | 0.935551 |
| ATXN2L | lightcyan | -0.074595 | 0.717233 | BST1 | NA | -0.311413194 | 0.113834532 | MLL4 | grey60 | 0.02395 | 0.907547 |

**Table 6. Gene module membership and correlation with growth rate (48/72h), with adjusted p values for each module**

| | | | |
|---|---|---|---|
| SCAF11 | lightcyan | -0.072951 | 0.723223 |
| MYEOV2 | lightcyan | -0.072526 | 0.724774 |
| ERH | lightcyan | -0.072513 | 0.724821 |
| KIAA1033 | lightcyan | 0.070064 | 0.733776 |
| XPNPEP3 | lightcyan | -0.06691 | 0.745361 |
| ATIC | lightcyan | -0.065135 | 0.751907 |
| ZNF592 | lightcyan | -0.061853 | 0.764052 |
| HLA-DPB1 | lightcyan | -0.061469 | 0.765475 |
| NDUFA1 | lightcyan | -0.060566 | 0.768826 |
| SNRPA1 | lightcyan | 0.057034 | 0.78198 |
| DIS3L | lightcyan | -0.053638 | 0.794681 |
| SERTAD2 | lightcyan | -0.053512 | 0.795151 |
| FBXW11 | lightcyan | -0.051589 | 0.802368 |
| C1QBP | lightcyan | 0.051486 | 0.802755 |
| CLCN4 | lightcyan | -0.051147 | 0.804031 |
| SLC4A11 | lightcyan | -0.049219 | 0.811285 |
| TMEM120A | lightcyan | -0.047927 | 0.816152 |
| ERLIN1 | lightcyan | -0.04725 | 0.818709 |
| CD63 | lightcyan | -0.047129 | 0.819165 |
| NDUFB5 | lightcyan | 0.046229 | 0.822563 |
| RPL21 | lightcyan | -0.045917 | 0.823741 |
| RIPK1 | lightcyan | -0.045337 | 0.825935 |
| C19orf53 | lightcyan | -0.044981 | 0.82728 |
| ZNF593 | lightcyan | -0.044684 | 0.828406 |
| SNRPE | lightcyan | 0.04435 | 0.829669 |
| YWHAZ | lightcyan | -0.041665 | 0.839846 |
| MRPL46 | lightcyan | 0.039453 | 0.84825 |
| SLC25A5 | lightcyan | -0.035786 | 0.862217 |
| MTMR9 | lightcyan | 0.034073 | 0.868755 |
| ARIH1 | lightcyan | -0.033797 | 0.869807 |
| ZZEF1 | lightcyan | -0.029353 | 0.88681 |
| PNMA1 | lightcyan | 0.028877 | 0.888635 |
| NPM3 | lightcyan | 0.02717 | 0.895183 |
| PUM1 | lightcyan | -0.026979 | 0.895914 |
| RAB34 | lightcyan | 0.025064 | 0.903268 |
| G3BP1 | lightcyan | -0.024395 | 0.905837 |
| CTAGE9 | lightcyan | 0.024389 | 0.905862 |
| VPS72 | lightcyan | 0.023697 | 0.908521 |
| YWHAE | lightcyan | 0.020872 | 0.919387 |
| AMZ2 | lightcyan | -0.018137 | 0.929927 |
| RC3H1 | lightcyan | 0.01519 | 0.941288 |
| OC1005068 | lightcyan | -0.014475 | 0.944047 |
| LINC00622 | lightcyan | 0.013167 | 0.949097 |
| SCARNA18 | lightcyan | 0.011298 | 0.956315 |
| ANKRD40 | lightcyan | 0.010029 | 0.96122 |
| GPNMB | lightcyan | 0.009914 | 0.961663 |
| HERC4 | lightcyan | 0.008727 | 0.96625 |
| ZFHX3 | lightcyan | -0.007181 | 0.972226 |
| PSMB10 | lightcyan | -0.006324 | 0.975542 |
| FBXW2 | lightcyan | 0.004991 | 0.980695 |
| RBM4B | lightcyan | -0.003852 | 0.985101 |
| RRAD | lightgreen | -0.453057 | 0.02011 |
| SLC9A1 | lightgreen | 0.387765 | 0.050308 |
| PELI2 | lightgreen | 0.33716 | 0.092112 |
| TMEM79 | lightgreen | 0.336828 | 0.092452 |
| TUSC1 | lightgreen | -0.324411 | 0.1059 |
| NADK | NA | -0.311122622 | 0.114190042 |
| OBFC1 | NA | -0.309216782 | 0.116542164 |
| RTN2 | NA | -0.308346184 | 0.117628426 |
| RRAGD | NA | -0.307845394 | 0.118256641 |
| CLPX | NA | -0.307592531 | 0.118574779 |
| ACADSB | NA | -0.307402856 | 0.118813832 |
| RCSD1 | NA | -0.306310321 | 0.120197689 |
| SYAP1 | NA | -0.30627079 | 0.120247982 |
| NPIP | NA | -0.305541554 | 0.121178519 |
| NLRP3 | NA | -0.305363758 | 0.121406193 |
| AP2A1 | NA | -0.305338966 | 0.121437965 |
| SERTAD2 | NA | -0.303203443 | 0.124197689 |
| TRIP12 | NA | -0.302170658 | 0.125548699 |
| TOR1AIP2 | NA | -0.302123967 | 0.125610029 |
| C10orf54 | NA | -0.301511291 | 0.126416834 |
| MME | NA | -0.301480891 | 0.126456964 |
| KIAA1731 | NA | -0.300229587 | 0.128116883 |
| HIVEP3 | NA | -0.299231757 | 0.129451902 |
| AAMDC | NA | -0.298791927 | 0.13004357 |
| PRCP | NA | -0.298607544 | 0.130292189 |
| ZNF702P | NA | -0.298415811 | 0.130551086 |
| TBCD | NA | -0.298248264 | 0.130777632 |
| NAA40 | NA | -0.298176997 | 0.130874082 |
| FBRS | NA | -0.297898947 | 0.131250874 |
| SPTBN1 | NA | -0.297667014 | 0.131565776 |
| BMS1 | NA | -0.297577788 | 0.131687067 |
| ZNF697 | NA | -0.297456164 | 0.131852529 |
| CXCL3 | NA | -0.296754002 | 0.132810738 |
| FUZ | NA | -0.296753156 | 0.132811895 |
| LOC1001323 | NA | -0.296659055 | 0.132940695 |
| SMG9 | NA | -0.296273932 | 0.133468772 |
| REV3L | NA | -0.296255728 | 0.133493771 |
| ZNF8 | NA | -0.295739345 | 0.134204316 |
| POLR2J2 | NA | -0.295204031 | 0.134943803 |
| EIF2B3 | NA | -0.293765155 | 0.136946113 |
| PLAGL2 | NA | -0.293583125 | 0.137200946 |
| CPPED1 | NA | -0.292959344 | 0.138076814 |
| FAM214B | NA | -0.292898611 | 0.138162306 |
| CCNJ | NA | -0.292485631 | 0.138744662 |
| RANBP1 | NA | -0.291443016 | 0.140222775 |
| GLT8D1 | NA | -0.291321971 | 0.140395114 |
| GATA3 | NA | -0.291119872 | 0.140683196 |
| RCL1 | NA | -0.290960608 | 0.140910517 |
| PPRC1 | NA | -0.290894032 | 0.141005622 |
| MSL1 | NA | -0.290820782 | 0.141110313 |
| REPS1 | NA | -0.290217718 | 0.141974367 |
| RAP1A | NA | -0.289650343 | 0.14279076 |
| MAPK14 | NA | -0.289368018 | 0.143198254 |
| SCRN3 | NA | -0.288224164 | 0.144857804 |
| TRIM21 | NA | -0.288079131 | 0.145069208 |
| TMEM176A | NA | -0.28740443 | 0.146055582 |
| SNX24 | NA | -0.287341971 | 0.146147137 |
| LRRC6 | NA | -0.286692329 | 0.147101848 |
| ECHDC2 | NA | -0.286683671 | 0.147114601 |
| KIAA1279 | NA | -0.28634993 | 0.147606832 |
| MIR155HG | NA | -0.286211534 | 0.147811297 |
| TMEM127 | grey60 | 0.024408 | 0.905787 |
| ANKRD40 | grey60 | 0.031548 | 0.878405 |
| CTAGE5 | grey60 | 0.035069 | 0.86495 |
| ARRB2 | grey60 | 0.036817 | 0.858283 |
| ZNF81 | grey60 | 0.046981 | 0.819724 |
| SMU1 | grey60 | 0.049879 | 0.808797 |
| SRPR | grey60 | 0.056751 | 0.783035 |
| ZNF426 | grey60 | 0.056767 | 0.782975 |
| TOPBP1 | grey60 | 0.070468 | 0.732296 |
| NAPG | grey60 | 0.071292 | 0.729281 |
| ZNF259 | grey60 | 0.078278 | 0.703874 |
| CEP97 | grey60 | 0.078291 | 0.703828 |
| ATF7IP | grey60 | 0.080627 | 0.695399 |
| CLCN5 | grey60 | 0.092867 | 0.651833 |
| RHOG | grey60 | 0.093297 | 0.65032 |
| EHBP1L1 | grey60 | 0.094868 | 0.644808 |
| LAMB1 | grey60 | 0.096285 | 0.639854 |
| KIAA0556 | grey60 | 0.096504 | 0.639088 |
| ZMIZ2 | grey60 | 0.09824 | 0.633043 |
| MED7 | grey60 | 0.101478 | 0.621824 |
| RAB11FIP5 | grey60 | 0.101947 | 0.620204 |
| KIAA0355 | grey60 | 0.108534 | 0.597663 |
| TMEM120A | grey60 | 0.111384 | 0.588021 |
| TECR | grey60 | 0.118376 | 0.56465 |
| IKZF2 | grey60 | 0.128277 | 0.532293 |
| SCAP | grey60 | 0.128428 | 0.531809 |
| GRK6 | grey60 | 0.138991 | 0.498303 |
| GLB1 | grey60 | 0.146007 | 0.476648 |
| ASXL1 | grey60 | 0.148715 | 0.46842 |
| DOCK1 | grey60 | 0.157553 | 0.442086 |
| ZXDC | grey60 | 0.158518 | 0.43926 |
| PNPLA2 | grey60 | 0.163029 | 0.426177 |
| TMEM110 | grey60 | 0.163049 | 0.426119 |
| SMCR7L | grey60 | 0.174304 | 0.394428 |
| CRTAP | grey60 | 0.178077 | 0.384111 |
| SLC35E1 | grey60 | 0.17918 | 0.381124 |
| MASTL | grey60 | 0.18204 | 0.373443 |
| CHD4 | grey60 | 0.182164 | 0.373111 |
| CTCF | grey60 | 0.182308 | 0.372728 |
| UBXN7 | grey60 | 0.184284 | 0.367478 |
| RAD21 | grey60 | 0.187488 | 0.359061 |
| PLXNB2 | grey60 | 0.194513 | 0.341004 |
| GPATCH1 | grey60 | 0.198777 | 0.330316 |
| ALG10 | grey60 | 0.221404 | 0.277045 |
| KBTBD7 | grey60 | 0.225769 | 0.267439 |
| B4GALT4 | grey60 | 0.230719 | 0.256809 |
| ADPGK | grey60 | 0.231046 | 0.256116 |
| RBM6 | grey60 | 0.24239 | 0.232847 |
| ZNF281 | grey60 | 0.242568 | 0.232492 |
| RPRD1B | grey60 | 0.250043 | 0.217967 |
| TOR1AIP1 | grey60 | 0.253699 | 0.211089 |
| TMEM87A | grey60 | 0.265126 | 0.190545 |
| MCM5 | grey60 | 0.344838 | 0.084486 |
| CMC4 | lightcyan | -0.47031 | 0.015325 |
| UQCR10 | lightcyan | -0.438199 | 0.025149 |
| LRRC37B | lightcyan | -0.42078 | 0.03231 |

**Table 6. Gene module membership and correlation with growth rate (48/72h), with adjusted p values for each module**

| | | | | | | | | | | | |
|---|---|---|---|---|---|---|---|---|---|---|---|
| DYNLL1 | lightgreen | -0.318491 | 0.112799 | DBF4B | NA | -0.286139748 | 0.147917433 | GPN1 | lightcyan | -0.366848 | 0.065265 |
| NBN | lightgreen | -0.312421 | 0.120212 | CD300E | NA | -0.286109954 | 0.147961499 | ISY1 | lightcyan | -0.345588 | 0.083768 |
| C4orf33 | lightgreen | -0.302542 | 0.13303 | TIMP3 | NA | -0.285596183 | 0.14872287 | ZNF702P | lightcyan | -0.343558 | 0.085722 |
| MRPS6 | lightgreen | 0.297017 | 0.140616 | SSBP4 | NA | -0.285527249 | 0.148825236 | HSD17B10 | lightcyan | -0.337508 | 0.091755 |
| MOCS3 | lightgreen | -0.293911 | 0.145015 | ARHGAP9 | NA | -0.284846487 | 0.149838891 | RPL36AL | lightcyan | -0.331432 | 0.098129 |
| HK2 | lightgreen | 0.288418 | 0.153035 | AKAP12 | NA | -0.284176988 | 0.150840587 | ZNF28 | lightcyan | -0.322522 | 0.108066 |
| SLC25A44 | lightgreen | 0.286595 | 0.155765 | PILRB | NA | -0.283908231 | 0.15124404 | PHF5A | lightcyan | -0.322143 | 0.108506 |
| SLC45A4 | lightgreen | 0.284986 | 0.158203 | CYB561D2 | NA | -0.283546358 | 0.151788496 | PLEKHO1 | lightcyan | -0.317673 | 0.113778 |
| CHAC1 | lightgreen | 0.282321 | 0.1623 | LSM14B | NA | -0.282539597 | 0.153310584 | PPIG | lightcyan | -0.31546 | 0.116458 |
| RCC1 | lightgreen | 0.280726 | 0.164787 | CHD6 | NA | -0.28166361 | 0.154643794 | SMYD3 | lightcyan | -0.312163 | 0.120536 |
| TUBB2A | lightgreen | 0.266602 | 0.187997 | SMOX | NA | -0.281417023 | 0.155020574 | C17orf80 | lightcyan | -0.307117 | 0.126976 |
| PIM2 | lightgreen | 0.26649 | 0.188188 | XYLT1 | NA | -0.279794104 | 0.157516689 | LOC285074 | lightcyan | -0.304734 | 0.130104 |
| GLCCI1 | lightgreen | 0.256745 | 0.205473 | NBR2 | NA | -0.279350779 | 0.158203479 | TPI1 | lightcyan | -0.304379 | 0.130575 |
| MRPS17 | lightgreen | -0.237516 | 0.242666 | AOC3 | NA | -0.279223471 | 0.158401097 | CREB3 | lightcyan | -0.30058 | 0.135689 |
| FAM117B | lightgreen | -0.236962 | 0.243799 | RNF34 | NA | -0.278867999 | 0.158953812 | RPL36 | lightcyan | -0.29955 | 0.1371 |
| FAN1 | lightgreen | 0.236867 | 0.243994 | CEP85 | NA | -0.278839776 | 0.158997754 | PARP8 | lightcyan | -0.295616 | 0.142588 |
| DBN1 | lightgreen | 0.226736 | 0.26534 | CYCS | NA | -0.277847873 | 0.160547588 | SREK1 | lightcyan | -0.293563 | 0.145515 |
| BCL2L11 | lightgreen | 0.226343 | 0.266193 | ABHD11 | NA | -0.277644235 | 0.16086709 | BST1 | lightcyan | -0.28443 | 0.159051 |
| MKNK2 | lightgreen | 0.222858 | 0.273821 | C2CD2 | NA | -0.277341138 | 0.161343474 | WBP1 | lightcyan | -0.282968 | 0.161299 |
| UNC13B | lightgreen | 0.2225 | 0.274612 | CXCL5 | NA | -0.277275693 | 0.161446466 | RNF185 | lightcyan | -0.282345 | 0.162263 |
| PCSK5 | lightgreen | -0.211651 | 0.299293 | CYBASC3 | NA | -0.276431205 | 0.162779641 | GNPTG | lightcyan | -0.281394 | 0.163742 |
| KLF2 | lightgreen | -0.20446 | 0.316388 | ZNF615 | NA | -0.276219076 | 0.163115745 | C1orf52 | lightcyan | -0.269151 | 0.18365 |
| TFPI2 | lightgreen | -0.202376 | 0.321453 | NKAP | NA | -0.276176632 | 0.163183053 | VRK1 | lightcyan | -0.268245 | 0.185187 |
| ZSWIM6 | lightgreen | 0.202246 | 0.321769 | CMAS | NA | -0.275802852 | 0.163776649 | RENBP | lightcyan | -0.266052 | 0.188944 |
| NEFH | lightgreen | -0.196863 | 0.335088 | EPAS1 | NA | -0.27568544 | 0.163963425 | GCDH | lightcyan | -0.258531 | 0.202227 |
| PCID2 | lightgreen | -0.191495 | 0.348693 | SRP68 | NA | -0.275496836 | 0.164263766 | NPL | lightcyan | -0.25735 | 0.20437 |
| FAM84B | lightgreen | 0.190141 | 0.352176 | COL9A2 | NA | -0.27361456 | 0.167282508 | DVL2 | lightcyan | -0.256908 | 0.205175 |
| TBX20 | lightgreen | 0.184764 | 0.36621 | TGFBI | NA | -0.272944646 | 0.168366274 | DDX59 | lightcyan | -0.249565 | 0.218876 |
| PCGF6 | lightgreen | -0.175057 | 0.392355 | ALG1 | NA | -0.272426873 | 0.169207292 | ZNF816 | lightcyan | -0.236966 | 0.243792 |
| EIF4A3 | lightgreen | -0.174619 | 0.393561 | TSEN34 | NA | -0.272305001 | 0.169405677 | FCGRT | lightcyan | -0.225023 | 0.269066 |
| TUBG2 | lightgreen | -0.159095 | 0.437575 | GGH | NA | -0.272109752 | 0.169723848 | ZNF92 | lightcyan | -0.222834 | 0.273874 |
| RAD51 | lightgreen | -0.157024 | 0.443641 | LAPTM4A | NA | -0.272097265 | 0.169744212 | SF3B2 | lightcyan | -0.221128 | 0.277659 |
| IKBIP | lightgreen | -0.156336 | 0.445665 | LAD1 | NA | -0.271399522 | 0.170884761 | SNHG7 | lightcyan | -0.219783 | 0.280667 |
| PNP | lightgreen | 0.152239 | 0.457824 | AP5B1 | NA | -0.270768868 | 0.171920271 | PNPLA4 | lightcyan | -0.219778 | 0.280679 |
| TSC22D3 | lightgreen | 0.141667 | 0.489986 | SRM | NA | -0.270476285 | 0.172402174 | SYNGAP1 | lightcyan | -0.215476 | 0.290437 |
| CD83 | lightgreen | -0.139015 | 0.498231 | ZFHX2 | NA | -0.270179097 | 0.172892632 | ACADSB | lightcyan | -0.210327 | 0.302395 |
| ATP1A1 | lightgreen | 0.135732 | 0.508528 | C17orf70 | NA | -0.268491967 | 0.175695504 | C9orf114 | lightcyan | -0.203235 | 0.319359 |
| BSCL2 | lightgreen | -0.134651 | 0.511942 | BHLHE40 | NA | -0.267251624 | 0.177776298 | TKT | lightcyan | -0.201005 | 0.324811 |
| DUSP28 | lightgreen | -0.132655 | 0.518274 | ZYG11B | NA | -0.266586562 | 0.178899066 | RARA | lightcyan | -0.200258 | 0.32665 |
| COIL | lightgreen | -0.131891 | 0.520708 | ZNF24 | NA | -0.266115954 | 0.179696537 | KBTBD4 | lightcyan | -0.195452 | 0.338632 |
| DLX2 | lightgreen | 0.13027 | 0.525889 | CEP85L | NA | -0.26504854 | 0.181514508 | ABHD14A | lightcyan | -0.192763 | 0.345451 |
| GADD45B | lightgreen | 0.12445 | 0.544697 | SRPR | NA | -0.264519869 | 0.182419643 | OTUB1 | lightcyan | -0.189651 | 0.353443 |
| ZNF227 | lightgreen | -0.122662 | 0.550535 | E4F1 | NA | -0.264273114 | 0.182843184 | AES | lightcyan | -0.189347 | 0.354227 |
| PTCD3 | lightgreen | -0.122278 | 0.551793 | ARID5A | NA | -0.264082516 | 0.183170803 | PLEKHB1 | lightcyan | -0.188511 | 0.356395 |
| TUBB3 | lightgreen | -0.122025 | 0.552623 | NDRG2 | NA | -0.263486967 | 0.184197121 | MPP5 | lightcyan | -0.179429 | 0.380453 |
| BAMBI | lightgreen | 0.117933 | 0.56612 | ERN1 | NA | -0.263101195 | 0.184864056 | NCF4 | lightcyan | -0.178829 | 0.382074 |
| DUSP2 | lightgreen | -0.117041 | 0.569079 | TXLNG | NA | -0.262854796 | 0.185290914 | CDK9 | lightcyan | -0.160303 | 0.434056 |
| DUSP7 | lightgreen | 0.113066 | 0.582359 | CNBP | NA | -0.262057073 | 0.186677571 | MAPK3 | lightcyan | -0.159309 | 0.43695 |
| HDGFRP3 | lightgreen | -0.111093 | 0.589002 | SUCLG1 | NA | -0.261799081 | 0.187127565 | COX10 | lightcyan | -0.151251 | 0.460782 |
| PACRGL | lightgreen | 0.108357 | 0.598266 | GPR141 | NA | -0.261296245 | 0.188006781 | PCYOX1 | lightcyan | -0.150024 | 0.464469 |
| MRPL37 | lightgreen | -0.107722 | 0.600425 | ZNF197 | NA | -0.260596154 | 0.189235656 | HAX1 | lightcyan | -0.149395 | 0.466365 |
| ARC | lightgreen | 0.102652 | 0.617777 | USP16 | NA | -0.260585438 | 0.189254508 | FOXN2 | lightcyan | -0.147021 | 0.473558 |
| UBE2S | lightgreen | -0.100881 | 0.623885 | COG3 | NA | -0.260036869 | 0.190221357 | REPIN1 | lightcyan | -0.139707 | 0.49607 |
| OSGIN2 | lightgreen | 0.096959 | 0.637501 | ZNF816 | NA | -0.258959833 | 0.192129543 | FLI1 | lightcyan | -0.138935 | 0.498479 |
| SPSB1 | lightgreen | 0.095892 | 0.641227 | MEAF6 | NA | -0.258824143 | 0.192370879 | TOP1MT | lightcyan | -0.128917 | 0.530234 |
| TUBA4A | lightgreen | -0.091061 | 0.658194 | CCDC134 | NA | -0.258243433 | 0.193406079 | PLA2G4B | lightcyan | -0.124808 | 0.543529 |

**Table 6. Gene module membership and correlation with growth rate (48/72h), with adjusted p values for each module**

| | | | | | | | | | | | |
|---|---|---|---|---|---|---|---|---|---|---|---|
| MAPKAPK2 | lightgreen | -0.091048 | 0.658239 | RTCA | NA | -0.2576226 | 0.194517043 | KCNE3 | lightcyan | -0.123446 | 0.547972 |
| FAM46C | lightgreen | 0.087053 | 0.6724 | KIAA0232 | NA | -0.256740688 | 0.196102744 | TOM1 | lightcyan | -0.115279 | 0.574951 |
| ZNF296 | lightgreen | -0.085908 | 0.676477 | GPR132 | NA | -0.256290226 | 0.196916106 | RCAN3 | lightcyan | -0.113145 | 0.582094 |
| KPNA2 | lightgreen | 0.085673 | 0.677313 | CBWD1 | NA | -0.254916098 | 0.199411566 | STK11 | lightcyan | -0.081212 | 0.693294 |
| LYL1 | lightgreen | -0.080056 | 0.697457 | SRSF9 | NA | -0.254562581 | 0.200057054 | SRA1 | lightcyan | -0.076901 | 0.708859 |
| EPAS1 | lightgreen | -0.079114 | 0.700855 | SLC12A2 | NA | -0.254240005 | 0.200647294 | ZNF558 | lightcyan | -0.072257 | 0.725755 |
| SCPEP1 | lightgreen | -0.079086 | 0.700955 | UQCR11 | NA | -0.252933962 | 0.203049242 | PPP1R13B | lightcyan | -0.070952 | 0.730525 |
| MPHOSPH1( | lightgreen | 0.078747 | 0.70218 | DCAF7 | NA | -0.252452923 | 0.203938853 | MLL5 | lightcyan | -0.057516 | 0.780181 |
| MRPL36 | lightgreen | -0.078303 | 0.703786 | PIP4K2A | NA | -0.252368843 | 0.204094618 | MRS2 | lightcyan | -0.034034 | 0.868902 |
| MARCKSL1 | lightgreen | 0.073187 | 0.722361 | CNRIP1 | NA | -0.252247534 | 0.204319499 | EML3 | lightcyan | -0.014597 | 0.943578 |
| EPHA2 | lightgreen | -0.071993 | 0.726718 | PPIL4 | NA | -0.25172032 | 0.2052988 | MRM1 | lightcyan | -0.009732 | 0.962368 |
| GRPEL2 | lightgreen | -0.07186 | 0.727206 | MAN1A2 | NA | -0.251435537 | 0.205829115 | TWSG1 | lightcyan | 0.047944 | 0.816088 |
| MDC1 | lightgreen | 0.068357 | 0.740042 | SHROOM3 | NA | -0.251091053 | 0.20647185 | UBR2 | lightcyan | 0.07262 | 0.72443 |
| FZD8 | lightgreen | -0.066857 | 0.745557 | QPCT | NA | -0.251090207 | 0.206473432 | GLS | lightcyan | 0.075826 | 0.712757 |
| LBH | lightgreen | 0.062969 | 0.759914 | RASAL2 | NA | -0.250748506 | 0.207112327 | PIK3CB | lightcyan | 0.076035 | 0.712001 |
| NDRG1 | lightgreen | 0.059995 | 0.77095 | CTNNB1 | NA | -0.25060182 | 0.207387006 | LDHAL6B | lightcyan | 0.078421 | 0.703358 |
| PDIA6 | lightgreen | -0.058846 | 0.775224 | ZNF565 | NA | -0.250244502 | 0.208057144 | BTAF1 | lightcyan | 0.079437 | 0.699688 |
| TRIM24 | lightgreen | 0.057677 | 0.77958 | CABIN1 | NA | -0.250147322 | 0.208239657 | CCDC90A | lightcyan | 0.097314 | 0.636265 |
| F11R | lightgreen | -0.053789 | 0.794115 | MTHFSD | NA | -0.249915737 | 0.208675034 | C5orf25 | lightcyan | 0.105656 | 0.607469 |
| CEBPZ | lightgreen | -0.044757 | 0.828128 | MMP19 | NA | -0.248716655 | 0.210939185 | PLCD1 | lightcyan | 0.118923 | 0.56284 |
| IQSEC3 | lightgreen | 0.043793 | 0.83178 | ZNF84 | NA | -0.248438711 | 0.211466381 | LINC00622 | lightcyan | 0.119127 | 0.562165 |
| ARF3 | lightgreen | -0.039471 | 0.848181 | MRPS11 | NA | -0.24835724 | 0.211621082 | ATIC | lightcyan | 0.130744 | 0.524372 |
| FAM117A | lightgreen | -0.039399 | 0.848453 | CD48 | NA | -0.247709034 | 0.212854661 | EIF4EBP2 | lightcyan | 0.131425 | 0.522195 |
| ADRB2 | lightgreen | 0.038115 | 0.853339 | ELK3 | NA | -0.247556618 | 0.213145425 | FBXL17 | lightcyan | 0.146059 | 0.47649 |
| CSRNP2 | lightgreen | -0.037167 | 0.856951 | LENG8 | NA | -0.247456852 | 0.213335893 | GDI2 | lightcyan | 0.158978 | 0.437915 |
| C2orf47 | lightgreen | -0.035418 | 0.863619 | MDN1 | NA | -0.247273416 | 0.213686402 | RPS6KA3 | lightcyan | 0.181133 | 0.375868 |
| DYNLL2 | lightgreen | -0.03438 | 0.867581 | MEGF9 | NA | -0.246533584 | 0.215104028 | SLC20A1 | lightcyan | 0.187392 | 0.35931 |
| ABTB2 | lightgreen | 0.034187 | 0.868319 | RHOG | NA | -0.246471474 | 0.215223329 | XPO7 | lightcyan | 0.196187 | 0.336783 |
| ZC3H12A | lightgreen | -0.032452 | 0.87495 | PLCG1 | NA | -0.246416157 | 0.215329618 | SNRPA1 | lightcyan | 0.198168 | 0.331828 |
| DNAJB1 | lightgreen | -0.030441 | 0.882643 | MAP3K10 | NA | -0.246339069 | 0.215477799 | MRC2 | lightcyan | 0.198999 | 0.329765 |
| KIAA0947 | lightgreen | -0.026895 | 0.896238 | BBS7 | NA | -0.246312114 | 0.21552963 | CAST | lightcyan | 0.200769 | 0.325393 |
| DDIT4 | lightgreen | 0.025217 | 0.902679 | SCO1 | NA | -0.245886854 | 0.216348454 | RFFL | lightcyan | 0.202578 | 0.320961 |
| EFNB1 | lightgreen | 0.023065 | 0.910949 | ACAA2 | NA | -0.245690318 | 0.216727587 | ACOT7 | lightcyan | 0.203284 | 0.319239 |
| DUSP10 | lightgreen | 0.021324 | 0.917648 | ZNRF1 | NA | -0.245062598 | 0.217941509 | TMLHE | lightcyan | 0.213145 | 0.295813 |
| HBEGF | lightgreen | 0.01613 | 0.937665 | CRYZ | NA | -0.244959472 | 0.218141378 | PYROXD1 | lightcyan | 0.218326 | 0.28395 |
| RAB7A | lightgreen | 0.014234 | 0.944979 | GUSBP3 | NA | -0.244647329 | 0.218747095 | HSD3BP4 | lightcyan | 0.223148 | 0.273182 |
| BIK | lightgreen | 0.013859 | 0.946428 | FBXL20 | NA | -0.243814713 | 0.220368342 | LOC1001314( | lightcyan | 0.236517 | 0.244712 |
| P2RX1 | lightgreen | 0.011717 | 0.954697 | SPATA5 | NA | -0.24353125 | 0.220922134 | KCTD20 | lightcyan | 0.236647 | 0.244445 |
| KLF4 | lightgreen | -0.009125 | 0.964714 | PSMC2 | NA | -0.243179028 | 0.221611561 | THOC5 | lightcyan | 0.241207 | 0.235204 |
| ULBP1 | lightgreen | 0.006528 | 0.974753 | PPARD | NA | -0.243038717 | 0.221886605 | MPZL3 | lightcyan | 0.243444 | 0.230758 |
| PFKFB3 | lightgreen | -0.004908 | 0.981015 | CNOT1 | NA | -0.241181273 | 0.225549262 | OR5B21 | lightcyan | 0.244279 | 0.229112 |
| EGR1 | lightgreen | 0.002748 | 0.98937 | TMEM201 | NA | -0.2409002 | 0.226107011 | PHF23 | lightcyan | 0.251623 | 0.214977 |
| DUSP5 | lightgreen | 7.39E-05 | 0.999714 | PMVK | NA | -0.240496279 | 0.22691015 | SNORA70B | lightcyan | 0.25278 | 0.212803 |
| SERGEF | lightyellow | -0.569905 | 0.002371 | GCNT1 | NA | -0.239893669 | 0.228111901 | SNX4 | lightcyan | 0.262177 | 0.19571 |
| AMIGO2 | lightyellow | -0.534349 | 0.004923 | ERVK13-1 | NA | -0.239370538 | 0.229158594 | RNF31 | lightcyan | 0.266412 | 0.188322 |
| CDC42SE2 | lightyellow | -0.467115 | 0.016132 | ASCC1 | NA | -0.23874503 | 0.230414331 | THOC1 | lightcyan | 0.277232 | 0.170331 |
| SNX1 | lightyellow | -0.4319 | 0.027573 | PGK1 | NA | -0.238667949 | 0.230569391 | CSNK2A2 | lightcyan | 0.280083 | 0.165799 |
| WDR61 | lightyellow | -0.422588 | 0.031498 | MTMR2 | NA | -0.238258347 | 0.231394538 | RPL23AP64 | lightcyan | 0.287866 | 0.153858 |
| CYB5A | lightyellow | -0.372592 | 0.060854 | BSCL2 | NA | -0.237615508 | 0.232693509 | MYOF | lightcyan | 0.291312 | 0.148771 |
| ETHE1 | lightyellow | -0.370661 | 0.06231 | ZC3H11A | NA | -0.236394924 | 0.235173252 | LOC1003350: | lightcyan | 0.304002 | 0.131076 |
| TMSB4X | lightyellow | -0.364858 | 0.06685 | PIK3CD | NA | -0.236300704 | 0.235365397 | SLC5A3 | lightcyan | 0.30846 | 0.125238 |
| RNASEK | lightyellow | -0.342407 | 0.086847 | SSX1 | NA | -0.235691162 | 0.236610966 | POLR2A | lightcyan | 0.309862 | 0.123442 |
| FAM173A | lightyellow | -0.328475 | 0.101348 | BCL6 | NA | -0.234510532 | 0.239035945 | KDELR1 | lightcyan | 0.311258 | 0.121672 |
| PSME2 | lightyellow | -0.319655 | 0.111417 | AKNA | NA | -0.234458038 | 0.239144148 | FLJ42627 | lightcyan | 0.311888 | 0.12088 |
| OTUD4 | lightyellow | -0.317875 | 0.113536 | STX17 | NA | -0.233959547 | 0.240173265 | ASCC3 | lightcyan | 0.314431 | 0.11772 |
| PIN1 | lightyellow | -0.317201 | 0.114346 | TMEM163 | NA | -0.233915032 | 0.240265306 | SOAT1 | lightcyan | 0.317038 | 0.114542 |

**Table 6. Gene module membership and correlation with growth rate (48/72h), with adjusted p values for each module**

| | | | | | | | | | | | |
|---|---|---|---|---|---|---|---|---|---|---|---|
| ANKRD44 | lightyellow | -0.301851 | 0.133963 | TGFA | NA | -0.233344079 | 0.241447909 | GAPDH | lightcyan | 0.321518 | 0.109232 |
| NDUFB7 | lightyellow | -0.300627 | 0.135625 | DCTN5 | NA | -0.233104961 | 0.241944329 | NDUFAF4 | lightcyan | 0.32244 | 0.108161 |
| PFDN6 | lightyellow | -0.293218 | 0.14601 | C5orf42 | NA | -0.233074795 | 0.242007004 | EID3 | lightcyan | 0.324948 | 0.10529 |
| H2AFY | lightyellow | -0.283197 | 0.160945 | TMEM14B | NA | -0.23300635 | 0.242149247 | PTPLB | lightcyan | 0.325145 | 0.105067 |
| SIL1 | lightyellow | -0.278338 | 0.168562 | TBC1D17 | NA | -0.23110602 | 0.246120588 | FXR2 | lightcyan | 0.331119 | 0.098466 |
| LOC645638 | lightyellow | -0.271878 | 0.179076 | SNAPC4 | NA | -0.231101278 | 0.246130552 | MORC2 | lightcyan | 0.334506 | 0.094864 |
| MAVS | lightyellow | -0.269925 | 0.182343 | MRPL49 | NA | -0.23107842 | 0.246178581 | GPD2 | lightcyan | 0.335494 | 0.093833 |
| NDUFA6 | lightyellow | -0.267221 | 0.186934 | JUP | NA | -0.230255252 | 0.24791234 | STX4 | lightcyan | 0.338448 | 0.090797 |
| C17orf62 | lightyellow | -0.251318 | 0.21555 | UCKL1 | NA | -0.230205976 | 0.248016378 | AKIRIN1 | lightcyan | 0.342776 | 0.086485 |
| C18orf8 | lightyellow | -0.247293 | 0.223238 | NAV1 | NA | -0.230135809 | 0.248164576 | ZNF430 | lightcyan | 0.345253 | 0.084088 |
| GNS | lightyellow | -0.239366 | 0.238907 | GLT25D1 | NA | -0.229945857 | 0.248566057 | KHNYN | lightcyan | 0.345326 | 0.084018 |
| PSMC2 | lightyellow | -0.238196 | 0.24128 | CYP27B1 | NA | -0.22899028 | 0.250592221 | COASY | lightcyan | 0.346832 | 0.082586 |
| PHB | lightyellow | -0.236919 | 0.243887 | EPB41L3 | NA | -0.228867032 | 0.250854336 | PARP6 | lightcyan | 0.358433 | 0.072168 |
| NUDT16L1 | lightyellow | -0.223474 | 0.272464 | VPS13A | NA | -0.228736294 | 0.251132578 | EIF4G1 | lightcyan | 0.363209 | 0.068185 |
| TMEM134 | lightyellow | -0.220806 | 0.278379 | SLTM | NA | -0.22823843 | 0.252193997 | SMARCD2 | lightcyan | 0.36683 | 0.065279 |
| NOP2 | lightyellow | -0.209734 | 0.303791 | ADPRM | NA | -0.22769055 | 0.253365435 | CSE1L | lightcyan | 0.368687 | 0.063827 |
| NEK9 | lightyellow | -0.199055 | 0.329626 | STON1 | NA | -0.227382138 | 0.25402642 | FBXW2 | lightcyan | 0.378271 | 0.056723 |
| CISD2 | lightyellow | -0.192141 | 0.347039 | CYTH4 | NA | -0.227372999 | 0.254046025 | PDE3B | lightcyan | 0.382244 | 0.053965 |
| TMED4 | lightyellow | -0.188596 | 0.356176 | ADD3 | NA | -0.227117365 | 0.254594779 | MAP4K3 | lightcyan | 0.399402 | 0.043237 |
| AP5B1 | lightyellow | 0.186184 | 0.362473 | TSPAN14 | NA | -0.225687457 | 0.257678532 | VPS4A | lightcyan | 0.40154 | 0.042028 |
| DNAJA4 | lightyellow | 0.183542 | 0.369445 | CXorf21 | NA | -0.223221231 | 0.263054121 | RPL13AP6 | lightcyan | 0.413608 | 0.035696 |
| RNF11 | lightyellow | 0.176166 | 0.389318 | DISC1 | NA | -0.223124689 | 0.263266017 | VCP | lightcyan | 0.421835 | 0.031834 |
| MLST8 | lightyellow | -0.170996 | 0.4036 | ALDH3A2 | NA | -0.222738638 | 0.264114453 | RASGEF1B | lightcyan | 0.427538 | 0.029359 |
| CXCL16 | lightyellow | -0.167437 | 0.413601 | TMEM11 | NA | -0.222699195 | 0.264201237 | LUC7L | lightcyan | 0.456653 | 0.019023 |
| MRPS24 | lightyellow | -0.165052 | 0.420381 | NCF4 | NA | -0.222605517 | 0.264407427 | SCAMP4 | lightcyan | 0.456766 | 0.01899 |
| AIP | lightyellow | -0.162065 | 0.428954 | TNFRSF10C | NA | -0.222192283 | 0.265318215 | SLC5A6 | lightcyan | 0.473153 | 0.014635 |
| NDUFB11 | lightyellow | -0.161431 | 0.430788 | CIDECP | NA | -0.222167869 | 0.265372087 | RTF1 | lightcyan | 0.47399 | 0.014437 |
| NUP43 | lightyellow | -0.158301 | 0.439896 | MFAP5 | NA | -0.221496513 | 0.266856303 | MYL9 | lightgreen | -0.257737 | 0.203665 |
| ARL5B | lightyellow | -0.157272 | 0.44291 | PTPN7 | NA | -0.221455585 | 0.266946958 | MREG | lightgreen | -0.257037 | 0.204938 |
| CLEC5A | lightyellow | -0.151719 | 0.459379 | LOC1005062 | NA | -0.221138299 | 0.267650419 | FCGR1A | lightgreen | -0.247075 | 0.223659 |
| QTRT1 | lightyellow | -0.14982 | 0.465083 | FAM102B | NA | -0.220667988 | 0.268695354 | PTRF | lightgreen | -0.246805 | 0.224182 |
| CD40 | lightyellow | -0.145292 | 0.478832 | SUSD3 | NA | -0.220603284 | 0.268839318 | BATF | lightgreen | -0.24569 | 0.226349 |
| NHP2 | lightyellow | -0.144583 | 0.481004 | DHRS7B | NA | -0.219319405 | 0.271706185 | TTF2 | lightgreen | -0.230117 | 0.258088 |
| C20orf111 | lightyellow | 0.135761 | 0.508437 | METTL16 | NA | -0.218915706 | 0.272611679 | C1orf54 | lightgreen | -0.217681 | 0.285408 |
| PNPO | lightyellow | -0.118104 | 0.565552 | LRBA | NA | -0.218641978 | 0.27322675 | TUBG1 | lightgreen | -0.204507 | 0.316274 |
| FCF1 | lightyellow | -0.114304 | 0.57821 | DPP4 | NA | -0.218411241 | 0.273745909 | ATP6V0D2 | lightgreen | -0.199628 | 0.328205 |
| CDK4 | lightyellow | 0.108525 | 0.597693 | TBL1XR1 | NA | -0.218273626 | 0.274055847 | FBP1 | lightgreen | -0.198821 | 0.330205 |
| PWP1 | lightyellow | -0.102677 | 0.61769 | LRSAM1 | NA | -0.218161221 | 0.274309172 | SSBP3 | lightgreen | -0.187453 | 0.359152 |
| )C1006309 | lightyellow | -0.102524 | 0.618217 | CPSF6 | NA | -0.217676063 | 0.275404285 | LYSMD2 | lightgreen | -0.18745 | 0.359159 |
| POLR2E | lightyellow | -0.101125 | 0.623044 | LOC1005057 | NA | -0.21694003 | 0.277071019 | DPYSL3 | lightgreen | -0.185461 | 0.364372 |
| SNORA71A | lightyellow | -0.098636 | 0.631665 | TSPAN13 | NA | -0.214534425 | 0.282563363 | SAMSN1 | lightgreen | -0.184994 | 0.365602 |
| PKD2 | lightyellow | 0.095466 | 0.642718 | ZNF12 | NA | -0.214340422 | 0.283009296 | MCOLN3 | lightgreen | -0.182176 | 0.373078 |
| ARFGAP1 | lightyellow | 0.094816 | 0.644992 | XPNPEP1 | NA | -0.214339471 | 0.283011482 | TLCD2 | lightgreen | -0.158727 | 0.438649 |
| IPCEF1 | lightyellow | -0.089506 | 0.663692 | SNX21 | NA | -0.214152469 | 0.283441749 | S100A9 | lightgreen | -0.157568 | 0.442042 |
| GRAMD1A | lightyellow | -0.089399 | 0.664073 | GSTZ1 | NA | -0.213976392 | 0.283847258 | NCAPD3 | lightgreen | -0.156092 | 0.446385 |
| AGAP3 | lightyellow | 0.084611 | 0.681106 | BMP2K | NA | -0.213039419 | 0.286011326 | GPR65 | lightgreen | -0.155629 | 0.447751 |
| C11orf10 | lightyellow | -0.084023 | 0.683209 | VAV2 | NA | -0.212940678 | 0.286239988 | ZNF366 | lightgreen | -0.140558 | 0.493425 |
| PIGV | lightyellow | -0.07357 | 0.720966 | PPID | NA | -0.212494935 | 0.287273677 | BTK | lightgreen | -0.139845 | 0.495642 |
| MYO9B | lightyellow | 0.072115 | 0.726272 | USP39 | NA | -0.212404388 | 0.287483946 | CECR1 | lightgreen | -0.138571 | 0.499615 |
| RILPL2 | lightyellow | -0.070925 | 0.730623 | SRD5A1 | NA | -0.211803349 | 0.288882156 | SLC37A2 | lightgreen | -0.131782 | 0.521055 |
| GPATCH2L | lightyellow | 0.063158 | 0.759217 | TMC6 | NA | -0.211593492 | 0.289371361 | TRPS1 | lightgreen | -0.119784 | 0.559995 |
| FAM199X | lightyellow | -0.060633 | 0.768579 | PPP2R3A | NA | -0.211517061 | 0.289549662 | CYP2U1 | lightgreen | -0.118617 | 0.563852 |
| RPS6KA5 | lightyellow | -0.058979 | 0.774729 | ZNF366 | NA | -0.211455383 | 0.289693597 | ANKS6 | lightgreen | -0.113003 | 0.582571 |
| SLC16A1 | lightyellow | 0.057687 | 0.779545 | DNAL4 | NA | -0.2111122 | 0.290495297 | LDLRAP1 | lightgreen | -0.111471 | 0.587728 |
| CDK2AP2 | lightyellow | 0.053024 | 0.796982 | TIMM8A | NA | -0.210923399 | 0.290936943 | CSPG4 | lightgreen | -0.108597 | 0.59745 |
| RNF138 | lightyellow | -0.05081 | 0.805296 | TBCC | NA | -0.210807783 | 0.291207605 | EHF | lightgreen | -0.107483 | 0.601238 |

**Table 6. Gene module membership and correlation with growth rate (48/72h), with adjusted p values for each module**

| | | | | | | | | | | | |
|---|---|---|---|---|---|---|---|---|---|---|---|
| C17orf85 | lightyellow | 0.049822 | 0.809012 | ACOT11 | NA | -0.210753906 | 0.291333788 | NRIP3 | lightgreen | -0.099544 | 0.628516 |
| MT1G | lightyellow | -0.049173 | 0.811456 | TDRKH | NA | -0.210350857 | 0.292278839 | RTN2 | lightgreen | -0.097477 | 0.635696 |
| CTBP2 | lightyellow | 0.046068 | 0.823171 | KYNU | NA | -0.210042202 | 0.293003865 | OSBPL8 | lightgreen | -0.092612 | 0.65273 |
| HNRNPU | lightyellow | -0.044263 | 0.829999 | GAB3 | NA | -0.209998412 | 0.293106819 | GFOD1 | lightgreen | -0.091566 | 0.656414 |
| TUBA1B | lightyellow | -0.029259 | 0.887172 | SIRPB2 | NA | -0.209807789 | 0.293555254 | ADRA2B | lightgreen | -0.090154 | 0.661398 |
| HNRNPUL2 | lightyellow | -0.028634 | 0.889567 | CUL4A | NA | -0.209796631 | 0.293581516 | TLR7 | lightgreen | -0.083485 | 0.685136 |
| ISOC1 | lightyellow | 0.028176 | 0.891322 | PSPC1 | NA | -0.209532152 | 0.294204445 | IL1RN | lightgreen | -0.077012 | 0.708456 |
| SLC39A9 | lightyellow | -0.025442 | 0.901815 | HAUS6 | NA | -0.208944933 | 0.295590496 | ANXA1 | lightgreen | -0.075948 | 0.712317 |
| PMF1 | lightyellow | -0.024954 | 0.903688 | MYPOP | NA | -0.208528126 | 0.296576799 | TP53BP2 | lightgreen | -0.06632 | 0.747537 |
| MED16 | lightyellow | -0.023796 | 0.908139 | CCDC111 | NA | -0.207793796 | 0.298319484 | MMP19 | lightgreen | -0.062934 | 0.760044 |
| MRPL14 | lightyellow | -0.02228 | 0.91397 | C15orf38 | NA | -0.207151601 | 0.299848769 | GIPC3 | lightgreen | -0.062093 | 0.763162 |
| SMIM3 | lightyellow | -0.013449 | 0.948011 | ZNF337 | NA | -0.206731675 | 0.300851402 | ST20 | lightgreen | -0.050733 | 0.805585 |
| CKAP4 | lightyellow | 0.008211 | 0.968245 | ASB7 | NA | -0.206381311 | 0.30168955 | FCGR1B | lightgreen | -0.049649 | 0.809664 |
| ABCD4 | lightyellow | 0.005874 | 0.977281 | RBM33 | NA | -0.206069285 | 0.30243721 | SLC29A3 | lightgreen | -0.04772 | 0.816935 |
| FRMD4A | lightyellow | -0.003366 | 0.986981 | GSTM4 | NA | -0.206029411 | 0.302532837 | TM4SF19 | lightgreen | -0.045059 | 0.826986 |
| PNPLA3 | lightyellow | -0.000462 | 0.998212 | TMEM216 | NA | -0.205996593 | 0.302611556 | FPR1 | lightgreen | -0.041616 | 0.840035 |
| IFT80 | magenta | -0.538691 | 0.004522 | ABTB2 | NA | -0.205684639 | 0.303360467 | FAM40B | lightgreen | -0.041187 | 0.84166 |
| GFM2 | magenta | -0.483914 | 0.012251 | LRRK1 | NA | -0.20545537 | 0.303911613 | DAPP1 | lightgreen | -0.035305 | 0.864049 |
| NTPCR | magenta | -0.478486 | 0.01341 | C12orf49 | NA | -0.205193628 | 0.304541586 | NR1H3 | lightgreen | -0.033705 | 0.87016 |
| LOC202181 | magenta | 0.444865 | 0.022775 | ZNF468 | NA | -0.204953615 | 0.305119973 | CCL24 | lightgreen | -0.03223 | 0.875798 |
| TK2 | magenta | -0.430725 | 0.028045 | RCN1 | NA | -0.204196647 | 0.306948602 | TIE1 | lightgreen | -0.026961 | 0.895983 |
| POC5 | magenta | -0.408826 | 0.038107 | TOP1MT | NA | -0.203484094 | 0.308676149 | PSME4 | lightgreen | -0.026584 | 0.897429 |
| KHDRBS3 | magenta | 0.382109 | 0.054057 | CRKL | NA | -0.203143406 | 0.309504256 | PLA2G4A | lightgreen | -0.025674 | 0.900924 |
| COQ5 | magenta | -0.378242 | 0.056743 | SLC48A1 | NA | -0.202309373 | 0.311537339 | DSP | lightgreen | -0.025159 | 0.902902 |
| MRPS11 | magenta | -0.373533 | 0.060153 | SARS2 | NA | -0.201825004 | 0.312721845 | RPRD2 | lightgreen | -0.022893 | 0.911612 |
| CCDC106 | magenta | -0.363849 | 0.067664 | FAM124A | NA | -0.201784169 | 0.312821833 | METTL1 | lightgreen | -0.021331 | 0.917623 |
| RBL1 | magenta | -0.357007 | 0.073392 | BUD31 | NA | -0.201520084 | 0.313468942 | TGM2 | lightgreen | -0.019862 | 0.92328 |
| NPC2 | magenta | -0.356426 | 0.073895 | TLK2 | NA | -0.201203883 | 0.314244844 | PODXL | lightgreen | -0.019519 | 0.9246 |
| ZNF460 | magenta | 0.356354 | 0.073957 | VNN1 | NA | -0.199843324 | 0.317596917 | ADM | lightgreen | -0.01566 | 0.939475 |
| MCEE | magenta | -0.345119 | 0.084216 | HADH | NA | -0.199648967 | 0.318077552 | GAL | lightgreen | -0.014402 | 0.944331 |
| C8orf40 | magenta | -0.342316 | 0.086936 | TTLL1 | NA | -0.199586063 | 0.318233207 | SCPEP1 | lightgreen | -0.011215 | 0.956638 |
| ZNF623 | magenta | -0.341307 | 0.087932 | ZXDC | NA | -0.1994841 | 0.318485611 | SPOCD1 | lightgreen | -0.010356 | 0.959957 |
| LAMTOR4 | magenta | -0.332257 | 0.097244 | YWHAZ | NA | -0.199319793 | 0.318892602 | HK3 | lightgreen | -0.004166 | 0.983886 |
| SAMD4B | magenta | 0.327408 | 0.102529 | FYCO1 | NA | -0.199190031 | 0.319214251 | ATP1A1 | lightgreen | -0.001107 | 0.995717 |
| ATG4B | magenta | 0.325921 | 0.104191 | PROCR | NA | -0.198613779 | 0.320645049 | EMP1 | lightgreen | 0.002311 | 0.99106 |
| MPDU1 | magenta | 0.325559 | 0.104599 | IL18BP | NA | -0.19831489 | 0.32138872 | EPS8 | lightgreen | 0.007557 | 0.970774 |
| ANP32B | magenta | -0.324829 | 0.105425 | ALPK1 | NA | -0.198289421 | 0.321452137 | ST6GALNAC | lightgreen | 0.007722 | 0.970135 |
| TSEN15 | magenta | -0.324721 | 0.105547 | ZNF624 | NA | -0.198195599 | 0.321685822 | S100A16 | lightgreen | 0.012727 | 0.950796 |
| CXCR2 | magenta | -0.324381 | 0.105934 | BCL9L | NA | -0.197279004 | 0.32397429 | LPL | lightgreen | 0.014613 | 0.943517 |
| SPATA5 | magenta | -0.323053 | 0.107454 | SPI1 | NA | -0.197274114 | 0.323986527 | P4HTM | lightgreen | 0.016282 | 0.937078 |
| FAM168B | magenta | 0.322663 | 0.107903 | SYK | NA | -0.197077745 | 0.324478104 | ALCAM | lightgreen | 0.024872 | 0.904005 |
| CRTC2 | magenta | 0.321649 | 0.109079 | USP38 | NA | -0.196696521 | 0.325433736 | SYNJ2 | lightgreen | 0.028311 | 0.890807 |
| SLC25A20 | magenta | -0.321573 | 0.109168 | ANKRD9 | NA | -0.196385425 | 0.326214846 | LOC646329 | lightgreen | 0.02955 | 0.886057 |
| CLASP1 | magenta | 0.319546 | 0.111546 | PPP1R37 | NA | -0.196222545 | 0.326624268 | GPC4 | lightgreen | 0.029604 | 0.885847 |
| LSMD1 | magenta | -0.316337 | 0.115391 | IL6 | NA | -0.195877336 | 0.327493033 | RBP1 | lightgreen | 0.034262 | 0.868033 |
| SHC1 | magenta | 0.314727 | 0.117355 | GBGT1 | NA | -0.195722716 | 0.32788261 | APBB2 | lightgreen | 0.037105 | 0.857188 |
| BRD7 | magenta | -0.314039 | 0.118202 | TSPYL5 | NA | -0.195585867 | 0.328227648 | ZNF486 | lightgreen | 0.038495 | 0.851895 |
| EDC3 | magenta | 0.312491 | 0.120124 | LOH12CR1 | NA | -0.19547685 | 0.32850267 | CLN8 | lightgreen | 0.040584 | 0.843952 |
| MBP | magenta | -0.311119 | 0.121848 | ZBTB10 | NA | -0.194948063 | 0.329838656 | PCSK6 | lightgreen | 0.050179 | 0.80767 |
| XPA | magenta | -0.310622 | 0.122476 | FLCN | NA | -0.194096463 | 0.331997164 | FNIP2 | lightgreen | 0.056924 | 0.782391 |
| EVI2B | magenta | -0.310224 | 0.122982 | CAT | NA | -0.194062483 | 0.332083467 | IQGAP3 | lightgreen | 0.058505 | 0.776497 |
| ACAA2 | magenta | -0.307667 | 0.126263 | CPNE8 | NA | -0.193923462 | 0.332436704 | TFPI | lightgreen | 0.061624 | 0.764901 |
| LAPTM4A | magenta | -0.304727 | 0.130113 | YES1 | NA | -0.193885098 | 0.332534223 | LASP1 | lightgreen | 0.06194 | 0.763729 |
| SCARNA10 | magenta | -0.303978 | 0.131107 | KIAA0753 | NA | -0.193775173 | 0.332813741 | MLXIPL | lightgreen | 0.062197 | 0.762774 |
| CALM2 | magenta | -0.30375 | 0.131412 | MCOLN2 | NA | -0.192699046 | 0.335557637 | RGCC | lightgreen | 0.063039 | 0.759657 |
| TXLNA | magenta | 0.301655 | 0.134227 | COL6A2 | NA | -0.192496638 | 0.336075256 | CRABP2 | lightgreen | 0.068546 | 0.739347 |

**Table 6. Gene module membership and correlation with growth rate (48/72h), with adjusted p values for each module**

| | | | | | | | | | | | |
|---|---|---|---|---|---|---|---|---|---|---|---|
| DCAF6 | magenta | 0.297737 | 0.139611 | SLC20A2 | NA | -0.192439474 | 0.336221528 | MAP7 | lightgreen | 0.068625 | 0.739055 |
| CDKN2C | magenta | -0.295177 | 0.143211 | URI1 | NA | -0.192409066 | 0.336299355 | ZNF697 | lightgreen | 0.071407 | 0.728861 |
| C1orf174 | magenta | -0.293511 | 0.145589 | B4GALT5 | NA | -0.192151007 | 0.336960256 | TNFRSF21 | lightgreen | 0.071586 | 0.728205 |
| SEC16A | magenta | 0.293364 | 0.1458 | LITAF | NA | -0.192074037 | 0.337157531 | SLC27A3 | lightgreen | 0.075742 | 0.713065 |
| TOX4 | magenta | 0.291688 | 0.148223 | GPX4 | NA | -0.192026671 | 0.337278965 | RASAL2 | lightgreen | 0.078006 | 0.70486 |
| NPC1 | magenta | 0.290604 | 0.149806 | SAMD9L | NA | -0.191622881 | 0.338315253 | NUMB | lightgreen | 0.078746 | 0.702185 |
| DYNC1H1 | magenta | 0.287215 | 0.154832 | CAMK1 | NA | -0.191548927 | 0.338505255 | AK8 | lightgreen | 0.088833 | 0.666075 |
| NT5DC1 | magenta | -0.286838 | 0.155398 | KIAA0907 | NA | -0.191340305 | 0.339041595 | RTN1 | lightgreen | 0.092469 | 0.653231 |
| DHX33 | magenta | -0.286289 | 0.156226 | ITGAM | NA | -0.191264647 | 0.33923623 | ENO2 | lightgreen | 0.09302 | 0.651294 |
| CCDC104 | magenta | -0.285113 | 0.158009 | LOC148413 | NA | -0.191076272 | 0.339721123 | DBN1 | lightgreen | 0.102703 | 0.617602 |
| SPCS3 | magenta | -0.283254 | 0.160857 | ABCF1 | NA | -0.191011017 | 0.339889192 | PHLDA2 | lightgreen | 0.106589 | 0.604284 |
| AFF4 | magenta | 0.28098 | 0.16439 | GP1BA | NA | -0.190958738 | 0.340023879 | DHRS11 | lightgreen | 0.111842 | 0.586477 |
| CRYZL1 | magenta | -0.279423 | 0.16684 | MYO18A | NA | -0.19047941 | 0.341260252 | RHOC | lightgreen | 0.11763 | 0.567125 |
| MPC2 | magenta | -0.276724 | 0.171148 | TRAF2 | NA | -0.189621689 | 0.343479381 | ITGB3 | lightgreen | 0.118705 | 0.563561 |
| DIP2B | magenta | -0.276025 | 0.172276 | BRCA2 | NA | -0.188910706 | 0.345325403 | PBX4 | lightgreen | 0.13542 | 0.509513 |
| ATP5H | magenta | -0.273021 | 0.177184 | ZAK | NA | -0.18880906 | 0.345589805 | SPATA13 | lightgreen | 0.148145 | 0.470146 |
| TRAPPC10 | magenta | 0.269148 | 0.183654 | LOC1002941 | NA | -0.188798553 | 0.345617143 | GREM1 | lightgreen | 0.149093 | 0.467276 |
| DLC1 | magenta | 0.268209 | 0.185247 | KIAA1468 | NA | -0.188357519 | 0.346765816 | TMEM241 | lightgreen | 0.158428 | 0.439524 |
| ALG10 | magenta | -0.265008 | 0.19075 | MGMT | NA | -0.18767264 | 0.3485541 | SEMA7A | lightgreen | 0.160916 | 0.432279 |
| SRCAP | magenta | 0.263457 | 0.193456 | PPP1R7 | NA | -0.187654185 | 0.348602363 | RTN4RL2 | lightgreen | 0.168844 | 0.409633 |
| PIGF | magenta | -0.26344 | 0.193486 | SNHG1 | NA | -0.187647333 | 0.348620284 | NOTCH3 | lightgreen | 0.17854 | 0.382854 |
| RPL23AP53 | magenta | 0.261018 | 0.197766 | S100A11 | NA | -0.186101671 | 0.352676767 | COL11A2 | lightgreen | 0.191424 | 0.348874 |
| LAG3 | magenta | -0.259244 | 0.200941 | SNURF | NA | -0.185654383 | 0.353855852 | AKR7A2 | lightgreen | 0.193779 | 0.342864 |
| SNX21 | magenta | -0.257969 | 0.203243 | MDM1 | NA | -0.185575712 | 0.354063476 | NPAS2 | lightgreen | 0.203285 | 0.319238 |
| ATRAID | magenta | -0.25273 | 0.212898 | UCP2 | NA | -0.18529698 | 0.35479967 | MRPS6 | lightgreen | 0.204899 | 0.315327 |
| NDUFAB1 | magenta | -0.252624 | 0.213096 | HIAT1 | NA | -0.185238147 | 0.354955178 | RNF122 | lightgreen | 0.212754 | 0.296721 |
| ANKRD50 | magenta | 0.250886 | 0.216366 | NBAS | NA | -0.184771669 | 0.356189599 | NCS1 | lightgreen | 0.218825 | 0.282823 |
| CLPTM1 | magenta | 0.250148 | 0.217766 | CHTOP | NA | -0.18467415 | 0.356447982 | OVOL1 | lightgreen | 0.22267 | 0.274238 |
| TBCC | magenta | -0.248519 | 0.220877 | TM4SF1 | NA | -0.184636054 | 0.35654895 | MATK | lightgreen | 0.234875 | 0.248099 |
| TOP3A | magenta | 0.247593 | 0.222659 | PHTF1 | NA | -0.184432241 | 0.357089409 | DCHS1 | lightgreen | 0.235087 | 0.24766 |
| PHF8 | magenta | 0.247165 | 0.223485 | FAM134C | NA | -0.184031039 | 0.358154708 | HSD3B7 | lightgreen | 0.237326 | 0.243055 |
| FAM200B | magenta | -0.246466 | 0.224839 | HBB | NA | -0.182924303 | 0.361103103 | HSD11B1 | lightgreen | 0.239648 | 0.238337 |
| MBD3 | magenta | 0.242638 | 0.232353 | IL8 | NA | -0.182801445 | 0.361431283 | KCNA10 | lightgreen | 0.240994 | 0.235632 |
| KPNA4 | magenta | 0.242591 | 0.232448 | WDR73 | NA | -0.182558846 | 0.362079827 | JARID2 | lightgreen | 0.254766 | 0.20911 |
| GEMIN2 | magenta | -0.241634 | 0.234353 | LIF | NA | -0.182272087 | 0.362847308 | SLC6A8 | lightgreen | 0.273296 | 0.176731 |
| SASH3 | magenta | -0.241289 | 0.235042 | PIK3C3 | NA | -0.182161534 | 0.363143445 | SH3RF3 | lightgreen | 0.277781 | 0.169452 |
| TNFSF13B | magenta | -0.238497 | 0.240668 | IDH2 | NA | -0.181983345 | 0.363621058 | ARAP3 | lightgreen | 0.309369 | 0.124071 |
| STAMBP | magenta | -0.238379 | 0.240909 | SNRPB2 | NA | -0.181292912 | 0.36547516 | ALDH1A2 | lightgreen | 0.319413 | 0.111704 |
| BPTF | magenta | 0.238325 | 0.241017 | CHML | NA | -0.180884919 | 0.366573389 | MYOZ1 | lightgreen | 0.345694 | 0.083666 |
| ZKSCAN2 | magenta | 0.235763 | 0.246265 | MKNK1 | NA | -0.180583386 | 0.367386294 | HMGN3 | lightyellow | -0.352367 | 0.077482 |
| SIK3 | magenta | 0.234714 | 0.248433 | ITGAV | NA | -0.180170418 | 0.36850132 | RPL9 | lightyellow | -0.291323 | 0.148755 |
| ELF4 | magenta | 0.234277 | 0.24934 | USP9X | NA | -0.179293853 | 0.370874618 | SLC44A1 | lightyellow | -0.285286 | 0.157746 |
| TRAFD1 | magenta | -0.234109 | 0.249691 | TMEM63A | NA | -0.178775986 | 0.372280913 | C6orf48 | lightyellow | -0.253736 | 0.21102 |
| GABARAPL | magenta | -0.232222 | 0.253638 | MRPL1 | NA | -0.178052809 | 0.374249922 | GMFG | lightyellow | -0.23747 | 0.24276 |
| UBE2D3 | magenta | 0.229237 | 0.259963 | SCARF1 | NA | -0.177832992 | 0.374849619 | OTUD4 | lightyellow | -0.22338 | 0.27267 |
| PXMP2 | magenta | -0.226188 | 0.266528 | LYPD3 | NA | -0.17774977 | 0.375076808 | GSTK1 | lightyellow | -0.223105 | 0.273275 |
| TOP1 | magenta | 0.226176 | 0.266556 | RFTN1 | NA | -0.17736126 | 0.376138461 | ZNF253 | lightyellow | -0.22194 | 0.275854 |
| GPRIN3 | magenta | 0.224931 | 0.269266 | MEX3C | NA | -0.177344725 | 0.376183681 | BRCA2 | lightyellow | -0.218007 | 0.28467 |
| RHOT2 | magenta | 0.223766 | 0.27182 | ACYP2 | NA | -0.177195137 | 0.376592943 | RPS13 | lightyellow | -0.217813 | 0.285111 |
| TWF2 | magenta | -0.218306 | 0.283994 | ZNF333 | NA | -0.177103742 | 0.376843119 | USO1 | lightyellow | -0.212458 | 0.29741 |
| DNAJC14 | magenta | 0.217945 | 0.284812 | SERPINB2 | NA | -0.176797178 | 0.377682978 | S100A4 | lightyellow | -0.209224 | 0.304997 |
| MAST2 | magenta | 0.21656 | 0.287959 | NEDD4L | NA | -0.176583715 | 0.378268421 | RPS3 | lightyellow | -0.204034 | 0.317418 |
| TBC1D8 | magenta | 0.215936 | 0.289385 | KPNA5 | NA | -0.17657534 | 0.378291398 | GTF2IRD2B | lightyellow | -0.191013 | 0.349931 |
| TCTN3 | magenta | -0.214 | 0.293835 | DFFB | NA | -0.17582377 | 0.380356886 | RPL7 | lightyellow | -0.19019 | 0.352051 |
| HINT2 | magenta | -0.210432 | 0.302148 | DOCK4 | NA | -0.17563293 | 0.380882392 | GLRX5 | lightyellow | -0.182202 | 0.37301 |
| RFT1 | magenta | -0.209562 | 0.304198 | CHRM3 | NA | -0.175499757 | 0.381249351 | ALAS1 | lightyellow | -0.181561 | 0.374723 |

**Table 6. Gene module membership and correlation with growth rate (48/72h), with adjusted p values for each module**

| | | | | | | | | | | | |
|---|---|---|---|---|---|---|---|---|---|---|---|
| UBXN7 | magenta | 0.209397 | 0.304587 | TAGLN2 | NA | -0.175265395 | 0.38189563 | ZNF302 | lightyellow | -0.177823 | 0.3848 |
| STARD3NL | magenta | -0.207319 | 0.30952 | CXADR | NA | -0.174946843 | 0.382775085 | TMEM57 | lightyellow | -0.172738 | 0.398755 |
| TATDN3 | magenta | -0.206166 | 0.31228 | COMMD4 | NA | -0.174907482 | 0.382883831 | ACN9 | lightyellow | -0.168264 | 0.411266 |
| FAM54B | magenta | 0.205336 | 0.314275 | SPDL1 | NA | -0.173969802 | 0.385479741 | C20orf196 | lightyellow | -0.167792 | 0.412598 |
| ANKRD52 | magenta | -0.204439 | 0.316439 | ZER1 | NA | -0.173427381 | 0.386985994 | RPL35 | lightyellow | -0.163243 | 0.425561 |
| ZNF674 | magenta | 0.204378 | 0.316586 | ICOSLG | NA | -0.173415687 | 0.387018503 | FRAT2 | lightyellow | -0.157945 | 0.440937 |
| NR2C1 | magenta | -0.202055 | 0.322237 | CSPG4 | NA | -0.17331853 | 0.38728867 | RPS11 | lightyellow | -0.157511 | 0.442208 |
| RNMTL1 | magenta | 0.199074 | 0.329579 | RNF213 | NA | -0.172418398 | 0.389796792 | LAMTOR4 | lightyellow | -0.156415 | 0.445431 |
| FAM92A1 | magenta | -0.19668 | 0.335546 | UGCG | NA | -0.172113412 | 0.390648701 | RPL19 | lightyellow | -0.156209 | 0.446039 |
| BLOC1S1 | magenta | -0.193751 | 0.342936 | MTRF1L | NA | -0.171265484 | 0.393022769 | DEF6 | lightyellow | -0.154155 | 0.452114 |
| HBS1L | magenta | -0.191123 | 0.349647 | TCTN2 | NA | -0.170874378 | 0.394120563 | ZCCHC7 | lightyellow | -0.150215 | 0.463895 |
| NDUFB1 | magenta | -0.190876 | 0.350283 | ZNF581 | NA | -0.170631681 | 0.394802663 | MICALL2 | lightyellow | -0.138516 | 0.499787 |
| INO80D | magenta | 0.190515 | 0.351211 | L3MBTL2 | NA | -0.170424847 | 0.3953845 | FAR2 | lightyellow | -0.136165 | 0.507165 |
| ENTPD4 | magenta | 0.18826 | 0.35705 | USP37 | NA | -0.170363815 | 0.39555628 | DESI1 | lightyellow | -0.132162 | 0.519844 |
| BFAR | magenta | 0.18701 | 0.360309 | LOC1002884 | NA | -0.170325033 | 0.395665455 | TAF9B | lightyellow | -0.120123 | 0.558879 |
| OC1001290 | magenta | 0.186977 | 0.360396 | ABCC10 | NA | -0.170107339 | 0.396278612 | RPL23 | lightyellow | -0.113357 | 0.581384 |
| PGLS | magenta | -0.18599 | 0.362983 | TCP11L2 | NA | -0.169565206 | 0.397807922 | SLC43A3 | lightyellow | -0.11183 | 0.586519 |
| RYBP | magenta | 0.183369 | 0.369904 | RILP | NA | -0.169520672 | 0.397933697 | RPL26 | lightyellow | -0.107841 | 0.600019 |
| KLHL21 | magenta | 0.181096 | 0.375968 | CLDN1 | NA | -0.16944279 | 0.39815371 | LSMD1 | lightyellow | -0.100939 | 0.623686 |
| ASB7 | magenta | -0.179508 | 0.380239 | LOC645513 | NA | -0.169134071 | 0.399026498 | ENDOD1 | lightyellow | -0.099823 | 0.627549 |
| ACAP2 | magenta | 0.17846 | 0.383072 | AP5M1 | NA | -0.169028131 | 0.399326252 | RPS10 | lightyellow | -0.099505 | 0.628651 |
| LYAR | magenta | -0.176445 | 0.388555 | IL10RB | NA | -0.168788917 | 0.400003569 | NXPE3 | lightyellow | -0.097159 | 0.636806 |
| HOOK2 | magenta | -0.176193 | 0.389243 | MTMR14 | NA | -0.168388134 | 0.401139811 | RGS10 | lightyellow | -0.094361 | 0.646587 |
| CHRNE | magenta | -0.175318 | 0.391641 | GALNT2 | NA | -0.168116309 | 0.401911481 | PTRHD1 | lightyellow | -0.09031 | 0.660849 |
| SCARNA13 | magenta | -0.174871 | 0.392867 | SLC37A2 | NA | -0.167775122 | 0.402881244 | KIF1B | lightyellow | -0.088636 | 0.666775 |
| LIG1 | magenta | -0.174734 | 0.393244 | PBXIP1 | NA | -0.167548468 | 0.403526196 | TAF1D | lightyellow | -0.088264 | 0.668095 |
| DCP1A | magenta | 0.174202 | 0.39471 | TCN2 | NA | -0.167499123 | 0.403666685 | RPL27A | lightyellow | -0.087781 | 0.669809 |
| DOCK7 | magenta | -0.173103 | 0.397743 | SOLH | NA | -0.167019395 | 0.405033942 | ZCCHC10 | lightyellow | -0.084264 | 0.682347 |
| TSC22D1 | magenta | -0.172605 | 0.399125 | ELK4 | NA | -0.166749592 | 0.40580404 | PTPN4 | lightyellow | -0.078102 | 0.70451 |
| CLEC4A | magenta | -0.171625 | 0.401847 | MRPS27 | NA | -0.166567712 | 0.40632364 | SCYL1 | lightyellow | -0.076448 | 0.710501 |
| PNPLA2 | magenta | 0.170629 | 0.404626 | PSAT1 | NA | -0.166384389 | 0.406847741 | GATAD2A | lightyellow | -0.073413 | 0.721535 |
| ORAOV1 | magenta | 0.169102 | 0.408905 | ZNF782 | NA | -0.166253558 | 0.407222006 | THYN1 | lightyellow | -0.071523 | 0.728438 |
| SFXN4 | magenta | -0.164109 | 0.423078 | NUP133 | NA | -0.165782101 | 0.408572283 | NOA1 | lightyellow | -0.064662 | 0.753653 |
| HSPA14 | magenta | -0.16359 | 0.424564 | NFAT5 | NA | -0.165723088 | 0.408741475 | ILKAP | lightyellow | -0.064251 | 0.755173 |
| NAA30 | magenta | 0.163148 | 0.425835 | C20orf112 | NA | -0.165559315 | 0.409211222 | AFF1 | lightyellow | -0.062544 | 0.76149 |
| CLPTM1L | magenta | -0.161791 | 0.429746 | RBAK | NA | -0.16537855 | 0.409730058 | MICALL1 | lightyellow | -0.057978 | 0.778459 |
| ACACB | magenta | 0.160724 | 0.432837 | MMP2 | NA | -0.165212215 | 0.4102078 | INF2 | lightyellow | -0.051598 | 0.802333 |
| SEC24C | magenta | 0.160716 | 0.432858 | SARNP | NA | -0.165182095 | 0.410294343 | ATF3 | lightyellow | -0.049489 | 0.810266 |
| HIPK3 | magenta | 0.159261 | 0.43709 | MAP7D3 | NA | -0.165091822 | 0.410553783 | LRP8 | lightyellow | -0.047626 | 0.817288 |
| MED11 | magenta | -0.158542 | 0.439189 | LZTFL1 | NA | -0.16434536 | 0.412702576 | LMBR1 | lightyellow | -0.044024 | 0.830902 |
| MTCP1 | magenta | 0.157514 | 0.442202 | CHD1 | NA | -0.164311498 | 0.412800203 | NCOR2 | lightyellow | -0.034505 | 0.867105 |
| MLLT1 | magenta | 0.154573 | 0.450876 | TRIM28 | NA | -0.163707928 | 0.414542454 | ARID5A | lightyellow | -0.033158 | 0.872248 |
| SMN2 | magenta | -0.152909 | 0.455822 | WDR59 | NA | -0.163615648 | 0.414809187 | RPS12 | lightyellow | -0.029624 | 0.885772 |
| LYRM7 | magenta | 0.15217 | 0.458029 | NBN | NA | -0.163163361 | 0.416117884 | AP5B1 | lightyellow | -0.026379 | 0.898215 |
| CFL2 | magenta | -0.151456 | 0.460166 | ETS2 | NA | -0.162996617 | 0.416600937 | VAMP8 | lightyellow | -0.023579 | 0.908973 |
| ACTR3 | magenta | -0.15101 | 0.461504 | FERMT2 | NA | -0.162723752 | 0.417392088 | RPS8 | lightyellow | -0.018752 | 0.927554 |
| XRCC3 | magenta | -0.150922 | 0.46177 | WHAMMP1 | NA | -0.162672283 | 0.417541411 | PHF6 | lightyellow | -0.012209 | 0.952797 |
| C1orf233 | magenta | -0.149382 | 0.466404 | KCTD17 | NA | -0.162480441 | 0.418098249 | CRLS1 | lightyellow | -0.00492 | 0.980971 |
| ALDH9A1 | magenta | -0.149355 | 0.466487 | AMIGO2 | NA | -0.162191284 | 0.418938329 | CDC42EP1 | lightyellow | -0.003297 | 0.987247 |
| SH3BGRL | magenta | -0.149198 | 0.46696 | ICA1 | NA | -0.16198047 | 0.419551387 | LIN37 | lightyellow | -0.001135 | 0.99561 |
| GBF1 | magenta | 0.148836 | 0.468054 | METTL15 | NA | -0.161846683 | 0.419940702 | DYNC1H1 | lightyellow | 0.00084 | 0.996751 |
| ANKRD28 | magenta | 0.147801 | 0.471189 | HHLA3 | NA | -0.161644781 | 0.42052861 | RPL15 | lightyellow | 0.004948 | 0.98086 |
| LAMB1 | magenta | 0.147193 | 0.473035 | CHPT1 | NA | -0.160246565 | 0.424612414 | KIAA1522 | lightyellow | 0.007727 | 0.970114 |
| COMMD3 | magenta | -0.146677 | 0.474605 | P2RX7 | NA | -0.160128925 | 0.424956995 | CREBL2 | lightyellow | 0.009149 | 0.964619 |
| MNF1 | magenta | -0.146511 | 0.475111 | METTL4 | NA | -0.159763972 | 0.426026966 | SLC25A25 | lightyellow | 0.009838 | 0.961958 |
| RBM19 | magenta | 0.145895 | 0.47699 | ZNF449 | NA | -0.159689435 | 0.426245675 | SF3B5 | lightyellow | 0.01427 | 0.944842 |

**Table 6. Gene module membership and correlation with growth rate (48/72h), with adjusted p values for each module**

| | | | | | | | | | | | |
|---|---|---|---|---|---|---|---|---|---|---|---|
| SNRNP200 | magenta | 0.14384 | 0.483284 | EIF5A | NA | -0.159577249 | 0.426574969 | RPS6 | lightyellow | 0.015832 | 0.938814 |
| RAPH1 | magenta | 0.140781 | 0.492733 | BAALC | NA | -0.159491032 | 0.426828134 | BTN2A2 | lightyellow | 0.017197 | 0.933549 |
| AKT2 | magenta | 0.140675 | 0.493062 | C1S | NA | -0.159328051 | 0.427306931 | TCHP | lightyellow | 0.021282 | 0.91781 |
| TMEM237 | magenta | -0.139931 | 0.495374 | TMEM129 | NA | -0.159287759 | 0.427425345 | GCN1L1 | lightyellow | 0.022621 | 0.912659 |
| CAPZA1 | magenta | -0.139448 | 0.496878 | FOXP4 | NA | -0.158914827 | 0.428522191 | PNP | lightyellow | 0.030345 | 0.88301 |
| NPLOC4 | magenta | 0.138297 | 0.500474 | HSD17B11 | NA | -0.158792282 | 0.42888295 | RPS23 | lightyellow | 0.041753 | 0.839512 |
| ZXDC | magenta | 0.138156 | 0.500914 | ECD | NA | -0.15864506 | 0.429316572 | KLHL5 | lightyellow | 0.045703 | 0.824552 |
| COPS4 | magenta | -0.136951 | 0.504692 | CYP1B1 | NA | -0.157979657 | 0.431279411 | PTPRJ | lightyellow | 0.047778 | 0.816714 |
| PLK1S1 | magenta | -0.135955 | 0.507823 | TMEM176E | NA | -0.157917905 | 0.431461815 | RHOD | lightyellow | 0.052426 | 0.799224 |
| ZFPL1 | magenta | 0.135885 | 0.508046 | LOC550112 | NA | -0.157792221 | 0.431833197 | FLNB | lightyellow | 0.055658 | 0.787121 |
| RTCA | magenta | -0.135297 | 0.5099 | RAB11FIP1 | NA | -0.157204557 | 0.433571981 | IFT74 | lightyellow | 0.06124 | 0.766327 |
| PDPR | magenta | -0.134511 | 0.512385 | AXL | NA | -0.157102437 | 0.433874521 | SLC9A7 | lightyellow | 0.073057 | 0.722836 |
| WAPAL | magenta | 0.132861 | 0.517618 | ANKMY1 | NA | -0.157029661 | 0.434090196 | PMEPA1 | lightyellow | 0.079737 | 0.698608 |
| CDAN1 | magenta | -0.132037 | 0.520241 | TNS1 | NA | -0.15630975 | 0.43622682 | MBNL1 | lightyellow | 0.080573 | 0.695593 |
| KDM3A | magenta | -0.131056 | 0.523375 | TSFM | NA | -0.156212457 | 0.436516011 | GAB1 | lightyellow | 0.09284 | 0.651925 |
| C15orf38 | magenta | -0.128045 | 0.533043 | UBE2O | NA | -0.155851656 | 0.437589352 | GLDN | lightyellow | 0.094508 | 0.646071 |
| GATC | magenta | 0.125896 | 0.539994 | CUX1 | NA | -0.155686371 | 0.438081533 | CPSF7 | lightyellow | 0.105913 | 0.606592 |
| NT5C2 | magenta | 0.125845 | 0.540161 | CCL22 | NA | -0.154996005 | 0.440140506 | CREBRF | lightyellow | 0.119876 | 0.559694 |
| PEX5 | magenta | 0.123495 | 0.54781 | HIVEP2 | NA | -0.154742263 | 0.440898584 | TBC1D9B | lightyellow | 0.12246 | 0.551198 |
| ZNHIT3 | magenta | -0.123423 | 0.548046 | DMAP1 | NA | -0.15466643 | 0.441125277 | AKAP17A | lightyellow | 0.123298 | 0.548456 |
| COPG2 | magenta | -0.12336 | 0.548253 | WNT5A | NA | -0.154519503 | 0.441564677 | SPATC1 | lightyellow | 0.129931 | 0.526978 |
| RWDD4 | magenta | -0.122537 | 0.550946 | VDAC1 | NA | -0.154234135 | 0.442418769 | HK2 | lightyellow | 0.133463 | 0.515706 |
| LRRC37A4P | magenta | 0.121754 | 0.553512 | MDH2 | NA | -0.15388576 | 0.443462639 | STAU2 | lightyellow | 0.143603 | 0.484014 |
| DDX6 | magenta | 0.121182 | 0.55539 | PRKRIR | NA | -0.153504798 | 0.444605663 | CYTIP | lightyellow | 0.15883 | 0.438349 |
| GRWD1 | magenta | 0.121112 | 0.555621 | LIG3 | NA | -0.153132813 | 0.445723276 | GIGYF1 | lightyellow | 0.163033 | 0.426165 |
| KIAA0319L | magenta | -0.118431 | 0.56447 | APPBP2 | NA | -0.152986545 | 0.446163141 | MFAP3 | lightyellow | 0.164511 | 0.421927 |
| SIK1 | magenta | 0.118221 | 0.565165 | LEMD2 | NA | -0.152593062 | 0.447347599 | RPL37A | lightyellow | 0.169699 | 0.40723 |
| DNAL4 | magenta | -0.117861 | 0.566357 | MRPL32 | NA | -0.152531657 | 0.447532589 | WDR67 | lightyellow | 0.173426 | 0.39685 |
| SDHA | magenta | -0.117138 | 0.568758 | TRERF1 | NA | -0.152313838 | 0.448189129 | ZNF438 | lightyellow | 0.176912 | 0.387279 |
| NT5C3L | magenta | -0.1168 | 0.56988 | SLC11A1 | NA | -0.151374397 | 0.451026627 | TNS3 | lightyellow | 0.179565 | 0.380085 |
| ARMC1 | magenta | -0.116438 | 0.571088 | VPS26B | NA | -0.151321266 | 0.451187392 | C20orf112 | lightyellow | 0.184318 | 0.367388 |
| BNIP3L | magenta | 0.116146 | 0.572059 | NARF | NA | -0.151225849 | 0.451476175 | EGR1 | lightyellow | 0.191291 | 0.349218 |
| NUP98 | magenta | 0.115736 | 0.573424 | OTUB2 | NA | -0.150963011 | 0.452272179 | TNIK | lightyellow | 0.214849 | 0.291877 |
| PLA2G4A | magenta | 0.113466 | 0.581018 | MPC2 | NA | -0.150699557 | 0.453070797 | HEATR2 | lightyellow | 0.218249 | 0.284123 |
| PEBP1 | magenta | -0.112676 | 0.58367 | LMNA | NA | -0.150698923 | 0.45307272 | VDR | lightyellow | 0.226359 | 0.266159 |
| INTS2 | magenta | 0.112612 | 0.583887 | ARL14EP | NA | -0.150606455 | 0.453353197 | UGDH | lightyellow | 0.254658 | 0.20931 |
| MLLT4 | magenta | 0.112458 | 0.584404 | ECE1 | NA | -0.150047977 | 0.455049157 | GXYLT1 | lightyellow | 0.269506 | 0.183049 |
| DCTN2 | magenta | -0.11037 | 0.591442 | TAPT1 | NA | -0.149929075 | 0.455410662 | PLEKHM3 | lightyellow | 0.295145 | 0.143256 |
| RCBTB1 | magenta | -0.108809 | 0.596729 | ATP2B4 | NA | -0.149297434 | 0.457333635 | SLC30A1 | lightyellow | 0.29924 | 0.137528 |
| PPARA | magenta | -0.107192 | 0.602228 | ZNF200 | NA | -0.148964256 | 0.458349686 | HCFC1 | lightyellow | 0.308442 | 0.125261 |
| WDR48 | magenta | -0.106204 | 0.605598 | IDH3A | NA | -0.148697484 | 0.459164083 | KDM6B | lightyellow | 0.308938 | 0.124624 |
| G6PD | magenta | 0.105811 | 0.606942 | VAMP5 | NA | -0.148631623 | 0.459365261 | PDE4A | lightyellow | 0.318669 | 0.112588 |
| HIGD2A | magenta | -0.105173 | 0.609121 | DYX1C1 | NA | -0.148613588 | 0.459420359 | RRN3 | magenta | -0.397247 | 0.044483 |
| PPP2R3B | magenta | -0.103262 | 0.615677 | CCL24 | NA | -0.148572642 | 0.459545462 | TGS1 | magenta | -0.383061 | 0.053411 |
| GTF2F1 | magenta | 0.100532 | 0.625092 | TOP1 | NA | -0.148555434 | 0.459598042 | GK5 | magenta | -0.354475 | 0.075602 |
| B4GALT1 | magenta | 0.09813 | 0.633423 | FAM105A | NA | -0.148414287 | 0.460029454 | SNX2 | magenta | -0.352939 | 0.076968 |
| EXOC3 | magenta | 0.093772 | 0.648651 | JAZF1 | NA | -0.148348031 | 0.46023204 | TIMM8A | magenta | -0.330774 | 0.098838 |
| DOCK5 | magenta | 0.093752 | 0.648723 | THUMPD1 | NA | -0.147929655 | 0.46151234 | PTPN9 | magenta | -0.325888 | 0.104228 |
| EXOC4 | magenta | -0.092954 | 0.651524 | NCF1C | NA | -0.147704724 | 0.462201438 | TM6SF1 | magenta | -0.320697 | 0.11019 |
| RNF121 | magenta | 0.090383 | 0.66059 | SNX20 | NA | -0.146891051 | 0.464698696 | NUBPL | magenta | -0.317369 | 0.114143 |
| FGR | magenta | 0.089442 | 0.663919 | HCAR3 | NA | -0.146662179 | 0.4654024 | FAM96A | magenta | -0.31224 | 0.120439 |
| CTAGE5 | magenta | -0.088984 | 0.665541 | CRLF2 | NA | -0.146449732 | 0.466056096 | ALDH1A1 | magenta | -0.308963 | 0.124592 |
| CYP2U1 | magenta | -0.087224 | 0.671792 | JKAMP | NA | -0.146345516 | 0.466376945 | RNASEH2A | magenta | -0.306577 | 0.127681 |
| NDUFA12 | magenta | -0.087003 | 0.672575 | ITGA2 | NA | -0.146318661 | 0.466459641 | AKAP7 | magenta | -0.299311 | 0.13743 |
| KBTBD7 | magenta | -0.085671 | 0.677323 | KIFC3 | NA | -0.146254029 | 0.466658696 | CD109 | magenta | -0.294533 | 0.144127 |
| RASSF7 | magenta | -0.085631 | 0.677463 | TMED10 | NA | -0.14572678 | 0.468284197 | ASPH | magenta | -0.294487 | 0.144193 |

**Table 6. Gene module membership and correlation with growth rate (48/72h), with adjusted p values for each module**

| | | | | | | | | | | | |
|---|---|---|---|---|---|---|---|---|---|---|---|
| C9orf91 | magenta | 0.084377 | 0.681942 | ANKRD17 | NA | -0.145572721 | 0.468759713 | NEDD1 | magenta | -0.289446 | 0.151511 |
| CYLD | magenta | 0.084049 | 0.683116 | EEF1G | NA | -0.145175191 | 0.46998788 | DBP | magenta | -0.287954 | 0.153726 |
| ACOT9 | magenta | -0.081529 | 0.692154 | TEX30 | NA | -0.145143083 | 0.470087153 | BROX | magenta | -0.284179 | 0.159435 |
| KANSL1L | magenta | -0.081029 | 0.693953 | MS4A6A | NA | -0.14489241 | 0.470862548 | CASP1 | magenta | -0.278986 | 0.167532 |
| CHD2 | magenta | 0.080067 | 0.697415 | TCEAL4 | NA | -0.144276227 | 0.472771369 | XPO4 | magenta | -0.275629 | 0.172917 |
| OTUD5 | magenta | 0.079015 | 0.70121 | PES1 | NA | -0.143953832 | 0.473771679 | C17orf61 | magenta | -0.274448 | 0.174841 |
| RAB31 | magenta | -0.077331 | 0.707301 | WDR34 | NA | -0.143744224 | 0.474422626 | CCNT1 | magenta | -0.272765 | 0.177607 |
| PPFIBP2 | magenta | -0.076023 | 0.712042 | TLCD2 | NA | -0.142008216 | 0.479831549 | ZDHHC17 | magenta | -0.270129 | 0.182001 |
| CA13 | magenta | 0.074153 | 0.718841 | RFX5 | NA | -0.141595096 | 0.48112335 | NLK | magenta | -0.26882 | 0.18421 |
| DZIP3 | magenta | -0.073762 | 0.720264 | RNASEK | NA | -0.14154351 | 0.481284779 | PPP3CC | magenta | -0.268489 | 0.184771 |
| HABP4 | magenta | -0.07365 | 0.720672 | SAMSN1 | NA | -0.141434626 | 0.481625607 | ANKRD49 | magenta | -0.258382 | 0.202496 |
| GLB1 | magenta | -0.069234 | 0.73682 | CDC42EP3 | NA | -0.141259816 | 0.482173053 | FIG4 | magenta | -0.257769 | 0.203607 |
| GSR | magenta | -0.067255 | 0.744094 | MAGED2 | NA | -0.141194322 | 0.482378239 | FEZ2 | magenta | -0.257423 | 0.204236 |
| CHD1 | magenta | -0.066888 | 0.745443 | SRPK1 | NA | -0.141012045 | 0.482949533 | MRPS35 | magenta | -0.257392 | 0.204293 |
| FNDC3B | magenta | 0.061621 | 0.764913 | AGAP6 | NA | -0.140652105 | 0.484078666 | PIGA | magenta | -0.25254 | 0.213253 |
| JUN | magenta | -0.061541 | 0.765207 | ZNF653 | NA | -0.140156455 | 0.485635722 | IKBKAP | magenta | -0.250634 | 0.216845 |
| GLA | magenta | 0.056755 | 0.783021 | RPAP1 | NA | -0.139897244 | 0.486451029 | PIK3CA | magenta | -0.250165 | 0.217734 |
| ZNF260 | magenta | 0.056684 | 0.783287 | SLMAP | NA | -0.139403649 | 0.48800547 | UQCRHL | magenta | -0.250042 | 0.217969 |
| C7orf29 | magenta | -0.055311 | 0.788417 | ITPA | NA | -0.139395433 | 0.488031369 | BAG1 | magenta | -0.248625 | 0.220674 |
| PDCD11 | magenta | -0.054774 | 0.790426 | ZNF736 | NA | -0.139332613 | 0.488229389 | FLJ31306 | magenta | -0.248458 | 0.220994 |
| NR3C1 | magenta | 0.053915 | 0.793641 | ZNF670 | NA | -0.139168323 | 0.488747451 | TXNDC12 | magenta | -0.247512 | 0.222814 |
| CBX4 | magenta | 0.05341 | 0.795534 | RPLP0P2 | NA | -0.138899497 | 0.489595754 | POLE3 | magenta | -0.246461 | 0.224849 |
| PIPOX | magenta | -0.05023 | 0.807479 | FAM149B1 | NA | -0.138703831 | 0.490213662 | HDAC8 | magenta | -0.245562 | 0.226599 |
| PDIA3P | magenta | -0.050191 | 0.807623 | AACS | NA | -0.138039813 | 0.492313541 | AHSA1 | magenta | -0.244738 | 0.228211 |
| ZC3H4 | magenta | 0.048961 | 0.812254 | TMEM45B | NA | -0.137931861 | 0.492655354 | GIN1 | magenta | -0.242185 | 0.233253 |
| DOCK1 | magenta | -0.048919 | 0.812416 | ISG20 | NA | -0.137859166 | 0.492885598 | NSDHL | magenta | -0.242114 | 0.233395 |
| MPV17 | magenta | -0.048794 | 0.812884 | CCDC109B | NA | -0.137359426 | 0.494469872 | IDI1 | magenta | -0.239178 | 0.239287 |
| CORO1C | magenta | -0.047821 | 0.816555 | TMTC1 | NA | -0.137132625 | 0.495189715 | SCOC | magenta | -0.238057 | 0.241562 |
| GNE | magenta | -0.047637 | 0.817247 | PPP1R26 | NA | -0.137002822 | 0.495601937 | PGM2 | magenta | -0.237372 | 0.242961 |
| CCNDBP1 | magenta | -0.046598 | 0.82117 | CSF1 | NA | -0.136583071 | 0.496936126 | TMEM159 | magenta | -0.234946 | 0.247953 |
| TMEM110 | magenta | 0.045287 | 0.826126 | UCK2 | NA | -0.136560288 | 0.497008594 | GALK2 | magenta | -0.234454 | 0.248974 |
| ZNF468 | magenta | -0.043593 | 0.832535 | PACS2 | NA | -0.136456057 | 0.497340199 | SNRNP40 | magenta | -0.233926 | 0.250071 |
| POLR2D | magenta | -0.041872 | 0.839061 | TLR2 | NA | -0.136329897 | 0.497741716 | PPP2R2D | magenta | -0.232167 | 0.253753 |
| VAMP2 | magenta | 0.038206 | 0.852996 | JOSD1 | NA | -0.135833651 | 0.499322638 | ZDHHC2 | magenta | -0.231025 | 0.256161 |
| KPNB1 | magenta | 0.037538 | 0.855537 | TMEM199 | NA | -0.135514515 | 0.500340651 | RABEPK | magenta | -0.230672 | 0.256909 |
| SP2 | magenta | 0.037323 | 0.856357 | C17orf58 | NA | -0.135434199 | 0.500597014 | IL1RAP | magenta | -0.229725 | 0.258922 |
| CYP1B1 | magenta | 0.037215 | 0.856767 | AMZ1 | NA | -0.135344029 | 0.50088491 | HAUS1 | magenta | -0.229301 | 0.259826 |
| AOC2 | magenta | 0.036261 | 0.860402 | PGBD2 | NA | -0.135270641 | 0.501119282 | NUDT12 | magenta | -0.226008 | 0.266921 |
| RDX | magenta | 0.035122 | 0.864749 | SERP1 | NA | -0.135180384 | 0.501407603 | ATF6 | magenta | -0.225715 | 0.267558 |
| TJP2 | magenta | -0.033307 | 0.871681 | ZNF552 | NA | -0.134876207 | 0.502379888 | MAN2A1 | magenta | -0.225016 | 0.269082 |
| OASL | magenta | -0.03172 | 0.877749 | DUSP7 | NA | -0.134854529 | 0.502449216 | MSANTD2 | magenta | -0.222559 | 0.274483 |
| CHCHD3 | magenta | 0.030627 | 0.881932 | LOC645638 | NA | -0.134802593 | 0.502615331 | IFT46 | magenta | -0.221995 | 0.275731 |
| PARP14 | magenta | 0.029744 | 0.885312 | ANKRD13A | NA | -0.134731463 | 0.502842883 | OXA1L | magenta | -0.221259 | 0.277368 |
| SLC19A2 | magenta | 0.029693 | 0.885509 | C16orf13 | NA | -0.134710775 | 0.502909076 | AP2B1 | magenta | -0.221245 | 0.277398 |
| GMPR | magenta | 0.029043 | 0.888 | COX5A | NA | -0.134591315 | 0.503291378 | TMX3 | magenta | -0.220811 | 0.278367 |
| PCNX | magenta | 0.028693 | 0.889342 | SERPINE1 | NA | -0.134587673 | 0.503303034 | RAB11A | magenta | -0.220712 | 0.278587 |
| TSN | magenta | 0.027954 | 0.892174 | VWA8 | NA | -0.134352736 | 0.504055322 | APPL1 | magenta | -0.218693 | 0.283119 |
| ENOX2 | magenta | -0.02768 | 0.893225 | GIGYF1 | NA | -0.134159086 | 0.504675822 | GCNT1 | magenta | -0.214636 | 0.292369 |
| AP2M1 | magenta | -0.025814 | 0.900386 | PCNX | NA | -0.133923316 | 0.505431798 | MLL3 | magenta | -0.213755 | 0.294401 |
| MRS2 | magenta | 0.018751 | 0.927558 | CELSR1 | NA | -0.133723658 | 0.506072418 | PSMD10 | magenta | -0.211533 | 0.299567 |
| FRYL | magenta | -0.009823 | 0.962016 | FBXL4 | NA | -0.133681476 | 0.506207814 | DLD | magenta | -0.211017 | 0.300775 |
| FBXL5 | magenta | -0.007954 | 0.969239 | ATP6V0D2 | NA | -0.13355152 | 0.506625059 | TNKS | magenta | -0.208127 | 0.307597 |
| SLC10A3 | magenta | 0.004726 | 0.981718 | AMICA1 | NA | -0.133488711 | 0.506826781 | MDH1 | magenta | -0.20687 | 0.310593 |
| AEBP2 | magenta | 0.003328 | 0.987125 | PARN | NA | -0.132447977 | 0.510175001 | TRAPPC1 | magenta | -0.20149 | 0.323622 |
| PRKCE | magenta | 0.002134 | 0.991745 | RWDD4 | NA | -0.132151228 | 0.511131679 | GOT1 | magenta | -0.198507 | 0.330986 |
| PRKAB2 | magenta | -0.001565 | 0.993945 | PPP5C | NA | -0.132012953 | 0.511577759 | AOC3 | magenta | -0.198114 | 0.331964 |

**Table 6. Gene module membership and correlation with growth rate (48/72h), with adjusted p values for each module**

| | | | | | | | | | | | |
|---|---|---|---|---|---|---|---|---|---|---|---|
| IL15RA | midnightblue | -0.463319 | 0.017136 | CSF2RB | NA | -0.131390571 | 0.513587941 | SAR1B | magenta | -0.197057 | 0.334602 |
| CST7 | midnightblue | -0.460033 | 0.018046 | DUSP14 | NA | -0.131320831 | 0.513813429 | CAMK1 | magenta | -0.193337 | 0.343988 |
| CCR5 | midnightblue | -0.454366 | 0.019709 | SLC45A3 | NA | -0.131246559 | 0.514053624 | EXTL2 | magenta | -0.191298 | 0.349199 |
| SELL | midnightblue | -0.440571 | 0.024282 | ZDHHC21 | NA | -0.131240991 | 0.514071635 | VPS13D | magenta | -0.19079 | 0.350505 |
| CD3D | midnightblue | -0.414821 | 0.035104 | HMBS | NA | -0.131211928 | 0.514165639 | DOCK11 | magenta | -0.190123 | 0.352223 |
| SGTB | midnightblue | -0.406471 | 0.039341 | DLG5 | NA | -0.131177887 | 0.514275758 | PRR14L | magenta | -0.188813 | 0.355612 |
| PTPN7 | midnightblue | -0.37956 | 0.055816 | TRIM62 | NA | -0.131011514 | 0.514814115 | PRUNE | magenta | -0.188117 | 0.35742 |
| CD5 | midnightblue | -0.374757 | 0.059252 | KCNE3 | NA | -0.130779111 | 0.515566598 | ALG5 | magenta | -0.186406 | 0.361889 |
| FYN | midnightblue | -0.364738 | 0.066946 | CERS6 | NA | -0.130667732 | 0.515927417 | ZKSCAN5 | magenta | -0.186038 | 0.362856 |
| MCOLN2 | midnightblue | -0.344368 | 0.084939 | WDFY3 | NA | -0.130179423 | 0.517510762 | ANAPC5 | magenta | -0.184699 | 0.366382 |
| ICAM4 | midnightblue | -0.343997 | 0.085297 | PPP1R3B | NA | -0.130046192 | 0.517943175 | RIPK1 | magenta | -0.184405 | 0.367158 |
| KLRB1 | midnightblue | -0.335882 | 0.09343 | FCHSD2 | NA | -0.12952222 | 0.519645469 | NPC2 | magenta | -0.183041 | 0.370776 |
| GNLY | midnightblue | -0.335868 | 0.093444 | IRF1 | NA | -0.128944368 | 0.521525945 | ANXA5 | magenta | -0.182139 | 0.373179 |
| CD52 | midnightblue | -0.332311 | 0.097187 | ICK | NA | -0.128703009 | 0.522312361 | HNRNPA0 | magenta | -0.180616 | 0.377257 |
| FAM22D | midnightblue | -0.331982 | 0.097539 | UBA5 | NA | -0.128681458 | 0.522382608 | CTR9 | magenta | -0.179068 | 0.381426 |
| APOBEC3G | midnightblue | -0.306242 | 0.128119 | HK2 | NA | -0.128640262 | 0.522516902 | ANKIB1 | magenta | -0.178808 | 0.382129 |
| CD3G | midnightblue | -0.299678 | 0.136925 | NSMCE4A | NA | -0.128555044 | 0.522794755 | IST1 | magenta | -0.177198 | 0.386502 |
| IRF4 | midnightblue | -0.299417 | 0.137284 | TMEM135 | NA | -0.128470474 | 0.523070564 | RBL2 | magenta | -0.176264 | 0.38905 |
| MSRB1 | midnightblue | -0.298916 | 0.137975 | PITPNC1 | NA | -0.128279041 | 0.523695149 | ESCO1 | magenta | -0.175789 | 0.390348 |
| CCL5 | midnightblue | -0.298589 | 0.138427 | FRYL | NA | -0.128045614 | 0.524457232 | PRKCH | magenta | -0.173915 | 0.395501 |
| MREG | midnightblue | -0.298228 | 0.138928 | RNASE6 | NA | -0.127760419 | 0.52538905 | EDEM1 | magenta | -0.173825 | 0.395748 |
| BIRC3 | midnightblue | -0.296819 | 0.140894 | LINC00152 | NA | -0.127596447 | 0.525925154 | VGLL3 | magenta | -0.173763 | 0.395919 |
| P2RY11 | midnightblue | -0.289647 | 0.151213 | LOC643837 | NA | -0.127534153 | 0.526128892 | YARS | magenta | -0.171645 | 0.401793 |
| CLEC2D | midnightblue | -0.279674 | 0.166444 | SERPINB6 | NA | -0.127444645 | 0.526421702 | MPC1 | magenta | -0.169651 | 0.407364 |
| SQLE | midnightblue | -0.278922 | 0.167633 | SPATA20 | NA | -0.12735021 | 0.526730717 | PTPN12 | magenta | -0.168561 | 0.410428 |
| DUSP4 | midnightblue | -0.276476 | 0.171548 | CYFIP1 | NA | -0.127279189 | 0.52696317 | GGH | magenta | -0.168434 | 0.410786 |
| PKDCC | midnightblue | -0.272636 | 0.17782 | LOC654433 | NA | -0.12709268 | 0.527573854 | TYW3 | magenta | -0.167521 | 0.413364 |
| CD86 | midnightblue | -0.269054 | 0.183814 | FBXW11 | NA | -0.126898497 | 0.528210026 | TPGS2 | magenta | -0.164906 | 0.420797 |
| COL9A2 | midnightblue | 0.268326 | 0.185048 | DNAJC15 | NA | -0.126639993 | 0.529057492 | SSB | magenta | -0.164821 | 0.421041 |
| TANK | midnightblue | -0.267966 | 0.185662 | GTF2IRD1 | NA | -0.126618563 | 0.529127777 | SRSF5 | magenta | -0.164418 | 0.422193 |
| ZC3H12D | midnightblue | -0.265725 | 0.189508 | TBXAS1 | NA | -0.126594777 | 0.52920579 | SREK1IP1 | magenta | -0.16413 | 0.423018 |
| NKG7 | midnightblue | -0.263741 | 0.192959 | RBP4 | NA | -0.126550508 | 0.529351006 | GNL1 | magenta | -0.160795 | 0.432631 |
| RSU1 | midnightblue | -0.241801 | 0.23402 | MTA1 | NA | -0.12632272 | 0.530098507 | BZW2 | magenta | -0.159795 | 0.435534 |
| RFTN1 | midnightblue | -0.236781 | 0.244171 | SPRED1 | NA | -0.126289983 | 0.530205979 | NCKAP1L | magenta | -0.158763 | 0.438543 |
| EIF4E3 | midnightblue | -0.236321 | 0.245115 | HPD | NA | -0.126217484 | 0.530444015 | VPS37B | magenta | -0.158666 | 0.438828 |
| INPP5D | midnightblue | 0.235299 | 0.247221 | CD300A | NA | -0.126203805 | 0.530488936 | FUBP1 | magenta | -0.155516 | 0.448085 |
| SPG21 | midnightblue | -0.234857 | 0.248137 | PLXNA3 | NA | -0.126182119 | 0.530560152 | HKR1 | magenta | -0.154111 | 0.452245 |
| ARID5B | midnightblue | -0.228588 | 0.261351 | ACAT2 | NA | -0.125856013 | 0.531631609 | HMGN2 | magenta | -0.153247 | 0.454815 |
| CD96 | midnightblue | -0.212134 | 0.298165 | GPR68 | NA | -0.125813695 | 0.531770728 | TGFBR1 | magenta | -0.151341 | 0.460511 |
| ANXA6 | midnightblue | -0.211247 | 0.300236 | MAPK1 | NA | -0.125772968 | 0.531904628 | EIF3L | magenta | -0.150624 | 0.462663 |
| FCRL1 | midnightblue | 0.210437 | 0.302136 | TNFRSF14 | NA | -0.125676079 | 0.532223245 | BMP2K | magenta | -0.148818 | 0.468109 |
| TAGAP | midnightblue | -0.207626 | 0.308789 | RETN | NA | -0.1254426 | 0.532991405 | NEDD4L | magenta | -0.148153 | 0.470122 |
| CD209 | midnightblue | -0.20514 | 0.314746 | LIMD2 | NA | -0.124953658 | 0.534601757 | TCEAL4 | magenta | -0.148008 | 0.470561 |
| TNFRSF9 | midnightblue | -0.204425 | 0.316472 | ATP6V1F | NA | -0.124864877 | 0.534894407 | ATP10D | magenta | -0.145984 | 0.476718 |
| CDCP1 | midnightblue | -0.197931 | 0.33242 | CA13 | NA | -0.124549952 | 0.535933114 | DYX1C1 | magenta | -0.14477 | 0.480431 |
| CRIP1 | midnightblue | -0.197435 | 0.333657 | CHD1L | NA | -0.124360562 | 0.536558231 | TARDBP | magenta | -0.143901 | 0.483096 |
| TAGLN2 | midnightblue | -0.195196 | 0.339278 | FGD6 | NA | -0.124296746 | 0.536768944 | HS1BP3 | magenta | -0.143428 | 0.484552 |
| CLSTN3 | midnightblue | -0.186111 | 0.362664 | GBA2 | NA | -0.123790809 | 0.53844088 | WAC | magenta | -0.142989 | 0.485903 |
| CD226 | midnightblue | -0.184678 | 0.366436 | PBX4 | NA | -0.123582738 | 0.539129186 | ATP2A3 | magenta | -0.139739 | 0.495972 |
| PXN | midnightblue | 0.182317 | 0.372702 | ZNF835 | NA | -0.123409247 | 0.53970342 | SAR1A | magenta | -0.13939 | 0.497058 |
| IL18 | midnightblue | -0.181799 | 0.374085 | THBS1 | NA | -0.123044506 | 0.540911601 | RAD17 | magenta | -0.138753 | 0.499048 |
| KLRK1 | midnightblue | -0.181455 | 0.375006 | GLT1D1 | NA | -0.122745519 | 0.541902922 | DNMT3A | magenta | -0.137025 | 0.504459 |
| SLAMF6 | midnightblue | -0.177143 | 0.386649 | ST3GAL4 | NA | -0.122340344 | 0.543247673 | CLP1 | magenta | -0.133843 | 0.514501 |
| C3 | midnightblue | -0.174159 | 0.394828 | PTPRC | NA | -0.122112208 | 0.544005529 | RPP14 | magenta | -0.133551 | 0.515428 |
| ZDHHC11 | midnightblue | -0.172463 | 0.399519 | YIPF1 | NA | -0.122109754 | 0.544013684 | TTC38 | magenta | -0.131797 | 0.521008 |
| ITK | midnightblue | -0.172187 | 0.400283 | ZFAND6 | NA | -0.121388121 | 0.546414171 | CHMP4C | magenta | -0.128703 | 0.530922 |

**Table 6. Gene module membership and correlation with growth rate (48/72h), with adjusted p values for each module**

| | | | |
|---|---|---|---|
| BATF | midnightblue | -0.169975 | 0.406455 |
| DHCR7 | midnightblue | -0.163333 | 0.425304 |
| RCSD1 | midnightblue | -0.163296 | 0.425411 |
| NLRP12 | midnightblue | -0.160739 | 0.432792 |
| DDB2 | midnightblue | 0.160335 | 0.433964 |
| SMAP2 | midnightblue | -0.15593 | 0.446862 |
| ZNF626 | midnightblue | -0.151801 | 0.459134 |
| SLC25A43 | midnightblue | -0.149821 | 0.465079 |
| PNKD | midnightblue | 0.14885 | 0.468012 |
| SMAD3 | midnightblue | -0.148596 | 0.468778 |
| APP | midnightblue | -0.147997 | 0.470594 |
| HMGCR | midnightblue | 0.140263 | 0.494341 |
| ZNF431 | midnightblue | -0.139144 | 0.497827 |
| COL6A1 | midnightblue | -0.138807 | 0.498879 |
| UBE3D | midnightblue | -0.138152 | 0.500925 |
| PLA2G7 | midnightblue | -0.135206 | 0.510188 |
| CAB39 | midnightblue | -0.132742 | 0.517997 |
| PDLIM2 | midnightblue | -0.129176 | 0.529402 |
| RASGRP1 | midnightblue | -0.125162 | 0.542379 |
| SYNE1 | midnightblue | -0.12467 | 0.54398 |
| C7orf41 | midnightblue | -0.123685 | 0.547192 |
| C15orf48 | midnightblue | -0.122767 | 0.550194 |
| CD84 | midnightblue | -0.118935 | 0.5628 |
| MLLT11 | midnightblue | 0.118444 | 0.564426 |
| PAFAH1B3 | midnightblue | -0.117269 | 0.568321 |
| ST6GAL1 | midnightblue | -0.113661 | 0.580363 |
| FAM186B | midnightblue | 0.109984 | 0.592748 |
| FANCE | midnightblue | -0.109378 | 0.5948 |
| RNASE6 | midnightblue | -0.105463 | 0.608131 |
| KMO | midnightblue | -0.104553 | 0.611247 |
| ARHGAP31 | midnightblue | -0.103639 | 0.614383 |
| ABCG2 | midnightblue | 0.098566 | 0.631908 |
| ITLN1 | midnightblue | -0.097655 | 0.635078 |
| ENPP2 | midnightblue | 0.096357 | 0.639603 |
| S100Z | midnightblue | 0.095731 | 0.641791 |
| KIAA1958 | midnightblue | 0.095514 | 0.642549 |
| IL32 | midnightblue | -0.088403 | 0.667601 |
| ANKRD22 | midnightblue | -0.084332 | 0.682106 |
| S100A13 | midnightblue | 0.083754 | 0.684172 |
| CHI3L1 | midnightblue | -0.081908 | 0.690792 |
| S100A1 | midnightblue | 0.080656 | 0.695294 |
| RNF125 | midnightblue | -0.076879 | 0.708941 |
| PLA1A | midnightblue | -0.07654 | 0.710169 |
| KCNJ5 | midnightblue | -0.076125 | 0.711675 |
| LILRB4 | midnightblue | 0.075918 | 0.712426 |
| SLAMF8 | midnightblue | 0.073911 | 0.719723 |
| ACTA2 | midnightblue | 0.07369 | 0.720527 |
| CD3E | midnightblue | -0.067348 | 0.74375 |
| ATP6V0D2 | midnightblue | -0.060585 | 0.768757 |
| SIGLEC5 | midnightblue | 0.055189 | 0.788873 |
| GPR82 | midnightblue | 0.053524 | 0.795106 |
| IFNG | midnightblue | -0.04865 | 0.813429 |
| FBN1 | midnightblue | 0.047978 | 0.815961 |
| PPBP | midnightblue | 0.047428 | 0.818035 |
| TMEM14A | midnightblue | 0.046904 | 0.820015 |
| DHCR24 | midnightblue | 0.046191 | 0.822708 |
| CRIP1 | NA | -0.12131378 | 0.546661744 |
| CAB39 | NA | -0.121130901 | 0.547270997 |
| IPO4 | NA | -0.12105881 | 0.547511247 |
| KIAA0355 | NA | -0.120935066 | 0.547923756 |
| AMPD3 | NA | -0.120859179 | 0.548176802 |
| ZNF107 | NA | -0.120840728 | 0.548238336 |
| TEX10 | NA | -0.119886529 | 0.551424896 |
| CPEB2 | NA | -0.119867233 | 0.551489423 |
| WDFY1 | NA | -0.119862767 | 0.551504358 |
| PPP1R2 | NA | -0.119721609 | 0.551976514 |
| ACOT9 | NA | -0.119351912 | 0.553213981 |
| ZNF768 | NA | -0.119294621 | 0.553405864 |
| ZNF211 | NA | -0.119280043 | 0.553454694 |
| CCT6P3 | NA | -0.119031317 | 0.554288124 |
| DPYSL3 | NA | -0.118584611 | 0.555786396 |
| ATXN7L1 | NA | -0.118492318 | 0.556096182 |
| NPHP3 | NA | -0.118360158 | 0.55653992 |
| CDK6 | NA | -0.11818435 | 0.557130462 |
| CCR7 | NA | -0.117924291 | 0.55800453 |
| ATP5SL | NA | -0.117392147 | 0.559795031 |
| BRD4 | NA | -0.117054438 | 0.560932672 |
| C11orf57 | NA | -0.116770099 | 0.56189134 |
| CA2 | NA | -0.116720008 | 0.562060303 |
| SUV420H1 | NA | -0.116640842 | 0.562327387 |
| MICB | NA | -0.116513617 | 0.562756724 |
| LILRB4 | NA | -0.116503249 | 0.56279172 |
| ANKLE2 | NA | -0.115911792 | 0.56478968 |
| TBPL1 | NA | -0.115138017 | 0.567408338 |
| NRBP2 | NA | -0.114931038 | 0.568109732 |
| TRIP4 | NA | -0.11481073 | 0.568517602 |
| SERPINB7 | NA | -0.114488288 | 0.569611397 |
| CSTA | NA | -0.114485488 | 0.5696209 |
| MFSD2A | NA | -0.114437324 | 0.569784364 |
| TIMM22 | NA | -0.11433461 | 0.570133037 |
| TEF | NA | -0.113778083 | 0.572023888 |
| NQO1 | NA | -0.11372017 | 0.572220814 |
| C14orf93 | NA | -0.113160404 | 0.574125775 |
| ENTPD7 | NA | -0.113045645 | 0.574516663 |
| FMO4 | NA | -0.112763627 | 0.575477766 |
| PKD1 | NA | -0.112343814 | 0.576909789 |
| ZRANB1 | NA | -0.112130697 | 0.577637353 |
| PCNXL2 | NA | -0.112046376 | 0.577925329 |
| WDR24 | NA | -0.111876293 | 0.578506397 |
| L3MBTL4 | NA | -0.111624782 | 0.579366126 |
| PRDX2 | NA | -0.111512712 | 0.579749393 |
| SHPK | NA | -0.11150692 | 0.579769201 |
| KDM4C | NA | -0.111457008 | 0.579939933 |
| MTX2 | NA | -0.111320259 | 0.580407816 |
| KIAA1462 | NA | -0.111171565 | 0.580916758 |
| ATP13A1 | NA | -0.110923534 | 0.581766139 |
| HEATR7A | NA | -0.110913906 | 0.581799122 |
| COMMD10 | NA | -0.110857073 | 0.581993827 |
| ELMOD2 | NA | -0.110351486 | 0.58372719 |
| ACVR1 | NA | -0.110213417 | 0.584200941 |
| KLHL24 | NA | -0.110103123 | 0.584579509 |
| PARP1 | NA | -0.109779234 | 0.585691825 |
| LEPRE1 | magenta | -0.128521 | 0.531509 |
| USP15 | magenta | -0.128325 | 0.532139 |
| ZMYND8 | magenta | -0.126588 | 0.537753 |
| PAFAH1B2 | magenta | -0.122319 | 0.55166 |
| SMARCA2 | magenta | -0.121694 | 0.553711 |
| RBM5 | magenta | -0.121524 | 0.554268 |
| PRMT6 | magenta | -0.121437 | 0.554554 |
| HDAC1 | magenta | -0.12075 | 0.556812 |
| MAN1A1 | magenta | -0.1204 | 0.557966 |
| SEL1L3 | magenta | -0.118818 | 0.563187 |
| LRRCC1 | magenta | -0.116883 | 0.569607 |
| MUT | magenta | -0.116334 | 0.571433 |
| TCEAL3 | magenta | -0.111648 | 0.587132 |
| ABHD11 | magenta | -0.108618 | 0.597378 |
| HCAR3 | magenta | -0.107711 | 0.60046 |
| GTF2H5 | magenta | -0.107574 | 0.600927 |
| SLC25A17 | magenta | -0.107075 | 0.602627 |
| SIK2 | magenta | -0.104882 | 0.61012 |
| HAUS4 | magenta | -0.102447 | 0.618482 |
| DDX5 | magenta | -0.101389 | 0.622133 |
| FCER1G | magenta | -0.097571 | 0.635371 |
| CDC25B | magenta | -0.08832 | 0.667896 |
| STK24 | magenta | -0.086722 | 0.673576 |
| NDFIP1 | magenta | -0.083297 | 0.685808 |
| SPATS2L | magenta | -0.079665 | 0.698867 |
| BCAT2 | magenta | -0.076402 | 0.710668 |
| THUMPD3 | magenta | -0.068572 | 0.739252 |
| MECP2 | magenta | -0.063276 | 0.758779 |
| ZNF768 | magenta | -0.036485 | 0.859551 |
| BPNT1 | magenta | -0.033651 | 0.870367 |
| SS18 | magenta | -0.028017 | 0.891932 |
| GBAP1 | magenta | -0.026608 | 0.897338 |
| WASH1 | magenta | -0.024306 | 0.90618 |
| SLC47A1 | magenta | -0.011919 | 0.953918 |
| CSRP1 | magenta | -0.010384 | 0.959846 |
| LOC1005067[illegible] | magenta | 0.018448 | 0.928727 |
| XPO6 | magenta | 0.019894 | 0.923153 |
| PILRA | magenta | 0.024938 | 0.90375 |
| QRICH1 | magenta | 0.026112 | 0.899243 |
| HYI | magenta | 0.026247 | 0.898722 |
| TOP1 | magenta | 0.030141 | 0.883792 |
| SEC61A2 | magenta | 0.038145 | 0.853227 |
| MRPL9 | magenta | 0.038425 | 0.852161 |
| HARS | magenta | 0.038897 | 0.850366 |
| NDUFAF7 | magenta | 0.041496 | 0.840488 |
| KLC4 | magenta | 0.053443 | 0.79541 |
| LSP1 | magenta | 0.053925 | 0.793605 |
| SNAPC2 | magenta | 0.054886 | 0.790006 |
| KLHDC3 | magenta | 0.058228 | 0.777526 |
| ANAPC13 | magenta | 0.063394 | 0.758342 |
| TMEM199 | magenta | 0.066837 | 0.74563 |
| ILVBL | magenta | 0.067856 | 0.74188 |
| IFFO2 | magenta | 0.075847 | 0.712681 |
| LOC728743 | magenta | 0.079989 | 0.6977 |
| CEP164 | magenta | 0.083996 | 0.683308 |
| RPAIN | magenta | 0.085972 | 0.67625 |

**Table 6. Gene module membership and correlation with growth rate (48/72h), with adjusted p values for each module**

| | | | | | | | | | | | |
|---|---|---|---|---|---|---|---|---|---|---|---|
| GPD1 | midnightblue | -0.044464 | 0.829238 | CD36 | NA | -0.109743374 | 0.585815037 | SNAP47 | magenta | 0.09001 | 0.661907 |
| GPR155 | midnightblue | 0.038244 | 0.85285 | RUNX3 | NA | -0.109383869 | 0.58705086 | FAM117A | magenta | 0.104126 | 0.612711 |
| SCARF1 | midnightblue | 0.037647 | 0.855122 | FAM151B | NA | -0.109271538 | 0.587437239 | ZWINT | magenta | 0.104323 | 0.612035 |
| SLC2A3 | midnightblue | 0.036728 | 0.858626 | DBF4 | NA | -0.10885723 | 0.588863263 | ARF5 | magenta | 0.109643 | 0.593904 |
| CNRIP1 | midnightblue | 0.033576 | 0.870654 | CTDSP1 | NA | -0.108446387 | 0.59027884 | PRPF19 | magenta | 0.109754 | 0.593528 |
| LIPN | midnightblue | -0.03003 | 0.884219 | TTF2 | NA | -0.108060248 | 0.591610638 | PHACTR4 | magenta | 0.11465 | 0.577052 |
| KDM3B | midnightblue | -0.027389 | 0.894342 | TEFM | NA | -0.108054709 | 0.591629751 | TM9SF4 | magenta | 0.115801 | 0.573208 |
| LPAR3 | midnightblue | -0.026049 | 0.899483 | CPSF7 | NA | -0.108018646 | 0.591754201 | BCAS2 | magenta | 0.121714 | 0.553644 |
| GRK6 | midnightblue | 0.02496 | 0.903668 | C5orf44 | NA | -0.107959157 | 0.591959516 | ANKRD11 | magenta | 0.128049 | 0.533031 |
| SPATS2 | midnightblue | 0.022937 | 0.911444 | DYRK3 | NA | -0.107632258 | 0.593088289 | FZD7 | magenta | 0.132909 | 0.517467 |
| NCS1 | midnightblue | 0.020518 | 0.920753 | CCT7 | NA | -0.107533863 | 0.593428227 | IER3 | magenta | 0.143351 | 0.484787 |
| PTPN1 | midnightblue | -0.019251 | 0.925633 | ZNF524 | NA | -0.107276547 | 0.594317603 | PTGFRN | magenta | 0.149463 | 0.466159 |
| RXRB | midnightblue | -0.018391 | 0.928947 | SLC43A2 | NA | -0.107246331 | 0.594422078 | PRMT1 | magenta | 0.149691 | 0.465472 |
| GLI3 | midnightblue | -0.01363 | 0.947311 | MPG | NA | -0.10712518 | 0.59484105 | CCDC51 | magenta | 0.151389 | 0.460366 |
| TREM2 | midnightblue | 0.012848 | 0.950328 | SCARB1 | NA | -0.106921612 | 0.595545323 | DIRC2 | magenta | 0.156149 | 0.446215 |
| RTN4 | midnightblue | -0.012099 | 0.953224 | CSF1R | NA | -0.106490217 | 0.597038979 | SLC7A5 | magenta | 0.168188 | 0.411481 |
| PAM | midnightblue | 0.010708 | 0.958594 | IRF9 | NA | -0.106132812 | 0.598277663 | UBFD1 | magenta | 0.179528 | 0.380184 |
| KIFC3 | midnightblue | 0.010188 | 0.960604 | IL15 | NA | -0.105887313 | 0.599129137 | RAB7L1 | magenta | 0.180461 | 0.377673 |
| IQGAP3 | midnightblue | -0.009378 | 0.963733 | UNC119B | NA | -0.105517168 | 0.600413904 | EPHB6 | magenta | 0.19316 | 0.344438 |
| FADS2 | midnightblue | 0.006541 | 0.974702 | IMPDH1 | NA | -0.105490619 | 0.600506101 | TJAP1 | magenta | 0.193818 | 0.342766 |
| PLS3 | midnightblue | -0.005679 | 0.978033 | RAB11FIP5 | NA | -0.105015517 | 0.602156987 | MAPK9 | magenta | 0.193836 | 0.342721 |
| STON2 | midnightblue | 0.005245 | 0.979712 | GXYLT1 | NA | -0.1049351 | 0.602436611 | PPAP2B | magenta | 0.195548 | 0.338392 |
| RASSF4 | midnightblue | 0.00312 | 0.98793 | NRP1 | NA | -0.104802863 | 0.602896542 | TRPM2 | magenta | 0.196233 | 0.336667 |
| STARD4 | midnightblue | -0.002382 | 0.990785 | RFX2 | NA | -0.104546622 | 0.603788187 | DEPDC5 | magenta | 0.20819 | 0.307446 |
| HPGD | midnightblue | -0.001461 | 0.99435 | MINA | NA | -0.104308472 | 0.604617377 | TMED1 | magenta | 0.209078 | 0.305342 |
| HMGCS1 | midnightblue | -0.000935 | 0.996384 | LZIC | NA | -0.104201298 | 0.604990695 | DNAJC7 | magenta | 0.212108 | 0.298226 |
| LDLR | midnightblue | -0.000626 | 0.997578 | ZNF23 | NA | -0.104166042 | 0.605113523 | FEM1B | magenta | 0.218263 | 0.284092 |
| NDUFS7 | NA | -0.557033 | 0.003117 | ATF3 | NA | -0.103929251 | 0.60593874 | COG1 | magenta | 0.224759 | 0.269642 |
| TRIM21 | NA | -0.536492 | 0.004721 | CPE | NA | -0.103784793 | 0.606442412 | GPANK1 | magenta | 0.227392 | 0.263924 |
| TRAIP | NA | -0.522403 | 0.006186 | PGS1 | NA | -0.103463745 | 0.607562413 | ZNF324B | magenta | 0.227943 | 0.262737 |
| PAXIP1 | NA | -0.521699 | 0.006268 | TLR4 | NA | -0.103421975 | 0.607708193 | APOE | magenta | 0.228909 | 0.260664 |
| SEMA7A | NA | 0.515936 | 0.006977 | ZNF273 | NA | -0.103384061 | 0.607840532 | FOXP4 | magenta | 0.229417 | 0.25958 |
| OBFC1 | NA | -0.511675 | 0.007542 | AP2A2 | NA | -0.103373457 | 0.607877546 | MRPS26 | magenta | 0.230673 | 0.256907 |
| FIS1 | NA | -0.485236 | 0.011982 | VTN | NA | -0.103042943 | 0.609031726 | CRTC1 | magenta | 0.234263 | 0.24937 |
| FPR2 | NA | -0.479201 | 0.013252 | WDR55 | NA | -0.102973789 | 0.609273329 | CSF3R | magenta | 0.23905 | 0.239546 |
| ERLIN2 | NA | -0.470227 | 0.015346 | TP53BP1 | NA | -0.101874888 | 0.613117962 | SH3RF3-AS1 | magenta | 0.244762 | 0.228164 |
| EPB41L5 | NA | 0.467723 | 0.015976 | FIGNL1 | NA | -0.101746271 | 0.613568601 | KLHL22 | magenta | 0.246413 | 0.224943 |
| MT1L | NA | -0.463543 | 0.017076 | ZNF671 | NA | -0.101592058 | 0.614109102 | SLC25A37 | magenta | 0.247441 | 0.222952 |
| RPS27 | NA | -0.461728 | 0.017572 | C1GALT1C1 | NA | -0.101469342 | 0.614539346 | TXNRD2 | magenta | 0.247672 | 0.222506 |
| CISD1 | NA | -0.458944 | 0.018356 | LLGL1 | NA | -0.101411603 | 0.614741827 | TIGD5 | magenta | 0.253505 | 0.21145 |
| OAZ1 | NA | -0.45236 | 0.020326 | SIGLEC9 | NA | -0.101166062 | 0.615603191 | STARD10 | magenta | 0.256391 | 0.20612 |
| ZNF367 | NA | -0.451138 | 0.02071 | ZKSCAN5 | NA | -0.101139957 | 0.615694798 | WIPI1 | magenta | 0.257008 | 0.204992 |
| ZNF428 | NA | -0.450383 | 0.02095 | ING5 | NA | -0.101135276 | 0.615711225 | ZBTB2 | magenta | 0.258544 | 0.202204 |
| KIAA1328 | NA | -0.449498 | 0.021235 | MOSPD3 | NA | -0.100534498 | 0.617821025 | ADRM1 | magenta | 0.266486 | 0.188195 |
| SEMA4B | NA | -0.446948 | 0.022071 | MSRB1 | NA | -0.100173975 | 0.619088523 | ZBTB5 | magenta | 0.271217 | 0.180178 |
| FAM111B | NA | -0.441259 | 0.024035 | LILRB3 | NA | -0.100168599 | 0.619107432 | LOC1005063 | magenta | 0.271429 | 0.179824 |
| OXNAD1 | NA | -0.440663 | 0.024249 | PMPCB | NA | -0.099845698 | 0.62024358 | CSF2RA | magenta | 0.276015 | 0.172293 |
| RFPL4A | NA | -0.438446 | 0.025057 | RBM27 | NA | -0.099815019 | 0.62035157 | RNF207 | magenta | 0.277958 | 0.169169 |
| C16orf13 | NA | -0.434912 | 0.026391 | UBE2R2 | NA | -0.099528534 | 0.621360374 | ARNT | magenta | 0.28146 | 0.16364 |
| PLAUR | NA | 0.434848 | 0.026416 | FKBP14 | NA | -0.0990909 | 0.622902705 | LRRC14 | magenta | 0.287899 | 0.153808 |
| ZNF622 | NA | -0.43227 | 0.027425 | RSU1 | NA | -0.098942628 | 0.623425604 | TP53 | magenta | 0.28937 | 0.151623 |
| UQCR11 | NA | -0.431661 | 0.027668 | COX7C | NA | -0.098938842 | 0.623438959 | PIAS3 | magenta | 0.29655 | 0.141272 |
| HMGN1 | NA | -0.428559 | 0.028933 | FNIP2 | NA | -0.098773733 | 0.624021454 | DDX3X | magenta | 0.297631 | 0.139759 |
| WDR73 | NA | -0.420753 | 0.032322 | TAP2 | NA | -0.09849932 | 0.624990054 | YIPF3 | magenta | 0.298759 | 0.138192 |
| HLA-A | NA | -0.4198 | 0.032757 | TRUB2 | NA | -0.098458127 | 0.625135505 | CUEDC2 | magenta | 0.300263 | 0.136122 |
| TDP1 | NA | -0.417774 | 0.033696 | KCNK13 | NA | -0.098272426 | 0.625791381 | ACSS1 | magenta | 0.302833 | 0.132639 |

**Table 6. Gene module membership and correlation with growth rate (48/72h), with adjusted p values for each module**

| | | | | | | | | | | | |
|---|---|---|---|---|---|---|---|---|---|---|---|
| C7orf49 | NA | -0.414459 | 0.03528 | TNS3 | NA | -0.09744849 | 0.628704789 | SNRPA | magenta | 0.304659 | 0.130203 |
| ZFAND2B | NA | -0.414138 | 0.035436 | TNFRSF10A | NA | -0.097364527 | 0.629001985 | TMEM219 | magenta | 0.308076 | 0.125733 |
| DDT | NA | -0.411825 | 0.03658 | KLHDC8B | NA | -0.097171702 | 0.629684725 | KCTD13 | magenta | 0.310697 | 0.122382 |
| WHAMMP1 | NA | -0.409291 | 0.037867 | GADD45G | NA | -0.09707432 | 0.630029639 | C14orf159 | magenta | 0.311662 | 0.121164 |
| SNAP47 | NA | -0.409035 | 0.037999 | ATP5J2 | NA | -0.096913412 | 0.630599721 | MAP2K7 | magenta | 0.31292 | 0.11959 |
| TACO1 | NA | -0.408109 | 0.03848 | MKS1 | NA | -0.096667411 | 0.631471679 | DVL1 | magenta | 0.314362 | 0.117804 |
| MCM6 | NA | -0.406298 | 0.039433 | CISD1 | NA | -0.096567779 | 0.631824965 | SH3BP2 | magenta | 0.315957 | 0.115852 |
| COA4 | NA | -0.40365 | 0.040861 | PRIM1 | NA | -0.096536688 | 0.631935226 | GPBP1L1 | magenta | 0.317711 | 0.113732 |
| MFSD11 | NA | -0.40266 | 0.041405 | ZNF227 | NA | -0.096523191 | 0.631983095 | TMEM52B | magenta | 0.331745 | 0.097793 |
| SSX1 | NA | -0.399116 | 0.043401 | RALGDS | NA | -0.096388304 | 0.632461565 | EYA3 | magenta | 0.341835 | 0.087409 |
| OC1001328 | NA | -0.396087 | 0.045166 | TSPYL2 | NA | -0.096085572 | 0.633535939 | PAG1 | magenta | 0.356491 | 0.073838 |
| USPL1 | NA | 0.395981 | 0.045228 | GPR183 | NA | -0.095979977 | 0.633910857 | USP28 | magenta | 0.361871 | 0.069283 |
| PRX | NA | -0.392565 | 0.047289 | TRIM4 | NA | -0.095780188 | 0.634620459 | STAT5A | magenta | 0.362108 | 0.069088 |
| WIBG | NA | -0.391663 | 0.047845 | PLIN3 | NA | -0.095751789 | 0.634721351 | AP1M1 | magenta | 0.362705 | 0.068597 |
| BLOC1S6 | NA | 0.389613 | 0.049128 | SH3BP5 | NA | -0.095358934 | 0.636117682 | CNPY3 | magenta | 0.378793 | 0.056354 |
| DNAJC9 | NA | -0.388667 | 0.049729 | ZBTB4 | NA | -0.094841098 | 0.637960086 | TMCC3 | magenta | 0.393476 | 0.046732 |
| SEMA4A | NA | 0.383989 | 0.052787 | IDNK | NA | -0.094674611 | 0.638552879 | COMT | magenta | 0.399694 | 0.04307 |
| MIS18A | NA | -0.383122 | 0.05337 | TBC1D30 | NA | -0.094458405 | 0.639323022 | RHBDD3 | magenta | 0.403972 | 0.040685 |
| BMS1 | NA | -0.380546 | 0.05513 | JAGN1 | NA | -0.09426171 | 0.640023981 | USP4 | magenta | 0.40399 | 0.040675 |
| RAD51AP1 | NA | -0.379186 | 0.056078 | NOTCH2 | NA | -0.094145971 | 0.640436579 | RRNAD1 | magenta | 0.405226 | 0.040006 |
| GUK1 | NA | -0.375868 | 0.058443 | LOC729852 | NA | -0.094139979 | 0.640457945 | CAMKK2 | magenta | 0.434917 | 0.026389 |
| GTF2E1 | NA | -0.375123 | 0.058985 | B9D2 | NA | -0.094069352 | 0.640709777 | H1F0 | magenta | 0.482883 | 0.012465 |
| ETFB | NA | -0.375034 | 0.05905 | PML | NA | -0.093392805 | 0.643124093 | ALG1 | midnightblue | -0.502489 | 0.008894 |
| IDE | NA | -0.373604 | 0.060101 | RUNDC1 | NA | -0.093046464 | 0.64436141 | ILF2 | midnightblue | -0.411874 | 0.036556 |
| LYSMD2 | NA | -0.373242 | 0.060369 | FUK | NA | -0.092861515 | 0.64502253 | DHRS7 | midnightblue | -0.393657 | 0.046622 |
| EMR1 | NA | -0.373139 | 0.060446 | COX19 | NA | -0.092614381 | 0.645906346 | DCTN3 | midnightblue | -0.356606 | 0.073738 |
| HKR1 | NA | -0.372193 | 0.061152 | CISH | NA | -0.092487154 | 0.646361524 | FIS1 | midnightblue | -0.350189 | 0.079461 |
| CDKN2AIP | NA | -0.370341 | 0.062554 | ANO8 | NA | -0.092346862 | 0.646863591 | ARRB1 | midnightblue | -0.342574 | 0.086683 |
| LBR | NA | -0.370272 | 0.062607 | BTN3A2 | NA | -0.092167494 | 0.64750572 | COX5A | midnightblue | -0.341967 | 0.087279 |
| ENSA | NA | -0.37012 | 0.062723 | MOCOS | NA | -0.091894934 | 0.648481943 | RNASEK | midnightblue | -0.332026 | 0.097491 |
| TBCE | NA | -0.369204 | 0.063426 | MED26 | NA | -0.091065202 | 0.651457265 | ANP32A | midnightblue | -0.32985 | 0.099842 |
| BPNT1 | NA | -0.368467 | 0.063997 | LINC00493 | NA | -0.090572273 | 0.653227326 | ZNF440 | midnightblue | -0.329568 | 0.10015 |
| TDRD7 | NA | -0.368156 | 0.064239 | HFE | NA | -0.090437576 | 0.653711329 | CYBASC3 | midnightblue | -0.298803 | 0.138131 |
| TRMT112 | NA | -0.368102 | 0.064282 | XIST | NA | -0.090094533 | 0.654944592 | ZNF100 | midnightblue | -0.293583 | 0.145486 |
| FABP3 | NA | 0.367792 | 0.064523 | SLC16A10 | NA | -0.089950014 | 0.655464414 | EXOC2 | midnightblue | -0.293307 | 0.145882 |
| SERP1 | NA | -0.366008 | 0.065931 | HMGB1 | NA | -0.089600028 | 0.656723928 | RILP | midnightblue | -0.288735 | 0.152564 |
| PARL | NA | -0.365809 | 0.066089 | RBPJ | NA | -0.089483934 | 0.657141924 | OR2A1 | midnightblue | -0.284706 | 0.158629 |
| MT2A | NA | -0.364989 | 0.066745 | TMTC2 | NA | -0.089361653 | 0.657582306 | FAM127A | midnightblue | -0.284184 | 0.159427 |
| ZNF187 | NA | -0.363901 | 0.067622 | FAM83G | NA | -0.089074146 | 0.65861817 | RRP9 | midnightblue | -0.279357 | 0.166944 |
| CENPO | NA | -0.363552 | 0.067905 | DNAJB11 | NA | -0.088962486 | 0.659020638 | TRAPPC2P1 | midnightblue | -0.277219 | 0.170352 |
| VPS41 | NA | -0.362364 | 0.068877 | LNX2 | NA | -0.088845047 | 0.659444037 | IL18BP | midnightblue | -0.275472 | 0.173173 |
| TEX14 | NA | 0.362219 | 0.068996 | AGL | NA | -0.088826094 | 0.659512378 | IFT57 | midnightblue | -0.270618 | 0.181181 |
| WDR76 | NA | -0.362151 | 0.069053 | PPP2R5D | NA | -0.088817541 | 0.659543219 | NBPF10 | midnightblue | -0.266117 | 0.188832 |
| HAVCR1 | NA | 0.360387 | 0.070518 | DEF6 | NA | -0.088755559 | 0.659766732 | MRPL54 | midnightblue | -0.265346 | 0.190164 |
| POLD3 | NA | -0.359724 | 0.071075 | IL27RA | NA | -0.088703164 | 0.659955698 | TUBB4B | midnightblue | -0.264166 | 0.192217 |
| UBL5 | NA | -0.358166 | 0.072396 | MGC2752 | NA | -0.088430288 | 0.660940168 | C1orf122 | midnightblue | -0.262368 | 0.195372 |
| SNHG9 | NA | 0.357564 | 0.072912 | CDADC1 | NA | -0.087968923 | 0.662605917 | LOC1001322 | midnightblue | -0.261735 | 0.196492 |
| LILRA5 | NA | -0.356871 | 0.073509 | C19orf24 | NA | -0.087842443 | 0.663062844 | STX6 | midnightblue | -0.261725 | 0.19651 |
| MS4A7 | NA | -0.355912 | 0.074342 | GBP2 | NA | -0.087441642 | 0.664511569 | TM9SF2 | midnightblue | -0.259183 | 0.201051 |
| MKKS | NA | -0.355718 | 0.074511 | DPH1 | NA | -0.087221545 | 0.665307628 | TPRA1 | midnightblue | -0.258308 | 0.202629 |
| C14orf142 | NA | -0.355435 | 0.074758 | RPAP3 | NA | -0.087171219 | 0.665489698 | PYCR2 | midnightblue | -0.257056 | 0.204904 |
| FUNDC1 | NA | -0.353756 | 0.07624 | OLR1 | NA | -0.087064304 | 0.665876563 | RELA | midnightblue | -0.255528 | 0.207704 |
| FCGR1A | NA | -0.353258 | 0.076683 | SERPINE2 | NA | -0.08701211 | 0.666065453 | ACOT8 | midnightblue | -0.254573 | 0.209467 |
| HNRNPC | NA | -0.352578 | 0.077293 | IDS | NA | -0.086937284 | 0.666336284 | CD74 | midnightblue | -0.253376 | 0.211691 |
| CBR1 | NA | -0.352212 | 0.077622 | TRIM68 | NA | -0.086838563 | 0.666693661 | ABCB7 | midnightblue | -0.253208 | 0.212004 |
| ZBP1 | NA | -0.351004 | 0.078716 | LOC1005066 | NA | -0.086795694 | 0.666848874 | LOC653501 | midnightblue | -0.252711 | 0.212932 |

**Table 6. Gene module membership and correlation with growth rate (48/72h), with adjusted p values for each module**

| | | | | | | | | | | | |
|---|---|---|---|---|---|---|---|---|---|---|---|
| SAYSD1 | NA | -0.350903 | 0.078808 | PIGM | NA | -0.086495394 | 0.66793652 | ZNF845 | midnightblue | -0.251002 | 0.216149 |
| FABP5 | NA | 0.350842 | 0.078864 | SMCR7L | NA | -0.086303573 | 0.66863161 | SLC2A8 | midnightblue | -0.248596 | 0.220729 |
| UROS | NA | -0.349909 | 0.079718 | PDSS1 | NA | -0.086212165 | 0.668962936 | ZNF625 | midnightblue | -0.245918 | 0.225906 |
| ZXDB | NA | 0.349171 | 0.080399 | KCNN4 | NA | -0.086185235 | 0.669060558 | MEF2D | midnightblue | -0.24589 | 0.225959 |
| KLHDC8B | NA | -0.347419 | 0.082033 | PLXND1 | NA | -0.08541266 | 0.671863453 | FAM21B | midnightblue | -0.245843 | 0.22605 |
| HLA-DRB1 | NA | 0.34677 | 0.082644 | GALM | NA | -0.085122249 | 0.672918171 | HIST2H2AC | midnightblue | -0.244556 | 0.228568 |
| AWAT2 | NA | 0.345997 | 0.083378 | IMMP2L | NA | -0.084864696 | 0.673854064 | SATB1 | midnightblue | -0.243959 | 0.229743 |
| CENPF | NA | -0.344749 | 0.084571 | FAM40B | NA | -0.084285268 | 0.675961305 | HERC3 | midnightblue | -0.243815 | 0.230026 |
| ASPHD2 | NA | -0.344645 | 0.084671 | CTSB | NA | -0.083962168 | 0.67713738 | PCSK5 | midnightblue | -0.241997 | 0.233628 |
| TAF1B | NA | -0.344572 | 0.084742 | CSF2RA | NA | -0.083850278 | 0.67754483 | TOMM7 | midnightblue | -0.236407 | 0.244939 |
| SESN3 | NA | -0.344258 | 0.085045 | CALM2 | NA | -0.083190328 | 0.679949846 | NOMO3 | midnightblue | -0.235852 | 0.246081 |
| ZBTB24 | NA | -0.344092 | 0.085205 | CSF2 | NA | -0.082819449 | 0.681302764 | ZNF598 | midnightblue | -0.233799 | 0.250335 |
| CCR1 | NA | -0.343984 | 0.08531 | SMPD1 | NA | -0.082792142 | 0.681402414 | MKRN2 | midnightblue | -0.226464 | 0.26593 |
| DHRS4 | NA | -0.343461 | 0.085817 | NIN | NA | -0.082577839 | 0.682184644 | C16orf88 | midnightblue | -0.223975 | 0.27136 |
| IL4I1 | NA | -0.342599 | 0.086658 | HES2 | NA | -0.082455152 | 0.682632608 | ZNF765 | midnightblue | -0.223117 | 0.27325 |
| FAM213A | NA | -0.3412 | 0.088037 | CALM1 | NA | -0.082273475 | 0.683296154 | LYPLA2 | midnightblue | -0.221529 | 0.276767 |
| EIF4G3 | NA | -0.33943 | 0.089806 | CD99 | NA | -0.08226952 | 0.683310602 | TECPR1 | midnightblue | -0.21434 | 0.29305 |
| SLC6A8 | NA | 0.335736 | 0.093581 | CEP135 | NA | -0.081920449 | 0.684586183 | PPP1R12C | midnightblue | -0.214312 | 0.293115 |
| PET117 | NA | -0.33527 | 0.094065 | C21orf7 | NA | -0.081520361 | 0.686049229 | RPL23AP7 | midnightblue | -0.213223 | 0.295633 |
| LINC00346 | NA | -0.33501 | 0.094337 | POLR3E | NA | -0.081194761 | 0.687240704 | CA2 | midnightblue | -0.212148 | 0.298132 |
| PRDM15 | NA | -0.334912 | 0.09444 | RRM2B | NA | -0.081098783 | 0.687592058 | CTSC | midnightblue | -0.210941 | 0.300953 |
| KIAA1731 | NA | -0.334433 | 0.094941 | ZNF232 | NA | -0.081034589 | 0.687827093 | CCDC58 | midnightblue | -0.209336 | 0.304731 |
| TPX2 | NA | -0.334227 | 0.095157 | CLCN3 | NA | -0.08079442 | 0.688706677 | LSM4 | midnightblue | -0.209262 | 0.304907 |
| NOD2 | NA | -0.333451 | 0.095975 | SLC45A4 | NA | -0.08056542 | 0.68954573 | LOC1001709: | midnightblue | -0.207428 | 0.309261 |
| DBT | NA | -0.331092 | 0.098495 | C20orf72 | NA | -0.080379626 | 0.690226734 | BASP1 | midnightblue | -0.198119 | 0.331951 |
| ERI3 | NA | -0.327839 | 0.102051 | DUSP4 | NA | -0.080197169 | 0.690895739 | FCGR2A | midnightblue | -0.189305 | 0.354336 |
| FOLR2 | NA | -0.327587 | 0.10233 | IL1A | NA | -0.079947763 | 0.691810592 | ARMC9 | midnightblue | -0.183382 | 0.369869 |
| HBB | NA | -0.326539 | 0.103498 | C20orf196 | NA | -0.079908268 | 0.691955502 | S100A8 | midnightblue | -0.179938 | 0.379079 |
| LOC439994 | NA | -0.326469 | 0.103576 | UNKL | NA | -0.079499257 | 0.693456825 | PRKCA | midnightblue | -0.178938 | 0.381779 |
| ANP32A | NA | -0.326292 | 0.103775 | BRD1 | NA | -0.079473348 | 0.693551966 | ERCC6 | midnightblue | -0.178868 | 0.381968 |
| RRM2 | NA | -0.325704 | 0.104435 | PDIA6 | NA | -0.079116163 | 0.694864042 | POLD3 | midnightblue | -0.1783 | 0.383506 |
| C7orf31 | NA | -0.325339 | 0.104847 | IRAK2 | NA | -0.078995852 | 0.695306186 | DHX9 | midnightblue | -0.162185 | 0.42861 |
| ACOT4 | NA | -0.324623 | 0.105659 | CAMKK1 | NA | -0.078987005 | 0.6953387 | RNU12 | midnightblue | -0.158832 | 0.438343 |
| PHKG2 | NA | 0.323969 | 0.106403 | VPS51 | NA | -0.078838101 | 0.695886067 | TIGD1 | midnightblue | -0.155714 | 0.4475 |
| ZSWIM7 | NA | -0.323183 | 0.107305 | LACTB | NA | -0.078318573 | 0.697796988 | HIST2H2BE | midnightblue | -0.155181 | 0.449076 |
| MVD | NA | 0.322962 | 0.107558 | CDKN2AIP | NA | -0.077843775 | 0.699544964 | ZNF844 | midnightblue | -0.152462 | 0.457156 |
| FPR3 | NA | -0.32162 | 0.109113 | RRAGC | NA | -0.077810073 | 0.699669095 | SRBD1 | midnightblue | -0.148175 | 0.470055 |
| PARP3 | NA | -0.320805 | 0.110065 | PABPC3 | NA | -0.077780894 | 0.699776571 | PEX11B | midnightblue | -0.146733 | 0.474435 |
| GNB4 | NA | -0.320325 | 0.110627 | LPAR1 | NA | -0.077748728 | 0.69989506 | TMTC2 | midnightblue | -0.146184 | 0.476108 |
| IBA57 | NA | 0.319998 | 0.111012 | RFX1 | NA | -0.077623675 | 0.700355769 | ACAA1 | midnightblue | -0.14026 | 0.49435 |
| UBP1 | NA | -0.319775 | 0.111275 | MAN2B2 | NA | -0.077224666 | 0.701826465 | LILRB2 | midnightblue | -0.137087 | 0.504263 |
| NARS2 | NA | -0.319252 | 0.111895 | DUS2L | NA | -0.077196472 | 0.70193042 | COG8 | midnightblue | -0.133036 | 0.517061 |
| SH2D4A | NA | 0.318744 | 0.112498 | ZSWIM4 | NA | -0.076720762 | 0.703685284 | ZSCAN18 | midnightblue | -0.131486 | 0.521999 |
| C17orf89 | NA | -0.31802 | 0.113362 | ZFP112 | NA | -0.07511588 | 0.709616469 | MYO1G | midnightblue | -0.127386 | 0.53517 |
| TNFSF10 | NA | -0.317676 | 0.113775 | TMEM55A | NA | -0.074357009 | 0.712426827 | CEP70 | midnightblue | -0.124141 | 0.545704 |
| RRAGD | NA | -0.31712 | 0.114443 | FAM210A | NA | -0.074338409 | 0.712495753 | BEX2 | midnightblue | -0.122422 | 0.551321 |
| ENTPD1 | NA | -0.317003 | 0.114585 | CHIT1 | NA | -0.073918204 | 0.714053547 | GAS5 | midnightblue | -0.112505 | 0.584246 |
| C11orf48 | NA | -0.31692 | 0.114685 | ACN9 | NA | -0.073742644 | 0.714704719 | IFITM2 | midnightblue | -0.109347 | 0.594905 |
| HIST1H4C | NA | -0.316133 | 0.115638 | ARHGEF7 | NA | -0.073574245 | 0.715329514 | FAM100A | midnightblue | -0.100287 | 0.62594 |
| ZNF33B | NA | -0.316066 | 0.11572 | CDK5RAP3 | NA | -0.073078022 | 0.717171638 | RNF149 | midnightblue | -0.093299 | 0.650313 |
| SEPW1 | NA | -0.315911 | 0.115907 | TRAPPC9 | NA | -0.072815594 | 0.718146472 | FCGR1C | midnightblue | -0.092436 | 0.653349 |
| TMEM201 | NA | -0.315908 | 0.115911 | GABARAPL1 | NA | -0.072774703 | 0.718298406 | VDAC1 | midnightblue | -0.09126 | 0.657492 |
| MRPL55 | NA | -0.314633 | 0.11747 | SDCCAG8 | NA | -0.072623333 | 0.718860935 | ZNF468 | midnightblue | -0.079859 | 0.698168 |
| EIF2B3 | NA | -0.314611 | 0.117498 | MB21D2 | NA | -0.072266021 | 0.720189352 | THOC6 | midnightblue | -0.0783 | 0.703796 |
| CCT7 | NA | -0.313591 | 0.118756 | NOTCH3 | NA | -0.072152099 | 0.72061306 | TSPAN4 | midnightblue | -0.07718 | 0.707847 |
| MAP7 | NA | 0.311345 | 0.121562 | GBP1P1 | NA | -0.071799893 | 0.721923519 | DAB2 | midnightblue | -0.073892 | 0.719791 |

**Table 6. Gene module membership and correlation with growth rate (48/72h), with adjusted p values for each module**

| | | | | | | | | | | | |
|---|---|---|---|---|---|---|---|---|---|---|---|
| IDH2 | NA | -0.311103 | 0.121868 | FES | NA | -0.071785702 | 0.721976336 | CAPZB | midnightblue | -0.066761 | 0.745913 |
| SERPINB6 | NA | -0.310542 | 0.122578 | LTV1 | NA | -0.071742444 | 0.722137346 | SNX11 | midnightblue | -0.063544 | 0.757788 |
| CCDC144A | NA | 0.310042 | 0.123213 | MITD1 | NA | -0.071485133 | 0.723095304 | PDHX | midnightblue | -0.062997 | 0.75981 |
| POLR2G | NA | -0.309085 | 0.124436 | ERMP1 | NA | -0.071005511 | 0.724882006 | NCEH1 | midnightblue | -0.057363 | 0.780752 |
| MMAA | NA | -0.309078 | 0.124444 | INPP4A | NA | -0.070703195 | 0.726008924 | YWHAZ | midnightblue | -0.055044 | 0.789417 |
| CCL2 | NA | -0.308369 | 0.125356 | LTBR | NA | -0.070541762 | 0.726610914 | RNF114 | midnightblue | -0.054913 | 0.789907 |
| TMEM51 | NA | -0.308036 | 0.125786 | NUP107 | NA | -0.070343342 | 0.727351046 | HMHA1 | midnightblue | -0.054276 | 0.792292 |
| )C1001320[illegible] | NA | -0.307478 | 0.126508 | MDFIC | NA | -0.070077967 | 0.728341302 | MGMT | midnightblue | -0.045925 | 0.823712 |
| NDUFV3 | NA | -0.306452 | 0.127843 | SLC22A5 | NA | -0.069648642 | 0.729944243 | SPRY2 | midnightblue | -0.043025 | 0.834687 |
| DDX26B | NA | -0.306307 | 0.128034 | PPP6R3 | NA | -0.069083245 | 0.732056915 | HRSP12 | midnightblue | -0.037549 | 0.855498 |
| STK17B | NA | -0.30562 | 0.128935 | LRRFIP1 | NA | -0.068983219 | 0.732430875 | MYO1E | midnightblue | -0.034341 | 0.867729 |
| ZNF512 | NA | -0.305507 | 0.129083 | CCL7 | NA | -0.068863145 | 0.732879863 | TFEC | midnightblue | -0.032024 | 0.876585 |
| CSRP2BP | NA | -0.304569 | 0.130323 | PSMD5 | NA | -0.06873537 | 0.733357743 | PHACTR1 | midnightblue | -0.03193 | 0.876944 |
| NSMCE4A | NA | -0.304207 | 0.130803 | ERCC2 | NA | -0.068389422 | 0.734652073 | PREP | midnightblue | -0.029637 | 0.885721 |
| CCBL2 | NA | -0.302221 | 0.133463 | ANKRD22 | NA | -0.068285015 | 0.735042844 | SECTM1 | midnightblue | -0.023479 | 0.90936 |
| CLK2 | NA | -0.301932 | 0.133853 | AHCTF1 | NA | -0.067671307 | 0.737341092 | ZSCAN22 | midnightblue | -0.023023 | 0.911114 |
| NDUFB2 | NA | -0.301908 | 0.133886 | SPIN1 | NA | -0.067655723 | 0.737399481 | TMEM42 | midnightblue | -0.017964 | 0.93059 |
| PLSCR1 | NA | -0.301692 | 0.134178 | C2orf49 | NA | -0.067635289 | 0.737476045 | NAPEPLD | midnightblue | -0.014636 | 0.943427 |
| FKBP4 | NA | 0.301469 | 0.13448 | TRIM11 | NA | -0.067578322 | 0.7376895 | DALRD3 | midnightblue | -0.007439 | 0.971229 |
| ZNF503 | NA | -0.300353 | 0.135999 | GPATCH4 | NA | -0.067021466 | 0.73977706 | GNE | midnightblue | -0.001108 | 0.995712 |
| HAGH | NA | -0.300076 | 0.136379 | ELMO2 | NA | -0.066970565 | 0.739967969 | ZNF692 | midnightblue | 0.001863 | 0.992795 |
| PKD2L1 | NA | -0.299297 | 0.137449 | DBT | NA | -0.066721065 | 0.740903963 | KIAA1147 | midnightblue | 0.004669 | 0.981941 |
| ELK3 | NA | -0.298154 | 0.13903 | CKB | NA | -0.06621174 | 0.742815796 | DZIP3 | midnightblue | 0.004928 | 0.98094 |
| URI1 | NA | -0.297821 | 0.139493 | CSTB | NA | -0.066207627 | 0.742831241 | ATP13A3 | midnightblue | 0.006634 | 0.974341 |
| MFGE8 | NA | 0.297308 | 0.140209 | ELOVL1 | NA | -0.065828017 | 0.744257145 | ZSWIM7 | midnightblue | 0.007799 | 0.969839 |
| OR6N1 | NA | 0.29703 | 0.140599 | NRDE2 | NA | -0.065344996 | 0.74607268 | FAM115A | midnightblue | 0.020092 | 0.922393 |
| SNX10 | NA | -0.296978 | 0.140671 | CLUAP1 | NA | -0.065047853 | 0.747190209 | RRP1B | midnightblue | 0.025085 | 0.903185 |
| RFXANK | NA | -0.296895 | 0.140788 | RGS1 | NA | -0.064820415 | 0.748045923 | SH3BP5 | midnightblue | 0.030417 | 0.882735 |
| TFCP2L1 | NA | 0.296819 | 0.140893 | ERV3-1 | NA | -0.064721734 | 0.748417292 | ARFGAP2 | midnightblue | 0.031381 | 0.879045 |
| WLS | NA | 0.295179 | 0.143207 | DDX56 | NA | -0.064100883 | 0.750755006 | SLC39A9 | midnightblue | 0.034514 | 0.867071 |
| DCTN5 | NA | -0.294957 | 0.143523 | NDUFA2 | NA | -0.063985166 | 0.751190959 | DGCR8 | midnightblue | 0.034974 | 0.865314 |
| NUDT15 | NA | -0.294173 | 0.144641 | FMO5 | NA | -0.063975227 | 0.751228405 | TMED2 | midnightblue | 0.045302 | 0.826068 |
| GSTM3 | NA | -0.294164 | 0.144653 | SLC2A3 | NA | -0.063644542 | 0.752474657 | CD164 | midnightblue | 0.048751 | 0.813047 |
| MUSTN1 | NA | 0.292852 | 0.146538 | HIPK2 | NA | -0.063358116 | 0.7535546 | LIN7C | midnightblue | 0.080383 | 0.696278 |
| RFC4 | NA | -0.292456 | 0.147109 | LYAR | NA | -0.063338141 | 0.753629932 | LUC7L3 | midnightblue | 0.08223 | 0.689637 |
| EIF1AD | NA | -0.291877 | 0.147949 | ELMSAN1 | NA | -0.063153997 | 0.754324486 | ODF3B | midnightblue | 0.083681 | 0.684433 |
| EIF2B4 | NA | -0.291597 | 0.148356 | UBA1 | NA | -0.063118151 | 0.754459713 | DEDD | midnightblue | 0.086078 | 0.67587 |
| ACRBP | NA | -0.291035 | 0.149176 | LRRC8C | NA | -0.062763708 | 0.755797202 | CNOT6L | midnightblue | 0.09302 | 0.651294 |
| RFWD3 | NA | 0.290001 | 0.150692 | MANBAL | NA | -0.062760924 | 0.755807709 | LRRC59 | midnightblue | 0.096202 | 0.640143 |
| GALNT6 | NA | -0.289591 | 0.151296 | FZR1 | NA | -0.062656023 | 0.756203687 | SAT2 | midnightblue | 0.102808 | 0.617239 |
| SPTY2D1 | NA | 0.289349 | 0.151654 | PARL | NA | -0.061924742 | 0.758965766 | UBASH3B | midnightblue | 0.102974 | 0.61667 |
| KDM5B | NA | -0.287348 | 0.154632 | ADCY3 | NA | -0.061876187 | 0.759149263 | TBC1D13 | midnightblue | 0.123697 | 0.547153 |
| LAMTOR5 | NA | -0.286813 | 0.155436 | CHD7 | NA | -0.061201554 | 0.761700118 | NUP188 | midnightblue | 0.124039 | 0.546038 |
| TBC1D3F | NA | 0.286451 | 0.155981 | RHOBTB1 | NA | -0.060889946 | 0.762879162 | TMX2 | midnightblue | 0.126146 | 0.539184 |
| REXO2 | NA | -0.286435 | 0.156006 | ULK1 | NA | -0.060698567 | 0.763603545 | PPP2R2A | midnightblue | 0.13128 | 0.522658 |
| COX7B | NA | -0.286403 | 0.156054 | BAG2 | NA | -0.060593389 | 0.764001738 | TARS | midnightblue | 0.131671 | 0.52141 |
| FAM149B1 | NA | -0.286394 | 0.156068 | MLL5 | NA | -0.060436758 | 0.764594831 | RAB35 | midnightblue | 0.14198 | 0.489019 |
| ZNF215 | NA | 0.28611 | 0.156496 | NDUFA10 | NA | -0.060281281 | 0.765183681 | C10orf2 | midnightblue | 0.162185 | 0.428608 |
| MCTP1 | NA | -0.28563 | 0.157224 | CRYBB2P1 | NA | -0.060197717 | 0.76550022 | FLJ23867 | midnightblue | 0.190577 | 0.351053 |
| GPR124 | NA | 0.284803 | 0.158482 | CHST3 | NA | -0.060153023 | 0.765669538 | TMED5 | midnightblue | 0.197676 | 0.333056 |
| TCP11L2 | NA | -0.28445 | 0.159021 | GBP3 | NA | -0.060143767 | 0.765704606 | ANAPC7 | midnightblue | 0.198938 | 0.329915 |
| C19orf66 | NA | -0.284333 | 0.159199 | OST4 | NA | -0.059935442 | 0.766493962 | FLJ43663 | midnightblue | 0.210837 | 0.301198 |
| PPM1N | NA | -0.28424 | 0.159341 | PPM1L | NA | -0.059739966 | 0.767234839 | BLOC1S6 | midnightblue | 0.232668 | 0.252701 |
| PMPCB | NA | -0.284236 | 0.159348 | MRC1 | NA | -0.059509352 | 0.768109152 | CHMP2B | midnightblue | 0.247239 | 0.223342 |
| MNAT1 | NA | 0.284121 | 0.159525 | TMED2 | NA | -0.05936552 | 0.768654595 | CPT1B | midnightblue | 0.266913 | 0.187462 |
| TMEM69 | NA | -0.283772 | 0.16006 | C19orf54 | NA | -0.059095393 | 0.769679265 | REXO1 | midnightblue | 0.271489 | 0.179725 |

**Table 6. Gene module membership and correlation with growth rate (48/72h), with adjusted p values for each module**

| | | | | | | | | | | | |
|---|---|---|---|---|---|---|---|---|---|---|---|
| TDRD3 | NA | -0.282708 | 0.1617 | GOLT1B | NA | -0.058781133 | 0.770871819 | C2orf43 | midnightblue | 0.293502 | 0.145601 |
| RPL13P5 | NA | -0.282645 | 0.161797 | CD40 | NA | -0.058762505 | 0.770942525 | PPP1R10 | midnightblue | 0.302054 | 0.133689 |
| TMEM50B | NA | -0.28205 | 0.162721 | PHGDH | NA | -0.058412169 | 0.77227261 | DRG2 | NA | -0.522195 | 0.00621 |
| MIF4GD | NA | -0.281551 | 0.163497 | SARM1 | NA | -0.057712646 | 0.774930301 | CENPO | NA | -0.504086 | 0.008646 |
| AR | NA | -0.28124 | 0.163983 | SLC25A46 | NA | -0.057672899 | 0.775081388 | MT1L | NA | -0.503469 | 0.008741 |
| PIK3C2B | NA | 0.279482 | 0.166746 | ASMTL | NA | -0.057525821 | 0.775640526 | CRYZ | NA | -0.432121 | 0.027485 |
| GPAA1 | NA | -0.279047 | 0.167435 | DNAJB12 | NA | -0.057321793 | 0.776416345 | HLA-A | NA | -0.40522 | 0.040009 |
| NADK | NA | -0.278719 | 0.167956 | RARG | NA | -0.056921069 | 0.77794072 | HM13 | NA | -0.389821 | 0.048997 |
| XPC | NA | -0.278087 | 0.168963 | DGCR8 | NA | -0.056874063 | 0.778119583 | LOC1001328: | NA | -0.38904 | 0.049492 |
| FAM165B | NA | -0.277886 | 0.169284 | ETV6 | NA | -0.056638613 | 0.779015676 | PSMD7 | NA | -0.379342 | 0.055969 |
| AP5M1 | NA | -0.277813 | 0.169401 | PSME2 | NA | -0.056546013 | 0.779368174 | SMOX | NA | -0.372722 | 0.060757 |
| GGTA1P | NA | -0.277708 | 0.169569 | SORL1 | NA | -0.056304959 | 0.780285991 | CCDC134 | NA | -0.369465 | 0.063225 |
| KDM2B | NA | -0.276924 | 0.170825 | ACTR1B | NA | -0.056291179 | 0.780338467 | REXO1L2P | NA | -0.359909 | 0.070919 |
| CCDC124 | NA | -0.276845 | 0.170952 | CCL20 | NA | -0.056279299 | 0.780383711 | IDH3A | NA | -0.3596 | 0.071179 |
| CEACAM21 | NA | 0.276748 | 0.171109 | LY6E | NA | -0.056250662 | 0.780492767 | SNX29P2 | NA | -0.356651 | 0.0737 |
| TIMD4 | NA | -0.276004 | 0.17231 | RFNG | NA | -0.056021955 | 0.781363901 | SLC25A51 | NA | -0.352508 | 0.077356 |
| ITPKC | NA | -0.275968 | 0.172368 | KANSL2 | NA | -0.055882684 | 0.781894501 | AACS | NA | -0.351202 | 0.078536 |
| C19orf40 | NA | -0.275785 | 0.172664 | ENTHD2 | NA | -0.055606152 | 0.782948328 | UCK2 | NA | -0.350214 | 0.079438 |
| ZBTB6 | NA | -0.275768 | 0.172693 | LRRC23 | NA | -0.05558678 | 0.783022167 | MTSS1 | NA | -0.35007 | 0.07957 |
| KIAA0430 | NA | -0.275679 | 0.172836 | CCDC53 | NA | -0.055582179 | 0.783039705 | GEMIN6 | NA | -0.348726 | 0.080812 |
| P4HA2 | NA | -0.275148 | 0.173699 | SNX29 | NA | -0.055571914 | 0.783078833 | PCGF1 | NA | -0.348326 | 0.081184 |
| SAMD1 | NA | -0.274878 | 0.174139 | DPH3 | NA | -0.055126338 | 0.784777726 | ZNF35 | NA | -0.345977 | 0.083397 |
| ORAI3 | NA | -0.273605 | 0.176222 | TCEAL3 | NA | -0.055059095 | 0.785034194 | ULBP2 | NA | -0.342041 | 0.087206 |
| RABL5 | NA | -0.273334 | 0.176669 | C16orf80 | NA | -0.054895495 | 0.785658265 | LOC401397 | NA | -0.341976 | 0.08727 |
| TRMT10C | NA | -0.273249 | 0.176809 | AMACR | NA | -0.054030281 | 0.788960857 | ELK3 | NA | -0.340094 | 0.089139 |
| TST | NA | -0.272918 | 0.177354 | GCFC2 | NA | -0.053961827 | 0.789222305 | EZR | NA | -0.337891 | 0.091364 |
| PSMB4 | NA | -0.27247 | 0.178094 | EHBP1L1 | NA | -0.053947101 | 0.789278551 | BNIP1 | NA | -0.334717 | 0.094643 |
| AGPAT5 | NA | -0.272383 | 0.178239 | PEX6 | NA | -0.052996098 | 0.792913092 | CCL2 | NA | -0.333861 | 0.095542 |
| LIMS2 | NA | -0.271498 | 0.179709 | CARHSP1 | NA | -0.052947464 | 0.793099072 | SRPX2 | NA | -0.332564 | 0.096917 |
| )C1001307( | NA | 0.271428 | 0.179827 | LIMK1 | NA | -0.052643433 | 0.794261982 | NAT1 | NA | -0.32709 | 0.102882 |
| SNAI2 | NA | 0.271305 | 0.180031 | CKAP4 | NA | -0.052560901 | 0.794577737 | CLUAP1 | NA | -0.325383 | 0.104798 |
| WWOX | NA | -0.270729 | 0.180993 | SLC28A3 | NA | -0.051353806 | 0.799199507 | EMR1 | NA | -0.323631 | 0.10679 |
| RNF111 | NA | 0.270356 | 0.181619 | E2F4 | NA | -0.0513468 | 0.799226353 | RAI14 | NA | -0.318605 | 0.112663 |
| ENC1 | NA | -0.268996 | 0.183911 | P4HA1 | NA | -0.051000751 | 0.800552554 | USP17L5 | NA | -0.318528 | 0.112756 |
| GGH | NA | -0.268709 | 0.184399 | HLA-DMA | NA | -0.050984553 | 0.800614643 | PIP4K2A | NA | -0.315889 | 0.115935 |
| RGPD1 | NA | -0.26759 | 0.186303 | PRKCE | NA | -0.050655374 | 0.801876718 | ZDHHC3 | NA | -0.314107 | 0.118118 |
| AKIP1 | NA | -0.267387 | 0.18665 | EPHX1 | NA | -0.049880125 | 0.804850938 | PRIM1 | NA | -0.313122 | 0.119339 |
| ECHS1 | NA | -0.266521 | 0.188135 | CD300LB | NA | -0.049701532 | 0.805536486 | TMEM53 | NA | -0.304076 | 0.130977 |
| HAAO | NA | 0.266428 | 0.188295 | DHX34 | NA | -0.049577144 | 0.806014041 | NBPF3 | NA | -0.303943 | 0.131154 |
| ATXN7L1 | NA | -0.265514 | 0.189874 | VARS | NA | -0.049395664 | 0.80671091 | TNFRSF10D | NA | -0.303374 | 0.131914 |
| TRMU | NA | 0.264989 | 0.190784 | PIK3IP1 | NA | -0.049373114 | 0.806797512 | ST6GAL1 | NA | -0.301558 | 0.134359 |
| CCDC127 | NA | -0.264483 | 0.191663 | SLC31A2 | NA | -0.049326304 | 0.806977286 | GCSH | NA | -0.299263 | 0.137496 |
| KCTD12 | NA | 0.264441 | 0.191738 | GM2A | NA | -0.04817945 | 0.811384771 | HIVEP3 | NA | -0.295312 | 0.143019 |
| TSTD1 | NA | -0.264431 | 0.191754 | UQCRB | NA | -0.047965778 | 0.81220656 | TIMM22 | NA | -0.295105 | 0.143313 |
| SPATA6 | NA | -0.263788 | 0.192877 | IL3RA | NA | -0.047706289 | 0.813204825 | APP | NA | -0.29091 | 0.149359 |
| CLDN7 | NA | -0.263706 | 0.193021 | ARL5B | NA | -0.047414298 | 0.814328465 | CD46 | NA | -0.289859 | 0.150902 |
| RARRES3 | NA | -0.263545 | 0.193303 | SPRY2 | NA | -0.047401517 | 0.814377659 | NOB1 | NA | -0.288367 | 0.15311 |
| BCLAF1 | NA | -0.263418 | 0.193526 | VDR | NA | -0.047311925 | 0.814722502 | FAM213A | NA | -0.287017 | 0.155129 |
| GOLIM4 | NA | -0.262889 | 0.194454 | ZNF326 | NA | -0.047227437 | 0.815047731 | SLC25A20 | NA | -0.286152 | 0.156433 |
| ZNF90 | NA | -0.261514 | 0.196885 | HAX1 | NA | -0.047106435 | 0.815513573 | CDK5 | NA | -0.283563 | 0.160382 |
| WBP1 | NA | -0.261213 | 0.197418 | MYO5A | NA | -0.04703841 | 0.815775487 | LAMTOR5 | NA | -0.282832 | 0.161509 |
| EEF2K | NA | -0.260278 | 0.199086 | BRD3 | NA | -0.046945877 | 0.816131792 | CTSO | NA | -0.281375 | 0.163772 |
| RNGTT | NA | -0.259433 | 0.200601 | DOCK3 | NA | -0.046719733 | 0.81700273 | C11orf82 | NA | -0.2755 | 0.173127 |
| POLR3E | NA | 0.25934 | 0.200768 | COA3 | NA | -0.046627311 | 0.817358732 | MFF | NA | -0.275339 | 0.173388 |
| DHRS7 | NA | -0.259312 | 0.200819 | AHCYL1 | NA | -0.04642486 | 0.818138682 | NSMCE4A | NA | -0.274727 | 0.174384 |
| MTMR14 | NA | -0.259021 | 0.201342 | SQRDL | NA | -0.046361586 | 0.818382481 | ZNF584 | NA | -0.272316 | 0.178351 |

**Table 6. Gene module membership and correlation with growth rate (48/72h), with adjusted p values for each module**

| | | | | | | | | | | | |
|---|---|---|---|---|---|---|---|---|---|---|---|
| SMG8 | NA | -0.258704 | 0.201913 | ARHGAP15 | NA | -0.045500746 | 0.821700987 | DUSP12 | NA | -0.271551 | 0.17962 |
| PIP4K2A | NA | -0.258466 | 0.202345 | AQP3 | NA | -0.04549709 | 0.821715086 | BMS1 | NA | -0.270904 | 0.180701 |
| GTF3C4 | NA | 0.25828 | 0.20268 | RPP14 | NA | -0.04547487 | 0.821800785 | MICA | NA | -0.26887 | 0.184125 |
| NDUFS3 | NA | -0.257437 | 0.204211 | PLCXD1 | NA | -0.045353454 | 0.822269095 | NRBP2 | NA | -0.267938 | 0.185709 |
| HLA-B | NA | 0.257094 | 0.204834 | TNFAIP2 | NA | -0.045125928 | 0.823146835 | METTL2A | NA | -0.267912 | 0.185754 |
| RFX3 | NA | 0.256515 | 0.205893 | TRAP1 | NA | -0.04462684 | 0.825072927 | CD52 | NA | -0.266131 | 0.188807 |
| HOMER3 | NA | 0.256497 | 0.205925 | RBFA | NA | -0.043780078 | 0.828343027 | CYSLTR2 | NA | -0.266009 | 0.189018 |
| GMFG | NA | -0.256239 | 0.206398 | PWP1 | NA | -0.04372537 | 0.8285544 | C5orf44 | NA | -0.265782 | 0.18941 |
| SNAP29 | NA | 0.256168 | 0.206529 | SPECC1 | NA | -0.043566553 | 0.829168083 | GLRX2 | NA | -0.251305 | 0.215575 |
| STX12 | NA | -0.255534 | 0.207694 | KDELR2 | NA | -0.042658192 | 0.832679922 | VNN1 | NA | -0.25078 | 0.216568 |
| SLC16A10 | NA | 0.255016 | 0.208648 | THOC6 | NA | -0.04262734 | 0.832799256 | LOC1002887 | NA | -0.250079 | 0.217898 |
| PTP4A2 | NA | -0.254902 | 0.208858 | RSAD2 | NA | -0.04254002 | 0.833137022 | C9orf89 | NA | -0.248663 | 0.2206 |
| ZNF160 | NA | -0.254797 | 0.209053 | PPARGC1B | NA | -0.042416749 | 0.833613901 | CLK4 | NA | -0.248604 | 0.220714 |
| TARDBP | NA | -0.253765 | 0.210967 | HAUS1 | NA | -0.042288051 | 0.83411184 | SRSF8 | NA | -0.248296 | 0.221306 |
| BZRAP1 | NA | 0.252679 | 0.212993 | COLEC12 | NA | -0.041466015 | 0.837293781 | TMEM176B | NA | -0.248104 | 0.221675 |
| PADI2 | NA | 0.252556 | 0.213222 | ZCCHC11 | NA | -0.041373248 | 0.837653022 | HNRNPC | NA | -0.247579 | 0.222685 |
| MSH2 | NA | -0.251938 | 0.214382 | LOC1005064 | NA | -0.041304874 | 0.837917822 | CPE | NA | -0.246833 | 0.224128 |
| HMGN4 | NA | -0.251904 | 0.214447 | LEPRE1 | NA | -0.041064909 | 0.838847297 | RASSF5 | NA | -0.246011 | 0.225725 |
| TAP2 | NA | -0.251805 | 0.214633 | TPST2 | NA | -0.040614305 | 0.840593225 | LIMS2 | NA | -0.244882 | 0.22793 |
| CCPG1 | NA | -0.250561 | 0.216983 | KIAA0226L | NA | -0.040556582 | 0.840816934 | TLE6 | NA | -0.241681 | 0.234258 |
| DCP1B | NA | -0.250548 | 0.217008 | AOC2 | NA | -0.040421881 | 0.841339025 | DPH5 | NA | -0.241457 | 0.234706 |
| KIAA1609 | NA | -0.250522 | 0.217056 | ZNF595 | NA | -0.040416488 | 0.841359929 | LCP1 | NA | -0.23901 | 0.239629 |
| TIAM1 | NA | 0.250501 | 0.217097 | ZMYM4 | NA | -0.040293886 | 0.841835181 | TMEM50B | NA | -0.238081 | 0.241514 |
| GLDN | NA | 0.250221 | 0.217629 | SLC7A2 | NA | -0.040289219 | 0.841853276 | LOC349196 | NA | -0.235992 | 0.245792 |
| CD38 | NA | -0.249889 | 0.21826 | ATL3 | NA | -0.039757451 | 0.843915251 | PPARG | NA | -0.235683 | 0.246428 |
| HIGD1A | NA | -0.248935 | 0.220081 | INSR | NA | -0.03953954 | 0.844760509 | FOXJ3 | NA | -0.23446 | 0.24896 |
| CBWD1 | NA | -0.248042 | 0.221793 | CERK | NA | -0.039510495 | 0.844873187 | SNCA | NA | -0.233253 | 0.251475 |
| GUF1 | NA | -0.247681 | 0.222489 | POLL | NA | -0.039503434 | 0.84490058 | TMTC1 | NA | -0.233147 | 0.251697 |
| DCUN1D3 | NA | -0.247681 | 0.22249 | UPP1 | NA | -0.039281419 | 0.845761957 | XRCC3 | NA | -0.231504 | 0.25515 |
| KCTD7 | NA | 0.24734 | 0.223147 | DSP | NA | -0.039196286 | 0.846092304 | ZNF528 | NA | -0.229302 | 0.259824 |
| FEN1 | NA | -0.247196 | 0.223426 | RHBDD2 | NA | -0.038938714 | 0.847091931 | CNRIP1 | NA | -0.228982 | 0.260508 |
| PROCR | NA | -0.245476 | 0.226768 | APH1B | NA | -0.038913039 | 0.847191589 | USP38 | NA | -0.228639 | 0.261241 |
| MSI2 | NA | -0.245243 | 0.227221 | GFOD1 | NA | -0.038845971 | 0.84745192 | GPATCH2L | NA | -0.227668 | 0.263328 |
| PCCA | NA | -0.245211 | 0.227284 | NEK6 | NA | -0.0382654 | 0.849706126 | TSC1 | NA | -0.226318 | 0.266247 |
| ITGAM | NA | -0.244891 | 0.227912 | SIK2 | NA | -0.038158917 | 0.8501197 | STON2 | NA | -0.218796 | 0.282888 |
| CTSC | NA | -0.244362 | 0.22895 | ZNF687 | NA | -0.037881883 | 0.851195856 | CTBS | NA | -0.218298 | 0.284011 |
| HLA-DQA1 | NA | 0.24407 | 0.229523 | LDLRAP1 | NA | -0.037250993 | 0.853647565 | TMEM55A | NA | -0.217477 | 0.285873 |
| ARMC10 | NA | -0.243964 | 0.229733 | BATF | NA | -0.03723998 | 0.853690375 | MMP2 | NA | -0.216056 | 0.28911 |
| C3orf58 | NA | -0.24382 | 0.230016 | LOC401093 | NA | -0.036989186 | 0.854665364 | GATM | NA | -0.215442 | 0.290515 |
| TAPBP | NA | -0.243608 | 0.230434 | TRHDE | NA | -0.03698196 | 0.854693462 | SEPP1 | NA | -0.214728 | 0.292156 |
| NOTCH2 | NA | 0.242909 | 0.231816 | NDUFV3 | NA | -0.036832035 | 0.855276415 | FKBP14 | NA | -0.214339 | 0.293053 |
| S100PBP | NA | -0.24284 | 0.231953 | PIGG | NA | -0.036748964 | 0.855599454 | RPS19BP1 | NA | -0.212456 | 0.297413 |
| GTF2A2 | NA | -0.242541 | 0.232546 | ESPL1 | NA | -0.036560499 | 0.85633242 | PHTF1 | NA | -0.207726 | 0.308552 |
| TAF4 | NA | -0.24242 | 0.232787 | TPM1 | NA | -0.036208042 | 0.857703486 | EBPL | NA | -0.206803 | 0.310753 |
| ICA1 | NA | -0.241517 | 0.234586 | NDEL1 | NA | -0.03567186 | 0.859790019 | RSF1 | NA | -0.205706 | 0.313384 |
| PARN | NA | -0.241399 | 0.234821 | ZNF652 | NA | -0.035568269 | 0.860193243 | ZNF350 | NA | -0.205617 | 0.313599 |
| OSBPL3 | NA | 0.241165 | 0.23529 | SHPRH | NA | -0.035483258 | 0.860524176 | LOC1005066 | NA | -0.20461 | 0.316026 |
| PHTF1 | NA | -0.240913 | 0.235794 | C10orf2 | NA | -0.035457258 | 0.86062539 | ALDH7A1 | NA | -0.203527 | 0.31865 |
| PSMG4 | NA | -0.239918 | 0.237794 | ELP6 | NA | -0.035259753 | 0.861394341 | MYL12A | NA | -0.201671 | 0.323177 |
| FAM109B | NA | -0.23986 | 0.237911 | SORD | NA | -0.035137189 | 0.861871585 | MOCOS | NA | -0.201292 | 0.324106 |
| LOC148413 | NA | -0.239721 | 0.23819 | HDAC4 | NA | -0.034816062 | 0.863122221 | SH3BGRL3 | NA | -0.200753 | 0.325431 |
| LIG3 | NA | -0.238852 | 0.239947 | INO80D | NA | -0.034377207 | 0.864831876 | SMPD1 | NA | -0.200141 | 0.32694 |
| ACRC | NA | 0.2387 | 0.240256 | DNAJC6 | NA | -0.034171888 | 0.865631942 | LY86 | NA | -0.199488 | 0.328553 |
| SMOX | NA | -0.238601 | 0.240457 | CPSF4 | NA | -0.033976597 | 0.866393056 | MPV17L2 | NA | -0.19719 | 0.33427 |
| ZBTB33 | NA | -0.237252 | 0.243206 | CDK19 | NA | -0.033939378 | 0.866538121 | ITGA4 | NA | -0.19664 | 0.335645 |
| PSMD7 | NA | -0.236997 | 0.243728 | XKR8 | NA | -0.033683117 | 0.867537059 | PPP4R1 | NA | -0.195483 | 0.338555 |

**Table 6. Gene module membership and correlation with growth rate (48/72h), with adjusted p values for each module**

| | | | | | | | | | | | |
|---|---|---|---|---|---|---|---|---|---|---|---|
| TDP2 | NA | 0.236964 | 0.243795 | ACOX3 | NA | -0.033672169 | 0.867579737 | DPH3 | NA | -0.194759 | 0.340382 |
| CLDN15 | NA | 0.236399 | 0.244954 | TSTD1 | NA | -0.033551576 | 0.868049898 | COLEC12 | NA | -0.194697 | 0.340539 |
| SMIM4 | NA | -0.236397 | 0.244958 | KDM4A | NA | -0.033059222 | 0.869969904 | TRIM35 | NA | -0.194545 | 0.340922 |
| AP3S2 | NA | -0.236149 | 0.245469 | HN1L | NA | -0.031838198 | 0.874734549 | SYNE3 | NA | -0.192817 | 0.345311 |
| SORD | NA | -0.23611 | 0.24555 | ENOX2 | NA | -0.031831374 | 0.874761187 | CHD1 | NA | -0.19211 | 0.347119 |
| DGKG | NA | 0.236085 | 0.245601 | LOC401397 | NA | -0.031482322 | 0.876124046 | METAP1 | NA | -0.190718 | 0.35069 |
| C2orf49 | NA | -0.235897 | 0.245987 | STAT2 | NA | -0.030350016 | 0.880547439 | SLC7A8 | NA | -0.190301 | 0.351763 |
| SH3D21 | NA | 0.235801 | 0.246186 | NEK7 | NA | -0.0299592 | 0.882074994 | ST8SIA4 | NA | -0.188706 | 0.355891 |
| HLA-DRB6 | NA | 0.235676 | 0.246442 | TRMT13 | NA | -0.029686551 | 0.883140924 | IL10 | NA | -0.187628 | 0.358696 |
| SNHG7 | NA | -0.235602 | 0.246595 | MIER3 | NA | -0.029313098 | 0.884601273 | SPA17 | NA | -0.187579 | 0.358822 |
| C11orf82 | NA | -0.234152 | 0.249601 | ABHD15 | NA | -0.029253873 | 0.884832899 | MRC1 | NA | -0.187239 | 0.359711 |
| FAM200A | NA | -0.233941 | 0.25004 | FAM225A | NA | -0.028856918 | 0.88638561 | LZIC | NA | -0.187068 | 0.360156 |
| PDLIM5 | NA | 0.233924 | 0.250074 | PSME1 | NA | -0.028816662 | 0.886543097 | ZNF626 | NA | -0.186033 | 0.362869 |
| ECSIT | NA | -0.233182 | 0.251624 | TBC1D7 | NA | -0.028758231 | 0.886771697 | NDUFA7 | NA | -0.184903 | 0.365843 |
| LCLAT1 | NA | -0.233084 | 0.251828 | IFI27 | NA | -0.028000248 | 0.889737918 | MIS18A | NA | -0.184754 | 0.366236 |
| L3MBTL2 | NA | -0.232711 | 0.25261 | ASNS | NA | -0.027870315 | 0.89024653 | PCOLCE2 | NA | -0.184477 | 0.366968 |
| TBXAS1 | NA | -0.232691 | 0.252653 | H1F0 | NA | -0.027768098 | 0.890646682 | PXDC1 | NA | -0.183123 | 0.370556 |
| CEP135 | NA | -0.232667 | 0.252702 | TPRG1L | NA | -0.027593041 | 0.891332043 | RAD51AP1 | NA | -0.178246 | 0.383652 |
| NCAPG2 | NA | -0.232063 | 0.253971 | DYNC1LI2 | NA | -0.02750411 | 0.891680246 | SIGLEC12 | NA | -0.177565 | 0.385501 |
| PRKAR2B | NA | -0.231975 | 0.254157 | FPR1 | NA | -0.027496437 | 0.891710289 | TPCN1 | NA | -0.176725 | 0.387789 |
| ALDH7A1 | NA | -0.231912 | 0.254289 | SLC25A6 | NA | -0.027361103 | 0.892240215 | ARHGAP15 | NA | -0.175826 | 0.390247 |
| GINS4 | NA | -0.230607 | 0.257048 | TK2 | NA | -0.027236647 | 0.892727587 | ARPC4 | NA | -0.173711 | 0.396065 |
| DYNC1LI2 | NA | 0.230436 | 0.25741 | SMG8 | NA | -0.027137861 | 0.893114462 | TSPAN15 | NA | -0.172987 | 0.398067 |
| USE1 | NA | 0.229919 | 0.258509 | APBA3 | NA | -0.027137727 | 0.893114987 | MRPS27 | NA | -0.172223 | 0.400184 |
| MME | NA | -0.22887 | 0.260747 | FAM134A | NA | -0.027002725 | 0.893643734 | INSIG1 | NA | -0.171802 | 0.401354 |
| SH2D3C | NA | -0.228227 | 0.262126 | SIN3B | NA | -0.026759029 | 0.894598302 | CST2 | NA | -0.168729 | 0.409955 |
| MFSD12 | NA | 0.227975 | 0.262669 | DNAJA4 | NA | -0.026179891 | 0.896867382 | ODC1 | NA | -0.16649 | 0.416286 |
| TDRD9 | NA | 0.227558 | 0.263565 | ZNF44 | NA | -0.026179097 | 0.896870491 | URI1 | NA | -0.165985 | 0.417723 |
| EPHA4 | NA | -0.227037 | 0.264689 | LOC1005067 | NA | -0.026137978 | 0.897031629 | PPHLN1 | NA | -0.165462 | 0.41921 |
| ZNF646 | NA | 0.227025 | 0.264716 | THAP3 | NA | -0.025765246 | 0.898492459 | ARHGAP4 | NA | -0.16499 | 0.420557 |
| RASA4 | NA | 0.226725 | 0.265365 | INHBA | NA | -0.025740944 | 0.898587717 | TBC1D14 | NA | -0.16245 | 0.427844 |
| PRSS21 | NA | 0.226549 | 0.265746 | HDDC2 | NA | -0.025677673 | 0.898835726 | PPP1R15B | NA | -0.161363 | 0.430985 |
| PIGU | NA | -0.226434 | 0.265996 | TPRN | NA | -0.025599335 | 0.89914281 | PMVK | NA | -0.16018 | 0.434415 |
| TFPT | NA | -0.226307 | 0.266272 | DNAJB5 | NA | -0.025596437 | 0.899154171 | MIF4GD | NA | -0.159689 | 0.435843 |
| LOC339894 | NA | 0.225888 | 0.267181 | RHOU | NA | -0.025248809 | 0.900517038 | GP1BA | NA | -0.159607 | 0.436082 |
| SYK | NA | -0.225798 | 0.267377 | TAP1 | NA | -0.025214132 | 0.900653006 | KHSRP | NA | -0.159508 | 0.436369 |
| GCNT2 | NA | -0.225418 | 0.268204 | SLC4A1AP | NA | -0.025167425 | 0.900836144 | CTSZ | NA | -0.157756 | 0.44149 |
| SAMSN1 | NA | -0.225226 | 0.268623 | ENC1 | NA | -0.023775426 | 0.90629642 | C1orf109 | NA | -0.154728 | 0.450417 |
| BRD3 | NA | -0.224691 | 0.269792 | MFSD12 | NA | -0.023495738 | 0.907394038 | WDR77 | NA | -0.154722 | 0.450435 |
| LOC550112 | NA | -0.224554 | 0.270092 | PDXK | NA | -0.023474508 | 0.907477361 | UBL5 | NA | -0.153867 | 0.45297 |
| L3MBTL4 | NA | -0.224526 | 0.270153 | KANSL1L | NA | -0.023256162 | 0.90833437 | SERPINE2 | NA | -0.153787 | 0.45321 |
| RNF25 | NA | -0.224523 | 0.27016 | PNMA1 | NA | -0.023193561 | 0.908580101 | FXYD6 | NA | -0.153778 | 0.453236 |
| AP2A1 | NA | -0.224232 | 0.270798 | FBXL12 | NA | -0.023167192 | 0.908683608 | CEP250 | NA | -0.152882 | 0.455905 |
| HP | NA | 0.224134 | 0.271012 | PTPMT1 | NA | -0.022930298 | 0.909613577 | FANCD2 | NA | -0.15279 | 0.456177 |
| PRKX | NA | -0.224127 | 0.271027 | FUNDC1 | NA | -0.02280683 | 0.91009832 | THBD | NA | -0.151079 | 0.461299 |
| RAP1A | NA | -0.224014 | 0.271275 | ABCA1 | NA | -0.022207397 | 0.912452165 | CSTA | NA | -0.151049 | 0.461388 |
| SIRPB1 | NA | 0.223853 | 0.271629 | IFT74 | NA | -0.02216991 | 0.912599394 | FPR2 | NA | -0.150861 | 0.461953 |
| C17orf47 | NA | 0.223532 | 0.272335 | SLC39A9 | NA | -0.022142514 | 0.912706989 | RAD9A | NA | -0.150378 | 0.463402 |
| PPM1A | NA | -0.223388 | 0.272652 | PCM1 | NA | -0.022037782 | 0.91311834 | VPS41 | NA | -0.149892 | 0.464865 |
| ZNF581 | NA | -0.223381 | 0.272668 | RCBTB2 | NA | -0.021991857 | 0.913298723 | OLFML2B | NA | -0.149387 | 0.466388 |
| PMVK | NA | -0.220689 | 0.27864 | NEK1 | NA | -0.021869143 | 0.913780737 | CSTB | NA | -0.14793 | 0.470798 |
| AKR1A1 | NA | -0.220623 | 0.278786 | ZNF692 | NA | -0.021685945 | 0.914500384 | THAP9-AS1 | NA | -0.146492 | 0.475168 |
| SERPINA1 | NA | 0.220542 | 0.278968 | BGLAP | NA | -0.020838084 | 0.91783184 | HSD17B7 | NA | -0.145764 | 0.477388 |
| INPP4A | NA | -0.220526 | 0.279004 | ITGA3 | NA | -0.02063439 | 0.918632411 | GNS | NA | -0.145648 | 0.477743 |
| IKZF5 | NA | -0.21992 | 0.280359 | ADCK2 | NA | -0.020559411 | 0.918927116 | GLT8D1 | NA | -0.144041 | 0.482667 |
| KLC4 | NA | -0.21982 | 0.280584 | RASA4CP | NA | -0.020488163 | 0.919207168 | SESN3 | NA | -0.144014 | 0.48275 |

**Table 6. Gene module membership and correlation with growth rate (48/72h), with adjusted p values for each module**

| | | | | | | | | | | | |
|---|---|---|---|---|---|---|---|---|---|---|---|
| CALML4 | NA | -0.219429 | 0.281461 | RNGTT | NA | -0.020443425 | 0.91938302 | PKDCC | NA | -0.143768 | 0.483507 |
| FBXL19 | NA | -0.218694 | 0.283118 | ARF4 | NA | -0.020323149 | 0.919855816 | HADHA | NA | -0.143509 | 0.484304 |
| TSTD2 | NA | 0.218464 | 0.283638 | TACO1 | NA | -0.019854531 | 0.921698159 | NPIP | NA | -0.143349 | 0.484794 |
| NEK1 | NA | -0.217979 | 0.284734 | RNF207 | NA | -0.01937122 | 0.923598684 | ZNF527 | NA | -0.143315 | 0.484898 |
| NEURL4 | NA | -0.217141 | 0.286636 | PPP4R1 | NA | -0.019153273 | 0.924455853 | LGALS2 | NA | -0.143314 | 0.484902 |
| MICALL2 | NA | 0.217013 | 0.286928 | LGALS3BP | NA | -0.019149927 | 0.924469011 | HAUS6 | NA | -0.142275 | 0.488108 |
| RNF135 | NA | -0.216989 | 0.286981 | TEX2 | NA | -0.018794267 | 0.925867972 | YTHDF2 | NA | -0.141852 | 0.489414 |
| TNS3 | NA | 0.216955 | 0.287058 | NARS2 | NA | -0.018310426 | 0.927771468 | SPRED1 | NA | -0.141691 | 0.489912 |
| MPPE1 | NA | -0.21678 | 0.287457 | ZNF26 | NA | -0.018098319 | 0.928606048 | XPR1 | NA | -0.141018 | 0.491996 |
| THAP9-AS1 | NA | -0.216608 | 0.287849 | CHI3L1 | NA | -0.018062207 | 0.928748144 | SERPINB6 | NA | -0.139432 | 0.49693 |
| SESTD1 | NA | -0.216413 | 0.288293 | UNC93B1 | NA | -0.017855693 | 0.9295608 | CBR1 | NA | -0.139056 | 0.4981 |
| IDNK | NA | -0.216064 | 0.289091 | RPS3 | NA | -0.017829578 | 0.929663568 | FCN1 | NA | -0.138359 | 0.500279 |
| CCDC58 | NA | -0.214822 | 0.291939 | TBL1X | NA | -0.017405569 | 0.931332325 | MRPL55 | NA | -0.138352 | 0.5003 |
| DAPP1 | NA | -0.214331 | 0.29307 | TOM1 | NA | -0.01707845 | 0.932619949 | ENSA | NA | -0.137899 | 0.50172 |
| PMP22 | NA | -0.214185 | 0.293408 | SLC12A9 | NA | -0.016875108 | 0.933420438 | LOC1005057: | NA | -0.137864 | 0.501829 |
| TSEN54 | NA | -0.213817 | 0.294257 | KIAA0195 | NA | -0.016652953 | 0.934295063 | TCTN2 | NA | -0.137424 | 0.503208 |
| WDR67 | NA | 0.213589 | 0.294786 | GCC1 | NA | -0.016450999 | 0.935090219 | PRDM15 | NA | -0.136875 | 0.504931 |
| KIAA0391 | NA | -0.213294 | 0.295468 | SDF4 | NA | -0.016435812 | 0.935150016 | HMGB3 | NA | -0.13525 | 0.510049 |
| RNF34 | NA | -0.213262 | 0.295542 | OSBPL8 | NA | -0.01573483 | 0.937910483 | IFT80 | NA | -0.133697 | 0.514963 |
| CCL13 | NA | -0.21304 | 0.296056 | SCCPDH | NA | -0.01545137 | 0.939026948 | PLAU | NA | -0.133087 | 0.5169 |
| TCEAL1 | NA | -0.212709 | 0.296825 | EML2 | NA | -0.015224049 | 0.939922383 | MSL2 | NA | -0.133018 | 0.517119 |
| DNA2 | NA | -0.212219 | 0.297967 | TMCO6 | NA | -0.015015741 | 0.940742982 | CAD | NA | -0.132489 | 0.518803 |
| SRBD1 | NA | -0.211937 | 0.298624 | FCN1 | NA | -0.014921549 | 0.94111406 | ZNF814 | NA | -0.130322 | 0.525723 |
| TEX2 | NA | -0.21155 | 0.299527 | SLC52A2 | NA | -0.014431706 | 0.943044023 | TXNDC17 | NA | -0.127563 | 0.534597 |
| AZU1 | NA | -0.210822 | 0.301234 | RANBP6 | NA | -0.014396225 | 0.94318383 | TMEM69 | NA | -0.127338 | 0.535324 |
| SOS1 | NA | -0.210788 | 0.301312 | FH | NA | -0.013872094 | 0.945249259 | GSTM3 | NA | -0.125705 | 0.540615 |
| DGCR6 | NA | -0.210494 | 0.302004 | GZMB | NA | -0.013047738 | 0.948498469 | OXSM | NA | -0.124622 | 0.544137 |
| ZNF614 | NA | 0.210422 | 0.302172 | AVEN | NA | -0.012837441 | 0.949327487 | MSI2 | NA | -0.124174 | 0.545596 |
| LOC283922 | NA | 0.210387 | 0.302254 | LSM10 | NA | -0.012359054 | 0.951213543 | KIAA0907 | NA | -0.123958 | 0.546301 |
| HAUS6 | NA | -0.21033 | 0.302387 | TMEM161A | NA | -0.012229846 | 0.951722996 | LOC729013 | NA | -0.122477 | 0.551141 |
| RUFY1 | NA | -0.209942 | 0.303302 | GPR64 | NA | -0.012150908 | 0.952034246 | GGT1 | NA | -0.122276 | 0.551802 |
| RASGEF1A | NA | -0.209871 | 0.303468 | KANK1 | NA | -0.011958409 | 0.952793298 | CEP85L | NA | -0.122133 | 0.552271 |
| VAMP5 | NA | -0.209779 | 0.303687 | CEP120 | NA | -0.011919017 | 0.952948634 | CD3EAP | NA | -0.121455 | 0.554495 |
| EIF4EBP3 | NA | 0.209395 | 0.304591 | HMGN4 | NA | -0.011847418 | 0.95323097 | PDPR | NA | -0.121289 | 0.555039 |
| HLX | NA | 0.20939 | 0.304603 | DOCK8 | NA | -0.011823726 | 0.953324398 | SLC20A2 | NA | -0.120267 | 0.558402 |
| CCNYL1 | NA | 0.209143 | 0.305189 | NAGLU | NA | -0.011400871 | 0.954991977 | LINC00341 | NA | -0.119929 | 0.559517 |
| GPD1L | NA | -0.209134 | 0.305209 | LOC652276 | NA | -0.011346677 | 0.955205709 | C15orf38 | NA | -0.119845 | 0.559797 |
| ARHGAP27 | NA | -0.209086 | 0.305323 | C6orf48 | NA | -0.010271339 | 0.959447305 | TGDS | NA | -0.119833 | 0.559833 |
| SOCS7 | NA | 0.207683 | 0.308653 | ORC3 | NA | -0.009678198 | 0.961787377 | CDKN3 | NA | -0.119458 | 0.561071 |
| TPST2 | NA | -0.207537 | 0.309001 | WDR41 | NA | -0.009545984 | 0.962309032 | TAF1B | NA | -0.118892 | 0.562943 |
| RNASE1 | NA | -0.206961 | 0.310375 | P4HTM | NA | -0.009520313 | 0.962410319 | SERP1 | NA | -0.11856 | 0.56404 |
| HDAC3 | NA | -0.206916 | 0.310483 | PET117 | NA | -0.009437806 | 0.962735862 | POLR2J2 | NA | -0.116716 | 0.570162 |
| DDX31 | NA | -0.206796 | 0.310769 | CXCL9 | NA | -0.009296216 | 0.963294543 | TNFAIP8L2 | NA | -0.11647 | 0.570981 |
| EZH1 | NA | -0.206087 | 0.312469 | KIAA0922 | NA | -0.009282846 | 0.963347298 | LPAR1 | NA | -0.11633 | 0.571447 |
| AAMDC | NA | -0.206075 | 0.312498 | AVPI1 | NA | -0.009154755 | 0.963852731 | MOSPD3 | NA | -0.115153 | 0.57537 |
| SPECC1L | NA | -0.205977 | 0.312733 | SECTM1 | NA | -0.009014141 | 0.964407592 | ING5 | NA | -0.114798 | 0.576559 |
| UFC1 | NA | -0.205522 | 0.313826 | CLEC4D | NA | -0.008973214 | 0.964569094 | NMRK1 | NA | -0.114566 | 0.577332 |
| UBB | NA | -0.205137 | 0.314753 | RAP1GAP2 | NA | -0.008567289 | 0.966170974 | CBWD6 | NA | -0.112827 | 0.583162 |
| SIGLEC12 | NA | -0.205026 | 0.315021 | KPTN | NA | -0.008511374 | 0.966391636 | CCDC85B | NA | -0.111791 | 0.586648 |
| DTL | NA | -0.20496 | 0.315181 | ITGB3 | NA | -0.008393311 | 0.96685757 | SH3GL1P1 | NA | -0.110198 | 0.592027 |
| ASAH1 | NA | -0.204956 | 0.315191 | P4HA2 | NA | -0.008253702 | 0.967408552 | ERCC2 | NA | -0.109356 | 0.594875 |
| SLC25A26 | NA | -0.204797 | 0.315575 | SCARNA10 | NA | -0.008175405 | 0.967717564 | LOC220906 | NA | -0.109291 | 0.595095 |
| ENDOG | NA | 0.204132 | 0.317182 | RP2 | NA | -0.007665054 | 0.969731863 | TK2 | NA | -0.108259 | 0.598599 |
| APMAP | NA | -0.203912 | 0.317714 | TTC13 | NA | -0.007574193 | 0.970090501 | UBL3 | NA | -0.107965 | 0.599598 |
| AXIN1 | NA | 0.203843 | 0.317883 | LAMB2 | NA | -0.0071514 | 0.97175938 | DIXDC1 | NA | -0.1072 | 0.6022 |
| FAM171B | NA | 0.202845 | 0.320308 | FBXO10 | NA | -0.007121247 | 0.97187841 | GABPB1 | NA | -0.105802 | 0.606971 |

**Table 6. Gene module membership and correlation with growth rate (48/72h), with adjusted p values for each module**

| | | | | | | | | | | | |
|---|---|---|---|---|---|---|---|---|---|---|---|
| MEGF9 | NA | -0.202399 | 0.321396 | BCS1L | NA | -0.007092618 | 0.971991418 | ZMYM6NB | NA | -0.105058 | 0.609515 |
| C11orf84 | NA | -0.202068 | 0.322205 | IRS2 | NA | -0.006821227 | 0.973062747 | NUDT6 | NA | -0.104999 | 0.609719 |
| BPIFA1 | NA | 0.201551 | 0.323472 | CARD8 | NA | -0.006675457 | 0.973638199 | HCAR2 | NA | -0.104951 | 0.609882 |
| OC1002887 | NA | -0.201462 | 0.32369 | PHF19 | NA | -0.006130905 | 0.97578802 | TEX30 | NA | -0.104183 | 0.612515 |
| TAF1 | NA | -0.201258 | 0.32419 | MAPRE1 | NA | -0.005964782 | 0.976443886 | NDUFS3 | NA | -0.102275 | 0.619076 |
| ZNF211 | NA | -0.201129 | 0.324506 | FBXL18 | NA | -0.005938563 | 0.976547405 | TTYH2 | NA | -0.101459 | 0.621891 |
| MTX3 | NA | -0.201069 | 0.324654 | TMEM86A | NA | -0.005817707 | 0.977024564 | RPL23AP53 | NA | -0.101371 | 0.622193 |
| ARHGAP26 | NA | -0.201017 | 0.324782 | RPL19 | NA | -0.005526723 | 0.97817345 | ERI3 | NA | -0.100932 | 0.623709 |
| GPA33 | NA | -0.200908 | 0.325051 | SDHAP2 | NA | -0.005448896 | 0.978480738 | IL6ST | NA | -0.099329 | 0.629262 |
| SLC31A2 | NA | -0.200639 | 0.325711 | ZNF778 | NA | -0.00536976 | 0.9787932 | ADCY3 | NA | -0.099178 | 0.629784 |
| OPN3 | NA | -0.200225 | 0.326732 | FANCG | NA | -0.005305618 | 0.979046461 | PET117 | NA | -0.099128 | 0.629959 |
| FDFT1 | NA | -0.200148 | 0.326921 | ZNF426 | NA | -0.004949589 | 0.980452255 | CLOCK | NA | -0.09852 | 0.632068 |
| CYGB | NA | -0.199841 | 0.32768 | CYTH3 | NA | -0.00492966 | 0.980530948 | TTBK2 | NA | -0.097941 | 0.634081 |
| CETN2 | NA | -0.199291 | 0.329041 | CCDC106 | NA | -0.0045053 | 0.982206632 | MED20 | NA | -0.097792 | 0.634599 |
| RABGGTB | NA | 0.19917 | 0.32934 | PLXNB2 | NA | -0.004398513 | 0.982628317 | NT5DC2 | NA | -0.09628 | 0.639871 |
| PCNXL2 | NA | -0.198666 | 0.33059 | ZNF498 | NA | -0.004314458 | 0.982960236 | MRPL37 | NA | -0.094909 | 0.644664 |
| TAGLN | NA | -0.198576 | 0.330814 | UTP6 | NA | -0.004295418 | 0.983035427 | NPM3 | NA | -0.094277 | 0.646882 |
| C10orf57 | NA | 0.198173 | 0.331816 | ATG2B | NA | -0.004015173 | 0.984142098 | KCNAB1 | NA | -0.094141 | 0.647357 |
| RPAP3 | NA | -0.197544 | 0.333384 | RPL23AP53 | NA | -0.004005761 | 0.984179268 | PHC1 | NA | -0.093704 | 0.648892 |
| NCF1C | NA | -0.197357 | 0.333851 | DCTN2 | NA | -0.003587685 | 0.985830284 | FANCF | NA | -0.092492 | 0.65315 |
| MS4A6A | NA | -0.196856 | 0.335106 | ANTXR2 | NA | -0.003385849 | 0.986627373 | CARD6 | NA | -0.091407 | 0.656972 |
| APOBEC3C | NA | -0.196834 | 0.335161 | SCD | NA | -0.003102488 | 0.987746438 | EHHADH | NA | -0.09127 | 0.657459 |
| VTN | NA | -0.196716 | 0.335455 | GEM | NA | -0.003044441 | 0.987975684 | SNHG5 | NA | -0.089825 | 0.662562 |
| EXOG | NA | -0.1964 | 0.336249 | TSC22D1 | NA | -0.002771144 | 0.989055031 | FAM198B | NA | -0.089286 | 0.664472 |
| VAV3 | NA | -0.19633 | 0.336425 | ASAH1 | NA | -0.002375052 | 0.990619369 | WWC3 | NA | -0.089097 | 0.665141 |
| MRPS23 | NA | -0.196259 | 0.336603 | RAP2A | NA | -0.002280634 | 0.990992274 | FLJ10038 | NA | -0.089074 | 0.665221 |
| GABRE | NA | 0.195758 | 0.337863 | RCOR1 | NA | -0.002041955 | 0.991934943 | PDS5B | NA | -0.088626 | 0.666809 |
| SLC24A4 | NA | 0.195266 | 0.339101 | ATP5H | NA | -0.001887888 | 0.992543439 | CAPN10 | NA | -0.088444 | 0.667455 |
| PKP2 | NA | -0.194923 | 0.339967 | CDC26 | NA | -0.001781908 | 0.992962017 | IWS1 | NA | -0.088228 | 0.668222 |
| ZNF117 | NA | 0.193725 | 0.343001 | ANKRD12 | NA | -0.001604663 | 0.993662066 | PRPSAP1 | NA | -0.087928 | 0.669289 |
| SPP1 | NA | -0.193613 | 0.343287 | APOL3 | NA | -0.001451797 | 0.994265827 | DEGS1 | NA | -0.087434 | 0.671043 |
| LOC285972 | NA | 0.192965 | 0.344936 | PSMB9 | NA | -0.001381588 | 0.994543128 | B3GNT5 | NA | -0.08672 | 0.673583 |
| CELF6 | NA | 0.19138 | 0.348988 | CPT1B | NA | -0.001034312 | 0.995914757 | TOR1AIP2 | NA | -0.08669 | 0.673691 |
| PSMB5 | NA | -0.19133 | 0.349116 | GPAM | NA | -0.000940099 | 0.99628687 | VPS13A | NA | -0.08612 | 0.675722 |
| HEXDC | NA | -0.191204 | 0.349439 | SERPINF1 | NA | -0.00060326 | 0.997617284 | LOC254100 | NA | -0.085973 | 0.676244 |
| HEG1 | NA | 0.190972 | 0.350036 | CBR1 | NA | -0.000488697 | 0.998069778 | KDM2B | NA | -0.085418 | 0.678224 |
| VNN1 | NA | -0.190606 | 0.350979 | RSBN1L | NA | 0.00080828 | 0.996807512 | BCAT1 | NA | -0.083828 | 0.683907 |
| PDCD5 | NA | -0.189919 | 0.352749 | WDR18 | NA | 0.000822475 | 0.996751449 | SCARNA13 | NA | -0.083733 | 0.684249 |
| TM4SF1 | NA | -0.189241 | 0.354502 | DEFB1 | NA | 0.000940278 | 0.996286162 | LOC389641 | NA | -0.083246 | 0.685991 |
| TSPAN32 | NA | 0.189215 | 0.354571 | ZNF814 | NA | 0.00107041 | 0.99577218 | ARL5B | NA | -0.083046 | 0.686708 |
| VPS52 | NA | -0.188939 | 0.355286 | LCLAT1 | NA | 0.0011523 | 0.995448741 | ENC1 | NA | -0.082274 | 0.689479 |
| OSBPL8 | NA | 0.188782 | 0.355691 | RGL3 | NA | 0.001295464 | 0.994883291 | SURF6 | NA | -0.082069 | 0.690216 |
| STXBP2 | NA | -0.188526 | 0.356358 | FTSJ3 | NA | 0.001348446 | 0.994674028 | PIP5K1A | NA | -0.081578 | 0.691977 |
| NR4A2 | NA | -0.188119 | 0.357415 | NBPF24 | NA | 0.001425248 | 0.994370688 | DHRS4L2 | NA | -0.081301 | 0.692974 |
| BMF | NA | -0.187974 | 0.357794 | C14orf142 | NA | 0.001726803 | 0.993179661 | KDM5B | NA | -0.080446 | 0.696052 |
| FIBP | NA | -0.187386 | 0.359326 | KDM3B | NA | 0.001749469 | 0.993090137 | ARFIP2 | NA | -0.079848 | 0.698207 |
| TTC27 | NA | -0.187287 | 0.359586 | PHF10 | NA | 0.00202195 | 0.992013951 | UROD | NA | -0.079387 | 0.699871 |
| MED20 | NA | -0.187135 | 0.359981 | EAF1 | NA | 0.002254006 | 0.991097441 | MRPL42 | NA | -0.079172 | 0.700647 |
| IL8 | NA | -0.186912 | 0.360565 | KIN | NA | 0.002536414 | 0.989982077 | RGL1 | NA | -0.078527 | 0.702975 |
| ATAD2 | NA | -0.186825 | 0.360791 | MARVELD1 | NA | 0.002674977 | 0.989434832 | STIP1 | NA | -0.078277 | 0.703879 |
| ACAT2 | NA | -0.186347 | 0.362045 | TBC1D22B | NA | 0.002736131 | 0.989193309 | SNORA1 | NA | -0.076935 | 0.708737 |
| OC1005067 | NA | -0.186159 | 0.362539 | TAF3 | NA | 0.002801801 | 0.988933954 | PPP1R37 | NA | -0.075746 | 0.713048 |
| DRAM2 | NA | -0.185831 | 0.363398 | NCF1B | NA | 0.003038782 | 0.987998033 | LIN54 | NA | -0.075557 | 0.713736 |
| THOC7 | NA | -0.185756 | 0.363596 | HECTD2 | NA | 0.003292614 | 0.98699558 | PCBP2 | NA | -0.075544 | 0.713781 |
| ORC3 | NA | -0.185359 | 0.364641 | RUSC1 | NA | 0.003430674 | 0.98645035 | STARD5 | NA | -0.075117 | 0.715333 |
| LOC728554 | NA | -0.18497 | 0.365666 | ADIPOR2 | NA | 0.003842064 | 0.984825714 | IQSEC2 | NA | -0.07472 | 0.716777 |

Table 6. Gene module membership and correlation with growth rate (48/72h), with adjusted p values for each module

| | | | | | | | | | | | |
|---|---|---|---|---|---|---|---|---|---|---|---|
| ETV6 | NA | -0.184851 | 0.36598 | EEFSEC | NA | 0.0038545 | 0.984776602 | TRIM68 | NA | -0.07413 | 0.718924 |
| NUP88 | NA | -0.184518 | 0.36686 | FAM214A | NA | 0.003956354 | 0.984374377 | FAM134A | NA | -0.073896 | 0.719778 |
| APOL1 | NA | -0.184274 | 0.367505 | SH3PXD2B | NA | 0.004103839 | 0.98379196 | ARHGEF35 | NA | -0.072557 | 0.724659 |
| BYSL | NA | -0.184052 | 0.368092 | CD300LF | NA | 0.00433688 | 0.982871695 | PFN1P2 | NA | -0.071837 | 0.727287 |
| OS9 | NA | -0.183215 | 0.370313 | METTL1 | NA | 0.004371806 | 0.982733777 | PUS1 | NA | -0.071349 | 0.729074 |
| RSBN1 | NA | -0.183086 | 0.370656 | DHX58 | NA | 0.004791652 | 0.981075897 | GSG2 | NA | -0.071283 | 0.729315 |
| NCSTN | NA | -0.182258 | 0.372861 | ARID5B | NA | 0.005536822 | 0.978133573 | FAM102B | NA | -0.071042 | 0.730197 |
| SUMO1 | NA | -0.182167 | 0.373103 | EIF3C | NA | 0.005868795 | 0.976822857 | CHD3 | NA | -0.07095 | 0.730532 |
| RNF149 | NA | -0.182013 | 0.373515 | EPDR1 | NA | 0.006273857 | 0.975223647 | RC3H2 | NA | -0.068713 | 0.738734 |
| LOC145783 | NA | -0.181782 | 0.374131 | FBXL3 | NA | 0.006559638 | 0.974095423 | KPNA3 | NA | -0.068251 | 0.740428 |
| POLL | NA | -0.18118 | 0.375743 | TNFRSF4 | NA | 0.006749991 | 0.97334396 | DUSP6 | NA | -0.066944 | 0.745238 |
| ZCCHC7 | NA | -0.180968 | 0.37631 | LOC220729 | NA | 0.006770713 | 0.973262157 | CIDEB | NA | -0.066801 | 0.745764 |
| ZNF544 | NA | -0.180567 | 0.377386 | FAM213B | NA | 0.006778385 | 0.973231871 | CELSR1 | NA | -0.066549 | 0.746692 |
| RHOBTB1 | NA | -0.180005 | 0.378899 | TRAPPC2P1 | NA | 0.00684476 | 0.972969848 | EIF4EBP3 | NA | -0.065995 | 0.748733 |
| ATXN7L3 | NA | 0.179658 | 0.379833 | CCDC115 | NA | 0.007200179 | 0.97156683 | UBXN4 | NA | -0.0645 | 0.754253 |
| PRKRA | NA | 0.179092 | 0.381363 | FAM125B | NA | 0.007231966 | 0.971441355 | KIAA0391 | NA | -0.061384 | 0.765791 |
| CDR1 | NA | 0.179018 | 0.381561 | NLE1 | NA | 0.007457063 | 0.970552832 | CALML4 | NA | -0.056887 | 0.782529 |
| GDPGP1 | NA | -0.178825 | 0.382084 | CAD | NA | 0.007585328 | 0.97004655 | DRAM2 | NA | -0.056721 | 0.783148 |
| PKM | NA | 0.178667 | 0.382513 | RLF | NA | 0.007918253 | 0.968732491 | PLD2 | NA | -0.056705 | 0.783207 |
| TMCC3 | NA | 0.17855 | 0.382828 | RBM22 | NA | 0.007935119 | 0.968665922 | YWHAE | NA | -0.056469 | 0.78409 |
| RARG | NA | -0.177698 | 0.385141 | GLS | NA | 0.008451314 | 0.966628661 | PKD2L1 | NA | -0.056442 | 0.784191 |
| ELMO2 | NA | -0.177564 | 0.385505 | TAF1 | NA | 0.009187518 | 0.96372345 | SCIN | NA | -0.05575 | 0.786778 |
| MYOZ1 | NA | 0.177517 | 0.385633 | MED11 | NA | 0.009364267 | 0.96302603 | CTSW | NA | -0.05564 | 0.787187 |
| USP22 | NA | -0.1774 | 0.385951 | MRPL19 | NA | 0.009406929 | 0.962857695 | BBS1 | NA | -0.055155 | 0.789001 |
| E4F1 | NA | -0.177337 | 0.386121 | PPARA | NA | 0.009493776 | 0.962515027 | LOC1002165 | NA | -0.054963 | 0.789718 |
| CNPPD1 | NA | 0.177279 | 0.386279 | SEMA6B | NA | 0.009685846 | 0.961757203 | FN3KRP | NA | -0.054811 | 0.79029 |
| NPEPPS | NA | -0.175915 | 0.390004 | THAP8 | NA | 0.009997175 | 0.960528901 | FN3K | NA | -0.054645 | 0.790909 |
| IL11RA | NA | -0.175222 | 0.391902 | CXorf40B | NA | 0.010258395 | 0.959498366 | APPBP2 | NA | -0.051891 | 0.801234 |
| OC1005057 | NA | -0.175197 | 0.391971 | NPC1 | NA | 0.010333074 | 0.959203763 | MAN1A2 | NA | -0.049588 | 0.809894 |
| PDHA1 | NA | -0.175068 | 0.392325 | DPH5 | NA | 0.010364068 | 0.959081496 | TXNDC9 | NA | -0.048626 | 0.813519 |
| HSPBAP1 | NA | -0.17506 | 0.392349 | ZNF382 | NA | 0.010466059 | 0.958679163 | KIAA1609 | NA | -0.048539 | 0.813848 |
| C15orf23 | NA | -0.174599 | 0.393615 | DAPP1 | NA | 0.010606681 | 0.958124452 | COMMD10 | NA | -0.048362 | 0.814515 |
| C1orf35 | NA | -0.174498 | 0.393894 | ZCCHC24 | NA | 0.010866126 | 0.957101068 | ZNF33B | NA | -0.047675 | 0.817106 |
| DUSP16 | NA | -0.173822 | 0.395757 | ZNF546 | NA | 0.010880173 | 0.957045663 | EIF5A | NA | -0.047641 | 0.817232 |
| CYSTM1 | NA | -0.173426 | 0.396852 | ATRN | NA | 0.011232271 | 0.955656921 | CCL18 | NA | -0.047303 | 0.818508 |
| OR6K6 | NA | 0.173328 | 0.397121 | TRAF3 | NA | 0.011404177 | 0.954978936 | CD101 | NA | -0.047279 | 0.818598 |
| ECE1 | NA | -0.173024 | 0.397963 | LANCL2 | NA | 0.012093408 | 0.952260976 | MAZ | NA | -0.047214 | 0.818845 |
| GEMIN5 | NA | -0.172796 | 0.398594 | XRCC6BP1 | NA | 0.01212984 | 0.952117322 | RFXANK | NA | -0.043909 | 0.831341 |
| C11orf83 | NA | -0.1723 | 0.39997 | ICMT | NA | 0.01223311 | 0.951710126 | EEF2K | NA | -0.043695 | 0.83215 |
| ACD | NA | -0.172091 | 0.400551 | SERINC5 | NA | 0.012292227 | 0.951477032 | KLHL24 | NA | -0.043144 | 0.834236 |
| DNASE2B | NA | -0.171845 | 0.401235 | SFXN1 | NA | 0.012356169 | 0.951224919 | CENPP | NA | -0.0422 | 0.837816 |
| C1orf122 | NA | -0.171669 | 0.401725 | CDKN1A | NA | 0.012587211 | 0.950313995 | SLC25A5 | NA | -0.041641 | 0.839936 |
| DIXDC1 | NA | -0.170952 | 0.403724 | SMC1A | NA | 0.013122926 | 0.94820208 | IRF9 | NA | -0.040832 | 0.843011 |
| OC1001282 | NA | -0.170858 | 0.403986 | FAM104A | NA | 0.013472127 | 0.946825629 | KLF5 | NA | -0.040336 | 0.844893 |
| NOC4L | NA | -0.170392 | 0.405288 | CIITA | NA | 0.013504394 | 0.94669845 | ZNF532 | NA | -0.040077 | 0.845877 |
| KCNAB1 | NA | -0.170042 | 0.406269 | METTL7A | NA | 0.014154841 | 0.944135002 | HAGHL | NA | -0.039215 | 0.849155 |
| IFITM10 | NA | 0.169403 | 0.408059 | TRIM65 | NA | 0.014213732 | 0.943902933 | TMBIM4 | NA | -0.037585 | 0.855358 |
| RBM28 | NA | 0.169242 | 0.408512 | ZFX | NA | 0.0144239 | 0.943074782 | ZNF8 | NA | -0.036859 | 0.858125 |
| KIAA0907 | NA | -0.168937 | 0.409369 | MYO9A | NA | 0.01449244 | 0.942804717 | POLD1 | NA | -0.035442 | 0.863528 |
| TMEM14E | NA | 0.168694 | 0.410053 | BCR | NA | 0.014598328 | 0.942387503 | MARCO | NA | -0.034241 | 0.86811 |
| ZNF12 | NA | 0.168629 | 0.410239 | FGD5 | NA | 0.014661796 | 0.942137439 | LOC652276 | NA | -0.034153 | 0.868447 |
| FAM225A | NA | -0.168532 | 0.41051 | KIF27 | NA | 0.014810516 | 0.941551495 | WDR83 | NA | -0.033444 | 0.871155 |
| AKIRIN2 | NA | 0.167751 | 0.412714 | TFCP2L1 | NA | 0.014815388 | 0.941532301 | LOC1001280 | NA | -0.032437 | 0.875007 |
| HDGF | NA | -0.167532 | 0.413333 | LIN54 | NA | 0.014854503 | 0.941378198 | ZNF785 | NA | -0.032119 | 0.876224 |
| VDAC1 | NA | -0.167509 | 0.413399 | GSTK1 | NA | 0.014938874 | 0.941045806 | CRYBB2P1 | NA | -0.032058 | 0.876457 |
| EMILIN2 | NA | 0.16732 | 0.413934 | VAMP4 | NA | 0.015142266 | 0.940244547 | RNPC3 | NA | -0.031188 | 0.879785 |

**Table 6. Gene module membership and correlation with growth rate (48/72h), with adjusted p values for each module**

| | | | | | | | | | | | |
|---|---|---|---|---|---|---|---|---|---|---|---|
| RNF40 | NA | 0.167118 | 0.414506 | FAM210B | NA | 0.01541813 | 0.939157879 | TRIB2 | NA | -0.030549 | 0.882231 |
| POMT2 | NA | -0.167089 | 0.414588 | PTS | NA | 0.01564071 | 0.938281181 | FAM134C | NA | -0.028564 | 0.889834 |
| OR5B21 | NA | 0.166778 | 0.415469 | FBXO21 | NA | 0.015800864 | 0.937650412 | BRD3 | NA | -0.027517 | 0.893849 |
| ZHX1 | NA | -0.166728 | 0.415611 | TUBG1 | NA | 0.016004986 | 0.936846523 | GPD1L | NA | -0.026542 | 0.897592 |
| SACS | NA | 0.166654 | 0.415822 | HLA-DQB2 | NA | 0.016262613 | 0.935832006 | PPBP | NA | -0.025673 | 0.900928 |
| AP3M2 | NA | -0.166544 | 0.416133 | CCNB2 | NA | 0.016419465 | 0.935214385 | PXK | NA | -0.025426 | 0.901878 |
| GIGYF1 | NA | -0.165948 | 0.417828 | HCAR2 | NA | 0.016547901 | 0.934708677 | MSMO1 | NA | -0.025388 | 0.902023 |
| CHPT1 | NA | -0.165892 | 0.417986 | GPA33 | NA | 0.016608973 | 0.934468218 | DNASE2B | NA | -0.025265 | 0.902496 |
| WARS2 | NA | 0.16576 | 0.418362 | DNMBP | NA | 0.01662325 | 0.934412007 | C11orf83 | NA | -0.02494 | 0.903742 |
| ZNF350 | NA | -0.165537 | 0.418996 | RAD9A | NA | 0.017161677 | 0.932292331 | NPM1 | NA | -0.023879 | 0.907821 |
| PLEKHA6 | NA | -0.165523 | 0.419038 | VLDLR | NA | 0.017398026 | 0.931362014 | HESX1 | NA | -0.023682 | 0.908577 |
| EFNA2 | NA | -0.165495 | 0.419116 | GLRX5 | NA | 0.017492492 | 0.930990203 | ALDOA | NA | -0.023089 | 0.910857 |
| O3FAR1 | NA | -0.165444 | 0.419263 | NVL | NA | 0.017718183 | 0.930101957 | TTTY15 | NA | -0.022788 | 0.912018 |
| C1orf109 | NA | -0.165281 | 0.419727 | ZFP92 | NA | 0.017780756 | 0.929855704 | AKR1C1 | NA | -0.022184 | 0.91434 |
| ITGA1 | NA | -0.165092 | 0.420268 | TTTY15 | NA | 0.01801114 | 0.92894909 | BAIAP2 | NA | -0.020935 | 0.919145 |
| HBA2 | NA | -0.165017 | 0.420481 | MMP15 | NA | 0.018495566 | 0.927043054 | ICAM2 | NA | -0.020562 | 0.920584 |
| NEK6 | NA | 0.164748 | 0.421249 | ZNF707 | NA | 0.018511504 | 0.926980352 | IMPDH1 | NA | -0.02002 | 0.922671 |
| CEACAM1 | NA | -0.164559 | 0.42179 | ARIH1 | NA | 0.018623513 | 0.926539695 | FTX | NA | -0.01964 | 0.924132 |
| RBM33 | NA | 0.164358 | 0.422364 | PTGER3 | NA | 0.018731293 | 0.926115698 | CLEC6A | NA | -0.019481 | 0.924747 |
| UNC119B | NA | -0.163509 | 0.424797 | HOXB6 | NA | 0.018770542 | 0.925961302 | XIST | NA | -0.019456 | 0.924844 |
| MKRN1 | NA | -0.163487 | 0.424861 | TMEM220 | NA | 0.018946496 | 0.925269164 | WARS2 | NA | -0.018654 | 0.927933 |
| ERCC6L2 | NA | -0.163422 | 0.425047 | PNP | NA | 0.019449339 | 0.923291469 | FRAT1 | NA | -0.017108 | 0.933892 |
| CBWD6 | NA | -0.163043 | 0.426138 | FAM222B | NA | 0.019569212 | 0.922820072 | RARS2 | NA | -0.016351 | 0.936813 |
| ZNF816 | NA | -0.16284 | 0.42672 | MIF4GD | NA | 0.020004412 | 0.921108869 | PSMD5 | NA | -0.015834 | 0.938804 |
| SUSD1 | NA | -0.162563 | 0.427519 | MFNG | NA | 0.020142424 | 0.920566277 | LY75 | NA | -0.015578 | 0.939794 |
| INTS12 | NA | -0.162416 | 0.427943 | BTBD2 | NA | 0.020361148 | 0.91970644 | RALGAPA2 | NA | -0.015286 | 0.94092 |
| SULT1B1 | NA | -0.162216 | 0.428518 | DHX8 | NA | 0.020493359 | 0.919186745 | RNF144B | NA | -0.014938 | 0.942262 |
| MIR22HG | NA | -0.16208 | 0.428912 | L1TD1 | NA | 0.020603193 | 0.918755029 | KCNQ1OT1 | NA | -0.014706 | 0.943159 |
| FDPS | NA | -0.161844 | 0.429593 | LDLR | NA | 0.020672902 | 0.918481043 | MAPK14 | NA | -0.014574 | 0.943666 |
| POLR2J2 | NA | -0.161814 | 0.429681 | ENGASE | NA | 0.020722158 | 0.918287449 | MPPE1 | NA | -0.01417 | 0.945227 |
| DIP2A | NA | 0.161635 | 0.430197 | PRKD3 | NA | 0.02084547 | 0.917802814 | PHF14 | NA | -0.013814 | 0.946602 |
| FAM198B | NA | 0.161566 | 0.430397 | MOB3B | NA | 0.021332311 | 0.915889731 | FADS2 | NA | -0.013404 | 0.948182 |
| IL10RB | NA | -0.16142 | 0.430821 | MAP2K1 | NA | 0.021571849 | 0.914948613 | SENP3 | NA | -0.013125 | 0.949262 |
| ARSK | NA | -0.160812 | 0.432579 | SNHG5 | NA | 0.021938841 | 0.913506964 | NEK1 | NA | -0.012128 | 0.953111 |
| C10orf88 | NA | 0.160805 | 0.432602 | GAS7 | NA | 0.021968643 | 0.913389904 | DDX18 | NA | -0.011857 | 0.954156 |
| ASTL | NA | 0.160637 | 0.433087 | PPIP5K1 | NA | 0.022213419 | 0.912428515 | L1TD1 | NA | -0.011764 | 0.954514 |
| PPTC7 | NA | 0.160214 | 0.434317 | SLAMF8 | NA | 0.022264905 | 0.912226313 | SOWAHD | NA | -0.010727 | 0.958521 |
| TM4SF19 | NA | 0.160171 | 0.43444 | SGK3 | NA | 0.022295905 | 0.912104568 | ACOT1 | NA | -0.01053 | 0.959282 |
| LGALS3 | NA | -0.159827 | 0.435442 | PYGO2 | NA | 0.02268141 | 0.910590755 | DDIT4 | NA | -0.010504 | 0.959384 |
| DCAF7 | NA | -0.159761 | 0.435635 | CCDC90B | NA | 0.023019312 | 0.909264125 | AKR1C3 | NA | -0.010154 | 0.960737 |
| MKL2 | NA | -0.159467 | 0.43649 | SRPX2 | NA | 0.023569989 | 0.907102628 | FUNDC1 | NA | -0.009863 | 0.961862 |
| TMED9 | NA | -0.159259 | 0.437097 | GDPGP1 | NA | 0.024069744 | 0.90514157 | C12orf43 | NA | -0.008143 | 0.968509 |
| UGDH | NA | 0.159054 | 0.437695 | ZNF432 | NA | 0.024094387 | 0.905044883 | LOC401321 | NA | -0.007062 | 0.972686 |
| SCAPER | NA | -0.158785 | 0.43848 | ZNF429 | NA | 0.024274786 | 0.904337132 | SFXN4 | NA | -0.00682 | 0.973621 |
| CD300C | NA | -0.158721 | 0.438667 | GALNT10 | NA | 0.025058464 | 0.901263399 | ACSL5 | NA | -0.00677 | 0.973816 |
| EED | NA | 0.1587 | 0.438728 | BCL2L11 | NA | 0.025097958 | 0.901108534 | PPP3R1 | NA | -0.006761 | 0.973852 |
| HMOX1 | NA | 0.158598 | 0.439026 | DNLZ | NA | 0.025235003 | 0.900571173 | CERK | NA | -0.005891 | 0.977213 |
| CDKN2D | NA | -0.157044 | 0.443582 | CPEB4 | NA | 0.025236855 | 0.900563912 | PIK3C2B | NA | -0.005431 | 0.978992 |
| SCCPDH | NA | -0.156495 | 0.445196 | FAM220A | NA | 0.025621318 | 0.899056638 | LINC00623 | NA | -0.003316 | 0.987172 |
| TSPAN4 | NA | 0.156262 | 0.445882 | RNF11 | NA | 0.025631374 | 0.899017215 | NGFRAP1 | NA | -0.003114 | 0.987953 |
| PRPSAP2 | NA | -0.1561 | 0.44636 | BCAT1 | NA | 0.02573729 | 0.898602038 | MYO18A | NA | -0.002947 | 0.988599 |
| TRPS1 | NA | -0.156041 | 0.446535 | LIN9 | NA | 0.02602859 | 0.897460315 | NNT | NA | -0.001844 | 0.992865 |
| TMEM135 | NA | -0.155472 | 0.448216 | VPS29 | NA | 0.02614745 | 0.89699451 | PIGU | NA | -0.001496 | 0.994213 |
| BMS1P4 | NA | -0.155432 | 0.448332 | MRPL15 | NA | 0.02630273 | 0.896386026 | FOLR3 | NA | 5.78E-05 | 0.999776 |
| IMPDH1 | NA | -0.15533 | 0.448634 | PROSC | NA | 0.026322822 | 0.896307298 | ANKRD20A9 | NA | 0.000185 | 0.999286 |
| B3GNT5 | NA | 0.155076 | 0.449387 | MSH2 | NA | 0.026453485 | 0.895795333 | C1orf35 | NA | 0.000312 | 0.998793 |

**Table 6. Gene module membership and correlation with growth rate (48/72h), with adjusted p values for each module**

| | | | | | | | | | | | |
|---|---|---|---|---|---|---|---|---|---|---|---|
| SLC38A10 | NA | -0.154953 | 0.449751 | ATP5C1 | NA | 0.026736949 | 0.894684798 | FCHSD2 | NA | 0.000816 | 0.996843 |
| TCEAL3 | NA | -0.154945 | 0.449774 | MOGS | NA | 0.027017875 | 0.893584395 | ZNF187 | NA | 0.001056 | 0.995916 |
| FAM104B | NA | -0.154547 | 0.450951 | DHRS4 | NA | 0.027082531 | 0.893331164 | ZMYM4 | NA | 0.001138 | 0.995599 |
| CYSLTR2 | NA | -0.154156 | 0.452112 | NDRG1 | NA | 0.02710858 | 0.893229141 | CBX6 | NA | 0.001252 | 0.995156 |
| TMEM176E | NA | -0.154015 | 0.45253 | CD22 | NA | 0.027269025 | 0.89260079 | PKIA | NA | 0.001349 | 0.994782 |
| TNFRSF10A | NA | -0.153894 | 0.452892 | MFF | NA | 0.0274072 | 0.892059709 | ZFX | NA | 0.001667 | 0.99355 |
| PSMD6-AS2 | NA | 0.153453 | 0.454202 | CADM1 | NA | 0.027783517 | 0.89058632 | DPCD | NA | 0.001828 | 0.99293 |
| ARID2 | NA | -0.153018 | 0.455498 | HARBI1 | NA | 0.02794411 | 0.88995766 | CYFIP1 | NA | 0.002201 | 0.991488 |
| PHF17 | NA | -0.152994 | 0.455569 | MUC20 | NA | 0.028321477 | 0.888480669 | DGCR2 | NA | 0.00245 | 0.990522 |
| ZNF777 | NA | -0.152962 | 0.455666 | FBXL2 | NA | 0.028520202 | 0.887703016 | VARS | NA | 0.002776 | 0.989263 |
| MFSD8 | NA | -0.152946 | 0.455711 | WDR91 | NA | 0.028577747 | 0.887477851 | ECHS1 | NA | 0.003128 | 0.987899 |
| ZNF845 | NA | -0.152883 | 0.455899 | APOBEC3C | NA | 0.028629676 | 0.887274668 | AVPI1 | NA | 0.003236 | 0.987485 |
| CHPF | NA | -0.152758 | 0.456273 | TRAM2 | NA | 0.028760935 | 0.886761116 | TBCK | NA | 0.003694 | 0.985712 |
| B4GALT5 | NA | -0.152627 | 0.456665 | LOC645212 | NA | 0.02881474 | 0.886550617 | CALM1 | NA | 0.0037 | 0.98569 |
| DDX3X | NA | 0.15232 | 0.457581 | MS4A4A | NA | 0.028997891 | 0.885834141 | EIF2B3 | NA | 0.0037 | 0.985686 |
| RGL3 | NA | -0.15228 | 0.457699 | UPK3BL | NA | 0.029070359 | 0.885550673 | ANKRD29 | NA | 0.003937 | 0.984771 |
| ATP9B | NA | -0.152188 | 0.457975 | KIAA0556 | NA | 0.029084854 | 0.885493976 | HIVEP2 | NA | 0.004278 | 0.983451 |
| FAM21C | NA | 0.152089 | 0.458273 | PRKX | NA | 0.029191269 | 0.885077749 | FUZ | NA | 0.004549 | 0.982403 |
| SDC3 | NA | 0.151937 | 0.458726 | PXN | NA | 0.029408291 | 0.884228996 | MYL12B | NA | 0.005099 | 0.980277 |
| CPNE8 | NA | -0.151182 | 0.460989 | ATP13A3 | NA | 0.030157251 | 0.881300837 | OR6K6 | NA | 0.00556 | 0.978493 |
| CD99 | NA | -0.151135 | 0.461128 | TCHP | NA | 0.030328503 | 0.880631514 | ATAD2 | NA | 0.005588 | 0.978386 |
| ACOX3 | NA | -0.150731 | 0.462343 | FBXW4 | NA | 0.030440172 | 0.880195112 | ARF1 | NA | 0.00661 | 0.974434 |
| QPCT | NA | -0.150581 | 0.462794 | GTPBP1 | NA | 0.030715268 | 0.879120182 | STARD4 | NA | 0.006688 | 0.974132 |
| ZNF366 | NA | -0.150388 | 0.463373 | ATP2B1 | NA | 0.030909155 | 0.878362698 | CYBRD1 | NA | 0.008047 | 0.968879 |
| EIF2A | NA | -0.150297 | 0.463646 | PTGS2 | NA | 0.032249782 | 0.873127991 | ZNF524 | NA | 0.0087 | 0.966353 |
| OR6K3 | NA | 0.149997 | 0.464551 | LOC283922 | NA | 0.032319048 | 0.872857672 | CD163 | NA | 0.008926 | 0.965481 |
| RABEP2 | NA | -0.149832 | 0.465047 | LINC00623 | NA | 0.032330464 | 0.872813118 | RASSF7 | NA | 0.009263 | 0.96418 |
| TSPYL2 | NA | 0.149815 | 0.465099 | RALGAPA2 | NA | 0.033268161 | 0.869155028 | GGA3 | NA | 0.010242 | 0.960394 |
| SLC17A5 | NA | -0.149474 | 0.466125 | ACIN1 | NA | 0.033896627 | 0.866704756 | SUPT3H | NA | 0.010852 | 0.95804 |
| PIK3AP1 | NA | -0.149369 | 0.466443 | CD2AP | NA | 0.03404896 | 0.866111018 | GTSF1 | NA | 0.011372 | 0.956031 |
| MAP3K10 | NA | -0.149287 | 0.466692 | CAMK1D | NA | 0.034060151 | 0.866067402 | ARFIP1 | NA | 0.011426 | 0.955821 |
| JAZF1 | NA | 0.14927 | 0.466743 | NFXL1 | NA | 0.034282402 | 0.865201287 | TMEM143 | NA | 0.01236 | 0.952213 |
| MAL | NA | 0.149245 | 0.466817 | GRWD1 | NA | 0.034420071 | 0.864664861 | PRKRIP1 | NA | 0.012737 | 0.95076 |
| RRN3P3 | NA | 0.148898 | 0.467864 | WBP2 | NA | 0.034686075 | 0.863628552 | HLA-DQB2 | NA | 0.014207 | 0.945082 |
| MT1A | NA | -0.148431 | 0.469278 | SGPP2 | NA | 0.034884588 | 0.86285532 | LOC90834 | NA | 0.01601 | 0.938127 |
| ZCCHC3 | NA | 0.148421 | 0.46931 | FGL2 | NA | 0.034922399 | 0.862708055 | ARHGEF1 | NA | 0.016708 | 0.935432 |
| KRT18 | NA | -0.148119 | 0.470222 | ZNF79 | NA | 0.034932269 | 0.862669614 | STIL | NA | 0.017264 | 0.933292 |
| ATF5 | NA | 0.148076 | 0.470355 | LAMB3 | NA | 0.035051362 | 0.862205809 | PLEK | NA | 0.018104 | 0.930051 |
| ZNF438 | NA | 0.147809 | 0.471165 | PAM | NA | 0.0352837 | 0.861301102 | GNGT2 | NA | 0.018575 | 0.928236 |
| TRAF3IP1 | NA | 0.147803 | 0.471181 | ANAPC13 | NA | 0.03533235 | 0.861111685 | ITFG2 | NA | 0.019866 | 0.923265 |
| CLEC4E | NA | 0.147457 | 0.472232 | PNPLA3 | NA | 0.035476808 | 0.860549283 | VAV2 | NA | 0.020268 | 0.921714 |
| NUP93 | NA | -0.147368 | 0.472503 | FKBP2 | NA | 0.035570859 | 0.860183161 | TLE2 | NA | 0.020838 | 0.919519 |
| B3GNTL1 | NA | -0.147344 | 0.472576 | SGCG | NA | 0.036136326 | 0.85798251 | TOM1L2 | NA | 0.021542 | 0.91681 |
| HPSE | NA | -0.146998 | 0.473627 | HEY1 | NA | 0.03621699 | 0.857668674 | ARHGAP31 | NA | 0.021629 | 0.916475 |
| ELMOD2 | NA | -0.146536 | 0.475035 | ITGA1 | NA | 0.0364538 | 0.856747439 | POFUT2 | NA | 0.021964 | 0.915188 |
| ARID5A | NA | 0.146377 | 0.475517 | SPIRE1 | NA | 0.03645752 | 0.856732967 | UQCRBP1 | NA | 0.022472 | 0.913232 |
| NARF | NA | -0.146061 | 0.476483 | ZNF486 | NA | 0.037075642 | 0.854329233 | PDE6G | NA | 0.024105 | 0.906952 |
| TGM2 | NA | -0.145862 | 0.47709 | ODF3B | NA | 0.037103571 | 0.854220654 | YES1 | NA | 0.024802 | 0.904274 |
| ETS2 | NA | 0.14564 | 0.477768 | FN3KRP | NA | 0.037389143 | 0.853110586 | REC8 | NA | 0.02494 | 0.903742 |
| SP110 | NA | -0.145527 | 0.478112 | HPGDS | NA | 0.037502851 | 0.852668657 | FAM98B | NA | 0.025497 | 0.901606 |
| INSR | NA | 0.145301 | 0.478804 | COPS7B | NA | 0.037808942 | 0.851479246 | USP9Y | NA | 0.027632 | 0.893411 |
| ZNF808 | NA | 0.145255 | 0.478945 | ALMS1 | NA | 0.038148869 | 0.850158726 | TNS1 | NA | 0.028442 | 0.890301 |
| FDX1 | NA | -0.145072 | 0.479504 | PTRHD1 | NA | 0.03818665 | 0.850011983 | FRYL | NA | 0.029138 | 0.887635 |
| SLC46A1 | NA | -0.14506 | 0.479541 | CD97 | NA | 0.038201133 | 0.84995573 | ZFY | NA | 0.029389 | 0.886672 |
| NCF1B | NA | -0.144828 | 0.480251 | CDKN3 | NA | 0.038426693 | 0.849079751 | PRADC1 | NA | 0.029768 | 0.885221 |
| RSC1A1 | NA | 0.144703 | 0.480635 | ARHGAP4 | NA | 0.038493649 | 0.848819757 | NUFIP1 | NA | 0.031419 | 0.878899 |

**Table 6. Gene module membership and correlation with growth rate (48/72h), with adjusted p values for each module**

| | | | | | | | | | | | |
|---|---|---|---|---|---|---|---|---|---|---|---|
| KIF20B | NA | -0.144512 | 0.481221 | HTRA2 | NA | 0.038565144 | 0.848542156 | LOC645212 | NA | 0.031863 | 0.877201 |
| EGLN3 | NA | -0.14445 | 0.481412 | C7orf43 | NA | 0.038608875 | 0.848372362 | POLR1E | NA | 0.03263 | 0.874269 |
| C17orf107 | NA | -0.144447 | 0.481421 | SPTY2D1 | NA | 0.038778283 | 0.847714675 | THAP11 | NA | 0.033911 | 0.869374 |
| ESPN | NA | -0.144023 | 0.482723 | UQCRH | NA | 0.039190529 | 0.846114647 | FKBP15 | NA | 0.034466 | 0.867254 |
| OLR1 | NA | -0.144001 | 0.482789 | TLE4 | NA | 0.039201872 | 0.84607063 | PRKAG2 | NA | 0.03472 | 0.866282 |
| C12orf73 | NA | -0.143933 | 0.482998 | CMSS1 | NA | 0.039483911 | 0.844976318 | JAZF1 | NA | 0.034938 | 0.865451 |
| PLA2G15 | NA | -0.143901 | 0.483097 | ITGA5 | NA | 0.039690479 | 0.844175011 | CNIH | NA | 0.034997 | 0.865227 |
| NBPF11 | NA | -0.142815 | 0.486441 | MECP2 | NA | 0.040143437 | 0.842418454 | FRG1B | NA | 0.035199 | 0.864456 |
| UBQLN4 | NA | -0.142768 | 0.486584 | MAP3K6 | NA | 0.040331286 | 0.8416902 | KIAA0141 | NA | 0.035413 | 0.863638 |
| AGPAT4 | NA | -0.14262 | 0.487041 | IL7 | NA | 0.040547916 | 0.840850523 | GOSR1 | NA | 0.037108 | 0.857177 |
| FANCG | NA | -0.142196 | 0.48835 | MPND | NA | 0.041187404 | 0.838372803 | ETS2 | NA | 0.037476 | 0.855773 |
| CHTOP | NA | -0.142165 | 0.488445 | TBX20 | NA | 0.041628762 | 0.836663619 | HSDL1 | NA | 0.038868 | 0.850474 |
| CABIN1 | NA | -0.14198 | 0.489018 | RASSF8 | NA | 0.041663753 | 0.836528146 | CEP128 | NA | 0.039781 | 0.847003 |
| ATRN | NA | -0.141741 | 0.489758 | METAP1 | NA | 0.041921176 | 0.835531629 | NUMA1 | NA | 0.040902 | 0.842743 |
| RSAD1 | NA | -0.14132 | 0.491062 | CASP9 | NA | 0.041972943 | 0.835331263 | ZNF107 | NA | 0.04092 | 0.842674 |
| RPS29 | NA | 0.141307 | 0.4911 | CEP128 | NA | 0.04227208 | 0.834173635 | NDUFAF6 | NA | 0.041124 | 0.8419 |
| C14orf159 | NA | 0.140786 | 0.492718 | PLCL1 | NA | 0.042344595 | 0.833893062 | SLC12A2 | NA | 0.04156 | 0.840247 |
| C3orf38 | NA | 0.140663 | 0.493098 | EEF2K | NA | 0.042417632 | 0.833610488 | SLC25A6 | NA | 0.042523 | 0.836592 |
| TMEM143 | NA | -0.140629 | 0.493206 | CEACAM8 | NA | 0.042528741 | 0.833180656 | ZNF678 | NA | 0.042563 | 0.836442 |
| SPEN | NA | -0.140011 | 0.495127 | HSD17B14 | NA | 0.043124319 | 0.830877419 | LPCAT1 | NA | 0.043103 | 0.834393 |
| TREML3P | NA | -0.139875 | 0.495547 | ATP5S | NA | 0.043280842 | 0.830272333 | ACBD4 | NA | 0.043332 | 0.833524 |
| ENG | NA | 0.139591 | 0.496433 | USP9Y | NA | 0.04369001 | 0.828691025 | ENO3 | NA | 0.0443 | 0.829858 |
| ZNF107 | NA | 0.139394 | 0.497048 | MAD1L1 | NA | 0.043952139 | 0.827678319 | ZNF441 | NA | 0.045216 | 0.826392 |
| METAP1 | NA | -0.138923 | 0.498515 | PHF17 | NA | 0.044358135 | 0.82611033 | SCAPER | NA | 0.046347 | 0.822119 |
| KIN | NA | -0.138876 | 0.498663 | LPCAT1 | NA | 0.04437824 | 0.8260327 | CADM1 | NA | 0.046537 | 0.821399 |
| GATAD2A | NA | 0.138832 | 0.498801 | REC8 | NA | 0.044483157 | 0.825627616 | EIF1AY | NA | 0.047252 | 0.818701 |
| PXDC1 | NA | -0.138685 | 0.49926 | DGKQ | NA | 0.044849221 | 0.824214589 | NPHP3 | NA | 0.048264 | 0.814883 |
| CRAMP1L | NA | -0.138404 | 0.500137 | DCAF8 | NA | 0.044948916 | 0.82382985 | MED19 | NA | 0.048757 | 0.813025 |
| FAM161B | NA | -0.138314 | 0.500418 | ARMCX1 | NA | 0.045127952 | 0.823139029 | ABI3 | NA | 0.049799 | 0.809101 |
| FN1 | NA | -0.138229 | 0.500686 | ZNF175 | NA | 0.04517813 | 0.822945437 | ATG14 | NA | 0.049893 | 0.808745 |
| GCH1 | NA | 0.138204 | 0.500764 | CCDC47 | NA | 0.045418558 | 0.822017977 | FLNA | NA | 0.049959 | 0.808499 |
| NFAM1 | NA | -0.138202 | 0.500768 | EOGT | NA | 0.045919714 | 0.820085502 | EFCAB13 | NA | 0.050325 | 0.80712 |
| GSTT1 | NA | 0.13749 | 0.503001 | AEBP2 | NA | 0.046125232 | 0.819293314 | SMIM7 | NA | 0.051047 | 0.804404 |
| ZNF582 | NA | -0.137389 | 0.503316 | AKR1C3 | NA | 0.046789892 | 0.81673251 | REXO4 | NA | 0.051499 | 0.802708 |
| RAI1 | NA | 0.136907 | 0.504831 | WIZ | NA | 0.047574204 | 0.813713068 | LINC00515 | NA | 0.052878 | 0.797531 |
| URB1 | NA | -0.136866 | 0.50496 | ARFIP2 | NA | 0.047767369 | 0.812969822 | GUSBP11 | NA | 0.053401 | 0.795571 |
| USP28 | NA | -0.136629 | 0.505702 | ASAP2 | NA | 0.047792502 | 0.812873127 | ZNF431 | NA | 0.05386 | 0.79385 |
| SNX15 | NA | -0.136325 | 0.50666 | KLHL5 | NA | 0.047954494 | 0.812249966 | NOTCH2 | NA | 0.054866 | 0.790083 |
| CD69 | NA | -0.136148 | 0.507216 | RNPC3 | NA | 0.048287502 | 0.810969274 | FNIP1 | NA | 0.055651 | 0.787144 |
| PEBP4 | NA | -0.135951 | 0.507839 | STXBP1 | NA | 0.048554849 | 0.809941444 | HLA-DPA1 | NA | 0.057287 | 0.781035 |
| KLHL24 | NA | -0.135696 | 0.508641 | NECAB3 | NA | 0.048921986 | 0.808530474 | LIG3 | NA | 0.057334 | 0.78086 |
| MOCOS | NA | -0.135036 | 0.510723 | ARSK | NA | 0.049061706 | 0.80799366 | SLC7A5P2 | NA | 0.057358 | 0.780771 |
| TFAP4 | NA | -0.134609 | 0.512075 | HCG18 | NA | 0.049361989 | 0.806840233 | MPV17L | NA | 0.05744 | 0.780464 |
| INPP5E | NA | -0.134586 | 0.512147 | ELL2 | NA | 0.049366381 | 0.806823368 | LEMD2 | NA | 0.057758 | 0.779277 |
| SYT11 | NA | 0.134548 | 0.512266 | ZMYND19 | NA | 0.049618984 | 0.8058534 | IL36B | NA | 0.058037 | 0.77824 |
| WDR41 | NA | -0.134059 | 0.513816 | C12orf43 | NA | 0.04965346 | 0.805721035 | RNF135 | NA | 0.059347 | 0.773361 |
| PPHLN1 | NA | -0.133923 | 0.514245 | APBB2 | NA | 0.049899225 | 0.804777631 | MNAT1 | NA | 0.059917 | 0.771241 |
| TSC1 | NA | -0.1333 | 0.516224 | PTK2B | NA | 0.049906913 | 0.804748124 | EXT1 | NA | 0.060392 | 0.769474 |
| PPP1CA | NA | -0.132707 | 0.518109 | GPANK1 | NA | 0.050312919 | 0.803190208 | LOC282997 | NA | 0.060631 | 0.768586 |
| TMEM167A | NA | -0.132655 | 0.518274 | PTGS1 | NA | 0.050536207 | 0.802333724 | OR6N1 | NA | 0.060669 | 0.768446 |
| CXADR | NA | -0.132609 | 0.51842 | USP49 | NA | 0.050607526 | 0.802060209 | TAS2R14 | NA | 0.060745 | 0.768164 |
| PPIL1 | NA | -0.132271 | 0.519497 | PELP1 | NA | 0.050808692 | 0.801288836 | POFUT1 | NA | 0.060931 | 0.767473 |
| LOC642852 | NA | 0.132163 | 0.519839 | PMS2P5 | NA | 0.051024421 | 0.800461821 | MAL | NA | 0.061127 | 0.766745 |
| USP13 | NA | -0.131861 | 0.520802 | GPBP1 | NA | 0.052144703 | 0.796170536 | SETD5 | NA | 0.062105 | 0.763116 |
| ZNF672 | NA | -0.131724 | 0.521242 | INPP5D | NA | 0.052170296 | 0.79607257 | SORL1 | NA | 0.062115 | 0.763081 |
| KIF3C | NA | 0.131628 | 0.521548 | PAK1 | NA | 0.052185433 | 0.796014626 | RPS4Y1 | NA | 0.06257 | 0.761394 |

**Table 6. Gene module membership and correlation with growth rate (48/72h), with adjusted p values for each module**

| | | | | | | | | | | | |
|---|---|---|---|---|---|---|---|---|---|---|---|
| SPOCD1 | NA | 0.131308 | 0.522569 | MLX | NA | 0.052934927 | 0.793147017 | GTF2E1 | NA | 0.062977 | 0.759887 |
| C1orf162 | NA | -0.131003 | 0.523543 | FAM200A | NA | 0.053131905 | 0.792393803 | ATP6V0E2 | NA | 0.063049 | 0.759619 |
| BBS1 | NA | -0.130897 | 0.523883 | PARP4 | NA | 0.053165737 | 0.792264452 | HMGCR | NA | 0.06343 | 0.758209 |
| ADAMTS15 | NA | 0.130673 | 0.5246 | BOD1 | NA | 0.053196608 | 0.792146428 | LRRC37A4P | NA | 0.063582 | 0.757647 |
| ADCY3 | NA | -0.130497 | 0.525164 | H2AFZ | NA | 0.053576952 | 0.790692677 | ELOF1 | NA | 0.06362 | 0.757506 |
| HMGB1 | NA | -0.130053 | 0.526584 | HIP1 | NA | 0.053615396 | 0.790545776 | SLC50A1 | NA | 0.06404 | 0.755952 |
| METTL7A | NA | -0.130026 | 0.526673 | OXLD1 | NA | 0.054453597 | 0.787344576 | ATXN7L3B | NA | 0.064118 | 0.755665 |
| CBWD2 | NA | -0.12988 | 0.52714 | NPRL2 | NA | 0.055258055 | 0.784275413 | GTF2E2 | NA | 0.065833 | 0.749331 |
| BNIP3 | NA | 0.129776 | 0.527474 | OS9 | NA | 0.055415738 | 0.783674193 | ERH | NA | 0.066408 | 0.747212 |
| EMP1 | NA | 0.129727 | 0.52763 | FAM195B | NA | 0.055429523 | 0.783621637 | ZBTB1 | NA | 0.067965 | 0.741483 |
| ZNF687 | NA | -0.129345 | 0.528859 | NFYA | NA | 0.056099629 | 0.781068016 | CLEC4E | NA | 0.067979 | 0.741429 |
| ABCD3 | NA | -0.129338 | 0.52888 | TMEM180 | NA | 0.056107825 | 0.781036795 | DEFB1 | NA | 0.068504 | 0.739501 |
| CEP72 | NA | -0.128529 | 0.531484 | FAH | NA | 0.056200123 | 0.780685245 | LOC606724 | NA | 0.070203 | 0.733269 |
| BANP | NA | 0.128094 | 0.532883 | SPTSSA | NA | 0.056270101 | 0.780418738 | NUTF2 | NA | 0.070221 | 0.733202 |
| NQO2 | NA | 0.127887 | 0.533554 | OPN3 | NA | 0.056472184 | 0.77964925 | PLA2G6 | NA | 0.07281 | 0.723737 |
| TM4SF4 | NA | 0.127668 | 0.534258 | CENPP | NA | 0.056531438 | 0.779423663 | UPK3BL | NA | 0.073754 | 0.720293 |
| ZNF528 | NA | -0.127485 | 0.53485 | NCAPD3 | NA | 0.056562508 | 0.77930538 | IFI16 | NA | 0.074004 | 0.719385 |
| MRPL50 | NA | 0.127286 | 0.535494 | ABCF3 | NA | 0.056641951 | 0.779002971 | FAM76B | NA | 0.074926 | 0.71603 |
| RASAL2 | NA | -0.12679 | 0.537098 | TNFRSF1B | NA | 0.056675482 | 0.778875339 | NQO2 | NA | 0.076921 | 0.708785 |
| C1orf123 | NA | 0.126774 | 0.537149 | TXNDC17 | NA | 0.056829653 | 0.778288582 | KDM5D | NA | 0.078309 | 0.703764 |
| GFER | NA | -0.126613 | 0.537669 | RBP1 | NA | 0.057192235 | 0.776909102 | LRRC47 | NA | 0.078487 | 0.703118 |
| ZNF615 | NA | -0.126558 | 0.53785 | WARS | NA | 0.057238204 | 0.776734257 | STYX | NA | 0.079448 | 0.69965 |
| DGUOK | NA | -0.126432 | 0.538257 | SRCAP | NA | 0.057469634 | 0.775854158 | LOC1005067 | NA | 0.079473 | 0.699558 |
| EEPD1 | NA | 0.126277 | 0.538759 | TPT1-AS1 | NA | 0.057770056 | 0.774712091 | DDX3Y | NA | 0.080872 | 0.694516 |
| MED28 | NA | -0.126166 | 0.53912 | QSOX1 | NA | 0.05786909 | 0.774335712 | LOC1005065 | NA | 0.081272 | 0.69308 |
| PIGL | NA | 0.126121 | 0.539265 | TANC2 | NA | 0.05796038 | 0.773988803 | ATP5L2 | NA | 0.082425 | 0.688938 |
| FOSL1 | NA | -0.125861 | 0.540109 | GAL | NA | 0.059275571 | 0.768995756 | FAM133B | NA | 0.083817 | 0.683948 |
| SIRT1 | NA | -0.125422 | 0.541535 | ZNF324 | NA | 0.059287411 | 0.768950846 | NIN | NA | 0.084042 | 0.68314 |
| ERCC4 | NA | -0.125064 | 0.542696 | CETN2 | NA | 0.059296262 | 0.768917275 | C19orf54 | NA | 0.084173 | 0.682675 |
| MRPS21 | NA | 0.124921 | 0.543163 | MMACHC | NA | 0.059394678 | 0.768544012 | ZNF518B | NA | 0.085342 | 0.678497 |
| BACE2 | NA | -0.124847 | 0.543404 | ZNF18 | NA | 0.059638256 | 0.767620412 | CASP2 | NA | 0.087092 | 0.672261 |
| DUSP12 | NA | -0.124256 | 0.54533 | ZNF213 | NA | 0.059732258 | 0.767264061 | OR6K3 | NA | 0.089495 | 0.663732 |
| ZC3H7B | NA | -0.12384 | 0.546687 | NPAS2 | NA | 0.059732531 | 0.767263023 | ADAMTSL4 | NA | 0.08962 | 0.663287 |
| NECAB3 | NA | -0.123373 | 0.548211 | ZNF17 | NA | 0.059815597 | 0.766948168 | KPTN | NA | 0.090287 | 0.660929 |
| TBKBP1 | NA | 0.123246 | 0.548626 | CASP7 | NA | 0.059851971 | 0.766810305 | ULK2 | NA | 0.091702 | 0.655933 |
| UACA | NA | -0.122469 | 0.551168 | ACVR2B | NA | 0.05994891 | 0.766442925 | GPR64 | NA | 0.094211 | 0.647113 |
| ERV3-1 | NA | -0.122463 | 0.551188 | MYO10 | NA | 0.060655887 | 0.763765119 | WBP11 | NA | 0.094449 | 0.646278 |
| UCKL1 | NA | -0.122116 | 0.552324 | TRIB2 | NA | 0.060703042 | 0.763586608 | GRAMD1B | NA | 0.095612 | 0.642204 |
| RNF122 | NA | 0.122018 | 0.552645 | CD38 | NA | 0.060741563 | 0.763440787 | BST2 | NA | 0.09569 | 0.641932 |
| NAGLU | NA | -0.121911 | 0.552999 | NUP210 | NA | 0.06078814 | 0.763264479 | C9orf3 | NA | 0.096158 | 0.640298 |
| DDR2 | NA | -0.12183 | 0.553263 | DFNA5 | NA | 0.060790304 | 0.763256288 | SEC22C | NA | 0.096819 | 0.637991 |
| LSM12 | NA | -0.121771 | 0.553457 | ZNF253 | NA | 0.062185216 | 0.757981609 | CRIM1 | NA | 0.097421 | 0.635894 |
| GNG5 | NA | -0.12151 | 0.554314 | PLCB3 | NA | 0.062956247 | 0.755070571 | CHIT1 | NA | 0.097748 | 0.634755 |
| CD44 | NA | 0.121287 | 0.555047 | ALDH2 | NA | 0.063567712 | 0.752764292 | CYSTM1 | NA | 0.09779 | 0.634606 |
| PTGR2 | NA | -0.12117 | 0.55543 | COTL1 | NA | 0.063844518 | 0.751720937 | VPS33B | NA | 0.098144 | 0.633375 |
| TMED8 | NA | -0.120429 | 0.557869 | FAM212B | NA | 0.063953691 | 0.751309549 | ARSK | NA | 0.098827 | 0.631003 |
| CEP128 | NA | -0.120406 | 0.557944 | AMMECR1 | NA | 0.064112118 | 0.750712684 | PEX5 | NA | 0.101791 | 0.620745 |
| BRD2 | NA | 0.120247 | 0.55847 | NCKAP5L | NA | 0.064863745 | 0.747882876 | TSPAN3 | NA | 0.102216 | 0.619278 |
| ACTR1A | NA | -0.120041 | 0.559147 | GOLGA1 | NA | 0.064892186 | 0.747775859 | EBP | NA | 0.1029 | 0.616925 |
| CAMP | NA | 0.119991 | 0.559314 | KIAA0141 | NA | 0.06496312 | 0.747508976 | FRY | NA | 0.10486 | 0.610194 |
| NUDT6 | NA | -0.119883 | 0.55967 | A4GALT | NA | 0.064977291 | 0.747455662 | UBXN6 | NA | 0.10509 | 0.609408 |
| PIGM | NA | -0.119871 | 0.55971 | PTRH1 | NA | 0.065533831 | 0.745362747 | GLB1L | NA | 0.106292 | 0.605297 |
| PPARGC1B | NA | -0.119726 | 0.560187 | FAM76A | NA | 0.065718219 | 0.744669728 | SH3PXD2B | NA | 0.106997 | 0.602893 |
| PRADC1 | NA | -0.119417 | 0.561209 | DNAJC21 | NA | 0.065927126 | 0.74388479 | PLEKHA2 | NA | 0.107112 | 0.6025 |
| USP7 | NA | 0.119275 | 0.561678 | F3 | NA | 0.065983359 | 0.743673545 | TREML3P | NA | 0.107971 | 0.599579 |
| PDGFB | NA | 0.119272 | 0.561688 | EBNA1BP2 | NA | 0.06635192 | 0.742289459 | MCM3 | NA | 0.108678 | 0.597176 |

**Table 6. Gene module membership and correlation with growth rate (48/72h), with adjusted p values for each module**

| | | | | | | | | | | | |
|---|---|---|---|---|---|---|---|---|---|---|---|
| SLC7A8 | NA | -0.119189 | 0.561962 | PTMA | NA | 0.066391691 | 0.742140151 | H2AFY2 | NA | 0.109979 | 0.592766 |
| GSDMD | NA | -0.11902 | 0.56252 | ZNF780A | NA | 0.066780943 | 0.7406793 | ZFYVE19 | NA | 0.110938 | 0.589526 |
| LOC654433 | NA | -0.118848 | 0.563088 | ZNF169 | NA | 0.067409844 | 0.738320907 | ZBED4 | NA | 0.111239 | 0.588511 |
| PGRMC1 | NA | 0.118295 | 0.564919 | CFD | NA | 0.067621682 | 0.73752703 | DUS2L | NA | 0.111448 | 0.587805 |
| HOMER2 | NA | -0.118278 | 0.564977 | USP20 | NA | 0.067701013 | 0.737229797 | LRRC16A | NA | 0.111536 | 0.587507 |
| PTPRC | NA | -0.118224 | 0.565155 | AATF | NA | 0.06780465 | 0.736841553 | ATP2B1 | NA | 0.111571 | 0.587388 |
| FFAR2 | NA | -0.117922 | 0.566156 | SLC6A6 | NA | 0.067909637 | 0.736448319 | PLXNA2 | NA | 0.112312 | 0.584895 |
| PRKACB | NA | -0.117749 | 0.566729 | TMED8 | NA | 0.06801259 | 0.736062762 | TTC30B | NA | 0.112515 | 0.584211 |
| RPL9 | NA | -0.117567 | 0.567333 | ZC3H13 | NA | 0.068139348 | 0.735588144 | SIGLEC5 | NA | 0.114655 | 0.577036 |
| ZNF860 | NA | 0.117211 | 0.568515 | USP6NL | NA | 0.068492533 | 0.734266221 | ANKMY1 | NA | 0.114781 | 0.576614 |
| TAF13 | NA | 0.11705 | 0.569051 | SLC9A1 | NA | 0.068649634 | 0.733678451 | HERC4 | NA | 0.11553 | 0.574111 |
| FBXL12 | NA | 0.11687 | 0.56965 | DDX60 | NA | 0.068889633 | 0.732780809 | CDR1 | NA | 0.115832 | 0.573105 |
| CIDECP | NA | -0.116846 | 0.569727 | CNNM4 | NA | 0.069286852 | 0.731295895 | C12orf65 | NA | 0.115856 | 0.573026 |
| OSBPL10 | NA | 0.116781 | 0.569946 | DGKD | NA | 0.069409618 | 0.73083715 | HABP4 | NA | 0.116249 | 0.571716 |
| LOC652276 | NA | 0.116721 | 0.570146 | FMR1 | NA | 0.069582841 | 0.730190017 | MED4 | NA | 0.117756 | 0.566708 |
| AACS | NA | -0.115837 | 0.573089 | BTN2A2 | NA | 0.069635886 | 0.729991884 | C1QA | NA | 0.119269 | 0.561698 |
| ZNF561 | NA | -0.115809 | 0.573182 | TM4SF19 | NA | 0.069920244 | 0.72893005 | BTD | NA | 0.11986 | 0.559746 |
| ZFP28 | NA | -0.114354 | 0.578044 | IGF1 | NA | 0.069995099 | 0.728650613 | BNIP3 | NA | 0.120266 | 0.558406 |
| SUMO1P1 | NA | -0.114352 | 0.578048 | ZCCHC3 | NA | 0.070519943 | 0.726692288 | CRELD1 | NA | 0.120596 | 0.55732 |
| PRKCA | NA | 0.11418 | 0.578625 | LSM3 | NA | 0.070632719 | 0.726271713 | PMS2P5 | NA | 0.120851 | 0.556482 |
| FBXL3 | NA | -0.114118 | 0.578832 | SLC1A4 | NA | 0.070841319 | 0.72549398 | NFRKB | NA | 0.123719 | 0.547081 |
| FGD6 | NA | 0.114103 | 0.578882 | HLA-E | NA | 0.071303754 | 0.723770815 | TAS2R19 | NA | 0.123947 | 0.546337 |
| RPL36A | NA | -0.113608 | 0.580543 | CDK2 | NA | 0.071332142 | 0.723665075 | NR2C2 | NA | 0.124404 | 0.544848 |
| GNG12 | NA | 0.113563 | 0.580692 | ZFY | NA | 0.071396547 | 0.723425199 | FDX1 | NA | 0.124763 | 0.543677 |
| LRRC57 | NA | -0.113215 | 0.581859 | DRAP1 | NA | 0.071468207 | 0.723158335 | RNF103 | NA | 0.125076 | 0.542657 |
| DDX10 | NA | -0.113182 | 0.581971 | PI4KAP1 | NA | 0.071803467 | 0.721910218 | HERPUD2 | NA | 0.125297 | 0.541941 |
| SLC8A1 | NA | 0.11315 | 0.58208 | FAM109B | NA | 0.072211819 | 0.720390936 | LINC00641 | NA | 0.127216 | 0.535719 |
| EXOSC7 | NA | -0.113124 | 0.582165 | CBWD2 | NA | 0.072236204 | 0.720300244 | ZNF326 | NA | 0.130059 | 0.526566 |
| ACVR2A | NA | -0.113008 | 0.582556 | DDX3Y | NA | 0.072405757 | 0.719669747 | SIDT2 | NA | 0.131158 | 0.523048 |
| CD300E | NA | -0.112976 | 0.582662 | METTL21A | NA | 0.072464489 | 0.719451387 | PDGFB | NA | 0.131607 | 0.521615 |
| GPR137B | NA | 0.112851 | 0.583083 | IPPK | NA | 0.072531593 | 0.719201931 | SNHG11 | NA | 0.131637 | 0.521517 |
| NABP2 | NA | -0.112744 | 0.583443 | ARHGDIB | NA | 0.072631358 | 0.718831106 | ALS2 | NA | 0.132785 | 0.51786 |
| SLC25A19 | NA | -0.112466 | 0.584377 | TCERG1 | NA | 0.073082711 | 0.717154222 | CSNK1D | NA | 0.133643 | 0.515135 |
| BUD13 | NA | -0.112313 | 0.584891 | EIF1AY | NA | 0.073250574 | 0.716530897 | CAMSAP3 | NA | 0.134692 | 0.511813 |
| IFI27 | NA | -0.111983 | 0.586002 | IFIT1 | NA | 0.073438045 | 0.715834971 | SARNP | NA | 0.135073 | 0.510606 |
| BDH2 | NA | -0.111722 | 0.586882 | RPL9 | NA | 0.073728126 | 0.714758577 | PTTG1IP | NA | 0.135207 | 0.510184 |
| CCDC53 | NA | -0.111671 | 0.587054 | FITM2 | NA | 0.073805759 | 0.714470595 | AVEN | NA | 0.136122 | 0.5073 |
| MEF2B | NA | 0.11149 | 0.587663 | SNORA81 | NA | 0.074116748 | 0.713317359 | PLCG2 | NA | 0.137192 | 0.503934 |
| DDX3Y | NA | 0.111484 | 0.587685 | SMARCA4 | NA | 0.074543108 | 0.711737297 | MSN | NA | 0.138248 | 0.500625 |
| NIN | NA | 0.110661 | 0.59046 | WDR6 | NA | 0.074597286 | 0.7115366 | SCD | NA | 0.139193 | 0.497675 |
| KITLG | NA | -0.110582 | 0.590726 | STAU2 | NA | 0.074942973 | 0.710256476 | LDLR | NA | 0.140622 | 0.493225 |
| HEATR2 | NA | 0.109942 | 0.592891 | KLHL7 | NA | 0.07507224 | 0.709777983 | RABGGTB | NA | 0.140799 | 0.492676 |
| LOC220906 | NA | -0.109675 | 0.593796 | OAT | NA | 0.075078635 | 0.709754315 | CST3 | NA | 0.142358 | 0.487851 |
| )C1002165[illegible] | NA | 0.109002 | 0.596077 | LOC1005057[illegible] | NA | 0.075250392 | 0.709118711 | LCLAT1 | NA | 0.142889 | 0.486211 |
| IL7 | NA | -0.108928 | 0.596326 | AIFM2 | NA | 0.075869533 | 0.706829101 | SIGLEC14 | NA | 0.14368 | 0.483777 |
| GBP3 | NA | -0.108792 | 0.596789 | EIF1AX | NA | 0.076090418 | 0.70601286 | STX17 | NA | 0.143973 | 0.482877 |
| ERRFI1 | NA | 0.106968 | 0.602993 | TEKT4P2 | NA | 0.076292432 | 0.70526663 | HPS6 | NA | 0.144414 | 0.481521 |
| SIGIRR | NA | 0.105315 | 0.608636 | ASAP1 | NA | 0.076364946 | 0.704998833 | OASL | NA | 0.145929 | 0.476886 |
| CRIM1 | NA | 0.105132 | 0.609261 | SGTA | NA | 0.077281986 | 0.701615126 | LARP4 | NA | 0.145983 | 0.47672 |
| MED4 | NA | 0.10493 | 0.609954 | KDM5D | NA | 0.077466363 | 0.700935474 | ZNF687 | NA | 0.146098 | 0.476369 |
| ZNF846 | NA | -0.104825 | 0.610313 | BHLHE41 | NA | 0.07794066 | 0.699188159 | GFER | NA | 0.147229 | 0.472926 |
| TBC1D14 | NA | -0.104743 | 0.610594 | CCL3 | NA | 0.077976109 | 0.699057622 | CCDC125 | NA | 0.147546 | 0.471963 |
| GNB5 | NA | -0.104357 | 0.611918 | HTRA4 | NA | 0.078049398 | 0.698787777 | WIZ | NA | 0.147567 | 0.471898 |
| MAD2L1 | NA | -0.104224 | 0.612376 | SFI1 | NA | 0.07807707 | 0.698685896 | ITGA3 | NA | 0.147669 | 0.471587 |
| ARFIP2 | NA | -0.104186 | 0.612504 | KLF11 | NA | 0.078281534 | 0.697933294 | ZNF77 | NA | 0.14873 | 0.468374 |
| USP49 | NA | -0.103979 | 0.613215 | DIS3 | NA | 0.078536426 | 0.696995463 | SNX7 | NA | 0.149172 | 0.467039 |

**Table 6. Gene module membership and correlation with growth rate (48/72h), with adjusted p values for each module**

| | | | | | | | | | | | |
|---|---|---|---|---|---|---|---|---|---|---|---|
| HAPLN3 | NA | -0.103453 | 0.615023 | CRIM1 | NA | 0.078676578 | 0.696479984 | P2RY11 | NA | 0.150649 | 0.462589 |
| LY75 | NA | 0.103379 | 0.615276 | TNFAIP6 | NA | 0.078922482 | 0.695575866 | EIF2C1 | NA | 0.152271 | 0.457726 |
| ACVR1 | NA | 0.103345 | 0.615394 | OCSTAMP | NA | 0.079194999 | 0.694574373 | ETV5 | NA | 0.152827 | 0.456066 |
| UBE2N | NA | 0.102932 | 0.616813 | TTLL12 | NA | 0.079629271 | 0.692979472 | AMICA1 | NA | 0.153485 | 0.454107 |
| PPP1R26 | NA | 0.10243 | 0.618542 | UBE2N | NA | 0.079748529 | 0.692541707 | WDR35 | NA | 0.154161 | 0.452096 |
| PPIP5K1 | NA | -0.102019 | 0.619957 | PVALB | NA | 0.079921251 | 0.691907867 | INPP5D | NA | 0.155705 | 0.447526 |
| ESR1 | NA | 0.101737 | 0.62093 | PCED1B-AS1 | NA | 0.080845883 | 0.688518169 | RBM33 | NA | 0.158308 | 0.439873 |
| CD93 | NA | 0.101625 | 0.621318 | FAM98B | NA | 0.081253684 | 0.687025032 | POMZP3 | NA | 0.158653 | 0.438865 |
| CYTH4 | NA | -0.101579 | 0.621474 | NFATC3 | NA | 0.08145266 | 0.686296909 | FAM104A | NA | 0.159277 | 0.437045 |
| AKR1C3 | NA | -0.101311 | 0.622401 | NMT1 | NA | 0.081479168 | 0.686199926 | GTPBP1 | NA | 0.15961 | 0.436073 |
| CCDC171 | NA | -0.100974 | 0.623567 | APOC2 | NA | 0.081627591 | 0.685656999 | LOC399753 | NA | 0.160778 | 0.432679 |
| LZIC | NA | -0.100963 | 0.623603 | DNA2 | NA | 0.081881949 | 0.684726923 | UNC5B | NA | 0.161814 | 0.42968 |
| MYO6 | NA | 0.100368 | 0.625659 | PDCD11 | NA | 0.082150972 | 0.683743706 | CCNYL1 | NA | 0.163937 | 0.423569 |
| ENTPD5 | NA | -0.100258 | 0.626042 | HCFC1 | NA | 0.082566662 | 0.682225448 | P2RX1 | NA | 0.165399 | 0.419389 |
| PBLD | NA | 0.100044 | 0.626781 | COPG2 | NA | 0.082617961 | 0.68203817 | KCNJ5 | NA | 0.168117 | 0.411681 |
| APBA1 | NA | 0.099508 | 0.628639 | NEO1 | NA | 0.082734384 | 0.681613204 | NANOG | NA | 0.1688 | 0.409755 |
| HECTD2 | NA | -0.099326 | 0.629271 | LRRC37A | NA | 0.082902531 | 0.680999608 | SPG7 | NA | 0.17094 | 0.403757 |
| ERVK13-1 | NA | 0.099304 | 0.629349 | OSBPL11 | NA | 0.08336546 | 0.679311324 | CHMP4A | NA | 0.174906 | 0.39277 |
| KRR1 | NA | -0.09927 | 0.629467 | TMEM42 | NA | 0.084299456 | 0.675909681 | NOL6 | NA | 0.175152 | 0.392096 |
| MVK | NA | -0.09923 | 0.629604 | ZNF43 | NA | 0.085266874 | 0.672392844 | CES1 | NA | 0.178079 | 0.384105 |
| FXYD6 | NA | -0.099095 | 0.630073 | GOLGA3 | NA | 0.085383338 | 0.671969916 | LOC644961 | NA | 0.181209 | 0.375665 |
| UBE2J2 | NA | 0.099051 | 0.630226 | TBC1D14 | NA | 0.085489621 | 0.671584046 | LILRB1 | NA | 0.18133 | 0.375342 |
| NSUN2 | NA | 0.099033 | 0.630289 | BCAS2 | NA | 0.086931136 | 0.666358536 | ZNHIT2 | NA | 0.18229 | 0.372774 |
| PVRL1 | NA | -0.098834 | 0.63098 | TMCO4 | NA | 0.087296306 | 0.665037189 | RNPEPL1 | NA | 0.184028 | 0.368155 |
| KLHDC4 | NA | 0.098562 | 0.631924 | PTGR2 | NA | 0.087342855 | 0.664868823 | HSPBAP1 | NA | 0.184047 | 0.368104 |
| FGD5 | NA | -0.098518 | 0.632076 | RPS4Y1 | NA | 0.087837677 | 0.663080062 | ZSCAN5A | NA | 0.185961 | 0.363059 |
| LRFN4 | NA | -0.09849 | 0.632172 | DHRS1 | NA | 0.088144978 | 0.661970089 | RBM12B-AS | NA | 0.187392 | 0.359312 |
| ZAK | NA | -0.098225 | 0.633095 | CDC42SE1 | NA | 0.088990998 | 0.658917863 | C3orf38 | NA | 0.18759 | 0.358793 |
| UQCC | NA | 0.097869 | 0.634334 | TMEM9 | NA | 0.089506984 | 0.657058925 | KDM2A | NA | 0.187894 | 0.358003 |
| BLVRB | NA | -0.09775 | 0.634748 | DDX54 | NA | 0.089759968 | 0.65614823 | CPEB2 | NA | 0.188732 | 0.355822 |
| NEK4 | NA | -0.097397 | 0.635976 | NAB1 | NA | 0.090191606 | 0.654595518 | NEAT1 | NA | 0.189761 | 0.353158 |
| ZDHHC13 | NA | -0.096844 | 0.637902 | DLST | NA | 0.090347739 | 0.654034211 | HERC5 | NA | 0.191458 | 0.348787 |
| ALKBH8 | NA | -0.095923 | 0.641119 | LIG1 | NA | 0.090371526 | 0.653948713 | VAV3 | NA | 0.19695 | 0.33487 |
| AIMP2 | NA | -0.095766 | 0.641666 | C17orf89 | NA | 0.090574314 | 0.653219991 | HERC6 | NA | 0.197731 | 0.332918 |
| TCN2 | NA | -0.095073 | 0.64409 | ISCU | NA | 0.090596085 | 0.653141777 | UBE3D | NA | 0.198545 | 0.330891 |
| TRPV4 | NA | 0.094665 | 0.645522 | SUN2 | NA | 0.090637146 | 0.652994269 | NBEAL2 | NA | 0.19883 | 0.330184 |
| TMTC1 | NA | -0.094663 | 0.645527 | NT5C3L | NA | 0.090732004 | 0.652653544 | CMSS1 | NA | 0.201644 | 0.323243 |
| CPEB2 | NA | 0.094449 | 0.646276 | IGF1R | NA | 0.090888705 | 0.652090839 | SNORA74B | NA | 0.201729 | 0.323034 |
| TMEM63A | NA | 0.094438 | 0.646316 | PKIA | NA | 0.091181144 | 0.651041197 | PPM1K | NA | 0.203523 | 0.318658 |
| RAB11FIP5 | NA | 0.093781 | 0.648621 | PPP2R5C | NA | 0.091196849 | 0.650984847 | RTN4R | NA | 0.205662 | 0.313489 |
| FAM70A | NA | -0.09346 | 0.649747 | CCND1 | NA | 0.091270015 | 0.650722345 | IMMP2L | NA | 0.20695 | 0.310403 |
| APPBP2 | NA | -0.093392 | 0.649985 | USPL1 | NA | 0.091567706 | 0.649654721 | PPDPF | NA | 0.209139 | 0.305196 |
| SLC12A6 | NA | 0.093255 | 0.650468 | C12orf65 | NA | 0.091656991 | 0.649334642 | GMPR | NA | 0.209762 | 0.303725 |
| SNX7 | NA | 0.093151 | 0.650833 | PAXIP1 | NA | 0.091898719 | 0.648468382 | DLG5 | NA | 0.210476 | 0.302045 |
| RERE | NA | -0.0931 | 0.651012 | NAALADL1 | NA | 0.092002555 | 0.64809641 | ATXN7L3 | NA | 0.212563 | 0.297166 |
| SENP5 | NA | 0.092977 | 0.651445 | PPIL1 | NA | 0.0920799 | 0.647819392 | C1orf123 | NA | 0.215667 | 0.290001 |
| FLJ27354 | NA | -0.092848 | 0.651898 | STPG1 | NA | 0.092304432 | 0.647015468 | MAP3K10 | NA | 0.216681 | 0.287682 |
| SUV420H1 | NA | -0.092687 | 0.652463 | U2AF2 | NA | 0.092372332 | 0.64677243 | PGM5P2 | NA | 0.218584 | 0.283366 |
| LIPA | NA | -0.092671 | 0.652522 | DOLK | NA | 0.092683018 | 0.645660835 | EP400 | NA | 0.219465 | 0.281382 |
| DPH5 | NA | -0.092543 | 0.652972 | TJP1 | NA | 0.093188308 | 0.643854557 | CXCR5 | NA | 0.220363 | 0.279367 |
| LOC387647 | NA | 0.092435 | 0.65335 | RNF25 | NA | 0.093218925 | 0.64374517 | GPR21 | NA | 0.222559 | 0.274481 |
| ATP2B4 | NA | -0.092145 | 0.654373 | MTPAP | NA | 0.093360751 | 0.643238568 | MAGEF1 | NA | 0.223488 | 0.272433 |
| ZNF473 | NA | -0.091966 | 0.655003 | ZNF791 | NA | 0.093531119 | 0.642630219 | LPCAT4 | NA | 0.224809 | 0.269534 |
| NFE2L3 | NA | -0.090986 | 0.658458 | CHN2 | NA | 0.09359412 | 0.642405311 | HLA-DMB | NA | 0.233388 | 0.251193 |
| MFSD3 | NA | -0.090946 | 0.658601 | PIK3AP1 | NA | 0.093608918 | 0.642352487 | FIBP | NA | 0.238761 | 0.240133 |
| CASP8 | NA | -0.089919 | 0.66223 | CYP3A5 | NA | 0.093694559 | 0.642046818 | ZNF543 | NA | 0.238842 | 0.239968 |

**Table 6. Gene module membership and correlation with growth rate (48/72h), with adjusted p values for each module**

| | | | | | | | | | | | |
|---|---|---|---|---|---|---|---|---|---|---|---|
| EPDR1 | NA | 0.089881 | 0.662365 | SURF6 | NA | 0.093702672 | 0.642017865 | VILL | NA | 0.240064 | 0.2375 |
| DLG5 | NA | 0.089875 | 0.662386 | SART3 | NA | 0.093906422 | 0.641290882 | CCDC12 | NA | 0.240951 | 0.235717 |
| ADO | NA | 0.089225 | 0.664687 | TMEM50B | NA | 0.094453766 | 0.63933955 | MIR1307 | NA | 0.245864 | 0.22601 |
| ZNF768 | NA | -0.089176 | 0.664859 | ZNF543 | NA | 0.094916753 | 0.637690785 | STK40 | NA | 0.249212 | 0.219551 |
| BARD1 | NA | -0.089064 | 0.665259 | DDX31 | NA | 0.095059031 | 0.63718445 | PPTC7 | NA | 0.25095 | 0.216246 |
| DSEL | NA | 0.089044 | 0.66533 | POLR3A | NA | 0.095233824 | 0.636562618 | C19orf66 | NA | 0.252505 | 0.213319 |
| GSK3A | NA | 0.088958 | 0.665635 | AK4 | NA | 0.095447527 | 0.635802688 | NUDCD2 | NA | 0.260038 | 0.199516 |
| XRCC6 | NA | -0.08894 | 0.665696 | DSTYK | NA | 0.096304667 | 0.632758313 | SLIT3 | NA | 0.26512 | 0.190557 |
| ZNF347 | NA | 0.088716 | 0.666493 | ARHGEF18 | NA | 0.096444403 | 0.632262553 | LPAR6 | NA | 0.268388 | 0.184944 |
| ZNF552 | NA | -0.088626 | 0.66681 | FOLR3 | NA | 0.096679261 | 0.631429667 | HLA-B | NA | 0.270286 | 0.181737 |
| FAM193B | NA | 0.088429 | 0.667509 | SLC35E2 | NA | 0.096722464 | 0.631276502 | ISG15 | NA | 0.27521 | 0.173597 |
| DDX39A | NA | 0.088416 | 0.667556 | LOC1005056 | NA | 0.09689423 | 0.630667694 | HIST2H3D | NA | 0.277505 | 0.169894 |
| LPL | NA | -0.088092 | 0.668704 | MORC4 | NA | 0.097093834 | 0.629960519 | E4F1 | NA | 0.278085 | 0.168966 |
| AVEN | NA | -0.087941 | 0.66924 | DNAJB9 | NA | 0.097313246 | 0.629183528 | NBPF14 | NA | 0.280451 | 0.165219 |
| ERAP2 | NA | 0.087376 | 0.671248 | PIM1 | NA | 0.097648874 | 0.627995736 | NHLRC1 | NA | 0.280945 | 0.164444 |
| PTRHD1 | NA | -0.087277 | 0.671603 | S100A1 | NA | 0.097718727 | 0.62774864 | BTBD19 | NA | 0.286864 | 0.15536 |
| TCHP | NA | -0.086871 | 0.673045 | MORC3 | NA | 0.098458153 | 0.625135414 | HLA-DQA1 | NA | 0.291807 | 0.14805 |
| PTBP3 | NA | -0.086593 | 0.674035 | MARCO | NA | 0.099017525 | 0.623161451 | MEP1A | NA | 0.300861 | 0.135306 |
| FMR1 | NA | 0.086582 | 0.674075 | NUP37 | NA | 0.09924025 | 0.622376184 | CEACAM21 | NA | 0.303006 | 0.132407 |
| AMD1 | NA | -0.086394 | 0.674743 | MRPL37 | NA | 0.099375049 | 0.621901115 | NFE2L2 | NA | 0.324286 | 0.106042 |
| TIPIN | NA | 0.086163 | 0.675568 | NOL7 | NA | 0.099454294 | 0.621621905 | UVSSA | NA | 0.332897 | 0.096563 |
| SERPINF1 | NA | -0.085727 | 0.677123 | MUT | NA | 0.100109606 | 0.619314941 | MX1 | NA | 0.336517 | 0.092773 |
| TPRN | NA | 0.085526 | 0.677839 | DIP2A | NA | 0.100127241 | 0.619252906 | SUCLG2 | NA | 0.339764 | 0.089469 |
| PCMT1 | NA | -0.084962 | 0.679854 | ZSWIM7 | NA | 0.100423401 | 0.618211496 | HLA-DRB1 | NA | 0.351082 | 0.078645 |
| LINC00623 | NA | -0.084942 | 0.679924 | O3FAR1 | NA | 0.100476858 | 0.618023599 | OAS3 | NA | 0.360867 | 0.070116 |
| ZNF35 | NA | 0.084549 | 0.681329 | PLXNA2 | NA | 0.100751167 | 0.617059789 | TYW1B | NA | 0.365226 | 0.066554 |
| GPR183 | NA | -0.084527 | 0.681408 | SBF2-AS1 | NA | 0.101122143 | 0.615757315 | PARP9 | NA | 0.379867 | 0.055602 |
| TMEM161A | NA | -0.084384 | 0.681918 | RFC4 | NA | 0.101245508 | 0.615324438 | SP110 | NA | 0.382062 | 0.054089 |
| CERS5 | NA | -0.084362 | 0.681996 | SPRYD7 | NA | 0.101427623 | 0.614685643 | LRRC37A | NA | 0.407282 | 0.038912 |
| ZNF37A | NA | 0.083961 | 0.683432 | ANKS3 | NA | 0.101758349 | 0.613526278 | C2orf42 | orange | -0.324525 | 0.105771 |
| CNOT6L | NA | -0.083838 | 0.683874 | FLJ27354 | NA | 0.102111736 | 0.612288474 | SEC23B | orange | -0.321886 | 0.108804 |
| ZNRD1 | NA | -0.083644 | 0.684566 | CLEC6A | NA | 0.102434913 | 0.611157394 | EZH2 | orange | -0.314532 | 0.117595 |
| SNORD15A | NA | -0.083343 | 0.685644 | ABCB10 | NA | 0.102453347 | 0.611092903 | IMMT | orange | -0.297995 | 0.139251 |
| XIST | NA | -0.083176 | 0.686242 | ZFP14 | NA | 0.10254696 | 0.610765446 | C2orf69 | orange | -0.295549 | 0.142683 |
| ARL16 | NA | 0.083005 | 0.686856 | FCGR3A | NA | 0.103132866 | 0.608717617 | AFG3L2 | orange | -0.294615 | 0.14401 |
| AKAP12 | NA | -0.082485 | 0.688721 | NPEPPS | NA | 0.103176075 | 0.608566708 | FAM200A | orange | -0.292223 | 0.147448 |
| ZNF3 | NA | -0.082425 | 0.688937 | FAM192A | NA | 0.103188989 | 0.608521609 | BTG3 | orange | -0.277004 | 0.170697 |
| ITGA4 | NA | 0.082313 | 0.689337 | RIN3 | NA | 0.103565348 | 0.607207868 | LOC1001282 | orange | -0.264415 | 0.191782 |
| SNORA1 | NA | 0.082223 | 0.689662 | DAAM1 | NA | 0.103626198 | 0.606995576 | SLC30A5 | orange | -0.263126 | 0.194038 |
| BEST1 | NA | 0.081918 | 0.690757 | SNX8 | NA | 0.104409563 | 0.604265341 | PNO1 | orange | -0.248991 | 0.219973 |
| GPKOW | NA | -0.081827 | 0.691085 | LONP1 | NA | 0.104452296 | 0.604116554 | YEATS4 | orange | -0.248352 | 0.221198 |
| NCF1 | NA | -0.081643 | 0.691744 | POC1A | NA | 0.104514974 | 0.603898352 | S100A11 | orange | -0.246901 | 0.223995 |
| METTL3 | NA | 0.081286 | 0.693027 | GNPDA2 | NA | 0.104527833 | 0.603853587 | TFB2M | orange | -0.238371 | 0.240924 |
| MILR1 | NA | -0.080552 | 0.69567 | RPS4X | NA | 0.104566158 | 0.603720186 | SNX10 | orange | -0.23253 | 0.25299 |
| PIK3R3 | NA | -0.080438 | 0.696081 | ALOX5AP | NA | 0.105010457 | 0.602174579 | NOL11 | orange | -0.229932 | 0.258481 |
| HS6ST1 | NA | -0.08037 | 0.696325 | BCCIP | NA | 0.105029913 | 0.602106937 | MMADHC | orange | -0.223833 | 0.271673 |
| HMGN2 | NA | -0.080337 | 0.696446 | FAM70B | NA | 0.105376838 | 0.600901293 | IFIH1 | orange | -0.223387 | 0.272655 |
| KIF27 | NA | -0.079879 | 0.698096 | ZNF224 | NA | 0.105437005 | 0.600692303 | ATP6V1H | orange | -0.220259 | 0.279602 |
| SLC35E3 | NA | -0.079587 | 0.699147 | MCM9 | NA | 0.106361022 | 0.597486611 | ERLEC1 | orange | -0.217483 | 0.285859 |
| MALSU1 | NA | -0.079536 | 0.699331 | PMS2L2 | NA | 0.106718895 | 0.596247009 | PON2 | orange | -0.214738 | 0.292134 |
| RBM27 | NA | -0.079465 | 0.699586 | AP1S2 | NA | 0.106852035 | 0.595786116 | PDHA1 | orange | -0.213755 | 0.2944 |
| ULK1 | NA | -0.079444 | 0.699664 | VILL | NA | 0.106963912 | 0.59539895 | STRN3 | orange | -0.210942 | 0.300952 |
| TTC39B | NA | -0.079399 | 0.699827 | POFUT2 | NA | 0.107212077 | 0.594540523 | MORF4L2 | orange | -0.200711 | 0.325535 |
| ALKBH5 | NA | -0.078308 | 0.703766 | TNFAIP8 | NA | 0.108111238 | 0.591434698 | CAND1 | orange | -0.197905 | 0.332483 |
| TEX10 | NA | -0.078238 | 0.704019 | ZBTB45 | NA | 0.108209711 | 0.591094984 | PPP2R3C | orange | -0.191188 | 0.349482 |
| DEFB1 | NA | 0.077744 | 0.705807 | USP30-AS1 | NA | 0.108231234 | 0.591020744 | MRPS36 | orange | -0.185636 | 0.363913 |

**Table 6. Gene module membership and correlation with growth rate (48/72h), with adjusted p values for each module**

| | | | | | | | | | | | |
|---|---|---|---|---|---|---|---|---|---|---|---|
| RBM17 | NA | 0.077414 | 0.707 | KLHDC4 | NA | 0.108562519 | 0.589878551 | BRIX1 | orange | -0.178475 | 0.383032 |
| BAZ2B | NA | -0.077204 | 0.707763 | EDN1 | NA | 0.109374229 | 0.587084012 | VDAC2 | orange | -0.17411 | 0.394963 |
| OR2A7 | NA | 0.07681 | 0.70919 | RNPEPL1 | NA | 0.110238723 | 0.584114096 | KIAA1279 | orange | -0.172926 | 0.398235 |
| MRPL28 | NA | -0.076612 | 0.709908 | NT5E | NA | 0.11033722 | 0.583776131 | PCGF6 | orange | -0.168439 | 0.410772 |
| HIP1 | NA | -0.076363 | 0.710808 | FAHD2A | NA | 0.111140011 | 0.581024786 | P2RX7 | orange | -0.167237 | 0.414168 |
| P4HA1 | NA | 0.076161 | 0.711543 | LUC7L2 | NA | 0.111531145 | 0.579686347 | C14orf119 | orange | -0.166159 | 0.417228 |
| ACADVL | NA | -0.07505 | 0.715578 | BPHL | NA | 0.111586169 | 0.579498167 | NEK7 | orange | -0.161617 | 0.430248 |
| CDK2 | NA | -0.075016 | 0.715703 | MPV17L | NA | 0.112499885 | 0.576377233 | PELO | orange | -0.159545 | 0.436262 |
| MAF | NA | -0.074136 | 0.718904 | BTBD9 | NA | 0.112672157 | 0.575789646 | PRPF18 | orange | -0.154839 | 0.450087 |
| ZNF527 | NA | -0.07377 | 0.720238 | DHDDS | NA | 0.112878694 | 0.57508554 | TBC1D22B | orange | -0.151208 | 0.460911 |
| EIF1AY | NA | 0.073555 | 0.721018 | INTS12 | NA | 0.113235266 | 0.573870845 | TAX1BP1 | orange | -0.148902 | 0.467855 |
| ZNF471 | NA | -0.073384 | 0.721643 | HMGCR | NA | 0.113370596 | 0.573410133 | SLMO2 | orange | -0.140174 | 0.494618 |
| HPGDS | NA | -0.073352 | 0.72176 | SLC25A28 | NA | 0.113450181 | 0.57313927 | UBE2B | orange | -0.109027 | 0.595992 |
| CIDEB | NA | -0.072461 | 0.725008 | EMC10 | NA | 0.113594676 | 0.572647642 | ANKRD37 | orange | -0.106756 | 0.603715 |
| PRDX5 | NA | -0.071937 | 0.726922 | TREML4 | NA | 0.114427951 | 0.569816178 | PUSL1 | orange | -0.091374 | 0.65709 |
| KDM5D | NA | 0.071664 | 0.72792 | ZNF473 | NA | 0.114573546 | 0.569322093 | CTDSPL2 | orange | -0.082272 | 0.689486 |
| OPLAH | NA | -0.071384 | 0.728943 | CD274 | NA | 0.114627514 | 0.569138995 | TMEM167B | orange | -0.074035 | 0.71927 |
| PLTP | NA | 0.071013 | 0.7303 | SCPEP1 | NA | 0.115249819 | 0.567029634 | TAGAP | orange | -0.053428 | 0.795467 |
| RALGAPA2 | NA | 0.070392 | 0.732574 | RGL2 | NA | 0.115421385 | 0.566448711 | SIGIRR | orange | -0.048977 | 0.812196 |
| KCNA3 | NA | 0.070275 | 0.733003 | TOPORS | NA | 0.115704933 | 0.565489209 | DHX33 | orange | -0.00974 | 0.962336 |
| RPS3 | NA | -0.069999 | 0.734016 | AMD1 | NA | 0.116250344 | 0.563645648 | KLHDC5 | orange | -0.006052 | 0.976592 |
| HEMK1 | NA | -0.069965 | 0.734141 | ASL | NA | 0.116361098 | 0.563271616 | TAOK3 | orange | 0.011392 | 0.955954 |
| BRK1 | NA | -0.069567 | 0.7356 | ABHD5 | NA | 0.116476996 | 0.562880334 | CFD | orange | 0.015782 | 0.939004 |
| ZNF407 | NA | -0.069417 | 0.73615 | PLS3 | NA | 0.116818874 | 0.561726839 | ZNF43 | orange | 0.033985 | 0.86909 |
| SNX29P2 | NA | -0.06931 | 0.73654 | RCBTB1 | NA | 0.11801749 | 0.557691212 | FKBP2 | orange | 0.036867 | 0.858093 |
| MB21D2 | NA | -0.069246 | 0.736776 | FBXO6 | NA | 0.118049062 | 0.557585093 | ZNF273 | orange | 0.054297 | 0.792212 |
| PER1 | NA | -0.068786 | 0.738464 | UVSSA | NA | 0.118343313 | 0.55659649 | HDAC6 | orange | 0.086259 | 0.675224 |
| THAP4 | NA | -0.068593 | 0.739173 | TNFSF10 | NA | 0.11843861 | 0.556276493 | KCTD5 | orange | 0.090986 | 0.658458 |
| TENM4 | NA | -0.068592 | 0.739175 | ZNF490 | NA | 0.118441599 | 0.556266457 | SIGLEC9 | orange | 0.101124 | 0.623048 |
| ETFDH | NA | -0.068575 | 0.739241 | ADK | NA | 0.118616762 | 0.555678499 | DSC2 | orange | 0.108136 | 0.599016 |
| SEC24B | NA | 0.068565 | 0.739275 | FAS | NA | 0.118710168 | 0.555365085 | NIPA1 | orange | 0.108848 | 0.596599 |
| CYB5D1 | NA | 0.068519 | 0.739444 | USP18 | NA | 0.118770832 | 0.555161578 | ADCY9 | orange | 0.124273 | 0.545275 |
| PRDM1 | NA | -0.068437 | 0.739748 | PLEKHA2 | NA | 0.119135067 | 0.55394041 | FAN1 | orange | 0.126072 | 0.539424 |
| MOSPD3 | NA | -0.068097 | 0.740997 | SLC25A35 | NA | 0.119148501 | 0.553895392 | RPL32P3 | orange | 0.139634 | 0.4963 |
| UBL3 | NA | -0.067926 | 0.741626 | CYSTM1 | NA | 0.119321553 | 0.553315656 | GPI | orange | 0.144779 | 0.480401 |
| NFX1 | NA | -0.067873 | 0.74182 | KIAA0196 | NA | 0.119400566 | 0.553051051 | PLOD1 | orange | 0.153204 | 0.454944 |
| SLC29A1 | NA | 0.067002 | 0.745023 | COX16 | NA | 0.119656087 | 0.552195739 | RPS6KA4 | orange | 0.161725 | 0.429936 |
| MORC3 | NA | -0.066494 | 0.746893 | VASH1 | NA | 0.119842891 | 0.55157083 | TMCO7 | orange | 0.167968 | 0.412101 |
| APOC2 | NA | -0.066413 | 0.747194 | SLC39A1 | NA | 0.120173148 | 0.55046683 | ANKRD36B | orange | 0.189683 | 0.353359 |
| CWC25 | NA | -0.066306 | 0.747587 | RPL36A | NA | 0.120209747 | 0.550344547 | STAT6 | orange | 0.265468 | 0.189952 |
| RPS4Y1 | NA | 0.066183 | 0.748042 | ZNF284 | NA | 0.120419551 | 0.549643807 | ATG4B | orange | 0.282562 | 0.161926 |
| CCT5 | NA | 0.065978 | 0.748795 | ZNF583 | NA | 0.120425206 | 0.549624925 | GTDC2 | orange | 0.285503 | 0.157416 |
| OSM | NA | -0.065457 | 0.75072 | SOD2 | NA | 0.120763544 | 0.548495773 | HIC2 | orange | 0.288926 | 0.15228 |
| ZNF333 | NA | -0.065211 | 0.751627 | IKBKE | NA | 0.121074209 | 0.547459925 | MRI1 | orange | 0.329159 | 0.100596 |
| UPK3BL | NA | 0.065188 | 0.75171 | MRPS6 | NA | 0.121144434 | 0.5472259 | TRAPPC10 | orange | 0.336217 | 0.093082 |
| POLA2 | NA | -0.065121 | 0.751958 | TSEN54 | NA | 0.1211826 | 0.54709873 | C15orf52 | orange | 0.358035 | 0.072508 |
| HPS5 | NA | -0.064975 | 0.752498 | RABAC1 | NA | 0.121703992 | 0.545362831 | PI4KAP2 | orange | 0.358718 | 0.071926 |
| PTGER3 | NA | 0.064676 | 0.753602 | MYO1E | NA | 0.121866602 | 0.544821971 | COA1 | orange | 0.379977 | 0.055525 |
| RPS4X | NA | -0.064662 | 0.753652 | FKBP1A | NA | 0.122046946 | 0.544222417 | ADRBK1 | orange | 0.418157 | 0.033517 |
| EML2 | NA | 0.064546 | 0.754083 | PDE7A | NA | 0.122262067 | 0.543507649 | DNM1L | orange | 0.435407 | 0.026201 |
| SMN1 | NA | 0.064245 | 0.755194 | CYP2S1 | NA | 0.122745783 | 0.541902044 | COMMD9 | paleturquoise | -0.481303 | 0.012798 |
| PLEKHA2 | NA | -0.064015 | 0.756043 | PFDN2 | NA | 0.122756597 | 0.541866177 | NOM1 | paleturquoise | -0.415236 | 0.034904 |
| MLL4 | NA | -0.063838 | 0.756699 | AP1B1 | NA | 0.122757513 | 0.541863138 | ANXA2 | paleturquoise | -0.294154 | 0.144667 |
| RNF103 | NA | 0.063797 | 0.756849 | PRDX4 | NA | 0.123207078 | 0.540372936 | MRPL48 | paleturquoise | -0.274438 | 0.174857 |
| GABPB2 | NA | 0.063727 | 0.757111 | ULK2 | NA | 0.123229446 | 0.540298841 | NAPSB | paleturquoise | -0.241324 | 0.234971 |
| PUS1 | NA | 0.063502 | 0.75794 | MED21 | NA | 0.123449581 | 0.539569895 | NAT10 | paleturquoise | -0.239791 | 0.238049 |

**Table 6. Gene module membership and correlation with growth rate (48/72h), with adjusted p values for each module**

| | | | | | | | | | | | |
|---|---|---|---|---|---|---|---|---|---|---|---|
| PLXND1 | NA | -0.063454 | 0.758121 | CC2D1A | NA | 0.123606099 | 0.539051886 | LAIR1 | paleturquoise | -0.217564 | 0.285675 |
| REST | NA | 0.063107 | 0.759403 | LAMP3 | NA | 0.124336166 | 0.536638778 | SLA | paleturquoise | -0.209645 | 0.304002 |
| METTL9 | NA | 0.062774 | 0.760636 | C11orf54 | NA | 0.124716004 | 0.53538531 | NUDT14 | paleturquoise | -0.204004 | 0.317493 |
| CCAR1 | NA | -0.062505 | 0.761635 | SIL1 | NA | 0.125027744 | 0.534357603 | RNF141 | paleturquoise | -0.195236 | 0.339178 |
| INPP5K | NA | 0.062275 | 0.762485 | NUB1 | NA | 0.125154756 | 0.533939152 | MSRB1 | paleturquoise | -0.180715 | 0.376989 |
| TTYH2 | NA | -0.062156 | 0.762926 | ZFYVE19 | NA | 0.125512232 | 0.532762256 | APMAP | paleturquoise | -0.166501 | 0.416254 |
| IGF1 | NA | 0.062019 | 0.763434 | CCDC69 | NA | 0.125684779 | 0.532194632 | PGAM5 | paleturquoise | -0.161151 | 0.431599 |
| TP53RK | NA | -0.061943 | 0.763718 | TATDN2 | NA | 0.126694136 | 0.528879937 | BDH1 | paleturquoise | -0.159531 | 0.436305 |
| USP9Y | NA | 0.061931 | 0.76376 | IFIT2 | NA | 0.126757643 | 0.528671713 | TNFSF12 | paleturquoise | -0.147333 | 0.47261 |
| IFT74 | NA | -0.061868 | 0.763995 | EHD1 | NA | 0.127227824 | 0.527131321 | CEMP1 | paleturquoise | -0.131564 | 0.521752 |
| RPS6KA2 | NA | 0.061521 | 0.765281 | SIGLEC14 | NA | 0.127262369 | 0.52701823 | NMB | paleturquoise | -0.115456 | 0.574361 |
| COQ6 | NA | -0.061383 | 0.765794 | BNIP1 | NA | 0.12739837 | 0.526573114 | DNAJC18 | paleturquoise | -0.084885 | 0.680127 |
| ZNF524 | NA | -0.061326 | 0.766007 | CD163 | NA | 0.127836126 | 0.525141615 | TMEM102 | paleturquoise | -0.080092 | 0.697327 |
| SRPRB | NA | -0.060982 | 0.767281 | HOMER3 | NA | 0.127886436 | 0.524977217 | CAMK1D | paleturquoise | -0.080001 | 0.697656 |
| ZMYM4 | NA | 0.060775 | 0.768051 | SRR | NA | 0.128116158 | 0.524226868 | VEGFB | paleturquoise | -0.07957 | 0.69921 |
| FBXL4 | NA | -0.060757 | 0.768118 | FRG1B | NA | 0.128327905 | 0.52353569 | IL7R | paleturquoise | -0.079151 | 0.700722 |
| MAN1A2 | NA | 0.060201 | 0.770185 | CCDC159 | NA | 0.128489093 | 0.523009835 | ZBTB38 | paleturquoise | -0.078662 | 0.702486 |
| QSER1 | NA | 0.060047 | 0.770755 | OAS3 | NA | 0.128955019 | 0.521491257 | SH2B2 | paleturquoise | -0.069948 | 0.734202 |
| TJP1 | NA | 0.05957 | 0.77253 | TAB3 | NA | 0.129187119 | 0.520735572 | GOLIM4 | paleturquoise | -0.049937 | 0.80858 |
| YPEL1 | NA | -0.058756 | 0.775561 | SIAH2 | NA | 0.129616987 | 0.519337387 | GZMB | paleturquoise | -0.049906 | 0.808696 |
| SH3PXD2B | NA | 0.058705 | 0.77575 | CAMP | NA | 0.129626244 | 0.519307297 | LAS1L | paleturquoise | -0.044923 | 0.827501 |
| CCDC111 | NA | -0.058439 | 0.776742 | NUP35 | NA | 0.12974519 | 0.518920749 | TRPV4 | paleturquoise | -0.040522 | 0.844188 |
| INO80C | NA | -0.058314 | 0.777208 | RUFY3 | NA | 0.129984008 | 0.518145057 | ZNRD1 | paleturquoise | -0.022227 | 0.914174 |
| SLC25A37 | NA | -0.058302 | 0.777251 | LARP4 | NA | 0.13034091 | 0.51698688 | FTSJ2 | paleturquoise | -0.013994 | 0.945904 |
| NUDCD3 | NA | -0.058216 | 0.777573 | TUBGCP5 | NA | 0.130464097 | 0.516587419 | LOC1005072[illegible] | paleturquoise | -0.009698 | 0.962498 |
| DYNC2H1 | NA | -0.058024 | 0.778287 | MED20 | NA | 0.130639782 | 0.516017979 | AMIGO2 | paleturquoise | -0.001588 | 0.993857 |
| ZC3HC1 | NA | -0.057925 | 0.778656 | NME6 | NA | 0.131454928 | 0.513379902 | CSK | paleturquoise | 0.001355 | 0.994758 |
| HLA-DQB2 | NA | 0.057754 | 0.779294 | TRA2A | NA | 0.131739766 | 0.512459629 | RGS19 | paleturquoise | 0.003325 | 0.98714 |
| FRG1B | NA | 0.057549 | 0.78006 | TRMU | NA | 0.131861198 | 0.512067544 | TOM1L1 | paleturquoise | 0.003765 | 0.985438 |
| ZFY | NA | 0.05728 | 0.781062 | ACACA | NA | 0.131951812 | 0.511775063 | KLF4 | paleturquoise | 0.005437 | 0.97897 |
| ING5 | NA | -0.056791 | 0.782889 | ZNF512 | NA | 0.132047937 | 0.51146488 | ELOVL7 | paleturquoise | 0.016048 | 0.937981 |
| LINC00263 | NA | 0.056721 | 0.783147 | FANCD2 | NA | 0.132232856 | 0.510868434 | KIAA0196 | paleturquoise | 0.018688 | 0.927802 |
| ACSF2 | NA | -0.05656 | 0.783748 | GRAMD1B | NA | 0.132462596 | 0.510127893 | BOD1 | paleturquoise | 0.019548 | 0.924487 |
| SORL1 | NA | -0.055877 | 0.786302 | UBE2Q1 | NA | 0.132817711 | 0.508984259 | PSTPIP2 | paleturquoise | 0.029185 | 0.887456 |
| TOPBP1 | NA | 0.055809 | 0.786557 | TMEM223 | NA | 0.132881962 | 0.508777479 | AQP9 | paleturquoise | 0.03597 | 0.861514 |
| CCDC12 | NA | -0.055587 | 0.787387 | SLC25A39 | NA | 0.132929308 | 0.508625126 | LAMTOR1 | paleturquoise | 0.045471 | 0.825428 |
| SARM1 | NA | -0.055295 | 0.788476 | CLYBL | NA | 0.133321872 | 0.507362799 | RIPK2 | paleturquoise | 0.048065 | 0.815634 |
| TOR1AIP2 | NA | -0.055238 | 0.78869 | NOL6 | NA | 0.133514596 | 0.50674364 | CD44 | paleturquoise | 0.071477 | 0.728605 |
| STON1 | NA | -0.055045 | 0.789413 | GIPC1 | NA | 0.134822269 | 0.502552395 | N4BP2L1 | paleturquoise | 0.075511 | 0.713904 |
| NR1D1 | NA | 0.055032 | 0.789462 | SLC46A2 | NA | 0.134921775 | 0.502234172 | TXLNA | paleturquoise | 0.086273 | 0.675175 |
| ACTR2 | NA | -0.05481 | 0.79029 | CBWD6 | NA | 0.136443326 | 0.497380708 | TRANK1 | paleturquoise | 0.090207 | 0.66121 |
| RNF6 | NA | 0.054215 | 0.792519 | ACSF2 | NA | 0.136537981 | 0.497079553 | EEFSEC | paleturquoise | 0.09472 | 0.645327 |
| ANO8 | NA | -0.054121 | 0.79287 | GVINP1 | NA | 0.13662843 | 0.496791865 | GBA2 | paleturquoise | 0.113651 | 0.580398 |
| DIEXF | NA | -0.053318 | 0.795881 | GEMIN6 | NA | 0.136734809 | 0.496453615 | CLEC2D | paleturquoise | 0.166666 | 0.415786 |
| RCBTB2 | NA | -0.053203 | 0.796313 | HSPBAP1 | NA | 0.136991807 | 0.495636924 | ASH1L-AS1 | paleturquoise | 0.209563 | 0.304195 |
| C5orf24 | NA | 0.053107 | 0.79667 | UPF3B | NA | 0.137604794 | 0.493691686 | LOC374443 | paleturquoise | 0.286446 | 0.15599 |
| GGCX | NA | 0.052826 | 0.797726 | ECM1 | NA | 0.138221435 | 0.491738731 | MSC | pink | -0.350028 | 0.079609 |
| ABI3 | NA | 0.052516 | 0.798888 | PTK2 | NA | 0.138422815 | 0.49110179 | HADH | pink | -0.328921 | 0.100857 |
| FPR1 | NA | -0.052445 | 0.799154 | TECPR2 | NA | 0.139057046 | 0.489098506 | CCND2 | pink | -0.32656 | 0.103474 |
| CTSH | NA | 0.051939 | 0.801055 | ZNF132 | NA | 0.139225116 | 0.48856833 | LOC730102 | pink | -0.295228 | 0.143138 |
| PROCA1 | NA | 0.051808 | 0.801548 | ZNF461 | NA | 0.139391549 | 0.488043609 | SEMA4B | pink | -0.272559 | 0.177947 |
| CC2D1A | NA | -0.051705 | 0.801932 | PCSK7 | NA | 0.139466095 | 0.487808675 | LILRA5 | pink | -0.264623 | 0.19142 |
| CDK19 | NA | -0.051699 | 0.801955 | SP110 | NA | 0.139696249 | 0.487083703 | SARDH | pink | -0.247983 | 0.221906 |
| DTNA | NA | 0.051293 | 0.803482 | DZIP3 | NA | 0.139813925 | 0.486713244 | AIF1 | pink | -0.244153 | 0.22936 |
| CTSW | NA | -0.051188 | 0.803875 | IREB2 | NA | 0.139908978 | 0.486414107 | SERPINE1 | pink | -0.234487 | 0.248904 |
| DRAM1 | NA | 0.051104 | 0.804191 | NCF1 | NA | 0.141074117 | 0.482754948 | HN1L | pink | -0.23199 | 0.254125 |

**Table 6. Gene module membership and correlation with growth rate (48/72h), with adjusted p values for each module**

| | | | | | | | | | | | |
|---|---|---|---|---|---|---|---|---|---|---|---|
| ZNF398 | NA | 0.051057 | 0.80437 | SELRC1 | NA | 0.141192401 | 0.482384259 | FBXL6 | pink | -0.228618 | 0.261286 |
| PKIA | NA | -0.050975 | 0.804677 | RCOR3 | NA | 0.141310324 | 0.482014847 | TUBA4A | pink | -0.224085 | 0.271121 |
| EIF1 | NA | 0.050544 | 0.806298 | POLR1B | NA | 0.141358418 | 0.481864224 | RNASE6 | pink | -0.210799 | 0.301286 |
| ELOF1 | NA | -0.05016 | 0.807741 | COX15 | NA | 0.141423611 | 0.481660092 | ENTPD7 | pink | -0.201971 | 0.322443 |
| GOLGA5 | NA | -0.050019 | 0.80827 | NOA1 | NA | 0.142149079 | 0.479391484 | C2orf49 | pink | -0.19687 | 0.335069 |
| PFDN2 | NA | 0.049944 | 0.808552 | SETD6 | NA | 0.142430674 | 0.478512386 | DHCR7 | pink | -0.187251 | 0.35968 |
| DOLPP1 | NA | 0.049805 | 0.809076 | NUMB | NA | 0.143078294 | 0.476493754 | MSRB2 | pink | -0.185807 | 0.363462 |
| MEX3C | NA | -0.049769 | 0.809214 | ZNF766 | NA | 0.143412822 | 0.47545275 | C18orf8 | pink | -0.185346 | 0.364676 |
| HNRNPD | NA | 0.049517 | 0.810161 | SH2D4A | NA | 0.143437016 | 0.475377504 | NDRG2 | pink | -0.182299 | 0.372752 |
| TMED2 | NA | 0.049117 | 0.811669 | IDUA | NA | 0.144430636 | 0.472292662 | GALM | pink | -0.170821 | 0.40409 |
| ARFIP1 | NA | -0.049109 | 0.811697 | TUBA1C | NA | 0.144966164 | 0.470634339 | ZNRF1 | pink | -0.170019 | 0.406333 |
| CDADC1 | NA | 0.048598 | 0.813624 | BRK1 | NA | 0.145102847 | 0.470211565 | LOC645638 | pink | -0.169345 | 0.408224 |
| ZBTB38 | NA | -0.04859 | 0.813652 | SYTL3 | NA | 0.145779903 | 0.468120288 | EVL | pink | -0.169096 | 0.408924 |
| RTN3 | NA | -0.048254 | 0.814919 | ANKFY1 | NA | 0.146761642 | 0.465096516 | PTPN7 | pink | -0.165799 | 0.418251 |
| PPP2R2D | NA | -0.048211 | 0.815083 | NAA25 | NA | 0.146920324 | 0.464608733 | TLR4 | pink | -0.16571 | 0.418504 |
| NPPC | NA | 0.048132 | 0.815381 | ECSCR | NA | 0.147089885 | 0.464087803 | PEX3 | pink | -0.165368 | 0.419479 |
| MARCO | NA | -0.047738 | 0.816865 | SRBD1 | NA | 0.14728549 | 0.463487237 | TRHDE | pink | -0.161012 | 0.432 |
| KCTD2 | NA | 0.047612 | 0.817343 | NT5C3 | NA | 0.147406093 | 0.463117154 | ECD | pink | -0.15988 | 0.435287 |
| BTG2 | NA | 0.04751 | 0.817727 | PRELID1 | NA | 0.148195742 | 0.460697851 | MEGF9 | pink | -0.159072 | 0.437642 |
| SWI5 | NA | -0.047358 | 0.818301 | PPIL3 | NA | 0.14861344 | 0.459420809 | CYP27B1 | pink | -0.158725 | 0.438653 |
| MUM1 | NA | -0.047298 | 0.818527 | SNAP47 | NA | 0.148772332 | 0.458935512 | FOSL1 | pink | -0.151693 | 0.459458 |
| GRHL1 | NA | -0.04709 | 0.819313 | SLC5A3 | NA | 0.149023319 | 0.458169483 | GTF2A2 | pink | -0.15049 | 0.463065 |
| ATL3 | NA | -0.046963 | 0.819793 | MAOA | NA | 0.149111857 | 0.457899421 | RGS1 | pink | -0.148507 | 0.46905 |
| IRS2 | NA | -0.04677 | 0.82052 | GLA | NA | 0.150327608 | 0.454199568 | BIRC3 | pink | -0.147564 | 0.471908 |
| JMJD4 | NA | -0.04666 | 0.820934 | C8B | NA | 0.150570431 | 0.453462492 | APOC1 | pink | -0.145653 | 0.477728 |
| SMYD5 | NA | -0.046629 | 0.821053 | TSPYL4 | NA | 0.150789292 | 0.452798696 | HSD17B11 | pink | -0.141677 | 0.489956 |
| CLEC6A | NA | -0.046235 | 0.822539 | SPIRE2 | NA | 0.150790673 | 0.452794509 | NES | pink | -0.140107 | 0.494827 |
| THAP1 | NA | 0.046161 | 0.822821 | PTGES | NA | 0.151064384 | 0.451965085 | MS4A4A | pink | -0.137825 | 0.50195 |
| KIAA1147 | NA | -0.046077 | 0.823137 | TIPIN | NA | 0.151341081 | 0.451127432 | SUSD1 | pink | -0.135507 | 0.509238 |
| ODF2L | NA | 0.046048 | 0.823248 | DTNA | NA | 0.151350363 | 0.451099345 | CHI3L1 | pink | -0.134125 | 0.513605 |
| SH3KBP1 | NA | -0.045681 | 0.824633 | PELI2 | NA | 0.151479173 | 0.450709689 | FAM165B | pink | -0.132139 | 0.519918 |
| ZNF8 | NA | 0.045586 | 0.824995 | LYZ | NA | 0.152652228 | 0.447169389 | DCP1B | pink | -0.12962 | 0.527973 |
| KAT5 | NA | -0.045526 | 0.825221 | AK8 | NA | 0.153150424 | 0.445670331 | BAALC | pink | -0.128906 | 0.53027 |
| FCN1 | NA | -0.045455 | 0.825489 | CPNE1 | NA | 0.153186083 | 0.445563136 | CCDC115 | pink | -0.12875 | 0.530772 |
| HDAC6 | NA | 0.04537 | 0.825812 | PLOD1 | NA | 0.153344332 | 0.445087593 | ADAM8 | pink | -0.128224 | 0.532464 |
| F3 | NA | -0.045274 | 0.826174 | HP | NA | 0.153967447 | 0.443217755 | PCED1B-AS1 | pink | -0.127314 | 0.535403 |
| MPG | NA | -0.045214 | 0.826402 | IFITM1 | NA | 0.153994855 | 0.443135606 | PVALB | pink | -0.127057 | 0.536233 |
| FHAD1 | NA | 0.04516 | 0.826607 | CCL4 | NA | 0.154157242 | 0.442649058 | OAT | pink | -0.125582 | 0.541015 |
| METTL1 | NA | -0.045105 | 0.826812 | PI3 | NA | 0.154308972 | 0.4421947 | DYRK4 | pink | -0.123654 | 0.547292 |
| POLR1B | NA | -0.044902 | 0.827582 | CIZ1 | NA | 0.154588948 | 0.441356964 | CASS4 | pink | -0.123275 | 0.54853 |
| MED7 | NA | -0.044878 | 0.827672 | PRKCA | NA | 0.154677202 | 0.441093072 | PARP1 | pink | -0.122945 | 0.54961 |
| RARRES1 | NA | 0.044637 | 0.828582 | CAST | NA | 0.154677399 | 0.441092483 | OTUD7B | pink | -0.12249 | 0.551099 |
| LOC646329 | NA | -0.044549 | 0.828918 | RPS12 | NA | 0.154878475 | 0.440491551 | E2F1 | pink | -0.122148 | 0.552221 |
| FANCC | NA | -0.044422 | 0.829398 | WDR81 | NA | 0.155075086 | 0.439904388 | ZNF552 | pink | -0.120143 | 0.558812 |
| TRAM2 | NA | -0.044227 | 0.830136 | RAI1 | NA | 0.155196561 | 0.439541825 | COL6A2 | pink | -0.119705 | 0.560257 |
| TXNDC9 | NA | -0.043855 | 0.831545 | TMEM187 | NA | 0.155562036 | 0.43845197 | THAP4 | pink | -0.112079 | 0.585679 |
| RGAG4 | NA | -0.043761 | 0.831899 | CKAP5 | NA | 0.155620558 | 0.438277591 | GCHFR | pink | -0.111392 | 0.587994 |
| RSBN1L-AS1 | NA | -0.043635 | 0.832376 | MEP1A | NA | 0.155635647 | 0.438232636 | TIRAP | pink | -0.110461 | 0.591135 |
| LRRC37A | NA | 0.0435 | 0.83289 | FOXN3-AS1 | NA | 0.156205678 | 0.436536167 | IGFBP6 | pink | -0.108421 | 0.598049 |
| EIF3D | NA | 0.042623 | 0.836211 | MRPS18A | NA | 0.156427363 | 0.435877369 | PMAIP1 | pink | -0.106726 | 0.603817 |
| MED21 | NA | -0.042112 | 0.838152 | G0S2 | NA | 0.156533876 | 0.435561028 | CARD16 | pink | -0.105722 | 0.607246 |
| EIF3L | NA | 0.041892 | 0.838987 | ZNF77 | NA | 0.156604869 | 0.43535025 | SELRC1 | pink | -0.105321 | 0.608615 |
| ADA | NA | 0.041633 | 0.839969 | RABGGTA | NA | 0.156644106 | 0.435233779 | LIF | pink | -0.10509 | 0.609408 |
| CCDC69 | NA | -0.041481 | 0.840545 | SETD2 | NA | 0.157365469 | 0.433095496 | TM4SF1 | pink | -0.105065 | 0.609493 |
| FKBP15 | NA | -0.041397 | 0.840863 | KRR1 | NA | 0.157660044 | 0.432223951 | TGFBI | pink | -0.101887 | 0.620412 |
| SPARC | NA | 0.041397 | 0.840864 | PGPEP1 | NA | 0.158345073 | 0.430200883 | TNIP3 | pink | -0.099216 | 0.629651 |

**Table 6. Gene module membership and correlation with growth rate (48/72h), with adjusted p values for each module**

| | | | | | | | | | | | |
|---|---|---|---|---|---|---|---|---|---|---|---|
| CDC42EP3 | NA | 0.041378 | 0.840936 | LSM1 | NA | 0.158705173 | 0.429139487 | GPAM | pink | -0.097438 | 0.635833 |
| ALKBH3 | NA | 0.040847 | 0.842953 | GPC4 | NA | 0.158847863 | 0.428719304 | SRGN | pink | -0.092682 | 0.652481 |
| ZNF324B | NA | -0.040783 | 0.843197 | ZBTB48 | NA | 0.159481225 | 0.426856936 | CYP4V2 | pink | -0.09167 | 0.656045 |
| CD72 | NA | -0.040561 | 0.844038 | TAF6L | NA | 0.160012165 | 0.425299152 | LILRB4 | pink | -0.091547 | 0.656479 |
| USP35 | NA | 0.040487 | 0.844319 | FLOT1 | NA | 0.160230688 | 0.42465891 | STK10 | pink | -0.087737 | 0.669967 |
| PIAS4 | NA | -0.040285 | 0.845089 | CAPN10 | NA | 0.160283043 | 0.424505596 | NPR1 | pink | -0.087646 | 0.670291 |
| TTTY15 | NA | 0.040017 | 0.846104 | HDGF | NA | 0.160321413 | 0.424393252 | DOCK2 | pink | -0.085671 | 0.677322 |
| LOC645513 | NA | -0.039957 | 0.846336 | SF3B4 | NA | 0.16055988 | 0.42369542 | CLU | pink | -0.084061 | 0.683072 |
| GLB1L | NA | 0.039819 | 0.846858 | MSRB2 | NA | 0.160661597 | 0.423397954 | LGR4 | pink | -0.082849 | 0.687416 |
| SLC2A1 | NA | -0.039353 | 0.848631 | KCNA3 | NA | 0.160860437 | 0.422816788 | SMAP1 | pink | -0.081442 | 0.692468 |
| FAM220A | NA | 0.039117 | 0.849527 | FBXO31 | NA | 0.161314462 | 0.421491421 | CCL3 | pink | -0.08092 | 0.694347 |
| IGSF10 | NA | -0.03902 | 0.849897 | CES2 | NA | 0.161534782 | 0.420849098 | BCL6 | pink | -0.08075 | 0.694958 |
| TGFB1 | NA | -0.038851 | 0.850538 | ILKAP | NA | 0.161563346 | 0.420765862 | ERV3-1 | pink | -0.079647 | 0.69893 |
| PIGG | NA | -0.038526 | 0.851775 | STX11 | NA | 0.16181414 | 0.420035433 | MAFB | pink | -0.078603 | 0.7027 |
| AHCTF1 | NA | -0.038517 | 0.85181 | SLC7A1 | NA | 0.161885499 | 0.419827728 | ZNF449 | pink | -0.078479 | 0.703149 |
| SMPD4 | NA | 0.038311 | 0.852594 | IFI6 | NA | 0.162218228 | 0.418860009 | PGAM1 | pink | -0.078379 | 0.703511 |
| EBLN2 | NA | -0.038097 | 0.853409 | PREB | NA | 0.16221824 | 0.418859974 | ZAK | pink | -0.077904 | 0.705228 |
| GFPT2 | NA | 0.038069 | 0.853517 | BAMBI | NA | 0.162428228 | 0.418249872 | PCED1B | pink | -0.076206 | 0.711379 |
| PDIA3 | NA | -0.037442 | 0.855905 | EIF2C1 | NA | 0.162446268 | 0.418197484 | BRPF3 | pink | -0.070422 | 0.732466 |
| BMS1P1 | NA | -0.037349 | 0.856257 | MIR1307 | NA | 0.16294286 | 0.416756735 | SPIN1 | pink | -0.069054 | 0.737482 |
| USP20 | NA | -0.037327 | 0.856341 | ELAC1 | NA | 0.163473813 | 0.415219343 | CLEC7A | pink | -0.068134 | 0.74086 |
| UBA5 | NA | -0.0373 | 0.856445 | STAC | NA | 0.164560754 | 0.412081896 | PDGFA | pink | -0.06509 | 0.752075 |
| MAP4K1 | NA | 0.037087 | 0.857258 | ATF2 | NA | 0.164888683 | 0.411137928 | SCCPDH | pink | -0.060082 | 0.770626 |
| NFKBIZ | NA | -0.036515 | 0.859435 | ARL8A | NA | 0.164893927 | 0.411122841 | PNPLA3 | pink | -0.059354 | 0.773336 |
| BNIP1 | NA | -0.036402 | 0.859865 | CEP97 | NA | 0.165003201 | 0.410808563 | SLC39A14 | pink | -0.056845 | 0.782686 |
| CD14 | NA | 0.036295 | 0.860275 | VAV3 | NA | 0.1655243 | 0.409311693 | CCL22 | pink | -0.056613 | 0.78355 |
| FN3KRP | NA | -0.036279 | 0.860335 | IRF7 | NA | 0.165896073 | 0.408245632 | APOL3 | pink | -0.055724 | 0.786871 |
| TACC1 | NA | 0.036203 | 0.860626 | ETV5 | NA | 0.16591123 | 0.408202203 | LAT | pink | -0.05475 | 0.790517 |
| PSMB1 | NA | 0.036092 | 0.861048 | ADA | NA | 0.166261317 | 0.407199806 | KIAA1462 | pink | -0.053484 | 0.795258 |
| KIF16B | NA | 0.035776 | 0.862256 | TRIM39 | NA | 0.166818312 | 0.405607815 | ITGAM | pink | -0.052779 | 0.797903 |
| G0S2 | NA | -0.035732 | 0.862421 | SCAP | NA | 0.166820893 | 0.405600446 | ASAP2 | pink | -0.051943 | 0.801039 |
| SLC25A6 | NA | 0.035692 | 0.862573 | CACTIN | NA | 0.166944396 | 0.40524793 | GRASP | pink | -0.049999 | 0.808346 |
| BATF3 | NA | -0.035186 | 0.864504 | CTNNBL1 | NA | 0.167677738 | 0.403158283 | SIGLEC15 | pink | -0.047347 | 0.818343 |
| GALNT12 | NA | 0.035183 | 0.864516 | FAM134B | NA | 0.168923723 | 0.399621796 | SCIMP | pink | -0.045231 | 0.826337 |
| UMODL1 | NA | -0.034982 | 0.865285 | DTX3L | NA | 0.168932428 | 0.39959715 | EPHX1 | pink | -0.045228 | 0.826349 |
| COLEC12 | NA | 0.034888 | 0.86564 | MX1 | NA | 0.168949925 | 0.399547617 | CSTF3 | pink | -0.044849 | 0.827782 |
| YES1 | NA | -0.034655 | 0.86653 | AARS2 | NA | 0.168989559 | 0.399435423 | FASTK | pink | -0.043898 | 0.831382 |
| GABARAPL1 | NA | -0.034403 | 0.867493 | TMEM170B | NA | 0.169277273 | 0.398621514 | MMP9 | pink | -0.040812 | 0.843086 |
| RNF13 | NA | 0.034222 | 0.868185 | GLB1L | NA | 0.169505353 | 0.397976967 | F3 | pink | -0.040138 | 0.845647 |
| ITGA3 | NA | -0.033955 | 0.869204 | PCBD1 | NA | 0.169510779 | 0.397961641 | MFHAS1 | pink | -0.03895 | 0.850163 |
| KDM2A | NA | 0.033923 | 0.869327 | OMA1 | NA | 0.169582023 | 0.397760434 | SPTSSA | pink | -0.037397 | 0.856076 |
| OFD1 | NA | -0.033828 | 0.86969 | NONO | NA | 0.16981594 | 0.397100211 | GPR68 | pink | -0.036175 | 0.860731 |
| ZER1 | NA | 0.033483 | 0.871006 | TBKBP1 | NA | 0.170221443 | 0.395957159 | TMEM170B | pink | -0.033153 | 0.872268 |
| KIAA0556 | NA | -0.03328 | 0.871781 | NDUFA7 | NA | 0.170417318 | 0.395405687 | LAMP3 | pink | -0.031968 | 0.8768 |
| LINC00515 | NA | -0.033255 | 0.871878 | RPS6KB2 | NA | 0.170637719 | 0.394785687 | DOCK6 | pink | -0.031759 | 0.877599 |
| PSENEN | NA | -0.032937 | 0.873094 | GNA12 | NA | 0.171004297 | 0.3937557 | NAIP | pink | -0.031558 | 0.878368 |
| L1TD1 | NA | -0.032778 | 0.8737 | SIKE1 | NA | 0.171160814 | 0.393316396 | ECM1 | pink | -0.028436 | 0.890326 |
| VPS16 | NA | 0.032744 | 0.873833 | HAUS5 | NA | 0.171546129 | 0.392236099 | SDHA | pink | -0.028157 | 0.891396 |
| SNX29 | NA | 0.032654 | 0.874174 | CD9 | NA | 0.172564747 | 0.389388376 | DNAJC30 | pink | -0.025813 | 0.900391 |
| GTPBP5 | NA | 0.032138 | 0.876149 | RPLP1 | NA | 0.17280464 | 0.388719437 | NDE1 | pink | -0.025545 | 0.90142 |
| SELRC1 | NA | 0.03208 | 0.876371 | LOC1005065[illegible] | NA | 0.172950308 | 0.388313563 | ADCK2 | pink | -0.024593 | 0.905075 |
| TAF15 | NA | -0.03203 | 0.876562 | MBD3 | NA | 0.173195422 | 0.38763115 | MS4A6A | pink | -0.023876 | 0.907833 |
| NPIP | NA | -0.031911 | 0.877018 | JHDM1D | NA | 0.173401832 | 0.387057025 | PTS | pink | -0.022507 | 0.913095 |
| ZBTB48 | NA | -0.03189 | 0.877099 | DCAF4 | NA | 0.173890584 | 0.385699512 | SLC1A4 | pink | -0.019879 | 0.923212 |
| GPR85 | NA | -0.031885 | 0.877117 | NUBP1 | NA | 0.17394415 | 0.385550896 | HIST1H2BK | pink | -0.018955 | 0.926774 |
| LINC00656 | NA | 0.031799 | 0.877447 | LPIN2 | NA | 0.174391269 | 0.38431169 | LIMD2 | pink | -0.018703 | 0.927744 |

**Table 6. Gene module membership and correlation with growth rate (48/72h), with adjusted p values for each module**

| | | | | | | | | | | | |
|---|---|---|---|---|---|---|---|---|---|---|---|
| ARHGAP6 | NA | -0.031748 | 0.877639 | RABL6 | NA | 0.174497347 | 0.384018026 | KCNN4 | pink | -0.016044 | 0.937997 |
| TMEM55A | NA | -0.031637 | 0.878067 | SPG7 | NA | 0.174769177 | 0.383266089 | RBP7 | pink | -0.015854 | 0.938727 |
| EHHADH | NA | 0.031336 | 0.879219 | CLK2 | NA | 0.174992121 | 0.382650011 | AMPD3 | pink | -0.014382 | 0.944409 |
| TNS1 | NA | -0.031045 | 0.880331 | GZF1 | NA | 0.175019512 | 0.382574359 | SYNJ2BP | pink | -0.013608 | 0.947394 |
| PMS2P5 | NA | 0.030923 | 0.880799 | OSBPL3 | NA | 0.175197599 | 0.382082702 | GRAMD1A | pink | -0.011263 | 0.956451 |
| BIN3 | NA | 0.03073 | 0.881538 | TSR3 | NA | 0.175797055 | 0.380430425 | CSF1R | pink | -0.008364 | 0.967653 |
| DDX18 | NA | -0.030559 | 0.882192 | AP2S1 | NA | 0.176600009 | 0.378223713 | HLX | pink | -0.007318 | 0.971696 |
| PRKRIP1 | NA | -0.030043 | 0.884168 | C10orf128 | NA | 0.176668716 | 0.378035233 | ZBTB10 | pink | -0.006061 | 0.976557 |
| PPP1R2 | NA | -0.029795 | 0.885116 | CALU | NA | 0.177096225 | 0.376863699 | NDUFA10 | pink | -0.004507 | 0.982567 |
| ZBTB17 | NA | 0.029743 | 0.885317 | TDRD9 | NA | 0.177245103 | 0.376456212 | RFTN1 | pink | -0.004264 | 0.983508 |
| BRF1 | NA | -0.029685 | 0.885539 | SGMS1 | NA | 0.179105649 | 0.371385337 | PKM | pink | -0.0015 | 0.994198 |
| CEP250 | NA | -0.029439 | 0.886481 | GPAA1 | NA | 0.180045096 | 0.368840085 | CCNB2 | pink | -0.001163 | 0.995501 |
| LST1 | NA | -0.029416 | 0.886568 | IMPDH2 | NA | 0.18036277 | 0.367981719 | BRK1 | pink | 0.001397 | 0.994597 |
| SMARCA4 | NA | -0.029166 | 0.887527 | SETDB2 | NA | 0.181410561 | 0.365158833 | C6orf106 | pink | 0.003804 | 0.985285 |
| PRSS36 | NA | -0.028953 | 0.888343 | SDHD | NA | 0.181556202 | 0.364767464 | O3FAR1 | pink | 0.003943 | 0.984748 |
| CMSS1 | NA | -0.02864 | 0.889545 | SFXN5 | NA | 0.182083208 | 0.363353344 | ZFP14 | pink | 0.004855 | 0.981223 |
| ZNF136 | NA | 0.02851 | 0.890043 | AXIN1 | NA | 0.182087233 | 0.363342554 | DRAM1 | pink | 0.006561 | 0.974624 |
| PPIL3 | NA | 0.028278 | 0.890932 | TRAPPC6A | NA | 0.182163546 | 0.363138054 | CCL4 | pink | 0.006768 | 0.973822 |
| C10orf10 | NA | 0.028072 | 0.891722 | ACYP1 | NA | 0.182459075 | 0.362346744 | TNFRSF10A | pink | 0.007488 | 0.971039 |
| TGIF1 | NA | -0.027814 | 0.89271 | DDRGK1 | NA | 0.182507792 | 0.362216397 | MRPL1 | pink | 0.007661 | 0.97037 |
| SMC3 | NA | -0.027481 | 0.893987 | SLC2A1 | NA | 0.182979633 | 0.360955363 | NEK6 | pink | 0.008536 | 0.96699 |
| SFI1 | NA | 0.027481 | 0.89399 | SNAP29 | NA | 0.183912341 | 0.358470242 | CYTH4 | pink | 0.009256 | 0.964205 |
| TLCD2 | NA | -0.02744 | 0.894146 | SLC16A1 | NA | 0.18395551 | 0.358355467 | KIDINS220 | pink | 0.009483 | 0.96333 |
| SKIV2L2 | NA | -0.026789 | 0.896644 | SLC25A22 | NA | 0.184358963 | 0.357283841 | STEAP3 | pink | 0.009743 | 0.962323 |
| FKBP2 | NA | -0.026645 | 0.897195 | IFITM3 | NA | 0.184671314 | 0.356455497 | CCR7 | pink | 0.010864 | 0.957994 |
| CCZ1 | NA | 0.02624 | 0.89875 | GSDMD | NA | 0.185210626 | 0.355027936 | GBP2 | pink | 0.014457 | 0.944117 |
| TDRKH | NA | -0.025864 | 0.900195 | SAMD4A | NA | 0.185224084 | 0.354992356 | SLC2A6 | pink | 0.014841 | 0.942636 |
| AFTPH | NA | -0.025655 | 0.900999 | CRABP2 | NA | 0.185296326 | 0.354801399 | DUSP7 | pink | 0.01581 | 0.938899 |
| ZBTB45 | NA | 0.025615 | 0.90115 | IER5 | NA | 0.185996982 | 0.352952528 | KLF9 | pink | 0.015954 | 0.93834 |
| MGC2752 | NA | 0.025566 | 0.901338 | PPAPDC1B | NA | 0.186237946 | 0.352318001 | SLC46A1 | pink | 0.016342 | 0.936845 |
| SLC35E2 | NA | -0.025276 | 0.902451 | TREML3P | NA | 0.186250882 | 0.352283956 | SLC28A3 | pink | 0.017059 | 0.93408 |
| WDR74 | NA | 0.025151 | 0.902934 | PIM3 | NA | 0.187056952 | 0.350166401 | TNIP1 | pink | 0.019144 | 0.926043 |
| CEP78 | NA | -0.025099 | 0.903131 | SUSD1 | NA | 0.187219571 | 0.34974012 | DGAT2 | pink | 0.021848 | 0.915633 |
| GON4L | NA | -0.025066 | 0.903261 | ECHS1 | NA | 0.188055794 | 0.347552971 | SCARF1 | pink | 0.023023 | 0.911112 |
| NAALADL1 | NA | 0.024802 | 0.904273 | KANSL1-AS1 | NA | 0.18810318 | 0.347429279 | USP12 | pink | 0.023864 | 0.90788 |
| WDR12 | NA | -0.024658 | 0.904827 | CD14 | NA | 0.188241829 | 0.347067508 | P2RY2 | pink | 0.024517 | 0.905367 |
| SERPINE1 | NA | 0.024461 | 0.905583 | TMCC3 | NA | 0.189067352 | 0.344918173 | SLC2A1 | pink | 0.025365 | 0.902111 |
| MSL1 | NA | -0.02432 | 0.906126 | MXI1 | NA | 0.191672591 | 0.338187574 | DUSP14 | pink | 0.025775 | 0.900535 |
| SP100 | NA | -0.02428 | 0.906281 | TOR2A | NA | 0.191826709 | 0.337791909 | TMEM132A | pink | 0.026924 | 0.896126 |
| MACF1 | NA | -0.024212 | 0.906539 | LILRB1 | NA | 0.192237746 | 0.336738025 | AGFG2 | pink | 0.027136 | 0.895314 |
| RICTOR | NA | -0.024167 | 0.906712 | INTS7 | NA | 0.193049126 | 0.334663509 | MYO10 | pink | 0.027579 | 0.893611 |
| SLC31A1 | NA | 0.024103 | 0.906961 | ZFYVE26 | NA | 0.193121589 | 0.334478616 | XPNPEP1 | pink | 0.028615 | 0.88964 |
| MEA1 | NA | -0.024082 | 0.907041 | LINC00263 | NA | 0.193822437 | 0.332693541 | KATNB1 | pink | 0.033352 | 0.871508 |
| CCNB2 | NA | 0.024018 | 0.907287 | GALNT12 | NA | 0.194538931 | 0.330874596 | CD300C | pink | 0.04043 | 0.844537 |
| CHAF1B | NA | 0.024003 | 0.907343 | RDX | NA | 0.194793173 | 0.330230613 | B3GALNT1 | pink | 0.040779 | 0.84321 |
| SYAP1 | NA | -0.023963 | 0.907498 | CASP8 | NA | 0.194978378 | 0.329761977 | DAG1 | pink | 0.042164 | 0.837954 |
| ZNF200 | NA | 0.023785 | 0.908183 | APOA1BP | NA | 0.195191794 | 0.329222457 | TP53INP2 | pink | 0.042292 | 0.837467 |
| MRRF | NA | -0.023211 | 0.91039 | RABL5 | NA | 0.195524806 | 0.32838167 | AIFM2 | pink | 0.042356 | 0.837226 |
| GGA1 | NA | -0.02307 | 0.910931 | BAGE2 | NA | 0.195648279 | 0.32807026 | ZC3H12A | pink | 0.042648 | 0.836119 |
| DUSP1 | NA | -0.022974 | 0.911302 | BORA | NA | 0.196433805 | 0.326093297 | CHD7 | pink | 0.044242 | 0.830079 |
| PTMA | NA | 0.022914 | 0.911533 | ITPR3 | NA | 0.196474371 | 0.325991401 | RASGRP1 | pink | 0.04603 | 0.823315 |
| GAS7 | NA | -0.022841 | 0.911812 | TUT1 | NA | 0.196557881 | 0.325781697 | MFSD12 | pink | 0.048133 | 0.815378 |
| ZNF486 | NA | -0.022589 | 0.912781 | SMYD5 | NA | 0.197163029 | 0.324264553 | DOCK4 | pink | 0.051755 | 0.801747 |
| RELT | NA | -0.022576 | 0.912833 | SNX7 | NA | 0.197352026 | 0.323791611 | CCND1 | pink | 0.055351 | 0.788268 |
| FAM108B1 | NA | -0.022524 | 0.913031 | BRWD1 | NA | 0.198059061 | 0.322026089 | ITGAX | pink | 0.059788 | 0.771718 |
| OAT | NA | 0.02235 | 0.913699 | HBEGF | NA | 0.198095338 | 0.321935662 | CPSF1 | pink | 0.060054 | 0.770729 |

Table 6. Gene module membership and correlation with growth rate (48/72h), with adjusted p values for each module

| | | | | | | | | | | | |
|---|---|---|---|---|---|---|---|---|---|---|---|
| ZNF124 | NA | 0.022038 | 0.914902 | MFAP1 | NA | 0.198313931 | 0.321391107 | CHD9 | pink | 0.06256 | 0.761431 |
| MCM9 | NA | -0.021923 | 0.915345 | SNRPF | NA | 0.198584518 | 0.320717809 | RAB32 | pink | 0.062782 | 0.760606 |
| SMCR7L | NA | 0.021795 | 0.915837 | NFKBIL1 | NA | 0.198763383 | 0.320273214 | WDR91 | pink | 0.063792 | 0.756868 |
| ZNF678 | NA | -0.021749 | 0.916014 | MDK | NA | 0.198764096 | 0.320271442 | FAM46A | pink | 0.065667 | 0.749943 |
| SORT1 | NA | 0.021672 | 0.91631 | SDSL | NA | 0.19969431 | 0.317965382 | MFI2 | pink | 0.066299 | 0.747612 |
| MBD5 | NA | -0.021279 | 0.917821 | C7orf13 | NA | 0.199936272 | 0.31736722 | NUB1 | pink | 0.066303 | 0.747598 |
| SUCNR1 | NA | 0.021105 | 0.918493 | GTF3C6 | NA | 0.200909123 | 0.314969197 | CLSTN3 | pink | 0.06944 | 0.736064 |
| MTPAP | NA | 0.021104 | 0.918497 | ISG15 | NA | 0.201360243 | 0.313861016 | ARHGAP6 | pink | 0.07262 | 0.724429 |
| HK3 | NA | 0.021018 | 0.918828 | MCCC2 | NA | 0.201652068 | 0.313145428 | B4GALT5 | pink | 0.074395 | 0.717961 |
| LOC645212 | NA | 0.020846 | 0.919489 | RALB | NA | 0.201703278 | 0.31301996 | TIFA | pink | 0.075199 | 0.715037 |
| HSPA1L | NA | 0.020185 | 0.922033 | ENPP2 | NA | 0.201835627 | 0.312695839 | TRPC6 | pink | 0.07533 | 0.71456 |
| RNPEPL1 | NA | -0.020073 | 0.922467 | ZMYM3 | NA | 0.201934905 | 0.312452843 | SHPK | pink | 0.077652 | 0.706139 |
| SLC26A11 | NA | -0.019975 | 0.922841 | CDR1 | NA | 0.202574122 | 0.310891078 | SYT11 | pink | 0.077695 | 0.705985 |
| NCEH1 | NA | -0.019899 | 0.923135 | GSTT1 | NA | 0.202918605 | 0.310051431 | ABCA1 | pink | 0.078188 | 0.7042 |
| PGM5P2 | NA | 0.019867 | 0.923259 | TNK2 | NA | 0.203381804 | 0.308924638 | FAM225A | pink | 0.080737 | 0.695004 |
| ADAM10 | NA | -0.019702 | 0.923895 | THAP1 | NA | 0.203599008 | 0.308397139 | BMF | pink | 0.082947 | 0.687065 |
| IQCE | NA | 0.019658 | 0.924064 | MRPS21 | NA | 0.203655491 | 0.308260055 | ACAD8 | pink | 0.083562 | 0.684859 |
| CCDC90B | NA | -0.019233 | 0.925701 | PPFIBP1 | NA | 0.203704404 | 0.308141375 | PTGR2 | pink | 0.083982 | 0.683358 |
| FAM35A | NA | -0.019174 | 0.925928 | ALDOA | NA | 0.203734154 | 0.308069206 | GMEB2 | pink | 0.085314 | 0.678594 |
| DHRS4L2 | NA | -0.019049 | 0.92641 | HERC5 | NA | 0.203849355 | 0.30778984 | CKB | pink | 0.088456 | 0.667412 |
| KIAA1683 | NA | 0.018999 | 0.926604 | C10orf118 | NA | 0.204206895 | 0.306923801 | MKL2 | pink | 0.093017 | 0.651304 |
| CAD | NA | 0.018981 | 0.926673 | BARD1 | NA | 0.204268694 | 0.306774264 | STX11 | pink | 0.094448 | 0.646282 |
| NPHP3 | NA | -0.018713 | 0.927705 | ADAL | NA | 0.204456333 | 0.306320505 | RCOR1 | pink | 0.09567 | 0.642003 |
| OSBPL2 | NA | 0.018685 | 0.927811 | CSF3R | NA | 0.204619656 | 0.305925889 | ASPHD1 | pink | 0.106183 | 0.605668 |
| KIAA0930 | NA | 0.018537 | 0.928384 | RPN1 | NA | 0.204957149 | 0.30511145 | CKAP4 | pink | 0.108824 | 0.59668 |
| RIN3 | NA | 0.018499 | 0.92853 | UXT | NA | 0.20548228 | 0.303846893 | OCSTAMP | pink | 0.11206 | 0.585743 |
| KTN1 | NA | 0.018435 | 0.928777 | ETFDH | NA | 0.205497716 | 0.303809771 | NCDN | pink | 0.112686 | 0.583636 |
| PPP4R1 | NA | -0.01841 | 0.928872 | SLC41A1 | NA | 0.205781115 | 0.303128734 | CCL3L3 | pink | 0.112833 | 0.583142 |
| SQRDL | NA | 0.018255 | 0.929469 | AGFG2 | NA | 0.206864538 | 0.300533946 | CDKN1A | pink | 0.114562 | 0.577347 |
| RBAK | NA | -0.017864 | 0.930978 | LOC220906 | NA | 0.209133155 | 0.295145779 | DUSP2 | pink | 0.117187 | 0.568596 |
| F13A1 | NA | 0.017461 | 0.932532 | MRPL42 | NA | 0.209624987 | 0.293985696 | NDRG1 | pink | 0.117838 | 0.566433 |
| LPXN | NA | -0.017384 | 0.932828 | MIS18A | NA | 0.210230892 | 0.292560501 | ADAMTS6 | pink | 0.121953 | 0.552861 |
| YTHDF2 | NA | 0.017308 | 0.933121 | DDX10 | NA | 0.210481409 | 0.291972516 | TNFRSF4 | pink | 0.12204 | 0.552576 |
| TMEM229B | NA | 0.017181 | 0.933612 | OSBPL1A | NA | 0.210531233 | 0.291855662 | LOH12CR1 | pink | 0.122281 | 0.551784 |
| DHX9 | NA | -0.016863 | 0.934837 | PPHLN1 | NA | 0.210638725 | 0.291603661 | CLCF1 | pink | 0.126649 | 0.537556 |
| BAX | NA | 0.016548 | 0.936053 | TMX1 | NA | 0.210752738 | 0.291336523 | A4GALT | pink | 0.126802 | 0.537059 |
| GTSF1 | NA | -0.016439 | 0.93647 | NDUFS1 | NA | 0.210754955 | 0.29133133 | TRIM14 | pink | 0.131194 | 0.522932 |
| NDUFAF6 | NA | 0.015749 | 0.939134 | APBA1 | NA | 0.211027775 | 0.290692733 | ELMSAN1 | pink | 0.1327 | 0.518131 |
| LMBR1 | NA | -0.015705 | 0.939301 | FARP2 | NA | 0.211131326 | 0.29045058 | ACSL4 | pink | 0.132776 | 0.517888 |
| LPCAT3 | NA | 0.015705 | 0.939304 | PROS1 | NA | 0.21131418 | 0.290023287 | GEM | pink | 0.134544 | 0.512279 |
| CHD9 | NA | 0.015595 | 0.939728 | KBTBD7 | NA | 0.211922397 | 0.288604871 | ABTB2 | pink | 0.138047 | 0.501255 |
| VAV2 | NA | 0.015457 | 0.940258 | KIAA1919 | NA | 0.212201829 | 0.287954683 | APOO | pink | 0.141227 | 0.49135 |
| THAP3 | NA | -0.015376 | 0.940572 | RRBP1 | NA | 0.212259839 | 0.287819823 | NFKBID | pink | 0.141553 | 0.490339 |
| NFRKB | NA | 0.015095 | 0.941657 | SLIT3 | NA | 0.212660943 | 0.286888425 | FAM83H | pink | 0.143954 | 0.482934 |
| ANKRD9 | NA | 0.014644 | 0.943398 | C1orf123 | NA | 0.212664067 | 0.286881179 | CYP2S1 | pink | 0.147645 | 0.471662 |
| EIF5B | NA | 0.014483 | 0.944017 | DNTTIP2 | NA | 0.21365225 | 0.284594729 | PXN | pink | 0.148875 | 0.467934 |
| FAM98B | NA | 0.014037 | 0.94574 | XRCC6 | NA | 0.213951233 | 0.28390523 | MTHFD2 | pink | 0.151907 | 0.458818 |
| PPM1L | NA | 0.013683 | 0.947104 | TPM3P9 | NA | 0.215012504 | 0.281466365 | C16orf58 | pink | 0.152921 | 0.455789 |
| EIF3I | NA | -0.01362 | 0.94735 | SEMA7A | NA | 0.219283926 | 0.271785687 | ITPR3 | pink | 0.153193 | 0.454977 |
| TRIM38 | NA | -0.013583 | 0.947494 | ATG2A | NA | 0.21933712 | 0.271666496 | FAM129B | pink | 0.153853 | 0.453014 |
| SLC52A2 | NA | -0.01331 | 0.948547 | GNG12 | NA | 0.219807659 | 0.270613622 | TOMM20 | pink | 0.155074 | 0.44939 |
| NUMBL | NA | 0.012862 | 0.950276 | LY6G5C | NA | 0.219905744 | 0.270394478 | MMP14 | pink | 0.161544 | 0.43046 |
| CSNK1D | NA | -0.012574 | 0.951386 | C1orf35 | NA | 0.220141767 | 0.269867621 | RAB13 | pink | 0.163575 | 0.424608 |
| SENP7 | NA | 0.012396 | 0.952074 | STK19 | NA | 0.220324784 | 0.26945954 | PDLIM4 | pink | 0.165828 | 0.418167 |
| VMO1 | NA | 0.012263 | 0.95259 | RPS6KA5 | NA | 0.221658235 | 0.266498283 | SLC38A2 | pink | 0.173634 | 0.396276 |
| RAD1 | NA | 0.011941 | 0.953831 | PRKCDBP | NA | 0.221705244 | 0.266394274 | PLAGL2 | pink | 0.176253 | 0.38908 |

**Table 6. Gene module membership and correlation with growth rate (48/72h), with adjusted p values for each module**

| | | | | | | | | | | | |
|---|---|---|---|---|---|---|---|---|---|---|---|
| FCGR1B | NA | -0.011569 | 0.955268 | TMEM18 | NA | 0.222091271 | 0.265541158 | TNKS1BP1 | pink | 0.178888 | 0.381913 |
| NAIP | NA | -0.011555 | 0.955324 | ZNF776 | NA | 0.222287259 | 0.265108704 | SPHK1 | pink | 0.18275 | 0.37155 |
| LRCH4 | NA | 0.011378 | 0.956008 | CTSL1 | NA | 0.222770805 | 0.264043691 | LONP1 | pink | 0.18786 | 0.358089 |
| ASNSD1 | NA | -0.01132 | 0.956232 | B4GALT4 | NA | 0.222826123 | 0.263922031 | LRRFIP2 | pink | 0.202475 | 0.321211 |
| ATG2B | NA | 0.011048 | 0.957282 | MBNL2 | NA | 0.223463088 | 0.262523761 | RELB | pink | 0.202813 | 0.320386 |
| PLEKHA7 | NA | 0.010429 | 0.959675 | HLA-DMB | NA | 0.223551469 | 0.262330127 | NRSN2 | pink | 0.204023 | 0.317446 |
| USP38 | NA | 0.010374 | 0.959887 | TSPAN3 | NA | 0.223819916 | 0.261742552 | RUSC2 | pink | 0.205019 | 0.315038 |
| WDR13 | NA | -0.010225 | 0.960462 | CFP | NA | 0.223925448 | 0.261511799 | IL17RA | pink | 0.219614 | 0.281046 |
| STEAP3 | NA | -0.010047 | 0.961149 | HLA-B | NA | 0.224084121 | 0.261165098 | ZSWIM4 | pink | 0.221368 | 0.277125 |
| TMEM158 | NA | -0.009643 | 0.962712 | MAP1LC3B2 | NA | 0.224127293 | 0.261070817 | PNKD | pink | 0.223045 | 0.273408 |
| IPO7 | NA | -0.009539 | 0.963111 | POLR3B | NA | 0.224406856 | 0.260460839 | POLR3D | pink | 0.225946 | 0.267054 |
| SIN3B | NA | -0.009522 | 0.963178 | FCER1G | NA | 0.224599512 | 0.260041021 | ITGA5 | pink | 0.230672 | 0.25691 |
| HCAR2 | NA | 0.009491 | 0.963298 | TGM2 | NA | 0.226377985 | 0.256186309 | EMR2 | pink | 0.234278 | 0.249339 |
| ST14 | NA | 0.009376 | 0.963744 | ATP5E | NA | 0.227709306 | 0.253325273 | NFKB2 | pink | 0.264755 | 0.191189 |
| FAM193A | NA | 0.009147 | 0.964626 | MKKS | NA | 0.228228957 | 0.252214221 | EHD1 | pink | 0.267004 | 0.187306 |
| LPAR1 | NA | -0.009108 | 0.964778 | MSH6 | NA | 0.228985492 | 0.250602402 | DENND3 | pink | 0.267301 | 0.186797 |
| NAA35 | NA | -0.009067 | 0.964937 | CD55 | NA | 0.229190761 | 0.250166235 | IFI6 | pink | 0.269286 | 0.183421 |
| OAS3 | NA | -0.009008 | 0.965165 | SLAIN2 | NA | 0.229702869 | 0.249080256 | SDC4 | pink | 0.269686 | 0.182746 |
| LEMD2 | NA | 0.008798 | 0.965976 | SLC27A3 | NA | 0.229978982 | 0.248496013 | AKIRIN2 | pink | 0.272144 | 0.178635 |
| NDUFS1 | NA | -0.00873 | 0.966241 | ALG14 | NA | 0.230018423 | 0.24841263 | FADS3 | pink | 0.277493 | 0.169912 |
| SLC35C1 | NA | -0.008479 | 0.967207 | ALKBH5 | NA | 0.23019464 | 0.248040316 | LTBP3 | pink | 0.305281 | 0.129381 |
| HLA-DOA | NA | -0.008246 | 0.968109 | CHD9 | NA | 0.231270799 | 0.245774543 | CTTN | pink | 0.310955 | 0.122055 |
| CEACAM8 | NA | -0.008236 | 0.968148 | NAMPT | NA | 0.231627735 | 0.245026058 | CFP | pink | 0.314717 | 0.117368 |
| ANKDD1A | NA | 0.008168 | 0.968411 | ITFG2 | NA | 0.232196449 | 0.243836585 | RSAD2 | pink | 0.31739 | 0.114118 |
| AKR1C1 | NA | -0.008035 | 0.968925 | AP4B1 | NA | 0.232373277 | 0.243467525 | TBKBP1 | pink | 0.343875 | 0.085416 |
| TCP1 | NA | -0.008028 | 0.968951 | SNRPG | NA | 0.232516629 | 0.243168602 | PLAUR | pink | 0.35423 | 0.07582 |
| CNTF | NA | 0.007995 | 0.969079 | PCSK6 | NA | 0.232734379 | 0.242715005 | DNAJB12 | pink | 0.367605 | 0.06467 |
| TMEM170E | NA | -0.007903 | 0.969435 | AGRN | NA | 0.23292005 | 0.242328675 | DENND5A | pink | 0.384571 | 0.052399 |
| MRPL42 | NA | 0.007449 | 0.971193 | CYB5R1 | NA | 0.234692173 | 0.238661794 | ELF4 | pink | 0.405529 | 0.039844 |
| BTD | NA | 0.007129 | 0.972428 | RPL3 | NA | 0.23516757 | 0.237684388 | TBC1D9 | pink | 0.459105 | 0.01831 |
| NCR3LG1 | NA | -0.007128 | 0.97243 | FLJ45340 | NA | 0.235312199 | 0.23738756 | BRCC3 | purple | -0.400306 | 0.042722 |
| MTF1 | NA | 0.006957 | 0.973092 | ZNF669 | NA | 0.235710442 | 0.236571501 | FH | purple | -0.391783 | 0.047771 |
| SRR | NA | 0.006771 | 0.973814 | TXNRD2 | NA | 0.237738097 | 0.232445421 | C12orf45 | purple | -0.379263 | 0.056024 |
| KIAA1704 | NA | -0.006766 | 0.97383 | CCDC125 | NA | 0.238040817 | 0.231833554 | IBTK | purple | -0.372516 | 0.06091 |
| TRA2A | NA | -0.006764 | 0.97384 | CTU2 | NA | 0.238665622 | 0.230574073 | MED28 | purple | -0.34731 | 0.082135 |
| SLC1A4 | NA | -0.006406 | 0.975223 | TMEM107 | NA | 0.238811099 | 0.230281477 | PAPD5 | purple | -0.340635 | 0.088599 |
| DC1002892 | NA | -0.006342 | 0.97547 | MAP7 | NA | 0.239253683 | 0.229392839 | OR2A9P | purple | -0.331832 | 0.0977 |
| ZNF77 | NA | -0.006154 | 0.976197 | DGCR14 | NA | 0.239543288 | 0.228812597 | CISD1 | purple | -0.331278 | 0.098295 |
| DOHH | NA | -0.00535 | 0.979307 | DENND3 | NA | 0.240163451 | 0.227573366 | ITGA6 | purple | -0.324611 | 0.105672 |
| PSMC3 | NA | -0.005167 | 0.980014 | PRSS21 | NA | 0.240635057 | 0.226633996 | TRIM33 | purple | -0.317264 | 0.11427 |
| WDR7 | NA | 0.005141 | 0.980115 | IGFBP6 | NA | 0.241389957 | 0.225135753 | RBBP8 | purple | -0.316494 | 0.1152 |
| ATP6V1F | NA | 0.005126 | 0.980171 | SYMPK | NA | 0.24210183 | 0.223729003 | OR2A20P | purple | -0.316299 | 0.115436 |
| MAPKBP1 | NA | -0.005035 | 0.980523 | GIMAP2 | NA | 0.242392646 | 0.223156016 | PRRG4 | purple | -0.311582 | 0.121264 |
| SECTM1 | NA | 0.005005 | 0.980639 | C10orf57 | NA | 0.242555933 | 0.222834727 | GRSF1 | purple | -0.309009 | 0.124532 |
| YPEL3 | NA | 0.004762 | 0.981581 | ZNF805 | NA | 0.24286612 | 0.22222525 | ATXN10 | purple | -0.306603 | 0.127647 |
| ITPRIPL2 | NA | -0.00427 | 0.983482 | MAK16 | NA | 0.244261494 | 0.219497381 | NSF | purple | -0.297024 | 0.140606 |
| SNORA42 | NA | -0.004104 | 0.984126 | PARP9 | NA | 0.244475938 | 0.219080165 | GMFB | purple | -0.294269 | 0.144503 |
| KLF9 | NA | -0.00403 | 0.984413 | PITPNB | NA | 0.245606028 | 0.216890325 | HSPA1B | purple | -0.293647 | 0.145393 |
| CHRM3 | NA | 0.00387 | 0.985032 | HSD17B7 | NA | 0.246762569 | 0.214664581 | PCNP | purple | -0.292417 | 0.147166 |
| LRP8 | NA | -0.003731 | 0.985568 | STIP1 | NA | 0.247047959 | 0.21411774 | SNRK | purple | -0.289703 | 0.15113 |
| MECR | NA | -0.003629 | 0.985962 | ALDH3B1 | NA | 0.24769652 | 0.212878524 | TMEM38B | purple | -0.286826 | 0.155416 |
| DGCR2 | NA | -0.003188 | 0.987667 | PCK2 | NA | 0.248221071 | 0.211879818 | RIC8B | purple | -0.282795 | 0.161566 |
| CAPNS2 | NA | 0.00313 | 0.987891 | DGCR2 | NA | 0.249612073 | 0.209246855 | EIF3J | purple | -0.281322 | 0.163855 |
| MBNL2 | NA | 0.002821 | 0.989087 | ZNF134 | NA | 0.250308406 | 0.207937187 | PTDSS1 | purple | -0.279358 | 0.166943 |
| NUFIP2 | NA | -0.002801 | 0.989164 | RNF4 | NA | 0.250336507 | 0.207884451 | TNFRSF12A | purple | -0.275407 | 0.173278 |
| CENPV | NA | -0.002668 | 0.98968 | S100A16 | NA | 0.25394852 | 0.201181671 | FNBP1 | purple | -0.272267 | 0.178431 |

**Table 6. Gene module membership and correlation with growth rate (48/72h), with adjusted p values for each module**

| | | | | | | | | | | | |
|---|---|---|---|---|---|---|---|---|---|---|---|
| SEC61A2 | NA | -0.002182 | 0.99156 | SDF2 | NA | 0.25566984 | 0.198040076 | ZNF621 | purple | -0.269078 | 0.183774 |
| MCM7 | NA | 0.002145 | 0.991704 | TOM1L1 | NA | 0.256027179 | 0.19739214 | DDX23 | purple | -0.26752 | 0.186422 |
| APBB2 | NA | 0.002074 | 0.991979 | ANKRD13C | NA | 0.25642481 | 0.196672857 | SAV1 | purple | -0.267346 | 0.18672 |
| CNIH | NA | -0.002036 | 0.992124 | ANKRD29 | NA | 0.257463698 | 0.194802099 | KIAA1715 | purple | -0.265422 | 0.190032 |
| ZMYND11 | NA | 0.002013 | 0.992213 | SMARCE1 | NA | 0.257967798 | 0.19389878 | C11orf57 | purple | -0.263829 | 0.192805 |
| GALNT10 | NA | 0.0019 | 0.99265 | FAM21A | NA | 0.258325735 | 0.19325913 | DCTN4 | purple | -0.263284 | 0.193761 |
| NRBP2 | NA | -0.001888 | 0.992696 | PFKFB2 | NA | 0.258499687 | 0.192948796 | CCDC92 | purple | -0.263086 | 0.194108 |
| WDR83 | NA | 0.001878 | 0.992735 | CCDC77 | NA | 0.258997304 | 0.192062934 | GCLM | purple | -0.262191 | 0.195685 |
| REXO4 | NA | -0.001644 | 0.993642 | C16orf58 | NA | 0.259089761 | 0.191898652 | MZT1 | purple | -0.260466 | 0.198749 |
| C1QA | NA | 0.001401 | 0.994582 | MVK | NA | 0.259291153 | 0.191541141 | RNF24 | purple | -0.255768 | 0.207263 |
| AHR | NA | -0.001331 | 0.994852 | ZBTB8OS | NA | 0.259595694 | 0.191001398 | ZC3H15 | purple | -0.245014 | 0.227669 |
| CRISPLD2 | NA | -0.001181 | 0.995433 | NBPF14 | NA | 0.260012994 | 0.190263515 | EIF3F | purple | -0.243541 | 0.230567 |
| SPTSSA | NA | -0.000993 | 0.996159 | NR4A3 | NA | 0.260123124 | 0.190069107 | SNX27 | purple | -0.243 | 0.231636 |
| TEX30 | NA | -0.000771 | 0.997017 | MRPS23 | NA | 0.260400403 | 0.189580251 | GATA3 | purple | -0.235947 | 0.245884 |
| DTX3L | NA | -0.00061 | 0.997641 | VPS39 | NA | 0.260996381 | 0.188532456 | GOLPH3 | purple | -0.23543 | 0.246951 |
| DNAJC11 | NA | -0.000101 | 0.999608 | LMF1 | NA | 0.261416439 | 0.18779636 | C17orf97 | purple | -0.232237 | 0.253606 |
| ZNF839 | NA | -1.7E-05 | 0.999934 | CHMP4A | NA | 0.261488533 | 0.187670225 | C12orf66 | purple | -0.229611 | 0.259164 |
| WEE1 | orange | -0.622682 | 0.000681 | FAM127C | NA | 0.262040299 | 0.186706805 | CPNE3 | purple | -0.225704 | 0.267581 |
| MTERFD3 | orange | -0.288508 | 0.1529 | DDX27 | NA | 0.263727402 | 0.183782296 | CAPRIN1 | purple | -0.224326 | 0.27059 |
| SPTBN1 | orange | -0.274721 | 0.174394 | HLA-DQA1 | NA | 0.264022183 | 0.183274595 | GNPTAB | purple | -0.223783 | 0.271783 |
| PHF19 | orange | -0.266434 | 0.188286 | ARL17A | NA | 0.264119071 | 0.183107937 | WDFY4 | purple | -0.223589 | 0.272209 |
| CD6 | orange | -0.245015 | 0.227669 | CNP | NA | 0.264120043 | 0.183106266 | VRK3 | purple | -0.221464 | 0.276911 |
| CLDN4 | orange | -0.22779 | 0.263066 | DAPK3 | NA | 0.264297536 | 0.182801235 | STIM1 | purple | -0.218716 | 0.283069 |
| SFXN1 | orange | -0.225333 | 0.26839 | MMAA | NA | 0.264765182 | 0.181999255 | TRIP12 | purple | -0.214782 | 0.292032 |
| DNAJC6 | orange | 0.219379 | 0.281575 | USP27X | NA | 0.265741314 | 0.180333154 | SOS1 | purple | -0.214044 | 0.293732 |
| JUP | orange | -0.207214 | 0.309772 | KCTD2 | NA | 0.267129729 | 0.177981714 | S100PBP | purple | -0.212173 | 0.298073 |
| RCOR1 | orange | 0.20151 | 0.323572 | ARL10 | NA | 0.268519108 | 0.175650165 | ADAM9 | purple | -0.211988 | 0.298504 |
| SLC28A3 | orange | -0.18992 | 0.352745 | GBE1 | NA | 0.268648268 | 0.17543451 | FAM161B | purple | -0.208836 | 0.305915 |
| PKN1 | orange | -0.175202 | 0.391959 | CECR5 | NA | 0.26906591 | 0.174738453 | UBE2F | purple | -0.206853 | 0.310635 |
| AGR2 | orange | -0.162078 | 0.428919 | CCNYL1 | NA | 0.269378288 | 0.174219098 | ATP6AP2 | purple | -0.201953 | 0.322487 |
| TSPAN1 | orange | -0.159642 | 0.435979 | BTD | NA | 0.269404095 | 0.17417624 | PCGF5 | purple | -0.199948 | 0.327416 |
| CCL22 | orange | -0.155131 | 0.449223 | UGDH | NA | 0.269465331 | 0.174074575 | KLHL12 | purple | -0.198158 | 0.331853 |
| WFDC2 | orange | -0.152499 | 0.457046 | CFLAR | NA | 0.27055642 | 0.172270092 | ALG11 | purple | -0.195832 | 0.337674 |
| CLU | orange | -0.143697 | 0.483725 | ZNF430 | NA | 0.2706888 | 0.172052054 | G3BP1 | purple | -0.193169 | 0.344415 |
| CD7 | orange | -0.132395 | 0.519101 | MED8 | NA | 0.270719919 | 0.172000828 | HTATIP2 | purple | -0.191443 | 0.348826 |
| LRIG1 | orange | 0.120863 | 0.556441 | NUDT15 | NA | 0.271644742 | 0.170483305 | ATP6V1A | purple | -0.187693 | 0.358524 |
| LMO7 | orange | -0.103416 | 0.615148 | CCL3L3 | NA | 0.272326811 | 0.169370163 | SPG21 | purple | -0.183026 | 0.370816 |
| SLPI | orange | -0.102864 | 0.617048 | STAT5A | NA | 0.272544853 | 0.169015398 | MTERFD1 | purple | -0.180169 | 0.378457 |
| TACSTD2 | orange | -0.101808 | 0.620684 | ZBTB17 | NA | 0.274676343 | 0.165574879 | UGCG | purple | -0.180138 | 0.378541 |
| CORO1A | orange | -0.097722 | 0.634845 | NCS1 | NA | 0.27503396 | 0.165002518 | CNIH3 | purple | -0.179732 | 0.379633 |
| DNALI1 | orange | -0.095801 | 0.641544 | PGRMC1 | NA | 0.276767607 | 0.16224764 | CLASP2 | purple | -0.174249 | 0.39458 |
| ESPL1 | orange | -0.092963 | 0.651495 | TTC7B | NA | 0.277578966 | 0.16096959 | SLC15A4 | purple | -0.171254 | 0.402883 |
| ANTXR2 | orange | -0.083901 | 0.683646 | ECI2 | NA | 0.278528649 | 0.159482737 | C11orf75 | purple | -0.166966 | 0.414936 |
| TPPP3 | orange | -0.076513 | 0.710264 | NDUFAF6 | NA | 0.279114938 | 0.158569707 | ATP11A | purple | -0.165942 | 0.417845 |
| SCGB1A1 | orange | 0.074357 | 0.718098 | ADAMTS6 | NA | 0.279145102 | 0.158522833 | MPP7 | purple | -0.165527 | 0.419026 |
| LCN2 | orange | 0.064403 | 0.75461 | MFGE8 | NA | 0.279440228 | 0.158064737 | RNGTT | purple | -0.162346 | 0.428144 |
| PIGK | orange | -0.060089 | 0.770599 | UHRF2 | NA | 0.279650887 | 0.157738324 | CCNI | purple | -0.161264 | 0.431269 |
| HMHA1 | orange | 0.05945 | 0.772976 | CABLES1 | NA | 0.279812881 | 0.157487645 | EIF1AD | purple | -0.160892 | 0.432348 |
| KRT17 | orange | -0.05655 | 0.783788 | ZNF673 | NA | 0.280403013 | 0.156576835 | XPOT | purple | -0.159033 | 0.437756 |
| CLDN1 | orange | -0.054346 | 0.792029 | CCRL2 | NA | 0.28049882 | 0.15642932 | TAOK2 | purple | -0.158424 | 0.439535 |
| IFITM3 | orange | -0.051858 | 0.80136 | LTBP3 | NA | 0.280980161 | 0.155689694 | MAPKAPK2 | purple | -0.158391 | 0.43963 |
| MUC20 | orange | 0.043635 | 0.832378 | PANK3 | NA | 0.284487603 | 0.150375255 | DBF4 | purple | -0.150491 | 0.463063 |
| IFITM1 | orange | -0.030988 | 0.880549 | MRPS16 | NA | 0.284823327 | 0.149873464 | MDN1 | purple | -0.145967 | 0.476768 |
| KRT19 | orange | 0.024026 | 0.907255 | SLX4 | NA | 0.285095274 | 0.149467876 | LOC648987 | purple | -0.139316 | 0.497289 |
| PIGR | orange | 0.023522 | 0.909195 | SAP30L | NA | 0.286170589 | 0.147871827 | TCP1 | purple | -0.127462 | 0.534923 |
| CTGF | orange | -0.008193 | 0.968315 | ATG10 | NA | 0.287682594 | 0.14564834 | MALT1 | purple | -0.124837 | 0.543437 |

**Table 6. Gene module membership and correlation with growth rate (48/72h), with adjusted p values for each module**

| | | | | | | | | | | | |
|---|---|---|---|---|---|---|---|---|---|---|---|
| SCGB3A1 | orange | -0.004025 | 0.984432 | ERCC4 | NA | 0.288850935 | 0.143946756 | ARID1A | purple | -0.122223 | 0.551974 |
| BPIFB1 | orange | 0.003397 | 0.986861 | HEG1 | NA | 0.289046493 | 0.143663347 | PBX3 | purple | -0.119906 | 0.559594 |
| CXCL17 | orange | -0.002326 | 0.991002 | SCYL3 | NA | 0.290664553 | 0.141333789 | RWDD3 | purple | -0.11947 | 0.561032 |
| HIST1H1C | pink | -0.6209 | 0.000713 | R3HCC1L | NA | 0.291926897 | 0.139535371 | ZNF347 | purple | -0.117406 | 0.567868 |
| HIST1H3F | pink | -0.592051 | 0.001441 | PGM5P2 | NA | 0.292128785 | 0.139249288 | PARP4 | purple | -0.115227 | 0.575123 |
| SURF2 | pink | -0.584037 | 0.001733 | ZDHHC14 | NA | 0.292752514 | 0.13836812 | KIAA1731 | purple | -0.112475 | 0.584347 |
| EPC2 | pink | 0.581034 | 0.001854 | KIAA0020 | NA | 0.292934356 | 0.138111984 | LRRC6 | purple | -0.106969 | 0.602989 |
| ALOX5 | pink | -0.56995 | 0.002368 | LILRA3 | NA | 0.296198819 | 0.133571944 | UROS | purple | -0.103612 | 0.614476 |
| PSMD11 | pink | 0.565427 | 0.002611 | KCNA10 | NA | 0.299914657 | 0.128537145 | FLOT1 | purple | -0.091188 | 0.657748 |
| SMG5 | pink | 0.508232 | 0.008028 | OSBPL5 | NA | 0.300148595 | 0.128224868 | WDR43 | purple | -0.088634 | 0.666783 |
| GBE1 | pink | 0.500723 | 0.009176 | FAIM3 | NA | 0.300741703 | 0.127435624 | EIF1 | purple | -0.087217 | 0.671815 |
| ZNF786 | pink | -0.4984 | 0.009558 | TMEM208 | NA | 0.302787617 | 0.124740355 | SERINC3 | purple | -0.072024 | 0.726604 |
| ARL8B | pink | 0.494905 | 0.010158 | MRFAP1 | NA | 0.303325935 | 0.124038162 | CXorf26 | purple | -0.065105 | 0.752016 |
| ARHGEF12 | pink | 0.492485 | 0.010591 | EXT1 | NA | 0.303519615 | 0.123786232 | COG5 | purple | -0.061631 | 0.764875 |
| YIPF5 | pink | 0.490112 | 0.011031 | ELOVL5 | NA | 0.303529868 | 0.123772906 | PSAP | purple | -0.060908 | 0.767557 |
| IFNGR2 | pink | -0.488934 | 0.011255 | ZDBF2 | NA | 0.304559637 | 0.122439823 | ATP2B4 | purple | -0.058589 | 0.776184 |
| DPCD | pink | -0.487673 | 0.011498 | DYNLT1 | NA | 0.304878563 | 0.122029101 | TRAPPC9 | purple | -0.056229 | 0.784986 |
| PIKFYVE | pink | 0.482122 | 0.012624 | ZNF252P | NA | 0.305486642 | 0.121248801 | CCDC109B | purple | -0.053936 | 0.793564 |
| PPP1R35 | pink | -0.476447 | 0.013868 | TMEM150A | NA | 0.307544786 | 0.118634921 | OSGIN2 | purple | -0.053655 | 0.794618 |
| SCO1 | pink | -0.47635 | 0.01389 | ABCD3 | NA | 0.308071331 | 0.11797291 | KDM3B | purple | -0.052568 | 0.798694 |
| GLT8D1 | pink | -0.469469 | 0.015534 | PHF20 | NA | 0.310578778 | 0.11485763 | HPGDS | purple | -0.048244 | 0.814956 |
| FOS | pink | -0.462007 | 0.017495 | SF3B14 | NA | 0.313044439 | 0.111853895 | NCOA1 | purple | -0.048175 | 0.815217 |
| FYTTD1 | pink | -0.461912 | 0.017521 | CCDC101 | NA | 0.317761166 | 0.106270451 | JOSD1 | purple | -0.041588 | 0.840137 |
| HIST1H2BF | pink | -0.459256 | 0.018267 | SLU7 | NA | 0.318195625 | 0.105766786 | BPHL | purple | -0.035939 | 0.861633 |
| MTMR4 | pink | 0.456087 | 0.019191 | BYSL | NA | 0.318412095 | 0.105516499 | ZFAT | purple | -0.03489 | 0.865635 |
| GDI1 | pink | 0.451976 | 0.020446 | CD63 | NA | 0.321383616 | 0.102125176 | TADA3 | purple | -0.028246 | 0.891052 |
| KIF13B | pink | 0.450515 | 0.020908 | P2RX1 | NA | 0.321667116 | 0.101805929 | TRIM24 | purple | -0.024645 | 0.904878 |
| PEX16 | pink | -0.449765 | 0.021148 | HLA-DRB1 | NA | 0.324927644 | 0.098187671 | PSMB4 | purple | -0.011366 | 0.956052 |
| STAT6 | pink | 0.4456 | 0.022525 | HSD3B7 | NA | 0.325051348 | 0.098052319 | CCL23 | purple | -0.00955 | 0.963068 |
| CPOX | pink | -0.440914 | 0.024159 | WDR12 | NA | 0.325246402 | 0.097839183 | NDUFV3 | purple | -0.000605 | 0.997659 |
| S100A6 | pink | -0.434978 | 0.026366 | DUSP10 | NA | 0.329958724 | 0.092794875 | UACA | purple | 0.006529 | 0.974746 |
| GPR75 | pink | 0.434602 | 0.026511 | METTL3 | NA | 0.330572549 | 0.092152507 | LST1 | purple | 0.009599 | 0.962879 |
| HIST1H1E | pink | -0.432044 | 0.027515 | RBM6 | NA | 0.330652908 | 0.09206866 | MSL1 | purple | 0.012114 | 0.953164 |
| PRDX3 | pink | -0.431186 | 0.027859 | GPD2 | NA | 0.331507517 | 0.091180509 | KTN1 | purple | 0.012816 | 0.950454 |
| PHF11 | pink | -0.430826 | 0.028004 | MCM3 | NA | 0.331939822 | 0.090733706 | C14orf28 | purple | 0.023558 | 0.909056 |
| PCYT1A | pink | 0.430672 | 0.028067 | CEP164 | NA | 0.332138055 | 0.090529379 | PICK1 | purple | 0.024864 | 0.904034 |
| TAF5L | pink | 0.430139 | 0.028283 | PLEKHO2 | NA | 0.336230059 | 0.086388701 | SUMF2 | purple | 0.026082 | 0.899356 |
| SH3GLB2 | pink | 0.428117 | 0.029117 | HOXB7 | NA | 0.338217753 | 0.084429915 | METTL18 | purple | 0.030583 | 0.8821 |
| PSMC5 | pink | -0.420819 | 0.032292 | MRS2 | NA | 0.340360428 | 0.082356361 | GPA33 | purple | 0.031737 | 0.877685 |
| IARS | pink | 0.419632 | 0.032834 | C15orf23 | NA | 0.340431115 | 0.082288621 | GRPEL1 | purple | 0.033414 | 0.871272 |
| TTC7B | pink | 0.41808 | 0.033553 | TMSB10 | NA | 0.342449891 | 0.080371835 | WDR11 | purple | 0.040458 | 0.84443 |
| ARRDC3 | pink | -0.417262 | 0.033937 | SP100 | NA | 0.347576469 | 0.075657197 | MVK | purple | 0.041718 | 0.839644 |
| AARS | pink | 0.414665 | 0.03518 | NUFIP1 | NA | 0.352335402 | 0.071472962 | ZDHHC18 | purple | 0.043098 | 0.834413 |
| RBMS1 | pink | 0.41382 | 0.035592 | DUSP23 | NA | 0.352404173 | 0.071413828 | RBM25 | purple | 0.050376 | 0.806929 |
| C20orf96 | pink | -0.413759 | 0.035622 | CHMP1B | NA | 0.353918499 | 0.07012123 | CD38 | purple | 0.056158 | 0.785252 |
| CCDC88A | pink | 0.408313 | 0.038373 | ZFP64 | NA | 0.359605013 | 0.065427545 | PRPF31 | purple | 0.060865 | 0.767718 |
| CLK3 | pink | 0.40829 | 0.038385 | METTL9 | NA | 0.363285013 | 0.062522107 | MSTO1 | purple | 0.071352 | 0.729061 |
| PRRC2B | pink | 0.405195 | 0.040023 | ACADS | NA | 0.366085913 | 0.060378625 | ZNF341 | purple | 0.07677 | 0.709335 |
| CUTC | pink | -0.404318 | 0.040497 | DAP | NA | 0.367186608 | 0.059552097 | FOXRED1 | purple | 0.078077 | 0.704603 |
| HIST1H1B | pink | -0.40316 | 0.041129 | GTF2E2 | NA | 0.368557985 | 0.058534662 | PRKAB1 | purple | 0.080371 | 0.69632 |
| MAD2L1BP | pink | 0.402832 | 0.04131 | MAPKAPK5-A | NA | 0.369708008 | 0.057691937 | MKRN1 | purple | 0.080868 | 0.694534 |
| CHST14 | pink | -0.400545 | 0.042587 | ACP1 | NA | 0.369811935 | 0.05761625 | SNORA47 | purple | 0.082338 | 0.689248 |
| ZNF93 | pink | -0.395086 | 0.045761 | FXR2 | NA | 0.37091548 | 0.056817328 | AAMP | purple | 0.08289 | 0.687269 |
| EIF2C2 | pink | 0.394427 | 0.046157 | TMEM143 | NA | 0.373771138 | 0.054790063 | OS9 | purple | 0.08291 | 0.687197 |
| LOC339803 | pink | -0.39379 | 0.046542 | PIKFYVE | NA | 0.373974271 | 0.054648042 | FLVCR2 | purple | 0.08466 | 0.680932 |
| TRUB1 | pink | -0.39377 | 0.046554 | SLC26A6 | NA | 0.375935601 | 0.053291537 | ATMIN | purple | 0.090691 | 0.659501 |

**Table 6. Gene module membership and correlation with growth rate (48/72h), with adjusted p values for each module**

| | | | | | | | | | | | |
|---|---|---|---|---|---|---|---|---|---|---|---|
| USP21 | pink | -0.393351 | 0.046808 | ZMYND11 | NA | 0.379350079 | 0.05099309 | ZNF585A | purple | 0.09678 | 0.638125 |
| TLE4 | pink | 0.392998 | 0.047024 | TYROBP | NA | 0.382297789 | 0.049072135 | CSTF2 | purple | 0.107666 | 0.600615 |
| RNASEH2A | pink | -0.392584 | 0.047277 | MASTL | NA | 0.383466293 | 0.048326608 | CNPPD1 | purple | 0.115223 | 0.575138 |
| IFIT1 | pink | -0.391287 | 0.048078 | TRAF3IP1 | NA | 0.383981093 | 0.048001002 | PHKA2 | purple | 0.116045 | 0.572396 |
| FBXW7 | pink | 0.391185 | 0.048142 | RABGGTB | NA | 0.384315021 | 0.047790722 | RIN3 | purple | 0.121414 | 0.554629 |
| RPS15A | pink | -0.390304 | 0.048693 | GTSF1 | NA | 0.393883112 | 0.042067864 | NRF1 | purple | 0.121592 | 0.554044 |
| PDZD11 | pink | -0.387432 | 0.050523 | ZNF397 | NA | 0.394230721 | 0.041870689 | DCAF5 | purple | 0.128399 | 0.531902 |
| HEATR6 | pink | -0.386517 | 0.051117 | ZNF804A | NA | 0.396012864 | 0.040871297 | ABCF3 | purple | 0.138152 | 0.500925 |
| HIST1H2AC | pink | -0.384656 | 0.052343 | WLS | NA | 0.396532064 | 0.040583735 | GPBAR1 | purple | 0.142529 | 0.487323 |
| CHRAC1 | pink | 0.38418 | 0.05266 | BTN2A1 | NA | 0.398024427 | 0.03976613 | MSL3 | purple | 0.144371 | 0.481654 |
| PIK3R5 | pink | 0.383652 | 0.053013 | SWI5 | NA | 0.399059638 | 0.039206723 | LOC148413 | purple | 0.145691 | 0.477613 |
| PSMB6 | pink | 0.383003 | 0.053451 | ZFYVE1 | NA | 0.401886473 | 0.037711036 | DEAF1 | purple | 0.147607 | 0.471777 |
| MRP63 | pink | -0.382507 | 0.053786 | PLA2G4A | NA | 0.404456107 | 0.036391279 | TRIB1 | purple | 0.149563 | 0.465858 |
| PSIMCT-1 | pink | 0.381276 | 0.054627 | TMBIM4 | NA | 0.410171459 | 0.033588141 | ZNF836 | purple | 0.15058 | 0.462797 |
| YEATS4 | pink | -0.376648 | 0.05788 | KIF13B | NA | 0.416094454 | 0.0308687 | TST | purple | 0.156629 | 0.444801 |
| DZIP1L | pink | -0.37242 | 0.060982 | ATP1B3 | NA | 0.427942079 | 0.025961049 | MAT2B | purple | 0.156955 | 0.443842 |
| FAF1 | pink | -0.37217 | 0.06117 | RPL29 | NA | 0.430649947 | 0.024933411 | PDCD6 | purple | 0.160002 | 0.434932 |
| KEAP1 | pink | 0.370289 | 0.062593 | MVD | NA | 0.434495925 | 0.023530881 | USPL1 | purple | 0.160912 | 0.432292 |
| MIIP | pink | 0.369234 | 0.063404 | TOMM20 | NA | 0.435041447 | 0.02333725 | SLC12A9 | purple | 0.164479 | 0.422019 |
| PSMD8 | pink | 0.364946 | 0.066779 | ZNF215 | NA | 0.440516943 | 0.021464659 | WDR20 | purple | 0.169391 | 0.408096 |
| ZNF134 | pink | 0.362917 | 0.068423 | EDEM2 | NA | 0.446348071 | 0.019607246 | ZNF673 | purple | 0.175846 | 0.390192 |
| SNORA27 | pink | -0.361808 | 0.069335 | RPLP2 | NA | 0.460882716 | 0.01554407 | PQLC3 | purple | 0.178671 | 0.382501 |
| UBE2G2 | pink | 0.361071 | 0.069947 | ALDH1A2 | NA | 0.470871104 | 0.013176428 | TMEM141 | purple | 0.179007 | 0.381591 |
| PDCD1LG2 | pink | -0.360062 | 0.07079 | SLC25A5 | NA | 0.487972145 | 0.009818507 | CDC23 | purple | 0.180388 | 0.377869 |
| ACVRL1 | pink | -0.353926 | 0.076089 | SRP14 | NA | 0.488702678 | 0.009692734 | TXN2 | purple | 0.185521 | 0.364216 |
| LOC440434 | pink | -0.352975 | 0.076937 | ANAPC16 | NA | 0.492819484 | 0.009008866 | GGCX | purple | 0.18933 | 0.354273 |
| SNX2 | pink | -0.352636 | 0.07724 | SCMH1 | NA | 0.511593676 | 0.006379945 | PDE1B | purple | 0.193193 | 0.344355 |
| SMUG1 | pink | -0.352204 | 0.077628 | IMPACT | NA | 0.522525582 | 0.00517207 | ZNF395 | purple | 0.198072 | 0.332068 |
| SCARB1 | pink | -0.348472 | 0.081048 | ZBTB8A | NA | 0.557763939 | 0.002503764 | CROT | purple | 0.201665 | 0.323191 |
| ZNF192 | pink | 0.346854 | 0.082565 | NUPR1 | NA | 0.559234424 | 0.0024249 | MFN2 | purple | 0.201832 | 0.322782 |
| S100A4 | pink | -0.346738 | 0.082675 | SLFN5 | NA | 0.560165495 | 0.002376075 | PRMT7 | purple | 0.203798 | 0.317991 |
| IAH1 | pink | -0.346628 | 0.082779 | MCU | orange | -0.475183644 | 0.012251008 | MON1A | purple | 0.210115 | 0.302893 |
| ZNF430 | pink | 0.344996 | 0.084335 | MED1 | orange | -0.44417938 | 0.020282028 | KBTBD2 | purple | 0.210255 | 0.302566 |
| COX14 | pink | -0.344102 | 0.085195 | FAM105B | orange | -0.421978025 | 0.028345747 | ILF3 | purple | 0.210997 | 0.300822 |
| ZKSCAN5 | pink | 0.34133 | 0.087908 | PPP1R15B | orange | -0.39911313 | 0.039177988 | USP36 | purple | 0.212913 | 0.296351 |
| SNIP1 | pink | 0.340302 | 0.088931 | SMYD2 | orange | -0.394524338 | 0.041704712 | PACSIN2 | purple | 0.21779 | 0.285162 |
| HSPA2 | pink | -0.340063 | 0.08917 | TMPPE | orange | -0.39309195 | 0.042519383 | RNF7 | purple | 0.220342 | 0.279415 |
| HINT1 | pink | -0.335182 | 0.094157 | PSD4 | orange | -0.390958844 | 0.043755934 | LPIN1 | purple | 0.220895 | 0.278179 |
| DSCR3 | pink | 0.334029 | 0.095365 | WDFY4 | orange | -0.382212147 | 0.04912713 | RAB1B | purple | 0.222666 | 0.274245 |
| OSBPL5 | pink | 0.333502 | 0.095922 | EP300 | orange | -0.376988082 | 0.052574574 | LOC253039 | purple | 0.223424 | 0.272574 |
| TPT1-AS1 | pink | 0.328974 | 0.100799 | GMFB | orange | -0.375913178 | 0.053306895 | BMP1 | purple | 0.22447 | 0.270276 |
| OGT | pink | 0.327581 | 0.102336 | KIAA1715 | orange | -0.358538927 | 0.066288453 | AKAP8 | purple | 0.227562 | 0.263558 |
| STAU1 | pink | 0.327233 | 0.102724 | NSF | orange | -0.356264107 | 0.068154716 | CD82 | purple | 0.230769 | 0.256704 |
| STARD9 | pink | 0.327051 | 0.102927 | ARID1A | orange | -0.330580084 | 0.092144642 | ZKSCAN1 | purple | 0.231019 | 0.256174 |
| MTERFD1 | pink | -0.326493 | 0.10355 | LRP1 | orange | -0.330027857 | 0.092722359 | URB2 | purple | 0.231659 | 0.254822 |
| GIMAP6 | pink | -0.32638 | 0.103676 | ADAM9 | orange | -0.326843075 | 0.096107523 | ASL | purple | 0.235204 | 0.247418 |
| SLAIN2 | pink | 0.324536 | 0.105758 | PHF2 | orange | -0.300666496 | 0.127535505 | COQ9 | purple | 0.235277 | 0.247268 |
| LDHD | pink | -0.323336 | 0.107129 | ZNF710 | orange | -0.283528478 | 0.151815434 | TOMM40L | purple | 0.239922 | 0.237786 |
| NDUFB10 | pink | -0.322314 | 0.108307 | SGK196 | orange | -0.282902286 | 0.152760997 | EDF1 | purple | 0.242206 | 0.233213 |
| CSNK1A1 | pink | 0.321387 | 0.109383 | TRIM33 | orange | -0.270609499 | 0.172182644 | EEF2 | purple | 0.243729 | 0.230195 |
| ZNF805 | pink | 0.320778 | 0.110096 | ATP11A | orange | -0.249532218 | 0.209397403 | SELO | purple | 0.246711 | 0.224364 |
| TMEM251 | pink | -0.320594 | 0.110311 | KAT2B | orange | -0.242631276 | 0.222686585 | TMEM229B | purple | 0.248599 | 0.220724 |
| PGK1 | pink | -0.320235 | 0.110734 | GNPTAB | orange | -0.238765773 | 0.230372614 | KAT7 | purple | 0.250893 | 0.216354 |
| KCTD5 | pink | 0.319009 | 0.112182 | POLR1A | orange | -0.22551071 | 0.258061386 | CACTIN | purple | 0.251061 | 0.216037 |
| C11orf75 | pink | -0.31804 | 0.113339 | CEP250 | orange | -0.183404789 | 0.359821317 | MED18 | purple | 0.255032 | 0.208618 |
| DCAF16 | pink | 0.315632 | 0.116248 | KLC2 | orange | -0.155227933 | 0.439448215 | APRT | purple | 0.256974 | 0.205053 |

**Table 6. Gene module membership and correlation with growth rate (48/72h), with adjusted p values for each module**

| | | | |
|---|---|---|---|
| HNRNPAB | pink | 0.315023 | 0.116993 |
| TM6SF1 | pink | -0.315001 | 0.117019 |
| STAG1 | pink | 0.312731 | 0.119825 |
| DENND5A | pink | 0.31159 | 0.121254 |
| SUB1 | pink | -0.310648 | 0.122443 |
| PSMD3 | pink | 0.310042 | 0.123212 |
| SEC22C | pink | 0.309397 | 0.124036 |
| BAP1 | pink | 0.306863 | 0.127307 |
| ZMYM5 | pink | 0.306007 | 0.128426 |
| PIGB | pink | -0.300488 | 0.135816 |
| POU2F1 | pink | -0.300466 | 0.135845 |
| MLL2 | pink | 0.300404 | 0.13593 |
| PSEN1 | pink | 0.299798 | 0.13676 |
| CSTF1 | pink | -0.29886 | 0.138052 |
| SLC35E1 | pink | 0.298354 | 0.138753 |
| C15orf61 | pink | -0.297672 | 0.1397 |
| YIPF6 | pink | 0.297531 | 0.139898 |
| KLHL20 | pink | 0.29563 | 0.142569 |
| FAM127B | pink | -0.295406 | 0.142886 |
| CBX7 | pink | -0.295122 | 0.143289 |
| PI4KAP2 | pink | 0.291922 | 0.147884 |
| POC1B | pink | 0.291406 | 0.148634 |
| WDR20 | pink | 0.288199 | 0.15336 |
| PACSIN2 | pink | 0.28717 | 0.154899 |
| STT3A | pink | 0.284755 | 0.158554 |
| CDR2 | pink | 0.284212 | 0.159385 |
| URM1 | pink | -0.283949 | 0.159789 |
| DLG1 | pink | -0.283736 | 0.160116 |
| RDH10 | pink | -0.2814 | 0.163733 |
| BAZ2A | pink | 0.279961 | 0.16599 |
| BBS9 | pink | -0.278913 | 0.167648 |
| MBD1 | pink | 0.278033 | 0.169049 |
| SKIV2L | pink | -0.27757 | 0.169788 |
| TMEM129 | pink | -0.276961 | 0.170766 |
| CDK10 | pink | -0.27593 | 0.17243 |
| CHMP1B | pink | 0.275433 | 0.173235 |
| VPS39 | pink | 0.275265 | 0.173508 |
| LATS1 | pink | 0.271275 | 0.180081 |
| UBE2C | pink | -0.269297 | 0.183402 |
| STARD8 | pink | 0.266778 | 0.187693 |
| OC1001348 | pink | 0.264124 | 0.192289 |
| EDEM3 | pink | 0.262029 | 0.195972 |
| IDI1 | pink | -0.260582 | 0.198542 |
| SF3B3 | pink | 0.260377 | 0.198909 |
| PEX7 | pink | -0.257557 | 0.203992 |
| URGCP | pink | 0.255617 | 0.207539 |
| DDA1 | pink | 0.254935 | 0.208797 |
| FRRS1 | pink | 0.254852 | 0.20895 |
| C17orf80 | pink | -0.254104 | 0.210335 |
| DCUN1D2 | pink | 0.254012 | 0.210506 |
| ZNF836 | pink | 0.253966 | 0.210593 |
| TUG1 | pink | 0.253377 | 0.211689 |
| YIPF1 | pink | -0.253231 | 0.211961 |
| CCDC22 | pink | -0.251103 | 0.215957 |
| SLC25A33 | pink | 0.250827 | 0.216478 |
| SRM | pink | -0.250353 | 0.217377 |
| UBXN4 | orange | -0.148610185 | 0.459430755 |
| CAMSAP1 | orange | -0.135079724 | 0.501729254 |
| UBQLN4 | orange | -0.114739437 | 0.568759363 |
| DDB2 | orange | -0.054865243 | 0.785773678 |
| MTX1 | orange | -0.02208544 | 0.912931155 |
| DPYD | orange | -0.013074711 | 0.948392142 |
| GFPT1 | orange | -0.012520581 | 0.950576689 |
| RPL18A | orange | 0.107352289 | 0.594055751 |
| USF1 | orange | 0.161706464 | 0.420348948 |
| RPL21 | orange | 0.172726584 | 0.388937023 |
| MNAT1 | orange | 0.180471635 | 0.36768783 |
| ISCA2 | orange | 0.207098187 | 0.299976185 |
| MRPL20 | orange | 0.215666148 | 0.279970911 |
| TCEB1 | orange | 0.228786531 | 0.251025638 |
| SOWAHD | orange | 0.230830254 | 0.246700425 |
| COQ7 | orange | 0.240239082 | 0.227422545 |
| DIS3L | orange | 0.257649644 | 0.194468557 |
| HCST | orange | 0.265288224 | 0.181105178 |
| SEL1L | orange | 0.27113497 | 0.17131861 |
| TIFA | orange | 0.275971558 | 0.16350854 |
| ANXA7 | orange | 0.280545778 | 0.156357055 |
| THRAP3 | orange | 0.284758819 | 0.149969788 |
| LPIN1 | orange | 0.28571599 | 0.148545073 |
| TIA1 | orange | 0.304157518 | 0.122959125 |
| RNF130 | orange | 0.304979902 | 0.121898805 |
| ZNF675 | orange | 0.33152569 | 0.091161694 |
| CHMP5 | orange | 0.33723654 | 0.085392599 |
| COX7A2L | orange | 0.348384702 | 0.074933656 |
| TMEM120A | orange | 0.358171855 | 0.066586896 |
| KTI12 | orange | 0.401390804 | 0.037969951 |
| ZNF518B | orange | 0.405461917 | 0.035884863 |
| MAX | orange | 0.40679573 | 0.035222012 |
| PDCL3 | orange | 0.413570721 | 0.032004893 |
| FAM100B | orange | 0.439454761 | 0.021817987 |
| CASP3 | orange | 0.441671401 | 0.02108595 |
| GIMAP4 | orange | 0.470382879 | 0.013284753 |
| LSM2 | orange | 0.47382035 | 0.012537526 |
| CLPP | orange | 0.500535293 | 0.007835722 |
| SUPT16H | orange | 0.576098277 | 0.00166207 |
| FLYWCH2 | pink | -0.624866848 | 0.000493153 |
| GUK1 | pink | -0.520146776 | 0.005416915 |
| SURF2 | pink | -0.514070317 | 0.006087292 |
| DENND1C | pink | -0.416574042 | 0.030656507 |
| SYNE3 | pink | -0.410387191 | 0.03348583 |
| USP34 | pink | -0.383259208 | 0.048458077 |
| MAP3K14 | pink | -0.37955257 | 0.050859271 |
| NUDT9 | pink | -0.376969245 | 0.052587339 |
| CALR | pink | -0.355834499 | 0.068511671 |
| TNFRSF9 | pink | -0.351470635 | 0.072219758 |
| NCBP1 | pink | -0.346399594 | 0.076720302 |
| TRAF7 | pink | -0.345937568 | 0.077140772 |
| EIF4G2 | pink | -0.331913636 | 0.090760723 |
| ARHGEF1 | pink | -0.32916802 | 0.093627311 |
| TMEM132A | pink | -0.328974 | 0.093832427 |
| C18orf8 | pink | -0.328370791 | 0.094472293 |
| SLC27A1 | pink | -0.327663837 | 0.095226378 |
| C11orf30 | purple | 0.259274 | 0.200886 |
| MOAP1 | purple | 0.260478 | 0.198728 |
| NDUFS5 | purple | 0.260762 | 0.198222 |
| PGRMC1 | purple | 0.263827 | 0.19281 |
| ELL | purple | 0.266296 | 0.188524 |
| CSNK1A1 | purple | 0.268054 | 0.185511 |
| CDC42SE1 | purple | 0.269464 | 0.183121 |
| MLF2 | purple | 0.272004 | 0.178868 |
| MVD | purple | 0.275436 | 0.17323 |
| PSMD3 | purple | 0.285755 | 0.157033 |
| MRPL46 | purple | 0.286623 | 0.155723 |
| HIRIP3 | purple | 0.28846 | 0.152972 |
| LEPROTL1 | purple | 0.288927 | 0.152279 |
| MMP15 | purple | 0.295577 | 0.142643 |
| WIPI2 | purple | 0.298911 | 0.137981 |
| CEBPA | purple | 0.300581 | 0.135688 |
| NOSIP | purple | 0.306875 | 0.127292 |
| MAEA | purple | 0.307685 | 0.126239 |
| FXC1 | purple | 0.31008 | 0.123165 |
| NFE2L1 | purple | 0.311016 | 0.121978 |
| YPEL5 | purple | 0.311641 | 0.121191 |
| ZNF618 | purple | 0.317123 | 0.11444 |
| LZTR1 | purple | 0.318034 | 0.113346 |
| FAM193B | purple | 0.322464 | 0.108134 |
| ABI1 | purple | 0.323386 | 0.107072 |
| IDH3G | purple | 0.326545 | 0.103491 |
| MPI | purple | 0.326946 | 0.103043 |
| TUBGCP4 | purple | 0.32933 | 0.100409 |
| CTNNBL1 | purple | 0.343425 | 0.085852 |
| SNIP1 | purple | 0.358954 | 0.071725 |
| ZNF410 | purple | 0.3608 | 0.070172 |
| SLC35C2 | purple | 0.363912 | 0.067613 |
| BAZ2A | purple | 0.36635 | 0.065659 |
| OSBPL5 | purple | 0.373596 | 0.060107 |
| ELMOD3 | purple | 0.392109 | 0.047569 |
| CASC3 | purple | 0.39328 | 0.046852 |
| TAB1 | purple | 0.402829 | 0.041312 |
| PSMB6 | purple | 0.409261 | 0.037883 |
| SMAD7 | purple | 0.435546 | 0.026148 |
| CHDH | purple | 0.456718 | 0.019004 |
| ZNF629 | purple | 0.458704 | 0.018426 |
| RINT1 | red | -0.379539 | 0.055831 |
| SACS | red | -0.373288 | 0.060335 |
| DNAJC9 | red | -0.365358 | 0.066449 |
| KIAA0430 | red | -0.339382 | 0.089853 |
| NTPCR | red | -0.333291 | 0.096145 |
| PEX19 | red | -0.330442 | 0.099198 |
| MRPL17 | red | -0.326657 | 0.103366 |
| METTL8 | red | -0.31878 | 0.112455 |
| TFG | red | -0.31479 | 0.117279 |
| RANBP2 | red | -0.299313 | 0.137427 |
| OXNAD1 | red | -0.29788 | 0.139411 |
| SNUPN | red | -0.284093 | 0.159567 |
| SEC61G | red | -0.272559 | 0.177948 |
| RIOK3 | red | -0.270145 | 0.181973 |
| CCDC6 | red | -0.26612 | 0.188826 |

**Table 6. Gene module membership and correlation with growth rate (48/72h), with adjusted p values for each module**

| | | | | | | | | | | | |
|---|---|---|---|---|---|---|---|---|---|---|---|
| CYB5B | pink | 0.248763 | 0.220409 | CDA | pink | -0.322515819 | 0.100854669 | GLMN | red | -0.265233 | 0.19036 |
| TOR1AIP1 | pink | 0.24653 | 0.224714 | IL4I1 | pink | -0.310953393 | 0.114397469 | REXO2 | red | -0.25973 | 0.200067 |
| FEM1B | pink | 0.245885 | 0.22597 | CNST | pink | -0.304594323 | 0.122395104 | PPP2R5B | red | -0.257254 | 0.204544 |
| RMND1 | pink | -0.245636 | 0.226456 | C2orf68 | pink | -0.303391156 | 0.123953285 | STYXL1 | red | -0.250509 | 0.217082 |
| CEPT1 | pink | -0.24537 | 0.226975 | LARP7 | pink | -0.298351024 | 0.130638653 | ZCRB1 | red | -0.250487 | 0.217123 |
| PNISR | pink | 0.244013 | 0.229635 | TRAPPC12 | pink | -0.292619358 | 0.138555895 | HSPH1 | red | -0.250025 | 0.218 |
| MAP3K2 | pink | 0.243924 | 0.22981 | BAZ1B | pink | -0.284832865 | 0.149859225 | TRUB1 | red | -0.248719 | 0.220493 |
| MSTO1 | pink | 0.241473 | 0.234674 | NDUFS2 | pink | -0.281357591 | 0.155111482 | TCEAL8 | red | -0.248675 | 0.220579 |
| SNORA64 | pink | -0.241145 | 0.235329 | RACGAP1 | pink | -0.280505495 | 0.156419046 | SSBP1 | red | -0.24825 | 0.221393 |
| FAM129A | pink | 0.24008 | 0.237467 | TDRD7 | pink | -0.277063361 | 0.16178094 | RRN3P2 | red | -0.248 | 0.221874 |
| ZNF526 | pink | 0.240043 | 0.237541 | CHTF8 | pink | -0.272594464 | 0.16893475 | ANKRD28 | red | -0.247021 | 0.223764 |
| PRPF8 | pink | 0.239539 | 0.238557 | ACP6 | pink | -0.27257227 | 0.168970826 | ZMAT2 | red | -0.246191 | 0.225374 |
| CHD4 | pink | 0.237692 | 0.242306 | TTLL5 | pink | -0.270120954 | 0.172988701 | POM121 | red | -0.245559 | 0.226604 |
| ING3 | pink | 0.236468 | 0.244812 | PAPD5 | pink | -0.267953524 | 0.176596691 | SUMO1P3 | red | -0.244929 | 0.227837 |
| BAGE2 | pink | 0.235222 | 0.247381 | ZC2HC1A | pink | -0.267292189 | 0.177707976 | ABHD10 | red | -0.244862 | 0.227968 |
| DCAF5 | pink | 0.233005 | 0.251995 | RBM10 | pink | -0.266886883 | 0.178391449 | SAMM50 | red | -0.244496 | 0.228686 |
| ALDH18A1 | pink | 0.230054 | 0.258221 | RCN3 | pink | -0.260362019 | 0.189647873 | GTF3C1 | red | -0.240411 | 0.236801 |
| DNM1L | pink | 0.227555 | 0.263573 | MST4 | pink | -0.259307514 | 0.191512119 | EXOSC10 | red | -0.236515 | 0.244716 |
| THG1L | pink | 0.226664 | 0.265497 | THEMIS2 | pink | -0.253914807 | 0.201243539 | LINC00189 | red | -0.225975 | 0.266993 |
| ATP11B | pink | 0.226395 | 0.266081 | HOOK2 | pink | -0.252880761 | 0.203147498 | FAM58A | red | -0.224093 | 0.271101 |
| ZNF432 | pink | 0.225059 | 0.268988 | DNAJA3 | pink | -0.249282551 | 0.20986857 | LYPLAL1 | red | -0.223954 | 0.271407 |
| ANKHD1 | pink | 0.224596 | 0.27 | POFUT1 | pink | -0.245834187 | 0.216450008 | LGALS1 | red | -0.220727 | 0.278555 |
| LDHB | pink | -0.221944 | 0.275846 | PABPC4 | pink | -0.24472672 | 0.21859293 | SPP1 | red | -0.220626 | 0.278781 |
| ILF3 | pink | 0.221693 | 0.276401 | CASKIN2 | pink | -0.243205599 | 0.221559503 | LYRM4 | red | -0.218145 | 0.284359 |
| NUMA1 | pink | 0.220687 | 0.278644 | NDOR1 | pink | -0.24233292 | 0.223273612 | FOCAD | red | -0.216098 | 0.289013 |
| LMAN2 | pink | 0.220675 | 0.27867 | C11orf24 | pink | -0.242166651 | 0.223601203 | HMBS | red | -0.215768 | 0.289769 |
| GID8 | pink | -0.220298 | 0.279513 | S100PBP | pink | -0.240861529 | 0.226183821 | ZMPSTE24 | red | -0.211285 | 0.300148 |
| HIF1AN | pink | 0.220002 | 0.280176 | TRIM24 | pink | -0.239964118 | 0.227971188 | JKAMP | red | -0.207717 | 0.308572 |
| SLC46A3 | pink | -0.218398 | 0.283787 | WWC3 | pink | -0.238869858 | 0.230163368 | SMARCB1 | red | -0.204594 | 0.316064 |
| MON1A | pink | 0.21792 | 0.284867 | CPSF1 | pink | -0.23829074 | 0.231329212 | VNN2 | red | -0.204156 | 0.317123 |
| C10orf118 | pink | 0.21763 | 0.285524 | PTDSS1 | pink | -0.232619614 | 0.242954005 | FNTB | red | -0.202552 | 0.321024 |
| PLEKHF2 | pink | 0.214671 | 0.292288 | UBE2K | pink | -0.229554961 | 0.249393592 | DDX6 | red | -0.200813 | 0.325284 |
| B4GALT4 | pink | 0.212669 | 0.296919 | NUMBL | pink | -0.223900878 | 0.261565511 | AKR1A1 | red | -0.193627 | 0.34325 |
| TTLL3 | pink | -0.211634 | 0.299332 | WIPF1 | pink | -0.220561021 | 0.268933378 | GMEB1 | red | -0.189764 | 0.35315 |
| SATB2 | pink | 0.209062 | 0.305379 | GDAP1 | pink | -0.220494315 | 0.269081882 | PABPC4 | red | -0.189645 | 0.353458 |
| ILKAP | pink | 0.207899 | 0.308139 | PRR14 | pink | -0.220122715 | 0.269910124 | C18orf32 | red | -0.188151 | 0.357333 |
| ASCC2 | pink | 0.205345 | 0.314253 | CHKA | pink | -0.219236418 | 0.271892168 | RBBP7 | red | -0.187188 | 0.359843 |
| BRAP | pink | 0.204287 | 0.316807 | ARHGEF35 | pink | -0.217047117 | 0.276828121 | TADA2A | red | -0.185343 | 0.364685 |
| TTC39C | pink | -0.203419 | 0.318912 | INTS2 | pink | -0.21536953 | 0.280648905 | GCC2 | red | -0.182678 | 0.371741 |
| DENND4A | pink | 0.202913 | 0.320142 | CTIF | pink | -0.214318062 | 0.28306072 | MTDH | red | -0.182615 | 0.371908 |
| RWDD1 | pink | -0.201996 | 0.322382 | PGAM1 | pink | -0.214152699 | 0.283441221 | IRAK4 | red | -0.180511 | 0.377537 |
| EYA3 | pink | 0.200542 | 0.325952 | MAT2A | pink | -0.213066045 | 0.285949684 | EXD2 | red | -0.180046 | 0.378789 |
| DNTTIP2 | pink | 0.199931 | 0.327457 | PKN2 | pink | -0.213019954 | 0.286056391 | FNTA | red | -0.176009 | 0.389747 |
| SLC25A32 | pink | 0.199324 | 0.328959 | MREG | pink | -0.211592854 | 0.289372849 | TOMM6 | red | -0.175728 | 0.390517 |
| SHB | pink | 0.198304 | 0.331491 | MUM1 | pink | -0.20877211 | 0.295999202 | UBE2H | red | -0.175083 | 0.392284 |
| SETD2 | pink | 0.197992 | 0.332268 | KLHDC10 | pink | -0.207246897 | 0.299621526 | ANP32B | red | -0.173653 | 0.396223 |
| NGDN | pink | 0.197711 | 0.332967 | PDPK1 | pink | -0.205776735 | 0.303139252 | P4HB | red | -0.173264 | 0.397299 |
| MUL1 | pink | 0.192157 | 0.346998 | RNF187 | pink | -0.203108358 | 0.309589525 | USP10 | red | -0.169007 | 0.409172 |
| SLBP | pink | 0.190391 | 0.351532 | SREBF1 | pink | -0.202856153 | 0.310203548 | SEC22A | red | -0.16758 | 0.413199 |
| STK32C | pink | 0.189406 | 0.354075 | ANKRD44 | pink | -0.202405296 | 0.311303093 | NIT2 | red | -0.16492 | 0.420758 |
| ZNF671 | pink | -0.188518 | 0.356379 | SACS | pink | -0.202202825 | 0.31179766 | RPN2 | red | -0.162005 | 0.42913 |
| SMAD4 | pink | 0.18729 | 0.359576 | CD86 | pink | -0.200387184 | 0.316254355 | HGSNAT | red | -0.161791 | 0.429746 |
| ABI1 | pink | 0.185106 | 0.365308 | FUCA1 | pink | -0.199690436 | 0.317974965 | USP39 | red | -0.157407 | 0.442516 |
| RBM3 | pink | 0.184333 | 0.367349 | TAF10 | pink | -0.199247305 | 0.31907226 | HTATSF1 | red | -0.15278 | 0.456208 |
| MED31 | pink | 0.183917 | 0.368451 | GPR107 | pink | -0.199145278 | 0.319325231 | ATP8A1 | red | -0.15031 | 0.463608 |
| DPM1 | pink | 0.18353 | 0.369475 | LIX1L | pink | -0.198757385 | 0.320288117 | FAM221A | red | -0.150128 | 0.464156 |

**Table 6. Gene module membership and correlation with growth rate (48/72h), with adjusted p values for each module**

| | | | | | | | | | | | |
|---|---|---|---|---|---|---|---|---|---|---|---|
| PGS1 | pink | 0.183459 | 0.369663 | ZC3H7B | pink | -0.198469553 | 0.321003768 | ACTR1A | red | -0.148781 | 0.468221 |
| GTPBP2 | pink | 0.181976 | 0.373614 | MTDH | pink | -0.198272204 | 0.321495012 | RPL7L1 | red | -0.147614 | 0.471756 |
| NANS | pink | 0.181365 | 0.375247 | ARHGEF3 | pink | -0.198271251 | 0.321497385 | RAB8A | red | -0.145949 | 0.476824 |
| MATR3 | pink | 0.179876 | 0.379246 | DDX19B | pink | -0.194238989 | 0.331635314 | PDPK1 | red | -0.144405 | 0.481548 |
| UAP1 | pink | -0.178804 | 0.382141 | RNF14 | pink | -0.193957943 | 0.33234907 | PDIA5 | red | -0.143784 | 0.483455 |
| BRAF | pink | 0.177878 | 0.384652 | PKD1P1 | pink | -0.192529372 | 0.335991512 | CDC42 | red | -0.138574 | 0.499605 |
| LNX2 | pink | 0.174267 | 0.394529 | ILF2 | pink | -0.188825954 | 0.345545853 | TAF2 | red | -0.137458 | 0.5031 |
| ACVR2B | pink | 0.172484 | 0.399461 | TFAP4 | pink | -0.187978413 | 0.347755019 | MRPL16 | red | -0.137181 | 0.503971 |
| FNIP1 | pink | 0.172364 | 0.399792 | PARP3 | pink | -0.18738488 | 0.349307103 | COPG2 | red | -0.135391 | 0.509603 |
| ARAP1 | pink | -0.167415 | 0.413663 | DIAPH1 | pink | -0.186923791 | 0.350515693 | HYAL2 | red | -0.134234 | 0.51326 |
| CHST11 | pink | 0.166751 | 0.415547 | ANKRD28 | pink | -0.186594449 | 0.351380474 | CD86 | red | -0.133918 | 0.514263 |
| PSMA5 | pink | 0.164619 | 0.421618 | SLC7A11 | pink | -0.183801264 | 0.358765667 | TRAPPC12 | red | -0.130722 | 0.524444 |
| SUDS3 | pink | -0.16286 | 0.426665 | ATP2A3 | pink | -0.17699313 | 0.377146026 | DNAJC5B | red | -0.127906 | 0.533492 |
| MTRF1L | pink | 0.160571 | 0.43328 | SPAG7 | pink | -0.174898421 | 0.38290887 | WDR41 | red | -0.127716 | 0.534105 |
| SIGLEC14 | pink | 0.154503 | 0.451082 | FOXN3 | pink | -0.17363133 | 0.386419251 | CTDP1 | red | -0.127102 | 0.536089 |
| MRPL12 | pink | -0.154257 | 0.451811 | MYC | pink | -0.172269274 | 0.390213205 | PRKAG1 | red | -0.126821 | 0.536998 |
| UBE2V2 | pink | 0.152464 | 0.457152 | PRKAG2 | pink | -0.171659722 | 0.391917944 | TRMT61B | red | -0.123763 | 0.546938 |
| CXorf40B | pink | -0.151981 | 0.458595 | GARS | pink | -0.170161652 | 0.396125585 | TRUB2 | red | -0.123608 | 0.547442 |
| ZNF14 | pink | 0.149158 | 0.467079 | IL6ST | pink | -0.167606747 | 0.403360306 | SH3BP5L | red | -0.121914 | 0.552987 |
| FUBP3 | pink | 0.147916 | 0.47084 | KDM5B | pink | -0.167157137 | 0.404641103 | SPCS1 | red | -0.121541 | 0.554211 |
| PDE8A | pink | 0.146141 | 0.476238 | ZDHHC6 | pink | -0.163007057 | 0.416570683 | FKRP | red | -0.12112 | 0.555595 |
| SMU1 | pink | 0.144831 | 0.480244 | TERF2IP | pink | -0.158541873 | 0.429620637 | NUDT9 | red | -0.120838 | 0.556522 |
| BZW1 | pink | 0.143324 | 0.484871 | PAPOLG | pink | -0.157195769 | 0.43359801 | ALKBH3 | red | -0.120184 | 0.558678 |
| DBP | pink | -0.139803 | 0.495774 | ZDHHC16 | pink | -0.156615655 | 0.435318229 | PPT2 | red | -0.118445 | 0.564423 |
| C19orf24 | pink | -0.138007 | 0.501378 | TOM1L2 | pink | -0.156288795 | 0.436289098 | GLIPR2 | red | -0.118134 | 0.565453 |
| COG4 | pink | 0.137192 | 0.503935 | MYO6 | pink | -0.154576836 | 0.441393188 | NAP1L4 | red | -0.115954 | 0.572699 |
| FBXO42 | pink | -0.130921 | 0.523807 | QKI | pink | -0.15320681 | 0.445500836 | CD97 | red | -0.114737 | 0.576763 |
| SLC25A30 | pink | 0.128749 | 0.530774 | C20orf96 | pink | -0.151955635 | 0.449269926 | TAF1C | red | -0.114457 | 0.577698 |
| TMEM185A | pink | 0.128054 | 0.533012 | PTER | pink | -0.151922958 | 0.449368592 | INTS2 | red | -0.114307 | 0.578201 |
| CPD | pink | 0.126621 | 0.537645 | ETNK1 | pink | -0.150916537 | 0.452413005 | UBE2M | red | -0.113517 | 0.580845 |
| TPRG1 | pink | 0.124586 | 0.544253 | NTPCR | pink | -0.149405311 | 0.457004911 | BHLHE40 | red | -0.108509 | 0.597749 |
| NOLC1 | pink | 0.119841 | 0.559808 | FAM70A | pink | -0.148920068 | 0.458484533 | ELAVL1 | red | -0.107966 | 0.599595 |
| MRFAP1 | pink | 0.11927 | 0.561695 | CD52 | pink | -0.148173798 | 0.460764994 | CRIP1 | red | -0.106808 | 0.603536 |
| ADTRP | pink | -0.116306 | 0.571526 | SAMM50 | pink | -0.145177458 | 0.469980874 | FAM185A | red | -0.105951 | 0.606462 |
| PSMB7 | pink | 0.113731 | 0.58013 | EEF1A1 | pink | -0.142590419 | 0.478014055 | UBE2E2 | red | -0.105385 | 0.608399 |
| NOP16 | pink | 0.108123 | 0.599059 | C16orf5 | pink | -0.14237302 | 0.478692307 | LYSMD3 | red | -0.104116 | 0.612743 |
| MOB3C | pink | 0.106482 | 0.604649 | G3BP1 | pink | -0.142363578 | 0.478721777 | AP4S1 | red | -0.098697 | 0.631454 |
| PDZD8 | pink | 0.105482 | 0.608064 | C10orf137 | pink | -0.141585925 | 0.481152047 | NUDT16 | red | -0.098232 | 0.633071 |
| ALKBH1 | pink | 0.105013 | 0.609671 | NDUFAF7 | pink | -0.141060364 | 0.482798058 | MAT2A | red | -0.098189 | 0.633218 |
| FADS3 | pink | 0.103662 | 0.614304 | BCAT2 | pink | -0.14073634 | 0.483814302 | ICK | red | -0.097727 | 0.634825 |
| CTNNB1 | pink | -0.102402 | 0.618636 | CREBBP | pink | -0.139618325 | 0.487329099 | PWWP2A | red | -0.096854 | 0.637867 |
| MASTL | pink | 0.097037 | 0.637232 | ERAL1 | pink | -0.138782551 | 0.489965017 | BPGM | red | -0.095457 | 0.642747 |
| MTHFD1 | pink | -0.096583 | 0.638815 | IRF4 | pink | -0.136181278 | 0.498214918 | TPR | red | -0.093651 | 0.649077 |
| PDHX | pink | -0.095474 | 0.642688 | ZSCAN29 | pink | -0.135396819 | 0.500716353 | KLHDC10 | red | -0.092805 | 0.652048 |
| HMGXB4 | pink | 0.093258 | 0.650456 | HN1 | pink | -0.135024033 | 0.501907255 | HCG18 | red | -0.092634 | 0.652652 |
| WBSCR22 | pink | 0.092572 | 0.652871 | PARP16 | pink | -0.134876492 | 0.502378978 | CPQ | red | -0.092328 | 0.653729 |
| PMS2CL | pink | 0.091425 | 0.656912 | CD3EAP | pink | -0.134310001 | 0.504192224 | STK11IP | red | -0.091453 | 0.656812 |
| GALNT1 | pink | 0.090712 | 0.659429 | MBNL1 | pink | -0.134001177 | 0.50518208 | KIF5B | red | -0.090237 | 0.661106 |
| THOC6 | pink | 0.089462 | 0.663847 | TAF9B | pink | -0.131993021 | 0.511642076 | MTHFS | red | -0.090001 | 0.66194 |
| ZSCAN29 | pink | 0.087687 | 0.670146 | EIF3F | pink | -0.131092323 | 0.514552597 | ZNF260 | red | -0.088535 | 0.667132 |
| SPAST | pink | 0.086442 | 0.674573 | SASH3 | pink | -0.129478472 | 0.519787721 | SNURF | red | -0.084356 | 0.682018 |
| NAGK | pink | 0.085608 | 0.677546 | TRMT10C | pink | -0.129039091 | 0.52121747 | VPS45 | red | -0.08075 | 0.694957 |
| GOSR1 | pink | 0.084873 | 0.680169 | TTYH2 | pink | -0.128487067 | 0.523016443 | MTMR9 | red | -0.079732 | 0.698623 |
| RAB4B | pink | 0.08121 | 0.693303 | KDM2B | pink | -0.126982527 | 0.527934686 | ZNF764 | red | -0.079283 | 0.700245 |
| MSH6 | pink | 0.081062 | 0.693836 | CUL7 | pink | -0.125596068 | 0.532486424 | STAMBP | red | -0.077538 | 0.706551 |
| PSMC1 | pink | 0.078697 | 0.70236 | TMEM138 | pink | -0.124857222 | 0.534919645 | SPAG7 | red | -0.077267 | 0.707532 |

**Table 6. Gene module membership and correlation with growth rate (48/72h), with adjusted p values for each module**

| | | | | | | | | | | | |
|---|---|---|---|---|---|---|---|---|---|---|---|
| ALS2 | pink | 0.075232 | 0.714918 | GEMIN8 | pink | -0.122285978 | 0.543428229 | SREBF2 | red | -0.077206 | 0.707753 |
| ANKRD17 | pink | 0.074805 | 0.71647 | RTN4 | pink | -0.119941238 | 0.551241963 | RHOT2 | red | -0.076751 | 0.709403 |
| LEPROT | pink | 0.069041 | 0.737527 | FAM198B | pink | -0.119868429 | 0.551485424 | PGM1 | red | -0.076444 | 0.710517 |
| KAL1 | pink | 0.068641 | 0.738998 | NRIP3 | pink | -0.11949597 | 0.55273163 | NISCH | red | -0.07592 | 0.712417 |
| UBXN6 | pink | 0.053849 | 0.793889 | RAB7A | pink | -0.118586741 | 0.555779248 | EIF3B | red | -0.072365 | 0.72536 |
| RNF139 | pink | 0.051557 | 0.802489 | UBE4B | pink | -0.117493874 | 0.55945255 | UCKL1 | red | -0.070067 | 0.733765 |
| ARHGAP25 | pink | 0.050957 | 0.804746 | C4orf34 | pink | -0.115546818 | 0.566024168 | UFD1L | red | -0.069877 | 0.734461 |
| SGSM2 | pink | -0.049862 | 0.808864 | TNKS2 | pink | -0.115437822 | 0.566393071 | ATP7A | red | -0.067301 | 0.743922 |
| ELMSAN1 | pink | 0.042926 | 0.835062 | PDRG1 | pink | -0.114411582 | 0.569871741 | ATP6V0D1 | red | -0.063785 | 0.756897 |
| UBE2O | pink | 0.033427 | 0.871223 | ZNF880 | pink | -0.114163097 | 0.57071547 | GCA | red | -0.062498 | 0.761658 |
| C12orf29 | pink | 0.019526 | 0.924571 | DIRC2 | pink | -0.114009505 | 0.571237273 | MAP3K14 | red | -0.062117 | 0.763071 |
| C19orf43 | pink | 0.017522 | 0.932297 | OGT | pink | -0.113334679 | 0.573532389 | ATXN7L1 | red | -0.059173 | 0.774009 |
| FAM49A | pink | -0.010354 | 0.959963 | ZNF25 | pink | -0.112694357 | 0.575713946 | AHCYL1 | red | -0.059078 | 0.77436 |
| SYNRG | pink | -0.004565 | 0.982341 | SLFN12 | pink | -0.112330611 | 0.57695485 | OSTC | red | -0.054647 | 0.790901 |
| UBIAD1 | pink | -0.004564 | 0.982346 | SMARCB1 | pink | -0.11162752 | 0.579356765 | MEX3C | red | -0.052752 | 0.798003 |
| LIN9 | purple | -0.509211 | 0.007887 | RPL32 | pink | -0.110800861 | 0.582186432 | TERF2IP | red | -0.051603 | 0.802318 |
| C20orf196 | purple | -0.43289 | 0.02718 | PLEKHG3 | pink | -0.109834611 | 0.585501581 | MRPL19 | red | -0.051327 | 0.803353 |
| NDEL1 | purple | 0.406832 | 0.03915 | ARHGAP30 | pink | -0.107797119 | 0.592518915 | SLFN12 | red | -0.050904 | 0.804943 |
| RTP4 | purple | -0.39488 | 0.045885 | THYN1 | pink | -0.105688261 | 0.599819897 | SNX17 | red | -0.050637 | 0.805949 |
| CORO6 | purple | -0.376161 | 0.058231 | TMEM101 | pink | -0.105264152 | 0.601292788 | QPRT | red | -0.050211 | 0.807549 |
| MTVR2 | purple | -0.375337 | 0.058828 | DUSP11 | pink | -0.105142774 | 0.601714607 | NIPSNAP3A | red | -0.048239 | 0.814976 |
| CCDC23 | purple | -0.375083 | 0.059013 | TP53BP2 | pink | -0.104505507 | 0.603931306 | ARMC6 | red | -0.04237 | 0.837173 |
| TNFAIP8L2 | purple | -0.362906 | 0.068432 | GTF3C2 | pink | -0.103334839 | 0.608012354 | TRMT5 | red | -0.039221 | 0.84913 |
| YME1L1 | purple | -0.361714 | 0.069413 | ZNF445 | pink | -0.103270656 | 0.608236436 | NUP153 | red | -0.034758 | 0.866137 |
| SNX20 | purple | -0.360227 | 0.070652 | SLC26A2 | pink | -0.101360556 | 0.614920861 | SDHB | red | -0.033856 | 0.869582 |
| CDCA5 | purple | -0.35152 | 0.078247 | VPS37B | pink | -0.098740683 | 0.624138077 | RSRC2 | red | -0.033384 | 0.871384 |
| CHMP4C | purple | -0.342814 | 0.086448 | PBX2 | pink | -0.09870583 | 0.624261074 | METTL7B | red | -0.033056 | 0.872638 |
| HIST1H2BB | purple | -0.337217 | 0.092053 | DSC2 | pink | -0.098333097 | 0.625577068 | DNAJB6 | red | -0.031948 | 0.876876 |
| CDKN1A | purple | 0.336936 | 0.092342 | SELO | pink | -0.098317754 | 0.625631263 | FGD5 | red | -0.030444 | 0.88263 |
| FAM83G | purple | 0.334362 | 0.095016 | ATG14 | pink | -0.098195879 | 0.626061818 | POLL | red | -0.027805 | 0.892744 |
| EGR2 | purple | 0.330765 | 0.098848 | FRG1 | pink | -0.096769802 | 0.631108693 | GTF3C2 | red | -0.026695 | 0.897004 |
| P2RY13 | purple | -0.327816 | 0.102076 | RNF5 | pink | -0.096024579 | 0.633752484 | DUSP1 | red | -0.025946 | 0.899878 |
| XIAP | purple | 0.327531 | 0.102392 | TRMT5 | pink | -0.094666874 | 0.638580431 | ADCK1 | red | -0.020734 | 0.919922 |
| THBD | purple | 0.317178 | 0.114373 | SSR1 | pink | -0.090989175 | 0.651730152 | EAF1 | red | -0.018825 | 0.927274 |
| WDR83OS | purple | -0.313566 | 0.118788 | PKM | pink | -0.090861926 | 0.652186988 | TRAF3IP2 | red | -0.018091 | 0.930101 |
| TRIB3 | purple | 0.30917 | 0.124326 | MAPK8IP3 | pink | -0.09067261 | 0.652866875 | PCNX | red | -0.01759 | 0.932032 |
| NUP107 | purple | -0.307644 | 0.126293 | PREP | pink | -0.09054454 | 0.653326965 | MGEA5 | red | -0.017521 | 0.932299 |
| BRAT1 | purple | -0.306957 | 0.127184 | KCTD12 | pink | -0.089697402 | 0.656373409 | DOLK | red | -0.016855 | 0.934867 |
| NBEAL2 | purple | -0.305016 | 0.129731 | YKT6 | pink | -0.088534316 | 0.660564796 | RC3H1 | red | -0.016541 | 0.936079 |
| HTRA4 | purple | 0.304573 | 0.130317 | ABI2 | pink | -0.087986375 | 0.662542878 | ZNF880 | red | -0.016516 | 0.936174 |
| LINC00094 | purple | -0.30208 | 0.133653 | FAM129B | pink | -0.087902697 | 0.66284515 | USP25 | red | -0.01511 | 0.941597 |
| PTPMT1 | purple | -0.298311 | 0.138812 | PCTP | pink | -0.086313854 | 0.668594348 | PPP6R3 | red | -0.01458 | 0.943643 |
| NIPAL3 | purple | -0.29671 | 0.141047 | UBXN7 | pink | -0.085141096 | 0.672849706 | EIF4A3 | red | -0.014174 | 0.945211 |
| CCDC71L | purple | -0.293681 | 0.145345 | PTPDC1 | pink | -0.083964887 | 0.677127482 | TNRC6A | red | -0.009663 | 0.962635 |
| GALK1 | purple | -0.293171 | 0.146077 | DCTN6 | pink | -0.081864783 | 0.684789675 | MAD2L2 | red | -0.009658 | 0.962651 |
| CXorf21 | purple | -0.289221 | 0.151843 | NOP14 | pink | -0.07883345 | 0.695903165 | YWHAH | red | -0.006981 | 0.972998 |
| THEMIS2 | purple | -0.288589 | 0.15278 | GORAB | pink | -0.078120199 | 0.698527122 | SCAF8 | red | -0.003914 | 0.984859 |
| ATF2 | purple | 0.282381 | 0.162207 | HIST1H2BK | pink | -0.076939559 | 0.702877973 | ARHGEF7 | red | -0.003752 | 0.985488 |
| LSM10 | purple | -0.279655 | 0.166474 | VIPAS39 | pink | -0.07469037 | 0.711191823 | SLC25A3 | red | 0.000109 | 0.999579 |
| ATP2B1 | purple | 0.273851 | 0.175818 | ACAD10 | pink | -0.072707373 | 0.718548604 | C1orf85 | red | 0.003482 | 0.986533 |
| C2orf43 | purple | -0.271962 | 0.178937 | GALNT11 | pink | -0.072117585 | 0.720741443 | STARD3NL | red | 0.003997 | 0.984538 |
| TP53INP2 | purple | 0.265743 | 0.189477 | RICTOR | pink | -0.071403402 | 0.72339967 | PDSS2 | red | 0.004373 | 0.983084 |
| MCFD2 | purple | 0.263908 | 0.192667 | MBOAT1 | pink | -0.070179524 | 0.727962288 | SLC7A11 | red | 0.005359 | 0.979273 |
| TNFRSF21 | purple | 0.256677 | 0.205597 | SPCS1 | pink | -0.068919429 | 0.73266939 | THAP2 | red | 0.006345 | 0.975458 |
| NDE1 | purple | -0.256427 | 0.206053 | UQCC | pink | -0.067470504 | 0.73809355 | FOXO3B | red | 0.006535 | 0.974724 |
| SENP3 | purple | -0.256064 | 0.206719 | RSRC2 | pink | -0.067194378 | 0.739128648 | OSTF1 | red | 0.013141 | 0.949197 |

**Table 6. Gene module membership and correlation with growth rate (48/72h), with adjusted p values for each module**

| | | | | | | | | | | | |
|---|---|---|---|---|---|---|---|---|---|---|---|
| TMEM170A | purple | 0.256062 | 0.206723 | C3orf37 | pink | -0.066267892 | 0.742604949 | ATP5G1 | red | 0.013441 | 0.948041 |
| CTSL1 | purple | 0.251355 | 0.215481 | PPIF | pink | -0.066010502 | 0.743571585 | BANP | red | 0.014174 | 0.945212 |
| CRY1 | purple | 0.251086 | 0.215989 | RUSC2 | pink | -0.060571968 | 0.764082841 | TRIM22 | red | 0.014621 | 0.943487 |
| RCN2 | purple | -0.25026 | 0.217555 | SFSWAP | pink | -0.059019277 | 0.769968064 | FOXRED2 | red | 0.016524 | 0.936145 |
| EIF2AK3 | purple | 0.248901 | 0.220145 | TOMM6 | pink | -0.058147682 | 0.773277181 | VPS28 | red | 0.021583 | 0.916652 |
| WIPF2 | purple | 0.245534 | 0.226653 | C1orf43 | pink | -0.057886179 | 0.77427077 | RBMS2 | red | 0.023193 | 0.91046 |
| PRIM1 | purple | -0.244561 | 0.228558 | LOC493754 | pink | -0.056625366 | 0.779066102 | ANKRD39 | red | 0.024095 | 0.906991 |
| SNX25 | purple | 0.242982 | 0.231671 | SMIM3 | pink | -0.055971004 | 0.781558005 | MINK1 | red | 0.024491 | 0.90547 |
| FAM133B | purple | 0.242055 | 0.233514 | USP25 | pink | -0.055164448 | 0.784632382 | TOX4 | red | 0.024569 | 0.905171 |
| STK40 | purple | 0.239929 | 0.237771 | MBTPS2 | pink | -0.052177476 | 0.796045084 | ELK4 | red | 0.027001 | 0.895831 |
| PAQR7 | purple | -0.239576 | 0.238483 | GRK5 | pink | -0.051884494 | 0.797166765 | KIAA0754 | red | 0.028559 | 0.889855 |
| OR52K1 | purple | -0.236943 | 0.243838 | PDIA3 | pink | -0.050370455 | 0.802969491 | OGT | red | 0.030964 | 0.880641 |
| FRAT2 | purple | -0.236854 | 0.244021 | PCOLCE2 | pink | -0.048277818 | 0.811006508 | CYC1 | red | 0.034616 | 0.866679 |
| GNA13 | purple | 0.236706 | 0.244324 | MSL2 | pink | -0.047374712 | 0.814480826 | TMEM107 | red | 0.036711 | 0.858688 |
| SET | purple | -0.233927 | 0.250068 | SLC38A6 | pink | -0.042877654 | 0.831831164 | ZNF224 | red | 0.037166 | 0.856955 |
| FAM105A | purple | -0.233233 | 0.251517 | HSPE1 | pink | -0.042443064 | 0.833512095 | ARIH2 | red | 0.037413 | 0.856013 |
| H2AFJ | purple | -0.232201 | 0.25368 | EIF2S1 | pink | -0.040736162 | 0.840121001 | CHTF8 | red | 0.037688 | 0.854968 |
| NRIP3 | purple | 0.231924 | 0.254264 | PDE4DIP | pink | -0.039515582 | 0.844853453 | LINC00263 | red | 0.0395 | 0.848073 |
| ZNF100 | purple | -0.224637 | 0.269909 | ZBTB40 | pink | -0.037944156 | 0.850953931 | IMP4 | red | 0.041519 | 0.840399 |
| TRIAP1 | purple | -0.223034 | 0.273434 | SYNJ1 | pink | -0.03653679 | 0.856424634 | PTPRC | red | 0.042432 | 0.836936 |
| SLC41A1 | purple | 0.222526 | 0.274555 | MED17 | pink | -0.033374234 | 0.868741387 | PRKCE | red | 0.043166 | 0.834154 |
| DAK | purple | -0.221063 | 0.277805 | CLCF1 | pink | -0.033333131 | 0.868901666 | FAM168B | red | 0.044022 | 0.830911 |
| TBC1D22A | purple | -0.21969 | 0.280877 | HSPA5 | pink | -0.031260991 | 0.876988403 | HCCS | red | 0.044911 | 0.827546 |
| SRSF8 | purple | -0.214092 | 0.293622 | PTGES2 | pink | -0.028389159 | 0.888215803 | EID1 | red | 0.046759 | 0.820563 |
| ISG20L2 | purple | 0.212063 | 0.298329 | UFD1L | pink | -0.028227522 | 0.888848369 | ZCCHC3 | red | 0.048832 | 0.812742 |
| ATXN1L | purple | 0.211573 | 0.299473 | LGALS3 | pink | -0.027212597 | 0.892821772 | USP5 | red | 0.049115 | 0.811674 |
| DNAJA3 | purple | -0.210482 | 0.30203 | INCENP | pink | -0.027111673 | 0.893217026 | ZFP112 | red | 0.051894 | 0.801223 |
| NRARP | purple | -0.209546 | 0.304236 | AGPAT9 | pink | -0.026878527 | 0.894130204 | APOA1BP | red | 0.052875 | 0.797543 |
| HEATR3 | purple | -0.208852 | 0.305875 | RNF40 | pink | -0.025987237 | 0.897622382 | WAPAL | red | 0.056797 | 0.782863 |
| TMOD2 | purple | -0.2086 | 0.306473 | RC3H1 | pink | -0.024910029 | 0.901845485 | GTF2F1 | red | 0.061037 | 0.767079 |
| SOX13 | purple | 0.206998 | 0.310288 | GPATCH3 | pink | -0.02278853 | 0.910170168 | THOC7 | red | 0.061133 | 0.766721 |
| C7orf25 | purple | -0.205684 | 0.313436 | ZNF295 | pink | -0.02178085 | 0.914127566 | PRELID1 | red | 0.061682 | 0.764686 |
| LINC00672 | purple | -0.204915 | 0.315288 | RNF13 | pink | -0.021036502 | 0.917052085 | FHL3 | red | 0.063916 | 0.75641 |
| ZNF649 | purple | -0.200205 | 0.326781 | SZRD1 | pink | -0.02073192 | 0.918249082 | ZNF264 | red | 0.064539 | 0.754106 |
| LAMTOR1 | purple | -0.19865 | 0.33063 | CEP170 | pink | -0.019751104 | 0.922104831 | RBM27 | red | 0.068267 | 0.74037 |
| STX7 | purple | -0.198518 | 0.330957 | USP12 | pink | -0.019412453 | 0.923436527 | VKORC1 | red | 0.069586 | 0.735528 |
| CISH | purple | -0.196855 | 0.335106 | POLR2G | pink | -0.01929878 | 0.923883577 | BIN2 | red | 0.070088 | 0.733689 |
| PARP4 | purple | -0.19661 | 0.335721 | HPGD | pink | -0.018774277 | 0.925946608 | KPNA4 | red | 0.072857 | 0.723564 |
| PLEKHM1 | purple | 0.196295 | 0.336511 | GDE1 | pink | -0.017448847 | 0.931161987 | NT5C2 | red | 0.073276 | 0.722036 |
| MLLT10 | purple | -0.195925 | 0.337442 | HIPK3 | pink | -0.017399416 | 0.931356543 | TMCC1 | red | 0.074166 | 0.718793 |
| SRXN1 | purple | 0.194649 | 0.34066 | ABHD10 | pink | -0.017274947 | 0.931846469 | LIG1 | red | 0.075886 | 0.712543 |
| NAPEPLD | purple | -0.191107 | 0.349689 | ATP10A | pink | -0.016887222 | 0.933372748 | MNDA | red | 0.081794 | 0.691202 |
| SNX32 | purple | -0.190499 | 0.351252 | EGR2 | pink | -0.015359441 | 0.939389056 | ZNF229 | red | 0.081914 | 0.69077 |
| BRPF3 | purple | 0.190461 | 0.351351 | RDBP | pink | -0.015326437 | 0.93951906 | FOXP1 | red | 0.083565 | 0.684848 |
| TAMM41 | purple | -0.189982 | 0.352588 | ATP5G3 | pink | -0.013724384 | 0.945831399 | KPNB1 | red | 0.088871 | 0.665943 |
| APH1B | purple | -0.187447 | 0.359168 | EFR3A | pink | -0.012983098 | 0.948753284 | CARKD | red | 0.09149 | 0.65668 |
| C16orf70 | purple | -0.186871 | 0.360672 | LARS | pink | -0.012829586 | 0.949358454 | ARID1B | red | 0.092558 | 0.652918 |
| ILF2 | purple | -0.186353 | 0.36203 | EIF3M | pink | -0.012813511 | 0.949421828 | PGLS | red | 0.092868 | 0.651827 |
| H2AFZ | purple | -0.184684 | 0.36642 | TRAFD1 | pink | -0.012501621 | 0.950651443 | DSCR3 | red | 0.095114 | 0.643949 |
| LYSMD3 | purple | 0.181927 | 0.373744 | ENO1 | pink | -0.011818493 | 0.953345033 | CLK2 | red | 0.096237 | 0.640023 |
| ZNF624 | purple | -0.181708 | 0.374329 | RAB33B | pink | -0.011734907 | 0.953674651 | C2CD3 | red | 0.097334 | 0.636197 |
| DUSP14 | purple | 0.180353 | 0.377963 | CELF1 | pink | -0.011328117 | 0.95527891 | C6orf120 | red | 0.09756 | 0.635408 |
| DNAJB9 | purple | 0.179152 | 0.3812 | DGAT2 | pink | -0.010543078 | 0.958375344 | NPRL2 | red | 0.100301 | 0.625891 |
| PPP1R3C | purple | -0.177724 | 0.385068 | TBC1D22A | pink | -0.010437068 | 0.958793525 | GPR180 | red | 0.100934 | 0.623702 |
| SCIMP | purple | -0.177644 | 0.385287 | ADPRHL2 | pink | -0.010321356 | 0.959249993 | FAM54B | red | 0.104104 | 0.612785 |
| DBR1 | purple | -0.173962 | 0.395371 | NHP2L1 | pink | -0.009813257 | 0.96125451 | YPEL2 | red | 0.106116 | 0.6059 |

**Table 6. Gene module membership and correlation with growth rate (48/72h), with adjusted p values for each module**

| | | | | | | | | | | | |
|---|---|---|---|---|---|---|---|---|---|---|---|
| VAMP4 | purple | -0.173626 | 0.396298 | ZNF780B | pink | -0.009524473 | 0.962393904 | ARMCX1 | red | 0.108056 | 0.599287 |
| SLC16A6 | purple | 0.172761 | 0.398691 | PPP2R5E | pink | -0.009342637 | 0.963111376 | RECQL5 | red | 0.108816 | 0.596706 |
| DNAJB11 | purple | 0.170378 | 0.405327 | NT5DC1 | pink | -0.008364757 | 0.966970262 | BBS12 | red | 0.114527 | 0.577464 |
| RRP1 | purple | 0.169797 | 0.406954 | ACSL5 | pink | -0.007639739 | 0.969831785 | SMAD2 | red | 0.115489 | 0.574249 |
| CNOT3 | purple | 0.167971 | 0.412093 | TNPO3 | pink | -0.007627447 | 0.969880301 | TOP3A | red | 0.118353 | 0.564728 |
| CTSL2 | purple | -0.166146 | 0.417265 | STARD8 | pink | -0.007065231 | 0.97209953 | BPTF | red | 0.120114 | 0.558908 |
| TIMM17B | purple | -0.163689 | 0.424283 | KIAA0391 | pink | -0.00669836 | 0.973547782 | CCDC107 | red | 0.122468 | 0.551171 |
| C22orf13 | purple | 0.162875 | 0.426622 | CCDC126 | pink | -0.005517966 | 0.978208023 | HOMER3 | red | 0.125373 | 0.541694 |
| CAMSAP3 | purple | -0.162363 | 0.428094 | TRIM22 | pink | -0.00511513 | 0.979798604 | NUDT17 | red | 0.126857 | 0.53688 |
| ZNF814 | purple | 0.16128 | 0.431223 | ACTR2 | pink | -0.003540205 | 0.986017789 | NUP133 | red | 0.128322 | 0.532148 |
| RRAGC | purple | -0.161176 | 0.431527 | NBEAL2 | pink | -0.003318445 | 0.986893567 | KLF6 | red | 0.135792 | 0.50834 |
| EPS8 | purple | 0.160116 | 0.4346 | STX7 | pink | -0.002199878 | 0.991311219 | RNF121 | red | 0.137439 | 0.503161 |
| NICN1 | purple | -0.157836 | 0.441255 | KLF6 | pink | -0.000975789 | 0.996145903 | HOXB7 | red | 0.139369 | 0.497124 |
| PPIF | purple | 0.155354 | 0.448562 | TP53RK | pink | 0.000499208 | 0.99802826 | RPUSD2 | red | 0.140034 | 0.495053 |
| PLEKHM1P | purple | 0.155145 | 0.44918 | NUP153 | pink | 0.001077222 | 0.995745273 | A2M | red | 0.143375 | 0.484715 |
| CCDC125 | purple | -0.154228 | 0.451898 | ACOT13 | pink | 0.002937723 | 0.988397146 | TMEM170A | red | 0.144378 | 0.481631 |
| ARG2 | purple | -0.153278 | 0.454722 | HLA-DRA | pink | 0.003475721 | 0.986272447 | CBX4 | red | 0.14458 | 0.481012 |
| SLC25A28 | purple | 0.151159 | 0.461056 | ZZZ3 | pink | 0.005868943 | 0.976822272 | NPLOC4 | red | 0.14971 | 0.465413 |
| PLK2 | purple | -0.150504 | 0.463024 | TCP1 | pink | 0.005919565 | 0.976622408 | FCHO2 | red | 0.157208 | 0.443098 |
| U2AF1 | purple | 0.149414 | 0.466308 | KIF5B | pink | 0.006270207 | 0.975238059 | CFLAR | red | 0.161768 | 0.429813 |
| NUDCD2 | purple | -0.14882 | 0.468102 | TNNT1 | pink | 0.007545495 | 0.970203778 | MPDU1 | red | 0.16748 | 0.41348 |
| C1GALT1C1 | purple | -0.146735 | 0.474428 | POLR1C | pink | 0.009317819 | 0.963209301 | PDCD10 | red | 0.16754 | 0.41331 |
| AGPAT6 | purple | 0.145497 | 0.478206 | FXYD6 | pink | 0.009588517 | 0.962141213 | NCOA2 | red | 0.17139 | 0.402503 |
| C12orf49 | purple | 0.145214 | 0.479071 | MUS81 | pink | 0.009995156 | 0.960536868 | ZNF44 | red | 0.174142 | 0.394875 |
| HIC2 | purple | 0.144544 | 0.481123 | ZSWIM6 | pink | 0.010636833 | 0.958005513 | ORAOV1 | red | 0.174287 | 0.394474 |
| VPS26B | purple | -0.143769 | 0.483502 | BZW1 | pink | 0.011281566 | 0.955462504 | AP4B1 | red | 0.178827 | 0.382078 |
| RNF169 | purple | -0.143593 | 0.484044 | SMAD2 | pink | 0.012617841 | 0.950193235 | ZNF23 | red | 0.18531 | 0.36477 |
| ZFAND5 | purple | -0.142421 | 0.487655 | SPRYD3 | pink | 0.01358432 | 0.94638343 | SMURF1 | red | 0.185894 | 0.363233 |
| MAML1 | purple | 0.139981 | 0.495217 | FAM76B | pink | 0.016568235 | 0.934628618 | RING1 | red | 0.186454 | 0.361764 |
| HIST1H3J | purple | -0.136646 | 0.50565 | POLR2J | pink | 0.016888549 | 0.933367523 | FAM96B | red | 0.190408 | 0.351487 |
| ZNF295 | purple | 0.135831 | 0.508216 | EVI5L | pink | 0.017404022 | 0.931338417 | UNC13B | red | 0.191151 | 0.349576 |
| PLEKHA3 | purple | -0.134834 | 0.511362 | SAP18 | pink | 0.019056106 | 0.924838027 | RBBP9 | red | 0.191275 | 0.349259 |
| TNFRSF12A | purple | 0.134725 | 0.511708 | B3GALNT1 | pink | 0.019961388 | 0.921278022 | FAM32A | red | 0.19186 | 0.347757 |
| MTIF3 | purple | -0.134034 | 0.513893 | PARP10 | pink | 0.020156153 | 0.920512303 | INTS4 | red | 0.197414 | 0.33371 |
| ALG2 | purple | -0.133515 | 0.515541 | ITPKC | pink | 0.020739056 | 0.918221038 | DUSP23 | red | 0.202646 | 0.320795 |
| FUK | purple | -0.133326 | 0.516142 | UBE2M | pink | 0.021117882 | 0.916732295 | VMAC | red | 0.202729 | 0.320592 |
| LINC00032 | purple | -0.132161 | 0.519848 | UBE2E2 | pink | 0.023517046 | 0.90731041 | RPS6KA2 | red | 0.203434 | 0.318875 |
| IL6ST | purple | -0.131921 | 0.520612 | KPNA4 | pink | 0.024145493 | 0.904844376 | UPRT | red | 0.204303 | 0.316769 |
| CLCF1 | purple | 0.131523 | 0.521882 | GLO1 | pink | 0.025832161 | 0.898230178 | EIF6 | red | 0.206825 | 0.3107 |
| TRIB1 | purple | 0.131079 | 0.523299 | PGGT1B | pink | 0.026355623 | 0.896178774 | DYNLT1 | red | 0.210058 | 0.303028 |
| RPL32P3 | purple | -0.130143 | 0.526296 | POLR2A | pink | 0.027774388 | 0.890622058 | ZKSCAN4 | red | 0.222412 | 0.274808 |
| SIAH2 | purple | 0.129571 | 0.528132 | ACAP2 | pink | 0.027964851 | 0.889876471 | CARM1 | red | 0.226537 | 0.265773 |
| ZNF79 | purple | -0.125687 | 0.540673 | HSBP1 | pink | 0.02831861 | 0.888491891 | ZNF460 | red | 0.227284 | 0.264156 |
| DALRD3 | purple | 0.124395 | 0.544875 | EIF1 | pink | 0.028844476 | 0.886434286 | SORBS3 | red | 0.232496 | 0.253062 |
| RPL7L1 | purple | -0.12437 | 0.544956 | PGLS | pink | 0.032314725 | 0.872874543 | PHB2 | red | 0.248478 | 0.220957 |
| TRMT5 | purple | 0.123827 | 0.546729 | C11orf10 | pink | 0.032752187 | 0.871167597 | ADD1 | red | 0.253534 | 0.211395 |
| CANT1 | purple | 0.119308 | 0.561566 | SKIV2L2 | pink | 0.032905551 | 0.870569314 | SHC1 | red | 0.253535 | 0.211393 |
| RGS2 | purple | -0.118027 | 0.565807 | TIMMDC1 | pink | 0.034786845 | 0.863236026 | TMEM9B | red | 0.257334 | 0.204399 |
| KLF6 | purple | 0.117416 | 0.567835 | LAMTOR5 | pink | 0.03645374 | 0.856747673 | NADSYN1 | red | 0.262223 | 0.195628 |
| HCG26 | purple | -0.116453 | 0.571037 | ANKMY2 | pink | 0.037072217 | 0.854342551 | GNA13 | red | 0.268324 | 0.185052 |
| HLA-DPA1 | purple | 0.115215 | 0.575163 | CIRBP | pink | 0.037230598 | 0.853726846 | MAN2A2 | red | 0.294898 | 0.143606 |
| NEDD4L | purple | 0.113943 | 0.57942 | INVS | pink | 0.037528858 | 0.852567588 | TRIM36 | red | 0.302004 | 0.133755 |
| TTC30A | purple | -0.113803 | 0.579887 | SPG21 | pink | 0.038019203 | 0.850662396 | NSMCE1 | red | 0.31124 | 0.121695 |
| SNX11 | purple | 0.11332 | 0.581509 | SDHA | pink | 0.039069731 | 0.84658343 | NGDN | red | 0.311353 | 0.121552 |
| OC1004994 | purple | -0.113288 | 0.581616 | NEMF | pink | 0.039751646 | 0.843937766 | RASA4 | red | 0.325304 | 0.104887 |
| MICAL1 | purple | -0.112888 | 0.582958 | GTF2A2 | pink | 0.042956456 | 0.831526446 | SPPL3 | red | 0.346384 | 0.08301 |

**Table 6. Gene module membership and correlation with growth rate (48/72h), with adjusted p values for each module**

| | | | | | | | | | | | |
|---|---|---|---|---|---|---|---|---|---|---|---|
| DUSP6 | purple | 0.110435 | 0.591225 | C16orf62 | pink | 0.043425058 | 0.829714912 | MYO1F | red | 0.374882 | 0.059161 |
| TFPI | purple | 0.109866 | 0.593149 | GTF2H2 | pink | 0.044016558 | 0.827429486 | SGPP1 | red | 0.384278 | 0.052594 |
| CKS2 | purple | -0.109137 | 0.595618 | GSR | pink | 0.044395837 | 0.825964755 | IMPACT | red | 0.38952 | 0.049187 |
| ARFGAP2 | purple | 0.108585 | 0.597491 | CLDN12 | pink | 0.044997937 | 0.823640689 | KIAA1958 | red | 0.407028 | 0.039046 |
| DANCR | purple | -0.108282 | 0.598521 | TCOF1 | pink | 0.050265186 | 0.803373328 | CWC15 | royalblue | -0.538583 | 0.004531 |
| SDF2L1 | purple | 0.107292 | 0.601889 | DCLRE1C | pink | 0.050619968 | 0.802012493 | SURF2 | royalblue | -0.521078 | 0.006342 |
| TIMM8A | purple | -0.102638 | 0.617827 | SC5DL | pink | 0.050854947 | 0.801111496 | METTL2B | royalblue | -0.471109 | 0.015129 |
| ATP1B2 | purple | -0.102577 | 0.618036 | USP14 | pink | 0.051750868 | 0.797678487 | TBCB | royalblue | -0.43985 | 0.024543 |
| BZW2 | purple | -0.099985 | 0.626988 | ACADVL | pink | 0.05208023 | 0.796417346 | COX7C | royalblue | -0.421017 | 0.032202 |
| GADD45A | purple | 0.099714 | 0.627926 | REEP5 | pink | 0.054316766 | 0.787866925 | TIMM13 | royalblue | -0.398674 | 0.043655 |
| SELK | purple | 0.099169 | 0.629814 | OSTC | pink | 0.054551758 | 0.786969908 | SERGEF | royalblue | -0.37889 | 0.056286 |
| HNRNPL | purple | -0.097583 | 0.635329 | PPARG | pink | 0.054941525 | 0.785482665 | HHLA3 | royalblue | -0.366029 | 0.065914 |
| ZNF175 | purple | -0.097537 | 0.635488 | MED7 | pink | 0.055746517 | 0.782413369 | NDUFA6 | royalblue | -0.341375 | 0.087864 |
| TNPO1 | purple | 0.095906 | 0.641178 | SEC31A | pink | 0.055775769 | 0.782301896 | HNRNPA1L2 | royalblue | -0.315491 | 0.11642 |
| RAB26 | purple | -0.095363 | 0.643075 | TFEC | pink | 0.057081713 | 0.777329525 | DYNLL1 | royalblue | -0.315369 | 0.116569 |
| MORF4L1 | purple | 0.093294 | 0.650331 | MKLN1 | pink | 0.05754029 | 0.775585515 | NASP | royalblue | -0.302467 | 0.133132 |
| FAM172A | purple | -0.092574 | 0.652861 | GAS2L3 | pink | 0.066148527 | 0.743053181 | SLC11A2 | royalblue | -0.298633 | 0.138366 |
| ITGAV | purple | 0.09216 | 0.654319 | CCDC107 | pink | 0.066228638 | 0.742752342 | MZT2A | royalblue | -0.279771 | 0.166291 |
| FAM217B | purple | -0.091306 | 0.65733 | EMC4 | pink | 0.066327035 | 0.742382886 | RALY | royalblue | -0.279435 | 0.16682 |
| SRP68 | purple | -0.09089 | 0.658799 | TBK1 | pink | 0.067336322 | 0.738596497 | RPL32 | royalblue | -0.273475 | 0.176436 |
| SDC2 | purple | -0.089398 | 0.664073 | RPS5 | pink | 0.068874907 | 0.732835878 | CTSB | royalblue | -0.266804 | 0.187649 |
| AVL9 | purple | 0.089391 | 0.6641 | ARL11 | pink | 0.069725153 | 0.729658495 | UFC1 | royalblue | -0.26029 | 0.199065 |
| GARS | purple | -0.089337 | 0.664289 | CERS5 | pink | 0.070149978 | 0.728072548 | OAZ1 | royalblue | -0.25954 | 0.200409 |
| CAHM | purple | -0.087588 | 0.670494 | PSME4 | pink | 0.070534597 | 0.726637634 | ARL6IP4 | royalblue | -0.253762 | 0.210971 |
| PXK | purple | 0.086796 | 0.673311 | SELK | pink | 0.074944212 | 0.710251889 | VPS51 | royalblue | -0.246271 | 0.225218 |
| FAM98A | purple | -0.084494 | 0.681526 | SENP7 | pink | 0.076449609 | 0.704686209 | MRPL18 | royalblue | -0.245458 | 0.226803 |
| NFKB1 | purple | -0.084417 | 0.681801 | MAML2 | pink | 0.081083449 | 0.687648196 | TRMT112 | royalblue | -0.235806 | 0.246174 |
| KIAA0355 | purple | -0.08178 | 0.691254 | FAM168B | pink | 0.082009565 | 0.684260454 | PNPO | royalblue | -0.232933 | 0.252146 |
| SLC4A1AP | purple | 0.080387 | 0.696265 | MTCH2 | pink | 0.085472042 | 0.671647864 | PSMB1 | royalblue | -0.224019 | 0.271264 |
| PHC1 | purple | -0.079882 | 0.698084 | MCTS1 | pink | 0.088270836 | 0.661515689 | C9orf123 | royalblue | -0.213867 | 0.294142 |
| CES4A | purple | -0.079791 | 0.698414 | ODC1 | pink | 0.088851999 | 0.659418972 | TBPL1 | royalblue | -0.204371 | 0.316604 |
| PPP1R15B | purple | -0.077961 | 0.705022 | TNFRSF10B | pink | 0.089764381 | 0.656132348 | ABHD8 | royalblue | -0.204066 | 0.317341 |
| MAGEH1 | purple | -0.077437 | 0.706919 | PLB1 | pink | 0.090960909 | 0.651831618 | SAP18 | royalblue | -0.202692 | 0.320682 |
| ACBD3 | purple | 0.077426 | 0.706959 | SEC61B | pink | 0.091291667 | 0.650644668 | CIB1 | royalblue | -0.199162 | 0.32936 |
| GAS5 | purple | -0.077134 | 0.708014 | SACM1L | pink | 0.091478453 | 0.649974741 | NAT9 | royalblue | -0.198848 | 0.330138 |
| TMEM214 | purple | -0.076209 | 0.711369 | PCNT | pink | 0.092132667 | 0.647630428 | CTNNBIP1 | royalblue | -0.187968 | 0.35781 |
| SWAP70 | purple | -0.075894 | 0.712511 | SEMA4A | pink | 0.0927755 | 0.645330088 | YIF1A | royalblue | -0.174479 | 0.393947 |
| RNF114 | purple | -0.075663 | 0.713352 | PSAP | pink | 0.094364481 | 0.639657699 | HIST1H4F | royalblue | -0.168687 | 0.410074 |
| DERL1 | purple | 0.073732 | 0.720373 | RSC1A1 | pink | 0.096254384 | 0.632936746 | MTMR14 | royalblue | -0.159815 | 0.435475 |
| TIGD2 | purple | -0.072212 | 0.72592 | TMEM209 | pink | 0.096577487 | 0.631790537 | PEF1 | royalblue | -0.150123 | 0.464171 |
| TADA2B | purple | -0.072005 | 0.726676 | ME3 | pink | 0.097571346 | 0.628270032 | ST13 | royalblue | -0.149896 | 0.464853 |
| MON1B | purple | -0.071996 | 0.726707 | UBA7 | pink | 0.097667276 | 0.627930638 | HS3ST2 | royalblue | -0.148405 | 0.469358 |
| CCRN4L | purple | -0.071219 | 0.729547 | PPA1 | pink | 0.100759818 | 0.617029402 | UQCRC1 | royalblue | -0.147439 | 0.472286 |
| TES | purple | 0.069591 | 0.735511 | AP3S2 | pink | 0.101795432 | 0.613396338 | ILK | royalblue | -0.142654 | 0.486938 |
| CSTB | purple | -0.06925 | 0.736761 | DGUOK | pink | 0.102272848 | 0.611724492 | RNF5 | royalblue | -0.14143 | 0.490721 |
| RAB21 | purple | 0.068999 | 0.737683 | AFTPH | pink | 0.102334923 | 0.611507255 | RANGRF | royalblue | -0.133799 | 0.514641 |
| PIP5K1A | purple | 0.067976 | 0.741439 | FRRS1 | pink | 0.103590384 | 0.607120518 | CLTB | royalblue | -0.133641 | 0.515141 |
| GOLGA6C | purple | -0.067391 | 0.743592 | POLR2D | pink | 0.103800339 | 0.606388201 | GPX1 | royalblue | -0.131345 | 0.522452 |
| AQP9 | purple | -0.065812 | 0.749408 | PCBP2 | pink | 0.104847825 | 0.602740141 | NDUFB7 | royalblue | -0.131301 | 0.522591 |
| ASB8 | purple | 0.06412 | 0.755657 | NT5C2 | pink | 0.105211225 | 0.601476706 | MRPS5 | royalblue | -0.129169 | 0.529423 |
| XPO6 | purple | -0.063367 | 0.758441 | ZNF597 | pink | 0.106822672 | 0.59588775 | TP53RK | royalblue | -0.122957 | 0.549571 |
| RAPGEF2 | purple | -0.063049 | 0.759619 | KIDINS220 | pink | 0.107482593 | 0.593605389 | ARHGAP30 | royalblue | -0.119534 | 0.560823 |
| TP53BP2 | purple | 0.0616 | 0.764988 | FADS3 | pink | 0.107894509 | 0.592182668 | GALT | royalblue | -0.109453 | 0.594547 |
| FUNDC2 | purple | -0.060506 | 0.769049 | BCAS3 | pink | 0.110216135 | 0.584191612 | UBXN1 | royalblue | -0.104483 | 0.611487 |
| TGIF2 | purple | -0.06008 | 0.770636 | RFT1 | pink | 0.110294245 | 0.583923579 | MRPL51 | royalblue | -0.097774 | 0.634662 |
| CYTH1 | purple | -0.059733 | 0.771925 | BUB3 | pink | 0.110475385 | 0.583302204 | FLOT2 | royalblue | -0.09631 | 0.639765 |

**Table 6. Gene module membership and correlation with growth rate (48/72h), with adjusted p values for each module**

| | | | | | | | | | | | |
|---|---|---|---|---|---|---|---|---|---|---|---|
| POLG2 | purple | -0.059647 | 0.772245 | MRPS31 | pink | 0.110934355 | 0.581729071 | UBN2 | royalblue | -0.087592 | 0.670483 |
| ARL8A | purple | -0.059365 | 0.773294 | ANO5 | pink | 0.112312155 | 0.577017843 | PQLC1 | royalblue | -0.082908 | 0.687206 |
| ZNFX1 | purple | -0.056795 | 0.782874 | CAPNS1 | pink | 0.112887184 | 0.575056603 | HERC2P9 | royalblue | -0.066418 | 0.747173 |
| CCSAP | purple | -0.056709 | 0.783191 | HIST2H2BE | pink | 0.11476834 | 0.568661344 | TSPO | royalblue | -0.060457 | 0.769231 |
| ZNF317 | purple | -0.047755 | 0.8168 | STRADB | pink | 0.117198044 | 0.560448778 | OGFR | royalblue | -0.056413 | 0.784298 |
| LIMS1 | purple | -0.047459 | 0.817919 | USP3 | pink | 0.117538291 | 0.559303043 | RNF181 | royalblue | -0.051845 | 0.801408 |
| COG3 | purple | -0.04667 | 0.820897 | TMEM14C | pink | 0.118116087 | 0.557359836 | SUN1 | royalblue | -0.043889 | 0.831416 |
| GPR32 | purple | -0.044503 | 0.82909 | STAU1 | pink | 0.119091444 | 0.554086601 | LBR | royalblue | -0.040423 | 0.844564 |
| SDHAF1 | purple | -0.042467 | 0.836806 | SLC9A6 | pink | 0.119098445 | 0.554063136 | TSR2 | royalblue | -0.037609 | 0.855266 |
| FLII | purple | 0.042097 | 0.838206 | GMEB1 | pink | 0.120838038 | 0.548247307 | UHMK1 | royalblue | -0.036666 | 0.85886 |
| SYNJ2 | purple | 0.037154 | 0.857 | CTTN | pink | 0.124314054 | 0.53671179 | B3GAT3 | royalblue | -0.036111 | 0.860975 |
| MAP7D1 | purple | -0.035545 | 0.863134 | SEC22C | pink | 0.12742936 | 0.526471713 | CXADR | royalblue | -0.028821 | 0.88885 |
| OTUB2 | purple | -0.035031 | 0.865098 | SNTB2 | pink | 0.128350758 | 0.523461117 | TNFRSF14 | royalblue | -0.024676 | 0.904756 |
| IBTK | purple | -0.034737 | 0.866217 | B3GALTL | pink | 0.130800331 | 0.515497869 | IGF1R | royalblue | -0.019998 | 0.922755 |
| CDV3 | purple | 0.033944 | 0.869246 | CRK | pink | 0.132270053 | 0.510748496 | LYL1 | royalblue | -0.019998 | 0.922756 |
| MAPK7 | purple | 0.031423 | 0.878885 | TMEM123 | pink | 0.133106886 | 0.508053917 | PHC3 | royalblue | -0.016575 | 0.935946 |
| NEDD9 | purple | -0.029764 | 0.885234 | APOOL | pink | 0.136637833 | 0.496761963 | RASA1 | royalblue | -0.009793 | 0.962129 |
| UBTD2 | purple | -0.029757 | 0.885265 | ATF7IP | pink | 0.137200473 | 0.494974319 | TOE1 | royalblue | -0.00948 | 0.963341 |
| USP12 | purple | 0.028075 | 0.89171 | EIF2C4 | pink | 0.137224433 | 0.494898262 | MEAF6 | royalblue | -0.008623 | 0.966654 |
| SDE2 | purple | -0.027992 | 0.892029 | TOX4 | pink | 0.138433867 | 0.491066845 | H2AFX | royalblue | -0.005267 | 0.979626 |
| OC10050658 | purple | 0.026626 | 0.897269 | HCCS | pink | 0.142079077 | 0.47961015 | REST | royalblue | 0.009879 | 0.961798 |
| WARS | purple | 0.025269 | 0.902481 | LSMD1 | pink | 0.143623301 | 0.474798366 | MRPL4 | royalblue | 0.013873 | 0.946373 |
| NRIP1 | purple | -0.023008 | 0.911169 | UPRT | pink | 0.143772352 | 0.474335245 | WSB1 | royalblue | 0.014595 | 0.943587 |
| ZNHIT2 | purple | 0.02207 | 0.914778 | WDR48 | pink | 0.145983277 | 0.467493053 | C5 | royalblue | 0.017597 | 0.932006 |
| ZHX2 | purple | -0.020442 | 0.921045 | AIFM1 | pink | 0.146083504 | 0.467184102 | POLR2E | royalblue | 0.019258 | 0.925606 |
| MAP4K4 | purple | 0.019963 | 0.922891 | BAZ2B | pink | 0.153969825 | 0.443210628 | ITGAL | royalblue | 0.019324 | 0.925349 |
| PRRC1 | purple | -0.019935 | 0.922998 | TSTA3 | pink | 0.156169808 | 0.436642812 | PMF1 | royalblue | 0.022688 | 0.912402 |
| ACSL5 | purple | 0.019127 | 0.92611 | GNG5 | pink | 0.157344613 | 0.433157238 | GEMIN8 | royalblue | 0.030508 | 0.882386 |
| ZNF697 | purple | 0.018981 | 0.926674 | PSMB10 | pink | 0.160424244 | 0.424092258 | FRMD4A | royalblue | 0.042287 | 0.837486 |
| DUS2L | purple | 0.018429 | 0.928799 | BST2 | pink | 0.163895508 | 0.414000552 | NIPBL | royalblue | 0.045583 | 0.825006 |
| SNORA41 | purple | 0.01835 | 0.929105 | DAK | pink | 0.165231293 | 0.410152989 | ZFPL1 | royalblue | 0.050766 | 0.80546 |
| IRF2BPL | purple | 0.018041 | 0.930294 | PIGN | pink | 0.16848987 | 0.400851211 | ZFAND2B | royalblue | 0.052024 | 0.800736 |
| ZNF416 | purple | -0.018017 | 0.930387 | DRG1 | pink | 0.169691798 | 0.39745052 | ITPR1 | royalblue | 0.060247 | 0.770015 |
| ZNF182 | purple | 0.017986 | 0.930507 | LRP11 | pink | 0.172895524 | 0.388466179 | PPIF | royalblue | 0.062493 | 0.761679 |
| ZNF202 | purple | -0.017747 | 0.931427 | MANF | pink | 0.173773015 | 0.386025812 | LONRF1 | royalblue | 0.065246 | 0.751495 |
| CXCL5 | purple | -0.016984 | 0.93437 | NAA30 | pink | 0.17383086 | 0.385865248 | ALDH16A1 | royalblue | 0.065359 | 0.75108 |
| SAFB | purple | -0.016906 | 0.934672 | PSMC4 | pink | 0.176188198 | 0.379354544 | SERPINF2 | royalblue | 0.068952 | 0.737856 |
| GABPB1 | purple | 0.012883 | 0.950194 | NIP7 | pink | 0.177149419 | 0.376718075 | PSMD9 | royalblue | 0.071945 | 0.726893 |
| JRKL | purple | -0.012438 | 0.951913 | NAPG | pink | 0.177268508 | 0.376392174 | TATDN2 | royalblue | 0.074339 | 0.718164 |
| IL1A | purple | -0.012377 | 0.952147 | EIF2AK3 | pink | 0.180071796 | 0.368767895 | TOPORS | royalblue | 0.079049 | 0.701091 |
| CH25H | purple | 0.00688 | 0.973391 | TDP2 | pink | 0.180611743 | 0.367309799 | NAA25 | royalblue | 0.090111 | 0.661552 |
| COQ10A | purple | 0.005252 | 0.979685 | IMP4 | pink | 0.183048437 | 0.360771696 | EPT1 | royalblue | 0.096525 | 0.639017 |
| MAP4K3 | red | 0.622744 | 0.00068 | FOXP1 | pink | 0.185799216 | 0.353473806 | STXBP5 | royalblue | 0.097678 | 0.634997 |
| PLAGL1 | red | 0.49803 | 0.00962 | DAD1 | pink | 0.187212259 | 0.349759281 | DNPEP | royalblue | 0.100571 | 0.62496 |
| CNP | red | 0.493756 | 0.010362 | SPG11 | pink | 0.188193735 | 0.347192973 | SDHAP2 | royalblue | 0.108222 | 0.598723 |
| FAM96A | red | -0.473245 | 0.014613 | ADIPOR1 | pink | 0.189122728 | 0.344774282 | SFXN3 | royalblue | 0.116201 | 0.571875 |
| KIFC1 | red | -0.469687 | 0.01548 | NUTF2 | pink | 0.189481405 | 0.343843151 | ABL1 | royalblue | 0.117539 | 0.567427 |
| SERPINB1 | red | -0.463083 | 0.0172 | S100A13 | pink | 0.190224779 | 0.341918142 | GRHL1 | royalblue | 0.118436 | 0.564452 |
| TPRKB | red | -0.455791 | 0.019279 | VPS11 | pink | 0.194800079 | 0.330213131 | SLC6A6 | royalblue | 0.130525 | 0.525072 |
| BTN2A1 | red | 0.453105 | 0.020095 | EIF2AK1 | pink | 0.195837996 | 0.327592126 | MGA | royalblue | 0.136315 | 0.506692 |
| KIAA0913 | red | 0.450242 | 0.020995 | GMPR | pink | 0.196330038 | 0.326354035 | MTMR10 | royalblue | 0.148919 | 0.467803 |
| TARSL2 | red | 0.428869 | 0.028805 | PHF8 | pink | 0.196781335 | 0.32522098 | GNG12 | royalblue | 0.150147 | 0.464098 |
| MTHFR | red | 0.422496 | 0.031539 | BMF | pink | 0.200760405 | 0.315335055 | TAOK1 | royalblue | 0.151768 | 0.459233 |
| PAIP1 | red | -0.418807 | 0.033215 | NOP56 | pink | 0.202616943 | 0.310786629 | PHF20L1 | royalblue | 0.169179 | 0.408691 |
| CYSLTR1 | red | -0.413246 | 0.035874 | BTRC | pink | 0.206654134 | 0.30103677 | SPAG9 | royalblue | 0.172895 | 0.398319 |
| PAQR3 | red | -0.413122 | 0.035935 | EIF6 | pink | 0.212022094 | 0.288372787 | RPIA | royalblue | 0.206868 | 0.310597 |

**Table 6. Gene module membership and correlation with growth rate (48/72h), with adjusted p values for each module**

| | | | | | | | | | | | |
|---|---|---|---|---|---|---|---|---|---|---|---|
| MAPK1IP1L | red | 0.406554 | 0.039297 | SYNCRIP | pink | 0.212918089 | 0.286292315 | C21orf2 | royalblue | 0.215882 | 0.289508 |
| RFC1 | red | -0.404572 | 0.040359 | OASL | pink | 0.213184778 | 0.285674914 | POGZ | royalblue | 0.221826 | 0.276106 |
| SNRPA | red | 0.402339 | 0.041583 | SEC62 | pink | 0.220694848 | 0.268635605 | PARP15 | royalblue | 0.257872 | 0.20342 |
| STK38 | red | -0.401601 | 0.041994 | ANKRD40 | pink | 0.225058523 | 0.259042562 | SLU7 | royalblue | 0.27385 | 0.17582 |
| OCRL | red | -0.398174 | 0.043944 | CDC34 | pink | 0.233444617 | 0.241239389 | PLCXD1 | royalblue | 0.342315 | 0.086937 |
| ZNF506 | red | 0.398087 | 0.043994 | C8orf82 | pink | 0.243232051 | 0.221507685 | ZNF37A | royalblue | 0.371863 | 0.0614 |
| TTLL1 | red | -0.393081 | 0.046973 | ORMDL2 | pink | 0.245144989 | 0.217781916 | HIST1H2BN | saddlebrown | -0.243531 | 0.230587 |
| MTCH1 | red | -0.387608 | 0.050409 | GMFG | pink | 0.24735065 | 0.213538777 | ZDHHC6 | saddlebrown | -0.211978 | 0.298529 |
| ATG4C | red | -0.387067 | 0.050759 | SLC35B1 | pink | 0.249168906 | 0.210083278 | EIF3D | saddlebrown | -0.161768 | 0.429812 |
| SDSL | red | 0.384396 | 0.052516 | FKBP4 | pink | 0.254158482 | 0.200796651 | C14orf142 | saddlebrown | -0.148295 | 0.469691 |
| CEP55 | red | -0.384209 | 0.052641 | FABP3 | pink | 0.254451292 | 0.200260553 | RPA2 | saddlebrown | -0.146525 | 0.475069 |
| CNOT8 | red | -0.381241 | 0.054651 | PDLIM4 | pink | 0.25813741 | 0.193595493 | CD99 | saddlebrown | -0.124145 | 0.545689 |
| EXTL2 | red | -0.380551 | 0.055127 | ASB8 | pink | 0.261220205 | 0.188139986 | SNORA71A | saddlebrown | -0.123077 | 0.549179 |
| ZCCHC9 | red | -0.380131 | 0.055418 | DIEXF | pink | 0.268140646 | 0.17628314 | KIAA0232 | saddlebrown | -0.104578 | 0.61116 |
| ARL4A | red | -0.378638 | 0.056463 | FAM96B | pink | 0.272125383 | 0.169698362 | SNORA65 | saddlebrown | -0.100171 | 0.626344 |
| MRPS14 | red | -0.378516 | 0.05655 | GPR137B | pink | 0.273112424 | 0.168094384 | IGFBP2 | saddlebrown | -0.097495 | 0.635634 |
| FBXO38 | red | -0.378136 | 0.056818 | VHL | pink | 0.276483311 | 0.162697157 | GEMIN5 | saddlebrown | -0.094044 | 0.647697 |
| CREBZF | red | 0.377531 | 0.057248 | DYNC1I2 | pink | 0.277657777 | 0.160845828 | RPS19 | saddlebrown | -0.078487 | 0.70312 |
| ALDH1A2 | red | 0.377245 | 0.057452 | URB2 | pink | 0.280144139 | 0.156975919 | RPPH1 | saddlebrown | -0.074933 | 0.716002 |
| SLC35D2 | red | -0.376401 | 0.058058 | KCMF1 | pink | 0.285760299 | 0.148479357 | TP53I13 | saddlebrown | -0.072018 | 0.726628 |
| USP1 | red | -0.374658 | 0.059325 | VWA5A | pink | 0.291856096 | 0.139635799 | SCARNA12 | saddlebrown | -0.064556 | 0.754046 |
| C12orf4 | red | -0.373274 | 0.060345 | UBE2D3 | pink | 0.299780903 | 0.128715939 | SNORA5C | saddlebrown | -0.054952 | 0.78976 |
| PSMB8 | red | -0.372866 | 0.060649 | ATP5G1 | pink | 0.305869173 | 0.120759811 | PPM1L | saddlebrown | -0.052698 | 0.798206 |
| FOXJ2 | red | 0.372159 | 0.061177 | WSB2 | pink | 0.31112978 | 0.114181275 | KLHL6 | saddlebrown | -0.050361 | 0.806984 |
| VTA1 | red | -0.371991 | 0.061304 | MCFD2 | pink | 0.313369572 | 0.11146219 | GOLT1B | saddlebrown | -0.049399 | 0.810607 |
| HUWE1 | red | 0.371727 | 0.061502 | ZNF330 | pink | 0.3157571 | 0.108616865 | SNORA63 | saddlebrown | -0.046825 | 0.820312 |
| TMEM194A | red | -0.371249 | 0.061864 | FCHO2 | pink | 0.337118433 | 0.085509035 | SNORA6 | saddlebrown | -0.032691 | 0.874035 |
| TMEM87B | red | -0.371217 | 0.061888 | SRP72 | pink | 0.340011757 | 0.082691119 | SNORA9 | saddlebrown | -0.030477 | 0.882507 |
| SELT | red | -0.369774 | 0.062988 | NDUFA8 | pink | 0.340083856 | 0.082621812 | SNORA67 | saddlebrown | -0.030375 | 0.882898 |
| SECISBP2 | red | -0.36914 | 0.063476 | LRRC28 | pink | 0.344499466 | 0.078460808 | RNF11 | saddlebrown | -0.012514 | 0.95162 |
| PIGX | red | -0.367599 | 0.064675 | VKORC1 | pink | 0.359559857 | 0.065463834 | PDZD7 | saddlebrown | -0.004829 | 0.98132 |
| SPNS1 | red | -0.365849 | 0.066057 | INTS4 | pink | 0.500433557 | 0.007850311 | HIST1H3D | saddlebrown | -0.004516 | 0.982531 |
| MEPCE | red | 0.365771 | 0.066119 | LOC10013289 | purple | -0.581450293 | 0.001468045 | FAM178A | saddlebrown | 0.005494 | 0.978751 |
| MLF1IP | red | -0.364405 | 0.067214 | ERI2 | purple | -0.477462443 | 0.011784236 | SCARNA16 | saddlebrown | 0.006404 | 0.975229 |
| SASS6 | red | -0.363891 | 0.06763 | PIP5K1C | purple | -0.475006212 | 0.012287987 | SQRDL | saddlebrown | 0.015039 | 0.941873 |
| TRIM68 | red | -0.363815 | 0.067692 | TAF8 | purple | -0.393569833 | 0.042246196 | SNORA71D | saddlebrown | 0.020187 | 0.922028 |
| ZDHHC2 | red | -0.3629 | 0.068438 | SCYL1 | purple | -0.384419674 | 0.047724969 | SNORA2A | saddlebrown | 0.025553 | 0.901388 |
| HCFC1 | red | 0.36248 | 0.068782 | KCNQ1 | purple | -0.377515776 | 0.052217961 | SNORA42 | saddlebrown | 0.031045 | 0.880331 |
| CCNH | red | -0.362445 | 0.068811 | EFTUD2 | purple | -0.367687981 | 0.059178541 | SNORD94 | saddlebrown | 0.036532 | 0.85937 |
| FARP2 | red | 0.361778 | 0.06936 | ASXL2 | purple | -0.354529755 | 0.069604607 | IFI27 | saddlebrown | 0.03833 | 0.852523 |
| FEM1A | red | -0.361624 | 0.069488 | C8orf33 | purple | -0.343919218 | 0.078998275 | SCARNA6 | saddlebrown | 0.041916 | 0.838896 |
| RRM1 | red | -0.352765 | 0.077125 | MED12 | purple | -0.33474591 | 0.087873576 | SCARNA3 | saddlebrown | 0.058867 | 0.775148 |
| PLIN2 | red | 0.351854 | 0.077944 | RGP1 | purple | -0.327841063 | 0.095036913 | SNORA74A | saddlebrown | 0.059868 | 0.771423 |
| PDCD2 | red | -0.35142 | 0.078338 | ANKS1A | purple | -0.307756636 | 0.118368239 | SNORD17 | saddlebrown | 0.061022 | 0.767136 |
| C19orf6 | red | 0.34994 | 0.07969 | ANAPC2 | purple | -0.30468018 | 0.122284465 | SNORA43 | saddlebrown | 0.062068 | 0.763253 |
| TBC1D10C | red | -0.348087 | 0.081407 | H1FX | purple | -0.283578376 | 0.151740268 | SNORA52 | saddlebrown | 0.072688 | 0.72418 |
| PRCP | red | -0.346837 | 0.082581 | WHSC2 | purple | -0.279837677 | 0.1574493 | ZNF688 | saddlebrown | 0.074251 | 0.718483 |
| SAMD9L | red | -0.346078 | 0.0833 | ZNF319 | purple | -0.277992585 | 0.160320812 | SNORA7B | saddlebrown | 0.076196 | 0.711417 |
| SOS2 | red | -0.345625 | 0.083732 | PER2 | purple | -0.268499527 | 0.175682876 | SNORA38 | saddlebrown | 0.083234 | 0.686034 |
| OXR1 | red | -0.343671 | 0.085613 | LETM1 | purple | -0.266216111 | 0.179526608 | SNORA5A | saddlebrown | 0.087071 | 0.672333 |
| BCKDHB | red | -0.34365 | 0.085633 | TPR | purple | -0.259552132 | 0.191078538 | SNORA46 | saddlebrown | 0.088076 | 0.668763 |
| SEC11A | red | -0.343581 | 0.085701 | PIEZO1 | purple | -0.258195221 | 0.193492196 | SNORA34 | saddlebrown | 0.090527 | 0.660082 |
| PSMG2 | red | -0.341316 | 0.087922 | DIS3L2 | purple | -0.254439535 | 0.20028206 | SNORA55 | saddlebrown | 0.097497 | 0.635627 |
| ASPH | red | -0.340907 | 0.088328 | FURIN | purple | -0.252824896 | 0.20325071 | GABARAPL1 | saddlebrown | 0.11612 | 0.572146 |
| PARG | red | -0.339988 | 0.089245 | SMC4 | purple | -0.240177743 | 0.22754486 | SNORA81 | saddlebrown | 0.12258 | 0.550806 |
| SUCLA2 | red | -0.337906 | 0.091349 | PSKH1 | purple | -0.239023886 | 0.229853949 | SNORA28 | saddlebrown | 0.130337 | 0.525675 |

**Table 6. Gene module membership and correlation with growth rate (48/72h), with adjusted p values for each module**

| | | | | | | | | | | | |
|---|---|---|---|---|---|---|---|---|---|---|---|
| PAPD4 | red | -0.33705 | 0.092225 | AP3D1 | purple | -0.238046123 | 0.231822839 | PIH1D1 | saddlebrown | 0.131553 | 0.521785 |
| TGFBR2 | red | -0.336936 | 0.092341 | CAPN1 | purple | -0.2354718 | 0.23706029 | SNORA16A | saddlebrown | 0.146697 | 0.474542 |
| CDK11A | red | -0.336301 | 0.092996 | ATP6V1A | purple | -0.232105785 | 0.244025955 | SNORA71B | saddlebrown | 0.17126 | 0.402864 |
| RPS20 | red | -0.336088 | 0.093216 | RALGAPA1 | purple | -0.225587089 | 0.257895896 | SNORD15B | saddlebrown | 0.17267 | 0.398945 |
| IFI44 | red | -0.335896 | 0.093415 | PGM3 | purple | -0.225314103 | 0.258487695 | HOXB6 | saddlebrown | 0.182766 | 0.371508 |
| TMEM5 | red | -0.335334 | 0.093999 | EXTL3 | purple | -0.215166267 | 0.281114118 | FAS | saddlebrown | 0.206503 | 0.311472 |
| GANC | red | -0.33513 | 0.094212 | SNTA1 | purple | -0.213026396 | 0.286041476 | SNORA50 | saddlebrown | 0.206776 | 0.310819 |
| GZF1 | red | 0.334003 | 0.095393 | PPP4R2 | purple | -0.210803013 | 0.291218776 | SNORA31 | saddlebrown | 0.222125 | 0.275444 |
| MDM1 | red | -0.33364 | 0.095775 | BAZ1A | purple | -0.207636989 | 0.298692442 | SNORD15A | saddlebrown | 0.228424 | 0.261703 |
| PGM2L1 | red | -0.333174 | 0.096269 | ZNF434 | purple | -0.199683789 | 0.317991407 | FDXR | saddlebrown | 0.27285 | 0.177466 |
| WSB2 | red | 0.332702 | 0.09677 | SND1 | purple | -0.198556581 | 0.320787283 | MIB2 | saddlebrown | 0.272858 | 0.177454 |
| LOC92249 | red | -0.332374 | 0.09712 | RAB3GAP2 | purple | -0.197322716 | 0.323864928 | PHF1 | saddlebrown | 0.287124 | 0.154969 |
| MAT2B | red | -0.331996 | 0.097524 | C2orf18 | purple | -0.195918812 | 0.327388579 | NR1H2 | saddlebrown | 0.380212 | 0.055362 |
| FAM3B | red | -0.331518 | 0.098037 | SBNO2 | purple | -0.194522629 | 0.330915914 | LMBR1L | saddlebrown | 0.42797 | 0.029178 |
| FRMD4B | red | -0.330605 | 0.099022 | MOV10 | purple | -0.194007797 | 0.332222393 | AKIP1 | salmon | -0.28589 | 0.156829 |
| UFL1 | red | -0.330493 | 0.099143 | EEF1B2 | purple | -0.191857816 | 0.337712081 | ALDH1L2 | salmon | -0.279823 | 0.166208 |
| TIMELESS | red | -0.330365 | 0.099282 | ATXN2L | purple | -0.183825179 | 0.358702049 | ZDHHC7 | salmon | -0.277103 | 0.170539 |
| COPB2 | red | -0.330047 | 0.099628 | SUGT1 | purple | -0.183001373 | 0.360897324 | AGPAT4 | salmon | -0.266074 | 0.188905 |
| SHPK | red | 0.329684 | 0.100023 | HELZ | purple | -0.180279013 | 0.36820792 | MPHOSPH8 | salmon | -0.241427 | 0.234766 |
| OPTN | red | -0.329512 | 0.100211 | NLN | purple | -0.179935173 | 0.369137374 | RPL10A | salmon | -0.240445 | 0.236732 |
| NKTR | red | -0.327641 | 0.102271 | TM9SF1 | purple | -0.17915407 | 0.3712539 | COL4A3BP | salmon | -0.236173 | 0.24542 |
| TYW3 | red | -0.327488 | 0.10244 | TPP1 | purple | -0.175412106 | 0.381490985 | WDR36 | salmon | -0.232966 | 0.252077 |
| MCM8 | red | -0.326558 | 0.103477 | SPCS3 | purple | -0.174228721 | 0.384761933 | ATP5O | salmon | -0.231616 | 0.254914 |
| SMAD1 | red | -0.326353 | 0.103706 | KIAA0247 | purple | -0.17402398 | 0.385329477 | STK38 | salmon | -0.215813 | 0.289666 |
| MARVELD1 | red | 0.32526 | 0.104937 | NOL8 | purple | -0.162616927 | 0.417702044 | TAPBP | salmon | -0.215077 | 0.291354 |
| MTO1 | red | -0.325017 | 0.105211 | GLRX3 | purple | -0.160093937 | 0.425059509 | GPHN | salmon | -0.211815 | 0.298909 |
| FEZ2 | red | -0.324512 | 0.105785 | TCTEX1D2 | purple | -0.158085465 | 0.430966968 | ZFAND5 | salmon | -0.210666 | 0.301598 |
| CHKB | red | -0.323988 | 0.106382 | ADAT1 | purple | -0.15724805 | 0.433443163 | SLC35D1 | salmon | -0.208662 | 0.306326 |
| TTI2 | red | -0.323626 | 0.106797 | GIT2 | purple | -0.152078854 | 0.448897984 | MTMR2 | salmon | -0.205876 | 0.312976 |
| FAM8A1 | red | -0.323461 | 0.106985 | ACTN1 | purple | -0.15040168 | 0.453974658 | POMT2 | salmon | -0.204194 | 0.317031 |
| SLC26A6 | red | 0.323227 | 0.107254 | ZNF592 | purple | -0.136684402 | 0.496613879 | SERF1A | salmon | -0.197063 | 0.334588 |
| SFMBT1 | red | -0.323106 | 0.107394 | UTP20 | purple | -0.134234204 | 0.504435083 | TMEM50A | salmon | -0.191367 | 0.34902 |
| CEP57 | red | -0.322898 | 0.107633 | KLHL9 | purple | -0.12693703 | 0.528083759 | TAF1 | salmon | -0.188796 | 0.355656 |
| CHSY1 | red | -0.322792 | 0.107755 | ADAM17 | purple | -0.123130104 | 0.540627949 | SPATA5 | salmon | -0.185332 | 0.364711 |
| NUSAP1 | red | -0.321367 | 0.109407 | RPN2 | purple | -0.121843736 | 0.544898013 | NRBF2 | salmon | -0.175127 | 0.392164 |
| ZNF207 | red | -0.320839 | 0.110024 | RANBP10 | purple | -0.119419123 | 0.552988914 | ASB3 | salmon | -0.171637 | 0.401814 |
| DNAJC24 | red | -0.319064 | 0.112117 | MGC12916 | purple | -0.115617936 | 0.565783521 | LOC440434 | salmon | -0.164496 | 0.421969 |
| MKNK1 | red | -0.319016 | 0.112175 | WBP5 | purple | -0.115509243 | 0.566151329 | TCTA | salmon | -0.163176 | 0.425755 |
| VBP1 | red | 0.316754 | 0.114885 | ZDHHC5 | purple | -0.114888353 | 0.568254429 | CCDC93 | salmon | -0.159337 | 0.436868 |
| INTS8 | red | -0.315954 | 0.115855 | TMEM237 | purple | -0.113525788 | 0.572882004 | ANKRD52 | salmon | -0.158363 | 0.439713 |
| RPL22L1 | red | -0.315668 | 0.116204 | RNF146 | purple | -0.110100308 | 0.58458917 | CDAN1 | salmon | -0.156405 | 0.445461 |
| ARPP19 | red | -0.315439 | 0.116483 | UPF3A | purple | -0.108587863 | 0.589791211 | DIP2B | salmon | -0.156239 | 0.445949 |
| CCDC101 | red | 0.314388 | 0.117772 | LAIR1 | purple | -0.107704272 | 0.592839548 | MED11 | salmon | -0.151993 | 0.458558 |
| CD302 | red | -0.313305 | 0.119112 | DENR | purple | -0.107530137 | 0.593441102 | RPL36A | salmon | -0.146659 | 0.474659 |
| ZBTB37 | red | -0.313241 | 0.119191 | RDH11 | purple | -0.106856588 | 0.595770359 | SMG7 | salmon | -0.146416 | 0.475399 |
| EI24 | red | -0.313234 | 0.119199 | N4BP2L2 | purple | -0.103661539 | 0.60687229 | SLTM | salmon | -0.144561 | 0.481069 |
| MOGAT1 | red | -0.31248 | 0.120138 | SEPT7P2 | purple | -0.102066833 | 0.612445698 | ETV6 | salmon | -0.143894 | 0.483117 |
| ACTG1 | red | 0.312035 | 0.120696 | SEC63 | purple | -0.100977535 | 0.61626489 | UQCRH | salmon | -0.139842 | 0.495652 |
| EZH2 | red | -0.311676 | 0.121146 | ESF1 | purple | -0.090801607 | 0.65240358 | HDAC2 | salmon | -0.137392 | 0.503307 |
| GMPPA | red | -0.311452 | 0.121428 | FNBP4 | purple | -0.09052133 | 0.653410361 | AUH | salmon | -0.136866 | 0.504959 |
| OVCA2 | red | -0.310546 | 0.122572 | SEMA3C | purple | -0.089925776 | 0.655551611 | SNHG1 | salmon | -0.13551 | 0.509226 |
| NOL8 | red | -0.31004 | 0.123215 | FAM172A | purple | -0.087415078 | 0.66460763 | ABL2 | salmon | -0.124578 | 0.54428 |
| FUCA1 | red | -0.309375 | 0.124064 | ITPR2 | purple | -0.075687647 | 0.707501465 | C2CD2 | salmon | -0.123279 | 0.548519 |
| BTF3L4 | red | -0.307621 | 0.126323 | SAP130 | purple | -0.072532727 | 0.719197716 | NAA60 | salmon | -0.114007 | 0.579205 |
| STRN4 | red | 0.307179 | 0.126896 | EHMT2 | purple | -0.07010133 | 0.728254103 | SCARNA10 | salmon | -0.11168 | 0.587023 |
| CDC34 | red | 0.306437 | 0.127863 | TLN1 | purple | -0.068303385 | 0.734974084 | PPP1CA | salmon | -0.111285 | 0.588356 |

**Table 6. Gene module membership and correlation with growth rate (48/72h), with adjusted p values for each module**

| | | | | | | | | | | | |
|---|---|---|---|---|---|---|---|---|---|---|---|
| ZCCHC2 | red | -0.30626 | 0.128095 | BRD2 | purple | -0.066487316 | 0.74178119 | ERC1 | salmon | -0.110414 | 0.591295 |
| ARID4B | red | -0.304695 | 0.130156 | UCHL5 | purple | -0.060544031 | 0.764188618 | HIGD2A | salmon | -0.108116 | 0.599084 |
| TUBD1 | red | -0.303038 | 0.132364 | APC | purple | -0.055884253 | 0.781888523 | EMC4 | salmon | -0.108076 | 0.599221 |
| RSL24D1 | red | -0.302507 | 0.133077 | PARP14 | purple | -0.051308959 | 0.799371348 | FYCO1 | salmon | -0.104056 | 0.612951 |
| EPSTI1 | red | -0.302255 | 0.133416 | SENP6 | purple | -0.050678732 | 0.801787146 | PRKAR1A | salmon | -0.099755 | 0.627785 |
| ATM | red | -0.302106 | 0.133618 | CCDC97 | purple | -0.042201357 | 0.834447295 | RPL37 | salmon | -0.098151 | 0.63335 |
| RFC2 | red | -0.301258 | 0.134766 | CMAHP | purple | -0.039102673 | 0.846455587 | PDCL | salmon | -0.097508 | 0.635588 |
| DECR1 | red | -0.300301 | 0.13607 | ZNF569 | purple | -0.039076724 | 0.846556292 | ATP5E | salmon | -0.097013 | 0.637315 |
| TRIT1 | red | -0.300224 | 0.136176 | HERC2P2 | purple | -0.035569977 | 0.860186596 | HNRNPL | salmon | -0.090557 | 0.659976 |
| NOMO2 | red | -0.299886 | 0.13664 | CLINT1 | purple | -0.034185344 | 0.865579506 | NHP2 | salmon | -0.08841 | 0.667578 |
| BROX | red | -0.299452 | 0.137236 | GAK | purple | -0.029107206 | 0.885406547 | FCGR3A | salmon | -0.087479 | 0.670882 |
| MTAP | red | -0.298603 | 0.138408 | LILRA6 | purple | -0.019972282 | 0.921235193 | PATL2 | salmon | -0.080472 | 0.695958 |
| PTPN11 | red | -0.298017 | 0.139221 | TRIM37 | purple | -0.019728728 | 0.922192814 | POM121C | salmon | -0.076961 | 0.708642 |
| HERC2 | red | 0.297677 | 0.139694 | RNF168 | purple | -0.018981152 | 0.925132847 | MNF1 | salmon | -0.07552 | 0.713869 |
| PRPF19 | red | -0.297634 | 0.139754 | TNRC6C | purple | -0.018448722 | 0.927227354 | RFC4 | salmon | -0.073424 | 0.721497 |
| XRCC1 | red | 0.297551 | 0.139869 | TRNT1 | purple | -0.016285488 | 0.935741932 | PSENEN | salmon | -0.071159 | 0.729769 |
| TBC1D15 | red | -0.296815 | 0.140899 | FARP1 | purple | -0.015881773 | 0.937331762 | TAX1BP3 | salmon | -0.068826 | 0.738316 |
| HNRPLL | red | -0.296333 | 0.141577 | HELB | purple | -0.011871289 | 0.95313684 | FAM20C | salmon | -0.066815 | 0.745714 |
| SPPL2B | red | 0.294896 | 0.14361 | ZNF322 | purple | -0.010665275 | 0.95789332 | ZNF416 | salmon | -0.06492 | 0.752699 |
| DERA | red | -0.294185 | 0.144623 | GNAI3 | purple | -0.009853404 | 0.961096115 | CRTC3 | salmon | -0.061391 | 0.765766 |
| ANKRD42 | red | -0.293946 | 0.144965 | HSCB | purple | -0.006710048 | 0.973501644 | SLC30A7 | salmon | -0.061112 | 0.766801 |
| SESN1 | red | -0.293789 | 0.145191 | ATPAF1 | purple | -0.005064148 | 0.979999908 | THUMPD1 | salmon | -0.060734 | 0.768202 |
| NCOA7 | red | 0.292831 | 0.146568 | APOC1 | purple | -0.002527123 | 0.990018771 | APIP | salmon | -0.059452 | 0.77297 |
| ERP29 | red | -0.292705 | 0.14675 | ANAPC10 | purple | -0.001846363 | 0.992707444 | ZNF14 | salmon | -0.058701 | 0.775765 |
| RNF115 | red | -0.291978 | 0.147803 | DYNLT3 | purple | 0.001754319 | 0.993070982 | EIF2S1 | salmon | -0.053847 | 0.793897 |
| ALG5 | red | -0.291556 | 0.148416 | PLXNA1 | purple | 0.004410716 | 0.982580125 | RNF169 | salmon | -0.052623 | 0.798489 |
| STK39 | red | -0.290928 | 0.149331 | MRPL50 | purple | 0.005347761 | 0.978880063 | UXT | salmon | -0.051986 | 0.800878 |
| UVRAG | red | -0.290855 | 0.149439 | HAUS2 | purple | 0.017486275 | 0.931014673 | LSM14A | salmon | -0.050961 | 0.804729 |
| TMEM131 | red | -0.290743 | 0.149602 | TMEM164 | purple | 0.018758381 | 0.926009138 | PEBP1 | salmon | -0.043733 | 0.832006 |
| TMEM219 | red | 0.290201 | 0.150397 | WRN | purple | 0.024677103 | 0.902758998 | ALOX5AP | salmon | -0.03461 | 0.866705 |
| SLC22A15 | red | -0.288868 | 0.152366 | LSG1 | purple | 0.028272268 | 0.888673249 | NIPSNAP1 | salmon | -0.026005 | 0.899654 |
| C18orf25 | red | 0.288193 | 0.15337 | PSMG1 | purple | 0.028386593 | 0.888225847 | COX5B | salmon | -0.024105 | 0.906953 |
| TTC37 | red | -0.287509 | 0.154391 | RPL7 | purple | 0.029813055 | 0.88264633 | PET100 | salmon | -0.022169 | 0.914396 |
| LANCL1 | red | -0.287346 | 0.154636 | DDHD1 | purple | 0.029844286 | 0.882524231 | SAMD8 | salmon | -0.017512 | 0.932333 |
| MORF4L2 | red | -0.286001 | 0.156661 | RAB18 | purple | 0.030340404 | 0.880585006 | ACVR1 | salmon | -0.017205 | 0.933517 |
| DHRS9 | red | -0.285533 | 0.157371 | SLC44A2 | purple | 0.038853107 | 0.84742422 | ZNF182 | salmon | -0.013848 | 0.946471 |
| WDR47 | red | -0.285293 | 0.157735 | AKAP10 | purple | 0.040105261 | 0.842566473 | TSN | salmon | -0.010489 | 0.959441 |
| KCTD9 | red | -0.28509 | 0.158044 | SLC4A2 | purple | 0.040266325 | 0.841942027 | PILRB | salmon | -0.010119 | 0.960871 |
| GPR108 | red | 0.284137 | 0.1595 | CCDC25 | purple | 0.040783674 | 0.839936895 | C19orf43 | salmon | -0.004716 | 0.981758 |
| LOC731275 | red | -0.283437 | 0.160574 | BDP1 | purple | 0.043718578 | 0.828580642 | CHCHD4 | salmon | -0.003726 | 0.985587 |
| SMARCD2 | red | 0.28276 | 0.16162 | CWF19L1 | purple | 0.050497328 | 0.80248284 | GSTP1 | salmon | -0.000563 | 0.997823 |
| FIG4 | red | -0.28251 | 0.162007 | CBWD5 | purple | 0.050561935 | 0.802235054 | PPARA | salmon | 0.003969 | 0.984648 |
| FGF10 | red | -0.282051 | 0.162719 | BMS1P4 | purple | 0.053992628 | 0.789104663 | TLK2 | salmon | 0.006291 | 0.97567 |
| TTC23 | red | -0.281151 | 0.164122 | ZNF148 | purple | 0.054172274 | 0.788418613 | NDUFA12 | salmon | 0.013237 | 0.94883 |
| ADAM9 | red | -0.280977 | 0.164394 | C14orf28 | purple | 0.056821733 | 0.77831872 | CCT7 | salmon | 0.021898 | 0.915441 |
| FAM82B | red | -0.280894 | 0.164525 | FAM82A2 | purple | 0.065888494 | 0.744029924 | ACD | salmon | 0.023813 | 0.908074 |
| CLIP1 | red | -0.280116 | 0.165746 | GTF2H2C | purple | 0.068254697 | 0.735156327 | NRIP1 | salmon | 0.028698 | 0.889322 |
| SECISBP2L | red | -0.279383 | 0.166903 | NAT10 | purple | 0.069311332 | 0.731204409 | RPS5 | salmon | 0.030445 | 0.882629 |
| ALG11 | red | -0.279276 | 0.167072 | BTBD3 | purple | 0.069569419 | 0.730240153 | NARS | salmon | 0.030991 | 0.880538 |
| WWP1 | red | -0.277104 | 0.170536 | PTPN2 | purple | 0.072143437 | 0.720645279 | FLII | salmon | 0.031112 | 0.880075 |
| ATP6V0A2 | red | -0.276911 | 0.170847 | TAF7 | purple | 0.07494932 | 0.710232983 | DOT1L | salmon | 0.032187 | 0.875961 |
| CLIC1 | red | 0.276879 | 0.170898 | PSTPIP2 | purple | 0.076034003 | 0.706221302 | PSMB3 | salmon | 0.032694 | 0.874024 |
| RGS18 | red | -0.276467 | 0.171561 | SH3YL1 | purple | 0.07703865 | 0.702512453 | DKC1 | salmon | 0.033011 | 0.872811 |
| AP3B1 | red | -0.274943 | 0.174033 | SUZ12P1 | purple | 0.08209801 | 0.683937232 | SSSCA1 | salmon | 0.033409 | 0.87129 |
| FADS1 | red | -0.27456 | 0.174658 | LRIG2 | purple | 0.083589837 | 0.678493573 | TMEM134 | salmon | 0.033797 | 0.869809 |
| IFI44L | red | -0.273231 | 0.176838 | SDAD1 | purple | 0.083711465 | 0.678050443 | BRD4 | salmon | 0.039134 | 0.849464 |

**Table 6. Gene module membership and correlation with growth rate (48/72h), with adjusted p values for each module**

| | | | |
|---|---|---|---|
| BLZF1 | red | -0.273023 | 0.177181 |
| SERAC1 | red | -0.272573 | 0.177924 |
| FUS | red | -0.272274 | 0.17842 |
| SF3A1 | red | 0.272099 | 0.17871 |
| GNAI1 | red | -0.271946 | 0.178964 |
| HELLS | red | -0.271753 | 0.179285 |
| FBXO8 | red | -0.271323 | 0.180002 |
| TMEM206 | red | -0.270735 | 0.180984 |
| GCHFR | red | -0.269728 | 0.182675 |
| ARAP2 | red | -0.269579 | 0.182926 |
| RIOK3 | red | -0.269578 | 0.182928 |
| DUSP8 | red | 0.269332 | 0.183343 |
| PIK3C2A | red | -0.268856 | 0.184149 |
| PER3 | red | -0.268597 | 0.184589 |
| RFC3 | red | -0.268497 | 0.184758 |
| GRB2 | red | -0.268209 | 0.185248 |
| HSP90AA1 | red | -0.268081 | 0.185465 |
| RAP1GDS1 | red | -0.267815 | 0.185919 |
| MMS22L | red | -0.26718 | 0.187005 |
| MAFF | red | 0.266938 | 0.187419 |
| ANGEL2 | red | -0.266537 | 0.188107 |
| PTPN9 | red | -0.266012 | 0.189012 |
| C3orf62 | red | -0.265775 | 0.189423 |
| METTL21A | red | -0.265402 | 0.190068 |
| C20orf72 | red | -0.265397 | 0.190075 |
| PI4KB | red | 0.265347 | 0.190162 |
| GPS1 | red | 0.265324 | 0.190202 |
| TRAPPC1 | red | -0.26458 | 0.191494 |
| TMEM68 | red | -0.26453 | 0.191582 |
| LRRC8B | red | -0.263641 | 0.193134 |
| PON2 | red | -0.263207 | 0.193895 |
| ACADM | red | -0.261927 | 0.196152 |
| SERF1A | red | -0.261695 | 0.196563 |
| UCHL3 | red | -0.261072 | 0.197669 |
| CNDP2 | red | -0.260882 | 0.198008 |
| LINC00174 | red | -0.260873 | 0.198024 |
| C5orf15 | red | -0.260142 | 0.199328 |
| REEP3 | red | -0.259113 | 0.201176 |
| DYX1C1 | red | -0.258545 | 0.202201 |
| TAX1BP1 | red | -0.258539 | 0.202212 |
| PTDSS2 | red | 0.258503 | 0.202277 |
| FAM83H | red | 0.258373 | 0.202513 |
| MAP3K1 | red | -0.258133 | 0.202947 |
| C4orf34 | red | -0.257636 | 0.203849 |
| RPL24 | red | -0.256901 | 0.205188 |
| MPHOSPH9 | red | -0.256193 | 0.206482 |
| ESD | red | -0.255082 | 0.208525 |
| CRBN | red | -0.254596 | 0.209423 |
| EPM2A | red | -0.254158 | 0.210237 |
| ST7 | red | -0.254137 | 0.210274 |
| DBNL | red | 0.253992 | 0.210545 |
| WBP2 | red | 0.253725 | 0.211041 |
| LARP7 | red | -0.253609 | 0.211256 |
| MDH1 | red | -0.251499 | 0.215209 |
| ADSL | red | -0.25076 | 0.216605 |
| HGSNAT | red | -0.250476 | 0.217143 |
| DDX17 | purple | 0.084645021 | 0.674652688 |
| DBNDD2 | purple | 0.090112695 | 0.654879277 |
| EXOSC9 | purple | 0.09581979 | 0.634479776 |
| BOLA3 | purple | 0.096361777 | 0.632555677 |
| GPR34 | purple | 0.101188249 | 0.615525339 |
| NUP54 | purple | 0.102616447 | 0.610522428 |
| DPY30 | purple | 0.106145339 | 0.598234229 |
| H6PD | purple | 0.110269275 | 0.584009256 |
| DHX38 | purple | 0.112460963 | 0.576510025 |
| TAF1A | purple | 0.114454765 | 0.569725171 |
| COPS2 | purple | 0.121027363 | 0.547616065 |
| UBE2Z | purple | 0.130925127 | 0.515093761 |
| ATP6V0C | purple | 0.133428501 | 0.50702019 |
| GPR180 | purple | 0.139883398 | 0.486494599 |
| C8orf76 | purple | 0.142828046 | 0.477273259 |
| FAM217B | purple | 0.144568415 | 0.471865727 |
| RGS10 | purple | 0.151506278 | 0.450627717 |
| COG7 | purple | 0.154923719 | 0.440356397 |
| HNRPDL | purple | 0.162332307 | 0.418528502 |
| ZCCHC4 | purple | 0.168146421 | 0.401825959 |
| BIRC2 | purple | 0.168412545 | 0.401070552 |
| SNRPE | purple | 0.170430661 | 0.395368137 |
| TAMM41 | purple | 0.176299104 | 0.379049805 |
| NFKBIZ | purple | 0.194571454 | 0.330792174 |
| ICT1 | purple | 0.19627631 | 0.326489088 |
| SBDSP1 | purple | 0.206069585 | 0.30243649 |
| MOB4 | purple | 0.206946588 | 0.300338005 |
| LOC388796 | purple | 0.211933782 | 0.288578362 |
| UBE2T | purple | 0.212099521 | 0.288192629 |
| DYNC2LI1 | purple | 0.213133995 | 0.285792415 |
| SBDS | purple | 0.215248813 | 0.280925133 |
| TM2D1 | purple | 0.222893143 | 0.26377468 |
| KATNA1 | purple | 0.233621622 | 0.24087256 |
| MTIF3 | purple | 0.242321241 | 0.223296611 |
| MINPP1 | purple | 0.255289012 | 0.198732209 |
| MRPL33 | purple | 0.280864844 | 0.155866663 |
| METTL5 | purple | 0.292525167 | 0.138688835 |
| TMEM126A | purple | 0.294847741 | 0.135437619 |
| DBI | purple | 0.300529485 | 0.127717612 |
| SLIRP | purple | 0.331248293 | 0.091449222 |
| FBXO22 | purple | 0.343005988 | 0.079849859 |
| RPL23 | purple | 0.353492874 | 0.070482701 |
| COMMD3 | purple | 0.369381429 | 0.057930283 |
| CCDC90A | purple | 0.382509682 | 0.048936276 |
| CCDC146 | purple | 0.401373754 | 0.037978882 |
| BCL3 | red | -0.621002635 | 0.000546981 |
| SGTB | red | -0.620828262 | 0.000549526 |
| RAF1 | red | -0.590509453 | 0.001184027 |
| SMARCD1 | red | -0.569985071 | 0.001910382 |
| CXXC5 | red | -0.567973317 | 0.00199879 |
| PRRC2C | red | -0.566174864 | 0.002080782 |
| ZFR | red | -0.562818776 | 0.00224154 |
| LRRC33 | red | -0.562016893 | 0.002281492 |
| TMEM43 | red | -0.560021389 | 0.002383576 |
| MAPKBP1 | red | -0.559950817 | 0.002387257 |
| TSPAN17 | red | -0.554443152 | 0.002689992 |
| ORMDL2 | salmon | 0.040247 | 0.84523 |
| RGL3 | salmon | 0.04027 | 0.845146 |
| SPIRE1 | salmon | 0.044423 | 0.829395 |
| C14orf2 | salmon | 0.046983 | 0.819717 |
| SCAMP3 | salmon | 0.052489 | 0.798989 |
| RAB4B | salmon | 0.053769 | 0.79419 |
| GADD45GIP | salmon | 0.055209 | 0.788798 |
| SENP5 | salmon | 0.058256 | 0.777422 |
| PINLYP | salmon | 0.058547 | 0.776337 |
| SEC24D | salmon | 0.063199 | 0.759064 |
| COX6A1 | salmon | 0.066656 | 0.746299 |
| VIPAS39 | salmon | 0.067186 | 0.744345 |
| SELK | salmon | 0.069554 | 0.735648 |
| TMEM223 | salmon | 0.069629 | 0.735372 |
| MRPL14 | salmon | 0.077374 | 0.707147 |
| RBM3 | salmon | 0.078284 | 0.703854 |
| UBA52 | salmon | 0.080833 | 0.694659 |
| BANF1 | salmon | 0.082024 | 0.690375 |
| ASNA1 | salmon | 0.083782 | 0.684073 |
| PARK7 | salmon | 0.086568 | 0.674126 |
| RANGAP1 | salmon | 0.089095 | 0.665148 |
| ZNF646 | salmon | 0.092031 | 0.654775 |
| ENY2 | salmon | 0.099191 | 0.629738 |
| RPL23A | salmon | 0.102491 | 0.618332 |
| SNRPG | salmon | 0.104573 | 0.611176 |
| DRG1 | salmon | 0.108216 | 0.598745 |
| RDH14 | salmon | 0.111253 | 0.588463 |
| EXOC4 | salmon | 0.113036 | 0.58246 |
| RNF40 | salmon | 0.11446 | 0.577687 |
| KLHDC4 | salmon | 0.115357 | 0.57469 |
| POLDIP2 | salmon | 0.116076 | 0.572291 |
| SERPINA1 | salmon | 0.117657 | 0.567033 |
| LYRM7 | salmon | 0.126303 | 0.538673 |
| NDEL1 | salmon | 0.12757 | 0.534576 |
| WBSCR22 | salmon | 0.129369 | 0.52878 |
| FUT4 | salmon | 0.132345 | 0.519261 |
| SUPT4H1 | salmon | 0.138305 | 0.500449 |
| NAALADL1 | salmon | 0.141739 | 0.489765 |
| CALM3 | salmon | 0.141933 | 0.489164 |
| SFT2D1 | salmon | 0.143085 | 0.485607 |
| TIMM23 | salmon | 0.148844 | 0.468029 |
| ARHGAP25 | salmon | 0.151232 | 0.460839 |
| NOP16 | salmon | 0.153046 | 0.455416 |
| ARL8A | salmon | 0.154071 | 0.452365 |
| AKT2 | salmon | 0.157159 | 0.443244 |
| MBD3 | salmon | 0.157466 | 0.442343 |
| TIMM10 | salmon | 0.158401 | 0.439601 |
| PSMC1 | salmon | 0.160569 | 0.433285 |
| COQ6 | salmon | 0.161412 | 0.430843 |
| KAT5 | salmon | 0.166637 | 0.415869 |
| CD151 | salmon | 0.169363 | 0.408174 |
| RABAC1 | salmon | 0.172059 | 0.400639 |
| VPS11 | salmon | 0.184527 | 0.366836 |
| TP53BP1 | salmon | 0.185125 | 0.365258 |
| DGCR14 | salmon | 0.186722 | 0.361063 |
| PSMG4 | salmon | 0.19506 | 0.339622 |

**Table 6. Gene module membership and correlation with growth rate (48/72h), with adjusted p values for each module**

| | | | | | | | | | | | |
|---|---|---|---|---|---|---|---|---|---|---|---|
| DNAJC25 | red | -0.24993 | 0.218182 | PSMD2 | red | -0.545643525 | 0.003241846 | CYLD | salmon | 0.195706 | 0.337993 |
| IL1RAP | red | -0.249583 | 0.218842 | RNF24 | red | -0.53561782 | 0.003985777 | TCEB2 | salmon | 0.198984 | 0.329801 |
| PAIP2 | red | -0.247953 | 0.221965 | HEATR6 | red | -0.534209478 | 0.004101093 | RBX1 | salmon | 0.201937 | 0.322526 |
| ME2 | red | -0.247808 | 0.222243 | SLC35B2 | red | -0.52907144 | 0.004546178 | CCDC71 | salmon | 0.20562 | 0.31359 |
| WDSUB1 | red | -0.247553 | 0.222735 | EML4 | red | -0.524961929 | 0.004931087 | HMGXB4 | salmon | 0.206788 | 0.310788 |
| SNF8 | red | 0.246632 | 0.224518 | PRKCSH | red | -0.523844861 | 0.005040368 | CHCHD5 | salmon | 0.208967 | 0.305605 |
| SSB | red | -0.246616 | 0.224547 | FLAD1 | red | -0.523589809 | 0.005065606 | OTUD5 | salmon | 0.214944 | 0.291659 |
| GART | red | -0.246568 | 0.224641 | DARS2 | red | -0.523152033 | 0.005109173 | LMAN2 | salmon | 0.21896 | 0.282518 |
| TCP11L1 | red | -0.245157 | 0.227391 | POLR3C | red | -0.522470889 | 0.005177592 | COX4I1 | salmon | 0.227569 | 0.263542 |
| BCOR | red | -0.244511 | 0.228656 | SLC23A2 | red | -0.520241893 | 0.005406939 | CLASP1 | salmon | 0.231816 | 0.254492 |
| DDX21 | red | -0.2444 | 0.228875 | NINJ1 | red | -0.518408396 | 0.005601985 | ANKZF1 | salmon | 0.234128 | 0.24965 |
| PTPN12 | red | -0.244351 | 0.228971 | RANBP9 | red | -0.515949558 | 0.00587287 | SUMO2 | salmon | 0.235267 | 0.247288 |
| RABEPK | red | -0.243673 | 0.230307 | GTF3C5 | red | -0.507928858 | 0.006834725 | UBE2D3 | salmon | 0.251393 | 0.215409 |
| TMEM189 | red | 0.243142 | 0.231355 | ARL15 | red | -0.50622344 | 0.007055491 | BAP1 | salmon | 0.251747 | 0.214743 |
| OC1005057( | red | -0.243055 | 0.231526 | UQCR10 | red | -0.503333722 | 0.007443267 | CPSF3L | salmon | 0.254526 | 0.209554 |
| MED29 | red | 0.242733 | 0.232166 | BCAP31 | red | -0.495888119 | 0.008525812 | INTS9 | salmon | 0.265581 | 0.189758 |
| WDR3 | red | -0.24272 | 0.232191 | ANPEP | red | -0.493305991 | 0.008930788 | ENDOG | salmon | 0.272258 | 0.178447 |
| SKA2 | red | -0.242668 | 0.232294 | NRBP1 | red | -0.492328446 | 0.00908825 | SLC25A28 | salmon | 0.275775 | 0.172681 |
| SEPSECS | red | -0.242629 | 0.232373 | PRPF38A | red | -0.491427831 | 0.009235371 | STX5 | salmon | 0.275817 | 0.172612 |
| APBB1IP | red | -0.241796 | 0.234029 | POLR3H | red | -0.491193951 | 0.009273901 | ERGIC3 | salmon | 0.286009 | 0.15665 |
| ZNF568 | red | -0.241672 | 0.234276 | SEC24A | red | -0.491152965 | 0.009280667 | VPS39 | salmon | 0.288229 | 0.153315 |
| CLNS1A | red | -0.240218 | 0.23719 | UBAC2 | red | -0.488168845 | 0.009784508 | CHMP6 | salmon | 0.303783 | 0.131368 |
| RBMX2 | red | -0.239816 | 0.237999 | BET1L | red | -0.486060597 | 0.010154079 | SYS1 | salmon | 0.316031 | 0.115761 |
| EMR2 | red | 0.23892 | 0.23981 | MSANTD3 | red | -0.485809632 | 0.010198839 | RNASEH2C | salmon | 0.325015 | 0.105213 |
| OC1001327( | red | -0.238878 | 0.239896 | NF1 | red | -0.484303228 | 0.010470981 | DOK1 | salmon | 0.330929 | 0.098671 |
| C9orf156 | red | -0.237809 | 0.242068 | DENND2D | red | -0.483617373 | 0.010596878 | PSMD8 | salmon | 0.336714 | 0.09257 |
| DAP | red | 0.236369 | 0.245017 | ORMDL1 | red | -0.483613964 | 0.010597507 | NFKBIL1 | salmon | 0.351635 | 0.078143 |
| TGFBRAP1 | red | -0.236069 | 0.245634 | LRCH3 | red | -0.480250052 | 0.01123345 | ASCC2 | salmon | 0.366799 | 0.065303 |
| DENND1A | red | -0.235978 | 0.24582 | ABHD13 | red | -0.479672465 | 0.011345776 | CSTF2T | salmon | 0.425153 | 0.030374 |
| SAR1B | red | -0.235798 | 0.246191 | UBE2A | red | -0.478157621 | 0.011644821 | LAMTOR2 | salmon | 0.465096 | 0.01666 |
| DVL1 | red | 0.235591 | 0.246618 | GPN1 | red | -0.475342608 | 0.012217956 | FAR1 | skyblue | -0.288172 | 0.153401 |
| DERL2 | red | -0.234595 | 0.248681 | XRN2 | red | -0.472745677 | 0.012767288 | AKAP11 | skyblue | -0.280902 | 0.164512 |
| MTHFD1L | red | -0.234383 | 0.24912 | FAM58A | red | -0.471758274 | 0.012981461 | DOCK10 | skyblue | -0.277278 | 0.170258 |
| ANKIB1 | red | -0.234233 | 0.249433 | PLK3 | red | -0.47116971 | 0.013110536 | EI24 | skyblue | -0.275014 | 0.173916 |
| GABARAP | red | -0.234008 | 0.2499 | BMI1 | red | -0.468811543 | 0.013638417 | RAD54L2 | skyblue | -0.229956 | 0.25843 |
| QPCTL | red | 0.232398 | 0.253268 | FOSL2 | red | -0.467413068 | 0.013959701 | GTDC1 | skyblue | -0.215615 | 0.290119 |
| CRNKL1 | red | -0.232031 | 0.254038 | ATG9A | red | -0.464872863 | 0.01455931 | MED27 | skyblue | -0.203732 | 0.318152 |
| ERCC1 | red | -0.231785 | 0.254558 | ATXN2 | red | -0.463291233 | 0.014943294 | CYCS | skyblue | -0.192868 | 0.345183 |
| UQCRHL | red | -0.231714 | 0.254707 | VPRBP | red | -0.463104538 | 0.014989166 | ZNF84 | skyblue | -0.188195 | 0.357219 |
| MPRIP | red | -0.231388 | 0.255394 | PTPN9 | red | -0.462411593 | 0.015160446 | MRPL20 | skyblue | -0.17518 | 0.392018 |
| SEC11C | red | -0.231015 | 0.256182 | ALG9 | red | -0.460064829 | 0.015752561 | ALG14 | skyblue | -0.168795 | 0.409769 |
| PPP4R1L | red | -0.230931 | 0.256361 | MGAT5 | red | -0.45995364 | 0.015781081 | ZC3H18 | skyblue | -0.166385 | 0.416584 |
| ZSWIM3 | red | -0.230574 | 0.257117 | ARFGEF2 | red | -0.457543593 | 0.016409822 | SCAMP1 | skyblue | -0.150507 | 0.463016 |
| PREX1 | red | -0.229903 | 0.258543 | ZBTB49 | red | -0.456464034 | 0.016698079 | SLC22A15 | skyblue | -0.150369 | 0.463431 |
| GDAP2 | red | -0.229634 | 0.259115 | APH1A | red | -0.454677185 | 0.017184345 | DESI2 | skyblue | -0.150158 | 0.464066 |
| ATF6 | red | -0.228846 | 0.260799 | SLC15A4 | red | -0.452463787 | 0.017802784 | NHLRC2 | skyblue | -0.139432 | 0.496928 |
| CDK17 | red | -0.22704 | 0.264684 | GNL3 | red | -0.451116034 | 0.018188233 | MED23 | skyblue | -0.133853 | 0.514469 |
| LYRM9 | red | -0.226882 | 0.265026 | QRSL1 | red | -0.450946828 | 0.018237106 | KAT2B | skyblue | -0.129691 | 0.527746 |
| POFUT1 | red | -0.225635 | 0.267732 | PITHD1 | red | -0.450864685 | 0.01826087 | IFI30 | skyblue | -0.116495 | 0.570898 |
| SYTL3 | red | -0.225412 | 0.268218 | STRN | red | -0.449368754 | 0.01869812 | METTL15 | skyblue | -0.110815 | 0.589941 |
| RAB7L1 | red | 0.225085 | 0.268931 | SAV1 | red | -0.448706304 | 0.018894475 | IDS | skyblue | -0.107079 | 0.602614 |
| ENPP5 | red | -0.224711 | 0.269748 | C10orf32 | red | -0.44821011 | 0.019042655 | NAV1 | skyblue | -0.101982 | 0.620085 |
| SMG7 | red | -0.22458 | 0.270035 | RUNX1 | red | -0.446061159 | 0.019695448 | ZNF624 | skyblue | -0.099635 | 0.628198 |
| VASH1 | red | 0.224162 | 0.270951 | POLR1D | red | -0.444342924 | 0.020230488 | MSR1 | skyblue | -0.097777 | 0.634652 |
| SKI | red | -0.223642 | 0.272093 | IBTK | red | -0.443996713 | 0.020339721 | SYNJ1 | skyblue | -0.097237 | 0.636533 |
| NSDHL | red | -0.223631 | 0.272116 | TTI1 | red | -0.443132154 | 0.020614607 | HECTD4 | skyblue | -0.097208 | 0.636636 |

**Table 6. Gene module membership and correlation with growth rate (48/72h), with adjusted p values for each module**

| | | | | | | | | | | | |
|---|---|---|---|---|---|---|---|---|---|---|---|
| FAM179B | red | -0.223422 | 0.272578 | RBM18 | red | -0.44040707 | 0.021500989 | C12orf73 | skyblue | -0.095963 | 0.64098 |
| CHMP6 | red | 0.222415 | 0.274801 | SPOP | red | -0.440223709 | 0.021561731 | TRIT1 | skyblue | -0.086274 | 0.675171 |
| LMBRD1 | red | -0.221487 | 0.276861 | EMD | red | -0.438201728 | 0.022240888 | ZC3H11A | skyblue | -0.066695 | 0.746155 |
| ARHGAP11E | red | -0.220233 | 0.279659 | TMEM64 | red | -0.43502261 | 0.023343914 | GFPT1 | skyblue | -0.055485 | 0.787767 |
| GORASP2 | red | -0.219772 | 0.280693 | SETD3 | red | -0.432889924 | 0.024108528 | GALK1 | skyblue | -0.053014 | 0.797021 |
| STK19 | red | -0.219325 | 0.281697 | PTBP3 | red | -0.430577879 | 0.024960326 | WDFY3 | skyblue | -0.043355 | 0.833437 |
| CDC5L | red | -0.219075 | 0.282259 | TNPO2 | red | -0.429638225 | 0.025313423 | ABT1 | skyblue | -0.038806 | 0.85071 |
| TELO2 | red | -0.218628 | 0.283266 | MED30 | red | -0.429572583 | 0.02533824 | BMPR2 | skyblue | -0.033301 | 0.871705 |
| XPOT | red | -0.21836 | 0.283871 | ZCCHC14 | red | -0.429281527 | 0.025448517 | NFAT5 | skyblue | -0.028717 | 0.889247 |
| MTX2 | red | -0.216773 | 0.287473 | RNF10 | red | -0.427887378 | 0.025982157 | ZNF652 | skyblue | -0.027283 | 0.894748 |
| DLD | red | -0.2157 | 0.289924 | PRDM10 | red | -0.425330941 | 0.026984247 | C21orf33 | skyblue | -0.016023 | 0.938077 |
| KIAA0586 | red | -0.215465 | 0.290464 | INTS6 | red | -0.423604243 | 0.027678627 | SMAD5 | skyblue | -0.015979 | 0.938244 |
| BAK1 | red | 0.215317 | 0.290802 | TINF2 | red | -0.423525964 | 0.027710445 | RGMB | skyblue | -0.012418 | 0.951989 |
| ELOVL5 | red | 0.214621 | 0.292403 | CNOT6 | red | -0.422146218 | 0.028276153 | CHMP2A | skyblue | -0.008413 | 0.967463 |
| MS4A4A | red | -0.213966 | 0.293914 | DOPEY2 | red | -0.42110104 | 0.028710869 | FBXO10 | skyblue | -0.000844 | 0.996733 |
| PPM1M | red | 0.213276 | 0.295509 | PER3 | red | -0.420626704 | 0.02890993 | ARPC2 | skyblue | 0.001673 | 0.993526 |
| PSMA6 | red | -0.212151 | 0.298124 | LMTK2 | red | -0.419947479 | 0.029196913 | DST | skyblue | 0.002253 | 0.991285 |
| PRMT1 | red | 0.211897 | 0.298718 | DAGLB | red | -0.419907025 | 0.029214078 | PDXP | skyblue | 0.003845 | 0.985127 |
| CDC37L1 | red | -0.210191 | 0.302715 | TADA2A | red | -0.419638126 | 0.02932838 | KANSL1 | skyblue | 0.015023 | 0.941934 |
| ZNF441 | red | 0.209087 | 0.305321 | BRPF1 | red | -0.418437147 | 0.029843298 | MRTO4 | skyblue | 0.020407 | 0.921179 |
| TH1L | red | -0.20831 | 0.307162 | ARNTL | red | -0.415100975 | 0.031312027 | LONRF3 | skyblue | 0.023217 | 0.910364 |
| PNKP | red | 0.207842 | 0.308273 | DTD1 | red | -0.413730035 | 0.031932189 | EXOSC7 | skyblue | 0.023322 | 0.909963 |
| PPP1R12A | red | 0.206162 | 0.312288 | RASSF3 | red | -0.411981206 | 0.032737589 | FAM207A | skyblue | 0.038577 | 0.851584 |
| DCUN1D5 | red | -0.205658 | 0.3135 | USP54 | red | -0.409884865 | 0.033724447 | CERS5 | skyblue | 0.054336 | 0.792066 |
| TEFM | red | -0.205572 | 0.313706 | MTRR | red | -0.407979269 | 0.034642084 | NDUFA1 | skyblue | 0.059245 | 0.773739 |
| MMGT1 | red | -0.20533 | 0.314289 | HNRNPH1 | red | -0.407222322 | 0.035012095 | PUM2 | skyblue | 0.065565 | 0.750322 |
| ICOSLG | red | 0.205189 | 0.314629 | TTC5 | red | -0.405879228 | 0.035676413 | PSMA5 | skyblue | 0.110957 | 0.589462 |
| TMPO | red | -0.205138 | 0.314752 | TM6SF1 | red | -0.402508373 | 0.03738818 | THRAP3 | skyblue | 0.130932 | 0.52377 |
| MX2 | red | -0.205053 | 0.314957 | FAM168A | red | -0.401182506 | 0.038079178 | TMSB10 | skyblue | 0.138263 | 0.50058 |
| SRSF10 | red | -0.204645 | 0.315941 | INPP5F | red | -0.399272154 | 0.039092662 | ISCA2 | skyblue | 0.139442 | 0.496898 |
| LRRC6 | red | -0.203882 | 0.317789 | ALG8 | red | -0.399260552 | 0.039098883 | TSR3 | skyblue | 0.149715 | 0.465401 |
| ATF6B | red | 0.203407 | 0.318941 | EIF2B2 | red | -0.397850022 | 0.039860997 | DPM2 | skyblue | 0.160852 | 0.432463 |
| SREK1IP1 | red | -0.202996 | 0.31994 | PRKCH | red | -0.397671796 | 0.039958128 | DAK | skyblue | 0.192579 | 0.34592 |
| C2CD2 | red | -0.202033 | 0.32229 | SOS1 | red | -0.3970908 | 0.040276075 | CDYL | skyblue | 0.193598 | 0.343325 |
| HIST2H2BA | red | -0.201968 | 0.322451 | MTMR9 | red | -0.396095963 | 0.040825163 | TMEM160 | skyblue | 0.199517 | 0.328482 |
| MGST1 | red | -0.201827 | 0.322795 | LOC339803 | red | -0.394342594 | 0.041807387 | ZNF593 | skyblue | 0.201747 | 0.322992 |
| C1RL | red | 0.201636 | 0.323262 | ZDHHC17 | red | -0.39341095 | 0.042336868 | CLIC1 | skyblue | 0.205252 | 0.314478 |
| CENPQ | red | -0.201462 | 0.32369 | POLE3 | red | -0.390828667 | 0.043832309 | COQ4 | skyblue | 0.211555 | 0.299515 |
| VWA8 | red | -0.201202 | 0.324328 | YWHAB | red | -0.38883331 | 0.045016261 | TEN1 | skyblue | 0.216138 | 0.288922 |
| FBXO3 | red | -0.200356 | 0.32641 | C5orf22 | red | -0.384434071 | 0.04771593 | CCRL2 | skyblue | 0.226155 | 0.266601 |
| TCAIM | red | -0.199974 | 0.327352 | STARD3 | red | -0.383188698 | 0.048502905 | STK16 | skyblue | 0.280272 | 0.165501 |
| PAG1 | red | 0.198266 | 0.331584 | PPP1R8 | red | -0.383186477 | 0.048504317 | B4GALT3 | skyblue | 0.435274 | 0.026252 |
| DDX60L | red | -0.198091 | 0.33202 | RAD54L2 | red | -0.383070648 | 0.04857803 | ZNF434 | steelblue | -0.38053 | 0.055142 |
| PSMD10 | red | -0.198086 | 0.332032 | ERCC6 | red | -0.382231627 | 0.049114617 | EFTUD2 | steelblue | -0.369736 | 0.063017 |
| MPC1 | red | -0.197908 | 0.332476 | ABCG1 | red | -0.381538338 | 0.049561513 | YLPM1 | steelblue | -0.320524 | 0.110394 |
| IFI35 | red | -0.195098 | 0.339526 | FLJ31306 | red | -0.381487627 | 0.049594327 | POMP | steelblue | -0.285501 | 0.157419 |
| SUPT4H1 | red | 0.195094 | 0.339535 | PEX26 | red | -0.381070281 | 0.04986503 | LETM1 | steelblue | -0.25698 | 0.205043 |
| TTC21B | red | -0.193952 | 0.342425 | FLI1 | red | -0.377692228 | 0.05209914 | MRPS31 | steelblue | -0.245446 | 0.226825 |
| RB1 | red | -0.193303 | 0.344074 | EIF2B1 | red | -0.377343354 | 0.052334274 | ATP5G3 | steelblue | -0.236674 | 0.244391 |
| TMX3 | red | -0.19284 | 0.345254 | FNBP1 | red | -0.37436714 | 0.054374182 | DNAJC16 | steelblue | -0.235633 | 0.246532 |
| USP15 | red | -0.192113 | 0.347111 | ZNF529 | red | -0.373021717 | 0.055316523 | SERINC1 | steelblue | -0.225091 | 0.268917 |
| CLDN23 | red | -0.190629 | 0.350918 | STAM | red | -0.371382127 | 0.056482107 | NABP2 | steelblue | -0.210182 | 0.302737 |
| NKIRAS1 | red | -0.190306 | 0.351752 | WRAP73 | red | -0.37008866 | 0.057415094 | TUBGCP2 | steelblue | -0.209647 | 0.303997 |
| TALDO1 | red | 0.189328 | 0.354278 | RBBP8 | red | -0.369447252 | 0.057882183 | AHNAK | steelblue | -0.208826 | 0.305939 |
| ITPK1 | red | 0.18932 | 0.354299 | RDH10 | red | -0.367010696 | 0.059683597 | EIF2AK4 | steelblue | -0.205223 | 0.314547 |
| DPP8 | red | -0.189312 | 0.354319 | SURF4 | red | -0.36581129 | 0.060586227 | AHCTF1 | steelblue | -0.203859 | 0.317843 |

**Table 6. Gene module membership and correlation with growth rate (48/72h), with adjusted p values for each module**

| | | | | | | | | | | | |
|---|---|---|---|---|---|---|---|---|---|---|---|
| ZRSR2 | red | -0.18804 | 0.357621 | CRAT | red | -0.363900898 | 0.06204579 | HS2ST1 | steelblue | -0.199557 | 0.328381 |
| N4BP2 | red | -0.187491 | 0.359053 | CPNE3 | red | -0.358228046 | 0.066541143 | GIT2 | steelblue | -0.198411 | 0.331224 |
| MMADHC | red | -0.187271 | 0.359627 | PRR13 | red | -0.357206172 | 0.06737699 | USP24 | steelblue | -0.188108 | 0.357444 |
| ALPK3 | red | 0.187211 | 0.359784 | CDC25B | red | -0.354008462 | 0.07004501 | VPS36 | steelblue | -0.178242 | 0.383663 |
| TOMM40 | red | 0.186867 | 0.360682 | ALG11 | red | -0.353374913 | 0.070583135 | HK1 | steelblue | -0.165065 | 0.420343 |
| UBXN2A | red | -0.186218 | 0.362384 | ACOT4 | red | -0.352551726 | 0.07128708 | VEZF1 | steelblue | -0.16475 | 0.421243 |
| SIRPB2 | red | -0.1862 | 0.36243 | IP6K1 | red | -0.352223274 | 0.071569457 | R3HDM2 | steelblue | -0.149639 | 0.46563 |
| CPEB3 | red | -0.185923 | 0.363159 | LDHA | red | -0.35125879 | 0.072403613 | SND1 | steelblue | -0.129681 | 0.527777 |
| RPAP2 | red | -0.18461 | 0.366616 | MET | red | -0.349791351 | 0.073687061 | KIAA1468 | steelblue | -0.123689 | 0.547177 |
| MOB3B | red | 0.183713 | 0.368991 | YRDC | red | -0.347840496 | 0.075420251 | SCAF11 | steelblue | -0.110353 | 0.5915 |
| CHCHD2 | red | 0.183634 | 0.369199 | CTNNA1 | red | -0.347487108 | 0.075737521 | RAB3GAP2 | steelblue | -0.106069 | 0.60606 |
| ERAP1 | red | -0.183237 | 0.370255 | FARS2 | red | -0.344759956 | 0.078220433 | ELP6 | steelblue | -0.104439 | 0.611636 |
| TIMP1 | red | 0.182709 | 0.371658 | MGAT4B | red | -0.343966561 | 0.078954318 | TNKS2 | steelblue | -0.103869 | 0.613591 |
| PPA2 | red | -0.182635 | 0.371855 | OTUD4 | red | -0.343957356 | 0.078962863 | PNN | steelblue | -0.102695 | 0.617628 |
| BRMS1 | red | 0.181571 | 0.374695 | FAM127B | red | -0.342906522 | 0.079943031 | COG2 | steelblue | -0.083509 | 0.685048 |
| QTRTD1 | red | -0.180849 | 0.376631 | TMBIM6 | red | -0.341999719 | 0.080796287 | FAM82A2 | steelblue | -0.082018 | 0.690396 |
| CALM3 | red | 0.18015 | 0.378509 | ANKRD49 | red | -0.34089906 | 0.081841252 | DNAL4 | steelblue | -0.074857 | 0.71628 |
| ARHGAP10 | red | -0.179785 | 0.379491 | INSIG1 | red | -0.339836239 | 0.082860024 | MYCBP2 | steelblue | -0.067745 | 0.742289 |
| PKD1P1 | red | 0.179762 | 0.379554 | LOC92249 | red | -0.33922673 | 0.083448606 | QKI | steelblue | -0.059454 | 0.772962 |
| SERTAD3 | red | -0.179305 | 0.380786 | TMEM230 | red | -0.336357338 | 0.08626225 | CD47 | steelblue | -0.056627 | 0.7835 |
| ZNF41 | red | 0.179159 | 0.381182 | DCTN4 | red | -0.334725633 | 0.087893997 | MRPL43 | steelblue | -0.048999 | 0.812114 |
| HS1BP3 | red | -0.178676 | 0.382486 | CLASP2 | red | -0.333773732 | 0.088856646 | CBWD5 | steelblue | -0.042975 | 0.834878 |
| DYRK4 | red | -0.17705 | 0.386904 | CHAMP1 | red | -0.332963722 | 0.089682061 | LAMP2 | steelblue | -0.01521 | 0.941214 |
| AKAP13 | red | -0.176953 | 0.387167 | TCF12 | red | -0.332297525 | 0.090365258 | TET2 | steelblue | -0.007685 | 0.970279 |
| ENOPH1 | red | -0.174629 | 0.393534 | MRPL48 | red | -0.329951548 | 0.092802405 | RNF41 | steelblue | 0.001207 | 0.995331 |
| SNRNP27 | red | -0.174443 | 0.394046 | ARPP19 | red | -0.329685994 | 0.093081367 | DCAF11 | steelblue | 0.013471 | 0.947925 |
| DDX27 | red | 0.174363 | 0.394265 | SNX27 | red | -0.329571538 | 0.093201797 | INVS | steelblue | 0.014132 | 0.945375 |
| EMC8 | red | 0.173545 | 0.396521 | PSMA3 | red | -0.329046326 | 0.093755925 | C1D | steelblue | 0.014467 | 0.944079 |
| SLC46A2 | red | 0.173361 | 0.397029 | TSC1 | red | -0.324349839 | 0.098821741 | TBCA | steelblue | 0.014928 | 0.942303 |
| SETD1B | red | 0.172651 | 0.398998 | MCL1 | red | -0.320634258 | 0.10297262 | MAF1 | steelblue | 0.01712 | 0.933845 |
| PIK3CA | red | -0.172057 | 0.400645 | SDHC | red | -0.316002095 | 0.108327974 | SDCCAG8 | steelblue | 0.026831 | 0.896482 |
| GIGYF2 | red | -0.171873 | 0.401157 | IGSF6 | red | -0.315684521 | 0.108702558 | DDX19B | steelblue | 0.028791 | 0.888963 |
| RAD17 | red | -0.171222 | 0.402972 | FKBP11 | red | -0.313685517 | 0.111082529 | TIMM17B | steelblue | 0.032228 | 0.875803 |
| CXorf38 | red | -0.169625 | 0.407438 | INPP5A | red | -0.313134102 | 0.111745772 | GOLGA7 | steelblue | 0.047668 | 0.81713 |
| FGFR1OP | red | -0.16937 | 0.408153 | BABAM1 | red | -0.313044631 | 0.111853663 | TDG | steelblue | 0.054122 | 0.792868 |
| SPRTN | red | -0.167392 | 0.413729 | RPL10 | red | -0.311302554 | 0.113969802 | GNB2L1 | steelblue | 0.062232 | 0.762644 |
| DDX54 | red | 0.166671 | 0.415774 | PLA2G7 | red | -0.310484162 | 0.114974068 | ACP6 | steelblue | 0.070462 | 0.732318 |
| PLEKHG3 | red | -0.166273 | 0.416904 | CD109 | red | -0.306287511 | 0.120226707 | SPRYD3 | steelblue | 0.091556 | 0.656449 |
| CTDSP1 | red | 0.164283 | 0.42258 | DDX60L | red | -0.303577886 | 0.123710509 | NUDT4 | steelblue | 0.093024 | 0.65128 |
| MAP2K2 | red | 0.163772 | 0.424043 | KLHL36 | red | -0.302993348 | 0.124471653 | ICT1 | steelblue | 0.094365 | 0.646573 |
| GTPBP8 | red | -0.163685 | 0.424294 | TAOK2 | red | -0.301280353 | 0.126721926 | MRPL50 | steelblue | 0.113302 | 0.581567 |
| NR1D2 | red | -0.163673 | 0.424326 | VRK3 | red | -0.300597706 | 0.127626913 | PQLC2 | steelblue | 0.134991 | 0.510866 |
| TBCD | red | -0.162671 | 0.427207 | LCMT1 | red | -0.299693372 | 0.128833044 | INPP5K | steelblue | 0.165055 | 0.420371 |
| MED13L | red | -0.162657 | 0.427248 | CSRNP1 | red | -0.298954831 | 0.1298242 | ASB8 | steelblue | 0.1708 | 0.404147 |
| EEF1D | red | 0.161658 | 0.430131 | METTL17 | red | -0.298412443 | 0.130555638 | CLEC16A | steelblue | 0.174691 | 0.393363 |
| GET4 | red | 0.161266 | 0.431265 | UBE2F | red | -0.295413535 | 0.134654041 | UBA7 | steelblue | 0.176588 | 0.388163 |
| NUP210 | red | -0.161114 | 0.431704 | DONSON | red | -0.294487233 | 0.135938613 | PCMTD1 | steelblue | 0.195737 | 0.337914 |
| KIAA1191 | red | 0.160278 | 0.434131 | MAPKAPK2 | red | -0.292343908 | 0.138944919 | U2AF2 | steelblue | 0.216335 | 0.288472 |
| MPZL1 | red | -0.160122 | 0.434585 | HERPUD1 | red | -0.287420846 | 0.146031526 | NUDCD3 | steelblue | 0.276313 | 0.17181 |
| C10orf128 | red | 0.160084 | 0.434695 | TPGS2 | red | -0.284577322 | 0.150241038 | MAPK8IP3 | steelblue | 0.284942 | 0.158269 |
| ZNF570 | red | -0.160076 | 0.434717 | RAPGEF1 | red | -0.283135329 | 0.152408607 | CD180 | tan | -0.367422 | 0.064814 |
| MBD2 | red | -0.15877 | 0.438523 | IL6R | red | -0.282574666 | 0.153257383 | MT1E | tan | -0.352584 | 0.077287 |
| AP2B1 | red | -0.157219 | 0.443068 | SNRK | red | -0.280811639 | 0.15594836 | MT2A | tan | -0.331261 | 0.098313 |
| AP1G2 | red | -0.157199 | 0.443126 | TMEM38B | red | -0.279827484 | 0.157465061 | MT1X | tan | -0.330605 | 0.099022 |
| UVSSA | red | 0.156738 | 0.444481 | SGPL1 | red | -0.279780319 | 0.157538012 | MT1M | tan | -0.329341 | 0.100398 |
| NUP37 | red | -0.156493 | 0.445201 | TRAK2 | red | -0.279374296 | 0.158166995 | RSPO3 | tan | -0.311899 | 0.120866 |

**Table 6. Gene module membership and correlation with growth rate (48/72h), with adjusted p values for each module**

| | | | | | | | | | | | |
|---|---|---|---|---|---|---|---|---|---|---|---|
| KCTD13 | red | 0.15489 | 0.449935 | CAPRIN1 | red | -0.27896164 | 0.158808079 | MT1F | tan | -0.305761 | 0.12875 |
| AHSA1 | red | -0.154538 | 0.450978 | MRAS | red | -0.277477567 | 0.161128922 | PEA15 | tan | -0.286432 | 0.15601 |
| INTS7 | red | -0.154026 | 0.452497 | NLRX1 | red | -0.276768897 | 0.162245603 | ARHGDIB | tan | -0.282657 | 0.161779 |
| TXK | red | -0.153423 | 0.454292 | KIAA0430 | red | -0.276359523 | 0.162893161 | SLC2A4RG | tan | 0.276913 | 0.170844 |
| PELP1 | red | 0.153353 | 0.4545 | VEZF1 | red | -0.273358207 | 0.167696646 | FXYD2 | tan | -0.273927 | 0.175694 |
| GLCE | red | -0.152825 | 0.456072 | FAM160B2 | red | -0.270459348 | 0.1724301 | RELT | tan | -0.272107 | 0.178696 |
| RPS6KA4 | red | 0.152485 | 0.457088 | ARPC1B | red | -0.269845571 | 0.173444224 | NFKBIA | tan | 0.271902 | 0.179036 |
| MTPN | red | -0.15201 | 0.458507 | GRB10 | red | -0.268814702 | 0.175156892 | MT1G | tan | -0.270303 | 0.181708 |
| RPL18 | red | 0.151537 | 0.459924 | PRUNE | red | -0.26184631 | 0.187045132 | MCOLN2 | tan | -0.264626 | 0.191414 |
| BTBD1 | red | -0.151369 | 0.460426 | PEF1 | red | -0.259909128 | 0.190446988 | CAMKK1 | tan | 0.256741 | 0.205479 |
| HSD17B12 | red | -0.151365 | 0.460439 | ZNF384 | red | -0.255358034 | 0.198606644 | ENTPD1 | tan | -0.255878 | 0.20706 |
| CENPP | red | -0.151332 | 0.460537 | CCNI | red | -0.254419962 | 0.200317867 | DCUN1D3 | tan | -0.252537 | 0.213258 |
| COG6 | red | -0.150709 | 0.462409 | TBL2 | red | -0.249516633 | 0.209426794 | SAMD4A | tan | 0.244039 | 0.229584 |
| CCDC18 | red | -0.150353 | 0.463478 | CDCP1 | red | -0.248646127 | 0.211072876 | NFE2L3 | tan | -0.233998 | 0.249921 |
| ASB1 | red | 0.149387 | 0.466389 | AKR1B1 | red | -0.248178588 | 0.211960583 | CA12 | tan | -0.230691 | 0.25687 |
| C17orf103 | red | 0.149224 | 0.46688 | DHRS7 | red | -0.247794091 | 0.212692514 | CCL8 | tan | -0.228343 | 0.261878 |
| ELF1 | red | -0.148783 | 0.468214 | RHCG | red | -0.241151603 | 0.225608094 | NINJ1 | tan | -0.228099 | 0.2624 |
| PRDX6 | red | 0.148224 | 0.469907 | ZDHHC9 | red | -0.240247472 | 0.227405817 | CENPV | tan | 0.222831 | 0.27388 |
| PQBP1 | red | -0.148118 | 0.470228 | SLC43A3 | red | -0.2398629 | 0.228173375 | MT1H | tan | -0.22064 | 0.278749 |
| MIPEP | red | -0.147657 | 0.471626 | MZT1 | red | -0.238818979 | 0.230265635 | PROCR | tan | -0.219157 | 0.282075 |
| G2E3 | red | -0.147327 | 0.472627 | PPP1R13B | red | -0.237737439 | 0.232446754 | DHRS4 | tan | -0.218361 | 0.28387 |
| RHOU | red | -0.145656 | 0.477719 | HENMT1 | red | -0.237107541 | 0.233723372 | ACP5 | tan | 0.212204 | 0.298002 |
| EDEM2 | red | 0.14524 | 0.47899 | DRAM1 | red | -0.234341795 | 0.239383865 | FAM124A | tan | -0.19846 | 0.331103 |
| RBM41 | red | -0.144939 | 0.479913 | GSPT1 | red | -0.233832561 | 0.24043589 | C1S | tan | -0.192902 | 0.345095 |
| UBE2Q2P1 | red | -0.143878 | 0.483168 | KLC1 | red | -0.231082749 | 0.246169485 | TEX2 | tan | -0.190929 | 0.350148 |
| PIK3IP1 | red | 0.1432 | 0.485252 | PLP2 | red | -0.227000977 | 0.254844877 | ESYT1 | tan | 0.183695 | 0.369039 |
| TFB2M | red | -0.142787 | 0.486526 | YWHAQ | red | -0.226945824 | 0.254963447 | CIZ1 | tan | 0.177968 | 0.384407 |
| MAN1A1 | red | -0.142667 | 0.486897 | SRC | red | -0.220225663 | 0.269680505 | LCP2 | tan | -0.171751 | 0.401498 |
| RRBP1 | red | 0.142085 | 0.488693 | LIMS1 | red | -0.219091334 | 0.272217509 | TNFRSF9 | tan | -0.171225 | 0.402963 |
| RREB1 | red | -0.141736 | 0.489774 | ATP6AP2 | red | -0.21906904 | 0.272267525 | ZNF589 | tan | 0.168532 | 0.41051 |
| HIATL1 | red | -0.141266 | 0.491229 | TKT | red | -0.218659532 | 0.273187279 | IRAK2 | tan | 0.1684 | 0.410882 |
| RGL2 | red | 0.140905 | 0.492349 | TIAL1 | red | -0.210540417 | 0.291834126 | PLA2G16 | tan | 0.165361 | 0.4195 |
| ZDHHC17 | red | -0.140738 | 0.492866 | MPZL1 | red | -0.208989664 | 0.29548477 | IL6 | tan | -0.16508 | 0.420301 |
| RAB11B | red | 0.139575 | 0.496484 | NMT2 | red | -0.207789877 | 0.298328802 | CYB5R3 | tan | -0.163991 | 0.423416 |
| ECM1 | red | -0.139312 | 0.497302 | CEP350 | red | -0.19643081 | 0.326100821 | PDE8A | tan | 0.163703 | 0.424241 |
| TTF2 | red | -0.139016 | 0.498225 | SLC7A8 | red | -0.193982967 | 0.332285481 | IL2RA | tan | -0.163554 | 0.424669 |
| CSF2RA | red | -0.138612 | 0.499488 | IKZF5 | red | -0.191588587 | 0.338403354 | KYNU | tan | -0.159835 | 0.435419 |
| THAP7 | red | -0.137062 | 0.504343 | TMEM222 | red | -0.190247875 | 0.341858437 | NNMT | tan | -0.159474 | 0.436469 |
| PI4KAP1 | red | 0.136574 | 0.505877 | MICALL1 | red | -0.188994014 | 0.345108792 | SLC9A9 | tan | -0.156597 | 0.444896 |
| PFKL | red | 0.135928 | 0.507911 | SHISA4 | red | -0.182717351 | 0.361656014 | PDCD1LG2 | tan | -0.154016 | 0.452529 |
| RPL35 | red | -0.133018 | 0.517118 | PACS1 | red | -0.182404991 | 0.362491484 | MYCL1 | tan | -0.152926 | 0.455771 |
| SAMD8 | red | -0.132625 | 0.518369 | DESI1 | red | -0.179604424 | 0.37003273 | TSFM | tan | -0.147096 | 0.473329 |
| SRD5A1 | red | -0.131666 | 0.521424 | DUSP5 | red | -0.175356829 | 0.381643416 | NEDD9 | tan | -0.145296 | 0.47882 |
| RPUSD2 | red | 0.131591 | 0.521663 | ZNHIT1 | red | -0.173681712 | 0.386279322 | CISH | tan | -0.145259 | 0.478934 |
| LIN52 | red | -0.129439 | 0.528557 | MAP4K4 | red | -0.165404143 | 0.409656579 | CABLES1 | tan | 0.144044 | 0.482658 |
| LTBR | red | -0.126701 | 0.537387 | PSMA2 | red | -0.152581086 | 0.447383673 | JAG1 | tan | 0.143638 | 0.483904 |
| SNRNP70 | red | 0.126305 | 0.53867 | ZNF217 | red | -0.151762528 | 0.449853158 | EPDR1 | tan | -0.143534 | 0.484225 |
| BCRP3 | red | -0.124842 | 0.543419 | C1orf52 | red | -0.150016867 | 0.455143725 | GPR84 | tan | -0.142333 | 0.487926 |
| CTR9 | red | -0.1232 | 0.548778 | ADAR | red | -0.140231547 | 0.485399662 | KIAA0930 | tan | 0.140037 | 0.495045 |
| SF3B14 | red | -0.122678 | 0.550485 | LRP8 | red | -0.136368896 | 0.497617578 | SEMA4A | tan | 0.136445 | 0.506283 |
| EFNA4 | red | 0.122 | 0.552706 | TNFSF15 | red | -0.130442112 | 0.516658699 | NFAM1 | tan | -0.133772 | 0.514724 |
| CES2 | red | -0.12193 | 0.552937 | FUT10 | red | -0.12589327 | 0.531509147 | CFB | tan | 0.132726 | 0.518046 |
| COMMD1 | red | -0.121415 | 0.554625 | NPRL3 | red | -0.122231295 | 0.543609866 | FABP4 | tan | -0.131773 | 0.521083 |
| CTNNBIP1 | red | -0.121375 | 0.554757 | ASF1A | red | -0.095595585 | 0.635276402 | MCTP1 | tan | -0.130464 | 0.525268 |
| MRPL4 | red | 0.120406 | 0.557945 | TBCCD1 | red | -0.05504348 | 0.785093755 | EIF5B | tan | 0.129122 | 0.529574 |
| CAMK1 | red | -0.120353 | 0.558122 | GABPB2 | red | -0.052038668 | 0.796576462 | GTPBP4 | tan | -0.128297 | 0.532229 |

**Table 6. Gene module membership and correlation with growth rate (48/72h), with adjusted p values for each module**

| | | | | | | | | | | | |
|---|---|---|---|---|---|---|---|---|---|---|---|
| SLC47A1 | red | -0.119361 | 0.561391 | ENDOD1 | red | -0.043697027 | 0.828663915 | IL3RA | tan | 0.126002 | 0.539651 |
| ERC1 | red | -0.118204 | 0.565222 | PCMTD2 | red | -0.038079121 | 0.850429646 | NCF1C | tan | -0.125617 | 0.540902 |
| LOC344887 | red | 0.116565 | 0.570663 | RAD52 | red | -0.029150776 | 0.88523613 | G0S2 | tan | 0.122486 | 0.551112 |
| NSUN6 | red | -0.116427 | 0.571124 | EPB41L1 | red | -0.024516447 | 0.903389146 | FFAR2 | tan | -0.120186 | 0.55867 |
| FASN | red | 0.112187 | 0.585316 | GOPC | red | -0.017610288 | 0.930526587 | TNFSF15 | tan | -0.117482 | 0.567616 |
| ERMP1 | red | -0.111968 | 0.586053 | CLEC2B | red | -0.011459558 | 0.954760526 | IDO1 | tan | -0.116915 | 0.569499 |
| CLCC1 | red | -0.110788 | 0.590031 | ZNF420 | red | -0.011416433 | 0.9549306 | CDKN2B | tan | -0.116768 | 0.569987 |
| CYB5D2 | red | -0.11019 | 0.592051 | POMT2 | red | -0.006958069 | 0.972522552 | C17orf49 | tan | 0.113089 | 0.582283 |
| LY96 | red | -0.109277 | 0.595145 | GRHL1 | red | -0.002621749 | 0.989645052 | CNBP | tan | -0.112895 | 0.582934 |
| COQ7 | red | 0.107941 | 0.59968 | SIRT6 | red | 0.003398667 | 0.986576751 | CAT | tan | -0.112885 | 0.582969 |
| NAA60 | red | -0.105909 | 0.606605 | HLA-DPB1 | red | 0.00581267 | 0.977044449 | GRK5 | tan | -0.112592 | 0.583954 |
| ITPA | red | -0.105131 | 0.609267 | LOC1001314[illegible] | red | 0.020221291 | 0.920256231 | PTGS2 | tan | 0.112505 | 0.584244 |
| KLHDC3 | red | -0.105118 | 0.609311 | PGAP3 | red | 0.023104102 | 0.908931269 | RCSD1 | tan | -0.109553 | 0.594209 |
| DOCK4 | red | 0.105005 | 0.609699 | GSKIP | red | 0.0241481 | 0.904834147 | CARD11 | tan | -0.108823 | 0.596685 |
| TMCO1 | red | 0.104312 | 0.612073 | TREM1 | red | 0.02457714 | 0.903151082 | IL8 | tan | -0.107425 | 0.601435 |
| PHPT1 | red | -0.104109 | 0.612771 | METTL13 | red | 0.026153702 | 0.896970009 | CD300E | tan | -0.106916 | 0.603167 |
| EEF1G | red | -0.103832 | 0.61372 | PRPSAP2 | red | 0.03445164 | 0.864541864 | MAP7D3 | tan | -0.105524 | 0.60792 |
| INCENP | red | -0.103541 | 0.61472 | ATP5O | red | 0.034821289 | 0.863101862 | IL1B | tan | -0.105113 | 0.609327 |
| PHACTR4 | red | 0.102784 | 0.617322 | SSR4 | red | 0.043909288 | 0.827843851 | EEPD1 | tan | -0.102441 | 0.618504 |
| CLDN12 | red | 0.101758 | 0.620858 | HMGCS1 | red | 0.050124103 | 0.803914635 | VEGFA | tan | -0.101527 | 0.621654 |
| TGFBR1 | red | -0.100885 | 0.623873 | RNASET2 | red | 0.052879313 | 0.79335971 | LGALS3 | tan | -0.101153 | 0.622948 |
| VPS8 | red | -0.099238 | 0.629575 | FAM40A | red | 0.057189387 | 0.776919935 | LAMB3 | tan | 0.10083 | 0.624064 |
| STT3B | red | -0.099038 | 0.630271 | GAS2L1 | red | 0.062354525 | 0.757342104 | PDK4 | tan | -0.100085 | 0.626639 |
| EIF4EBP2 | red | 0.098368 | 0.632596 | ZNF542 | red | 0.064806832 | 0.748097038 | RCN1 | tan | -0.099555 | 0.628477 |
| PIGQ | red | -0.096783 | 0.638116 | SERPINF2 | red | 0.076445727 | 0.704700545 | MGC12916 | tan | -0.097969 | 0.633986 |
| SSRP1 | red | 0.096367 | 0.639569 | N4BP2L1 | red | 0.079816939 | 0.692290637 | AKR1C2 | tan | -0.096971 | 0.63746 |
| GTPBP6 | red | -0.09635 | 0.639629 | LEPROTL1 | red | 0.080463935 | 0.689917682 | ADAMTS15 | tan | -0.095319 | 0.643231 |
| HEATR7A | red | 0.095973 | 0.640945 | ADH5 | red | 0.083172146 | 0.68001615 | LMTK2 | tan | -0.092977 | 0.651444 |
| LASP1 | red | 0.094724 | 0.645312 | TMEM229[illegible] | red | 0.087643945 | 0.663780181 | ANKMY2 | tan | -0.092823 | 0.651986 |
| POMGNT1 | red | 0.093555 | 0.649413 | PMM1 | red | 0.092277586 | 0.647111566 | MAP1LC3A | tan | 0.092086 | 0.654582 |
| ARHGEF2 | red | 0.091354 | 0.657161 | ZNF75A | red | 0.093881492 | 0.641379816 | CD274 | tan | 0.08998 | 0.662015 |
| SPG7 | red | 0.090697 | 0.659481 | RPS6KB1 | red | 0.094082833 | 0.640661706 | HMOX1 | tan | -0.087923 | 0.669305 |
| CECR5 | red | 0.090582 | 0.659885 | FAM203A | red | 0.094687207 | 0.638508022 | EDN1 | tan | -0.087214 | 0.671825 |
| TAOK3 | red | -0.09054 | 0.660033 | RNF135 | red | 0.094993955 | 0.637416018 | ADORA2B | tan | -0.086399 | 0.674728 |
| SF3B5 | red | -0.089447 | 0.6639 | NICN1 | red | 0.100561522 | 0.617726059 | IL18 | tan | -0.083305 | 0.685782 |
| SNX33 | red | 0.088739 | 0.666412 | KARS | red | 0.104907556 | 0.602532399 | GAS7 | tan | -0.081899 | 0.690824 |
| DNAJC7 | red | 0.086675 | 0.673745 | TRAPPC5 | red | 0.106077142 | 0.5984707 | SLC39A8 | tan | -0.081856 | 0.690978 |
| SNRNP40 | red | -0.084234 | 0.682453 | THAP10 | red | 0.106557941 | 0.596804388 | ABRACL | tan | -0.079542 | 0.699308 |
| RAP2B | red | 0.083784 | 0.684065 | DAPK1 | red | 0.108236521 | 0.591002509 | PXMP2 | tan | 0.079208 | 0.700516 |
| SFXN5 | red | 0.08339 | 0.685478 | PRPF38B | red | 0.111024904 | 0.581418932 | ZFYVE26 | tan | 0.078207 | 0.704133 |
| SCAMP4 | red | 0.081649 | 0.691724 | C1QA | red | 0.114919986 | 0.568147195 | DOK2 | tan | -0.078187 | 0.704206 |
| SLC25A17 | red | -0.079258 | 0.700335 | RAB11FIP3 | red | 0.115341861 | 0.566717945 | NFKB1 | tan | 0.077935 | 0.705114 |
| ATP5C1 | red | -0.077244 | 0.707615 | MXD4 | red | 0.116479692 | 0.562871233 | LOC1002895[illegible] | tan | -0.075594 | 0.713601 |
| MTFP1 | red | -0.074736 | 0.716721 | C1QB | red | 0.119515199 | 0.55266726 | SOBP | tan | 0.068032 | 0.741236 |
| NAT9 | red | 0.073777 | 0.720209 | PTPLAD1 | red | 0.119724468 | 0.551966947 | CCL20 | tan | -0.067964 | 0.741485 |
| PMS2L2 | red | -0.07116 | 0.729765 | ZNF839 | red | 0.121087794 | 0.547414648 | TREM2 | tan | 0.067345 | 0.743762 |
| LMBR1L | red | 0.070309 | 0.732881 | S100P | red | 0.121550711 | 0.545872894 | ABHD6 | tan | 0.066679 | 0.746212 |
| ARHGEF28 | red | -0.069739 | 0.734969 | MLKL | red | 0.123342613 | 0.539924046 | ST3GAL4 | tan | -0.062301 | 0.762389 |
| WAC | red | -0.068152 | 0.740794 | C14orf159 | red | 0.123794517 | 0.538428616 | ZNF117 | tan | 0.06196 | 0.763652 |
| RFNG | red | -0.067776 | 0.742175 | PRADC1 | red | 0.126021787 | 0.53108681 | CD14 | tan | 0.061672 | 0.764722 |
| TAPT1 | red | -0.066909 | 0.745367 | TRMT1 | red | 0.130163167 | 0.517563512 | GSR | tan | 0.061329 | 0.765994 |
| KIF3A | red | -0.065693 | 0.749848 | RPP21 | red | 0.131184256 | 0.514255153 | FABP5 | tan | 0.060625 | 0.768609 |
| YTHDF1 | red | 0.065438 | 0.750789 | THTPA | red | 0.134518446 | 0.503524647 | MEMO1 | tan | 0.059063 | 0.774416 |
| AP5S1 | red | -0.063273 | 0.758788 | RPAIN | red | 0.135823834 | 0.499353939 | ISG20 | tan | 0.058334 | 0.777131 |
| TMUB2 | red | 0.063125 | 0.759336 | CCDC102B | red | 0.136764168 | 0.496360283 | SEMA4D | tan | -0.058222 | 0.777548 |
| C7orf55 | red | 0.060792 | 0.767987 | ZWINT | red | 0.137204212 | 0.49496245 | NR4A2 | tan | -0.057515 | 0.780186 |

**Table 6. Gene module membership and correlation with growth rate (48/72h), with adjusted p values for each module**

| | | | | | | | | | | | |
|---|---|---|---|---|---|---|---|---|---|---|---|
| EIF2B5 | red | -0.059627 | 0.772319 | CDR2 | red | 0.13754304 | 0.493887482 | DDX10 | tan | 0.057183 | 0.781424 |
| EEA1 | red | -0.0589 | 0.775024 | CSNK2A1 | red | 0.137714671 | 0.493343412 | CDA | tan | -0.05655 | 0.783787 |
| RIN2 | red | 0.052165 | 0.800205 | SSH2 | red | 0.142920338 | 0.4769857 | RAB42 | tan | 0.056535 | 0.783843 |
| NFKBIL1 | red | -0.051621 | 0.802249 | C1orf54 | red | 0.148593861 | 0.459480629 | SLC22A5 | tan | -0.056276 | 0.78481 |
| IER3IP1 | red | -0.050481 | 0.806534 | PPM1N | red | 0.150043955 | 0.455061381 | CEP135 | tan | 0.054528 | 0.791348 |
| PPP1R16A | red | -0.04921 | 0.811317 | TIMM17B | red | 0.152502721 | 0.447619776 | IL1A | tan | -0.053079 | 0.796777 |
| CUEDC2 | red | 0.048848 | 0.812683 | ZNF90 | red | 0.15271445 | 0.44698202 | PRPF8 | tan | 0.052161 | 0.800221 |
| ANGEL1 | red | 0.048725 | 0.813146 | LMAN2L | red | 0.154431421 | 0.441828206 | C15orf48 | tan | -0.050981 | 0.804654 |
| SCAP | red | 0.04851 | 0.813954 | SPPL2B | red | 0.155755294 | 0.43787626 | PIK3AP1 | tan | -0.050438 | 0.806697 |
| TNIP2 | red | -0.047518 | 0.817697 | EPHB6 | red | 0.158727919 | 0.429072492 | SLAMF7 | tan | -0.049781 | 0.809168 |
| NUP35 | red | -0.047506 | 0.817741 | APOE | red | 0.160609204 | 0.42355116 | CLCN4 | tan | -0.049317 | 0.810913 |
| TREM1 | red | -0.045252 | 0.826258 | KIAA1147 | red | 0.162395333 | 0.418345415 | MARCKS | tan | -0.049212 | 0.811309 |
| PPP1R18 | red | 0.044388 | 0.829525 | ZBTB5 | red | 0.167351842 | 0.404086168 | NEURL3 | tan | -0.048503 | 0.81398 |
| WDR81 | red | 0.043706 | 0.83211 | HINT2 | red | 0.167575953 | 0.403447953 | COQ10A | tan | -0.047696 | 0.817024 |
| DDX56 | red | 0.042383 | 0.837124 | LOC728431 | red | 0.169376471 | 0.398341112 | CD22 | tan | 0.04736 | 0.818292 |
| ZNF736 | red | -0.038014 | 0.853724 | MANBA | red | 0.171335636 | 0.392826043 | PELI3 | tan | 0.045317 | 0.826011 |
| FBXO21 | red | 0.034975 | 0.865311 | THOC1 | red | 0.178009083 | 0.374369172 | KIAA0040 | tan | -0.045086 | 0.826885 |
| NMRAL1 | red | -0.032735 | 0.873868 | SLC25A19 | red | 0.17865326 | 0.372614638 | BRI3BP | tan | -0.044601 | 0.828721 |
| UBTF | red | 0.031562 | 0.878352 | KCTD18 | red | 0.178967809 | 0.371759646 | SLC7A2 | tan | -0.043744 | 0.831963 |
| CAMKK2 | red | 0.031122 | 0.880034 | TOR3A | red | 0.179404208 | 0.370575344 | TAP2 | tan | 0.043743 | 0.831968 |
| HMOX2 | red | 0.030866 | 0.881018 | EXOSC2 | red | 0.181155708 | 0.365844268 | IL15RA | tan | -0.043476 | 0.832979 |
| CIR1 | red | -0.030556 | 0.882203 | ERH | red | 0.181258959 | 0.365566479 | SOD2 | tan | -0.043189 | 0.834067 |
| PPP1R9B | red | 0.02973 | 0.885367 | ACAD11 | red | 0.186033394 | 0.3528566 | CCL15 | tan | -0.04301 | 0.834744 |
| ZDHHC14 | red | 0.029306 | 0.886991 | L3HYPDH | red | 0.186803844 | 0.350830501 | FAM212B | tan | -0.039763 | 0.847071 |
| ZBTB8A | red | 0.028346 | 0.89067 | PDE1B | red | 0.187893858 | 0.347975877 | MGST3 | tan | 0.039571 | 0.847802 |
| KLHL22 | red | 0.02746 | 0.894071 | FAM129A | red | 0.191381471 | 0.338935724 | ABCG2 | tan | 0.039314 | 0.848779 |
| OGG1 | red | 0.027109 | 0.895416 | SLC7A6OS | red | 0.193828582 | 0.332677914 | BLVRB | tan | -0.038761 | 0.850882 |
| KIAA0195 | red | 0.027084 | 0.89551 | OPHN1 | red | 0.194024695 | 0.332179461 | GAS2L3 | tan | -0.0381 | 0.853398 |
| RPL26L1 | red | 0.025995 | 0.899693 | ZNF174 | red | 0.194411288 | 0.331198196 | PTGES | tan | 0.037387 | 0.856114 |
| ZNF641 | red | 0.025307 | 0.902332 | XPC | red | 0.194961244 | 0.329805314 | GCH1 | tan | 0.035793 | 0.86219 |
| ARHGAP33 | red | -0.024922 | 0.903811 | LUC7L3 | red | 0.197942371 | 0.322317066 | NCF1B | tan | -0.033798 | 0.869804 |
| C16orf58 | red | -0.022946 | 0.91141 | GPX1 | red | 0.198047223 | 0.322055601 | JAK3 | tan | 0.031371 | 0.879085 |
| GLT1D1 | red | -0.022403 | 0.913499 | CEBPA | red | 0.198218943 | 0.321627668 | B4GALT1 | tan | 0.030247 | 0.883387 |
| ABHD16A | red | -0.021139 | 0.91836 | RAB35 | red | 0.199384753 | 0.318731657 | SNTA1 | tan | 0.029431 | 0.886512 |
| RASAL3 | red | 0.020743 | 0.919887 | PHF20L1 | red | 0.199739378 | 0.317853914 | TNFAIP6 | tan | 0.029209 | 0.887363 |
| PDXP | red | -0.020691 | 0.920085 | TAX1BP3 | red | 0.201018435 | 0.31470045 | SLC2A3 | tan | -0.024245 | 0.906413 |
| HSD17B7P2 | red | 0.019341 | 0.925286 | ZNF230 | red | 0.204302426 | 0.30669266 | PLSCR1 | tan | -0.02343 | 0.909547 |
| TRIM62 | red | 0.018602 | 0.928132 | PTGFRN | red | 0.207513427 | 0.298986536 | RCBTB2 | tan | 0.021359 | 0.917513 |
| NSUN4 | red | 0.017573 | 0.932097 | LAGE3 | red | 0.207961391 | 0.297921191 | SOCS3 | tan | -0.019809 | 0.923483 |
| ADCK3 | red | 0.015531 | 0.939973 | BECN1 | red | 0.209321154 | 0.294702005 | SERINC5 | tan | -0.019363 | 0.925199 |
| CPNE2 | red | -0.01496 | 0.942177 | TXN2 | red | 0.210428879 | 0.292095747 | CEP78 | tan | 0.018663 | 0.927897 |
| FERMT2 | red | -0.012815 | 0.950457 | PCYOX1L | red | 0.214489748 | 0.282666017 | COTL1 | tan | -0.018042 | 0.930292 |
| PEX13 | red | -0.012305 | 0.952425 | TELO2 | red | 0.215929074 | 0.2793708 | NBN | tan | -0.017514 | 0.932328 |
| C6orf89 | red | -0.011829 | 0.954264 | NAA38 | red | 0.218162911 | 0.274305361 | NCF1 | tan | 0.017483 | 0.932444 |
| GOLM1 | red | -0.011547 | 0.955354 | CBR4 | red | 0.218875247 | 0.272702535 | APOBEC3A | tan | 0.016519 | 0.936163 |
| XPO4 | red | -0.010471 | 0.95951 | DPEP2 | red | 0.219449983 | 0.271413715 | TSC22D1 | tan | -0.015331 | 0.940747 |
| CCDC93 | red | 0.009 | 0.965195 | SERPINA1 | red | 0.219765137 | 0.27070866 | PLA2G15 | tan | -0.014946 | 0.942233 |
| RUNX3 | red | 0.008924 | 0.96549 | VAPA | red | 0.219935183 | 0.270328728 | FBN1 | tan | 0.014618 | 0.943497 |
| SUMF2 | red | -0.007245 | 0.971978 | DNTTIP1 | red | 0.223209677 | 0.263079474 | PDE4B | tan | 0.013004 | 0.949728 |
| PMS2P3 | red | 0.006375 | 0.975341 | HMG20A | red | 0.223863766 | 0.261646654 | PLTP | tan | -0.012533 | 0.951545 |
| AGTRAP | red | 0.005958 | 0.976955 | RAB13 | red | 0.225341515 | 0.25842823 | AK4 | tan | -0.011095 | 0.957099 |
| CARHSP1 | red | 0.005322 | 0.979416 | BACE1 | red | 0.225473335 | 0.258142393 | EREG | tan | -0.010411 | 0.959744 |
| ATG2A | red | 0.004808 | 0.981401 | LOC253039 | red | 0.225829909 | 0.257370231 | CP | tan | -0.010131 | 0.960823 |
| CHML | red | -0.00456 | 0.982362 | CAPG | red | 0.233948978 | 0.240195115 | ECE1 | tan | 0.010093 | 0.96097 |
| ANKS3 | red | -0.003506 | 0.986438 | POGZ | red | 0.234413002 | 0.239237004 | SNHG12 | tan | 0.009028 | 0.965087 |
| SHROOM3 | royalblue | -0.371484 | 0.061686 | ZNF593 | red | 0.239092091 | 0.229717023 | CD9 | tan | 0.006771 | 0.973811 |

**Table 6. Gene module membership and correlation with growth rate (48/72h), with adjusted p values for each module**

| | | | | | | | | | | | |
|---|---|---|---|---|---|---|---|---|---|---|---|
| LYPD3 | royalblue | -0.364492 | 0.067144 | LPXN | red | 0.240868377 | 0.226170216 | HAMP | tan | 0.006451 | 0.975051 |
| CCL20 | royalblue | -0.364393 | 0.067224 | TRIP6 | red | 0.24211061 | 0.22371169 | COQ2 | tan | -0.006199 | 0.976025 |
| C4orf27 | royalblue | -0.358092 | 0.07246 | HNMT | red | 0.243810187 | 0.220377177 | INHBA | tan | -0.006058 | 0.976568 |
| SERPINB2 | royalblue | -0.301196 | 0.13485 | PQLC3 | red | 0.24432867 | 0.219366628 | RAB21 | tan | -0.005798 | 0.977575 |
| NFE2 | royalblue | -0.288215 | 0.153337 | STK16 | red | 0.24679274 | 0.214606726 | STAP2 | tan | 0.005066 | 0.980406 |
| IRF1 | royalblue | -0.281437 | 0.163675 | UBE2V1 | red | 0.249045963 | 0.210315721 | OSM | tan | -0.003739 | 0.985537 |
| ZNF267 | royalblue | 0.27081 | 0.180859 | EPS8L1 | red | 0.249690106 | 0.20909981 | OSBPL11 | tan | 0.0036 | 0.986074 |
| KCNN4 | royalblue | -0.262527 | 0.195092 | KHDRBS1 | red | 0.250956273 | 0.206723695 | MOSPD2 | tan | 0.002843 | 0.989003 |
| PTX3 | royalblue | -0.241581 | 0.234457 | MYNN | red | 0.251540433 | 0.205633673 | TLR2 | tan | -0.002751 | 0.98936 |
| KIAA0922 | royalblue | -0.239882 | 0.237866 | HCLS1 | red | 0.252202526 | 0.204402976 | SERPINF1 | tan | 0.002207 | 0.991463 |
| TRAF3 | royalblue | 0.231652 | 0.254837 | COQ4 | red | 0.252547133 | 0.203764416 | CSF2RB | tan | -0.001102 | 0.995737 |
| CCDC134 | royalblue | -0.213476 | 0.295047 | ELMOD3 | red | 0.252662245 | 0.203551416 | RNF8 | turquoise | -0.580801 | 0.001864 |
| TNFAIP8 | royalblue | -0.211664 | 0.29926 | CNNM3 | red | 0.253510671 | 0.201986203 | NCBP1 | turquoise | -0.49953 | 0.009371 |
| KYNU | royalblue | 0.203726 | 0.318167 | BBS2 | red | 0.255097665 | 0.199080597 | ACO2 | turquoise | -0.472544 | 0.014781 |
| IL6 | royalblue | -0.19005 | 0.352412 | RBCK1 | red | 0.255505002 | 0.198339457 | AGAP2 | turquoise | -0.47231 | 0.014837 |
| CCL4 | royalblue | -0.188085 | 0.357504 | SUPT5H | red | 0.256198852 | 0.197081375 | PRKAB2 | turquoise | -0.452425 | 0.020306 |
| BHLHE41 | royalblue | -0.185914 | 0.363182 | MBNL3 | red | 0.256458134 | 0.196612659 | FAM89A | turquoise | -0.442796 | 0.023491 |
| SLC11A2 | royalblue | -0.185519 | 0.36422 | ALKBH8 | red | 0.257326025 | 0.195049304 | MMP12 | turquoise | -0.407205 | 0.038953 |
| PDE4B | royalblue | -0.182204 | 0.373004 | CUL2 | red | 0.260722299 | 0.189013824 | RANBP1 | turquoise | -0.383377 | 0.053198 |
| ABCA1 | royalblue | 0.179357 | 0.380647 | NDUFS3 | red | 0.261012108 | 0.188504861 | FLVCR1 | turquoise | -0.373352 | 0.060288 |
| CA2 | royalblue | -0.168276 | 0.411232 | ING4 | red | 0.262093713 | 0.186613723 | LPP | turquoise | -0.37009 | 0.062745 |
| CCL3L3 | royalblue | 0.164993 | 0.420548 | FTO | red | 0.262746445 | 0.185478837 | TMEM14B | turquoise | -0.36892 | 0.063646 |
| AMPD3 | royalblue | -0.150866 | 0.461938 | SYT11 | red | 0.264146323 | 0.18306108 | NUP155 | turquoise | -0.367071 | 0.065089 |
| LAMC1 | royalblue | -0.149961 | 0.464659 | BATF3 | red | 0.26417488 | 0.183011989 | FAM89B | turquoise | -0.363526 | 0.067927 |
| TMEM53 | royalblue | -0.149487 | 0.466087 | CASC4 | red | 0.264677551 | 0.18214935 | FUOM | turquoise | -0.361923 | 0.069241 |
| FBXL2 | royalblue | 0.144441 | 0.481438 | OGFOD1 | red | 0.265660824 | 0.180470134 | C11orf31 | turquoise | -0.360293 | 0.070596 |
| C5orf45 | royalblue | -0.141982 | 0.489012 | RNF7 | red | 0.266410585 | 0.179196979 | PPCS | turquoise | -0.356069 | 0.074205 |
| IL18BP | royalblue | -0.1386 | 0.499525 | HPS6 | red | 0.269066742 | 0.174737069 | C7orf41 | turquoise | -0.355076 | 0.075074 |
| KIAA1009 | royalblue | -0.138593 | 0.499546 | ASPSCR1 | red | 0.269632541 | 0.173797182 | C12orf44 | turquoise | -0.353994 | 0.076028 |
| GPR132 | royalblue | -0.136041 | 0.507553 | C22orf28 | red | 0.27180537 | 0.170220698 | GOSR2 | turquoise | -0.351654 | 0.078126 |
| CCRL2 | royalblue | 0.135592 | 0.508968 | KCTD20 | red | 0.271952054 | 0.169981135 | KLHL18 | turquoise | -0.349209 | 0.080364 |
| JAKMIP2 | royalblue | -0.129473 | 0.528448 | MIS18BP1 | red | 0.27610671 | 0.16329398 | TMEM214 | turquoise | -0.345532 | 0.083821 |
| SNX9 | royalblue | 0.129287 | 0.529044 | EDF1 | red | 0.27781223 | 0.160603478 | HNRNPUL2 | turquoise | -0.343657 | 0.085626 |
| CCR7 | royalblue | -0.121746 | 0.553538 | C9orf78 | red | 0.281500128 | 0.154893519 | ATPBD4 | turquoise | -0.342205 | 0.087046 |
| MFSD2A | royalblue | 0.114441 | 0.577751 | TMEM14A | red | 0.281683621 | 0.154613248 | STX2 | turquoise | -0.341145 | 0.088092 |
| SPDL1 | royalblue | -0.111769 | 0.586724 | ZNF642 | red | 0.281746468 | 0.154517336 | UBL4A | turquoise | -0.339047 | 0.090191 |
| PLEKHM3 | royalblue | 0.111067 | 0.58909 | FAM156B | red | 0.282594442 | 0.153227386 | SUMO1 | turquoise | -0.338274 | 0.090974 |
| CCL3 | royalblue | -0.106834 | 0.603449 | LINC00641 | red | 0.286294541 | 0.147688639 | OSGIN1 | turquoise | -0.328552 | 0.101263 |
| TNFSF15 | royalblue | -0.103949 | 0.613317 | MLF2 | red | 0.286923019 | 0.146762315 | SRSF4 | turquoise | -0.323551 | 0.106882 |
| KLHL5 | royalblue | -0.102894 | 0.616944 | MTMR1 | red | 0.292813404 | 0.138282314 | TAF10 | turquoise | -0.319266 | 0.111878 |
| CHD3 | royalblue | 0.102733 | 0.617499 | RNF20 | red | 0.293674134 | 0.137073496 | CPT1A | turquoise | -0.318979 | 0.112219 |
| MSR1 | royalblue | -0.100097 | 0.6266 | ZNF629 | red | 0.294550952 | 0.135849967 | CCDC23 | turquoise | -0.31765 | 0.113806 |
| KANK1 | royalblue | -0.099992 | 0.626961 | CUL1 | red | 0.296401819 | 0.133293246 | ZNF197 | turquoise | -0.317284 | 0.114246 |
| RPSAP58 | royalblue | 0.092699 | 0.652423 | MLH1 | red | 0.300964914 | 0.12713952 | NECAB3 | turquoise | -0.317064 | 0.114511 |
| C19orf10 | royalblue | 0.086284 | 0.675137 | VPS33B | red | 0.302176165 | 0.125541467 | MTERF | turquoise | -0.316448 | 0.115256 |
| DDX43 | royalblue | -0.083757 | 0.68416 | TYMP | red | 0.302207823 | 0.125499897 | PHC2 | turquoise | -0.316256 | 0.115489 |
| PDE7A | royalblue | 0.083172 | 0.686257 | ATP5I | red | 0.305593824 | 0.121111644 | MYC | turquoise | -0.316086 | 0.115694 |
| MIR155HG | royalblue | -0.07914 | 0.700762 | ACSF3 | red | 0.305600069 | 0.121103656 | FLT1 | turquoise | -0.311796 | 0.120996 |
| RAB20 | royalblue | 0.077471 | 0.706796 | TRIM66 | red | 0.30703945 | 0.119272832 | LFNG | turquoise | -0.310517 | 0.122609 |
| IRAK2 | royalblue | 0.077026 | 0.708405 | NUCKS1 | red | 0.307107173 | 0.119187196 | KATNAL1 | turquoise | -0.309431 | 0.123992 |
| KDM6B | royalblue | 0.074532 | 0.717463 | IRF3 | red | 0.307951114 | 0.118123815 | CPSF2 | turquoise | -0.308904 | 0.124667 |
| CSF2 | royalblue | -0.07431 | 0.718269 | C9orf142 | red | 0.30928242 | 0.116460566 | RAC1 | turquoise | -0.308804 | 0.124796 |
| ZC3H12C | royalblue | -0.072973 | 0.723143 | MORC2 | red | 0.315622082 | 0.10877632 | TPM1 | turquoise | -0.308457 | 0.125242 |
| TNFAIP6 | royalblue | -0.070686 | 0.731498 | VMO1 | red | 0.315990764 | 0.108341322 | EEF1B2 | turquoise | -0.306168 | 0.128215 |
| NFKB2 | royalblue | -0.067233 | 0.744172 | CAMLG | red | 0.317622621 | 0.106431439 | GTPBP8 | turquoise | -0.305603 | 0.128958 |
| RGS1 | royalblue | -0.064969 | 0.752519 | MRPL46 | red | 0.32074734 | 0.102844401 | RACGAP1 | turquoise | -0.302978 | 0.132445 |

**Table 6. Gene module membership and correlation with growth rate (48/72h), with adjusted p values for each module**

| | | | | | | | | | | | |
|---|---|---|---|---|---|---|---|---|---|---|---|
| COL6A2 | royalblue | -0.064786 | 0.753195 | IFT52 | red | 0.321161404 | 0.102375929 | LRFN4 | turquoise | -0.3015 | 0.134438 |
| EBP | royalblue | 0.06047 | 0.769185 | TOMM22 | red | 0.322099094 | 0.101320917 | VPS26A | turquoise | -0.298302 | 0.138825 |
| GEM | royalblue | -0.059241 | 0.773753 | TUBG2 | red | 0.322994854 | 0.100320689 | XPA | turquoise | -0.29667 | 0.141102 |
| GNGT2 | royalblue | 0.058167 | 0.777755 | PARVG | red | 0.323462074 | 0.09980192 | MICU1 | turquoise | -0.29586 | 0.142244 |
| SPIRE2 | royalblue | -0.053064 | 0.796832 | SLC25A11 | red | 0.323893908 | 0.099324228 | HAUS8 | turquoise | -0.295628 | 0.142571 |
| TMEM63B | royalblue | 0.051503 | 0.802691 | HSPD1 | red | 0.325046145 | 0.09805801 | PHLDA3 | turquoise | -0.295563 | 0.142663 |
| VWA7 | royalblue | -0.051175 | 0.803923 | FBXL15 | red | 0.325724814 | 0.097317888 | GCFC2 | turquoise | -0.292193 | 0.147491 |
| GPX3 | royalblue | 0.050409 | 0.806803 | SCAMP4 | red | 0.328099378 | 0.094761267 | SNRPB | turquoise | -0.291525 | 0.148461 |
| RELB | royalblue | -0.046981 | 0.819725 | ZMYM5 | red | 0.329100325 | 0.093698838 | FAM214B | turquoise | -0.291484 | 0.148521 |
| HHEX | royalblue | -0.040485 | 0.844326 | FAM207A | red | 0.329882157 | 0.092875239 | MTHFSD | turquoise | -0.29059 | 0.149827 |
| MMP9 | royalblue | -0.039216 | 0.849151 | HDHD2 | red | 0.329919643 | 0.092835887 | RNASEL | turquoise | -0.289445 | 0.151512 |
| MSMO1 | royalblue | -0.038468 | 0.851998 | TMEM141 | red | 0.331993607 | 0.090678234 | CREB3L2 | turquoise | -0.287625 | 0.154218 |
| ANKMY2 | royalblue | -0.037423 | 0.855976 | CSTF2 | red | 0.338176338 | 0.08447038 | CHRNA3 | turquoise | -0.286389 | 0.156075 |
| MFHAS1 | royalblue | 0.034194 | 0.86829 | VPS4A | red | 0.34020427 | 0.082506161 | TMEM2 | turquoise | -0.282623 | 0.161832 |
| TMEM88 | royalblue | -0.02965 | 0.885674 | APEH | red | 0.343282646 | 0.07959114 | GUSBP3 | turquoise | -0.280724 | 0.16479 |
| PTPRJ | royalblue | -0.028133 | 0.891487 | FAM204A | red | 0.343581856 | 0.079312053 | ERGIC1 | turquoise | -0.27954 | 0.166654 |
| SOD2 | royalblue | 0.025356 | 0.902144 | RHBDD3 | red | 0.350828544 | 0.072778121 | MRPS10 | turquoise | -0.279254 | 0.167108 |
| CYP51A1 | royalblue | -0.022145 | 0.914488 | EXOSC3 | red | 0.351933473 | 0.071819318 | MRPS33 | turquoise | -0.279016 | 0.167485 |
| PDE6G | royalblue | 0.018678 | 0.92784 | MAEA | red | 0.354711605 | 0.069451477 | ZNF576 | turquoise | -0.278348 | 0.168546 |
| LIF | royalblue | 0.015323 | 0.940778 | DCPS | red | 0.354985777 | 0.069221098 | LRRC37BP1 | turquoise | -0.275727 | 0.172758 |
| TNIP1 | royalblue | -0.014545 | 0.943779 | GDI1 | red | 0.355099188 | 0.069125974 | PUS10 | turquoise | -0.274717 | 0.174402 |
| ZSWIM4 | royalblue | -0.012783 | 0.950582 | BRI3BP | red | 0.36193281 | 0.063577829 | TM7SF3 | turquoise | -0.273935 | 0.17568 |
| LINC00152 | royalblue | -0.009984 | 0.961392 | RBMX | red | 0.362099008 | 0.063447332 | ATP9A | turquoise | -0.273876 | 0.175778 |
| RHOH | royalblue | -0.007532 | 0.970868 | PLAUR | red | 0.366087406 | 0.060377498 | LOC10050703 | turquoise | -0.273844 | 0.175829 |
| CD9 | royalblue | -0.005792 | 0.977597 | AMDHD2 | red | 0.366282104 | 0.060230652 | THUMPD2 | turquoise | -0.272806 | 0.177539 |
| ZFX | royalblue | 0.004861 | 0.981197 | SOAT1 | red | 0.366611609 | 0.059982766 | GPS2 | turquoise | -0.272405 | 0.178202 |
| HIST1H2BH | salmon | -0.613952 | 0.00085 | RPL37 | red | 0.372604508 | 0.055611317 | PTPN18 | turquoise | -0.271728 | 0.179327 |
| HIST1H2AG | salmon | -0.482635 | 0.012516 | ZSCAN21 | red | 0.372702837 | 0.055541729 | FSCN1 | turquoise | -0.270882 | 0.180737 |
| PSME1 | salmon | -0.480812 | 0.012903 | UBE2E1 | red | 0.373958883 | 0.05465879 | LIMA1 | turquoise | -0.270621 | 0.181175 |
| HIST3H2A | salmon | -0.454417 | 0.019693 | DCUN1D2 | red | 0.377915837 | 0.051948869 | ZNF720 | turquoise | -0.270116 | 0.182022 |
| HIST1H2AM | salmon | -0.448054 | 0.021705 | SLC20A1 | red | 0.384328972 | 0.047781952 | ZNF623 | turquoise | -0.26943 | 0.183178 |
| FIGNL1 | salmon | -0.443459 | 0.02326 | GPNMB | red | 0.385794795 | 0.046867608 | OXSR1 | turquoise | -0.268093 | 0.185446 |
| PCMTD2 | salmon | -0.426091 | 0.029972 | ZNRD1 | red | 0.386427292 | 0.046477364 | SNRNP25 | turquoise | -0.267681 | 0.186147 |
| SLC7A1 | salmon | 0.411873 | 0.036557 | SCNM1 | red | 0.390544252 | 0.043999544 | MCEE | turquoise | -0.266798 | 0.18766 |
| HIST1H3C | salmon | -0.398288 | 0.043878 | CHMP2B | red | 0.393814359 | 0.042106949 | DECR1 | turquoise | -0.266171 | 0.188739 |
| TXNL4A | salmon | -0.386001 | 0.051455 | RAPGEFL1 | red | 0.395719135 | 0.041034696 | UBXN2B | turquoise | -0.265873 | 0.189253 |
| HIST1H3G | salmon | -0.385298 | 0.051917 | EMG1 | red | 0.397730365 | 0.039926188 | FTSJD1 | turquoise | -0.265108 | 0.190578 |
| ZYG11B | salmon | -0.384557 | 0.052408 | EIF2D | red | 0.398172494 | 0.03968573 | SGOL2 | turquoise | -0.263222 | 0.193869 |
| HIST1H3B | salmon | -0.372946 | 0.060589 | PLA2G16 | red | 0.399474494 | 0.03898431 | AGPAT3 | turquoise | -0.261423 | 0.197046 |
| TMEM223 | salmon | -0.37114 | 0.061946 | LOC10050638 | red | 0.399948304 | 0.038731522 | ANP32E | turquoise | -0.260346 | 0.198965 |
| HIST1H2AJ | salmon | -0.370799 | 0.062205 | CCDC51 | red | 0.404551972 | 0.036342767 | SLC11A1 | turquoise | -0.25861 | 0.202083 |
| SNORA12 | salmon | -0.364722 | 0.066959 | RPL22 | red | 0.40704365 | 0.035099894 | PPT1 | turquoise | -0.258011 | 0.203168 |
| CDT1 | salmon | -0.363412 | 0.068019 | C7orf50 | red | 0.40842401 | 0.03442615 | SRD5A3 | turquoise | -0.257342 | 0.204383 |
| FICD | salmon | 0.357635 | 0.072851 | SETDB1 | red | 0.409244246 | 0.034030735 | NCOR1 | turquoise | -0.256448 | 0.206015 |
| ECHDC2 | salmon | -0.35279 | 0.077102 | PRMT1 | red | 0.414688581 | 0.03149755 | PITPNC1 | turquoise | -0.256239 | 0.206398 |
| COMMD10 | salmon | -0.349182 | 0.080389 | PRRC2B | red | 0.416471383 | 0.030701829 | MSH3 | turquoise | -0.256198 | 0.206474 |
| SLC30A1 | salmon | 0.347479 | 0.081977 | CNPY3 | red | 0.417819267 | 0.03011104 | LYRM5 | turquoise | -0.255266 | 0.208187 |
| PDE4DIP | salmon | 0.342451 | 0.086804 | BIN3 | red | 0.417838604 | 0.030102631 | LILRB5 | turquoise | -0.254788 | 0.209069 |
| CCNB1IP1 | salmon | -0.341584 | 0.087657 | FASTKD2 | red | 0.422508774 | 0.028126606 | PRPF6 | turquoise | -0.25454 | 0.209527 |
| HIST3H2BB | salmon | -0.334462 | 0.09491 | DDX49 | red | 0.424252559 | 0.027416236 | ARPC1B | turquoise | -0.25381 | 0.210883 |
| BLOC1S5 | salmon | -0.33038 | 0.099266 | SNRNP48 | red | 0.426659489 | 0.026459639 | GNPAT | turquoise | -0.253734 | 0.211024 |
| NOB1 | salmon | -0.327781 | 0.102115 | FAM162A | red | 0.430172378 | 0.025112208 | RARS | turquoise | -0.252697 | 0.212959 |
| C2orf71 | salmon | -0.323859 | 0.106529 | HNRNPAB | red | 0.430430122 | 0.025015582 | VAMP7 | turquoise | -0.251972 | 0.214319 |
| CASP6 | salmon | -0.316207 | 0.115549 | ATOX1 | red | 0.43816749 | 0.022252536 | PAGR1 | turquoise | -0.251367 | 0.215458 |
| HIST2H2AB | salmon | -0.293173 | 0.146075 | VCP | red | 0.438561499 | 0.022118786 | ECT2 | turquoise | -0.251196 | 0.215781 |
| AAK1 | salmon | 0.292871 | 0.14651 | IFT43 | red | 0.439709083 | 0.021732959 | C2orf68 | turquoise | -0.251064 | 0.21603 |

**Table 6. Gene module membership and correlation with growth rate (48/72h), with adjusted p values for each module**

| | | | | | | | | | | | |
|---|---|---|---|---|---|---|---|---|---|---|---|
| SESN2 | salmon | 0.291445 | 0.148577 | RPL11 | red | 0.458208768 | 0.016234263 | RPL22L1 | turquoise | -0.251004 | 0.216144 |
| PPAP2B | salmon | 0.278945 | 0.167597 | PSMC1 | red | 0.459350966 | 0.015936414 | LDHA | turquoise | -0.250403 | 0.217283 |
| LRRC59 | salmon | 0.275286 | 0.173475 | MRPL24 | red | 0.464379324 | 0.014678244 | TMEM68 | turquoise | -0.250144 | 0.217775 |
| NDUFA7 | salmon | -0.271243 | 0.180135 | WDFY2 | red | 0.465135467 | 0.014496353 | DDX46 | turquoise | -0.249116 | 0.219733 |
| RRS1 | salmon | -0.262515 | 0.195113 | ADAM15 | red | 0.472496478 | 0.012821062 | TMTC4 | turquoise | -0.24906 | 0.219842 |
| IFIT5 | salmon | -0.258444 | 0.202383 | COMT | red | 0.47270702 | 0.012775617 | COX11 | turquoise | -0.248107 | 0.221669 |
| ARCN1 | salmon | 0.25776 | 0.203623 | MAGEH1 | red | 0.476230293 | 0.012034748 | SMPDL3A | turquoise | -0.247294 | 0.223236 |
| PDSS1 | salmon | -0.254989 | 0.208698 | PHF23 | red | 0.479136075 | 0.011450925 | HINT1 | turquoise | -0.247153 | 0.223509 |
| TARS | salmon | 0.253553 | 0.211361 | CHDH | red | 0.48142924 | 0.011007004 | EXTL3 | turquoise | -0.246154 | 0.225446 |
| BAIAP2 | salmon | 0.252184 | 0.21392 | APRT | red | 0.484099059 | 0.010508328 | IPO9 | turquoise | -0.24499 | 0.227717 |
| TYMS | salmon | -0.245174 | 0.227358 | NDUFA1 | red | 0.487814367 | 0.009845849 | TPP1 | turquoise | -0.244629 | 0.228425 |
| DYNLT1 | salmon | -0.244596 | 0.228489 | FGGY | red | 0.498729374 | 0.008098101 | SH3YL1 | turquoise | -0.244449 | 0.228778 |
| FNIP2 | salmon | 0.24143 | 0.234759 | ROMO1 | red | 0.502085117 | 0.00761628 | POLR2C | turquoise | -0.24444 | 0.228795 |
| SNX8 | salmon | 0.241196 | 0.235228 | C10orf11 | red | 0.505496907 | 0.007151348 | ATP5C1 | turquoise | -0.243706 | 0.23024 |
| MICALL1 | salmon | 0.241171 | 0.235277 | TMEM219 | red | 0.510548927 | 0.00650691 | FAM114A1 | turquoise | -0.243375 | 0.230895 |
| TSPAN13 | salmon | -0.240081 | 0.237465 | DDX55 | red | 0.517670884 | 0.005682102 | RPLP0 | turquoise | -0.242676 | 0.232278 |
| C17orf59 | salmon | -0.23412 | 0.249667 | PFDN5 | red | 0.520895676 | 0.005338796 | FAM104B | turquoise | -0.242448 | 0.232732 |
| NINJ1 | salmon | -0.230384 | 0.25752 | RPS23 | red | 0.545493522 | 0.003252034 | PYURF | turquoise | -0.240908 | 0.235803 |
| PRRG4 | salmon | -0.230077 | 0.258173 | RPL37A | red | 0.571062572 | 0.00186443 | GBAS | turquoise | -0.240891 | 0.235839 |
| SNHG11 | salmon | -0.227382 | 0.263945 | SNRPA1 | red | 0.571670601 | 0.001838922 | ATG5 | turquoise | -0.240336 | 0.236952 |
| HYOU1 | salmon | 0.222048 | 0.275614 | DEM1 | red | 0.580399305 | 0.001504521 | RHOBTB1 | turquoise | -0.240002 | 0.237624 |
| NRP2 | salmon | 0.220506 | 0.279048 | LAMTOR2 | red | 0.602093428 | 0.000891058 | ZDHHC20 | turquoise | -0.239942 | 0.237745 |
| COX5B | salmon | -0.213937 | 0.29398 | NDUFA3 | red | 0.603910873 | 0.000851352 | SDF2L1 | turquoise | -0.239618 | 0.238399 |
| ADORA2B | salmon | 0.212546 | 0.297204 | NDUFB5 | red | 0.607988403 | 0.00076782 | SNORA10 | turquoise | -0.23868 | 0.240298 |
| DISP1 | salmon | 0.21003 | 0.303094 | SSH1 | royalblue | -0.574110122 | 0.001739578 | MRPL30 | turquoise | -0.237482 | 0.242735 |
| SGTA | salmon | 0.207461 | 0.309182 | KIFAP3 | royalblue | -0.46350785 | 0.014890215 | ETFA | turquoise | -0.237386 | 0.242931 |
| C8B | salmon | -0.206319 | 0.311911 | MAMLD1 | royalblue | -0.442650156 | 0.020769171 | EIF4E | turquoise | -0.236613 | 0.244516 |
| KIAA1804 | salmon | 0.205477 | 0.313935 | GTF3C1 | royalblue | -0.428774176 | 0.025641677 | OFD1 | turquoise | -0.235912 | 0.245957 |
| HIST1H2AB | salmon | -0.202553 | 0.32102 | YTHDC2 | royalblue | -0.401013767 | 0.038167845 | PPP2R5A | turquoise | -0.235904 | 0.245974 |
| CHRM4 | salmon | -0.200966 | 0.324908 | NOL10 | royalblue | -0.330904084 | 0.091806952 | NCKIPSD | turquoise | -0.235081 | 0.247673 |
| FAM83D | salmon | -0.194477 | 0.341095 | SPRED2 | royalblue | -0.315892764 | 0.108456823 | GORASP2 | turquoise | -0.234666 | 0.248533 |
| MSRB2 | salmon | -0.193589 | 0.343347 | MS4A7 | royalblue | -0.311658345 | 0.11353523 | ACADM | turquoise | -0.234512 | 0.248853 |
| MANF | salmon | 0.189839 | 0.352957 | USP42 | royalblue | -0.302761522 | 0.124774467 | CFL2 | turquoise | -0.233748 | 0.250441 |
| HIST1H2AH | salmon | -0.186442 | 0.361796 | SMARCC2 | royalblue | -0.298019087 | 0.131087973 | EIF4E3 | turquoise | -0.232824 | 0.252372 |
| KLC1 | salmon | 0.1829 | 0.37115 | DPP3 | royalblue | -0.297229475 | 0.13216133 | C11orf74 | turquoise | -0.232781 | 0.252463 |
| TMEM183A | salmon | -0.181869 | 0.3739 | HMBOX1 | royalblue | -0.239445363 | 0.229008687 | ATP6V1B2 | turquoise | -0.231725 | 0.254683 |
| PNPLA1 | salmon | -0.180366 | 0.377926 | MYADM | royalblue | -0.210090791 | 0.292889656 | HSD17B12 | turquoise | -0.231485 | 0.255189 |
| AP2S1 | salmon | -0.17718 | 0.38655 | STX16 | royalblue | -0.175297808 | 0.38180621 | NAA15 | turquoise | -0.23101 | 0.256193 |
| C17orf79 | salmon | -0.172634 | 0.399043 | ISOC1 | royalblue | -0.161322938 | 0.421466699 | FASTKD1 | turquoise | -0.23072 | 0.256808 |
| HIST1H4A | salmon | -0.171748 | 0.401504 | PRRG1 | royalblue | -0.145609049 | 0.46864756 | MRPS28 | turquoise | -0.230321 | 0.257653 |
| HIST1H2BI | salmon | -0.171498 | 0.4022 | ZFP1 | royalblue | -0.111460467 | 0.579928102 | NUP85 | turquoise | -0.22933 | 0.259764 |
| RETN | salmon | -0.171498 | 0.402202 | METTL14 | royalblue | -0.109267664 | 0.587450567 | DTWD1 | turquoise | -0.229281 | 0.25987 |
| C1orf63 | salmon | -0.171295 | 0.402767 | LANCL1 | royalblue | -0.104024122 | 0.605608056 | ARL2 | turquoise | -0.229016 | 0.260435 |
| ABL1 | salmon | 0.169769 | 0.407033 | NSD1 | royalblue | -0.07347402 | 0.715701452 | PAIP2 | turquoise | -0.22837 | 0.261819 |
| HIST1H3A | salmon | -0.167286 | 0.414029 | PPP1R14B | royalblue | -0.061566983 | 0.760318089 | TRIM37 | turquoise | -0.227934 | 0.262756 |
| SNHG12 | salmon | -0.161462 | 0.430698 | CDCA7L | royalblue | -0.05872451 | 0.771086745 | FAM60A | turquoise | -0.226233 | 0.266431 |
| MMP19 | salmon | 0.160517 | 0.433435 | SNORA67 | royalblue | -0.053865406 | 0.789590605 | HCFC1R1 | turquoise | -0.226163 | 0.266583 |
| HBP1 | salmon | -0.156654 | 0.444727 | HIST1H2BE | royalblue | -0.051241295 | 0.799630632 | TFAM | turquoise | -0.226038 | 0.266855 |
| DOT1L | salmon | 0.156302 | 0.445764 | NOMO1 | royalblue | -0.045219096 | 0.82278739 | TMEM65 | turquoise | -0.225578 | 0.267856 |
| RANBP6 | salmon | -0.151676 | 0.459509 | FTSJD2 | royalblue | -0.033134517 | 0.869676233 | CMC1 | turquoise | -0.225354 | 0.268343 |
| UBE2J1 | salmon | 0.14888 | 0.46792 | STX8 | royalblue | -0.02645078 | 0.895805931 | B3GALT4 | turquoise | -0.225265 | 0.268537 |
| NFE2L2 | salmon | 0.142828 | 0.486399 | PEX10 | royalblue | -0.01437999 | 0.943247801 | RAB5A | turquoise | -0.224126 | 0.27103 |
| ZNF841 | salmon | 0.141171 | 0.491523 | USP30 | royalblue | -0.01361269 | 0.946271611 | ZNF284 | turquoise | -0.223975 | 0.271362 |
| PTPRE | salmon | 0.139064 | 0.498077 | TMEM206 | royalblue | 0.057875117 | 0.774312807 | UEVLD | turquoise | -0.223057 | 0.273383 |
| PRPF3 | salmon | -0.135067 | 0.510627 | ASNSD1 | royalblue | 0.058936139 | 0.770283537 | ZNF548 | turquoise | -0.222821 | 0.273903 |
| JOSD1 | salmon | 0.134718 | 0.511728 | H3F3A | royalblue | 0.068124942 | 0.735642081 | TUBA1B | turquoise | -0.222784 | 0.273985 |

**Table 6. Gene module membership and correlation with growth rate (48/72h), with adjusted p values for each module**

| | | | | | | | | | | | |
|---|---|---|---|---|---|---|---|---|---|---|---|
| ENTPD7 | salmon | 0.134687 | 0.511829 | RANGRF | royalblue | 0.102014937 | 0.612627426 | PIK3R4 | turquoise | -0.222274 | 0.275114 |
| PHF23 | salmon | -0.131006 | 0.523535 | RPS16 | royalblue | 0.112568106 | 0.576144511 | CLN5 | turquoise | -0.22202 | 0.275677 |
| PAFAH1B1 | salmon | -0.129947 | 0.526926 | FAM114A1 | royalblue | 0.118673457 | 0.555488255 | TTC1 | turquoise | -0.22192 | 0.275898 |
| EDN1 | salmon | 0.129294 | 0.529021 | SNORA71C | royalblue | 0.153199988 | 0.445521339 | BSCL2 | turquoise | -0.221476 | 0.276885 |
| CFLAR | salmon | 0.128627 | 0.531167 | SNORA70 | royalblue | 0.155412007 | 0.438899182 | FAM111A | turquoise | -0.220908 | 0.27815 |
| TMEM11 | salmon | 0.127285 | 0.535496 | NDUFB11 | royalblue | 0.163264777 | 0.415824237 | GSN | turquoise | -0.220782 | 0.278431 |
| ABCF1 | salmon | -0.126635 | 0.537598 | C17orf79 | royalblue | 0.164167552 | 0.413215344 | SRF | turquoise | -0.22049 | 0.279085 |
| PINLYP | salmon | -0.124067 | 0.545945 | LSM7 | royalblue | 0.170430613 | 0.395368273 | C6orf162 | turquoise | -0.220053 | 0.280061 |
| TMEM217 | salmon | 0.12256 | 0.55087 | GADD45GIP | royalblue | 0.186756765 | 0.350954108 | CCT2 | turquoise | -0.219746 | 0.280751 |
| HSPA6 | salmon | 0.122212 | 0.55201 | MRPS9 | royalblue | 0.190616458 | 0.340906477 | NUDT19 | turquoise | -0.21945 | 0.281416 |
| TMBIM4 | salmon | -0.120733 | 0.556868 | ANKRD10 | royalblue | 0.199927355 | 0.317389252 | NDUFA5 | turquoise | -0.219042 | 0.282332 |
| TMCO6 | salmon | -0.119274 | 0.56168 | DDX28 | royalblue | 0.201725471 | 0.312965596 | SF3A3 | turquoise | -0.218738 | 0.283019 |
| HIST1H2BK | salmon | -0.119143 | 0.562113 | CYC1 | royalblue | 0.205804555 | 0.303072447 | RAP1A | turquoise | -0.218626 | 0.28327 |
| MARCKS | salmon | -0.112372 | 0.584693 | H3F3B | royalblue | 0.210506962 | 0.291912581 | CAMTA1 | turquoise | -0.218502 | 0.283551 |
| IL1RN | salmon | 0.110929 | 0.589556 | NTAN1 | royalblue | 0.211945278 | 0.288551596 | TMEM126B | turquoise | -0.216962 | 0.287043 |
| CLN8 | salmon | 0.110046 | 0.59254 | FDX1L | royalblue | 0.224940866 | 0.259298257 | ARAF | turquoise | -0.216511 | 0.28807 |
| IMPA1 | salmon | 0.106776 | 0.603646 | RING1 | royalblue | 0.249983527 | 0.208547526 | RHOF | turquoise | -0.2164 | 0.288324 |
| PDIA5 | salmon | -0.103947 | 0.613325 | CMC2 | royalblue | 0.256558105 | 0.196432141 | PLAC8 | turquoise | -0.21587 | 0.289534 |
| RAPGEF1 | salmon | 0.101607 | 0.621379 | NUDC | royalblue | 0.259140773 | 0.191808052 | POP4 | turquoise | -0.215569 | 0.290224 |
| SLC7A11 | salmon | 0.099943 | 0.627134 | IRF2BP1 | royalblue | 0.267072863 | 0.1780776 | IL13RA1 | turquoise | -0.214396 | 0.292921 |
| RAB35 | salmon | 0.09352 | 0.649536 | PFN1 | royalblue | 0.273010049 | 0.168260249 | ACAT2 | turquoise | -0.214164 | 0.293456 |
| BRD1 | salmon | 0.092503 | 0.653111 | CHCHD2 | royalblue | 0.274093714 | 0.16651037 | ANAPC10 | turquoise | -0.214029 | 0.293767 |
| DHDDS | salmon | 0.085671 | 0.677322 | CCDC12 | royalblue | 0.277368163 | 0.161300958 | TCEANC2 | turquoise | -0.213983 | 0.293873 |
| EAF1 | salmon | 0.08277 | 0.687698 | MYEOV2 | royalblue | 0.282584964 | 0.153241762 | ATAD2B | turquoise | -0.213911 | 0.29404 |
| CXCL3 | salmon | -0.081037 | 0.693923 | TRMT2A | royalblue | 0.289684842 | 0.142741024 | TGFA | turquoise | -0.213665 | 0.294608 |
| ZNF546 | salmon | 0.079938 | 0.697884 | SCARNA7 | royalblue | 0.290041424 | 0.142227674 | PIP5K1C | turquoise | -0.213399 | 0.295225 |
| TMEM19 | salmon | -0.078455 | 0.703234 | SPATA5L1 | royalblue | 0.321315898 | 0.102201543 | C5orf42 | turquoise | -0.213139 | 0.295828 |
| CSF1 | salmon | 0.076322 | 0.710959 | RRS1 | royalblue | 0.323714057 | 0.099522969 | BCKDHA | turquoise | -0.212984 | 0.296188 |
| OSGIN1 | salmon | 0.075605 | 0.71356 | POLR2F | royalblue | 0.325831434 | 0.097201997 | IKBIP | turquoise | -0.212343 | 0.297677 |
| AKR1C2 | salmon | 0.072859 | 0.723556 | C6orf1 | royalblue | 0.340456798 | 0.08226402 | ZNF451 | turquoise | -0.212275 | 0.297837 |
| SURF4 | salmon | -0.072159 | 0.726112 | RPL18 | royalblue | 0.343344382 | 0.079533495 | CENPC1 | turquoise | -0.212229 | 0.297944 |
| SOWAHC | salmon | 0.069728 | 0.735009 | FAU | royalblue | 0.344132295 | 0.078800582 | KIAA1551 | turquoise | -0.212002 | 0.298471 |
| IL6R | salmon | -0.066317 | 0.747548 | NDUFB9 | royalblue | 0.346378905 | 0.076739093 | BTF3L4 | turquoise | -0.211873 | 0.298773 |
| UBE2Z | salmon | 0.062737 | 0.760773 | SEPHS1 | royalblue | 0.350442704 | 0.07311524 | C18orf21 | turquoise | -0.211563 | 0.299498 |
| APBA3 | salmon | -0.061764 | 0.764379 | EXOSC1 | royalblue | 0.352591437 | 0.071252999 | HUS1 | turquoise | -0.210588 | 0.301782 |
| SRSF7 | salmon | -0.058873 | 0.775123 | EMC9 | royalblue | 0.353456794 | 0.070513408 | NOC2L | turquoise | -0.21053 | 0.301919 |
| PDGFC | salmon | -0.056107 | 0.785443 | AIMP1 | royalblue | 0.358095407 | 0.066649181 | VAMP3 | turquoise | -0.210522 | 0.301937 |
| LOC730101 | salmon | -0.054664 | 0.790837 | PRDX5 | royalblue | 0.380373818 | 0.050319361 | MPHOSPH6 | turquoise | -0.210031 | 0.303091 |
| YRDC | salmon | -0.051634 | 0.802198 | TCEB2 | royalblue | 0.386002703 | 0.046739047 | GRHPR | turquoise | -0.209585 | 0.304144 |
| SLC39A14 | salmon | -0.050534 | 0.806335 | ZNF688 | royalblue | 0.397232904 | 0.040198124 | CDK7 | turquoise | -0.209353 | 0.30469 |
| CD274 | salmon | 0.047415 | 0.818085 | COX5B | royalblue | 0.405723748 | 0.035753963 | CRIPT | turquoise | -0.209095 | 0.305302 |
| PDRG1 | salmon | -0.046602 | 0.821154 | PET100 | royalblue | 0.416759662 | 0.030574695 | HNRPLL | turquoise | -0.208944 | 0.305658 |
| PRMT3 | salmon | 0.042152 | 0.837999 | EIF3K | royalblue | 0.45557129 | 0.016939591 | EVI2B | turquoise | -0.208862 | 0.305854 |
| GCLM | salmon | -0.040639 | 0.843742 | RNF113A | royalblue | 0.470544082 | 0.013248904 | LOC728554 | turquoise | -0.208522 | 0.306657 |
| SNORA33 | salmon | 0.038015 | 0.853722 | PSMA7 | royalblue | 0.490198247 | 0.009439445 | ELP4 | turquoise | -0.208331 | 0.307112 |
| PIM3 | salmon | -0.035473 | 0.863411 | RPUSD2 | royalblue | 0.504747615 | 0.007251353 | SP140L | turquoise | -0.208121 | 0.307611 |
| LOC643837 | salmon | -0.033977 | 0.869121 | PRDX6 | royalblue | 0.529243152 | 0.004530667 | UBE2L6 | turquoise | -0.207919 | 0.308091 |
| SHMT2 | salmon | 0.033119 | 0.8724 | INTS8 | salmon | -0.508397121 | 0.006775138 | LRRC40 | turquoise | -0.207825 | 0.308316 |
| ARFGAP3 | salmon | -0.03142 | 0.878895 | GDAP2 | salmon | -0.460395974 | 0.015667871 | NSFP1 | turquoise | -0.207556 | 0.308955 |
| YKT6 | salmon | -0.028142 | 0.891455 | APPL1 | salmon | -0.428563017 | 0.02572242 | HLA-DMA | turquoise | -0.207437 | 0.309239 |
| KLHL15 | salmon | -0.02808 | 0.891691 | ANGEL2 | salmon | -0.419055052 | 0.02957747 | IFNAR1 | turquoise | -0.207419 | 0.309283 |
| LPCAT1 | salmon | 0.027847 | 0.892585 | TXNDC12 | salmon | -0.414655809 | 0.031512331 | MAP3K7 | turquoise | -0.207295 | 0.309577 |
| TOM1L2 | salmon | -0.027766 | 0.892895 | MRPS35 | salmon | -0.401796735 | 0.037757807 | MRPS17 | turquoise | -0.207037 | 0.310194 |
| TUBB6 | salmon | -0.027034 | 0.895704 | PTAR1 | salmon | -0.39031586 | 0.044134203 | FAM193A | turquoise | -0.20697 | 0.310353 |
| CBLL1 | salmon | -0.023366 | 0.909794 | DDHD2 | salmon | -0.382092953 | 0.049203752 | NSL1 | turquoise | -0.20688 | 0.31057 |
| PELI1 | salmon | 0.022188 | 0.914325 | PIK3CA | salmon | -0.379866962 | 0.050652046 | DTNBP1 | turquoise | -0.205895 | 0.312928 |

**Table 6. Gene module membership and correlation with growth rate (48/72h), with adjusted p values for each module**

| | | | | | | | | | | | |
|---|---|---|---|---|---|---|---|---|---|---|---|
| FBXL15 | salmon | -0.021941 | 0.915275 | TFB2M | salmon | -0.373970695 | 0.054650539 | ECH1 | turquoise | -0.205247 | 0.314489 |
| BTG1 | salmon | 0.021321 | 0.91766 | LEO1 | salmon | -0.36569276 | 0.060676001 | SMC4 | turquoise | -0.204931 | 0.315249 |
| TNFRSF1B | salmon | 0.020522 | 0.920738 | UBA6 | salmon | -0.365198382 | 0.061051557 | AKR1B1 | turquoise | -0.204443 | 0.316429 |
| PLEK | salmon | 0.018726 | 0.927656 | PPP3CC | salmon | -0.363876893 | 0.062064303 | TM9SF1 | turquoise | -0.203277 | 0.319256 |
| HIST1H4I | salmon | 0.017795 | 0.931244 | NSDHL | salmon | -0.360564776 | 0.06465992 | UBR7 | turquoise | -0.202686 | 0.320697 |
| BCL6 | salmon | -0.016599 | 0.935854 | GRB2 | salmon | -0.358799137 | 0.066077519 | HMGB2 | turquoise | -0.202595 | 0.320917 |
| TULP3 | salmon | -0.016235 | 0.937258 | REEP3 | salmon | -0.357661126 | 0.067003863 | DICER1 | turquoise | -0.201911 | 0.32259 |
| ZCCHC14 | salmon | -0.014732 | 0.943056 | BROX | salmon | -0.356503003 | 0.067956842 | C8orf40 | turquoise | -0.201455 | 0.323707 |
| FDXACB1 | salmon | -0.013291 | 0.948618 | PSMD10 | salmon | -0.352853229 | 0.071028628 | RPF2 | turquoise | -0.201106 | 0.324563 |
| DGCR8 | salmon | 0.00894 | 0.965428 | NPAT | salmon | -0.352406817 | 0.071411556 | C9orf41 | turquoise | -0.200941 | 0.32497 |
| POLR1E | salmon | -0.007298 | 0.971775 | FAR1 | salmon | -0.351138611 | 0.072508074 | TRIM27 | turquoise | -0.199173 | 0.329331 |
| OC1002870 | salmon | 0.004006 | 0.984505 | DOCK10 | salmon | -0.346998808 | 0.076177596 | AIM2 | turquoise | -0.199144 | 0.329405 |
| NCOA5 | salmon | -0.003327 | 0.987129 | C17orf61 | salmon | -0.332745918 | 0.089904993 | FTSJ1 | turquoise | -0.198316 | 0.33146 |
| TXNDC5 | salmon | -0.002486 | 0.990385 | UTP14A | salmon | -0.330872284 | 0.091840054 | EIF3I | turquoise | -0.197896 | 0.332506 |
| MOGS | salmon | -0.000709 | 0.997256 | CRBN | salmon | -0.33080522 | 0.091909894 | DTX4 | turquoise | -0.197454 | 0.333611 |
| IER5L | salmon | 0.000391 | 0.998488 | IFT46 | salmon | -0.329134051 | 0.093663198 | ZNF473 | turquoise | -0.197029 | 0.334672 |
| CRAT | skyblue | -0.436877 | 0.025643 | NAT1 | salmon | -0.32219005 | 0.101219015 | DCAF17 | turquoise | -0.196599 | 0.335749 |
| ERGIC1 | skyblue | -0.397633 | 0.044258 | TNKS | salmon | -0.321237229 | 0.102290313 | TTLL1 | turquoise | -0.195826 | 0.33769 |
| CHMP7 | skyblue | -0.379365 | 0.055952 | FRS2 | salmon | -0.315594506 | 0.108808908 | ZNF7 | turquoise | -0.195823 | 0.337699 |
| CYBASC3 | skyblue | -0.340513 | 0.088721 | PBRM1 | salmon | -0.312418379 | 0.11261101 | SUGT1 | turquoise | -0.195556 | 0.338372 |
| ETFA | skyblue | -0.269876 | 0.182426 | TAX1BP1 | salmon | -0.302898186 | 0.12459589 | APPL2 | turquoise | -0.195439 | 0.338666 |
| TP53I13 | skyblue | -0.237166 | 0.243381 | PTP4A2 | salmon | -0.300593984 | 0.12763186 | PRR11 | turquoise | -0.195111 | 0.339492 |
| RCAN1 | skyblue | -0.230157 | 0.258002 | CCNT1 | salmon | -0.295613749 | 0.13437755 | LOC728377 | turquoise | -0.194899 | 0.340027 |
| ARMCX3 | skyblue | -0.199933 | 0.327453 | PRMT6 | salmon | -0.292684502 | 0.138464007 | DNM2 | turquoise | -0.194377 | 0.341348 |
| IFITM2 | skyblue | -0.195813 | 0.337723 | ATP10D | salmon | -0.283482676 | 0.151884454 | ORC2 | turquoise | -0.194076 | 0.342112 |
| ARHGEF5 | skyblue | -0.178011 | 0.384291 | RABEPK | salmon | -0.276899559 | 0.162039302 | ZNF207 | turquoise | -0.194031 | 0.342226 |
| CREB3L2 | skyblue | -0.177478 | 0.385739 | BTBD1 | salmon | -0.273425341 | 0.16758812 | PIK3R1 | turquoise | -0.193464 | 0.343665 |
| ZNF138 | skyblue | 0.173479 | 0.396705 | RAP2C | salmon | -0.269336117 | 0.174289149 | TMEM237 | turquoise | -0.193329 | 0.344008 |
| RELA | skyblue | -0.172561 | 0.399248 | DDX21 | salmon | -0.268955983 | 0.174921472 | UBE2W | turquoise | -0.193038 | 0.34475 |
| GNA11 | skyblue | -0.166731 | 0.415602 | SP4 | salmon | -0.267819616 | 0.176821311 | PSMA4 | turquoise | -0.192962 | 0.344942 |
| FAM103A1 | skyblue | -0.156656 | 0.444723 | ZNF383 | salmon | -0.267272612 | 0.177740947 | NUDT16P1 | turquoise | -0.192509 | 0.346097 |
| QSOX2 | skyblue | -0.147159 | 0.473139 | CDK17 | salmon | -0.264344851 | 0.182719982 | NAE1 | turquoise | -0.192245 | 0.346772 |
| DOM3Z | skyblue | -0.135973 | 0.507769 | MMGT1 | salmon | -0.264207838 | 0.182955342 | CCNC | turquoise | -0.192164 | 0.346981 |
| NOC2L | skyblue | -0.135921 | 0.507933 | YARS2 | salmon | -0.235865053 | 0.236255183 | ICA1 | turquoise | -0.191399 | 0.34894 |
| ACO2 | skyblue | -0.135059 | 0.510652 | DESI2 | salmon | -0.223191044 | 0.263120364 | RBM48 | turquoise | -0.190651 | 0.350863 |
| C6orf170 | skyblue | -0.134995 | 0.510853 | NHLRC2 | salmon | -0.221686959 | 0.266434727 | SIGLEC10 | turquoise | -0.190281 | 0.351816 |
| MYO9A | skyblue | 0.124531 | 0.544433 | C14orf119 | salmon | -0.219107652 | 0.272180905 | CCRN4L | turquoise | -0.190162 | 0.352123 |
| RFWD2 | skyblue | -0.116009 | 0.572515 | MTPN | salmon | -0.214527212 | 0.282579935 | KDELC2 | turquoise | -0.190146 | 0.352163 |
| ATG10 | skyblue | 0.115503 | 0.574202 | SUCO | salmon | -0.214186955 | 0.283362371 | DNAJC13 | turquoise | -0.190143 | 0.352172 |
| ARMC5 | skyblue | -0.107651 | 0.600665 | GTF3C3 | salmon | -0.212467201 | 0.28733807 | LPGAT1 | turquoise | -0.190041 | 0.352435 |
| USP48 | skyblue | -0.075363 | 0.714439 | XPOT | salmon | -0.212435739 | 0.287411131 | RPAP2 | turquoise | -0.189946 | 0.352679 |
| RAB40C | skyblue | 0.062128 | 0.763031 | ZMYND8 | salmon | -0.193527305 | 0.333444543 | IGHMBP2 | turquoise | -0.189899 | 0.352801 |
| CUL4B | skyblue | -0.060792 | 0.767989 | SRSF10 | salmon | -0.184258389 | 0.357550802 | CALM2 | turquoise | -0.189717 | 0.35327 |
| ERO1LB | skyblue | 0.057701 | 0.779492 | DHRSX | salmon | -0.180195554 | 0.368433397 | CNST | turquoise | -0.189632 | 0.353492 |
| CD276 | skyblue | 0.041464 | 0.840611 | CTR9 | salmon | -0.175658563 | 0.380811783 | ACER3 | turquoise | -0.189301 | 0.354347 |
| C6orf106 | skyblue | 0.039471 | 0.848181 | AGPS | salmon | -0.175502505 | 0.381241777 | TXNDC15 | turquoise | -0.189265 | 0.354441 |
| CYTH2 | skyblue | -0.035947 | 0.861601 | LPAR2 | salmon | -0.174945382 | 0.38277912 | NDUFAF1 | turquoise | -0.188904 | 0.355375 |
| RBM14 | skyblue | -0.033627 | 0.870458 | HEATR2 | salmon | -0.168784074 | 0.40001729 | UCP2 | turquoise | -0.188494 | 0.35644 |
| DDX42 | skyblue | -0.03302 | 0.872779 | KATNB1 | salmon | -0.167143766 | 0.404679226 | RAB27A | turquoise | -0.188125 | 0.3574 |
| SLC38A7 | skyblue | -0.029782 | 0.885169 | TMEM192 | salmon | -0.162214985 | 0.418869435 | SPDL1 | turquoise | -0.18761 | 0.358742 |
| UBA3 | skyblue | -0.021093 | 0.918539 | RBM5 | salmon | -0.158274977 | 0.430407659 | MRPL27 | turquoise | -0.187439 | 0.359189 |
| SLC9A3R1 | skyblue | -0.020547 | 0.920642 | GIN1 | salmon | -0.155699849 | 0.438041387 | RHOQ | turquoise | -0.186876 | 0.360659 |
| PCED1B-AS | skyblue | -0.013448 | 0.948013 | UBR1 | salmon | -0.155639843 | 0.438220135 | PTPN22 | turquoise | -0.186782 | 0.360905 |
| SOGA1 | skyblue | -0.008985 | 0.965254 | SLC25A24 | salmon | -0.152238251 | 0.448417081 | HEXB | turquoise | -0.186657 | 0.361233 |
| NOM1 | tan | -0.482387 | 0.012568 | TJAP1 | salmon | -0.151578573 | 0.450409124 | NDUFB1 | turquoise | -0.185857 | 0.363332 |
| SNX18 | tan | -0.44525 | 0.022643 | ARFGAP1 | salmon | -0.148782298 | 0.458905084 | ADNP | turquoise | -0.185801 | 0.363479 |

**Table 6. Gene module membership and correlation with growth rate (48/72h), with adjusted p values for each module**

| | | | | | | | | | | | |
|---|---|---|---|---|---|---|---|---|---|---|---|
| TAF10 | tan | -0.425849 | 0.030075 | SS18 | salmon | -0.148557495 | 0.459591745 | NKIRAS1 | turquoise | -0.185788 | 0.363513 |
| ZNF510 | tan | -0.398392 | 0.043818 | TRAPPC1 | salmon | -0.136087804 | 0.498512654 | PGP | turquoise | -0.185151 | 0.365188 |
| ZDBF2 | tan | 0.386856 | 0.050896 | OSGIN2 | salmon | -0.129944992 | 0.518271747 | CNOT10 | turquoise | -0.185014 | 0.365552 |
| ZNF408 | tan | 0.382999 | 0.053453 | GRPEL2 | salmon | -0.128150234 | 0.524115607 | BCL10 | turquoise | -0.18482 | 0.366062 |
| HELB | tan | -0.371243 | 0.061868 | HS1BP3 | salmon | -0.126024798 | 0.531076917 | XPO1 | turquoise | -0.184756 | 0.366231 |
| CASP2 | tan | -0.362618 | 0.068668 | MAN2A1 | salmon | -0.123913849 | 0.53803405 | GFM2 | turquoise | -0.184675 | 0.366446 |
| KCNQ1 | tan | -0.35877 | 0.071881 | ERCC6L2 | salmon | -0.123504061 | 0.539389561 | TLR5 | turquoise | -0.184467 | 0.366994 |
| PAGR1 | tan | -0.355086 | 0.075065 | DDX5 | salmon | -0.123476173 | 0.53948187 | NMI | turquoise | -0.184425 | 0.367107 |
| ZCWPW1 | tan | 0.351283 | 0.078462 | HMGN2 | salmon | -0.114478198 | 0.569645639 | CEP41 | turquoise | -0.184335 | 0.367344 |
| FBXO32 | tan | -0.335097 | 0.094246 | ANKRD36B | salmon | -0.109195493 | 0.587698868 | PDE6D | turquoise | -0.184251 | 0.367564 |
| C11orf31 | tan | -0.331507 | 0.098048 | ORMDL3 | salmon | -0.099931308 | 0.619942273 | YTHDF3 | turquoise | -0.183462 | 0.369656 |
| MSANTD4 | tan | -0.330901 | 0.098702 | SAMHD1 | salmon | -0.099001537 | 0.623217834 | C4orf27 | turquoise | -0.183212 | 0.37032 |
| TMEM230 | tan | -0.328698 | 0.101103 | LYN | salmon | -0.094494751 | 0.639193528 | USMG5 | turquoise | -0.182895 | 0.371162 |
| OR2A20P | tan | -0.322805 | 0.10774 | SLC25A17 | salmon | -0.093491887 | 0.64277029 | CXorf38 | turquoise | -0.182414 | 0.372444 |
| CGGBP1 | tan | -0.322488 | 0.108106 | DTX2P1-UPK3BP1-P | salmon | -0.092434552 | 0.646549754 | CGRRF1 | turquoise | -0.182003 | 0.373542 |
| COG8 | tan | -0.321254 | 0.10954 | ARL16 | salmon | -0.090448031 | 0.653673754 | CDC73 | turquoise | -0.181905 | 0.373802 |
| SYTL1 | tan | -0.319468 | 0.111639 | DCUN1D1 | salmon | -0.085896827 | 0.670106396 | FASTKD3 | turquoise | -0.181262 | 0.375524 |
| CLINT1 | tan | -0.319218 | 0.111935 | CUEDC2 | salmon | -0.08569171 | 0.670850567 | ITGB1 | turquoise | -0.180178 | 0.378433 |
| SIRT7 | tan | -0.316697 | 0.114954 | SENP2 | salmon | -0.078627112 | 0.696661906 | LSM1 | turquoise | -0.180057 | 0.378758 |
| PEX11B | tan | -0.308497 | 0.12519 | SIRT1 | salmon | -0.078595711 | 0.696777396 | C11orf73 | turquoise | -0.177883 | 0.384636 |
| ZNF26 | tan | 0.307169 | 0.126909 | ARMC7 | salmon | -0.076389214 | 0.704909216 | RAB14 | turquoise | -0.177805 | 0.384849 |
| SEC23A | tan | 0.302823 | 0.132653 | NPC2 | salmon | -0.074061983 | 0.713520402 | MOB3A | turquoise | -0.177249 | 0.386361 |
| ZNF280D | tan | -0.302601 | 0.132951 | CREB1 | salmon | -0.065550714 | 0.745299283 | RBM14 | turquoise | -0.176584 | 0.388175 |
| ZC2HC1A | tan | -0.295466 | 0.142801 | BMPR2 | salmon | -0.058849505 | 0.770612317 | ASMTL | turquoise | -0.176408 | 0.388654 |
| AGPAT3 | tan | -0.29322 | 0.146007 | HEATR1 | salmon | -0.052134289 | 0.796210402 | CFDP1 | turquoise | -0.176299 | 0.388952 |
| FGGY | tan | 0.292017 | 0.147745 | KCTD13 | salmon | -0.041717557 | 0.836319843 | TSSC1 | turquoise | -0.176229 | 0.389145 |
| TMEM109 | tan | -0.290913 | 0.149355 | KLHDC3 | salmon | -0.025492364 | 0.899562158 | B3GALT6 | turquoise | -0.175469 | 0.391226 |
| USF2 | tan | -0.286943 | 0.15524 | CLP1 | salmon | -0.024305934 | 0.904214937 | TPT1 | turquoise | -0.175208 | 0.391941 |
| RHOA | tan | -0.282533 | 0.161971 | STK24 | salmon | -0.018026096 | 0.928890242 | ITGB3BP | turquoise | -0.175082 | 0.392287 |
| AFG3L1P | tan | 0.279337 | 0.166976 | NSMAF | salmon | -0.012019102 | 0.952553973 | LACTB | turquoise | -0.174913 | 0.392751 |
| GTF2I | tan | -0.277066 | 0.170597 | BCLAF1 | salmon | -0.002335146 | 0.99077698 | TRAF3IP1 | turquoise | -0.174731 | 0.393251 |
| PPCS | tan | -0.27611 | 0.172138 | C8orf59 | salmon | 0.002350246 | 0.99071734 | SBF1 | turquoise | -0.174509 | 0.393863 |
| FGD3 | tan | -0.272291 | 0.178391 | DOHH | salmon | 0.005134244 | 0.979723132 | RPL6 | turquoise | -0.174076 | 0.395055 |
| TBL2 | tan | -0.271271 | 0.180088 | CLCC1 | salmon | 0.005508993 | 0.978243453 | INTU | turquoise | -0.174071 | 0.39507 |
| FUZ | tan | -0.271023 | 0.180502 | ABCA11P | salmon | 0.007177906 | 0.971654752 | ZNF83 | turquoise | -0.173505 | 0.396634 |
| TCF12 | tan | -0.267282 | 0.18683 | MED6 | salmon | 0.014829139 | 0.941478127 | CAPZA2 | turquoise | -0.173241 | 0.397363 |
| PARP1 | tan | -0.262664 | 0.19485 | MGC72080 | salmon | 0.016044348 | 0.936691512 | ZNHIT6 | turquoise | -0.172926 | 0.398234 |
| SLC25A40 | tan | -0.259995 | 0.199592 | SPATC1 | salmon | 0.018542119 | 0.926859906 | SRSF10 | turquoise | -0.172877 | 0.39837 |
| TBRG4 | tan | -0.259233 | 0.200961 | ANKRD13D | salmon | 0.02534614 | 0.900135427 | MRPL47 | turquoise | -0.172217 | 0.400202 |
| UNC93B1 | tan | -0.258276 | 0.202687 | SULT1A4 | salmon | 0.026612465 | 0.895172466 | ZNF746 | turquoise | -0.172125 | 0.400456 |
| STARD3 | tan | -0.25623 | 0.206414 | DUSP22 | salmon | 0.027964083 | 0.889879477 | GMPS | turquoise | -0.172081 | 0.40058 |
| PRR12 | tan | -0.255321 | 0.208085 | SNAPC2 | salmon | 0.028285883 | 0.888619966 | PMS1 | turquoise | -0.171851 | 0.401218 |
| CALHM2 | tan | -0.254703 | 0.209226 | ZNF345 | salmon | 0.029567116 | 0.883607923 | WIBG | turquoise | -0.171166 | 0.403128 |
| PBX3 | tan | -0.25095 | 0.216246 | SLC17A5 | salmon | 0.030024686 | 0.881819008 | TMEM109 | turquoise | -0.171093 | 0.403331 |
| FBL | tan | -0.249668 | 0.21868 | RBM39 | salmon | 0.031134436 | 0.877482696 | DMXL1 | turquoise | -0.170931 | 0.403783 |
| RBBP7 | tan | -0.244963 | 0.22777 | CAMKK2 | salmon | 0.031341156 | 0.87667532 | UBTD2 | turquoise | -0.170766 | 0.404244 |
| TAF8 | tan | -0.243996 | 0.22967 | CIRH1A | salmon | 0.03551509 | 0.860400256 | VPS35 | turquoise | -0.170441 | 0.405152 |
| RNF141 | tan | -0.241885 | 0.233851 | PAG1 | salmon | 0.043211905 | 0.830538819 | RBM34 | turquoise | -0.17043 | 0.405181 |
| C12orf57 | tan | 0.241601 | 0.234418 | ANKRD11 | salmon | 0.04646414 | 0.81798734 | DND1 | turquoise | -0.169269 | 0.408438 |
| USB1 | tan | -0.240454 | 0.236715 | PACSIN2 | salmon | 0.051903827 | 0.797092738 | PROSC | turquoise | -0.16882 | 0.4097 |
| MAGED2 | tan | -0.229345 | 0.259732 | FOXRED1 | salmon | 0.053849609 | 0.789650949 | API5 | turquoise | -0.168683 | 0.410086 |
| ATG16L2 | tan | -0.226159 | 0.266591 | RFC3 | salmon | 0.063876178 | 0.751601627 | POLG | turquoise | -0.168605 | 0.410304 |
| DHODH | tan | 0.225892 | 0.267172 | HERC2 | salmon | 0.06388409 | 0.751571815 | SEH1L | turquoise | -0.16852 | 0.410543 |
| IFT140 | tan | -0.224549 | 0.270103 | YPEL3 | salmon | 0.091946649 | 0.648296673 | LSAMP | turquoise | -0.168462 | 0.410707 |
| NFKBID | tan | 0.222536 | 0.274532 | LSP1 | salmon | 0.110599491 | 0.582876645 | CNOT2 | turquoise | -0.168111 | 0.411696 |
| CCDC47 | tan | 0.218647 | 0.283223 | LRRC14 | salmon | 0.121684774 | 0.545426771 | GPR160 | turquoise | -0.167743 | 0.412738 |

**Table 6. Gene module membership and correlation with growth rate (48/72h), with adjusted p values for each module**

| | | | | | | | | | | | |
|---|---|---|---|---|---|---|---|---|---|---|---|
| TMEM41A | tan | 0.21821 | 0.284211 | HNRNPU | salmon | 0.124932234 | 0.534672369 | SMURF2 | turquoise | -0.167629 | 0.413058 |
| PIP5K1C | tan | -0.214344 | 0.29304 | C15orf40 | salmon | 0.134606923 | 0.50324142 | PPA2 | turquoise | -0.167392 | 0.41373 |
| FASTKD2 | tan | 0.212876 | 0.296438 | TULP4 | salmon | 0.135054449 | 0.501810034 | PUS7L | turquoise | -0.166943 | 0.415001 |
| CMAHP | tan | -0.211115 | 0.300546 | IPO13 | salmon | 0.139891635 | 0.486468679 | SGK223 | turquoise | -0.16636 | 0.416654 |
| BTBD19 | tan | 0.210539 | 0.301898 | GLYCTK | salmon | 0.148887385 | 0.458584281 | OCIAD1 | turquoise | -0.16636 | 0.416656 |
| NUTF2 | tan | -0.210206 | 0.302679 | RAD21 | salmon | 0.150282875 | 0.45433542 | CLNS1A | turquoise | -0.166209 | 0.417084 |
| OR2A9P | tan | -0.208209 | 0.307402 | DDX3X | salmon | 0.154272267 | 0.44230459 | ACTR3 | turquoise | -0.166172 | 0.417191 |
| AGA | tan | -0.205938 | 0.312826 | GTPBP3 | salmon | 0.162686229 | 0.417500947 | FXR1 | turquoise | -0.1658 | 0.418248 |
| UBE2G1 | tan | -0.204991 | 0.315105 | CACNB3 | salmon | 0.173665 | 0.386325734 | SNX9 | turquoise | -0.165689 | 0.418564 |
| ZC3H15 | tan | -0.20478 | 0.315614 | CTDNEP1 | salmon | 0.181665623 | 0.364473588 | CCDC71L | turquoise | -0.165277 | 0.419738 |
| MAST3 | tan | -0.201186 | 0.324367 | LOC1005067[illegible] | salmon | 0.192598448 | 0.335814835 | HAGH | turquoise | -0.165112 | 0.42021 |
| MEAF6 | tan | -0.199866 | 0.327618 | QPCTL | salmon | 0.200871605 | 0.315061469 | ZC3H6 | turquoise | -0.164217 | 0.422769 |
| GPATCH2 | tan | -0.199531 | 0.328446 | BRMS1 | salmon | 0.201070171 | 0.314573306 | PRPF40A | turquoise | -0.164053 | 0.423238 |
| APOO | tan | 0.194968 | 0.339853 | NOSIP | salmon | 0.202487366 | 0.311102761 | RP9 | turquoise | -0.163444 | 0.424984 |
| VOPP1 | tan | -0.193573 | 0.343389 | RSAD1 | salmon | 0.212026523 | 0.28836248 | IRGQ | turquoise | -0.163418 | 0.425061 |
| USP32P1 | tan | 0.190755 | 0.350595 | B4GALT3 | salmon | 0.224023782 | 0.261296904 | PFDN4 | turquoise | -0.163264 | 0.425501 |
| EIF3F | tan | -0.187488 | 0.35906 | PMS2P1 | salmon | 0.228359246 | 0.251936154 | CCR5 | turquoise | -0.163231 | 0.425598 |
| PPARD | tan | -0.184019 | 0.368181 | MAP2K7 | salmon | 0.238719326 | 0.230466031 | BLOC1S2 | turquoise | -0.1632 | 0.425685 |
| BTBD6 | tan | -0.181851 | 0.373946 | BLOC1S3 | salmon | 0.296522521 | 0.133127734 | IQCG | turquoise | -0.162766 | 0.426934 |
| BABAM1 | tan | -0.180949 | 0.376362 | ABCD1 | salmon | 0.312450489 | 0.112572086 | METAP2 | turquoise | -0.162299 | 0.428279 |
| LPCAT4 | tan | -0.180737 | 0.376932 | LILRA2 | salmon | 0.3160053 | 0.108324197 | KIAA1586 | turquoise | -0.162299 | 0.42828 |
| TRIM14 | tan | -0.180541 | 0.377456 | CORO1B | salmon | 0.32559633 | 0.097457685 | ADAP1 | turquoise | -0.162288 | 0.428312 |
| TUBGCP4 | tan | 0.177586 | 0.385444 | C2CD2L | salmon | 0.34464093 | 0.078330198 | ARV1 | turquoise | -0.162242 | 0.428446 |
| PTER | tan | 0.174464 | 0.393986 | ABHD16A | salmon | 0.347828286 | 0.075431196 | RNF168 | turquoise | -0.162097 | 0.428863 |
| SBDS | tan | 0.167562 | 0.41325 | NDUFS5 | salmon | 0.374238574 | 0.054463684 | NFX1 | turquoise | -0.162076 | 0.428924 |
| PTGS2 | tan | 0.166783 | 0.415455 | SERBP1 | tan | -0.543333275 | 0.003401809 | KRR1 | turquoise | -0.161984 | 0.429189 |
| UBAP2 | tan | -0.166342 | 0.416708 | BRCC3 | tan | -0.522934691 | 0.005130921 | SIRPA | turquoise | -0.16193 | 0.429344 |
| RAB22A | tan | 0.163799 | 0.423966 | HIST1H1E | tan | -0.516931607 | 0.005763381 | ME1 | turquoise | -0.161897 | 0.42944 |
| ZNF253 | tan | -0.162659 | 0.427244 | CCDC84 | tan | -0.514967637 | 0.005984097 | CCDC112 | turquoise | -0.161778 | 0.429783 |
| AUP1 | tan | -0.161267 | 0.431262 | LOC729737 | tan | -0.500686534 | 0.007814076 | SURF1 | turquoise | -0.161476 | 0.430657 |
| FTSJ2 | tan | -0.159614 | 0.436062 | IK | tan | -0.496197193 | 0.008478386 | AMN1 | turquoise | -0.161024 | 0.431965 |
| SARNP | tan | 0.1589 | 0.438143 | REEP4 | tan | -0.493425711 | 0.008911662 | RFX7 | turquoise | -0.161007 | 0.432016 |
| APOBR | tan | -0.157872 | 0.44115 | KIAA1143 | tan | -0.48997034 | 0.009477681 | C3orf33 | turquoise | -0.160972 | 0.432118 |
| CAMTA2 | tan | -0.157163 | 0.443232 | FAM103A1 | tan | -0.488925813 | 0.009654587 | SLC44A2 | turquoise | -0.160869 | 0.432415 |
| SPPL2A | tan | 0.155918 | 0.446898 | UHRF1BP1 | tan | -0.472465745 | 0.012827707 | APTX | turquoise | -0.160255 | 0.434198 |
| KIAA0146 | tan | -0.155039 | 0.449495 | UIMC1 | tan | -0.435954933 | 0.023015918 | LARS2 | turquoise | -0.160218 | 0.434305 |
| PPP6C | tan | -0.152749 | 0.456301 | PRRG4 | tan | -0.430835498 | 0.024864221 | RSRC1 | turquoise | -0.160155 | 0.434487 |
| HIST1H4B | tan | -0.151325 | 0.460559 | PEA15 | tan | -0.422427314 | 0.028160151 | TTC3 | turquoise | -0.159555 | 0.436234 |
| RPL7 | tan | -0.150446 | 0.463198 | ELF1 | tan | -0.416380026 | 0.030742207 | BBS5 | turquoise | -0.159542 | 0.436271 |
| SERPINH1 | tan | 0.148883 | 0.46791 | ATP6V1G1 | tan | -0.407980798 | 0.034641339 | HOOK3 | turquoise | -0.15942 | 0.436628 |
| ZDHHC5 | tan | -0.147733 | 0.471393 | PPM1A | tan | -0.405682968 | 0.035774326 | NFKBIE | turquoise | -0.15893 | 0.438055 |
| PIEZO1 | tan | -0.143032 | 0.485772 | MRP63 | tan | -0.403870377 | 0.03668881 | EPG5 | turquoise | -0.158292 | 0.439921 |
| GABPA | tan | 0.138915 | 0.49854 | RABGEF1 | tan | -0.400931329 | 0.038211223 | USP34 | turquoise | -0.158254 | 0.440031 |
| CD97 | tan | -0.138718 | 0.499156 | MT1L | tan | -0.397860925 | 0.039855061 | C7orf23 | turquoise | -0.158232 | 0.440096 |
| GBA2 | tan | 0.136721 | 0.505414 | RHOA | tan | -0.390230934 | 0.044184359 | OR2A7 | turquoise | -0.157819 | 0.441307 |
| NUDT4 | tan | 0.133579 | 0.515336 | C11orf48 | tan | -0.389997723 | 0.044322319 | OGFRL1 | turquoise | -0.157504 | 0.442231 |
| IKBKE | tan | -0.132578 | 0.518517 | PSMD7 | tan | -0.382408628 | 0.049001031 | SUSD3 | turquoise | -0.157447 | 0.442399 |
| CLEC16A | tan | 0.128445 | 0.531754 | SSU72 | tan | -0.375819649 | 0.053370991 | LRPPRC | turquoise | -0.157115 | 0.443373 |
| BIRC2 | tan | 0.122729 | 0.550318 | POLD3 | tan | -0.369970692 | 0.05750078 | AP3S1 | turquoise | -0.156827 | 0.444218 |
| TBCCD1 | tan | -0.121229 | 0.555237 | AAR2 | tan | -0.351466071 | 0.072223714 | ATP5S | turquoise | -0.156193 | 0.446087 |
| RPL11 | tan | 0.117217 | 0.568497 | CHAF1A | tan | -0.349235961 | 0.074177342 | PRR13 | turquoise | -0.156189 | 0.446097 |
| MAGOH | tan | -0.116984 | 0.569272 | ZNF649 | tan | -0.345472876 | 0.077565443 | B4GALT7 | turquoise | -0.155475 | 0.448204 |
| TSC2 | tan | -0.116475 | 0.570963 | PHF5A | tan | -0.331122489 | 0.091579845 | NAP1L1 | turquoise | -0.155275 | 0.448798 |
| ANKMY1 | tan | -0.114954 | 0.576036 | PPIG | tan | -0.328798727 | 0.094018014 | MBIP | turquoise | -0.154603 | 0.450787 |
| TRAPPC9 | tan | -0.112788 | 0.583295 | MALT1 | tan | -0.307770329 | 0.118351018 | COX7B | turquoise | -0.15458 | 0.450854 |
| SEPN1 | tan | -0.111367 | 0.588076 | SNORA49 | tan | -0.293799911 | 0.136897496 | TBCEL | turquoise | -0.154407 | 0.451367 |

**Table 6. Gene module membership and correlation with growth rate (48/72h), with adjusted p values for each module**

| | | | |
|---|---|---|---|
| SLFN11 | tan | -0.110487 | 0.591048 |
| CYP4V2 | tan | -0.109399 | 0.594731 |
| RGS19 | tan | -0.103935 | 0.613366 |
| ZBTB1 | tan | -0.102153 | 0.619496 |
| TUBGCP6 | tan | -0.100361 | 0.625686 |
| SLC25A25 | tan | -0.098447 | 0.632322 |
| MGAM | tan | 0.097431 | 0.635857 |
| CPT1B | tan | 0.095353 | 0.643112 |
| DDX17 | tan | 0.093093 | 0.651039 |
| MCF2L2 | tan | 0.09283 | 0.651963 |
| TLR2 | tan | 0.090571 | 0.659926 |
| THAP10 | tan | -0.090335 | 0.660759 |
| SUZ12 | tan | -0.088978 | 0.665561 |
| SPATA13 | tan | 0.087105 | 0.672212 |
| EXOSC9 | tan | 0.082294 | 0.689408 |
| RAB11FIP1 | tan | -0.07778 | 0.705676 |
| TRIOBP | tan | -0.077722 | 0.705887 |
| TUBGCP2 | tan | -0.076869 | 0.708976 |
| GPR34 | tan | -0.069312 | 0.736532 |
| PLEKHA8 | tan | 0.068802 | 0.738407 |
| TRMT10B | tan | -0.065224 | 0.751577 |
| RPS6 | tan | 0.063988 | 0.756144 |
| NUP54 | tan | 0.063168 | 0.759177 |
| TCOF1 | tan | -0.062304 | 0.762379 |
| NEK11 | tan | -0.06002 | 0.770857 |
| ZCCHC4 | tan | -0.057306 | 0.780966 |
| RAB18 | tan | -0.055682 | 0.78703 |
| PIAS1 | tan | -0.052814 | 0.797769 |
| ATP6V0C | tan | -0.050904 | 0.804942 |
| DYNLT3 | tan | -0.050382 | 0.806907 |
| CMTM6 | tan | -0.049354 | 0.810775 |
| FERMT3 | tan | -0.046866 | 0.820158 |
| AGL | tan | -0.046824 | 0.820315 |
| DLAT | tan | -0.044306 | 0.829838 |
| RPL4 | tan | 0.041571 | 0.840203 |
| ADAM17 | tan | -0.035761 | 0.862312 |
| RNPC3 | tan | 0.035063 | 0.864975 |
| TOMM34 | tan | 0.034951 | 0.8654 |
| AP1S2 | tan | -0.033011 | 0.87281 |
| NUDT21 | tan | -0.031707 | 0.877798 |
| TCEA3 | tan | -0.0304 | 0.882801 |
| TAB2 | tan | 0.026225 | 0.89881 |
| CCDC109B | tan | 0.023042 | 0.911038 |
| CCDC126 | tan | 0.020265 | 0.921725 |
| ABCF2 | tan | 0.019727 | 0.923797 |
| RPS6KB1 | tan | 0.017992 | 0.930482 |
| GAK | tan | 0.011817 | 0.954311 |
| RAET1K | tan | -0.011122 | 0.956996 |
| ACP1 | tan | 0.007763 | 0.969976 |
| NOD1 | tan | 0.004144 | 0.983971 |
| KIAA1429 | tan | 0.003345 | 0.987061 |
| TAF7 | tan | -0.002452 | 0.990514 |
| KATNAL1 | turquoise | -0.560353 | 0.002908 |
| PPT2 | turquoise | -0.538935 | 0.0045 |
| ELF2 | turquoise | -0.517214 | 0.006814 |
| ARL17A | turquoise | 0.492664 | 0.010559 |
| EMC1 | tan | -0.286183243 | 0.147853118 |
| MRGBP | tan | -0.278678901 | 0.159248395 |
| SNORA53 | tan | -0.274953565 | 0.165131066 |
| FUNDC2 | tan | -0.27131684 | 0.171020273 |
| TFPT | tan | -0.268350986 | 0.175931151 |
| UBXN11 | tan | -0.262785564 | 0.185410973 |
| PTGR1 | tan | -0.260711678 | 0.189032494 |
| PNN | tan | -0.250110732 | 0.208308405 |
| TMOD3 | tan | -0.239484582 | 0.228930139 |
| FAF1 | tan | -0.233014251 | 0.242132825 |
| EIF3J | tan | -0.213529233 | 0.284878732 |
| ATP6V0B | tan | -0.187855413 | 0.348076324 |
| SNHG8 | tan | -0.149271265 | 0.457413398 |
| LRRC42 | tan | -0.123426934 | 0.539644866 |
| RPL28 | tan | -0.102915109 | 0.609478375 |
| RPL12 | tan | -0.095276386 | 0.636411239 |
| SLC39A4 | tan | -0.055174321 | 0.784594729 |
| B4GALT1 | tan | -0.010734632 | 0.957619738 |
| LRCH4 | tan | 0.018028401 | 0.928881168 |
| RASGRP1 | tan | 0.027953284 | 0.889921751 |
| UBE3B | tan | 0.046236269 | 0.818865386 |
| HVCN1 | tan | 0.058298762 | 0.772703308 |
| RPS28 | tan | 0.069900344 | 0.729004344 |
| MSN | tan | 0.080546238 | 0.689616028 |
| NUP188 | tan | 0.089173263 | 0.658260989 |
| RPS29 | tan | 0.096344325 | 0.632617597 |
| MAF1 | tan | 0.106899092 | 0.595623258 |
| SCIMP | tan | 0.1132524 | 0.573812507 |
| MCMBP | tan | 0.117234021 | 0.560327582 |
| DDB1 | tan | 0.125290191 | 0.533493122 |
| DMXL2 | tan | 0.127668771 | 0.525688659 |
| DHODH | tan | 0.13029938 | 0.517121583 |
| ZC3HAV1 | tan | 0.130692092 | 0.515848491 |
| IFITM10 | tan | 0.135315345 | 0.500976509 |
| ZNF341 | tan | 0.135879989 | 0.499174911 |
| RNF111 | tan | 0.143373172 | 0.475576074 |
| LAPTM5 | tan | 0.155656965 | 0.438169129 |
| PLA2G6 | tan | 0.156839148 | 0.434655066 |
| LZTR1 | tan | 0.159087518 | 0.42801409 |
| FAM193B | tan | 0.1599927 | 0.425356209 |
| PQLC2 | tan | 0.162907571 | 0.416859028 |
| DOPEY1 | tan | 0.165844409 | 0.408393686 |
| NECAP1 | tan | 0.167522072 | 0.403601343 |
| NIPAL3 | tan | 0.168289896 | 0.401418598 |
| PIGU | tan | 0.168576817 | 0.400604659 |
| DPY19L3 | tan | 0.17579111 | 0.380446792 |
| CCDC144A | tan | 0.17630124 | 0.379043938 |
| LST1 | tan | 0.178134853 | 0.374026237 |
| GGCX | tan | 0.179445699 | 0.370462858 |
| ELL | tan | 0.181973094 | 0.363648547 |
| C2orf47 | tan | 0.182171128 | 0.363117739 |
| CCDC15 | tan | 0.184409763 | 0.357149044 |
| ACTA2 | tan | 0.189313043 | 0.344280033 |
| BAZ2A | tan | 0.189628946 | 0.343460569 |
| GALNT7 | tan | 0.19780469 | 0.322660593 |
| CRTAP | tan | 0.201963324 | 0.312383307 |
| CEBPD | turquoise | -0.154222 | 0.451916 |
| AQR | turquoise | -0.15342 | 0.454302 |
| HARS2 | turquoise | -0.152624 | 0.456673 |
| HSD17B14 | turquoise | -0.152545 | 0.45691 |
| LMBRD1 | turquoise | -0.152424 | 0.457269 |
| BCL2A1 | turquoise | -0.152239 | 0.457823 |
| NQO1 | turquoise | -0.152118 | 0.458186 |
| GLE1 | turquoise | -0.151922 | 0.458773 |
| NLE1 | turquoise | -0.151831 | 0.459044 |
| ARPC5L | turquoise | -0.151763 | 0.459246 |
| CUL5 | turquoise | -0.151665 | 0.45954 |
| DNASE1 | turquoise | -0.150383 | 0.463387 |
| TXLNG | turquoise | -0.150147 | 0.464098 |
| DNAJC8 | turquoise | -0.150037 | 0.46443 |
| TNFSF13B | turquoise | -0.149958 | 0.464666 |
| XBP1 | turquoise | -0.14939 | 0.466379 |
| STRAP | turquoise | -0.149052 | 0.467399 |
| KRCC1 | turquoise | -0.148691 | 0.468491 |
| PCIF1 | turquoise | -0.148452 | 0.469216 |
| IL10RB | turquoise | -0.14833 | 0.469584 |
| CTU2 | turquoise | -0.148294 | 0.469692 |
| EMC2 | turquoise | -0.148166 | 0.470081 |
| RPL34 | turquoise | -0.147777 | 0.471261 |
| SRCIN1 | turquoise | -0.147064 | 0.473426 |
| RNF14 | turquoise | -0.147019 | 0.473564 |
| TIPARP | turquoise | -0.146943 | 0.473796 |
| SNORA14B | turquoise | -0.146813 | 0.474191 |
| PAK1IP1 | turquoise | -0.146657 | 0.474666 |
| PRDM4 | turquoise | -0.146563 | 0.474952 |
| DIS3 | turquoise | -0.146526 | 0.475066 |
| FBXO5 | turquoise | -0.145909 | 0.476945 |
| B3GALTL | turquoise | -0.145689 | 0.477619 |
| BOLA3 | turquoise | -0.145189 | 0.479146 |
| RNF115 | turquoise | -0.144968 | 0.479823 |
| PSMA6 | turquoise | -0.144645 | 0.480813 |
| COMMD8 | turquoise | -0.144578 | 0.481017 |
| BTF3 | turquoise | -0.144379 | 0.481627 |
| ALG2 | turquoise | -0.144151 | 0.482329 |
| DNAJC2 | turquoise | -0.143779 | 0.483472 |
| DANCR | turquoise | -0.143725 | 0.483639 |
| C6orf62 | turquoise | -0.143668 | 0.483812 |
| B4GALT2 | turquoise | -0.143472 | 0.484416 |
| ARHGEF5 | turquoise | -0.143463 | 0.484442 |
| MUM1 | turquoise | -0.142894 | 0.486195 |
| GRIPAP1 | turquoise | -0.142891 | 0.486204 |
| ZNF329 | turquoise | -0.142634 | 0.486999 |
| C5orf51 | turquoise | -0.142577 | 0.487175 |
| FAM92A1 | turquoise | -0.14248 | 0.487475 |
| GABARAPL2 | turquoise | -0.141967 | 0.489058 |
| TRPM7 | turquoise | -0.141921 | 0.489199 |
| TATDN3 | turquoise | -0.141901 | 0.489262 |
| ZC3H8 | turquoise | -0.141689 | 0.489918 |
| ZCCHC16 | turquoise | -0.140996 | 0.492067 |
| DCAF13 | turquoise | -0.140851 | 0.492515 |
| TSNAX | turquoise | -0.140432 | 0.493816 |
| GALNS | turquoise | -0.140379 | 0.493981 |

**Table 6. Gene module membership and correlation with growth rate (48/72h), with adjusted p values for each module**

| | | | | | | | | | | | |
|---|---|---|---|---|---|---|---|---|---|---|---|
| DDX41 | turquoise | -0.485888 | 0.011851 | ASPHD2 | tan | 0.202541177 | 0.310971453 | C11orf80 | turquoise | -0.139909 | 0.495444 |
| CRCP | turquoise | 0.465423 | 0.016574 | ARCN1 | tan | 0.204898286 | 0.305253401 | MGLL | turquoise | -0.139344 | 0.497203 |
| PARP2 | turquoise | -0.454958 | 0.019529 | ABI1 | tan | 0.209028729 | 0.295392455 | ELAC1 | turquoise | -0.139306 | 0.497321 |
| FLVCR1 | turquoise | -0.445277 | 0.022634 | C12orf29 | tan | 0.209832902 | 0.293496151 | MRE11A | turquoise | -0.139222 | 0.497584 |
| ARRB1 | turquoise | -0.434541 | 0.026535 | PIGT | tan | 0.21104752 | 0.29064655 | GFM1 | turquoise | -0.139046 | 0.498132 |
| PVALB | turquoise | -0.427995 | 0.029168 | TMEM19 | tan | 0.21266176 | 0.28688653 | CDKN2AIP | turquoise | -0.13896 | 0.498401 |
| AGAP2 | turquoise | -0.422067 | 0.03173 | SSRP1 | tan | 0.214672672 | 0.282245863 | DYNLL2 | turquoise | -0.138546 | 0.499694 |
| ZSCAN18 | turquoise | -0.421811 | 0.031845 | WIPI1 | tan | 0.227576832 | 0.253609023 | CDKN2AIPN | turquoise | -0.138477 | 0.49991 |
| IFT57 | turquoise | -0.421678 | 0.031905 | MEN1 | tan | 0.228259666 | 0.252148664 | FAM177A1 | turquoise | -0.138263 | 0.50058 |
| KBTBD4 | turquoise | -0.419975 | 0.032676 | DPM2 | tan | 0.238547404 | 0.230812027 | USP40 | turquoise | -0.137272 | 0.503684 |
| CIAPIN1 | turquoise | -0.419289 | 0.032992 | EIF2AK2 | tan | 0.253999236 | 0.201088624 | DTD2 | turquoise | -0.137233 | 0.503806 |
| P2RY12 | turquoise | -0.418791 | 0.033222 | SIDT2 | tan | 0.256478346 | 0.196576152 | PDCD11 | turquoise | -0.136904 | 0.50484 |
| MSC | turquoise | -0.407307 | 0.038899 | SFT2D2 | tan | 0.265818828 | 0.180201307 | COPS4 | turquoise | -0.1365 | 0.50611 |
| TIMM13 | turquoise | -0.399348 | 0.043268 | ESYT1 | tan | 0.273216436 | 0.167925986 | TDP1 | turquoise | -0.136413 | 0.506382 |
| POLA1 | turquoise | -0.398363 | 0.043834 | CAPRIN2 | tan | 0.283101703 | 0.152459419 | ACSL3 | turquoise | -0.136393 | 0.506447 |
| ZNF639 | turquoise | -0.395302 | 0.045633 | SMAD4 | tan | 0.283839663 | 0.151347097 | PIGF | turquoise | -0.136085 | 0.507414 |
| SGPP1 | turquoise | 0.393908 | 0.04647 | EIF4G1 | tan | 0.285144346 | 0.149394773 | AMACR | turquoise | -0.13599 | 0.507716 |
| HTATSF1 | turquoise | -0.387827 | 0.050268 | PRPF31 | tan | 0.286876231 | 0.146831133 | YAE1D1 | turquoise | -0.135916 | 0.507948 |
| PUS10 | turquoise | -0.386076 | 0.051406 | ASCC3 | tan | 0.293585051 | 0.137198248 | RBBP6 | turquoise | -0.135875 | 0.508076 |
| LYPLAL1 | turquoise | -0.383401 | 0.053182 | MARS | tan | 0.29917969 | 0.129521842 | C9orf72 | turquoise | -0.135642 | 0.508812 |
| PRKCD | turquoise | -0.379601 | 0.055788 | C1RL | tan | 0.302556348 | 0.125042917 | PIEZO1 | turquoise | -0.135569 | 0.509041 |
| DHX58 | turquoise | -0.378939 | 0.056252 | ZKSCAN1 | tan | 0.305573389 | 0.121137786 | TMPO | turquoise | -0.135136 | 0.510408 |
| SLA | turquoise | -0.377541 | 0.057241 | MTMR4 | tan | 0.308890156 | 0.116948832 | RANBP10 | turquoise | -0.135106 | 0.510503 |
| KIAA0101 | turquoise | -0.376846 | 0.057738 | SLC35C2 | tan | 0.311564229 | 0.113650066 | LAMB2 | turquoise | -0.135098 | 0.510529 |
| DPAGT1 | turquoise | -0.376548 | 0.057952 | CLTC | tan | 0.312080245 | 0.113021499 | TANK | turquoise | -0.134472 | 0.512507 |
| ABHD11 | turquoise | -0.376354 | 0.058092 | PPAP2B | tan | 0.312488368 | 0.112526183 | PUS7 | turquoise | -0.134467 | 0.512523 |
| ORAI2 | turquoise | -0.374914 | 0.059137 | TAB1 | tan | 0.3138588 | 0.110874707 | STIM2 | turquoise | -0.134335 | 0.512942 |
| C6orf162 | turquoise | -0.374443 | 0.059482 | UBTF | tan | 0.313917947 | 0.110803837 | CCDC7 | turquoise | -0.133768 | 0.514739 |
| HSPA1A | turquoise | 0.37444 | 0.059485 | G6PC3 | tan | 0.323413793 | 0.099855434 | FAM175A | turquoise | -0.133752 | 0.514788 |
| CEP290 | turquoise | -0.373215 | 0.060389 | SLC29A1 | tan | 0.323905357 | 0.099311586 | TSEN15 | turquoise | -0.133568 | 0.515373 |
| TTC13 | turquoise | -0.370223 | 0.062644 | DCAF5 | tan | 0.326612302 | 0.096356374 | VPS54 | turquoise | -0.133264 | 0.516337 |
| BTN3A1 | turquoise | -0.370125 | 0.062719 | LRRC25 | tan | 0.328440344 | 0.094398347 | EPB41L4A | turquoise | -0.132847 | 0.517663 |
| SSH3 | turquoise | -0.369521 | 0.063182 | RPA1 | tan | 0.333950132 | 0.088677655 | C20orf194 | turquoise | -0.132722 | 0.518062 |
| LDB1 | turquoise | -0.368948 | 0.063624 | RNF31 | tan | 0.334827136 | 0.087791814 | ZNF383 | turquoise | -0.132565 | 0.518561 |
| NMB | turquoise | -0.367229 | 0.064965 | MED18 | tan | 0.338600712 | 0.084056431 | NOC3L | turquoise | -0.13253 | 0.518672 |
| UBE2L6 | turquoise | -0.365982 | 0.065951 | MPI | tan | 0.341645814 | 0.081131169 | MFN1 | turquoise | -0.132444 | 0.518945 |
| ZMYND19 | turquoise | -0.364572 | 0.06708 | ZNF658 | tan | 0.344565562 | 0.078399763 | ZNHIT3 | turquoise | -0.132158 | 0.519856 |
| NMI | turquoise | -0.358757 | 0.071893 | NAGA | tan | 0.347119218 | 0.076068896 | UTP23 | turquoise | -0.132138 | 0.51992 |
| LRPAP1 | turquoise | -0.358419 | 0.07218 | PRR3 | tan | 0.348223327 | 0.075077695 | HELB | turquoise | -0.131399 | 0.522277 |
| ABHD14A | turquoise | -0.357891 | 0.072632 | PIAS3 | tan | 0.349101981 | 0.074295988 | IPO4 | turquoise | -0.131189 | 0.522948 |
| NIPSNAP1 | turquoise | -0.357103 | 0.073309 | TPRG1 | tan | 0.350683618 | 0.072904607 | C1orf131 | turquoise | -0.131186 | 0.522957 |
| HFE | turquoise | -0.356161 | 0.074125 | DPAGT1 | tan | 0.354068319 | 0.069994332 | PTPLAD2 | turquoise | -0.131075 | 0.523314 |
| NDRG3 | turquoise | -0.351811 | 0.077984 | PPP1R21 | tan | 0.354731542 | 0.069434705 | SLC27A1 | turquoise | -0.130592 | 0.524857 |
| MRPS28 | turquoise | -0.35136 | 0.078392 | IDH3G | tan | 0.355627688 | 0.068684021 | WBP4 | turquoise | -0.129934 | 0.526968 |
| NHLRC2 | turquoise | -0.351086 | 0.078641 | C11orf49 | tan | 0.356379375 | 0.068059186 | C1orf63 | turquoise | -0.129461 | 0.528484 |
| MRPL1 | turquoise | -0.349801 | 0.079817 | FAM208A | tan | 0.365667019 | 0.060695511 | PWP2 | turquoise | -0.129407 | 0.52866 |
| UQCRB | turquoise | -0.348852 | 0.080695 | RAB2B | tan | 0.378345666 | 0.051660968 | TMEM70 | turquoise | -0.129258 | 0.529137 |
| ZWILCH | turquoise | -0.348003 | 0.081485 | RTF1 | tan | 0.378931827 | 0.051270375 | ARL2BP | turquoise | -0.129161 | 0.52945 |
| SEC16B | turquoise | -0.347659 | 0.081808 | HDLBP | tan | 0.380611469 | 0.050163967 | RAB28 | turquoise | -0.129155 | 0.529469 |
| PLAC8 | turquoise | -0.347495 | 0.081961 | KPNA6 | tan | 0.390559059 | 0.043990825 | NAA20 | turquoise | -0.129013 | 0.529925 |
| RPL13A | turquoise | -0.347076 | 0.082356 | NABP2 | tan | 0.399989135 | 0.038709799 | CHTOP | turquoise | -0.128895 | 0.530306 |
| FAM168A | turquoise | -0.345678 | 0.083681 | KAT7 | tan | 0.40023356 | 0.038579962 | ZNF513 | turquoise | -0.128875 | 0.53037 |
| NAT1 | turquoise | -0.344248 | 0.085054 | YIF1B | tan | 0.413695501 | 0.031947938 | CYP20A1 | turquoise | -0.12881 | 0.530579 |
| UBE2W | turquoise | -0.342971 | 0.086294 | TRMT10B | tan | 0.423219176 | 0.027835433 | ZNF398 | turquoise | -0.128434 | 0.531788 |
| ZNF688 | turquoise | -0.342089 | 0.087159 | MIIP | tan | 0.441721172 | 0.021069746 | SMG8 | turquoise | -0.128222 | 0.532473 |
| TM7SF3 | turquoise | -0.340296 | 0.088937 | VAMP8 | tan | 0.443873584 | 0.020378686 | HSCB | turquoise | -0.127957 | 0.533325 |

**Table 6. Gene module membership and correlation with growth rate (48/72h), with adjusted p values for each module**

| | | | | | | | | | | | |
|---|---|---|---|---|---|---|---|---|---|---|---|
| KIF3B | turquoise | -0.33666 | 0.092625 | CDS2 | tan | 0.447978904 | 0.019112026 | EPC1 | turquoise | -0.127945 | 0.533366 |
| NIF3L1 | turquoise | -0.336541 | 0.092749 | SLC5A6 | tan | 0.47197066 | 0.012935144 | SFR1 | turquoise | -0.127836 | 0.533717 |
| MRPL54 | turquoise | -0.335452 | 0.093876 | COG1 | tan | 0.482672848 | 0.010772321 | HMBOX1 | turquoise | -0.127755 | 0.53398 |
| ATG5 | turquoise | -0.334644 | 0.094719 | TMEM106C | tan | 0.527729162 | 0.00466899 | SCYL2 | turquoise | -0.127719 | 0.534095 |
| CDK1 | turquoise | -0.333663 | 0.095751 | UNG | tan | 0.563346068 | 0.002215598 | POLR3B | turquoise | -0.127638 | 0.534358 |
| FAM53B | turquoise | -0.332422 | 0.097068 | ARRDC4 | turquoise | -0.575510033 | 0.001684686 | DNAJA4 | turquoise | -0.127308 | 0.535421 |
| TAZ | turquoise | -0.332402 | 0.09709 | HELQ | turquoise | -0.552747816 | 0.002789557 | FAM219B | turquoise | -0.127079 | 0.536163 |
| ATP5O | turquoise | -0.331955 | 0.097567 | DDX41 | turquoise | -0.544941386 | 0.003289769 | TOMM70A | turquoise | -0.126687 | 0.537429 |
| GPR133 | turquoise | 0.331554 | 0.097997 | MTX3 | turquoise | -0.531331633 | 0.00434558 | SLC17A9 | turquoise | -0.126598 | 0.537718 |
| SPI1 | turquoise | -0.330098 | 0.099571 | ZNF131 | turquoise | -0.525614609 | 0.004868171 | HAUS2 | turquoise | -0.126584 | 0.537765 |
| UFSP2 | turquoise | -0.329526 | 0.100195 | HAUS8 | turquoise | -0.51713543 | 0.005740875 | LINC00493 | turquoise | -0.126303 | 0.538675 |
| PRR11 | turquoise | -0.328965 | 0.100809 | LOC731275 | turquoise | -0.515558523 | 0.005916953 | PRKAA1 | turquoise | -0.126239 | 0.538881 |
| SNRPB | turquoise | -0.328514 | 0.101305 | GTPBP8 | turquoise | -0.504844886 | 0.007238305 | EFHA1 | turquoise | -0.126008 | 0.539631 |
| ACTR6 | turquoise | -0.327689 | 0.102217 | CCNH | turquoise | -0.502344164 | 0.00758011 | C17orf70 | turquoise | -0.125987 | 0.5397 |
| PCNA | turquoise | -0.327451 | 0.102481 | RANBP2 | turquoise | -0.4925132 | 0.009058313 | DDX47 | turquoise | -0.125863 | 0.540102 |
| CHCHD4 | turquoise | -0.323463 | 0.106983 | WIBG | turquoise | -0.491641592 | 0.009200273 | SLCO2B1 | turquoise | -0.125579 | 0.541024 |
| MRPL16 | turquoise | -0.322548 | 0.108037 | PLEKHG2 | turquoise | -0.483338949 | 0.010648345 | GPCPD1 | turquoise | -0.125322 | 0.541858 |
| CMC4 | turquoise | -0.321865 | 0.108828 | WWP1 | turquoise | -0.475486826 | 0.012188034 | ZFP106 | turquoise | -0.124637 | 0.544089 |
| NMNAT1 | turquoise | -0.321576 | 0.109164 | C1orf198 | turquoise | -0.471829048 | 0.012966011 | AHR | turquoise | -0.124551 | 0.544368 |
| SCARNA16 | turquoise | -0.320107 | 0.110884 | PPP1CC | turquoise | -0.470847854 | 0.01318157 | NR2C1 | turquoise | -0.124056 | 0.54598 |
| TMEM42 | turquoise | -0.318366 | 0.112949 | RINT1 | turquoise | -0.469474239 | 0.013488323 | RHOBTB3 | turquoise | -0.124037 | 0.546043 |
| LACTB | turquoise | -0.313477 | 0.118897 | TMEM106B | turquoise | -0.464788007 | 0.014579702 | ID2 | turquoise | -0.124004 | 0.546152 |
| SOCS3 | turquoise | -0.312888 | 0.119629 | SOS2 | turquoise | -0.464274906 | 0.01470351 | C20orf94 | turquoise | -0.123993 | 0.546186 |
| XPNPEP1 | turquoise | -0.31271 | 0.119851 | ITGB8 | turquoise | -0.459780481 | 0.015825583 | TRIM44 | turquoise | -0.123801 | 0.546812 |
| ERI2 | turquoise | -0.312338 | 0.120316 | BCL7B | turquoise | -0.459528595 | 0.015890502 | CORO1C | turquoise | -0.123677 | 0.547217 |
| ITFG3 | turquoise | -0.312219 | 0.120465 | VPS37C | turquoise | -0.457360931 | 0.016458305 | SPATA7 | turquoise | -0.123671 | 0.547238 |
| GCC2 | turquoise | -0.311 | 0.121998 | FAM53B | turquoise | -0.453176286 | 0.017601742 | DENR | turquoise | -0.123533 | 0.547689 |
| RNPEP | turquoise | -0.310393 | 0.122767 | PEX19 | turquoise | -0.452518993 | 0.01778714 | IDNK | turquoise | -0.123108 | 0.549078 |
| STK36 | turquoise | -0.310225 | 0.12298 | PIK3C2A | turquoise | -0.451923311 | 0.017956543 | ZNF569 | turquoise | -0.122651 | 0.550571 |
| XAF1 | turquoise | -0.309461 | 0.123954 | IL13RA1 | turquoise | -0.442634607 | 0.020774173 | VOPP1 | turquoise | -0.12248 | 0.551132 |
| GALNS | turquoise | -0.308297 | 0.125449 | GORASP2 | turquoise | -0.441774821 | 0.021052292 | AP3D1 | turquoise | -0.122451 | 0.551229 |
| ZNF823 | turquoise | -0.308169 | 0.125614 | SUCLA2 | turquoise | -0.440169318 | 0.021579776 | XRN1 | turquoise | -0.122437 | 0.551275 |
| CCDC86 | turquoise | -0.307414 | 0.126591 | XPO4 | turquoise | -0.437780596 | 0.022384512 | CISD2 | turquoise | -0.122046 | 0.552554 |
| SMC4 | turquoise | -0.307297 | 0.126743 | MSI2 | turquoise | -0.433526705 | 0.023878126 | COPB1 | turquoise | -0.12192 | 0.552968 |
| UBE3C | turquoise | -0.307215 | 0.126849 | RXRA | turquoise | -0.430386463 | 0.025031929 | GLYR1 | turquoise | -0.121893 | 0.553056 |
| ANP32E | turquoise | -0.305977 | 0.128466 | NOP2 | turquoise | -0.425271123 | 0.027008064 | PTEN | turquoise | -0.121692 | 0.553717 |
| LGALS8 | turquoise | -0.305473 | 0.129128 | DUT | turquoise | -0.421175148 | 0.028679869 | DAP3 | turquoise | -0.121606 | 0.554 |
| GPR160 | turquoise | -0.304508 | 0.130404 | ARPC3 | turquoise | -0.421003983 | 0.028751509 | ADI1 | turquoise | -0.121571 | 0.554114 |
| DNAJC1 | turquoise | -0.304401 | 0.130546 | DR1 | turquoise | -0.420422836 | 0.028995827 | SNX13 | turquoise | -0.121312 | 0.554964 |
| GHDC | turquoise | 0.303777 | 0.131377 | MMADHC | turquoise | -0.416775067 | 0.030567914 | DEK | turquoise | -0.121005 | 0.555974 |
| HAT1 | turquoise | -0.302666 | 0.132863 | CDC73 | turquoise | -0.416733397 | 0.030586261 | SYF2 | turquoise | -0.120936 | 0.556202 |
| ANKRD26 | turquoise | -0.302447 | 0.133159 | METTL2A | turquoise | -0.416379794 | 0.03074231 | IGBP1 | turquoise | -0.120874 | 0.556405 |
| GSTM2 | turquoise | -0.301381 | 0.1346 | ARNTL2 | turquoise | -0.415261838 | 0.031239898 | MCTS1 | turquoise | -0.120778 | 0.556719 |
| RAB11FIP4 | turquoise | -0.301175 | 0.134879 | ZMIZ2 | turquoise | -0.414793078 | 0.031450457 | RNF213 | turquoise | -0.120768 | 0.556754 |
| ZNF436 | turquoise | -0.300606 | 0.135654 | MT1M | turquoise | -0.414565251 | 0.031553204 | MDM2 | turquoise | -0.120764 | 0.556767 |
| GSTK1 | turquoise | -0.297234 | 0.140312 | MS4A14 | turquoise | -0.414213594 | 0.031712328 | POLR2B | turquoise | -0.120571 | 0.557401 |
| STK4 | turquoise | -0.297011 | 0.140625 | MAN2C1 | turquoise | -0.410862207 | 0.033261433 | CEBPZ | turquoise | -0.119893 | 0.559637 |
| DHFR | turquoise | -0.296893 | 0.140789 | PWWP2A | turquoise | -0.407088663 | 0.035077758 | C15orf37 | turquoise | -0.119634 | 0.560491 |
| TMEM205 | turquoise | -0.296825 | 0.140886 | SREK1 | turquoise | -0.405759502 | 0.035736118 | BLMH | turquoise | -0.119334 | 0.561484 |
| SCYL3 | turquoise | -0.296629 | 0.14116 | SRF | turquoise | -0.400914625 | 0.038220018 | FER | turquoise | -0.11921 | 0.561893 |
| C14orf101 | turquoise | -0.296066 | 0.141953 | LBX2-AS1 | turquoise | -0.400576336 | 0.038398468 | ACTR10 | turquoise | -0.119031 | 0.562482 |
| HMBS | turquoise | -0.295761 | 0.142383 | LPP | turquoise | -0.400234496 | 0.038579466 | GINM1 | turquoise | -0.118927 | 0.562827 |
| PAAF1 | turquoise | -0.295458 | 0.142812 | LRRC8B | turquoise | -0.399106003 | 0.039181815 | EIF5 | turquoise | -0.11885 | 0.563082 |
| ITGB2 | turquoise | -0.294949 | 0.143535 | PTPN1 | turquoise | -0.396071272 | 0.040838866 | RMI1 | turquoise | -0.118728 | 0.563486 |
| BCKDHA | turquoise | -0.294702 | 0.143885 | NOL11 | turquoise | -0.395980224 | 0.040889428 | SNORA75 | turquoise | -0.118695 | 0.563595 |
| CAAP1 | turquoise | -0.293544 | 0.145541 | TMEM131 | turquoise | -0.395008096 | 0.041432392 | SCFD1 | turquoise | -0.118582 | 0.563969 |

**Table 6. Gene module membership and correlation with growth rate (48/72h), with adjusted p values for each module**

| | | | | | | | | | | | |
|---|---|---|---|---|---|---|---|---|---|---|---|
| SCARNA9 | turquoise | 0.292201 | 0.147478 | CD302 | turquoise | -0.392827027 | 0.042671433 | S1PR4 | turquoise | -0.118489 | 0.564277 |
| OC10050645 | turquoise | 0.292085 | 0.147647 | FLVCR1 | turquoise | -0.392415169 | 0.042908669 | TMEM180 | turquoise | -0.118478 | 0.564313 |
| OC10019098 | turquoise | 0.291659 | 0.148266 | EXOC2 | turquoise | -0.391883946 | 0.043216205 | C3orf14 | turquoise | -0.118367 | 0.56468 |
| C4orf29 | turquoise | -0.290974 | 0.149265 | ME2 | turquoise | -0.391658504 | 0.043347245 | AGPAT5 | turquoise | -0.11835 | 0.564736 |
| C15orf39 | turquoise | -0.290801 | 0.149517 | FOCAD | turquoise | -0.391624731 | 0.043366903 | ZFYVE28 | turquoise | -0.118323 | 0.564827 |
| NAP1L4 | turquoise | -0.290146 | 0.150479 | ZNF570 | turquoise | -0.390595274 | 0.043969506 | C5orf28 | turquoise | -0.118293 | 0.564924 |
| SEPT7P2 | turquoise | -0.290111 | 0.15053 | CDC37L1 | turquoise | -0.388563096 | 0.045178519 | ZNF382 | turquoise | -0.118129 | 0.565469 |
| SMG1P1 | turquoise | 0.289643 | 0.15122 | KLF2 | turquoise | -0.386408988 | 0.046488621 | DTNA | turquoise | -0.118001 | 0.565894 |
| SPATA20 | turquoise | -0.289114 | 0.152001 | TMEM194B | turquoise | -0.385204829 | 0.047233939 | TARBP1 | turquoise | -0.117591 | 0.567253 |
| SLC29A3 | turquoise | -0.28869 | 0.152631 | GNA11 | turquoise | -0.385140403 | 0.04727408 | MBP | turquoise | -0.117483 | 0.567611 |
| TOP2A | turquoise | -0.288171 | 0.153402 | LINC00674 | turquoise | -0.38495809 | 0.047387816 | SETX | turquoise | -0.117271 | 0.568317 |
| GPATCH8 | turquoise | -0.286011 | 0.156646 | NFIC | turquoise | -0.384906844 | 0.047419826 | UBE3B | turquoise | -0.117229 | 0.568454 |
| AMZ2P1 | turquoise | -0.285863 | 0.156871 | RAP1GDS1 | turquoise | -0.382617874 | 0.04886702 | CSRNP1 | turquoise | -0.117122 | 0.568813 |
| VEGFA | turquoise | -0.285129 | 0.157984 | MMP12 | turquoise | -0.381426934 | 0.049633622 | HBS1L | turquoise | -0.117106 | 0.568864 |
| C15orf40 | turquoise | -0.284032 | 0.15966 | POLM | turquoise | -0.381100478 | 0.049845404 | ZNF227 | turquoise | -0.11695 | 0.569382 |
| RWDD3 | turquoise | -0.283955 | 0.15978 | CHERP | turquoise | -0.380835108 | 0.05001808 | TMA16 | turquoise | -0.116829 | 0.569785 |
| SFPQ | turquoise | 0.283554 | 0.160395 | SH2B1 | turquoise | -0.379914575 | 0.050620721 | AKT1S1 | turquoise | -0.116477 | 0.570957 |
| FXN | turquoise | -0.283239 | 0.16088 | AMN1 | turquoise | -0.379535089 | 0.050870812 | UBE4B | turquoise | -0.116356 | 0.57136 |
| HAUS2 | turquoise | -0.283222 | 0.160906 | KATNAL1 | turquoise | -0.378368094 | 0.05164598 | SEPT7P2 | turquoise | -0.116337 | 0.571424 |
| RPPH1 | turquoise | -0.280985 | 0.164383 | ST13 | turquoise | -0.378057884 | 0.051853586 | AIDA | turquoise | -0.11607 | 0.572313 |
| FBXO48 | turquoise | -0.280544 | 0.165073 | EVI2B | turquoise | -0.376553781 | 0.052869503 | CACYBP | turquoise | -0.115878 | 0.572952 |
| GOLGA1 | turquoise | -0.280513 | 0.165122 | DDX46 | turquoise | -0.375268994 | 0.053749586 | GTF2H3 | turquoise | -0.115764 | 0.573332 |
| TLR7 | turquoise | -0.280352 | 0.165375 | ZDHHC2 | turquoise | -0.373960295 | 0.054657804 | APOBEC3C | turquoise | -0.115488 | 0.574253 |
| MAFK | turquoise | 0.28014 | 0.165708 | SLC9A3R1 | turquoise | -0.372818797 | 0.055459751 | ACAT1 | turquoise | -0.11518 | 0.575282 |
| UTP18 | turquoise | -0.279754 | 0.166316 | PRR12 | turquoise | -0.371059896 | 0.05671342 | SMC2 | turquoise | -0.114704 | 0.576872 |
| PLAC2 | turquoise | -0.279382 | 0.166905 | FAM89B | turquoise | -0.37028313 | 0.057274059 | NAF1 | turquoise | -0.114624 | 0.57714 |
| WDR60 | turquoise | -0.279321 | 0.167002 | ACTR6 | turquoise | -0.369177118 | 0.058079784 | BTN3A1 | turquoise | -0.11432 | 0.578155 |
| SEC23B | turquoise | -0.279226 | 0.167151 | RNF8 | turquoise | -0.368106837 | 0.058867866 | NUP35 | turquoise | -0.114293 | 0.578246 |
| WDR44 | turquoise | -0.279152 | 0.167268 | MYO1G | turquoise | -0.367945151 | 0.058987641 | EAPP | turquoise | -0.114255 | 0.578374 |
| TFAM | turquoise | -0.27904 | 0.167447 | FGFR1OP | turquoise | -0.366769391 | 0.059864348 | WASF2 | turquoise | -0.113873 | 0.579654 |
| ITFG1 | turquoise | -0.278763 | 0.167887 | FASTKD1 | turquoise | -0.364800874 | 0.061354834 | ANKRD26 | turquoise | -0.113528 | 0.580811 |
| COPS7A | turquoise | -0.277434 | 0.170008 | PMS1 | turquoise | -0.363930345 | 0.062023088 | MPST | turquoise | -0.113272 | 0.581668 |
| MAGOHB | turquoise | 0.277302 | 0.170218 | HPS1 | turquoise | -0.363290162 | 0.062518113 | AKAP9 | turquoise | -0.113218 | 0.581849 |
| MGLL | turquoise | -0.276986 | 0.170727 | CLIP4 | turquoise | -0.36317239 | 0.062609513 | ATPAF1 | turquoise | -0.113175 | 0.581995 |
| C11orf80 | turquoise | -0.276849 | 0.170947 | MOB3A | turquoise | -0.362593261 | 0.063060474 | FCF1 | turquoise | -0.112992 | 0.582608 |
| TUBGCP5 | turquoise | -0.27591 | 0.172463 | PPP2R5A | turquoise | -0.36239946 | 0.063211946 | TMEM86A | turquoise | -0.112808 | 0.583226 |
| NLN | turquoise | -0.275633 | 0.172911 | GRIPAP1 | turquoise | -0.360487168 | 0.064721731 | ZNF766 | turquoise | -0.112666 | 0.583703 |
| SLC20A2 | turquoise | -0.275618 | 0.172936 | PMEPA1 | turquoise | -0.359142269 | 0.065800162 | OSBPL9 | turquoise | -0.112546 | 0.584107 |
| TMEM126B | turquoise | -0.274957 | 0.17401 | LYPLAL1 | turquoise | -0.355665215 | 0.068652722 | ZNF385A | turquoise | -0.112463 | 0.584387 |
| PANK2 | turquoise | -0.274181 | 0.175277 | MRPL3 | turquoise | -0.355314108 | 0.068945986 | TMEM41B | turquoise | -0.112364 | 0.584721 |
| IGHMBP2 | turquoise | -0.273653 | 0.176144 | TTC21B | turquoise | -0.354997386 | 0.069211356 | NDUFB6 | turquoise | -0.112351 | 0.584764 |
| PIK3R4 | turquoise | -0.273575 | 0.176271 | LYRM4 | turquoise | -0.352193044 | 0.071595489 | SNX14 | turquoise | -0.112315 | 0.584884 |
| RNH1 | turquoise | -0.272823 | 0.177511 | DECR1 | turquoise | -0.349685576 | 0.073780244 | FOXK1 | turquoise | -0.11227 | 0.585038 |
| SNHG16 | turquoise | -0.272509 | 0.17803 | CPSF2 | turquoise | -0.349602992 | 0.07385306 | RNF219 | turquoise | -0.112132 | 0.5855 |
| EXOC6 | turquoise | -0.271696 | 0.179379 | UBE2E3 | turquoise | -0.34909085 | 0.074305853 | GTF2H1 | turquoise | -0.112011 | 0.585909 |
| GPAM | turquoise | -0.27109 | 0.18039 | PAIP2 | turquoise | -0.346318726 | 0.076793769 | PGRMC2 | turquoise | -0.111919 | 0.586219 |
| OC10050699 | turquoise | -0.270111 | 0.18203 | MTERF | turquoise | -0.345997106 | 0.07708649 | PCNA | turquoise | -0.111682 | 0.587015 |
| TUFM | turquoise | -0.269904 | 0.182379 | MDC1 | turquoise | -0.345985974 | 0.077096637 | ZNF121 | turquoise | -0.111619 | 0.587227 |
| AMIGO3 | turquoise | 0.269133 | 0.18368 | ELMO1 | turquoise | -0.344661354 | 0.078311356 | RARG | turquoise | -0.111455 | 0.58778 |
| FBXW8 | turquoise | -0.268932 | 0.18402 | TUBD1 | turquoise | -0.342605025 | 0.080225958 | DHX40 | turquoise | -0.111303 | 0.588293 |
| MED27 | turquoise | -0.267386 | 0.186651 | HDGFRP2 | turquoise | -0.340434429 | 0.082285446 | PHAX | turquoise | -0.111224 | 0.58856 |
| C2orf68 | turquoise | -0.267341 | 0.18673 | BLZF1 | turquoise | -0.340310222 | 0.082404501 | FAM45A | turquoise | -0.111198 | 0.588647 |
| MPHOSPH6 | turquoise | -0.267234 | 0.186912 | UFM1 | turquoise | -0.33872578 | 0.08393473 | TMEM209 | turquoise | -0.110769 | 0.590094 |
| ZNF271 | turquoise | 0.26723 | 0.186919 | DHX37 | turquoise | -0.338229224 | 0.08441871 | AMOTL1 | turquoise | -0.110728 | 0.590235 |
| VPS35 | turquoise | -0.267221 | 0.186935 | GOLGB1 | turquoise | -0.337676933 | 0.084959498 | FURIN | turquoise | -0.110661 | 0.590459 |
| SMC2 | turquoise | -0.266958 | 0.187386 | CDC42BPB | turquoise | -0.337068218 | 0.085558576 | GNPDA2 | turquoise | -0.110619 | 0.590601 |

**Table 6. Gene module membership and correlation with growth rate (48/72h), with adjusted p values for each module**

| | | | |
|---|---|---|---|
| PI4K2B | turquoise | -0.266917 | 0.187455 |
| ING2 | turquoise | -0.266353 | 0.188425 |
| ZCRB1 | turquoise | -0.265988 | 0.189054 |
| NDUFA8 | turquoise | -0.265381 | 0.190103 |
| LOC440944 | turquoise | -0.265011 | 0.190746 |
| CPE | turquoise | -0.264312 | 0.191961 |
| CD2 | turquoise | -0.263714 | 0.193007 |
| FBXO4 | turquoise | -0.263445 | 0.193477 |
| TPCN2 | turquoise | -0.263365 | 0.193618 |
| C1orf131 | turquoise | -0.263165 | 0.19397 |
| RPRD2 | turquoise | -0.262781 | 0.194645 |
| PBXIP1 | turquoise | -0.261415 | 0.197059 |
| IFNAR1 | turquoise | -0.260647 | 0.198426 |
| CDKL1 | turquoise | -0.260455 | 0.198769 |
| ADAP1 | turquoise | -0.259886 | 0.199787 |
| IQCG | turquoise | -0.259379 | 0.200697 |
| LOC550643 | turquoise | -0.258781 | 0.201775 |
| ATPBD4 | turquoise | -0.258722 | 0.201882 |
| POLR2J | turquoise | -0.257999 | 0.20319 |
| TNFSF12 | turquoise | -0.257405 | 0.204269 |
| SLC25A36 | turquoise | -0.257355 | 0.20436 |
| FAM177A1 | turquoise | -0.257051 | 0.204913 |
| SLC7A5 | turquoise | 0.256531 | 0.205863 |
| FAM89B | turquoise | -0.256527 | 0.205871 |
| LLGL1 | turquoise | -0.256277 | 0.206328 |
| UQCRH | turquoise | -0.255524 | 0.207712 |
| TSPAN14 | turquoise | -0.255331 | 0.208067 |
| DDX23 | turquoise | -0.255238 | 0.208238 |
| ANKS1A | turquoise | -0.254886 | 0.208888 |
| NAE1 | turquoise | -0.254878 | 0.208902 |
| CAPZA2 | turquoise | -0.25451 | 0.209583 |
| MSH3 | turquoise | -0.253459 | 0.211536 |
| LAMP2 | turquoise | -0.253182 | 0.212053 |
| FBXO5 | turquoise | 0.253148 | 0.212117 |
| XRN1 | turquoise | -0.25305 | 0.212299 |
| DDX46 | turquoise | -0.252295 | 0.213712 |
| RPRD1A | turquoise | -0.252235 | 0.213825 |
| PGAM5 | turquoise | -0.251959 | 0.214343 |
| GBAS | turquoise | -0.251933 | 0.214393 |
| SNRPC | turquoise | -0.251854 | 0.21454 |
| ABCC5 | turquoise | -0.251833 | 0.21458 |
| GRIPAP1 | turquoise | -0.250886 | 0.216366 |
| MLPH | turquoise | -0.250089 | 0.21788 |
| MTM1 | turquoise | -0.249694 | 0.21863 |
| SEC22A | turquoise | -0.249429 | 0.219135 |
| MPEG1 | turquoise | -0.249187 | 0.219598 |
| RBM34 | turquoise | -0.248742 | 0.22045 |
| FAM118B | turquoise | -0.248547 | 0.220823 |
| APKAPK5-A | turquoise | -0.247969 | 0.221935 |
| SUGP2 | turquoise | -0.247885 | 0.222095 |
| LACC1 | turquoise | -0.247336 | 0.223155 |
| ZNF329 | turquoise | -0.247105 | 0.223602 |
| GNPAT | turquoise | -0.246986 | 0.223832 |
| FLNB | turquoise | 0.246643 | 0.224496 |
| CUL4A | turquoise | -0.24631 | 0.225143 |
| TRPC4AP | turquoise | -0.24623 | 0.225298 |
| PHRF1 | turquoise | -0.336952308 | 0.085673013 |
| ADRBK2 | turquoise | -0.336864888 | 0.085759399 |
| SERPINB1 | turquoise | -0.336630096 | 0.08599174 |
| UBE3C | turquoise | -0.335774718 | 0.086842229 |
| RENBP | turquoise | -0.335262252 | 0.087354807 |
| PCBD2 | turquoise | -0.334258885 | 0.088365024 |
| BTF3L4 | turquoise | -0.334097346 | 0.088528488 |
| ZNF746 | turquoise | -0.333836634 | 0.088792789 |
| TRPC4AP | turquoise | -0.332836405 | 0.089812325 |
| PHKB | turquoise | -0.332259138 | 0.090404744 |
| INF2 | turquoise | -0.331093741 | 0.091609715 |
| DYNLL1 | turquoise | -0.33017697 | 0.092566095 |
| HIST1H4F | turquoise | -0.329839952 | 0.092919558 |
| ZNF35 | turquoise | -0.329711247 | 0.093054812 |
| ZNF394 | turquoise | -0.329543628 | 0.093231181 |
| FAM179B | turquoise | -0.329409438 | 0.093372557 |
| NOL9 | turquoise | -0.328558395 | 0.094272937 |
| LRRC37BP1 | turquoise | -0.32843186 | 0.094407364 |
| ARL2 | turquoise | -0.327788728 | 0.095092834 |
| GTF2F2 | turquoise | -0.326209465 | 0.096791924 |
| ZBTB38 | turquoise | -0.326031709 | 0.096984584 |
| ZUFSP | turquoise | -0.325685624 | 0.097360514 |
| KIAA2026 | turquoise | -0.325482819 | 0.097581314 |
| KIAA1551 | turquoise | -0.325421512 | 0.097648134 |
| PHLDA3 | turquoise | -0.325000349 | 0.098108104 |
| TMEM109 | turquoise | -0.324671513 | 0.098468367 |
| PDZD11 | turquoise | -0.324330452 | 0.098843068 |
| WAS | turquoise | -0.324290906 | 0.098886583 |
| RBM45 | turquoise | -0.323600107 | 0.099649042 |
| HNRPLL | turquoise | -0.323349828 | 0.099926366 |
| LOC728407 | turquoise | -0.323345443 | 0.099931231 |
| LIAS | turquoise | -0.323020876 | 0.100291744 |
| PRR11 | turquoise | -0.32257385 | 0.100789869 |
| WDSUB1 | turquoise | -0.321705484 | 0.10176278 |
| SLC30A5 | turquoise | -0.320678774 | 0.10292213 |
| UTRN | turquoise | -0.320424342 | 0.10321095 |
| USP10 | turquoise | -0.320025063 | 0.103665412 |
| TRIQK | turquoise | -0.319916852 | 0.103788835 |
| MRPL30 | turquoise | -0.319597441 | 0.104153786 |
| ZMPSTE24 | turquoise | -0.318516518 | 0.10539592 |
| C6orf162 | turquoise | -0.318237083 | 0.105718818 |
| TCF25 | turquoise | -0.318017837 | 0.105972679 |
| CKS1B | turquoise | -0.317786583 | 0.106240936 |
| PPIP5K2 | turquoise | -0.314388248 | 0.110241515 |
| P4HB | turquoise | -0.313597455 | 0.111188254 |
| ZNF623 | turquoise | -0.313046151 | 0.11185183 |
| ZNF764 | turquoise | -0.311716915 | 0.113463808 |
| NAA15 | turquoise | -0.311185087 | 0.114113548 |
| MAPK7 | turquoise | -0.310638867 | 0.114783727 |
| VPS26A | turquoise | -0.310098676 | 0.115449362 |
| DNAJC24 | turquoise | -0.309796344 | 0.115823142 |
| CGGBP1 | turquoise | -0.30928433 | 0.116458192 |
| TMEM2 | turquoise | -0.308053255 | 0.117995591 |
| RBBP6 | turquoise | -0.307990509 | 0.118074348 |
| HKR1 | turquoise | -0.307964025 | 0.118107601 |
| VGLL4 | turquoise | -0.307360273 | 0.11886755 |
| DHTKD1 | turquoise | -0.110519 | 0.590939 |
| RBL1 | turquoise | -0.110483 | 0.591062 |
| TRIO | turquoise | -0.110201 | 0.592014 |
| TRABD | turquoise | -0.110137 | 0.592233 |
| RBM8A | turquoise | -0.10903 | 0.595979 |
| POT1 | turquoise | -0.109011 | 0.596045 |
| ZC3H3 | turquoise | -0.1089 | 0.596423 |
| ZDHHC8 | turquoise | -0.108698 | 0.597108 |
| FAM13B | turquoise | -0.10831 | 0.598426 |
| PSMG1 | turquoise | -0.107934 | 0.599703 |
| PFDN1 | turquoise | -0.10792 | 0.599751 |
| GOLGA4 | turquoise | -0.107919 | 0.599755 |
| HAT1 | turquoise | -0.107819 | 0.600092 |
| SPOPL | turquoise | -0.107544 | 0.601031 |
| KRAS | turquoise | -0.107432 | 0.601411 |
| EPS15 | turquoise | -0.107407 | 0.601497 |
| SENP6 | turquoise | -0.107201 | 0.602197 |
| LPCAT2 | turquoise | -0.107071 | 0.60264 |
| SHOC2 | turquoise | -0.10677 | 0.603667 |
| CEP72 | turquoise | -0.106764 | 0.603688 |
| PAXIP1 | turquoise | -0.106574 | 0.604336 |
| HYOU1 | turquoise | -0.106222 | 0.605535 |
| DMTF1 | turquoise | -0.106159 | 0.605752 |
| SKAP2 | turquoise | -0.106146 | 0.605797 |
| SRPK2 | turquoise | -0.105981 | 0.60636 |
| SMIM4 | turquoise | -0.105621 | 0.60759 |
| METTL6 | turquoise | -0.105586 | 0.607711 |
| TTC19 | turquoise | -0.105554 | 0.607818 |
| GOLGA1 | turquoise | -0.105554 | 0.607821 |
| NUP107 | turquoise | -0.105168 | 0.609141 |
| CWF19L2 | turquoise | -0.105155 | 0.609184 |
| BSDC1 | turquoise | -0.104714 | 0.610694 |
| MGAT2 | turquoise | -0.104696 | 0.610757 |
| DYRK1A | turquoise | -0.104649 | 0.610918 |
| GNA11 | turquoise | -0.104603 | 0.611077 |
| HELZ | turquoise | -0.104484 | 0.611481 |
| POR | turquoise | -0.104201 | 0.612454 |
| KDELR3 | turquoise | -0.103854 | 0.613645 |
| TSR1 | turquoise | -0.10369 | 0.614209 |
| ARSB | turquoise | -0.103288 | 0.61559 |
| TAF12 | turquoise | -0.102947 | 0.616762 |
| ANAPC4 | turquoise | -0.102883 | 0.616981 |
| TADA2B | turquoise | -0.102609 | 0.617924 |
| C5orf15 | turquoise | -0.102527 | 0.618208 |
| HSPB1 | turquoise | -0.102477 | 0.618381 |
| YIPF2 | turquoise | -0.102112 | 0.619639 |
| CBX3 | turquoise | -0.102037 | 0.619895 |
| RNF167 | turquoise | -0.101616 | 0.621349 |
| ME3 | turquoise | -0.101496 | 0.621761 |
| HSP90B1 | turquoise | -0.101454 | 0.621907 |
| PRRC1 | turquoise | -0.101313 | 0.622392 |
| TMEM179E | turquoise | -0.101061 | 0.623265 |
| CAPN7 | turquoise | -0.100729 | 0.624411 |
| ADAP2 | turquoise | -0.100636 | 0.624734 |
| CCDC91 | turquoise | -0.1005 | 0.625204 |
| KIAA0528 | turquoise | -0.100094 | 0.62661 |

**Table 6. Gene module membership and correlation with growth rate (48/72h), with adjusted p values for each module**

| | | | |
|---|---|---|---|
| TMEM164 | turquoise | 0.246078 | 0.225594 |
| MYC | turquoise | -0.24571 | 0.22631 |
| ACAA1 | turquoise | -0.244667 | 0.228351 |
| DND1 | turquoise | -0.244389 | 0.228896 |
| METTL2A | turquoise | -0.244297 | 0.229077 |
| MYCBP | turquoise | -0.244115 | 0.229435 |
| USMG5 | turquoise | -0.243782 | 0.23009 |
| C3orf14 | turquoise | -0.243714 | 0.230226 |
| CNTLN | turquoise | -0.243596 | 0.230458 |
| BRIX1 | turquoise | -0.24352 | 0.230607 |
| GLMN | turquoise | -0.243144 | 0.23135 |
| HDAC2 | turquoise | -0.243055 | 0.231527 |
| FNBP4 | turquoise | -0.24249 | 0.232647 |
| MGMT | turquoise | -0.242334 | 0.232957 |
| MAZ | turquoise | -0.241952 | 0.233717 |
| EIF3M | turquoise | -0.241596 | 0.234428 |
| HSCB | turquoise | -0.241353 | 0.234913 |
| SUMF1 | turquoise | -0.240791 | 0.236039 |
| MBIP | turquoise | -0.240565 | 0.236492 |
| CPSF2 | turquoise | -0.240305 | 0.237015 |
| KXD1 | turquoise | 0.240105 | 0.237417 |
| ATPAF1 | turquoise | -0.239874 | 0.237881 |
| IGBP1 | turquoise | -0.239781 | 0.23807 |
| C10orf54 | turquoise | -0.239083 | 0.239481 |
| CWC15 | turquoise | -0.23876 | 0.240135 |
| PIK3C3 | turquoise | -0.23801 | 0.241658 |
| ZNF385A | turquoise | -0.23791 | 0.241862 |
| FAM45A | turquoise | -0.237821 | 0.242044 |
| SLK | turquoise | -0.237476 | 0.242748 |
| VAMP7 | turquoise | -0.237246 | 0.243218 |
| CCP110 | turquoise | -0.23709 | 0.243538 |
| POP4 | turquoise | -0.236465 | 0.244819 |
| ANLN | turquoise | -0.236276 | 0.245208 |
| MOV10 | turquoise | -0.236051 | 0.24567 |
| C7orf23 | turquoise | -0.235517 | 0.246771 |
| TRIM27 | turquoise | -0.235471 | 0.246867 |
| CRIPT | turquoise | -0.234713 | 0.248436 |
| C8orf59 | turquoise | -0.234555 | 0.248763 |
| NTMT1 | turquoise | -0.234027 | 0.249862 |
| FADD | turquoise | -0.233995 | 0.249927 |
| IDH1 | turquoise | -0.23337 | 0.25123 |
| SCYL1 | turquoise | -0.233365 | 0.251242 |
| SLC25A24 | turquoise | -0.233147 | 0.251696 |
| ANAPC4 | turquoise | -0.233026 | 0.25195 |
| SYF2 | turquoise | -0.232411 | 0.25324 |
| PUS7L | turquoise | -0.23228 | 0.253516 |
| INTS10 | turquoise | -0.232179 | 0.253728 |
| LIX1L | turquoise | -0.232142 | 0.253806 |
| METTL2B | turquoise | -0.231946 | 0.254217 |
| RAD51C | turquoise | -0.231897 | 0.254321 |
| PCIF1 | turquoise | -0.231775 | 0.254577 |
| IGF1R | turquoise | -0.231768 | 0.254593 |
| CORO7 | turquoise | -0.231636 | 0.25487 |
| SLC39A7 | turquoise | 0.23154 | 0.255072 |
| SLC9A8 | turquoise | -0.231394 | 0.255381 |
| FAM160B1 | turquoise | -0.231279 | 0.255623 |
| TMEFF1 | turquoise | -0.30639869 | 0.120085319 |
| SMNDC1 | turquoise | -0.306287314 | 0.120226957 |
| XPNPEP3 | turquoise | -0.306128113 | 0.12042963 |
| ATPBD4 | turquoise | -0.30560592 | 0.121096173 |
| C17orf62 | turquoise | -0.305507275 | 0.12122239 |
| AGPAT3 | turquoise | -0.305382199 | 0.121382565 |
| CSNK1G2 | turquoise | -0.30364354 | 0.123625232 |
| SUMF1 | turquoise | -0.302524734 | 0.125084319 |
| DROSHA | turquoise | -0.301933971 | 0.125859823 |
| RHOQ | turquoise | -0.301726405 | 0.126133131 |
| ZDHHC8 | turquoise | -0.301402154 | 0.126560948 |
| PLGRKT | turquoise | -0.299329336 | 0.129320904 |
| UBR7 | turquoise | -0.299156674 | 0.129552767 |
| FAM3B | turquoise | -0.298347622 | 0.130643253 |
| TCEANC2 | turquoise | -0.298198447 | 0.130845047 |
| USP15 | turquoise | -0.298024209 | 0.131081031 |
| DDX24 | turquoise | -0.296994244 | 0.132482323 |
| DICER1 | turquoise | -0.296486936 | 0.133176514 |
| ARV1 | turquoise | -0.296413415 | 0.133277338 |
| COL8A2 | turquoise | -0.295745817 | 0.134195393 |
| DERA | turquoise | -0.294495052 | 0.135927733 |
| CWC15 | turquoise | -0.293585825 | 0.137197164 |
| ADNP | turquoise | -0.293527496 | 0.137278893 |
| NSL1 | turquoise | -0.290866041 | 0.141045621 |
| NDUFAF1 | turquoise | -0.290299557 | 0.141856886 |
| ALAS1 | turquoise | -0.289956669 | 0.14234957 |
| TFAM | turquoise | -0.288024434 | 0.145148992 |
| DYRK1A | turquoise | -0.287023902 | 0.146614012 |
| SPTAN1 | turquoise | -0.286630419 | 0.147193064 |
| H2AFY | turquoise | -0.285828532 | 0.148378198 |
| ATG5 | turquoise | -0.285616624 | 0.148692524 |
| EIF4E | turquoise | -0.285302044 | 0.149160022 |
| SH2B2 | turquoise | -0.284780087 | 0.149938025 |
| TIRAP | turquoise | -0.284278824 | 0.150687912 |
| CEPT1 | turquoise | -0.284021162 | 0.151074416 |
| SLC35F5 | turquoise | -0.283362847 | 0.152065133 |
| PUS7 | turquoise | -0.283151181 | 0.152384658 |
| ARAF | turquoise | -0.28121097 | 0.15533592 |
| C18orf32 | turquoise | -0.280999766 | 0.155659622 |
| CDK7 | turquoise | -0.280952059 | 0.155732807 |
| RAB14 | turquoise | -0.280600884 | 0.156272281 |
| ZNF513 | turquoise | -0.280349505 | 0.156659265 |
| SAT1 | turquoise | -0.280251259 | 0.156810692 |
| MCM8 | turquoise | -0.279299297 | 0.158283373 |
| BCKDHA | turquoise | -0.279140005 | 0.158530753 |
| SELPLG | turquoise | -0.278968332 | 0.158797668 |
| TMEM65 | turquoise | -0.278730277 | 0.159168321 |
| ZNF451 | turquoise | -0.278481988 | 0.159555563 |
| ITGB5 | turquoise | -0.277267024 | 0.161460113 |
| NR2F6 | turquoise | -0.277201977 | 0.161562531 |
| AP2B1 | turquoise | -0.276575082 | 0.162551956 |
| NCOR1 | turquoise | -0.275675177 | 0.163979758 |
| TNFAIP1 | turquoise | -0.275301295 | 0.164575564 |
| SEPSECS | turquoise | -0.275028614 | 0.165011063 |
| LIMA1 | turquoise | -0.273565784 | 0.167361249 |
| FBXW8 | turquoise | -0.273044538 | 0.168204359 |
| OTUD6B | turquoise | -0.09999 | 0.62697 |
| KAT6B | turquoise | -0.099739 | 0.627839 |
| MRPS18A | turquoise | -0.099642 | 0.628176 |
| CBLL1 | turquoise | -0.099526 | 0.628577 |
| ATP5L | turquoise | -0.099291 | 0.629392 |
| STT3B | turquoise | -0.099285 | 0.629412 |
| KATNBL1 | turquoise | -0.099137 | 0.629925 |
| RDH11 | turquoise | -0.099056 | 0.630208 |
| GPSM3 | turquoise | -0.098724 | 0.63136 |
| STK38L | turquoise | -0.098585 | 0.631844 |
| GCSHP3 | turquoise | -0.098452 | 0.632306 |
| GALC | turquoise | -0.098245 | 0.633024 |
| DUSP5 | turquoise | -0.09817 | 0.633284 |
| LGALS8 | turquoise | -0.09757 | 0.635372 |
| PPP2R1B | turquoise | -0.097564 | 0.635393 |
| ERBB2IP | turquoise | -0.097543 | 0.635467 |
| TRIM25 | turquoise | -0.09745 | 0.63579 |
| BMPR1A | turquoise | -0.097285 | 0.636366 |
| FBXO30 | turquoise | -0.096901 | 0.637706 |
| NAIF1 | turquoise | -0.096855 | 0.637863 |
| KCTD2 | turquoise | -0.096777 | 0.638137 |
| PTPRM | turquoise | -0.096739 | 0.638269 |
| PACS1 | turquoise | -0.096602 | 0.638746 |
| HYPK | turquoise | -0.096519 | 0.639038 |
| EXOC1 | turquoise | -0.096449 | 0.63928 |
| WRN | turquoise | -0.095756 | 0.641701 |
| NFYB | turquoise | -0.095439 | 0.64281 |
| ZFP161 | turquoise | -0.095349 | 0.643124 |
| ERGIC2 | turquoise | -0.095186 | 0.643696 |
| ZC3H4 | turquoise | -0.095138 | 0.643865 |
| OSGEPL1 | turquoise | -0.09512 | 0.643928 |
| ACBD6 | turquoise | -0.095022 | 0.644271 |
| EARS2 | turquoise | -0.094937 | 0.644567 |
| RB1CC1 | turquoise | -0.094858 | 0.644845 |
| CIRH1A | turquoise | -0.094622 | 0.645672 |
| MDM1 | turquoise | -0.094609 | 0.645715 |
| EHMT2 | turquoise | -0.094096 | 0.647516 |
| RLF | turquoise | -0.092988 | 0.651407 |
| EXOC5 | turquoise | -0.092883 | 0.651774 |
| SMEK1 | turquoise | -0.092535 | 0.653001 |
| ZNF76 | turquoise | -0.092512 | 0.653079 |
| PMM2 | turquoise | -0.092401 | 0.653472 |
| NIPA2 | turquoise | -0.092259 | 0.653971 |
| WBP5 | turquoise | -0.092245 | 0.654021 |
| POLH | turquoise | -0.0922 | 0.654179 |
| ARID2 | turquoise | -0.092054 | 0.654694 |
| DHX32 | turquoise | -0.091982 | 0.654947 |
| KIAA1107 | turquoise | -0.091964 | 0.655011 |
| CCT3 | turquoise | -0.091652 | 0.65611 |
| RFC3 | turquoise | -0.091204 | 0.657691 |
| TMEM165 | turquoise | -0.090995 | 0.658429 |
| CTBP2 | turquoise | -0.090705 | 0.659452 |
| NUDT15 | turquoise | -0.090702 | 0.659461 |
| APC | turquoise | -0.090636 | 0.659697 |
| ZNF292 | turquoise | -0.090532 | 0.660064 |
| PLS3 | turquoise | -0.090503 | 0.660166 |

**Table 6. Gene module membership and correlation with growth rate (48/72h), with adjusted p values for each module**

| | | | |
|---|---|---|---|
| KIAA0528 | turquoise | -0.231212 | 0.255766 |
| FUOM | turquoise | -0.230268 | 0.257766 |
| ITGB5 | turquoise | -0.230217 | 0.257876 |
| C5orf28 | turquoise | -0.229906 | 0.258537 |
| HSPB11 | turquoise | -0.229874 | 0.258604 |
| GPCPD1 | turquoise | -0.229662 | 0.259056 |
| EXOSC5 | turquoise | -0.22964 | 0.259104 |
| ZNF451 | turquoise | -0.229421 | 0.259571 |
| CCDC7 | turquoise | -0.229416 | 0.259582 |
| SEH1L | turquoise | -0.229385 | 0.259648 |
| COX16 | turquoise | -0.228944 | 0.26059 |
| ZNF576 | turquoise | -0.2288 | 0.260898 |
| OTUD3 | turquoise | 0.228183 | 0.262221 |
| AHSA2 | turquoise | -0.228001 | 0.262612 |
| NDUFA5 | turquoise | -0.227238 | 0.264256 |
| SIRPA | turquoise | -0.226861 | 0.26507 |
| B3GALT6 | turquoise | -0.226467 | 0.265924 |
| HARS2 | turquoise | -0.22616 | 0.266589 |
| UBXN2B | turquoise | -0.226016 | 0.266903 |
| CENPN | turquoise | -0.225965 | 0.267013 |
| FOXO1 | turquoise | -0.22539 | 0.268265 |
| DTWD1 | turquoise | -0.225379 | 0.26829 |
| THAP6 | turquoise | -0.225164 | 0.268758 |
| SPRY2 | turquoise | -0.224905 | 0.269324 |
| LILRB1 | turquoise | 0.22473 | 0.269706 |
| COG5 | turquoise | -0.224506 | 0.270196 |
| VPS26A | turquoise | -0.224453 | 0.270313 |
| LDHAL6B | turquoise | 0.224346 | 0.270546 |
| AK1 | turquoise | 0.224277 | 0.270699 |
| TNIK | turquoise | 0.223877 | 0.271577 |
| OCIAD1 | turquoise | -0.223745 | 0.271867 |
| RNF146 | turquoise | -0.223735 | 0.271888 |
| NDUFAF1 | turquoise | -0.222591 | 0.274412 |
| RGS10 | turquoise | -0.222196 | 0.275286 |
| C17orf58 | turquoise | -0.221615 | 0.276577 |
| OSGEPL1 | turquoise | -0.221573 | 0.276669 |
| EMD | turquoise | -0.221336 | 0.277197 |
| IQSEC2 | turquoise | -0.221095 | 0.277733 |
| ZNF217 | turquoise | -0.220679 | 0.278662 |
| ZNF788 | turquoise | -0.220547 | 0.278957 |
| ENO3 | turquoise | -0.220129 | 0.279892 |
| C1orf27 | turquoise | -0.219933 | 0.280331 |
| HCFC2 | turquoise | -0.219913 | 0.280376 |
| AHCY | turquoise | 0.219597 | 0.281085 |
| PCBD2 | turquoise | -0.219557 | 0.281174 |
| PROSER1 | turquoise | -0.219311 | 0.281728 |
| ZNF652 | turquoise | -0.219221 | 0.28193 |
| LETM1 | turquoise | -0.219028 | 0.282365 |
| LMNA | turquoise | 0.218887 | 0.282682 |
| TESK1 | turquoise | -0.218733 | 0.283029 |
| GNL2 | turquoise | -0.218547 | 0.28345 |
| DNAJC18 | turquoise | -0.218415 | 0.283748 |
| TRAK2 | turquoise | -0.218348 | 0.283898 |
| MRPL43 | turquoise | -0.2181 | 0.284459 |
| CCDC107 | turquoise | -0.217943 | 0.284815 |
| DIAPH1 | turquoise | -0.217915 | 0.28488 |
| RNF216P1 | turquoise | -0.272548857 | 0.169008888 |
| RBM14 | turquoise | -0.270549922 | 0.1722808 |
| RNF219 | turquoise | -0.27025009 | 0.172775381 |
| OGFR | turquoise | -0.270059627 | 0.173090072 |
| LACC1 | turquoise | -0.269963518 | 0.173249019 |
| CCDC6 | turquoise | -0.269420843 | 0.174148431 |
| SUMO1 | turquoise | -0.268178494 | 0.176219768 |
| TLE1 | turquoise | -0.267896121 | 0.176692956 |
| MARCKS | turquoise | -0.267244917 | 0.177787597 |
| FHOD1 | turquoise | -0.266766838 | 0.178594233 |
| CAMTA1 | turquoise | -0.266689833 | 0.178724398 |
| IDE | turquoise | -0.26639802 | 0.179218263 |
| C7orf23 | turquoise | -0.266234075 | 0.179496141 |
| SSB | turquoise | -0.265419143 | 0.180881868 |
| UBE2B | turquoise | -0.265264065 | 0.181146409 |
| SLC4A8 | turquoise | -0.265258541 | 0.181155836 |
| ERBB2IP | turquoise | -0.26477183 | 0.181987871 |
| MDM2 | turquoise | -0.264512359 | 0.182432524 |
| STRAP | turquoise | -0.264337002 | 0.182733459 |
| POLR2M | turquoise | -0.264192519 | 0.182981669 |
| NTMT1 | turquoise | -0.263563148 | 0.184065615 |
| UEVLD | turquoise | -0.263499494 | 0.184175492 |
| CGRRF1 | turquoise | -0.2627899 | 0.185403454 |
| BAD | turquoise | -0.262763969 | 0.185448434 |
| USP53 | turquoise | -0.262667731 | 0.18561544 |
| FTSJD1 | turquoise | -0.26219303 | 0.186440734 |
| IFNAR1 | turquoise | -0.261382418 | 0.187855903 |
| WBP4 | turquoise | -0.260885114 | 0.188727771 |
| VRK2 | turquoise | -0.259059324 | 0.191952723 |
| XPO1 | turquoise | -0.259010018 | 0.192040338 |
| FYB | turquoise | -0.258966985 | 0.192116828 |
| CEBPD | turquoise | -0.258253701 | 0.19338774 |
| HBS1L | turquoise | -0.258028551 | 0.193790109 |
| RGS19 | turquoise | -0.257966919 | 0.193900353 |
| ARHGEF5 | turquoise | -0.25776632 | 0.19425947 |
| LOC728377 | turquoise | -0.257573268 | 0.19460551 |
| FOXK1 | turquoise | -0.257056268 | 0.195534306 |
| NAP1L1 | turquoise | -0.256437633 | 0.196649691 |
| SSBP1 | turquoise | -0.256023403 | 0.197398979 |
| SNX1 | turquoise | -0.255776574 | 0.19784639 |
| ZNF613 | turquoise | -0.254388202 | 0.20037598 |
| SP140L | turquoise | -0.253121574 | 0.202703002 |
| HSPB11 | turquoise | -0.253062208 | 0.202812518 |
| DCUN1D4 | turquoise | -0.253006952 | 0.202914489 |
| TCF20 | turquoise | -0.252209652 | 0.204389758 |
| ZNF689 | turquoise | -0.252076834 | 0.204636225 |
| APTX | turquoise | -0.252062689 | 0.204662484 |
| RND3 | turquoise | -0.251798048 | 0.205154219 |
| LRRCC1 | turquoise | -0.25140298 | 0.205889801 |
| LRP10 | turquoise | -0.251370443 | 0.205950462 |
| EMR1 | turquoise | -0.251219266 | 0.206232472 |
| RAB11FIP2 | turquoise | -0.250241023 | 0.208063677 |
| FAM173B | turquoise | -0.249552834 | 0.209358529 |
| ADSL | turquoise | -0.249337651 | 0.209764525 |
| R3HDM2 | turquoise | -0.248924434 | 0.21054566 |
| SF3A2 | turquoise | -0.248907399 | 0.210577906 |
| AASDHPPT | turquoise | -0.090252 | 0.661052 |
| SH3BGRL | turquoise | -0.09013 | 0.661483 |
| TEKT4P2 | turquoise | -0.090095 | 0.661606 |
| TMEM126A | turquoise | -0.08967 | 0.663111 |
| BCAP29 | turquoise | -0.089655 | 0.663163 |
| PITPNA | turquoise | -0.089634 | 0.663238 |
| IDH1 | turquoise | -0.089606 | 0.663337 |
| ERP44 | turquoise | -0.089304 | 0.664408 |
| ARG2 | turquoise | -0.089016 | 0.665426 |
| STXBP2 | turquoise | -0.089011 | 0.665445 |
| FYB | turquoise | -0.088336 | 0.667839 |
| ATP9B | turquoise | -0.088221 | 0.668246 |
| TXNIP | turquoise | -0.088049 | 0.66886 |
| SBNO1 | turquoise | -0.087886 | 0.669437 |
| LLPH | turquoise | -0.087505 | 0.670791 |
| SKP1 | turquoise | -0.087418 | 0.671099 |
| TRAFD1 | turquoise | -0.087045 | 0.672427 |
| LOC728730 | turquoise | -0.086724 | 0.673569 |
| IRF2 | turquoise | -0.08637 | 0.67483 |
| KLHL9 | turquoise | -0.086294 | 0.6751 |
| BTN3A3 | turquoise | -0.086155 | 0.675595 |
| ERI1 | turquoise | -0.086148 | 0.675622 |
| RABEP1 | turquoise | -0.086034 | 0.676027 |
| NME4 | turquoise | -0.085807 | 0.676837 |
| MAPK7 | turquoise | -0.085799 | 0.676867 |
| AIM1 | turquoise | -0.085405 | 0.67827 |
| AMZ1 | turquoise | -0.085043 | 0.679563 |
| PIAS2 | turquoise | -0.084733 | 0.680673 |
| PIR | turquoise | -0.084557 | 0.681299 |
| MBD4 | turquoise | -0.084436 | 0.681731 |
| PTBP2 | turquoise | -0.084421 | 0.681787 |
| BICD2 | turquoise | -0.084284 | 0.682276 |
| KLHL7 | turquoise | -0.084011 | 0.683251 |
| RAB30 | turquoise | -0.083969 | 0.683405 |
| LAMTOR3 | turquoise | -0.083796 | 0.684022 |
| SLC35A5 | turquoise | -0.08258 | 0.688381 |
| CNTLN | turquoise | -0.082522 | 0.688588 |
| ZNF407 | turquoise | -0.082499 | 0.688669 |
| FES | turquoise | -0.082388 | 0.689069 |
| SF3B14 | turquoise | -0.082199 | 0.689748 |
| FANCL | turquoise | -0.082131 | 0.689991 |
| MON2 | turquoise | -0.081548 | 0.692086 |
| ZFC3H1 | turquoise | -0.080685 | 0.695193 |
| DHDDS | turquoise | -0.080626 | 0.695404 |
| BCKDHB | turquoise | -0.08057 | 0.695604 |
| KBTBD8 | turquoise | -0.080415 | 0.696162 |
| PAFAH1B3 | turquoise | -0.08007 | 0.697406 |
| ING2 | turquoise | -0.080007 | 0.697633 |
| CAPN1 | turquoise | -0.079741 | 0.698592 |
| ROCK1 | turquoise | -0.07974 | 0.698595 |
| SLC25A43 | turquoise | -0.079563 | 0.699235 |
| CDC27 | turquoise | -0.078884 | 0.701685 |
| NUCB2 | turquoise | -0.078856 | 0.701786 |
| MAN2C1 | turquoise | -0.078589 | 0.702752 |
| TIMP3 | turquoise | -0.078526 | 0.702979 |
| RNFT1 | turquoise | -0.078483 | 0.703136 |

**Table 6. Gene module membership and correlation with growth rate (48/72h), with adjusted p values for each module**

| | | | |
|---|---|---|---|
| CENPW | turquoise | -0.217831 | 0.28507 |
| HERC2P9 | turquoise | -0.216909 | 0.287163 |
| NARG2 | turquoise | -0.216316 | 0.288516 |
| CENPB | turquoise | -0.216109 | 0.288987 |
| ORC6 | turquoise | -0.215886 | 0.289498 |
| TACC3 | turquoise | -0.215707 | 0.289908 |
| ZC3H3 | turquoise | -0.21565 | 0.290038 |
| FBP1 | turquoise | -0.215282 | 0.290884 |
| LRRC40 | turquoise | -0.214978 | 0.291581 |
| TBL3 | turquoise | -0.214934 | 0.291681 |
| ZBTB25 | turquoise | -0.214725 | 0.292164 |
| RGP1 | turquoise | -0.214433 | 0.292835 |
| SLC2A6 | turquoise | -0.214156 | 0.293474 |
| C3orf37 | turquoise | -0.213253 | 0.295563 |
| RFK | turquoise | -0.213205 | 0.295675 |
| PPP2R3C | turquoise | -0.212966 | 0.296228 |
| FES | turquoise | -0.212675 | 0.296906 |
| ABCC3 | turquoise | -0.212594 | 0.297094 |
| PRPF6 | turquoise | -0.212443 | 0.297445 |
| PPIP5K2 | turquoise | -0.21224 | 0.297918 |
| MIR1307 | turquoise | 0.21218 | 0.298056 |
| AASDHPPT | turquoise | -0.211887 | 0.298741 |
| SLC12A4 | turquoise | -0.211692 | 0.299195 |
| FAM221A | turquoise | -0.211646 | 0.299304 |
| PSMA4 | turquoise | -0.211618 | 0.299369 |
| TRMT13 | turquoise | -0.211162 | 0.300435 |
| RPL19 | turquoise | -0.211035 | 0.300734 |
| ZDHHC20 | turquoise | -0.210406 | 0.302209 |
| C11orf74 | turquoise | -0.210184 | 0.302732 |
| GLYR1 | turquoise | -0.209904 | 0.30339 |
| TYK2 | turquoise | -0.20966 | 0.303967 |
| RBM48 | turquoise | -0.209288 | 0.304845 |
| TRIQK | turquoise | -0.208971 | 0.305594 |
| MPV17L2 | turquoise | -0.208928 | 0.305696 |
| IRF2BP2 | turquoise | -0.208635 | 0.306391 |
| TMEM192 | turquoise | -0.208626 | 0.306411 |
| ZDHHC16 | turquoise | -0.208363 | 0.307036 |
| LOC388796 | turquoise | 0.208208 | 0.307404 |
| CHCHD10 | turquoise | -0.20812 | 0.307613 |
| CCDC25 | turquoise | -0.208013 | 0.307867 |
| MCAT | turquoise | -0.20792 | 0.308089 |
| SMIM7 | turquoise | -0.207644 | 0.308745 |
| LINC00674 | turquoise | -0.207567 | 0.308929 |
| BEX4 | turquoise | -0.20704 | 0.310186 |
| ACOT8 | turquoise | -0.206826 | 0.310698 |
| RANBP10 | turquoise | -0.206634 | 0.311157 |
| RPL32 | turquoise | -0.206144 | 0.312331 |
| MAN2B1 | turquoise | -0.20608 | 0.312484 |
| ANKRA2 | turquoise | -0.205659 | 0.313498 |
| FAM13B | turquoise | -0.205442 | 0.31402 |
| KIAA1586 | turquoise | -0.205072 | 0.314911 |
| CTSA | turquoise | -0.20507 | 0.314915 |
| NIPA2 | turquoise | -0.205055 | 0.314952 |
| RAB27A | turquoise | -0.20456 | 0.316147 |
| GINM1 | turquoise | -0.20451 | 0.316268 |
| DNAJC13 | turquoise | -0.203992 | 0.317522 |
| DNAJC3 | turquoise | -0.248302449 | 0.211725165 |
| METTL8 | turquoise | -0.247865629 | 0.212556205 |
| CENPC1 | turquoise | -0.246990745 | 0.214227292 |
| UBE2W | turquoise | -0.245919682 | 0.216285171 |
| VOPP1 | turquoise | -0.244270391 | 0.219480061 |
| TMED4 | turquoise | -0.243996211 | 0.220014249 |
| GGPS1 | turquoise | -0.24344466 | 0.221091489 |
| NUDT16 | turquoise | -0.243334072 | 0.221307907 |
| SLC25A20 | turquoise | -0.242669124 | 0.222612191 |
| SLMO2 | turquoise | -0.24261318 | 0.22272216 |
| SEC22A | turquoise | -0.242098512 | 0.223735546 |
| HSPB1 | turquoise | -0.242061194 | 0.223809147 |
| DPH2 | turquoise | -0.241734695 | 0.224453771 |
| ACAP3 | turquoise | -0.24167119 | 0.224579295 |
| VPS35 | turquoise | -0.241444495 | 0.22502777 |
| LFNG | turquoise | -0.240662523 | 0.226579367 |
| RBM48 | turquoise | -0.240632336 | 0.226639407 |
| ZNF292 | turquoise | -0.240435838 | 0.227030492 |
| GTF2H1 | turquoise | -0.240395052 | 0.227111726 |
| EXOC6 | turquoise | -0.240258828 | 0.227383179 |
| RAD17 | turquoise | -0.240166462 | 0.22756736 |
| AIP | turquoise | -0.24014255 | 0.227615058 |
| TESK1 | turquoise | -0.239887203 | 0.228124819 |
| PLEKHM3 | turquoise | -0.239236758 | 0.229426779 |
| DTX4 | turquoise | -0.239104702 | 0.229691712 |
| KIAA1522 | turquoise | -0.23799406 | 0.231927991 |
| ZBED5 | turquoise | -0.237498827 | 0.232929803 |
| MCM3AP | turquoise | -0.237491065 | 0.232945527 |
| FOXO1 | turquoise | -0.237338837 | 0.233254063 |
| IKBIP | turquoise | -0.237255181 | 0.233423731 |
| FAM160B1 | turquoise | -0.236679822 | 0.234592887 |
| TP53INP2 | turquoise | -0.235932969 | 0.236116323 |
| RANBP3 | turquoise | -0.235103678 | 0.237815593 |
| RHOF | turquoise | -0.235023161 | 0.237981009 |
| ATXN1L | turquoise | -0.234818843 | 0.238401103 |
| CDK13 | turquoise | -0.234180816 | 0.239716101 |
| ALKBH6 | turquoise | -0.233993395 | 0.240103293 |
| CD47 | turquoise | -0.233909009 | 0.240277761 |
| ACADM | turquoise | -0.233584405 | 0.240949659 |
| ZNF441 | turquoise | -0.233128027 | 0.241896414 |
| COX11 | turquoise | -0.233035278 | 0.242089121 |
| ZNF516 | turquoise | -0.232816049 | 0.242545024 |
| KCTD9 | turquoise | -0.232706153 | 0.242773773 |
| ESYT2 | turquoise | -0.232630866 | 0.242930565 |
| NRAS | turquoise | -0.231876148 | 0.24450603 |
| GPS2 | turquoise | -0.23166557 | 0.244946808 |
| CPM | turquoise | -0.231593863 | 0.245097024 |
| KIAA0513 | turquoise | -0.230690523 | 0.246994573 |
| ATP6V1H | turquoise | -0.230533215 | 0.247325999 |
| SNX9 | turquoise | -0.229891907 | 0.248680162 |
| CCP110 | turquoise | -0.229516325 | 0.249475484 |
| RARS | turquoise | -0.229102075 | 0.250354618 |
| DYNLL2 | turquoise | -0.229050892 | 0.250463382 |
| SH3BP5L | turquoise | -0.228531029 | 0.251569836 |
| B4GALT2 | turquoise | -0.228145962 | 0.252391456 |
| GALT | turquoise | -0.228143019 | 0.252397743 |
| AZIN1 | turquoise | -0.078443 | 0.70328 |
| TAF9 | turquoise | -0.078325 | 0.703706 |
| ST7 | turquoise | -0.078102 | 0.704512 |
| KIAA0947 | turquoise | -0.077916 | 0.705186 |
| RAP2B | turquoise | -0.077911 | 0.705202 |
| PUS3 | turquoise | -0.077188 | 0.707819 |
| ARL5A | turquoise | -0.077159 | 0.707926 |
| LOC643837 | turquoise | -0.076898 | 0.70887 |
| SLC4A7 | turquoise | -0.076221 | 0.711326 |
| COIL | turquoise | -0.075213 | 0.714987 |
| ARFGEF1 | turquoise | -0.075128 | 0.715295 |
| TNRC6C | turquoise | -0.074981 | 0.71583 |
| MAP2K4 | turquoise | -0.074959 | 0.71591 |
| ZFYVE21 | turquoise | -0.074766 | 0.71661 |
| ATG7 | turquoise | -0.074758 | 0.716639 |
| NHLRC3 | turquoise | -0.074624 | 0.717126 |
| HSDL2 | turquoise | -0.07452 | 0.717504 |
| DNAJB4 | turquoise | -0.074269 | 0.71842 |
| SLC25A36 | turquoise | -0.073893 | 0.719787 |
| ICOS | turquoise | -0.073813 | 0.72008 |
| CLK1 | turquoise | -0.073699 | 0.720495 |
| KLHL15 | turquoise | -0.073628 | 0.720753 |
| TWISTNB | turquoise | -0.073581 | 0.720924 |
| MINA | turquoise | -0.073342 | 0.721797 |
| ATP5A1 | turquoise | -0.073315 | 0.721895 |
| EXOC6 | turquoise | -0.073199 | 0.722315 |
| ERO1L | turquoise | -0.073138 | 0.722541 |
| SEPSECS | turquoise | -0.072782 | 0.723837 |
| RAD50 | turquoise | -0.072103 | 0.726316 |
| SNX16 | turquoise | -0.072064 | 0.726461 |
| CCNL1 | turquoise | -0.071468 | 0.728638 |
| SLC25A16 | turquoise | -0.071252 | 0.729426 |
| ZDHHC4 | turquoise | -0.071193 | 0.729642 |
| SIGLEC11 | turquoise | -0.070844 | 0.730921 |
| NUDT3 | turquoise | -0.070275 | 0.733002 |
| APLF | turquoise | -0.070227 | 0.733181 |
| RHBDD2 | turquoise | -0.069959 | 0.734161 |
| DCAF7 | turquoise | -0.069712 | 0.735067 |
| PPIB | turquoise | -0.069487 | 0.735893 |
| HLTF | turquoise | -0.068886 | 0.738097 |
| RAB8B | turquoise | -0.068834 | 0.738287 |
| CABLES2 | turquoise | -0.068828 | 0.738312 |
| SEPN1 | turquoise | -0.068791 | 0.738447 |
| ALAD | turquoise | -0.06854 | 0.73937 |
| SMEK2 | turquoise | -0.068395 | 0.7399 |
| AGTPBP1 | turquoise | -0.06838 | 0.739956 |
| AP1G1 | turquoise | -0.067643 | 0.742665 |
| IFNGR1 | turquoise | -0.067498 | 0.743198 |
| ASB13 | turquoise | -0.067476 | 0.74328 |
| VGLL4 | turquoise | -0.067295 | 0.743947 |
| MTM1 | turquoise | -0.067181 | 0.744364 |
| SLAMF6 | turquoise | -0.067137 | 0.744525 |
| MLF1 | turquoise | -0.066371 | 0.747347 |
| PTOV1 | turquoise | -0.066221 | 0.7479 |
| CD84 | turquoise | -0.066176 | 0.748066 |
| RECK | turquoise | -0.066116 | 0.748288 |

**Table 6. Gene module membership and correlation with growth rate (48/72h), with adjusted p values for each module**

| | | | |
|---|---|---|---|
| ASXL2 | turquoise | -0.203791 | 0.318008 |
| CEP350 | turquoise | -0.2031 | 0.319688 |
| SMARCAL1 | turquoise | -0.202803 | 0.320411 |
| CNEP1R1 | turquoise | -0.202714 | 0.320629 |
| DHX40 | turquoise | -0.202364 | 0.321481 |
| FGD4 | turquoise | 0.202284 | 0.321676 |
| RGCC | turquoise | 0.201443 | 0.323735 |
| CMC1 | turquoise | -0.200947 | 0.324955 |
| CNBP | turquoise | -0.200936 | 0.324981 |
| HERC5 | turquoise | -0.200279 | 0.326599 |
| RANBP17 | turquoise | -0.200274 | 0.32661 |
| DHX32 | turquoise | -0.200234 | 0.326709 |
| MRPS9 | turquoise | 0.199845 | 0.327669 |
| APLF | turquoise | -0.199728 | 0.327959 |
| RAB28 | turquoise | -0.199184 | 0.329305 |
| ZNF883 | turquoise | -0.199124 | 0.329453 |
| KANSL3 | turquoise | -0.199042 | 0.329657 |
| AARS2 | turquoise | -0.198232 | 0.331669 |
| LOC729737 | turquoise | -0.197858 | 0.332601 |
| LOC374443 | turquoise | 0.19782 | 0.332695 |
| CLYBL | turquoise | -0.197405 | 0.333731 |
| ALG6 | turquoise | -0.197311 | 0.333967 |
| COX11 | turquoise | -0.197117 | 0.334451 |
| NAF1 | turquoise | -0.197104 | 0.334486 |
| BCL10 | turquoise | -0.196815 | 0.335209 |
| HSF2 | turquoise | -0.196376 | 0.336308 |
| TMEM191E | turquoise | -0.196273 | 0.336568 |
| BLOC1S2 | turquoise | -0.196088 | 0.337032 |
| C9orf72 | turquoise | -0.196048 | 0.337133 |
| ZNF567 | turquoise | 0.195777 | 0.337814 |
| RCN1 | turquoise | -0.195513 | 0.338479 |
| NR1H2 | turquoise | -0.195333 | 0.338932 |
| LAMB2 | turquoise | -0.195093 | 0.339537 |
| TBC1D3 | turquoise | 0.195043 | 0.339665 |
| LGALSL | turquoise | -0.194773 | 0.340346 |
| PYURF | turquoise | -0.194238 | 0.341701 |
| IPO5 | turquoise | -0.194192 | 0.341817 |
| ADCK2 | turquoise | -0.193603 | 0.343312 |
| PTGER2 | turquoise | -0.193565 | 0.343409 |
| KLHL9 | turquoise | -0.193504 | 0.343564 |
| VPS33A | turquoise | -0.193416 | 0.343787 |
| SLC35E2B | turquoise | -0.192795 | 0.345368 |
| POLD1 | turquoise | -0.192603 | 0.345859 |
| API5 | turquoise | -0.192561 | 0.345965 |
| CDC42 | turquoise | -0.192332 | 0.346549 |
| LRP10 | turquoise | -0.192204 | 0.346877 |
| WRN | turquoise | -0.192182 | 0.346934 |
| CACYBP | turquoise | -0.192129 | 0.347069 |
| DNAJB14 | turquoise | -0.191914 | 0.347619 |
| GPAT2 | turquoise | -0.191735 | 0.348077 |
| HS2ST1 | turquoise | -0.191575 | 0.348488 |
| DENR | turquoise | -0.191529 | 0.348606 |
| ITGB1 | turquoise | -0.191513 | 0.348646 |
| C1D | turquoise | -0.191492 | 0.348702 |
| ZMIZ1 | turquoise | 0.191428 | 0.348866 |
| GOLGA7 | turquoise | -0.191263 | 0.349289 |
| GPSM3 | turquoise | -0.227102882 | 0.254625891 |
| RP9 | turquoise | -0.226946629 | 0.254961715 |
| PUS7L | turquoise | -0.226784152 | 0.255311225 |
| BCL2 | turquoise | -0.226761845 | 0.255359235 |
| CNOT2 | turquoise | -0.226516682 | 0.255887269 |
| DOCK7 | turquoise | -0.225718458 | 0.257611417 |
| GNL2 | turquoise | -0.225080762 | 0.25899425 |
| CENPW | turquoise | -0.225008111 | 0.259152098 |
| COG8 | turquoise | -0.224671125 | 0.259885083 |
| ILK | turquoise | -0.224487972 | 0.260284026 |
| SETX | turquoise | -0.224215886 | 0.260877417 |
| TCAIM | turquoise | -0.224105009 | 0.261119479 |
| RASSF7 | turquoise | -0.22365623 | 0.262100725 |
| S1PR4 | turquoise | -0.223530767 | 0.262375474 |
| IPO5 | turquoise | -0.222595057 | 0.264430457 |
| CAPZA2 | turquoise | -0.222239696 | 0.265213612 |
| BBS12 | turquoise | -0.221325948 | 0.267234235 |
| TANK | turquoise | -0.221082405 | 0.267774468 |
| HCFC1R1 | turquoise | -0.220395672 | 0.269301584 |
| CYGB | turquoise | -0.219353803 | 0.271629121 |
| LONRF1 | turquoise | -0.219078258 | 0.272246844 |
| IRAK4 | turquoise | -0.218860681 | 0.27273525 |
| CDC42 | turquoise | -0.218821761 | 0.272822674 |
| ZNF800 | turquoise | -0.218049161 | 0.274561868 |
| HUS1 | turquoise | -0.217976466 | 0.274725876 |
| ODF2 | turquoise | -0.217305543 | 0.276242515 |
| MAP4K5 | turquoise | -0.216952293 | 0.277043197 |
| UBE2L6 | turquoise | -0.216460574 | 0.2781602 |
| UCHL1 | turquoise | -0.215964068 | 0.279290991 |
| GTF2I | turquoise | -0.214965684 | 0.281573678 |
| CMTM3 | turquoise | -0.213541914 | 0.284849448 |
| ZNF385A | turquoise | -0.213535424 | 0.284864434 |
| SLC35C1 | turquoise | -0.212866847 | 0.28641104 |
| EIF3I | turquoise | -0.212283198 | 0.287765527 |
| FAM193A | turquoise | -0.211871003 | 0.288724556 |
| ABCC3 | turquoise | -0.21140885 | 0.28980222 |
| PPP3CB | turquoise | -0.211179906 | 0.290337019 |
| TNNI2 | turquoise | -0.211072306 | 0.290588581 |
| TRPV2 | turquoise | -0.211047826 | 0.290645835 |
| KIAA0368 | turquoise | -0.210934701 | 0.290910494 |
| NOC2L | turquoise | -0.210798113 | 0.29123025 |
| TSEN15 | turquoise | -0.21074504 | 0.291354556 |
| TCEAL8 | turquoise | -0.210216057 | 0.292595343 |
| RMRP | turquoise | -0.209794215 | 0.293587203 |
| SLC9A9 | turquoise | -0.208870151 | 0.295767303 |
| TBC1D15 | turquoise | -0.208553327 | 0.296517105 |
| CCND3 | turquoise | -0.208494416 | 0.296656657 |
| MOB1B | turquoise | -0.208451577 | 0.296758163 |
| WDR44 | turquoise | -0.20793199 | 0.297991038 |
| KLF8 | turquoise | -0.207817792 | 0.298262436 |
| RRP8 | turquoise | -0.206200624 | 0.302122362 |
| RUFY2 | turquoise | -0.20572635 | 0.303260266 |
| C18orf21 | turquoise | -0.204945393 | 0.305139798 |
| ZWILCH | turquoise | -0.204047157 | 0.307310534 |
| PLCL2 | turquoise | -0.203596403 | 0.308403462 |
| LMBRD1 | turquoise | -0.20334004 | 0.30902613 |
| COPS7B | turquoise | -0.066062 | 0.748487 |
| TMEM30A | turquoise | -0.065994 | 0.748739 |
| UFSP2 | turquoise | -0.065854 | 0.749252 |
| CCDC117 | turquoise | -0.065532 | 0.75044 |
| RAB2A | turquoise | -0.065465 | 0.750688 |
| CEP152 | turquoise | -0.064621 | 0.753805 |
| CCDC25 | turquoise | -0.064082 | 0.755797 |
| C1GALT1C1 | turquoise | -0.063992 | 0.75613 |
| FAM49B | turquoise | -0.063889 | 0.756512 |
| MAP3K5 | turquoise | -0.063658 | 0.757365 |
| IQGAP2 | turquoise | -0.063211 | 0.75902 |
| CRAMP1L | turquoise | -0.063029 | 0.759693 |
| KDM4C | turquoise | -0.062599 | 0.761286 |
| PPP6C | turquoise | -0.062516 | 0.761594 |
| N4BP2L2 | turquoise | -0.062341 | 0.762242 |
| VPS33A | turquoise | -0.062299 | 0.762397 |
| UTP18 | turquoise | -0.062228 | 0.762659 |
| OMA1 | turquoise | -0.062082 | 0.7632 |
| FERMT3 | turquoise | -0.061987 | 0.763554 |
| TRIQK | turquoise | -0.061935 | 0.763746 |
| SOGA1 | turquoise | -0.061845 | 0.764082 |
| EVI2A | turquoise | -0.061795 | 0.764264 |
| SLC12A7 | turquoise | -0.061656 | 0.76478 |
| FAM98A | turquoise | -0.061478 | 0.765443 |
| RAD1 | turquoise | -0.061453 | 0.765534 |
| COPS3 | turquoise | -0.0614 | 0.765733 |
| PGM3 | turquoise | -0.061323 | 0.766017 |
| FAM160B1 | turquoise | -0.060979 | 0.767294 |
| ASXL2 | turquoise | -0.06044 | 0.769297 |
| PCGF3 | turquoise | -0.059966 | 0.771057 |
| CCNG1 | turquoise | -0.059667 | 0.772168 |
| ITPR2 | turquoise | -0.059258 | 0.773692 |
| RNF214 | turquoise | -0.059225 | 0.773814 |
| PAPOLA | turquoise | -0.059179 | 0.773987 |
| ICOSLG | turquoise | -0.059162 | 0.774047 |
| SPRED2 | turquoise | -0.059149 | 0.774096 |
| ZNF331 | turquoise | -0.059107 | 0.774254 |
| DKFZP586I14 | turquoise | -0.058867 | 0.775148 |
| DPY30 | turquoise | -0.058783 | 0.775459 |
| NUPL1 | turquoise | -0.058775 | 0.77549 |
| C2orf18 | turquoise | -0.058731 | 0.775654 |
| PTGER4 | turquoise | -0.058592 | 0.77617 |
| MALAT1 | turquoise | -0.058555 | 0.776309 |
| MON1B | turquoise | -0.058553 | 0.776317 |
| RWDD2A | turquoise | -0.058528 | 0.776408 |
| TOP2B | turquoise | -0.058125 | 0.777911 |
| TACC3 | turquoise | -0.057213 | 0.781311 |
| UCHL5 | turquoise | -0.056938 | 0.782338 |
| RAB3IP | turquoise | -0.056622 | 0.783517 |
| PRKAR2A | turquoise | -0.056619 | 0.78353 |
| LRRC8D | turquoise | -0.056596 | 0.783616 |
| DYNC1LI1 | turquoise | -0.056372 | 0.78445 |
| FADD | turquoise | -0.056245 | 0.784926 |
| DDX42 | turquoise | -0.056097 | 0.785477 |
| ZNF638 | turquoise | -0.055855 | 0.786385 |
| HSPA13 | turquoise | -0.054801 | 0.790325 |

**Table 6. Gene module membership and correlation with growth rate (48/72h), with adjusted p values for each module**

| | | | | | | | | | | | |
|---|---|---|---|---|---|---|---|---|---|---|---|
| CLMN | turquoise | -0.190366 | 0.351595 | LRPPRC | turquoise | -0.203219362 | 0.309319512 | LRRC58 | turquoise | -0.054776 | 0.790418 |
| FAM53C | turquoise | 0.190275 | 0.351831 | PPM1F | turquoise | -0.203176283 | 0.309424282 | SDAD1 | turquoise | -0.054702 | 0.790696 |
| C22orf32 | turquoise | -0.190156 | 0.352139 | GPC1 | turquoise | -0.203092083 | 0.309629124 | LRIF1 | turquoise | -0.054702 | 0.790698 |
| ABHD8 | turquoise | -0.19011 | 0.352257 | TMPO | turquoise | -0.202769636 | 0.310414357 | FASN | turquoise | -0.054649 | 0.790894 |
| ISOC2 | turquoise | -0.189947 | 0.352676 | SPATA7 | turquoise | -0.202332346 | 0.311481229 | HIVEP1 | turquoise | -0.054512 | 0.791408 |
| DENND6A | turquoise | -0.189738 | 0.353216 | NOTCH1 | turquoise | -0.202232393 | 0.311725405 | CASP8AP2 | turquoise | -0.054508 | 0.791421 |
| GIPC1 | turquoise | 0.18973 | 0.353237 | KIAA1586 | turquoise | -0.202012766 | 0.312262351 | TRAPPC6B | turquoise | -0.054426 | 0.791731 |
| GLRX3 | turquoise | -0.189532 | 0.353748 | TRIM25 | turquoise | -0.201808881 | 0.312761322 | PER1 | turquoise | -0.054294 | 0.792223 |
| ABCE1 | turquoise | -0.189092 | 0.354889 | DCAF17 | turquoise | -0.200217897 | 0.316671878 | MTMR12 | turquoise | -0.053988 | 0.793369 |
| TCF19 | turquoise | -0.18909 | 0.354895 | AHR | turquoise | -0.200130852 | 0.316886695 | USP8 | turquoise | -0.053975 | 0.793419 |
| CSRNP1 | turquoise | -0.189068 | 0.354951 | AGXT2L2 | turquoise | -0.198866774 | 0.320016395 | TESK1 | turquoise | -0.053721 | 0.79437 |
| SBF1 | turquoise | -0.188878 | 0.355445 | LPGAT1 | turquoise | -0.198745295 | 0.320318158 | ZNF148 | turquoise | -0.053655 | 0.794617 |
| PRMT10 | turquoise | -0.188393 | 0.356703 | HMHA1 | turquoise | -0.198329225 | 0.321353027 | SNW1 | turquoise | -0.053615 | 0.794766 |
| PUF60 | turquoise | -0.188116 | 0.357423 | DNAJC13 | turquoise | -0.197497667 | 0.323427453 | RASA2 | turquoise | -0.053322 | 0.795866 |
| SMURF2 | turquoise | -0.188061 | 0.357566 | SLC35E2B | turquoise | -0.197432942 | 0.323589259 | CENPN | turquoise | -0.053258 | 0.796105 |
| CXCR7 | turquoise | -0.187923 | 0.357926 | UBXN2B | turquoise | -0.19690242 | 0.324917386 | TM2D1 | turquoise | -0.053131 | 0.796583 |
| ARL4C | turquoise | -0.187473 | 0.3591 | SRD5A3 | turquoise | -0.195738781 | 0.327842119 | MCM3AP | turquoise | -0.052968 | 0.797193 |
| KDELC2 | turquoise | -0.187445 | 0.359171 | SNX33 | turquoise | -0.195269758 | 0.329025496 | FGL2 | turquoise | -0.052769 | 0.79794 |
| THOP1 | turquoise | -0.187436 | 0.359197 | ITFG1 | turquoise | -0.19489875 | 0.329963414 | BNIP2 | turquoise | -0.05224 | 0.799924 |
| C5orf51 | turquoise | -0.187419 | 0.359241 | RWDD1 | turquoise | -0.194580558 | 0.330769104 | FAM206A | turquoise | -0.052061 | 0.800595 |
| HNRNPA1L2 | turquoise | -0.187098 | 0.360079 | NCKIPSD | turquoise | -0.194440648 | 0.331123747 | EGLN1 | turquoise | -0.051697 | 0.801963 |
| LIN7C | turquoise | 0.186806 | 0.360843 | MAP7D1 | turquoise | -0.194099377 | 0.331989763 | FTSJ3 | turquoise | -0.051664 | 0.802087 |
| RNF168 | turquoise | -0.18656 | 0.361488 | GCC2 | turquoise | -0.193827066 | 0.332681769 | NMD3 | turquoise | -0.05129 | 0.803492 |
| HIPK2 | turquoise | 0.186243 | 0.362319 | ZSCAN18 | turquoise | -0.193801377 | 0.332747095 | SUZ12 | turquoise | -0.05117 | 0.803942 |
| SLC35A5 | turquoise | -0.18596 | 0.363062 | FAM199X | turquoise | -0.193386537 | 0.333803108 | VPS4B | turquoise | -0.051062 | 0.804349 |
| NOL11 | turquoise | -0.18547 | 0.36435 | LRIF1 | turquoise | -0.192840727 | 0.3351956 | C4orf52 | turquoise | -0.050652 | 0.805891 |
| CD55 | turquoise | 0.185232 | 0.364977 | SCYL2 | turquoise | -0.19259446 | 0.335825033 | CLIC2 | turquoise | -0.049749 | 0.809287 |
| LAMTOR3 | turquoise | -0.185137 | 0.365227 | GNPAT | turquoise | -0.192381057 | 0.336371047 | JMJD1C | turquoise | -0.049537 | 0.810086 |
| MRC1 | turquoise | -0.184972 | 0.365661 | RBBP7 | turquoise | -0.192274726 | 0.336643308 | ADAMDEC1 | turquoise | -0.049455 | 0.810396 |
| EIF1B | turquoise | 0.184938 | 0.365751 | PANK2 | turquoise | -0.191851437 | 0.33772845 | HCG26 | turquoise | -0.04945 | 0.810415 |
| SCARNA6 | turquoise | -0.184512 | 0.366876 | C6orf70 | turquoise | -0.191538469 | 0.33853213 | ISCU | turquoise | -0.048782 | 0.812929 |
| ZNF331 | turquoise | -0.184011 | 0.368202 | SNRNP25 | turquoise | -0.191414292 | 0.338851327 | ABCC3 | turquoise | -0.048586 | 0.813669 |
| PEG10 | turquoise | -0.183833 | 0.368672 | RILPL2 | turquoise | -0.191108385 | 0.339638433 | DHFR | turquoise | -0.048434 | 0.814241 |
| TTC1 | turquoise | -0.183519 | 0.369505 | PHC3 | turquoise | -0.19107867 | 0.339714949 | CAAP1 | turquoise | -0.048183 | 0.815189 |
| PPP1R37 | turquoise | -0.183509 | 0.369532 | VAMP7 | turquoise | -0.190796139 | 0.340442982 | CCDC82 | turquoise | -0.047856 | 0.81642 |
| TRAPPC5 | turquoise | -0.183301 | 0.370083 | C9orf72 | turquoise | -0.19043093 | 0.341385451 | RGP1 | turquoise | -0.047845 | 0.816461 |
| LARS2 | turquoise | -0.183104 | 0.370608 | GPATCH2L | turquoise | -0.190180544 | 0.342032509 | NOP58 | turquoise | -0.047551 | 0.817573 |
| TARBP1 | turquoise | -0.182754 | 0.371539 | RAP2B | turquoise | -0.189953592 | 0.342619646 | CASP4 | turquoise | -0.047376 | 0.818233 |
| ST5 | turquoise | -0.182566 | 0.372039 | ACTR3 | turquoise | -0.189603185 | 0.34352735 | ARPC5 | turquoise | -0.047264 | 0.818656 |
| VPRBP | turquoise | -0.182428 | 0.372408 | WDR61 | turquoise | -0.189571163 | 0.343610372 | NDUFB4 | turquoise | -0.047086 | 0.819326 |
| MRPS10 | turquoise | -0.182011 | 0.373521 | PPP6R1 | turquoise | -0.188467597 | 0.346478906 | ZDHHC5 | turquoise | -0.046938 | 0.819885 |
| RHOB | turquoise | -0.181835 | 0.37399 | AP3S1 | turquoise | -0.188294136 | 0.346931084 | ZNF461 | turquoise | -0.046325 | 0.822199 |
| PPA1 | turquoise | -0.181824 | 0.37402 | ZFYVE27 | turquoise | -0.18773972 | 0.348378704 | RAB3GAP1 | turquoise | -0.046322 | 0.822212 |
| CAND1 | turquoise | -0.181624 | 0.374555 | COPB1 | turquoise | -0.187388658 | 0.349297212 | HOMER1 | turquoise | -0.046177 | 0.822761 |
| GTPBP10 | turquoise | -0.181075 | 0.376025 | C11orf80 | turquoise | -0.187278369 | 0.349586066 | AKAP10 | turquoise | -0.045639 | 0.824794 |
| SMEK2 | turquoise | -0.181033 | 0.376138 | ECT2 | turquoise | -0.186810607 | 0.350812749 | ATG3 | turquoise | -0.045494 | 0.825342 |
| ASAP1 | turquoise | 0.180877 | 0.376555 | ARIH2 | turquoise | -0.186237623 | 0.352318852 | C6orf70 | turquoise | -0.045427 | 0.825594 |
| PPP4R2 | turquoise | -0.180877 | 0.376556 | TM7SF3 | turquoise | -0.186042447 | 0.352832753 | ZNF213 | turquoise | -0.045167 | 0.826579 |
| CCDC14 | turquoise | -0.180658 | 0.377143 | OFD1 | turquoise | -0.185498205 | 0.3542681 | U2SURP | turquoise | -0.044811 | 0.827924 |
| PSKH1 | turquoise | -0.180351 | 0.377969 | TAF2 | turquoise | -0.185449774 | 0.354395994 | GATAD2B | turquoise | -0.044747 | 0.828168 |
| SLC4A8 | turquoise | -0.180326 | 0.378036 | TMEM116 | turquoise | -0.185439117 | 0.354424142 | ANKS1A | turquoise | -0.044124 | 0.830525 |
| BTBD3 | turquoise | -0.180293 | 0.378124 | ZNF747 | turquoise | -0.185392418 | 0.354547495 | NDUFA4 | turquoise | -0.043987 | 0.831043 |
| TECR | turquoise | -0.179843 | 0.379335 | PCGF3 | turquoise | -0.185318252 | 0.354743455 | PCM1 | turquoise | -0.04342 | 0.833192 |
| MOSPD2 | turquoise | -0.179651 | 0.379853 | CENPQ | turquoise | -0.1852162 | 0.355013198 | COG7 | turquoise | -0.043083 | 0.834471 |
| EEF1B2 | turquoise | -0.178905 | 0.381866 | EFHA1 | turquoise | -0.184830136 | 0.356034742 | UAP1L1 | turquoise | -0.042929 | 0.835054 |
| GPSM2 | turquoise | -0.178716 | 0.382378 | TOMM70A | turquoise | -0.184739969 | 0.356273577 | ZXDB | turquoise | -0.04284 | 0.835392 |

**Table 6. Gene module membership and correlation with growth rate (48/72h), with adjusted p values for each module**

| | | | | | | | | | | | |
|---|---|---|---|---|---|---|---|---|---|---|---|
| BOLA3 | turquoise | -0.178544 | 0.382846 | APMAP | turquoise | -0.184691592 | 0.356401759 | PHB | turquoise | -0.04266 | 0.836071 |
| MFN1 | turquoise | -0.178479 | 0.38302 | SNHG16 | turquoise | -0.184632052 | 0.356559557 | PDLIM2 | turquoise | -0.042137 | 0.838054 |
| SERPINF2 | turquoise | -0.178268 | 0.383593 | HERC2P9 | turquoise | -0.183226733 | 0.360296001 | TRNT1 | turquoise | -0.041765 | 0.839466 |
| LSG1 | turquoise | -0.177997 | 0.384328 | ZNF641 | turquoise | -0.182805069 | 0.361421599 | MAK16 | turquoise | -0.041483 | 0.840537 |
| MAN2B2 | turquoise | -0.177909 | 0.384568 | SMARCA2 | turquoise | -0.182752982 | 0.361560784 | IMPA1 | turquoise | -0.041289 | 0.841272 |
| PRKAG1 | turquoise | -0.177377 | 0.386013 | MICU1 | turquoise | -0.182510498 | 0.362209158 | PSMD12 | turquoise | -0.041234 | 0.841482 |
| NPHP4 | turquoise | -0.177047 | 0.386912 | AGA | turquoise | -0.182471323 | 0.362313972 | ZNF280C | turquoise | -0.041008 | 0.842342 |
| NADKD1 | turquoise | -0.176673 | 0.387931 | FAM8A1 | turquoise | -0.182292306 | 0.362793161 | MYSM1 | turquoise | -0.040994 | 0.842395 |
| POT1 | turquoise | -0.176231 | 0.38914 | YIPF2 | turquoise | -0.182270639 | 0.362851185 | ESF1 | turquoise | -0.040564 | 0.844025 |
| HELZ | turquoise | -0.176057 | 0.389614 | MPHOSPH9 | turquoise | -0.181162786 | 0.365825223 | HNRNPR | turquoise | -0.040371 | 0.844759 |
| ZNF692 | turquoise | -0.175834 | 0.390225 | EIF5 | turquoise | -0.17996317 | 0.369061641 | NRP1 | turquoise | -0.040114 | 0.845736 |
| -UPK3BP1-P | turquoise | -0.175545 | 0.391016 | CTSA | turquoise | -0.179444873 | 0.370465097 | UBQLN1 | turquoise | -0.039715 | 0.847254 |
| EFHA1 | turquoise | -0.17495 | 0.39265 | ROCK1 | turquoise | -0.179424933 | 0.370519154 | FMO4 | turquoise | -0.039584 | 0.847753 |
| CASP8AP2 | turquoise | -0.174865 | 0.392884 | MAP3K7 | turquoise | -0.179274121 | 0.370928144 | MAGOH | turquoise | -0.039572 | 0.847797 |
| CYP20A1 | turquoise | -0.174604 | 0.393603 | IMPA1 | turquoise | -0.179028804 | 0.371593985 | ZFAND6 | turquoise | -0.039443 | 0.848288 |
| SUSD3 | turquoise | -0.174531 | 0.393803 | TRAF6 | turquoise | -0.178860525 | 0.372051134 | MAP4 | turquoise | -0.038832 | 0.850611 |
| ARL2BP | turquoise | -0.174469 | 0.393974 | AIM2 | turquoise | -0.17879744 | 0.372222594 | SNX20 | turquoise | -0.038807 | 0.850706 |
| MEIS3P1 | turquoise | 0.17435 | 0.394301 | AUP1 | turquoise | -0.17869042 | 0.372513573 | PLP2 | turquoise | -0.038603 | 0.851483 |
| CBWD5 | turquoise | -0.174238 | 0.394611 | SYVN1 | turquoise | -0.178567871 | 0.372846934 | RFWD2 | turquoise | -0.038172 | 0.853123 |
| DTD2 | turquoise | -0.173703 | 0.396084 | GRAMD1A | turquoise | -0.17772718 | 0.375138489 | PCBP1 | turquoise | -0.038115 | 0.853341 |
| EHD1 | turquoise | 0.173694 | 0.396111 | CCDC14 | turquoise | -0.177543572 | 0.375640053 | MYO9B | turquoise | -0.037984 | 0.85384 |
| HACE1 | turquoise | -0.172487 | 0.399453 | GLTSCR1 | turquoise | -0.176985294 | 0.37716749 | ZNF322 | turquoise | -0.037908 | 0.854129 |
| EIF3B | turquoise | -0.172393 | 0.399712 | VAPB | turquoise | -0.176851398 | 0.37753436 | SAP130 | turquoise | -0.037783 | 0.854606 |
| DNAJC5 | turquoise | -0.172316 | 0.399925 | NKIRAS1 | turquoise | -0.176775552 | 0.377742267 | CCND3 | turquoise | -0.037686 | 0.854975 |
| LYRM1 | turquoise | -0.172313 | 0.399934 | TACC3 | turquoise | -0.176081683 | 0.37964735 | MTF1 | turquoise | -0.037628 | 0.855194 |
| DYNC2LI1 | turquoise | -0.17229 | 0.399998 | SMC2 | turquoise | -0.17600089 | 0.379869537 | ZNF516 | turquoise | -0.037613 | 0.855253 |
| SMNDC1 | turquoise | -0.17145 | 0.402335 | PHB | turquoise | -0.175695426 | 0.380710256 | CHN2 | turquoise | -0.037594 | 0.855323 |
| CUX1 | turquoise | 0.171258 | 0.40287 | RNF41 | turquoise | -0.175389289 | 0.3815539 | CAMSAP2 | turquoise | -0.03759 | 0.855339 |
| LYRM5 | turquoise | -0.170596 | 0.404718 | PTOV1 | turquoise | -0.175053004 | 0.382481868 | RIF1 | turquoise | -0.037314 | 0.856393 |
| IFNAR2 | turquoise | -0.170468 | 0.405076 | QSOX2 | turquoise | -0.174486671 | 0.384047575 | INTS10 | turquoise | -0.037295 | 0.856465 |
| RSRC1 | turquoise | -0.170343 | 0.405425 | TTC7A | turquoise | -0.173802822 | 0.385943069 | DDX50 | turquoise | -0.037193 | 0.856852 |
| PPP3CB | turquoise | -0.170147 | 0.405974 | COG6 | turquoise | -0.173192273 | 0.387639913 | COX15 | turquoise | -0.037005 | 0.857568 |
| OC1005056 | turquoise | -0.170068 | 0.406194 | PRKAR2A | turquoise | -0.173037347 | 0.388071162 | MAPK6 | turquoise | -0.036908 | 0.85794 |
| COG2 | turquoise | -0.16973 | 0.407142 | C9orf41 | turquoise | -0.172889725 | 0.388482334 | EIF2S3 | turquoise | -0.036626 | 0.859015 |
| RNU12 | turquoise | -0.169714 | 0.407188 | RCAN1 | turquoise | -0.172778027 | 0.388793614 | RGPD8 | turquoise | -0.036594 | 0.859135 |
| AKAP9 | turquoise | -0.169677 | 0.407292 | ATAD3B | turquoise | -0.172341693 | 0.390010952 | ZNF248 | turquoise | -0.036543 | 0.859328 |
| PLA2G16 | turquoise | 0.169503 | 0.407779 | RAB11FIP4 | turquoise | -0.170833762 | 0.394234667 | STXBP1 | turquoise | -0.036408 | 0.859844 |
| PRKRIR | turquoise | -0.169432 | 0.407978 | B3GALT6 | turquoise | -0.170741517 | 0.394493885 | FKBP5 | turquoise | -0.036043 | 0.861234 |
| PTPLAD1 | turquoise | -0.169429 | 0.407986 | RAB5A | turquoise | -0.170514122 | 0.395133303 | TTC14 | turquoise | -0.035698 | 0.86255 |
| IFNGR1 | turquoise | -0.16932 | 0.408294 | PDPR | turquoise | -0.170029149 | 0.396498975 | NUDT16L1 | turquoise | -0.03554 | 0.863155 |
| UBXN4 | turquoise | -0.168914 | 0.409436 | SNX13 | turquoise | -0.169636187 | 0.397607502 | TRAM1 | turquoise | -0.035277 | 0.864158 |
| MEMO1 | turquoise | 0.168883 | 0.409523 | TSR1 | turquoise | -0.169389777 | 0.398303509 | CWF19L1 | turquoise | -0.034928 | 0.865488 |
| TTC28 | turquoise | 0.168507 | 0.41058 | CCNC | turquoise | -0.16902175 | 0.399344312 | AKTIP | turquoise | -0.03472 | 0.866282 |
| ALAS1 | turquoise | -0.168415 | 0.410841 | CUL3 | turquoise | -0.168384031 | 0.401151451 | MCCC2 | turquoise | -0.034218 | 0.8682 |
| CUL7 | turquoise | -0.168386 | 0.410921 | MVP | turquoise | -0.168201715 | 0.401668937 | MTFR1 | turquoise | -0.033801 | 0.869793 |
| AGPS | turquoise | -0.16829 | 0.411192 | THAP4 | turquoise | -0.167217003 | 0.404470432 | IQSEC3 | turquoise | -0.033756 | 0.869964 |
| NBPF24 | turquoise | 0.168231 | 0.41136 | CRIPT | turquoise | -0.167085811 | 0.404844496 | PPAT | turquoise | -0.033645 | 0.87039 |
| CBX3 | turquoise | -0.168127 | 0.411653 | NMD3 | turquoise | -0.166785967 | 0.405700167 | GTPBP10 | turquoise | -0.033447 | 0.871145 |
| MED12 | turquoise | -0.168034 | 0.411914 | PUS10 | turquoise | -0.166566106 | 0.40632823 | ALKBH4 | turquoise | -0.033182 | 0.872159 |
| APPL2 | turquoise | -0.168033 | 0.411918 | C8orf40 | turquoise | -0.166290046 | 0.407117606 | FBXL4 | turquoise | -0.032035 | 0.876543 |
| STXBP5 | turquoise | 0.167893 | 0.412312 | WASF2 | turquoise | -0.165488255 | 0.409415136 | APOC2 | turquoise | -0.032004 | 0.876661 |
| LIN37 | turquoise | 0.167511 | 0.413394 | NIT2 | turquoise | -0.165237845 | 0.410134165 | DNAJB14 | turquoise | -0.031901 | 0.877057 |
| ZC3H6 | turquoise | -0.167341 | 0.413873 | FAM60A | turquoise | -0.164792692 | 0.411414121 | PCID2 | turquoise | -0.03171 | 0.877786 |
| USP40 | turquoise | -0.167339 | 0.413878 | PTPN4 | turquoise | -0.164777391 | 0.411458154 | C14orf166 | turquoise | -0.031471 | 0.8787 |
| UBXN11 | turquoise | 0.166814 | 0.415367 | TNFSF13B | turquoise | -0.164703022 | 0.411672218 | DENND6A | turquoise | -0.031267 | 0.879483 |
| FGFR1OP2 | turquoise | 0.166412 | 0.416509 | SMN2 | turquoise | -0.164250144 | 0.41297712 | FAM18B1 | turquoise | -0.031261 | 0.879503 |

**Table 6. Gene module membership and correlation with growth rate (48/72h), with adjusted p values for each module**

| | | | |
|---|---|---|---|
| SS18L1 | turquoise | 0.166195 | 0.417124 |
| ARL5A | turquoise | -0.166078 | 0.417457 |
| AHCYL2 | turquoise | -0.166012 | 0.417644 |
| PPIA | turquoise | -0.165997 | 0.417689 |
| TRMT1L | turquoise | -0.165991 | 0.417705 |
| ARRDC4 | turquoise | -0.165868 | 0.418055 |
| ATL2 | turquoise | -0.165838 | 0.418139 |
| TAF1D | turquoise | -0.165806 | 0.41823 |
| C3orf17 | turquoise | -0.165516 | 0.419057 |
| HEXB | turquoise | -0.165429 | 0.419305 |
| STIM2 | turquoise | -0.165298 | 0.41968 |
| TBC1D12 | turquoise | -0.165109 | 0.420219 |
| TXNDC15 | turquoise | -0.164722 | 0.421323 |
| CLK1 | turquoise | -0.164606 | 0.421656 |
| AZIN1 | turquoise | -0.164529 | 0.421874 |
| CEP57L1 | turquoise | -0.164488 | 0.421992 |
| CUL5 | turquoise | -0.16423 | 0.42273 |
| MLKL | turquoise | -0.164208 | 0.422795 |
| DHX8 | turquoise | 0.163989 | 0.423421 |
| VPS45 | turquoise | -0.163889 | 0.423707 |
| ZNF664 | turquoise | 0.163541 | 0.424705 |
| ZNF518A | turquoise | -0.163246 | 0.425553 |
| NDUFA4 | turquoise | -0.163152 | 0.425825 |
| SLC7A7 | turquoise | -0.163123 | 0.425908 |
| RPS11 | turquoise | -0.163084 | 0.42602 |
| RNFT1 | turquoise | -0.162988 | 0.426296 |
| NAA20 | turquoise | -0.162512 | 0.427667 |
| CD47 | turquoise | -0.162366 | 0.428086 |
| BCAP29 | turquoise | -0.162255 | 0.428408 |
| SULT1A4 | turquoise | -0.162205 | 0.428551 |
| CALCOCO1 | turquoise | 0.1622 | 0.428566 |
| ABCC10 | turquoise | -0.162146 | 0.428723 |
| WHSC2 | turquoise | -0.162115 | 0.428811 |
| DEK | turquoise | -0.16181 | 0.429691 |
| TBP | turquoise | -0.161775 | 0.429792 |
| HLTF | turquoise | -0.161619 | 0.430244 |
| KLHL7 | turquoise | -0.161141 | 0.431627 |
| PTPN18 | turquoise | -0.161074 | 0.431821 |
| FURIN | turquoise | -0.161037 | 0.431927 |
| GOSR2 | turquoise | -0.160741 | 0.432786 |
| CEBPD | turquoise | -0.160458 | 0.433608 |
| ELAC2 | turquoise | -0.16045 | 0.433631 |
| ARHGAP30 | turquoise | -0.160366 | 0.433875 |
| KRCC1 | turquoise | -0.160027 | 0.43486 |
| MINA | turquoise | -0.159756 | 0.435648 |
| SCYL2 | turquoise | -0.159651 | 0.435955 |
| RAD50 | turquoise | -0.159606 | 0.436085 |
| SRF | turquoise | -0.1595 | 0.436393 |
| RHEB | turquoise | -0.159472 | 0.436475 |
| ACER3 | turquoise | -0.158989 | 0.437884 |
| C14orf93 | turquoise | -0.158739 | 0.438613 |
| NFKBIE | turquoise | -0.158623 | 0.438953 |
| SIGMAR1 | turquoise | -0.158452 | 0.439452 |
| STRAP | turquoise | -0.158363 | 0.439712 |
| HMG20B | turquoise | -0.157886 | 0.441111 |
| RHOQ | turquoise | -0.157819 | 0.441307 |

| | | | |
|---|---|---|---|
| ING2 | turquoise | -0.164182731 | 0.413171557 |
| CCNB1IP1 | turquoise | -0.16406554 | 0.413509688 |
| API5 | turquoise | -0.163679843 | 0.414623622 |
| OGFRL1 | turquoise | -0.163454255 | 0.415275917 |
| UQCRC1 | turquoise | -0.163176528 | 0.416079754 |
| SCAF8 | turquoise | -0.16295481 | 0.416722098 |
| DMTF1 | turquoise | -0.162953119 | 0.416726999 |
| NMI | turquoise | -0.161711985 | 0.420332869 |
| GINM1 | turquoise | -0.161287344 | 0.421570517 |
| ZC3H4 | turquoise | -0.160409293 | 0.424136012 |
| BTN3A1 | turquoise | -0.160187308 | 0.424785966 |
| ZNF414 | turquoise | -0.159523091 | 0.426733988 |
| SOCS7 | turquoise | -0.159392284 | 0.427118196 |
| STIM2 | turquoise | -0.158700959 | 0.429151901 |
| MRPS10 | turquoise | -0.15858707 | 0.42948744 |
| ZNHIT6 | turquoise | -0.158536214 | 0.429637318 |
| ATAD3A | turquoise | -0.157551653 | 0.432544531 |
| PWP2 | turquoise | -0.157518391 | 0.432642935 |
| ARPC4 | turquoise | -0.156548741 | 0.435516889 |
| RNF214 | turquoise | -0.156406987 | 0.435937898 |
| PRPF6 | turquoise | -0.155800827 | 0.437740677 |
| ECH1 | turquoise | -0.155362788 | 0.439045952 |
| NPLOC4 | turquoise | -0.155138538 | 0.439714985 |
| IL1B | turquoise | -0.154720889 | 0.440962473 |
| CTNNBIP1 | turquoise | -0.153348945 | 0.445073733 |
| POLG2 | turquoise | -0.15308915 | 0.445854557 |
| ZNF136 | turquoise | -0.152395661 | 0.447942443 |
| CHORDC1 | turquoise | -0.151979036 | 0.449199278 |
| ACE | turquoise | -0.151930444 | 0.449345987 |
| LTN1 | turquoise | -0.151868236 | 0.449533845 |
| FAM63B | turquoise | -0.151616097 | 0.450295685 |
| TXNIP | turquoise | -0.151580419 | 0.450403541 |
| ZYX | turquoise | -0.151373631 | 0.451028946 |
| NAE1 | turquoise | -0.1508936 | 0.452482515 |
| PI4K2B | turquoise | -0.150555164 | 0.453508814 |
| APLF | turquoise | -0.150332538 | 0.454184596 |
| CLIC4 | turquoise | -0.149843431 | 0.455671148 |
| EIF2B5 | turquoise | -0.149537179 | 0.456603252 |
| MLF1 | turquoise | -0.149273792 | 0.457405696 |
| CRNKL1 | turquoise | -0.149201917 | 0.457624802 |
| RYBP | turquoise | -0.148697167 | 0.459165051 |
| CCDC50 | turquoise | -0.148278071 | 0.460445997 |
| CIC | turquoise | -0.147691989 | 0.462240467 |
| TMEM126E | turquoise | -0.146667583 | 0.465385777 |
| NR1H2 | turquoise | -0.146457888 | 0.466030994 |
| SCAF1 | turquoise | -0.146400145 | 0.466208746 |
| SLC12A7 | turquoise | -0.146333674 | 0.466413411 |
| SMURF2 | turquoise | -0.145575691 | 0.468750543 |
| EDC4 | turquoise | -0.145489994 | 0.469015162 |
| UAP1L1 | turquoise | -0.14508948 | 0.470252903 |
| ATAD2B | turquoise | -0.145003684 | 0.470518266 |
| SMEK1 | turquoise | -0.144910106 | 0.470807786 |
| TRPM7 | turquoise | -0.144405987 | 0.472369063 |
| NOC3L | turquoise | -0.144193627 | 0.473027552 |
| DEK | turquoise | -0.143644559 | 0.474732301 |
| DENND4B | turquoise | -0.143618713 | 0.474812627 |

| | | | |
|---|---|---|---|
| CD276 | turquoise | -0.031108 | 0.880089 |
| EIF2S2 | turquoise | -0.031023 | 0.880417 |
| IMPA2 | turquoise | -0.030957 | 0.880667 |
| NLN | turquoise | -0.030669 | 0.881771 |
| ACBD5 | turquoise | -0.030304 | 0.883168 |
| ACVR1B | turquoise | -0.029845 | 0.884925 |
| COPS2 | turquoise | -0.029746 | 0.885304 |
| SF3B1 | turquoise | -0.029661 | 0.88563 |
| FOPNL | turquoise | -0.029518 | 0.886178 |
| TRMT1L | turquoise | -0.029388 | 0.886675 |
| POLA2 | turquoise | -0.029206 | 0.887375 |
| TMEM39A | turquoise | -0.029143 | 0.887615 |
| TBL3 | turquoise | -0.029054 | 0.887955 |
| RRP1 | turquoise | -0.028845 | 0.888759 |
| CINP | turquoise | -0.02873 | 0.889199 |
| RNF4 | turquoise | -0.028512 | 0.890035 |
| CSNK1E | turquoise | -0.028284 | 0.89091 |
| TXNL1 | turquoise | -0.028003 | 0.891986 |
| JAK2 | turquoise | -0.027558 | 0.893693 |
| PPP2R5C | turquoise | -0.027264 | 0.89482 |
| TWF1 | turquoise | -0.027105 | 0.895429 |
| PAAF1 | turquoise | -0.027051 | 0.895637 |
| SS18L1 | turquoise | -0.026803 | 0.896591 |
| MRPL33 | turquoise | -0.026772 | 0.89671 |
| HECTD3 | turquoise | -0.026523 | 0.897666 |
| SLC45A3 | turquoise | -0.026333 | 0.898394 |
| MLLT4 | turquoise | -0.026209 | 0.898869 |
| C1orf50 | turquoise | -0.026188 | 0.898949 |
| CMTM7 | turquoise | -0.026026 | 0.899572 |
| PLEKHM1P | turquoise | -0.025742 | 0.900665 |
| FKBP3 | turquoise | -0.025711 | 0.900782 |
| LRIG2 | turquoise | -0.025508 | 0.901562 |
| SYNGR2 | turquoise | -0.025306 | 0.902337 |
| HPS1 | turquoise | -0.025167 | 0.902871 |
| TLN1 | turquoise | -0.024304 | 0.906186 |
| PDCD7 | turquoise | -0.024153 | 0.906766 |
| DDHD1 | turquoise | -0.024091 | 0.907005 |
| GNAI3 | turquoise | -0.023878 | 0.907826 |
| PRCC | turquoise | -0.0238 | 0.908124 |
| SNORA21 | turquoise | -0.023424 | 0.90957 |
| YTHDF1 | turquoise | -0.023122 | 0.910733 |
| NETO2 | turquoise | -0.023114 | 0.910764 |
| SNAPC1 | turquoise | -0.023034 | 0.911071 |
| FMNL1 | turquoise | -0.022706 | 0.91233 |
| AK1 | turquoise | -0.02257 | 0.912853 |
| BCAR3 | turquoise | -0.022324 | 0.9138 |
| STX18 | turquoise | -0.022056 | 0.914831 |
| PHTF2 | turquoise | -0.021836 | 0.91568 |
| CLINT1 | turquoise | -0.021788 | 0.915863 |
| FAM208B | turquoise | -0.021298 | 0.917748 |
| PTK2 | turquoise | -0.021194 | 0.91815 |
| AHCYL2 | turquoise | -0.021085 | 0.918569 |
| EXOC7 | turquoise | -0.020327 | 0.921487 |
| CCDC132 | turquoise | -0.020062 | 0.922509 |
| TOR1A | turquoise | -0.019893 | 0.923158 |
| TMEM164 | turquoise | -0.019868 | 0.923256 |

**Table 6. Gene module membership and correlation with growth rate (48/72h), with adjusted p values for each module**

| | | | |
|---|---|---|---|
| PAPSS1 | turquoise | -0.157773 | 0.44144 |
| BTN3A3 | turquoise | -0.1577 | 0.441654 |
| ZFYVE27 | turquoise | 0.15763 | 0.441859 |
| EIF3E | turquoise | -0.15753 | 0.442153 |
| LPGAT1 | turquoise | -0.157505 | 0.442226 |
| CHKA | turquoise | 0.157402 | 0.442528 |
| PUS7 | turquoise | -0.157347 | 0.442692 |
| FCHO2 | turquoise | 0.156948 | 0.443862 |
| HSD17B4 | turquoise | -0.156564 | 0.444993 |
| DCAF11 | turquoise | -0.156137 | 0.446252 |
| FAM125B | turquoise | -0.156054 | 0.446497 |
| EIF3G | turquoise | -0.155985 | 0.4467 |
| DGCR14 | turquoise | 0.155941 | 0.446828 |
| SLC25A39 | turquoise | -0.155736 | 0.447434 |
| PPP2R2A | turquoise | 0.155657 | 0.447668 |
| EOGT | turquoise | 0.155452 | 0.448272 |
| C18orf21 | turquoise | -0.15544 | 0.44831 |
| ME1 | turquoise | -0.155283 | 0.448773 |
| EXOC1 | turquoise | -0.155039 | 0.449496 |
| C1orf43 | turquoise | -0.15499 | 0.44964 |
| DROSHA | turquoise | -0.154967 | 0.449707 |
| KIAA1107 | turquoise | -0.15477 | 0.450291 |
| PRAM1 | turquoise | -0.154466 | 0.451192 |
| KDELR3 | turquoise | -0.154433 | 0.45129 |
| DDX50 | turquoise | -0.154331 | 0.451593 |
| PDXK | turquoise | -0.154275 | 0.451759 |
| RRAGB | turquoise | -0.154206 | 0.451965 |
| TIPARP | turquoise | -0.154186 | 0.452023 |
| PLCB3 | turquoise | 0.153992 | 0.452599 |
| EGLN1 | turquoise | -0.153398 | 0.454365 |
| CCNC | turquoise | -0.153167 | 0.455053 |
| EIF2S2 | turquoise | -0.152839 | 0.456031 |
| EHF | turquoise | -0.152283 | 0.457691 |
| MRPS15 | turquoise | -0.152078 | 0.458304 |
| KIAA1551 | turquoise | -0.151904 | 0.458825 |
| PHAX | turquoise | -0.151585 | 0.459779 |
| MRPL47 | turquoise | -0.15149 | 0.460065 |
| RNF20 | turquoise | -0.151462 | 0.460148 |
| PFAS | turquoise | -0.151037 | 0.461424 |
| OGFRL1 | turquoise | -0.150939 | 0.461718 |
| FAM70B | turquoise | -0.15086 | 0.461954 |
| GTF2H3 | turquoise | -0.150486 | 0.463079 |
| TRIP4 | turquoise | -0.150412 | 0.4633 |
| MAP1S | turquoise | -0.150193 | 0.46396 |
| ZNF569 | turquoise | -0.150084 | 0.464288 |
| PIK3R1 | turquoise | -0.149993 | 0.464562 |
| PYGL | turquoise | -0.149963 | 0.464653 |
| RARS | turquoise | -0.14957 | 0.465837 |
| GNPDA1 | turquoise | -0.14945 | 0.466199 |
| ZDHHC8 | turquoise | -0.148816 | 0.468112 |
| HERC6 | turquoise | -0.148794 | 0.468181 |
| SNX14 | turquoise | -0.148675 | 0.46854 |
| TNFRSF14 | turquoise | -0.148622 | 0.468699 |
| N4BP2L2 | turquoise | -0.148541 | 0.468945 |
| TOMM70A | turquoise | -0.148525 | 0.468995 |
| RPLP0 | turquoise | -0.148384 | 0.469421 |
| XPR1 | turquoise | -0.142747662 | 0.477523787 |
| CDK18 | turquoise | -0.142180722 | 0.47929266 |
| TBRG4 | turquoise | -0.14203471 | 0.479748763 |
| HERC1 | turquoise | -0.141742072 | 0.480663561 |
| SPOPL | turquoise | -0.141332515 | 0.481945345 |
| CCDC71L | turquoise | -0.141314913 | 0.482000474 |
| FBXO30 | turquoise | -0.140994741 | 0.483003785 |
| RBL1 | turquoise | -0.140652939 | 0.48407605 |
| GGNBP2 | turquoise | -0.140507416 | 0.484532937 |
| EVI5 | turquoise | -0.139676539 | 0.487145767 |
| FAM13B | turquoise | -0.13945242 | 0.487851768 |
| GCA | turquoise | -0.139414193 | 0.487972241 |
| TIPARP | turquoise | -0.139022282 | 0.489208203 |
| ITGB2 | turquoise | -0.137711531 | 0.493353363 |
| STK4 | turquoise | -0.137280967 | 0.494718832 |
| AKAP9 | turquoise | -0.137238834 | 0.494852555 |
| FAM111A | turquoise | -0.13624628 | 0.498007925 |
| ZNF280C | turquoise | -0.136127045 | 0.498387653 |
| TRIOBP | turquoise | -0.136102573 | 0.498465605 |
| IGBP1 | turquoise | -0.136054949 | 0.498617327 |
| METTL21D | turquoise | -0.135751105 | 0.499585854 |
| GNAI2 | turquoise | -0.135717862 | 0.499691875 |
| PCCB | turquoise | -0.135642779 | 0.499931377 |
| NCOA5 | turquoise | -0.135433201 | 0.5006002 |
| NOC4L | turquoise | -0.135088 | 0.501702806 |
| FASTK | turquoise | -0.135076805 | 0.501738582 |
| GUF1 | turquoise | -0.134922498 | 0.502231863 |
| DLG4 | turquoise | -0.13469364 | 0.502963904 |
| MAZ | turquoise | -0.134570618 | 0.503357628 |
| ADSS | turquoise | -0.134384262 | 0.503954341 |
| HDAC2 | turquoise | -0.134041945 | 0.505051356 |
| SLCO2B1 | turquoise | -0.133868384 | 0.505608009 |
| BCKDHB | turquoise | -0.133784039 | 0.505878637 |
| ELP4 | turquoise | -0.13374369 | 0.506008123 |
| AGAP3 | turquoise | -0.133708994 | 0.506119483 |
| SGOL2 | turquoise | -0.133559215 | 0.506600349 |
| HIST1H4B | turquoise | -0.133353433 | 0.507261379 |
| DRG2 | turquoise | -0.133270987 | 0.507526336 |
| VDAC2 | turquoise | -0.133000723 | 0.508395372 |
| NDRG3 | turquoise | -0.132899924 | 0.508719678 |
| TAF9 | turquoise | -0.132672904 | 0.509450455 |
| TRMT11 | turquoise | -0.13197149 | 0.511711558 |
| CEP70 | turquoise | -0.131911172 | 0.511906231 |
| CCDC82 | turquoise | -0.131208433 | 0.514176944 |
| SFPQ | turquoise | -0.131015871 | 0.514800015 |
| FAM200B | turquoise | -0.13099192 | 0.514877538 |
| NCOA1 | turquoise | -0.13070425 | 0.515809101 |
| TSNAX | turquoise | -0.130504786 | 0.516455508 |
| MRPS33 | turquoise | -0.129865437 | 0.518530112 |
| ARL5A | turquoise | -0.12985451 | 0.518565603 |
| RASA2 | turquoise | -0.12937147 | 0.520135733 |
| FAM3A | turquoise | -0.12921848 | 0.520633507 |
| GNPDA1 | turquoise | -0.128536629 | 0.522854805 |
| TPM4 | turquoise | -0.128147232 | 0.524125408 |
| GSN | turquoise | -0.128067168 | 0.524386841 |
| TRAF3IP2 | turquoise | -0.127527002 | 0.526152282 |
| GGNBP2 | turquoise | -0.018103 | 0.930054 |
| BCR | turquoise | -0.018102 | 0.93006 |
| POLG2 | turquoise | -0.017909 | 0.930803 |
| ACVR2A | turquoise | -0.017804 | 0.931209 |
| SDHAF2 | turquoise | -0.017768 | 0.931347 |
| RCN2 | turquoise | -0.017478 | 0.932464 |
| CAP1 | turquoise | -0.017343 | 0.932984 |
| UNC50 | turquoise | -0.01705 | 0.934117 |
| SAP30 | turquoise | -0.016963 | 0.934453 |
| ZNF436 | turquoise | -0.016818 | 0.935009 |
| ATF1 | turquoise | -0.01515 | 0.941445 |
| RNF146 | turquoise | -0.015117 | 0.941573 |
| METTL21D | turquoise | -0.014967 | 0.942148 |
| DHX36 | turquoise | -0.014481 | 0.944024 |
| GPR89A | turquoise | -0.014128 | 0.945388 |
| TM9SF3 | turquoise | -0.013764 | 0.946794 |
| STK3 | turquoise | -0.013468 | 0.947938 |
| N6AMT1 | turquoise | -0.013421 | 0.948116 |
| MFSD8 | turquoise | -0.01299 | 0.949783 |
| RPL23AP82 | turquoise | -0.012776 | 0.950608 |
| RPRD1A | turquoise | -0.01266 | 0.951057 |
| GPD1 | turquoise | -0.012643 | 0.951121 |
| CCDC88B | turquoise | -0.012209 | 0.952798 |
| ARRDC1 | turquoise | -0.012027 | 0.9535 |
| EBAG9 | turquoise | -0.011691 | 0.9548 |
| CLYBL | turquoise | -0.01139 | 0.955959 |
| EMILIN2 | turquoise | -0.010884 | 0.957916 |
| ELMOD2 | turquoise | -0.010005 | 0.96131 |
| SNRPC | turquoise | -0.009987 | 0.961381 |
| NDRG3 | turquoise | -0.009691 | 0.962527 |
| C7orf29 | turquoise | -0.009585 | 0.962935 |
| MAML1 | turquoise | -0.009115 | 0.96475 |
| CERS6 | turquoise | -0.008977 | 0.965286 |
| HDAC3 | turquoise | -0.008796 | 0.965983 |
| INO80C | turquoise | -0.008294 | 0.967924 |
| PPP6R1 | turquoise | -0.008164 | 0.968427 |
| LACE1 | turquoise | -0.008151 | 0.968478 |
| ATAD3B | turquoise | -0.008133 | 0.968546 |
| E2F6 | turquoise | -0.007502 | 0.970987 |
| CHKA | turquoise | -0.0075 | 0.970993 |
| ADAT1 | turquoise | -0.007243 | 0.971985 |
| EIF4E2 | turquoise | -0.00693 | 0.973196 |
| EEF1E1 | turquoise | -0.006912 | 0.973265 |
| ZYX | turquoise | -0.006661 | 0.974236 |
| GALNT7 | turquoise | -0.006586 | 0.974527 |
| SUZ12P1 | turquoise | -0.00615 | 0.976213 |
| CNOT7 | turquoise | -0.006049 | 0.976605 |
| ARHGAP21 | turquoise | -0.005829 | 0.977456 |
| ARID4A | turquoise | -0.005657 | 0.978121 |
| ERI2 | turquoise | -0.004804 | 0.981419 |
| NPHP4 | turquoise | -0.004746 | 0.981642 |
| GMIP | turquoise | -0.004685 | 0.981877 |
| GPR107 | turquoise | -0.004637 | 0.982065 |
| C14orf101 | turquoise | -0.004623 | 0.98212 |
| PTPN2 | turquoise | -0.004528 | 0.982484 |
| XRCC4 | turquoise | -0.004439 | 0.98283 |

**Table 6. Gene module membership and correlation with growth rate (48/72h), with adjusted p values for each module**

| | | | | | | | | | | | |
|---|---|---|---|---|---|---|---|---|---|---|---|
| AP3S1 | turquoise | -0.148257 | 0.469805 | PIN1 | turquoise | -0.127478358 | 0.526311408 | TBK1 | turquoise | -0.004249 | 0.983565 |
| GSN | turquoise | -0.148243 | 0.469848 | DNAJC2 | turquoise | -0.127178302 | 0.527293463 | RBM22 | turquoise | -0.00402 | 0.984452 |
| CNTRL | turquoise | 0.148167 | 0.470079 | PIGF | turquoise | -0.127067818 | 0.527655285 | ZNF268 | turquoise | -0.003909 | 0.984881 |
| SYNE3 | turquoise | -0.148158 | 0.470106 | GGA2 | turquoise | -0.126920001 | 0.528139558 | ZC3H13 | turquoise | -0.003814 | 0.985246 |
| DHRS1 | turquoise | -0.14802 | 0.470523 | POR | turquoise | -0.126636271 | 0.529069699 | COPS8 | turquoise | -0.003791 | 0.985334 |
| JUNB | turquoise | -0.147942 | 0.470759 | NUDT16L1 | turquoise | -0.126469716 | 0.529616072 | WRNIP1 | turquoise | -0.003575 | 0.986171 |
| MAPK6 | turquoise | 0.147269 | 0.472803 | RNF138 | turquoise | -0.126148784 | 0.530669631 | SLC25A32 | turquoise | -0.003377 | 0.986938 |
| JDP2 | turquoise | -0.147043 | 0.47349 | NHLRC3 | turquoise | -0.125794291 | 0.53183452 | MGAT4A | turquoise | -0.003327 | 0.987132 |
| VGLL4 | turquoise | -0.146925 | 0.473849 | ADPRH | turquoise | -0.125291618 | 0.533488425 | VAPB | turquoise | -0.003127 | 0.987902 |
| C14orf2 | turquoise | -0.146708 | 0.474509 | COPS4 | turquoise | -0.125250753 | 0.533622986 | NDUFV2 | turquoise | -0.002952 | 0.988582 |
| DTX4 | turquoise | -0.146637 | 0.474727 | ATP11C | turquoise | -0.12509206 | 0.534145688 | SIPA1L2 | turquoise | -0.002078 | 0.991963 |
| PHF13 | turquoise | -0.146599 | 0.474843 | CCNG1 | turquoise | -0.12473856 | 0.53531092 | HMG20B | turquoise | -0.001738 | 0.993278 |
| EEF1E1 | turquoise | -0.146128 | 0.476278 | EZH1 | turquoise | -0.124732346 | 0.535331411 | KATNA1 | turquoise | -0.001347 | 0.994791 |
| PTPLAD2 | turquoise | -0.146005 | 0.476652 | C6orf62 | turquoise | -0.123944608 | 0.53793237 | MTIF3 | turquoise | -0.001173 | 0.995461 |
| TRANK1 | turquoise | -0.145901 | 0.476971 | CASP8AP2 | turquoise | -0.123859908 | 0.538212387 | LGMN | turquoise | -0.000561 | 0.997831 |
| COMMD8 | turquoise | -0.145769 | 0.477373 | CASP6 | turquoise | -0.123397345 | 0.539742824 | LMBRD2 | turquoise | -0.000412 | 0.998406 |
| B9D2 | turquoise | -0.145732 | 0.477487 | CLDN23 | turquoise | -0.122912321 | 0.541349768 | MINPP1 | turquoise | 0.00016 | 0.999382 |
| SLC35B3 | turquoise | -0.145666 | 0.477688 | PFKP | turquoise | -0.122255444 | 0.54352965 | AHCY | turquoise | 0.000194 | 0.999248 |
| HOXA1 | turquoise | -0.145645 | 0.477753 | YY1 | turquoise | -0.122028975 | 0.544282148 | CES2 | turquoise | 0.000259 | 0.998998 |
| BBS4 | turquoise | -0.145438 | 0.478386 | DTD2 | turquoise | -0.121986985 | 0.544421723 | NARG2 | turquoise | 0.000287 | 0.99889 |
| SLC35A2 | turquoise | -0.145423 | 0.478432 | HIF1A | turquoise | -0.121792449 | 0.545068583 | CYP51A1 | turquoise | 0.000505 | 0.998046 |
| QPRT | turquoise | -0.14533 | 0.478716 | TRAPPC6B | turquoise | -0.120861553 | 0.548168886 | UBE2S | turquoise | 0.000852 | 0.996704 |
| NAA16 | turquoise | -0.145326 | 0.478726 | IRF2 | turquoise | -0.120704049 | 0.548694251 | TUBGCP3 | turquoise | 0.001075 | 0.99584 |
| PITPNC1 | turquoise | -0.145325 | 0.478729 | EMILIN2 | turquoise | -0.12004011 | 0.550911434 | TMEM59 | turquoise | 0.001351 | 0.994774 |
| PKNOX1 | turquoise | 0.145218 | 0.479058 | RNFT1 | turquoise | -0.119118354 | 0.553996416 | BBS4 | turquoise | 0.0018 | 0.993037 |
| NCBP2 | turquoise | -0.145149 | 0.479268 | OSGEPL1 | turquoise | -0.119012641 | 0.554350728 | SLC35B3 | turquoise | 0.001958 | 0.992425 |
| KIAA0319 | turquoise | 0.145051 | 0.479571 | BTF3 | turquoise | -0.118936036 | 0.554607547 | SRSF1 | turquoise | 0.002045 | 0.99209 |
| TTC3 | turquoise | -0.144959 | 0.479851 | HNRNPA1L2 | turquoise | -0.118805063 | 0.55504676 | CSNK2A1 | turquoise | 0.002132 | 0.991754 |
| TMEM106C | turquoise | -0.144908 | 0.480007 | NR2C1 | turquoise | -0.118511546 | 0.556031635 | HEATR3 | turquoise | 0.002796 | 0.989185 |
| SAP130 | turquoise | -0.144833 | 0.480236 | VDAC3 | turquoise | -0.11800453 | 0.557734776 | SLC38A7 | turquoise | 0.002895 | 0.988801 |
| NEU3 | turquoise | 0.144656 | 0.480778 | CLN5 | turquoise | -0.117553718 | 0.559251117 | AP4E1 | turquoise | 0.003057 | 0.988174 |
| C1orf198 | turquoise | -0.144656 | 0.480781 | USMG5 | turquoise | -0.117064933 | 0.560897304 | ZFAND4 | turquoise | 0.003085 | 0.988068 |
| ZDHHC4 | turquoise | -0.144458 | 0.481385 | CDK2AP1 | turquoise | -0.116735379 | 0.562008454 | TCF20 | turquoise | 0.003721 | 0.985607 |
| YEATS2 | turquoise | -0.144068 | 0.482583 | TNIP1 | turquoise | -0.116347446 | 0.563317715 | TIMP2 | turquoise | 0.004233 | 0.983627 |
| RAB42 | turquoise | -0.143778 | 0.483474 | UBR4 | turquoise | -0.116346423 | 0.563321171 | CUL4B | turquoise | 0.0043 | 0.983368 |
| STAM | turquoise | -0.143633 | 0.48392 | CTNND1 | turquoise | -0.116047827 | 0.564329864 | CAMK2G | turquoise | 0.004355 | 0.983155 |
| CCDC112 | turquoise | -0.143603 | 0.484014 | RHOBTB3 | turquoise | -0.11570431 | 0.565491317 | UNC119 | turquoise | 0.005056 | 0.980445 |
| GRN | turquoise | 0.143533 | 0.484228 | SNHG6 | turquoise | -0.115674659 | 0.565591619 | TMED7 | turquoise | 0.005232 | 0.979763 |
| MORC4 | turquoise | -0.143414 | 0.484595 | LOC550643 | turquoise | -0.115581059 | 0.5659083 | SFMBT2 | turquoise | 0.005284 | 0.979564 |
| YIPF2 | turquoise | -0.143359 | 0.484764 | SLC25A43 | turquoise | -0.115519727 | 0.566115849 | SMCHD1 | turquoise | 0.005606 | 0.978317 |
| ZMYM2 | turquoise | -0.142998 | 0.485876 | SNHG4 | turquoise | -0.115420965 | 0.566450134 | AIFM1 | turquoise | 0.005614 | 0.978286 |
| EMC2 | turquoise | -0.14229 | 0.488061 | CXCL16 | turquoise | -0.11458234 | 0.569292256 | NBEAL1 | turquoise | 0.005736 | 0.977812 |
| ADAMTS6 | turquoise | 0.142079 | 0.488712 | PSMA6 | turquoise | -0.114574876 | 0.569317579 | ZFYVE16 | turquoise | 0.006851 | 0.973504 |
| ZFAND6 | turquoise | -0.141596 | 0.490207 | ZNF548 | turquoise | -0.114566721 | 0.569345247 | NUDT21 | turquoise | 0.007029 | 0.972814 |
| CCDC159 | turquoise | 0.141515 | 0.490456 | KDELC2 | turquoise | -0.114256175 | 0.57039936 | TTC30A | turquoise | 0.007269 | 0.971885 |
| UCHL5 | turquoise | -0.141078 | 0.49181 | LRRC8D | turquoise | -0.11407664 | 0.571009167 | LRP10 | turquoise | 0.007373 | 0.971486 |
| DGKD | turquoise | 0.140876 | 0.492437 | RAD23A | turquoise | -0.114012555 | 0.571226906 | LILRA6 | turquoise | 0.007516 | 0.970932 |
| DYRK1A | turquoise | -0.140751 | 0.492827 | LYST | turquoise | -0.11370933 | 0.572257677 | FBXL20 | turquoise | 0.007744 | 0.970051 |
| UBE2B | turquoise | -0.140622 | 0.493227 | ZZEF1 | turquoise | -0.11308189 | 0.574393194 | NIPAL2 | turquoise | 0.007873 | 0.969552 |
| ADI1 | turquoise | -0.140529 | 0.493516 | PGAM5 | turquoise | -0.112847914 | 0.575190448 | TMEM62 | turquoise | 0.008221 | 0.968206 |
| C6orf48 | turquoise | -0.140119 | 0.49479 | CAPZB | turquoise | -0.112707129 | 0.575670393 | ANKRD27 | turquoise | 0.008753 | 0.966151 |
| EAF2 | turquoise | -0.140068 | 0.494948 | SMEK2 | turquoise | -0.112666242 | 0.575809815 | OSBPL10 | turquoise | 0.008798 | 0.965975 |
| ZNF83 | turquoise | -0.139758 | 0.495914 | TTYH3 | turquoise | -0.11261237 | 0.575993538 | CENPB | turquoise | 0.009417 | 0.963585 |
| KCTD10 | turquoise | 0.139405 | 0.497013 | ATG12 | turquoise | -0.11123981 | 0.580683148 | SDF2 | turquoise | 0.009548 | 0.963079 |
| RABGAP1 | turquoise | -0.139292 | 0.497365 | FAM35A | turquoise | -0.111040079 | 0.581366965 | MVP | turquoise | 0.009857 | 0.961882 |
| ACAT1 | turquoise | -0.139281 | 0.4974 | MBD4 | turquoise | -0.110560569 | 0.583010094 | CHID1 | turquoise | 0.010006 | 0.961307 |

**Table 6. Gene module membership and correlation with growth rate (48/72h), with adjusted p values for each module**

| | | | |
|---|---|---|---|
| PALB2 | turquoise | -0.138946 | 0.498445 |
| OST4 | turquoise | -0.138651 | 0.499367 |
| GFM1 | turquoise | -0.138314 | 0.50042 |
| YY1 | turquoise | -0.138089 | 0.501123 |
| ATP2C1 | turquoise | -0.137477 | 0.503042 |
| ITGAX | turquoise | 0.137396 | 0.503295 |
| TBCK | turquoise | -0.136917 | 0.504797 |
| OPHN1 | turquoise | 0.136791 | 0.505194 |
| ZMIZ2 | turquoise | -0.136418 | 0.506366 |
| TSR1 | turquoise | -0.136306 | 0.506719 |
| TPM4 | turquoise | -0.135719 | 0.508568 |
| NDNL2 | turquoise | -0.135587 | 0.508985 |
| GPSM3 | turquoise | -0.135244 | 0.510066 |
| NSUN3 | turquoise | -0.135195 | 0.510221 |
| FXR1 | turquoise | -0.134986 | 0.510883 |
| DPH2 | turquoise | -0.134635 | 0.511991 |
| ZFP106 | turquoise | -0.134445 | 0.512595 |
| CWF19L2 | turquoise | -0.134236 | 0.513256 |
| CDKN3 | turquoise | -0.1341 | 0.513686 |
| FHOD1 | turquoise | -0.13409 | 0.513717 |
| FCRL2 | turquoise | 0.134032 | 0.513901 |
| ACACA | turquoise | -0.133927 | 0.514234 |
| ACP6 | turquoise | -0.133718 | 0.514898 |
| FAM178A | turquoise | -0.133677 | 0.515027 |
| NFYB | turquoise | -0.13335 | 0.516066 |
| CHORDC1 | turquoise | 0.133167 | 0.516645 |
| COPB1 | turquoise | -0.132975 | 0.517255 |
| HNRNPM | turquoise | -0.132809 | 0.517785 |
| SGOL2 | turquoise | -0.132795 | 0.517829 |
| RNMT | turquoise | -0.132765 | 0.517924 |
| LPP | turquoise | -0.132746 | 0.517984 |
| FOPNL | turquoise | -0.132583 | 0.518504 |
| LRIF1 | turquoise | 0.13159 | 0.52167 |
| EIF3H | turquoise | -0.131561 | 0.521759 |
| MAN1C1 | turquoise | -0.130816 | 0.524142 |
| SH3BGRL3 | turquoise | -0.130377 | 0.525546 |
| XPR1 | turquoise | 0.130151 | 0.52627 |
| GM2A | turquoise | -0.130126 | 0.526352 |
| MCM4 | turquoise | -0.129946 | 0.526928 |
| DLG4 | turquoise | 0.12977 | 0.527492 |
| ASH1L | turquoise | 0.12955 | 0.528198 |
| CPQ | turquoise | -0.129377 | 0.528754 |
| MKI67 | turquoise | -0.129366 | 0.528789 |
| PSMG1 | turquoise | -0.128795 | 0.530627 |
| MRPL30 | turquoise | -0.128729 | 0.530837 |
| FKTN | turquoise | -0.128604 | 0.531241 |
| SDAD1 | turquoise | -0.128582 | 0.531314 |
| SPOPL | turquoise | -0.128561 | 0.531379 |
| ENDOD1 | turquoise | -0.128375 | 0.53198 |
| ATP5A1 | turquoise | -0.128286 | 0.532267 |
| PPIB | turquoise | -0.128264 | 0.532337 |
| RHOF | turquoise | -0.128119 | 0.532804 |
| SETX | turquoise | -0.128119 | 0.532805 |
| C20orf94 | turquoise | -0.127902 | 0.533505 |
| ZSCAN21 | turquoise | 0.127832 | 0.533728 |
| KCNK5 | turquoise | 0.127651 | 0.534313 |
| BLOC1S2 | turquoise | -0.110390756 | 0.583592474 |
| MRE11A | turquoise | -0.109778454 | 0.585694506 |
| CCL18 | turquoise | -0.109535109 | 0.586530823 |
| GABARAP | turquoise | -0.109326946 | 0.587246641 |
| PMM2 | turquoise | -0.109311184 | 0.587300857 |
| GLMN | turquoise | -0.109187288 | 0.587727099 |
| CLTB | turquoise | -0.109175636 | 0.587767193 |
| TXNDC15 | turquoise | -0.1091695 | 0.587788306 |
| ANXA11 | turquoise | -0.109103955 | 0.588013869 |
| PTGER4 | turquoise | -0.109003639 | 0.58835916 |
| HAT1 | turquoise | -0.108607625 | 0.589723111 |
| LRRC40 | turquoise | -0.108308061 | 0.590755778 |
| CCT6A | turquoise | -0.10824865 | 0.590960676 |
| ATP9A | turquoise | -0.107929629 | 0.592061436 |
| ARL2BP | turquoise | -0.107599846 | 0.593200259 |
| LRRC58 | turquoise | -0.107044054 | 0.595121673 |
| MIER2 | turquoise | -0.106675179 | 0.596398374 |
| SNORD94 | turquoise | -0.106637466 | 0.596528965 |
| RECK | turquoise | -0.10623796 | 0.59791313 |
| KITLG | turquoise | -0.106184781 | 0.598097481 |
| FAM188A | turquoise | -0.10572353 | 0.599697483 |
| MLLT4 | turquoise | -0.105316603 | 0.60111055 |
| KRAS | turquoise | -0.105127978 | 0.601766032 |
| DMXL1 | turquoise | -0.104751033 | 0.603076849 |
| PBX3 | turquoise | -0.104345722 | 0.604487648 |
| PVR | turquoise | -0.104060192 | 0.60548235 |
| PDCD7 | turquoise | -0.103712697 | 0.606693849 |
| DTWD1 | turquoise | -0.103665371 | 0.606858923 |
| GLIPR2 | turquoise | -0.103369631 | 0.607890901 |
| PPP1CB | turquoise | -0.103255995 | 0.608287628 |
| RAB28 | turquoise | -0.103111574 | 0.608791986 |
| MRPL16 | turquoise | -0.102808695 | 0.609850288 |
| IL28RA | turquoise | -0.10254784 | 0.610762367 |
| LYRM7 | turquoise | -0.101860749 | 0.613167496 |
| CIAPIN1 | turquoise | -0.101350293 | 0.614956858 |
| MYCBP2 | turquoise | -0.101274682 | 0.61522209 |
| ABCE1 | turquoise | -0.100776624 | 0.616970373 |
| PTBP2 | turquoise | -0.100702622 | 0.61723031 |
| P2RX4 | turquoise | -0.100466051 | 0.618061582 |
| B3GAT3 | turquoise | -0.100424037 | 0.618209263 |
| ACER3 | turquoise | -0.100163325 | 0.619125983 |
| TOP2B | turquoise | -0.100154736 | 0.619156191 |
| TMED7 | turquoise | -0.100065006 | 0.619471839 |
| NAF1 | turquoise | -0.099619464 | 0.621040108 |
| AIM1 | turquoise | -0.099578579 | 0.621184101 |
| AP4E1 | turquoise | -0.099234775 | 0.622395481 |
| C20orf194 | turquoise | -0.098907526 | 0.623549425 |
| TMEM41B | turquoise | -0.098544189 | 0.624831634 |
| PTAFR | turquoise | -0.098355008 | 0.625499674 |
| GPR77 | turquoise | -0.09758672 | 0.628215635 |
| LY96 | turquoise | -0.097462476 | 0.628655292 |
| SFR1 | turquoise | -0.097123751 | 0.629854554 |
| GOLGA7B | turquoise | -0.097087468 | 0.629983067 |
| SSH3 | turquoise | -0.096923821 | 0.630562839 |
| BBS5 | turquoise | -0.096842781 | 0.630850027 |
| MFN1 | turquoise | -0.096695314 | 0.631372751 |
| WDR34 | turquoise | 0.010366 | 0.959916 |
| SLC25A1 | turquoise | 0.010449 | 0.959596 |
| MEF2A | turquoise | 0.011156 | 0.956864 |
| ZNF337 | turquoise | 0.011562 | 0.955296 |
| CCDC146 | turquoise | 0.012255 | 0.952619 |
| RCHY1 | turquoise | 0.012509 | 0.951641 |
| CCDC85C | turquoise | 0.012523 | 0.951585 |
| ZNF498 | turquoise | 0.012623 | 0.951199 |
| ZBTB25 | turquoise | 0.012774 | 0.950615 |
| SGSM3 | turquoise | 0.012774 | 0.950614 |
| HBP1 | turquoise | 0.012796 | 0.95053 |
| TNFRSF1B | turquoise | 0.013341 | 0.948426 |
| KLF16 | turquoise | 0.013477 | 0.947902 |
| ATN1 | turquoise | 0.013537 | 0.947669 |
| SIRT3 | turquoise | 0.013749 | 0.946852 |
| HRAS | turquoise | 0.014198 | 0.945117 |
| PPAP2A | turquoise | 0.014267 | 0.944851 |
| FMNL2 | turquoise | 0.01436 | 0.944493 |
| FAM172A | turquoise | 0.014755 | 0.942969 |
| UBE2D1 | turquoise | 0.015003 | 0.94201 |
| MOSPD1 | turquoise | 0.01528 | 0.940942 |
| RCAN1 | turquoise | 0.015336 | 0.940725 |
| GTF3C6 | turquoise | 0.015869 | 0.938668 |
| CMPK1 | turquoise | 0.016246 | 0.937217 |
| DBT | turquoise | 0.016588 | 0.935896 |
| NUP98 | turquoise | 0.017465 | 0.932516 |
| MTOR | turquoise | 0.017911 | 0.930794 |
| POLR1C | turquoise | 0.018285 | 0.929353 |
| DNAJA2 | turquoise | 0.018291 | 0.92933 |
| FKTN | turquoise | 0.018312 | 0.929249 |
| FPGS | turquoise | 0.018572 | 0.928248 |
| SAFB2 | turquoise | 0.018599 | 0.928144 |
| HSPA9 | turquoise | 0.018603 | 0.928131 |
| MAP4K4 | turquoise | 0.018659 | 0.927913 |
| ABHD14B | turquoise | 0.018907 | 0.926956 |
| TTYH3 | turquoise | 0.018972 | 0.926705 |
| KDM3A | turquoise | 0.019174 | 0.92593 |
| ANKH | turquoise | 0.019427 | 0.924954 |
| GTF2I | turquoise | 0.019441 | 0.924899 |
| AKT3 | turquoise | 0.019478 | 0.924757 |
| STK4 | turquoise | 0.020136 | 0.922221 |
| ZNF791 | turquoise | 0.020378 | 0.92129 |
| PCYT2 | turquoise | 0.020544 | 0.920651 |
| PACS2 | turquoise | 0.020803 | 0.919656 |
| RRP15 | turquoise | 0.020958 | 0.91906 |
| CARS2 | turquoise | 0.021595 | 0.916607 |
| SOX13 | turquoise | 0.022219 | 0.914203 |
| DPH2 | turquoise | 0.022472 | 0.913233 |
| STAM2 | turquoise | 0.02249 | 0.913163 |
| GPR89C | turquoise | 0.023151 | 0.910621 |
| FBXO33 | turquoise | 0.023277 | 0.910134 |
| SLC6A12 | turquoise | 0.023433 | 0.909536 |
| FRA10AC1 | turquoise | 0.023906 | 0.907717 |
| MIER1 | turquoise | 0.024232 | 0.906463 |
| WDR82 | turquoise | 0.024493 | 0.905459 |
| MYBBP1A | turquoise | 0.024603 | 0.905038 |

**Table 6. Gene module membership and correlation with growth rate (48/72h), with adjusted p values for each module**

| | | | | | | | | | | | |
|---|---|---|---|---|---|---|---|---|---|---|---|
| ARHGAP32 | turquoise | 0.127482 | 0.534859 | APOBR | turquoise | -0.096337252 | 0.632642694 | ERO1LB | turquoise | 0.024706 | 0.904641 |
| KPNA3 | turquoise | -0.127464 | 0.534917 | RBM34 | turquoise | -0.095946749 | 0.634028855 | HIBCH | turquoise | 0.024733 | 0.904539 |
| ADNP | turquoise | -0.127435 | 0.535012 | EPG5 | turquoise | -0.095942525 | 0.634043855 | PFKP | turquoise | 0.024864 | 0.904036 |
| STPG1 | turquoise | -0.127407 | 0.535101 | POLR2B | turquoise | -0.095851289 | 0.634367891 | PELI1 | turquoise | 0.025259 | 0.902519 |
| ATP6V1B2 | turquoise | -0.12727 | 0.535544 | PGM1 | turquoise | -0.095821715 | 0.634472938 | KIAA2026 | turquoise | 0.025357 | 0.902142 |
| ACBD5 | turquoise | -0.127243 | 0.535634 | GBAS | turquoise | -0.095634936 | 0.635136556 | DTX2P1-UPK3BP1-P | turquoise | 0.025465 | 0.901727 |
| GMPS | turquoise | -0.126877 | 0.536815 | PTPRM | turquoise | -0.095492584 | 0.635642511 | GOLPH3L | turquoise | 0.026662 | 0.89713 |
| UBR3 | turquoise | 0.12668 | 0.537455 | SNX14 | turquoise | -0.095485683 | 0.635667044 | CCDC97 | turquoise | 0.026936 | 0.89608 |
| NINJ2 | turquoise | 0.125923 | 0.539906 | ACVR2A | turquoise | -0.094733943 | 0.638341596 | CEP104 | turquoise | 0.026987 | 0.895883 |
| RXRA | turquoise | -0.125764 | 0.540425 | PPA2 | turquoise | -0.094640028 | 0.638676042 | AMFR | turquoise | 0.027097 | 0.895461 |
| ARID4A | turquoise | -0.12575 | 0.54047 | FPGS | turquoise | -0.094494716 | 0.639193653 | GTPBP6 | turquoise | 0.027375 | 0.894395 |
| ATHL1 | turquoise | -0.125672 | 0.540724 | SGSM3 | turquoise | -0.094266032 | 0.640008577 | CYTH3 | turquoise | 0.027638 | 0.893384 |
| EPB41L4A | turquoise | -0.125449 | 0.541446 | NMB | turquoise | -0.093849931 | 0.641492409 | UBA3 | turquoise | 0.027732 | 0.893027 |
| DCAF13 | turquoise | -0.12534 | 0.541802 | EXOC7 | turquoise | -0.093840328 | 0.641526671 | TFDP2 | turquoise | 0.027937 | 0.89224 |
| COTL1 | turquoise | -0.1252 | 0.542257 | ZNF229 | turquoise | -0.093536644 | 0.642610494 | IFNAR2 | turquoise | 0.0282 | 0.891232 |
| GGNBP2 | turquoise | -0.12479 | 0.54359 | MBP | turquoise | -0.093254433 | 0.643618323 | GRWD1 | turquoise | 0.028526 | 0.889979 |
| CDKN2B | turquoise | -0.124721 | 0.543815 | USP5 | turquoise | -0.093007654 | 0.644500121 | PRPF39 | turquoise | 0.028607 | 0.889671 |
| C11orf73 | turquoise | -0.124523 | 0.54446 | TTC17 | turquoise | -0.092954011 | 0.644691859 | DNAAF2 | turquoise | 0.029476 | 0.88634 |
| KHSRP | turquoise | -0.124261 | 0.545313 | C9orf123 | turquoise | -0.092951919 | 0.644699339 | TMX1 | turquoise | 0.029864 | 0.884852 |
| B4GALT2 | turquoise | -0.124113 | 0.545793 | LMF2 | turquoise | -0.092847332 | 0.645073239 | SLC26A2 | turquoise | 0.029886 | 0.88477 |
| CCDC132 | turquoise | -0.123761 | 0.546944 | PICALM | turquoise | -0.092792062 | 0.645270865 | SSH3 | turquoise | 0.030068 | 0.884073 |
| STX16 | turquoise | -0.123496 | 0.547808 | RAB27A | turquoise | -0.092747362 | 0.645430713 | TMED3 | turquoise | 0.030514 | 0.882363 |
| DAP3 | turquoise | -0.123465 | 0.54791 | CABLES2 | turquoise | -0.092449684 | 0.646495607 | KIN | turquoise | 0.030793 | 0.881297 |
| MYO7B | turquoise | -0.123431 | 0.548023 | OCIAD1 | turquoise | -0.092446434 | 0.646507235 | DYNLT3 | turquoise | 0.03093 | 0.880771 |
| PARP11 | turquoise | -0.123426 | 0.548037 | FAM45A | turquoise | -0.091891617 | 0.648493829 | DHRS3 | turquoise | 0.031598 | 0.878217 |
| LRRC34 | turquoise | -0.122881 | 0.54982 | PTPLAD2 | turquoise | -0.090183595 | 0.654624322 | KDM5A | turquoise | 0.032023 | 0.876587 |
| SKIL | turquoise | -0.122845 | 0.549937 | TSSC1 | turquoise | -0.090123629 | 0.654839955 | FBXL18 | turquoise | 0.032642 | 0.874224 |
| KRAS | turquoise | -0.122613 | 0.550698 | FOXO4 | turquoise | -0.090075718 | 0.655012258 | FOXN3 | turquoise | 0.033024 | 0.872762 |
| TAF12 | turquoise | -0.122497 | 0.551076 | TIMP2 | turquoise | -0.089958906 | 0.655432425 | JMJD8 | turquoise | 0.033158 | 0.87225 |
| HERC1 | turquoise | 0.122243 | 0.551909 | SMPDL3A | turquoise | -0.089667382 | 0.656481464 | ZNF280D | turquoise | 0.033197 | 0.872102 |
| RPL37A | turquoise | 0.122232 | 0.551946 | ACTR10 | turquoise | -0.088943514 | 0.659089032 | IQCB1 | turquoise | 0.033807 | 0.869771 |
| FOXN3 | turquoise | 0.122089 | 0.552415 | ITGAE | turquoise | -0.088744422 | 0.659806898 | ST14 | turquoise | 0.03395 | 0.869222 |
| RPAIN | turquoise | -0.121989 | 0.55274 | ERICH1 | turquoise | -0.088618553 | 0.660260895 | STBD1 | turquoise | 0.033954 | 0.869209 |
| TRAM1 | turquoise | -0.121986 | 0.552751 | SEPN1 | turquoise | -0.088549374 | 0.660510469 | PIAS1 | turquoise | 0.033974 | 0.869132 |
| GBP5 | turquoise | -0.121831 | 0.553259 | TMEM68 | turquoise | -0.088157527 | 0.661924775 | LXN | turquoise | 0.034263 | 0.868028 |
| NUCB2 | turquoise | -0.121779 | 0.55343 | TNIP3 | turquoise | -0.087855409 | 0.663015993 | TEF | turquoise | 0.034431 | 0.867386 |
| STK10 | turquoise | -0.121742 | 0.553552 | ANO10 | turquoise | -0.087545025 | 0.664137771 | DBR1 | turquoise | 0.034949 | 0.865408 |
| TMA16 | turquoise | -0.121109 | 0.555632 | BPTF | turquoise | -0.087516606 | 0.664240515 | PDCD2L | turquoise | 0.035041 | 0.865059 |
| RSPRY1 | turquoise | -0.120552 | 0.557464 | GNB5 | turquoise | -0.087293033 | 0.665049027 | MANBAL | turquoise | 0.035326 | 0.863969 |
| MSL2 | turquoise | -0.120492 | 0.557663 | ANKRA2 | turquoise | -0.087267251 | 0.665142285 | HERC1 | turquoise | 0.035418 | 0.86362 |
| SH3PXD2A | turquoise | -0.12015 | 0.55879 | FOPNL | turquoise | -0.087126917 | 0.665649993 | RABGAP1L | turquoise | 0.036782 | 0.858419 |
| COX6B1 | turquoise | -0.120143 | 0.558813 | IGF2R | turquoise | -0.086950717 | 0.66628766 | ZNF613 | turquoise | 0.037695 | 0.854942 |
| RBM11 | turquoise | 0.119978 | 0.559357 | KLF16 | turquoise | -0.086859946 | 0.666616249 | PLK1S1 | turquoise | 0.038254 | 0.852813 |
| LFNG | turquoise | -0.119882 | 0.559674 | ZBTB41 | turquoise | -0.08672457 | 0.667106416 | LTV1 | turquoise | 0.038378 | 0.852339 |
| TUBE1 | turquoise | -0.119782 | 0.560001 | ACSL3 | turquoise | -0.086554234 | 0.667723356 | MFSD2A | turquoise | 0.038574 | 0.851593 |
| MRPL40 | turquoise | -0.119765 | 0.560059 | C4orf27 | turquoise | -0.086401529 | 0.668276618 | PARP16 | turquoise | 0.038863 | 0.850493 |
| ROGDI | turquoise | -0.119586 | 0.560651 | EPC1 | turquoise | -0.08591703 | 0.670033116 | ALPK3 | turquoise | 0.039134 | 0.849464 |
| PTEN | turquoise | -0.119486 | 0.56098 | ALDH16A1 | turquoise | -0.085829583 | 0.670350325 | ZMYM2 | turquoise | 0.039431 | 0.848334 |
| EPG5 | turquoise | -0.119411 | 0.561228 | EIF4E2 | turquoise | -0.085694047 | 0.670842085 | SNX33 | turquoise | 0.039552 | 0.847872 |
| ERGIC2 | turquoise | -0.119353 | 0.561418 | ATG16L2 | turquoise | -0.085623702 | 0.671097366 | SWI5 | turquoise | 0.039594 | 0.847713 |
| DDX47 | turquoise | -0.119226 | 0.561838 | BCL2A1 | turquoise | -0.085550478 | 0.671363137 | HNRPDL | turquoise | 0.040081 | 0.845863 |
| ME3 | turquoise | 0.119205 | 0.56191 | AMPD2 | turquoise | -0.085290634 | 0.672306555 | RPS15 | turquoise | 0.04015 | 0.845601 |
| PRPF40A | turquoise | -0.119141 | 0.562121 | SLC35A5 | turquoise | -0.084988415 | 0.673404435 | C7orf60 | turquoise | 0.040155 | 0.845581 |
| GOLPH3 | turquoise | -0.119104 | 0.562243 | C5orf28 | turquoise | -0.084854464 | 0.673891254 | NUCB1 | turquoise | 0.040553 | 0.844068 |
| THYN1 | turquoise | -0.11877 | 0.563346 | RB1 | turquoise | -0.084787621 | 0.674134227 | SDHAF1 | turquoise | 0.040685 | 0.843568 |
| CXCL11 | turquoise | -0.118674 | 0.563665 | KRCC1 | turquoise | -0.084783495 | 0.674149228 | ATF5 | turquoise | 0.040904 | 0.842734 |

**Table 6. Gene module membership and correlation with growth rate (48/72h), with adjusted p values for each module**

| | | | | | | | | | | | |
|---|---|---|---|---|---|---|---|---|---|---|---|
| RFX1 | turquoise | 0.118558 | 0.564048 | ROGDI | turquoise | -0.084661366 | 0.674593255 | ZFP30 | turquoise | 0.041203 | 0.841602 |
| UBE2M | turquoise | -0.118486 | 0.564287 | CUL5 | turquoise | -0.084363788 | 0.675675606 | SARS2 | turquoise | 0.041266 | 0.84136 |
| PDE2A | turquoise | 0.118331 | 0.564799 | DTWD2 | turquoise | -0.08391444 | 0.677311172 | TAF1A | turquoise | 0.041496 | 0.840489 |
| WASH5P | turquoise | -0.118289 | 0.564939 | ACBD6 | turquoise | -0.08378418 | 0.677785569 | ALG6 | turquoise | 0.041499 | 0.840478 |
| SPATA7 | turquoise | -0.1182 | 0.565233 | GAPVD1 | turquoise | -0.083643455 | 0.678298212 | SMC5 | turquoise | 0.042004 | 0.838559 |
| CCT3 | turquoise | -0.117895 | 0.566246 | TOE1 | turquoise | -0.082796629 | 0.681386038 | SAP30L | turquoise | 0.042158 | 0.837977 |
| WIPF1 | turquoise | -0.117892 | 0.566255 | SUZ12 | turquoise | -0.082162782 | 0.683700555 | CLEC12A | turquoise | 0.04339 | 0.833306 |
| ITPKB | turquoise | -0.117703 | 0.566882 | CBX4 | turquoise | -0.082134564 | 0.683803661 | ACTN1 | turquoise | 0.043838 | 0.831609 |
| RASA2 | turquoise | -0.117511 | 0.56752 | TGFB1 | turquoise | -0.081991502 | 0.684326471 | CIAPIN1 | turquoise | 0.044006 | 0.830973 |
| SNURF | turquoise | -0.117489 | 0.567592 | NELF | turquoise | -0.081594154 | 0.685779298 | NCK1 | turquoise | 0.044147 | 0.830437 |
| TNIP3 | turquoise | -0.117287 | 0.568263 | DENND6A | turquoise | -0.081427663 | 0.686388364 | ATG10 | turquoise | 0.044377 | 0.829566 |
| VDAC2 | turquoise | -0.117203 | 0.568541 | XRCC4 | turquoise | -0.081347174 | 0.686682886 | ACIN1 | turquoise | 0.044806 | 0.827945 |
| TRAPPC6B | turquoise | -0.116978 | 0.569291 | PITPNA | turquoise | -0.081340566 | 0.686707067 | SLC15A3 | turquoise | 0.04482 | 0.827892 |
| MTA3 | turquoise | -0.116906 | 0.56953 | WBP1L | turquoise | -0.081261223 | 0.686997437 | ANKRA2 | turquoise | 0.045178 | 0.826538 |
| LATS2 | turquoise | -0.116758 | 0.57002 | EXOC1 | turquoise | -0.081161216 | 0.687363495 | STK36 | turquoise | 0.045192 | 0.826484 |
| ZNF282 | turquoise | -0.116641 | 0.570411 | ENSA | turquoise | -0.080849255 | 0.688505819 | ZMYM6 | turquoise | 0.045331 | 0.825957 |
| CLEC12A | turquoise | -0.116457 | 0.571024 | PGD | turquoise | -0.080746935 | 0.688880633 | TSPYL5 | turquoise | 0.04546 | 0.825469 |
| TNFAIP2 | turquoise | 0.116408 | 0.571185 | ZNF202 | turquoise | -0.080712002 | 0.689008616 | MCPH1 | turquoise | 0.04565 | 0.824752 |
| ZCCHC16 | turquoise | 0.116279 | 0.571617 | PGRMC2 | turquoise | -0.08063646 | 0.689285403 | CDC42BPA | turquoise | 0.04635 | 0.822106 |
| H3F3B | turquoise | 0.115484 | 0.574266 | RELB | turquoise | -0.08063402 | 0.689294342 | CMTM6 | turquoise | 0.047128 | 0.81917 |
| KATNBL1 | turquoise | -0.115467 | 0.574324 | SHOC2 | turquoise | -0.080520487 | 0.689710404 | BDP1 | turquoise | 0.048256 | 0.814912 |
| GRHPR | turquoise | -0.115402 | 0.574538 | C5orf51 | turquoise | -0.080465737 | 0.689911077 | FGR | turquoise | 0.049558 | 0.810009 |
| C1orf50 | turquoise | -0.115223 | 0.575138 | FTSJ1 | turquoise | -0.080291322 | 0.690550488 | PIGK | turquoise | 0.050001 | 0.808341 |
| SAMD9 | turquoise | -0.114842 | 0.576412 | ZFP106 | turquoise | -0.080232207 | 0.690767253 | LARS | turquoise | 0.050142 | 0.80781 |
| PSPC1 | turquoise | 0.11482 | 0.576484 | NSUN4 | turquoise | -0.079172715 | 0.69465625 | DHX38 | turquoise | 0.050875 | 0.805052 |
| BAHD1 | turquoise | 0.114796 | 0.576563 | RAB5B | turquoise | -0.079116348 | 0.694863362 | CORO7 | turquoise | 0.051188 | 0.803875 |
| PMAIP1 | turquoise | 0.114775 | 0.576633 | TXN | turquoise | -0.079030949 | 0.695177194 | RTTN | turquoise | 0.051441 | 0.802926 |
| DYRK1B | turquoise | -0.114703 | 0.576876 | CCDC88B | turquoise | -0.078976163 | 0.695378552 | FAM118B | turquoise | 0.051663 | 0.802089 |
| NXF3 | turquoise | -0.114526 | 0.577468 | ANKRD50 | turquoise | -0.078550535 | 0.696943566 | TAMM41 | turquoise | 0.051814 | 0.801523 |
| DIS3 | turquoise | -0.113506 | 0.580882 | IFNGR1 | turquoise | -0.078186887 | 0.698281642 | CCT8 | turquoise | 0.052385 | 0.799379 |
| MRPS27 | turquoise | -0.113223 | 0.581834 | GLCE | turquoise | -0.077851745 | 0.699515609 | SUMF1 | turquoise | 0.052814 | 0.797769 |
| MIER1 | turquoise | -0.113164 | 0.582031 | GMPS | turquoise | -0.077629434 | 0.700334552 | RPE | turquoise | 0.053009 | 0.797038 |
| EIF4E2 | turquoise | -0.113036 | 0.58246 | ITGAX | turquoise | -0.077595585 | 0.700459274 | TRIM28 | turquoise | 0.05329 | 0.795984 |
| SHPRH | turquoise | 0.112494 | 0.584282 | NIPSNAP1 | turquoise | -0.077572212 | 0.700545398 | ADNP2 | turquoise | 0.053566 | 0.794952 |
| SBNO2 | turquoise | 0.11233 | 0.584833 | NCDN | turquoise | -0.077333794 | 0.701424127 | CNTRL | turquoise | 0.053971 | 0.793433 |
| GAA | turquoise | 0.112297 | 0.584944 | CPVL | turquoise | -0.076650029 | 0.703946339 | TAZ | turquoise | 0.05411 | 0.792913 |
| CINP | turquoise | -0.112263 | 0.58506 | CENPB | turquoise | -0.076182511 | 0.705672638 | NUDCD1 | turquoise | 0.054534 | 0.791324 |
| CAPNS1 | turquoise | -0.112002 | 0.585938 | SLC2A13 | turquoise | -0.076128777 | 0.705871141 | S100A10 | turquoise | 0.054962 | 0.789723 |
| ODF3B | turquoise | -0.111738 | 0.586825 | CLMN | turquoise | -0.076030055 | 0.706235889 | E2F3 | turquoise | 0.055374 | 0.788183 |
| ORC2 | turquoise | -0.111386 | 0.588012 | PIGV | turquoise | -0.075870718 | 0.706824719 | ANKRD50 | turquoise | 0.055434 | 0.787958 |
| SLC19A3 | turquoise | -0.111384 | 0.58802 | VPS54 | turquoise | -0.07538731 | 0.708612168 | GADD45A | turquoise | 0.055682 | 0.78703 |
| UBXN1 | turquoise | -0.111318 | 0.588242 | PTGER2 | turquoise | -0.075273826 | 0.709032006 | DLAT | turquoise | 0.055834 | 0.786463 |
| ADAT1 | turquoise | -0.111258 | 0.588446 | CTNNAL1 | turquoise | -0.074955616 | 0.710209675 | LMNA | turquoise | 0.055942 | 0.786058 |
| STX18 | turquoise | -0.111126 | 0.588891 | FEM1C | turquoise | -0.074498645 | 0.711902018 | ATP13A1 | turquoise | 0.056021 | 0.785762 |
| RP2 | turquoise | 0.111086 | 0.589025 | INO80E | turquoise | -0.07422613 | 0.712911889 | DNAJC1 | turquoise | 0.056802 | 0.782845 |
| FAM63B | turquoise | -0.110893 | 0.589677 | U2SURP | turquoise | -0.074002114 | 0.713742384 | KIAA1429 | turquoise | 0.056887 | 0.782529 |
| HNRPDL | turquoise | -0.110874 | 0.589741 | ATG4A | turquoise | -0.073958099 | 0.7139056 | HDAC4 | turquoise | 0.056927 | 0.782378 |
| TMEM59 | turquoise | -0.110868 | 0.589762 | DDX47 | turquoise | -0.073885242 | 0.714175791 | ERCC8 | turquoise | 0.057222 | 0.781279 |
| MBNL3 | turquoise | -0.11053 | 0.590902 | CLSTN1 | turquoise | -0.073793006 | 0.714517901 | EIF3H | turquoise | 0.057569 | 0.779983 |
| RNF24 | turquoise | -0.110471 | 0.591102 | PWWP2B | turquoise | -0.073641938 | 0.715078337 | SUPT7L | turquoise | 0.057803 | 0.77911 |
| HNRNPA0 | turquoise | -0.11018 | 0.592085 | NEDD4 | turquoise | -0.073491724 | 0.715635746 | CDH23 | turquoise | 0.057975 | 0.778471 |
| WAS | turquoise | 0.110154 | 0.592175 | SPATA2L | turquoise | -0.072609782 | 0.718911299 | ZNF562 | turquoise | 0.058057 | 0.778165 |
| BOD1 | turquoise | -0.109943 | 0.592889 | C3orf17 | turquoise | -0.07163889 | 0.722522825 | SLC45A4 | turquoise | 0.058304 | 0.777242 |
| YTHDF3 | turquoise | -0.109866 | 0.59315 | KCTD7 | turquoise | -0.071527016 | 0.722939346 | BLNK | turquoise | 0.05866 | 0.775919 |
| GALC | turquoise | -0.109502 | 0.59438 | SMIM4 | turquoise | -0.071421598 | 0.723331906 | AVL9 | turquoise | 0.058809 | 0.775364 |
| WDR6 | turquoise | 0.108005 | 0.599461 | TBC1D24 | turquoise | -0.070810521 | 0.725608789 | AQP3 | turquoise | 0.059297 | 0.773546 |

**Table 6. Gene module membership and correlation with growth rate (48/72h), with adjusted p values for each module**

| | | | |
|---|---|---|---|
| DNTTIP1 | turquoise | -0.107934 | 0.599703 |
| PLCG1 | turquoise | -0.107775 | 0.600245 |
| PEX3 | turquoise | -0.107703 | 0.600488 |
| TEX261 | turquoise | -0.10752 | 0.601111 |
| ZFYVE21 | turquoise | -0.107296 | 0.601873 |
| VAPA | turquoise | 0.107189 | 0.602239 |
| UTP20 | turquoise | -0.107024 | 0.602802 |
| IQCB1 | turquoise | -0.106991 | 0.602912 |
| MAP4K5 | turquoise | -0.106983 | 0.60294 |
| CLEC1A | turquoise | -0.106972 | 0.602977 |
| EXOC7 | turquoise | -0.106909 | 0.603194 |
| APAF1 | turquoise | -0.106798 | 0.603571 |
| ELOVL7 | turquoise | -0.106367 | 0.605042 |
| METTL21D | turquoise | 0.105816 | 0.606922 |
| TRAPPC8 | turquoise | -0.105696 | 0.607334 |
| ATG7 | turquoise | -0.105049 | 0.609546 |
| HINFP | turquoise | 0.104365 | 0.611891 |
| METTL25 | turquoise | -0.104131 | 0.612693 |
| USP32 | turquoise | -0.104004 | 0.61313 |
| CMTM7 | turquoise | -0.103704 | 0.61416 |
| TNNT1 | turquoise | -0.103357 | 0.615351 |
| FRA10AC1 | turquoise | -0.102949 | 0.616756 |
| LRPPRC | turquoise | -0.102902 | 0.616915 |
| CSNK1E | turquoise | -0.102882 | 0.616986 |
| HIVEP1 | turquoise | -0.102826 | 0.617176 |
| VPS54 | turquoise | -0.102786 | 0.617314 |
| WHSC1L1 | turquoise | -0.102591 | 0.617986 |
| C1GALT1 | turquoise | -0.102381 | 0.618709 |
| FAM91A1 | turquoise | 0.102261 | 0.619125 |
| ZNF322 | turquoise | -0.102132 | 0.619569 |
| PRRT4 | turquoise | -0.101926 | 0.620277 |
| ARGLU1 | turquoise | 0.101819 | 0.620646 |
| PLEKHM2 | turquoise | -0.101674 | 0.621146 |
| RGPD6 | turquoise | 0.101647 | 0.621242 |
| N4BP2L1 | turquoise | 0.101585 | 0.621454 |
| RFX5 | turquoise | -0.101536 | 0.621624 |
| PGRMC2 | turquoise | -0.101484 | 0.621805 |
| ZNF76 | turquoise | -0.10112 | 0.623061 |
| PPAP2A | turquoise | 0.10071 | 0.624478 |
| ZNF25 | turquoise | -0.10062 | 0.62479 |
| CELF1 | turquoise | -0.1006 | 0.624857 |
| ELP2 | turquoise | -0.100413 | 0.625506 |
| RAB8B | turquoise | -0.100306 | 0.625877 |
| MBNL1 | turquoise | 0.100132 | 0.626477 |
| ARL6IP6 | turquoise | -0.099704 | 0.627962 |
| LYST | turquoise | -0.099534 | 0.62855 |
| PROSC | turquoise | -0.099434 | 0.628896 |
| S100A16 | turquoise | -0.099247 | 0.629547 |
| WNK1 | turquoise | -0.099218 | 0.629645 |
| PLEC | turquoise | -0.09861 | 0.631756 |
| SPATA2 | turquoise | 0.098545 | 0.631981 |
| GPR107 | turquoise | -0.098517 | 0.63208 |
| TRNT1 | turquoise | -0.098492 | 0.632165 |
| SH3TC1 | turquoise | -0.098411 | 0.632446 |
| MSRA | turquoise | 0.098312 | 0.632794 |
| RTTN | turquoise | -0.098255 | 0.632992 |
| RABGAP1 | turquoise | -0.070710428 | 0.725981956 |
| MX2 | turquoise | -0.070461949 | 0.726908597 |
| RIF1 | turquoise | -0.070442372 | 0.72698162 |
| TRIP11 | turquoise | -0.07022802 | 0.727781321 |
| SDHB | turquoise | -0.069470264 | 0.730610566 |
| PIK3R4 | turquoise | -0.069196802 | 0.731632444 |
| HIVEP1 | turquoise | -0.069137239 | 0.731855079 |
| PPP6R2 | turquoise | -0.068644234 | 0.733698649 |
| PTPN22 | turquoise | -0.068306237 | 0.734963412 |
| CROCCP2 | turquoise | -0.068100771 | 0.735732577 |
| CYP20A1 | turquoise | -0.067468126 | 0.738102465 |
| DYNC1LI1 | turquoise | -0.067344154 | 0.738567139 |
| ERO1L | turquoise | -0.066774835 | 0.740702216 |
| ARSB | turquoise | -0.066510668 | 0.74169354 |
| EMC2 | turquoise | -0.066464288 | 0.741867629 |
| SLC25A1 | turquoise | -0.065936681 | 0.743848894 |
| AK1 | turquoise | -0.065749137 | 0.744553542 |
| TRIM41 | turquoise | -0.065726775 | 0.744637577 |
| ATF5 | turquoise | -0.065091324 | 0.747026687 |
| ABCA5 | turquoise | -0.0645034 | 0.749239149 |
| NDUFB7 | turquoise | -0.064279899 | 0.750080728 |
| THAP2 | turquoise | -0.063316858 | 0.753710199 |
| ADI1 | turquoise | -0.063307437 | 0.753745726 |
| SRPK2 | turquoise | -0.062820781 | 0.755581793 |
| ZNF788 | turquoise | -0.062708292 | 0.756006374 |
| ZDHHC20 | turquoise | -0.062701958 | 0.756030286 |
| HTATIP2 | turquoise | -0.062682258 | 0.75610465 |
| ACAA1 | turquoise | -0.06250669 | 0.756767488 |
| ZFP161 | turquoise | -0.062362524 | 0.757311896 |
| SLC25A36 | turquoise | -0.062248658 | 0.757741964 |
| GPCPD1 | turquoise | -0.062210855 | 0.75788476 |
| ANO6 | turquoise | -0.061578729 | 0.760273678 |
| ATP6V1B2 | turquoise | -0.061276014 | 0.761418455 |
| DRAM2 | turquoise | -0.060902643 | 0.762831108 |
| PPP6C | turquoise | -0.060864084 | 0.76297704 |
| SGMS2 | turquoise | -0.060275412 | 0.76520591 |
| MTF1 | turquoise | -0.058986402 | 0.770092805 |
| MLLT10 | turquoise | -0.058869171 | 0.77053768 |
| ZNF121 | turquoise | -0.058726175 | 0.771080425 |
| RNF103 | turquoise | -0.058108499 | 0.773426037 |
| NUCB2 | turquoise | -0.058066811 | 0.773584417 |
| MTMR12 | turquoise | -0.057946227 | 0.774042584 |
| SH3BGRL | turquoise | -0.057898688 | 0.774223231 |
| SNRNP70 | turquoise | -0.057401043 | 0.776114971 |
| AQR | turquoise | -0.057329997 | 0.776385147 |
| ATP6V0D1 | turquoise | -0.057317055 | 0.776434365 |
| RASA1 | turquoise | -0.057256596 | 0.776664303 |
| NIPA2 | turquoise | -0.056571828 | 0.7792699 |
| MRPS17 | turquoise | -0.056339372 | 0.780154946 |
| ITGAL | turquoise | -0.056186656 | 0.780736537 |
| CDK2AP2 | turquoise | -0.056080485 | 0.781140938 |
| CYB5D2 | turquoise | -0.055903893 | 0.781813692 |
| AHCYL2 | turquoise | -0.055705399 | 0.78257007 |
| ARMC6 | turquoise | -0.055473482 | 0.783454051 |
| HOMER1 | turquoise | -0.055418272 | 0.78366453 |
| PTEN | turquoise | -0.054955838 | 0.785428065 |
| CTNND1 | turquoise | 0.059625 | 0.772326 |
| PRKCI | turquoise | 0.060313 | 0.769769 |
| RBM19 | turquoise | 0.06083 | 0.767846 |
| LMAN2L | turquoise | 0.060843 | 0.767799 |
| RAD52 | turquoise | 0.061408 | 0.765701 |
| ACAP3 | turquoise | 0.06169 | 0.764657 |
| RPTOR | turquoise | 0.061693 | 0.764643 |
| SF3A2 | turquoise | 0.062183 | 0.762828 |
| DYNC1I2 | turquoise | 0.062688 | 0.760956 |
| FAM18B2 | turquoise | 0.063448 | 0.758143 |
| HIPK2 | turquoise | 0.063652 | 0.757389 |
| TBC1D5 | turquoise | 0.064537 | 0.754114 |
| GNB2 | turquoise | 0.065019 | 0.752334 |
| PICALM | turquoise | 0.065149 | 0.751856 |
| FNBP4 | turquoise | 0.065159 | 0.751819 |
| ZBTB11 | turquoise | 0.065238 | 0.751527 |
| PJA2 | turquoise | 0.06527 | 0.751408 |
| DDX17 | turquoise | 0.066165 | 0.748107 |
| RMND5B | turquoise | 0.066271 | 0.747718 |
| PPARGC1B | turquoise | 0.066784 | 0.745828 |
| PTGER3 | turquoise | 0.067236 | 0.744162 |
| IL27RA | turquoise | 0.067471 | 0.743297 |
| TAF13 | turquoise | 0.067484 | 0.743249 |
| WWP2 | turquoise | 0.068047 | 0.741179 |
| USP6NL | turquoise | 0.068177 | 0.740702 |
| G6PD | turquoise | 0.068223 | 0.740534 |
| C6orf136 | turquoise | 0.068233 | 0.740496 |
| TRPV2 | turquoise | 0.069331 | 0.736465 |
| MLPH | turquoise | 0.069432 | 0.736093 |
| RETSAT | turquoise | 0.069522 | 0.735766 |
| CXorf40B | turquoise | 0.069653 | 0.735284 |
| ZFAND2A | turquoise | 0.069728 | 0.73501 |
| GEMIN2 | turquoise | 0.069923 | 0.734292 |
| CEP57L1 | turquoise | 0.070201 | 0.733276 |
| PTER | turquoise | 0.071364 | 0.729017 |
| RXRA | turquoise | 0.071797 | 0.727434 |
| INO80E | turquoise | 0.071884 | 0.727117 |
| DHX58 | turquoise | 0.072044 | 0.726533 |
| RAB12 | turquoise | 0.072613 | 0.724455 |
| CWC22 | turquoise | 0.072634 | 0.724376 |
| THAP6 | turquoise | 0.073104 | 0.722665 |
| YBEY | turquoise | 0.073887 | 0.719811 |
| SLC25A30 | turquoise | 0.07415 | 0.718851 |
| DNAJB9 | turquoise | 0.074779 | 0.716565 |
| PTPLA | turquoise | 0.075117 | 0.715335 |
| PKNOX1 | turquoise | 0.075168 | 0.715151 |
| BAX | turquoise | 0.075531 | 0.713828 |
| RRN3P1 | turquoise | 0.076007 | 0.712102 |
| NFKBIZ | turquoise | 0.076801 | 0.709223 |
| FBXL3 | turquoise | 0.07695 | 0.708682 |
| RBM7 | turquoise | 0.076981 | 0.708569 |
| FAM217B | turquoise | 0.077446 | 0.706885 |
| HDAC10 | turquoise | 0.07777 | 0.705713 |
| CHD2 | turquoise | 0.079083 | 0.700968 |
| CHIC2 | turquoise | 0.079666 | 0.698864 |
| PIGO | turquoise | 0.079812 | 0.698338 |

**Table 6. Gene module membership and correlation with growth rate (48/72h), with adjusted p values for each module**

| | | | |
|---|---|---|---|
| PGP | turquoise | -0.097912 | 0.634183 |
| RWDD2A | turquoise | -0.09766 | 0.635061 |
| MAGT1 | turquoise | 0.097627 | 0.635174 |
| CFL1 | turquoise | 0.09761 | 0.635233 |
| RAB12 | turquoise | -0.097563 | 0.635398 |
| MRPL18 | turquoise | 0.097498 | 0.635625 |
| MRPS36 | turquoise | -0.096942 | 0.63756 |
| CD300LF | turquoise | -0.096875 | 0.637794 |
| AKAP10 | turquoise | -0.096712 | 0.638365 |
| ARPC5L | turquoise | -0.096411 | 0.639413 |
| RFX7 | turquoise | -0.096317 | 0.639741 |
| NFKBIA | turquoise | -0.095833 | 0.641434 |
| VPS4B | turquoise | -0.095815 | 0.641497 |
| PIK3CG | turquoise | -0.095664 | 0.642024 |
| COPZ1 | turquoise | -0.095554 | 0.642407 |
| GRK5 | turquoise | -0.095493 | 0.64262 |
| DICER1 | turquoise | -0.095293 | 0.643321 |
| RNF148 | turquoise | 0.095147 | 0.643831 |
| CLIC2 | turquoise | -0.094972 | 0.644446 |
| MDFIC | turquoise | -0.094719 | 0.645333 |
| FAM173B | turquoise | -0.094307 | 0.646775 |
| MRPL19 | turquoise | -0.094293 | 0.646823 |
| C19orf54 | turquoise | 0.094251 | 0.646971 |
| G3BP2 | turquoise | -0.094089 | 0.647541 |
| ACE | turquoise | -0.094084 | 0.647559 |
| EPB41L3 | turquoise | -0.093908 | 0.648174 |
| AIM1 | turquoise | -0.093902 | 0.648195 |
| GEN1 | turquoise | 0.09384 | 0.648412 |
| RPL29 | turquoise | -0.093591 | 0.649286 |
| PLXDC2 | turquoise | 0.093508 | 0.64958 |
| METTL21B | turquoise | -0.093383 | 0.650019 |
| SNHG4 | turquoise | -0.093357 | 0.650111 |
| LLPH | turquoise | -0.093064 | 0.651138 |
| DPY30 | turquoise | -0.092929 | 0.651613 |
| RPTOR | turquoise | 0.092699 | 0.652422 |
| NBEAL1 | turquoise | -0.092511 | 0.653084 |
| MGST2 | turquoise | -0.092392 | 0.653504 |
| TXNRD3 | turquoise | -0.092381 | 0.653541 |
| OGFOD1 | turquoise | -0.092321 | 0.653753 |
| UBR5 | turquoise | 0.092293 | 0.653853 |
| VAPB | turquoise | -0.092147 | 0.654366 |
| NDUFV2 | turquoise | -0.091938 | 0.655103 |
| DCUN1D4 | turquoise | -0.091757 | 0.655741 |
| CPVL | turquoise | -0.091755 | 0.655747 |
| SCFD1 | turquoise | -0.091729 | 0.655838 |
| CEP152 | turquoise | -0.091522 | 0.656569 |
| DLGAP4 | turquoise | 0.091108 | 0.658028 |
| SLC25A16 | turquoise | -0.090944 | 0.658608 |
| TMEM180 | turquoise | -0.090788 | 0.659157 |
| SDHAP2 | turquoise | -0.090544 | 0.660021 |
| MCPH1 | turquoise | -0.090311 | 0.660843 |
| FZR1 | turquoise | -0.090248 | 0.661066 |
| MCM3AP | turquoise | -0.090153 | 0.661404 |
| CXCL10 | turquoise | -0.089749 | 0.662833 |
| C7orf60 | turquoise | -0.089674 | 0.663098 |
| PCED1A | turquoise | 0.089192 | 0.664802 |
| NCBP2 | turquoise | -0.054265229 | 0.788063686 |
| TWISTNB | turquoise | -0.054089576 | 0.78873441 |
| MAU2 | turquoise | -0.053888168 | 0.789503655 |
| HAUS4 | turquoise | -0.053709198 | 0.790187368 |
| RMI1 | turquoise | -0.053380627 | 0.791442985 |
| MON2 | turquoise | -0.053289649 | 0.791790745 |
| ACOT8 | turquoise | -0.053074085 | 0.792614881 |
| DISP1 | turquoise | -0.052343073 | 0.795411272 |
| ZNF714 | turquoise | -0.052064331 | 0.796478213 |
| SNX16 | turquoise | -0.051858151 | 0.79726764 |
| USP11 | turquoise | -0.051541827 | 0.798479166 |
| GTPBP10 | turquoise | -0.051518116 | 0.798569998 |
| CDKL1 | turquoise | -0.051422408 | 0.798936663 |
| VPS8 | turquoise | -0.051387677 | 0.799069731 |
| HAGH | turquoise | -0.051295316 | 0.799423624 |
| ITPKB | turquoise | -0.050827021 | 0.801218561 |
| SNAPC1 | turquoise | -0.050427783 | 0.802749588 |
| ITGB1 | turquoise | -0.050180901 | 0.803696703 |
| CD226 | turquoise | -0.050161898 | 0.803769617 |
| MLLT6 | turquoise | -0.050089306 | 0.80404816 |
| NDUFA5 | turquoise | -0.050035963 | 0.804252857 |
| FBXO46 | turquoise | -0.049969037 | 0.804509694 |
| FAM208B | turquoise | -0.049625032 | 0.805830178 |
| RHOD | turquoise | -0.049425252 | 0.806597285 |
| SH3KBP1 | turquoise | -0.049422963 | 0.806606073 |
| VMAC | turquoise | -0.048459864 | 0.810306585 |
| ZBTB25 | turquoise | -0.04838484 | 0.810595018 |
| SLC39A8 | turquoise | -0.048363488 | 0.81067711 |
| MTMR3 | turquoise | -0.046842851 | 0.816528548 |
| ZNF654 | turquoise | -0.046472747 | 0.817954181 |
| AGTPBP1 | turquoise | -0.046414507 | 0.818178569 |
| SRP54 | turquoise | -0.04635649 | 0.818402115 |
| PHAX | turquoise | -0.046306876 | 0.818593296 |
| FNTB | turquoise | -0.045876539 | 0.820251947 |
| RRP1 | turquoise | -0.045690244 | 0.820970219 |
| ST14 | turquoise | -0.045497265 | 0.821714413 |
| PFDN6 | turquoise | -0.045364625 | 0.822226003 |
| RXRB | turquoise | -0.045251577 | 0.822662087 |
| ELOVL7 | turquoise | -0.045184852 | 0.822919502 |
| TBL3 | turquoise | -0.044907481 | 0.82398975 |
| FMN1 | turquoise | -0.044856566 | 0.82418624 |
| PSMD12 | turquoise | -0.044736912 | 0.82464805 |
| SLC25A16 | turquoise | -0.0445981 | 0.825183873 |
| ATAD2 | turquoise | -0.044262455 | 0.826479796 |
| TGFBRAP1 | turquoise | -0.043807369 | 0.82823759 |
| RAD1 | turquoise | -0.043414267 | 0.829756618 |
| PRKAA1 | turquoise | -0.043017383 | 0.831290864 |
| UNC13B | turquoise | -0.042806744 | 0.832105386 |
| ADAMDEC1 | turquoise | -0.04262794 | 0.832796936 |
| ZNF664 | turquoise | -0.042449447 | 0.833487404 |
| HEXB | turquoise | -0.042373568 | 0.833780966 |
| PPIB | turquoise | -0.0421225 | 0.834752453 |
| FAM175B | turquoise | -0.041323415 | 0.837846016 |
| TEC | turquoise | -0.040688233 | 0.840306729 |
| TIMM9 | turquoise | -0.040479219 | 0.841116781 |
| EGLN1 | turquoise | -0.040466642 | 0.841165527 |
| ETFDH | turquoise | 0.080134 | 0.697177 |
| KCNAB2 | turquoise | 0.08054 | 0.695711 |
| WDR5B | turquoise | 0.081365 | 0.692745 |
| RAB10 | turquoise | 0.081463 | 0.692392 |
| POLR2J3 | turquoise | 0.081695 | 0.691558 |
| SLC9A1 | turquoise | 0.082201 | 0.689741 |
| IQSEC1 | turquoise | 0.082688 | 0.687991 |
| RANBP3 | turquoise | 0.083593 | 0.684749 |
| MOV10 | turquoise | 0.084066 | 0.683057 |
| KIAA1737 | turquoise | 0.084271 | 0.682324 |
| APOBR | turquoise | 0.084578 | 0.681224 |
| GEN1 | turquoise | 0.085302 | 0.678639 |
| PION | turquoise | 0.085977 | 0.676229 |
| TMEM48 | turquoise | 0.086265 | 0.675205 |
| SMN1 | turquoise | 0.086458 | 0.674516 |
| TUBE1 | turquoise | 0.086797 | 0.673309 |
| IL28RA | turquoise | 0.086797 | 0.673308 |
| R3HCC1L | turquoise | 0.087356 | 0.671319 |
| SCARB2 | turquoise | 0.087562 | 0.670587 |
| ZBTB8OS | turquoise | 0.087727 | 0.670001 |
| ZNRF2 | turquoise | 0.087801 | 0.669738 |
| OGFOD1 | turquoise | 0.088595 | 0.666919 |
| SRSF11 | turquoise | 0.088664 | 0.666676 |
| SNRPE | turquoise | 0.08877 | 0.666299 |
| RAI1 | turquoise | 0.089869 | 0.662408 |
| SYVN1 | turquoise | 0.090919 | 0.658696 |
| DENND4B | turquoise | 0.0915 | 0.656644 |
| NUP54 | turquoise | 0.091773 | 0.655683 |
| RAB18 | turquoise | 0.091932 | 0.655124 |
| FAM73B | turquoise | 0.092789 | 0.652105 |
| DUSP11 | turquoise | 0.092823 | 0.651987 |
| SLC25A45 | turquoise | 0.093313 | 0.650264 |
| TREX1 | turquoise | 0.093969 | 0.647959 |
| FKSG29 | turquoise | 0.094096 | 0.647515 |
| PIGG | turquoise | 0.09419 | 0.647184 |
| MGC2752 | turquoise | 0.09499 | 0.644382 |
| AARS2 | turquoise | 0.095079 | 0.64407 |
| ZNF26 | turquoise | 0.095213 | 0.6436 |
| C7orf55 | turquoise | 0.095223 | 0.643567 |
| C1GALT1 | turquoise | 0.095406 | 0.642928 |
| CYP1B1 | turquoise | 0.096473 | 0.639198 |
| GDPGP1 | turquoise | 0.096531 | 0.638996 |
| SCAF1 | turquoise | 0.097647 | 0.635104 |
| USP14 | turquoise | 0.09887 | 0.630855 |
| UBE2D2 | turquoise | 0.098934 | 0.630631 |
| MOB4 | turquoise | 0.100681 | 0.624577 |
| STX16 | turquoise | 0.101228 | 0.622687 |
| POP5 | turquoise | 0.101633 | 0.621288 |
| ARL1 | turquoise | 0.10188 | 0.620438 |
| ABHD5 | turquoise | 0.101978 | 0.620098 |
| CSNK2B | turquoise | 0.102281 | 0.619053 |
| MBOAT1 | turquoise | 0.102308 | 0.618961 |
| SHISA5 | turquoise | 0.102797 | 0.617279 |
| WBSCR16 | turquoise | 0.103702 | 0.614165 |
| SPTLC1 | turquoise | 0.10403 | 0.613041 |
| SP2 | turquoise | 0.104063 | 0.612929 |

**Table 6. Gene module membership and correlation with growth rate (48/72h), with adjusted p values for each module**

| | | | | | | | | | | | |
|---|---|---|---|---|---|---|---|---|---|---|---|
| C8orf76 | turquoise | 0.089172 | 0.664875 | R3HDM4 | turquoise | -0.040401917 | 0.841416408 | SMPD2 | turquoise | 0.105296 | 0.608703 |
| MALAT1 | turquoise | 0.088997 | 0.665496 | ORC2 | turquoise | -0.040008066 | 0.842943344 | RABL2A | turquoise | 0.105301 | 0.608684 |
| ZNF229 | turquoise | -0.088962 | 0.665621 | ALKBH4 | turquoise | -0.039994261 | 0.842996874 | CCNY | turquoise | 0.105693 | 0.607343 |
| SPRYD7 | turquoise | -0.088891 | 0.66587 | RGPD8 | turquoise | -0.039992292 | 0.843004509 | MARVELD1 | turquoise | 0.106401 | 0.604927 |
| SGSM3 | turquoise | 0.088375 | 0.667701 | TDG | turquoise | -0.039410755 | 0.845260137 | TOLLIP | turquoise | 0.107113 | 0.602498 |
| ESRRA | turquoise | 0.088232 | 0.668207 | AGAP2 | turquoise | -0.03915087 | 0.846268546 | PTGES3 | turquoise | 0.107175 | 0.602287 |
| A4GALT | turquoise | 0.088205 | 0.668303 | PHLPP2 | turquoise | -0.038581854 | 0.848477273 | TOR3A | turquoise | 0.10735 | 0.601691 |
| THUMPD2 | turquoise | -0.088135 | 0.668553 | ALPK3 | turquoise | -0.038072823 | 0.850454109 | SHARPIN | turquoise | 0.107441 | 0.601379 |
| UBR4 | turquoise | -0.087972 | 0.669131 | AASDHPPT | turquoise | -0.037997082 | 0.850748329 | ALMS1 | turquoise | 0.107719 | 0.600433 |
| MLF1 | turquoise | -0.087937 | 0.669256 | RNF144B | turquoise | -0.037387869 | 0.853115535 | PDS5A | turquoise | 0.107723 | 0.600421 |
| SLC26A2 | turquoise | 0.087429 | 0.671062 | LOC1001299( | turquoise | -0.037267968 | 0.853581581 | ZBTB7A | turquoise | 0.107937 | 0.599694 |
| ANXA11 | turquoise | -0.087409 | 0.671132 | FAM221A | turquoise | -0.037235264 | 0.853708706 | RAD51C | turquoise | 0.108403 | 0.598109 |
| FAM98C | turquoise | -0.087383 | 0.671226 | PIH1D1 | turquoise | -0.037088817 | 0.854278013 | UBE2G1 | turquoise | 0.108837 | 0.596634 |
| ATP5L | turquoise | -0.087207 | 0.671851 | FAM91A1 | turquoise | -0.036928188 | 0.854902533 | SAYSD1 | turquoise | 0.109335 | 0.594946 |
| ASMTL | turquoise | 0.087162 | 0.67201 | USP40 | turquoise | -0.036718704 | 0.85571713 | RABGAP1 | turquoise | 0.109359 | 0.594867 |
| CYFIP1 | turquoise | -0.087155 | 0.672035 | MLST8 | turquoise | -0.03641025 | 0.856916843 | GMCL1 | turquoise | 0.110075 | 0.592442 |
| GGA2 | turquoise | -0.087133 | 0.672115 | TTLL4 | turquoise | -0.035845072 | 0.85911587 | CLCN6 | turquoise | 0.110104 | 0.592344 |
| PGM3 | turquoise | -0.086593 | 0.674036 | VPS45 | turquoise | -0.035045226 | 0.862229704 | TRIM62 | turquoise | 0.111178 | 0.588716 |
| SMAD6 | turquoise | 0.086577 | 0.674092 | RRP12 | turquoise | -0.034889511 | 0.862836147 | HEMK1 | turquoise | 0.111285 | 0.588354 |
| MBOAT7 | turquoise | -0.086499 | 0.674372 | UNC5B | turquoise | -0.034158872 | 0.865682668 | DLST | turquoise | 0.112083 | 0.585665 |
| ZNF689 | turquoise | -0.086409 | 0.674689 | TRIM8 | turquoise | -0.033582825 | 0.867928061 | USP37 | turquoise | 0.112111 | 0.585571 |
| BCL7C | turquoise | -0.086361 | 0.67486 | EMC7 | turquoise | -0.033468186 | 0.868375039 | MED16 | turquoise | 0.112653 | 0.583748 |
| ASB6 | turquoise | 0.086314 | 0.675028 | UBQLN1 | turquoise | -0.033266303 | 0.869162274 | ING4 | turquoise | 0.113926 | 0.579474 |
| ZNF292 | turquoise | -0.085674 | 0.677311 | PLSCR1 | turquoise | -0.033169247 | 0.869540781 | CD81 | turquoise | 0.114645 | 0.577069 |
| PIGH | turquoise | 0.085671 | 0.677323 | DKFZP586I14 | turquoise | -0.032921891 | 0.870505577 | SRFBP1 | turquoise | 0.114783 | 0.576607 |
| PPAPDC1B | turquoise | 0.085575 | 0.677665 | CCDC112 | turquoise | -0.032456685 | 0.872320562 | SFPQ | turquoise | 0.115155 | 0.575366 |
| ZNF148 | turquoise | -0.085139 | 0.679221 | CLK1 | turquoise | -0.032285554 | 0.872988386 | AGAP3 | turquoise | 0.11522 | 0.575146 |
| TTYH3 | turquoise | -0.085042 | 0.679568 | SAYSD1 | turquoise | -0.032242467 | 0.873156539 | CASC4 | turquoise | 0.115528 | 0.574119 |
| PUS3 | turquoise | -0.085024 | 0.67963 | PI4KB | turquoise | -0.032106508 | 0.873687185 | NTMT1 | turquoise | 0.115811 | 0.573175 |
| TPP1 | turquoise | -0.085004 | 0.679702 | AZIN1 | turquoise | -0.032005094 | 0.874083036 | KCTD7 | turquoise | 0.115963 | 0.57267 |
| RPLP2 | turquoise | 0.084902 | 0.680068 | RAVER1 | turquoise | -0.031601942 | 0.875656955 | HMG20A | turquoise | 0.115967 | 0.572656 |
| FAM49B | turquoise | -0.08488 | 0.680145 | YPEL2 | turquoise | -0.031564641 | 0.875802603 | MRPL39 | turquoise | 0.11628 | 0.571613 |
| ERI1 | turquoise | -0.084603 | 0.681137 | MINK1 | turquoise | -0.0309202 | 0.87831955 | USP48 | turquoise | 0.116472 | 0.570972 |
| ZBTB44 | turquoise | -0.08456 | 0.681289 | BCAP29 | turquoise | -0.030883918 | 0.878461287 | POLR1B | turquoise | 0.116633 | 0.570437 |
| GAPT | turquoise | -0.084541 | 0.681357 | JMJD1C | turquoise | -0.030737578 | 0.879033017 | SUV39H2 | turquoise | 0.116941 | 0.569414 |
| TMEM2 | turquoise | -0.084325 | 0.682128 | C6orf106 | turquoise | -0.030157742 | 0.881298916 | ZNF202 | turquoise | 0.117443 | 0.567744 |
| CCT8 | turquoise | -0.084229 | 0.682474 | IL7R | turquoise | -0.030010861 | 0.881873047 | ATP13A2 | turquoise | 0.118973 | 0.562674 |
| PFDN4 | turquoise | -0.083838 | 0.683873 | FLOT2 | turquoise | -0.029831616 | 0.882573765 | SIGMAR1 | turquoise | 0.119495 | 0.56095 |
| CCNL1 | turquoise | -0.083819 | 0.683941 | SAMD4B | turquoise | -0.029598895 | 0.883483662 | GAK | turquoise | 0.119516 | 0.560882 |
| CAPN7 | turquoise | -0.083424 | 0.685354 | RAB2A | turquoise | -0.029435873 | 0.884121132 | TAF7 | turquoise | 0.11997 | 0.559384 |
| RRN3P1 | turquoise | 0.08304 | 0.686732 | ZNF282 | turquoise | -0.029298788 | 0.884657235 | PPAPDC1B | turquoise | 0.120446 | 0.557812 |
| SGK223 | turquoise | -0.082871 | 0.687337 | LXN | turquoise | -0.029141871 | 0.885270957 | CUX1 | turquoise | 0.121115 | 0.555613 |
| PRKCI | turquoise | 0.082476 | 0.688752 | ANKRD39 | turquoise | -0.028808196 | 0.88657622 | SBNO2 | turquoise | 0.121381 | 0.554739 |
| POLR2J3 | turquoise | -0.0824 | 0.689025 | ST7 | turquoise | -0.028311425 | 0.888520009 | ADIPOR2 | turquoise | 0.12235 | 0.551558 |
| PHF12 | turquoise | -0.082172 | 0.689844 | ARID1B | turquoise | -0.027446113 | 0.891907335 | PPP1R3B | turquoise | 0.123158 | 0.548914 |
| EXOC8 | turquoise | 0.082102 | 0.690095 | GNB2 | turquoise | -0.027196449 | 0.892885011 | GAS5-AS1 | turquoise | 0.123172 | 0.548868 |
| ZNRF2 | turquoise | 0.081904 | 0.690807 | TEX264 | turquoise | -0.026811696 | 0.894391992 | CCDC126 | turquoise | 0.123189 | 0.548811 |
| MED19 | turquoise | 0.081776 | 0.691268 | ZNF576 | turquoise | -0.026763054 | 0.894582535 | SRCRB4D | turquoise | 0.123369 | 0.548222 |
| POLR2H | turquoise | 0.081737 | 0.691408 | CHCHD3 | turquoise | -0.026624357 | 0.895125878 | POLA1 | turquoise | 0.123854 | 0.546638 |
| RASGRP4 | turquoise | -0.081685 | 0.691593 | ZNF460 | turquoise | -0.026319034 | 0.896322141 | EXOC3 | turquoise | 0.123933 | 0.546383 |
| RANGAP1 | turquoise | -0.081668 | 0.691654 | MAGT1 | turquoise | -0.026251202 | 0.896587941 | APAF1 | turquoise | 0.12399 | 0.546197 |
| SLC12A7 | turquoise | -0.081483 | 0.69232 | GALC | turquoise | -0.025302429 | 0.900306808 | POLE | turquoise | 0.124301 | 0.545183 |
| RPL23 | turquoise | -0.081376 | 0.692706 | CCDC132 | turquoise | -0.024281933 | 0.904309096 | NDST1 | turquoise | 0.124472 | 0.544625 |
| ZC3H13 | turquoise | -0.081303 | 0.692966 | ESRRA | turquoise | -0.024140304 | 0.904864732 | ACOX3 | turquoise | 0.125174 | 0.54234 |
| ARSD | turquoise | 0.081302 | 0.692969 | GPATCH2 | turquoise | -0.023525521 | 0.907277148 | ARHGAP22 | turquoise | 0.125695 | 0.540648 |
| COPS8 | turquoise | -0.08122 | 0.693266 | CCT3 | turquoise | -0.023438524 | 0.907618592 | ABHD12 | turquoise | 0.126174 | 0.539094 |

**Table 6. Gene module membership and correlation with growth rate (48/72h), with adjusted p values for each module**

| | | | | | | | | | | | |
|---|---|---|---|---|---|---|---|---|---|---|---|
| KLHDC2 | turquoise | 0.081122 | 0.693617 | MYCBP | turquoise | -0.022525448 | 0.911203156 | TCF7L2 | turquoise | 0.126869 | 0.53684 |
| SMARCA2 | turquoise | -0.081095 | 0.693714 | HLTF | turquoise | -0.022178908 | 0.912564053 | PABPN1 | turquoise | 0.127146 | 0.535945 |
| RPL37 | turquoise | 0.080706 | 0.695115 | JMJD8 | turquoise | -0.022138979 | 0.912720873 | SEC24B | turquoise | 0.127258 | 0.535585 |
| UBE2K | turquoise | 0.080667 | 0.695254 | GTPBP6 | turquoise | -0.021812644 | 0.914002673 | BAG6 | turquoise | 0.128803 | 0.5306 |
| RBM6 | turquoise | 0.080642 | 0.695345 | POLR2J3 | turquoise | -0.020798575 | 0.917987117 | CIC | turquoise | 0.129048 | 0.529812 |
| C9orf41 | turquoise | -0.080557 | 0.695652 | GPAT2 | turquoise | -0.020521415 | 0.919076465 | IRF2BP1 | turquoise | 0.130326 | 0.52571 |
| RPL26 | turquoise | 0.080493 | 0.695884 | POLE | turquoise | -0.020179246 | 0.920421519 | SLC7A7 | turquoise | 0.130747 | 0.524362 |
| TFDP2 | turquoise | -0.08023 | 0.69683 | FBXO4 | turquoise | -0.0200373 | 0.920979566 | ZNF277 | turquoise | 0.131534 | 0.521848 |
| PWWP2A | turquoise | -0.079856 | 0.698178 | FTSJ2 | turquoise | -0.019876645 | 0.92161121 | NAA16 | turquoise | 0.131642 | 0.521502 |
| CLSTN1 | turquoise | -0.079552 | 0.699274 | UHMK1 | turquoise | -0.019682365 | 0.92237512 | FGFR1OP2 | turquoise | 0.132968 | 0.517277 |
| NARS | turquoise | 0.079539 | 0.699322 | GCN1L1 | turquoise | -0.019109784 | 0.924626901 | CLCN7 | turquoise | 0.134477 | 0.512491 |
| SPINT1 | turquoise | 0.079425 | 0.699731 | HRAS | turquoise | -0.018541321 | 0.926863045 | EXOSC9 | turquoise | 0.135308 | 0.509865 |
| TXNL1 | turquoise | 0.079406 | 0.6998 | LOC1005072 | turquoise | -0.017811331 | 0.92973538 | SLC35A1 | turquoise | 0.135672 | 0.508718 |
| NDUFB4 | turquoise | -0.079201 | 0.700539 | RBM19 | turquoise | -0.016283122 | 0.935751249 | TMEM104 | turquoise | 0.136499 | 0.506113 |
| VMA21 | turquoise | -0.079182 | 0.700609 | ASB13 | turquoise | -0.016220848 | 0.935996467 | ATR | turquoise | 0.136635 | 0.505686 |
| KIAA0226L | turquoise | -0.079125 | 0.700816 | ATF1 | turquoise | -0.015886846 | 0.937311783 | IBA57 | turquoise | 0.136667 | 0.505585 |
| SDHB | turquoise | -0.079058 | 0.701055 | PLK1S1 | turquoise | -0.015688482 | 0.938093026 | ROGDI | turquoise | 0.137077 | 0.504296 |
| TTC19 | turquoise | 0.079004 | 0.701253 | YTHDF1 | turquoise | -0.015364695 | 0.93936836 | MBTD1 | turquoise | 0.138062 | 0.501207 |
| PCMTD1 | turquoise | -0.078986 | 0.701318 | ADCY9 | turquoise | -0.015329686 | 0.939506263 | SOLH | turquoise | 0.138207 | 0.500753 |
| GNAI2 | turquoise | 0.07893 | 0.701519 | POLRMT | turquoise | -0.015088325 | 0.940457043 | SMPD4 | turquoise | 0.138235 | 0.500666 |
| KATNA1 | turquoise | 0.078896 | 0.701643 | HYAL2 | turquoise | -0.014507227 | 0.942746454 | SLC25A39 | turquoise | 0.13866 | 0.499338 |
| GOLGA4 | turquoise | 0.07885 | 0.701807 | DUS3L | turquoise | -0.014298518 | 0.943568833 | ZNFX1 | turquoise | 0.142194 | 0.488358 |
| MVP | turquoise | -0.07859 | 0.702748 | BAG6 | turquoise | -0.013826779 | 0.945427845 | PSMC6 | turquoise | 0.143288 | 0.484982 |
| PSMC6 | turquoise | 0.078571 | 0.702817 | ELF4 | turquoise | -0.013539589 | 0.946559731 | YEATS2 | turquoise | 0.143554 | 0.484164 |
| TRPM7 | turquoise | -0.078563 | 0.702846 | KLF13 | turquoise | -0.013420603 | 0.947028715 | NELF | turquoise | 0.143574 | 0.484101 |
| HOOK3 | turquoise | -0.078419 | 0.703364 | NDUFA6 | turquoise | -0.013296861 | 0.947516458 | TRRAP | turquoise | 0.143627 | 0.48394 |
| WRNIP1 | turquoise | -0.07834 | 0.703651 | FAM92A1 | turquoise | -0.013244362 | 0.947723397 | WIPF2 | turquoise | 0.143702 | 0.483709 |
| PTRH1 | turquoise | 0.078193 | 0.704182 | C15orf48 | turquoise | -0.012892306 | 0.949111199 | RASA4CP | turquoise | 0.144301 | 0.481867 |
| PITPNA | turquoise | 0.078177 | 0.704241 | NUP43 | turquoise | -0.012817616 | 0.949405645 | RBMX | turquoise | 0.144962 | 0.479843 |
| PTGER4 | turquoise | -0.077945 | 0.70508 | S100A10 | turquoise | -0.01241731 | 0.950983854 | FBXO22 | turquoise | 0.145695 | 0.477601 |
| FN3K | turquoise | -0.077913 | 0.705194 | PFDN1 | turquoise | -0.012112361 | 0.952186243 | LOC1001909 | turquoise | 0.146012 | 0.476631 |
| LRRC23 | turquoise | 0.077687 | 0.706012 | SLPI | turquoise | -0.012094538 | 0.95225652 | SAMD4B | turquoise | 0.146699 | 0.474537 |
| PIGC | turquoise | 0.077626 | 0.706233 | METTL6 | turquoise | -0.012024638 | 0.952532143 | PAQR4 | turquoise | 0.147702 | 0.471487 |
| SLC43A2 | turquoise | -0.077388 | 0.707096 | FAM98C | turquoise | -0.011719963 | 0.953733582 | DCAF4 | turquoise | 0.147911 | 0.470854 |
| MTMR2 | turquoise | -0.076952 | 0.708676 | CPQ | turquoise | -0.011445252 | 0.954816946 | INPP5E | turquoise | 0.148058 | 0.470409 |
| LMF2 | turquoise | -0.076924 | 0.708777 | PRKCI | turquoise | -0.011410946 | 0.954952239 | TMEM115 | turquoise | 0.149171 | 0.467041 |
| PLA2G6 | turquoise | -0.076441 | 0.710529 | MRPS5 | turquoise | -0.011235644 | 0.955643617 | ECHDC2 | turquoise | 0.150015 | 0.464495 |
| CEACAM4 | turquoise | 0.07614 | 0.711618 | SETD1A | turquoise | -0.011039672 | 0.956416554 | SIPA1 | turquoise | 0.151751 | 0.459282 |
| HTATIP2 | turquoise | -0.075952 | 0.712302 | EPB41L4A | turquoise | -0.010887551 | 0.957016561 | BCL9L | turquoise | 0.152215 | 0.457896 |
| TTBK2 | turquoise | -0.075876 | 0.712576 | CIAO1 | turquoise | -0.010783341 | 0.957427606 | NEU3 | turquoise | 0.152602 | 0.45674 |
| UGGT1 | turquoise | 0.075664 | 0.713346 | RPTOR | turquoise | -0.010594666 | 0.958171847 | ACOT2 | turquoise | 0.152808 | 0.456126 |
| SETDB2 | turquoise | -0.075601 | 0.713577 | ANKRD6 | turquoise | -0.010586994 | 0.958202108 | FGD4 | turquoise | 0.15418 | 0.452042 |
| KCTD17 | turquoise | -0.075555 | 0.713745 | VPS4B | turquoise | -0.010433365 | 0.958808134 | LOC1001320 | turquoise | 0.154238 | 0.45187 |
| COX17 | turquoise | -0.075407 | 0.714279 | RMND5B | turquoise | -0.009033083 | 0.964332847 | USP11 | turquoise | 0.154626 | 0.450718 |
| RBM4 | turquoise | 0.075312 | 0.714624 | ARHGAP21 | turquoise | -0.008806975 | 0.965225095 | CDK18 | turquoise | 0.155417 | 0.448375 |
| OC1005072 | turquoise | -0.074999 | 0.715762 | CCDC28A | turquoise | -0.008618305 | 0.965969643 | EGFL7 | turquoise | 0.157879 | 0.441131 |
| HMGB2 | turquoise | -0.074895 | 0.71614 | PDIK1L | turquoise | -0.008539792 | 0.966279488 | NCKAP5L | turquoise | 0.158438 | 0.439494 |
| TRAP1 | turquoise | -0.074645 | 0.717051 | PNPO | turquoise | -0.008435362 | 0.966691616 | PDGFC | turquoise | 0.158874 | 0.43822 |
| LDLRAD3 | turquoise | -0.074521 | 0.717502 | SLC9A8 | turquoise | -0.008013889 | 0.968355029 | QSOX2 | turquoise | 0.158951 | 0.437995 |
| ERCC8 | turquoise | 0.074378 | 0.718024 | FAM98A | turquoise | -0.00707789 | 0.972049557 | EWSR1 | turquoise | 0.159296 | 0.436988 |
| UTP23 | turquoise | -0.07422 | 0.718598 | PRPF4 | turquoise | -0.007007604 | 0.97232701 | CANT1 | turquoise | 0.159602 | 0.436098 |
| PPM1K | turquoise | 0.07412 | 0.718962 | GFER | turquoise | -0.00595861 | 0.976468255 | FAM21A | turquoise | 0.159738 | 0.435702 |
| ACSL3 | turquoise | -0.073791 | 0.720161 | LOC1002895 | turquoise | -0.00517041 | 0.979580327 | SLC10A3 | turquoise | 0.160879 | 0.432387 |
| HMGCL | turquoise | 0.073698 | 0.720498 | TOLLIP | turquoise | -0.005009267 | 0.980216613 | BIRC7 | turquoise | 0.160969 | 0.432125 |
| CSNK2A1 | turquoise | 0.073661 | 0.720632 | NARG2 | turquoise | -0.005004686 | 0.9802347 | PLXNA3 | turquoise | 0.162307 | 0.428256 |
| MRPL39 | turquoise | -0.073487 | 0.721268 | DSTN | turquoise | -0.004978626 | 0.9803376 | HAAO | turquoise | 0.163455 | 0.424955 |

**Table 6. Gene module membership and correlation with growth rate (48/72h), with adjusted p values for each module**

| | | | | | | | | | | | |
|---|---|---|---|---|---|---|---|---|---|---|---|
| ZFP91 | turquoise | -0.073326 | 0.721854 | CYBA | turquoise | -0.004877093 | 0.980738519 | GPRIN3 | turquoise | 0.163793 | 0.423982 |
| IWS1 | turquoise | 0.072897 | 0.72342 | DHX40 | turquoise | -0.004534513 | 0.982091273 | C9orf91 | turquoise | 0.165037 | 0.420423 |
| ABHD6 | turquoise | 0.0726 | 0.724502 | IQGAP2 | turquoise | -0.004415436 | 0.98256149 | STARD8 | turquoise | 0.165166 | 0.420056 |
| ZNF197 | turquoise | -0.07199 | 0.726729 | TMEM117 | turquoise | -0.004084471 | 0.983868442 | RAD23A | turquoise | 0.165505 | 0.419089 |
| IL27RA | turquoise | 0.07179 | 0.72746 | UXS1 | turquoise | -0.00397378 | 0.98430556 | ZCWPW1 | turquoise | 0.165587 | 0.418855 |
| HSBP1L1 | turquoise | -0.071743 | 0.727631 | SP2 | turquoise | -0.003597642 | 0.985790961 | UNK | turquoise | 0.165935 | 0.417864 |
| PRKAR2A | turquoise | -0.071543 | 0.728364 | EIF2AK4 | turquoise | -0.003384634 | 0.98663217 | RAB6A | turquoise | 0.166931 | 0.415035 |
| GAPVD1 | turquoise | -0.071215 | 0.729562 | EARS2 | turquoise | -0.002488342 | 0.990171936 | CARD9 | turquoise | 0.168723 | 0.409973 |
| PPCDC | turquoise | -0.071153 | 0.72979 | AGPAT5 | turquoise | -0.001533074 | 0.993944812 | ZFAND1 | turquoise | 0.169719 | 0.407174 |
| OSBPL9 | turquoise | -0.071071 | 0.730089 | ZNF331 | turquoise | -0.001478656 | 0.994159744 | PARP14 | turquoise | 0.170916 | 0.403823 |
| PHRF1 | turquoise | -0.071042 | 0.730197 | MGAT2 | turquoise | -0.001425902 | 0.994368105 | TFE3 | turquoise | 0.172777 | 0.398646 |
| TMCO3 | turquoise | -0.070902 | 0.730707 | SDC2 | turquoise | -0.001369493 | 0.994590901 | DUSP8 | turquoise | 0.176144 | 0.389376 |
| MRPS18B | turquoise | 0.070698 | 0.731454 | NUDT18 | turquoise | -0.000817813 | 0.996769861 | GABPA | turquoise | 0.176544 | 0.388284 |
| SLC4A7 | turquoise | 0.070646 | 0.731644 | TYK2 | turquoise | -0.000797443 | 0.996850316 | ATAT1 | turquoise | 0.177487 | 0.385713 |
| USP11 | turquoise | -0.070507 | 0.732156 | STRN4 | turquoise | -0.000558907 | 0.997792466 | CUL7 | turquoise | 0.177559 | 0.385517 |
| WWC3 | turquoise | -0.070436 | 0.732413 | CCT2 | turquoise | -0.000302387 | 0.998805649 | PSMD14 | turquoise | 0.178261 | 0.383612 |
| GXYLT1 | turquoise | -0.070229 | 0.733173 | MAPK6 | turquoise | -0.000102109 | 0.999596698 | POLDIP3 | turquoise | 0.178443 | 0.383118 |
| MYBBP1A | turquoise | -0.070189 | 0.733317 | CCAR1 | turquoise | -9.84654E-05 | 0.999611087 | IRF3 | turquoise | 0.178872 | 0.381957 |
| MBD4 | turquoise | -0.070187 | 0.733328 | OTUD6B | turquoise | 0.000270816 | 0.998930349 | UBR3 | turquoise | 0.17998 | 0.378966 |
| PTGES3 | turquoise | 0.07008 | 0.733718 | NBR1 | turquoise | 0.00049095 | 0.998060878 | SARM1 | turquoise | 0.180355 | 0.377957 |
| TMEM70 | turquoise | -0.070068 | 0.733762 | DCTN1 | turquoise | 0.000543723 | 0.997852439 | ESRRA | turquoise | 0.181315 | 0.375381 |
| DHX36 | turquoise | -0.070009 | 0.733979 | WDR19 | turquoise | 0.001161896 | 0.99541084 | LRP11 | turquoise | 0.181396 | 0.375165 |
| OSBPL1A | turquoise | -0.069867 | 0.734499 | PAPOLA | turquoise | 0.001545462 | 0.993895887 | PML | turquoise | 0.182994 | 0.370901 |
| PURB | turquoise | 0.069688 | 0.735155 | MYSM1 | turquoise | 0.001960933 | 0.992254944 | ACTN4 | turquoise | 0.183124 | 0.370555 |
| RBM7 | turquoise | 0.069605 | 0.735459 | TRAM1 | turquoise | 0.002060019 | 0.991863598 | TATDN1 | turquoise | 0.1843 | 0.367436 |
| DNASE2 | turquoise | -0.069573 | 0.735578 | SLC4A7 | turquoise | 0.002420295 | 0.990440685 | TBC1D8 | turquoise | 0.184757 | 0.366228 |
| GTF3C6 | turquoise | -0.069412 | 0.736168 | GLUD1 | turquoise | 0.003125045 | 0.987657354 | ZBTB45 | turquoise | 0.185171 | 0.365136 |
| PCGF1 | turquoise | -0.069406 | 0.73619 | RFWD2 | turquoise | 0.004082371 | 0.983876735 | TAF6L | turquoise | 0.186394 | 0.361921 |
| KCNAB2 | turquoise | -0.069181 | 0.737016 | XAF1 | turquoise | 0.004088627 | 0.983852029 | PAK4 | turquoise | 0.187426 | 0.359223 |
| CASZ1 | turquoise | -0.068957 | 0.737836 | SCFD1 | turquoise | 0.004882373 | 0.980717669 | JHDM1D | turquoise | 0.188702 | 0.3559 |
| MAP4 | turquoise | -0.068861 | 0.73819 | ZFYVE21 | turquoise | 0.004949782 | 0.980451496 | TMCO4 | turquoise | 0.189483 | 0.353875 |
| APC | turquoise | -0.068788 | 0.738458 | ADCK1 | turquoise | 0.005368686 | 0.978797441 | TJP2 | turquoise | 0.189623 | 0.353514 |
| SPATA2L | turquoise | 0.06863 | 0.739039 | CLIC2 | turquoise | 0.007166253 | 0.971700752 | ATXN1L | turquoise | 0.189626 | 0.353507 |
| MTOR | turquoise | -0.068542 | 0.739363 | KIAA1107 | turquoise | 0.007569542 | 0.970108859 | KIAA0247 | turquoise | 0.189746 | 0.353196 |
| SFMBT2 | turquoise | -0.068239 | 0.740473 | GAB1 | turquoise | 0.008019582 | 0.968332562 | LPAR2 | turquoise | 0.190422 | 0.351453 |
| LSM1 | turquoise | -0.068231 | 0.740502 | ZFYVE28 | turquoise | 0.00831366 | 0.967171921 | GLA | turquoise | 0.190704 | 0.350725 |
| GHITM | turquoise | -0.06821 | 0.74058 | SBF2 | turquoise | 0.009194406 | 0.963696272 | PKD1L1 | turquoise | 0.193039 | 0.344747 |
| ERP44 | turquoise | -0.0679 | 0.741719 | CASP4 | turquoise | 0.009264255 | 0.963420656 | KCTD10 | turquoise | 0.193583 | 0.343363 |
| TWF1 | turquoise | -0.067895 | 0.74174 | CAPN7 | turquoise | 0.009268096 | 0.963405499 | TBC1D25 | turquoise | 0.194303 | 0.341536 |
| ZBED5 | turquoise | -0.067865 | 0.741849 | QTRT1 | turquoise | 0.009681353 | 0.961774929 | SIAH2 | turquoise | 0.194321 | 0.341489 |
| LUC7L3 | turquoise | 0.067746 | 0.742287 | ID2 | turquoise | 0.009992781 | 0.960546236 | HEATR7A | turquoise | 0.1948 | 0.340277 |
| RPS8 | turquoise | -0.067715 | 0.742399 | TPT1 | turquoise | 0.010027481 | 0.960409338 | PPCDC | turquoise | 0.196349 | 0.336376 |
| SERPING1 | turquoise | -0.067443 | 0.743399 | UHRF1BP1L | turquoise | 0.010450482 | 0.958740611 | TSPYL2 | turquoise | 0.197142 | 0.33439 |
| CCDC82 | turquoise | -0.067253 | 0.744101 | NUP98 | turquoise | 0.010698821 | 0.957760997 | MYO19 | turquoise | 0.197943 | 0.332389 |
| CCDC28A | turquoise | -0.066731 | 0.746023 | SLC4A11 | turquoise | 0.010766184 | 0.957495281 | KIAA0195 | turquoise | 0.198099 | 0.332 |
| ZCCHC10 | turquoise | -0.066628 | 0.7464 | EPT1 | turquoise | 0.010887846 | 0.957015396 | MCFD2 | turquoise | 0.19899 | 0.329787 |
| ZNF574 | turquoise | -0.06651 | 0.746834 | STT3B | turquoise | 0.010888776 | 0.957011731 | ADPRH | turquoise | 0.199288 | 0.329047 |
| PIGO | turquoise | 0.066294 | 0.747631 | BBC3 | turquoise | 0.0109127 | 0.956917363 | MAFK | turquoise | 0.199855 | 0.327645 |
| SCP2 | turquoise | -0.066281 | 0.74768 | CIB1 | turquoise | 0.011419521 | 0.954918424 | ZNF274 | turquoise | 0.201288 | 0.324117 |
| TCERG1 | turquoise | -0.066274 | 0.747707 | JAK2 | turquoise | 0.012070856 | 0.952349898 | GRIN2D | turquoise | 0.202099 | 0.32213 |
| MAN2A1 | turquoise | -0.065844 | 0.749291 | NEK9 | turquoise | 0.012244995 | 0.951663265 | SMAD6 | turquoise | 0.20226 | 0.321735 |
| TMEM9 | turquoise | -0.06574 | 0.749676 | AHI1 | turquoise | 0.012326751 | 0.951340911 | MAVS | turquoise | 0.202382 | 0.321438 |
| ELK4 | turquoise | 0.065704 | 0.749805 | C14orf101 | turquoise | 0.012579899 | 0.950342822 | FICD | turquoise | 0.204002 | 0.317496 |
| ZNF543 | turquoise | 0.065645 | 0.750025 | E2F3 | turquoise | 0.012605008 | 0.950243829 | SNRNP70 | turquoise | 0.20482 | 0.315518 |
| ZC3H14 | turquoise | 0.065636 | 0.750057 | RETSAT | turquoise | 0.012837098 | 0.94932884 | NEO1 | turquoise | 0.205238 | 0.314509 |
| CSDA | turquoise | 0.065408 | 0.7509 | CARKD | turquoise | 0.012846178 | 0.949293046 | NAGLU | turquoise | 0.205376 | 0.314177 |

**Table 6. Gene module membership and correlation with growth rate (48/72h), with adjusted p values for each module**

| | | | |
|---|---|---|---|
| C2orf18 | turquoise | -0.065367 | 0.751051 |
| ADH5 | turquoise | -0.065315 | 0.751241 |
| SUZ12P1 | turquoise | -0.065279 | 0.751377 |
| GTF2H1 | turquoise | -0.065229 | 0.75156 |
| TRIM39 | turquoise | 0.065173 | 0.751767 |
| TTC14 | turquoise | 0.064619 | 0.753812 |
| ANO10 | turquoise | -0.064532 | 0.754135 |
| ACOT7 | turquoise | 0.064488 | 0.754295 |
| HCG18 | turquoise | -0.064358 | 0.754775 |
| SKAP2 | turquoise | -0.064358 | 0.754778 |
| TCTEX1D2 | turquoise | -0.064183 | 0.755423 |
| BICD2 | turquoise | -0.063971 | 0.756208 |
| WBSCR16 | turquoise | 0.063744 | 0.757047 |
| C20orf194 | turquoise | -0.063542 | 0.757796 |
| LYN | turquoise | 0.062764 | 0.760673 |
| HDAC4 | turquoise | 0.062639 | 0.761136 |
| RIF1 | turquoise | -0.06262 | 0.761208 |
| SENP6 | turquoise | -0.062555 | 0.761449 |
| IL7R | turquoise | -0.062431 | 0.761909 |
| LZTR1 | turquoise | -0.062023 | 0.763422 |
| TUT1 | turquoise | 0.061942 | 0.763722 |
| GTPBP3 | turquoise | 0.061601 | 0.764984 |
| COPS3 | turquoise | -0.061525 | 0.765268 |
| ZC3H8 | turquoise | -0.061107 | 0.766817 |
| CHN2 | turquoise | -0.061076 | 0.766932 |
| FAM18B1 | turquoise | 0.060979 | 0.767295 |
| WBP5 | turquoise | -0.060867 | 0.767709 |
| CASP4 | turquoise | -0.060847 | 0.767783 |
| DPH3 | turquoise | -0.060692 | 0.768359 |
| ERICH1 | turquoise | -0.060577 | 0.768785 |
| XRCC4 | turquoise | -0.060517 | 0.769009 |
| LOC286367 | turquoise | 0.059877 | 0.771388 |
| PRKAA1 | turquoise | -0.059634 | 0.772291 |
| UNC50 | turquoise | -0.05959 | 0.772455 |
| C17orf70 | turquoise | 0.059587 | 0.772467 |
| MIS18BP1 | turquoise | -0.05907 | 0.77439 |
| ERO1L | turquoise | -0.058987 | 0.774702 |
| PPP1R3B | turquoise | 0.058985 | 0.774707 |
| EXOC5 | turquoise | -0.058935 | 0.774893 |
| MAN2C1 | turquoise | 0.058926 | 0.774926 |
| BCL2L13 | turquoise | 0.05861 | 0.776105 |
| SHOC2 | turquoise | -0.058412 | 0.77684 |
| RPS14 | turquoise | -0.058315 | 0.777203 |
| DDX51 | turquoise | -0.058208 | 0.7776 |
| CLN5 | turquoise | -0.057919 | 0.77868 |
| FBXW4 | turquoise | -0.057781 | 0.779192 |
| FARP1 | turquoise | 0.057686 | 0.779548 |
| EPB41L1 | turquoise | -0.057594 | 0.779892 |
| WASF2 | turquoise | -0.057588 | 0.779914 |
| LPAR2 | turquoise | -0.057574 | 0.779963 |
| SRSF11 | turquoise | -0.057562 | 0.780008 |
| TEF | turquoise | 0.05733 | 0.780877 |
| ANAPC16 | turquoise | 0.057312 | 0.780941 |
| PSMF1 | turquoise | 0.057273 | 0.781089 |
| ARRB2 | turquoise | -0.057211 | 0.781319 |
| ZNF746 | turquoise | -0.057012 | 0.782062 |
| IPO8 | turquoise | 0.013126456 | 0.948188166 |
| CD80 | turquoise | 0.013602012 | 0.946313698 |
| MRPL47 | turquoise | 0.014004931 | 0.944725759 |
| FCGR1A | turquoise | 0.014090679 | 0.944387845 |
| PPIC | turquoise | 0.014510865 | 0.942732118 |
| PFAS | turquoise | 0.014748124 | 0.941797312 |
| ANAPC4 | turquoise | 0.014944601 | 0.941023242 |
| LOC728613 | turquoise | 0.015220981 | 0.939934465 |
| PPP1R9B | turquoise | 0.015826561 | 0.937549204 |
| ZNF142 | turquoise | 0.015978331 | 0.936951495 |
| GLE1 | turquoise | 0.015978845 | 0.936949472 |
| TOMM34 | turquoise | 0.016421058 | 0.93520811 |
| C16orf87 | turquoise | 0.017172479 | 0.93224981 |
| UBR5 | turquoise | 0.017248946 | 0.931948814 |
| ZNF280D | turquoise | 0.017392667 | 0.931383109 |
| ATF4 | turquoise | 0.017729592 | 0.930057057 |
| FBXO33 | turquoise | 0.017864099 | 0.92952772 |
| DHX36 | turquoise | 0.018157542 | 0.928373012 |
| ADAP1 | turquoise | 0.018352416 | 0.927606256 |
| NME7 | turquoise | 0.018892553 | 0.925481349 |
| GPR89A | turquoise | 0.019003151 | 0.925046317 |
| LOC387647 | turquoise | 0.019241931 | 0.924107159 |
| ATP9B | turquoise | 0.01994164 | 0.921355666 |
| CMPK1 | turquoise | 0.020487722 | 0.919208901 |
| ANKHD1 | turquoise | 0.020499196 | 0.919163798 |
| U2AF1 | turquoise | 0.020560822 | 0.91892157 |
| STAM2 | turquoise | 0.020775844 | 0.918076454 |
| NFKBIB | turquoise | 0.021034286 | 0.917060795 |
| LMBRD2 | turquoise | 0.021351415 | 0.915814673 |
| PHTF2 | turquoise | 0.021596576 | 0.914851472 |
| NUDT22 | turquoise | 0.022455893 | 0.911476285 |
| EIF4A3 | turquoise | 0.022488568 | 0.911347975 |
| SCARNA3 | turquoise | 0.023216566 | 0.908489796 |
| TARBP1 | turquoise | 0.023431604 | 0.90764575 |
| FAR2 | turquoise | 0.023837181 | 0.90605409 |
| CDC42EP1 | turquoise | 0.023900346 | 0.905806233 |
| TTC14 | turquoise | 0.024416915 | 0.903779577 |
| POT1 | turquoise | 0.024528749 | 0.903340894 |
| CDKN2AIPN | turquoise | 0.024894179 | 0.901907643 |
| OSM | turquoise | 0.025040297 | 0.901334639 |
| ARFGEF1 | turquoise | 0.025668782 | 0.89887058 |
| SNORA71D | turquoise | 0.026767096 | 0.894566702 |
| CCNL1 | turquoise | 0.026827683 | 0.894329367 |
| IFRD2 | turquoise | 0.027131263 | 0.893140304 |
| MCOLN1 | turquoise | 0.027400334 | 0.892086594 |
| TOR1A | turquoise | 0.027446286 | 0.891906658 |
| TTC3 | turquoise | 0.027590409 | 0.891342348 |
| ABCD4 | turquoise | 0.02820481 | 0.888937256 |
| MOB3C | turquoise | 0.028512423 | 0.887733459 |
| ZFC3H1 | turquoise | 0.029155594 | 0.885217283 |
| CFL1 | turquoise | 0.029179877 | 0.885122305 |
| ATP2C1 | turquoise | 0.030067019 | 0.88165353 |
| TMED9 | turquoise | 0.030216215 | 0.88107037 |
| SLC25A37 | turquoise | 0.030431201 | 0.88023017 |
| RNASEH1 | turquoise | 0.032422583 | 0.872453636 |
| ISOC2 | turquoise | 0.033023489 | 0.870109278 |
| EOGT | turquoise | 0.207195 | 0.309817 |
| DCAF8 | turquoise | 0.207949 | 0.30802 |
| SIK3 | turquoise | 0.21015 | 0.302812 |
| CNOT3 | turquoise | 0.210706 | 0.301504 |
| LOC388796 | turquoise | 0.21089 | 0.301074 |
| ZNF271 | turquoise | 0.211484 | 0.299682 |
| GAA | turquoise | 0.21163 | 0.299342 |
| TRMT1 | turquoise | 0.212276 | 0.297833 |
| MSH6 | turquoise | 0.21229 | 0.297802 |
| RAVER1 | turquoise | 0.212519 | 0.297267 |
| MED13 | turquoise | 0.213985 | 0.29387 |
| ZNF296 | turquoise | 0.215246 | 0.290965 |
| DGKD | turquoise | 0.215306 | 0.290827 |
| PPM1G | turquoise | 0.215742 | 0.289828 |
| DIS3L | turquoise | 0.215847 | 0.289588 |
| UCK1 | turquoise | 0.216033 | 0.289163 |
| MS4A6E | turquoise | 0.217191 | 0.286523 |
| LZTS2 | turquoise | 0.217588 | 0.285621 |
| FIZ1 | turquoise | 0.219578 | 0.281127 |
| ZBTB48 | turquoise | 0.220228 | 0.279669 |
| C3orf18 | turquoise | 0.223218 | 0.273028 |
| ZSCAN29 | turquoise | 0.224488 | 0.270237 |
| PLBD2 | turquoise | 0.226819 | 0.26516 |
| NACC2 | turquoise | 0.227411 | 0.263882 |
| PER2 | turquoise | 0.228766 | 0.260971 |
| ABCC1 | turquoise | 0.229391 | 0.259633 |
| NSUN4 | turquoise | 0.2312 | 0.25579 |
| BCKDK | turquoise | 0.232192 | 0.253701 |
| PIGH | turquoise | 0.232316 | 0.25344 |
| DAGLA | turquoise | 0.232322 | 0.253426 |
| NOTCH1 | turquoise | 0.233512 | 0.250934 |
| JUND | turquoise | 0.234856 | 0.24814 |
| ATAD1 | turquoise | 0.235179 | 0.247469 |
| TRIM5 | turquoise | 0.235852 | 0.246081 |
| CCS | turquoise | 0.238345 | 0.240977 |
| EDC4 | turquoise | 0.238757 | 0.24014 |
| ANO8 | turquoise | 0.238955 | 0.239739 |
| TNFAIP1 | turquoise | 0.238965 | 0.239718 |
| DDX51 | turquoise | 0.242019 | 0.233584 |
| TULP4 | turquoise | 0.242473 | 0.232682 |
| DPH1 | turquoise | 0.243884 | 0.229891 |
| GSTO1 | turquoise | 0.245447 | 0.226823 |
| PLXNA1 | turquoise | 0.24569 | 0.226349 |
| IFIT5 | turquoise | 0.245809 | 0.226118 |
| DCP1A | turquoise | 0.246494 | 0.224786 |
| TRIM41 | turquoise | 0.246515 | 0.224744 |
| ANKRD6 | turquoise | 0.249311 | 0.219362 |
| SLC4A2 | turquoise | 0.251643 | 0.214938 |
| DYRK1B | turquoise | 0.251878 | 0.214496 |
| ARL6IP6 | turquoise | 0.255724 | 0.207343 |
| CLDN12 | turquoise | 0.258445 | 0.202382 |
| C17orf103 | turquoise | 0.260033 | 0.199524 |
| SPINT1 | turquoise | 0.260136 | 0.199341 |
| ATF6B | turquoise | 0.260844 | 0.198076 |
| WLS | turquoise | 0.270136 | 0.181989 |
| HMGCS1 | turquoise | 0.270587 | 0.181231 |

**Table 6. Gene module membership and correlation with growth rate (48/72h), with adjusted p values for each module**

| | | | |
|---|---|---|---|
| CDK18 | turquoise | -0.05697 | 0.78222 |
| ZNF548 | turquoise | 0.056725 | 0.783133 |
| EVI2A | turquoise | -0.056224 | 0.785006 |
| FAM135A | turquoise | 0.055867 | 0.786339 |
| C9orf123 | turquoise | 0.055495 | 0.787731 |
| TMEM39A | turquoise | 0.05536 | 0.788235 |
| DTWD2 | turquoise | -0.055353 | 0.788261 |
| ANKRD6 | turquoise | 0.055066 | 0.789335 |
| TOLLIP | turquoise | 0.054634 | 0.790949 |
| ALG13 | turquoise | 0.054564 | 0.791211 |
| SAR1A | turquoise | -0.054395 | 0.791846 |
| THRA | turquoise | 0.054352 | 0.792005 |
| FAM134A | turquoise | -0.05434 | 0.792051 |
| CMPK1 | turquoise | -0.054213 | 0.792526 |
| DDHD1 | turquoise | -0.053654 | 0.794619 |
| EAPP | turquoise | -0.053574 | 0.794921 |
| SNORD94 | turquoise | -0.053534 | 0.795071 |
| CLCN6 | turquoise | 0.053369 | 0.795688 |
| CAMTA1 | turquoise | 0.053368 | 0.795694 |
| CERS6 | turquoise | 0.053151 | 0.796507 |
| DYNC1I2 | turquoise | -0.053016 | 0.797014 |
| [illegible]OC1001323[illegible] | turquoise | -0.052958 | 0.797231 |
| ZNF589 | turquoise | -0.052936 | 0.797311 |
| FBXO33 | turquoise | -0.052876 | 0.797537 |
| FAM134B | turquoise | -0.052864 | 0.797582 |
| RASA1 | turquoise | -0.052673 | 0.7983 |
| TMEM126A | turquoise | -0.052667 | 0.798323 |
| RAB3GAP2 | turquoise | -0.052632 | 0.798453 |
| TM2D1 | turquoise | -0.052537 | 0.798809 |
| KIF1B | turquoise | 0.052529 | 0.798839 |
| SDHAF2 | turquoise | 0.051974 | 0.800924 |
| TBC1D24 | turquoise | -0.051971 | 0.800934 |
| NUP50 | turquoise | -0.051923 | 0.801115 |
| FKBP3 | turquoise | -0.05181 | 0.80154 |
| SDF2 | turquoise | 0.051788 | 0.801622 |
| VASN | turquoise | -0.051602 | 0.802318 |
| NCK1 | turquoise | -0.051593 | 0.802353 |
| PION | turquoise | -0.051204 | 0.803815 |
| CCDC163P | turquoise | -0.050855 | 0.805126 |
| STXBP3 | turquoise | -0.050756 | 0.805499 |
| FAM210A | turquoise | 0.050691 | 0.805742 |
| MGAT2 | turquoise | -0.050683 | 0.805776 |
| DMXL1 | turquoise | -0.050505 | 0.806443 |
| EBAG9 | turquoise | -0.050131 | 0.80785 |
| EIF2AK4 | turquoise | 0.050082 | 0.808037 |
| PRPF39 | turquoise | -0.049789 | 0.809137 |
| SNX13 | turquoise | -0.049583 | 0.809914 |
| SNW1 | turquoise | 0.049232 | 0.811237 |
| RAB2A | turquoise | -0.049195 | 0.811374 |
| CYP2S1 | turquoise | 0.048873 | 0.812588 |
| NFXL1 | turquoise | -0.048855 | 0.812655 |
| PICALM | turquoise | 0.048464 | 0.814127 |
| ASXL1 | turquoise | 0.048261 | 0.814895 |
| CD4 | turquoise | -0.048219 | 0.815054 |
| ELL3 | turquoise | -0.048203 | 0.815114 |
| VRK2 | turquoise | -0.047917 | 0.81619 |
| RRN3P1 | turquoise | 0.033159225 | 0.869579869 |
| IQSEC3 | turquoise | 0.033572922 | 0.867966674 |
| ZNF616 | turquoise | 0.034177515 | 0.865610013 |
| HNRNPA0 | turquoise | 0.0346268 | 0.863859459 |
| PDLIM2 | turquoise | 0.034657861 | 0.863738458 |
| PDXP | turquoise | 0.037687508 | 0.851951077 |
| CHIC2 | turquoise | 0.037760528 | 0.851667351 |
| LAS1L | turquoise | 0.039286728 | 0.845741359 |
| TMEM39A | turquoise | 0.039600317 | 0.844524745 |
| NUDCD1 | turquoise | 0.040144051 | 0.842416075 |
| SLC15A3 | turquoise | 0.04023636 | 0.842058196 |
| VPS33A | turquoise | 0.04064749 | 0.84046462 |
| ARRDC2 | turquoise | 0.040704171 | 0.840244969 |
| MTM1 | turquoise | 0.041149956 | 0.838517855 |
| ERI1 | turquoise | 0.041260386 | 0.838090128 |
| PIAS1 | turquoise | 0.04144642 | 0.837369663 |
| UTP23 | turquoise | 0.041524825 | 0.837066057 |
| HSDL2 | turquoise | 0.042236514 | 0.834311254 |
| PDSS2 | turquoise | 0.042341045 | 0.833906795 |
| FAM118B | turquoise | 0.042525893 | 0.833191672 |
| NLRP1 | turquoise | 0.04260802 | 0.832873985 |
| POLDIP3 | turquoise | 0.042644852 | 0.83273152 |
| ATP13A2 | turquoise | 0.04279763 | 0.832140631 |
| UBE2D1 | turquoise | 0.043004519 | 0.831340604 |
| TFDP2 | turquoise | 0.043074859 | 0.83106864 |
| GPR157 | turquoise | 0.043137164 | 0.830827761 |
| CEP104 | turquoise | 0.043318925 | 0.830125128 |
| NUPL1 | turquoise | 0.043786504 | 0.828318202 |
| GTF2H3 | turquoise | 0.043850201 | 0.828072116 |
| UBE2J2 | turquoise | 0.043942615 | 0.827715112 |
| TMCO3 | turquoise | 0.044062413 | 0.827252369 |
| SNAI3 | turquoise | 0.044461571 | 0.825710955 |
| PPP2R1A | turquoise | 0.044645553 | 0.825000694 |
| CNEP1R1 | turquoise | 0.045459992 | 0.821858167 |
| BIN2 | turquoise | 0.045596334 | 0.82133235 |
| SPATS2 | turquoise | 0.046304261 | 0.818603371 |
| LYL1 | turquoise | 0.046678481 | 0.817161625 |
| DNAAF2 | turquoise | 0.046706206 | 0.817054833 |
| DNPEP | turquoise | 0.046859247 | 0.816465401 |
| CMTM6 | turquoise | 0.047827002 | 0.812740403 |
| ZXDB | turquoise | 0.04945782 | 0.806472221 |
| VAC14 | turquoise | 0.050065965 | 0.804137725 |
| MALAT1 | turquoise | 0.050223298 | 0.803534037 |
| TAF13 | turquoise | 0.050291909 | 0.803270808 |
| KIAA0528 | turquoise | 0.05033185 | 0.803117584 |
| MED29 | turquoise | 0.05047977 | 0.802550183 |
| ZNF266 | turquoise | 0.050508836 | 0.802438701 |
| SLAMF7 | turquoise | 0.051309261 | 0.799370189 |
| OGDH | turquoise | 0.051822515 | 0.797404106 |
| ERP44 | turquoise | 0.051914946 | 0.797050162 |
| DAP3 | turquoise | 0.053110219 | 0.792476719 |
| IMPA2 | turquoise | 0.053236056 | 0.791995617 |
| MRPS24 | turquoise | 0.053548609 | 0.790800987 |
| OSCAR | turquoise | 0.054004521 | 0.789059239 |
| SLC25A25 | turquoise | 0.054467599 | 0.787291129 |
| GOLPH3L | turquoise | 0.054840585 | 0.785867752 |
| ZNF317 | turquoise | 0.275603 | 0.172959 |
| MYD88 | turquoise | 0.277392 | 0.170074 |
| QPCTL | turquoise | 0.280322 | 0.165423 |
| ZNF776 | turquoise | 0.281037 | 0.1643 |
| IFITM3 | turquoise | 0.283218 | 0.160913 |
| INTS1 | turquoise | 0.287946 | 0.153739 |
| IFITM1 | turquoise | 0.291346 | 0.148721 |
| CXorf56 | turquoise | 0.294923 | 0.143571 |
| SLC12A4 | turquoise | 0.296533 | 0.141296 |
| MED22 | turquoise | 0.306261 | 0.128093 |
| FBXO6 | turquoise | 0.307575 | 0.126382 |
| MIER2 | turquoise | 0.316971 | 0.114624 |
| RIT1 | turquoise | 0.317699 | 0.113747 |
| SLC16A13 | turquoise | 0.333477 | 0.095948 |
| FBXL2 | turquoise | 0.334482 | 0.09489 |
| HEXDC | turquoise | 0.334899 | 0.094453 |
| KIAA1432 | turquoise | 0.335594 | 0.093728 |
| WDR6 | turquoise | 0.338709 | 0.090533 |
| SNF8 | turquoise | 0.349024 | 0.080535 |
| BRAT1 | turquoise | 0.354972 | 0.075165 |
| CROCCP2 | turquoise | 0.357707 | 0.072789 |
| TRMT10B | turquoise | 0.360376 | 0.070527 |
| LENG9 | turquoise | 0.369158 | 0.063462 |
| ANKRD54 | turquoise | 0.38346 | 0.053142 |
| VPS16 | turquoise | 0.395704 | 0.045393 |
| CNP | turquoise | 0.398766 | 0.043602 |
| PCK2 | turquoise | 0.409217 | 0.037905 |
| FCHSD1 | turquoise | 0.431307 | 0.02781 |
| TNK2 | turquoise | 0.478064 | 0.013504 |
| LOC339803 | violet | -0.275927 | 0.172434 |
| COMMD2 | violet | -0.235284 | 0.247252 |
| ALDH3A2 | violet | -0.231675 | 0.25479 |
| CCDC84 | violet | -0.223457 | 0.272501 |
| P2RY1 | violet | -0.21753 | 0.285752 |
| MACC1 | violet | -0.211046 | 0.300707 |
| JAKMIP2 | violet | -0.205913 | 0.312886 |
| C10orf137 | violet | -0.19884 | 0.330158 |
| NBPF1 | violet | -0.197379 | 0.333798 |
| SLC39A4 | violet | -0.196554 | 0.335862 |
| DNAH1 | violet | -0.195661 | 0.338106 |
| PER3 | violet | -0.191799 | 0.347913 |
| SLC38A6 | violet | -0.185983 | 0.363 |
| GRB10 | violet | -0.184145 | 0.367845 |
| FARS2 | violet | -0.164445 | 0.422115 |
| MITF | violet | -0.163738 | 0.424142 |
| QTRTD1 | violet | -0.161847 | 0.429584 |
| LGALSL | violet | -0.154461 | 0.451206 |
| FAM149B1 | violet | -0.14912 | 0.467194 |
| PPIP5K1 | violet | -0.148312 | 0.469638 |
| PDXK | violet | -0.129266 | 0.52911 |
| SNX24 | violet | -0.090359 | 0.660673 |
| LYZ | violet | -0.088841 | 0.666048 |
| CDC42EP3 | violet | -0.078575 | 0.702801 |
| TCEA3 | violet | -0.065049 | 0.752223 |
| MICAL1 | violet | -0.037458 | 0.855841 |
| TMEM9 | violet | 0.009322 | 0.96395 |

**Table 6. Gene module membership and correlation with growth rate (48/72h), with adjusted p values for each module**

| | | | | | | | | | | | |
|---|---|---|---|---|---|---|---|---|---|---|---|
| SFXN3 | turquoise | -0.04776 | 0.816784 | BNIP2 | turquoise | 0.054944974 | 0.785469509 | GADD45G | violet | 0.011672 | 0.95487 |
| CLCN7 | turquoise | -0.047548 | 0.817585 | CLPTM1L | turquoise | 0.055056684 | 0.785043389 | SIL1 | violet | 0.023938 | 0.907594 |
| POP5 | turquoise | -0.046988 | 0.819695 | FAM18B1 | turquoise | 0.055832432 | 0.782085978 | KIF16B | violet | 0.049237 | 0.811216 |
| TRMT1 | turquoise | 0.046603 | 0.821152 | SLC19A3 | turquoise | 0.056348936 | 0.780118528 | ITM2B | violet | 0.060322 | 0.769733 |
| SKP1 | turquoise | -0.046393 | 0.821946 | CYB5A | turquoise | 0.056587135 | 0.779211633 | SNORA68 | violet | 0.089402 | 0.664062 |
| CNOT1 | turquoise | -0.046389 | 0.821961 | ZCCHC10 | turquoise | 0.056727482 | 0.778677421 | MOB3B | violet | 0.097063 | 0.63714 |
| FOLR3 | turquoise | -0.046378 | 0.821999 | GRN | turquoise | 0.056819493 | 0.778327247 | SMAP2 | violet | 0.099108 | 0.630027 |
| CRLF3 | turquoise | -0.046154 | 0.822845 | ANKRD54 | turquoise | 0.057209125 | 0.776844859 | FBXO25 | violet | 0.183511 | 0.369527 |
| SLIRP | turquoise | 0.04577 | 0.824298 | RGPD6 | turquoise | 0.057710858 | 0.774937099 | LRCH4 | violet | 0.184422 | 0.367113 |
| FOXO3B | turquoise | -0.045527 | 0.825218 | SFXN3 | turquoise | 0.057960264 | 0.773989246 | SNORA84 | violet | 0.185651 | 0.363872 |
| SIRT3 | turquoise | -0.045353 | 0.825874 | PELI1 | turquoise | 0.058274317 | 0.772796155 | PELI2 | violet | 0.208243 | 0.307321 |
| HSPA9 | turquoise | 0.045303 | 0.826063 | LOC728730 | turquoise | 0.058481445 | 0.772009548 | FAM129A | violet | 0.214109 | 0.293584 |
| CHIC2 | turquoise | 0.04527 | 0.826189 | GATAD1 | turquoise | 0.058614856 | 0.771503011 | SCARNA7 | violet | 0.227238 | 0.264255 |
| SLC33A1 | turquoise | 0.045237 | 0.826312 | HPCAL1 | turquoise | 0.059178392 | 0.769364385 | PRKCDBP | violet | 0.243674 | 0.230304 |
| B3GALNT1 | turquoise | 0.04509 | 0.826869 | KDM5A | turquoise | 0.060614318 | 0.763922499 | ABTB1 | violet | 0.259834 | 0.19988 |
| BCL7B | turquoise | -0.045074 | 0.826928 | NAA50 | turquoise | 0.061838881 | 0.759290255 | SNORA33 | violet | 0.295794 | 0.142337 |
| ZNF417 | turquoise | -0.044924 | 0.827497 | OPA1 | turquoise | 0.062620883 | 0.756336344 | DCLRE1B | violet | 0.31508 | 0.116922 |
| ZBTB8OS | turquoise | -0.044899 | 0.827594 | SLC39A7 | turquoise | 0.063398408 | 0.753402657 | FAM100B | violet | 0.507589 | 0.008121 |
| HCFC1R1 | turquoise | -0.044888 | 0.827635 | ERCC8 | turquoise | 0.063646371 | 0.752467762 | PDHB | white | -0.390673 | 0.048461 |
| TMCC1 | turquoise | 0.044757 | 0.82813 | CTBP1 | turquoise | 0.063909623 | 0.751475601 | ANAPC15 | white | -0.356 | 0.074265 |
| ITIH5 | turquoise | -0.044628 | 0.828616 | PIGC | turquoise | 0.065027198 | 0.747267912 | KLF7 | white | -0.31276 | 0.11979 |
| VKORC1 | turquoise | -0.044596 | 0.82874 | PLBD2 | turquoise | 0.065744875 | 0.744569557 | MAF | white | -0.285149 | 0.157955 |
| NOTCH1 | turquoise | 0.044294 | 0.829882 | ZNF638 | turquoise | 0.065750185 | 0.744549606 | FOS | white | -0.284395 | 0.159105 |
| SNRPG | turquoise | -0.044188 | 0.830284 | RAB10 | turquoise | 0.065755121 | 0.744531058 | C1orf162 | white | -0.282234 | 0.162435 |
| RUSC1 | turquoise | 0.044185 | 0.830295 | KAT6B | turquoise | 0.065801416 | 0.744357097 | ABHD13 | white | -0.268635 | 0.184523 |
| TMEM30A | turquoise | -0.044057 | 0.830778 | TBC1D23 | turquoise | 0.066327677 | 0.742380477 | PABPC1 | white | -0.259002 | 0.201376 |
| PSTPIP2 | turquoise | -0.043593 | 0.832535 | NEU1 | turquoise | 0.067135843 | 0.739348133 | PIGM | white | -0.255425 | 0.207893 |
| PPAT | turquoise | 0.043501 | 0.832884 | CDC123 | turquoise | 0.067734613 | 0.73710392 | KIAA1143 | white | -0.218147 | 0.284353 |
| PMM2 | turquoise | -0.043483 | 0.832954 | APAF1 | turquoise | 0.067958819 | 0.736264126 | LRP1 | white | -0.205104 | 0.314833 |
| BDP1 | turquoise | -0.043424 | 0.833178 | ZNF226 | turquoise | 0.067963385 | 0.736247025 | RPL13P5 | white | -0.203615 | 0.318437 |
| BNIP2 | turquoise | -0.043349 | 0.83346 | MEF2A | turquoise | 0.068280158 | 0.735061023 | ZNF710 | white | -0.201025 | 0.324763 |
| MRPL2 | turquoise | -0.043244 | 0.83386 | NETO2 | turquoise | 0.068472648 | 0.734340627 | TET3 | white | -0.193947 | 0.342437 |
| SOLH | turquoise | 0.043167 | 0.834152 | ZFYVE16 | turquoise | 0.068548214 | 0.734057881 | TAPBPL | white | -0.187807 | 0.358229 |
| SRCRB4D | turquoise | 0.042853 | 0.83534 | SPINT1 | turquoise | 0.068728111 | 0.733384892 | SH2B1 | white | -0.186781 | 0.360909 |
| NUDCD1 | turquoise | 0.042348 | 0.837255 | TUBGCP3 | turquoise | 0.068772887 | 0.733217418 | CNN2 | white | -0.185918 | 0.363172 |
| NDUFB8 | turquoise | -0.042169 | 0.837933 | RPAP2 | turquoise | 0.068799319 | 0.733118561 | ELMO1 | white | -0.172231 | 0.400163 |
| SHISA5 | turquoise | -0.04183 | 0.839219 | PURB | turquoise | 0.068961207 | 0.732513178 | EIF4G3 | white | -0.168779 | 0.409815 |
| ZNF498 | turquoise | 0.041764 | 0.83947 | RBM3 | turquoise | 0.069101699 | 0.731987929 | AXL | white | -0.161158 | 0.431577 |
| TCF20 | turquoise | -0.041751 | 0.83952 | LAMTOR3 | turquoise | 0.06928556 | 0.731300721 | GAB3 | white | -0.156663 | 0.4447 |
| PQLC1 | turquoise | -0.041594 | 0.840117 | JUNB | turquoise | 0.06944339 | 0.730710969 | SKP2 | white | -0.147259 | 0.472834 |
| ATE1 | turquoise | 0.041369 | 0.840968 | LAMTOR4 | turquoise | 0.070216369 | 0.727824796 | ZNF510 | white | -0.142874 | 0.486257 |
| KNTC1 | turquoise | 0.041273 | 0.841335 | TBC1D8 | turquoise | 0.070375561 | 0.727230846 | AP2A2 | white | -0.140616 | 0.493245 |
| UBE2L3 | turquoise | 0.041178 | 0.841696 | HINT3 | turquoise | 0.07080336 | 0.725635486 | ZNF217 | white | -0.138575 | 0.499604 |
| POLE | turquoise | 0.041127 | 0.84189 | CXorf38 | turquoise | 0.070925793 | 0.725179107 | DOK3 | white | -0.121299 | 0.555009 |
| PTPN2 | turquoise | -0.040816 | 0.843068 | TRMT1L | turquoise | 0.071246171 | 0.723985314 | RPS28 | white | -0.117341 | 0.568084 |
| MGEA5 | turquoise | -0.040457 | 0.844432 | DHRS11 | turquoise | 0.07137647 | 0.723499974 | ZC2HC1A | white | -0.117122 | 0.568812 |
| ZFAND3 | turquoise | -0.04041 | 0.844612 | NHP2 | turquoise | 0.072528496 | 0.719213441 | STRBP | white | -0.111883 | 0.58634 |
| CES1 | turquoise | -0.040408 | 0.844619 | DDA1 | turquoise | 0.072631407 | 0.718830924 | UQCC | white | -0.104028 | 0.613046 |
| PHLPP2 | turquoise | -0.040397 | 0.84466 | NEU3 | turquoise | 0.073039197 | 0.717315831 | GGA2 | white | -0.101084 | 0.623183 |
| FKBP5 | turquoise | 0.040285 | 0.845088 | ARID4A | turquoise | 0.073174014 | 0.716815166 | ARHGAP18 | white | -0.095275 | 0.643384 |
| CTNNAL1 | turquoise | 0.040227 | 0.845309 | PIK3R1 | turquoise | 0.07340612 | 0.715953467 | ZZEF1 | white | -0.080565 | 0.695624 |
| PLBD2 | turquoise | 0.040216 | 0.84535 | ICOS | turquoise | 0.073712864 | 0.714815194 | PCTP | white | -0.07542 | 0.714232 |
| RAC2 | turquoise | 0.040213 | 0.84536 | PIP4K2C | turquoise | 0.073969469 | 0.713863437 | ZFHX3 | white | -0.065056 | 0.752199 |
| KLHL26 | turquoise | -0.040205 | 0.84539 | IFNAR2 | turquoise | 0.073983811 | 0.713810252 | ZNF417 | white | -0.055435 | 0.787952 |
| GATAD1 | turquoise | -0.040161 | 0.845558 | ASB6 | turquoise | 0.074089028 | 0.71342013 | ZNF592 | white | -0.048461 | 0.81414 |
| PPP6R1 | turquoise | 0.040049 | 0.845986 | PPCDC | turquoise | 0.074372224 | 0.712370443 | SNX29 | white | -0.045868 | 0.823926 |

**Table 6. Gene module membership and correlation with growth rate (48/72h), with adjusted p values for each module**

| | | | | | | | | | | | |
|---|---|---|---|---|---|---|---|---|---|---|---|
| PER2 | turquoise | -0.039981 | 0.846243 | CRCP | turquoise | 0.074997354 | 0.710055168 | RASGRP4 | white | -0.045734 | 0.824435 |
| ATG4A | turquoise | -0.039856 | 0.846717 | ZDHHC4 | turquoise | 0.076678626 | 0.70384079 | CCPG1 | white | -0.042185 | 0.837873 |
| CAMSAP2 | turquoise | -0.039635 | 0.847557 | RPS6KA1 | turquoise | 0.077367744 | 0.701298978 | AMD1 | white | -0.039016 | 0.849912 |
| NCOA3 | turquoise | -0.039371 | 0.848561 | UBQLN2 | turquoise | 0.077798329 | 0.699712349 | TSC2 | white | -0.036923 | 0.857882 |
| ATAD3B | turquoise | 0.03935 | 0.84864 | C17orf103 | turquoise | 0.078810138 | 0.695988871 | PDIA3P | white | -0.0334 | 0.871326 |
| GDF15 | turquoise | -0.039346 | 0.848657 | C5orf24 | turquoise | 0.07885693 | 0.695816842 | LOC152217 | white | -0.030304 | 0.883169 |
| USP8 | turquoise | -0.038633 | 0.851369 | TSPAN31 | turquoise | 0.079731458 | 0.692604364 | SLC4A11 | white | -0.015796 | 0.938951 |
| ACAP3 | turquoise | -0.038481 | 0.851948 | CD101 | turquoise | 0.080079022 | 0.691329065 | PLCB2 | white | -0.013491 | 0.947849 |
| PAPOLA | turquoise | -0.038083 | 0.853462 | TAF12 | turquoise | 0.080753876 | 0.688855205 | VAPA | white | -0.003291 | 0.987269 |
| ANKRD13D | turquoise | -0.038067 | 0.853522 | MTA3 | turquoise | 0.081132665 | 0.687468017 | ARHGAP27 | white | -0.002796 | 0.989186 |
| AGTPBP1 | turquoise | -0.037938 | 0.854015 | EID1 | turquoise | 0.081894309 | 0.684681738 | GALNT10 | white | 0.005869 | 0.977298 |
| UBE2D2 | turquoise | 0.037932 | 0.854038 | RIPK2 | turquoise | 0.082788476 | 0.681415792 | ARF4 | white | 0.009533 | 0.963134 |
| RCHY1 | turquoise | -0.037857 | 0.854321 | ATG7 | turquoise | 0.083093296 | 0.680303713 | KCTD12 | white | 0.015142 | 0.941476 |
| R3HDM1 | turquoise | -0.037591 | 0.855335 | ABCC1 | turquoise | 0.083122768 | 0.680196224 | CD63 | white | 0.027973 | 0.892102 |
| TMEM41B | turquoise | -0.037131 | 0.857089 | SLC12A4 | turquoise | 0.083149671 | 0.68009811 | DCTN2 | white | 0.036094 | 0.861041 |
| FBXL18 | turquoise | -0.037004 | 0.857573 | SF3B1 | turquoise | 0.083394154 | 0.679206729 | PUM1 | white | 0.041677 | 0.839799 |
| RGPD8 | turquoise | 0.036988 | 0.857632 | INTS10 | turquoise | 0.083418954 | 0.679116331 | ARHGAP32 | white | 0.049911 | 0.80868 |
| SPTLC1 | turquoise | 0.03684 | 0.858197 | PPP2R1B | turquoise | 0.083876255 | 0.677450229 | CTSK | white | 0.057596 | 0.779883 |
| SART3 | turquoise | -0.036829 | 0.858238 | DNAJC18 | turquoise | 0.084862185 | 0.673863188 | SDF4 | white | 0.084308 | 0.682192 |
| DNAJC2 | turquoise | -0.036719 | 0.85866 | ARHGAP5 | turquoise | 0.084958328 | 0.67351377 | CCDC28A | white | 0.103923 | 0.613408 |
| FAM206A | turquoise | -0.036466 | 0.859623 | FAM49B | turquoise | 0.085180735 | 0.672705712 | FBXW11 | white | 0.109258 | 0.595207 |
| SFSWAP | turquoise | 0.035967 | 0.861527 | CINP | turquoise | 0.085414849 | 0.671855505 | TOMM34 | white | 0.111239 | 0.58851 |
| CACUL1 | turquoise | 0.035809 | 0.862129 | CRLF3 | turquoise | 0.085527873 | 0.671445187 | ATP5B | white | 0.138056 | 0.501228 |
| SETD5 | turquoise | 0.035521 | 0.863228 | PCNA | turquoise | 0.08703976 | 0.665965386 | C1QBP | white | 0.146121 | 0.476298 |
| ARHGAP18 | turquoise | -0.03543 | 0.863572 | ZMYM1 | turquoise | 0.087374049 | 0.664756005 | C12orf57 | white | 0.149758 | 0.46527 |
| BTBD10 | turquoise | 0.035237 | 0.864308 | PIAS2 | turquoise | 0.08748258 | 0.664363541 | KHDRBS1 | white | 0.172207 | 0.400229 |
| C6orf62 | turquoise | -0.034663 | 0.866503 | VPS28 | turquoise | 0.087563399 | 0.664071343 | SPIRE2 | white | 0.177977 | 0.384383 |
| CRYBB2P1 | turquoise | -0.034544 | 0.866957 | CBL | turquoise | 0.087636097 | 0.66380855 | ANGPT4 | white | 0.190676 | 0.350799 |
| ODF2 | turquoise | 0.034542 | 0.866964 | HRSP12 | turquoise | 0.087932035 | 0.662739165 | EIF1B | white | 0.218062 | 0.284547 |
| ZNF337 | turquoise | 0.034317 | 0.867823 | FGR | turquoise | 0.088110848 | 0.662093331 | PNRC2 | white | 0.218988 | 0.282456 |
| ATXN3 | turquoise | -0.034254 | 0.868061 | TBC1D25 | turquoise | 0.088987244 | 0.658931395 | ANXA7 | white | 0.257209 | 0.204625 |
| ITIH4 | turquoise | -0.033997 | 0.869045 | WBSCR16 | turquoise | 0.089115711 | 0.658468376 | MARS2 | white | 0.261437 | 0.19702 |
| KIAA0020 | turquoise | -0.033782 | 0.869867 | DNAJB6 | turquoise | 0.089387331 | 0.657489819 | ZSCAN16 | white | 0.269938 | 0.182322 |
| INO80E | turquoise | -0.033236 | 0.871953 | TMEM165 | turquoise | 0.089924505 | 0.655556181 | KCTD18 | white | 0.384111 | 0.052705 |
| PSMD12 | turquoise | -0.033056 | 0.87264 | ZFP36L2 | turquoise | 0.090590917 | 0.653160343 | PDCD6IP | yellow | -0.618209 | 0.000763 |
| VPS13C | turquoise | -0.033052 | 0.872656 | EVI2A | turquoise | 0.09063002 | 0.653019867 | TUBB | yellow | -0.49652 | 0.009877 |
| KIAA1522 | turquoise | -0.032998 | 0.87286 | POLR3GL | turquoise | 0.091172178 | 0.651073369 | SLC7A6 | yellow | -0.49535 | 0.01008 |
| POLR2B | turquoise | -0.032997 | 0.872865 | ZMYM6 | turquoise | 0.091233871 | 0.650852016 | SNHG8 | yellow | -0.466854 | 0.0162 |
| ABHD12 | turquoise | -0.032238 | 0.875766 | HMOX2 | turquoise | 0.091234525 | 0.650849667 | ACOT4 | yellow | -0.443847 | 0.023125 |
| SMAD5 | turquoise | 0.032178 | 0.875998 | RCHY1 | turquoise | 0.091520841 | 0.64982275 | CKAP2 | yellow | -0.438852 | 0.024908 |
| ZNF513 | turquoise | -0.032092 | 0.876323 | NADSYN1 | turquoise | 0.091859652 | 0.648608356 | EMC1 | yellow | -0.426619 | 0.029747 |
| KLF3 | turquoise | -0.031538 | 0.878444 | ZNF83 | turquoise | 0.091944677 | 0.64830374 | YWHAB | yellow | -0.423189 | 0.031232 |
| CXCL9 | turquoise | -0.031215 | 0.879681 | TUBA1A | turquoise | 0.092125051 | 0.647657701 | CBY1 | yellow | -0.420652 | 0.032368 |
| JMJD1C | turquoise | -0.031193 | 0.879764 | KPNB1 | turquoise | 0.092584242 | 0.646014161 | POLR3F | yellow | -0.418334 | 0.033434 |
| RUFY3 | turquoise | -0.03088 | 0.880962 | ZMYM2 | turquoise | 0.092859434 | 0.645029969 | VTI1B | yellow | -0.416963 | 0.034078 |
| TMEM117 | turquoise | -0.030829 | 0.881157 | HSPA13 | turquoise | 0.093842592 | 0.641518594 | CAPN2 | yellow | -0.416837 | 0.034138 |
| DENND4B | turquoise | 0.030467 | 0.882542 | ERGIC2 | turquoise | 0.094067468 | 0.640716497 | MZT2B | yellow | -0.414148 | 0.035432 |
| DNAJA2 | turquoise | 0.030466 | 0.882549 | CCDC7 | turquoise | 0.096081285 | 0.633551158 | GLUL | yellow | -0.405043 | 0.040105 |
| ALAD | turquoise | -0.030444 | 0.882633 | CYTIP | turquoise | 0.096384449 | 0.63247524 | DNMBP | yellow | -0.402245 | 0.041635 |
| ZNF668 | turquoise | 0.030246 | 0.88339 | USP8 | turquoise | 0.097024044 | 0.630207742 | GLIPR1 | yellow | -0.401089 | 0.042281 |
| SRFBP1 | turquoise | 0.030201 | 0.883563 | AGPAT6 | turquoise | 0.097072315 | 0.630036741 | INTS6 | yellow | -0.397154 | 0.044538 |
| AP4E1 | turquoise | -0.029208 | 0.887366 | BAX | turquoise | 0.097204711 | 0.629567829 | TMEM222 | yellow | -0.397007 | 0.044624 |
| BLMH | turquoise | -0.029198 | 0.887404 | MAN2A2 | turquoise | 0.09826276 | 0.625825527 | RPS15A | yellow | -0.393607 | 0.046653 |
| SLC27A1 | turquoise | 0.029045 | 0.88799 | LACE1 | turquoise | 0.098672156 | 0.624379919 | RPL35A | yellow | -0.392895 | 0.047086 |
| LOC152217 | turquoise | -0.028991 | 0.888196 | HIC2 | turquoise | 0.098872761 | 0.623672061 | MRPL52 | yellow | -0.38876 | 0.04967 |
| KDM5A | turquoise | 0.028758 | 0.88909 | KBTBD8 | turquoise | 0.099300383 | 0.62216424 | HIST1H3E | yellow | -0.381487 | 0.054482 |

**Table 6. Gene module membership and correlation with growth rate (48/72h), with adjusted p values for each module**

| | | | |
|---|---|---|---|
| GSTO1 | turquoise | -0.028406 | 0.890441 |
| RCE1 | turquoise | -0.028073 | 0.891717 |
| BUB3 | turquoise | -0.028069 | 0.891734 |
| KLF16 | turquoise | -0.02805 | 0.891805 |
| GPATCH4 | turquoise | 0.027912 | 0.892334 |
| PGAP1 | turquoise | -0.027858 | 0.892544 |
| NETO2 | turquoise | -0.027402 | 0.894293 |
| OMA1 | turquoise | -0.027307 | 0.894654 |
| HINT3 | turquoise | -0.027078 | 0.895533 |
| TRIM25 | turquoise | -0.026949 | 0.896028 |
| TASP1 | turquoise | -0.0269 | 0.896217 |
| GUSBP11 | turquoise | 0.026556 | 0.897538 |
| MEF2A | turquoise | -0.026509 | 0.897719 |
| SLC35A1 | turquoise | -0.026468 | 0.897877 |
| JHDM1D | turquoise | -0.026203 | 0.898892 |
| NMD3 | turquoise | -0.026018 | 0.899605 |
| MOB4 | turquoise | 0.025893 | 0.900084 |
| DAGLA | turquoise | -0.025671 | 0.900936 |
| EPB41L2 | turquoise | -0.02559 | 0.901247 |
| EMR4P | turquoise | 0.02555 | 0.901401 |
| DNPEP | turquoise | -0.0255 | 0.901593 |
| TNFRSF10C | turquoise | 0.025407 | 0.901951 |
| FRY | turquoise | -0.025346 | 0.902183 |
| DPH1 | turquoise | -0.025274 | 0.902461 |
| GPI | turquoise | -0.025197 | 0.902755 |
| ODC1 | turquoise | 0.025004 | 0.903497 |
| PTK2 | turquoise | 0.024893 | 0.903924 |
| ARF4 | turquoise | 0.024418 | 0.90575 |
| PDCD7 | turquoise | 0.02432 | 0.906124 |
| SMC5 | turquoise | 0.024164 | 0.906725 |
| KIAA0196 | turquoise | 0.023935 | 0.907606 |
| LRRK2 | turquoise | -0.023825 | 0.90803 |
| ESF1 | turquoise | -0.023701 | 0.908507 |
| CRLS1 | turquoise | -0.023698 | 0.908519 |
| KIAA0247 | turquoise | 0.023642 | 0.908732 |
| RRP15 | turquoise | -0.023544 | 0.90911 |
| MTFR1 | turquoise | -0.02349 | 0.909316 |
| CWF19L1 | turquoise | -0.023463 | 0.90942 |
| RAB30 | turquoise | 0.023448 | 0.90948 |
| SNX16 | turquoise | -0.023444 | 0.909493 |
| UBE2D1 | turquoise | -0.023418 | 0.909594 |
| INTS1 | turquoise | 0.02338 | 0.90974 |
| FAM208A | turquoise | -0.023349 | 0.909859 |
| PHF3 | turquoise | 0.023336 | 0.909909 |
| CWC22 | turquoise | -0.023091 | 0.910849 |
| PHF6 | turquoise | -0.022343 | 0.913726 |
| ROMO1 | turquoise | -0.022294 | 0.913917 |
| UBE3B | turquoise | -0.02226 | 0.914048 |
| TCTEX1D4 | turquoise | 0.022218 | 0.914208 |
| GTF2H5 | turquoise | -0.021975 | 0.915144 |
| FTO | turquoise | -0.021869 | 0.915552 |
| AMPD2 | turquoise | -0.021772 | 0.915924 |
| HCK | turquoise | 0.021656 | 0.916371 |
| UNK | turquoise | 0.021515 | 0.916913 |
| SLC25A35 | turquoise | 0.021464 | 0.917112 |
| CFD | turquoise | 0.021121 | 0.91843 |
| COPS7A | turquoise | 0.099391581 | 0.621842861 |
| CUL4B | turquoise | 0.099668264 | 0.620868256 |
| FAM135A | turquoise | 0.100145283 | 0.619189441 |
| MBD1 | turquoise | 0.100550436 | 0.617765015 |
| RTTN | turquoise | 0.100682457 | 0.617301148 |
| ARHGAP22 | turquoise | 0.100807305 | 0.616862619 |
| CSTF3 | turquoise | 0.100872813 | 0.616632571 |
| FAM206A | turquoise | 0.100989051 | 0.616224463 |
| EIF2S3 | turquoise | 0.10116841 | 0.615594953 |
| SMCHD1 | turquoise | 0.101500491 | 0.614430126 |
| PDCD2L | turquoise | 0.101561719 | 0.61421546 |
| BCL7C | turquoise | 0.101631486 | 0.613970893 |
| ALAD | turquoise | 0.101788147 | 0.613421864 |
| SRSF11 | turquoise | 0.10194733 | 0.612864207 |
| DCAF13 | turquoise | 0.102075201 | 0.612416396 |
| MAD2L2 | turquoise | 0.102200403 | 0.611978062 |
| DGCR6L | turquoise | 0.102217854 | 0.611916976 |
| SERPINH1 | turquoise | 0.102270607 | 0.611732337 |
| GBF1 | turquoise | 0.102530476 | 0.610823101 |
| HSPA9 | turquoise | 0.103597316 | 0.607096333 |
| SCARB2 | turquoise | 0.104276757 | 0.60472784 |
| CWF19L2 | turquoise | 0.104299538 | 0.604648496 |
| FICD | turquoise | 0.104453964 | 0.604110746 |
| SQSTM1 | turquoise | 0.104521434 | 0.603875864 |
| ZNRF2 | turquoise | 0.104723246 | 0.603173526 |
| HMG20B | turquoise | 0.104772508 | 0.603002137 |
| C7orf73 | turquoise | 0.104908124 | 0.602530422 |
| HSBP1L1 | turquoise | 0.105187258 | 0.601559999 |
| STK38L | turquoise | 0.105449597 | 0.600648568 |
| GOLM1 | turquoise | 0.105464452 | 0.600596976 |
| UNC45A | turquoise | 0.105531049 | 0.600365701 |
| MTFR1 | turquoise | 0.106670994 | 0.596412865 |
| HEATR5A | turquoise | 0.106703208 | 0.596301322 |
| CYLD | turquoise | 0.10691966 | 0.59555208 |
| C20orf94 | turquoise | 0.107580252 | 0.593267951 |
| GORASP1 | turquoise | 0.107767439 | 0.592621402 |
| CDC14A | turquoise | 0.107972862 | 0.591912212 |
| SLC35A1 | turquoise | 0.108793823 | 0.589081639 |
| RPS19 | turquoise | 0.109016579 | 0.588314616 |
| HDAC3 | turquoise | 0.109487888 | 0.58669317 |
| ZNF7 | turquoise | 0.109512463 | 0.586608679 |
| RAP1B | turquoise | 0.109922573 | 0.585199448 |
| TAB2 | turquoise | 0.110684173 | 0.582586346 |
| EBAG9 | turquoise | 0.111453944 | 0.579950414 |
| KLHDC2 | turquoise | 0.111754534 | 0.57892253 |
| MKNK2 | turquoise | 0.112777158 | 0.575431639 |
| ACAT1 | turquoise | 0.113110628 | 0.574295307 |
| PAK4 | turquoise | 0.113790651 | 0.571981157 |
| KCTD5 | turquoise | 0.115074969 | 0.56762195 |
| ARID2 | turquoise | 0.1151467 | 0.567378921 |
| VMA21 | turquoise | 0.116646395 | 0.562308651 |
| NCLN | turquoise | 0.11675162 | 0.56195367 |
| FGFR1OP2 | turquoise | 0.118282676 | 0.556800147 |
| ALCAM | turquoise | 0.118540435 | 0.555934665 |
| SPSB1 | turquoise | 0.118641739 | 0.555594683 |
| DBR1 | turquoise | 0.119097739 | 0.554065504 |
| SRSF2 | yellow | -0.381217 | 0.054667 |
| KIAA0494 | yellow | -0.381037 | 0.054791 |
| RPUSD1 | yellow | -0.379924 | 0.055562 |
| DDT | yellow | -0.37583 | 0.058471 |
| UBE4A | yellow | -0.375269 | 0.058878 |
| ABR | yellow | -0.371302 | 0.061824 |
| DNAJB1 | yellow | -0.370415 | 0.062497 |
| ORAI2 | yellow | -0.363216 | 0.068179 |
| HIST1H2BC | yellow | -0.362874 | 0.068459 |
| INPP1 | yellow | -0.362821 | 0.068502 |
| DARS2 | yellow | -0.361426 | 0.069652 |
| CPPED1 | yellow | -0.361361 | 0.069706 |
| REEP4 | yellow | -0.360692 | 0.070263 |
| TMEM64 | yellow | -0.357949 | 0.072582 |
| NEIL2 | yellow | -0.356879 | 0.073502 |
| MCU | yellow | -0.355597 | 0.074617 |
| LOC541471 | yellow | -0.3549 | 0.075229 |
| ARPP19 | yellow | -0.352567 | 0.077303 |
| DTD1 | yellow | -0.350653 | 0.079036 |
| NRBP1 | yellow | -0.350441 | 0.07923 |
| RPL10 | yellow | -0.348411 | 0.081104 |
| LOC92249 | yellow | -0.347885 | 0.081596 |
| C5orf22 | yellow | -0.347573 | 0.081888 |
| DAGLB | yellow | -0.346991 | 0.082436 |
| RNASEH2B | yellow | -0.346949 | 0.082475 |
| RRAS | yellow | -0.346795 | 0.082621 |
| MRP63 | yellow | -0.345283 | 0.084059 |
| SNX18 | yellow | -0.344792 | 0.08453 |
| BCAP31 | yellow | -0.343011 | 0.086255 |
| NDUFB10 | yellow | -0.342002 | 0.087245 |
| KLF10 | yellow | -0.34107 | 0.088166 |
| RIC8A | yellow | -0.338452 | 0.090794 |
| RASSF2 | yellow | -0.337499 | 0.091765 |
| WRAP73 | yellow | -0.334229 | 0.095155 |
| SEPW1 | yellow | -0.334143 | 0.095245 |
| ALG8 | yellow | -0.333825 | 0.09558 |
| C10orf54 | yellow | -0.333183 | 0.096259 |
| C1QC | yellow | -0.333129 | 0.096316 |
| APLP2 | yellow | -0.33237 | 0.097124 |
| IFT140 | yellow | -0.331498 | 0.098058 |
| PTP4A2 | yellow | -0.331217 | 0.09836 |
| PHF21A | yellow | -0.330918 | 0.098683 |
| RPS27 | yellow | -0.330021 | 0.099656 |
| TPMT | yellow | -0.329827 | 0.099867 |
| WDR74 | yellow | -0.329769 | 0.09993 |
| UBE2A | yellow | -0.329443 | 0.100286 |
| FAM103A1 | yellow | -0.329089 | 0.100673 |
| CHD1L | yellow | -0.328326 | 0.101512 |
| RAF1 | yellow | -0.325694 | 0.104447 |
| TCEA1 | yellow | -0.325497 | 0.104668 |
| RPL5 | yellow | -0.324789 | 0.10547 |
| CHAMP1 | yellow | -0.323763 | 0.10664 |
| MGAT5 | yellow | -0.322566 | 0.108016 |
| DHX57 | yellow | -0.322402 | 0.108205 |
| RPL13AP20 | yellow | -0.32156 | 0.109183 |
| STUB1 | yellow | -0.320239 | 0.110728 |

**Table 6. Gene module membership and correlation with growth rate (48/72h), with adjusted p values for each module**

| | | | | | | | | | | | |
|---|---|---|---|---|---|---|---|---|---|---|---|
| ARPC5 | turquoise | 0.020919 | 0.91921 | MRPL21 | turquoise | 0.119544271 | 0.552569949 | SNORA48 | yellow | -0.320155 | 0.110827 |
| VAMP8 | turquoise | -0.020711 | 0.92001 | TCIRG1 | turquoise | 0.119606706 | 0.552360984 | PLEKHJ1 | yellow | -0.319853 | 0.111184 |
| FMNL2 | turquoise | 0.020636 | 0.920296 | GNA13 | turquoise | 0.120026239 | 0.550957798 | RASSF3 | yellow | -0.319374 | 0.11175 |
| CAP1 | turquoise | -0.020636 | 0.920297 | TXNL1 | turquoise | 0.120396355 | 0.549721262 | BRPF1 | yellow | -0.318328 | 0.112994 |
| SEMA3C | turquoise | 0.020522 | 0.920735 | KIAA0930 | turquoise | 0.121591258 | 0.545737945 | SLC35B2 | yellow | -0.317932 | 0.113468 |
| TRRAP | turquoise | 0.020433 | 0.921079 | TSPO | turquoise | 0.121805173 | 0.545026263 | RPS9 | yellow | -0.317666 | 0.113786 |
| C11orf58 | turquoise | 0.020344 | 0.921421 | HDAC5 | turquoise | 0.122719787 | 0.541988276 | VPRBP | yellow | -0.317666 | 0.113787 |
| EIF2AK2 | turquoise | -0.020324 | 0.921498 | SNX17 | turquoise | 0.123442397 | 0.539593677 | THOC3 | yellow | -0.317172 | 0.11438 |
| GPR89C | turquoise | -0.02027 | 0.921705 | YWHAH | turquoise | 0.124762599 | 0.535231642 | MIDN | yellow | -0.316634 | 0.115031 |
| DAB2 | turquoise | -0.020233 | 0.921851 | PTTG1IP | turquoise | 0.125965053 | 0.531273233 | RPS14 | yellow | -0.316207 | 0.115548 |
| SMCHD1 | turquoise | 0.020228 | 0.92187 | GPR153 | turquoise | 0.126206485 | 0.530480134 | TPD52 | yellow | -0.316141 | 0.115628 |
| EIF2S3 | turquoise | -0.020204 | 0.921963 | SMPD4 | turquoise | 0.126654218 | 0.52901084 | TMEM106A | yellow | -0.315808 | 0.116034 |
| TNF | turquoise | 0.019776 | 0.923608 | SEPHS2 | turquoise | 0.126751523 | 0.52869178 | RNPS1 | yellow | -0.314689 | 0.117402 |
| SYVN1 | turquoise | 0.01974 | 0.923747 | MLPH | turquoise | 0.126907336 | 0.52818106 | SETD9 | yellow | -0.31463 | 0.117474 |
| THBS1 | turquoise | -0.019686 | 0.923955 | PATL1 | turquoise | 0.126957573 | 0.528016448 | TMBIM6 | yellow | -0.314333 | 0.11784 |
| PLCG2 | turquoise | 0.019269 | 0.925564 | YAE1D1 | turquoise | 0.127122552 | 0.527476024 | MCRS1 | yellow | -0.314186 | 0.118021 |
| AMFR | turquoise | -0.018872 | 0.927094 | TREM2 | turquoise | 0.128017614 | 0.524548682 | RHOA | yellow | -0.312175 | 0.12052 |
| SUGP1 | turquoise | 0.018557 | 0.928306 | PUS3 | turquoise | 0.12812501 | 0.524197963 | ACP2 | yellow | -0.312159 | 0.120541 |
| MED8 | turquoise | -0.018532 | 0.928401 | ORAOV1 | turquoise | 0.128486175 | 0.523019353 | GLRX | yellow | -0.311074 | 0.121905 |
| RAP1B | turquoise | 0.018401 | 0.928905 | RBMS2 | turquoise | 0.128523006 | 0.522899232 | PPP1R35 | yellow | -0.310855 | 0.122182 |
| ATG12 | turquoise | 0.018356 | 0.929082 | UGGT1 | turquoise | 0.128559939 | 0.522778792 | COPS5 | yellow | -0.309716 | 0.123628 |
| NUDT19 | turquoise | -0.018179 | 0.929761 | ZNF589 | turquoise | 0.129654032 | 0.51921698 | GID8 | yellow | -0.308843 | 0.124746 |
| RPL22 | turquoise | 0.018161 | 0.929834 | TBC1D20 | turquoise | 0.129839955 | 0.518612879 | ERLIN2 | yellow | -0.308786 | 0.124818 |
| HEATR5A | turquoise | 0.018009 | 0.930417 | FKTN | turquoise | 0.12992672 | 0.51833108 | TMEM230 | yellow | -0.308212 | 0.125558 |
| PJA2 | turquoise | -0.017928 | 0.930729 | RAB3IP | turquoise | 0.12993254 | 0.51831218 | XRN2 | yellow | -0.308177 | 0.125603 |
| MST4 | turquoise | 0.017879 | 0.93092 | HSF1 | turquoise | 0.130141821 | 0.517632784 | CALHM2 | yellow | -0.307923 | 0.125931 |
| FER | turquoise | -0.017866 | 0.930971 | OSBPL2 | turquoise | 0.132477544 | 0.510079728 | ARMCX3 | yellow | -0.307436 | 0.126562 |
| NCDN | turquoise | -0.017827 | 0.931121 | TSR2 | turquoise | 0.132809848 | 0.50900957 | LRR1 | yellow | -0.307227 | 0.126834 |
| ASH1L-AS1 | turquoise | 0.017603 | 0.931984 | PHF6 | turquoise | 0.132850325 | 0.508879292 | COX8A | yellow | -0.30693 | 0.12722 |
| MRPL33 | turquoise | -0.017597 | 0.932005 | ZADH2 | turquoise | 0.133064164 | 0.508191311 | HPS4 | yellow | -0.305997 | 0.128439 |
| IPO4 | turquoise | 0.017334 | 0.933021 | MED13 | turquoise | 0.13312384 | 0.507999398 | EMD | yellow | -0.30494 | 0.129831 |
| MFAP1 | turquoise | 0.017262 | 0.933299 | RCC1 | turquoise | 0.133460541 | 0.506917265 | ANPEP | yellow | -0.304746 | 0.130088 |
| LMBRD2 | turquoise | -0.017235 | 0.9334 | HSD17B4 | turquoise | 0.13407195 | 0.50495515 | PSMC5 | yellow | -0.304229 | 0.130774 |
| H6PD | turquoise | -0.017209 | 0.933504 | CCDC66 | turquoise | 0.134972129 | 0.502073177 | H2AFY | yellow | -0.302219 | 0.133465 |
| CCNT2 | turquoise | -0.01718 | 0.933615 | LRWD1 | turquoise | 0.13515746 | 0.501480847 | POLR3H | yellow | -0.301865 | 0.133944 |
| TBCA | turquoise | -0.016988 | 0.934356 | C19orf25 | turquoise | 0.136237307 | 0.498036495 | ALG12 | yellow | -0.301635 | 0.134255 |
| LILRB2 | turquoise | -0.016887 | 0.934743 | PQLC1 | turquoise | 0.136550665 | 0.497039204 | FBXW8 | yellow | -0.301485 | 0.134458 |
| CD58 | turquoise | -0.01684 | 0.934926 | ADNP2 | turquoise | 0.136729142 | 0.496471633 | PEX14 | yellow | -0.301284 | 0.134731 |
| SIPA1L2 | turquoise | -0.01657 | 0.935967 | TM9SF3 | turquoise | 0.136860715 | 0.496053427 | POLR1D | yellow | -0.301279 | 0.134737 |
| HOMER1 | turquoise | -0.016418 | 0.936552 | PAPSS1 | turquoise | 0.137619727 | 0.493644348 | MFAP5 | yellow | -0.300138 | 0.136294 |
| COPS2 | turquoise | 0.016239 | 0.937243 | PTPLA | turquoise | 0.137769514 | 0.493169623 | USP46 | yellow | -0.299943 | 0.136561 |
| PLAU | turquoise | 0.015958 | 0.938326 | BLMH | turquoise | 0.138203926 | 0.491794132 | UNC13D | yellow | -0.299777 | 0.136789 |
| PPP2R1B | turquoise | 0.015678 | 0.939407 | CD58 | turquoise | 0.139339441 | 0.488207863 | ACTL6A | yellow | -0.298497 | 0.138555 |
| RABGAP1L | turquoise | -0.015575 | 0.939806 | TMEM62 | turquoise | 0.139509943 | 0.487670514 | ACYP2 | yellow | -0.298242 | 0.138908 |
| TAF9 | turquoise | -0.01556 | 0.939862 | IKZF2 | turquoise | 0.139572641 | 0.487472993 | C6orf57 | yellow | -0.297768 | 0.139567 |
| CD74 | turquoise | 0.015443 | 0.940312 | PLAU | turquoise | 0.141915744 | 0.480120548 | FAM168A | yellow | -0.297392 | 0.140092 |
| EIF5 | turquoise | -0.015433 | 0.940353 | TSNARE1 | turquoise | 0.142538174 | 0.478177005 | RPL12 | yellow | -0.297 | 0.14064 |
| PHC2 | turquoise | 0.014982 | 0.942091 | BCL2L1 | turquoise | 0.143015683 | 0.476688722 | CD68 | yellow | -0.296366 | 0.141531 |
| STK38L | turquoise | -0.014939 | 0.942259 | LZTS2 | turquoise | 0.143300676 | 0.4758016 | ATP1B1 | yellow | -0.295486 | 0.142772 |
| TLN1 | turquoise | 0.014923 | 0.942321 | STBD1 | turquoise | 0.143956216 | 0.473764278 | C8orf33 | yellow | -0.295375 | 0.14293 |
| CEP85L | turquoise | -0.014884 | 0.94247 | KIAA1429 | turquoise | 0.14507038 | 0.470311971 | ARSA | yellow | -0.29348 | 0.145633 |
| PPM1G | turquoise | -0.014877 | 0.942497 | RAB6A | turquoise | 0.145497832 | 0.468990956 | TRIM47 | yellow | -0.292147 | 0.147557 |
| MFAP3 | turquoise | -0.014797 | 0.942807 | OTUD5 | turquoise | 0.145998209 | 0.467447018 | ATG9A | yellow | -0.292065 | 0.147676 |
| UBE2V1 | turquoise | -0.014689 | 0.943222 | CFDP1 | turquoise | 0.146129535 | 0.467042244 | C12orf52 | yellow | -0.291079 | 0.149111 |
| ANAPC7 | turquoise | 0.014638 | 0.943418 | LMLN | turquoise | 0.146873739 | 0.464751904 | KCNQ1 | yellow | -0.290926 | 0.149335 |
| XBP1 | turquoise | -0.014452 | 0.944137 | C7orf60 | turquoise | 0.147774995 | 0.461986096 | COPA | yellow | -0.290535 | 0.149908 |

**Table 6. Gene module membership and correlation with growth rate (48/72h), with adjusted p values for each module**

| | | | | | | | | | | | |
|---|---|---|---|---|---|---|---|---|---|---|---|
| REPS1 | turquoise | 0.01432 | 0.944648 | YEATS2 | turquoise | 0.148862175 | 0.45866123 | HIST1H4B | yellow | -0.289746 | 0.151068 |
| FARSA | turquoise | -0.014219 | 0.945037 | G6PD | turquoise | 0.149288427 | 0.457361089 | CTTNBP2NL | yellow | -0.289251 | 0.151799 |
| ATAD2B | turquoise | -0.014171 | 0.945223 | GATC | turquoise | 0.149924069 | 0.455425885 | MSANTD4 | yellow | -0.288506 | 0.152903 |
| RAB10 | turquoise | -0.013951 | 0.94607 | EFHD2 | turquoise | 0.150124455 | 0.454816712 | TCF3 | yellow | -0.288146 | 0.153439 |
| SGK3 | turquoise | 0.013951 | 0.946072 | MAPKAPK3 | turquoise | 0.150734959 | 0.452963438 | ZNF649 | yellow | -0.287155 | 0.154922 |
| VKORC1L1 | turquoise | 0.013562 | 0.947573 | SLC35B3 | turquoise | 0.150837296 | 0.45265317 | ATXN2 | yellow | -0.285388 | 0.157591 |
| CYP3A5 | turquoise | 0.013509 | 0.947778 | TJP2 | turquoise | 0.150903287 | 0.452453158 | COL8A2 | yellow | -0.284847 | 0.158414 |
| SEC31A | turquoise | -0.01326 | 0.94874 | ATF6B | turquoise | 0.151252922 | 0.451394228 | GTF3C5 | yellow | -0.284184 | 0.159427 |
| ZNF319 | turquoise | 0.01319 | 0.949011 | SMC5 | turquoise | 0.152008928 | 0.449109039 | RNF170 | yellow | -0.283434 | 0.16058 |
| CDC42EP1 | turquoise | 0.013158 | 0.949133 | AKAP17A | turquoise | 0.15208271 | 0.448886349 | SNAPC4 | yellow | -0.282841 | 0.161495 |
| ITPR2 | turquoise | -0.01314 | 0.949203 | IRF5 | turquoise | 0.152948145 | 0.446278659 | IRF8 | yellow | -0.282457 | 0.162089 |
| ZNF274 | turquoise | -0.013018 | 0.949673 | AMFR | turquoise | 0.155125984 | 0.439752454 | LMNB2 | yellow | -0.282325 | 0.162293 |
| CEP104 | turquoise | 0.013015 | 0.949684 | MAGOH | turquoise | 0.155245336 | 0.439396293 | TTI1 | yellow | -0.281449 | 0.163657 |
| SH3YL1 | turquoise | 0.01267 | 0.951017 | MGAT4A | turquoise | 0.155551991 | 0.438481905 | ARPC1A | yellow | -0.281109 | 0.164187 |
| COG7 | turquoise | 0.012546 | 0.951497 | DHRS3 | turquoise | 0.155776177 | 0.437814077 | SLC38A9 | yellow | -0.280409 | 0.165285 |
| UBE2Q1 | turquoise | -0.012471 | 0.951786 | RNF121 | turquoise | 0.156178197 | 0.43661787 | MRPL35 | yellow | -0.280024 | 0.165891 |
| DAZAP1 | turquoise | -0.011911 | 0.953949 | NSUN2 | turquoise | 0.156364123 | 0.436065249 | STRN | yellow | -0.279099 | 0.167352 |
| SLC39A8 | turquoise | -0.011668 | 0.954888 | FRA10AC1 | turquoise | 0.156851736 | 0.43461773 | TMUB1 | yellow | -0.2785 | 0.168305 |
| TRMT11 | turquoise | -0.011641 | 0.954992 | PSMD9 | turquoise | 0.158378771 | 0.430101497 | PITHD1 | yellow | -0.277339 | 0.170159 |
| DLST | turquoise | -0.011593 | 0.955177 | SUN1 | turquoise | 0.159590431 | 0.42653627 | FAM109A | yellow | -0.275927 | 0.172435 |
| CCNK | turquoise | -0.011249 | 0.956505 | WRNIP1 | turquoise | 0.159648118 | 0.426366935 | MED30 | yellow | -0.275916 | 0.172453 |
| TMEM208 | turquoise | -0.011241 | 0.956537 | STAT5B | turquoise | 0.160089863 | 0.425071447 | ZNF529 | yellow | -0.274741 | 0.174362 |
| PGM5 | turquoise | 0.011225 | 0.956596 | TOMM40 | turquoise | 0.160726509 | 0.423208181 | C15orf61 | yellow | -0.274491 | 0.174771 |
| HMG20A | turquoise | 0.010995 | 0.957487 | POP4 | turquoise | 0.160815807 | 0.422947196 | JMJD7 | yellow | -0.274245 | 0.175173 |
| CRABP2 | turquoise | -0.010979 | 0.957548 | HNRNPF | turquoise | 0.161691481 | 0.420392584 | CDKN1B | yellow | -0.274114 | 0.175387 |
| SNX30 | turquoise | -0.010953 | 0.957649 | AKTIP | turquoise | 0.162239949 | 0.418796879 | AGPAT2 | yellow | -0.273152 | 0.176968 |
| DSTYK | turquoise | -0.010725 | 0.95853 | AMZ2 | turquoise | 0.162936953 | 0.416773857 | RIN1 | yellow | -0.272591 | 0.177894 |
| STK3 | turquoise | 0.01067 | 0.958741 | MSMO1 | turquoise | 0.163881153 | 0.414042009 | ZMAT3 | yellow | -0.272429 | 0.178162 |
| DPYD | turquoise | -0.010421 | 0.959703 | MOSPD1 | turquoise | 0.16474675 | 0.411546344 | PLEKHM2 | yellow | -0.271419 | 0.179842 |
| SSX2IP | turquoise | -0.010379 | 0.959867 | GBA | turquoise | 0.165384847 | 0.409711979 | LBX2-AS1 | yellow | -0.271229 | 0.180158 |
| CSNK2B | turquoise | 0.009727 | 0.962387 | UNC50 | turquoise | 0.167993946 | 0.402259126 | POLE4 | yellow | -0.271009 | 0.180525 |
| DYNC1LI1 | turquoise | 0.009396 | 0.963665 | TMED1 | turquoise | 0.168351932 | 0.401242535 | HACL1 | yellow | -0.270281 | 0.181746 |
| KFZP586I14 | turquoise | 0.009349 | 0.963847 | HDAC10 | turquoise | 0.168393311 | 0.401125121 | METTL17 | yellow | -0.270164 | 0.181941 |
| STRN3 | turquoise | -0.009205 | 0.964403 | CD99L2 | turquoise | 0.168912082 | 0.399654756 | CCDC137 | yellow | -0.268164 | 0.185324 |
| ZBTB41 | turquoise | -0.009133 | 0.96468 | EIF3H | turquoise | 0.168935619 | 0.399588115 | TTC5 | yellow | -0.267841 | 0.185874 |
| ARHGAP21 | turquoise | -0.00909 | 0.964849 | EXOSC7 | turquoise | 0.169521148 | 0.397932354 | DNAJC14 | yellow | -0.267808 | 0.18593 |
| PPP3R1 | turquoise | 0.009039 | 0.965046 | PDE4A | turquoise | 0.169590104 | 0.397737616 | POLR3C | yellow | -0.267418 | 0.186597 |
| ZNF880 | turquoise | 0.008622 | 0.966658 | SH3BP1 | turquoise | 0.169804926 | 0.397131282 | NUP88 | yellow | -0.266307 | 0.188505 |
| CSK | turquoise | 0.007848 | 0.969648 | NOP58 | turquoise | 0.170579561 | 0.394949233 | NR2F6 | yellow | -0.264844 | 0.191035 |
| CHM | turquoise | -0.007843 | 0.969668 | RAB43 | turquoise | 0.170758164 | 0.394447098 | RPUSD4 | yellow | -0.264747 | 0.191205 |
| LDLRAP1 | turquoise | -0.007756 | 0.970004 | ZMIZ1 | turquoise | 0.172744936 | 0.38888586 | TTC12 | yellow | -0.264393 | 0.191821 |
| RASGRP3 | turquoise | -0.007727 | 0.970116 | KIAA1737 | turquoise | 0.173353643 | 0.387191018 | FGD2 | yellow | -0.263551 | 0.193291 |
| TMEM184B | turquoise | 0.007665 | 0.970355 | MRPL40 | turquoise | 0.173678118 | 0.386289303 | FAM50A | yellow | -0.263076 | 0.194125 |
| DESI1 | turquoise | 0.007649 | 0.970417 | TAZ | turquoise | 0.174928737 | 0.382825106 | LDHD | yellow | -0.262938 | 0.194369 |
| ARL1 | turquoise | 0.007533 | 0.970865 | NUDT21 | turquoise | 0.175327389 | 0.381724613 | KLHL36 | yellow | -0.262464 | 0.195204 |
| GPR180 | turquoise | 0.007501 | 0.97099 | CWC22 | turquoise | 0.175774146 | 0.380493494 | LRPAP1 | yellow | -0.262002 | 0.196019 |
| CLCN3 | turquoise | 0.007413 | 0.971331 | HIBCH | turquoise | 0.176463158 | 0.378599289 | SRD5A1 | yellow | -0.261763 | 0.196442 |
| HIBCH | turquoise | 0.006952 | 0.973114 | MID2 | turquoise | 0.177509167 | 0.37573408 | RPUSD3 | yellow | -0.261518 | 0.196878 |
| BCL2L1 | turquoise | -0.006851 | 0.973503 | FMNL3 | turquoise | 0.178058481 | 0.374234457 | ZFP90 | yellow | -0.261078 | 0.197659 |
| PILRA | turquoise | 0.006789 | 0.973744 | SLC33A1 | turquoise | 0.181468819 | 0.365002251 | UBXN11 | yellow | -0.260765 | 0.198217 |
| SPIRE1 | turquoise | 0.006715 | 0.974027 | PJA2 | turquoise | 0.182430603 | 0.362422937 | IRAK1 | yellow | -0.260604 | 0.198504 |
| E2F3 | turquoise | 0.006574 | 0.974573 | C4orf52 | turquoise | 0.18360476 | 0.359288647 | DHRS7B | yellow | -0.260554 | 0.198592 |
| USP37 | turquoise | 0.00633 | 0.975519 | PPP1R16A | turquoise | 0.184733446 | 0.35629086 | TFPT | yellow | -0.260439 | 0.198799 |
| MGAT4A | turquoise | 0.006242 | 0.975858 | ACOT2 | turquoise | 0.185103799 | 0.355310439 | PUF60 | yellow | -0.259786 | 0.199967 |
| EDC4 | turquoise | 0.006126 | 0.976305 | USP48 | turquoise | 0.185396531 | 0.35453663 | NRD1 | yellow | -0.259585 | 0.200327 |
| TRIM44 | turquoise | -0.006092 | 0.976439 | FCHSD1 | turquoise | 0.186978315 | 0.350372648 | ZBTB49 | yellow | -0.259535 | 0.200417 |

**Table 6. Gene module membership and correlation with growth rate (48/72h), with adjusted p values for each module**

| | | | | | | | | | | | |
|---|---|---|---|---|---|---|---|---|---|---|---|
| SLC9B2 | turquoise | 0.006019 | 0.976719 | TBCK | turquoise | 0.187351563 | 0.349394352 | SNORA24 | yellow | -0.2579 | 0.203369 |
| ARFGEF1 | turquoise | 0.005925 | 0.977082 | MTMR10 | turquoise | 0.187440074 | 0.349162598 | AP5Z1 | yellow | -0.256256 | 0.206368 |
| CCDC41 | turquoise | 0.005901 | 0.977176 | ALYREF | turquoise | 0.188050869 | 0.347565828 | UST | yellow | -0.256178 | 0.206509 |
| SMEK1 | turquoise | -0.005805 | 0.977547 | DAB2 | turquoise | 0.18887712 | 0.345412754 | RIPK3 | yellow | -0.256136 | 0.206586 |
| SLC35A4 | turquoise | 0.005801 | 0.977562 | ATPAF2 | turquoise | 0.190334278 | 0.341635136 | ZNF689 | yellow | -0.256073 | 0.206703 |
| NME7 | turquoise | -0.00575 | 0.977759 | DCP1B | turquoise | 0.191222513 | 0.339344648 | CS | yellow | -0.255902 | 0.207017 |
| GPANK1 | turquoise | 0.005655 | 0.978126 | DBNL | turquoise | 0.191796792 | 0.337868693 | SIVA1 | yellow | -0.255854 | 0.207105 |
| U2SURP | turquoise | 0.00558 | 0.978419 | SSSCA1 | turquoise | 0.192354108 | 0.336440038 | SNX12 | yellow | -0.255156 | 0.20839 |
| DMC1 | turquoise | 0.005409 | 0.979077 | SDHAF2 | turquoise | 0.192657219 | 0.335664561 | NAGPA | yellow | -0.253447 | 0.211558 |
| PHACTR1 | turquoise | -0.005301 | 0.979497 | INO80B | turquoise | 0.19457923 | 0.33077247 | AKAP1 | yellow | -0.253254 | 0.211918 |
| TAF1A | turquoise | 0.005151 | 0.980075 | STARD3NL | turquoise | 0.195479865 | 0.328495063 | TRMT10C | yellow | -0.25245 | 0.213423 |
| DYM | turquoise | 0.004619 | 0.982132 | CCDC71 | turquoise | 0.196606671 | 0.325659218 | PRKCD | yellow | -0.252277 | 0.213746 |
| NUP153 | turquoise | 0.004223 | 0.983666 | FKBP3 | turquoise | 0.197095779 | 0.32443294 | ANKRD32 | yellow | -0.251005 | 0.216142 |
| AVPI1 | turquoise | 0.004045 | 0.984354 | RPIA | turquoise | 0.198388358 | 0.321205823 | CXCL2 | yellow | -0.249702 | 0.218615 |
| FAM208B | turquoise | -0.003847 | 0.98512 | CNTRL | turquoise | 0.198478504 | 0.320981498 | STARD3 | yellow | -0.248106 | 0.221671 |
| RPS23 | turquoise | 0.003574 | 0.986176 | SPR | turquoise | 0.19867031 | 0.320504514 | KAT8 | yellow | -0.247972 | 0.221928 |
| DCTD | turquoise | 0.003524 | 0.98637 | LIN7C | turquoise | 0.199166303 | 0.319273091 | SOX4 | yellow | -0.247309 | 0.223206 |
| NEDD4 | turquoise | 0.003445 | 0.986674 | PPAP2A | turquoise | 0.199209187 | 0.319166756 | NARS2 | yellow | -0.246286 | 0.225188 |
| TIMP2 | turquoise | 0.003419 | 0.986774 | FAM73B | turquoise | 0.20089468 | 0.315004718 | CDK11B | yellow | -0.24468 | 0.228325 |
| CLK4 | turquoise | 0.002872 | 0.988889 | CHFR | turquoise | 0.2009255 | 0.314928924 | ZNF470 | yellow | -0.243067 | 0.231503 |
| ZFAND1 | turquoise | -0.002776 | 0.989261 | PFKL | turquoise | 0.20127134 | 0.314079217 | USB1 | yellow | -0.242117 | 0.233389 |
| PABPC3 | turquoise | -0.002333 | 0.990975 | ZNF674 | turquoise | 0.204616653 | 0.305933142 | STAC3 | yellow | -0.241637 | 0.234345 |
| UBQLN1 | turquoise | 0.001875 | 0.992746 | THOP1 | turquoise | 0.204904746 | 0.305237822 | ASAH1 | yellow | -0.24086 | 0.2359 |
| SLC30A7 | turquoise | 0.001695 | 0.993445 | CAP1 | turquoise | 0.204914084 | 0.3052153 | ARHGEF40 | yellow | -0.240729 | 0.236163 |
| JAK2 | turquoise | 0.00161 | 0.993773 | C2orf29 | turquoise | 0.204921274 | 0.305197959 | ERAP1 | yellow | -0.240598 | 0.236426 |
| ATAD1 | turquoise | 0.001569 | 0.993932 | CCDC94 | turquoise | 0.206203518 | 0.302115425 | TMEM173 | yellow | -0.240027 | 0.237574 |
| LINC00597 | turquoise | 0.001421 | 0.994503 | SAT2 | turquoise | 0.206892957 | 0.300466071 | PMPCB | yellow | -0.239893 | 0.237843 |
| GNAI3 | turquoise | 0.001346 | 0.994795 | KLF3 | turquoise | 0.207314726 | 0.299459848 | PRDM10 | yellow | -0.238836 | 0.23998 |
| RAB5A | turquoise | -0.000929 | 0.996405 | PKD1L1 | turquoise | 0.207764236 | 0.298389769 | PARL | yellow | -0.237185 | 0.243343 |
| UBD | turquoise | 0.000518 | 0.997995 | TSPAN33 | turquoise | 0.209212367 | 0.294958746 | TRADD | yellow | -0.236636 | 0.244468 |
| RNF144B | turquoise | -0.000481 | 0.998138 | HHEX | turquoise | 0.210274123 | 0.292458981 | CSRP2BP | yellow | -0.236626 | 0.244488 |
| ENGASE | turquoise | -0.000424 | 0.99836 | MYL12B | turquoise | 0.211875541 | 0.288713988 | PTGR1 | yellow | -0.236292 | 0.245174 |
| NCKIPSD | turquoise | 0.000276 | 0.998931 | ASB1 | turquoise | 0.212002971 | 0.288417295 | RNH1 | yellow | -0.236158 | 0.245449 |
| TM9SF3 | turquoise | -5.66E-06 | 0.999978 | GABPA | turquoise | 0.213279338 | 0.285456205 | MLXIP | yellow | -0.23539 | 0.247034 |
| SNORA24 | white | -0.332973 | 0.096482 | CD83 | turquoise | 0.213338531 | 0.285319351 | GGT7 | yellow | -0.235373 | 0.24707 |
| SNORA5C | white | -0.27475 | 0.174348 | NBEAL1 | turquoise | 0.213388312 | 0.285204291 | BCL11A | yellow | -0.234842 | 0.248167 |
| ACIN1 | white | -0.224869 | 0.269403 | C6orf228 | turquoise | 0.214751586 | 0.282064728 | SPI1 | yellow | -0.234708 | 0.248445 |
| TUBG1 | white | -0.224798 | 0.269557 | PUSL1 | turquoise | 0.21524191 | 0.280940936 | PRR12 | yellow | -0.234067 | 0.249777 |
| MFF | white | -0.207685 | 0.308649 | GDI2 | turquoise | 0.215413312 | 0.280548764 | FUK | yellow | -0.233534 | 0.250888 |
| SNORA23 | white | -0.206328 | 0.311891 | PTPN23 | turquoise | 0.215612502 | 0.280093455 | SLC39A3 | yellow | -0.233105 | 0.251784 |
| SNORA9 | white | -0.196117 | 0.33696 | C9orf91 | turquoise | 0.216079517 | 0.279027795 | TAF4 | yellow | -0.232307 | 0.253458 |
| TMEM99 | white | -0.191875 | 0.347718 | SF3A1 | turquoise | 0.217032372 | 0.276861559 | BEST1 | yellow | -0.231309 | 0.25556 |
| SNORA57 | white | -0.180892 | 0.376515 | TMCC1 | turquoise | 0.218702975 | 0.273089611 | IP6K1 | yellow | -0.230525 | 0.257221 |
| SNORD17 | white | -0.175024 | 0.392446 | LPCAT2 | turquoise | 0.218956209 | 0.272520745 | ENTPD6 | yellow | -0.230338 | 0.257617 |
| RBMX | white | 0.174318 | 0.394388 | CYP51A1 | turquoise | 0.220585301 | 0.268879338 | PPM1B | yellow | -0.228717 | 0.261076 |
| SNORA52 | white | -0.173831 | 0.395732 | KIAA1009 | turquoise | 0.222303016 | 0.265073956 | TFEB | yellow | -0.22593 | 0.267089 |
| SNORA2A | white | -0.16715 | 0.414415 | ZBTB9 | turquoise | 0.222313535 | 0.265050759 | PEX26 | yellow | -0.225152 | 0.268783 |
| ELAC1 | white | -0.137482 | 0.503024 | C3orf14 | turquoise | 0.223750242 | 0.261894973 | CHCHD10 | yellow | -0.224918 | 0.269296 |
| SNORA65 | white | -0.135521 | 0.509194 | FRMD8 | turquoise | 0.224165723 | 0.260986913 | FBXL19 | yellow | -0.224455 | 0.270308 |
| HHLA3 | white | -0.129507 | 0.528339 | PARP15 | turquoise | 0.226393969 | 0.256151836 | BID | yellow | -0.223591 | 0.272205 |
| SNORA62 | white | -0.12662 | 0.537649 | GDF15 | turquoise | 0.226503596 | 0.255915473 | CISD3 | yellow | -0.223578 | 0.272234 |
| SNORA5A | white | -0.116265 | 0.571661 | UBTD1 | turquoise | 0.226756202 | 0.25537138 | HENMT1 | yellow | -0.223575 | 0.272242 |
| SNORA74A | white | -0.111806 | 0.586599 | ISG20L2 | turquoise | 0.230114124 | 0.248210388 | QARS | yellow | -0.221984 | 0.275755 |
| SNORA71D | white | -0.107794 | 0.600179 | CALCOCO1 | turquoise | 0.230208069 | 0.248011958 | MARK4 | yellow | -0.221747 | 0.276283 |
| CD2AP | white | -0.10475 | 0.610573 | ADRBK1 | turquoise | 0.230494539 | 0.247407528 | TUFM | yellow | -0.221725 | 0.27633 |
| KIDINS220 | white | 0.090939 | 0.658626 | WWC2 | turquoise | 0.239911493 | 0.228076294 | NIT1 | yellow | -0.221696 | 0.276396 |

**Table 6. Gene module membership and correlation with growth rate (48/72h), with adjusted p values for each module**

| | | | | | | | | | | | |
|---|---|---|---|---|---|---|---|---|---|---|---|
| SNORA6 | white | -0.086456 | 0.674525 | SEC24B | turquoise | 0.240436359 | 0.227029456 | DOPEY2 | yellow | -0.22143 | 0.276988 |
| SNORA71C | white | 0.085939 | 0.676364 | RUVBL2 | turquoise | 0.241159098 | 0.22559323 | SQLE | yellow | -0.22075 | 0.278502 |
| ZBTB11 | white | -0.069918 | 0.734313 | LCORL | turquoise | 0.241710165 | 0.224502252 | SLC2A9 | yellow | -0.21971 | 0.280831 |
| SNORA48 | white | -0.066275 | 0.747702 | CRLS1 | turquoise | 0.241955298 | 0.224018085 | USP22 | yellow | -0.218389 | 0.283806 |
| NOA1 | white | -0.059954 | 0.771102 | RPL26 | turquoise | 0.242438884 | 0.223065005 | SGK196 | yellow | -0.217623 | 0.285541 |
| SNORA46 | white | -0.058778 | 0.775478 | EIF1B | turquoise | 0.246907972 | 0.214385854 | PRRC2A | yellow | -0.217253 | 0.28638 |
| USP47 | white | 0.055894 | 0.786236 | PIK3R5 | turquoise | 0.247851063 | 0.212583955 | PPP1R26 | yellow | -0.216394 | 0.288337 |
| SNORD15B | white | 0.036421 | 0.859795 | MPV17 | turquoise | 0.247905453 | 0.212480349 | NF1 | yellow | -0.216285 | 0.288587 |
| SNORA16A | white | 0.035046 | 0.865039 | FOXO3 | turquoise | 0.248319439 | 0.211692886 | INTS8 | yellow | -0.21591 | 0.289443 |
| SNORA71B | white | -0.034441 | 0.867349 | EEF1E1 | turquoise | 0.251682265 | 0.20536961 | SRR | yellow | -0.215136 | 0.291218 |
| SNORA31 | white | 0.024646 | 0.904873 | CDK4 | turquoise | 0.253109176 | 0.202725869 | MYPOP | yellow | -0.214773 | 0.292053 |
| SLX4 | white | 0.021952 | 0.915234 | NARFL | turquoise | 0.254910577 | 0.199421636 | TAF8 | yellow | -0.213654 | 0.294633 |
| SNORA28 | white | -0.020945 | 0.919106 | CLN3 | turquoise | 0.256484017 | 0.19656591 | RPL28 | yellow | -0.213604 | 0.29475 |
| SNORA74B | white | -0.009266 | 0.964166 | KIAA0226 | turquoise | 0.258810524 | 0.192395111 | LIMS1 | yellow | -0.213315 | 0.295419 |
| SLC6A6 | white | 0.002428 | 0.990606 | ARL6IP6 | turquoise | 0.259554127 | 0.191075006 | FAM213B | yellow | -0.212691 | 0.296867 |
| SNORA34 | white | 0.001105 | 0.995727 | TUBE1 | turquoise | 0.262074419 | 0.186647343 | FBL | yellow | -0.212546 | 0.297204 |
| SNORA21 | white | -0.001083 | 0.995812 | RAB11B | turquoise | 0.262883132 | 0.185241791 | UTP14C | yellow | -0.209625 | 0.304049 |
| ZNF564 | yellow | -0.517814 | 0.006739 | NOL12 | turquoise | 0.26514924 | 0.181342457 | CMAHP | yellow | -0.209267 | 0.304894 |
| ZFP90 | yellow | -0.512869 | 0.00738 | EDEM3 | turquoise | 0.267594007 | 0.177200207 | WDR83OS | yellow | -0.2073 | 0.309565 |
| HNRNPA3 | yellow | -0.449191 | 0.021334 | DCAF10 | turquoise | 0.268885962 | 0.175038124 | CAMTA2 | yellow | -0.207092 | 0.310062 |
| TSNARE1 | yellow | 0.437112 | 0.025555 | UQCRFS1 | turquoise | 0.270884597 | 0.17172992 | GON4L | yellow | -0.206459 | 0.311577 |
| TSFM | yellow | -0.435092 | 0.026322 | DUSP3 | turquoise | 0.271176198 | 0.171250949 | MRAS | yellow | -0.206274 | 0.312019 |
| CRELD2 | yellow | 0.427053 | 0.029564 | USP32 | turquoise | 0.27163236 | 0.170503561 | LAMP1 | yellow | -0.205837 | 0.31307 |
| LRRCC1 | yellow | -0.413487 | 0.035755 | PLEKHB2 | turquoise | 0.273898195 | 0.166825138 | HERPUD1 | yellow | -0.205816 | 0.313119 |
| JAGN1 | yellow | -0.400059 | 0.042863 | CCM2 | turquoise | 0.277414537 | 0.161228019 | PHRF1 | yellow | -0.205002 | 0.315079 |
| KDELR1 | yellow | 0.39956 | 0.043146 | SPINT2 | turquoise | 0.27851725 | 0.159500526 | TBP | yellow | -0.204628 | 0.315981 |
| SEC61A1 | yellow | 0.39119 | 0.048139 | CCNDBP1 | turquoise | 0.280679505 | 0.156151388 | PPP1R11 | yellow | -0.204488 | 0.31632 |
| SLC39A6 | yellow | -0.389828 | 0.048992 | KRT79 | turquoise | 0.282069772 | 0.154024612 | CEBPG | yellow | -0.203248 | 0.319328 |
| CSNK2A2 | yellow | 0.386725 | 0.050981 | LHFPL2 | turquoise | 0.284944189 | 0.149693111 | PFAS | yellow | -0.203007 | 0.319913 |
| C14orf169 | yellow | 0.386101 | 0.051389 | CEACAM21 | turquoise | 0.285146058 | 0.149392223 | MED15 | yellow | -0.202943 | 0.32007 |
| BLOC1S3 | yellow | 0.385783 | 0.051598 | CSNK2B | turquoise | 0.286667289 | 0.147138736 | FHOD1 | yellow | -0.202786 | 0.320452 |
| CCDC75 | yellow | -0.38383 | 0.052894 | SUV39H2 | turquoise | 0.287483058 | 0.145940385 | PLCG1 | yellow | -0.201603 | 0.323343 |
| GPD2 | yellow | 0.380189 | 0.055378 | BNIP3L | turquoise | 0.287595235 | 0.14577615 | MED1 | yellow | -0.200751 | 0.325437 |
| LENG9 | yellow | 0.379791 | 0.055655 | FXC1 | turquoise | 0.288125131 | 0.145002133 | GTF2H4 | yellow | -0.199968 | 0.327366 |
| KLHL12 | yellow | -0.376225 | 0.058185 | NDUFA4 | turquoise | 0.292904088 | 0.138154595 | NENF | yellow | -0.199563 | 0.328366 |
| POLK | yellow | -0.372276 | 0.06109 | BCKDK | turquoise | 0.298786937 | 0.130050294 | SGSH | yellow | -0.199489 | 0.32855 |
| FARS2 | yellow | -0.368264 | 0.064156 | MRFAP1L1 | turquoise | 0.298909261 | 0.129885538 | C16orf72 | yellow | -0.19939 | 0.328796 |
| PHIP | yellow | -0.366959 | 0.065178 | GPR108 | turquoise | 0.300541964 | 0.127701018 | SLC16A5 | yellow | -0.199272 | 0.329087 |
| CCDC9 | yellow | 0.363894 | 0.067628 | PPAT | turquoise | 0.301957442 | 0.125828946 | C4orf48 | yellow | -0.197583 | 0.333286 |
| ITGB7 | yellow | -0.359655 | 0.071133 | ATP6V1D | turquoise | 0.305927762 | 0.120685044 | TMC8 | yellow | -0.196786 | 0.335281 |
| PPP2R3A | yellow | -0.359224 | 0.071497 | ENY2 | turquoise | 0.308162902 | 0.117858059 | ITPA | yellow | -0.196779 | 0.335297 |
| CPPED1 | yellow | -0.356138 | 0.074145 | RIT1 | turquoise | 0.311471861 | 0.113762852 | ZDHHC13 | yellow | -0.196468 | 0.336079 |
| LOC285074 | yellow | -0.352595 | 0.077277 | CCT5 | turquoise | 0.318058424 | 0.10592565 | KCNC4 | yellow | -0.196157 | 0.336859 |
| PCED1B | yellow | -0.351568 | 0.078203 | C19orf6 | turquoise | 0.319121029 | 0.1046999 | LRSAM1 | yellow | -0.195718 | 0.337964 |
| TSSC4 | yellow | 0.351481 | 0.078282 | ATAD1 | turquoise | 0.319829328 | 0.103888742 | ASF1A | yellow | -0.194685 | 0.34057 |
| TCEAL4 | yellow | -0.349049 | 0.080512 | PRICKLE3 | turquoise | 0.320014795 | 0.103677119 | FGD3 | yellow | -0.193523 | 0.343515 |
| PLCD1 | yellow | 0.34686 | 0.082559 | CHMP6 | turquoise | 0.323197376 | 0.100095576 | CDV3 | yellow | -0.193133 | 0.344507 |
| TBC1D20 | yellow | 0.345694 | 0.083667 | SNF8 | turquoise | 0.325542011 | 0.09751683 | KIAA0513 | yellow | -0.192737 | 0.345516 |
| TMEM102 | yellow | -0.342687 | 0.086572 | KIAA0913 | turquoise | 0.339736283 | 0.082956331 | SIRT7 | yellow | -0.192063 | 0.347237 |
| CTBS | yellow | -0.340392 | 0.088841 | C1GALT1 | turquoise | 0.341184729 | 0.081569055 | CMBL | yellow | -0.191827 | 0.347842 |
| TMEM242 | yellow | -0.339022 | 0.090216 | BANF1 | turquoise | 0.350435153 | 0.073121849 | BABAM1 | yellow | -0.191595 | 0.348438 |
| C16orf80 | yellow | -0.338185 | 0.091064 | ASTE1 | turquoise | 0.351401532 | 0.072279691 | HP1BP3 | yellow | -0.189483 | 0.353877 |
| FAM111A | yellow | -0.335719 | 0.093598 | CDKAL1 | turquoise | 0.354040259 | 0.070018086 | IL4R | yellow | -0.189281 | 0.354399 |
| CASP3 | yellow | 0.335707 | 0.093611 | SUPT4H1 | turquoise | 0.374790613 | 0.054080194 | LOC1002941 | yellow | -0.188083 | 0.357509 |
| CNTROB | yellow | -0.334544 | 0.094824 | ZDHHC13 | turquoise | 0.378049193 | 0.051859412 | SLC39A13 | yellow | -0.187796 | 0.358256 |
| GPR65 | yellow | -0.334095 | 0.095296 | UBE2L3 | turquoise | 0.385419634 | 0.047100299 | TCTEX1D2 | yellow | -0.186925 | 0.36053 |

**Table 6. Gene module membership and correlation with growth rate (48/72h), with adjusted p values for each module**

| | | | | | | | | | | | |
|---|---|---|---|---|---|---|---|---|---|---|---|
| WDFY2 | yellow | 0.333005 | 0.096448 | RPL27A | turquoise | 0.386734257 | 0.046288896 | C7orf26 | yellow | -0.186818 | 0.360811 |
| GLIPR1 | yellow | -0.332813 | 0.096652 | ATG4B | turquoise | 0.39555454 | 0.041126486 | CDC37 | yellow | -0.186619 | 0.361332 |
| GAPDH | yellow | 0.332424 | 0.097067 | COPE | turquoise | 0.396949483 | 0.040353713 | GSKIP | yellow | -0.186599 | 0.361385 |
| MAP3K12 | yellow | -0.328964 | 0.100811 | NDUFAF2 | turquoise | 0.397078886 | 0.040282616 | ZNF254 | yellow | -0.183252 | 0.370215 |
| NUBPL | yellow | -0.328813 | 0.100976 | COPS8 | turquoise | 0.41489258 | 0.031405667 | EPB41 | yellow | -0.18097 | 0.376307 |
| PKD1 | yellow | 0.326195 | 0.103883 | PSMC3 | turquoise | 0.42680947 | 0.026400938 | TRPC4AP | yellow | -0.180181 | 0.378426 |
| SYMPK | yellow | 0.324706 | 0.105565 | CENPV | turquoise | 0.460492255 | 0.015643318 | ATP2C1 | yellow | -0.17962 | 0.379937 |
| MRPS35 | yellow | -0.323038 | 0.107472 | EFNB1 | turquoise | 0.493222119 | 0.008944208 | VPS37C | yellow | -0.179495 | 0.380272 |
| ZNF493 | yellow | -0.32101 | 0.109824 | ARHGEF12 | turquoise | 0.504924172 | 0.007227684 | PARP2 | yellow | -0.179345 | 0.380679 |
| MMP15 | yellow | 0.320814 | 0.110054 | VAT1 | turquoise | 0.548596359 | 0.003046781 | GNB4 | yellow | -0.179102 | 0.381336 |
| MIB2 | yellow | 0.32071 | 0.110175 | CWC25 | white | -0.368645018 | 0.058470552 | PBXIP1 | yellow | -0.178612 | 0.382661 |
| SNRPD3 | yellow | -0.319224 | 0.111928 | GLRX | white | -0.317487667 | 0.106588429 | CCNG2 | yellow | -0.176823 | 0.387523 |
| ASF1A | yellow | -0.318333 | 0.112988 | CUTA | white | -0.276190825 | 0.163160543 | ZNF93 | yellow | -0.17512 | 0.392184 |
| MESDC1 | yellow | 0.318172 | 0.11318 | TBCB | white | -0.26710121 | 0.178029799 | HNMT | yellow | -0.174498 | 0.393894 |
| STK35 | yellow | -0.316637 | 0.115027 | RPL13A | white | -0.266617277 | 0.178847103 | GNPDA1 | yellow | -0.173417 | 0.396875 |
| FAM120B | yellow | 0.314977 | 0.117049 | C12orf44 | white | -0.2501615 | 0.208213022 | ALPK1 | yellow | -0.173329 | 0.39712 |
| HAUS4 | yellow | -0.313414 | 0.118975 | UBXN1 | white | -0.243462866 | 0.221055874 | ZNF740 | yellow | -0.172992 | 0.398052 |
| DARS2 | yellow | -0.312131 | 0.120575 | POLR2C | white | -0.238270549 | 0.231369929 | ST3GAL5 | yellow | -0.172938 | 0.3982 |
| ZCCHC6 | yellow | -0.311055 | 0.121929 | MRPL54 | white | -0.206499089 | 0.301407635 | MAN2B1 | yellow | -0.171745 | 0.401513 |
| ZNF335 | yellow | 0.310324 | 0.122854 | NDUFS8 | white | -0.201122908 | 0.314443732 | CYGB | yellow | -0.171261 | 0.402861 |
| POLR2A | yellow | 0.310264 | 0.12293 | RPLP0 | white | -0.172337117 | 0.390023728 | AAAS | yellow | -0.170921 | 0.40381 |
| C19orf38 | yellow | 0.306322 | 0.128014 | UBB | white | -0.145929856 | 0.46765777 | BLOC1S5 | yellow | -0.170919 | 0.403815 |
| ADRM1 | yellow | 0.306164 | 0.128221 | RPS18 | white | -0.121921831 | 0.544638329 | TMEM163 | yellow | -0.170306 | 0.405528 |
| APPL1 | yellow | -0.306156 | 0.128231 | C11orf31 | white | -0.110315784 | 0.583849677 | LCMT1 | yellow | -0.169718 | 0.407177 |
| GOT1 | yellow | -0.304644 | 0.130224 | EIF3B | white | -0.099513866 | 0.621412041 | DBF4B | yellow | -0.169656 | 0.407351 |
| C17orf61 | yellow | -0.304635 | 0.130235 | OSTF1 | white | -0.070097029 | 0.728270155 | ZBTB6 | yellow | -0.166895 | 0.415137 |
| SCAMP1 | yellow | -0.303427 | 0.131843 | RPS19BP1 | white | -0.05598647 | 0.781499082 | MAPKBP1 | yellow | -0.166816 | 0.415362 |
| CCNG2 | yellow | -0.301998 | 0.133764 | RPL36 | white | -0.040415457 | 0.841363924 | UPF2 | yellow | -0.166335 | 0.416726 |
| OXSR1 | yellow | -0.30169 | 0.13418 | NIPAL2 | white | -0.036713127 | 0.855738818 | PLA2G7 | yellow | -0.166271 | 0.416909 |
| GLYCTK | yellow | 0.301348 | 0.134645 | KLHL6 | white | -0.033987818 | 0.866349318 | STAG3 | yellow | -0.165733 | 0.41844 |
| GMFB | yellow | -0.301096 | 0.134986 | DIDO1 | white | -0.028147594 | 0.889161192 | CDKL1 | yellow | -0.16542 | 0.419331 |
| KDELR2 | yellow | 0.300429 | 0.135896 | MRPS18C | white | -0.026301642 | 0.896390292 | AUP1 | yellow | -0.164544 | 0.421833 |
| AMDHD2 | yellow | 0.298953 | 0.137924 | ARPC5L | white | -0.025683719 | 0.898812029 | MIPEP | yellow | -0.164505 | 0.421942 |
| EIF2B1 | yellow | -0.298892 | 0.138008 | RPA2 | white | -0.020885185 | 0.917646735 | TRERF1 | yellow | -0.164345 | 0.422403 |
| ZNHIT1 | yellow | -0.298528 | 0.138512 | RNF181 | white | 0.023673725 | 0.906695521 | MTERFD2 | yellow | -0.16431 | 0.4225 |
| EHD4 | yellow | -0.298441 | 0.138632 | BPGM | white | 0.036537352 | 0.856422448 | GSE1 | yellow | -0.163905 | 0.423662 |
| PARP8 | yellow | -0.295631 | 0.142567 | CLEC16A | white | 0.08535645 | 0.672067549 | SNORA57 | yellow | -0.163359 | 0.42523 |
| LY6G5B | yellow | 0.295405 | 0.142887 | AKT3 | white | 0.091044541 | 0.651531419 | TTL | yellow | -0.163282 | 0.425451 |
| CHRNA5 | yellow | -0.293777 | 0.145207 | TBC1D13 | white | 0.105003963 | 0.602197161 | PSMA2 | yellow | -0.161703 | 0.430001 |
| PLAGL2 | yellow | 0.293695 | 0.145324 | GSTO1 | white | 0.106560293 | 0.596796239 | MED12 | yellow | -0.16126 | 0.431283 |
| CNST | yellow | -0.293284 | 0.145915 | UBL5 | white | 0.126036654 | 0.531037966 | CEP350 | yellow | -0.158178 | 0.440256 |
| COX7C | yellow | -0.291666 | 0.148256 | FAM178A | white | 0.147303699 | 0.46343135 | NFATC1 | yellow | -0.157723 | 0.441588 |
| FLOT1 | yellow | -0.291458 | 0.148559 | RPS26 | white | 0.204720036 | 0.30568351 | IL6R | yellow | -0.157038 | 0.443599 |
| SCRN3 | yellow | -0.291002 | 0.149224 | RAD51C | white | 0.282760884 | 0.152975097 | LHPP | yellow | -0.155156 | 0.44915 |
| DECR2 | yellow | -0.290397 | 0.150109 | TCF3 | yellow | -0.680964937 | 9.24536E-05 | WHSC2 | yellow | -0.154373 | 0.451468 |
| KIFAP3 | yellow | -0.290345 | 0.150186 | PDK1 | yellow | -0.676074565 | 0.000108492 | SUGP2 | yellow | -0.153813 | 0.453133 |
| CASD1 | yellow | -0.288279 | 0.153241 | BID | yellow | -0.638182273 | 0.000341436 | SSR4 | yellow | -0.152518 | 0.45699 |
| TPI1 | yellow | -0.286121 | 0.15648 | LDB1 | yellow | -0.637528141 | 0.000347797 | EIF4G2 | yellow | -0.152403 | 0.457332 |
| SLC16A3 | yellow | 0.285682 | 0.157144 | MBD6 | yellow | -0.627192547 | 0.000463038 | MTX2 | yellow | -0.151762 | 0.459251 |
| ADAMTS1 | yellow | 0.285449 | 0.157498 | CMIP | yellow | -0.616320686 | 0.000619015 | SDHD | yellow | -0.15173 | 0.459345 |
| TAPBPL | yellow | -0.285159 | 0.157939 | STK25 | yellow | -0.611386801 | 0.000703743 | SLFN11 | yellow | -0.151475 | 0.460111 |
| WIZ | yellow | 0.28479 | 0.158501 | ARSA | yellow | -0.606785699 | 0.000791683 | DHX37 | yellow | -0.150826 | 0.462057 |
| SNORA84 | yellow | 0.284197 | 0.159408 | FASTKD5 | yellow | -0.601944381 | 0.000894384 | SNORA62 | yellow | -0.150586 | 0.462776 |
| EPHB6 | yellow | 0.283999 | 0.159711 | NSRP1 | yellow | -0.588545218 | 0.001241193 | PEX16 | yellow | -0.150215 | 0.463893 |
| C12orf23 | yellow | 0.281641 | 0.163357 | FYTTD1 | yellow | -0.585824954 | 0.001324299 | TBC1D4 | yellow | -0.148798 | 0.468169 |
| PSIP1 | yellow | -0.281239 | 0.163984 | RAE1 | yellow | -0.580829073 | 0.001489511 | IKZF5 | yellow | -0.14833 | 0.469583 |

**Table 6. Gene module membership and correlation with growth rate (48/72h), with adjusted p values for each module**

| | | | | | | | | | | | |
|---|---|---|---|---|---|---|---|---|---|---|---|
| IFT46 | yellow | -0.281205 | 0.164037 | DAXX | yellow | -0.579954746 | 0.001520184 | TNRC18 | yellow | -0.147509 | 0.472074 |
| TIA1 | yellow | 0.280992 | 0.164371 | KLF10 | yellow | -0.578422699 | 0.001575246 | SLC35E2B | yellow | -0.146041 | 0.476543 |
| MRTO4 | yellow | -0.280878 | 0.164549 | TAF11 | yellow | -0.576809926 | 0.00163506 | ZNF140 | yellow | -0.145643 | 0.47776 |
| NUPR1 | yellow | 0.279836 | 0.166188 | UBN1 | yellow | -0.5761644 | 0.001659544 | NADK | yellow | -0.145128 | 0.479332 |
| STAG2 | yellow | -0.279826 | 0.166204 | PHIP | yellow | -0.575979692 | 0.001666608 | C17orf51 | yellow | -0.144341 | 0.481745 |
| PPP2CB | yellow | 0.278873 | 0.167712 | CDYL2 | yellow | -0.573465141 | 0.001765383 | CBLB | yellow | -0.142841 | 0.48636 |
| NVL | yellow | -0.278708 | 0.167973 | SIRT2 | yellow | -0.571313703 | 0.001853858 | NFIC | yellow | -0.139868 | 0.49557 |
| DIAPH2 | yellow | 0.275224 | 0.173575 | COA4 | yellow | -0.570380421 | 0.00189341 | AZI2 | yellow | -0.139541 | 0.49659 |
| TRAPPC2L | yellow | -0.274953 | 0.174017 | CCR1 | yellow | -0.568681226 | 0.001967288 | CCDC86 | yellow | -0.139226 | 0.49757 |
| LTV1 | yellow | -0.274693 | 0.17444 | ATXN1 | yellow | -0.565933954 | 0.002091982 | DPYD | yellow | -0.137128 | 0.504136 |
| SMARCC1 | yellow | -0.273833 | 0.175847 | CEBPB | yellow | -0.565902101 | 0.002093467 | NLRX1 | yellow | -0.135694 | 0.508647 |
| PITRM1 | yellow | -0.272952 | 0.177298 | ZNF431 | yellow | -0.563594865 | 0.002203447 | CBX7 | yellow | -0.134828 | 0.511381 |
| ZNF383 | yellow | -0.272731 | 0.177663 | AFG3L2 | yellow | -0.562278774 | 0.002268377 | C16orf54 | yellow | -0.134624 | 0.512027 |
| PTP4A1 | yellow | -0.27263 | 0.17783 | DNAJB2 | yellow | -0.560494743 | 0.002359013 | TSC22D4 | yellow | -0.13443 | 0.512642 |
| ASH2L | yellow | -0.272324 | 0.178336 | PDIA4 | yellow | -0.55987316 | 0.002391313 | SNORA14A | yellow | -0.134421 | 0.51267 |
| CTPS1 | yellow | -0.271002 | 0.180538 | CXCL2 | yellow | -0.558698126 | 0.002453411 | CUEDC1 | yellow | -0.13428 | 0.513115 |
| SELPLG | yellow | -0.270296 | 0.181719 | CHMP1A | yellow | -0.557523963 | 0.00251684 | MPND | yellow | -0.133907 | 0.514298 |
| RHBDD3 | yellow | 0.26822 | 0.18523 | HIST1H2BC | yellow | -0.556631877 | 0.002565966 | TBL2 | yellow | -0.133677 | 0.515026 |
| SHQ1 | yellow | -0.26793 | 0.185722 | DGAT1 | yellow | -0.556139047 | 0.002593457 | ELF2 | yellow | -0.133466 | 0.515695 |
| MZT1 | yellow | -0.267728 | 0.186067 | SUPV3L1 | yellow | -0.555979612 | 0.002602404 | TNPO1 | yellow | -0.133032 | 0.517076 |
| LRBA | yellow | -0.267162 | 0.187035 | IDH3B | yellow | -0.555526478 | 0.002627978 | BCL7B | yellow | -0.132894 | 0.517515 |
| ITCH | yellow | -0.265928 | 0.189158 | RPUSD1 | yellow | -0.555165187 | 0.002648522 | ZBTB44 | yellow | -0.13265 | 0.518289 |
| TTC38 | yellow | -0.265422 | 0.190032 | RBM42 | yellow | -0.555101232 | 0.002652173 | TP53I11 | yellow | -0.132342 | 0.519271 |
| CDYL | yellow | 0.264113 | 0.192309 | GLTPD1 | yellow | -0.554182425 | 0.002705103 | DNAJC17 | yellow | -0.131988 | 0.520398 |
| BAG1 | yellow | -0.263837 | 0.192791 | TMF1 | yellow | -0.551585053 | 0.002859653 | GPR85 | yellow | -0.131039 | 0.523428 |
| ALDH5A1 | yellow | -0.263297 | 0.193737 | GPN2 | yellow | -0.55052374 | 0.002924945 | TMOD3 | yellow | -0.130322 | 0.525723 |
| C9orf114 | yellow | -0.262509 | 0.195123 | UBE4A | yellow | -0.54974947 | 0.002973378 | ZFP3 | yellow | -0.126931 | 0.536642 |
| FBXL6 | yellow | -0.262199 | 0.195671 | MTO1 | yellow | -0.549540676 | 0.002986556 | LSM3 | yellow | -0.126097 | 0.539344 |
| GCNT1 | yellow | -0.262007 | 0.19601 | ZNF564 | yellow | -0.547821565 | 0.003096966 | GCNT2 | yellow | -0.124424 | 0.544782 |
| CKAP2 | yellow | -0.261989 | 0.196043 | DPP9 | yellow | -0.547717264 | 0.003103776 | CDK10 | yellow | -0.122215 | 0.552001 |
| TOR2A | yellow | 0.261335 | 0.197202 | TNFAIP3 | yellow | -0.544901984 | 0.003292476 | WDR26 | yellow | -0.121612 | 0.553979 |
| TPGS2 | yellow | -0.260763 | 0.198219 | CDAN1 | yellow | -0.543794302 | 0.00336936 | ZFYVE9 | yellow | -0.120918 | 0.55626 |
| NEDD1 | yellow | -0.260384 | 0.198896 | ST3GAL1 | yellow | -0.543631922 | 0.003380759 | LILRB3 | yellow | -0.1197 | 0.560274 |
| N4BP1 | yellow | -0.25916 | 0.201091 | LOC440434 | yellow | -0.542845559 | 0.003436427 | ABI2 | yellow | -0.118928 | 0.562825 |
| ZBTB5 | yellow | 0.258958 | 0.201456 | STUB1 | yellow | -0.541901758 | 0.00350427 | TTC7A | yellow | -0.118806 | 0.563229 |
| PSMA1 | yellow | -0.258855 | 0.201641 | CXCL1 | yellow | -0.540823615 | 0.003583166 | MIA3 | yellow | -0.118477 | 0.564318 |
| UBA7 | yellow | -0.2586 | 0.202102 | TSG101 | yellow | -0.540435811 | 0.003611912 | CIAO1 | yellow | -0.117439 | 0.567758 |
| LMF1 | yellow | 0.258214 | 0.202801 | SCARNA5 | yellow | -0.540230245 | 0.00362723 | SON | yellow | -0.116915 | 0.569499 |
| MPP7 | yellow | -0.256953 | 0.205092 | PAPSS2 | yellow | -0.539973103 | 0.003646468 | MECR | yellow | -0.114314 | 0.578177 |
| SRRM1 | yellow | -0.256952 | 0.205094 | MGRN1 | yellow | -0.539619869 | 0.003673038 | TUBGCP6 | yellow | -0.113091 | 0.582275 |
| APOL4 | yellow | -0.25645 | 0.206012 | SCRIB | yellow | -0.538756071 | 0.003738707 | MCAT | yellow | -0.112488 | 0.584301 |
| ZNF644 | yellow | 0.256115 | 0.206626 | FUS | yellow | -0.53867382 | 0.003745012 | MED25 | yellow | -0.111974 | 0.586032 |
| PLD3 | yellow | 0.255367 | 0.208001 | TBC1D2B | yellow | -0.537277266 | 0.003853455 | USF2 | yellow | -0.111309 | 0.588272 |
| GALK2 | yellow | -0.254636 | 0.20935 | COBRA1 | yellow | -0.53283176 | 0.004216635 | NDFIP2 | yellow | -0.11041 | 0.591309 |
| TMEM159 | yellow | -0.253724 | 0.211043 | LOC541471 | yellow | -0.532272528 | 0.004264318 | TDRKH | yellow | -0.110189 | 0.592054 |
| PRR14L | yellow | -0.253456 | 0.211541 | TRMT61A | yellow | -0.531349039 | 0.004344064 | SCARNA17 | yellow | -0.109785 | 0.593424 |
| TXNDC12 | yellow | -0.253242 | 0.211941 | MLL | yellow | -0.531321977 | 0.00434642 | MAP3K11 | yellow | -0.109271 | 0.595163 |
| VPS29 | yellow | -0.253145 | 0.212122 | TAF1B | yellow | -0.529930974 | 0.004468985 | ZNF319 | yellow | -0.108782 | 0.596821 |
| UIMC1 | yellow | -0.251715 | 0.214802 | MED28 | yellow | -0.528911471 | 0.004560669 | USP35 | yellow | -0.108664 | 0.597223 |
| RBM26 | yellow | -0.251491 | 0.215224 | ICAM1 | yellow | -0.528311418 | 0.004615375 | NHP2L1 | yellow | -0.104765 | 0.61052 |
| DDO | yellow | 0.251044 | 0.216068 | N4BP2 | yellow | -0.524605383 | 0.004965747 | RPS4X | yellow | -0.104657 | 0.610889 |
| RBL2 | yellow | -0.250843 | 0.216448 | ELF2 | yellow | -0.52379112 | 0.005045677 | NAA40 | yellow | -0.104412 | 0.611728 |
| UBA6 | yellow | -0.249268 | 0.219444 | ETFB | yellow | -0.521375272 | 0.005289271 | C22orf29 | yellow | -0.104087 | 0.612845 |
| STX11 | yellow | -0.24882 | 0.220301 | TBC1D10B | yellow | -0.520996933 | 0.005328307 | TMEM41A | yellow | -0.103357 | 0.615351 |
| SH3BP2 | yellow | 0.248282 | 0.221331 | MAP3K4 | yellow | -0.520721202 | 0.00535691 | YME1L1 | yellow | -0.102314 | 0.618941 |
| DENND3 | yellow | 0.24822 | 0.221451 | TRIP10 | yellow | -0.520585981 | 0.005370984 | TMED10 | yellow | -0.102246 | 0.619174 |

**Table 6. Gene module membership and correlation with growth rate (48/72h), with adjusted p values for each module**

| | | | |
|---|---|---|---|
| CSF3R | yellow | 0.248043 | 0.221793 |
| EPS8L1 | yellow | 0.247376 | 0.223077 |
| MAPK13 | yellow | 0.246986 | 0.223832 |
| MCM3 | yellow | -0.246039 | 0.225669 |
| KIAA1715 | yellow | -0.245788 | 0.226158 |
| TXNRD2 | yellow | 0.245727 | 0.226278 |
| DBF4 | yellow | -0.244949 | 0.227797 |
| TXNDC17 | yellow | -0.244672 | 0.228341 |
| IKBKAP | yellow | -0.244393 | 0.228888 |
| MUT | yellow | -0.244258 | 0.229154 |
| DTNBP1 | yellow | -0.243868 | 0.229921 |
| RAB11FIP2 | yellow | -0.243825 | 0.230006 |
| ABTB1 | yellow | 0.243758 | 0.230138 |
| COA3 | yellow | -0.243451 | 0.230744 |
| PHF1 | yellow | 0.242939 | 0.231757 |
| DUSP22 | yellow | 0.242732 | 0.232167 |
| LILRA2 | yellow | 0.242636 | 0.232358 |
| CTDSP2 | yellow | 0.242215 | 0.233194 |
| NOP9 | yellow | -0.241683 | 0.234254 |
| FAR1 | yellow | -0.241675 | 0.234271 |
| FAM105B | yellow | -0.241029 | 0.235562 |
| MTERFD2 | yellow | -0.240389 | 0.236845 |
| CLPX | yellow | -0.240103 | 0.23742 |
| DOCK11 | yellow | -0.239813 | 0.238005 |
| ZMYND8 | yellow | -0.239546 | 0.238545 |
| MAP3K8 | yellow | 0.239415 | 0.238808 |
| BCL2L12 | yellow | 0.239167 | 0.239311 |
| PRKCH | yellow | -0.238982 | 0.239684 |
| ELL | yellow | 0.238306 | 0.241056 |
| AKAP5 | yellow | -0.237859 | 0.241967 |
| ARHGEF17 | yellow | 0.237446 | 0.242809 |
| ASL | yellow | 0.237396 | 0.242912 |
| MAP2K7 | yellow | 0.237381 | 0.242942 |
| FAM175B | yellow | -0.236326 | 0.245104 |
| NLK | yellow | -0.236088 | 0.245595 |
| SLC30A5 | yellow | -0.235885 | 0.246012 |
| TLE1 | yellow | -0.235692 | 0.24641 |
| IARS2 | yellow | -0.235263 | 0.247296 |
| RASSF5 | yellow | -0.234826 | 0.2482 |
| ICK | yellow | -0.234558 | 0.248756 |
| ERCC3 | yellow | -0.233584 | 0.250785 |
| PCBD1 | yellow | 0.233423 | 0.25112 |
| PMS2P1 | yellow | 0.233019 | 0.251965 |
| B2M | yellow | -0.232287 | 0.253499 |
| FAM114A2 | yellow | -0.23216 | 0.253766 |
| HEXA | yellow | 0.231561 | 0.25503 |
| JMJD6 | yellow | 0.231482 | 0.255196 |
| B4GALT3 | yellow | 0.231081 | 0.256042 |
| SEC24A | yellow | -0.230247 | 0.25781 |
| RNASET2 | yellow | -0.22921 | 0.26002 |
| COPS7B | yellow | -0.228826 | 0.260841 |
| CST3 | yellow | -0.228747 | 0.261011 |
| AOC3 | yellow | -0.227782 | 0.263083 |
| GOLGA8B | yellow | 0.22731 | 0.2641 |
| CAPN3 | yellow | -0.2272 | 0.264338 |
| CNPY2 | yellow | -0.226689 | 0.265443 |
| HIST1H1C | yellow | -0.518816761 | 0.005558036 |
| CCDC137 | yellow | -0.517142968 | 0.005740044 |
| STAG2 | yellow | -0.516887437 | 0.005768268 |
| MPHOSPH8 | yellow | -0.5159705 | 0.005870517 |
| TMEM106A | yellow | -0.512401694 | 0.006283188 |
| LMAN1 | yellow | -0.512377094 | 0.006286116 |
| RPS6KC1 | yellow | -0.512320439 | 0.006292862 |
| WFS1 | yellow | -0.510734304 | 0.006484228 |
| IRF8 | yellow | -0.510431049 | 0.006521368 |
| SART1 | yellow | -0.510218416 | 0.006547517 |
| PHYH | yellow | -0.509275402 | 0.006664554 |
| ZCCHC2 | yellow | -0.509143719 | 0.006681036 |
| SIRT7 | yellow | -0.508795873 | 0.006724741 |
| TTC31 | yellow | -0.507993567 | 0.006826465 |
| RALBP1 | yellow | -0.507063737 | 0.006945976 |
| SF1 | yellow | -0.50681542 | 0.006978189 |
| SETD8 | yellow | -0.506761145 | 0.006985247 |
| TRAPPC11 | yellow | -0.505668264 | 0.007128641 |
| HTT | yellow | -0.505410921 | 0.007162765 |
| NUP205 | yellow | -0.505276571 | 0.007180634 |
| CORO2A | yellow | -0.505228041 | 0.007187098 |
| SMS | yellow | -0.504593647 | 0.007272048 |
| RARA | yellow | -0.50305397 | 0.007481741 |
| YBX1 | yellow | -0.502325367 | 0.00758273 |
| C15orf39 | yellow | -0.501670331 | 0.007674498 |
| C21orf59 | yellow | -0.500079595 | 0.007901246 |
| C3 | yellow | -0.499903852 | 0.007926639 |
| NLRC4 | yellow | -0.499370547 | 0.008004114 |
| STRA13 | yellow | -0.499093559 | 0.008044604 |
| UBAC1 | yellow | -0.496849894 | 0.008378959 |
| ZCCHC9 | yellow | -0.496170222 | 0.008482516 |
| COX14 | yellow | -0.49528711 | 0.008618671 |
| ARPC1A | yellow | -0.493993281 | 0.008821453 |
| SPNS1 | yellow | -0.492832204 | 0.009006818 |
| IL10RA | yellow | -0.49232651 | 0.009088564 |
| TRUB1 | yellow | -0.492120876 | 0.009121983 |
| MAP2K3 | yellow | -0.490999958 | 0.009305962 |
| SZT2 | yellow | -0.490191625 | 0.009440554 |
| FKBP8 | yellow | -0.489102854 | 0.00962441 |
| HGS | yellow | -0.487074986 | 0.009974831 |
| WDTC1 | yellow | -0.486611304 | 0.010056435 |
| MED24 | yellow | -0.486566835 | 0.010064291 |
| NCL | yellow | -0.486470097 | 0.010081397 |
| HPS4 | yellow | -0.485094087 | 0.010327362 |
| TNRC18 | yellow | -0.48374707 | 0.010572974 |
| CRADD | yellow | -0.479624747 | 0.011355097 |
| EIF4B | yellow | -0.478097695 | 0.011656785 |
| MLXIP | yellow | -0.476743509 | 0.011929872 |
| XPA | yellow | -0.476684029 | 0.011941988 |
| TEP1 | yellow | -0.475856679 | 0.012111577 |
| KIAA0100 | yellow | -0.475116259 | 0.012265041 |
| UBE2Q2 | yellow | -0.474286235 | 0.012438993 |
| DHRS9 | yellow | -0.474196014 | 0.012458024 |
| PITPNM1 | yellow | -0.473418727 | 0.012622986 |
| DDX52 | yellow | -0.473156114 | 0.012679129 |
| PCNP | yellow | -0.472501287 | 0.012820022 |
| LATS2 | yellow | -0.101858 | 0.620514 |
| TMC6 | yellow | -0.101607 | 0.62138 |
| HELZ2 | yellow | -0.099809 | 0.627597 |
| UPF3A | yellow | -0.098042 | 0.633731 |
| BTBD6 | yellow | -0.097901 | 0.634222 |
| CCT5 | yellow | -0.097385 | 0.636018 |
| C3orf37 | yellow | -0.096636 | 0.638629 |
| DVL3 | yellow | -0.096361 | 0.63959 |
| FANCG | yellow | -0.095649 | 0.642077 |
| SEC13 | yellow | -0.095337 | 0.643168 |
| PPARD | yellow | -0.09358 | 0.649326 |
| NMT1 | yellow | -0.093293 | 0.650334 |
| MNT | yellow | -0.092429 | 0.653372 |
| AGA | yellow | -0.09172 | 0.655869 |
| FAM86A | yellow | -0.091501 | 0.656643 |
| CKS2 | yellow | -0.09048 | 0.660247 |
| DIS3L2 | yellow | -0.086117 | 0.67573 |
| KIFC2 | yellow | -0.084966 | 0.679837 |
| C12orf29 | yellow | -0.083282 | 0.685863 |
| TRIM56 | yellow | -0.082664 | 0.688078 |
| ERMAP | yellow | -0.081717 | 0.691478 |
| H1FX | yellow | -0.079891 | 0.69805 |
| ICMT | yellow | -0.078883 | 0.701688 |
| PI4K2A | yellow | -0.077779 | 0.705679 |
| TAGLN2 | yellow | -0.07572 | 0.713143 |
| GNAQ | yellow | -0.075477 | 0.714025 |
| SLC35B4 | yellow | -0.072173 | 0.72606 |
| DGCR11 | yellow | -0.070102 | 0.733638 |
| ATXN2L | yellow | -0.067329 | 0.74382 |
| DNAJC15 | yellow | -0.067126 | 0.744566 |
| TREM1 | yellow | -0.066132 | 0.748229 |
| LOC440173 | yellow | -0.065679 | 0.749901 |
| CXCL3 | yellow | -0.065676 | 0.749911 |
| ZNF318 | yellow | -0.065477 | 0.750644 |
| TSTD2 | yellow | -0.064123 | 0.755646 |
| GNB5 | yellow | -0.06376 | 0.756987 |
| UNC93B1 | yellow | -0.063013 | 0.759752 |
| NPRL3 | yellow | -0.062453 | 0.761827 |
| SLC35E3 | yellow | -0.06189 | 0.763914 |
| PVRL1 | yellow | -0.061783 | 0.764311 |
| CAMSAP1 | yellow | -0.061408 | 0.7657 |
| MCM9 | yellow | -0.060969 | 0.76733 |
| LSG1 | yellow | -0.059349 | 0.773351 |
| PLXND1 | yellow | -0.058812 | 0.775352 |
| NAA10 | yellow | -0.058001 | 0.778374 |
| SLC25A40 | yellow | -0.057712 | 0.779452 |
| NMNAT1 | yellow | -0.056535 | 0.783841 |
| CEBPA-AS1 | yellow | -0.056437 | 0.784208 |
| GALNT2 | yellow | -0.054641 | 0.790924 |
| RRP8 | yellow | -0.054638 | 0.790936 |
| FAM53B | yellow | -0.054396 | 0.791841 |
| NOL7 | yellow | -0.053114 | 0.796646 |
| TRIOBP | yellow | -0.049484 | 0.810285 |
| HSBP1L1 | yellow | -0.049093 | 0.811758 |
| PHF13 | yellow | -0.048232 | 0.815002 |
| C11orf54 | yellow | -0.045085 | 0.826888 |

**Table 6. Gene module membership and correlation with growth rate (48/72h), with adjusted p values for each module**

| | | | | | | | | | | | |
|---|---|---|---|---|---|---|---|---|---|---|---|
| RAPGEF3 | yellow | 0.226626 | 0.26558 | RPUSD4 | yellow | -0.472287379 | 0.012866328 | POLR3E | yellow | -0.042863 | 0.835301 |
| ISCA2 | yellow | -0.224042 | 0.271213 | DNAH1 | yellow | -0.472087483 | 0.012909725 | TLE3 | yellow | -0.041822 | 0.839251 |
| BOP1 | yellow | 0.22367 | 0.272031 | QDPR | yellow | -0.470071239 | 0.013354283 | FNDC3B | yellow | -0.041633 | 0.839967 |
| DDX1 | yellow | -0.2221 | 0.275499 | HNRNPA3 | yellow | -0.468924938 | 0.013612636 | CDKN2D | yellow | -0.041194 | 0.841636 |
| TIGD5 | yellow | 0.221379 | 0.277101 | CCDC59 | yellow | -0.46855667 | 0.01369651 | SYTL3 | yellow | -0.040887 | 0.842799 |
| STK16 | yellow | 0.220313 | 0.279479 | PLA2G4C | yellow | -0.465417907 | 0.014428892 | SBDSP1 | yellow | -0.040028 | 0.846064 |
| KIAA0232 | yellow | -0.218735 | 0.283025 | ZNF417 | yellow | -0.464837782 | 0.014567738 | ANKLE2 | yellow | -0.038493 | 0.851901 |
| TUBA1C | yellow | 0.218482 | 0.283597 | ZNF45 | yellow | -0.462363499 | 0.015172393 | NME7 | yellow | -0.037288 | 0.85649 |
| DHX16 | yellow | 0.21811 | 0.284437 | TMEM175 | yellow | -0.461801913 | 0.015312478 | ITPKB | yellow | -0.036158 | 0.860796 |
| GPX1 | yellow | -0.217979 | 0.284734 | RNF2 | yellow | -0.461558262 | 0.015373587 | VMA21 | yellow | -0.035435 | 0.863554 |
| TET2 | yellow | -0.217788 | 0.285167 | UTP11L | yellow | -0.460832795 | 0.01555673 | TMEM201 | yellow | -0.03343 | 0.871211 |
| LRP1 | yellow | -0.217673 | 0.285427 | CLIP1 | yellow | -0.460000435 | 0.015769073 | PKN1 | yellow | -0.033139 | 0.872322 |
| PRMT2 | yellow | -0.217433 | 0.285971 | TRIM26 | yellow | -0.459945848 | 0.015783082 | TGFB1 | yellow | -0.032896 | 0.87325 |
| RNF31 | yellow | 0.217077 | 0.286782 | WTAP | yellow | -0.459881978 | 0.015799486 | PDIA6 | yellow | -0.031918 | 0.876992 |
| IST1 | yellow | -0.216796 | 0.28742 | HIST1H2AC | yellow | -0.458298989 | 0.01621057 | SLC26A11 | yellow | -0.029821 | 0.885016 |
| ZCCHC24 | yellow | 0.21643 | 0.288256 | AK2 | yellow | -0.458197272 | 0.016237283 | PIK3C3 | yellow | -0.028502 | 0.890074 |
| DBNDD2 | yellow | -0.216424 | 0.288268 | CS | yellow | -0.458042783 | 0.016277926 | DCTN1 | yellow | -0.028263 | 0.890991 |
| NET1 | yellow | -0.216196 | 0.288789 | ZC3H7A | yellow | -0.45787648 | 0.016321769 | ZNF671 | yellow | -0.027362 | 0.894445 |
| ATP6V0E1 | yellow | 0.216072 | 0.289074 | ACTR5 | yellow | -0.457091429 | 0.016530053 | ATG4A | yellow | -0.027345 | 0.894512 |
| NCF2 | yellow | -0.214858 | 0.291858 | BLVRA | yellow | -0.456997426 | 0.016555139 | NCSTN | yellow | -0.02567 | 0.900941 |
| RRN3 | yellow | -0.214615 | 0.292415 | CD2BP2 | yellow | -0.456473437 | 0.01669555 | HINT2 | yellow | -0.024683 | 0.904729 |
| LOC253039 | yellow | 0.214088 | 0.293632 | TBC1D10A | yellow | -0.456410726 | 0.01671242 | PTGES2 | yellow | -0.024672 | 0.904774 |
| PPP5C | yellow | -0.2138 | 0.294296 | CDK11A | yellow | -0.455788239 | 0.016880637 | PGBD2 | yellow | -0.023472 | 0.909385 |
| ABCA11P | yellow | 0.213397 | 0.295229 | YOD1 | yellow | -0.455783171 | 0.016882012 | SLC9A3R2 | yellow | -0.022898 | 0.911593 |
| ENTHD2 | yellow | -0.213326 | 0.295394 | MRPL52 | yellow | -0.455682292 | 0.016909406 | CUL2 | yellow | -0.022442 | 0.913346 |
| ADD3 | yellow | -0.213182 | 0.295729 | KANSL3 | yellow | -0.455520413 | 0.01695344 | PTPN1 | yellow | -0.022112 | 0.914617 |
| STX1B | yellow | 0.213063 | 0.296003 | CUTC | yellow | -0.455137028 | 0.017058106 | NSMCE2 | yellow | -0.022054 | 0.914841 |
| SYNGAP1 | yellow | -0.212898 | 0.296386 | PQBP1 | yellow | -0.454122112 | 0.017337751 | FAM135A | yellow | -0.021772 | 0.915924 |
| TMEM185B | yellow | -0.212134 | 0.298164 | ARRDC3 | yellow | -0.453874627 | 0.01740651 | NOP14 | yellow | -0.021689 | 0.916243 |
| PHF5A | yellow | -0.212127 | 0.298182 | MTG1 | yellow | -0.452784888 | 0.017711949 | NOC4L | yellow | -0.02166 | 0.916357 |
| HDAC8 | yellow | -0.210902 | 0.301045 | PSIP1 | yellow | -0.452005694 | 0.017933035 | HARBI1 | yellow | -0.02134 | 0.917589 |
| LRRC25 | yellow | 0.210403 | 0.302217 | HNRNPA2B1 | yellow | -0.451821267 | 0.017985695 | NIF3L1 | yellow | -0.02106 | 0.918664 |
| TXN2 | yellow | 0.209707 | 0.303856 | CANX | yellow | -0.451745024 | 0.018007502 | DUSP16 | yellow | -0.017885 | 0.930897 |
| C1orf159 | yellow | -0.209644 | 0.304005 | WHAMM | yellow | -0.451041418 | 0.018209772 | DENND4A | yellow | -0.017422 | 0.932681 |
| MARK3 | yellow | -0.209365 | 0.304663 | DDX26B | yellow | -0.450553029 | 0.018351266 | ZNF132 | yellow | -0.016943 | 0.934529 |
| TOB2 | yellow | 0.208745 | 0.306131 | MT1X | yellow | -0.449849939 | 0.018556546 | OPHN1 | yellow | -0.016733 | 0.935339 |
| TRIM11 | yellow | 0.207693 | 0.30863 | MT1E | yellow | -0.445456881 | 0.019882276 | ATL2 | yellow | -0.016707 | 0.935438 |
| MPP5 | yellow | -0.207528 | 0.309023 | CCDC174 | yellow | -0.444872203 | 0.020064424 | TES | yellow | -0.01531 | 0.940826 |
| ANPEP | yellow | -0.206945 | 0.310414 | EIF2B4 | yellow | -0.444657783 | 0.020131564 | TRIM65 | yellow | -0.013386 | 0.948251 |
| PSAT1 | yellow | -0.206534 | 0.311398 | PLOD3 | yellow | -0.442711093 | 0.020749578 | LRRC45 | yellow | -0.013182 | 0.94904 |
| C21orf7 | yellow | -0.205655 | 0.313506 | TET3 | yellow | -0.441884904 | 0.021016514 | PLEC | yellow | -0.012249 | 0.952645 |
| SCARNA12 | yellow | -0.205256 | 0.314466 | PPP2R5B | yellow | -0.440976244 | 0.02131333 | FARP1 | yellow | -0.012021 | 0.953524 |
| SLC5A6 | yellow | 0.204654 | 0.31592 | SH3GL1 | yellow | -0.440521882 | 0.021463027 | SBDS | yellow | -0.011508 | 0.955503 |
| RRP12 | yellow | 0.204346 | 0.316663 | GLTSCR2 | yellow | -0.438553247 | 0.02212158 | TNPO2 | yellow | -0.009619 | 0.962801 |
| LRRC14 | yellow | 0.203865 | 0.317828 | PARG | yellow | -0.437313045 | 0.022544853 | OPLAH | yellow | -0.009408 | 0.96362 |
| GTF3C3 | yellow | -0.203802 | 0.317981 | MRPS25 | yellow | -0.437078056 | 0.022625792 | SLIRP | yellow | -0.00934 | 0.963881 |
| DDHD2 | yellow | -0.203511 | 0.318689 | SERF1A | yellow | -0.436474834 | 0.022834651 | ATP6V0C | yellow | -0.007742 | 0.970058 |
| ING4 | yellow | 0.202772 | 0.320486 | CNDP2 | yellow | -0.435862999 | 0.023048093 | ORC5 | yellow | -0.007302 | 0.971757 |
| CERK | yellow | -0.201799 | 0.322864 | CBFB | yellow | -0.434762741 | 0.023436013 | LRRK2 | yellow | -0.004433 | 0.982852 |
| TRIP11 | yellow | -0.201348 | 0.323968 | VIMP | yellow | -0.432598599 | 0.024214537 | RAP1B | yellow | -0.004215 | 0.983698 |
| TJAP1 | yellow | -0.201187 | 0.324365 | TUSC2 | yellow | -0.431057975 | 0.024781469 | ZDHHC9 | yellow | -0.001717 | 0.993357 |
| PLEKHO1 | yellow | -0.200908 | 0.325049 | POLK | yellow | -0.430751817 | 0.024895405 | ZNF804A | yellow | 9.8E-05 | 0.999621 |
| ZNF673 | yellow | 0.200553 | 0.325924 | AP5Z1 | yellow | -0.429386824 | 0.025408577 | KAT6A | yellow | 0.003238 | 0.987474 |
| PRPF38A | yellow | -0.20008 | 0.32709 | OPTN | yellow | -0.429013658 | 0.025550353 | DPAGT1 | yellow | 0.004215 | 0.983694 |
| MDN1 | yellow | -0.200001 | 0.327285 | SLK | yellow | -0.427854499 | 0.02599485 | BTBD10 | yellow | 0.004368 | 0.983104 |
| DNMT3A | yellow | -0.19963 | 0.3282 | CD68 | yellow | -0.427706624 | 0.026052003 | C16orf87 | yellow | 0.005563 | 0.978481 |

**Table 6. Gene module membership and correlation with growth rate (48/72h), with adjusted p values for each module**

| | | | | | | | | | | | |
|---|---|---|---|---|---|---|---|---|---|---|---|
| DGCR6L | yellow | -0.198657 | 0.330613 | TMEM158 | yellow | -0.427053073 | 0.02630582 | EPB41L1 | yellow | 0.006702 | 0.974079 |
| TPD52 | yellow | -0.198598 | 0.330759 | SNX12 | yellow | -0.425051326 | 0.027095725 | DNAJC11 | yellow | 0.007275 | 0.971862 |
| MTRR | yellow | -0.19724 | 0.334143 | SERTAD3 | yellow | -0.423920718 | 0.027550288 | ELL2 | yellow | 0.008041 | 0.968901 |
| ZC3HAV1 | yellow | 0.197115 | 0.334456 | MPRIP | yellow | -0.421459386 | 0.02856122 | CCDC106 | yellow | 0.009869 | 0.961836 |
| TMOD3 | yellow | -0.196695 | 0.335509 | UNC13D | yellow | -0.421375278 | 0.028596287 | NT5C3 | yellow | 0.011324 | 0.956216 |
| MED23 | yellow | -0.196577 | 0.335805 | VTI1A | yellow | -0.420374519 | 0.029016215 | PPM1N | yellow | 0.011397 | 0.955935 |
| WDR43 | yellow | -0.196425 | 0.336184 | PTPRA | yellow | -0.419925275 | 0.029206334 | PTPRE | yellow | 0.011916 | 0.953928 |
| BMP2K | yellow | -0.196056 | 0.337111 | PHF11 | yellow | -0.419696991 | 0.029303327 | PHF3 | yellow | 0.013078 | 0.94944 |
| MACC1 | yellow | -0.1952 | 0.339269 | ERP29 | yellow | -0.418463678 | 0.029831845 | C9orf64 | yellow | 0.015902 | 0.938543 |
| GCFC2 | yellow | -0.195073 | 0.339589 | RNF166 | yellow | -0.418283662 | 0.029909627 | LPCAT3 | yellow | 0.017945 | 0.930666 |
| MEN1 | yellow | -0.194563 | 0.340879 | AKAP1 | yellow | -0.417096344 | 0.030426753 | LINC00094 | yellow | 0.019431 | 0.924939 |
| ZNF529 | yellow | -0.194311 | 0.341514 | RIOK1 | yellow | -0.415868082 | 0.030969268 | LY6E | yellow | 0.019563 | 0.924428 |
| PPP3CC | yellow | -0.193659 | 0.343169 | DDX39B | yellow | -0.414679596 | 0.031501602 | ACP1 | yellow | 0.019777 | 0.923607 |
| ADAM15 | yellow | 0.193611 | 0.343291 | NR1D2 | yellow | -0.413987954 | 0.031814768 | TOMM22 | yellow | 0.020341 | 0.921435 |
| C7orf50 | yellow | 0.192422 | 0.346321 | TPM3 | yellow | -0.412370674 | 0.032556827 | SLAIN2 | yellow | 0.021119 | 0.918438 |
| TIPRL | yellow | -0.191889 | 0.347684 | PSMD6 | yellow | -0.410436312 | 0.033462569 | TTC4 | yellow | 0.022886 | 0.911638 |
| RBP4 | yellow | -0.191104 | 0.349698 | IDI1 | yellow | -0.406227435 | 0.035503223 | GPR34 | yellow | 0.024826 | 0.90418 |
| MFSD1 | yellow | 0.190633 | 0.350908 | SNORA27 | yellow | -0.406107018 | 0.035563039 | UTP15 | yellow | 0.025534 | 0.901461 |
| ARHGEF10L | yellow | -0.190556 | 0.351106 | NKTR | yellow | -0.406049451 | 0.035591664 | NOD1 | yellow | 0.028637 | 0.889556 |
| DDRGK1 | yellow | -0.190364 | 0.351602 | LOC285074 | yellow | -0.404839104 | 0.036197774 | RRAGB | yellow | 0.029447 | 0.886451 |
| IFT20 | yellow | -0.188894 | 0.355401 | GNB4 | yellow | -0.404795334 | 0.036219847 | CLEC2B | yellow | 0.032263 | 0.87567 |
| CYB5R1 | yellow | -0.188852 | 0.355512 | SLC39A11 | yellow | -0.404292684 | 0.036474097 | TRAPPC5 | yellow | 0.033372 | 0.871432 |
| CMC2 | yellow | 0.188703 | 0.355898 | PSMA1 | yellow | -0.402445669 | 0.037420632 | COPG1 | yellow | 0.035521 | 0.863228 |
| DCTN4 | yellow | -0.187964 | 0.35782 | HDHD3 | yellow | -0.402149416 | 0.037574261 | UBE2K | yellow | 0.03557 | 0.863041 |
| NDFIP1 | yellow | -0.187645 | 0.358651 | ERC1 | yellow | -0.402002534 | 0.037650616 | CNTROB | yellow | 0.036662 | 0.858875 |
| CSTA | yellow | -0.187373 | 0.359361 | TMEM51 | yellow | -0.400292807 | 0.038548543 | TRA2B | yellow | 0.037047 | 0.857407 |
| YPEL5 | yellow | 0.187128 | 0.360001 | ZFYVE20 | yellow | -0.39959656 | 0.03891906 | POLR3A | yellow | 0.03778 | 0.854617 |
| GCC1 | yellow | 0.186073 | 0.362765 | GPRC5A | yellow | -0.397263719 | 0.040181236 | CRLF3 | yellow | 0.038848 | 0.85055 |
| WDR45L | yellow | 0.185553 | 0.36413 | PLRG1 | yellow | -0.396934121 | 0.04036216 | GOPC | yellow | 0.039088 | 0.849637 |
| SEL1L | yellow | 0.185218 | 0.365012 | ERLIN2 | yellow | -0.396538138 | 0.04058038 | NEK11 | yellow | 0.03926 | 0.848983 |
| RBBP5 | yellow | 0.185038 | 0.365486 | CARS | yellow | -0.396235504 | 0.040747788 | CBL | yellow | 0.039482 | 0.848141 |
| CDC42SE1 | yellow | 0.184529 | 0.366829 | ACP2 | yellow | -0.393617458 | 0.042219047 | IFT172 | yellow | 0.039868 | 0.846673 |
| RPS19BP1 | yellow | -0.184146 | 0.367843 | AAED1 | yellow | -0.392640607 | 0.042778684 | ST5 | yellow | 0.040652 | 0.843694 |
| ATN1 | yellow | 0.183918 | 0.368449 | LMNB2 | yellow | -0.391910779 | 0.043200629 | SLC35C1 | yellow | 0.041779 | 0.839412 |
| ST3GAL1 | yellow | -0.183664 | 0.36912 | KIAA1609 | yellow | -0.39147654 | 0.043453242 | C8orf76 | yellow | 0.042113 | 0.838146 |
| CYBA | yellow | -0.183643 | 0.369176 | USP13 | yellow | -0.390013351 | 0.044313063 | HES2 | yellow | 0.042623 | 0.836213 |
| TNKS | yellow | -0.183109 | 0.370595 | NDUFS4 | yellow | -0.389835722 | 0.044418355 | SMYD5 | yellow | 0.043213 | 0.833978 |
| GK5 | yellow | -0.18288 | 0.371204 | ACVRL1 | yellow | -0.38481985 | 0.047474202 | MRPS22 | yellow | 0.043884 | 0.831435 |
| TMEM56 | yellow | -0.182775 | 0.371483 | OSBP | yellow | -0.383469896 | 0.048324323 | AP1S2 | yellow | 0.044713 | 0.828298 |
| KANSL1 | yellow | -0.181425 | 0.375086 | PMP22 | yellow | -0.382792256 | 0.048755559 | C15orf40 | yellow | 0.045709 | 0.824531 |
| LYZ | yellow | 0.181163 | 0.375788 | ZNF276 | yellow | -0.380943243 | 0.04994766 | MAP7D1 | yellow | 0.046142 | 0.822891 |
| PTAR1 | yellow | -0.181134 | 0.375865 | VCAN | yellow | -0.379975241 | 0.05058083 | LOC10050564 | yellow | 0.04678 | 0.820483 |
| COMT | yellow | 0.180097 | 0.378651 | CDC5L | yellow | -0.379604935 | 0.050824709 | METTL5 | yellow | 0.047299 | 0.818523 |
| MED10 | yellow | 0.179837 | 0.379352 | RBMX2 | yellow | -0.378740161 | 0.051397836 | SAMD1 | yellow | 0.049205 | 0.811336 |
| TMEM199 | yellow | -0.179762 | 0.379553 | TRPM4 | yellow | -0.375579981 | 0.053535514 | ACAD10 | yellow | 0.049243 | 0.811194 |
| CEBPA | yellow | 0.179617 | 0.379943 | DVL3 | yellow | -0.373745702 | 0.054807867 | NBPF11 | yellow | 0.053519 | 0.795126 |
| SYNJ1 | yellow | -0.179479 | 0.380318 | MNT | yellow | -0.372363786 | 0.055781966 | UBE2L3 | yellow | 0.053953 | 0.793499 |
| TGS1 | yellow | -0.177961 | 0.384426 | CHPF | yellow | -0.372187881 | 0.055906924 | KIAA0146 | yellow | 0.054243 | 0.792416 |
| NCKAP1L | yellow | -0.177878 | 0.384651 | PRKAR1A | yellow | -0.370461162 | 0.057145182 | LRRK1 | yellow | 0.055216 | 0.788774 |
| APRT | yellow | 0.177559 | 0.385518 | MAP2K5 | yellow | -0.370448338 | 0.057154457 | SMARCA4 | yellow | 0.05574 | 0.786814 |
| SGK196 | yellow | -0.177174 | 0.386566 | ABL2 | yellow | -0.370432177 | 0.057166148 | FAM91A1 | yellow | 0.056587 | 0.783648 |
| GATM | yellow | -0.177152 | 0.386625 | BCOR | yellow | -0.37023336 | 0.057310128 | DDB1 | yellow | 0.056905 | 0.782459 |
| EML3 | yellow | -0.17664 | 0.388021 | FAM109A | yellow | -0.368343397 | 0.058692967 | MAST3 | yellow | 0.057261 | 0.781133 |
| OSCAR | yellow | -0.176468 | 0.38849 | SRI | yellow | -0.368059041 | 0.058903253 | AFG3L1P | yellow | 0.058956 | 0.774816 |
| RABL2B | yellow | 0.175612 | 0.390833 | M6PR | yellow | -0.364931602 | 0.061254967 | CUL1 | yellow | 0.058963 | 0.774788 |
| MRPL9 | yellow | -0.175495 | 0.391154 | SEPW1 | yellow | -0.364554137 | 0.061543669 | KIAA1109 | yellow | 0.059643 | 0.772258 |

**Table 6. Gene module membership and correlation with growth rate (48/72h), with adjusted p values for each module**

| | | | |
|---|---|---|---|
| IVD | yellow | 0.175099 | 0.392241 |
| NEO1 | yellow | 0.174765 | 0.39316 |
| MITF | yellow | -0.174704 | 0.393326 |
| MED1 | yellow | -0.174446 | 0.394035 |
| ZFP36L2 | yellow | 0.173564 | 0.39647 |
| SNRPD2 | yellow | -0.173519 | 0.396594 |
| HERC2P2 | yellow | -0.173402 | 0.396917 |
| CCNT1 | yellow | -0.173273 | 0.397273 |
| TEN1 | yellow | 0.172706 | 0.398843 |
| NMT1 | yellow | -0.172644 | 0.399015 |
| MRPL23 | yellow | -0.171456 | 0.402318 |
| ZDHHC21 | yellow | -0.171409 | 0.402449 |
| ZNF675 | yellow | 0.170444 | 0.405144 |
| TNNI2 | yellow | -0.170277 | 0.40561 |
| SMC6 | yellow | -0.170144 | 0.405984 |
| SPHK1 | yellow | 0.170127 | 0.40603 |
| FAM195B | yellow | -0.168666 | 0.410133 |
| ITGA5 | yellow | 0.168078 | 0.411791 |
| WDR46 | yellow | 0.167959 | 0.412128 |
| FOXN2 | yellow | -0.167829 | 0.412494 |
| DESI2 | yellow | -0.167825 | 0.412506 |
| TMEM18 | yellow | 0.167575 | 0.413211 |
| AIFM3 | yellow | 0.167452 | 0.413561 |
| SRSF5 | yellow | -0.166938 | 0.415015 |
| LRRC42 | yellow | -0.166919 | 0.415069 |
| RAB11A | yellow | -0.166765 | 0.415506 |
| MGC72080 | yellow | -0.16576 | 0.418361 |
| PTGFRN | yellow | 0.16565 | 0.418674 |
| ATOX1 | yellow | -0.1649 | 0.420814 |
| PPP2R5C | yellow | -0.164884 | 0.420861 |
| KPNA5 | yellow | -0.163948 | 0.42354 |
| ATP5B | yellow | -0.163165 | 0.425786 |
| TMEM234 | yellow | 0.162945 | 0.426419 |
| ASF1B | yellow | -0.162582 | 0.427463 |
| ZNF740 | yellow | -0.161997 | 0.429152 |
| TTC17 | yellow | -0.161768 | 0.429814 |
| HSBP1 | yellow | -0.159891 | 0.435256 |
| WDR37 | yellow | 0.159811 | 0.43549 |
| AKAP11 | yellow | -0.159709 | 0.435786 |
| ANAPC5 | yellow | -0.159161 | 0.437382 |
| TRHDE | yellow | -0.158984 | 0.437899 |
| FAM126B | yellow | -0.15888 | 0.438201 |
| TOR1B | yellow | -0.158447 | 0.439468 |
| RPL27 | yellow | 0.158318 | 0.439845 |
| SETD7 | yellow | -0.15825 | 0.440045 |
| DOCK10 | yellow | -0.157482 | 0.442293 |
| MKL1 | yellow | 0.157479 | 0.442304 |
| PDCL3 | yellow | 0.156803 | 0.44429 |
| FAM122B | yellow | 0.156178 | 0.44613 |
| HIPK1 | yellow | 0.156027 | 0.446575 |
| RBM5 | yellow | -0.155779 | 0.447307 |
| CAMLG | yellow | 0.155126 | 0.449237 |
| KPTN | yellow | 0.154723 | 0.45043 |
| KIAA1967 | yellow | 0.154594 | 0.450814 |
| KLF11 | yellow | -0.154529 | 0.451005 |
| AAGAB | yellow | -0.154354 | 0.451524 |
| HEATR5B | yellow | -0.364153036 | 0.061851606 |
| NUSAP1 | yellow | -0.364115701 | 0.061880329 |
| C16orf72 | yellow | -0.363738958 | 0.062170759 |
| TROVE2 | yellow | -0.362360154 | 0.063242702 |
| HMGA1 | yellow | -0.361389244 | 0.064006088 |
| DNMT1 | yellow | -0.359576314 | 0.065450607 |
| ZNF428 | yellow | -0.358728674 | 0.066134587 |
| STIM1 | yellow | -0.357983591 | 0.066740363 |
| SEC11C | yellow | -0.357858534 | 0.066842458 |
| ARID4B | yellow | -0.352466013 | 0.071360687 |
| QARS | yellow | -0.350322644 | 0.073220383 |
| ACSL1 | yellow | -0.349535138 | 0.073912929 |
| UBE2H | yellow | -0.348224263 | 0.07507686 |
| TP53I3 | yellow | -0.348170121 | 0.075125233 |
| MITF | yellow | -0.347585135 | 0.075649411 |
| UBE2Q2P1 | yellow | -0.346241459 | 0.076864016 |
| CEBPG | yellow | -0.345748733 | 0.077313129 |
| ELK1 | yellow | -0.345542932 | 0.077501306 |
| EZH2 | yellow | -0.345039492 | 0.07796311 |
| SUGP2 | yellow | -0.344895432 | 0.078095641 |
| GPHN | yellow | -0.344776835 | 0.078204877 |
| BRPF3 | yellow | -0.342986776 | 0.079867849 |
| ZNF777 | yellow | -0.34208021 | 0.080720269 |
| GNAI1 | yellow | -0.34051769 | 0.082205713 |
| VPS9D1 | yellow | -0.339155143 | 0.083517943 |
| HNRNPH2 | yellow | -0.336667399 | 0.085954795 |
| RNF170 | yellow | -0.335447179 | 0.087169576 |
| SNX29P2 | yellow | -0.334423044 | 0.088199143 |
| ZNF510 | yellow | -0.33319142 | 0.08944945 |
| UFC1 | yellow | -0.330247364 | 0.092492394 |
| AGPAT4 | yellow | -0.328861867 | 0.093951127 |
| ATG16L1 | yellow | -0.326641153 | 0.096325236 |
| AEN | yellow | -0.32567057 | 0.09737689 |
| TIMM8B | yellow | -0.325651439 | 0.097397705 |
| TTC33 | yellow | -0.324169884 | 0.099019843 |
| RSF1 | yellow | -0.32284259 | 0.100490188 |
| FAM126A | yellow | -0.321330197 | 0.102185414 |
| SRSF5 | yellow | -0.318637509 | 0.105256339 |
| MT2A | yellow | -0.318047558 | 0.105938239 |
| TMEM140 | yellow | -0.317830441 | 0.106190021 |
| RRN3P2 | yellow | -0.315160102 | 0.109323225 |
| RREB1 | yellow | -0.312287139 | 0.112770202 |
| GSE1 | yellow | -0.311675172 | 0.113514707 |
| INIP | yellow | -0.308634063 | 0.117268412 |
| MPP5 | yellow | -0.308031379 | 0.118023045 |
| HMGN1 | yellow | -0.307840691 | 0.118262552 |
| ZNF775 | yellow | -0.30767298 | 0.118473495 |
| TIMM17A | yellow | -0.30374579 | 0.123492508 |
| MMP10 | yellow | -0.301130259 | 0.126920502 |
| GCDH | yellow | -0.300997289 | 0.127096615 |
| C17orf75 | yellow | -0.300757294 | 0.127414926 |
| APP | yellow | -0.300355203 | 0.127949533 |
| SFMBT1 | yellow | -0.296701137 | 0.132883084 |
| ZDHHC7 | yellow | -0.29520846 | 0.134937672 |
| TBC1D4 | yellow | -0.29491201 | 0.135348446 |
| SLA | yellow | -0.293894784 | 0.136764848 |
| RNF139 | yellow | 0.059837 | 0.771536 |
| KIF3C | yellow | 0.060271 | 0.769925 |
| UHRF1BP1L | yellow | 0.061185 | 0.76653 |
| PMM1 | yellow | 0.061622 | 0.764909 |
| ACACA | yellow | 0.062078 | 0.763216 |
| SLC16A1 | yellow | 0.062326 | 0.762295 |
| MTR | yellow | 0.063079 | 0.759507 |
| HSPA4 | yellow | 0.063085 | 0.759486 |
| COPZ1 | yellow | 0.063395 | 0.758337 |
| RPL4 | yellow | 0.064018 | 0.756033 |
| ELP3 | yellow | 0.067037 | 0.744897 |
| DCK | yellow | 0.067249 | 0.744115 |
| SLC19A1 | yellow | 0.067324 | 0.743838 |
| SIRPD | yellow | 0.068995 | 0.737698 |
| RNMTL1 | yellow | 0.06912 | 0.73724 |
| TBC1D24 | yellow | 0.069344 | 0.736415 |
| AURKAPS1 | yellow | 0.069566 | 0.735604 |
| ARL14EPL | yellow | 0.070142 | 0.733493 |
| HS6ST1 | yellow | 0.071167 | 0.729738 |
| SLC33A1 | yellow | 0.071627 | 0.728057 |
| STAU1 | yellow | 0.072383 | 0.725296 |
| ERVK13-1 | yellow | 0.07418 | 0.718742 |
| ATXN3 | yellow | 0.074536 | 0.717445 |
| ZBTB4 | yellow | 0.074927 | 0.716026 |
| ALKBH5 | yellow | 0.075495 | 0.71396 |
| SNORA30 | yellow | 0.075773 | 0.712952 |
| NSUN2 | yellow | 0.075915 | 0.712437 |
| ABCF2 | yellow | 0.076852 | 0.709037 |
| SLC31A1 | yellow | 0.077949 | 0.705066 |
| FBRS | yellow | 0.078116 | 0.704461 |
| KNTC1 | yellow | 0.078318 | 0.703729 |
| PITPNB | yellow | 0.078574 | 0.702806 |
| CRKL | yellow | 0.079105 | 0.700887 |
| CC2D1A | yellow | 0.080987 | 0.694105 |
| SCARNA18 | yellow | 0.081881 | 0.690891 |
| AIG1 | yellow | 0.08399 | 0.683327 |
| TMEM19 | yellow | 0.084031 | 0.683183 |
| TACC1 | yellow | 0.084681 | 0.680856 |
| KIAA1033 | yellow | 0.086006 | 0.676126 |
| IER5 | yellow | 0.086352 | 0.674895 |
| NECAP1 | yellow | 0.088943 | 0.665688 |
| TMEM245 | yellow | 0.089864 | 0.662425 |
| GLTSCR1 | yellow | 0.090122 | 0.661513 |
| NFXL1 | yellow | 0.090664 | 0.659597 |
| UBE2V1 | yellow | 0.091198 | 0.657712 |
| HSPA6 | yellow | 0.091241 | 0.657558 |
| FBXO4 | yellow | 0.091663 | 0.65607 |
| HDLBP | yellow | 0.093017 | 0.651304 |
| RPS6KB1 | yellow | 0.093064 | 0.651138 |
| PURB | yellow | 0.093372 | 0.650056 |
| YIPF6 | yellow | 0.09338 | 0.650029 |
| UBE2O | yellow | 0.0937 | 0.648903 |
| MCMBP | yellow | 0.093729 | 0.648803 |
| ASPHD2 | yellow | 0.094769 | 0.645157 |
| TXNRD3 | yellow | 0.095213 | 0.643601 |
| PAN3 | yellow | 0.097901 | 0.634222 |

**Table 6. Gene module membership and correlation with growth rate (48/72h), with adjusted p values for each module**

| | | | | | | | | | | | |
|---|---|---|---|---|---|---|---|---|---|---|---|
| IP6K1 | yellow | -0.153297 | 0.454666 | STX12 | yellow | -0.291800244 | 0.13971506 | FBXO21 | yellow | 0.098248 | 0.633015 |
| BRI3BP | yellow | 0.153145 | 0.45512 | CCDC22 | yellow | -0.28866738 | 0.144213136 | RNF130 | yellow | 0.09827 | 0.632936 |
| AIG1 | yellow | 0.152463 | 0.457153 | METTL23 | yellow | -0.287082062 | 0.146528562 | XPC | yellow | 0.100289 | 0.625936 |
| TYROBP | yellow | 0.151867 | 0.458935 | PMPCA | yellow | -0.285420458 | 0.148983923 | FUT8 | yellow | 0.101279 | 0.62251 |
| DOCK8 | yellow | -0.151616 | 0.459689 | KIF3B | yellow | -0.283752538 | 0.151478117 | TCOF1 | yellow | 0.101322 | 0.622364 |
| SERPINA9 | yellow | -0.151572 | 0.45982 | SLC2A9 | yellow | -0.282912322 | 0.152745809 | DHX15 | yellow | 0.101779 | 0.620786 |
| LRCH3 | yellow | -0.151444 | 0.460201 | KIF13A | yellow | -0.280243956 | 0.156821953 | TMEM187 | yellow | 0.102782 | 0.61733 |
| HARS | yellow | -0.151242 | 0.460808 | VGLL3 | yellow | -0.27820732 | 0.159984725 | PRPSAP2 | yellow | 0.103548 | 0.614697 |
| GTPBP4 | yellow | -0.151019 | 0.461478 | SGSH | yellow | -0.274696174 | 0.165543102 | CLTC | yellow | 0.10454 | 0.611291 |
| ZNF10 | yellow | -0.150913 | 0.461795 | SLC25A26 | yellow | -0.274193965 | 0.166349139 | CETN2 | yellow | 0.105045 | 0.60956 |
| GYS1 | yellow | 0.150754 | 0.462273 | ERCC5 | yellow | -0.268046272 | 0.17644123 | ZGPAT | yellow | 0.105239 | 0.608898 |
| MAPK3 | yellow | -0.150413 | 0.463297 | TUBGCP6 | yellow | -0.267017642 | 0.178170748 | TIFAB | yellow | 0.106302 | 0.605263 |
| TMEM52B | yellow | 0.150384 | 0.463386 | BANP | yellow | -0.266179727 | 0.179588325 | IREB2 | yellow | 0.108325 | 0.598373 |
| CCDC57 | yellow | -0.149999 | 0.464545 | C9orf114 | yellow | -0.261602754 | 0.187470506 | ATP5I | yellow | 0.108987 | 0.596127 |
| NDUFA3 | yellow | 0.149962 | 0.464654 | NUP93 | yellow | -0.261418081 | 0.187793487 | CDC14A | yellow | 0.110578 | 0.590741 |
| ZNF143 | yellow | -0.149262 | 0.466767 | PPP4R1L | yellow | -0.260554686 | 0.189308619 | FTO | yellow | 0.111547 | 0.58747 |
| C9orf78 | yellow | 0.148737 | 0.468353 | AURKAIP1 | yellow | -0.253320361 | 0.202336577 | CD58 | yellow | 0.112342 | 0.584794 |
| NUDC | yellow | 0.147396 | 0.472416 | RPS25 | yellow | -0.242422357 | 0.223097531 | PGPEP1 | yellow | 0.113458 | 0.581046 |
| PLBD1 | yellow | -0.147038 | 0.473506 | USP21 | yellow | -0.238381288 | 0.231146667 | DOPEY1 | yellow | 0.113712 | 0.580194 |
| CPSF1 | yellow | -0.146758 | 0.474358 | TMEM173 | yellow | -0.234626991 | 0.238796013 | EIF4ENIF1 | yellow | 0.114695 | 0.576901 |
| WIPI2 | yellow | 0.146694 | 0.474553 | STK11 | yellow | -0.231837231 | 0.244587451 | L3HYPDH | yellow | 0.117863 | 0.566353 |
| ECSCR | yellow | -0.146511 | 0.47511 | NBPF9 | yellow | -0.231420954 | 0.24545949 | CWC27 | yellow | 0.118299 | 0.564905 |
| ADAMTSL4 | yellow | 0.146468 | 0.475241 | RAI14 | yellow | -0.228179507 | 0.252319811 | ANTXR2 | yellow | 0.118463 | 0.564361 |
| MPZL3 | yellow | 0.1463 | 0.475753 | PRPF3 | yellow | -0.224530622 | 0.26019109 | ARHGAP17 | yellow | 0.118818 | 0.563187 |
| ZNF642 | yellow | 0.146259 | 0.475879 | RBM41 | yellow | -0.223462917 | 0.262524135 | C19orf53 | yellow | 0.120027 | 0.559195 |
| GNPTAB | yellow | -0.146203 | 0.47605 | HECTD4 | yellow | -0.223051375 | 0.263427006 | UBE2T | yellow | 0.121895 | 0.553051 |
| C1orf52 | yellow | -0.146185 | 0.476105 | TTC12 | yellow | -0.220377861 | 0.269341265 | LYRM2 | yellow | 0.122134 | 0.552266 |
| VARS | yellow | -0.146041 | 0.476544 | MAST3 | yellow | -0.213312212 | 0.285380196 | BATF3 | yellow | 0.123744 | 0.546998 |
| ELP5 | yellow | 0.145563 | 0.478002 | SAFB | yellow | -0.212893807 | 0.286348573 | ZNF658 | yellow | 0.125359 | 0.541739 |
| ZDHHC3 | yellow | -0.145439 | 0.478382 | CPEB3 | yellow | -0.209390103 | 0.294539357 | MGST2 | yellow | 0.125725 | 0.540549 |
| GCFC1 | yellow | -0.145182 | 0.479167 | RNASEH2A | yellow | -0.207625048 | 0.298720857 | ROCK2 | yellow | 0.128057 | 0.533004 |
| GIMAP4 | yellow | 0.145137 | 0.479307 | CRTC3 | yellow | -0.20743408 | 0.299175488 | GATAD1 | yellow | 0.128289 | 0.532258 |
| LEPRE1 | yellow | -0.14508 | 0.47948 | SOWAHC | yellow | -0.201657961 | 0.313130988 | C9orf78 | yellow | 0.128914 | 0.530243 |
| ZNF706 | yellow | -0.145018 | 0.479671 | EHF | yellow | -0.198102027 | 0.321918989 | SEC23A | yellow | 0.131404 | 0.522262 |
| EVI5 | yellow | -0.144852 | 0.480179 | TNFRSF12A | yellow | -0.191258529 | 0.339251969 | PPWD1 | yellow | 0.131625 | 0.521556 |
| NUP160 | yellow | -0.144654 | 0.480785 | RPS14 | yellow | -0.187900236 | 0.347959216 | PAM | yellow | 0.131758 | 0.521133 |
| OSBPL11 | yellow | -0.144301 | 0.481869 | SNX18 | yellow | -0.18403391 | 0.358147078 | ZNF684 | yellow | 0.132169 | 0.519821 |
| PTGS1 | yellow | 0.143903 | 0.483089 | PSMG3 | yellow | -0.182037575 | 0.363475664 | HIBADH | yellow | 0.132651 | 0.518288 |
| FBXO31 | yellow | 0.143866 | 0.483205 | SAR1B | yellow | -0.17999753 | 0.368968711 | CCDC47 | yellow | 0.132733 | 0.518027 |
| HLA-DMA | yellow | -0.143334 | 0.484841 | ARG2 | yellow | -0.178871899 | 0.372020224 | ARSD | yellow | 0.133186 | 0.516585 |
| ALG14 | yellow | -0.142734 | 0.486691 | DOT1L | yellow | -0.178394594 | 0.373318586 | TPP2 | yellow | 0.135064 | 0.510635 |
| CHD8 | yellow | 0.142386 | 0.487764 | CLLU1OS | yellow | -0.1666423 | 0.406110509 | MIB1 | yellow | 0.135904 | 0.507987 |
| PPP3CA | yellow | -0.140985 | 0.492098 | CD37 | yellow | -0.163380642 | 0.415488892 | KARS | yellow | 0.137913 | 0.501675 |
| USP6NL | yellow | -0.140665 | 0.493093 | TCTN1 | yellow | -0.160220192 | 0.42468965 | SCNM1 | yellow | 0.139452 | 0.496867 |
| FCHSD2 | yellow | 0.140467 | 0.493707 | MICAL1 | yellow | -0.155377223 | 0.439002904 | THAP10 | yellow | 0.141016 | 0.492004 |
| SNAPC3 | yellow | -0.140359 | 0.494042 | SMYD4 | yellow | -0.147041334 | 0.464236931 | BIRC2 | yellow | 0.141395 | 0.490828 |
| STRBP | yellow | -0.139672 | 0.49618 | WAC | yellow | -0.139678888 | 0.487138372 | CEP76 | yellow | 0.141788 | 0.489613 |
| VGLL3 | yellow | -0.139284 | 0.49739 | GTPBP4 | yellow | -0.134681122 | 0.503003958 | GPBP1 | yellow | 0.141796 | 0.489586 |
| OSBP | yellow | -0.138813 | 0.498861 | ARHGAP12 | yellow | -0.133787898 | 0.505866255 | RABIF | yellow | 0.142318 | 0.487974 |
| NOSIP | yellow | 0.138784 | 0.498951 | ADCK4 | yellow | -0.128635622 | 0.522532029 | TRIM66 | yellow | 0.142526 | 0.487333 |
| CD36 | yellow | -0.138169 | 0.500872 | SOX4 | yellow | -0.121470595 | 0.546139577 | HPS5 | yellow | 0.142942 | 0.486049 |
| PGAP3 | yellow | 0.13803 | 0.501307 | HELZ2 | yellow | -0.119918267 | 0.551318769 | SCYL3 | yellow | 0.143774 | 0.483486 |
| NDUFS5 | yellow | 0.137976 | 0.501478 | BLOC1S1 | yellow | -0.119479209 | 0.552787743 | DDX39A | yellow | 0.146324 | 0.47568 |
| SFT2D3 | yellow | -0.136663 | 0.505597 | KIF16B | yellow | -0.110999754 | 0.581505067 | PDLIM5 | yellow | 0.146832 | 0.474133 |
| PPDPF | yellow | 0.136469 | 0.506205 | EVL | yellow | -0.098100409 | 0.626399178 | G6PC3 | yellow | 0.147094 | 0.473334 |
| HIRA | yellow | -0.136379 | 0.506489 | TBX19 | yellow | -0.059953192 | 0.766426699 | HSPD1 | yellow | 0.14796 | 0.470707 |

**Table 6. Gene module membership and correlation with growth rate (48/72h), with adjusted p values for each module**

| | | | |
|---|---|---|---|
| GNB1 | yellow | 0.136034 | 0.507576 |
| SEL1L3 | yellow | -0.135824 | 0.508238 |
| MAP1LC3A | yellow | 0.13569 | 0.508661 |
| C10orf137 | yellow | -0.135436 | 0.509461 |
| FOXP4 | yellow | 0.135411 | 0.509539 |
| SHKBP1 | yellow | 0.134994 | 0.510858 |
| SLC9A3R2 | yellow | -0.134681 | 0.511845 |
| GBP4 | yellow | -0.134436 | 0.512621 |
| VLDLR | yellow | 0.132791 | 0.51784 |
| BRCA2 | yellow | -0.132717 | 0.518077 |
| SNORA47 | yellow | -0.132662 | 0.518253 |
| HAX1 | yellow | -0.13182 | 0.520934 |
| CXorf26 | yellow | 0.131044 | 0.523411 |
| FBXO10 | yellow | -0.130438 | 0.525352 |
| TIMM21 | yellow | -0.130203 | 0.526105 |
| ARMC8 | yellow | 0.130154 | 0.526263 |
| SETD9 | yellow | -0.13008 | 0.526498 |
| TBC1D1 | yellow | -0.129844 | 0.527256 |
| IQGAP2 | yellow | -0.129691 | 0.527746 |
| MSANTD2 | yellow | -0.129497 | 0.528369 |
| HAUS5 | yellow | -0.129114 | 0.529599 |
| CNPY3 | yellow | 0.129099 | 0.52965 |
| LHPP | yellow | -0.128697 | 0.530943 |
| TECPR2 | yellow | 0.128696 | 0.530947 |
| CD163 | yellow | 0.12765 | 0.534318 |
| RAD23B | yellow | 0.127598 | 0.534486 |
| ZNF302 | yellow | -0.127132 | 0.53599 |
| CCNJ | yellow | 0.125941 | 0.539849 |
| RNF123 | yellow | 0.125791 | 0.540337 |
| CORO1B | yellow | 0.124919 | 0.543171 |
| ASB3 | yellow | -0.124383 | 0.544916 |
| SPATS2L | yellow | -0.123934 | 0.546378 |
| NCAPH2 | yellow | -0.123402 | 0.548117 |
| RNF2 | yellow | -0.122835 | 0.549969 |
| GPX7 | yellow | 0.12281 | 0.550051 |
| LGR4 | yellow | 0.121881 | 0.553095 |
| KIAA1524 | yellow | -0.121736 | 0.553572 |
| DHX29 | yellow | -0.121515 | 0.554298 |
| CDC6 | yellow | -0.12124 | 0.5552 |
| KIAA1598 | yellow | -0.121147 | 0.555506 |
| NBPF9 | yellow | 0.120961 | 0.55612 |
| PTPLB | yellow | 0.120224 | 0.558546 |
| C9orf64 | yellow | -0.120186 | 0.558671 |
| ABCA2 | yellow | 0.119851 | 0.559775 |
| LOC728377 | yellow | -0.119587 | 0.560647 |
| VASP | yellow | 0.119452 | 0.561091 |
| SERPINB8 | yellow | -0.119409 | 0.561234 |
| RBFOX2 | yellow | 0.119331 | 0.561493 |
| MED22 | yellow | -0.118161 | 0.565364 |
| CHMP2A | yellow | -0.117538 | 0.567429 |
| MALT1 | yellow | -0.117361 | 0.568019 |
| ASUN | yellow | -0.116849 | 0.569717 |
| COX7A2L | yellow | 0.116718 | 0.570154 |
| ATP10D | yellow | -0.115773 | 0.573304 |
| ZNF254 | yellow | -0.115332 | 0.574772 |
| SH3GL1 | yellow | -0.115053 | 0.575706 |
| PDCD5 | yellow | -0.059026644 | 0.769940111 |
| PSMB1 | yellow | -0.037408187 | 0.853036567 |
| SRA1 | yellow | -0.023411307 | 0.907725415 |
| FHL3 | yellow | -0.019874789 | 0.921618508 |
| STOML1 | yellow | -0.015218264 | 0.939945169 |
| PSMF1 | yellow | 0.001212748 | 0.995209988 |
| PXK | yellow | 0.002962258 | 0.98830025 |
| USP7 | yellow | 0.006892051 | 0.972783161 |
| TAF15 | yellow | 0.010932944 | 0.956837514 |
| IL17RA | yellow | 0.024351619 | 0.904035718 |
| ARL3 | yellow | 0.034893724 | 0.862819736 |
| STIL | yellow | 0.037015726 | 0.854562177 |
| CASP2 | yellow | 0.054954903 | 0.78543163 |
| PDK2 | yellow | 0.057361313 | 0.776266055 |
| BRAP | yellow | 0.058746699 | 0.771002518 |
| NFATC2IP | yellow | 0.064229278 | 0.75027138 |
| PNPLA6 | yellow | 0.066951233 | 0.740040479 |
| C5orf25 | yellow | 0.067910227 | 0.736446108 |
| MCM6 | yellow | 0.073757785 | 0.71464855 |
| MIPEP | yellow | 0.075017029 | 0.709982338 |
| SYNGR2 | yellow | 0.079543384 | 0.693294797 |
| AUH | yellow | 0.088368177 | 0.661164329 |
| ALG13 | yellow | 0.088985686 | 0.65893701 |
| DCLRE1B | yellow | 0.089210346 | 0.658127378 |
| DNAJC4 | yellow | 0.091697336 | 0.649190031 |
| CCNY | yellow | 0.093176787 | 0.643895719 |
| ABTB1 | yellow | 0.097556866 | 0.628321266 |
| MTF2 | yellow | 0.101337358 | 0.615002227 |
| METTL18 | yellow | 0.106112532 | 0.598347979 |
| NXPE3 | yellow | 0.106271368 | 0.597797327 |
| TMEM184C | yellow | 0.108503538 | 0.590081836 |
| COX10 | yellow | 0.115507202 | 0.566158236 |
| CRTC2 | yellow | 0.116818199 | 0.561729115 |
| CLN8 | yellow | 0.119725679 | 0.551962897 |
| BTAF1 | yellow | 0.120251615 | 0.550204676 |
| ANXA4 | yellow | 0.123568323 | 0.539176888 |
| LGALS9 | yellow | 0.123652502 | 0.538898358 |
| TMEM134 | yellow | 0.12366588 | 0.538854101 |
| MRPL2 | yellow | 0.123920178 | 0.53801313 |
| PLCB2 | yellow | 0.125494188 | 0.532821632 |
| CCZ1 | yellow | 0.129559343 | 0.519524774 |
| SLC29A3 | yellow | 0.138500713 | 0.490855517 |
| TEN1 | yellow | 0.138869089 | 0.489691757 |
| ATXN7L3B | yellow | 0.139576051 | 0.487462254 |
| SNAPIN | yellow | 0.142577002 | 0.4780559 |
| ZNF140 | yellow | 0.150401515 | 0.45397516 |
| C5orf43 | yellow | 0.150524051 | 0.453603228 |
| WDR35 | yellow | 0.150863565 | 0.452573547 |
| C22orf32 | yellow | 0.151471039 | 0.450734291 |
| INTS9 | yellow | 0.152773813 | 0.446803297 |
| CD33 | yellow | 0.153651873 | 0.444164199 |
| FAN1 | yellow | 0.157550515 | 0.432547898 |
| IVNS1ABP | yellow | 0.160624631 | 0.423506045 |
| MFSD7 | yellow | 0.160627393 | 0.423497969 |
| FUCA2 | yellow | 0.168338687 | 0.401280122 |
| PMS2CL | yellow | 0.169475288 | 0.398061895 |
| LOC84989 | yellow | 0.148546 | 0.46893 |
| PBX2 | yellow | 0.148654 | 0.468604 |
| PRPF38B | yellow | 0.148955 | 0.467693 |
| PTDSS2 | yellow | 0.151036 | 0.461427 |
| BTBD9 | yellow | 0.151475 | 0.460109 |
| BECN1 | yellow | 0.152638 | 0.456631 |
| CIR1 | yellow | 0.152834 | 0.456048 |
| ADAM17 | yellow | 0.1531 | 0.455255 |
| MTPAP | yellow | 0.153569 | 0.453857 |
| FAM204A | yellow | 0.154213 | 0.451944 |
| ZER1 | yellow | 0.15453 | 0.451004 |
| FASTKD2 | yellow | 0.156138 | 0.446248 |
| POLR3K | yellow | 0.157748 | 0.441513 |
| LCORL | yellow | 0.158668 | 0.438821 |
| FNDC3A | yellow | 0.160722 | 0.432841 |
| SPPL2A | yellow | 0.162088 | 0.428889 |
| TPK1 | yellow | 0.162186 | 0.428605 |
| BMS1P4 | yellow | 0.162616 | 0.427367 |
| NR2C2AP | yellow | 0.162818 | 0.426784 |
| ATP5G2 | yellow | 0.163034 | 0.426164 |
| TNFAIP8 | yellow | 0.163157 | 0.42581 |
| FAM21C | yellow | 0.164396 | 0.422254 |
| STT3A | yellow | 0.165498 | 0.419108 |
| KIF27 | yellow | 0.165645 | 0.418691 |
| UBE2E1 | yellow | 0.166469 | 0.416346 |
| C1orf233 | yellow | 0.166889 | 0.415155 |
| LRMP | yellow | 0.167597 | 0.41315 |
| VPS25 | yellow | 0.168655 | 0.410164 |
| MMD | yellow | 0.169471 | 0.407871 |
| BRAF | yellow | 0.16959 | 0.407536 |
| THAP1 | yellow | 0.171201 | 0.403028 |
| APEH | yellow | 0.173201 | 0.397472 |
| SETD6 | yellow | 0.174414 | 0.394124 |
| CCNT2 | yellow | 0.174574 | 0.393685 |
| GTPBP5 | yellow | 0.176489 | 0.388433 |
| SRSF6 | yellow | 0.176819 | 0.387533 |
| IARS | yellow | 0.176887 | 0.387347 |
| GALNT4 | yellow | 0.17846 | 0.383073 |
| DNA2 | yellow | 0.1799 | 0.379182 |
| CPD | yellow | 0.180304 | 0.378093 |
| GSK3A | yellow | 0.180735 | 0.376937 |
| TAB2 | yellow | 0.181882 | 0.373863 |
| ARID3B | yellow | 0.182327 | 0.372676 |
| SNAP23 | yellow | 0.182698 | 0.371688 |
| METTL9 | yellow | 0.183026 | 0.370814 |
| ATP11B | yellow | 0.183491 | 0.369578 |
| THTPA | yellow | 0.183574 | 0.36936 |
| SORT1 | yellow | 0.183957 | 0.368343 |
| PDE12 | yellow | 0.184105 | 0.367953 |
| MXI1 | yellow | 0.184827 | 0.366045 |
| FBXO48 | yellow | 0.184944 | 0.365735 |
| POGK | yellow | 0.186174 | 0.362499 |
| ZNF33A | yellow | 0.186438 | 0.361806 |
| LOC1005061 | yellow | 0.188787 | 0.355681 |
| KEL | yellow | 0.189147 | 0.354745 |
| SPCS2 | yellow | 0.189947 | 0.352678 |

**Table 6. Gene module membership and correlation with growth rate (48/72h), with adjusted p values for each module**

| | | | | | | | | | | | |
|---|---|---|---|---|---|---|---|---|---|---|---|
| CRTC1 | yellow | 0.114764 | 0.57667 | C10orf12 | yellow | 0.172230319 | 0.390322025 | CAMLG | yellow | 0.190655 | 0.350852 |
| CDIPT | yellow | -0.114517 | 0.577499 | SMAP1 | yellow | 0.181966288 | 0.363666797 | UPF3B | yellow | 0.191173 | 0.349518 |
| IFI30 | yellow | -0.114506 | 0.577534 | DECR2 | yellow | 0.183354866 | 0.359954371 | HSPA1A | yellow | 0.191219 | 0.349401 |
| COASY | yellow | 0.114018 | 0.579169 | MRPL10 | yellow | 0.185354801 | 0.354646877 | MRPS15 | yellow | 0.192382 | 0.346423 |
| IGFBP2 | yellow | -0.11239 | 0.584631 | EXOC4 | yellow | 0.193399151 | 0.333770967 | TMEM161A | yellow | 0.192484 | 0.346163 |
| COX4I1 | yellow | 0.111483 | 0.587687 | C21orf91 | yellow | 0.194267104 | 0.331563963 | C21orf91 | yellow | 0.192528 | 0.34605 |
| RPL13 | yellow | 0.110989 | 0.589353 | WDR70 | yellow | 0.196346301 | 0.326313163 | IMPAD1 | yellow | 0.192822 | 0.345301 |
| YIPF3 | yellow | 0.11072 | 0.590261 | C14orf166 | yellow | 0.19962682 | 0.31813235 | DCPS | yellow | 0.193178 | 0.344393 |
| UMPS | yellow | -0.110538 | 0.590875 | LOC1001290 | yellow | 0.20079561 | 0.315248424 | LEMD3 | yellow | 0.193561 | 0.343419 |
| CTDNEP1 | yellow | 0.110435 | 0.591225 | FAM21C | yellow | 0.201140646 | 0.314400158 | FAM134B | yellow | 0.194035 | 0.342215 |
| FCGR3A | yellow | 0.110044 | 0.592547 | CDC40 | yellow | 0.205259558 | 0.304382826 | LNX2 | yellow | 0.194661 | 0.340629 |
| ICAM2 | yellow | -0.109625 | 0.593965 | MEPCE | yellow | 0.213300149 | 0.285408086 | GABPB2 | yellow | 0.195666 | 0.338093 |
| IRF7 | yellow | 0.10938 | 0.594794 | CDK8 | yellow | 0.215996556 | 0.27921691 | UHRF2 | yellow | 0.196625 | 0.335683 |
| LOC728613 | yellow | -0.109063 | 0.595871 | XRCC1 | yellow | 0.216589803 | 0.27786636 | SNX5 | yellow | 0.198497 | 0.331012 |
| LGALS3BP | yellow | 0.108673 | 0.597193 | ABT1 | yellow | 0.216677594 | 0.277666855 | USP31 | yellow | 0.198632 | 0.330674 |
| NOTCH2NL | yellow | 0.107748 | 0.600336 | AP5S1 | yellow | 0.216827055 | 0.277327418 | C22orf28 | yellow | 0.198781 | 0.330305 |
| CD3EAP | yellow | -0.107521 | 0.601108 | RPL35 | yellow | 0.216849784 | 0.277275821 | ALKBH8 | yellow | 0.199915 | 0.327497 |
| HEATR1 | yellow | -0.10748 | 0.601247 | PLXDC2 | yellow | 0.216990901 | 0.276955615 | TMEM234 | yellow | 0.202611 | 0.320879 |
| NELF | yellow | -0.10731 | 0.601826 | ITPK1 | yellow | 0.217279011 | 0.276302602 | ATG16L2 | yellow | 0.204548 | 0.316175 |
| TMX1 | yellow | -0.105353 | 0.608505 | MR1 | yellow | 0.217484867 | 0.275836625 | NICN1 | yellow | 0.205428 | 0.314053 |
| RTEL1 | yellow | 0.105205 | 0.609012 | TMEM41A | yellow | 0.218045837 | 0.274569366 | LOC387647 | yellow | 0.206434 | 0.311636 |
| ZNF84 | yellow | -0.105186 | 0.609079 | GPX3 | yellow | 0.218974163 | 0.272480441 | TBC1D23 | yellow | 0.206633 | 0.31116 |
| HVCN1 | yellow | -0.10513 | 0.60927 | SEC14L1 | yellow | 0.221418749 | 0.267028566 | MAP1LC3B2 | yellow | 0.207202 | 0.309799 |
| WDFY3 | yellow | 0.103701 | 0.614169 | SLC30A1 | yellow | 0.223848705 | 0.26167959 | NFATC2IP | yellow | 0.20788 | 0.308183 |
| ATP6AP1 | yellow | 0.103675 | 0.614258 | PCED1A | yellow | 0.224697944 | 0.259826697 | SMG1 | yellow | 0.208944 | 0.305658 |
| UBR1 | yellow | -0.103631 | 0.614411 | EDC3 | yellow | 0.225082445 | 0.258990595 | FZD1 | yellow | 0.211418 | 0.299837 |
| E2F1 | yellow | -0.102501 | 0.618296 | TARSL2 | yellow | 0.226904199 | 0.255052958 | PIGC | yellow | 0.212563 | 0.297165 |
| KIF1C | yellow | 0.102386 | 0.618695 | INO80 | yellow | 0.227023797 | 0.254795827 | MESDC2 | yellow | 0.214399 | 0.292914 |
| TRA2B | yellow | 0.101808 | 0.620684 | XYLT2 | yellow | 0.228087611 | 0.252516113 | ZNF75A | yellow | 0.214633 | 0.292375 |
| ALDH3A2 | yellow | -0.101489 | 0.621787 | DNASE2B | yellow | 0.230115631 | 0.248207204 | DBNL | yellow | 0.214706 | 0.292206 |
| LOC440173 | yellow | 0.101123 | 0.623049 | ACOT7 | yellow | 0.230605518 | 0.247173631 | HBEGF | yellow | 0.215329 | 0.290775 |
| SRA1 | yellow | -0.100425 | 0.625463 | TRANK1 | yellow | 0.231918676 | 0.244417076 | ZNF546 | yellow | 0.215966 | 0.289316 |
| CAPG | yellow | -0.099861 | 0.627415 | MRPL4 | yellow | 0.232584295 | 0.243027587 | MRPS21 | yellow | 0.21654 | 0.288005 |
| COQ4 | yellow | -0.09951 | 0.628632 | MRPS22 | yellow | 0.232704001 | 0.242778253 | RBM47 | yellow | 0.217524 | 0.285766 |
| HSD3B7 | yellow | 0.099451 | 0.628837 | GPS1 | yellow | 0.241479933 | 0.224957623 | RP2 | yellow | 0.217895 | 0.284925 |
| KIAA1467 | yellow | 0.099145 | 0.6299 | EIF4EBP2 | yellow | 0.248827646 | 0.21072891 | MLH1 | yellow | 0.218036 | 0.284606 |
| DNAJC19 | yellow | -0.098172 | 0.633279 | ARPC2 | yellow | 0.249811154 | 0.20887185 | CELF6 | yellow | 0.221426 | 0.276995 |
| ZNF165 | yellow | -0.098083 | 0.63359 | MED31 | yellow | 0.24995322 | 0.208604523 | VLDLR | yellow | 0.221562 | 0.276694 |
| SNORA68 | yellow | -0.097871 | 0.634324 | QSER1 | yellow | 0.252502658 | 0.203846751 | ABCA3 | yellow | 0.222538 | 0.274529 |
| TMEM160 | yellow | -0.097443 | 0.635816 | SPIN4 | yellow | 0.253040327 | 0.202852894 | OSBPL3 | yellow | 0.224465 | 0.270287 |
| SOAT1 | yellow | 0.096783 | 0.638116 | POC1B | yellow | 0.253338046 | 0.202304 | HTRA2 | yellow | 0.225106 | 0.268884 |
| STAT5A | yellow | -0.096646 | 0.638595 | ZNF271 | yellow | 0.257381673 | 0.194949357 | SSH2 | yellow | 0.225785 | 0.267405 |
| LONRF3 | yellow | 0.096433 | 0.639336 | KIAA0146 | yellow | 0.258649566 | 0.192681684 | PPP1R3D | yellow | 0.225854 | 0.267254 |
| QRICH1 | yellow | 0.09643 | 0.639347 | PPP2CB | yellow | 0.26410909 | 0.183125101 | TMEM106C | yellow | 0.22612 | 0.266678 |
| ANKRD11 | yellow | 0.096137 | 0.640371 | FAM32A | yellow | 0.264327635 | 0.182749543 | MAX | yellow | 0.228984 | 0.260503 |
| KCTD18 | yellow | -0.096114 | 0.64045 | MMS19 | yellow | 0.265001488 | 0.181594939 | EXOSC3 | yellow | 0.231489 | 0.25518 |
| FAM115A | yellow | -0.096075 | 0.640586 | ZNF585A | yellow | 0.270067761 | 0.173076623 | ENPP4 | yellow | 0.232165 | 0.253756 |
| GTF2E2 | yellow | -0.09517 | 0.643751 | PDE3B | yellow | 0.272436159 | 0.169192183 | MANBA | yellow | 0.234973 | 0.247897 |
| DCAF15 | yellow | 0.094878 | 0.644773 | PIP4K2B | yellow | 0.273194413 | 0.167961633 | MEF2B | yellow | 0.235809 | 0.246168 |
| MED18 | yellow | 0.094816 | 0.644991 | MTFP1 | yellow | 0.27498243 | 0.165084904 | EIF4A2 | yellow | 0.236662 | 0.244415 |
| VSIG4 | yellow | -0.094401 | 0.646447 | RAB38 | yellow | 0.275072595 | 0.164940765 | TFDP1 | yellow | 0.237645 | 0.242403 |
| FTH1 | yellow | 0.094074 | 0.647592 | RRAGA | yellow | 0.275470538 | 0.164305675 | AMDHD2 | yellow | 0.240001 | 0.237626 |
| SAT1 | yellow | -0.093777 | 0.648635 | CHUK | yellow | 0.275957073 | 0.163531547 | DCSTAMP | yellow | 0.240383 | 0.236857 |
| PGBD2 | yellow | 0.092951 | 0.651535 | RNLS | yellow | 0.278471327 | 0.159572205 | BET1 | yellow | 0.240892 | 0.235835 |
| LOC399753 | yellow | -0.092277 | 0.653907 | PARP6 | yellow | 0.280800327 | 0.155965735 | DUSP10 | yellow | 0.241568 | 0.234483 |
| AATF | yellow | -0.091755 | 0.655748 | HSPBP1 | yellow | 0.281679697 | 0.154619237 | NAMPT | yellow | 0.243333 | 0.230978 |

**Table 6. Gene module membership and correlation with growth rate (48/72h), with adjusted p values for each module**

| | | | | | | | | | | | |
|---|---|---|---|---|---|---|---|---|---|---|---|
| SERF2 | yellow | 0.091645 | 0.656133 | RNMTL1 | yellow | 0.282005471 | 0.15412252 | ABCA2 | yellow | 0.243457 | 0.230733 |
| TUFT1 | yellow | 0.090428 | 0.66043 | PHPT1 | yellow | 0.282527327 | 0.153329203 | CASP10 | yellow | 0.244991 | 0.227715 |
| SPTLC2 | yellow | 0.090423 | 0.660448 | PEX13 | yellow | 0.282618763 | 0.153190503 | PDCD4 | yellow | 0.245473 | 0.226773 |
| PBRM1 | yellow | -0.089338 | 0.664287 | EED | yellow | 0.288349662 | 0.144675054 | KLHDC2 | yellow | 0.247104 | 0.223603 |
| PYCARD | yellow | -0.089269 | 0.664533 | NUDCD3 | yellow | 0.289062089 | 0.143640761 | SNRNP48 | yellow | 0.247149 | 0.223516 |
| LRRC45 | yellow | 0.088822 | 0.666115 | RAB5C | yellow | 0.295031105 | 0.135183314 | CD55 | yellow | 0.247995 | 0.221885 |
| ZNF511 | yellow | -0.088587 | 0.666948 | AIG1 | yellow | 0.296099393 | 0.133708599 | PKI55 | yellow | 0.251462 | 0.21528 |
| YTHDC1 | yellow | 0.088071 | 0.668779 | IMPAD1 | yellow | 0.30660169 | 0.119827475 | CSGALNACT | yellow | 0.251642 | 0.21494 |
| OC1001306 | yellow | -0.087416 | 0.671106 | BAG4 | yellow | 0.307674904 | 0.118471073 | ZNF230 | yellow | 0.25288 | 0.212617 |
| SEPHS2 | yellow | -0.086862 | 0.673077 | NF2 | yellow | 0.309240549 | 0.116512612 | SAP30BP | yellow | 0.255466 | 0.207819 |
| ESYT1 | yellow | 0.086196 | 0.675452 | KDSR | yellow | 0.310454304 | 0.115010829 | SATB2 | yellow | 0.257818 | 0.203519 |
| TRAF4 | yellow | 0.085204 | 0.67899 | TMEM234 | yellow | 0.312167421 | 0.112915563 | DMXL2 | yellow | 0.259157 | 0.201098 |
| SIGLEC9 | yellow | -0.084368 | 0.681975 | TP53 | yellow | 0.314287887 | 0.110361335 | URGCP | yellow | 0.25964 | 0.200229 |
| MAD1L1 | yellow | 0.084065 | 0.683059 | PPP2R4 | yellow | 0.317225013 | 0.106894464 | RAB2B | yellow | 0.262697 | 0.194793 |
| SRGN | yellow | -0.084052 | 0.683106 | IARS | yellow | 0.317356149 | 0.106741588 | MMAA | yellow | 0.265395 | 0.19008 |
| PNPT1 | yellow | 0.083896 | 0.683666 | STRADA | yellow | 0.317428099 | 0.106657778 | CHMP5 | yellow | 0.266501 | 0.18817 |
| DISC1 | yellow | -0.08353 | 0.684976 | SPPL3 | yellow | 0.320843305 | 0.102735684 | HNRNPA1 | yellow | 0.268548 | 0.184672 |
| NT5C | yellow | 0.083159 | 0.686302 | GNPNAT1 | yellow | 0.325089052 | 0.098011093 | AKAP8L | yellow | 0.268963 | 0.183968 |
| SERINC3 | yellow | -0.082173 | 0.68984 | MATK | yellow | 0.328471834 | 0.094364881 | SCARNA9 | yellow | 0.269499 | 0.183062 |
| MAPRE1 | yellow | 0.081539 | 0.692119 | UBA3 | yellow | 0.328934459 | 0.093874271 | RAB5C | yellow | 0.269645 | 0.182815 |
| DYNLRB1 | yellow | 0.081433 | 0.692501 | BSG | yellow | 0.330301949 | 0.092435275 | PPP1R21 | yellow | 0.271157 | 0.180278 |
| DUS1L | yellow | 0.081352 | 0.69279 | SPTLC2 | yellow | 0.332205883 | 0.090459546 | ZDBF2 | yellow | 0.271751 | 0.179287 |
| TNKS2 | yellow | -0.081099 | 0.693702 | OXSM | yellow | 0.333363442 | 0.089274019 | SLC4A1AP | yellow | 0.272507 | 0.178034 |
| RAP1GAP2 | yellow | 0.080712 | 0.695093 | ZNF192 | yellow | 0.333860316 | 0.088768756 | HNRNPAB | yellow | 0.276137 | 0.172095 |
| C3orf18 | yellow | 0.080315 | 0.696524 | C11orf73 | yellow | 0.335279695 | 0.087337323 | FAM210B | yellow | 0.277492 | 0.169914 |
| SRC | yellow | -0.080146 | 0.697134 | LIPA | yellow | 0.336246435 | 0.086372423 | NUDT5 | yellow | 0.277634 | 0.169687 |
| FAU | yellow | 0.080046 | 0.697493 | HNRNPD | yellow | 0.337520974 | 0.085112684 | CHMP1B | yellow | 0.281236 | 0.163989 |
| B3GALNT2 | yellow | -0.080002 | 0.697651 | RNF216 | yellow | 0.337838327 | 0.084801194 | ING3 | yellow | 0.281396 | 0.163739 |
| ARL11 | yellow | -0.079988 | 0.6977 | COX6B1 | yellow | 0.339573167 | 0.083113675 | TDRD9 | yellow | 0.281495 | 0.163585 |
| ZNF558 | yellow | -0.079617 | 0.699039 | RPL15 | yellow | 0.342351879 | 0.080464102 | DCUN1D2 | yellow | 0.282628 | 0.161824 |
| BMPR2 | yellow | -0.079538 | 0.699324 | RPL26L1 | yellow | 0.34262698 | 0.080205331 | NOP10 | yellow | 0.285156 | 0.157944 |
| SP4 | yellow | -0.07935 | 0.700001 | ADCY7 | yellow | 0.345346551 | 0.077681196 | C10orf57 | yellow | 0.286217 | 0.156334 |
| PUM2 | yellow | 0.078646 | 0.702546 | R3HDM1 | yellow | 0.345762966 | 0.077300128 | C10orf11 | yellow | 0.291184 | 0.148958 |
| GID4 | yellow | -0.078528 | 0.702972 | SUMO2 | yellow | 0.349626086 | 0.073832691 | ZCCHC8 | yellow | 0.292709 | 0.146744 |
| SNTB1 | yellow | 0.077693 | 0.70599 | THAP7 | yellow | 0.34989691 | 0.073594159 | LOC1005067 | yellow | 0.294364 | 0.144367 |
| EP300 | yellow | -0.077168 | 0.707893 | MRPL39 | yellow | 0.34991757 | 0.073575987 | FBXO31 | yellow | 0.294429 | 0.144275 |
| PGD | yellow | 0.07713 | 0.70803 | LMBR1L | yellow | 0.352659889 | 0.071194279 | HSD17B7P2 | yellow | 0.29488 | 0.143632 |
| GLUD1 | yellow | 0.076817 | 0.709164 | FDXACB1 | yellow | 0.352918331 | 0.070972916 | ZNF669 | yellow | 0.295509 | 0.14274 |
| TMSB10 | yellow | 0.076168 | 0.711517 | IFT172 | yellow | 0.353321963 | 0.070628253 | MIS18BP1 | yellow | 0.299535 | 0.137121 |
| VPS25 | yellow | -0.075595 | 0.713598 | DPF2 | yellow | 0.356248517 | 0.068167644 | FAM162A | yellow | 0.299813 | 0.136739 |
| ATF7IP2 | yellow | -0.074735 | 0.716722 | SFT2D1 | yellow | 0.362596707 | 0.063057784 | USF1 | yellow | 0.299967 | 0.136528 |
| DCUN1D1 | yellow | -0.074159 | 0.718817 | PRMT5 | yellow | 0.362729292 | 0.062954323 | TUG1 | yellow | 0.301557 | 0.13436 |
| ZC3H11A | yellow | -0.073848 | 0.719953 | PCCA | yellow | 0.364519381 | 0.061570305 | MFAP1 | yellow | 0.301862 | 0.133948 |
| COX20 | yellow | 0.073396 | 0.721598 | CDK5RAP2 | yellow | 0.364763195 | 0.061383642 | ZNF267 | yellow | 0.304363 | 0.130596 |
| RMRP | yellow | -0.073203 | 0.722302 | HIF1AN | yellow | 0.365474008 | 0.060841955 | LOC1005060 | yellow | 0.308063 | 0.12575 |
| NOL6 | yellow | 0.072941 | 0.723257 | ARRB2 | yellow | 0.36610091 | 0.060367303 | SCO2 | yellow | 0.308364 | 0.125362 |
| TMBIM1 | yellow | 0.072719 | 0.724067 | COMMD1 | yellow | 0.366616545 | 0.059979059 | TRIM39 | yellow | 0.308569 | 0.125098 |
| NSUN5P1 | yellow | -0.072587 | 0.724549 | MIEN1 | yellow | 0.367083855 | 0.059628881 | PORCN | yellow | 0.309271 | 0.124197 |
| MYH9 | yellow | 0.072562 | 0.72464 | PPM1D | yellow | 0.368686581 | 0.058439955 | FAM40A | yellow | 0.309903 | 0.123389 |
| STAT3 | yellow | -0.072549 | 0.72469 | GTF3C4 | yellow | 0.372233585 | 0.055874436 | C7orf50 | yellow | 0.311496 | 0.121373 |
| NEDD8 | yellow | 0.071883 | 0.727121 | KDELR1 | yellow | 0.378098476 | 0.051826383 | ATP6V0E1 | yellow | 0.312229 | 0.120452 |
| LSM7 | yellow | -0.071277 | 0.729337 | RGCC | yellow | 0.382175047 | 0.049150969 | PRR3 | yellow | 0.31266 | 0.119914 |
| HOXB6 | yellow | -0.070521 | 0.732102 | MTFMT | yellow | 0.385984384 | 0.046750364 | SHKBP1 | yellow | 0.314249 | 0.117943 |
| MAP2K1 | yellow | -0.069972 | 0.734115 | PORCN | yellow | 0.386244266 | 0.046590024 | CTDSP2 | yellow | 0.317457 | 0.114037 |
| CYB5R4 | yellow | -0.069955 | 0.734175 | MAPK1IP1L | yellow | 0.38644103 | 0.046468916 | RTP4 | yellow | 0.318202 | 0.113144 |
| SLC25A51 | yellow | 0.068701 | 0.738777 | ELP5 | yellow | 0.388420614 | 0.045264263 | MIEN1 | yellow | 0.318811 | 0.112418 |

**Table 6. Gene module membership and correlation with growth rate (48/72h), with adjusted p values for each module**

| | | | |
|---|---|---|---|
| FCGR3B | yellow | -0.067537 | 0.743055 |
| PQLC3 | yellow | 0.067177 | 0.74438 |
| SCARNA17 | yellow | 0.06708 | 0.744736 |
| MRPS2 | yellow | 0.06702 | 0.744956 |
| MRPL20 | yellow | -0.066658 | 0.746292 |
| C15orf52 | yellow | -0.066323 | 0.747524 |
| COQ9 | yellow | 0.066109 | 0.748315 |
| ANAPC1 | yellow | -0.066086 | 0.748399 |
| SIPA1L3 | yellow | -0.065675 | 0.749913 |
| METTL15 | yellow | 0.065023 | 0.75232 |
| SGMS2 | yellow | -0.064998 | 0.752413 |
| IFI27L2 | yellow | 0.064908 | 0.752745 |
| UCK1 | yellow | -0.063447 | 0.758146 |
| MUS81 | yellow | 0.06342 | 0.758247 |
| PSAP | yellow | -0.063398 | 0.758327 |
| C11orf63 | yellow | -0.063372 | 0.758424 |
| ZNF737 | yellow | 0.062354 | 0.762192 |
| EIF3J | yellow | -0.061889 | 0.763917 |
| MAK16 | yellow | -0.06188 | 0.763951 |
| PPAN | yellow | 0.061777 | 0.764332 |
| LTA4H | yellow | -0.0617 | 0.764617 |
| SOWAHD | yellow | -0.061605 | 0.764971 |
| DFFA | yellow | -0.060583 | 0.768765 |
| LRRFIP2 | yellow | 0.059856 | 0.771467 |
| RPS2 | yellow | 0.05939 | 0.773202 |
| SLC25A29 | yellow | 0.058986 | 0.774704 |
| ACSF3 | yellow | 0.058821 | 0.775318 |
| ZNF714 | yellow | -0.058366 | 0.777013 |
| PPP1R15A | yellow | 0.058275 | 0.777352 |
| NSD1 | yellow | -0.058188 | 0.777675 |
| GTF2IRD1 | yellow | -0.058123 | 0.777917 |
| LIG4 | yellow | -0.057891 | 0.778784 |
| CSTF3 | yellow | 0.057005 | 0.782087 |
| IFT43 | yellow | 0.056918 | 0.782413 |
| MMD | yellow | 0.056807 | 0.782829 |
| VPS13D | yellow | -0.056763 | 0.782991 |
| APOL3 | yellow | -0.056472 | 0.78408 |
| SIRT6 | yellow | 0.056211 | 0.785052 |
| PIM1 | yellow | 0.055929 | 0.786106 |
| PCYOX1 | yellow | -0.055825 | 0.786494 |
| BBS2 | yellow | 0.055789 | 0.786631 |
| TBC1D17 | yellow | -0.055595 | 0.787357 |
| EMP3 | yellow | 0.055428 | 0.78798 |
| LONP1 | yellow | -0.05478 | 0.790405 |
| SNORA50 | yellow | -0.054436 | 0.791693 |
| TESK2 | yellow | 0.054386 | 0.791877 |
| USP32P2 | yellow | 0.054209 | 0.792543 |
| FRS3 | yellow | -0.053655 | 0.794616 |
| TMEM60 | yellow | -0.053563 | 0.794962 |
| WDR1 | yellow | 0.053269 | 0.796066 |
| PICK1 | yellow | -0.053175 | 0.796416 |
| MLL3 | yellow | -0.052895 | 0.797465 |
| GFPT1 | yellow | 0.052509 | 0.798914 |
| FDXR | yellow | -0.052135 | 0.800318 |
| OGFOD3 | yellow | 0.051901 | 0.801196 |
| ORMDL3 | yellow | 0.051801 | 0.801573 |
| PRKRA | yellow | 0.389749347 | 0.044469625 |
| H2AFV | yellow | 0.390649513 | 0.043937593 |
| MTHFR | yellow | 0.390753413 | 0.043876509 |
| CADPS2 | yellow | 0.390980103 | 0.043743471 |
| MATR3 | yellow | 0.404638732 | 0.036298907 |
| PSEN1 | yellow | 0.405219993 | 0.036006151 |
| DIAPH2 | yellow | 0.407971408 | 0.03464591 |
| RPF1 | yellow | 0.408327248 | 0.034473039 |
| RPS6KA4 | yellow | 0.40909476 | 0.034102525 |
| PANK4 | yellow | 0.409488372 | 0.033913753 |
| CWC27 | yellow | 0.412719028 | 0.032395828 |
| COQ6 | yellow | 0.414239079 | 0.031700774 |
| ACP5 | yellow | 0.418425162 | 0.029848473 |
| CDK14 | yellow | 0.420571461 | 0.028933185 |
| AARS | yellow | 0.421829526 | 0.028407308 |
| NUDT5 | yellow | 0.422479063 | 0.028138837 |
| TMEM147 | yellow | 0.424829846 | 0.02718429 |
| ENDOG | yellow | 0.425026448 | 0.027105661 |
| PSMD11 | yellow | 0.425191349 | 0.027039853 |
| CALM3 | yellow | 0.426435522 | 0.026547493 |
| DNAJA1 | yellow | 0.428615863 | 0.025702193 |
| PINLYP | yellow | 0.431725427 | 0.024534545 |
| ZCWPW1 | yellow | 0.431868521 | 0.024481868 |
| RPS15 | yellow | 0.439786213 | 0.021707225 |
| PNPT1 | yellow | 0.441500988 | 0.021141506 |
| ATP6AP1 | yellow | 0.442735671 | 0.02074168 |
| RAB32 | yellow | 0.448125018 | 0.019068162 |
| HUWE1 | yellow | 0.450548179 | 0.018352675 |
| OLA1 | yellow | 0.452242395 | 0.017865636 |
| FBXW7 | yellow | 0.452481949 | 0.017797636 |
| ITGB1BP1 | yellow | 0.45330657 | 0.017565183 |
| SYS1 | yellow | 0.454813494 | 0.017146845 |
| SOD1 | yellow | 0.45510146 | 0.017067843 |
| IVD | yellow | 0.456909443 | 0.016578647 |
| MAD2L1BP | yellow | 0.457897903 | 0.016316116 |
| UTP15 | yellow | 0.459272451 | 0.015956743 |
| ARL8B | yellow | 0.460681382 | 0.01559518 |
| DHPS | yellow | 0.463167633 | 0.01497365 |
| MYOF | yellow | 0.46688321 | 0.014083055 |
| MGST3 | yellow | 0.470951463 | 0.013158668 |
| EXOG | yellow | 0.479249947 | 0.011428535 |
| NANS | yellow | 0.481042698 | 0.011080811 |
| RPE | yellow | 0.482012918 | 0.010896332 |
| MKI67IP | yellow | 0.49175981 | 0.00918091 |
| EIF4EBP1 | yellow | 0.496701982 | 0.008401405 |
| ATP6V0E1 | yellow | 0.499389666 | 0.008001326 |
| ATP5L | yellow | 0.499943355 | 0.007920925 |
| ACTR8 | yellow | 0.506054445 | 0.007077691 |
| COX6A1 | yellow | 0.506351492 | 0.007038709 |
| NDUFB8 | yellow | 0.52279804 | 0.005144634 |
| CLN6 | yellow | 0.523843401 | 0.005040513 |
| COA1 | yellow | 0.523888212 | 0.005036089 |
| TRIAP1 | yellow | 0.524548187 | 0.004971326 |
| NDUFA13 | yellow | 0.54521045 | 0.003271333 |
| JOSD2 | yellow | 0.546781765 | 0.003165426 |
| CHCHD5 | yellow | 0.549877768 | 0.002965306 |
| LOC441454 | yellow | 0.319253 | 0.111893 |
| KCTD3 | yellow | 0.319431 | 0.111683 |
| LOC1002718[illegible] | yellow | 0.320789 | 0.110083 |
| SHMT1 | yellow | 0.3213 | 0.109486 |
| CREBZF | yellow | 0.325698 | 0.104442 |
| BIN3 | yellow | 0.330059 | 0.099614 |
| ZNF642 | yellow | 0.331909 | 0.097618 |
| DSN1 | yellow | 0.332682 | 0.096791 |
| POLR2H | yellow | 0.333419 | 0.096009 |
| C11orf49 | yellow | 0.334786 | 0.09457 |
| PDLIM7 | yellow | 0.335172 | 0.094167 |
| PLEKHA1 | yellow | 0.336225 | 0.093074 |
| FAM151B | yellow | 0.336313 | 0.092984 |
| KIAA1430 | yellow | 0.337408 | 0.091857 |
| MAPKAPK5 | yellow | 0.337599 | 0.091662 |
| GOLGA8B | yellow | 0.339518 | 0.089717 |
| SRSF7 | yellow | 0.340775 | 0.088459 |
| CASP3 | yellow | 0.342333 | 0.086919 |
| LRRC25 | yellow | 0.344994 | 0.084336 |
| TIAF1 | yellow | 0.348631 | 0.0809 |
| TTC7B | yellow | 0.350865 | 0.078843 |
| ARMC8 | yellow | 0.352509 | 0.077354 |
| TUFT1 | yellow | 0.35277 | 0.07712 |
| SYMPK | yellow | 0.352896 | 0.077007 |
| FGGY | yellow | 0.357304 | 0.073136 |
| RAB22A | yellow | 0.357538 | 0.072935 |
| MAGOHB | yellow | 0.358631 | 0.072 |
| PLEKHA8 | yellow | 0.359315 | 0.07142 |
| RPL7A | yellow | 0.360249 | 0.070633 |
| ZNF138 | yellow | 0.360511 | 0.070414 |
| PSMB2 | yellow | 0.363413 | 0.068019 |
| THG1L | yellow | 0.363643 | 0.067832 |
| ELP5 | yellow | 0.363969 | 0.067567 |
| ZNF211 | yellow | 0.366728 | 0.065359 |
| SUPT5H | yellow | 0.37015 | 0.0627 |
| SESN2 | yellow | 0.375145 | 0.058968 |
| PCF11 | yellow | 0.378905 | 0.056276 |
| SF3B3 | yellow | 0.383281 | 0.053263 |
| MAD2L1BP | yellow | 0.388162 | 0.050052 |
| ZNF215 | yellow | 0.405318 | 0.039957 |
| TUT1 | yellow | 0.406266 | 0.03945 |
| NFS1 | yellow | 0.410146 | 0.037429 |
| FBXL15 | yellow | 0.412723 | 0.036133 |
| FAM156B | yellow | 0.422672 | 0.031461 |
| NUPR1 | yellow | 0.42675 | 0.029692 |
| PSEN1 | yellow | 0.427462 | 0.029391 |
| IVD | yellow | 0.442896 | 0.023456 |
| CDS2 | yellow | 0.456126 | 0.01918 |
| LIMD1 | yellow | 0.485354 | 0.011958 |
| EPS8L1 | yellow | 0.540823 | 0.004335 |
| SLFN5 | yellow | 0.575916 | 0.002078 |

**Table 6. Gene module membership and correlation with growth rate (48/72h), with adjusted p values for each module**

| | | | |
|---|---|---|---|
| CKB | yellow | -0.051115 | 0.80415 |
| FAM131A | yellow | 0.050936 | 0.804822 |
| NUCB1 | yellow | -0.050835 | 0.805202 |
| CSF2RB | yellow | -0.050566 | 0.806215 |
| RPL10A | yellow | -0.048903 | 0.812474 |
| EIF4A2 | yellow | 0.048025 | 0.815783 |
| DEM1 | yellow | -0.047478 | 0.817847 |
| ZNF767 | yellow | -0.047461 | 0.817911 |
| ASPSCR1 | yellow | -0.047263 | 0.818659 |
| PTS | yellow | -0.046978 | 0.819733 |
| TMEM66 | yellow | -0.046701 | 0.820782 |
| FCER1G | yellow | -0.04634 | 0.822146 |
| CDC42EP4 | yellow | 0.046301 | 0.822292 |
| C1QB | yellow | 0.046183 | 0.822738 |
| MPLKIP | yellow | -0.045704 | 0.824548 |
| RPL18A | yellow | -0.043755 | 0.831921 |
| CYCS | yellow | -0.043536 | 0.832753 |
| TXNDC11 | yellow | -0.041823 | 0.839246 |
| CSAD | yellow | 0.040388 | 0.844697 |
| NMRK1 | yellow | -0.040341 | 0.844873 |
| LSP1 | yellow | -0.04009 | 0.845828 |
| PCGF5 | yellow | -0.040082 | 0.845859 |
| INVS | yellow | -0.039992 | 0.846202 |
| DPM2 | yellow | 0.039531 | 0.847955 |
| GSS | yellow | -0.039201 | 0.849207 |
| LOC285359 | yellow | 0.038945 | 0.850183 |
| ZFYVE26 | yellow | 0.038523 | 0.851786 |
| SEC22B | yellow | -0.038052 | 0.853581 |
| RPL15 | yellow | 0.03789 | 0.854198 |
| LOC646762 | yellow | 0.037868 | 0.854282 |
| MAPK1 | yellow | -0.037371 | 0.856175 |
| ZBTB4 | yellow | 0.036967 | 0.857712 |
| IER3 | yellow | -0.03687 | 0.858084 |
| CD80 | yellow | -0.036347 | 0.860077 |
| NFATC3 | yellow | -0.036271 | 0.860366 |
| GLG1 | yellow | 0.036255 | 0.860428 |
| PPP4C | yellow | 0.035745 | 0.862372 |
| ZGLP1 | yellow | -0.035473 | 0.86341 |
| MDK | yellow | -0.035185 | 0.864509 |
| CLPP | yellow | 0.034876 | 0.865688 |
| PCYT2 | yellow | -0.034552 | 0.866924 |
| THTPA | yellow | 0.034404 | 0.86749 |
| RFFL | yellow | 0.034231 | 0.86815 |
| SPSB2 | yellow | 0.033754 | 0.869973 |
| NONO | yellow | -0.033409 | 0.871291 |
| NAA38 | yellow | -0.033277 | 0.871793 |
| CCDC117 | yellow | -0.033165 | 0.872221 |
| FTL | yellow | 0.032841 | 0.87346 |
| THAP11 | yellow | -0.032807 | 0.87359 |
| SLC35B4 | yellow | -0.032733 | 0.873873 |
| SLC37A2 | yellow | -0.032212 | 0.875866 |
| AP1M1 | yellow | 0.031652 | 0.878009 |
| IMMP2L | yellow | 0.031089 | 0.880164 |
| CCDC90A | yellow | -0.031029 | 0.880392 |
| ATAT1 | yellow | 0.030842 | 0.881109 |
| PARP10 | yellow | -0.030416 | 0.882737 |

| | | | |
|---|---|---|---|
| C14orf2 | yellow | 0.554498065 | 0.002686819 |
| YIPF5 | yellow | 0.568334964 | 0.001982643 |
| C14orf1 | yellow | 0.570789924 | 0.001875967 |
| RBFOX2 | yellow | 0.578737003 | 0.001563811 |
| CHRAC1 | yellow | 0.595241543 | 0.00105558 |
| UBE2V2 | yellow | 0.61652539 | 0.0006157 |

**Table 6. Gene module membership and correlation with growth rate (48/72h), with adjusted p values for each module**

| | | | |
|---|---|---|---|
| ANXA7 | yellow | 0.030204 | 0.883551 |
| ENOSF1 | yellow | 0.029827 | 0.884996 |
| BAG2 | yellow | -0.029504 | 0.886233 |
| ARHGEF18 | yellow | 0.029051 | 0.887968 |
| ACSS1 | yellow | 0.028873 | 0.88865 |
| ACOT1 | yellow | -0.028063 | 0.891757 |
| HNMT | yellow | -0.028031 | 0.891879 |
| SS18 | yellow | -0.027889 | 0.892424 |
| ZNF259 | yellow | 0.026883 | 0.896284 |
| FAS | yellow | -0.026664 | 0.897124 |
| C11orf57 | yellow | 0.025922 | 0.899972 |
| ARPC2 | yellow | 0.02571 | 0.900784 |
| RPP21 | yellow | 0.025481 | 0.901665 |
| ZNF654 | yellow | 0.025291 | 0.902397 |
| SLC38A6 | yellow | 0.024413 | 0.905768 |
| RTN4IP1 | yellow | -0.024354 | 0.905996 |
| MTX1 | yellow | -0.02398 | 0.907432 |
| CTSD | yellow | 0.023642 | 0.908731 |
| BIRC6 | yellow | -0.023628 | 0.908786 |
| HYI | yellow | -0.023118 | 0.910749 |
| TMLHE | yellow | -0.022849 | 0.91178 |
| TM9SF4 | yellow | -0.022488 | 0.913171 |
| HLA-DRA | yellow | 0.022329 | 0.913782 |
| NRF1 | yellow | 0.021882 | 0.915502 |
| ZNF766 | yellow | 0.021403 | 0.917344 |
| FBXO25 | yellow | -0.020768 | 0.91979 |
| SNHG5 | yellow | 0.020404 | 0.92119 |
| ABT1 | yellow | -0.019907 | 0.923104 |
| TRAPPC6A | yellow | 0.019197 | 0.925838 |
| ZNF318 | yellow | 0.019164 | 0.925968 |
| DDX19B | yellow | 0.018378 | 0.928996 |
| DCK | yellow | -0.017984 | 0.930514 |
| C21orf33 | yellow | -0.017662 | 0.931755 |
| TCTA | yellow | -0.015702 | 0.939315 |
| HIST1H2AK | yellow | 0.014461 | 0.944104 |
| HIST1H4H | yellow | 0.01441 | 0.944302 |
| KATNB1 | yellow | 0.014181 | 0.945184 |
| QSOX1 | yellow | -0.013696 | 0.947054 |
| USF1 | yellow | -0.013518 | 0.947741 |
| THOC5 | yellow | 0.01294 | 0.949975 |
| CASP9 | yellow | -0.012669 | 0.95102 |
| TMEM150A | yellow | 0.011925 | 0.953893 |
| WBP11 | yellow | 0.011916 | 0.953928 |
| AP3M1 | yellow | -0.011885 | 0.954048 |
| BTRC | yellow | -0.011838 | 0.954231 |
| LANCL2 | yellow | 0.011678 | 0.954847 |
| FAM120A | yellow | -0.010562 | 0.959159 |
| NR2C2AP | yellow | 0.00994 | 0.961562 |
| TCEB1 | yellow | -0.009601 | 0.962871 |
| C9orf142 | yellow | 0.009507 | 0.963236 |
| PPP1R13L | yellow | 0.008622 | 0.966657 |
| CYB5R2 | yellow | 0.008114 | 0.968622 |
| FOXRED1 | yellow | -0.00808 | 0.96875 |
| THRAP3 | yellow | -0.008012 | 0.969015 |
| EPRS | yellow | 0.007998 | 0.969068 |
| TMEM141 | yellow | 0.007933 | 0.969321 |

**Table 6. Gene module membership and correlation with growth rate (48/72h), with adjusted p values for each module**

| | | | |
|---|---|---|---|
| LAGE3 | yellow | -0.007285 | 0.971824 |
| FSD1L | yellow | -0.006513 | 0.974811 |
| UBA52 | yellow | -0.006465 | 0.974996 |
| SIKE1 | yellow | 0.00642 | 0.975167 |
| LENG1 | yellow | 0.005039 | 0.980508 |
| TP53 | yellow | -0.005028 | 0.980553 |
| TOR3A | yellow | 0.004697 | 0.981832 |
| DEF6 | yellow | -0.004562 | 0.982353 |
| NSMCE2 | yellow | -0.003858 | 0.985075 |
| TPT1 | yellow | -0.003534 | 0.98633 |
| PCM1 | yellow | -0.003422 | 0.986763 |
| DPP7 | yellow | 0.002389 | 0.990761 |
| DCAF4 | yellow | -0.002201 | 0.991487 |
| BCAS2 | yellow | -0.001363 | 0.994727 |
| AKT1 | yellow | 0.001202 | 0.995351 |
| OR6N2 | yellow | 0.000599 | 0.997681 |
| APOE | yellow | -0.000247 | 0.999046 |

**Supplementary File 6B. Genes with a significant association between growth rate and expression after FDR correction within modules**

| Hour | GeneID | Module | Rsq | Pvalue | Padj |
|---|---|---|---|---|---|
| 24Hr | RASSF2 | black | -0.72918 | 1.60E-05 | 0.007295 |
| 24Hr | PDCD6IP | black | -0.68535 | 7.99E-05 | 0.016896 |
| 24Hr | PHF21A | black | -0.67532 | 0.000111 | 0.016896 |
| 2Hr | WEE1 | orange | -0.62268 | 0.000681 | 0.028587 |
| 24Hr | TCF3 | yellow | -0.68096 | 9.25E-05 | 0.031354 |
| 24Hr | PDK1 | yellow | -0.67607 | 0.000108 | 0.031354 |
| 24Hr | BID | yellow | -0.63818 | 0.000341 | 0.034359 |
| 24Hr | LDB1 | yellow | -0.63753 | 0.000348 | 0.034359 |
| 24Hr | MBD6 | yellow | -0.62719 | 0.000463 | 0.034359 |
| 24Hr | CMIP | yellow | -0.61632 | 0.000619 | 0.034359 |
| 24Hr | STK25 | yellow | -0.61139 | 0.000704 | 0.034359 |
| 24Hr | ARSA | yellow | -0.60679 | 0.000792 | 0.034359 |
| 24Hr | FASTKD5 | yellow | -0.60194 | 0.000894 | 0.034359 |
| 24Hr | NSRP1 | yellow | -0.58855 | 0.001241 | 0.034359 |
| 24Hr | FYTTD1 | yellow | -0.58582 | 0.001324 | 0.034359 |
| 24Hr | RAE1 | yellow | -0.58083 | 0.00149 | 0.034359 |
| 24Hr | DAXX | yellow | -0.57995 | 0.00152 | 0.034359 |
| 24Hr | KLF10 | yellow | -0.57842 | 0.001575 | 0.034359 |
| 24Hr | TAF11 | yellow | -0.57681 | 0.001635 | 0.034359 |
| 24Hr | UBN1 | yellow | -0.57616 | 0.00166 | 0.034359 |
| 24Hr | PHIP | yellow | -0.57598 | 0.001667 | 0.034359 |
| 24Hr | CDYL2 | yellow | -0.57347 | 0.001765 | 0.034359 |
| 24Hr | SIRT2 | yellow | -0.57131 | 0.001854 | 0.034359 |
| 24Hr | COA4 | yellow | -0.57038 | 0.001893 | 0.034359 |
| 24Hr | CCR1 | yellow | -0.56868 | 0.001967 | 0.034359 |
| 24Hr | ATXN1 | yellow | -0.56593 | 0.002092 | 0.034359 |
| 24Hr | CEBPB | yellow | -0.5659 | 0.002093 | 0.034359 |
| 24Hr | ZNF431 | yellow | -0.56359 | 0.002203 | 0.034359 |
| 24Hr | AFG3L2 | yellow | -0.56228 | 0.002268 | 0.034359 |
| 24Hr | DNAJB2 | yellow | -0.56049 | 0.002359 | 0.034359 |
| 24Hr | PDIA4 | yellow | -0.55987 | 0.002391 | 0.034359 |
| 24Hr | CXCL2 | yellow | -0.5587 | 0.002453 | 0.034359 |
| 24Hr | CHMP1A | yellow | -0.55752 | 0.002517 | 0.034359 |
| 24Hr | HIST1H2BC | yellow | -0.55663 | 0.002566 | 0.034359 |
| 24Hr | DGAT1 | yellow | -0.55614 | 0.002593 | 0.034359 |
| 24Hr | SUPV3L1 | yellow | -0.55598 | 0.002602 | 0.034359 |
| 24Hr | IDH3B | yellow | -0.55553 | 0.002628 | 0.034359 |
| 24Hr | RPUSD1 | yellow | -0.55517 | 0.002649 | 0.034359 |
| 24Hr | RBM42 | yellow | -0.5551 | 0.002652 | 0.034359 |
| 24Hr | GLTPD1 | yellow | -0.55418 | 0.002705 | 0.034359 |
| 24Hr | TMF1 | yellow | -0.55159 | 0.00286 | 0.034359 |
| 24Hr | GPN2 | yellow | -0.55052 | 0.002925 | 0.034359 |
| 24Hr | UBE4A | yellow | -0.54975 | 0.002973 | 0.034359 |

| | | | | | |
|---|---|---|---|---|---|
| 24Hr | MTO1 | yellow | -0.54954 | 0.002987 | 0.034359 |
| 24Hr | ZNF564 | yellow | -0.54782 | 0.003097 | 0.034359 |
| 24Hr | DPP9 | yellow | -0.54772 | 0.003104 | 0.034359 |
| 24Hr | TNFAIP3 | yellow | -0.5449 | 0.003292 | 0.034359 |
| 24Hr | CDAN1 | yellow | -0.54379 | 0.003369 | 0.034359 |
| 24Hr | ST3GAL1 | yellow | -0.54363 | 0.003381 | 0.034359 |
| 24Hr | LOC440434 | yellow | -0.54285 | 0.003436 | 0.034359 |
| 24Hr | STUB1 | yellow | -0.5419 | 0.003504 | 0.034359 |
| 24Hr | CXCL1 | yellow | -0.54082 | 0.003583 | 0.034359 |
| 24Hr | TSG101 | yellow | -0.54044 | 0.003612 | 0.034359 |
| 24Hr | SCARNA5 | yellow | -0.54023 | 0.003627 | 0.034359 |
| 24Hr | PAPSS2 | yellow | -0.53997 | 0.003646 | 0.034359 |
| 24Hr | MGRN1 | yellow | -0.53962 | 0.003673 | 0.034359 |
| 24Hr | SCRIB | yellow | -0.53876 | 0.003739 | 0.034359 |
| 24Hr | FUS | yellow | -0.53867 | 0.003745 | 0.034359 |
| 24Hr | TBC1D2B | yellow | -0.53728 | 0.003853 | 0.034802 |
| 24Hr | COBRA1 | yellow | -0.53283 | 0.004217 | 0.036945 |
| 24Hr | LOC541471 | yellow | -0.53227 | 0.004264 | 0.036945 |
| 24Hr | TRMT61A | yellow | -0.53135 | 0.004344 | 0.036945 |
| 24Hr | MLL | yellow | -0.53132 | 0.004346 | 0.036945 |
| 24Hr | TAF1B | yellow | -0.52993 | 0.004469 | 0.037436 |
| 24Hr | MED28 | yellow | -0.52891 | 0.004561 | 0.037573 |
| 24Hr | ICAM1 | yellow | -0.52831 | 0.004615 | 0.037573 |
| 24Hr | N4BP2 | yellow | -0.52461 | 0.004966 | 0.038326 |
| 24Hr | ELF2 | yellow | -0.52379 | 0.005046 | 0.038326 |
| 24Hr | ETFB | yellow | -0.52138 | 0.005289 | 0.038326 |
| 24Hr | TBC1D10B | yellow | -0.521 | 0.005328 | 0.038326 |
| 24Hr | MAP3K4 | yellow | -0.52072 | 0.005357 | 0.038326 |
| 24Hr | TRIP10 | yellow | -0.52059 | 0.005371 | 0.038326 |
| 24Hr | HIST1H1C | yellow | -0.51882 | 0.005558 | 0.039177 |
| 24Hr | CCDC137 | yellow | -0.51714 | 0.00574 | 0.039691 |
| 24Hr | STAG2 | yellow | -0.51689 | 0.005768 | 0.039691 |
| 24Hr | MPHOSPH8 | yellow | -0.51597 | 0.005871 | 0.03992 |
| 24Hr | TMEM106A | yellow | -0.5124 | 0.006283 | 0.039944 |
| 24Hr | LMAN1 | yellow | -0.51238 | 0.006286 | 0.039944 |
| 24Hr | RPS6KC1 | yellow | -0.51232 | 0.006293 | 0.039944 |
| 24Hr | WFS1 | yellow | -0.51073 | 0.006484 | 0.039944 |
| 24Hr | IRF8 | yellow | -0.51043 | 0.006521 | 0.039944 |
| 24Hr | SART1 | yellow | -0.51022 | 0.006548 | 0.039944 |
| 24Hr | PHYH | yellow | -0.50928 | 0.006665 | 0.039944 |
| 24Hr | ZCCHC2 | yellow | -0.50914 | 0.006681 | 0.039944 |
| 24Hr | SIRT7 | yellow | -0.5088 | 0.006725 | 0.039944 |
| 24Hr | TTC31 | yellow | -0.50799 | 0.006826 | 0.039944 |
| 24Hr | RALBP1 | yellow | -0.50706 | 0.006946 | 0.039944 |
| 24Hr | SF1 | yellow | -0.50682 | 0.006978 | 0.039944 |
| 24Hr | SETD8 | yellow | -0.50676 | 0.006985 | 0.039944 |
| 24Hr | TRAPPC11 | yellow | -0.50567 | 0.007129 | 0.039944 |

| | | | | | |
|---|---|---|---|---|---|
| 24Hr | HTT | yellow | -0.50541 | 0.007163 | 0.039944 |
| 24Hr | NUP205 | yellow | -0.50528 | 0.007181 | 0.039944 |
| 24Hr | CORO2A | yellow | -0.50523 | 0.007187 | 0.039944 |
| 24Hr | SMS | yellow | -0.50459 | 0.007272 | 0.040031 |
| 24Hr | RARA | yellow | -0.50305 | 0.007482 | 0.040788 |
| 24Hr | YBX1 | yellow | -0.50233 | 0.007583 | 0.040788 |
| 24Hr | C15orf39 | yellow | -0.50167 | 0.007674 | 0.040788 |
| 24Hr | C21orf59 | yellow | -0.50008 | 0.007901 | 0.040788 |
| 24Hr | C3 | yellow | -0.4999 | 0.007927 | 0.040788 |
| 24Hr | NLRC4 | yellow | -0.49937 | 0.008004 | 0.040788 |
| 24Hr | STRA13 | yellow | -0.49909 | 0.008045 | 0.040788 |
| 24Hr | UBAC1 | yellow | -0.49685 | 0.008379 | 0.041862 |
| 24Hr | ZCCHC9 | yellow | -0.49617 | 0.008483 | 0.041905 |
| 24Hr | COX14 | yellow | -0.49529 | 0.008619 | 0.042217 |
| 24Hr | ARPC1A | yellow | -0.49399 | 0.008821 | 0.042847 |
| 24Hr | SPNS1 | yellow | -0.49283 | 0.009007 | 0.043143 |
| 24Hr | IL10RA | yellow | -0.49233 | 0.009089 | 0.043143 |
| 24Hr | TRUB1 | yellow | -0.49212 | 0.009122 | 0.043143 |
| 24Hr | MAP2K3 | yellow | -0.491 | 0.009306 | 0.043378 |
| 24Hr | SZT2 | yellow | -0.49019 | 0.009441 | 0.043653 |
| 24Hr | FKBP8 | yellow | -0.4891 | 0.009624 | 0.04415 |
| 24Hr | HGS | yellow | -0.48707 | 0.009975 | 0.044823 |
| 24Hr | WDTC1 | yellow | -0.48661 | 0.010056 | 0.044823 |
| 24Hr | MED24 | yellow | -0.48657 | 0.010064 | 0.044823 |
| 24Hr | NCL | yellow | -0.48647 | 0.010081 | 0.044823 |
| 24Hr | HPS4 | yellow | -0.48509 | 0.010327 | 0.045567 |
| 24Hr | TNRC18 | yellow | -0.48375 | 0.010573 | 0.046297 |
| 24Hr | CRADD | yellow | -0.47962 | 0.011355 | 0.048571 |
| 24Hr | EIF4B | yellow | -0.4781 | 0.011657 | 0.04918 |
| 24Hr | MLXIP | yellow | -0.47674 | 0.01193 | 0.049658 |
| 24Hr | XPA | yellow | -0.47668 | 0.011942 | 0.049658 |
| 24Hr | EXOG | yellow | 0.47925 | 0.011429 | 0.048571 |
| 24Hr | NANS | yellow | 0.481043 | 0.011081 | 0.047796 |
| 24Hr | RPE | yellow | 0.482013 | 0.010896 | 0.047354 |
| 24Hr | MKI67IP | yellow | 0.49176 | 0.009181 | 0.043143 |
| 24Hr | EIF4EBP1 | yellow | 0.496702 | 0.008401 | 0.041862 |
| 24Hr | ATP6V0E1 | yellow | 0.49939 | 0.008001 | 0.040788 |
| 24Hr | ATP5L | yellow | 0.499943 | 0.007921 | 0.040788 |
| 24Hr | ACTR8 | yellow | 0.506054 | 0.007078 | 0.039944 |
| 24Hr | COX6A1 | yellow | 0.506351 | 0.007039 | 0.039944 |
| 24Hr | NDUFB8 | yellow | 0.522798 | 0.005145 | 0.038326 |
| 24Hr | CLN6 | yellow | 0.523843 | 0.005041 | 0.038326 |
| 24Hr | COA1 | yellow | 0.523888 | 0.005036 | 0.038326 |
| 24Hr | TRIAP1 | yellow | 0.524548 | 0.004971 | 0.038326 |
| 24Hr | NDUFA13 | yellow | 0.54521 | 0.003271 | 0.034359 |
| 24Hr | JOSD2 | yellow | 0.546782 | 0.003165 | 0.034359 |
| 24Hr | CHCHD5 | yellow | 0.549878 | 0.002965 | 0.034359 |

| | | | | | |
|---|---|---|---|---|---|
| 24Hr | C14orf2 | yellow | 0.554498 | 0.002687 | 0.034359 |
| 24Hr | YIPF5 | yellow | 0.568335 | 0.001983 | 0.034359 |
| 24Hr | C14orf1 | yellow | 0.57079 | 0.001876 | 0.034359 |
| 24Hr | RBFOX2 | yellow | 0.578737 | 0.001564 | 0.034359 |
| 24Hr | CHRAC1 | yellow | 0.595242 | 0.001056 | 0.034359 |
| 24Hr | UBE2V2 | yellow | 0.616525 | 0.000616 | 0.034359 |
