## Supplemental Table 7 for "Human alveolar macrophage response to *Mycobacterium tuberculosis*: immune characteristics underlying large inter-individual variability"

**Supplementary File 7. Summary of significant correlations between protein levels and gene modules in control and infected AMs at 2, 24 and 72 h**

| Network | Module | Protein | Growth Rate | Correlation | p value |
|---|---|---|---|---|---|
| 2h control | purple | CXCL5 | No | 0.71720538 | 3.74E-05 |
| 2h control | royalblue | CCL3 | No | 0.803094138 | 7.89E-07 |
| 2h control | royalblue | IL6 | No | 0.760285808 | 6.58E-06 |
| 2h infected | darkred | CCL19 | No | 0.794648468 | 1.25E-06 |
| 2h infected | lightcyan | IL1B | No | 0.786457043 | 1.90E-06 |
| 2h infected | purple | CXCL5 | No | 0.740813659 | 1.50E-05 |
| 2h infected | royalblue | CCL3 | No | 0.78146984 | 2.44E-06 |
| 24h control | darkgreen | IL1B | No | -0.741845421 | 1.44E-05 |
| 24h control | yellow | IL16 | No | 0.733784429 | 1.99E-05 |
| 24h infected | blue | CCL3 | No | 0.767235062 | 4.81E-06 |
| 24h infected | blue | | **Yes** | -0.521591493 | 0.00628 |
| 24h infected | darkorange | CCL3 | No | 0.728430885 | 2.45E-05 |
| 24h infected | green | CCL13 | No | -0.742658317 | 1.39E-05 |
| 24h infected | green | IL15 | No | -0.770469814 | 4.14E-06 |
| 24h infected | lightcyan | CCL3 | No | -0.706304363 | 5.52E-05 |
| 24h infected | orange | IL10 | No | 0.783608785 | 2.20E-06 |
| 24h infected | red | CCL3 | No | 0.796979009 | 1.10E-06 |
| 24h infected | red | IL10 | No | -0.723175818 | 2.99E-05 |
| 24h infected | red | | **Yes** | -0.545775692 | 0.00393 |
| 24h infected | salmon | CCL3 | No | 0.754510651 | 8.47E-06 |
| 24h infected | tan | | **Yes** | 0.52391146 | 0.00601 |
| 24h infected | yellow | CCL3 | No | 0.756589968 | 7.74E-06 |
| 24h infected | yellow | | **Yes** | -0.607881927 | 0.00099 |
| 72h control | darkred | CXCL10 | No | 0.740180104 | 1.54E-05 |
| 72h infected | darkgrey | MMP9 | No | -0.726631475 | 2.63E-05 |
| 72h infected | grey | IL15 | No | -0.705979049 | 5.59E-05 |
| 72h infected | grey | MMP9 | No | -0.752202091 | 9.36E-06 |
| 72h infected | pink | CSF2 | No | 0.708702121 | 5.08E-05 |
| 72h infected | tan | CCL19 | No | 0.760456781 | 6.53E-06 |
| 72h infected | tan | IL15 | No | 0.758051202 | 7.26E-06 |
| 72h infected | tan | IL18 | No | 0.74201705 | 1.43E-05 |
| 72h infected | tan | IL1A | No | 0.735941955 | 1.83E-05 |
| 72h infected | tan | TNF | No | 0.78475801 | 2.07E-06 |
