## Supplemental Table 8 for "Human alveolar macrophage response to *Mycobacterium tuberculosis*: immune characteristics underlying large inter-individual variability"

**Supplementary File 8. GO term enrichment for 72 hr modules**

| Module | geneSet | description | link | size | overlap | expect | enrichmer | pValue | FDR | overlapId | userId |
|---|---|---|---|---|---|---|---|---|---|---|---|
| Blue | R-HSA-8953854 | Metabolism of RNA | http://reactome.org/PathwayBrowser/#/R-HSA-8953854 | 674 | 54 | 27.90771 | 1.934949 | 1.87E-06 | 0.00323 | 10073;10250;10605;10907;11017;11097;11157;11218;11340;134430;2071;2107;23283;25904;26168;27292;3190;4850;50628;51002;5441;54802;55006;55039;55131;5515;55339;55644;56902;56915;5696;5707;5901;6132;6147;6165;6170;6227;6232;6233;6434;6626;677;7311;7884;80145;81605;83443;8563;8741;8780;88745;9337;9652 | CNOT8;RPL35A;PPP2CA;RPL8;TPRKB;TRIT1;PSMD1;THOC5;RPS21;EXOSC8;RPL39;PAIP1;UBA52;ETF1;RBM28;RAN;TTC37;PSMB8;WDR33;SNRPA;WDR36;RIOK3;CNOT4;TNFSF13;SNUPN;RPS27;SLBP;RRP36;HNRNPK;TRMT12;DIMT1;SRRM1;CNOT10;GEMIN4;TRMT61B;SNRNP27;SF3B5;OSGEP;THOC7;CSTF2T;SENP3;POLR2L;URM1;PNO1;RPL23A;RPS27A;ERCC3;NUPL2;EXOSC5;TXNL4A;DDX20;TRA2B;LSM6;ZFP36L1 |
| Yellow | R-HSA-72203 | Processing of Capped Intron-Containing Pre-mRNA | http://reactome.org/PathwayBrowser/#/R-HSA-72203 | 243 | 22 | 7.989483 | 2.75362 | 1.63E-05 | 0.02811 | 10084;10283;10421;220988;23165;2521;3181;3184;3188;4904;51362;5356;6430;7536;7919;84248;8480;9092;9129;9589;9688;988 | FYTTD1;RAE1;FUS;SART1;SF1;NUP205;YBX1;HNRNPA3;WTAP;CD2BP2;PQBP1;HNRNPA2B1;DDX39B;CWC27;PLRG1;CDC5L;HNRNPD;HNRNPH2;SRSF5;NUP93;PRPF3;CDC40 |
| Yellow | R-HSA-8953854 | Metabolism of RNA | http://reactome.org/PathwayBrowser/#/R-HSA-8953854 | 674 | 42 | 22.16013 | 1.895296 | 4.36E-05 | 0.03769 | 10084;10283;10419;10421;11056;11224;115708;1975;220988;23165;2521;25821;3181;3184;3188;4691;4904;51121;51362;5356;5682;5689;5717;6138;6208;6209;6230;6430;7536;7919;80153;83732;84135;84248;8480;9092;9129;9491;9589;9688;9861;988 | FYTTD1;RAE1;MTO1;FUS;TRMT61A;SART1;SF1;NUP205;YBX1;NCL;EIF4B;DDX52;HNRNPA3;WTAP;UTP15;CD2BP2;PQBP1;HNRNPA2B1;RPS15;PSMD11;RIOK1;DDX39B;CWC27;PSMD6;PSMA1;PLRG1;CDC5L;PRMT5;RPL26L1;RPL15;HNRNPD;HNRNPH2;SRSF5;NUP93;RPS25;EDC3;PRPF3;RPL35;CDC40;RPS14;PSMB1;PSMF1 |
| Tan | R-HSA-170834 | Signaling by TGF-beta Receptor Complex | http://reactome.org/PathwayBrowser/#/R-HSA-170834 | 73 | 6 | 0.615596 | 9.746652 | 3.28E-05 | 0.03144 | 387;4089;4221;54778;5494;9110 | PPM1A;RHOA;MTMR4;SMAD4;MEN1;RNF111 |
| Tan | R-HSA-5696398 | Nucleotide Excision Repair | http://reactome.org/PathwayBrowser/#/R-HSA-5696398 | 111 | 7 | 0.936043 | 7.478287 | 3.99E-05 | 0.03144 | 10401;10714;1642;29844;54778;6117;8178 | POLD3;PIAS3;RPA1;TFPT;ELL;RNF111;DDB1 |
| Tan | R-HSA-73894 | DNA Repair | http://reactome.org/PathwayBrowser/#/R-HSA-73894 | 316 | 11 | 2.664772 | 4.127933 | 6.63E-05 | 0.03144 | 10401;10714;10973;1642;29844;51720;54778;6117;7374;79184;8178 | UNG;BRCC3;UIMC1;POLD3;PIAS3;RPA1;ASCC3;TFPT;ELL;RNF111;DDB1 |
| Tan | R-HSA-5696399 | Global Genome Nucleotide Excision Repair (GG-NER) | http://reactome.org/PathwayBrowser/#/R-HSA-5696399 | 84 | 6 | 0.708357 | 8.470305 | 7.28E-05 | 0.03144 | 10401;10714;1642;29844;54778;6117 | POLD3;PIAS3;RPA1;TFPT;RNF111;DDB1 |
