## Supplemental Table 9 for "Human alveolar macrophage response to *Mycobacterium tuberculosis*: immune characteristics underlying large inter-individual variability"

**Supplementary File 9A.  Protein levels with gene modules correlations, 2hr control**

**Control, 2hr protein vs. module correlation**

| | CCL13 | CCL17 | CCL19 | CCL2 | CCL20 | CCL22 | CCL3 | CCL4 | CSF2 | CXCL10 | CXCL5 | IL10 | IL15 | IL16 | IL18 | IL1A | IL1B | IL6 | IL7 | IL8 | MMP1 | MMP10 | MMP2 | MMP3 | MMP9 | TNF | VEGFA |
|---|---|---|---|---|---|---|---|---|---|---|---|---|---|---|---|---|---|---|---|---|---|---|---|---|---|---|---|
| MEblack | -0.06 | -0.10 | 0.09 | -0.20 | -0.21 | -0.04 | -0.12 | -0.20 | -0.27 | -0.28 | -0.10 | 0.04 | -0.05 | -0.15 | -0.47 | -0.38 | -0.08 | -0.25 | 0.22 | 0.03 | 0.08 | -0.21 | 0.14 | 0.34 | 0.26 | -0.29 | -0.02 |
| MEblue | -0.02 | 0.05 | 0.22 | -0.06 | -0.43 | 0.08 | -0.19 | -0.20 | -0.31 | -0.30 | -0.13 | -0.12 | -0.13 | 0.11 | -0.51 | -0.27 | -0.21 | -0.43 | -0.01 | -0.16 | -0.01 | -0.19 | -0.06 | -0.02 | -0.19 | -0.38 | -0.20 |
| MEbrown | -0.28 | 0.27 | 0.24 | 0.32 | 0.21 | 0.13 | 0.22 | 0.35 | 0.11 | 0.31 | 0.27 | -0.68 | 0.24 | 0.10 | 0.29 | 0.22 | -0.55 | -0.17 | -0.26 | -0.18 | -0.02 | 0.19 | -0.37 | -0.23 | -0.47 | -0.01 | -0.33 |
| MEcyan | -0.01 | -0.29 | -0.05 | -0.19 | 0.45 | -0.16 | 0.17 | 0.13 | 0.18 | 0.15 | 0.23 | 0.28 | -0.07 | -0.37 | 0.19 | -0.05 | 0.20 | 0.38 | 0.27 | 0.32 | 0.07 | 0.09 | 0.19 | 0.36 | 0.31 | 0.25 | 0.23 |
| MEdarkgreen | -0.27 | 0.27 | 0.24 | 0.34 | 0.05 | 0.15 | 0.17 | 0.32 | 0.09 | 0.27 | 0.26 | -0.66 | 0.12 | 0.24 | 0.28 | 0.22 | -0.56 | -0.19 | -0.30 | -0.16 | -0.08 | 0.19 | -0.41 | -0.26 | -0.59 | 0.06 | -0.39 |
| MEdarkgrey | -0.42 | 0.07 | 0.36 | 0.14 | 0.26 | 0.03 | 0.28 | 0.34 | 0.00 | 0.23 | 0.36 | -0.69 | 0.21 | -0.14 | -0.08 | -0.11 | -0.67 | -0.29 | 0.02 | 0.02 | 0.17 | 0.08 | -0.22 | 0.30 | -0.10 | -0.11 | -0.30 |
| MEdarkorange | 0.01 | -0.13 | -0.40 | -0.17 | -0.23 | -0.18 | -0.29 | -0.39 | -0.10 | -0.21 | -0.62 | 0.43 | 0.08 | 0.13 | 0.33 | -0.03 | 0.40 | 0.02 | 0.05 | 0.01 | -0.10 | -0.04 | 0.09 | 0.09 | 0.25 | 0.02 | 0.05 |
| MEdarkred | -0.03 | 0.23 | 0.51 | 0.32 | 0.22 | 0.33 | 0.32 | 0.43 | 0.11 | 0.34 | 0.40 | -0.64 | 0.03 | 0.13 | -0.15 | 0.12 | -0.51 | -0.23 | -0.16 | 0.06 | 0.33 | 0.15 | -0.31 | -0.07 | -0.23 | -0.06 | -0.18 |
| MEdarkturquoise | 0.18 | -0.19 | 0.01 | -0.30 | -0.41 | -0.12 | -0.25 | -0.47 | -0.22 | -0.45 | -0.67 | 0.44 | -0.09 | 0.16 | 0.08 | -0.18 | 0.40 | -0.04 | -0.03 | -0.10 | -0.25 | -0.05 | 0.04 | -0.05 | -0.05 | -0.14 | -0.12 |
| MEgreen | 0.28 | 0.07 | -0.16 | -0.07 | -0.60 | 0.08 | -0.34 | -0.42 | -0.28 | -0.39 | -0.50 | 0.38 | -0.10 | 0.28 | -0.23 | -0.11 | 0.31 | -0.19 | -0.01 | -0.13 | -0.13 | -0.26 | 0.09 | -0.13 | 0.05 | -0.26 | 0.00 |
| MEgreenyellow | 0.27 | -0.01 | -0.24 | -0.07 | -0.16 | 0.02 | -0.10 | -0.24 | -0.10 | -0.13 | -0.31 | 0.42 | 0.13 | 0.03 | -0.05 | -0.03 | 0.41 | 0.13 | 0.15 | 0.14 | 0.02 | -0.19 | 0.28 | 0.14 | 0.46 | -0.06 | 0.24 |
| MEgrey | -0.23 | -0.07 | 0.01 | -0.14 | -0.05 | -0.08 | -0.08 | -0.10 | -0.19 | -0.10 | 0.01 | -0.20 | 0.17 | -0.26 | -0.50 | -0.33 | -0.26 | -0.27 | 0.24 | -0.06 | 0.18 | -0.12 | 0.17 | 0.32 | 0.38 | -0.25 | 0.09 |
| MEgrey60 | 0.26 | -0.22 | -0.08 | -0.18 | 0.24 | -0.07 | -0.03 | 0.01 | 0.13 | 0.01 | 0.24 | 0.54 | -0.28 | -0.28 | -0.10 | -0.02 | 0.41 | 0.42 | 0.23 | 0.26 | -0.06 | -0.01 | 0.33 | 0.04 | 0.19 | 0.23 | 0.43 |
| MElightcyan | 0.46 | -0.05 | -0.25 | -0.06 | -0.13 | 0.02 | -0.08 | -0.18 | 0.13 | -0.07 | -0.24 | 0.61 | -0.12 | 0.09 | 0.03 | 0.12 | 0.62 | 0.35 | 0.09 | 0.16 | -0.01 | -0.05 | 0.29 | -0.05 | 0.37 | 0.23 | 0.37 |
| MElightgreen | -0.03 | -0.07 | 0.25 | -0.15 | -0.29 | -0.02 | -0.16 | -0.20 | -0.20 | -0.33 | -0.23 | 0.06 | -0.16 | 0.09 | -0.04 | -0.20 | -0.04 | -0.23 | -0.04 | -0.08 | -0.21 | -0.10 | -0.18 | 0.02 | -0.38 | -0.22 | -0.34 |
| MElightyellow | 0.15 | 0.05 | 0.33 | 0.14 | -0.12 | 0.15 | 0.18 | 0.20 | 0.10 | 0.10 | 0.18 | -0.18 | -0.26 | 0.28 | -0.20 | 0.10 | -0.12 | -0.06 | -0.16 | 0.05 | 0.10 | 0.03 | -0.21 | -0.12 | -0.29 | 0.13 | -0.14 |
| MEmagenta | -0.11 | -0.08 | 0.35 | -0.09 | 0.09 | 0.04 | 0.12 | 0.12 | -0.11 | -0.05 | 0.29 | -0.19 | -0.15 | -0.20 | -0.32 | -0.25 | -0.29 | -0.17 | 0.09 | 0.10 | 0.07 | -0.07 | -0.06 | 0.22 | -0.11 | -0.18 | -0.15 |
| MEmidnightblue | 0.56 | 0.22 | 0.09 | 0.34 | 0.56 | 0.38 | 0.61 | 0.56 | 0.54 | 0.54 | 0.35 | 0.21 | -0.03 | 0.15 | 0.49 | 0.51 | 0.32 | 0.62 | -0.09 | 0.49 | 0.01 | 0.09 | -0.15 | -0.11 | 0.05 | 0.32 | 0.10 |
| MEorange | 0.50 | 0.36 | 0.00 | 0.18 | 0.03 | 0.45 | 0.28 | 0.16 | 0.27 | 0.08 | 0.06 | 0.09 | 0.09 | 0.12 | 0.01 | 0.35 | 0.17 | 0.21 | -0.15 | 0.01 | 0.12 | 0.04 | -0.08 | -0.32 | -0.11 | -0.06 | -0.04 |
| MEpink | 0.12 | -0.28 | 0.02 | -0.29 | 0.13 | -0.10 | -0.03 | -0.08 | -0.03 | -0.12 | 0.17 | 0.38 | -0.23 | -0.37 | -0.36 | -0.26 | 0.23 | 0.18 | 0.35 | 0.25 | 0.08 | -0.09 | 0.35 | 0.31 | 0.34 | 0.03 | 0.31 |
| MEpurple | -0.07 | -0.09 | 0.36 | 0.11 | 0.51 | 0.08 | 0.38 | 0.51 | 0.25 | 0.41 | 0.72 | -0.25 | -0.18 | -0.14 | -0.05 | 0.06 | -0.29 | 0.23 | 0.06 | 0.37 | 0.10 | 0.19 | -0.07 | 0.16 | -0.12 | 0.31 | 0.07 |
| MEred | -0.39 | 0.09 | 0.01 | 0.16 | 0.34 | -0.08 | 0.19 | 0.25 | 0.12 | 0.29 | 0.15 | -0.52 | 0.33 | -0.09 | 0.38 | 0.13 | -0.39 | -0.03 | -0.11 | -0.10 | 0.05 | 0.21 | -0.17 | 0.02 | -0.07 | 0.09 | -0.15 |
| MEroyalblue | 0.60 | 0.04 | 0.34 | 0.31 | 0.68 | 0.36 | 0.80 | 0.69 | 0.66 | 0.61 | 0.58 | 0.18 | -0.14 | 0.22 | 0.42 | 0.53 | 0.32 | 0.76 | -0.15 | 0.64 | -0.03 | 0.27 | -0.17 | -0.07 | 0.04 | 0.57 | 0.11 |
| MEsalmon | -0.05 | -0.10 | 0.20 | 0.12 | 0.64 | 0.04 | 0.35 | 0.49 | 0.32 | 0.48 | 0.67 | -0.11 | -0.05 | -0.17 | 0.16 | 0.18 | -0.13 | 0.41 | 0.06 | 0.39 | 0.02 | 0.27 | 0.03 | 0.06 | -0.03 | 0.42 | 0.23 |
| MEskyblue | -0.09 | 0.09 | 0.45 | 0.18 | 0.13 | 0.11 | 0.22 | 0.35 | 0.17 | 0.24 | 0.43 | -0.47 | -0.12 | 0.07 | -0.16 | 0.09 | -0.43 | -0.12 | -0.15 | 0.00 | 0.09 | 0.10 | -0.30 | -0.13 | -0.48 | 0.10 | -0.21 |
| MEtan | 0.21 | -0.30 | -0.36 | -0.19 | 0.24 | -0.17 | 0.02 | -0.07 | 0.21 | 0.08 | -0.03 | 0.59 | -0.17 | -0.15 | 0.26 | 0.07 | 0.57 | 0.53 | 0.22 | 0.33 | 0.05 | 0.13 | 0.35 | 0.20 | 0.49 | 0.43 | 0.43 |
| MEturquoise | -0.07 | -0.19 | -0.39 | -0.10 | 0.24 | -0.21 | -0.04 | -0.07 | 0.13 | 0.13 | -0.12 | 0.26 | 0.12 | -0.10 | 0.32 | 0.07 | 0.29 | 0.31 | 0.13 | 0.18 | 0.05 | 0.13 | 0.22 | 0.18 | 0.46 | 0.29 | 0.32 |
| MEwhite | 0.54 | 0.34 | 0.13 | 0.38 | -0.16 | 0.46 | 0.22 | 0.20 | 0.09 | 0.16 | -0.04 | 0.18 | -0.08 | 0.47 | 0.16 | 0.30 | 0.21 | 0.16 | -0.23 | 0.17 | -0.13 | 0.07 | -0.14 | -0.30 | -0.12 | 0.12 | 0.00 |
| MEyellow | 0.49 | -0.11 | -0.17 | -0.13 | -0.30 | 0.02 | -0.20 | -0.27 | -0.01 | -0.23 | -0.26 | 0.69 | -0.34 | 0.17 | -0.12 | 0.04 | 0.63 | 0.24 | 0.04 | 0.12 | -0.08 | -0.14 | 0.23 | -0.16 | 0.11 | 0.13 | 0.28 |

**Control, 2hr protein vs. module correlation p value**

| | CCL13 | CCL17 | CCL19 | CCL2 | CCL20 | CCL22 | CCL3 | CCL4 | CSF2 | CXCL10 | CXCL5 | IL10 | IL15 | IL16 | IL18 | IL1A | IL1B | IL6 | IL7 | IL8 | MMP1 | MMP10 | MMP2 | MMP3 | MMP9 | TNF | VEGFA |
|---|---|---|---|---|---|---|---|---|---|---|---|---|---|---|---|---|---|---|---|---|---|---|---|---|---|---|---|
| MEblack | 0.77 | 0.61 | 0.68 | 0.32 | 0.29 | 0.84 | 0.57 | 0.33 | 0.19 | 0.17 | 0.63 | 0.83 | 0.82 | 0.45 | 0.02 | 0.06 | 0.71 | 0.21 | 0.28 | 0.89 | 0.68 | 0.31 | 0.48 | 0.09 | 0.20 | 0.15 | 0.91 |
| MEblue | 0.92 | 0.80 | 0.29 | 0.75 | 0.03 | 0.70 | 0.34 | 0.34 | 0.12 | 0.14 | 0.53 | 0.57 | 0.54 | 0.60 | 0.01 | 0.19 | 0.29 | 0.03 | 0.94 | 0.43 | 0.96 | 0.34 | 0.75 | 0.90 | 0.35 | 0.06 | 0.33 |
| MEbrown | 0.17 | 0.19 | 0.23 | 0.11 | 0.31 | 0.52 | 0.28 | 0.08 | 0.58 | 0.12 | 0.19 | 0.00 | 0.23 | 0.61 | 0.15 | 0.29 | 0.00 | 0.42 | 0.20 | 0.37 | 0.93 | 0.36 | 0.06 | 0.26 | 0.01 | 0.97 | 0.10 |
| MEcyan | 0.94 | 0.15 | 0.80 | 0.36 | 0.02 | 0.43 | 0.39 | 0.51 | 0.38 | 0.47 | 0.27 | 0.17 | 0.72 | 0.07 | 0.36 | 0.81 | 0.32 | 0.06 | 0.18 | 0.11 | 0.74 | 0.66 | 0.37 | 0.07 | 0.13 | 0.22 | 0.25 |
| MEdarkgreen | 0.18 | 0.17 | 0.24 | 0.09 | 0.80 | 0.47 | 0.39 | 0.12 | 0.66 | 0.18 | 0.20 | 0.00 | 0.56 | 0.25 | 0.16 | 0.29 | 0.00 | 0.36 | 0.14 | 0.44 | 0.70 | 0.36 | 0.04 | 0.21 | 0.00 | 0.76 | 0.05 |
| MEdarkgrey | 0.03 | 0.72 | 0.07 | 0.50 | 0.19 | 0.89 | 0.17 | 0.09 | 0.99 | 0.25 | 0.07 | 0.00 | 0.30 | 0.50 | 0.70 | 0.60 | 0.00 | 0.16 | 0.90 | 0.91 | 0.41 | 0.69 | 0.29 | 0.14 | 0.62 | 0.58 | 0.14 |
| MEdarkorange | 0.97 | 0.54 | 0.04 | 0.40 | 0.26 | 0.39 | 0.15 | 0.05 | 0.62 | 0.29 | 0.00 | 0.03 | 0.69 | 0.52 | 0.10 | 0.87 | 0.04 | 0.91 | 0.82 | 0.96 | 0.61 | 0.83 | 0.66 | 0.67 | 0.21 | 0.94 | 0.82 |
| MEdarkred | 0.87 | 0.26 | 0.01 | 0.11 | 0.29 | 0.10 | 0.11 | 0.03 | 0.61 | 0.09 | 0.05 | 0.00 | 0.88 | 0.52 | 0.46 | 0.57 | 0.01 | 0.25 | 0.44 | 0.76 | 0.10 | 0.45 | 0.13 | 0.75 | 0.26 | 0.76 | 0.37 |
| MEdarkturquoise | 0.37 | 0.36 | 0.96 | 0.14 | 0.04 | 0.55 | 0.23 | 0.02 | 0.27 | 0.02 | 0.00 | 0.03 | 0.66 | 0.43 | 0.68 | 0.39 | 0.04 | 0.84 | 0.89 | 0.64 | 0.22 | 0.82 | 0.86 | 0.80 | 0.82 | 0.50 | 0.54 |
| MEgreen | 0.17 | 0.73 | 0.45 | 0.73 | 0.00 | 0.71 | 0.09 | 0.03 | 0.17 | 0.05 | 0.01 | 0.06 | 0.62 | 0.17 | 0.25 | 0.58 | 0.13 | 0.36 | 0.97 | 0.54 | 0.52 | 0.21 | 0.65 | 0.52 | 0.80 | 0.21 | 0.99 |
| MEgreenyellow | 0.18 | 0.95 | 0.24 | 0.72 | 0.45 | 0.92 | 0.63 | 0.24 | 0.63 | 0.52 | 0.13 | 0.03 | 0.53 | 0.87 | 0.80 | 0.87 | 0.04 | 0.51 | 0.46 | 0.50 | 0.92 | 0.35 | 0.16 | 0.50 | 0.02 | 0.77 | 0.24 |
| MEgrey | 0.25 | 0.74 | 0.97 | 0.48 | 0.79 | 0.69 | 0.69 | 0.62 | 0.34 | 0.63 | 0.97 | 0.32 | 0.41 | 0.21 | 0.01 | 0.10 | 0.20 | 0.19 | 0.23 | 0.76 | 0.37 | 0.57 | 0.40 | 0.11 | 0.06 | 0.21 | 0.68 |
| MEgrey60 | 0.19 | 0.27 | 0.69 | 0.38 | 0.25 | 0.73 | 0.89 | 0.97 | 0.54 | 0.97 | 0.24 | 0.00 | 0.17 | 0.16 | 0.61 | 0.91 | 0.04 | 0.03 | 0.25 | 0.20 | 0.78 | 0.97 | 0.10 | 0.85 | 0.35 | 0.27 | 0.03 |
| MElightcyan | 0.02 | 0.80 | 0.22 | 0.78 | 0.53 | 0.94 | 0.71 | 0.37 | 0.53 | 0.73 | 0.25 | 0.00 | 0.56 | 0.65 | 0.88 | 0.55 | 0.00 | 0.08 | 0.66 | 0.42 | 0.97 | 0.79 | 0.15 | 0.81 | 0.06 | 0.26 | 0.06 |
| MElightgreen | 0.90 | 0.73 | 0.22 | 0.45 | 0.15 | 0.92 | 0.45 | 0.32 | 0.32 | 0.10 | 0.26 | 0.76 | 0.43 | 0.68 | 0.86 | 0.33 | 0.86 | 0.25 | 0.86 | 0.69 | 0.30 | 0.64 | 0.38 | 0.93 | 0.06 | 0.29 | 0.09 |
| MElightyellow | 0.46 | 0.82 | 0.10 | 0.51 | 0.56 | 0.47 | 0.37 | 0.34 | 0.62 | 0.61 | 0.38 | 0.39 | 0.20 | 0.17 | 0.32 | 0.62 | 0.57 | 0.78 | 0.44 | 0.81 | 0.62 | 0.87 | 0.29 | 0.57 | 0.15 | 0.52 | 0.49 |
| MEmagenta | 0.58 | 0.70 | 0.08 | 0.67 | 0.65 | 0.86 | 0.57 | 0.56 | 0.58 | 0.80 | 0.15 | 0.34 | 0.47 | 0.33 | 0.11 | 0.22 | 0.15 | 0.42 | 0.65 | 0.62 | 0.74 | 0.72 | 0.77 | 0.28 | 0.58 | 0.38 | 0.46 |
| MEmidnightblue | 0.00 | 0.29 | 0.67 | 0.09 | 0.00 | 0.05 | 0.00 | 0.00 | 0.00 | 0.00 | 0.08 | 0.31 | 0.88 | 0.47 | 0.01 | 0.01 | 0.11 | 0.00 | 0.66 | 0.01 | 0.97 | 0.67 | 0.46 | 0.61 | 0.81 | 0.11 | 0.62 |
| MEorange | 0.01 | 0.07 | 0.99 | 0.38 | 0.88 | 0.02 | 0.17 | 0.45 | 0.18 | 0.69 | 0.78 | 0.67 | 0.68 | 0.55 | 0.94 | 0.08 | 0.40 | 0.30 | 0.46 | 0.96 | 0.56 | 0.86 | 0.69 | 0.11 | 0.60 | 0.76 | 0.83 |
| MEpink | 0.57 | 0.16 | 0.91 | 0.16 | 0.52 | 0.64 | 0.88 | 0.70 | 0.88 | 0.54 | 0.39 | 0.06 | 0.27 | 0.06 | 0.07 | 0.19 | 0.26 | 0.39 | 0.08 | 0.21 | 0.71 | 0.65 | 0.08 | 0.12 | 0.09 | 0.89 | 0.13 |
| MEpurple | 0.74 | 0.67 | 0.07 | 0.58 | 0.01 | 0.71 | 0.05 | 0.01 | 0.22 | 0.04 | 0.00 | 0.22 | 0.37 | 0.49 | 0.81 | 0.76 | 0.16 | 0.25 | 0.75 | 0.06 | 0.62 | 0.34 | 0.73 | 0.45 | 0.56 | 0.12 | 0.72 |
| MEred | 0.05 | 0.67 | 0.96 | 0.43 | 0.09 | 0.70 | 0.35 | 0.22 | 0.55 | 0.15 | 0.45 | 0.01 | 0.10 | 0.67 | 0.05 | 0.53 | 0.05 | 0.88 | 0.59 | 0.64 | 0.80 | 0.29 | 0.41 | 0.90 | 0.73 | 0.65 | 0.46 |
| MEroyalblue | 0.00 | 0.85 | 0.09 | 0.12 | 0.00 | 0.07 | 0.00 | 0.00 | 0.00 | 0.00 | 0.00 | 0.38 | 0.50 | 0.28 | 0.03 | 0.01 | 0.11 | 0.00 | 0.47 | 0.00 | 0.89 | 0.18 | 0.39 | 0.73 | 0.84 | 0.00 | 0.61 |
| MEsalmon | 0.81 | 0.63 | 0.32 | 0.56 | 0.00 | 0.86 | 0.08 | 0.01 | 0.11 | 0.01 | 0.00 | 0.60 | 0.80 | 0.42 | 0.42 | 0.37 | 0.54 | 0.04 | 0.79 | 0.05 | 0.91 | 0.18 | 0.88 | 0.76 | 0.88 | 0.03 | 0.25 |
| MEskyblue | 0.65 | 0.68 | 0.02 | 0.38 | 0.52 | 0.58 | 0.28 | 0.08 | 0.42 | 0.24 | 0.03 | 0.02 | 0.55 | 0.74 | 0.43 | 0.67 | 0.03 | 0.56 | 0.46 | 1.00 | 0.66 | 0.64 | 0.14 | 0.53 | 0.01 | 0.64 | 0.31 |
| MEtan | 0.31 | 0.14 | 0.07 | 0.35 | 0.24 | 0.41 | 0.93 | 0.73 | 0.31 | 0.71 | 0.87 | 0.00 | 0.40 | 0.47 | 0.20 | 0.74 | 0.00 | 0.01 | 0.27 | 0.10 | 0.81 | 0.52 | 0.08 | 0.32 | 0.01 | 0.03 | 0.03 |
| MEturquoise | 0.74 | 0.36 | 0.05 | 0.63 | 0.24 | 0.31 | 0.84 | 0.74 | 0.52 | 0.52 | 0.58 | 0.19 | 0.55 | 0.64 | 0.12 | 0.72 | 0.15 | 0.13 | 0.54 | 0.39 | 0.80 | 0.51 | 0.29 | 0.39 | 0.02 | 0.14 | 0.11 |
| MEwhite | 0.00 | 0.09 | 0.53 | 0.06 | 0.44 | 0.02 | 0.28 | 0.32 | 0.65 | 0.44 | 0.84 | 0.38 | 0.69 | 0.01 | 0.44 | 0.14 | 0.30 | 0.42 | 0.26 | 0.40 | 0.53 | 0.74 | 0.51 | 0.14 | 0.56 | 0.54 | 0.98 |
| MEyellow | 0.01 | 0.61 | 0.40 | 0.52 | 0.13 | 0.91 | 0.32 | 0.18 | 0.96 | 0.27 | 0.20 | 0.00 | 0.09 | 0.40 | 0.55 | 0.86 | 0.00 | 0.24 | 0.85 | 0.57 | 0.68 | 0.50 | 0.27 | 0.43 | 0.58 | 0.53 | 0.16 |

**Supplementary File 9B. Protein levels with gene modules correlations, 2hr infected**
**Infected, 2hr protein vs. module correlation**

| | GT2 | CCL13 | CCL17 | CCL19 | CCL2 | CCL20 | CCL22 | CCL3 | CCL4 | CSF2 | CXCL10 | CXCL5 | IL10 | IL15 | IL16 | IL18 | IL1A | IL1B | IL6 | IL7 | IL8 | MMP1 | MMP10 | MMP2 | MMP3 | MMP9 | TNF | VEGFA |
|---|---|---|---|---|---|---|---|---|---|---|---|---|---|---|---|---|---|---|---|---|---|---|---|---|---|---|---|---|
| MEblack | 0.27 | 0.16 | 0.02 | 0.22 | -0.10 | 0.17 | 0.07 | 0.21 | 0.08 | -0.19 | -0.11 | 0.18 | 0.07 | 0.04 | -0.18 | -0.20 | -0.17 | 0.09 | 0.19 | 0.11 | 0.18 | -0.16 | 0.15 | -0.24 | 0.16 | -0.06 | 0.15 | -0.05 |
| MEblue | 0.13 | 0.06 | 0.28 | 0.26 | 0.11 | 0.05 | 0.16 | 0.19 | 0.19 | -0.12 | -0.11 | 0.25 | -0.04 | 0.00 | -0.07 | -0.28 | -0.11 | -0.10 | 0.10 | -0.02 | 0.03 | -0.32 | 0.11 | -0.12 | 0.01 | -0.24 | 0.10 | -0.05 |
| MEbrown | -0.30 | -0.31 | 0.30 | 0.10 | 0.41 | -0.06 | -0.19 | -0.04 | 0.20 | 0.25 | 0.22 | 0.02 | -0.56 | -0.06 | 0.25 | 0.03 | 0.19 | -0.58 | -0.47 | -0.22 | -0.33 | 0.04 | -0.15 | -0.07 | -0.07 | -0.10 | -0.24 | -0.17 |
| MEcyan | 0.30 | -0.01 | -0.50 | -0.10 | -0.33 | 0.15 | -0.11 | -0.06 | -0.17 | -0.05 | 0.10 | 0.04 | 0.24 | 0.06 | -0.30 | 0.16 | -0.05 | 0.21 | 0.16 | 0.18 | 0.29 | 0.30 | 0.15 | -0.13 | 0.14 | 0.29 | 0.03 | 0.18 |
| MEdarkgreen | -0.28 | -0.20 | 0.37 | 0.21 | 0.47 | 0.28 | -0.11 | 0.25 | 0.42 | 0.27 | 0.31 | 0.42 | -0.46 | -0.12 | 0.21 | 0.14 | 0.21 | -0.46 | -0.16 | -0.40 | -0.12 | -0.18 | 0.16 | -0.11 | -0.11 | -0.20 | -0.10 | -0.08 |
| MEdarkgrey | -0.03 | -0.22 | 0.20 | 0.35 | 0.31 | 0.21 | -0.25 | 0.20 | 0.33 | 0.13 | 0.25 | 0.22 | -0.58 | 0.00 | 0.10 | -0.09 | 0.08 | -0.54 | -0.32 | -0.11 | -0.04 | 0.11 | -0.01 | -0.45 | 0.12 | -0.14 | -0.07 | -0.23 |
| MEdarkorange | -0.11 | -0.08 | -0.16 | -0.44 | -0.30 | -0.43 | -0.08 | -0.41 | -0.45 | -0.23 | -0.32 | -0.66 | 0.24 | 0.05 | 0.10 | 0.21 | -0.02 | 0.24 | -0.14 | -0.02 | -0.09 | 0.22 | -0.31 | 0.26 | -0.07 | 0.13 | -0.11 | -0.11 |
| MEdarkred | -0.27 | 0.19 | 0.29 | 0.79 | 0.35 | 0.30 | 0.14 | 0.43 | 0.46 | 0.19 | 0.35 | 0.30 | -0.60 | -0.05 | 0.14 | -0.18 | 0.20 | -0.47 | -0.08 | -0.20 | 0.09 | 0.28 | 0.15 | -0.21 | 0.21 | -0.35 | 0.13 | -0.17 |
| MEdarkturquoise | -0.03 | 0.08 | -0.10 | -0.17 | -0.40 | -0.48 | -0.04 | -0.26 | -0.45 | -0.38 | -0.53 | -0.61 | 0.25 | 0.02 | -0.10 | -0.07 | -0.12 | 0.18 | -0.02 | 0.01 | -0.04 | -0.09 | -0.07 | 0.34 | 0.04 | 0.00 | -0.10 | -0.19 |
| MEgreen | 0.06 | 0.18 | 0.24 | 0.01 | 0.01 | -0.06 | 0.29 | 0.14 | 0.04 | -0.14 | -0.25 | -0.04 | 0.31 | 0.03 | 0.06 | -0.04 | -0.07 | 0.32 | 0.27 | 0.02 | 0.07 | -0.31 | 0.03 | 0.18 | -0.05 | -0.13 | 0.14 | -0.04 |
| MEgreenyellow | 0.12 | 0.03 | -0.05 | -0.22 | -0.12 | -0.15 | 0.10 | -0.05 | -0.16 | -0.06 | -0.16 | -0.32 | 0.43 | 0.18 | 0.14 | 0.10 | -0.09 | 0.53 | 0.18 | 0.35 | 0.11 | 0.04 | -0.18 | 0.09 | 0.06 | 0.19 | 0.07 | 0.05 |
| MEgrey | -0.08 | -0.15 | 0.05 | -0.08 | -0.06 | -0.51 | -0.14 | -0.36 | -0.25 | -0.14 | -0.24 | -0.55 | -0.30 | -0.07 | 0.14 | -0.29 | -0.07 | -0.31 | -0.48 | 0.07 | -0.37 | 0.14 | -0.48 | 0.02 | -0.05 | -0.16 | -0.16 | -0.26 |
| MEgrey60 | 0.45 | 0.14 | -0.31 | -0.07 | -0.20 | 0.27 | 0.18 | 0.12 | 0.01 | 0.02 | 0.10 | 0.41 | 0.44 | 0.12 | -0.37 | 0.01 | -0.07 | 0.35 | 0.46 | 0.26 | 0.37 | -0.10 | 0.34 | -0.10 | 0.03 | 0.22 | 0.18 | 0.41 |
| MElightcyan | -0.02 | 0.40 | -0.10 | -0.21 | -0.12 | 0.10 | 0.36 | 0.15 | -0.06 | 0.10 | -0.07 | -0.15 | 0.65 | 0.07 | 0.17 | 0.34 | 0.12 | 0.79 | 0.54 | 0.16 | 0.24 | 0.01 | 0.09 | 0.43 | -0.09 | 0.19 | 0.26 | 0.16 |
| MElightgreen | 0.10 | 0.08 | 0.07 | 0.11 | -0.12 | -0.07 | 0.00 | 0.06 | -0.04 | -0.22 | -0.26 | 0.00 | -0.01 | -0.04 | -0.15 | -0.10 | -0.03 | -0.11 | 0.02 | -0.17 | 0.07 | -0.24 | 0.14 | 0.04 | -0.01 | -0.18 | -0.04 | -0.18 |
| MElightyellow | -0.09 | 0.30 | 0.17 | 0.41 | 0.18 | 0.40 | 0.26 | 0.46 | 0.36 | 0.15 | 0.17 | 0.40 | 0.01 | -0.15 | 0.08 | 0.02 | 0.14 | 0.08 | 0.34 | -0.15 | 0.21 | -0.07 | 0.30 | 0.06 | 0.02 | -0.20 | 0.23 | 0.01 |
| MEmagenta | 0.31 | 0.04 | -0.01 | 0.38 | -0.01 | 0.30 | 0.02 | 0.25 | 0.20 | -0.07 | 0.09 | 0.47 | -0.10 | 0.00 | -0.35 | -0.22 | -0.13 | -0.17 | 0.14 | 0.05 | 0.24 | -0.08 | 0.29 | -0.36 | 0.21 | -0.07 | 0.08 | 0.05 |
| MEmidnightblue | -0.06 | 0.16 | 0.16 | 0.23 | 0.29 | 0.36 | 0.40 | 0.56 | 0.44 | 0.49 | 0.53 | 0.20 | 0.14 | -0.06 | 0.03 | 0.49 | 0.30 | 0.21 | 0.38 | 0.10 | 0.40 | 0.35 | 0.08 | 0.05 | 0.11 | 0.20 | 0.09 | 0.09 |
| MEorange | -0.10 | 0.12 | 0.34 | 0.14 | 0.20 | 0.03 | 0.50 | 0.25 | 0.18 | 0.13 | 0.10 | 0.20 | 0.25 | 0.13 | -0.06 | 0.05 | -0.02 | 0.30 | 0.29 | 0.11 | 0.01 | -0.11 | 0.00 | 0.22 | -0.06 | -0.02 | 0.11 | 0.06 |
| MEpink | 0.46 | 0.17 | -0.29 | 0.12 | -0.26 | 0.28 | 0.08 | 0.16 | 0.01 | -0.13 | 0.01 | 0.35 | 0.27 | 0.09 | -0.40 | -0.14 | -0.16 | 0.23 | 0.36 | 0.23 | 0.36 | -0.06 | 0.31 | -0.26 | 0.16 | 0.09 | 0.18 | 0.22 |
| MEpurple | 0.27 | 0.09 | -0.06 | 0.36 | 0.17 | 0.58 | 0.07 | 0.44 | 0.43 | 0.22 | 0.45 | 0.74 | -0.10 | -0.05 | -0.20 | 0.02 | 0.06 | -0.14 | 0.30 | -0.01 | 0.40 | 0.01 | 0.47 | -0.44 | 0.06 | 0.02 | 0.19 | 0.27 |
| MEred | -0.30 | -0.22 | -0.04 | -0.11 | 0.09 | -0.09 | -0.34 | -0.20 | -0.08 | 0.09 | 0.11 | -0.23 | -0.39 | -0.01 | 0.16 | 0.13 | 0.10 | -0.34 | -0.42 | -0.16 | -0.26 | 0.22 | -0.19 | -0.04 | 0.01 | 0.08 | -0.22 | -0.15 |
| MEroyalblue | 0.06 | 0.29 | 0.05 | 0.34 | 0.31 | 0.69 | 0.30 | 0.78 | 0.62 | 0.55 | 0.57 | 0.61 | 0.20 | 0.02 | 0.01 | 0.42 | 0.33 | 0.25 | 0.63 | 0.12 | 0.60 | 0.10 | 0.49 | 0.01 | 0.03 | 0.26 | 0.24 | 0.23 |
| MEsalmon | 0.34 | -0.13 | -0.14 | 0.05 | 0.19 | 0.38 | 0.00 | 0.21 | 0.33 | 0.35 | 0.50 | 0.57 | -0.06 | 0.08 | -0.08 | 0.11 | 0.17 | -0.16 | 0.17 | 0.15 | 0.32 | 0.07 | 0.32 | -0.42 | -0.11 | 0.22 | 0.09 | 0.42 |
| MEskyblue | -0.07 | 0.10 | 0.22 | 0.47 | 0.28 | 0.30 | 0.12 | 0.36 | 0.39 | 0.19 | 0.25 | 0.51 | -0.29 | -0.08 | -0.05 | -0.16 | 0.10 | -0.31 | 0.06 | -0.19 | 0.08 | -0.07 | 0.25 | -0.11 | 0.03 | -0.26 | 0.07 | 0.05 |
| MEtan | 0.16 | 0.33 | -0.43 | -0.17 | -0.35 | 0.27 | 0.11 | 0.12 | -0.14 | -0.01 | 0.05 | 0.06 | 0.50 | 0.01 | -0.17 | 0.34 | 0.00 | 0.57 | 0.47 | 0.08 | 0.40 | 0.15 | 0.23 | 0.11 | 0.07 | 0.31 | 0.16 | 0.24 |
| MEturquoise | -0.06 | 0.02 | -0.29 | -0.37 | -0.21 | -0.07 | -0.10 | -0.21 | -0.25 | 0.00 | 0.01 | -0.30 | 0.19 | 0.05 | 0.12 | 0.29 | 0.06 | 0.26 | 0.00 | 0.04 | 0.05 | 0.27 | -0.12 | 0.08 | -0.05 | 0.27 | -0.02 | 0.08 |
| MEwhite | -0.07 | 0.17 | 0.33 | 0.03 | 0.38 | 0.18 | 0.34 | 0.42 | 0.41 | 0.22 | 0.16 | 0.24 | 0.33 | 0.17 | 0.31 | 0.20 | 0.19 | 0.33 | 0.50 | 0.01 | 0.28 | -0.32 | 0.32 | 0.11 | -0.31 | 0.04 | 0.38 | 0.18 |
| MEyellow | 0.14 | 0.34 | -0.07 | -0.04 | -0.15 | 0.10 | 0.36 | 0.16 | -0.02 | -0.04 | -0.13 | 0.08 | 0.56 | 0.01 | -0.06 | 0.09 | 0.00 | 0.56 | 0.52 | 0.09 | 0.27 | -0.16 | 0.20 | 0.29 | -0.06 | 0.00 | 0.23 | 0.19 |

**Infected, 2hr protein vs. module correlation p value**

| | GT2 | CCL13 | CCL17 | CCL19 | CCL2 | CCL20 | CCL22 | CCL3 | CCL4 | CSF2 | CXCL10 | CXCL5 | IL10 | IL15 | IL16 | IL18 | IL1A | IL1B | IL6 | IL7 | IL8 | MMP1 | MMP10 | MMP2 | MMP3 | MMP9 | TNF | VEGFA |
|---|---|---|---|---|---|---|---|---|---|---|---|---|---|---|---|---|---|---|---|---|---|---|---|---|---|---|---|---|
| MEblack | 0.18 | 0.44 | 0.92 | 0.28 | 0.61 | 0.42 | 0.75 | 0.29 | 0.70 | 0.35 | 0.58 | 0.38 | 0.75 | 0.84 | 0.39 | 0.33 | 0.42 | 0.67 | 0.34 | 0.60 | 0.37 | 0.43 | 0.46 | 0.24 | 0.42 | 0.78 | 0.47 | 0.81 |
| MEblue | 0.52 | 0.78 | 0.17 | 0.20 | 0.59 | 0.81 | 0.43 | 0.35 | 0.36 | 0.56 | 0.61 | 0.22 | 0.83 | 0.98 | 0.73 | 0.16 | 0.59 | 0.64 | 0.63 | 0.93 | 0.88 | 0.11 | 0.59 | 0.56 | 0.95 | 0.25 | 0.63 | 0.80 |
| MEbrown | 0.13 | 0.12 | 0.13 | 0.63 | 0.04 | 0.77 | 0.34 | 0.83 | 0.32 | 0.21 | 0.27 | 0.92 | 0.00 | 0.77 | 0.23 | 0.87 | 0.35 | 0.00 | 0.02 | 0.29 | 0.10 | 0.83 | 0.46 | 0.73 | 0.74 | 0.61 | 0.25 | 0.42 |
| MEcyan | 0.14 | 0.95 | 0.01 | 0.63 | 0.10 | 0.46 | 0.58 | 0.79 | 0.41 | 0.81 | 0.62 | 0.83 | 0.24 | 0.78 | 0.13 | 0.44 | 0.82 | 0.31 | 0.43 | 0.37 | 0.15 | 0.14 | 0.46 | 0.52 | 0.49 | 0.14 | 0.87 | 0.38 |
| MEdarkgreen | 0.17 | 0.33 | 0.06 | 0.30 | 0.02 | 0.16 | 0.61 | 0.21 | 0.03 | 0.18 | 0.13 | 0.03 | 0.02 | 0.55 | 0.31 | 0.50 | 0.30 | 0.02 | 0.42 | 0.04 | 0.55 | 0.38 | 0.42 | 0.60 | 0.59 | 0.33 | 0.64 | 0.68 |
| MEdarkgrey | 0.89 | 0.27 | 0.32 | 0.08 | 0.12 | 0.31 | 0.21 | 0.33 | 0.10 | 0.54 | 0.23 | 0.28 | 0.00 | 0.99 | 0.63 | 0.67 | 0.70 | 0.00 | 0.11 | 0.60 | 0.86 | 0.59 | 0.96 | 0.02 | 0.57 | 0.48 | 0.72 | 0.26 |
| MEdarkorange | 0.59 | 0.68 | 0.44 | 0.02 | 0.14 | 0.03 | 0.70 | 0.04 | 0.02 | 0.25 | 0.11 | 0.00 | 0.24 | 0.79 | 0.61 | 0.31 | 0.94 | 0.23 | 0.49 | 0.94 | 0.66 | 0.29 | 0.12 | 0.19 | 0.75 | 0.54 | 0.58 | 0.60 |
| MEdarkred | 0.18 | 0.34 | 0.15 | 0.00 | 0.08 | 0.13 | 0.51 | 0.03 | 0.02 | 0.36 | 0.08 | 0.14 | 0.00 | 0.80 | 0.48 | 0.38 | 0.32 | 0.02 | 0.70 | 0.32 | 0.66 | 0.16 | 0.46 | 0.30 | 0.31 | 0.08 | 0.51 | 0.41 |
| MEdarkturquoise | 0.88 | 0.71 | 0.64 | 0.41 | 0.04 | 0.01 | 0.83 | 0.20 | 0.02 | 0.06 | 0.00 | 0.00 | 0.22 | 0.93 | 0.63 | 0.74 | 0.56 | 0.37 | 0.92 | 0.98 | 0.84 | 0.67 | 0.74 | 0.08 | 0.83 | 0.99 | 0.63 | 0.34 |
| MEgreen | 0.76 | 0.39 | 0.23 | 0.98 | 0.97 | 0.76 | 0.16 | 0.50 | 0.85 | 0.50 | 0.22 | 0.85 | 0.12 | 0.89 | 0.76 | 0.85 | 0.73 | 0.11 | 0.17 | 0.92 | 0.72 | 0.12 | 0.88 | 0.39 | 0.80 | 0.54 | 0.49 | 0.85 |
| MEgreenyellow | 0.57 | 0.90 | 0.81 | 0.28 | 0.55 | 0.47 | 0.64 | 0.79 | 0.43 | 0.78 | 0.43 | 0.11 | 0.03 | 0.38 | 0.51 | 0.63 | 0.65 | 0.01 | 0.37 | 0.08 | 0.60 | 0.85 | 0.38 | 0.68 | 0.77 | 0.34 | 0.75 | 0.79 |
| MEgrey | 0.70 | 0.48 | 0.82 | 0.71 | 0.76 | 0.01 | 0.49 | 0.07 | 0.22 | 0.51 | 0.24 | 0.00 | 0.14 | 0.74 | 0.48 | 0.15 | 0.75 | 0.13 | 0.01 | 0.74 | 0.06 | 0.50 | 0.01 | 0.92 | 0.81 | 0.43 | 0.43 | 0.19 |
| MEgrey60 | 0.02 | 0.49 | 0.12 | 0.73 | 0.34 | 0.19 | 0.38 | 0.57 | 0.94 | 0.94 | 0.64 | 0.04 | 0.02 | 0.56 | 0.06 | 0.97 | 0.72 | 0.08 | 0.02 | 0.20 | 0.06 | 0.61 | 0.09 | 0.64 | 0.90 | 0.27 | 0.39 | 0.04 |
| MElightcyan | 0.93 | 0.05 | 0.62 | 0.31 | 0.57 | 0.64 | 0.07 | 0.46 | 0.78 | 0.61 | 0.74 | 0.46 | 0.00 | 0.75 | 0.41 | 0.09 | 0.56 | 0.00 | 0.00 | 0.43 | 0.23 | 0.95 | 0.67 | 0.03 | 0.66 | 0.35 | 0.21 | 0.45 |
| MElightgreen | 0.63 | 0.71 | 0.74 | 0.58 | 0.56 | 0.74 | 1.00 | 0.78 | 0.86 | 0.29 | 0.20 | 0.99 | 0.95 | 0.85 | 0.46 | 0.62 | 0.89 | 0.58 | 0.92 | 0.41 | 0.75 | 0.23 | 0.51 | 0.84 | 0.97 | 0.38 | 0.86 | 0.39 |
| MElightyellow | 0.64 | 0.13 | 0.39 | 0.04 | 0.37 | 0.05 | 0.21 | 0.02 | 0.07 | 0.45 | 0.39 | 0.04 | 0.96 | 0.48 | 0.69 | 0.92 | 0.50 | 0.70 | 0.09 | 0.47 | 0.31 | 0.74 | 0.14 | 0.76 | 0.91 | 0.33 | 0.25 | 0.98 |
| MEmagenta | 0.12 | 0.86 | 0.96 | 0.06 | 0.98 | 0.14 | 0.93 | 0.21 | 0.32 | 0.72 | 0.66 | 0.02 | 0.63 | 1.00 | 0.08 | 0.29 | 0.53 | 0.41 | 0.51 | 0.80 | 0.24 | 0.69 | 0.14 | 0.07 | 0.30 | 0.75 | 0.70 | 0.81 |
| MEmidnightblue | 0.78 | 0.43 | 0.42 | 0.25 | 0.15 | 0.07 | 0.04 | 0.00 | 0.02 | 0.01 | 0.01 | 0.34 | 0.51 | 0.77 | 0.88 | 0.01 | 0.13 | 0.29 | 0.05 | 0.61 | 0.04 | 0.08 | 0.69 | 0.82 | 0.58 | 0.33 | 0.67 | 0.66 |
| MEorange | 0.64 | 0.55 | 0.09 | 0.49 | 0.33 | 0.89 | 0.01 | 0.21 | 0.37 | 0.52 | 0.62 | 0.33 | 0.23 | 0.52 | 0.77 | 0.81 | 0.93 | 0.14 | 0.14 | 0.58 | 0.96 | 0.58 | 1.00 | 0.29 | 0.79 | 0.91 | 0.58 | 0.78 |
| MEpink | 0.02 | 0.39 | 0.15 | 0.55 | 0.21 | 0.17 | 0.69 | 0.43 | 0.97 | 0.53 | 0.97 | 0.08 | 0.18 | 0.68 | 0.04 | 0.49 | 0.43 | 0.26 | 0.07 | 0.27 | 0.07 | 0.79 | 0.13 | 0.21 | 0.44 | 0.65 | 0.37 | 0.29 |
| MEpurple | 0.19 | 0.68 | 0.77 | 0.07 | 0.42 | 0.00 | 0.75 | 0.02 | 0.03 | 0.27 | 0.02 | 0.00 | 0.64 | 0.81 | 0.33 | 0.94 | 0.77 | 0.51 | 0.13 | 0.96 | 0.04 | 0.96 | 0.01 | 0.03 | 0.77 | 0.93 | 0.35 | 0.18 |
| MEred | 0.14 | 0.29 | 0.85 | 0.59 | 0.68 | 0.65 | 0.09 | 0.33 | 0.70 | 0.67 | 0.60 | 0.27 | 0.05 | 0.97 | 0.44 | 0.52 | 0.63 | 0.09 | 0.03 | 0.45 | 0.20 | 0.28 | 0.35 | 0.85 | 0.96 | 0.69 | 0.29 | 0.47 |
| MEroyalblue | 0.77 | 0.15 | 0.80 | 0.09 | 0.12 | 0.00 | 0.14 | 0.00 | 0.00 | 0.00 | 0.00 | 0.00 | 0.33 | 0.91 | 0.96 | 0.03 | 0.10 | 0.22 | 0.00 | 0.57 | 0.00 | 0.63 | 0.01 | 0.97 | 0.87 | 0.21 | 0.24 | 0.27 |
| MEsalmon | 0.09 | 0.53 | 0.49 | 0.81 | 0.35 | 0.06 | 0.98 | 0.31 | 0.10 | 0.08 | 0.01 | 0.00 | 0.78 | 0.70 | 0.69 | 0.61 | 0.41 | 0.43 | 0.41 | 0.46 | 0.11 | 0.72 | 0.11 | 0.03 | 0.58 | 0.28 | 0.67 | 0.03 |
| MEskyblue | 0.73 | 0.64 | 0.28 | 0.02 | 0.17 | 0.14 | 0.54 | 0.07 | 0.05 | 0.35 | 0.22 | 0.01 | 0.15 | 0.69 | 0.81 | 0.43 | 0.62 | 0.13 | 0.75 | 0.35 | 0.70 | 0.74 | 0.23 | 0.60 | 0.89 | 0.20 | 0.75 | 0.82 |
| MEtan | 0.43 | 0.10 | 0.03 | 0.40 | 0.08 | 0.18 | 0.59 | 0.55 | 0.49 | 0.95 | 0.80 | 0.76 | 0.01 | 0.96 | 0.41 | 0.09 | 1.00 | 0.00 | 0.01 | 0.70 | 0.04 | 0.47 | 0.27 | 0.60 | 0.72 | 0.13 | 0.45 | 0.24 |
| MEturquoise | 0.78 | 0.92 | 0.15 | 0.06 | 0.30 | 0.74 | 0.63 | 0.30 | 0.22 | 0.99 | 0.95 | 0.14 | 0.36 | 0.79 | 0.57 | 0.16 | 0.76 | 0.21 | 0.98 | 0.83 | 0.80 | 0.19 | 0.57 | 0.69 | 0.82 | 0.18 | 0.93 | 0.69 |
| MEwhite | 0.74 | 0.41 | 0.10 | 0.88 | 0.05 | 0.37 | 0.09 | 0.03 | 0.04 | 0.27 | 0.42 | 0.24 | 0.10 | 0.40 | 0.13 | 0.33 | 0.35 | 0.10 | 0.01 | 0.95 | 0.17 | 0.11 | 0.11 | 0.59 | 0.13 | 0.84 | 0.05 | 0.37 |
| MEyellow | 0.49 | 0.09 | 0.74 | 0.84 | 0.45 | 0.64 | 0.07 | 0.44 | 0.91 | 0.84 | 0.53 | 0.70 | 0.00 | 0.96 | 0.78 | 0.67 | 0.99 | 0.00 | 0.01 | 0.66 | 0.18 | 0.45 | 0.34 | 0.15 | 0.79 | 0.98 | 0.25 | 0.36 |

**Supplementary File 9C. Protein levels with gene module correlations, 24hr control**

**Control, 24hr protein vs. module correlation**

| | CCL13 | CCL17 | CCL19 | CCL2 | CCL20 | CCL22 | CCL3 | CCL4 | CSF2 | CXCL10 | CXCL5 | IL10 | IL15 | IL16 | IL18 | IL1A | IL1B | IL6 | IL7 | IL8 | MMP1 | MMP10 | MMP2 | MMP3 | MMP9 | TNF | VEGFA |
|---|---|---|---|---|---|---|---|---|---|---|---|---|---|---|---|---|---|---|---|---|---|---|---|---|---|---|---|
| MEblack | -0.16 | 0.27 | 0.13 | 0.37 | 0.07 | 0.42 | 0.11 | -0.15 | -0.20 | 0.17 | 0.21 | -0.57 | -0.24 | 0.51 | 0.00 | 0.17 | -0.60 | -0.19 | 0.23 | 0.32 | 0.04 | 0.15 | 0.36 | 0.47 | -0.40 | 0.16 | 0.12 |
| MEblue | 0.34 | 0.27 | 0.32 | 0.44 | 0.66 | 0.35 | 0.40 | 0.15 | 0.13 | 0.41 | 0.37 | -0.31 | 0.49 | 0.45 | 0.57 | 0.68 | -0.11 | 0.20 | -0.02 | 0.54 | 0.50 | 0.53 | 0.12 | 0.23 | 0.03 | 0.10 | -0.12 |
| MEbrown | -0.28 | -0.10 | -0.45 | -0.08 | -0.39 | -0.25 | -0.27 | -0.21 | 0.04 | -0.34 | -0.36 | 0.51 | -0.10 | 0.01 | -0.36 | -0.09 | 0.41 | 0.07 | -0.14 | -0.50 | -0.04 | -0.20 | -0.52 | -0.15 | 0.14 | 0.02 | 0.11 |
| MEcyan | 0.18 | 0.10 | -0.02 | 0.03 | -0.07 | -0.19 | 0.01 | -0.15 | 0.11 | -0.10 | 0.00 | 0.22 | 0.35 | -0.03 | -0.05 | 0.01 | 0.21 | -0.09 | -0.34 | -0.09 | -0.06 | -0.03 | -0.47 | -0.07 | 0.26 | -0.27 | 0.00 |
| MEdarkgreen | 0.07 | 0.13 | 0.18 | 0.08 | -0.15 | 0.32 | 0.17 | 0.04 | -0.14 | 0.36 | 0.38 | -0.69 | 0.06 | -0.22 | -0.09 | -0.14 | -0.74 | -0.31 | 0.16 | 0.50 | -0.10 | 0.09 | 0.15 | 0.16 | -0.52 | -0.13 | 0.24 |
| MEdarkgrey | 0.43 | 0.35 | 0.21 | 0.42 | 0.52 | 0.36 | 0.33 | -0.02 | 0.69 | 0.56 | 0.14 | 0.21 | 0.20 | 0.12 | 0.31 | 0.46 | 0.32 | 0.62 | -0.42 | 0.13 | 0.03 | -0.17 | -0.06 | -0.04 | 0.35 | 0.00 | -0.03 |
| MEdarkorange | 0.07 | 0.35 | 0.17 | 0.47 | 0.27 | 0.33 | 0.24 | -0.29 | -0.02 | 0.24 | 0.28 | -0.35 | 0.19 | 0.55 | 0.08 | 0.39 | -0.31 | -0.23 | -0.04 | 0.28 | 0.38 | 0.29 | 0.05 | 0.46 | 0.06 | -0.09 | 0.26 |
| MEdarkred | -0.04 | -0.20 | 0.00 | -0.13 | 0.24 | -0.04 | -0.08 | 0.22 | -0.04 | 0.15 | -0.06 | 0.11 | 0.11 | -0.15 | 0.29 | 0.18 | 0.22 | 0.23 | 0.26 | 0.04 | 0.29 | 0.24 | 0.10 | -0.17 | 0.14 | 0.12 | -0.11 |
| MEdarkturquoise | -0.30 | 0.12 | -0.20 | 0.12 | -0.31 | 0.02 | -0.23 | -0.33 | 0.00 | -0.22 | -0.20 | 0.29 | -0.32 | 0.20 | -0.38 | -0.08 | 0.13 | 0.05 | -0.25 | -0.40 | -0.27 | -0.23 | -0.37 | 0.14 | 0.00 | 0.11 | 0.26 |
| MEgreen | -0.67 | -0.25 | -0.58 | -0.25 | -0.57 | -0.36 | -0.50 | -0.17 | -0.42 | -0.68 | -0.63 | 0.34 | -0.54 | 0.32 | -0.42 | -0.22 | 0.20 | -0.19 | 0.28 | -0.53 | -0.09 | -0.06 | -0.12 | 0.03 | -0.09 | 0.26 | -0.20 |
| MEgreenyellow | -0.39 | -0.23 | -0.47 | -0.28 | -0.52 | -0.30 | -0.40 | -0.11 | -0.02 | -0.42 | -0.57 | 0.58 | -0.57 | -0.16 | -0.42 | -0.41 | 0.42 | 0.20 | 0.02 | -0.62 | -0.43 | -0.43 | -0.30 | -0.21 | -0.07 | 0.21 | -0.06 |
| MEgrey | -0.17 | 0.19 | 0.04 | 0.26 | 0.00 | 0.21 | -0.07 | -0.31 | 0.00 | 0.06 | 0.01 | -0.01 | -0.27 | 0.30 | -0.19 | 0.07 | -0.10 | -0.02 | -0.10 | -0.18 | -0.16 | -0.26 | 0.02 | 0.23 | 0.06 | 0.01 | 0.32 |
| MEgrey60 | 0.10 | -0.27 | 0.01 | -0.43 | -0.13 | -0.26 | -0.02 | 0.28 | -0.08 | -0.10 | -0.01 | -0.05 | 0.02 | -0.53 | 0.00 | -0.38 | -0.04 | -0.14 | 0.13 | 0.05 | -0.18 | -0.10 | 0.15 | -0.28 | -0.10 | -0.15 | -0.11 |
| MElightcyan | -0.22 | -0.41 | -0.35 | -0.54 | -0.37 | -0.43 | -0.25 | 0.20 | -0.17 | -0.46 | -0.43 | 0.33 | -0.43 | -0.37 | -0.11 | -0.46 | 0.31 | 0.00 | 0.28 | -0.32 | -0.21 | -0.15 | 0.12 | -0.29 | -0.09 | 0.09 | -0.41 |
| MElightgreen | 0.10 | -0.35 | 0.05 | -0.42 | 0.13 | -0.21 | -0.02 | 0.41 | 0.01 | 0.00 | -0.08 | 0.11 | -0.02 | -0.48 | 0.22 | -0.18 | 0.21 | 0.16 | 0.22 | 0.03 | 0.06 | 0.04 | 0.33 | -0.34 | 0.06 | 0.06 | -0.27 |
| MElightyellow | 0.21 | -0.03 | 0.23 | 0.09 | 0.49 | 0.24 | 0.30 | 0.40 | 0.02 | 0.37 | 0.25 | -0.40 | 0.27 | 0.07 | 0.47 | 0.37 | -0.22 | 0.17 | 0.28 | 0.53 | 0.50 | 0.48 | 0.45 | 0.02 | -0.16 | 0.20 | -0.17 |
| MEmagenta | 0.22 | 0.14 | 0.11 | 0.23 | 0.33 | 0.05 | 0.11 | -0.10 | 0.30 | 0.21 | 0.06 | 0.29 | 0.43 | 0.14 | 0.17 | 0.40 | 0.36 | 0.28 | -0.36 | -0.04 | 0.33 | 0.14 | -0.34 | -0.02 | 0.45 | -0.02 | 0.12 |
| MEmidnightblue | -0.31 | 0.03 | -0.02 | 0.05 | -0.18 | 0.20 | -0.06 | -0.10 | -0.29 | -0.06 | 0.00 | -0.41 | -0.55 | 0.22 | -0.15 | -0.19 | -0.46 | -0.28 | 0.38 | 0.09 | -0.10 | -0.02 | 0.50 | 0.32 | -0.37 | 0.14 | -0.03 |
| MEorange | -0.15 | -0.41 | -0.35 | -0.55 | -0.47 | -0.48 | -0.38 | 0.12 | 0.12 | -0.31 | -0.48 | 0.66 | -0.16 | -0.68 | -0.37 | -0.52 | 0.54 | 0.24 | -0.16 | -0.59 | -0.38 | -0.46 | -0.38 | -0.57 | 0.13 | 0.02 | 0.03 |
| MEpink | -0.13 | -0.16 | -0.15 | -0.31 | -0.46 | -0.17 | -0.09 | 0.03 | -0.27 | -0.23 | -0.07 | -0.25 | -0.16 | -0.30 | -0.21 | -0.48 | -0.32 | -0.40 | 0.25 | 0.09 | -0.16 | 0.02 | 0.12 | -0.02 | -0.35 | -0.11 | -0.17 |
| MEpurple | 0.02 | -0.17 | -0.15 | -0.17 | -0.01 | -0.22 | -0.14 | 0.00 | 0.20 | -0.03 | -0.20 | 0.52 | 0.20 | -0.28 | 0.01 | 0.02 | 0.55 | 0.25 | -0.15 | -0.28 | 0.22 | 0.04 | -0.34 | -0.29 | 0.41 | -0.02 | -0.02 |
| MEred | 0.13 | 0.25 | 0.17 | 0.34 | 0.33 | 0.34 | 0.33 | 0.09 | -0.12 | 0.23 | 0.28 | -0.55 | 0.16 | 0.49 | 0.39 | 0.41 | -0.43 | -0.09 | 0.23 | 0.58 | 0.28 | 0.46 | 0.29 | 0.36 | -0.35 | 0.13 | -0.23 |
| MEroyalblue | -0.40 | -0.14 | -0.35 | -0.24 | -0.64 | -0.19 | -0.32 | -0.28 | -0.18 | -0.35 | -0.34 | 0.17 | -0.52 | -0.20 | -0.61 | -0.55 | -0.03 | -0.23 | 0.05 | -0.45 | -0.39 | -0.44 | -0.14 | -0.02 | -0.14 | -0.04 | 0.22 |
| MEsalmon | 0.35 | 0.11 | 0.21 | 0.16 | 0.37 | 0.07 | 0.31 | 0.16 | 0.07 | 0.26 | 0.30 | -0.27 | 0.62 | 0.13 | 0.30 | 0.38 | -0.14 | -0.06 | -0.10 | 0.43 | 0.41 | 0.38 | -0.04 | 0.04 | 0.08 | -0.15 | 0.02 |
| MEtan | 0.24 | -0.24 | 0.03 | -0.36 | 0.06 | -0.32 | -0.05 | 0.29 | 0.22 | -0.05 | -0.07 | 0.40 | 0.19 | -0.57 | 0.12 | -0.18 | 0.44 | 0.25 | -0.20 | -0.18 | -0.21 | -0.22 | -0.16 | -0.48 | 0.26 | -0.14 | -0.11 |
| MEturquoise | 0.33 | -0.04 | 0.09 | 0.00 | 0.38 | -0.09 | 0.18 | 0.18 | 0.24 | 0.18 | 0.06 | 0.25 | 0.53 | -0.10 | 0.36 | 0.35 | 0.40 | 0.28 | -0.20 | 0.11 | 0.37 | 0.26 | -0.22 | -0.20 | 0.37 | -0.07 | -0.11 |
| MEwhite | -0.40 | -0.08 | -0.24 | -0.10 | -0.47 | 0.00 | -0.20 | -0.16 | -0.31 | -0.20 | -0.19 | -0.24 | -0.44 | 0.02 | -0.44 | -0.38 | -0.39 | -0.28 | 0.24 | -0.10 | -0.12 | -0.10 | 0.16 | 0.19 | -0.40 | 0.12 | 0.21 |
| MEyellow | 0.06 | 0.35 | 0.22 | 0.56 | 0.48 | 0.44 | 0.27 | -0.10 | -0.01 | 0.28 | 0.26 | -0.36 | 0.10 | 0.73 | 0.33 | 0.62 | -0.25 | 0.07 | 0.07 | 0.37 | 0.39 | 0.40 | 0.21 | 0.46 | -0.06 | 0.19 | 0.01 |

**Control, 24hr protein vs. module correlation p value**

| | CCL13 | CCL17 | CCL19 | CCL2 | CCL20 | CCL22 | CCL3 | CCL4 | CSF2 | CXCL10 | CXCL5 | IL10 | IL15 | IL16 | IL18 | IL1A | IL1B | IL6 | IL7 | IL8 | MMP1 | MMP10 | MMP2 | MMP3 | MMP9 | TNF | VEGFA |
|---|---|---|---|---|---|---|---|---|---|---|---|---|---|---|---|---|---|---|---|---|---|---|---|---|---|---|---|
| MEblack | 0.44 | 0.19 | 0.53 | 0.06 | 0.74 | 0.03 | 0.58 | 0.48 | 0.32 | 0.41 | 0.31 | 0.00 | 0.24 | 0.01 | 0.99 | 0.42 | 0.00 | 0.35 | 0.25 | 0.11 | 0.85 | 0.46 | 0.07 | 0.02 | 0.05 | 0.43 | 0.57 |
| MEblue | 0.08 | 0.18 | 0.11 | 0.03 | 0.00 | 0.08 | 0.04 | 0.46 | 0.51 | 0.04 | 0.06 | 0.13 | 0.01 | 0.02 | 0.00 | 0.00 | 0.58 | 0.33 | 0.92 | 0.00 | 0.01 | 0.01 | 0.55 | 0.26 | 0.88 | 0.61 | 0.55 |
| MEbrown | 0.16 | 0.62 | 0.02 | 0.69 | 0.05 | 0.21 | 0.18 | 0.31 | 0.86 | 0.09 | 0.07 | 0.01 | 0.62 | 0.94 | 0.07 | 0.67 | 0.04 | 0.74 | 0.50 | 0.01 | 0.84 | 0.33 | 0.01 | 0.47 | 0.51 | 0.92 | 0.59 |
| MEcyan | 0.39 | 0.63 | 0.91 | 0.89 | 0.72 | 0.35 | 0.97 | 0.46 | 0.58 | 0.61 | 1.00 | 0.28 | 0.08 | 0.89 | 0.82 | 0.96 | 0.31 | 0.66 | 0.09 | 0.66 | 0.78 | 0.90 | 0.02 | 0.72 | 0.19 | 0.18 | 0.99 |
| MEdarkgreen | 0.74 | 0.53 | 0.39 | 0.71 | 0.45 | 0.11 | 0.41 | 0.84 | 0.51 | 0.07 | 0.06 | 0.00 | 0.77 | 0.27 | 0.66 | 0.48 | 0.00 | 0.13 | 0.43 | 0.01 | 0.64 | 0.66 | 0.47 | 0.45 | 0.01 | 0.54 | 0.24 |
| MEdarkgrey | 0.03 | 0.08 | 0.31 | 0.03 | 0.01 | 0.07 | 0.10 | 0.92 | 0.00 | 0.00 | 0.48 | 0.30 | 0.33 | 0.56 | 0.12 | 0.02 | 0.11 | 0.00 | 0.03 | 0.54 | 0.87 | 0.41 | 0.77 | 0.84 | 0.08 | 0.99 | 0.88 |
| MEdarkorange | 0.74 | 0.08 | 0.39 | 0.02 | 0.18 | 0.10 | 0.24 | 0.15 | 0.92 | 0.23 | 0.17 | 0.08 | 0.34 | 0.00 | 0.70 | 0.05 | 0.12 | 0.27 | 0.83 | 0.16 | 0.05 | 0.15 | 0.81 | 0.02 | 0.79 | 0.67 | 0.20 |
| MEdarkred | 0.86 | 0.32 | 0.98 | 0.52 | 0.24 | 0.84 | 0.71 | 0.28 | 0.85 | 0.45 | 0.79 | 0.60 | 0.60 | 0.47 | 0.15 | 0.39 | 0.28 | 0.27 | 0.21 | 0.83 | 0.15 | 0.23 | 0.62 | 0.42 | 0.49 | 0.55 | 0.61 |
| MEdarkturquoise | 0.13 | 0.55 | 0.33 | 0.57 | 0.12 | 0.92 | 0.26 | 0.09 | 0.98 | 0.27 | 0.32 | 0.14 | 0.11 | 0.32 | 0.06 | 0.71 | 0.54 | 0.81 | 0.22 | 0.04 | 0.18 | 0.26 | 0.06 | 0.48 | 0.98 | 0.59 | 0.21 |
| MEgreen | 0.00 | 0.22 | 0.00 | 0.22 | 0.00 | 0.07 | 0.01 | 0.41 | 0.03 | 0.00 | 0.00 | 0.09 | 0.00 | 0.11 | 0.03 | 0.27 | 0.32 | 0.34 | 0.16 | 0.01 | 0.66 | 0.79 | 0.55 | 0.89 | 0.65 | 0.21 | 0.33 |
| MEgreenyellow | 0.05 | 0.25 | 0.02 | 0.17 | 0.01 | 0.14 | 0.04 | 0.60 | 0.93 | 0.03 | 0.00 | 0.00 | 0.00 | 0.43 | 0.03 | 0.04 | 0.03 | 0.33 | 0.94 | 0.00 | 0.03 | 0.03 | 0.14 | 0.30 | 0.72 | 0.30 | 0.78 |
| MEgrey | 0.41 | 0.35 | 0.83 | 0.19 | 0.98 | 0.31 | 0.74 | 0.13 | 0.98 | 0.77 | 0.95 | 0.96 | 0.19 | 0.13 | 0.34 | 0.73 | 0.63 | 0.91 | 0.64 | 0.37 | 0.43 | 0.20 | 0.92 | 0.25 | 0.78 | 0.96 | 0.11 |
| MEgrey60 | 0.61 | 0.19 | 0.95 | 0.03 | 0.54 | 0.20 | 0.91 | 0.17 | 0.70 | 0.64 | 0.95 | 0.82 | 0.92 | 0.01 | 0.99 | 0.06 | 0.83 | 0.48 | 0.54 | 0.82 | 0.39 | 0.63 | 0.46 | 0.16 | 0.64 | 0.47 | 0.58 |
| MElightcyan | 0.28 | 0.04 | 0.08 | 0.00 | 0.07 | 0.03 | 0.22 | 0.34 | 0.39 | 0.02 | 0.03 | 0.10 | 0.03 | 0.06 | 0.59 | 0.02 | 0.13 | 0.99 | 0.16 | 0.12 | 0.30 | 0.46 | 0.57 | 0.14 | 0.65 | 0.65 | 0.04 |
| MElightgreen | 0.63 | 0.08 | 0.80 | 0.03 | 0.53 | 0.30 | 0.91 | 0.04 | 0.97 | 1.00 | 0.69 | 0.60 | 0.91 | 0.01 | 0.27 | 0.38 | 0.31 | 0.43 | 0.29 | 0.90 | 0.77 | 0.84 | 0.10 | 0.09 | 0.79 | 0.76 | 0.19 |
| MElightyellow | 0.30 | 0.87 | 0.25 | 0.66 | 0.01 | 0.24 | 0.14 | 0.04 | 0.93 | 0.06 | 0.22 | 0.04 | 0.18 | 0.74 | 0.02 | 0.06 | 0.29 | 0.41 | 0.16 | 0.01 | 0.01 | 0.01 | 0.02 | 0.91 | 0.44 | 0.32 | 0.42 |
| MEmagenta | 0.28 | 0.50 | 0.60 | 0.27 | 0.10 | 0.80 | 0.60 | 0.63 | 0.14 | 0.31 | 0.77 | 0.15 | 0.03 | 0.50 | 0.42 | 0.04 | 0.07 | 0.17 | 0.07 | 0.86 | 0.10 | 0.48 | 0.09 | 0.93 | 0.02 | 0.92 | 0.55 |
| MEmidnightblue | 0.13 | 0.87 | 0.90 | 0.80 | 0.38 | 0.34 | 0.78 | 0.62 | 0.15 | 0.77 | 0.98 | 0.04 | 0.00 | 0.29 | 0.46 | 0.36 | 0.02 | 0.17 | 0.06 | 0.67 | 0.63 | 0.94 | 0.01 | 0.12 | 0.06 | 0.50 | 0.90 |
| MEorange | 0.47 | 0.04 | 0.08 | 0.00 | 0.02 | 0.01 | 0.05 | 0.55 | 0.55 | 0.12 | 0.01 | 0.00 | 0.44 | 0.00 | 0.06 | 0.01 | 0.00 | 0.23 | 0.43 | 0.00 | 0.06 | 0.02 | 0.05 | 0.00 | 0.53 | 0.92 | 0.88 |
| MEpink | 0.53 | 0.44 | 0.45 | 0.12 | 0.02 | 0.40 | 0.65 | 0.89 | 0.18 | 0.25 | 0.72 | 0.21 | 0.44 | 0.14 | 0.30 | 0.01 | 0.11 | 0.04 | 0.21 | 0.67 | 0.44 | 0.94 | 0.57 | 0.90 | 0.08 | 0.60 | 0.41 |
| MEpurple | 0.93 | 0.40 | 0.47 | 0.39 | 0.97 | 0.28 | 0.48 | 0.99 | 0.32 | 0.89 | 0.32 | 0.01 | 0.33 | 0.17 | 0.97 | 0.91 | 0.00 | 0.22 | 0.47 | 0.16 | 0.28 | 0.86 | 0.09 | 0.15 | 0.04 | 0.92 | 0.92 |
| MEred | 0.51 | 0.22 | 0.40 | 0.09 | 0.10 | 0.09 | 0.10 | 0.66 | 0.55 | 0.27 | 0.17 | 0.00 | 0.42 | 0.01 | 0.05 | 0.04 | 0.03 | 0.67 | 0.25 | 0.00 | 0.17 | 0.02 | 0.15 | 0.07 | 0.08 | 0.54 | 0.26 |
| MEroyalblue | 0.04 | 0.49 | 0.08 | 0.24 | 0.00 | 0.35 | 0.11 | 0.16 | 0.37 | 0.08 | 0.09 | 0.39 | 0.01 | 0.32 | 0.00 | 0.00 | 0.87 | 0.26 | 0.82 | 0.02 | 0.05 | 0.03 | 0.51 | 0.93 | 0.49 | 0.83 | 0.28 |
| MEsalmon | 0.08 | 0.59 | 0.31 | 0.43 | 0.06 | 0.74 | 0.12 | 0.44 | 0.74 | 0.20 | 0.14 | 0.19 | 0.00 | 0.53 | 0.13 | 0.06 | 0.49 | 0.76 | 0.63 | 0.03 | 0.04 | 0.05 | 0.86 | 0.83 | 0.68 | 0.46 | 0.91 |
| MEtan | 0.23 | 0.23 | 0.88 | 0.07 | 0.79 | 0.11 | 0.82 | 0.14 | 0.29 | 0.82 | 0.73 | 0.04 | 0.35 | 0.00 | 0.57 | 0.39 | 0.02 | 0.22 | 0.34 | 0.37 | 0.29 | 0.28 | 0.42 | 0.01 | 0.21 | 0.49 | 0.59 |
| MEturquoise | 0.09 | 0.86 | 0.68 | 0.98 | 0.05 | 0.66 | 0.37 | 0.39 | 0.24 | 0.38 | 0.75 | 0.22 | 0.01 | 0.62 | 0.07 | 0.08 | 0.04 | 0.17 | 0.33 | 0.60 | 0.06 | 0.20 | 0.29 | 0.34 | 0.06 | 0.72 | 0.60 |
| MEwhite | 0.04 | 0.71 | 0.25 | 0.63 | 0.02 | 0.98 | 0.33 | 0.42 | 0.12 | 0.32 | 0.36 | 0.23 | 0.02 | 0.91 | 0.02 | 0.06 | 0.05 | 0.16 | 0.23 | 0.62 | 0.55 | 0.63 | 0.42 | 0.36 | 0.04 | 0.55 | 0.30 |
| MEyellow | 0.79 | 0.08 | 0.29 | 0.00 | 0.01 | 0.02 | 0.18 | 0.64 | 0.95 | 0.16 | 0.20 | 0.07 | 0.62 | 0.00 | 0.09 | 0.00 | 0.22 | 0.75 | 0.75 | 0.06 | 0.05 | 0.04 | 0.29 | 0.02 | 0.76 | 0.34 | 0.95 |

**Supplementary File 9D. Protein levels with gene module correlations, 24hr infected**

**Infected, 24hr protein vs. module correlation**

| | GT2 | CCL13 | CCL17 | CCL19 | CCL2 | CCL20 | CCL22 | CCL3 | CCL4 | CSF2 | CXCL10 | CXCL5 | IL10 | IL15 | IL16 | IL18 | IL1A | IL1B | IL6 | IL7 | IL8 | MMP1 | MMP10 | MMP2 | MMP3 | MMP9 | TNF | VEGFA |
|---|---|---|---|---|---|---|---|---|---|---|---|---|---|---|---|---|---|---|---|---|---|---|---|---|---|---|---|---|
| MEblack | -0.50 | -0.29 | 0.10 | -0.08 | 0.07 | -0.23 | 0.09 | 0.63 | 0.22 | -0.11 | 0.01 | 0.27 | -0.66 | -0.21 | 0.19 | -0.16 | -0.07 | -0.34 | 0.04 | -0.19 | 0.53 | -0.02 | -0.04 | 0.28 | 0.31 | -0.37 | 0.10 | -0.26 |
| MEblue | -0.52 | -0.03 | 0.28 | 0.26 | 0.43 | 0.26 | 0.25 | 0.77 | 0.35 | 0.10 | 0.16 | 0.23 | -0.41 | 0.20 | 0.48 | 0.35 | 0.26 | -0.06 | 0.11 | -0.11 | 0.57 | 0.37 | 0.11 | 0.21 | 0.29 | -0.14 | 0.14 | -0.10 |
| MEbrown | 0.01 | -0.25 | -0.16 | -0.32 | -0.14 | -0.45 | -0.31 | -0.16 | -0.12 | 0.04 | -0.40 | -0.41 | 0.51 | -0.20 | -0.13 | -0.26 | -0.23 | 0.26 | -0.02 | 0.16 | -0.51 | -0.28 | -0.08 | -0.47 | -0.37 | 0.03 | -0.12 | 0.05 |
| MEcyan | 0.17 | 0.03 | 0.25 | 0.15 | 0.28 | -0.08 | 0.00 | 0.20 | -0.33 | 0.07 | -0.16 | -0.01 | -0.02 | 0.19 | 0.26 | -0.13 | -0.04 | -0.22 | -0.34 | 0.19 | -0.04 | -0.26 | 0.34 | -0.48 | 0.28 | 0.42 | -0.30 | 0.19 |
| MEdarkgreen | -0.25 | 0.09 | 0.11 | 0.10 | 0.00 | -0.13 | 0.26 | 0.41 | 0.20 | -0.04 | 0.23 | 0.41 | -0.69 | 0.09 | -0.22 | -0.01 | 0.07 | -0.37 | -0.06 | -0.04 | 0.62 | -0.04 | -0.05 | 0.22 | 0.21 | -0.58 | -0.06 | -0.06 |
| MEdarkgrey | -0.06 | 0.39 | 0.36 | 0.40 | 0.47 | 0.37 | 0.32 | 0.07 | 0.03 | 0.41 | 0.41 | 0.00 | 0.36 | 0.48 | 0.23 | 0.49 | 0.56 | 0.56 | 0.31 | 0.06 | 0.04 | 0.13 | -0.02 | -0.21 | 0.04 | 0.18 | 0.28 | -0.07 |
| MEdarkorange | -0.28 | -0.09 | 0.16 | 0.14 | 0.25 | -0.03 | 0.07 | 0.73 | 0.23 | -0.15 | 0.04 | 0.23 | -0.64 | 0.15 | 0.27 | 0.03 | -0.05 | -0.54 | -0.35 | 0.04 | 0.52 | 0.07 | 0.17 | 0.15 | 0.30 | -0.09 | -0.28 | -0.05 |
| MEdarkred | -0.21 | -0.07 | -0.17 | -0.05 | -0.05 | 0.21 | -0.02 | -0.04 | 0.31 | -0.02 | 0.12 | -0.20 | 0.25 | -0.01 | 0.00 | 0.25 | 0.06 | 0.28 | 0.19 | -0.09 | -0.09 | 0.37 | -0.24 | 0.20 | -0.28 | -0.13 | 0.14 | 0.01 |
| MEdarkturquoise | 0.00 | -0.34 | 0.04 | -0.19 | -0.07 | -0.44 | -0.09 | -0.14 | -0.35 | 0.07 | -0.39 | -0.26 | 0.31 | -0.26 | 0.04 | -0.38 | -0.17 | 0.18 | 0.09 | 0.00 | -0.37 | -0.47 | 0.00 | -0.50 | -0.01 | 0.13 | 0.07 | -0.05 |
| MEgreen | -0.22 | -0.74 | -0.39 | -0.65 | -0.44 | -0.66 | -0.47 | 0.03 | -0.03 | -0.34 | -0.61 | -0.49 | 0.02 | -0.77 | 0.01 | -0.69 | -0.67 | -0.27 | -0.16 | -0.13 | -0.36 | -0.37 | -0.26 | 0.06 | -0.17 | -0.09 | -0.15 | -0.25 |
| MEgreenyellow | 0.21 | -0.26 | -0.30 | -0.38 | -0.37 | -0.36 | -0.33 | -0.54 | -0.22 | 0.02 | -0.35 | -0.56 | 0.67 | -0.45 | -0.22 | -0.34 | -0.22 | 0.45 | 0.21 | -0.10 | -0.68 | -0.36 | -0.22 | -0.31 | -0.35 | 0.03 | 0.14 | -0.10 |
| MEgrey | 0.20 | 0.45 | 0.34 | 0.48 | 0.41 | 0.43 | 0.34 | -0.10 | -0.06 | 0.37 | 0.38 | 0.07 | 0.39 | 0.54 | 0.15 | 0.47 | 0.58 | 0.63 | 0.42 | 0.03 | 0.02 | 0.09 | 0.16 | -0.25 | 0.08 | 0.20 | 0.42 | 0.06 |
| MEgrey60 | 0.45 | 0.21 | -0.09 | 0.08 | -0.22 | 0.13 | 0.02 | -0.38 | -0.13 | -0.16 | 0.06 | 0.10 | -0.15 | 0.06 | -0.29 | -0.02 | -0.07 | -0.30 | -0.19 | 0.13 | 0.00 | -0.13 | -0.05 | 0.27 | 0.02 | 0.04 | -0.17 | 0.04 |
| MElightcyan | 0.24 | -0.13 | -0.42 | -0.47 | -0.54 | -0.21 | -0.31 | -0.71 | -0.20 | -0.15 | -0.23 | -0.26 | 0.27 | -0.50 | -0.38 | -0.35 | -0.38 | -0.04 | -0.06 | -0.05 | -0.49 | -0.13 | -0.32 | 0.20 | -0.30 | -0.05 | -0.10 | -0.10 |
| MElightgreen | 0.28 | 0.19 | -0.24 | -0.02 | -0.26 | 0.30 | -0.03 | -0.56 | 0.08 | -0.04 | 0.09 | -0.07 | 0.27 | -0.01 | -0.30 | 0.22 | 0.05 | 0.19 | 0.12 | 0.01 | -0.22 | 0.22 | -0.23 | 0.33 | -0.31 | -0.02 | 0.09 | -0.02 |
| MElightyellow | -0.43 | 0.13 | 0.01 | 0.16 | 0.15 | 0.30 | 0.18 | 0.47 | 0.61 | 0.07 | 0.32 | 0.21 | -0.30 | 0.17 | 0.08 | 0.42 | 0.24 | 0.08 | 0.24 | -0.22 | 0.52 | 0.54 | -0.13 | 0.50 | -0.08 | -0.41 | 0.24 | -0.28 |
| MEmagenta | -0.11 | 0.09 | 0.14 | 0.24 | 0.31 | 0.26 | 0.04 | 0.23 | 0.13 | 0.17 | 0.08 | -0.16 | 0.30 | 0.30 | 0.28 | 0.30 | 0.20 | 0.29 | 0.06 | 0.03 | -0.07 | 0.21 | 0.18 | -0.27 | -0.08 | 0.29 | 0.02 | 0.18 |
| MEmidnightblue | -0.19 | -0.16 | -0.22 | -0.32 | -0.34 | -0.19 | -0.06 | -0.04 | 0.12 | -0.20 | 0.01 | 0.26 | -0.49 | -0.40 | -0.25 | -0.26 | -0.25 | -0.39 | -0.02 | -0.17 | 0.25 | 0.05 | -0.29 | 0.54 | 0.04 | -0.41 | 0.02 | -0.26 |
| MEorange | 0.42 | 0.05 | -0.22 | -0.15 | -0.28 | -0.14 | -0.23 | -0.64 | -0.22 | 0.12 | -0.18 | -0.49 | 0.78 | -0.11 | -0.34 | -0.12 | -0.05 | 0.49 | 0.13 | 0.07 | -0.69 | -0.30 | -0.09 | -0.42 | -0.44 | 0.19 | 0.04 | 0.05 |
| MEpink | 0.09 | -0.06 | -0.08 | -0.24 | -0.22 | -0.42 | -0.11 | 0.05 | -0.17 | -0.07 | -0.26 | 0.19 | -0.46 | -0.26 | -0.27 | -0.43 | -0.32 | -0.60 | -0.39 | 0.08 | 0.15 | -0.33 | 0.01 | 0.04 | 0.12 | -0.25 | -0.35 | -0.21 |
| MEpurple | 0.10 | 0.08 | -0.14 | -0.04 | -0.03 | 0.13 | -0.12 | -0.27 | 0.07 | 0.12 | -0.03 | -0.34 | 0.58 | 0.09 | -0.08 | 0.17 | 0.04 | 0.37 | 0.03 | 0.07 | -0.40 | 0.18 | -0.06 | -0.21 | -0.39 | 0.13 | -0.04 | 0.10 |
| MEred | -0.55 | -0.24 | 0.14 | -0.04 | 0.18 | -0.16 | 0.09 | 0.80 | 0.27 | -0.06 | 0.02 | 0.27 | -0.72 | -0.14 | 0.30 | -0.08 | -0.06 | -0.44 | -0.09 | -0.16 | 0.62 | 0.08 | 0.01 | 0.29 | 0.36 | -0.35 | -0.02 | -0.27 |
| MEroyalblue | 0.36 | -0.11 | -0.17 | -0.34 | -0.32 | -0.52 | -0.23 | -0.42 | -0.34 | -0.04 | -0.36 | -0.26 | 0.27 | -0.28 | -0.36 | -0.48 | -0.29 | -0.01 | -0.11 | 0.11 | -0.44 | -0.55 | -0.13 | -0.36 | -0.21 | -0.03 | -0.13 | -0.14 |
| MEsalmon | -0.22 | 0.05 | 0.20 | 0.28 | 0.31 | 0.07 | 0.10 | 0.75 | 0.29 | -0.06 | 0.10 | 0.23 | -0.55 | 0.30 | 0.24 | 0.14 | 0.07 | -0.40 | -0.26 | 0.04 | 0.54 | 0.09 | 0.24 | 0.05 | 0.23 | -0.01 | -0.22 | -0.01 |
| MEtan | 0.52 | 0.36 | -0.01 | 0.24 | -0.02 | 0.41 | 0.08 | -0.63 | -0.19 | 0.10 | 0.17 | -0.10 | 0.52 | 0.31 | -0.16 | 0.29 | 0.25 | 0.40 | 0.17 | 0.15 | -0.35 | 0.06 | 0.04 | -0.12 | -0.16 | 0.34 | 0.13 | 0.32 |
| MEturquoise | -0.04 | 0.18 | 0.06 | 0.18 | 0.24 | 0.32 | 0.02 | 0.14 | 0.18 | 0.16 | 0.15 | -0.11 | 0.26 | 0.32 | 0.16 | 0.34 | 0.19 | 0.22 | -0.01 | 0.06 | -0.04 | 0.32 | 0.09 | -0.13 | -0.15 | 0.19 | -0.04 | 0.13 |
| MEwhite | -0.15 | -0.32 | -0.11 | -0.28 | -0.21 | -0.54 | -0.18 | 0.31 | 0.08 | -0.14 | -0.29 | -0.09 | -0.31 | -0.37 | -0.15 | -0.39 | -0.27 | -0.30 | -0.11 | -0.03 | 0.10 | -0.43 | -0.11 | 0.01 | -0.04 | -0.32 | -0.12 | -0.35 |
| MEyellow | -0.61 | -0.20 | 0.22 | 0.11 | 0.34 | 0.10 | 0.18 | 0.76 | 0.33 | 0.01 | 0.08 | 0.19 | -0.45 | 0.01 | 0.49 | 0.20 | 0.14 | -0.09 | 0.14 | -0.17 | 0.53 | 0.29 | 0.05 | 0.27 | 0.31 | -0.22 | 0.17 | -0.15 |

**Infected, 24hr protein vs. module correlation p value**

| | GT2 | CCL13 | CCL17 | CCL19 | CCL2 | CCL20 | CCL22 | CCL3 | CCL4 | CSF2 | CXCL10 | CXCL5 | IL10 | IL15 | IL16 | IL18 | IL1A | IL1B | IL6 | IL7 | IL8 | MMP1 | MMP10 | MMP2 | MMP3 | MMP9 | TNF | VEGFA |
|---|---|---|---|---|---|---|---|---|---|---|---|---|---|---|---|---|---|---|---|---|---|---|---|---|---|---|---|---|
| MEblack | 0.01 | 0.15 | 0.64 | 0.69 | 0.75 | 0.25 | 0.65 | 0.00 | 0.29 | 0.59 | 0.97 | 0.18 | 0.00 | 0.31 | 0.36 | 0.43 | 0.73 | 0.09 | 0.84 | 0.34 | 0.00 | 0.91 | 0.84 | 0.17 | 0.13 | 0.06 | 0.64 | 0.19 |
| MEblue | 0.01 | 0.87 | 0.16 | 0.20 | 0.03 | 0.21 | 0.22 | 0.00 | 0.08 | 0.64 | 0.44 | 0.26 | 0.04 | 0.32 | 0.01 | 0.08 | 0.20 | 0.78 | 0.58 | 0.60 | 0.00 | 0.07 | 0.60 | 0.30 | 0.16 | 0.49 | 0.48 | 0.62 |
| MEbrown | 0.97 | 0.23 | 0.43 | 0.11 | 0.48 | 0.02 | 0.12 | 0.45 | 0.56 | 0.83 | 0.04 | 0.04 | 0.01 | 0.33 | 0.52 | 0.20 | 0.27 | 0.19 | 0.91 | 0.44 | 0.01 | 0.16 | 0.70 | 0.02 | 0.06 | 0.88 | 0.58 | 0.80 |
| MEcyan | 0.41 | 0.87 | 0.22 | 0.48 | 0.17 | 0.70 | 1.00 | 0.34 | 0.10 | 0.72 | 0.42 | 0.98 | 0.93 | 0.36 | 0.21 | 0.51 | 0.85 | 0.27 | 0.09 | 0.35 | 0.84 | 0.20 | 0.09 | 0.01 | 0.17 | 0.03 | 0.14 | 0.34 |
| MEdarkgreen | 0.22 | 0.66 | 0.59 | 0.62 | 0.99 | 0.53 | 0.20 | 0.04 | 0.33 | 0.85 | 0.25 | 0.04 | 0.00 | 0.67 | 0.28 | 0.94 | 0.74 | 0.06 | 0.76 | 0.85 | 0.00 | 0.86 | 0.82 | 0.29 | 0.30 | 0.00 | 0.75 | 0.76 |
| MEdarkgrey | 0.78 | 0.05 | 0.07 | 0.04 | 0.02 | 0.06 | 0.12 | 0.73 | 0.87 | 0.04 | 0.04 | 1.00 | 0.07 | 0.01 | 0.25 | 0.01 | 0.00 | 0.00 | 0.13 | 0.77 | 0.84 | 0.54 | 0.94 | 0.31 | 0.86 | 0.38 | 0.17 | 0.73 |
| MEdarkorange | 0.16 | 0.67 | 0.42 | 0.49 | 0.22 | 0.88 | 0.73 | 0.00 | 0.26 | 0.47 | 0.86 | 0.27 | 0.00 | 0.47 | 0.18 | 0.89 | 0.79 | 0.00 | 0.08 | 0.85 | 0.01 | 0.74 | 0.40 | 0.47 | 0.14 | 0.67 | 0.17 | 0.80 |
| MEdarkred | 0.31 | 0.75 | 0.42 | 0.81 | 0.80 | 0.30 | 0.91 | 0.83 | 0.12 | 0.92 | 0.55 | 0.33 | 0.22 | 0.96 | 0.99 | 0.22 | 0.78 | 0.16 | 0.35 | 0.65 | 0.67 | 0.06 | 0.25 | 0.32 | 0.17 | 0.52 | 0.48 | 0.95 |
| MEdarkturquoise | 0.99 | 0.09 | 0.84 | 0.35 | 0.75 | 0.02 | 0.65 | 0.51 | 0.08 | 0.74 | 0.05 | 0.20 | 0.12 | 0.20 | 0.84 | 0.06 | 0.42 | 0.38 | 0.66 | 0.99 | 0.07 | 0.02 | 0.99 | 0.01 | 0.95 | 0.52 | 0.73 | 0.79 |
| MEgreen | 0.28 | 0.00 | 0.05 | 0.00 | 0.02 | 0.00 | 0.02 | 0.88 | 0.88 | 0.09 | 0.00 | 0.01 | 0.90 | 0.00 | 0.95 | 0.00 | 0.00 | 0.18 | 0.44 | 0.52 | 0.07 | 0.06 | 0.21 | 0.77 | 0.39 | 0.67 | 0.47 | 0.22 |
| MEgreenyellow | 0.30 | 0.20 | 0.13 | 0.06 | 0.06 | 0.07 | 0.10 | 0.00 | 0.27 | 0.93 | 0.08 | 0.00 | 0.00 | 0.02 | 0.28 | 0.09 | 0.29 | 0.02 | 0.31 | 0.63 | 0.00 | 0.07 | 0.28 | 0.12 | 0.08 | 0.90 | 0.49 | 0.64 |
| MEgrey | 0.33 | 0.02 | 0.09 | 0.01 | 0.04 | 0.03 | 0.09 | 0.63 | 0.77 | 0.06 | 0.05 | 0.75 | 0.05 | 0.00 | 0.46 | 0.01 | 0.00 | 0.00 | 0.03 | 0.90 | 0.93 | 0.67 | 0.45 | 0.22 | 0.70 | 0.32 | 0.03 | 0.78 |
| MEgrey60 | 0.02 | 0.31 | 0.65 | 0.69 | 0.28 | 0.54 | 0.91 | 0.05 | 0.52 | 0.43 | 0.79 | 0.62 | 0.47 | 0.77 | 0.15 | 0.93 | 0.74 | 0.14 | 0.35 | 0.53 | 0.98 | 0.53 | 0.83 | 0.19 | 0.91 | 0.86 | 0.41 | 0.85 |
| MElightcyan | 0.24 | 0.52 | 0.03 | 0.02 | 0.00 | 0.29 | 0.12 | 0.00 | 0.34 | 0.46 | 0.25 | 0.21 | 0.19 | 0.01 | 0.06 | 0.08 | 0.06 | 0.83 | 0.75 | 0.83 | 0.01 | 0.51 | 0.11 | 0.32 | 0.13 | 0.81 | 0.64 | 0.63 |
| MElightgreen | 0.17 | 0.35 | 0.24 | 0.92 | 0.19 | 0.14 | 0.88 | 0.00 | 0.69 | 0.86 | 0.66 | 0.72 | 0.18 | 0.97 | 0.14 | 0.28 | 0.79 | 0.35 | 0.55 | 0.95 | 0.28 | 0.27 | 0.26 | 0.10 | 0.12 | 0.91 | 0.67 | 0.92 |
| MElightyellow | 0.03 | 0.54 | 0.97 | 0.42 | 0.46 | 0.13 | 0.39 | 0.02 | 0.00 | 0.72 | 0.12 | 0.30 | 0.13 | 0.42 | 0.71 | 0.03 | 0.23 | 0.69 | 0.25 | 0.28 | 0.01 | 0.00 | 0.54 | 0.01 | 0.69 | 0.04 | 0.23 | 0.16 |
| MEmagenta | 0.59 | 0.67 | 0.51 | 0.24 | 0.13 | 0.21 | 0.86 | 0.27 | 0.53 | 0.42 | 0.71 | 0.44 | 0.13 | 0.13 | 0.16 | 0.14 | 0.32 | 0.14 | 0.78 | 0.87 | 0.74 | 0.29 | 0.38 | 0.18 | 0.68 | 0.15 | 0.90 | 0.39 |
| MEmidnightblue | 0.35 | 0.44 | 0.28 | 0.11 | 0.09 | 0.35 | 0.76 | 0.84 | 0.57 | 0.33 | 0.96 | 0.20 | 0.01 | 0.04 | 0.23 | 0.20 | 0.22 | 0.05 | 0.93 | 0.41 | 0.22 | 0.82 | 0.16 | 0.00 | 0.84 | 0.04 | 0.93 | 0.20 |
| MEorange | 0.03 | 0.83 | 0.27 | 0.46 | 0.17 | 0.49 | 0.26 | 0.00 | 0.27 | 0.57 | 0.38 | 0.01 | 0.00 | 0.61 | 0.09 | 0.55 | 0.80 | 0.01 | 0.52 | 0.73 | 0.00 | 0.13 | 0.67 | 0.03 | 0.02 | 0.36 | 0.83 | 0.80 |
| MEpink | 0.67 | 0.77 | 0.71 | 0.24 | 0.27 | 0.03 | 0.59 | 0.79 | 0.40 | 0.72 | 0.21 | 0.35 | 0.02 | 0.20 | 0.19 | 0.03 | 0.11 | 0.00 | 0.05 | 0.71 | 0.46 | 0.10 | 0.95 | 0.86 | 0.57 | 0.22 | 0.08 | 0.30 |
| MEpurple | 0.64 | 0.71 | 0.50 | 0.85 | 0.87 | 0.52 | 0.55 | 0.19 | 0.75 | 0.56 | 0.90 | 0.09 | 0.00 | 0.68 | 0.69 | 0.39 | 0.86 | 0.06 | 0.89 | 0.75 | 0.04 | 0.37 | 0.76 | 0.30 | 0.05 | 0.52 | 0.85 | 0.64 |
| MEred | 0.00 | 0.24 | 0.49 | 0.85 | 0.38 | 0.44 | 0.65 | 0.00 | 0.18 | 0.75 | 0.94 | 0.17 | 0.00 | 0.49 | 0.14 | 0.69 | 0.77 | 0.03 | 0.67 | 0.43 | 0.00 | 0.71 | 0.95 | 0.15 | 0.07 | 0.08 | 0.91 | 0.18 |
| MEroyalblue | 0.07 | 0.58 | 0.41 | 0.09 | 0.11 | 0.01 | 0.26 | 0.03 | 0.09 | 0.84 | 0.07 | 0.20 | 0.18 | 0.17 | 0.07 | 0.01 | 0.15 | 0.95 | 0.58 | 0.60 | 0.02 | 0.00 | 0.52 | 0.07 | 0.30 | 0.87 | 0.53 | 0.50 |
| MEsalmon | 0.29 | 0.81 | 0.32 | 0.16 | 0.12 | 0.74 | 0.62 | 0.00 | 0.15 | 0.77 | 0.64 | 0.27 | 0.00 | 0.13 | 0.25 | 0.50 | 0.73 | 0.04 | 0.19 | 0.85 | 0.00 | 0.66 | 0.25 | 0.82 | 0.26 | 0.95 | 0.29 | 0.96 |
| MEtan | 0.01 | 0.07 | 0.98 | 0.23 | 0.91 | 0.04 | 0.68 | 0.00 | 0.35 | 0.63 | 0.39 | 0.62 | 0.01 | 0.13 | 0.43 | 0.14 | 0.21 | 0.04 | 0.41 | 0.46 | 0.08 | 0.77 | 0.84 | 0.55 | 0.44 | 0.09 | 0.54 | 0.11 |
| MEturquoise | 0.83 | 0.38 | 0.76 | 0.38 | 0.23 | 0.11 | 0.91 | 0.49 | 0.38 | 0.44 | 0.48 | 0.58 | 0.19 | 0.12 | 0.44 | 0.09 | 0.35 | 0.29 | 0.95 | 0.76 | 0.84 | 0.11 | 0.65 | 0.53 | 0.48 | 0.34 | 0.85 | 0.53 |
| MEwhite | 0.48 | 0.11 | 0.59 | 0.16 | 0.29 | 0.00 | 0.39 | 0.13 | 0.69 | 0.48 | 0.16 | 0.67 | 0.12 | 0.06 | 0.46 | 0.05 | 0.19 | 0.13 | 0.60 | 0.87 | 0.64 | 0.03 | 0.61 | 0.98 | 0.85 | 0.12 | 0.55 | 0.08 |
| MEyellow | 0.00 | 0.33 | 0.28 | 0.58 | 0.09 | 0.63 | 0.38 | 0.00 | 0.10 | 0.96 | 0.70 | 0.35 | 0.02 | 0.96 | 0.01 | 0.32 | 0.48 | 0.67 | 0.51 | 0.42 | 0.01 | 0.15 | 0.82 | 0.19 | 0.13 | 0.28 | 0.41 | 0.47 |

**Supplementary File 9E. Protein levels with gene module correlations, 72hr control**

**Control, 72hr protein vs. module correlation**

| | CCL13 | CCL17 | CCL19 | CCL2 | CCL20 | CCL22 | CCL3 | CCL4 | CSF2 | CXCL10 | CXCL5 | IL10 | IL15 | IL16 | IL18 | IL1A | IL1B | IL6 | IL7 | IL8 | MMP1 | MMP10 | MMP2 | MMP3 | MMP9 | TNF | VEGFA |
|---|---|---|---|---|---|---|---|---|---|---|---|---|---|---|---|---|---|---|---|---|---|---|---|---|---|---|---|
| MEblack | 0.09 | 0.50 | 0.29 | 0.51 | 0.12 | 0.52 | 0.17 | -0.13 | 0.02 | 0.35 | 0.23 | -0.40 | -0.02 | 0.31 | 0.24 | 0.24 | -0.37 | -0.10 | -0.07 | 0.34 | -0.26 | 0.09 | 0.13 | 0.50 | 0.05 | 0.01 | 0.01 |
| MEblue | -0.38 | 0.23 | -0.24 | 0.33 | -0.45 | 0.00 | -0.02 | -0.10 | -0.08 | -0.15 | -0.22 | 0.11 | -0.48 | 0.37 | -0.38 | 0.11 | 0.01 | -0.27 | -0.19 | -0.31 | 0.00 | -0.27 | -0.01 | 0.20 | -0.32 | 0.09 | -0.25 |
| MEbrown | 0.52 | 0.46 | 0.41 | 0.39 | 0.67 | 0.50 | 0.34 | -0.01 | 0.20 | 0.34 | 0.34 | -0.15 | 0.08 | 0.24 | 0.63 | 0.29 | 0.02 | 0.40 | -0.16 | 0.40 | 0.07 | 0.38 | 0.19 | 0.33 | 0.18 | 0.08 | 0.09 |
| MEcyan | -0.22 | 0.01 | -0.01 | 0.09 | -0.32 | -0.06 | -0.30 | -0.30 | -0.24 | -0.02 | 0.14 | -0.32 | 0.15 | -0.04 | -0.22 | -0.19 | -0.43 | -0.41 | 0.21 | 0.04 | -0.30 | -0.23 | -0.09 | 0.15 | -0.38 | -0.29 | 0.18 |
| MEdarkgreen | 0.09 | -0.15 | -0.08 | -0.33 | 0.18 | -0.05 | 0.20 | 0.13 | -0.07 | -0.16 | -0.19 | 0.31 | 0.07 | -0.01 | 0.13 | 0.22 | 0.44 | 0.26 | 0.05 | -0.05 | 0.22 | 0.36 | -0.10 | -0.17 | 0.22 | 0.29 | 0.04 |
| MEdarkgrey | -0.11 | 0.27 | -0.25 | 0.32 | -0.11 | -0.03 | 0.22 | -0.01 | 0.11 | -0.24 | -0.23 | 0.43 | -0.60 | 0.41 | -0.17 | 0.20 | 0.43 | 0.16 | -0.36 | -0.37 | 0.32 | -0.09 | 0.02 | 0.06 | -0.32 | 0.23 | -0.29 |
| MEdarkolivegreen | -0.16 | -0.03 | -0.12 | 0.07 | -0.31 | -0.17 | -0.21 | -0.16 | -0.06 | -0.17 | 0.06 | 0.03 | -0.23 | 0.03 | -0.34 | -0.12 | -0.07 | -0.21 | -0.19 | -0.18 | -0.13 | -0.20 | -0.17 | 0.06 | -0.27 | -0.04 | -0.07 |
| MEdarkorange | -0.29 | -0.59 | -0.33 | -0.65 | -0.33 | -0.58 | -0.20 | 0.12 | -0.01 | -0.31 | -0.38 | 0.40 | 0.09 | -0.45 | -0.36 | -0.32 | 0.32 | 0.00 | 0.18 | -0.39 | 0.19 | -0.19 | -0.21 | -0.61 | 0.00 | -0.04 | -0.08 |
| MEdarkred | 0.48 | 0.40 | 0.57 | 0.17 | 0.39 | 0.51 | 0.27 | -0.03 | 0.46 | 0.74 | 0.09 | 0.09 | 0.26 | -0.28 | 0.36 | -0.10 | 0.01 | 0.46 | -0.48 | 0.24 | 0.02 | 0.00 | -0.46 | -0.07 | 0.41 | -0.11 | -0.01 |
| MEdarkturquoise | -0.30 | 0.00 | -0.18 | 0.04 | -0.45 | -0.27 | -0.28 | -0.30 | -0.07 | -0.24 | -0.07 | 0.13 | -0.17 | 0.02 | -0.43 | -0.22 | -0.02 | -0.25 | -0.10 | -0.33 | 0.04 | -0.36 | -0.22 | -0.04 | -0.51 | -0.14 | -0.07 |
| MEgreen | -0.39 | -0.47 | -0.37 | -0.35 | -0.42 | -0.52 | -0.43 | -0.08 | -0.34 | -0.43 | -0.08 | -0.09 | 0.15 | -0.14 | -0.44 | -0.25 | -0.15 | -0.42 | 0.39 | -0.22 | 0.00 | -0.22 | 0.03 | -0.22 | -0.33 | -0.22 | 0.21 |
| MEgreenyellow | 0.07 | -0.14 | -0.15 | -0.12 | 0.11 | -0.21 | -0.08 | -0.06 | -0.11 | -0.25 | 0.01 | 0.17 | 0.05 | 0.06 | -0.01 | -0.02 | 0.22 | 0.08 | 0.09 | -0.13 | 0.22 | 0.07 | -0.03 | -0.08 | -0.18 | 0.02 | 0.15 |
| MEgrey | -0.03 | -0.35 | 0.04 | -0.34 | -0.10 | -0.24 | -0.38 | -0.11 | -0.20 | 0.00 | 0.15 | -0.15 | 0.43 | -0.45 | -0.11 | -0.37 | -0.27 | -0.20 | 0.27 | 0.05 | -0.27 | -0.09 | -0.23 | -0.24 | -0.13 | -0.36 | 0.32 |
| MEgrey60 | 0.03 | -0.21 | 0.16 | -0.21 | -0.04 | -0.02 | -0.17 | -0.12 | -0.01 | 0.04 | 0.25 | -0.42 | 0.35 | -0.30 | 0.05 | -0.22 | -0.42 | -0.18 | 0.23 | 0.31 | -0.36 | -0.05 | 0.08 | -0.01 | 0.14 | -0.11 | 0.11 |
| MElightcyan | -0.32 | -0.03 | -0.39 | -0.05 | -0.32 | -0.32 | -0.05 | -0.06 | -0.10 | -0.36 | -0.38 | 0.49 | -0.30 | 0.23 | -0.38 | 0.05 | 0.47 | -0.04 | -0.08 | -0.53 | 0.50 | -0.13 | -0.17 | -0.13 | -0.31 | 0.13 | -0.13 |
| MElightgreen | 0.20 | 0.43 | -0.01 | 0.21 | 0.16 | 0.31 | 0.62 | -0.10 | 0.36 | -0.09 | -0.25 | 0.40 | -0.49 | 0.31 | 0.19 | 0.31 | 0.56 | 0.53 | -0.42 | -0.07 | 0.42 | 0.12 | -0.03 | 0.21 | 0.12 | 0.63 | -0.28 |
| MElightyellow | -0.07 | -0.57 | -0.03 | -0.59 | -0.11 | -0.40 | -0.32 | 0.10 | -0.01 | 0.00 | -0.10 | 0.10 | 0.39 | -0.69 | -0.11 | -0.48 | -0.03 | -0.01 | 0.25 | -0.07 | -0.25 | -0.16 | -0.22 | -0.60 | 0.05 | -0.35 | 0.11 |
| MEmagenta | 0.40 | -0.07 | 0.30 | -0.12 | 0.38 | 0.06 | -0.01 | -0.06 | 0.18 | 0.17 | 0.30 | -0.15 | 0.33 | -0.31 | 0.33 | -0.21 | -0.11 | 0.23 | 0.01 | 0.29 | -0.10 | 0.08 | -0.01 | -0.06 | 0.17 | -0.11 | 0.17 |
| MEmidnightblue | 0.09 | 0.11 | 0.32 | 0.14 | -0.03 | 0.28 | -0.03 | 0.00 | -0.05 | 0.34 | 0.37 | -0.59 | 0.15 | -0.23 | 0.06 | -0.11 | -0.66 | -0.21 | -0.04 | 0.44 | -0.55 | -0.04 | 0.06 | 0.21 | 0.09 | -0.29 | 0.16 |
| MEorange | 0.35 | 0.19 | 0.18 | 0.00 | 0.39 | 0.12 | 0.26 | 0.01 | 0.23 | 0.16 | 0.07 | 0.28 | 0.06 | 0.01 | 0.27 | 0.18 | 0.40 | 0.43 | -0.19 | 0.02 | 0.46 | 0.32 | -0.19 | 0.00 | 0.18 | 0.18 | 0.01 |
| MEpaleturquoise | -0.35 | 0.00 | -0.14 | 0.30 | -0.40 | -0.09 | -0.36 | -0.09 | -0.21 | -0.13 | 0.10 | -0.30 | -0.19 | 0.17 | -0.31 | -0.15 | -0.47 | -0.52 | 0.06 | -0.07 | -0.47 | -0.38 | 0.22 | 0.12 | -0.44 | -0.31 | -0.09 |
| MEpink | 0.34 | 0.60 | 0.22 | 0.34 | 0.35 | 0.51 | 0.64 | -0.08 | 0.34 | 0.22 | -0.05 | 0.19 | -0.27 | 0.35 | 0.36 | 0.43 | 0.39 | 0.50 | -0.39 | 0.14 | 0.44 | 0.32 | -0.05 | 0.39 | 0.32 | 0.55 | -0.19 |
| MEpurple | -0.08 | -0.27 | -0.04 | -0.32 | 0.03 | -0.13 | -0.04 | 0.20 | 0.04 | 0.03 | -0.27 | 0.17 | 0.20 | -0.21 | 0.07 | -0.08 | 0.19 | 0.07 | 0.20 | -0.08 | 0.14 | 0.05 | 0.03 | -0.26 | 0.33 | 0.05 | -0.05 |
| MEred | -0.19 | 0.04 | 0.01 | 0.14 | -0.33 | 0.01 | -0.22 | -0.17 | -0.09 | 0.00 | 0.16 | -0.35 | -0.04 | 0.00 | -0.24 | -0.14 | -0.46 | -0.38 | 0.02 | 0.07 | -0.34 | -0.27 | 0.04 | 0.19 | -0.17 | -0.14 | -0.01 |
| MEroyalblue | -0.15 | 0.25 | 0.10 | 0.32 | -0.26 | 0.19 | -0.04 | -0.08 | 0.01 | 0.13 | 0.08 | -0.22 | -0.26 | 0.05 | -0.13 | -0.05 | -0.34 | -0.22 | -0.22 | 0.06 | -0.36 | -0.20 | 0.01 | 0.24 | -0.15 | -0.11 | -0.16 |
| MEsaddlebrown | 0.22 | 0.38 | 0.44 | 0.17 | 0.11 | 0.37 | 0.19 | -0.11 | 0.29 | 0.42 | 0.18 | -0.14 | 0.02 | -0.26 | 0.25 | -0.09 | -0.20 | 0.21 | -0.26 | 0.23 | -0.20 | -0.07 | -0.26 | 0.17 | 0.03 | -0.13 | -0.08 |
| MEsalmon | -0.15 | -0.28 | 0.03 | -0.33 | -0.27 | -0.14 | -0.13 | 0.00 | 0.06 | 0.02 | -0.05 | -0.14 | 0.13 | -0.42 | -0.16 | -0.26 | -0.21 | -0.15 | 0.06 | 0.07 | -0.28 | -0.16 | -0.06 | -0.17 | 0.23 | -0.06 | -0.10 |
| MEskyblue | -0.18 | 0.04 | -0.15 | -0.06 | -0.21 | -0.06 | 0.16 | 0.10 | 0.13 | -0.07 | -0.42 | 0.43 | -0.35 | -0.03 | -0.15 | 0.01 | 0.38 | 0.14 | -0.20 | -0.33 | 0.19 | -0.11 | -0.16 | -0.10 | 0.04 | 0.20 | -0.29 |
| MEsteelblue | -0.04 | 0.13 | 0.08 | 0.24 | -0.15 | 0.11 | -0.11 | -0.18 | -0.14 | -0.01 | 0.33 | -0.48 | -0.03 | 0.09 | -0.08 | -0.05 | -0.52 | -0.31 | 0.01 | 0.24 | -0.37 | -0.11 | 0.17 | 0.29 | -0.24 | -0.16 | 0.08 |
| MEtan | 0.59 | 0.59 | 0.51 | 0.26 | 0.62 | 0.62 | 0.62 | -0.11 | 0.33 | 0.47 | 0.32 | -0.12 | 0.14 | 0.13 | 0.63 | 0.44 | 0.18 | 0.51 | -0.21 | 0.50 | 0.38 | 0.62 | -0.08 | 0.50 | 0.57 | 0.46 | 0.04 |
| MEturquoise | 0.11 | -0.35 | -0.02 | -0.50 | 0.14 | -0.28 | -0.07 | 0.02 | 0.05 | -0.03 | -0.14 | 0.32 | 0.32 | -0.39 | 0.04 | -0.16 | 0.35 | 0.24 | 0.11 | -0.12 | 0.26 | 0.12 | -0.29 | -0.38 | 0.19 | -0.02 | 0.15 |
| MEviolet | -0.17 | -0.19 | -0.41 | 0.18 | -0.12 | -0.22 | -0.15 | 0.21 | -0.23 | -0.44 | 0.00 | 0.00 | -0.32 | 0.40 | -0.23 | 0.13 | -0.02 | -0.24 | 0.12 | -0.18 | -0.24 | -0.19 | 0.36 | -0.13 | -0.43 | -0.05 | -0.11 |
| MEwhite | 0.07 | 0.04 | 0.16 | 0.26 | 0.07 | 0.16 | -0.21 | -0.05 | -0.15 | 0.13 | 0.35 | -0.51 | 0.15 | 0.05 | 0.12 | -0.11 | -0.57 | -0.27 | 0.16 | 0.32 | -0.53 | -0.10 | 0.27 | 0.19 | -0.13 | -0.29 | 0.16 |
| MEyellow | 0.24 | 0.50 | 0.28 | 0.60 | 0.24 | 0.50 | 0.18 | -0.05 | 0.07 | 0.28 | 0.34 | -0.36 | -0.19 | 0.34 | 0.28 | 0.20 | -0.35 | 0.01 | -0.22 | 0.33 | -0.30 | 0.04 | 0.22 | 0.46 | -0.08 | -0.03 | -0.02 |

**Control, 72hr protein vs. module correlation p value**

| | CCL13 | CCL17 | CCL19 | CCL2 | CCL20 | CCL22 | CCL3 | CCL4 | CSF2 | CXCL10 | CXCL5 | IL10 | IL15 | IL16 | IL18 | IL1A | IL1B | IL6 | IL7 | IL8 | MMP1 | MMP10 | MMP2 | MMP3 | MMP9 | TNF | VEGFA |
|---|---|---|---|---|---|---|---|---|---|---|---|---|---|---|---|---|---|---|---|---|---|---|---|---|---|---|---|
| MEblack | 0.65 | 0.01 | 0.15 | 0.01 | 0.58 | 0.01 | 0.42 | 0.53 | 0.91 | 0.08 | 0.25 | 0.04 | 0.91 | 0.13 | 0.24 | 0.23 | 0.06 | 0.62 | 0.73 | 0.09 | 0.21 | 0.68 | 0.52 | 0.01 | 0.79 | 0.94 | 0.96 |
| MEblue | 0.06 | 0.26 | 0.23 | 0.10 | 0.02 | 0.99 | 0.91 | 0.63 | 0.70 | 0.46 | 0.27 | 0.59 | 0.01 | 0.07 | 0.05 | 0.60 | 0.97 | 0.18 | 0.36 | 0.13 | 1.00 | 0.18 | 0.97 | 0.32 | 0.12 | 0.65 | 0.22 |
| MEbrown | 0.01 | 0.02 | 0.04 | 0.05 | 0.00 | 0.01 | 0.09 | 0.96 | 0.32 | 0.09 | 0.09 | 0.45 | 0.71 | 0.23 | 0.00 | 0.15 | 0.93 | 0.04 | 0.44 | 0.04 | 0.73 | 0.06 | 0.35 | 0.10 | 0.37 | 0.69 | 0.67 |
| MEcyan | 0.29 | 0.97 | 0.97 | 0.67 | 0.12 | 0.77 | 0.14 | 0.14 | 0.23 | 0.94 | 0.50 | 0.11 | 0.46 | 0.86 | 0.28 | 0.34 | 0.03 | 0.04 | 0.31 | 0.86 | 0.14 | 0.25 | 0.67 | 0.48 | 0.05 | 0.15 | 0.38 |
| MEdarkgreen | 0.66 | 0.46 | 0.68 | 0.10 | 0.38 | 0.81 | 0.34 | 0.53 | 0.73 | 0.45 | 0.35 | 0.12 | 0.73 | 0.96 | 0.51 | 0.29 | 0.03 | 0.20 | 0.80 | 0.79 | 0.28 | 0.07 | 0.62 | 0.42 | 0.27 | 0.16 | 0.84 |
| MEdarkgrey | 0.60 | 0.19 | 0.21 | 0.12 | 0.61 | 0.89 | 0.29 | 0.96 | 0.59 | 0.24 | 0.27 | 0.03 | 0.00 | 0.04 | 0.41 | 0.33 | 0.03 | 0.43 | 0.07 | 0.07 | 0.11 | 0.66 | 0.92 | 0.76 | 0.11 | 0.25 | 0.14 |
| MEdarkolivegreen | 0.44 | 0.89 | 0.55 | 0.74 | 0.12 | 0.41 | 0.31 | 0.43 | 0.77 | 0.40 | 0.78 | 0.88 | 0.26 | 0.89 | 0.09 | 0.56 | 0.73 | 0.30 | 0.35 | 0.37 | 0.53 | 0.32 | 0.40 | 0.77 | 0.19 | 0.84 | 0.72 |
| MEdarkorange | 0.16 | 0.00 | 0.10 | 0.00 | 0.11 | 0.00 | 0.33 | 0.57 | 0.97 | 0.12 | 0.06 | 0.04 | 0.67 | 0.02 | 0.07 | 0.12 | 0.12 | 0.99 | 0.39 | 0.05 | 0.35 | 0.35 | 0.30 | 0.00 | 0.99 | 0.84 | 0.69 |
| MEdarkred | 0.01 | 0.04 | 0.00 | 0.41 | 0.05 | 0.01 | 0.18 | 0.90 | 0.02 | 0.00 | 0.66 | 0.65 | 0.20 | 0.16 | 0.07 | 0.62 | 0.96 | 0.02 | 0.01 | 0.24 | 0.93 | 1.00 | 0.02 | 0.75 | 0.04 | 0.58 | 0.97 |
| MEdarkturquoise | 0.14 | 1.00 | 0.37 | 0.83 | 0.02 | 0.19 | 0.17 | 0.13 | 0.73 | 0.23 | 0.74 | 0.54 | 0.42 | 0.91 | 0.03 | 0.28 | 0.93 | 0.22 | 0.62 | 0.10 | 0.83 | 0.07 | 0.29 | 0.84 | 0.01 | 0.49 | 0.72 |
| MEgreen | 0.05 | 0.02 | 0.06 | 0.08 | 0.03 | 0.01 | 0.03 | 0.69 | 0.08 | 0.03 | 0.70 | 0.68 | 0.46 | 0.50 | 0.02 | 0.21 | 0.46 | 0.03 | 0.05 | 0.28 | 0.99 | 0.28 | 0.87 | 0.29 | 0.10 | 0.29 | 0.30 |
| MEgreenyellow | 0.74 | 0.49 | 0.47 | 0.56 | 0.60 | 0.31 | 0.69 | 0.76 | 0.60 | 0.21 | 0.97 | 0.42 | 0.82 | 0.76 | 0.97 | 0.93 | 0.28 | 0.70 | 0.67 | 0.53 | 0.29 | 0.73 | 0.90 | 0.69 | 0.37 | 0.91 | 0.48 |
| MEgrey | 0.90 | 0.08 | 0.85 | 0.09 | 0.61 | 0.25 | 0.05 | 0.60 | 0.32 | 0.98 | 0.46 | 0.45 | 0.03 | 0.02 | 0.60 | 0.06 | 0.18 | 0.33 | 0.19 | 0.82 | 0.18 | 0.65 | 0.27 | 0.25 | 0.53 | 0.07 | 0.11 |
| MEgrey60 | 0.88 | 0.31 | 0.44 | 0.30 | 0.83 | 0.91 | 0.41 | 0.55 | 0.97 | 0.85 | 0.22 | 0.03 | 0.08 | 0.13 | 0.81 | 0.28 | 0.03 | 0.38 | 0.26 | 0.12 | 0.07 | 0.79 | 0.69 | 0.96 | 0.50 | 0.61 | 0.59 |
| MElightcyan | 0.11 | 0.87 | 0.05 | 0.81 | 0.11 | 0.11 | 0.82 | 0.75 | 0.62 | 0.07 | 0.06 | 0.01 | 0.14 | 0.27 | 0.06 | 0.82 | 0.02 | 0.86 | 0.71 | 0.01 | 0.01 | 0.54 | 0.41 | 0.53 | 0.12 | 0.51 | 0.53 |
| MElightgreen | 0.32 | 0.03 | 0.96 | 0.31 | 0.43 | 0.13 | 0.00 | 0.63 | 0.07 | 0.66 | 0.22 | 0.04 | 0.01 | 0.12 | 0.36 | 0.13 | 0.00 | 0.01 | 0.03 | 0.72 | 0.03 | 0.54 | 0.87 | 0.29 | 0.57 | 0.00 | 0.17 |
| MElightyellow | 0.74 | 0.00 | 0.89 | 0.00 | 0.59 | 0.04 | 0.11 | 0.63 | 0.97 | 0.98 | 0.64 | 0.61 | 0.05 | 0.00 | 0.60 | 0.01 | 0.90 | 0.95 | 0.23 | 0.72 | 0.21 | 0.44 | 0.28 | 0.00 | 0.81 | 0.08 | 0.59 |
| MEmagenta | 0.04 | 0.73 | 0.14 | 0.55 | 0.06 | 0.77 | 0.96 | 0.78 | 0.37 | 0.40 | 0.13 | 0.47 | 0.10 | 0.12 | 0.10 | 0.30 | 0.60 | 0.25 | 0.95 | 0.15 | 0.62 | 0.71 | 0.96 | 0.78 | 0.42 | 0.58 | 0.41 |
| MEmidnightblue | 0.67 | 0.58 | 0.12 | 0.51 | 0.87 | 0.17 | 0.90 | 0.99 | 0.81 | 0.09 | 0.06 | 0.00 | 0.48 | 0.26 | 0.77 | 0.60 | 0.00 | 0.30 | 0.83 | 0.02 | 0.00 | 0.85 | 0.77 | 0.30 | 0.68 | 0.15 | 0.43 |
| MEorange | 0.08 | 0.36 | 0.37 | 1.00 | 0.05 | 0.57 | 0.21 | 0.97 | 0.27 | 0.44 | 0.72 | 0.17 | 0.75 | 0.95 | 0.18 | 0.38 | 0.04 | 0.03 | 0.34 | 0.93 | 0.02 | 0.11 | 0.35 | 1.00 | 0.37 | 0.38 | 0.95 |
| MEpaleturquoise | 0.08 | 0.98 | 0.50 | 0.14 | 0.04 | 0.67 | 0.07 | 0.68 | 0.29 | 0.52 | 0.64 | 0.14 | 0.35 | 0.41 | 0.12 | 0.45 | 0.01 | 0.01 | 0.79 | 0.72 | 0.01 | 0.06 | 0.27 | 0.57 | 0.02 | 0.12 | 0.68 |
| MEpink | 0.09 | 0.00 | 0.29 | 0.09 | 0.08 | 0.01 | 0.00 | 0.70 | 0.08 | 0.28 | 0.81 | 0.35 | 0.18 | 0.08 | 0.07 | 0.03 | 0.05 | 0.01 | 0.05 | 0.50 | 0.03 | 0.11 | 0.82 | 0.05 | 0.11 | 0.00 | 0.36 |
| MEpurple | 0.70 | 0.18 | 0.84 | 0.11 | 0.88 | 0.52 | 0.85 | 0.32 | 0.83 | 0.90 | 0.18 | 0.40 | 0.33 | 0.30 | 0.74 | 0.71 | 0.35 | 0.74 | 0.32 | 0.69 | 0.51 | 0.80 | 0.87 | 0.19 | 0.10 | 0.82 | 0.82 |
| MEred | 0.35 | 0.86 | 0.94 | 0.50 | 0.10 | 0.97 | 0.28 | 0.41 | 0.66 | 0.98 | 0.45 | 0.08 | 0.84 | 0.99 | 0.24 | 0.49 | 0.02 | 0.05 | 0.93 | 0.72 | 0.09 | 0.19 | 0.85 | 0.36 | 0.40 | 0.50 | 0.97 |
| MEroyalblue | 0.48 | 0.21 | 0.61 | 0.11 | 0.20 | 0.36 | 0.86 | 0.69 | 0.97 | 0.52 | 0.68 | 0.29 | 0.20 | 0.80 | 0.52 | 0.81 | 0.09 | 0.28 | 0.29 | 0.78 | 0.07 | 0.32 | 0.96 | 0.24 | 0.45 | 0.61 | 0.45 |
| MEsaddlebrown | 0.28 | 0.06 | 0.03 | 0.40 | 0.60 | 0.06 | 0.35 | 0.59 | 0.15 | 0.03 | 0.38 | 0.50 | 0.93 | 0.20 | 0.21 | 0.67 | 0.32 | 0.31 | 0.21 | 0.26 | 0.32 | 0.72 | 0.20 | 0.41 | 0.88 | 0.53 | 0.71 |
| MEsalmon | 0.46 | 0.17 | 0.88 | 0.10 | 0.18 | 0.49 | 0.53 | 1.00 | 0.75 | 0.92 | 0.83 | 0.49 | 0.54 | 0.03 | 0.45 | 0.19 | 0.29 | 0.46 | 0.76 | 0.73 | 0.16 | 0.43 | 0.78 | 0.40 | 0.25 | 0.78 | 0.64 |
| MEskyblue | 0.38 | 0.85 | 0.46 | 0.77 | 0.30 | 0.79 | 0.45 | 0.64 | 0.53 | 0.73 | 0.03 | 0.03 | 0.08 | 0.88 | 0.47 | 0.97 | 0.06 | 0.49 | 0.32 | 0.10 | 0.35 | 0.60 | 0.45 | 0.61 | 0.84 | 0.32 | 0.15 |
| MEsteelblue | 0.84 | 0.52 | 0.71 | 0.24 | 0.46 | 0.58 | 0.60 | 0.38 | 0.49 | 0.96 | 0.10 | 0.01 | 0.89 | 0.68 | 0.69 | 0.82 | 0.01 | 0.13 | 0.95 | 0.24 | 0.06 | 0.59 | 0.42 | 0.15 | 0.24 | 0.43 | 0.69 |
| MEtan | 0.00 | 0.00 | 0.01 | 0.20 | 0.00 | 0.00 | 0.00 | 0.58 | 0.10 | 0.01 | 0.11 | 0.56 | 0.49 | 0.53 | 0.00 | 0.02 | 0.38 | 0.01 | 0.29 | 0.01 | 0.06 | 0.00 | 0.69 | 0.01 | 0.00 | 0.02 | 0.85 |
| MEturquoise | 0.60 | 0.08 | 0.92 | 0.01 | 0.51 | 0.17 | 0.74 | 0.91 | 0.80 | 0.87 | 0.50 | 0.11 | 0.12 | 0.05 | 0.86 | 0.44 | 0.08 | 0.24 | 0.59 | 0.55 | 0.20 | 0.55 | 0.15 | 0.05 | 0.35 | 0.92 | 0.46 |
| MEviolet | 0.42 | 0.36 | 0.04 | 0.39 | 0.57 | 0.28 | 0.48 | 0.31 | 0.25 | 0.02 | 1.00 | 0.99 | 0.11 | 0.04 | 0.26 | 0.53 | 0.91 | 0.24 | 0.54 | 0.39 | 0.25 | 0.36 | 0.07 | 0.52 | 0.03 | 0.82 | 0.61 |
| MEwhite | 0.73 | 0.85 | 0.44 | 0.21 | 0.72 | 0.43 | 0.30 | 0.82 | 0.46 | 0.54 | 0.08 | 0.01 | 0.48 | 0.81 | 0.56 | 0.61 | 0.00 | 0.18 | 0.45 | 0.11 | 0.01 | 0.62 | 0.19 | 0.35 | 0.52 | 0.15 | 0.43 |
| MEyellow | 0.24 | 0.01 | 0.16 | 0.00 | 0.23 | 0.01 | 0.39 | 0.80 | 0.73 | 0.17 | 0.09 | 0.07 | 0.36 | 0.09 | 0.17 | 0.34 | 0.08 | 0.96 | 0.29 | 0.10 | 0.13 | 0.86 | 0.28 | 0.02 | 0.69 | 0.87 | 0.92 |

**Supplementary File 9F. Protein levels with gene module correlations, 72hr infected**

**Infected, 72hr protein vs. module correlation**

| | GT2 | CCL13 | CCL17 | CCL19 | CCL2 | CCL20 | CCL22 | CCL3 | CCL4 | CSF2 | CXCL10 | CXCL5 | IL10 | IL15 | IL16 | IL18 | IL1A | IL1B | IL6 | IL7 | IL8 | MMP1 | MMP10 | MMP2 | MMP3 | MMP9 | TNF | VEGFA |
|---|---|---|---|---|---|---|---|---|---|---|---|---|---|---|---|---|---|---|---|---|---|---|---|---|---|---|---|---|
| MEblack | 0.07 | 0.25 | 0.25 | 0.22 | 0.14 | 0.22 | 0.34 | -0.06 | -0.21 | 0.32 | 0.06 | 0.06 | 0.03 | 0.24 | 0.04 | 0.27 | 0.33 | 0.25 | 0.32 | 0.17 | 0.15 | -0.06 | 0.03 | 0.06 | 0.16 | 0.30 | 0.38 | 0.11 |
| MEblue | -0.02 | 0.02 | 0.01 | -0.14 | -0.09 | -0.18 | 0.06 | -0.20 | -0.26 | 0.17 | -0.12 | -0.15 | 0.05 | -0.12 | -0.13 | -0.12 | -0.03 | 0.04 | 0.08 | 0.09 | -0.13 | 0.03 | -0.10 | 0.04 | 0.03 | 0.03 | 0.07 | -0.21 |
| MEbrown | -0.08 | -0.12 | 0.30 | 0.21 | 0.40 | 0.23 | 0.03 | 0.32 | 0.05 | 0.05 | -0.09 | 0.27 | -0.08 | -0.01 | 0.49 | 0.22 | 0.33 | 0.15 | -0.02 | 0.01 | 0.22 | -0.11 | 0.19 | 0.08 | 0.24 | -0.26 | 0.04 | 0.21 |
| MEcyan | 0.00 | -0.28 | -0.17 | -0.06 | 0.05 | -0.13 | -0.29 | 0.17 | 0.32 | -0.27 | 0.00 | -0.04 | 0.01 | -0.16 | 0.09 | -0.15 | -0.15 | -0.14 | -0.38 | -0.21 | -0.05 | 0.01 | 0.05 | -0.01 | -0.13 | -0.22 | -0.27 | 0.12 |
| MEdarkgreen | 0.09 | -0.09 | -0.18 | -0.02 | -0.15 | 0.02 | -0.16 | -0.06 | 0.30 | -0.16 | 0.04 | -0.15 | 0.21 | 0.05 | -0.08 | -0.02 | -0.17 | -0.02 | -0.12 | -0.17 | -0.15 | 0.07 | 0.00 | -0.21 | -0.26 | -0.01 | -0.11 | 0.22 |
| MEdarkgrey | -0.21 | -0.47 | 0.00 | -0.30 | 0.11 | -0.39 | -0.34 | 0.07 | -0.11 | -0.07 | -0.42 | -0.01 | -0.06 | -0.57 | 0.23 | -0.35 | -0.14 | -0.11 | -0.37 | -0.09 | -0.18 | -0.05 | -0.01 | 0.07 | 0.08 | -0.73 | -0.36 | -0.23 |
| MEdarkolivegre | -0.17 | -0.34 | 0.02 | -0.16 | 0.08 | -0.31 | -0.22 | 0.22 | 0.04 | -0.21 | -0.11 | 0.20 | -0.32 | -0.36 | 0.16 | -0.33 | -0.19 | -0.31 | -0.30 | -0.09 | 0.09 | -0.32 | 0.07 | 0.14 | 0.11 | -0.51 | -0.37 | -0.24 |
| MEdarkorange | 0.08 | 0.11 | -0.51 | -0.24 | -0.50 | -0.19 | -0.24 | -0.19 | 0.29 | -0.23 | 0.14 | -0.36 | 0.18 | 0.06 | -0.58 | -0.21 | -0.43 | -0.19 | -0.07 | -0.15 | -0.24 | 0.35 | -0.09 | -0.07 | -0.48 | 0.29 | -0.12 | -0.07 |
| MEdarkred | 0.19 | 0.43 | 0.37 | 0.61 | 0.49 | 0.41 | 0.38 | 0.22 | 0.13 | 0.52 | 0.19 | 0.04 | 0.39 | 0.56 | 0.06 | 0.50 | 0.52 | 0.50 | 0.29 | 0.03 | 0.26 | -0.02 | 0.16 | -0.03 | 0.02 | 0.30 | 0.59 | 0.61 |
| MEdarkturquoi | -0.05 | 0.11 | 0.10 | -0.10 | -0.07 | -0.10 | 0.09 | -0.15 | -0.32 | 0.25 | -0.13 | 0.05 | -0.02 | -0.06 | -0.10 | -0.09 | -0.03 | 0.01 | 0.23 | 0.14 | -0.06 | 0.06 | -0.04 | 0.07 | 0.09 | -0.13 | 0.10 | -0.30 |
| MEgreen | 0.05 | 0.11 | -0.17 | -0.26 | -0.38 | -0.10 | 0.04 | -0.34 | -0.26 | -0.06 | 0.06 | -0.15 | 0.00 | 0.01 | -0.32 | -0.13 | -0.27 | -0.15 | 0.14 | 0.13 | -0.22 | 0.20 | -0.21 | -0.09 | -0.05 | 0.22 | -0.04 | -0.35 |
| MEgreenyellow | -0.06 | -0.18 | 0.05 | -0.11 | 0.05 | -0.02 | -0.13 | 0.00 | -0.10 | -0.08 | -0.19 | 0.06 | 0.00 | -0.18 | 0.24 | -0.06 | -0.06 | -0.03 | -0.09 | 0.01 | -0.13 | 0.03 | -0.07 | -0.11 | 0.11 | -0.35 | -0.17 | -0.11 |
| MEgrey | -0.28 | -0.43 | -0.17 | -0.54 | -0.23 | -0.60 | -0.40 | -0.25 | -0.23 | 0.05 | -0.62 | -0.34 | 0.18 | -0.71 | -0.16 | -0.51 | -0.37 | -0.05 | -0.30 | -0.01 | -0.50 | -0.01 | -0.15 | -0.01 | -0.15 | -0.75 | -0.37 | -0.42 |
| MEgrey60 | 0.04 | 0.15 | -0.04 | 0.00 | -0.22 | 0.03 | 0.19 | 0.08 | -0.04 | -0.17 | 0.43 | 0.23 | -0.46 | 0.13 | -0.24 | -0.05 | -0.13 | -0.34 | 0.20 | 0.01 | 0.30 | -0.24 | 0.07 | 0.22 | 0.11 | 0.39 | -0.06 | -0.29 |
| MElightcyan | 0.03 | 0.12 | -0.12 | -0.23 | -0.25 | -0.11 | 0.00 | -0.52 | -0.34 | 0.24 | -0.28 | -0.35 | 0.43 | -0.03 | -0.24 | -0.06 | -0.14 | 0.20 | 0.12 | 0.15 | -0.48 | 0.42 | -0.28 | -0.25 | -0.15 | 0.00 | 0.13 | -0.16 |
| MElightgreen | -0.08 | -0.14 | -0.10 | -0.35 | -0.35 | -0.26 | 0.04 | -0.41 | -0.38 | 0.22 | -0.35 | -0.26 | 0.15 | -0.44 | -0.23 | -0.29 | -0.24 | -0.01 | 0.06 | 0.03 | -0.41 | 0.20 | -0.15 | -0.04 | 0.03 | -0.10 | -0.15 | -0.42 |
| MElightyellow | 0.00 | -0.01 | -0.31 | -0.02 | -0.15 | -0.05 | -0.27 | 0.16 | 0.47 | -0.40 | 0.17 | -0.11 | 0.02 | 0.08 | -0.24 | -0.08 | -0.27 | -0.21 | -0.26 | -0.17 | 0.00 | -0.01 | 0.03 | -0.08 | -0.33 | 0.05 | -0.24 | 0.18 |
| MEmagenta | -0.06 | -0.17 | 0.07 | 0.04 | 0.15 | 0.02 | -0.10 | 0.31 | 0.14 | -0.20 | 0.05 | 0.25 | -0.25 | -0.09 | 0.23 | -0.04 | -0.01 | -0.17 | -0.15 | -0.07 | 0.19 | -0.19 | 0.13 | 0.10 | 0.14 | -0.25 | -0.22 | 0.00 |
| MEmidnightblu | -0.11 | 0.11 | 0.29 | 0.36 | 0.30 | 0.12 | 0.30 | 0.43 | 0.12 | 0.01 | 0.41 | 0.41 | -0.49 | 0.24 | 0.12 | 0.15 | 0.26 | -0.13 | 0.08 | -0.01 | 0.64 | -0.56 | 0.27 | 0.28 | 0.25 | 0.17 | 0.09 | 0.18 |
| MEorange | -0.06 | -0.02 | 0.11 | 0.20 | 0.29 | 0.15 | -0.07 | 0.17 | 0.19 | 0.10 | -0.15 | 0.14 | 0.14 | 0.08 | 0.29 | 0.15 | 0.20 | 0.18 | -0.07 | -0.04 | 0.08 | 0.07 | 0.12 | -0.06 | -0.02 | -0.25 | 0.09 | 0.29 |
| MEpaleturquois | -0.19 | -0.42 | 0.05 | -0.27 | 0.10 | -0.38 | -0.19 | 0.30 | -0.10 | -0.35 | -0.01 | 0.24 | -0.62 | -0.43 | 0.25 | -0.38 | -0.17 | -0.45 | -0.30 | -0.01 | 0.21 | -0.39 | 0.05 | 0.33 | 0.30 | -0.39 | -0.42 | -0.44 |
| MEpink | 0.09 | 0.36 | 0.27 | 0.31 | 0.16 | 0.29 | 0.41 | -0.26 | -0.27 | 0.71 | -0.23 | -0.11 | 0.51 | 0.27 | 0.01 | 0.37 | 0.43 | 0.58 | 0.45 | 0.18 | -0.09 | 0.19 | -0.03 | -0.12 | 0.04 | 0.27 | 0.61 | 0.24 |
| MEpurple | 0.22 | 0.20 | -0.33 | -0.07 | -0.36 | 0.05 | -0.01 | -0.25 | 0.12 | -0.09 | 0.15 | -0.37 | 0.28 | 0.20 | -0.40 | 0.05 | -0.17 | 0.05 | 0.08 | -0.07 | -0.24 | 0.32 | -0.13 | -0.14 | -0.31 | 0.49 | 0.08 | 0.09 |
| MEred | -0.02 | 0.09 | 0.09 | -0.03 | -0.05 | -0.08 | 0.18 | 0.01 | -0.22 | 0.06 | 0.15 | 0.15 | -0.32 | 0.00 | -0.10 | -0.09 | -0.02 | -0.16 | 0.17 | 0.09 | 0.19 | -0.22 | 0.02 | 0.20 | 0.18 | 0.14 | 0.03 | -0.29 |
| MEroyalblue | -0.06 | -0.03 | 0.14 | 0.09 | 0.15 | -0.13 | 0.07 | 0.22 | 0.04 | 0.13 | 0.09 | 0.15 | -0.27 | -0.05 | 0.00 | -0.08 | 0.12 | -0.05 | 0.02 | -0.06 | 0.30 | -0.34 | 0.18 | 0.29 | 0.08 | -0.07 | 0.05 | -0.03 |
| MEsaddlebrowr | 0.00 | 0.34 | 0.20 | 0.34 | 0.09 | 0.21 | 0.25 | -0.09 | -0.13 | 0.46 | -0.14 | 0.00 | 0.23 | 0.22 | -0.12 | 0.32 | 0.37 | 0.42 | 0.26 | 0.11 | 0.08 | -0.07 | 0.03 | -0.06 | -0.04 | 0.01 | 0.33 | 0.40 |
| MEsalmon | 0.10 | 0.12 | -0.28 | -0.06 | -0.34 | -0.13 | 0.02 | -0.03 | 0.19 | -0.18 | 0.37 | -0.08 | -0.17 | 0.12 | -0.47 | -0.15 | -0.26 | -0.27 | 0.02 | -0.11 | 0.10 | -0.08 | 0.03 | 0.13 | -0.21 | 0.43 | -0.07 | -0.12 |
| MEskyblue | -0.08 | 0.00 | 0.27 | 0.16 | 0.26 | 0.22 | 0.13 | 0.18 | -0.12 | -0.06 | 0.08 | 0.46 | -0.32 | 0.08 | 0.38 | 0.15 | 0.17 | -0.09 | 0.06 | 0.11 | 0.28 | -0.14 | 0.08 | 0.04 | 0.32 | -0.13 | -0.01 | -0.04 |
| MEsteelblue | -0.16 | -0.09 | 0.29 | 0.05 | 0.19 | -0.08 | 0.17 | 0.22 | -0.23 | 0.05 | 0.11 | 0.47 | -0.58 | -0.14 | 0.21 | -0.11 | 0.10 | -0.24 | 0.09 | 0.06 | 0.41 | -0.46 | 0.14 | 0.30 | 0.40 | -0.18 | -0.08 | -0.31 |
| MEtan | 0.22 | 0.61 | 0.47 | 0.76 | 0.47 | 0.70 | 0.55 | 0.06 | 0.01 | 0.61 | 0.18 | 0.27 | 0.34 | 0.76 | 0.24 | 0.74 | 0.74 | 0.64 | 0.53 | 0.16 | 0.36 | 0.00 | 0.10 | -0.17 | 0.10 | 0.43 | 0.78 | 0.66 |
| MEturquoise | 0.13 | 0.09 | -0.20 | 0.03 | -0.16 | 0.10 | -0.15 | -0.08 | 0.28 | -0.09 | 0.06 | -0.18 | 0.31 | 0.15 | -0.21 | 0.04 | -0.15 | 0.05 | -0.05 | -0.12 | -0.19 | 0.26 | 0.00 | -0.21 | -0.30 | 0.06 | -0.04 | 0.25 |
| MEviolet | -0.26 | -0.45 | 0.01 | -0.27 | 0.15 | -0.28 | -0.28 | 0.26 | 0.06 | -0.34 | -0.11 | 0.15 | -0.35 | -0.42 | 0.32 | -0.33 | -0.17 | -0.34 | -0.37 | -0.03 | 0.06 | -0.21 | 0.06 | 0.17 | 0.18 | -0.44 | -0.39 | -0.34 |
| MEwhite | -0.16 | -0.30 | 0.21 | -0.06 | 0.24 | -0.07 | -0.03 | 0.35 | -0.11 | -0.30 | 0.05 | 0.36 | -0.55 | -0.26 | 0.39 | -0.10 | 0.03 | -0.28 | -0.18 | 0.04 | 0.28 | -0.44 | 0.07 | 0.21 | 0.39 | -0.29 | -0.30 | -0.26 |
| MEyellow | -0.18 | -0.26 | 0.37 | 0.11 | 0.45 | 0.00 | 0.04 | 0.40 | -0.07 | 0.02 | -0.09 | 0.38 | -0.37 | -0.23 | 0.52 | 0.01 | 0.28 | -0.03 | -0.12 | 0.02 | 0.34 | -0.41 | 0.20 | 0.25 | 0.40 | -0.41 | -0.09 | -0.02 |

**Infected, 72hr protein vs. module correlation p value**

| | GT2 | CCL13 | CCL17 | CCL19 | CCL2 | CCL20 | CCL22 | CCL3 | CCL4 | CSF2 | CXCL10 | CXCL5 | IL10 | IL15 | IL16 | IL18 | IL1A | IL1B | IL6 | IL7 | IL8 | MMP1 | MMP10 | MMP2 | MMP3 | MMP9 | TNF | VEGFA |
|---|---|---|---|---|---|---|---|---|---|---|---|---|---|---|---|---|---|---|---|---|---|---|---|---|---|---|---|---|
| MEblack | 0.74 | 0.21 | 0.22 | 0.29 | 0.50 | 0.29 | 0.09 | 0.79 | 0.29 | 0.11 | 0.77 | 0.79 | 0.88 | 0.24 | 0.84 | 0.18 | 0.10 | 0.23 | 0.11 | 0.40 | 0.46 | 0.78 | 0.89 | 0.77 | 0.45 | 0.14 | 0.06 | 0.60 |
| MEblue | 0.93 | 0.94 | 0.98 | 0.48 | 0.65 | 0.39 | 0.76 | 0.32 | 0.20 | 0.42 | 0.56 | 0.46 | 0.80 | 0.57 | 0.52 | 0.56 | 0.87 | 0.86 | 0.71 | 0.67 | 0.51 | 0.89 | 0.61 | 0.85 | 0.90 | 0.89 | 0.73 | 0.31 |
| MEbrown | 0.71 | 0.56 | 0.14 | 0.31 | 0.04 | 0.26 | 0.89 | 0.11 | 0.80 | 0.81 | 0.67 | 0.18 | 0.69 | 0.95 | 0.01 | 0.29 | 0.09 | 0.48 | 0.91 | 0.96 | 0.29 | 0.61 | 0.36 | 0.71 | 0.23 | 0.20 | 0.84 | 0.29 |
| MEcyan | 0.99 | 0.17 | 0.41 | 0.76 | 0.82 | 0.53 | 0.15 | 0.40 | 0.11 | 0.18 | 0.99 | 0.86 | 0.96 | 0.43 | 0.66 | 0.46 | 0.45 | 0.51 | 0.05 | 0.30 | 0.82 | 0.95 | 0.79 | 0.96 | 0.52 | 0.29 | 0.18 | 0.57 |
| MEdarkgreen | 0.65 | 0.67 | 0.37 | 0.92 | 0.46 | 0.94 | 0.45 | 0.78 | 0.14 | 0.43 | 0.84 | 0.46 | 0.31 | 0.80 | 0.68 | 0.92 | 0.40 | 0.92 | 0.55 | 0.40 | 0.46 | 0.73 | 0.99 | 0.31 | 0.19 | 0.96 | 0.60 | 0.28 |
| MEdarkgrey | 0.30 | 0.02 | 0.99 | 0.14 | 0.60 | 0.05 | 0.09 | 0.73 | 0.61 | 0.73 | 0.03 | 0.98 | 0.76 | 0.00 | 0.25 | 0.08 | 0.48 | 0.59 | 0.06 | 0.66 | 0.37 | 0.80 | 0.95 | 0.72 | 0.69 | 0.00 | 0.07 | 0.25 |
| MEdarkolivegre | 0.41 | 0.09 | 0.92 | 0.45 | 0.69 | 0.13 | 0.27 | 0.29 | 0.86 | 0.31 | 0.58 | 0.33 | 0.11 | 0.07 | 0.42 | 0.10 | 0.35 | 0.13 | 0.14 | 0.66 | 0.67 | 0.11 | 0.74 | 0.50 | 0.60 | 0.01 | 0.06 | 0.24 |
| MEdarkorange | 0.70 | 0.60 | 0.01 | 0.23 | 0.01 | 0.35 | 0.25 | 0.34 | 0.15 | 0.27 | 0.49 | 0.07 | 0.39 | 0.76 | 0.00 | 0.30 | 0.03 | 0.36 | 0.72 | 0.46 | 0.23 | 0.08 | 0.66 | 0.73 | 0.01 | 0.16 | 0.55 | 0.72 |
| MEdarkred | 0.36 | 0.03 | 0.06 | 0.00 | 0.01 | 0.04 | 0.05 | 0.28 | 0.54 | 0.01 | 0.36 | 0.86 | 0.05 | 0.00 | 0.77 | 0.01 | 0.01 | 0.01 | 0.15 | 0.87 | 0.19 | 0.93 | 0.44 | 0.89 | 0.94 | 0.13 | 0.00 | 0.00 |
| MEdarkturquoi | 0.80 | 0.58 | 0.62 | 0.62 | 0.73 | 0.61 | 0.67 | 0.45 | 0.11 | 0.22 | 0.52 | 0.82 | 0.93 | 0.78 | 0.62 | 0.65 | 0.88 | 0.95 | 0.26 | 0.50 | 0.78 | 0.78 | 0.84 | 0.72 | 0.68 | 0.53 | 0.62 | 0.13 |
| MEgreen | 0.81 | 0.59 | 0.40 | 0.20 | 0.05 | 0.63 | 0.84 | 0.09 | 0.20 | 0.76 | 0.76 | 0.46 | 0.98 | 0.98 | 0.11 | 0.52 | 0.18 | 0.47 | 0.48 | 0.52 | 0.28 | 0.32 | 0.31 | 0.66 | 0.81 | 0.28 | 0.84 | 0.08 |
| MEgreenyellow | 0.79 | 0.39 | 0.82 | 0.59 | 0.82 | 0.91 | 0.52 | 0.99 | 0.63 | 0.68 | 0.35 | 0.76 | 0.98 | 0.39 | 0.24 | 0.77 | 0.77 | 0.89 | 0.66 | 0.96 | 0.53 | 0.89 | 0.75 | 0.60 | 0.58 | 0.08 | 0.41 | 0.60 |
| MEgrey | 0.16 | 0.03 | 0.42 | 0.00 | 0.25 | 0.00 | 0.05 | 0.22 | 0.26 | 0.79 | 0.00 | 0.09 | 0.39 | 0.00 | 0.44 | 0.01 | 0.07 | 0.79 | 0.14 | 0.95 | 0.01 | 0.96 | 0.46 | 0.95 | 0.47 | 0.00 | 0.06 | 0.03 |
| MEgrey60 | 0.84 | 0.45 | 0.86 | 0.99 | 0.29 | 0.88 | 0.34 | 0.71 | 0.85 | 0.41 | 0.03 | 0.25 | 0.02 | 0.54 | 0.23 | 0.82 | 0.52 | 0.09 | 0.32 | 0.98 | 0.14 | 0.25 | 0.72 | 0.28 | 0.59 | 0.05 | 0.78 | 0.15 |
| MElightcyan | 0.90 | 0.57 | 0.55 | 0.26 | 0.22 | 0.59 | 0.99 | 0.01 | 0.09 | 0.24 | 0.17 | 0.08 | 0.03 | 0.87 | 0.23 | 0.78 | 0.50 | 0.33 | 0.55 | 0.46 | 0.01 | 0.03 | 0.17 | 0.21 | 0.46 | 0.98 | 0.52 | 0.43 |
| MElightgreen | 0.68 | 0.51 | 0.63 | 0.08 | 0.08 | 0.20 | 0.84 | 0.04 | 0.05 | 0.28 | 0.08 | 0.21 | 0.45 | 0.02 | 0.26 | 0.15 | 0.24 | 0.98 | 0.76 | 0.89 | 0.04 | 0.32 | 0.47 | 0.85 | 0.89 | 0.63 | 0.48 | 0.03 |
| MElightyellow | 0.99 | 0.96 | 0.13 | 0.93 | 0.47 | 0.80 | 0.19 | 0.45 | 0.02 | 0.04 | 0.40 | 0.60 | 0.94 | 0.71 | 0.25 | 0.71 | 0.19 | 0.31 | 0.20 | 0.40 | 0.99 | 0.95 | 0.90 | 0.69 | 0.10 | 0.82 | 0.23 | 0.37 |
| MEmagenta | 0.75 | 0.42 | 0.72 | 0.84 | 0.47 | 0.93 | 0.64 | 0.13 | 0.51 | 0.33 | 0.80 | 0.22 | 0.21 | 0.65 | 0.26 | 0.84 | 0.95 | 0.41 | 0.47 | 0.75 | 0.36 | 0.36 | 0.52 | 0.62 | 0.50 | 0.22 | 0.28 | 1.00 |
| MEmidnightblu | 0.59 | 0.58 | 0.15 | 0.07 | 0.13 | 0.56 | 0.13 | 0.03 | 0.57 | 0.98 | 0.04 | 0.04 | 0.01 | 0.24 | 0.56 | 0.46 | 0.21 | 0.53 | 0.68 | 0.97 | 0.00 | 0.00 | 0.18 | 0.17 | 0.22 | 0.40 | 0.66 | 0.38 |
| MEorange | 0.76 | 0.94 | 0.58 | 0.32 | 0.15 | 0.46 | 0.74 | 0.40 | 0.34 | 0.61 | 0.45 | 0.51 | 0.49 | 0.70 | 0.14 | 0.46 | 0.33 | 0.39 | 0.74 | 0.85 | 0.71 | 0.72 | 0.57 | 0.76 | 0.93 | 0.23 | 0.65 | 0.15 |
| MEpaleturquois | 0.35 | 0.03 | 0.80 | 0.18 | 0.62 | 0.06 | 0.35 | 0.13 | 0.63 | 0.08 | 0.96 | 0.25 | 0.00 | 0.03 | 0.23 | 0.05 | 0.41 | 0.02 | 0.13 | 0.96 | 0.30 | 0.05 | 0.80 | 0.10 | 0.14 | 0.05 | 0.03 | 0.03 |
| MEpink | 0.65 | 0.07 | 0.18 | 0.13 | 0.43 | 0.15 | 0.04 | 0.20 | 0.19 | 0.00 | 0.25 | 0.59 | 0.01 | 0.18 | 0.96 | 0.06 | 0.03 | 0.00 | 0.02 | 0.38 | 0.68 | 0.36 | 0.88 | 0.57 | 0.85 | 0.19 | 0.00 | 0.24 |
| MEpurple | 0.29 | 0.34 | 0.10 | 0.74 | 0.07 | 0.79 | 0.95 | 0.23 | 0.56 | 0.68 | 0.45 | 0.06 | 0.16 | 0.33 | 0.04 | 0.81 | 0.40 | 0.79 | 0.71 | 0.73 | 0.23 | 0.11 | 0.53 | 0.50 | 0.13 | 0.01 | 0.70 | 0.67 |
| MEred | 0.94 | 0.67 | 0.65 | 0.90 | 0.80 | 0.69 | 0.38 | 0.98 | 0.28 | 0.78 | 0.46 | 0.45 | 0.12 | 0.99 | 0.64 | 0.67 | 0.93 | 0.42 | 0.41 | 0.66 | 0.36 | 0.28 | 0.94 | 0.33 | 0.37 | 0.51 | 0.88 | 0.15 |
| MEroyalblue | 0.78 | 0.88 | 0.51 | 0.65 | 0.47 | 0.53 | 0.73 | 0.28 | 0.83 | 0.54 | 0.65 | 0.45 | 0.18 | 0.79 | 1.00 | 0.71 | 0.57 | 0.80 | 0.90 | 0.77 | 0.13 | 0.09 | 0.38 | 0.15 | 0.70 | 0.74 | 0.81 | 0.90 |
| MEsaddlebrowr | 0.99 | 0.09 | 0.33 | 0.09 | 0.64 | 0.31 | 0.23 | 0.65 | 0.54 | 0.02 | 0.49 | 0.98 | 0.25 | 0.28 | 0.57 | 0.11 | 0.06 | 0.03 | 0.19 | 0.60 | 0.71 | 0.72 | 0.88 | 0.75 | 0.84 | 0.94 | 0.10 | 0.04 |
| MEsalmon | 0.64 | 0.57 | 0.17 | 0.77 | 0.09 | 0.54 | 0.91 | 0.89 | 0.34 | 0.39 | 0.06 | 0.69 | 0.40 | 0.57 | 0.02 | 0.45 | 0.19 | 0.18 | 0.92 | 0.59 | 0.63 | 0.68 | 0.88 | 0.52 | 0.31 | 0.03 | 0.73 | 0.58 |
| MEskyblue | 0.69 | 1.00 | 0.19 | 0.45 | 0.21 | 0.29 | 0.52 | 0.38 | 0.56 | 0.76 | 0.70 | 0.02 | 0.11 | 0.71 | 0.05 | 0.46 | 0.40 | 0.65 | 0.76 | 0.58 | 0.16 | 0.48 | 0.69 | 0.84 | 0.11 | 0.53 | 0.96 | 0.83 |
| MEsteelblue | 0.44 | 0.65 | 0.15 | 0.81 | 0.36 | 0.69 | 0.39 | 0.29 | 0.26 | 0.80 | 0.58 | 0.02 | 0.00 | 0.48 | 0.30 | 0.60 | 0.62 | 0.23 | 0.67 | 0.78 | 0.04 | 0.02 | 0.50 | 0.13 | 0.04 | 0.37 | 0.71 | 0.13 |
| MEtan | 0.29 | 0.00 | 0.02 | 0.00 | 0.02 | 0.00 | 0.00 | 0.78 | 0.97 | 0.00 | 0.38 | 0.18 | 0.09 | 0.00 | 0.24 | 0.00 | 0.00 | 0.00 | 0.01 | 0.43 | 0.07 | 0.99 | 0.63 | 0.40 | 0.63 | 0.03 | 0.00 | 0.00 |
| MEturquoise | 0.54 | 0.67 | 0.32 | 0.87 | 0.45 | 0.63 | 0.47 | 0.70 | 0.17 | 0.67 | 0.78 | 0.39 | 0.13 | 0.45 | 0.31 | 0.83 | 0.47 | 0.82 | 0.82 | 0.56 | 0.36 | 0.20 | 1.00 | 0.30 | 0.13 | 0.77 | 0.87 | 0.22 |
| MEviolet | 0.20 | 0.02 | 0.96 | 0.18 | 0.47 | 0.16 | 0.17 | 0.20 | 0.79 | 0.09 | 0.61 | 0.47 | 0.08 | 0.03 | 0.11 | 0.10 | 0.41 | 0.09 | 0.06 | 0.89 | 0.77 | 0.31 | 0.78 | 0.41 | 0.38 | 0.03 | 0.05 | 0.09 |
| MEwhite | 0.43 | 0.13 | 0.29 | 0.77 | 0.24 | 0.75 | 0.90 | 0.08 | 0.60 | 0.14 | 0.80 | 0.07 | 0.00 | 0.19 | 0.05 | 0.63 | 0.88 | 0.17 | 0.38 | 0.84 | 0.16 | 0.02 | 0.72 | 0.31 | 0.05 | 0.16 | 0.14 | 0.21 |
| MEyellow | 0.38 | 0.19 | 0.06 | 0.58 | 0.02 | 0.99 | 0.85 | 0.05 | 0.73 | 0.92 | 0.67 | 0.06 | 0.06 | 0.26 | 0.01 | 0.96 | 0.16 | 0.88 | 0.57 | 0.91 | 0.09 | 0.04 | 0.34 | 0.23 | 0.04 | 0.04 | 0.66 | 0.92 |
