## Supplemental Table 10 for "Human alveolar macrophage response to *Mycobacterium tuberculosis*: immune characteristics underlying large inter-individual variability"

**Supplementary File 10. Differentially expressed (DE) genes selected from a curated M1/M2 macrophage gene list**

| CLASS | SUB | CATEGORY | GENES | INCLUDE | Module_2hr | Module_24h | Module_72h | Fold_change_pval_2h | Fold_change_2h | Fold_change_pval_24h | Fold_change_24h | Fold_change_pval_72h | Fold_change_72h |
|---|---|---|---|---|---|---|---|---|---|---|---|---|---|
| M2 | M2A | Transcription Factors | GATA3 | N | blue | | purple | 1.00 | 0.07 | 1.00 | 0.03 | 0.20 | 0.30 |
| M2 | M2D | Transcription Factors | IRF3 | N | brown | red | turquoise | 1.00 | -0.04 | 1.00 | 0.04 | 0.33 | -0.13 |
| M2 | | Transcription Factors | STAT6 | N | pink | blue | orange | 1.00 | -0.03 | 0.08 | 0.20 | 0.09 | 0.32 |
| M1 | M1 | Surface Markers | CD86 | N | midnightblue | pink | red | 1.00 | -0.02 | 0.24 | 0.17 | 0.81 | 0.05 |
| M2 | M2C | Surface Markers | TLR1 | N | black | blue | blue | 1.00 | 0.00 | 1.00 | 0.04 | 0.08 | 0.36 |
| M2 | M2D | Surface Markers | MSR1 | N | royalblue | green | skyblue | 1.00 | -0.03 | 0.13 | -0.13 | 1.00 | 0.00 |
| M1 | M1 | Released Cytokines | IL12A | N | | | | 1.00 | 0.37 | 0.80 | 1.35 | 0.55 | -1.12 |
| M2 | | Released Cytokines | TGFB2 | N | | | | 1.00 | 0.26 | 1.00 | -0.04 | 0.86 | -0.22 |
| M1 | GLYCO | Metabolic Enzymes | TPI1 | N | yellow | blue | lightcyan | 1.00 | 0.03 | 0.65 | 0.16 | 1.00 | 0.08 |
| M1 | GLYCO | Metabolic Enzymes | GAPDH | N | yellow | greenyellow | lightcyan | 1.00 | 0.02 | 1.00 | 0.00 | 0.18 | 0.24 |
| M1 | GLYCO | Metabolic Enzymes | GPI | N | turquoise | magenta | orange | 1.00 | 0.03 | 1.00 | -0.02 | 1.00 | 0.03 |
| M1 | GLYCO | Metabolic Enzymes | PGK1 | N | pink | | green | 1.00 | -0.04 | 1.00 | 0.00 | 0.21 | 0.09 |
| M1 | GLYCO | Metabolic Enzymes | ALDOA | N | | | | 1.00 | 0.38 | 1.00 | -0.04 | 1.00 | 0.02 |
| M1 | GLYCO | Metabolic Enzymes | PGK2 | N | | | | 1.00 | -0.02 | 1.00 | -0.22 | 1.00 | 0.03 |
| M1 | GLYCO | Metabolic Enzymes | CAD | N | | | | 1.00 | -0.03 | 0.17 | -0.20 | 0.13 | -0.16 |
| M1 | GLYCO | Metabolic Enzymes | NOS2 | N | | | | 1.00 | 0.41 | 1.00 | 0.38 | 1.00 | 0.35 |
| M2 | OXPHOS | Metabolic Enzymes | UQCRQ | N | blue | midnightblue | blue | 1.00 | -0.02 | 0.06 | -0.23 | 0.38 | -0.18 |
| M2 | OXPHOS | Metabolic Enzymes | NDUFA1 | N | lightcyan | red | skyblue | 1.00 | -0.03 | 0.27 | -0.12 | 0.32 | 0.10 |
| M2 | OXPHOS | Metabolic Enzymes | TLCD2 | N | | | lightgreen | 1.00 | -0.01 | 0.53 | -0.17 | 0.16 | 0.27 |
| M2 | OXPHOS | Metabolic Enzymes | ARG1 | N | | | | 1.00 | 0.73 | 1.00 | -0.03 | NA | NA |
| M2 | OXPHOS | Metabolic Enzymes | CYP2F1 | N | | | | 1.00 | -0.56 | 1.00 | 0.00 | 1.00 | -0.05 |
| M2 | OXPHOS | Metabolic Enzymes | GSTA1 | N | | | | 1.00 | -0.17 | 1.00 | -0.40 | 1.00 | 0.16 |
| M1 | GLYCO | Transcription Factors | HIF1A | Y | green | turquoise | blue | 1.00 | 0.03 | 0.00 | -0.25 | 0.00 | -0.48 |
| M1 | M1 | Transcription Factors | STAT3 | Y | yellow | blue | brown | 1.00 | -0.01 | 0.00 | 0.30 | 0.25 | 0.22 |
| M1 | M1 | Transcription Factors | STAT1 | Y | black | darkgrey | darkred | 1.00 | 0.01 | 0.00 | 1.08 | 0.00 | 1.68 |
| M1 | M1 | Transcription Factors | NFKB1 | Y | purple | darkturquoise | tan | 0.97 | 0.24 | 0.06 | 0.18 | 0.00 | 0.61 |
| M1 | M1 | Transcription Factors | IRF4 | Y | midnightblue | pink | | 1.00 | -0.06 | 0.00 | 0.89 | 0.00 | 1.76 |
| M1 | M1 | Transcription Factors | JUN | Y | magenta | | | 0.00 | 0.63 | 0.99 | 0.17 | 0.00 | 1.23 |
| M2 | M2A | Transcription Factors | PPARG | Y | darkred | pink | | 1.00 | -0.05 | 0.73 | 0.12 | 0.00 | -0.32 |
| M1 | M1 | Surface Markers | CD80 | Y | yellow | turquoise | brown | 1.00 | 0.10 | 0.00 | 0.55 | 0.00 | 1.22 |
| M1 | M1 | Surface Markers | CIITA | Y | black | | darkgrey | 1.00 | 0.10 | 0.00 | 0.55 | 0.04 | -0.33 |
| M2 | M2A | Surface Markers | IL1RN | Y | salmon | brown | lightgreen | 1.00 | -0.07 | 0.00 | 0.97 | 0.00 | 1.18 |
| M2 | M2A | Surface Markers | CD36 | Y | yellow | | black | 1.00 | 0.04 | 0.00 | -0.51 | 0.00 | -0.82 |
| M2 | M2C | Surface Markers | TLR8 | Y | darkred | green | darkgrey | 1.00 | -0.02 | 0.00 | -0.54 | 0.00 | -0.70 |
| M2 | | Surface Markers | MRC1 | Y | turquoise | | | 1.00 | 0.01 | 0.00 | 1.13 | 0.00 | 0.63 |
| M2 | | Surface Markers | CD163 | Y | yellow | | | 1.00 | -0.02 | 0.00 | -0.26 | 0.01 | -0.33 |
| M1 | M1 | Released Cytokines | TNF | Y | turquoise | green | black | 0.03 | 0.30 | 0.00 | 1.42 | 0.00 | 1.77 |
| M1 | M1 | Released Cytokines | IL1B | Y | blue | turquoise | tan | 0.00 | 1.28 | 0.00 | 2.29 | 0.00 | 1.83 |
| M1 | M1 | Released Cytokines | IL23A | Y | | | brown | 0.29 | 0.91 | 0.00 | 2.53 | 0.00 | 2.67 |
| M1 | M1 | Released Cytokines | IL6 | Y | royalblue | | tan | 0.00 | 1.77 | 0.00 | 3.73 | 0.00 | 5.53 |
| M2 | M2D | Released Cytokines | VEGFA | Y | turquoise | midnightblue | tan | 1.00 | -0.10 | 0.01 | -0.41 | 0.00 | 1.51 |
| M2 | | Released Cytokines | IL10 | Y | | | | 0.00 | 1.58 | 0.44 | 0.40 | 0.00 | -0.98 |
| M1 | GLYCO | Metabolic Enzymes | HK1 | Y | blue | black | steelblue | 1.00 | -0.05 | 1.00 | 0.03 | 0.00 | -0.31 |
| M1 | GLYCO | Metabolic Enzymes | ENO2 | Y | darkturquoise | green | lightgreen | 1.00 | 0.03 | 1.00 | 0.00 | 0.00 | 0.91 |
| M1 | GLYCO | Metabolic Enzymes | ENO1 | Y | black | pink | grey60 | 1.00 | -0.09 | 0.88 | 0.05 | 0.00 | 0.19 |
| M1 | GLYCO | Metabolic Enzymes | PKM | Y | | pink | pink | 1.00 | 0.07 | 0.93 | 0.09 | 0.00 | 0.75 |
| M1 | GLYCO | Metabolic Enzymes | SLC2A1 | Y | | | pink | 1.00 | 0.07 | 0.89 | 0.10 | 0.00 | 1.00 |
| M1 | GLYCO | Metabolic Enzymes | GP1BA | Y | | | | 1.00 | 0.59 | 0.00 | 0.75 | 0.00 | 3.59 |
| M1 | GLYCO | Metabolic Enzymes | PFKFB3 | Y | lightgreen | | black | 1.00 | 0.06 | 0.13 | 0.21 | 0.05 | -0.38 |
| M2 | OXPHOS | Metabolic Enzymes | GLUL | Y | cyan | black | yellow | 1.00 | 0.00 | 0.87 | -0.08 | 0.00 | -0.57 |
| M2 | OXPHOS | Metabolic Enzymes | IDO1 | Y | | darkgrey | tan | 1.00 | 0.01 | 0.00 | 3.86 | 0.00 | 5.28 |
| M2 | OXPHOS | Metabolic Enzymes | NDUFB1 | Y | magenta | magenta | turquoise | 1.00 | -0.05 | 0.01 | -0.27 | 0.00 | -0.46 |
| M2 | OXPHOS | Metabolic Enzymes | NDUFS3 | Y | | red | | 1.00 | 0.02 | 1.00 | -0.03 | 0.00 | 0.24 |
| M2 | OXPHOS | Metabolic Enzymes | CACNB3 | Y | green | salmon | green | 1.00 | 0.09 | 0.01 | 0.32 | 0.06 | 0.32 |
| M2 | OXPHOS | Metabolic Enzymes | HADH | Y | grey60 | | pink | 1.00 | -0.06 | 0.00 | -0.52 | 0.00 | -1.03 |
| M2 | OXPHOS | Metabolic Enzymes | CAT | Y | greenyellow | | tan | 1.00 | -0.02 | 0.01 | -0.37 | 0.00 | -0.68 |
| M2 | OXPHOS | Metabolic Enzymes | SLC11A1 | Y | blue | | turquoise | 1.00 | 0.02 | 0.02 | 0.37 | 0.00 | 0.37 |
| M2 | OXPHOS | Metabolic Enzymes | ACYP2 | Y | brown | | yellow | 1.00 | -0.06 | 0.05 | 0.34 | 0.42 | -0.18 |
| M2 | OXPHOS | Metabolic Enzymes | SHPK | Y | red | | pink | 1.00 | 0.14 | 0.00 | 0.51 | 0.00 | 0.56 |
